# Dual chromosomal origins of replication (*oriC*) in the genomes of the Halomonadaceae – a prediction study

**DOI:** 10.1101/2025.08.05.668638

**Authors:** Christoph Weigel

## Abstract

Chromosome replication is a key process in the bacterial cell cycle. Most, but not all, bacteria follow the "Replicon Model" for the initiation step: Their genomes encode an initiator and contain a replicator locus at which the initiator acts as activator. The single chromosomes of the model organisms *E. coli* and *B. subtilis* each contain one *dnaA* gene encoding the initiator protein DnaA, and one replication origin, *oriC*. The single chromosome of *P. aeruginosa* is the only exception known to date that contains two functional replication origins, *oriC1* and *oriC2*. Since a preliminary study indicated that this unusual dualorigin configuration is also found in other Pseudomonadaceae, it seemed interesting to examine whether it can also be predicted for the Halomonadaceae, a sister clade of the Pseudomonadaceae. Here I show that of 163 Halomonadaceae genomes with an annotated *dnaA* gene, 23 have a single predicted origin, *oriC2*, and 138 have dual predicted origins, *oriC1* and *oriC2*, with the genomic localizations of both corresponding to those in the *P. aeruginosa* genome. The distribution of species/strains with single or dual predicted *oriC*·s in the taxogenomic tree of the Halomonadaceae suggests that a demise of *oriC1* or its "downgrading" occurred repeatedly during their diversification. Experimental studies of chromosome replication in individual species/strains of the Halomonadaceae, and likewise computational cell cycle simulations, can benefit from taking the single/dual origin predictions presented here into account.

## 1 Introduction

A theoretical framework for the initiation of chromosome replication in *Escherichia coli* was outlined by Jacob, Brenner and Cuzin in their "Replicon Model" of 1963: *"A unit capable of independent replication or replicon would carry two specific determinants. I. A structural gene controlling the synthesis of a specific initiator. 2. An operator of replication, or replicator, i.e., a specific element of recognition upon which the corresponding initiator would act, allowing the replication of the DNA attached to the replicator."* (1).

Regardless of known backup mechanisms such as ’Stable DNA Replication’ (SDR) (2), it is safe to state that the *"specific initiator"* of bi-directional chromosome replication in *E. coli* is a protein, DnaA, and the *"replicator"* is conventionally referred to as the origin of chromosome replication, *oriC* (3). The origin features multiple DnaA binding sites (DnaA boxes), while DnaA contains ATP/ADP-binding (domain 3) and DNA-binding domains (domain 4: dsDNA; domain 3: ssDNA) (3, 4). When during growth enough DnaA-ATP has accumulated in the cell, an active initiation complex can be formed at the origin resulting in strand opening and recruitment of the replicative helicase, DnaB, by interaction with *oriC*-bound DnaA (5, 6). Strand opening requires *oriC* to be negatively supercoiled and occurs within the DNA unwinding element (DUE), an AT-rich region flanking the array of DnaA boxes (7, 8). Recently, DnaA-ATP was shown to assemble into a continuous oligomer in *Bacillus subtilis oriC* at the site of DNA strand opening, which extends from a dsDNA anchor at the DUE-proximal DnaA box(es) to capture a ssDNA strand at the DnaA-trio motifs, (NAN)n, via its binding to DnaA domain 3 (**Figure 3**) (8, 9).

The "Replicon Model" applies to all experimentally confirmed DnaA-dependent chromosomal replication origins of species from various bacteria phyla, all of which contain an AT-rich region flanked by an array of DnaA boxes (10–19). This "array of DnaA boxes" is a hallmark of bacterial *oriC*·s and used as one criterion for their prediction by bioinformatics methods (20, 21). However, these "arrays…" vary greatly in length and are extremely heterogeneous in terms of the number of DnaA boxes, their degree of deviation from the consensus sequence 5’-TTWTNCACA (22), and their respective distances and orientations, forward (fwd) or reverse (rev). Furthermore, these "arrays…" can be contiguous, as in *E. coli*, or split into two, as in *B. subtilis*, with one sub-array flanked by the DUE and the second sub-array distal at a genomic distance of ∼1.5 kb (10, 11). The more strictly-defined conserved structural *oriC* features are: 1. one DNA unwinding element (DUE) (8), which is 2. flanked downstream at a distance of ∼2 helical turns by one or two closely spaced reverse-oriented (rev) consensus DnaA box(es) designated "R1" in reference to the DnaA box R1 in *E. coli oriC* (5’-TGTGNAWAA) (22), and which is 3. flanked upstream by a gene with counter-clockwise (ccw) transcriptional direction. DnaA-trio motifs located between the DUE and the DUE-proximal, conserved "R1" DnaA box(es) can be assigned to most experimentally confirmed and the majority of predicted bacterial *oriC* sequences (9).

The "Replicon Model" is generally understood to mean that in bacterial chromosomes one single gene encodes the cis-acting *"specific initiator"* and one single DNA segment represents the *"replicator"* (1). Indeed, the chromosomes of the model organisms *E. coli* (10), *B. subtilis* (*11*), *Caulobacter crescentus* (13), and of other bacteria whose replication origin has been experimentally confirmed (14–19) contain one single *dnaA* gene and, with one exception, one single *oriC*. This exception is *Pseudomonas aeruginosa* PAO2003 for which Yee & Smith (1990) found two sequences pae310 (*oriC1*) and pae301 (*oriC2*), both of which could drive autonomous replication of minichromosomes in *P. aeruginosa* PAO2003 and *P. putida* KT2440 (12). Later, Jiang *et al.* (2006) demonstrated that plasmids carrying *oriC1* and *oriC2* of *P. aeruginosa* PAO116 both form pre-priming complexes *in vitro* (DnaA-dependent strand opening and helicase loading), albeit *oriC2* to a lesser extent, and thus both are *bona fide* replication origins (23). It has not been investigated whether these two origins may be differentially regulated or whether it is irrelevant for cell cycle regulation which origin "fires", or both, as they are located only ∼5 kb apart on the 5.5–7 Mbp chromosomes of *P. aeruginosa* strains (**Figure 1**) (24).

**Figure 1.**
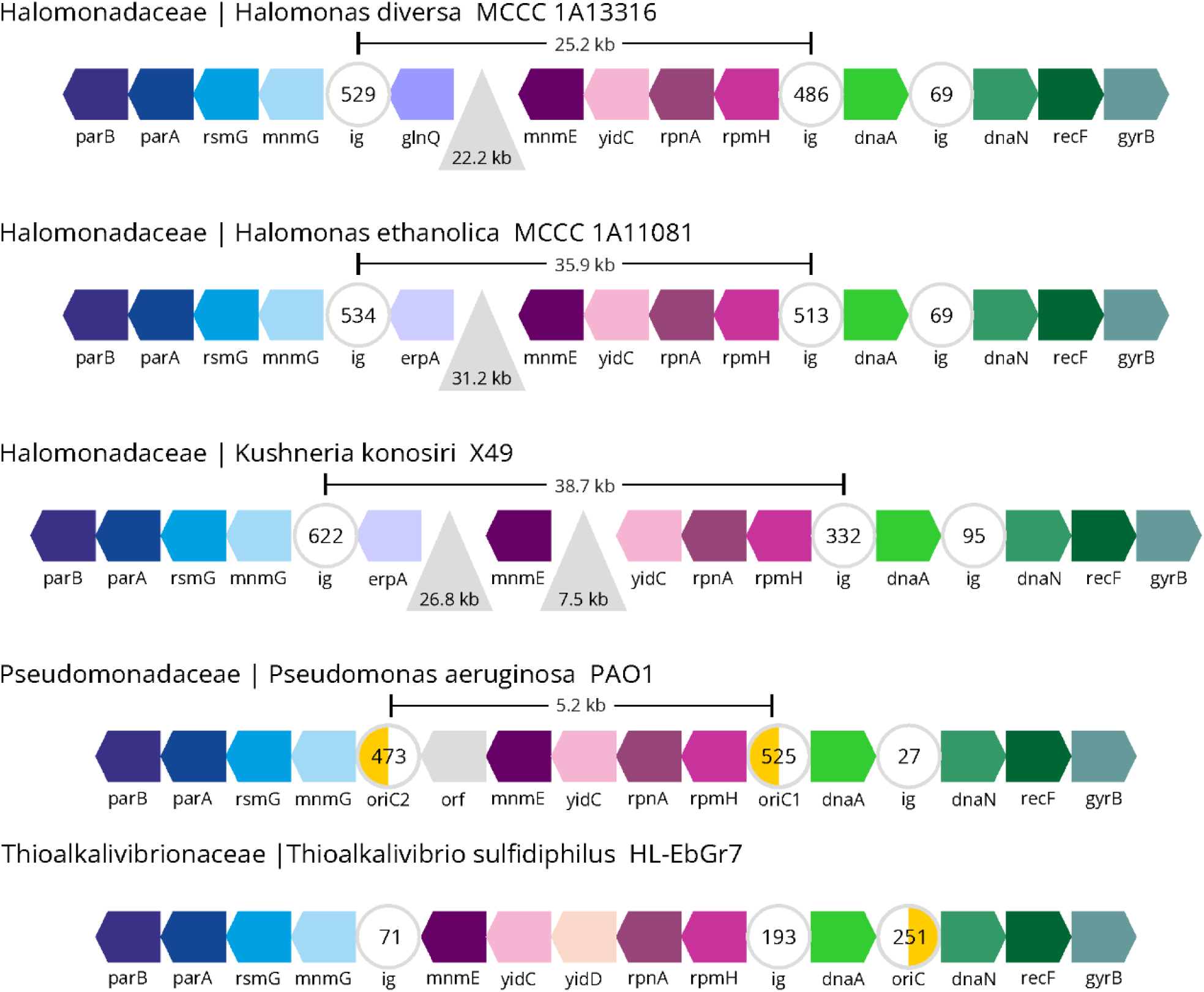
Gene order in the *parB—gyrB* region of three genomes of the Halomonadaceae, *H. diversa* MCCC 1A13316 [NZ_CP053382.1], *H. ethanolica* MCCC 1A11081 [NZ_JABFTX010000004.1], *K. konosiri* X49 [NZ_CP021323.1], *P. aeruginosa* PAO1 [NC_002516.2] (Pseudomonadaceae), and *T. sulfidiphilus* HL-EbGr7 [NC_011901.1] (Thioalkalivibrionaceae). Genes are not shown to scale, orthologous genes are shown in the same color and the direction of transcription is indicated by arrowhead. The positions and the lengths in kb of insertions in the three Halomonadaceae genomes are indicated by light-gray triangles. Only the lengths of the intergenic regions (ig) relevant here are given in base pairs (bp); the relative positions of predicted DUE·s in the ig·s labeled "oriC" are indicated by the orange half-circle.

In a pilot study on the prediction of *oriC*·s in the Gammaproteobacteria, I found this dual-origin configuration, *oriC1* and *oriC2*, in 88 of 91 examined genomes of species of the GTDB-family Pseudomonadaceae (25), with *oriC1* not detectable only in *P. pohangensis* DSM 17875 and presumably inactivated by insertion in *P. syringae* CFBP6109 and *P. savastanoi* 1448A. The situation was less clear for a few examples from the (phylogenetic) sister family of the Pseudomonadaceae, the Halomonadaceae: while *oriC1* and *oriC2* could be predicted for *Halomonas elongata* DSM 2581, only *oriC2* could be predicted for *Kushneria marisflavi* SW32 (EMBO Workshop in Kyllini, Greece, 2018; Suppl. Poster 1).

The recently published taxogenomic investigation of the family Halomonadaceae comprising 168 species/strains by de la Haba *et al.* (2023) was my incentive to 1. contribute predictions for the *oriC* in these genomes, and 2. to examine the distribution of predicted *oriC1*·s and *oriC2*·s in this extended group of comparatively closely related genomes (26). Since the methodology I use here to predict *oriC* in bacterial genomes has been successfully applied in a few individual cases for identification and subsequent experimental confirmation (14–17, 19, 27), but has not yet been thoroughly documented and evaluated on a larger data set, this is done in the present study.

## 2 Materials and Methods

### 2.1 Genome sequences and sequence analysis

All Genome sequences used in this study were retrieved from the NCBI RefSeq or Nucleotide databases, the accession numbers are listed in Suppl. Table 1 (28). Retrieved sequences were further processed with LibreOffice 24.2.1.2. Writer software. Graphics were generated with CorelDRAW^®^X8 (Version 18.1.0.661). GC-skew analyses for complete genomes or larger contigs (>50 kb) were performed online with the Webskew server (https://genskew.csb.univie.ac.at/webskew) (29). SIDD analyses of genomic sequence segments were performed with locally installed SIST software and further processed with Libre-Office 24.2.1.2. Calc (30, 31).

### 2.2 A manual method for the prediction of bacterial replication origins (*oriC*)

The sequential steps required to predict a DnaA-dependent origin of replication (*oriC*) in a bacterial genome of interest are detailed below.

#### 2.2.1

The *dnaA* gene is to be identified in a closed/complete and annotated genome in question. In un-annotated genome sequences, the *dnaA* gene can be identified by tBLASTn using a verified DnaA protein as query, e.g., DnaA of *E. coli* K-12 [AAC76725.1] (32). The *dnaA* gene is located on the main chromosome in most bacterial genomes, and only exceptionally on a secondary chromosome/megaplasmid as in *Paracoccus denitrificans* PD1222 [*oriC* NC_008686.1; *dnaA* NC_008687.1]. Occasionally genomes contain two complete *dnaA* genes, e.g. in species from the GTDB-classes Desulfovibrionia and Chlamydiales. *dnaA*-less genomes are known from some Cyanobacteria and from reduced genomes of endosymbionts, so prediction of *oriC* in these cases is pointless. The DnaA protein of a genome in question should have a length of ∼450 residues and over its entire length >30% BLASTp-identity with DnaA of *E. coli* K-12 [AAC76725.1] (32). Please note that annotation software occasionally does not recognize the GTG start codon commonly found in *dnaA* genes..

#### 2.2.2

The minimum of the cumulative GC-skew (inflection point) in a closed genome is determined with appropriate software, e.g., SkewIT (29). Larger contigs (>50 kb) can be screened for their GC-skew in the case of un-closed complete genomes, but often do not give clear results.

#### 2.2.3

Since the cumulative GC-skew curve is in many cases not "V"-shaped but rather "U"-shaped, or jagged with secondary minima, the inflection point cannot always be unambiguously determined. It must also be kept in mind that the GC-skew minimum indicates the larger "origin region", but does not define *oriC* per se. It is therefore necessary to screen all >100 bp-long intergenic regions (ig) on both sides of a GC-skew minimum for structural features of an *oriC*. Based on previous experience, intergenic regions with *oriC* features are usually located at a distance of 0–20 kb from the GC-skew minimum, but larger distances have been found, e.g., in the Alphaproteobacteria (unpubl. observation).

#### 2.2.4

Intergenic regions in the vicinity of the GC-ske minimum that contain at least one consensus DnaA box (5’-TTWTNCACA) are analysed for the presence of a DNA unwinding element (DUE), a key structural element of a bacterial *oriC* (22, 8). For the SIDD analysis to identify a DUE, the intergenic region in question is prepared so that it is located central in a 2.5 kb-long sequence segment.

#### 2.2.5

A 2.5 kb-long sequence segment (Section 2.2.4) is used as input for the SIDD subprogram of SIST (31). Briefly, the SIDD algorithm calculates for each basepair in a stretch of dsDNA the base-context dependent Gibbs energy G (kcal·mol^−1^) required for its transition from dsDNA to ssDNA at a given superhelicity σ. Fixed parameters are salt concentration (0.1 M) and temperature (37°C), and the DNA is assumed to be circular DNA (linear input DNA is artificially "circularized" by adding 50 G/C basepairs at the ends). For a complete cycle, (negative) superhelicity at physiologically relevant levels is sequentially assayed in the range of σ=–0.040 (low) to σ=–0.060 (high) by increments of 0.005.

#### 2.2.6

Strong SIDD signals for the program run at σ=–0.060 (high neg. superhelicity) are found mostly within the intergenic region in question, located centrally in the 2.5 kb-long sequence segment (Section 2.2.4). This can be checked in the SIDD output (.txt file), and if it is the case, further analysis can be limited to evaluating the central 500 bp sequence stretch (most *oriC*·s have a length of 250–450 bp and "fit" into window of 500 bp). The SIDD output (.txt files) is transferred to a spreadsheet program for all tested σ-values (σ=–0.040 – σ=–0.060) and displayed in a diagram (G-values over sequence position) as line graphs (**Figure 2**, for examples). SIDD signals are identified in the diagrams as stretches of 20–50 bp in which the G required for strand separation decreases continuously with increasing negative superhelicities. Sequence stretches that do not show this dependency, i.e., show high strand-separation proficiency for each tested superhelicity are not considered. Note that the SIDD program does not support simulating the influence of bound protein(s) on the unwindability of a sequence segment under investigation. This must be taken into account, but affects the expected accuracy in an unquantifiable way. I designate SIDD signals as "strong" that have G values of 0–3 at high negative superhelicity (σ=-0.06) in a stretch of 20-50 bp, as "weak" for values of 3-6, and do not consider signals if the values are >6. This classification is arbitrary, but has proven itself in practice, e.g., in a proof-of-concept application on the known *oriC*·s of *E. coli* and *B. subtilis* (**Figure 2**, **Figure 3**), and in the experimental mapping of the unwinding region in the *oriC* of *H. pylori* (14). "Strong" and "weak" SIDD signals are marked with their width in the sequence file (Suppl. Table 2, for examples).

**Figure 2.**
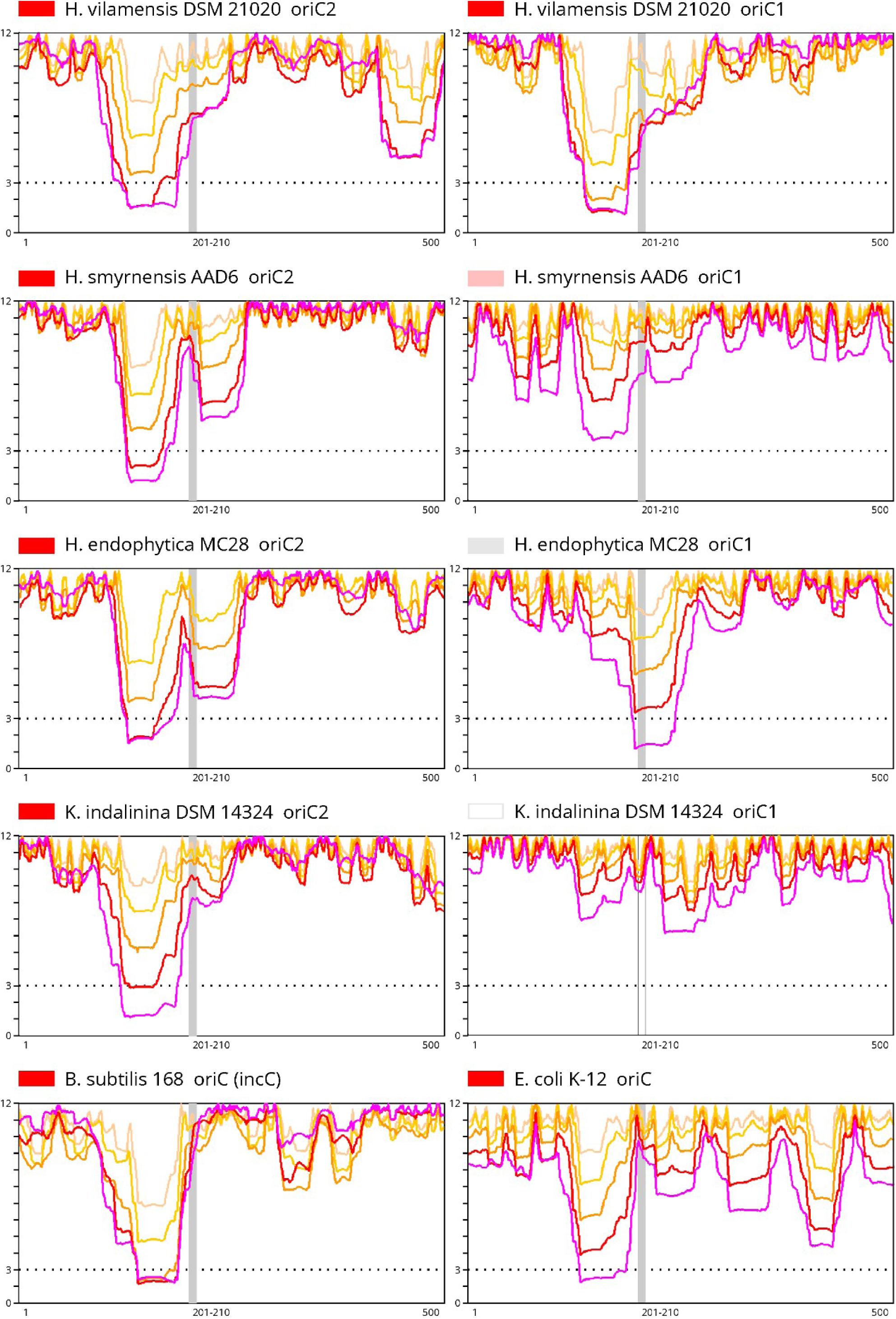
SIDD profiles for "*oriC2*" and "*oriC1*" of four genomes of the Halomonadaceae, and SIDD profiles of the *oriC* of *E. coli* MG1655 and *B. subtilis* 168 for comparison. X axis: 500 bp long "central" segment (see text); the vertical gray strip e (pos. 201–210) indicates the position of the "R1" DnaA box. Y axis: G(x) (kcal·mol^−1^), 0–3 strong SIDD signal for strand separation, 3–6 weak SIDD signal for strand separation, 3–12 no SIDD signal for strand separation. In theory, a value of 0 indicates complete strand separation, a value of 12 indicates fully double-stranded DNA. Color code: SIDD at σ=–0.060 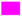, at σ=–0.055 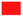, at σ=–0.50 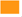, at σ=–0.45 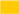, at σ=–0.040 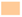 (σ: superhelicity).

#### 2.2.7

If the SIDD analysis results in a "strong" or "weak" signal for a sequence stretch of ∼50 bp at a distance of ∼2 helical turns from the previously (Section 2.2.4) detected DnaA box, it can safely be considered as DNA unwinding element (DUE) but additional criteria must be met to designate the intergenic region (ig) in question as predicted *oriC* (Sections 2.2.9 – 2.2.12). It is helpful for further analysis and the comparison with known *oriC*·s to arrange the entire sequence – if necessary as reverse-complement – such that the DnaA box is in reverse orientation (5’-TGTGNAWAA). See Section 2.2.12 in case no strong SIDD signal is found.

#### 2.2.8

The previously (2.2.4) detected DnaA box in the intergenic region in rev orientation (or arranged for by reverse-complement) at a distance of ∼2 helical turns from the right flank of the SIDD signal can be 1. a single DnaA box as in *E. coli oriC* ("R1"), 2. twin DnaA boxes that are closely spaced, or directly adjacent as in *oriC·*s of Actinomycetota, or overlapping by –1 as in *oriC·*s of Campylobacterota, or 3. even a triple box as in *B. subtilis* 168 *oriC*(*incC*) (**Figure 3**). In the case of twin or triple DnaA boxes only a comparison with known *oriC·*s can distinguish which box corresponds to the "R1" DnaA box in *E. coli oriC* (**Figure 3**).

**Figure 3.**
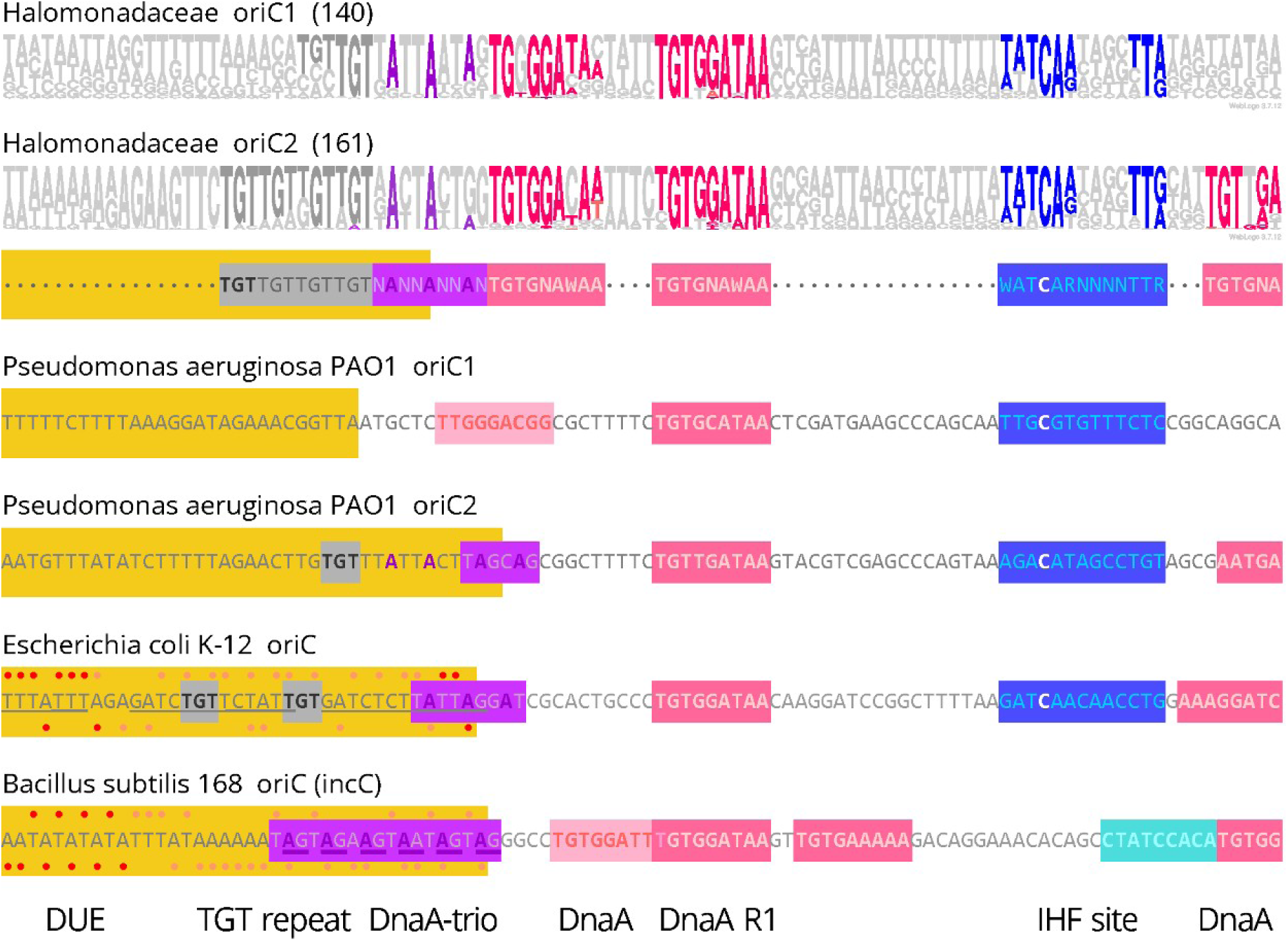
WebLogo consensus for 140 *oriC1* and 161 *oriC2* sequences for the Halomonadaceae from pos. 150–249 of the 500 bp long "central" segment containing the conserved R1-type DnaA box at pos. 201–210 (45). The corresponding sequences of *oriC1* and *oriC2* of *P. aeruginosa* PAO1 [NC_002516.2], and the *oriC* sequences of *E. coli* K-12 [NC_000913.3] and *B. subtilis* 168 [NC_000964.3] are included for comparison and likewise aligned on the R1 DnaA box. Orange: DUE according to SIDD analysis. Red: consensus DnaA box, rev orientation 5’-TGTGNAWAA (22). Pale red: relaxed DnaA box, rev. orientation. Sky blue: relaxed DnaA box, fwd orientation 5’-TTWTNCACA. Dark blue: IHF-binding site (white: conserved C) (41). Magenta: DnaA-trio motif (NAN)_n_ (9); bases bound by DnaA domain 3 are underlined (4). Gray: TGT repeats. In the *E. coli* and the *B. subtilis* DUE sequences, single-stranded bases found by KMnO_4_-footprinting for the upper strand (above) and the lower strand (below) are indicated by red dots; pale red dots demark bases that were found as single-stranded when SSB was included in the experiment (38). Underline: 13-mer repeats in *E. coli oriC* (42).

#### 2.2.9

The intergenic region in question is examined for the direction of transcription of the flanking genes. If the sequence is arranged such that the DUE is flanked downstream (to the right) at a distance of ∼2 helical turns by the rev-oriented DnaA box(es) (5’-TGTGNAWAA), the upstream flanking gene (to the left) should have counter-clockwise (ccw) direction of transcription, either driven by its own promoter as, e.g., *mnmG* in *E. coli* (<*mnmG·oriC*) or as part of a larger transcription unit/operon as, e.g., *dnaN* in *B. subtilis* (<*dnaN·oriC·<dnaA*). Ccw transcriptional direction of the gene to the left/upstream of *oriC* was found without exception in all experimentally confirmed *oriC*·s (10–19) and, in addition, in virtually all >1 k *oriC* predictions obtained with the methodology described here (unpubl. observation).

#### 2.2.10

The intergenic region in question is examined for the presence of DnaA-trio motifs (NAN)n between the DUE and the DUE-proximal, conserved DnaA box(es). DnaA-trio motifs are found in most experimentally confirmed and the majority of predicted bacterial *oriC* sequences (9).

#### 2.2.11

The assignment of DnaA boxes in the intergenic region in question is made by eye or using suitable software, e.g., programs from the MEME suite (33). No expectation can be specified for the number of DnaA boxes in an *oriC*, it can be anything between the (predicted) five in *Koribacter versatilis* Ellin345 [NC_008009.1] and the (predicted) 18 in *Anaplasma phagocytophilum* HZ [NC_007797.1] (unpubl. observation). The ∼30 DnaA boxes predicted for the *oriC*·s of *Streptomyces* spp. are exceptional for these unusually long replication origins (∼0.9 kb) (17). DnaA boxes can be found on both sides of the DUE, but the majority of them will usually be clustered on one side of the DUE. For the DnaA box assignment, both orientations, forward (fwd) and reverse (rev), of the 9 bp-long DnaA box are taken into account, as well as the consensus sequence (5’-TTWTNCACA) containing 1 mismatch, and, in addition, relaxed DnaA boxes with up to 4 mismatches (22). With one exception, the 9-mer DnaA boxes found so far for bacteria of all phyla match well with the *E. coli* consensus sequence (5’-TTWTNCACA), allowing for one mismatch. A 12-mer DnaA box (5’-AAACCTACCACC) has been experimentally confirmed only for *Thermotoga maritima* (34). It is especially advisable to consider relaxed variants when assigning DnaA boxes, as this allows to check whether they point to arrays of regularly spaced DnaA boxes that serve as anchor for DnaA filaments as in *E. coli oriC* (35).

#### 2.2.12

The intergenic region in question is considered a predicted *oriC* if it matches the following criteria: 1. one DNA unwinding element (DUE) (8), that is 2. flanked downstream at a distance of ∼2 helical turns by at least one reverse-oriented (rev) consensus DnaA box(es) (22), and that is 3. flanked upstream by a gene with counter-clockwise (ccw) transcriptional direction, 4. an array of DnaA boxes contiguous with the DUE-flanking DnaA box(es), and 5. DnaA-trio motifs located between the DUE and the DUE-proximal, conserved DnaA box(es) (9). Criterion 2 must be met in any case, criterion 5 is optional, but if criterion 1 is not met, an *oriC* with reduced functionality can still be assumed if all others are met.

#### 2.2.13

If the intergenic region in question can be considered a predicted *oriC* because it meets all criteria (Section 2.2.12) but is shorter than ∼250 bp, it has to be clarified whether it is part of a bipartite *oriC* with an additional DnaA box array in an intergenic region at a distance of ∼1.5 kb.

#### 2.2.14

It is optional to screen the predicted *oriC* for binding sites of proteins known to bind to *oriC·s*, such as Fis, IHF, IciA, HU, Rob in *E. coli*, CtrA in several Alphaproteobacteria, and Spo0A in Bacillota. As these binding sites usually have a degenerate sequences, it is advisable to always screen several *oriC* sequences together using suitable software, e.g., programs of the MEME suite (33).

#### 2.2.15

When several more closely related *oriC* sequences are compared in an alignment, the detection of degenerate/relaxed binding sites of replication factors is typically more reliable. This is especially true for relaxed DnaA boxes (with up to 4 mismatches), but also for the detection of (NAN)n DnaA-trio motifs.

## 3 Results

### 3.1 Sizes of and distances between putative replication origins (*oriC*) in the genomes of the Halomonadaceae

Assuming that the gene synteny of the region encompassing *oriC2* and *oriC1* in the genome of *P. aeru-ginosa* PAO1 (12, 23) is largely conserved in the genomes of the Halomonadaceae, I extracted genome segments spanning the *parB* and *gyrB* genes from the 161 genomes in the RefSeq database (28) for which de la Haba *et al*. (2023) published a taxogenomic study (26). RefSeq sequences for *Halomonas rifensis* CECT 7698 (GCM10020179) were not available (January 2024), and I added the RefSeq sequences of *Halomonas campaniensis* LS21, *Halotalea alkalilenta* IHB B 13600, and *Chromohalobacter canadensis* 85B to the set, giving a total of 163 genomes for this study (Suppl. Table 1). All 163 genomes have annotated *dnaA* genes and the DnaA proteins show 88.7–99% BLASTp similarity to each other (32); the two *dnaA* genes annotated as pseudogenes do not fall outside this range upon tentative reconstruction (Suppl. Table 1).

A comparison of the gene order in the *parB—gyrB* segment in 149 genomes of the Halomonadaceae with that in the corresponding segment of the *P. aeruginosa* PAO1 genome shows that the gene synteny is largely conserved (**Figure 1**). In 13 genomes, *parA* and *gyrB* are contained in separate contigs, whose gene orders do not deviate from that of the 149 contiguous segments, which, however, does not allow an estimate of their distance (marked ’blank’ in Suppl. Table 1).

The position of *oriC2* in the *P. aeruginosa* PAO1 genome corresponds to a comparatively large ig upstream of *mnmG* in the Halomonadaceae genomes (med 564 bp; max 1388 bp; min 286 bp). The position of *oriC1* in *P. aeruginosa* corresponds to a similarly large ig between *rpmH* and *dnaA* in the Halomonadaceae genomes (med 534 bp; max 943 bp; min 330 bp). The comparatively small ig between *dnaA* and *dnaN*, as found in *P. aeruginosa* PAO1, is conserved in the Halomonadaceae (med 65; max 331; min 23). The exact sizes (in base pairs) of the three intergenic regions (ig) are listed individually for each genome in Suppl. Table 1.

An indication of the position(s) of the putative replication origin(s) in the genomes of the Halomonadaceae can be obtained by comparison with the replication origin region of the phylogenetically distantly related bacterium *Thioalkalivibrio sulfidiphilus* HL-EbGr7 (**Figure 1**; see also Suppl. Poster 1). The gene order in the *parB—gyrB* segment of the *T. sulfidiphilus* genome is identical to that in the *P. aeruginosa* PAO1 genome and in the genomes of the Halomonadaceae. However, the intergenic regions corresponding to the positions of *oriC2* and *oriC1* in the *P. aeruginosa* PAO1 genome are relatively short (71 bp and 193 bp, respectively), and *oriC* could be predicted for the somewhat longer *dnaA·dnaN* ig (251 bp). The location of *oriC* in *T. sulfidiphilus* corresponds to the known *oriC* location in the *dnaA·dnaN* ig in the genomes of *B. subtilis* 168 (11), *Clostridioides difficile* Δ*erm* (19), *Campylobacter jejuni* 81116 (16), and *Streptomyces coelicolor* A3(2) (17).

Since the length of the intergenic regions upstream of *mnmG* and between *rpmH* and *dnaA* in the genomes of the Halomonadaceae corresponds to those of *oriC2* and *oriC1* in the genome of *P. aeruginosa* PAO1, respectively, I tentatively refer to them as "*oriC2*" and "*oriC1*" in the following, i.e., without anticipating their verification as DnaA-dependent chromosomal origin of replication (*oriC*).

The synteny of the origin regions in the genomes of 149 Halomonadaceae and the origin region in the genome of *P. aeruginosa* PAO1 is interrupted by the insertion of about two dozen genes, which in all cases are inserted between *mnmG* and *glnQ*/*erpA* (**Figure 1**). In the genomes of the nine *Kushneria* sp. and few others genomes, an additional, shorter insertion is located between *mnmE* and *yidC* (**Figure 1**). Together, these insertions result in significantly larger distance between "*oriC2*" and "*oriC1*" (med 23.6 kb, max 102.3 kb, min 9.3 kb) compared to *P. aeruginosa* PAO1 (5.2 kb).

In three genomes I find a notable deviation from the common arrangement of genes in the origin region encompassing "*oriC2*" and "*oriC1*".

1. In the *Halomonas jeotgali* Hwa genome, intragenomic recombination has resulted in "*oriC2*" and "*oriC1*" and several of their flanking genes being present in reverse order, but without significantly changing the distance between them (33.7 kb) (marked "☓" in Suppl. Table 1).
2. In the *Halomonas chromatireducens* AGD 8-3 genome, intragenomic recombination split "*oriC2*" at the right-flanking *glnQ* gene from the contiguous segment encompassing *erpA*, *rpmH*, "*oriC1*", and *dnaA.* In this new configuration, which is otherwise not found in the genomes of the Halomonadaceae, "*oriC2*" and "*oriC1*" are 1.2 Mbp apart. Both have all the features of a fully functional *oriC* (Section 3.4), but two functional origins at this distance in one genome are unprecedented (except for artificial strain constructs, see (36)). I therefore suspect, but cannot prove, that an error occurred during assembly of the complete genome from contigs.
3. In the *Zymobacter palmae* IAM 14233 = DSM 1049 genome, a recombination event split "*oriC1*" such that "*oriC1*" and the 3’-adjacent *dnaA* gene were retained while the 5’-adjacent *mnmE—rpmH* genes were transposed to another genomic location and being replaced by the *sdiA* gene. In this configuration, "*oriC2*" and "*oriC1*" are in reverse order without significantly changing the distance between them (22.5 kb) (Suppl. Table 1).

### 3.2 Location of the GC-skew minimum in the genomes of the Halomonadaceae

The cumulative GC-skew of a bacterial genome can be used to narrow down the region in which the origin of replication (*oriC*) is located (37, 20). In most cases, *oriC* is located directly at or at a distance of a several kilo basepairs (kb) from the inflection point at the minimum of the cumulative GC skew curve. I determined the GC-skew minimum for the 149 Halomonadaceae genomes in which "*oriC2*" and "*oriC1*" are located on a single genomic segment of the complete genome or on a larger contig (>50 kb) (Section 2.2). With few exceptions, the GC-skew minimum (inflection point) is located closer to "*oriC2*" (med 5,8 kb, max 96,9 kb, min 0 kb), and distal to "*oriC1*" (med 29,1 kb, max 185,7 kb, min 0 kb) (listed in Suppl. Table 1). Since the GC-skew minimum is not used in the present study as a criterion to localize the origin(s) of replication, I leave it with this note.

### 3.3 Detection of R1-type DnaA boxes in putative origins ("*oriC*") and DUE prediction

To prepare for the assessment of whether a DUE can be predicted for the intergenic sequences tentatively designated "*oriC2*" and "*oriC1*" in the 163 Halomonadaceae genomes, I arranged the sequences such that the direction of transcription of the left-flanking genes *mnmG* and *rpmH* (*sdiA*), respectively, is counter-clockwise.

On visual inspection, I detected a consensus DnaA box in reverse orientation ("R1") approximately in the centre of 161 of 162 "*oriC2*" sequences and in 140 of the 163 "*oriC1*" sequences. This prompted me to further align all "*oriC*" sequences, so that the DnaA box R1 is located at pos. 201–210 of a 500 bp long "central" sequence segment in a 2.5 kb long sequence stretch containing "*oriC*" together with flanking gene sequences (Suppl. Table 2). No specific alignment was performed for the 24 "*oriC*" sequences lacking a detectable DnaA box R1, and the respective "*oriC*" sequences were kept as 2.5 kb sequence stretches with the flanking genes.

The SIST software (31) was applied to obtain predictions of approximately 10–60 bp long regions of superhelical ’stress-induced duplex destabilization’ (SIDD) (30) in the "*oriC*" sequences of the Halomonadaceae contained in the above mentioned 2.5 kb sequence stretches. Only the SIDD profiles obtained for the 500 bp long "central" segment were analyzed further as there were no repeatedly found SIDD predictions for other positions in the entire sequence set (except for pairs of sequences with significant nucleotide sequence identity). In all obtained SIDD profiles, SIDD signals were mostly confined to pos. 140–190 and/or pos. 210–260 in the 500 bp long "central" segment.

As outlined in Section 2.2, I refer to SIDD signals with values from 0 to 3 as "strong", those with values from 3 to 6 as "weak" and those above 6 as "absent" on a coarse qualitative scale. Since "strong" and "weak" SIDD signals at pos. 140–190 are located at a distance of ∼2 helical turns from the DnaA box R1 (pos. 201–210), I posit that these are the DUEs of the corresponding *oriC*.

I have juxtaposed the SIDD profiles for "*oriC2*" and "*oriC1*" of four species as examples of the most frequently found combinations in **Figure 2**. For comparison, I included the SIDD profiles of *E. coli* K-12 MG1655 *oriC* and of *B. subtilis* 168 *oriC* (*incC*), for which it has been experimentally shown that the SIDD signal at positions 140–190 corresponds to the DUE of these origins (38).

1. The SIDD profiles for *Halomonas vilamensis* DSM 21020 reveal a "strong" DUE for both "*oriC2*" and "*oriC1*" (labeled red/red).
2. The SIDD profiles for *Halomonas smyrnensis* AAD6 reveal a "strong" DUE for "*oriC2*" and a "weak" DUE for "*oriC1*" (labeled red/pale red).
3. The SIDD profiles for *Halomonas endophytica* MC28 reveal a "strong" DUE for "*oriC2*" and no DUE for "*oriC1*" (labeled red/gray)
4. The SIDD profiles for *Kushneria indalinina* DSM 14324 reveal a "strong" DUE for "*oriC2*" and no DUE for "*oriC1*" that also lacks the conserved DnaA box R1 at pos. 201–210 (labeled red/white)

SIDD signals at pos. 210–260 were found in about two thirds of the SIDD profiles for both "*oriC2*" and "*oriC1*" that overlap with the position of the conserved IHF-binding site (**Figure 3**). In **Figure 2**, this SIDD signal is prominent in "*oriC1*" of *H. endophytica* MC28, but also seen in "*oriC2*" of *H. smyrnensis* AAD6 and*H. endophytica* MC28. Because the SIDD signals at pos. 210–260 lack a conserved R1-type DnaA box ∼2 helical turns downstream, they are not considered as DUE and will not be discussed further. The entire set of SIDD predictions is available from the author upon request.

### 3.4 Conservation of other structural elements in putative replication origins ("*oriC*") and *oriC* assignment

In addition to the three structural elements of bacterial *oriC·*s mentioned in the introduction (ccw transcription of the left-flanking gene, a DNA unwinding element (DUE), and a R1-type DnaA box down-stream of the DUE at a distance of ∼2 helical turns), others are known which are frequently but not generally found in bacterial *oriC·*s.

The DnaA-trio motif, (NAN)n, is located in the *oriC (incC*) of *B. subtilis* between the DUE and the conserved DnaA box R1 (DnaA box#6 in the *B. subtilis* convention) (9). DnaA protein binds to dsDNA at DnaA boxes via its C-terminal domain 4 (39), Recently, Pellicari *et al*. (2024) demonstrated that oligomerized DnaA binds to adjacent DnaA-trio motifs via its central domain 3 during strand opening (4). Apart from the replication origins of *E. coli* and *B. subtilis*, DnaA-trio motifs were detected by sequence comparisons in the (putative) chromosomal replication origins of most bacterial genomes of various phyla (**Figure 3**).

The *E. coli* K-12 *oriC* contains a binding site for integration host factor (IHF) with the consensus sequence 5’-WAT**C**ARNNNNTTR (conserved C in bold), located downstream of DnaA box R1 (40, 41). As a ’histone-like’ or nucleoid-associated protein (NAP), IHF strongly bends DNA upon binding and is considered a positive regulator of the initiation of chromosome replication from *oriC* in *E. coli* K-12 (3). IHF-binding sites can be detected in the majority of *oriC·*s from the Gammaproteobacteria, and (almost) invariably with the conserved C at a distance of 21 bp downstream of DnaA box R1 (**Figure 3**).

Three AT-rich 13·mer repeats (L, M, R) with the consensus sequence 5’-GATCTNTTNWWWN, located in the DUE of *E. coli* K-12 *oriC*, were proposed as entry sites for DnaA during initiation of chromosome replication (42). In *E. coli*, 5’-GATC sites are methylated by *dam* methylase (5’-G^m^ATC), and are bound by SeqA protein when hemimethylated shortly after replication, (43). Accordingly, these 13-mer sequences have so far only been found in bacterial genomes containing *dam* and *seqA* genes, the "*dam* clade" of the Gammaproteobacteria (44) (**Figure 3**).

From the collection of 2.5 kb sequence segments of the Halomonadaceae genomes (Suppl. Table 2), I extracted 161 sequences for "*oriC2*" and separately 140 sequences for "*oriC1*" each from pos. 150–249 of the 500 bp long "central" segment containing the conserved R1-type DnaA box at pos. 201–210 to obtain a consensus sequence. For both collections, I did not distinguish here whether a "strong" or "weak" DUE or no DUE was predicted for the individual sequences (Section 3.3).

The consensus sequences for "*oriC1*" and "*oriC2*" are shown as WebLogos in **Figure 3** (45). For comparison with *oriC* sequences with confirmed function, the corresponding sequences from *oriC1* and *oriC2* of *P. aeruginosa* PAO1, *E. coli* K-12 *oriC* and *B. subtilis* 168 *oriC* (*incC*) are included in **Figure 3**. All sequences are aligned with the conserved R1-type DnaA box and the structural elements highlighted. In the DUE sequences of *E. coli* K-12 and *B. subtilis*, the bases in the upper and lower strand are marked (red dots), which were experimentally confirmed by *in vitro* KMnO4 footprinting as single-stranded upon strand-opening (38).

For both "*oriC1*" and "*oriC2*" individually and in direct comparison with each other, the WebLogo con-ensus sequences show that the arrangement of the structural elements and their spacing from each other is highly conserved. The distance between the right flank of the DUE and the DnaA box R1 is approx. 2 helical turns. Between the short DnaA-trio motifs (2–3 repeats) and DnaA box R1 an additional DnaA box in the same rev orientation as R1 but deviating from the consensus sequence is found at a distance of 4 bp from R1. Such "twin R1" DnaA boxes are found in the *B. subtilis oriC* (*incC*) and also in the *oriC·*s of *S. coelicolor* A3(2) (17) and *C. jejuni* 81116 (16) but is here positionally specific for the Halomonadaceae as is not found in *E. coli* K-12 *oriC* or both *oriC·*s of *P. aeruginosa*.

In both the "*oriC1*" and "*oriC2*" WebLogo consensus sequences IHF-binding sites are found at a distance of 21 bp of the conserved C downstream of DnaA box R1 as is common among *oriC·*s of the Gammaproteobacteria (Suppl. Poster 1). Unlike in the "*oriC1*" consensus sequence, a DnaA box is conserved downstream of the IHF-binding site in the "*oriC2*" WebLogo consensus sequence. The other difference between the "*oriC1*" and "*oriC2*" consensus sequences is that a TGT motif directly upstream of the DnaA-trio motifs occurs only once or twice in the former, whereas it is found four times as direct repeat in the latter. The occurrence of a TGT motif, but not as a direct repeat, in *oriC2* of *P. aeruginosa* PAO1 and *E. coli* K-12 *oriC* suggests that TGT repeats are specific for *oriC·*s of the Halomonadaceae. Since a possible function of this motif is not known, I will leave it here with its mention. Not unexpected is the absence of identifiable AT-rich 13·mers in "*oriC1*" and "*oriC2*" consensus sequences, as the Halomonadaceae, like *P. aeruginosa* PAO1, do not belong to the "*dam* clade" of the Gammaproteobacteria.

Without accounting for the distribution of all DnaA boxes in the 500 bp "central" segments in the assessment (Section 3.5), I consider the conservation of key structural elements of a chromosomal origin of replication sufficient to assign the status of predicted *oriC1* and *oriC2* sequences to the preliminary "*oriC1*" and "*oriC2*" sequences, respectively.

I propose a further coarse classification of the (predicted) *oriC·*s based on their predicted DUE·s (Section 3.3). WebLogo consensus sequences obtained separately for *oriC1* sequences with a predicted "strong" DUE (61), "weak" DUE (39) and no DUE (40) reveal the complete conservation of the structural elements, but differences in the GC content of the first 26 bp (DUE region) up to the conserved TGT repeat preceding the DnaA-trio motifs (Suppl. Figure 3.2). The calculated average GC content for pos. 150– 175 of the 500 bp long "central" segment gradually increases from 24 % in the "strong" DUE samples to 33.3 % in the "weak" DUE samples and to 50.3 % in the samples lacking a predicted DUE. It is known that GC-rich dsDNA sequences are less likely to be unwould than AT-rich ones, and in this respect this observation is consistent with the SIDD prediction. However, the SIDD prediction also calculates the "unwindability" depending on the degree of superhelicity of a DNA sequence. A comparably detailed analysis was not meaningful for the *oriC2* sequences, as this set contains too few samples with "weak" (10) or no DUE (1).

I propose the following classification:

1. *oriC1* and *oriC2* sequences, for which a "strong" DUE is predicted, are fully functional replication origins (red label).
2. *oriC1* and *oriC2* sequences, for which a "weak" DUE is predicted, have a slightly reduced functionality as replication origins (pale red label).
3. *oriC1* and *oriC2* sequences, for which no DUE is predicted, have reduced functionality as replication origins (gray label).
4. "*oriC1*" and "*oriC2*" sequences, for which no DUE is predicted and that lack the R1-type DnaA box and, in addition, other structural elements, are not replication origins but intergenic regions (ig), regardless of whether they contain DnaA boxes (white label).

These four labels are assigned to the *oriC2* and *oriC1* sequences of the individual species of the Halomonadaceae in **Figure 5** and listed in Suppl. Table 1.

### 3.5 Distribution of DnaA boxes in the *oriC2* and *oriC1* sequences

I manually examined the "*oriC1*" and "*oriC2*" sequences of the Halomonadaceae for the distribution of DnaA boxes in addition to the R1-type DnaA box (Section 3.3). Intergenic regions were included that could not be classified as *oriC* due to the absence of the DnaA box R1 and a DUE (Section 3.4). The search was for consensus *E. coli* DnaA boxes (5’-TTWTNCACA) (22) containing 1 mismatch, and for relaxed DnaA boxes with up to 4 mismatches, as reasoned in Section 2.2.11.

In Suppl. Table 2, the multiple DnaA boxes detected in the "*oriC2*" and "*oriC1*" sequences in the 2.5 kb sequence context used for the prediction of DUE (Section 3.3) are annotated for the complete intergenic regions, i.e. for the "central" 500 bp long sequence segment and the flanking sequences. A one-to-one comparison of the individual "*oriC1*" and "*oriC2*" sequences is not possible in Suppl. Table 2, but can be realized in the synopsis in Suppl. Figure 4.2, which is limited to the "central" 500 bp long sequence segment (with the DnaA box R1 at pos. 201–210 used for the alignment, see Section 3.3); the positions of the DUE and the DnaA boxes are shown exactly to scale.

Because it is not possible to display the synopsis in Suppl. Figure 4.2 on a single printed page such that species names and the DNA sequences remain legible, the scaled-down synopsis in **Figure 4** does not show these but otherwise all sequences for *oriC1* and *oriC2* separately in the order in which they are displayed in Suppl. Figure 4.2 and in **Figure 5**; the positions of the DUE and the DnaA boxes are shown exactly to scale.

**Figure 4.**
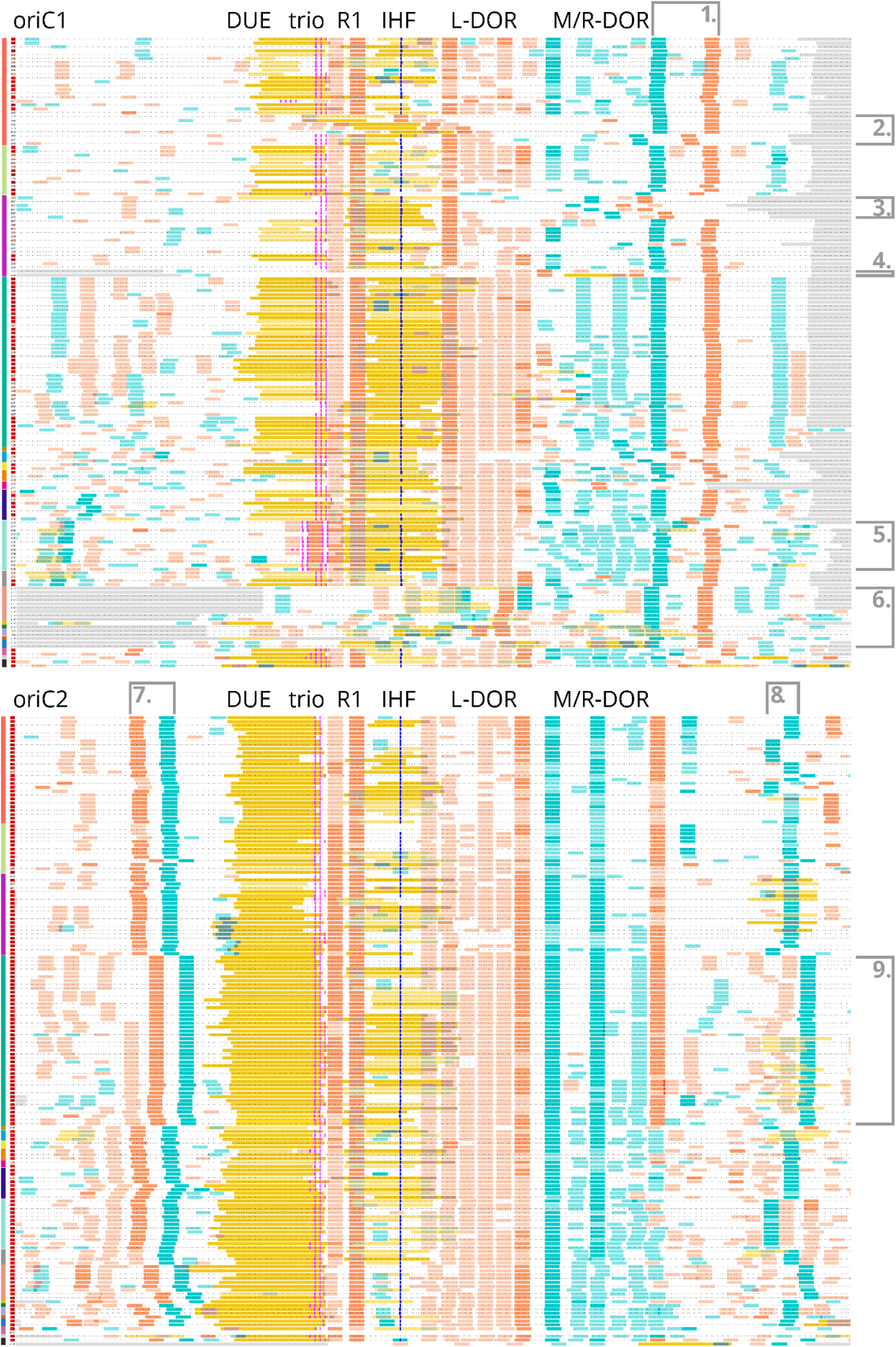
Distribution of DnaA boxes in the *oriC1* (upper panel) and *oriC2* sequences (kower panel) of the Halomonadaceae. Structural features of *oriC*·s are indicated in the header lines: DUE, DNA unwinding element; trio, DnaA-trio motif; R1, DnaA box R1; IHF, IHF-binding site; L-DOR, M/R-DOR, DnaA initiator-oligomerization regions acc. to Hayashi *et al*. (2020) (35). The vertical bar on the left shows the same color code for the clades as in Figure 5: 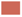 Halomonas (sensu stricto), 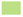 Halomonas_G, 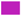 Halomonas_H, 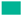 Halomonas_I, 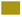 Halomonas_F, 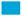 Halomonas_B, 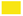 Halomonas_E, 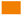 Halomonas_A, 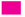 Aidingimonas, 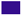 Modicisalibacter, 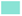 Salinicola, 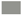 Chromohalobacter, 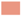 Kushneria, 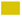 "Phytohalomas", 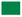 Carnimonas, 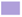 Halotalea, 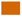 Zymobacter, 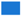 Larsenimonas, 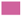 Cobetia, 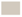 uncertain classification, 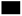 not Halomonadaceae. *Halomonas campaniensis* LS21. All *oriC* sequences were aligned with the DnaA box R1 at pos. 201–210 of the "central" 500 bp segment (Section 3.3). On the left of each line, the *oriC* quality is indicated: 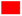 oriC, strong DUE, 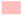 oriC, weak DUE, 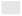 oriC, no DUE, ▭ no oriC structure. Color code for the *oriC* sequences: ATATATAAT SIDD strong (0–3); CTTTTATGA SIDD weak (3–6); GAATATATA SIDD strong / DnaA box overlap; GAAT-TAATA SIDD weak / DnaA box overlap; **TGTGGATAA** DnaA box R1-type; TGTGGATAA DnaA box cons rev; TGTGGATAA DnaA box relaxed rev; TTTCACAAG DnaA box overlap; CTGTCCACA DnaA box cons fwd; TCCGGGGCA DnaA box relaxed fwd; TATTACTAG DnaA-trio; `ATAT`**`C`**`AATA` central C of IHF site; `ATG` orf with start/stop codon. See text for brackets **1.]** – **9.]**.

As shown in **Figure 3**, the structural elements ’DUE–DnaA-trio–DnaA box R1–IHF-binding site’ align well also in **Figure 4** for both the *oriC1* and *oriC2* sequences, which can be distinguished from each other by the different configurations of the L-DOR and M/R-DOR elements. According to Hayashi *et al*. (2020), DnaA initiator-oligomerization regions (DOR·s) downstream of the IHF-binding site in the *E. coli oriC* are arrays of regularly spaced, rev-oriented DnaA boxes (5’-TGTGNAWAA) in L-DOR and with regularly spaced, fwd-oriented DnaA boxes (5’-TTWTNCACA) in M/R-DOR (35). DnaA boxes in DOR·s are mostly "low-affinity", or relaxed, and serve as binding sites for filamentous DnaA oligomers (46). The L-DOR in the *oriC1* sequences of the Halomonadaceae comprise (mostly) five regularly spaced (2–3 bp), rev-oriented, relaxed DnaA boxes while (mostly) six in the *oriC2* L-DOR, the latter with a closer distance to the IHF-binding site. In both *oriC1* and *oriC2*, the rightmost DnaA box of the L-DOR is (almost) invariably at a distance of 90 bp from DnaA box R1.

Unlike in *E. coli oriC,* no M/R-DOR motif can be clearly detected in the *oriC1* sequences of the Halomonadaceae (**Figure 4**, upper panel). The segment of about 64 bp downstream of the L-DOR comprises up to six fwd-oriented, relaxed DnaA boxes, but a consistent, regularly-spaced arrangement is not recognizable. Also unlike in *E. coli oriC,* no M/R-DOR motif can be clearly detected in the *oriC2* sequences of the Halomonadaceae as no consistent, regularly-spaced arrangement is reconizable (**Figure 4**, lower panel). One fwd-oriented DnaA box is fully conserved at a distance of 108±1 bp from DnaA box R1 that is also partially present at this position in the *oriC1* sequences and may represent the left border of an unusual M/R-DOR motif. A second fwd-oriented DnaA box is conserved at a distance of 135±2 bp from DnaA box R1. A third, rev-oriented DnaA box is conserved exclusively in the subfamilies ’Halomonas_sensu stricto’ to ’Halomonas_I’ at a distance of 171±1 bp from DnaA box R1.

The DUE in the *oriC1* sequences is ∼10 bp shorter than in the *oriC2* sequences, and the DUE-upstream sequences are completely different though with a reasonably well conserved DnaA box pattern among the *oriC2* sequences. It is therefore unlikely that one of the origin sequences was the result of a duplication of the other during speciation of the family Halomonadaceae.

Some notable features of the synopsis are described here, as indicated in numbered square brackets in **Figure 4**:

1.] downstream of the M/R-DOR, a pair of DnaA boxes with opposite orientation and a distance of 23±1 bp (∼2 helical turns) is conserved in most *oriC1* sequences and not found in the *oriC2* sequences. Since it is also present in *oriC*·s with reduced functionality and in non-functional *oriC*·s (2.], 3.], 4.], 5.], 6.]), it is more likely that this pair of DnaA boxes plays a role in regulating the downstream *dnaA* promoter than that it has a function for *oriC*.

2.] All eight *oriC1*·s lack a detectable DUE and have "scrambled" L-DOR and M/R-DOR motifs. The *oriC1*·s of *H. urmiana, H. saccharevitans, H. denitrificans, H. aestuarii,* and *H. ventosae* lack, in addition, the ’DnaA-trio–DnaA box R1–IHF-binding site’ motifs and were classified as non-functional (ig) (Section 3.4). These eight species share a twig within the branch of the ’Halomonas_sensu stricto’ subfamily in the phylogenomic tree (**Figure 5**).

**Figure 5.**
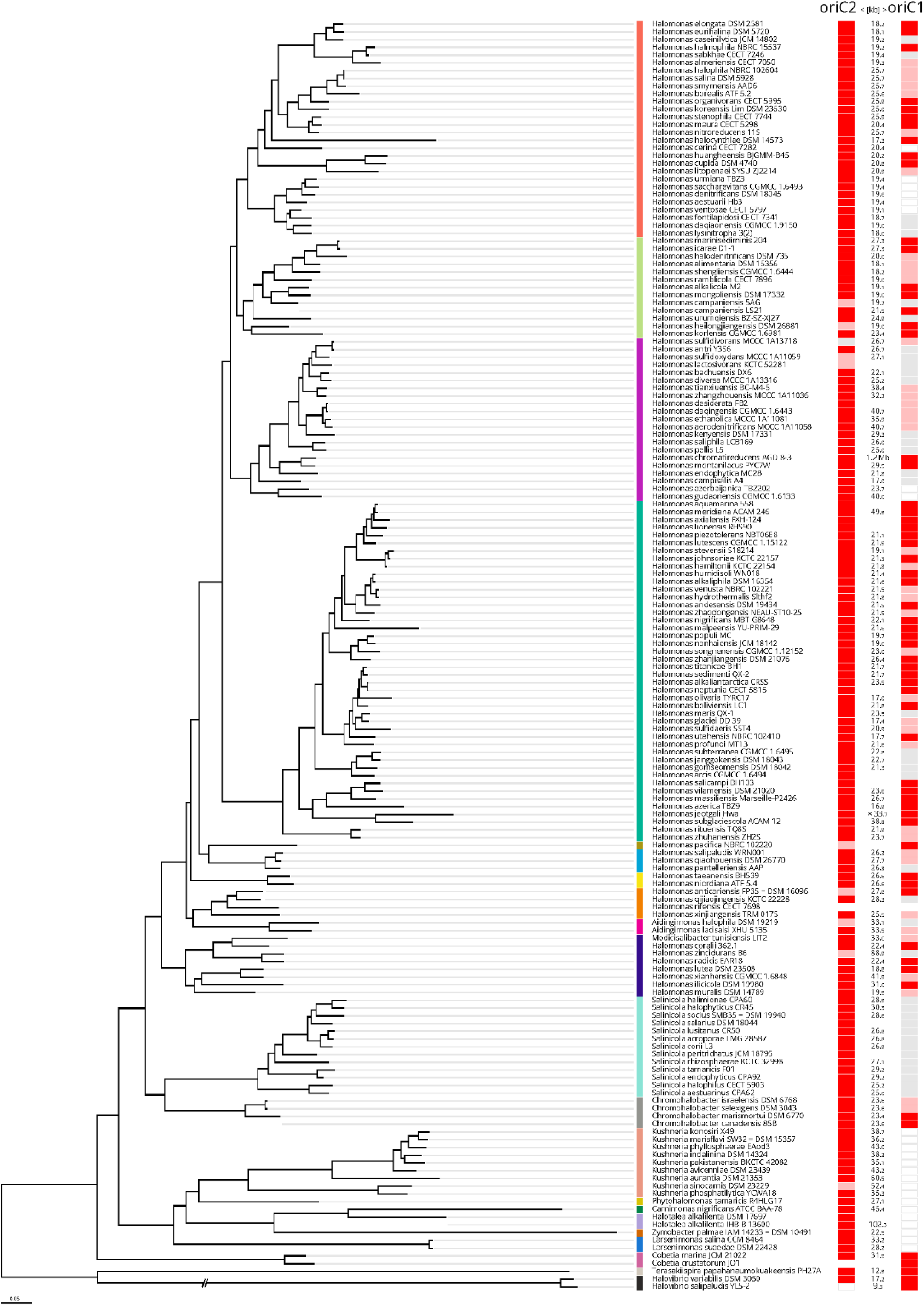
Projection of the *oriC* classifiactions for Halomonadaceae species/strains onto the phylogenomic tree of de la Haba *et al*. (2023), Suppl. Figure 1 (26). Clade expanded maximum-likelihood phylogenomic tree based on the concatenation of the translated sequence of the 189 single-copy genes shared at least by 90% of the members of the family Halomonadaceae under study (“core90” set). Bar, 0.05 changes per nucleotide position. Clade color code: 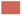 Halomonas (sensu stricto), 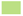 Halomonas_G, 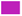 Halomonas_H, 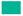 Halomonas_I, 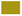 Halomonas_F, 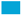 Halomonas_B, 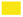 Halomonas_E, 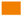 Halomonas_A, 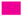 Aidingimonas, 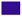 Modicisalibacter, 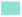 Salinicola, 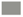 Chromohalobacter, 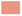 Kushneria, 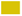 "Phytohalomas", 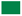 Carnimonas, 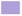 Halotalea, 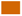 Zymobacter, 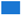 Larsenimonas, 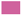 Cobetia, 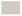 uncertain classification, 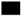 not Halomonadaceae. *Halomonas campaniensis* LS21, *Chromohalobacter canadensis* 85B, and *Halotalea alkalilenta* IHB B 13600 were tentatively placed in the tree. 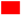 oriC, strong DUE, 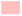 oriC, weak DUE, 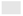 oriC, no DUE, ▭ no oriC structure.

3.] The *oriC1*·s of this group of six species *H. sulfidivorans*, *H. antri*, *H. sulfidoxydans*, *H. lactosivorans*, *H. bachuensis*, and *H*. *diversa* retain the ’DnaA-trio–DnaA box R1–IHF-binding site’ motifs, but, except for *H. sulfidivorans,* lack a detectable DUE. One or more deletion(s) downstream of the IHF-binding site render the L-DOR and M/R-DOR motifs unrecognizable and have reduced the distance to the downstream *dnaA* gene by 30–40 bp. These six species share a twig within the branch of the ’Halomonas_H’ subfamily in the phylogenomic tree (**Figure 5**).

4.] The *oriC1*·s of *H. azerbaijanica* and *H. gudaonensis* have unusually short *rpmH*–*dnaA* intergenic regions that lack a DUE and the ’DnaA-trio–DnaA box R1–IHF-binding site’ signature motifs despite the presence of multiple DnaA boxes. These ig·s are thus not considered *oriC1* sequences. For the arbitrary alignment with other *oriC1* sequences, I used the conserved DnaA box pair in the 3’-part (1.]), which then brings the 5’-flanking gene *rpmH* approximately 80 bp closer. These two species share a twig within the branch of the ’Halomonas_H’ subfamily in the phylogenomic tree (**Figure 5**).

5.] The 13 *oriC1*·s of the genus *Salinicola* lack a detectable DUE but have ’DnaA box R1–IHF-binding site– L-DOR–M/R-DOR’ motifs that align almost perfectly with other *oriC1* sequences. Unusual because not found in any other *oriC1* is the arrangement of two rev-oriented DnaA boxes upstream of the DnaA box R1, both 5’-flanked by DnaA-trio motifs. These 13 species share a branch with the genus *Chromohalobacter* in the phylogenomic tree of the Halomonadaceae (**Figure 5**).

6.] All 15 genomes of the genera *Kushneria*, *Phytohalomonas*, *Carnimonas*, *Halotalea*, *Zymobacter*, and *Larsenimonas* have unusually short *rpmH*–*dnaA* intergenic regions (*sdiA*–*dnaA* in *Zymobacter palmae*) that lack a DUE and the ’DnaA-trio–DnaA box R1–IHF-binding site’ signature motifs despite the presence of multiple DnaA boxes. These ig·s are thus not considered *oriC1* sequences. For the arbitrary alignment with other *oriC1* sequences, I used the conserved DnaA box pair in the 3’-part (1.]), which then brings the 5’-flanking gene *rpmH* (*sdiA*) approximately 100 bp closer. These 15 species share a branch in the phylogenomic tree of the Halomonadaceae (**Figure 5**).

7.] upstream of the DUE, a pair of DnaA boxes with opposite orientation and a distance of 9±1 bp (∼1 helical turn) is conserved in most *oriC2* sequences. Despite its conservation it is unlikely to play a role in *oriC* function as it is not present in any of the *oriC1* sequences.

8.] approximately 50 bp downstream of the M/R-DOR, a consensus fwd DnaA box (5’-TTWTNCACA) is found in almost all *oriC2* sequences, whose position in individual clades is shifted by +1 or –1 helical turns.

9.] Consistently for all members of the ’Halomonas_I’ and ’*Kushneria*’ subfamilies, a shift to the right by one helical turn is found for the DnaA box pair [6.] and of the single box [7.] supporting the validity of the phylogenomic tree (**Figure 5**).

Neither in the *oriC1* nor in the *oriC2* sequences could I detect by eye any other conspicuously conserved motifs that could have hinted at the binding of other replication factors.

To summarize, after determining the DnaA box distribution I did not have to correct the classification of any of the *oriC1*·s and *oriC2*·s based on their (presumed) functionality (Section 3.4). Both *oriC1*·s and *oriC2·*s have the same overall architecture as *E. coli oriC* with the constraint that a M/R-DOR motif cannot be clearly assigned.

### 3.6 *oriC2* and *oriC1* in the phylogenomic tree of the Halomonadaceae

For the Halomonadaceae genomes analyzed here, functional *oriC·*s are located at the genomic positions and in a conserved context that correspond to the positions and context of *oriC2* and *oriC1* in the *P. aeruginosa* PAO1 genome, albeit at a greater distance from each other. As stated in Section 3.4, these *oriC2* and *oriC1* sequences can be roughly classified according to different degrees of (assumed) functionality. This made it interesting to examine in which combinations *oriC2* and *oriC1* are distributed among the individual species in the phylogenomic tree of the Halomonadaceae proposed by de la Haba *et al*. (2023) (26).

A cursory glance at **Figure 5** reveals that fully functional *oriC*-s (red label) predominate in both *oriC2* and *oriC1* across the species tree, with *oriC2* outnumbering *oriC1* by a ratio of 2.5:1 (150:61). This suggests that dual chromosomal origins of replication (*oriC*) were already present in the ancestor(s) of the Halomonadaceae (see the two *Cobetia* sp. at the root of the tree) and that one origin, *oriC1*, was repeatedly and independently lost or "downgraded" during speciation in the different lineages.

This first impression is confirmed by a closer look at the various combinations of *oriC2* and *oriC1* in the 21 species of the subfamily ’Halomonas_H’ (**Figure 5**), orchid vertical line). A fully functional *oriC2* (red label) is found 1. together with a fully functional *oriC1* in *H. montanilacus*, 2. together with an *oriC1* of slightly reduced functionality (pale red label) in *H. desiderata*, 3. together with an *oriC1* lacking the DUE (gray label) in *H. bachuensis*, or 4. together with an *rpmH·dnaA* ig that lacks the structural features of a replication origin in *H. azerbaijanica* (white label). *H. sulfidivorans* is the only example for a species in the entire set that has a combination of an DUE-lacking *oriC2* (gray label) with an *oriC1* of slightly reduced functionality (pale red label). The various combinations of *oriC2* and *oriC1* as in the ’Halomonas_H’ sub-family are likewise found in the other subfamilies of the Halomonadaceae: ’_sensu stricto’, ’_G’, ’_I’, ’_F’, ’_B’, ’_E’, and ’_A’.

The question of whether different combinations of *oriC2* and *oriC1* occur not only in closely related species within a subfamily, or whether this is also found for strains of a species, cannot be answered here due to a lack of sufficient examples. Both *H. alkalilenta* IHB B 13600 and *H. alkalilenta* DSM 17697 (genus Halotalea) have the same *oriC2*/*oriC1* combination. The impression that different *oriC2*/*oriC1* combinations can occur at the strain level as in *H. campaniensis* 5AG and *H. campaniensis* LS21 (’Halomonas_G’ subfamily) is probably misleading. The DnaN protein of *H. campaniensis* LS21 is 86 % identical to DnaN of *H. campaniensis* 5AG, but 100 % identical to DnaN of *H. alkaliphila* DSM 16354 (’Halomonas_I’ subfamily). *H. campaniensis* LS21 may need a taxonomic re-classification, which was beyond the scope of this study.

That the choice for a preferred *oriC* can occur on the genus level can be seen in a branch containing the genera *Salinicola* and *Chromohalobacter*. One sub-branch comprises 13 spezies of the genus *Salinicola* that all have a fully functional *oriC2* (red label) but their *oriC1* lack the DUE (gray label). The other sub-branch comprises four spezies of the genus *Chromohalobacter* that each have two functional *oriC*·s.

In another large branch, which includes the genera *Kushneria* through *Larsenimonas*, all species have a fully functional *oriC2* combined with an *rpmH·dnaA* ig that lacks the structural features of a replication origin. In these species, the choice for a unique *oriC*, here *oriC2*, apparently occurred during the split into separate genera.

A clear correlation between certain *oriC2*/*oriC1* combinations and the genomic distances between the two origins cannot be detected. Only in the genus *Kushneria* do the distances between the fully functional *oriC2* (red label) and the inactive *oriC1* (*rpmH·dnaA* ig) tend to be larger than in the genomes of most other Halomonadaceae (med 38.7 kb, as compared to med 23.6 kb for the complete set). In any case, the 49.9 kb distance between two fully functional origins (red label) in the *Halomonas meridian-a* ACAM 246 genome (’Halomonas_I’ subfamily) remains surprising. As mentioned in Section 3.1, the extremely large distance of >1.2 Mb between the two fully functional origins of *Halomonas chromatireducens* AGD 8-3 is probably due to an genome assembly error.

To summarize, of 163 Halomonadaceae genomes with an annotated *dnaA* gene, 23 have a single origin, *oriC2*, and 138 have a dual origins, *oriC1* and *oriC2*. In 77 of the 138 genomes with dual origins, *oriC1* may have a reduced functionality as compared to the corresponding *oriC2*. In four cases, all from different subfamilies, *oriC1* may have a higher functionality than the corredponding *oriC2*.

## 4 Discussion

The predictions of DnaA-dependent replication origins (*oriC*) in bacterial genomes obtained in the present study were generated using a manual procedure developed through an interplay of prediction and experimental validation (10–19, 27). Of 163 Halomonadaceae genomes with an annotated *dnaA* gene, 23 have a single predicted origin, *oriC2*, and 138 have a dual predicted origins, *oriC1* and *oriC2*.

The most frequently cited database for bacterial replication origins, DoriC12 by Dong et al. (2023) (21), gives identical predicted *oriC* positions in the Halomonadaceae genomes, *oriC2* (<*mnmG[gidA]·oriC2*) and *oriC1* (<*rpmH·oriC1*), based on a scoring of algorithm-driven detection of structural elements as described in the Ori-Finder 2022 method of Dong et al. (2022) (47). Of the 163 Halomonadaceae gemomes studied here, 133 are contained in the DoriC 12.1 database, while 30 are not (Suppl. Table 3). For 129 of these 133 genomes. DoriC 12.1 predicts a single *oriC* at the position of *oriC2*, and a single *oriC* at the position of *oriC1* in the genome of *H. lutescens* CGMCC 1.15122 (for 3 cases, the database was not accessible). By contrast, the manual method clealy predicts a dual-origin configuration for *H. lutescens* CGMCC 1.15122 (#067 in Suppl. Tables 1+3). For the additional 238 Halomonadaceae genomes in the DoriC 12.1 database (3 duplications not counted), 216 predictions are given for a single *oriC* at the position of *oriC2*, and 13 for a single *oriC* at the position of *oriC1*, while for 9 genomes data were not accessible (Suppl. Table 3). Without conducting a genuine benchmarking of the manual method and the algorithm-driven Ori-Finder 2022 method of Dong et al. (2022) (47), I limit myself to noting that the differing results obtained by the two methods are likely due to the fact that OriFinder 2022 does not detect dual *oriC*·s in a chromosome even when both sequences contain all structural elements of a bona fide *oriC*.

Both the predicted 140 *oriC1* (<*rpmH·oriC1*) and the 162 *oriC2* (<*mnmG·oriC2*) sequences in the present study are located 20–30 kb apart and in a conserved genomic context that the Halomonadaceae largely share with the Pseudomonadaceae. Upon alignment, the *oriC1* and *oriC2* sequences show an almost identical arrangement of the structural elements in the "left" part: ccw transcription of the left-flanking gene, the DUE, the DnaA-trio motif, the DnaA box R1, and the IHF binding site (**Figure 3**). By contrast, the *oriC1* and *oriC2* sequences show a highly diverging arrangement of the structural elements in the "right" part: while the L-DOR motif in the *oriC1* sequences contains five rev-oriented DnaA boxes, there are 6 rev-oriented DnaA boxes in the *oriC2* sequences, which are also located closer to the IHF-binding site (**Figure 4**). Overall, but without attempting to quantify, the L-DOR motifs in the *oriC1* and *oriC2* sequences are significantly more similar among each other than to the others. This also applies to the M/R-DOR motifs that cannot be clearly assigned. It can therefore be safely ruled out that the *oriC1* and *oriC2* sequences are duplications at later stages of diversification of the family Halomonadaceae and the dual-origin configuration was in all likelihood inherited from an ancestral lineage.

In 77 of the 138 genomes with dual origins, *oriC1* may have a reduced functionality as compared to the corresponding *oriC2*. In four cases, all from different subfamilies, *oriC1* may have a higher functionality than the corredponding *oriC2*. The distribution of species/strains with single or dual predicted *oriC*·s in the taxogenomic tree of the Halomonadaceae suggests that a demise of *oriC1* or its "downgrading" occurred repeatedly during their diversification (**Figure 5**).

The *oriC* predictions obtained in this study are sufficiently plausible to inform experimental cell-cycle studies that involve a manipulation of the origin(s) of replication or factors interacting with *oriC* for individual species of the Halomonadaceae. Likewise, computational cell cycle simulations for individual species can benefit from taking the single/dual origin predictions presented here into account. Despite their plausibility, however, I emphasise that *oriC* predictions cannot substitute for their experimental validation in order to publish experimental results or to annotate the corresponding genomes. Three accepted methods are available for validation: 1. establishing a selectable "minichromosome" plasmid that carries the predicted *oriC* as the sole replication origin in the target organism, as was pioneered for *E. coli* by von Meyenburg et al. (1979) (48), 2. the in vitro "unwinding assay" using single strand-specific P1 nuclease with cognate DnaA (heterologously expressed, purified) bound to a negatively-supercoiled plasmid (ccc form) carrying the predicted *oriC* (42), and 3. whole genome sequencing (WGS) for growing cells of the target organism and genome copy number analysis by mapping to the reference sequence, as described by Skovgaard et al. (2011) (49), and later applied by Dimude et al. (2018) to study *E. coli* strains with additional ectopic *oriC* copies (36). The latter method is particularly well-suited to investigating dual origins with differing functionality simultaneously. In the Halomonadaceae genomes, the predicted dual origins are >20 kb apart and therefore lie well within the resolution range. However, none of these methods is high-throughput, so the experimental validation of predicted *oriC*·s will have to be limited to particularlily interesting Halomonadaceae species.

If the predicted dual-origin configuration, with possibly differing functionality of *oriC1* and *oriC2*, is experimentally confirmed for individual Halomonadaceae species, I do not expect any significant effects on cell cycle regulation. For one, I could not find any clues in the *oriC1* and *oriC2* sequences, such as additional (putative) recognition sites for DNA-binding factors, which could hint at their differential regulation under certain growth conditions. Also, it is known from studies of *E. coli* strains with additional ectopic *oriC* copies that such strains are viable even when the inter-origin distance is >1 Mb and "origin firing" occurs from both *oriC* copies (36). Strains with dual *oriC* show reduced growth rates, which can be attributed to replication-transcription conflicts with nearby located *rrn* operons and replichore imbalance, which, in one known case, is partly alleviated by tweaking the replication fork speed (36, 50). Since there is no *rrn* operon located in the ∼20–30 kb intervening sequence between *oriC1* and *oriC2* in the genomes of the Halomonadaceae, replication-transcription conflicts and those caused by very large inter-origin distances can be excluded here.

In conclusion, the result of this study, the prediction of a dual-origin (*oriC*) configuration in many of the presently known genomes of the Halomonadaceae is interesting but does not mean that the "Replicon Model" by Jacob et al. (1963) needs to be rewritten (1). It merely adds the footnote to the model, which is always found in biology: the exception to the rule.

## Supporting information

Figure 1_v02.zip

Figure 2_v01.zip

Figure 3_v02.zip

Figure 4_v01.zip

Figure 5_v01.zip

Legends to Figures.zip

Suppl. Figure 3.2_v03.zip

Suppl. Figure 4.2_v01.zip

Suppl. Poster 1.zip

Suppl. Table 1_v02.zip

Suppl. Table 2_v01.zip

Suppl. Table 3_v02.zip

## Abbreviations

BLAST: Basic Local Alignment Search Tool;
GTDB: Genome Taxonomy Database;
RefSeq: NCBI Reference Sequence Database;
DUE: DNA unwinding element;
IHF: Integration host factor (binding site);
ig: intergenic region;
max: maximum;
min: minimum;
med: median;
bp: base pair(s);
kb: kilo base pairs;
Mbp: mega basepairs;
dsDNA: double-stranded DNA;
ssDNA: single-stranded DNA;
cw: clockwise;
ccw: counter-clockwise.

## Acknowledgments

I thank Monika Glinkowska, Torsten Waldminghaus, Theodor Sperlea, Rafał Donczew, Anna Pawlik, Jolanta Zakrzewska-Czerwińska, Kirsten Skarstad and Ole Skovgaard for continuous support in the past, and David Ussery for suggesting, back in 2006, to use Craig Benham’s WebSIDD server/the SIST software for the prediction of DNA-unwinding elements in bacterial replication origins,.

## Conflict of interest

I declare that the research was conducted in the absence of any commercial or financial relationships that could be construed as a potential conflict of interest.

## Supplementary material

Figure 1_v02.pdf (zip)

Figure 2_v01.pdf (zip)

Figure 3_v02.pdf (zip)

Figure 4_v01.pdf (zip)

Figure 5_v01.pdf (zip)

Legends to Figures.pdf (zip)

Suppl. Figure 3.2_v03.pdf (zip)

Suppl. Figure 4.2_v01.pdf (zip)

Suppl. Poster 1.pdf (zip)

Suppl. Table 1_v02.pdf (zip)

Suppl. Table 2_v02.pdf (zip)

Suppl. Table 3_v02.pdf (zip)

## Notes

### Competing Interest Statement

The authors have declared no competing interest.

