## Supplementary figures and images for "Dual chromosomal origins of replication (*oriC*) in the genomes of the Halomonadaceae – a prediction study"

### Figure 1_v02.pdf

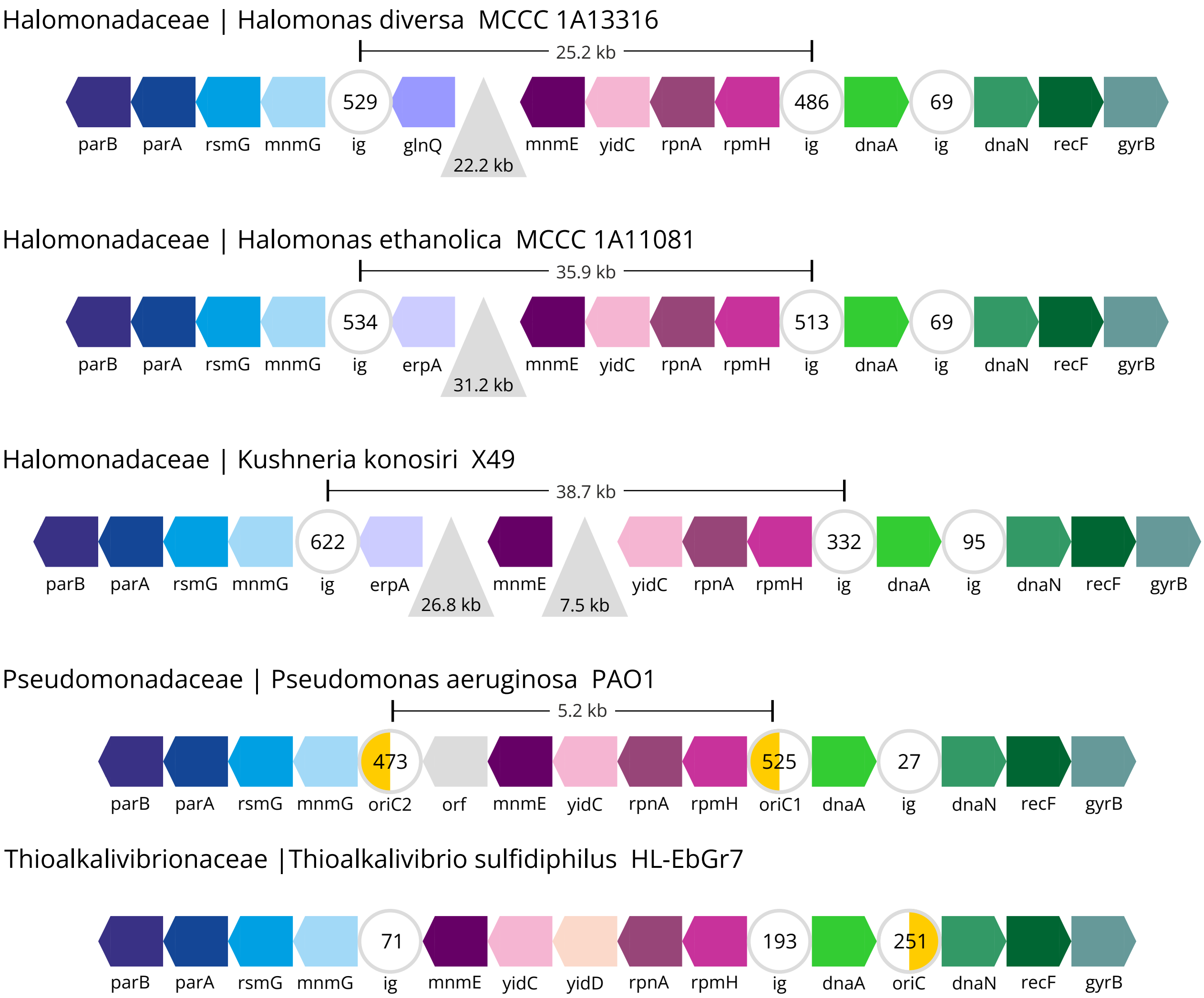

### Figure 2_v01.pdf

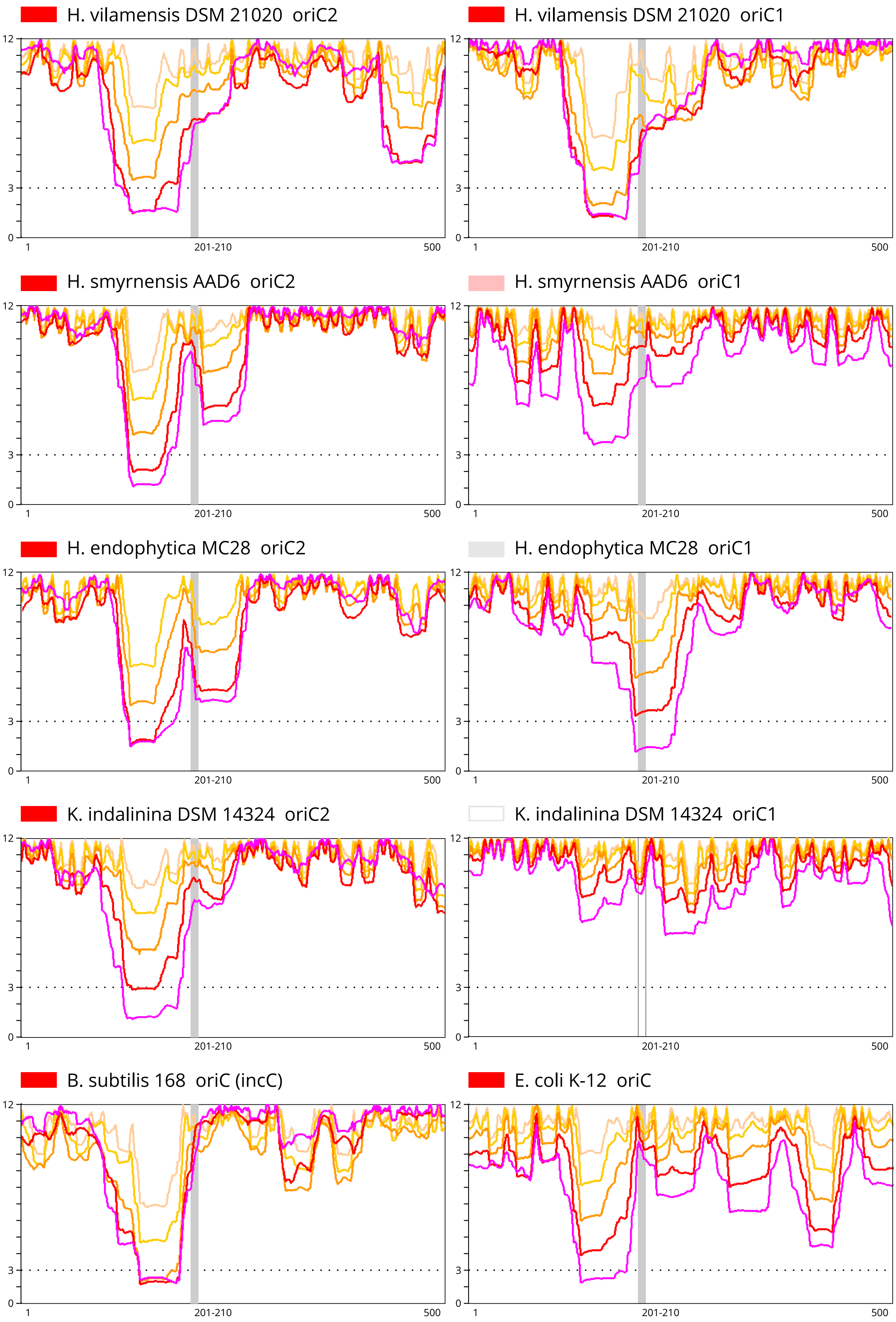

### Figure 3_v02.pdf

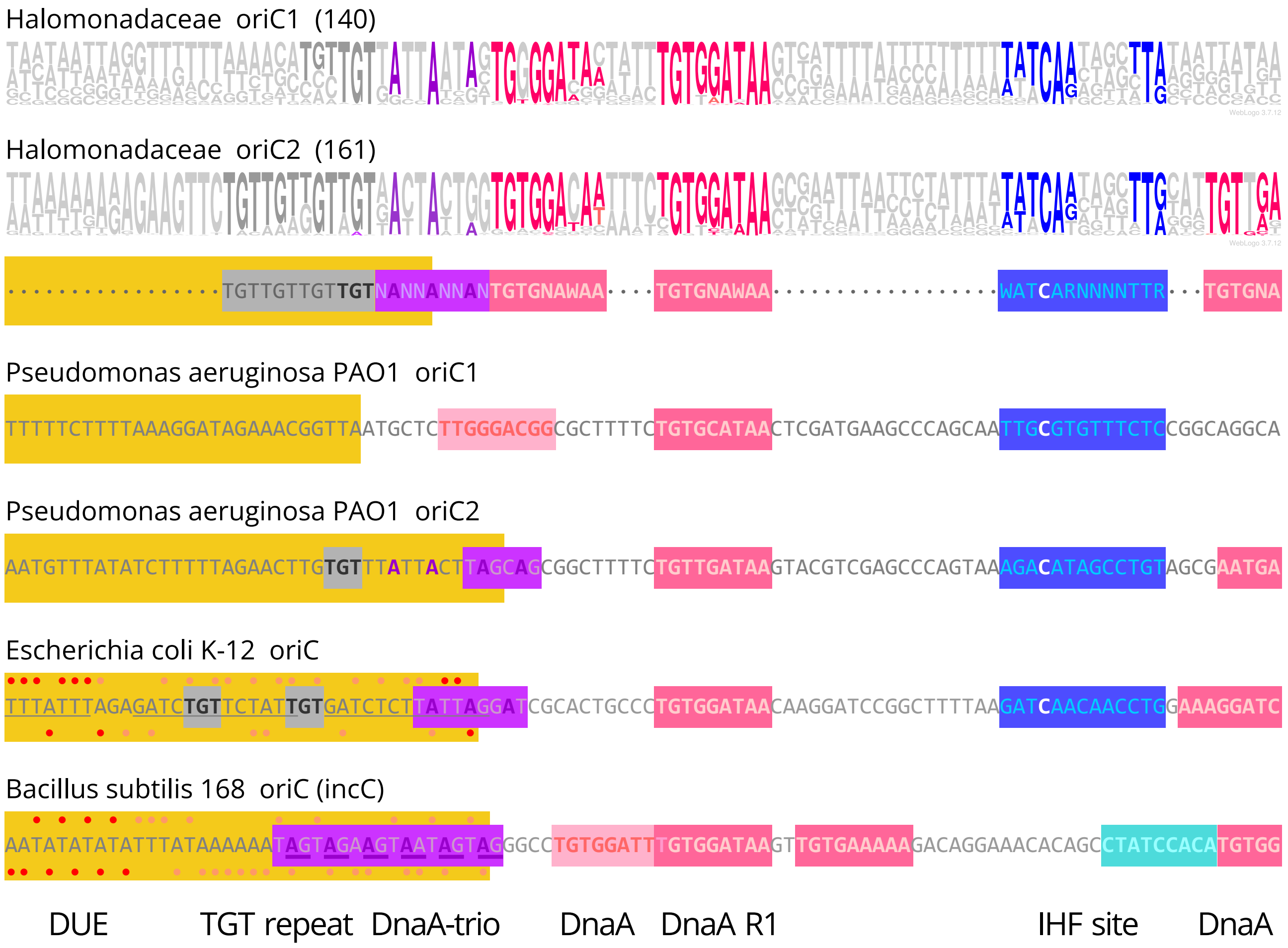

### Figure 5_v01.pdf

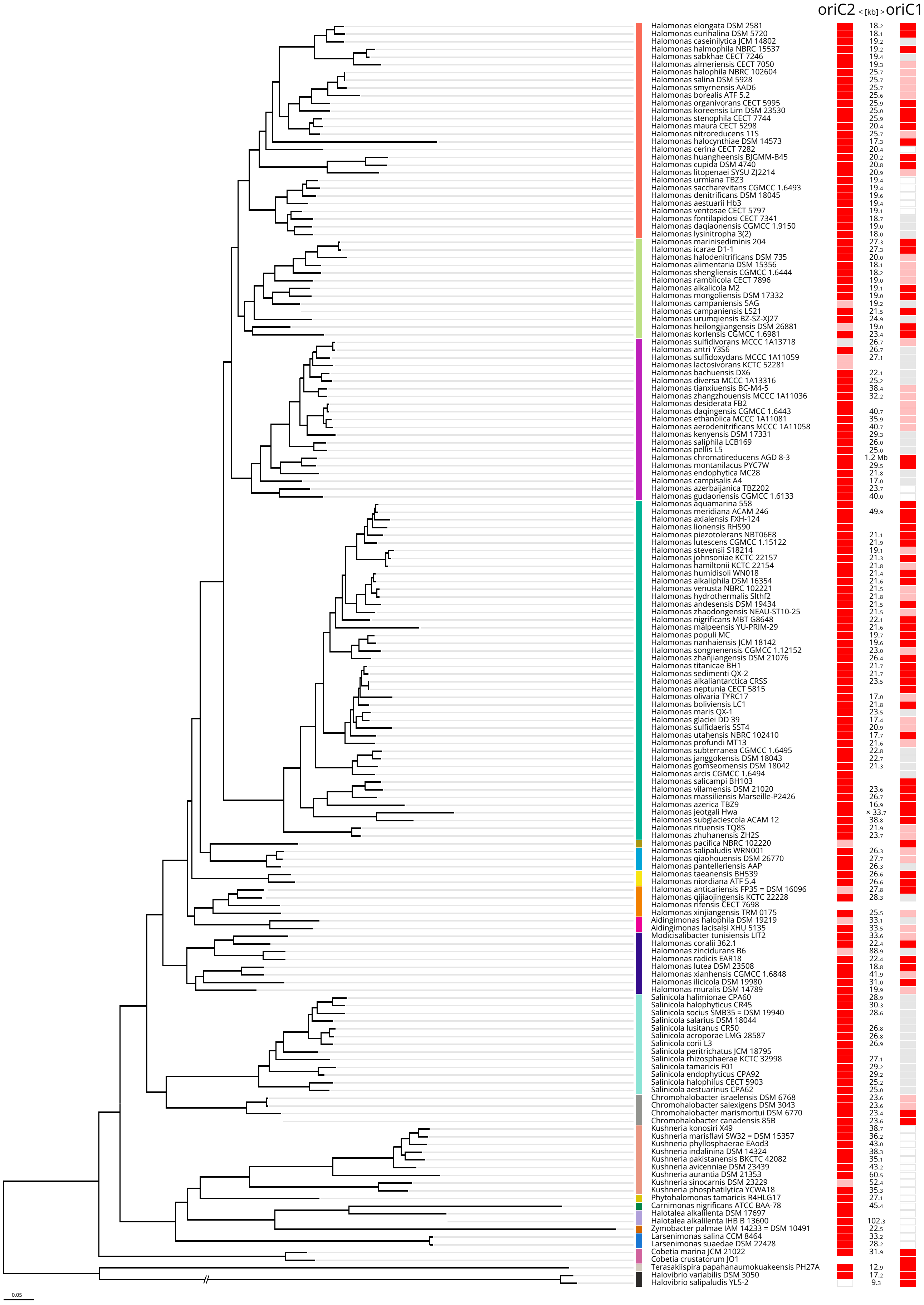

### Suppl. Figure 3.2_v03.pdf

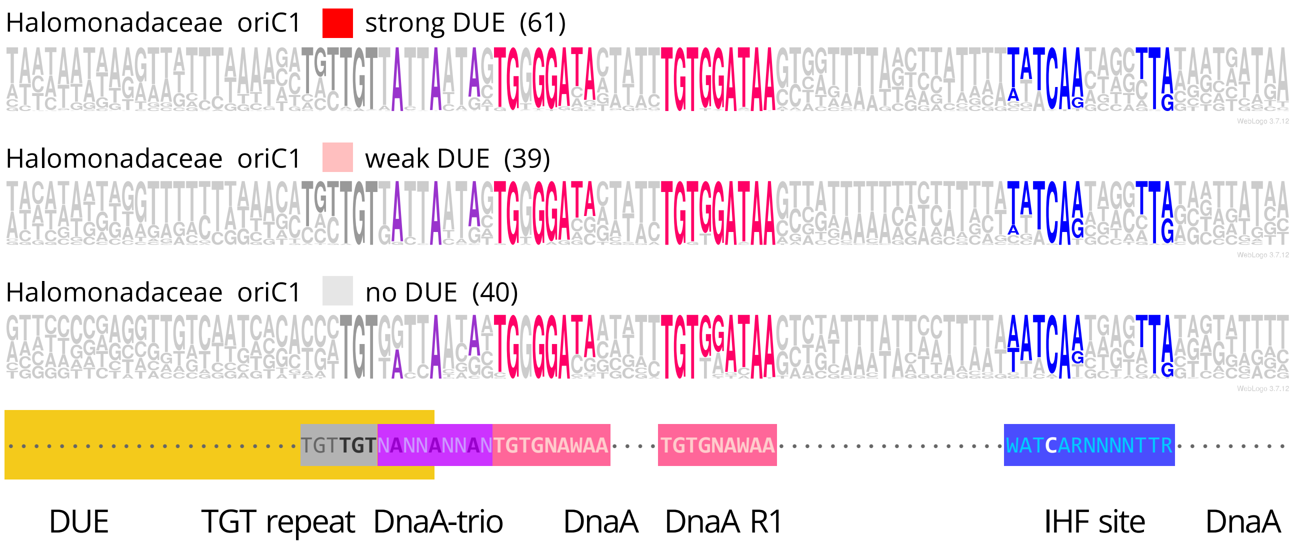


Legend as for Figure 3

Suppl. Figure 3.2

### Suppl. Figure 4.2_v01.pdf

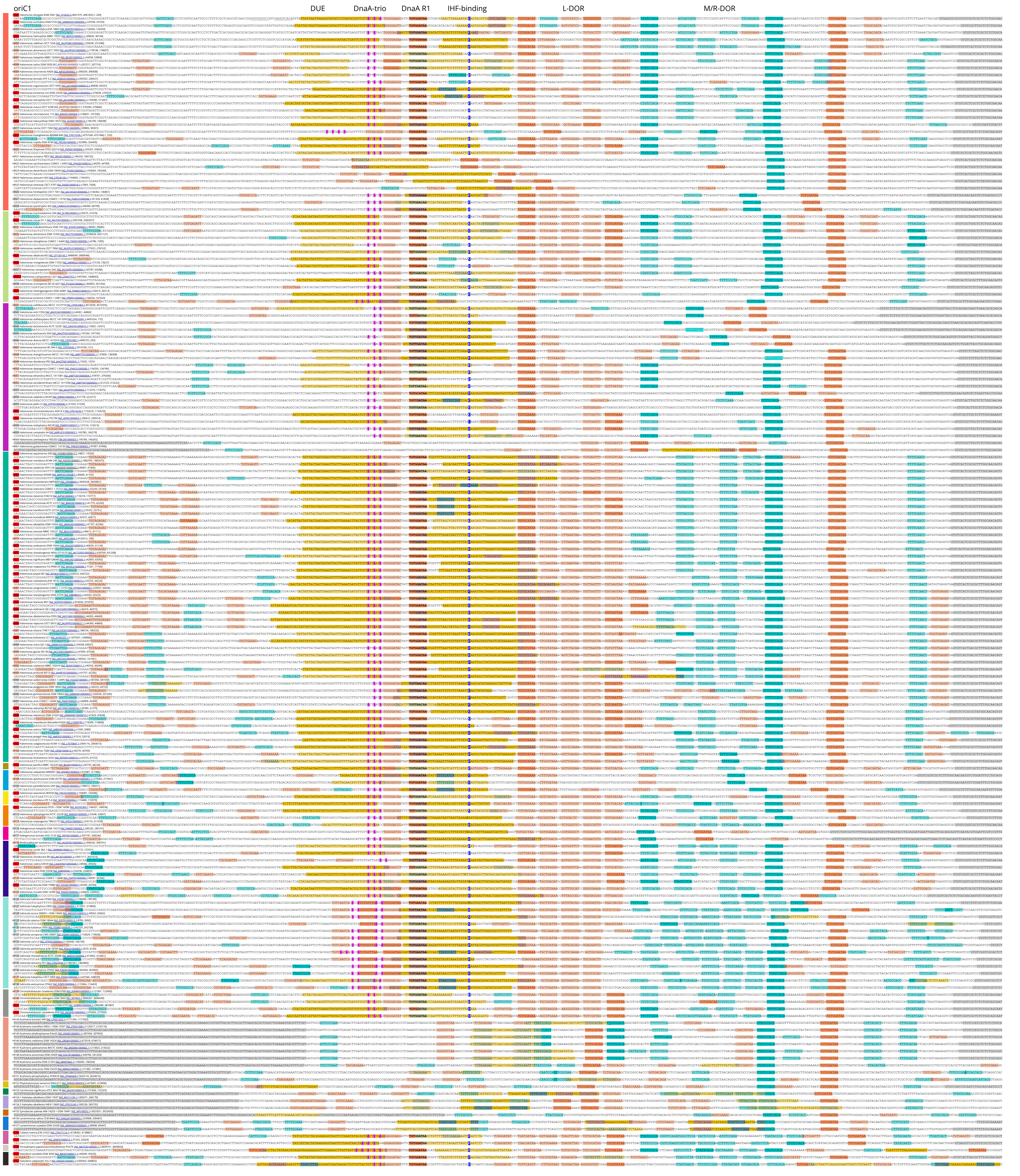


Suppl. Figure 4.2

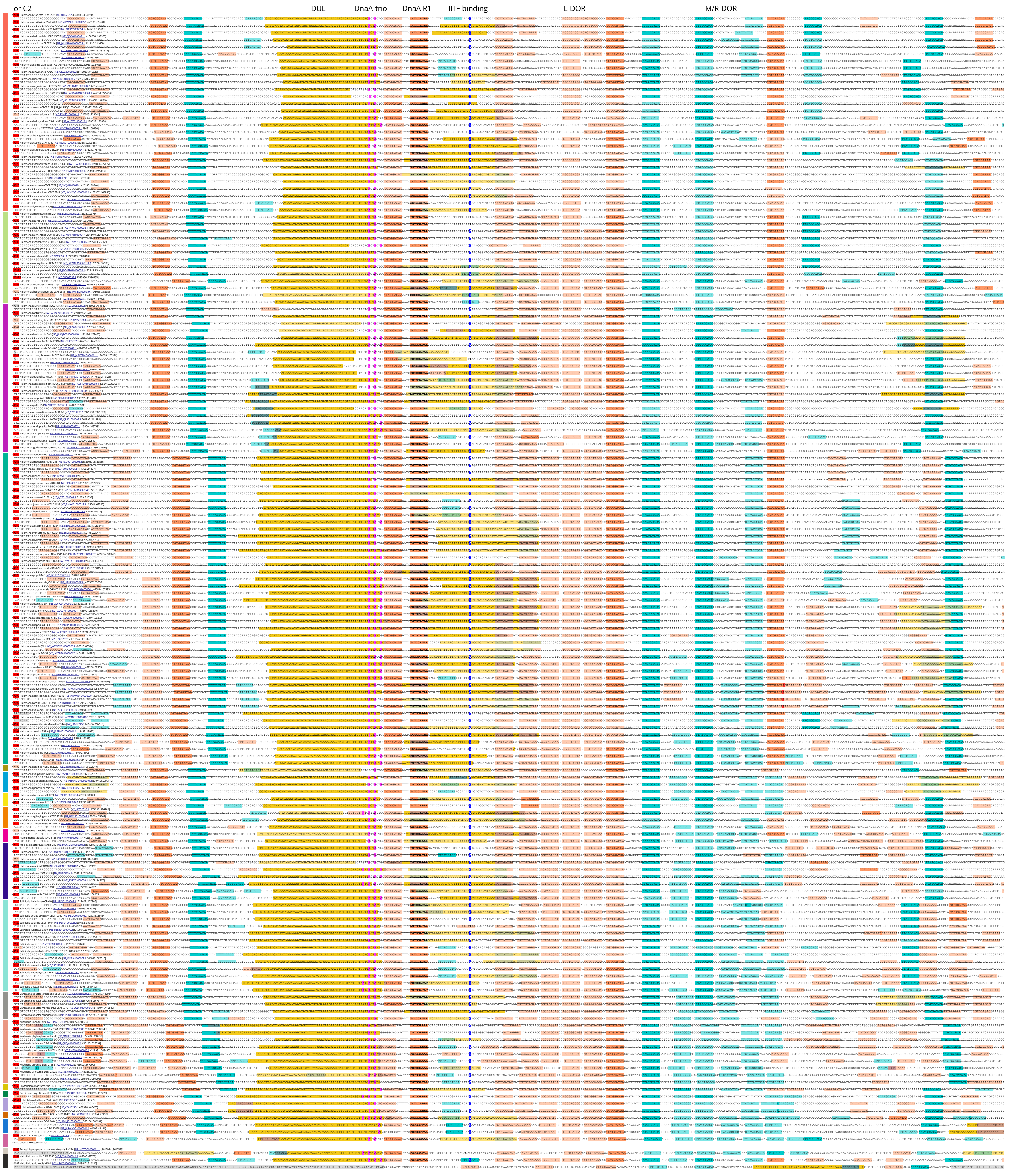


Suppl. Figure 4.2
