## Supplementary material for "Dual chromosomal origins of replication (*oriC*) in the genomes of the Halomonadaceae – a prediction study": Suppl. Poster 1.zip: Suppl. Poster 1.pdf

### A comprehensive set of oriC predictions for the Gammaproteobacteria

Christoph Weigel   Institute of Biotechnology, Faculty III, Technische Universität Berlin (TUB),
Straße des 17. Juni 135, 10623 Berlin, Germany,

**Summary** Comparison of 665 replication origins (oriC) predicted for 593 complete genomes of the Gammaproteobacteria suggests that the present model for the oriC structure of E. coli (Rozgaja et al. (2011), Sakiyama et al. (2017)) can be generalized for this phylum with respect to arrangement of five structural elements: 1. counterclockwise transcription of the left-flanking gene, 2. presence of a ~60 bp long DNA unwinding element (DUE) encompassing the DnaA trio motif on its 3' side (Richardson et al. (2016)), 3. distance of 1–2 helical turns between the DnaA trio motif and the R1-type DnaA box, 4. an IHF-binding site two helical turns downstream of the R1 DnaA box, 5. a ~150 bp long bipartite DnaA oligomerization region (DOR) downstream of the IHF-binding site. Numbers and arrangements of the (mostly) degenerate DnaA boxes and their preferential distances of 2±1 bp in DORL (3–7 rev boxes) and in DORR (4–9 fwd boxes) are conserved in most oriCs on the taxonomic family level. With exceptions, the chromosomal location of oriC is conserved on the taxonomic order level. Dual oriCs at distances of <20 kb from each other as in P. aeruginosa are predicted for 88 of 92 Pseudomonadaceae and 12 of 36 Oceanospirillales. The consequences of dual oriCs for cell cycle regulation remain to be explored. The variability observed for the "DnaA boxR1 region" in the predicted oriCs with respect to the distance of the DnaA trio motif to DnaA box R1 suggests that not one single solution exists for the unwinding reaction, and that this oriC subregion holds the key for its understanding.

#### A oriC location vs. phylogeny

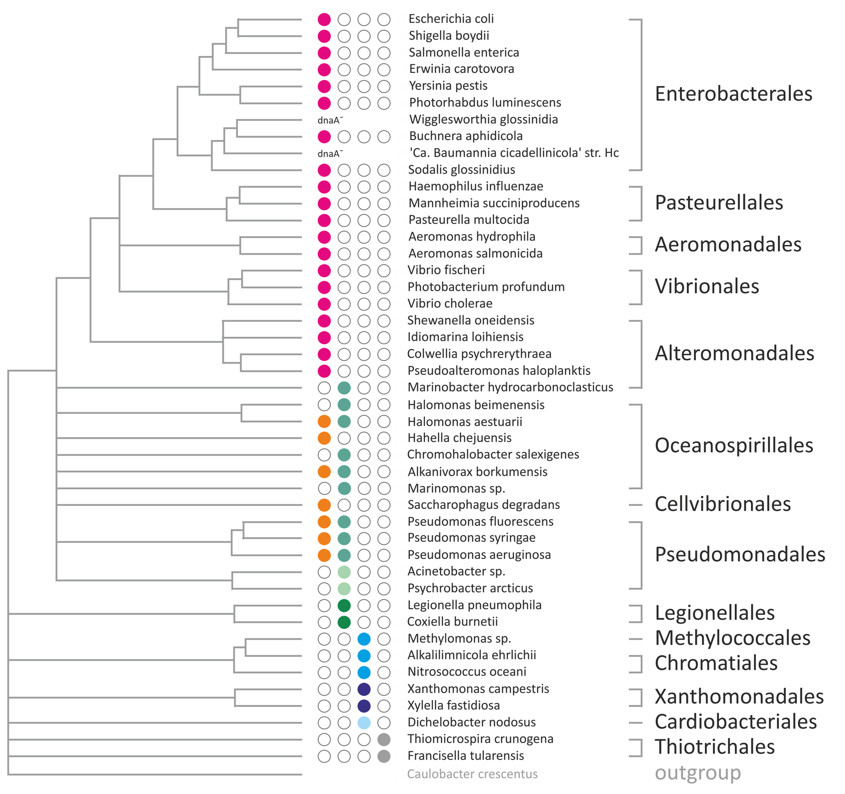


#### B oriC structure, schematical

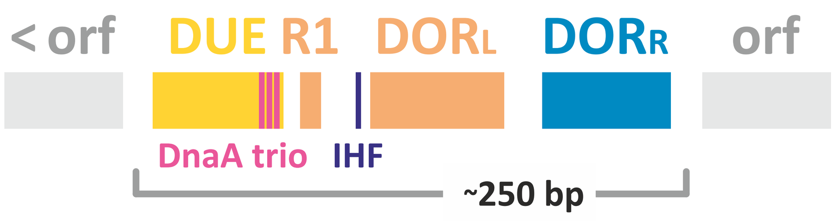


#### C oriC-flanking genes

- < mnmG oriC [< mioC]
- < mnmG oriC [< mnmE]
- < rpmH oriC dnaA >
- < rpmH oriC dnaA >
- < rpmH oriC dnaA >
- < dnaN oriC < dnaA
- < dnaN oriC < dnaA
- < xxx oriC xxx

**Legend A** (above) Maximum-likelihood tree for 45 Gammaproteobacteria based on concatenated sequences of 36 proteins; modified from Gao et al. (2009). Predicted and known (E. coli, P. aeruginosa) oriC locations are indicated by colored circles; color code see panel C. **B** Schematic representation of the generalized oriC structure in the Gammaproteobacteria. Structural motifs are shown in the same colors as in the alignment (right side of poster). DUE, DNA unwinding element (orange); R1, DnaA box R1 (E. coli oriC, red) used for the alignment; DORL (DnaA boxes rev orientation, red), DORR (DnaA boxes fwd orientation, blue), left and right part, respectively, of the DOR (DnaA oligomerization region) acc. to Sakiyama et al. (2017); DnaA trio, motif acc. to Richardson et al. (2016) (pink); IHF, IHF-binding site in oriC (blue). **C** Genes flanking oriCs in the Gammaproteobacteria; the direction of transcription is indicated by < or >. Different color hues reflect different numbers/arrangements of DnaA boxes in the respective DORs. Flanking genes found only in a subset of the sequences are shown in square brackets. **D** (below) OriC predictions for selected genomes of the Actino-, Epsilonproteo-, CPR-, and Cyanobacteria were obtained with the same method as used for the Gammaproteobacteria. Structural motifs are shown in the same colors as in the alignment (right side of poster). A variant of the DnaA trio motif (adjacent 5'-TAG repeats) prevalent in the oriCs of Actino-, Epsilonproteo-, CPR-, and Cyanobacteria is shown in green.

#### D DnaA box R1 regions in various oriCs

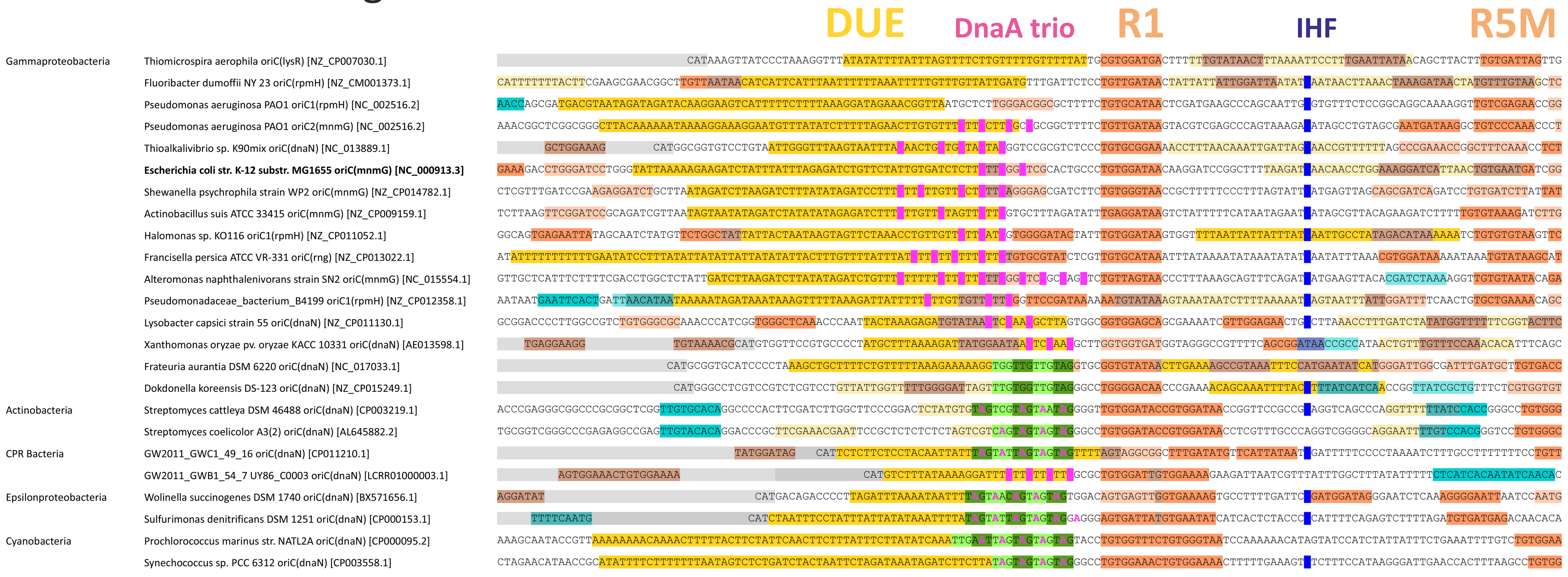


**Method** OriC predictions for complete genomes of the Gammaproteobacteria [ncbi.nlm.nih.gov/genome/] were performed by a stepwise procedure: 1. detection of a dnaA gene (tBLASTn), 2. detection of a minimum in the cumulative GC-skew (GenSkew [http://genskew.csb.univie.ac.at/]), 3. detection of DNA-unwinding element(s) (DUEs) in intergenic regions using a local SIDD server (Benham + Bi (2004)), 4. manual assignment of DnaA boxes (5'-TTWTNCACA, allowing for 3 mismatches) to intergenic regions (Schaper + Messer (1995)) and the flanking orfs, and 5. synteny analysis of the presumed oriC region. More degenerate DnaA boxes as described by Rozgaja et al. (2011) and Sakiyama et al. (2017) were included in step 4 for the intergenic regions (pale red).

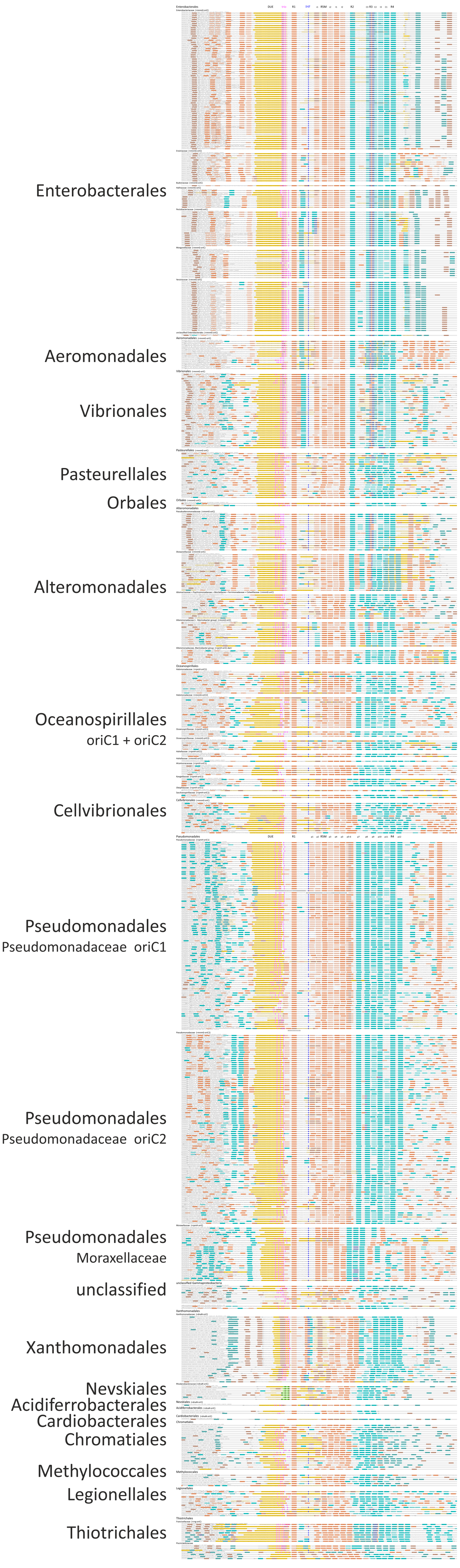
