## Supplementary material for "Dual chromosomal origins of replication (*oriC*) in the genomes of the Halomonadaceae – a prediction study": Suppl. Table 1_v02.zip: Suppl. Table 1_v02.pdf

| # | Species / Strain | RefSeq / Nucleotide | 5' gene | ig | oriC2 | 3' gene | oriC2 Δ oriC1 [kb] | 5' gene | ig | oriC1 | 3' gene | DnaA | ig dnaA·dnaN [bp] | GC$_{min}$ ←oriC2 [kb] | GC$_{min}$ ←oriC1 [kb] |
|---|---|---|---|---|---|---|---|---|---|---|---|---|---|---|---|
| | Halomonas (sensu stricto) | | | | | | | | | | | | | | |
| 001 | Halomonas elongata DSM 2581 | NC_014532.2 | < mnmG | 535 | | < glnQ | 18.2 | < rpmH | 504 | | dnaA > | WP_013330759.1 | 70 | 7.0 | 25.2 |
| 002 | Halomonas eurihalina DSM 5720 | NZ_JARWAH010000005.1 | < mnmG | 536 | | < glnQ | 18.1 | < rpmH | 659 | | dnaA > | WP_149322984.1 | 75 | 2.2 | 20.3 |
| 003 | Halomonas caseinilytica JCM 14802 | NZ_BDEP01000014.1 | < mnmG | 536 | | < glnQ | 19.2 | < rpmH | 514 | | dnaA > | WP_064699906.1 | 74 | 1.2 | 20.3 |
| 004 | Halomonas halmophila NBRC 15537 | NZ_BJOC01000021.1 | < mnmG | 582 | | < glnQ | 19.2 | < rpmH | 515 | | dnaA > | WP_141319605.1 | 214 | 1.7 | 20.9 |
| 005 | Halomonas sabkhae CECT 7246 | NZ_JAUFPM010000008.1 | < mnmG | 583 | | < glnQ | 19.4 | < rpmH | 505 | | dnaA > | WP_290271253.1 | 213 | 3.4 | 22.9 |
| 006 | Halomonas almeriensis CECT 7050 | NZ_JAUFPO010000005.1 | < mnmG | 537 | | < glnQ | 19.3 | < rpmH | 597 | | dnaA > | WP_264826042.1 | 65 | 1.3 | 20.6 |
| 007 | Halomonas halophila NBRC 102604 | NZ_BJUS01000022.1 | < mnmG | 537 | | < glnQ | 25.7 | < rpmH | 596 | | dnaA > | WP_046078887.1 | 64 | 8.8 | 34.6 |
| 008 | Halomonas salina DSM 5928 | NZ_JASFAS010000003.1 | < mnmG | 537 | | < glnQ | 25.7 | < rpmH | 598 | | dnaA > | WP_046078887.1 | 64 | 8.6 | 34.3 |
| 009 | Halomonas smyrnensis AAD6 | NZ_AJKS02000002.1 | < mnmG | 537 | | < glnQ | 25.7 | < rpmH | 515 | | dnaA > | WP_026068327.1 | 70 | 8.6 | 34.2 |
| 010 | Halomonas borealis ATF 5.2 | NZ_SDMO01000002.1 | < mnmG | 525 | | < glnQ | 25.6 | < rpmH | 515 | | dnaA > | WP_136247198.1 | 70 | 4.2 | 29.8 |
| 011 | Halomonas organivorans CECT 5995 | NZ_JACHXM010000012.1 | < mnmG | 529 | | < glnQ | 25.9 | < rpmH | 516 | | dnaA > | WP_183388125.1 | 71 | 8.9 | 34.8 |
| 012 | Halomonas koreensis Lim DSM 23530 | NZ_JARWAK010000004.1 | < mnmG | 527 | | < glnQ | 25.0 | < rpmH | 557 | | dnaA > | WP_309652160.1 | 99 | 22.6 | 47.6 |
| 013 | Halomonas stenophila CECT 7744 | NZ_JACHXR010000003.1 | < mnmG | 537 | | < glnQ | 25.9 | < rpmH | 515 | | dnaA > | WP_183383095.1 | 70 | 8.5 | 34.4 |
| 014 | Halomonas maura CECT 5298 | NZ_JAUFPQ010000013.1 | < mnmG | 536 | | < glnQ | 20.4 | < rpmH | 514 | | dnaA > | WP_290310641.1 | 70 | 10.2 | 30.6 |
| 015 | Halomonas nitroreducens 11S | NZ_RXNS01000004.1 | < mnmG | 536 | | < glnQ | 25.7 | < rpmH | 515 | | dnaA > | WP_126482028.1 | 70 | 25.5 | 51.3 |
| 016 | Halomonas halocynthiae DSM 14573 | NZ_AUDZ01000013.1 | < mnmG | 542 | | < glnQ | 17.3 | < rpmH | 521 | | dnaA > | WP_027965680.1 | 82 | 1.8 | 19.1 |
| 017 | Halomonas cerina CECT 7282 | NZ_JACHXP010000005.1 | < mnmG | 531 | | < glnQ | 20.4 | < rpmH | 505 | | dnaA > | WP_183324828.1 | 72 | 9.4 | 29.8 |
| 018 | Halomonas huangheensis BJGMM-B45 | NZ_CP013106.1 | < mnmG | 643 | | < glnQ | 20.2 | < rpmH | 616 | | dnaA > | WP_021819523.1 | 74 | 20.2 | 0.0 |
| 019 | Halomonas cupida DSM 4740 | NZ_FRCA01000005.1 | < mnmG | 546 | | < glnQ | 20.8 | < rpmH | 632 | | dnaA > | WP_073435379.1 | 123 | 1.6 | 22.4 |
| 020 | Halomonas litopenaei SYSU ZJ2214 | NZ_PXNS01000004.1 | < mnmG | 540 | | < glnQ | 20.9 | < rpmH | 601 | | dnaA > | WP_045995088.1 | 60 | 8.5 | 29.4 |
| 021 | Halomonas urmiana TBZ3 | NZ_VBUI01000001.1 | < mnmG | 565 | | < glnQ | 19.4 | < rpmH | 563 | | dnaA > | WP_138178790.1 | 61 | 25.9 | 45.3 |
| 022 | Halomonas saccharevitans CGMCC 1.6493 | NZ_FPAQ01000016.1 | < mnmG | 576 | | < glnQ | 19.4 | < rpmH | 573 | | dnaA > | WP_089849293.1 | 61 | 8.8 | 28.2 |
| 023 | Halomonas denitrificans DSM 18045 | NZ_PYVX01000003.1 | < mnmG | 553 | | < glnQ | 19.6 | < rpmH | 553 | | dnaA > | WP_108446183.1 | 61 | 27.7 | 47.2 |
| 024 | Halomonas aestuarii Hb3 | NZ_CP018139.1 | < mnmG | 532 | | < glnQ | 19.4 | < rpmH | 530 | | dnaA > | WP_071943402.1 | 61 | 1.9 | 21.4 |
| 025 | Halomonas ventosae CECT 5797 | NZ_SNZJ01000018.1 | < mnmG | 531 | | < glnQ | 19.1 | < rpmH | 510 | | dnaA > | WP_106231148.1 | 61 | 8.6 | 27.7 |
| 026 | Halomonas fontilapidosi CECT 7341 | NZ_JACHXQ010000006.1 | < mnmG | 535 | | < glnQ | 18.7 | < rpmH | 508 | | dnaA > | WP_183314339.1 | 61 | 8.7 | 27.4 |
| 027 | Halomonas daqiaonensis CGMCC 1.9150 | NZ_FOBC01000008.1 | < mnmG | 533 | | < glnQ | 19.0 | < rpmH | 507 | | dnaA > | WP_089712429.1 | 61 | 8.4 | 27.6 |
| 028 | Halomonas lysinitropha 3(2) | NZ_CABVOU010000015.1 | < mnmG | 530 | | < glnQ | 18.0 | < rpmH | 508 | | dnaA > | WP_151442000.1 | 61 | 8.6 | 26.6 |
| | Halomonas_G | | | | | | | | | | | | | | |
| 029 | Halomonas marinisediminis 204 | NZ_SLTR01000012.1 | < mnmG | 569 | | < glnQ | 27.3 | < rpmH | 578 | | dnaA > | WP_132043426.1 | 289 | 8.9 | 36.2 |
| 030 | Halomonas icarae D1-1 | NZ_WUTS01000001.1 | < mnmG | 569 | | < glnQ | 27.3 | < rpmH | 578 | | dnaA > | WP_161423690.1 | 107 | 8.3 | 35.6 |
| 031 | Halomonas halodenitrificans DSM 735 | NZ_JHVH01000002.1 | < mnmG | 554 | | < glnQ | 20.0 | < rpmH | 568 | | dnaA > | WP_027960518.1 | 106 | 9.2 | 29.2 |
| 032 | Halomonas alimentaria DSM 15356 | NZ_WUTT01000001.1 | < mnmG | 543 | | < glnQ | 18.1 | < rpmH | 569 | | dnaA > | WP_161431934.1 | 106 | 12.7 | 30.9 |
| 033 | Halomonas shengliensis CGMCC 1.6444 | NZ_FNIV01000006.1 | < mnmG | 532 | | < glnQ | 18.2 | < rpmH | 580 | | dnaA > | WP_089678848.1 | 103 | 26.5 | 44.7 |
| 034 | Halomonas ramblicola CECT 7896 | NZ_JAUFPL010000032.1 | < mnmG | 535 | | < glnQ | 19.0 | < rpmH | 521 | | dnaA > | WP_290233281.1 | 111 | 9.4 | 28.4 |
| 035 | Halomonas alkalicola M2 | NZ_CP130143.1 | < mnmG | 530 | | < glnQ | 19.1 | < rpmH | 545 | | dnaA > | WP_302140028.1 | 181 | 19.6 | 0.5 |
| 036 | Halomonas mongoliensis DSM 17332 | NZ_JARWAL010000011.1 | < mnmG | 532 | | < glnQ | 19.0 | < rpmH | 556 | | dnaA > | WP_309637217.1 | 122 | 7.7 | 26.6 |
| 037.1 | Halomonas campaniensis 5AG | NZ_JACHZF010000004.1 | < mnmG | 528 | | < glnQ | 19.2 | < rpmH | 520 | | dnaA > | WP_183329889.1 | 116 | 8.6 | 27.8 |
| 037.2 | Halomonas campaniensis LS21 | NZ_CP007757.1 | < mnmG | 565 | | < glnQ | 21.5 | < rpmH | 534 | | dnaA > | WP_038479938.1 | 55 | 5.1 | 26.7 |
| 038 | Halomonas urumqiensis BZ-SZ-XJ27 | NZ_PYUD01000002.1 | < mnmG | 574 | | < glnQ | 24.9 | < rpmH | 516 | | dnaA > | WP_102587617.1 | 115 | 3.0 | 27.9 |
| 039 | Halomonas heilongjiangensis DSM 26881 | NZ_PNRE01000019.1 | < mnmG | 563 | | < glnQ | 19.0 | < rpmH | 519 | | dnaA > | WP_102626523.1 | 106 | 8.5 | 27.6 |
| 040 | Halomonas korlensis CGMCC 1.6981 | NZ_FPBP01000003.1 | < mnmG | 546 | | < glnQ | 23.4 | < rpmH | 565 | | dnaA > | WP_089793768.1 | 74 | 6.7 | 30.1 |
| | Halomonas_H | | | | | | | | | | | | | | |
| 041 | Halomonas sulfidivorans MCCC 1A13718 | NZ_CP053383.1 | < mnmG | 536 | | < glnQ | 26.7 | < rpmH | 472 | | dnaA > | WP_209475102.1 | 63 | 10.7 | 37.5 |
| 042 | Halomonas antri Y3S6 | NZ_JAHYCA010000007.1 | < mnmG | 534 | | < glnQ | 26.7 | < rpmH | 473 | | dnaA > | WP_219793242.1 | 63 | | |
| 043 | Halomonas sulfidoxydans MCCC 1A11059 | NZ_CP053381.1 | < mnmG | 538 | | < glnQ | 27.1 | < rpmH | 462 | | dnaA > | WP_197449037.1 | 64 | 0.0 | 27.1 |
| 044 | Halomonas lactosivorans KCTC 52281 | NZ_QHGY01000015.1 rpmH<br>NZ_QHGY01000013.1 mnmG | < mnmG | 529 | | < glnQ | | < rpmH | 473 | | dnaA > | WP_111415265.1 | | | |
| 045 | Halomonas bachuensis DX6 | NZ_JAAQTO010000018.1 | < mnmG | 530 | | < glnQ | 22.1 | < rpmH | 485 | | dnaA > | WP_167112993.1 | 68 | 8.8 | 33.9 |
| 046 | Halomonas diversa MCCC 1A13316 | NZ_CP053382.1 | < mnmG | 529 | | < glnQ | 25.2 | < rpmH | 486 | | dnaA > | WP_197567058.1 | 69 | 11.7 | 36.8 |
| 047 | Halomonas tianxiuensis BC-M4-5 | NZ_CP035042.1 | < mnmG | 539 | | < glnQ | 38.4 | < rpmH | 499 | | dnaA > | WP_159547782.1 | 67 | 38.4 | 0.0 |
| 048 | Halomonas zhangzhouensis MCCC 1A11036 | NZ_JABFTT010000001.1 | < mnmG | 537 | | < glnQ | 32.2 | < rpmH | 499 | | dnaA > | WP_234272019.1 | 45 | 8.0 | 40.3 |
| 049 | Halomonas desiderata FB2 | NZ_JAAQTN010000031.1 rpmH<br>NZ_JAAQTN010000055.1 mnmG | < mnmG | 536 | | < erpA | | < rpmH | 513 | | dnaA > | WP_086510199.1 | 69 | | |
| 050 | Halomonas daqingensis CGMCC 1.6443 | NZ_FNVC01000006.1 | < mnmG | 536 | | < erpA | 40.7 | < rpmH | 513 | | dnaA > | WP_086510199.1 | 69 | 8.8 | 49.5 |
| 051 | Halomonas ethanolica MCCC 1A11081 | NZ_JABFTX010000004.1 | < mnmG | 534 | | < erpA | 35.9 | < rpmH | 513 | | dnaA > | WP_234271340.1 | 69 | 6.1 | 42.0 |
| 052 | Halomonas aerodenitrificans MCCC 1A11058 | NZ_JABFTV010000003.1 | < mnmG | 533 | | < erpA | 40.7 | < rpmH | 513 | | dnaA > | WP_010626004.1 | 69 | 8.3 | 49.0 |
| 053 | Halomonas kenyensis DSM 17331 | NZ_JACEFT010000002.1 | < mnmG | 541 | | < erpA | 29.3 | < rpmH | 461 | | dnaA > | WP_181513381.1 | 63 | 9.0 | 38.3 |
| 054 | Halomonas saliphila LCB169 | NZ_PJRN01000005.1 | < mnmG | 555 | | < glnQ | 26.0 | < rpmH | 509 | | dnaA > | WP_104203154.1 | 62 | 9.1 | 35.1 |
| 055 | Halomonas pellis L5 | NZ_VTPY01000006.1 | < mnmG | 521 | | < glnQ | 25.0 | < rpmH | 508 | | dnaA > | WP_149329305.1 | 64 | 8.5 | 33.5 |
| 056 | Halomonas chromatireducens AGD 8-3 | NZ_CP014226.1 | < mnmG | 542 | | < glnQ | >1.2 Mb | < rpmH | 501 | | dnaA > | WP_066446042.1 | 69 | 62.0 | > 1 Mb |
| 057 | Halomonas montanilacus PYC7W | NZ_QPII01000003.1 | < mnmG | 542 | | < glnQ | 29.5 | < rpmH | 502 | | dnaA > | WP_114478220.1 | 68 | 8.9 | 38.4 |
| 058 | Halomonas endophytica MC28 | NZ_PNRF01000027.1 | < mnmG | 528 | | < glnQ | 21.8 | < rpmH | 500 | | dnaA > | WP_102653758.1 | 70 | 8.3 | 30.1 |
| 059 | Halomonas campisalis A4 | NZ_JABFUC010000002.1 | < mnmG | 733 | | < erpA | 17.0 | < rpmH | 506 | | dnaA > | WP_238975739.1 | 69 | 9.1 | 26.1 |
| 060 | Halomonas azerbaijanica TBZ202 | SBLD01000003.1 | < mnmG | 525 | | < glnQ | 23.7 | < rpmH | 388 | | dnaA > | TFH87569.1 | 85 | 9.3 | 33.0 |
| 061 | Halomonas gudaonensis CGMCC 1.6133 | NZ_FNES01000002.1 | < mnmG | 523 | | < glnQ | 40.0 | < rpmH | 377 | | dnaA > | WP_089682774.1 | 58 | | |
| | Halomonas_I | | | | | | | | | | | | | | |
| 062 | Halomonas aquamarina 558 | NZ_FODB01000017.1 rpmH<br>NZ_FODB01000057.1 mnmG | < mnmG | 522 | | contig end | | < rpmH | 534 | | dnaA > | WP_058577312.1 | 58 | | |
| 063 | Halomonas meridiana ACAM 246 | NZ_FSQY01000001.1 | < mnmG | 579 | | < glnQ | 49.9 | < rpmH | 534 | | dnaA > | WP_058577312.1 | 58 | 5.5 | 55.5 |
| 064 | Halomonas axialensis FXH-124 | JAJGND010000025.1 rpmH<br>JAJGND010000012.1 mnmG | < mnmG | 579 | | < glnQ | | < rpmH | 534 | | dnaA > | WP_058577312.1 | 61 | | |
| 065 | Halomonas lionensis RHS90 | NZ_MWVI01000087.1 rpmH<br>NZ_MWVI01000003.1 mnmG | < mnmG | 579 | | contig end | | < rpmH | 534 | | dnaA > | WP_083025674.1 | 58 | | |
| 066 | Halomonas piezotolerans NBT06E8 | NZ_CP048602.1 | < mnmG | 578 | | < glnQ | 21.1 | < rpmH | 534 | | dnaA > | WP_153843622.1 | 60 | 21.1 | 0.0 |
| 067 | Halomonas lutescens CGMCC 1.15122 | NZ_BMHM01000004.1 | < mnmG | 576 | | < glnQ | 21.9 | < rpmH | 534 | | dnaA > | WP_188639516.1 | 60 | 1.1 | 23.0 |
| 068 | Halomonas stevensii S18214 | NZ_AJTS01000041.1 | < mnmG | 542 | | < glnQ | 19.1 | < rpmH | 533 | | dnaA > | WP_016915608.1 | 55 | 2.1 | 21.2 |
| 069 | Halomonas johnsoniae KCTC 22157 | NZ_BMXO01000019.1 | < mnmG | 542 | | < glnQ | 21.3 | < rpmH | 533 | | dnaA > | WP_193462054.1 | 56 | 1.1 | 22.4 |
| 070 | Halomonas hamiltonii KCTC 22154 | NZ_BMXN01000011.1 | < mnmG | 542 | | < glnQ | 21.8 | < rpmH | 533 | | dnaA > | WP_039179107.1 | 56 | 1.6 | 23.4 |
| 071 | Halomonas humidisoli WN018 | NZ_NSKA01000003.1 | < mnmG | 565 | | < glnQ | 21.4 | < rpmH | 534 | | dnaA > | WP_038479938.1 | 55 | 0.6 | 22.0 |
| 072 | Halomonas alkaliphila DSM 16354 | NZ_JABASV010000002.1 | < mnmG | 565 | | < glnQ | 21.6 | < rpmH | 534 | | dnaA > | WP_038479938.1 | 55 | 1.0 | 22.6 |
| 073 | Halomonas venusta NBRC 102221 | NZ_BJUL01000002.1 | < mnmG | 566 | | < glnQ | 21.5 | < rpmH | 535 | | dnaA > | WP_146942203.1 | 55 | 0.0 | 21.6 |
| 074 | Halomonas hydrothermalis Slthf2 | NZ_AP022843.1 | < mnmG | 565 | | < glnQ | 21.8 | < rpmH | 535 | | dnaA > | WP_125744652.1 | 55 | 3.7 | 25.5 |
| 075 | Halomonas andesensis DSM 19434 | NZ_RZHG01000019.1 | < mnmG | 578 | | < glnQ | 21.5 | < rpmH | 533 | | dnaA > | WP_126947390.1 | 54 | 0.8 | 21.5 |

Suppl. Table 1



| # | Species / Strain | RefSeq / Nucleotide | 5' gene | ig | | 3' gene | oriC2 Δ oriC1 [kb] | 5' gene | ig | | 3' gene | DnaA | ig dnaA-dnaN [bp] | $GC_{min}$ ←oriC2 [kb] | $GC_{min}$ ←oriC1 [kb] |
|---|---|---|---|---|---|---|---|---|---|---|---|---|---|---|---|
| | Halomonas_I (cont.) | | | | | | | | | | | | | | |
| 076 | Halomonas zhaodongensis NEAU-ST10-25 | NZ_JACCDD010000002.1 | < mnmG | 564 | | < glnQ | 21.5 | < rpmH | 534 | | dnaA > | WP_162219879.1 | 56 | 0.3 | 21.2 |
| 077 | Halomonas nigrificans MBT G8648 | NZ_NWUX01000004.1 | < mnmG | 554 | | < glnQ | 22.1 | < rpmH | 534 | | dnaA > | WP_078089687.1 | 57 | 0.0 | 22.1 |
| 078 | Halomonas malpeensis YU-PRIM-29 | NZ_WHVL01000008.1 | < mnmG | 565 | | < glnQ | 21.6 | < rpmH | 532 | | dnaA > | WP_227391296.1 | 65 | 8.9 | 30.5 |
| 079 | Halomonas populi MC | NZ_RZHE01000015.1 | < mnmG | 555 | | < glnQ | 19.7 | < rpmH | 534 | | dnaA > | WP_126952542.1 | 57 | 2.8 | 22.5 |
| 080 | Halomonas nanhaiensis JCM 18142 | NZ_RZHF01000015.1 | < mnmG | 554 | | < glnQ | 19.6 | < rpmH | 534 | | dnaA > | WP_127062055.1 | 57 | 1.8 | 17.8 |
| 081 | Halomonas songnenensis CGMCC 1.12152 | NZ_PVTK01000004.1 | < mnmG | 556 | | < glnQ | 23.0 | < rpmH | 534 | | dnaA > | WP_127062055.1 | 58 | 1.1 | 24.0 |
| 082 | Halomonas zhanjiangensis DSM 21076 | NZ_KB898653.1 | < mnmG | 574 | | < glnQ | 26.4 | < rpmH | 534 | | dnaA > | WP_018918395.1 | 45 | 1.2 | 25.1 |
| 083 | Halomonas titanicae BH1 | NZ_AOPO01000001.1 | < mnmG | 617 | | < glnQ | 21.7 | < rpmH | 533 | | dnaA > | WP_009286359.1 | 60 | 2.9 | 24.6 |
| 084 | Halomonas sedimenti QX-2 | NZ_JACCGK010000004.1 | < mnmG | 617 | | < glnQ | 21.7 | < rpmH | 533 | | dnaA > | WP_009286359.1 | 60 | 1.1 | 22.8 |
| 085 | Halomonas alkaliantarctica CRSS | NZ_SOCZ01000018.1 | < mnmG | 616 | | < glnQ | 23.5 | < rpmH | 533 | | dnaA > | WP_035582418.1 | 60 | 1.3 | 24.8 |
| 086 | Halomonas neptunia CECT 5815 | NZ_JAUFPP010000021.1 rpmH<br>NZ_JAUFPP010000005.1 mnmG | < mnmG | 616 | | < glnQ | | < rpmH | 533 | | dnaA > | WP_035582418.1 | 60 | | |
| 087 | Halomonas olivaria TYRC17 | NZ_JALPZO010000066.1 | < mnmG | 552 | | < glnQ | 17.0 | < rpmH | 534 | | dnaA > | WP_249976387.1 | 50 | 0.3 | 17.3 |
| 088 | Halomonas boliviensis LC1 | NZ_JH393257.1 | < mnmG | 595 | | < glnQ | 21.8 | < rpmH | 534 | | dnaA > | WP_007112186.1 | 60 | | |
| 089 | Halomonas maris QX-1 | NZ_JABWCV010000006.1 | < mnmG | 591 | | < glnQ | 23.5 | < rpmH | 535 | | dnaA > | WP_008959106.1 | 60 | 1.3 | 24.8 |
| 090 | Halomonas glaciei DD 39 | NZ_JACCDE010000017.1 | < mnmG | 594 | | < glnQ | 17.4 | < rpmH | 534 | | dnaA > | WP_035556369.1 | 60 | 1.5 | 18.9 |
| 091 | Halomonas sulfidaeris SST4 | NZ_QNTU01000008.1 | < mnmG | 593 | | < glnQ | 20.9 | < rpmH | 533 | | dnaA > | WP_113270223.1 | 64 | 1.6 | 22.6 |
| 092 | Halomonas utahensis NBRC 102410 | NZ_BJXV01000011.1* | < mnmG | 617 | | < glnQ | 17.7 | < rpmH | 534 | | dnaA > | WP_146875402.1 | 60 | 1.4 | 19.0 |
| 093 | Halomonas profundi MT13 | NZ_JAHRFY010000004.1 | < mnmG | 592 | | < glnQ | 21.6 | < rpmH | 534 | | dnaA > | WP_235040921.1 | 60 | 0.9 | 20.7 |
| 094 | Halomonas subterranea CGMCC 1.6495 | NZ_FOGS01000001.1 | < mnmG | 557 | | < glnQ | 22.8 | < rpmH | 532 | | dnaA > | WP_066315837.1 | 54 | 3.0 | 25.8 |
| 095 | Halomonas janggokensis DSM 18043 | NZ_JARWAJ010000002.1 | < mnmG | 557 | | < glnQ | 22.7 | < rpmH | 532 | | dnaA > | WP_085919166.1 | 55 | 0.8 | 23.6 |
| 096 | Halomonas gomseomensis DSM 18042 | NZ_JARWAI010000003.1 | < mnmG | 558 | | < glnQ | 21.3 | < rpmH | 532 | | dnaA > | WP_230445400.1 | 56 | 2.8 | 24.1 |
| 097 | Halomonas arcis CGMCC 1.6494 | NZ_FNII01000021.1 rpmH<br>NZ_FNII01000001.1 mnmG | < mnmG | 557 | | < glnQ | | < rpmH | 532 | | dnaA > | WP_089708083.1 | 57 | | |
| 098 | Halomonas salicampi BH103 | NZ_JACCDF010000018.1 rpmH<br>NZ_JACCDF010000008.1 mnmG | < mnmG | 551 | | < glnQ | | < rpmH | 524 | | dnaA > | WP_179931582.1 | 54 | | |
| 099 | Halomonas vilamensis DSM 21020 | NZ_JARWAN010000010.1 | < mnmG | 550 | | < glnQ | 23.6 | < rpmH | 534 | | dnaA > | WP_309655818.1 | 78 | 0.2 | 23.8 |
| 100 | Halomonas massiliensis Marseille-P2426 | NZ_LT699745.1 | < mnmG | 551 | | < glnQ | 26.7 | < rpmH | 522 | | dnaA > | WP_075880493.1 | 43 | 5.2 | 31.9 |
| 101 | Halomonas azerica TBZ9 | NZ_JABFHI010000006.1 | < mnmG | 584 | | < glnQ | 16,9 | < rpmH | 577 | | dnaA > | [pseudo] | 23 | 1.4 | 18.3 |
| 102 | Halomonas jeotgali Hwa | NZ_AMQY01000003.1 | < mnmG | 707 | | ydfG > | × 33.7 | < rpmH | 728 | | dnaA > | WP_017429126.1 | 61 | 1.7 | 32.0 |
| 103 | Halomonas subglaciescola ACAM 12 | NZ_LT670847.1 | < mnmG | 652 | | ydfG > | 38.8 | < rpmH | 544 | | dnaA > | WP_079553303.1 | 51 | 11.0 | 49.0 |
| 104 | Halomonas rituensis TQ8S | NZ_QPIJ01000013.1 | < mnmG | 566 | | < glnQ | 21.9 | < rpmH | 543 | | dnaA > | WP_114486371.1 | 70 | 1.6 | 23.4 |
| 105 | Halomonas zhuhanensis ZH2S | NZ_WTKP01000010.1 | < mnmG | 565 | | < glnQ | 23.7 | < rpmH | 544 | | dnaA > | WP_160419533.1 | 71 | 1.3 | 25.0 |
| | Halomonas_F | | | | | | | | | | | | | | |
| 106 | Halomonas pacifica NBRC 102220 | NZ_BJUK01000023.1 rpmH<br>NZ_BJUK01000013.1 mnmG | < mnmG | 544 | | < glnQ | | < rpmH | 556 | | dnaA > | WP_146803234.1 | 64 | | |
| | Halomonas_B | | | | | | | | | | | | | | |
| 107 | Halomonas salipaludis WRN001 | NZ_NSKB01000003.1 | < mnmG | 544 | | < glnQ | 26.3 | < rpmH | 501 | | dnaA > | WP_095620543.1 | 48 | 9.5 | 35.8 |
| 108 | Halomonas qiaohouensis DSM 26770 | NZ_JARWAM010000001.1 | < mnmG | 544 | | < glnQ | 27.7 | < rpmH | 502 | | dnaA > | WP_309715924.1 | 48 | 8.3 | 36.0 |
| 109 | Halomonas pantelleriensis AAP | NZ_FNGH01000005.1 | < mnmG | 547 | | < glnQ | 26.3 | < rpmH | 501 | | dnaA > | WP_089658023.1 | 47 | 3.9 | 30.2 |
| | Halomonas_E | | | | | | | | | | | | | | |
| 110 | Halomonas taeanensis BH539 | NZ_FNCI01000005.1 | < mnmG | 615 | | < glnQ | 26.6 | < rpmH | 568 | | dnaA > | WP_148252394.1 | 61 | 9.3 | 35.9 |
| 111 | Halomonas niordiana ATF 5.4 | NZ_SDSD01000004.1 | < mnmG | 617 | | < glnQ | 26.6 | < rpmH | 557 | | dnaA > | WP_136254741.1 | 62 | 1.7 | 28.4 |
| | Halomonas_A | | | | | | | | | | | | | | |
| 112 | Halomonas anticariensis FP35 = DSM 16096 | NZ_KE332393.1 | < mnmG | 549 | | < glnQ | 27.8 | < rpmH | 547 | | dnaA > | WP_016418290.1 | 82 | 1.1 | 28.9 |
| 113 | Halomonas qijiaojingensis KCTC 22228 | NZ_BMXS01000003.1 | < mnmG | 549 | | < glnQ | 28.3 | < rpmH | 565 | | dnaA > | WP_189466589.1 | 90 | 1.8 | 30.1 |
| 114 | Halomonas rifensis CECT 7698 | not yet in RefSeq | | | | | | | | | | | | | |
| 115 | Halomonas xinjiangensis TRM 0175 | NZ_JPZL01000003.1 | < mnmG | 537 | | < glnQ | 25.5 | < rpmH | 531 | | dnaA > | WP_043531287.1 | 85 | 2.4 | 27,9 |
| | Aidingimonas | | | | | | | | | | | | | | |
| 116 | Aidingimonas halophila DSM 19219 | NZ_FNNI01000003.1 | < mnmG | 550 | | < glnQ | 33.1 | < rpmH | 447 | | dnaA > | WP_148252394.1 | 54 | 2.0 | 35.1 |
| 117 | Aidingimonas lacisalsi XHU 5135 | NZ_VRYH01000001.1 | < mnmG | 559 | | < glnQ | 33.5 | < rpmH | 459 | | dnaA > | WP_148252394.1 | 55 | 3.3 | 36.5 |
| | Modicisalibacter | | | | | | | | | | | | | | |
| 118 | Modicisalibacter tunisiensis LIT2 | NZ_JAGXFD010000001.1 | < mnmG | 554 | | < glnQ | 33.6 | < rpmH | 607 | | dnaA > | WP_224416154.1 | 71 | 32.3 | 65.9 |
| 119 | Halomonas coralii 362.1 | NZ_QWBW01000012.1 | < mnmG | 564 | | < glnQ | 22.4 | < rpmH | 567 | | dnaA > | WP_129139778.1 | 64 | 8.8 | 31.2 |
| 120 | Halomonas zincidurans B6 | NZ_JNCK01000001.1 | < mnmG | 587 | | < glnQ | 88.9 | < rpmH | 569 | | dnaA > | WP_031384876.1 | 57 | 96.9 | 185.7 |
| 121 | Halomonas radicis EAR18 | NZ_CAAHFN010000048.1 | < mnmG | 638 | | < glnQ | 22.4 | < rpmH | 572 | | dnaA > | WP_136065006.1 | 61 | 6.0 | 28.4 |
| 122 | Halomonas lutea DSM 23508 | NZ_KB899996.1 | < mnmG | 573 | | < glnQ | 18.8 | < rpmH | 570 | | dnaA > | WP_019019262.1 | 67 | 6.0 | 24.8 |
| 123 | Halomonas xianhensis CGMCC 1.6848 | NZ_FOPY01000006.1 | < mnmG | 586 | | < glnQ | 41.9 | < rpmH | 570 | | dnaA > | WP_092845625.1 | 68 | 6.8 | 48.6 |
| 124 | Halomonas ilicicola DSM 19980 | NZ_FQUJ01000004.1 | < mnmG | 591 | | < glnQ | 31.0 | < rpmH | 571 | | dnaA > | WP_072820265.1 | 67 | 8.2 | 39.2 |
| 125 | Halomonas muralis DSM 14789 | NZ_FNGI01000004.1 | < mnmG | 611 | | < glnQ | 19.9 | < rpmH | 564 | | dnaA > | WP_089727921.1 | 65 | 8.9 | 28.9 |
| | Salinicola | | | | | | | | | | | | | | |
| 126 | Salinicola halimionae CPA60 | NZ_PZJQ01000002.1 | < mnmG | 1026 | | < glnQ | 28.9 | < rpmH | 594 | | dnaA > | WP_110654280.1 | 185 | 0.3 | 29.2 |
| 127 | Salinicola halophyticus CR45 | NZ_PZJN01000009.1 | < mnmG | 975 | | < glnQ | 30.3 | < rpmH | 617 | | dnaA > | WP_075561441.1 | 184 | 1.9 | 32.2 |
| 128 | Salinicola socius SMB35 = DSM 19940 | NZ_MSDO01000012.1 | < mnmG | 911 | | < glnQ | 28.6 | < rpmH | 604 | | dnaA > | WP_075570063.1 | 331 | 1.5 | 30.1 |
| 129 | Salinicola salarius DSM 18044 | NZ_PZJT01000012.1 rpmH<br>NZ_PZJT01000021.1 mnmG | < mnmG | 761 | | < glnQ | | < rpmH | 606 | | dnaA > | WP_071232451.1 | 169 | | |
| 130 | Salinicola lusitanus CR50 | NZ_PZJM01000009.1 | < mnmG | 749 | | < glnQ | 26.8 | < rpmH | 605 | | dnaA > | WP_110601628.1 | 166 | 8,4 | 35.2 |
| 131 | Salinicola acroporae LMG 28587 | NZ_PZJW01000003.1 | < mnmG | 748 | | < glnQ | 26.8 | < rpmH | 604 | | dnaA > | WP_110715226.1 | 166 | 8.6 | 35.4 |
| 132 | Salinicola corii L3 | NZ_VTPX01000004.1 | < mnmG | 761 | | < glnQ | 26.9 | < rpmH | 610 | | dnaA > | WP_149435093.1 | 167 | 1,1 | 28.0 |
| 133 | Salinicola peritrichatus JCM 18795 | NZ_PZJU01000083.1 rpmH<br>NZ_PZJU01000033.1 mnmG | < mnmG | 588 | | < glnQ | | < rpmH | 618 | | dnaA > | WP_110651634.1 | 182 | | |
| 134 | Salinicola rhizosphaerae KCTC 32998 | NZ_BMZI01000002.1 | < mnmG | 734 | | < glnQ | 27.1 | < rpmH | 621 | | dnaA > | WP_189443599.1 | 128 | 7.1 | 34.3 |
| 135 | Salinicola tamaricis F01 | NZ_CP023559.1 | < mnmG | 795 | | < glnQ | 29.2 | < rpmH | 622 | | dnaA > | [pseudo] | | 1.9 | |
| 136 | Salinicola endophyticus CPA92 | NZ_PZJO01000003.1 | < mnmG | 778 | | < glnQ | 29.2 | < rpmH | 624 | | dnaA > | WP_110673717.1 | 140 | 7.1 | 36.2 |
| 137 | Salinicola halophilus CECT 5903 | NZ_PZJV01000006.1 | < mnmG | 850 | | < glnQ | 25.2 | < rpmH | 684 | | dnaA > | WP_110669649.1 | 142 | 13.7 | 38.9 |
| 138 | Salinicola aestuarinus CPA62 | NZ_PZJP01000004.1 | < mnmG | 800 | | < glnQ | 25.0 | < rpmH | 683 | | dnaA > | WP_110685640.1 | 142 | 6.1 | 31.1 |
| | Chromohalobacter | | | | | | | | | | | | | | |
| 139 | Chromohalobacter israelensis DSM 6768 | NZ_JQNW01000005.1 | < mnmG | 601 | | < glnQ | 23.6 | < rpmH | 589 | | dnaA > | WP_011505312.1 | 49 | 1.7 | 25.4 |
| 140 | Chromohalobacter salexigens DSM 3043 | NC_007963.1 | < mnmG | 602 | | < glnQ | 23.6 | < rpmH | 589 | | dnaA > | WP_011505312.1 | 49 | 23,8 | 0.1 |
| 141 | Chromohalobacter marismortui DSM 6770 | NZ_SOBR01000004.1 | < mnmG | 614 | | < glnQ | 23.4 | < rpmH | 589 | | dnaA > | WP_133697561.1 | 48 | 1.8 | 25.3 |
| 142 | Chromohalobacter canadensis 85B | NZ_JAJQJH010000002.1 | < mnmG | 605 | | < glnQ | 23.6 | < rpmH | 590 | | dnaA > | WP_277810369.1 | 49 | 2.7 | 26.3 |
| | Kushneria | | | | | | | | | | | | | | |
| 143 | Kushneria konosiri X49 | NZ_CP021323.1 | < mnmG | 622 | | < erpA | 38.7 | < rpmH | 332 | | dnaA > | WP_086621096.1 | 95 | 2.5 | 36.2 |
| 144 | Kushneria marisflavi SW32 = DSM 15357 | NZ_CP021358.1 | < mnmG | 622 | | < erpA | 36.2 | < rpmH | 332 | | dnaA > | WP_086900485.1 | 94 | 10.5 | 46.7 |
| 145 | Kushneria phyllosphaerae EAod3 | NZ_ONZI01000001.1 | < mnmG | 620 | | < erpA | 43.0 | < rpmH | 332 | | dnaA > | WP_108841163.1 | 97 | 4.9 | 47.9 |

Suppl. Table 1

| # | Species / Strain | RefSeq / Nucleotide | 5' gene | ig | | 3' gene | oriC2 Δ oriC1 [kb] | 5' gene | ig | | 3' gene | DnaA | ig dnaA·dnaN [bp] | $GC_{min}$ ←oriC2 [kb] | $GC_{min}$ ←oriC1 [kb] |
|---|---|---|---|---|---|---|---|---|---|---|---|---|---|---|---|
| | Kushneria (cont.)) | | | | | | | | | | | | | | |
| 146 | Kushneria indalinina DSM 14324 | NZ_QRDJ01000007.1 | < mnmG | 622 | | < erpA | 38.3 | < rpmH | 330 | | dnaA > | WP_115854213.1 | 97 | 7.7 | 46.0 |
| 147 | Kushneria pakistanensis BKCTC 42082 | NZ_BMZM01000002.1 | < mnmG | 620 | | < erpA | 35.1 | < rpmH | 330 | | dnaA > | WP_189516301.1 | 97 | 0.3 | 35.4 |
| 148 | Kushneria avicenniae DSM 23439 | NZ_FOLY01000003.1 | < mnmG | 623 | | < erpA | 43.2 | < rpmH | 330 | | dnaA > | WP_090133075.1 | 97 | 7.2 | 50.4 |
| 149 | Kushneria aurantia DSM 21353 | NZ_KB907862.1 | < mnmG | 964 | | < erpA | 60.5 | < rpmH | 396 | | dnaA > | WP_019949883.1 | 87 | 6.4 | 66.9 |
| 150 | Kushneria sinocarnis DSM 23229 | NZ_RBIN01000001.1 | < mnmG | 607 | | < erpA | 52.4 | < rpmH | 390 | | dnaA > | WP_121170259.1 | 83 | 1.8 | 54.3 |
| 151 | Kushneria phosphatilytica YCWA18 | NZ_CP043420.1 | < mnmG | 601 | | < erpA | 35.3 | < rpmH | 396 | | dnaA > | WP_070980264.1 | 83 | 4.9 | 40.2 |
| | Phytohalomonas | | | | | | | | | | | | | | |
| 152 | Phytohalomonas tamaricis R4HLG17 | NZ_PVRV01000019.1 | < mnmG | 623 | | < erpA | 27.1 | < rpmH | 458 | | dnaA > | WP_106476778.1 | 81 | 0.4 | 26.7 |
| | Carnimonas | | | | | | | | | | | | | | |
| 153 | Carnimonas nigrificans ATCC BAA-78 | NZ_JAGO01000013.1 | < mnmG | 855 | | < erpA | 45.4 | < rpmH | 494 | | dnaA > | WP_025733208.1 | 113 | 1.2 | 46.7 |
| | Halotalea | | | | | | | | | | | | | | |
| 154.1 | Halotalea alkalilenta DSM 17697 | NZ_KK211226.1 rpmH<br>NZ_KK211229.1 mnmG | < mnmG | 661 | | < erpA | | < rpmH | 428 | | dnaA > | WP_027350303.1 | 73 | | |
| 154.2 | Halotalea alkalilenta IHB B 13600 | NZ_CP015243.1 | < mnmG | 663 | | < erpA | 102.3 | < rpmH | 428 | | dnaA > | WP_064121474.1 | 73 | 13.8 | 116.1 |
| | Zymobacter | | | | | | | | | | | | | | |
| 155 | Zymobacter palmae IAM 14233 = DSM 10491 | NZ_AP018933.1 | < mnmG | 678 | | < COG2859 | 22.5 | < sdiA | 345 | | dnaA > | WP_027704352.1 | 188 | 3.2 | 25.7 |
| | Larsenimonas | | | | | | | | | | | | | | |
| 156 | Larsenimonas salina CCM 8464 | NZ_JAMLJJ010000003.1 | < mnmG | 562 | | < erpA | 33.2 | < rpmH | 405 | | dnaA > | WP_251927440.1 | 59 | 1.4 | 34.6 |
| 157 | Larsenimonas suaedae DSM 22428 | NZ_JARWAO010000001.1 | < mnmG | 568 | | < erpA | 28.2 | < rpmH | 392 | | dnaA > | WP_251593434.1 | 59 | 1.7 | 30.0 |
| | Cobetia | | | | | | | | | | | | | | |
| 158 | Cobetia marina JCM 21022 | NZ_CP017114.1 | < mnmG | 1388 | | < glnQ | 31.9 | < rpmH | 841 | | dnaA > | WP_077379738.1 | 148 | 37.8 | 5.9 |
| 159 | Cobetia crustatorum JO1 | NZ_JEMF01000015.1 rpmH<br>NZ_JEMF01000037.1 mnmG | < mnmG | seq gap | | < glnQ | | < rpmH | 943 | | dnaA > | WP_024951037.1 | 151 | | |
| | uncertain | | | | | | | | | | | | | | |
| 160 | Terasakiispira papahanaumokuakeensis PH27A | NZ_MDTQ01000001.1 | < mnmG | 541 | | ompV > | 12.9 | < rpmH | 568 | | dnaA > | WP_068999754.1 | 35 | | |
| | not Halomonadaceae | | | | | | | | | | | | | | |
| 161 | Halovibrio variabilis DSM 3050 | NZ_BJXV01000011.1 | < mnmG | 617 | | < glnQ | 17.2 | < rpmH | 534 | | dnaA > | WP_146875402.1 | 60 | 18.6 | 1.4 |
| 162 | Halovibrio salipaludis YL5-2 | NZ_NSKD01000001.1 | < mnmG | 286 | | < DUF3833 | 9.3 | < rpmH | 626 | | dnaA > | WP_233128829.1 | 44 | 5.4 | 3.9 |

* see de la Haba *et al.* (2023) Supplementary Fig 1

× inverted gene order

- strong DUE
- weak DUE
- no DUE
- no oriC

135 Salinicola tamaricis F01 DnaA (reconstructed)

```
  1 VSVALWQQCL NSLQDDARSS STPGSALCRR RMARRTSSVC WRPTAFVRDW VSDKYLKRIN
 61 ELLREVASSS KPPKVRLVVG SRRAAAVSPR IWARRYRPPA SRRSAQCARV KTSPRSPEVA
121 EEQALDQNRE REEGGRRGGE RQVQVEGSLK HQSGLNPNFT FETFVEGKSN QLARAASRQV
181 SENPGGAYNP LFLYGGVGLG KTHLMHAVGN HLAGLRDNAK VVYLHSERFV ADMVKALQLN
241 AINDFKRFYR SVDALLIDDI QFFGGKERSQ EEFFHTFNAL LEGGQQMILT SDRYPKEISG
301 VEERLKSRFG WGLTVAIEPP ELETRVAILM KKADQAKVDL PHDAAFFIAQ KIRSNVRELE
361 GALKKVIADS HFMGKPITQD FIRESLKDLL ALQDKQVGIE NIQRTVAEYY KIKMADLLSK
421 RRSRSVARPR QVAMALAKEL TNHSLPEIGD AFGGRDHTTV LHACRKVVAL KEESADIRED
481 YKNLMRLLTS
```

101 Halomonas azerica TBZ9 DnaA (reconstructed)

```
  1 MSLALWQQCL DYLQDELNSQ QFNTWIRPLQ AEEGESNELC LLAPNRFVRD WVNDKYAKRI
 61 SELVRELSPA KPPKVSLTVG SRRVAAPAPQ PRQLGTPFQP QPTGNPVSAS ANRSPSPAVS
121 TPSRTHMNDD REIDQLREES ASKRSGERQV QVEGSLKHTS GLNPNFTFET FVEGKSNQLA
181 RAASRQVSEN PGGAYNPLFL YGGVGLGKTH LMHAVGNALA ERRENARVVY LHSERFVADM
241 VKALQLNAIN DFKRFYRSVD ALLIDDIQFF AGKERSQEEF FHTFNALLEG GQQMILTSDR
301 YPKEISGVEE RLKSRFGWGL TVAIEPPELE TRVAILMKKA DQSKVDLPHD AAFFIAQKIR
361 SNVRELEGAL KKEKVIADSH FMGKPITQDF IRESLKDLLA LQDKQVGVDN IQRTVAEYYK
421 IKVADLLSKR RSRSVARPRQ VAMALAKELT NHSLPEIGDA FGGRDHTTVL HACRKVKALQ
481 EENADIREDY KNLLRL
```
