## Supplementary material for "Dual chromosomal origins of replication (*oriC*) in the genomes of the Halomonadaceae – a prediction study": Suppl. Table 2_v01.zip: Suppl. Table 2_v01.pdf

color code: ATATATAAT SIDD strong (0–3); CTTTTATGA SIDD weak (3–6); GAATATATA SIDD strong / DnaA box overlap; GAATTAATA SIDD weak / DnaA box overlap; **TGTGGATAA** DnaA box R1-type; TGTGGATAA DnaA box cons rev; TGTGGATAA DnaA box relaxed rev; TTTCACAAG DnaA box overlap; CTGTCCACA DnaA box cons fwd; TCCGGGGCA DnaA box relaxed fwd; TATTACTAG DnaA trio; ATATCAATA central C of IHF site; ATG orf with start/stop codon.

(numbering as in Suppl. Table 1)

#001 Halomonas elongata DSM 2581
#002 Halomonas eurihalina DSM 5720
#003 Halomonas caseinilytica JCM 14802
#004 Halomonas halmophila NBRC 15537
#005 Halomonas sabkhae CECT 7246
#006 Halomonas almeriensis CECT 7050
#007 Halomonas halophila NBRC 102604
#008 Halomonas salina DSM 5928
#009 Halomonas smyrnensis AAD6
#010 Halomonas borealis ATF 5.2
#011 Halomonas organivorans CECT 5995
#012 Halomonas koreensis Lim DSM 23530
#013 Halomonas stenophila CECT 7744
#014 Halomonas maura CECT 5298
#015 Halomonas nitroreducens 11S
#016 Halomonas halocynthiae DSM 14573
#017 Halomonas cerina CECT 7282
#018 Halomonas huangheensis BJGMM-B45
#019 Halomonas cupida DSM 4740
#020 Halomonas litopenaei SYSU ZJ2214
#021 Halomonas urmiana TBZ3
#022 Halomonas saccharevitans CGMCC 1.6493
#023 Halomonas denitrificans DSM 18045
#024 Halomonas aestuarii Hb3
#025 Halomonas ventosae CECT 5797
#026 Halomonas fontilapidosi CECT 7341
#027 Halomonas daqiaonensis CGMCC 1.9150
#028 Halomonas lysinitroph a 3(2)
#029 Halomonas marinisediminis 204
#030 Halomonas icarae D1-1
#031 Halomonas halodenitrificans DSM 735
#032 Halomonas alimentaria DSM 15356
#033 Halomonas shengliensis CGMCC 1.6444
#034 Halomonas ramblicola CECT 7896
#035 Halomonas alkalicola M2
#036 Halomonas mongolie nsis DSM 17332
#037.1 Halomonas campaniensis 5AG
#037.2 Halomonas campaniensis LS21
#038 Halomonas urumqiensis BZ-SZ-XJ27
#039 Halomonas heilongjiangensis DSM 26881
#040 Halomonas korlensis CGMCC 1.6981
#041 Halomonas sulfidivorans MCCC 1A13718
#042 Halomonas antri Y3S6
#043 Halomonas sulfidoxydans MCCC 1A11059
#044 Halomonas lactosivorans KCTC 52281
#045 Halomonas bachuensis DX6
#046 Halomonas diversa MCCC 1A13316
#047 Halomonas tianxiuensis BC-M4-5
#048 Halomonas zhangzhouensis MCCC 1A11036
#049 Halomonas desiderata FB2
#050 Halomonas daqingensis CGMCC 1.6443
#051 Halomonas ethanolica MCCC 1A11081
#052 Halomonas aerodenitrificans MCCC 1A11058
#053 Halomonas kenyensis DSM 17331
#054 Halomonas saliphila LCB169
#055 Halomonas pellis L5
#056 Halomonas chromatireducens AGD 8-3
#057 Halomonas montanilacus PYC7W
#058 Halomonas endophytica MC28
#059 Halomonas campisalis A4
#060 Halomonas azerbaijanica TBZ202
#061 Halomonas gudaonensis CGMCC 1.6133
#062 Halomonas aquamarina 558
#063 Halomonas meridiana ACAM 246
#064 Halomonas axialensis FXH-124
#065 Halomonas lionensis RHS90
#066 Halomonas piezotolerans NBT06E8
#067 Halomonas lutescens CGMCC 1.15122
#068 Halomonas stevensii S18214
#069 Halomonas johnsoniae KCTC 22157
#070 Halomonas hamiltonii KCTC 22154
#071 Halomonas humidisoli WN018
#072 Halomonas alkaliphila DSM 16354
#073 Halomonas venusta NBRC 102221
#074 Halomonas hydrothermalis Slthf2
#075 Halomonas andesensis DSM 19434
#076 Halomonas zhaodongensis NEAU-ST10-25
#077 Halomonas nigrificans MBT G8648
#078 Halomonas malpeensis YU-PRIM-29
#079 Halomonas populi MC
#080 Halomonas nanhaiensis JCM 18142
#081 Halomonas songnenensis CGMCC 1.12152
#082 Halomonas zhanjiangensis DSM 21076
#083 Halomonas titanicae BH1
#084 Halomonas sedimenti QX-2
#085 Halomonas alkaliantarctica CRSS
#086 Halomonas neptunia CECT 5815
#087 Halomonas olivaria TYRC17
#088 Halomonas boliviensis LC1
#089 Halomonas maris QX-1
#090 Halomonas glaciei DD 39
#091 Halomonas sulfidaeris SST4
#092 Halomonas utahensis NBRC 102410
#093 Halomonas profundi MT13
#094 Halomonas subterranea CGMCC 1.6495
#095 Halomonas janggokensis DSM 18043
#096 Halomonas gomseomensis DSM 18042
#097 Halomonas arcis CGMCC 1.6494
#098 Halomonas salicampi BH103
#099 Halomonas vilamensis DSM 21020
#100 Halomonas massiliensis Marseille-P2426
#101 Halomonas azerica TBZ9
#102 Halomonas jeotgali Hwa
#103 Halomonas subglaciescola ACAM 12
#104 Halomonas rituensis TQ8S
#105 Halomonas zhuhanensis ZH2S
#106 Halomonas pacifica NBRC 102220
#107 Halomonas salipaludis WRN001
#108 Halomonas qiaohouensis DSM 26770
#109 Halomonas pantelleriensis AAP
#110 Halomonas taeanensis BH539
#111 Halomonas niordiana ATF 5.4
#112 Halomonas anticariensis FP35 = DSM 16096
#113 Halomonas qijiaojingensis KCTC 22228
#114 Halomonas rifensis CECT 7698
#115 Halomonas xinjiangensis TRM 0175
#116 Aidingimonas halophila DSM 19219
#117 Aidingimonas lacisalsi XHU 5135
#118 Modicisalibacter tunisiensis LIT2
#119 Halomonas coralii 362.1
#120 Halomonas zincidurans B6
#121 Halomonas radicis EAR18
#122 Halomonas lutea DSM 23508
#123 Halomonas xianhensis CGMCC 1.6848
#124 Halomonas ilicicola DSM 19980
#125 Halomonas muralis DSM 14789
#126 Salinicola halimionae CPA60
#127 Salinicola halophyticus CR45
#128 Salinicola socius SMB35 = DSM 19940
#129 Salinicola salarius DSM 18044
#130 Salinicola lusitanus CR50
#131 Salinicola acroporae LMG 28587
#132 Salinicola corii L3
#133 Salinicola peritrichatus JCM 18795
#134 Salinicola rhizosphaerae KCTC 32998
#135 Salinicola tamaricis F01
#136 Salinicola endophyticus CPA92
#137 Salinicola halophilus CECT 5903
#138 Salinicola aestuarinus CPA62
#139 Chromohalobacter israelensis DSM 6768
#140 Chromohalobacter salexigens DSM 3043
#141 Chromohalobacter marismortui DSM 6770
#142 Chromohalobacter canadensis 85B
#143 Kushneria konosiri X49
#144 Kushneria marisflavi SW32 = DSM 15357
#145 Kushneria phyllosphaerae EAod3
#146 Kushneria indalinina DSM 14324
#147 Kushneria pakistanensis BKCTC 42082
#148 Kushneria avicenniae DSM 23439
#149 Kushneria aurantia DSM 21353
#150 Kushneria sinocarnis DSM 23229
#151 Kushneria phosphatilytica YCWA18
#152 Phytohalomonas tamaricis R4HLG17
#153 Carnimonas nigrificans ATCC BAA-78
#154.1 Halotalea alkalilenta DSM 17697
#154.2 Halotalea alkalilenta IHB B 13600
#155 Zymobacter palmae IAM 14233 = DSM 10491
#156 Larsenimonas salina CCM 8464
#157 Larsenimonas suaedae DSM 22428
#158 Cobetia marina JCM 21022
#159 Cobetia crustatorum JO1
#160 Terasakiispira papahanaumokuakeensis PH27A
#161 Halovibrio variabilis DSM 3050
#162 Halovibrio salipaludis YL5-2

GenSkew GCmin pos. 4036634 (GCmin<>oriC1 25.2 kb; GCmin<>oriC2 7.0 kb; oriC2<>oriC1 18.2 kb)

>Halomonas elongata DSM 2581 [NC_014532.2:4061575..4061825;1..249], <rpmH oriC1 dnaA>, 504 bp

AGCGTTACCGGCGTTGGCGGTGCCGCTGGACGGAACCGACAGATCGCTACTGTCCGTATCCGCGGCATCCTGGGATGACGCTGGCGAGTCGCTGGTGATG
GCCGGGGCCTCCGGCGCCTGGGTCGCCTGGCCGTAGTCCTGATTCCACTGCACGACGATCAGATAGGCGAGTATCGCCAGGGGAATCAACAGGAGTAGTC
GTTTTACGTCCATGAGGGGTCTGCCCAATGGGTGGTGAACATTCCGATGGCGCAGCGTGAGTCGCACACCCGAGATACGCGAAAGCCTGCCATCGGCAGG
CCGGCTTTCATCAGCGGAGCGTCCGCTTTCGGGGGGGCGGCGTCAAGCTTCGCGACGAGAGGCGATAACAGGCTCATCCGAACTCCGATCGATGGCGCTG
GCAACGGTGCGCCCGACCGGTCATCGACGGTCTGTCGAGGCGGTAGTGGGGCGGCCGGAGGTCTTCTGCGCTTCGCGCTCCAGGCGGCGCCACATGCCAT
GTAATTGGCGATGGACAACCTCATTGTCCAGCTCCTGTACGCCACGCCGGGCGAGTAACACGATGTCCACGGCGGGGAGTCGATGCTGGCGAAGCCGAAT
GGATTCGCGCGTGATTCGCTTGAGGCGATTGCGGTCAACGGCCCGACGGACGTTCTTCTTGCTGAAGATGAGGCCGAGACGAGGGTGGCCCAATTCGTTG
GGCGAGGCCAGGGCCATCAATCCCTTGCCGTGAAGCTTGAAAGCCGCCGTATCGAAGACGCGTCGATAGTCCTCGGCGGTCAGTAGGCGCAGCCGCCGAG
GGAAGCCCTGATAGGACACGCGTTCGCGATCAGGCGCTCAGACGCTTGCGGCCCTTGGCGCGACGACGCGCCAGGACGGCACGACCATTCTTGGTGGCCA
TGCGAGCGCGGAAGCCATGTGCGCGCTTGCGCTTGAGTACGCTGGGTTGGAAAGTGCGTTTCATAACTGATTATTCCCACGTGGTTTTTTGGAACGGATC

GATAACTTTTCAGAGGCGCCCGGTTTCCGCTGGAAACCTGGGGGATTCGGCTCAAGACCGGGCATTGTAGTGATTTCACGGCCTCGGTGCAATTGCCCT
GTTCGCTCAGCGTCAACCGGGGTGGGGCCATCGAGGTCACAGTTATTATTAATTAAAGGTATATCCTGATACGGTTGTTATTATAACTGGGGAAACATTC
**TGTGGATAA**GCTGATATAACTTAAAAATATCAGTACGTTACTTCGATAATTATGGTGTGGACATCCTTGGGGGCAGTGGTGTATCCACTGTGGGTATCCG
TTGCATGGAAAGGAGTCTTGTCCACAGCGCTGGATGATCACCATATTCCTCCCGGTCATTTCCCGGCCTTCGGGACAGGGTTTTCCACAAAGGCAATCGA
ATCGCTGCTTCATGTGGATAACCCGGAGGTTGGGCGATACAATGTGCGGTCGTTCTTGAATCTACGATGTGAGGTCGCGTGTCGCTCGCTCTCTGGCAAC

AGTGTCTGGCTACCCTGCAGGATGAATTGAGCTCCCAGCAGTTCAACACCTGGATTCGCCCATTGCAGGCGGAAGAGGAGGGGGAGTCCAACCAGTTGCG
CCTGCTGGCGCCCAACCGGTTCGTGCGCGATTGGGTCAGTGACAAGTATGCCAAGCGTATCAGCGAACTGATGCGCGAGCTGGCCCCTGCCAAGCCGCCC
AAGGTGGTGTTGACGGTGGGTAGTCGCCGTGCGGCTCCCGCTCCCCAGCCGCGAGAGCTCGGCGATCCCGTGTCCGCGGCGCCTCGGTCGCCCGCCGTGC
ATACGCCGCCGCGGGGTGATGTGGCGGATGAATCCGAAATCGACCAGTTACGCGAGGAAGGCGAGGCGCAGCAGCGTCGTCCGACGGGCGAACGCCAGGT
CCAGGTCGAGGGTAGCCTCAAGCACAACAGTGGTCTGAACCATAATTTCACCTTCGAAACCTTCGTCGAGGGCAAGTCCAACCAATTGGCCCGGGCTGCG
TCGCGGCAGGTTTCCGAAAATCCCGGTGGCGCCTATAACCCGCTGTTTCTCTATGGCGGCGTCGGCCTGGGCAAGACCCACCTGATGCATGCCGTCGGCA
ATGCCCTGGATGCCCGCCGCGAGAACGCACGGGTGGTGTATCTGCACTCCGAGCGCTTCGTCGCCGACATGGTCAAGGCGCTGCAGCTCAATGCCATCAA
TGACTTCAAGCGCTTTTACCGTAGTGTCGATGCCTTGCTGATCGACGATATCCAGTTCTTCGCCGGCAAGGAACGCTCTCAGGAAGAGTTCTTCCATACC
TTCAATGCCTTGCTCGAAGGCGGTCAGCAGATGATCCTGACCTCGGACCGTTATCCCAAGGAAATCAGTGGGGTAGAGGAGCGCCTCAAGTCGCGCTTCG
GCTGGGGGCTGACAGTGGCGATCGAGCCGCCGGAGCTCGAGACCCGGGTCGCGATCCTGATGAAGAAGGCCGATCAGGCCAAGGTCAATCTGCCCCATGA

>Halomonas elongata DSM 2581 [NC_014532.2:4043405..4043904], <mnmG oriC2 <COG1126 (glnQ), 535 bp

CCAGTCCCTGGATCGAACGGACGACCTGCAACTGCACATCGAACGGCAGTGACGTGGAAATGCCGTTGGGATACAGCTCGTGGGTGTCCAGCCCCTCCGG
CTCGACGAAGATCTGGTGGCTCGACTTGTCGGCGAAGCGATGCACCTTGTCCTCGATGGAGGGGCAATAGCGGGGACCCACACCCTCGATGACCCCGGAG
AACATCGGCGAACGCTCCAGGTTGGCGTGGATGATCTCGTGGGTCCGCTCATTGGTGTGCGCAATATGGCAGCTCACCTGGCGCGGATGCTGGCTCCGTT
CGCCCAGATAGGACATGACCGGCGTGGGCTCATCGCCGGGCTGCTCGTCGAGCTTCGAGAAATCCACGCTCTTGGCATCGAGACGCGGCGGGGTGCCGGT
CTTCAGGCGATCGACGCGAAACGGCAAGGCCCGGAGGCGTTCGGCGAGGGCATTGGAGGGCGGATCGCCGGCACGGCCCCCACGGCTGTTGTCCAGGCCG
ATGTGGATAACACCGCCGAGGAAAGTGCCTGTGCACAACACCACGGTCTCGCTCATGAAGCGAATTCCCGTTTCGGTGCATACACCGCGAACCTGGTCAC
CCTCGACGATGAGATCGCCGGCTGCCTGCTGGAAGATCGTCAGGTTGGCCTGGTTTTCCAGCATGCCGCGAATGGCGGCCTTGTAGCGAACGCGATCCGC
CTGGGCACGGGTGGCACGCACGGCCGGTCCTTTCCGGGCGTTGAGCACACGGAACTGGATACCGCCCATGTCGGTGGCCAGCCCCATGGCACCACCGAGG
GCATCGATTTCCTTGACCAGATGGCTCTTGCCGATCCCGCCGATGGCGGGATTGCACGACATCTGGCCGAGCGTCTCGATGTTGTGGGTCAAAAGCAACG
TCCGGCAGCCCGTGCGGGCCGAGGCCAGAGCGGCTTCGGTGCCCGCATGACCACCACCGATGACGATGACGTCGAAGCGGTCGGGGTACTCCAAGGTTCA

CCTCGTTCGGCGCTGCGGCGCCGATATTGCGGATCGGGGATGAAATCAGCCAGCCAGTATAAACCTCCTGTGGGTAATCGTCAGGGTTTTCCACACCCGG
ATTTTCATCTTCGTCCCCTTTTCACACCACTAGCTATTAAATAAACAATAAAAGAATAGGAAGTTCTGTTGTTGTAGTGACTACTGGTGTGGACGTTTTC
**CGTGGATAA**ACCGTTTTATTGCCATAATATCAATATCTTGCAATGTTGATCGAAAGTAGGAAAAGCAGCGGGCGAGCTGCGGACGATGTGTTCGGGGCCCT
GTGGATGGAAAGCCGGTTTGCCCACAAGCTGACACGAACACGGCTTTTCCACCGATCACTACCGGGCTTGTCACCGCAGCTGTGAACAACCGGGGCGATT
TGCCCACATCCCCGGGATGGCGGTGGGTGTGGCGTGATTTTGGTTCATTTGGCAATCGGGTTTTCCACAGCCGAAAAAAAGGCCCTGTCGATAACGACAG

GGCCTTCGGGCATGCCTGAAAAGGTGGTATCAGTGCTTGAGAACTCGAGACAGGAAGCTTTTTGTCGGTGCCTCCGCAGGGGTATCGAAGATCGTCTCGG
GGGGACCTTCTTCGACGATCCGGCCCTTGTCGATGAAGATGACCCGATCGGCGACTTCTCGGGCGAAGCCCATCTCGTGGGTCACGATCATCATGGTCAT
GCCGTCGTTGGCCAGCTCGCGCATGGCATCGAGTACTTCGCCGATCATCTCCGGGTCCAGTGCCGAGGTGGGCTCGTCGAAGAGCATCAGGCGAGGTTCC
ATGGCCAGGGCTCGTGCCAGTGCCACGCGCTGTTGCTGGCCGCCGGACAGCTGGGTGGGATACTTGTGTGCCTGATCGCTGATGCCGACCCGGGCCAGCA
GGCGCTCGCCGGTCTCGGTGGCGGTGGATCGGTTCAAGCCCCGCACCTTCATGGGGGCCAGAATCACGTTGTCGAGCACGCTCAGGTGCGGAAAGAGGTT
GAACTGCTGGAAGACCATGCCGACCTCGGTACGGATCTTCTGCAGCGCCTGAGTGGCCTTGCCGTCGGGAAGCAGCGGATTGCCATCGACTTCCAGCTCG
CCTTGCTGGAACTCTTCCAGGCCGTTGACGCAGCGGATCAGCGTCGACTTGCCGGAACCGCTGGCGCCGATGATCACCACCACCTCTCCGGGCGTCACCG
TCAGGTCGATGTCCTGGAGGACGTGAAGACTGCCGAAATGCTTGTTGAGCTTGTTCAGGCGCACGATGGGCTCGGCGCCGGTCGTGGACTTCTCGTGGTT
CATGTCGTGCGTTCCTTTATTGTCCGACCAGGCCCTTGCGCTCGAGCAGGCGCAGCGCAAGCGAGAGCGTCAGGGTGATCGCCAGGTAGAGCATGGCGAC
CATCAGGTAAACCTCCAGGGCGGTGAAGGTGGTGGCGATGTAGATCTGCCCCTGGCGGACCAGCTCGCCGACGCCGATGACCGAGAACAACGAGGTGTCC

>Halomonas elongata DSM 2581 [NC_014532.2:1695..1765], dnaA> ig dnaN>, 70 bp

WebSkew GCmin pos. 67591 (GCmin<>oriC1 20.3 kb; GCmin<>oriC2 2.2 kb; oriC2<>oriC1 18.1 kb)

>Halomonas eurihalina DSM 5720 [NZ_JARWAH010000005.1:c47030..47529], <rpmH oriC1 dnaA>, 659 bp

CGGCGCTATCAGGGGTGGCGGCGCCGCTGGACGGAACCGACAGATCGCTACCGTTCGTATCCCCAGTATCCTGGGATGACGCTGGCGAGTTGCTGGTGAT
GCCCGGGGCCTCCGGCGCCTGAGTCGCCTGGCCGTAGTCCTGATTCCACTGTACGACGAGCAAGTAGGCGAGTATCGCCAGGGGAATCAACAGGAGTAGT
CGTTTTACGTCCATGAGGGGTCTGCCCAATGGGTGGTGAACATTCCGATGGCGCAGCGTGAGTCGCACACCCGACATACGCGAAAGCCTGCCATCGGCAG
GCTGGCTTTCATCAGCGGAGCGTCCGCTTTCGGGGGGGCGGCGTCAAGCTTCGCGACGAGTGGCGGAAACAGGCTCATCCGAACTCCGAGCGATGACGGC
GGCAACGGTACGCTGGACCGGTCATCGACGATCTGTCGAGGCGGTAGTGGAGCGCTCCGAGGCTTTCTGCGCTTCGCGCTCCAGACGTCGCCACATGCCA
TGTAATTGGCGGTGGACAACTTCATTGTCCAGCTCCTGTACTCCACGCCGGGCGAGCAATACGATGTCCACGGCGGGGAGTCGATGCTGGCGAAGCCGAA
TGGATTCGCGCGTGATTCGCTTGAGGCGATTACGGTCAACGGCTCGGCGGACGTTCTTCTTGCTGAAGATCAGGCCGAGACGAGGGTGACCCAACTCATT
GGGCGAGGCCAGGGCCATCAACCCCTTGCCGTGAAGCTTGAAAGCCGCCGTATCGAAGACACGTCGATAATCCTCGGCGGTCAGTAGGCGTAGCCGCCGA
GGAAAGCCCTGATAGGACACGCGTTTGTGTTCAGGCGCTCAGACGCTTGCGGCCCTTGGCGCGACGACGCGCCAGGACAGCACGTCCATTCTTGGTGGCC
ATGCGAGCACGGAAACCGTGTGCGCGCTTGCGCTTGAGGACGCTGGGCTGGAAGGTGCGTTTCATAACTAATTATTCCCACGTGGTTTTTGGAACGGATC

GATAACTTTTCAGAGGCGCCCGGTTTCCGTTGGAAACCCGGGGGATTCGGCTCAAGACCGGGCATTGTAGTGATTTCACGGCCTCGGTGCAATTGCCCTG
TTTGCCTAGAGTCGGCGATGTCCGGGCGTATTGGTGCCAGAATTATTACTAATTAAAAACATAGTTTGATATAGCTGTTATTATAGTTGGGGAGTTAATT
**TGTGGATAA**GTGGACATACACCATAAAAAACAATGGGATATATTCATAATTCTGGTGTGGACATCCTTTGGGATGGCAGTGTATCCACTGTGAGTATCCA
TTGCATGGAAGTGAACCTTGTCCACAACGCCAGGTGATCACCATATTGCTCCCGGTCATTTCCCGGCCTTTAGGACAGGGTTTTCCACAAAGCTCATTGA
AGTGCGACTTCATGTGGATAACCAGAGGCTTGGGCGATACAATGTGCGGTCGTTCTTGAATCTACGATGGTGAGGTCGCGTGTCGCTTGCTCTCTGGCAA

CAGTGTCTGGCTACCCTGCAGGATGAATTGAGCTCTCAGCAGTTCAACACCTGGATTCGCCCATTGCAGGCGGAAGAAGAGGGAGAGTCCAACCAGTTGC
GCCTGCTGGCGCCCAACCGGTTCGTGCGTGATTGGGTCAGCGACAAGTATGCCAAGCGTATCAGTGAACTGATGCGCGAGTTGGCCCCTGCCAAGCCGCC
CAAGGTGGTACTGACCGTGGGCAGCCGCCGAGCGGCTCCCGCTCCCCAGCCGCGTGAACTTGGTGATCCGGTGTCTGCGGCGCCTCGCTCGCCCGCCGTT
CATACCCCTCCGCGGGGCGATGTGGCGGATGAATCCGAAATCGATCAACTGCGAGAGGAAGGTGAGGCGCCGCGTCGCCCCACGGGCGAGCGTCAGGTCC
AGGTCGAGGGCAGTCTCAAGCACAACAGTGGACTGAACCATAACTTCACCTTCGAGACCTTTGTCGAGGGCAAGTCGAACCAGTTGGCGCGGGCTGCGTC
GCGACAGGTTTCCGAGAATCCCGGCGGCGCCTACAACCCGCTGTTTCTCTATGGCGGCGTCGGCCTGGGCAAGACCCACCTGATGCATGCCGTCGGCAAT
GCCCTGGATGCACGGCGCGAGAATGCACGGGTGGTGTATCTGCACTCCGAGCGCTTCGTCGCCGACATGGTCAAGGCGCTGCAGCTCAACGCCATCAATG
ACTTCAAGCGCTTCTATCGCAGTGTCGACGCTTTGCTGATCGATGATATCCAGTTCTTCGCCGGCAAGGAGCGCTCCCAGGAAGAGTTCTTCCATACCTT
CAATGCCTTGCTCGAAGGCGGTCAGCAGATGATCCTGACATCGGACCGTTATCCCAAGGAAATCAGTGGGGTAGAGGAGCGTCTCAAGTCGCGCTTCGGT
TGGGGGCTGACCGTGGCGATCGAGCCGCCGGAGCTCGAGACCCGGGTCGCGATCCTGATGAAGAAGGCCGACCAGGCCAAGGTCAATCTGCCGCACGACG

>Halomonas eurihalina DSM 5720 [NZ_JARWAH010000005.1:c65149..65648], <mnmG oriC2 <COG1126 (glnQ), 536 bp

TCCAGCCCCTGGATCGAACGGACGACCTGCAGTTGCACGTCAAACGGCAAGGAGGTCGAAATGCCGTTGGGATACAGCTCGTGCGTGTCCAGCCCCTCCG
GCTCGATGAAGATCTGGTGGCTCGACTTGTCGGCGAAGCGGTGCACCTTGTCCTCGATGGAGGGGCAATAGCGAGGCCCCACGCCTTCGATGGCTCCGGA
GAACATCGGCGAACGCTCGAGGTTGGCGCGGATGATCTCGTGGGTCCGCTCGTTGGTATGAGCGATGTGACAGCTCACCTGCCGCGGGTGCTGGCTCCGT
TCGCCAAGGTAGGACATGACCGGGGTGGGATCGTCACCGGGCTGCTCGTCGAGCTTCGAAAAGTCCACGCTCTTGGCATCGATACGTGGTGGGGTACCGG
TCTTCAGACGATCGACGCGGAACGGCAGGGCACGAAGACGTTCGGCCAGAGCATTGGAAGGTGGATCTCCGGCACGCCCCCCACGGCTGTTGTCCAGGCC
GATGTGGATAACACCACCCAGGAACGTGCCTGTGCACAACACCACGGTCTCGCTCATGAAGCGAATCCCCGTCTCGGTGCGCACACCGCGAACCTGGTCG
CCCTCGACAATGAGATCGCCGGCTGCCTGCTGGAAGAGAGTCAGGTTGGGCTGGTTTTCCAGCATGCCGCGGATGGCGGCCTTGTAGCGAATGCGGTCCG
CCTGGGCCCGAGTCGCGCGTACGGCCGGCCCTTTCCGGGCGTTGAGCACACGGAACTGGATACCGCCCATGTCGGTGGCCAGCCCCATGGCGCCACCAAG
GGCATCGATTTCCTTGACCAAATGGCTCTTGCCGATCCCGCCAATGGCGGGATTGCACGACATCTGGCCGAGAGTCTCGATGTTGTGGGTCAAAAGCAGC
GTCCGGCAGCCCGTACGGGCCGAGGCCAGGGCGGCTTCGGTGCCCGCATGACCACCACCGATGACAATGACGTCGAAACGGTCGGGGTACTCCAAGGTTC

ACCTCGTTCGGTGCTACGGCACCGATATTGCGGATCGGGGATGAAATCGGCCAGCCAGTATAAACCTCCTGTGGGTAATCGTCAGGGTTTTCCACACCCG
GATTTTCATTCGCAGTCCGTCTCGCATCATTAGCTATTACATAACTAATAAAAGAATAGGAAGTTCTGTTGTTGTTGTAACTACTGGTGTGGATGTTTTT
**CGTGGATAA**ACCGTTTTATTGTTTTACTATCAATAGCTTGCAATGTTAACCGAAAGTAGGAAAAAGCGCGTACTGGCTGCGGACGATGTGTTCGGGGCCT
GTGGATGAAAGCTCGGGTTGCCCACAAGCTGACACAGACACGCCTTTTCCACCGATCCCTACCTGGCTTGTCACCACGGCTGTGAACAACCGAGCCAAAT
TGCCCACATCCTTGCAATGGCGGCGGTTGTGGCATGATTCAGGCGCTTCGGGCGATTTAGTTTTCCACAGGCAGAAAAAAGGCCCTGTCGATAACGACAG

GGCCCCTGAAGGCACGCTTGAAACGGCGCTATCAGTGCTTGAGCACTCGGGACAGGAAGCTTTGTGTCGGCGCCTCCGTGGGGGTATCGAAGATTGTTTC
GGGTGGGCCTTCTTCGACGATGCGGCCTTTGTCGATGAAAATGACCCGATCGGCGACCTCACGGGCAAAACCCATCTCGTGGGTCACGATCATCATGGTC
ATGCCGTCGTTGGCCAGTTCTCGCATGGCATCAAGTACCTCGCCGATCATCTCCGGGTCGAGTGCCGAGGTGGGCTCGTCGAAGAGCATCAGGCGGGGTT
CCATGGCCAGGGCCCGTGCCAGCGCCACACGCTGCTGCTGGCCGCCGGACAGCTGGGCCGGATACTTGTGCGCCTGATCGCTGATACCGACCCGAGCCAG
CAGGCGCTCGCCGGTCTCGGTAGCGGTGGATCGGTTCAAGCCCCGCACCTTCATGGGTGCCAGAATCACGTTGTCGAGCACGCTCAGATGCGGGAATAGG
TTGAACTGCTGGAAGACCATGCCGACCTCGGTACGGATCTTCTGCAGCGCCTGAGTGGCCTTGCCGTTGGGCAGGAGCGGGTTGCTGTCGACTTCCAGTT
CGCCCTGCTGGAACTCTTCCAGGCCATTGACGCAGCGGATCAGTGTCGACTTGCCGGAACCGCTGGCACCGATGATCACCACCACCTCGCCGGGCGTCAC
CGTCAGGTCGATGTCCTGGAGGACGTGAAGACTGCCGAAATGCTTGTTGAGCTTGTTCAGGCGCACGATGGGCTCGGCGCCGGACGCGGACTTCTCGTGG
TTCATGTCGTGCGTTCCTTCTATTGTCCGACCAGGCCCTTGCGCTCGAGCAGACGCAGCGCAAGCGAGAGTGTCAGGGTGATCACCAAGTAGAGCATGGC
GACCATCAGGTACACCTCCAGGGCGGTGAAGGTGGTGGCGATGTAGATCTGCCCCTGGCGGACCAGCTCGCCGACGCCGATGACCGAGAACAACGAGGTA

> Halomonas eurihalina DSM 5720 [NZ_JARWAH010000005.1:c45512..45588], dnaA> ig dnaN>, 75 bp

WebSkew GCmin pos. 146640 (GCmin<>oriC1 20.3 kb; GCmin<>oriC2 1.2 kb; oriC2<>oriC1 19.2 kb)

>Halomonas caseinilytica JCM 14802 [NZ_BDEP01000014.1:c126044..126543], <rpmH oriC1 dnaA>, 514 bp

TCGCTGCCCATCTCGGCGTTGCCGGTGCCGCCGGACGGCACCGACAGGTTGCCACTGTCAGCCTCCCCAGTCTCCCGGGACGAAGCTGGCGAGTCGTCGG
TGATGGCCGGGGCCTCCGGCGCTTGGGTCGCCTGGCCGTAGTCCTGATTCCACTGCACGACGATCAGGTAGGCGAGTATCGCCAGGGGAATCAGCAGGAG
TAGTCGTTTTACGTCCATGAGGGGTCTGCCCGATGGGTGGTGAACATTCCGATGGCGCGGCGTGAGGTGCTCTCCCGACATACGCGAAAGCCTGCCAAGG
GCAGGCTGGCATTCGTCAGCGGAGCGTCCGCTTTCGAGGGGGCGGCGTCAAGCTCCGCGGCAAGAGGCGAAATCAGGCGCATGCAAGCTCCGGGCGATGA
CGGCGGCGAGGCGCCGCATCGGTCATCGGCGGTCTGTCGAGGGGCTGGTGGAGCCGCCCGTCGATTTTTGCGCTTCGCGCTCCAGGCGTCGCCACATGCC
ATGCAATTGGCGATGGACAACCTCATTGTCCAGCTCCTGTACGCCACGCCGGGCGAGCAAAACGATATCCACGGCGGGGAGTCGATGCTGGCGAAGCCGA
ACGGATTCGCGCGTGATTCGCTTGAGGCGATTGCGATCAACGGCCCGACGAACGTTCTTCTTGCTGAAGATGAGGCCGAGTCGAGGGTGACCCAATGCGT
TGGGCGAGGCCAGGGCCATCAACCCCTTGCCGTGAAGCTTGAAAGCTGCCGTATCGAAGACACGTCGATAGTCCTCGGCGGTCAGCAGGCGCAGCCGCCG
AGGAAAGCCCTGATAGGACACGCGCTCGCGATCAGGCGCTCAAGCGCTTGCGGCCCTTGGCGCGACGGCGCGCCAGGACGGCACGACCATTCTTGGTGGC
CATGCGAGCGCGGAAGCCGTGTGCGCGCTTGCGCTTGAGAACGCTGGGCTGGAAAGTGCGTTTCATAACTGTTTATTCCCACGTGGTTTTTGGAACGGAT

CGATAATTTCAGAGGCACCCGGTTTCCATCGGAAACCCGGGGAGATTCGGCTCAAGACCGGGAATTGTAGTGATTTCGCGGCCTCGGCGCAATTCCCTCT
TGACTCACGGGCAACGCACGACGCGAGGGGCTCATGCCTCAATCATTATTATTTAGAACGATAGATTGATGAGGTTGTTATTATAGCTGGGGAGAAATTT
**TGTGGATAA**GTATATTTATATTTTTTATATCAAATGGATACGGTATTGATCGTGGTGTGGACATCCTTGGGGGTGATGGTGTATCCACTGTGGCTTTCCG
TTGCATGGAAGCTGCAATTGTCCACAGTGCTGATGCATCCCCATATCGCTCCCCGTCATTTCCTGACGTGGGGGAAGAGGTTTTCCACAAAGTTCATCGA
AACGCGGCTTCATGTGGATAACCCGAGGGCTGGACGATACAATGTGCGGTCGTTCTTGAAACTATGATTGGTGAGGTTGCGTGTCGCTCGCTCTCTGGCA

ACAGTGTCTGGCTACCCTGCAGGATGAATTGAGCTCCCAGCAGTTCAACACCTGGATCCGCCCATTGCAGGCGGAGGAGGAGGGAGAGTCCAACCAGTTG
CGCCTGTTGGCGCCTAACCGGTTCGTGCGCGATTGGGTCAGTGACAAGTATGCCAAGCGCATCAGTGAGCTGATGCGCGAGTTGGCGCCTGCCAAGCCCC
CAAAGGTGGTGCTGACCGTGGGCAGTCGCCGAGCGGCTCCTGCTCCCCAGCCGCGGGAACTCGGCGATCCGGTCTCGGCGGCGCCTCGAGCGCCTGCCGT
GCACACGCCACCACGGGGCGATGTGGCGGATGAATCCGAGATCGACCAGTTGAGGGAAGAAGGGGAGGCATCGCGTCGTTCCTCGGGCGAACGCCAGGTA
CAGGTCGAGGGCAGCCTCAAACACAACAGTGGCCTGAATCATAATTTCACCTTCGAAACCTTCGTCGAGGGCAAGTCCAACCAATTGGCACGGGCTGCTT
CGCGGCAGGTCTCCGAGAACCCCGGAGGCGCTTATAATCCGTTGTTTCTCTATGGTGGGGTCGGCCTGGGCAAGACTCACCTGATGCATGCCGTCGGCAA
CGCCCTGGATGTGCGTCGCGAGAACGCTCGGGTGGTGTATCTGCATTCCGAGCGTTTCGTCGCCGATATGGTGAAGGCGCTGCAACTCAACGCCATCAAC
GATTTCAAGCGCTTCTACCGCAGTGTCGATGCCTTGCTGATCGACGATATCCAGTTCTTCGCCGGCAAGGAGCGTTCCCAGGAAGAATTCTTCCATACCT
TCAATGCCTTGCTTGAAGGTGGTCAGCAAATGATCCTGACCTCGGATCGTTATCCCAAGGAAATCAGCGGGGTAGAGGAGCGTCTCAAGTCGCGCTTCGG
TTGGGGGCTGACCGTGGCGATCGAGCCTCCGGAGCTCGAGACCCGGGTAGCGATCCTGATGAAGAAGGCCGACCAGGCCAAGGTCAATCTGCCGCATGAT

>Halomonas caseinilytica JCM 14802 [NZ_BDEP01000014.1:c145212..145711], <mnmG oriC2 <COG1126 (glnQ), 536 bp

TCCAGCCCTCGGATCGAGCGGACCACCTGCAGCTGGACGTCGAACGGCAAGGACGTGGAGATGCCATTGGGATACAACTCGTGCGTGTCCAGTCCCTCCG
GCTCGACGAAGATCTGGTGGCTTGCCTTGTCGGCGAAGCGATGCACCTTGTCCTCGATGGAGGGGCAGTAGCGGGGACCGACACCCTCGATGGCCCCGGA
GAACATCGGCGAACGCTCCAGATTGGCGTGGATGATCTCGTGGGTCCGCTCGTTGGTGTGCGCGATATGGCAGCTCACCTGGCGCGGATGCTGGCTGCGT
TCGCCGAGATAGGACATGACCGGCGTTGGCTCATCACCAGGCTGCTCGTCGAGCTTCGAAAAGTCCACGCTCTTGGCATCGAGACGCGGTGGGGTGCCGG
TCTTCAGGCGATCGACGCGAAACGGCAAGGCACGCAAGCGCTCGGCCAGGGCATTGGAGGGCGGATCGCCAGCACGGCCTCCACGACTGTTGTCCAGGCC
GATGTGGATAACGCCGCCGAGGAAGGTGCCTGTGCACAACACCACGGTCTCGCTCATGAAGCGAATCCCCGTTTCGGTGCATACACCGCGAACCTGGTCG
CCCTCGACGATGAGATCGCCGGCTGCCTGCTGGAAGAGCGTCAGGTTGGGCTGGTTTTCCAGCATGCCGCGAATGGCGGCCTTGTAGCGAACGCGATCCG
CCTGGGCACGAGTCGCGCGCACGGCTGGTCCCTTTCGAGCGTTGAGCACACGAAACTGGATACCGCCCATGTCGGTGGCCAGCCCCATGGCGCCCCCCAG
GGCATCGATTTCCTTGACCAGGTGGCTCTTGCCGATCCCGCCAATGGCGGGATTGCACGACATCTGACCGAGGGTCTCGATGTTGTGGGTCAGAAGCAGT
GTCCGGCAGCCCGTGCGGGCCGAGGCCAATGCGGCTTCGGTGCCCGCGTGACCACCACCGATGACGATGACATCGAAACGGTCGGGGTACTCCAAGGTTC

ACCTCGTTCGGCGCCAGGCGCCGATATTGCGGATCGGGGATGAAATCAGCCAGCCAGTATAACCCTCCTGTGGGTAATCGTCAGGGTTTTCCACACCCAG
ATTTTCATCCTCGCCCTTCTGCTGAGTCATTACCTATTACATAAACAATATAAGAATAAGAAGTTCTGTTGCTGTAGTAACTAACTGGTGTGGACGTTTTC
**CGTGGATAA**ACCGTTTTATTTCTTTTCTTTCAATAGATTGCAGTGTTGATCGAAGGTAGGAAAAGACGCTGGCGGCCTGCGGACGATGTGTTCGGGGCCT
GTGGATGAAAGATCGTGTTGCCCACAGGCGAGCGCGAGCACGGCTTTTCCACCGATGTCTACCGTGATTGTCACCGTGACTGTGAACAACCCACCCTATT
TTTCCACATCCCGCCAATGGCGGGCGTCGTGGCGTGATAAAAGCGCTTGTGCCGCTTGGGTTTTCCACAGGCAGAAAAAAGGCCCTGTCGGTAACGACAG

GGCCTTTCAAGGTATGCTGAGAGAGGTGATATCAGTGCTTGAGCACGCGAGACAGGAAGCTTTGCGTGGGCGCTTCAGAGGGGGAATCGAAGATCGTCTC
GGGCGGGCCTTCCTCGACGATCCGACCCTTGTCGATGAAGATGACCCGATCGGCCACCTCGCGGGCAAACCCCATCTCGTGGGTCACGATCATCATGGTC
ATGCCCTCGTTGGCTAGCTCGCGCATGGCATCGAGCACTTCACCGATCATTTCCGGGTCCAGCGCCGAGGTGGGCTCATCGAAGAGCATCAGCCGAGGTT
CCATGGCCAGGGCGCGTGCTAGTGCCACCCGCTGCTGCTGGCCGCCGGACAGCTGTGTGGGGTACTTGTCTATCTGGTCGCTGATGCCGACCCGAGCCAG
CAGGCGTTCCCCTGTCTCGACGGCGGCATCACGGCTCAGGCCACGGACCTTCATCGGCGCCAGGATCACGTTGTCGAGCACGCTCAGGTGCGGGAACAGG
TTGAATTGCTGGAAGACCATGCCGACTTCGGTACGGATCTTCTGCAGCGCCTGAGTTGCCTTGCCGTCGGGCAGCAGGGGATTGCCATCGACGTCCAGTT
CGCCTTGCTGGAACTCCTCGAGGCCATTGATGCAGCGGATCAGGGTCGATTTGCCGGAGCCGCTGGCACCGATGATCACTACCACCTCACCTGGCGTTAC
CGTCAGGTCGATGTCCTGAAGAACGTGAAAGTCGCCGAAGTGCTTGTTGAGCTTGTTCAGCCGCACGATGGCCTCGCCATCGGACGTGGACTTCTCGTGA
TTCATGTCGTGCATTCCTTTCATTGTCCGACCAGGCCCTTGCGCTCGAGCAAGCGCAACGCAAGCGAGAGCGTCAGGGTGATCGCCAGGTAGAGCATGGC
GACCATCAGGTAGACCTCCAGGGCGGTGAAGGTGGTGGCGATGTAGATCTGTCCCTGGCGGACCAGTTCACCGACGCCGATGACTGAGAACAGCGAAGTG

>Halomonas caseinilytica JCM 14802 [NZ_BDEP01000014.1:c124526..124599], dnaA> ig dnaN>, 74 bp

WebSkew GCmin pos. 110798 (2nd min.) (GCmin<>oriC1 20.9 kb; GCmin<>oriC2 1.7 kb; oriC2<>oriC1 19.2 kb)

>Halomonas halmophila NBRC 15537 [NZ_BJOC01000021.1:c89645..90144], <rpmH oriC1 dnaA>, 515 bp

CATCTGATTGTCCATGTCCGCACTGGAAGGGCTGCTGGCAGGAACCGAGAGGTCACTGCTCTTTTCGGTGCCCTCGGGCGCTGTCGAGCTCTCGCTCGCA
ATGCCCGGGGCTTTCGGCGCCTGTGTGGCCTGGCCATAGTCCTGGTTCCACTGCACTACGATCAAGTAGGCGAGCACTGCCAGGGGAATCAGCAGGAGGA
GTCGTTTTACGTCCATGAGGGGTCTGCCCGATGGGTGGTGAACATTCCGATGGCGTAGCGTGATTCGTACACCCTGCACAAACGAAAGCCCGCCAATGGC
AGGCCGGCTCTCGTCAGAGGAGCGTCCGCTTTCGGCGGGGTGACGTCAAGCTTGGCGGCAAGCGGAGAAAACAAGCTCATCAGAGCTCCAGACGATGATG
ACAGCCGCAGGGTGGATGCTTCATCGATGCTCTGTCGAGGCGTCAGAGGAGCGACGTGAGGCCTTCTGCGCTTCACGCTCAAGGCGACGCCACATGCCAT
GTAACTGGCGGTGGACAACATCATTGTCCAGCTCCTGTACGCCACGTCGGGCGAGCAGCACGATATCCACGGCGGGAAGTCGGTGCTGTCGATGCCGTAT
GGATTCACGCGTGATTCGCTTGAGACGATTGCGATCAACGGCCCGACGGACATTTTTCTTGCTGAAGATCAGGCCAATACGCGGGTGGCCCAGGGTATTG
GGTGTGGCCAGCGCCATCATCCCTTTGCCGTGCAGCTTGAAGGCTGCCGAATCGAAGACACGTCGATAGTCCTCGGCGGTCAGCAGTCGCAGCCGCCGAG
GAAAGCCCTGACAGGACACGTGTTCTCGATCAAGCGCTTAGACGCTTGCGGCCCTTGGCGCGACGGCGCGCCAGGACGGCACGACCATTCTTGGTGGCCA
TGCGAGCGCGGAAACCATGCATGCGCTTGCGCTTGAGAACGCTGGGTTGAAAGGTGCGTTTCATAACTGATTATTCCCACGTGGTGTTGGTACGGAGTCG

AAAACTGTTTGAGGCGCTCGGCTGCTGCATCGGCGCCAAGCAAAACCGGCTCAAGACCGGGCATTGTAGTGATTTCCAGGCGCTGGCGCAATTCCTGCTC
GACCTGGGCCCGACAAGCCTCGTTTCTCGGGTCGCCATGACAATCATTATTACAAGAAGAAAGGATAACTGATTCTGTTATTATTGTTGGGGAGAAAAAT
**TGTGGAAAA**CCTGGATTAAAGTCTTATCTTCAAATAGTTAGCTTTTTATTGCTGCTGTGCACAACCTTGGAGAAAAGCGGGAGCTGGCTGTGAATAGCCT
CTGCAAGGATTCCAGAACTGTACACAGGGGCCGATGCTGCCACCTGTTGTGCGCCGCTTGACACTCAGCCTGAAAACAGGGTTTTCCACATTGTCAAATC
AGGCATGCTGTCATGTGGATAACCCGCAGGTTGAGCGCTACAATGTTCGGCCGTTCTTGAATCTACGATGTGAGGTTGTGTGTCACTCGCTCTCTGGCAA

CAGTGTCTGGCTACCCTTCAGGATGAGTTGAGCTCCCAGCAGTTCAATACCTGGATCCGCCCCTTGCAGGCGGAAGAGGGGGAGGCCAACCAGCTGCGCC
TGCTGGCGCCGAACCGCTTTGTGCGCGACTGGGTCAGTGACAAGTACGCCAAGCGCATCAGCGAACTGATGCGCGAGCTGGCGCCTGCCAAGCCGCCCAA
GGTCGTGCTGGTGGTGGGCAGCCGTCGTGCCGCTCCTGCCCCGCAACCGAGAGAGCTCGGTGACCCTGTTTCGGCAACGCCCCAGGCGCCCAATGTGCAC
ACGCCACCGCGCGGAGATGTGGCGGATGAATCGGAGATCGACCAGGCCGGGAATGAGGCCGAGGAAGCCCGTGCCCCACGTCGGACCTCAGGAGAGCGTC
AGGTGCAGGTCGAGGGCAGCCTCAAGCACAACAGCGGGCTGAATCAGAACTTTACCTTCGACACCTTCGTCGAGGGCAAGTCGAATCAACTGGCCAGGGC
CGCCTCCCGCCAGGTTTCCGAGAACCCCGGGGGCGCCTACAATCCGCTGTTCCTGTATGGCGGAGTCGGCCTCGGCAAGACGCACCTGATGCATGCGGTG
GGCAATGCGCTGGACATGCGGCGCGAGAATGCGCGCGTGGTCTACCTGCACTCGGAACGCTTCGTGGCCGACATGGTCAAGGCACTGCAGCTCAACGCCA
TCAATGACTTCAAGCGCTTCTACCGCAGTGTCGATGCACTCCTGATCGACGACATCCAGTTCTTTGCCGGCAAGGAGCGCTCCCAGGAAGAGTTCTTCCA
CACCTTCAACGCCTTGCTCGAGGGCGGTCAGCAGATGATCATGACCTCAGACCGCTATCCCAAGGAAATCAGTGGGGTAGAGGAACGTCTGAAGTCGCGC
TTCGGCTGGGGGCTGACCGTGGCGATCGAGCCGCCGGAGCTCGAGACGCGGGTTGCCATCCTGATGAAGAAGGCCGATCAGGCCAAGGTCAACCTGCCCC

>Halomonas halmophila NBRC 15537 [NZ_BJOC01000021.1:c108858..109357], <mnmG oriC2 <COG1126 (glnQ), 582 bp

CCATGCCGCGGATGGAGCGCACGACCTGCAGCTGCACATCGAAGGGAAGCGACGTGGAGATGCCGTTGGGATACAGCTCATGGGTCTCCAGGCCCTCCGG
CTCGACGAAGATCTGATGGCTGTTCTTGTCCGCGAAGCGATGCACCTTGTCCTCGATGGACGGGCAATAGCGCGGTCCCACGCTGTCGATGGTGCCGGAA
AACATCGGCGACCGATCCAGGTTGGCGTGAATGATCTCGTGGGTGCGCTCGTTGGTGTGCGCGATGTGGCAGCTCACCTGACGCGGATGCAGATCCCGAC
CGCCCATGTAGGACATGACCGGGGTCGGGTCATCACCCGGTTGGGCATCGAGCTGCGAGAAATCGACGCTCTTGGCATCCAGCCGCGGGGGCGTCCCGGT
CTTGAGTCGATCGACGCTGAACGGCTGGGCCCGCAGACGCTCTGCCAGCCCATTGGAGGGCGGATCGCCGGCTCGACCGCCGCGGCTGTTGTCCATCCCG
ATGTGGATAACTCCGCCGAGGAAGGTGCCGGTACACAAGACCACGCTTTCGCTGTGGAAGCGGATTCCGGTTTCGGTGGTGACGCCGCGTACCTGGTTGC
CGTCGACGATCAGGTCGCCGGCAGCCTGCTGAAAGATTGTCAGATTGGGCTGGTTTTCCAGCATGCCGCGGATGGCGGCCTTGTACAGCACGCGGTCCGC
CTGGGCACGGGTCGCCCGCACGGCGGGGCCCTTGCGGGCATTGAGCACGCGAAACTGGATGCCGGCCATGTCGGTGGCCAGGCCCATGGCACCGCCCAGG
GCATCGATCTCCTTGACCAGATGACTCTTGCCGATCCCGCCGATGGCGGGATTGCACGACATCTGACCGAGGGTCTCGATGTTGTGGGTCAGTAGCAGCG
TACGGCAGCCCGTGCGAGCCGAGGCCAGAGCGGCTTCGGTGCCCGCATGGCCACCACCGATGACGATGACGTCGAAACGGTCGGGGTAATCCAAGGTTCA

CCTCGTTCGGCGCATCGGCGCCGATATTGCGAATCGGGGATGAAATCAGCCAGCCAGTATAAACCTCCTGTGGGTAATCGTCAGGGTTTTCCACACCTAG
ATTTTGGTCCGCGCCCTTTCACACACCCTTAGACAACAATATAAATAATATATAGATAAGAAGTTCTGTTGTTGTTGTTATTACTGGTGTGGAGCTTTTC
**TGTTGATAA**TCGGTCATAAGCCTTTTACATCAACAGCATGCAGTGTTGACTGATATACGCAAAGCTTCCGGGCGAGCTGCGCACGAGCTGTTGGAGGGCT
GTGGATGAAACCTCGACTTGCCCACAAGCCGGAAAAATCAGCCCTTTTCCACCGGTCTCTACCCGAGTTGTCACCGCGGCTGTGAACAACCTGCCTTCTT
GTTCAGCATTACCGGCGTGGCAGTGCCTGCAAGCGAATTTCGGGGTCATCGTCCGGTTGGTTATCCACAGGCAGAAAAAAGGCCCTGTCGATAACGACAG

AGCCTTGTGGAAGTGCCTGATTGGCACTCGATGAGATCCTGGAAGGCGGTAAATGCCTCCCGCACGTGCGCGACGGTCAGTGCTTGAGCACCCTCGAAAG
GAAACTCTGGGTGGCAGCCTCGACAGGCTGATCGAAGATCGTCTCGGGCGGGCCTTCCTCGACGATGCGGCCCTTGTCGATGAAGATCACGCGGTCGGCG
ACCTCGCGGGCAAAGCCCATTTCGTGGGTAACGATCATCATGGTCATGCCGTCGTTGGCCAGCTCGCGCATGGCATCCAGGACTTCACCGATCATCTCGG
GATCCAGCGCCGAGGTGGGCTCATCGAAGAGCATCAGGCGCGGTTCCATGGCCAGGGCGCGGGCCAGGGCAACACGCTGTTGTTGACCGCCGGAGAGCTG
GGTGGGGTATTTGTGGATCTGGTCGCTGATACCCACGCGTTCGAGCAGGCGTTCACCGGTCGCCACCGCCGTTTCGCGGTCCTGACCACGCACCTTCATG
GGCGCCAGAATCACGTTGTCGAGCACGCTGAGGTGCGGAAAGAGGTTGAATTGCTGGAAGACCATGCCAACTTCGGTGCGGATCTTCTGCAGGGCCTGAG
TGGCCTTGCCGTAGGGCAGCAAGGGGCTGTCGTCGACATCCAGTTCGCCCTGCTGGAATTCTTCCAGGCCGTTGACGCAGCGGATCAGTGTCGATTTGCC
GGAACCACTGGCGCCGATGACCACGACGACCTCGCCGGTGGCGACCCTAAGGTCGATATCCTGGAGCACGTGAAAGTCGCCGAAATGCTTGTTGAGCTTG
TTCATGCGCACGATGGGCTCGGCGCCTGCCTTGGACTTCTGCTGGTTCATGAGGTGCCTTCCTTATTGTCCGACGAGGCCCTTGCGCTCGAGCATGCGCA
GCGCAAAAGATAGGGTCAGGGTGATCGCCAGGTACAGCAGGGCAACCATCAGGTAGACCTCCAGGGCGGTGAAGGTGGTGGCGATGTAGATCTGTCCCTG

>Halomonas halmophila NBRC 15537 [NZ_BJOC01000021.1:c87982..88195], dnaA> ig dnaN>, 214 bp

< page 1 | #005 | Halomonas sabkhae CECT 7246

WebSkew GCmin pos. 207900 (2nd min.) (GCmin<>oriC1 22.9 kb; GCmin<>oriC2 3.4 kb; oriC2<>oriC1 19.4 kb)

>Halomonas sabkhae CECT 7246 [NZ_JAUFPM010000008.1:230549..231048], <rpmH oriC1 dnaA , 505 bp

CTGACTTTCCATGTCCGCACTGGAAGGGCTGCTGGCAGGGACCGAGAGGTCATTGCCCTGTTCTTCGCTGCCCTCGGACGCTGTCGAGCTCTCGCTCGCA
ATGCCCGGAGCTTCCGGCGCCTGTGTGGCCTGGCCATAGTCCTGATTCCACTGCACTACGATCAAGTAGGCGAGCACTGCCAGGGGAATCAGCAGGAGGA
GTCGTTTTACGTCCATGAGGGGTCTGCCCGATGGGTGGTGAACATTCCGATGGCGTAGCGTGAGTCGTACACCCTGCACAAACGAAAGCCCGCCAATGGC
AGGCCGGCTCTCGTCAGAGGAGCGTCCGCTTTCGGCGGGGTGACGTCAAGCTTGGCGGCAAGCGGAGAAAACAAGCTCATCAGAGCTCCAGGCGATGATG
ACAGCCGCAGGGTGGATGCTTCATCGATGCTCTGTCGAGGCGTCAGAGGAGTGACGTGAGGCCTTCTGCGCTTCACGCTCGAGGCGACGCCACATGCCAT
GTAACTGGCGGTGGACAACATCATTGTCCAGCTCCTGTACGCCACGTCGGGCGAGCAGCACGATATCCACGGCGGGAAGTCGGTGCTGTCGATGCCGTAT
GGATTCACGCGTGATTCGCTTGAGGCGATTGCGATCAACGGCCCGACGGACATTTTTCTTGCTGAAGATCAGGCCAATACGCGGGTGGCCCAGGGTATTG
GGTGTGGCCAGCGCCATCATCCCTTTGCCGTGCAGCTTGAAGGCTGCCGAATCGAAGACACGTCGATAGTCCTCGGCGGTCAGCAGTCGCAGCCGCCGAG
GAAAGCCCTGACAGGACACGTGTTCTCGATCAAGCGCTTAGACGCTTGCGGCCCTTGGCGCGACGACGCGCCAGGACGGCACGACCATTCTTGGTGGCCA
TGCGAGCGCGGAAACCATGCATGCGCTTGCGCTTGAGAACGCTGGGTTGAAAGGTGCGTTTCATAACTGATTATTCCCACGTGGTGTTGGTACGGAGTCG

AAAGCTGTCTGAGGCGCTCGGCTGCTGCATTGGCGCCGAGCAAAACCGGCTCAAGACCGGGCATTGTAGTGATTTCCAGACGCTGGCGCAATTCCTGCTC
GACCTGAGCCCGACAAGCCTCGTTTCTCGGGTTGCCATGACAATCATTATTACAAGAAGAGAGGATAACTGATGCTGTTATTATTGTTGGGGAGAAAAAT
**TGTGGAAAA**CCCGGTTTAAAACATTATCTTCAAATAGTTAGCTTTTTATTGCTGCTGTGCACAACCTTGGAGGAAAGTGGGCGTTGGCTGTGAATGGCCT
CTGCAGGGATTCCTTAGCTGTACACAGGGGCCGATGCTGCCACCTGTTGTGCGCCGCTTGACACCCAGCCTGAAAACAGGGTTTTCCACATTACCAAATC
AGACATGCTGTCATGTGGATAACCCGCAGGTTGAGCGCTACAATGTTCGGCCGTTCTTGAATCTACGATGTGAGGTCGTGTGTCACTCGCTCTCTGGCAA

CAGTGTCTGGCTACCCTTCAGGATGAGTTGAGCTCCCAGCAGTTCAATACCTGGATCCGCCCCCTGCAGGCGGAAGAGGGGGAGGCCAACCAGCTTCGCC
TGCTGGCGCCGAACCGCTTTGTGCGCGATTGGGTCAGTGACAAGTACGCCAAGCGCATCAGCGAACTGATGCGCGAGCTGGCGCCTGCCAAACCACCCAA
GGTCGTGCTGGTAGTGGGCAGCCGTCGTGCCGCTCCTGCCCCGCAACCCAGAGAGCTCGGCGATCCCGTCTCGGCAACGCCTCAGGCGCCCAATGTGCAC
ACGCCGCCGCGCGGTGATGTGGCGGACGAGTCGGAGATCGATCAGGCCGGGGATGAGGCCGAGGAAGCCCGCGCTCCGCGTCGGCCCTCGGGAGAGCGCC
AGGTGCAGGTCGAGGGTAGCCTCAAGCACAACAGCGGGCTGAATCAGAACTTTACCTTCGACACCTTCGTCGAGGGCAAGTCGAACCAGCTGGCCAGGGC
CGCCTCCCGCCAGGTATCCGAGAACCCCGGGGGCGCCTACAACCCGCTGTTCCTGTATGGCGGGGTAGGCCTCGGCAAGACTCACCTGATGCATGCGGTG
GGCAATGCGCTGGACATGCGGCGCGAGAATGCGCGCGTGGTCTACCTGCACTCGGAACGCTTCGTGGCCGATATGGTCAAGGCGCTGCAGCTCAACGCCA
TCAATGACTTCAAGCGCTTCTATCGCAGTGTCGATGCACTCCTGATCGATGACATCCAGTTCTTTGCCGGCAAGGAGCGCTCCCAGGAAGAGTTCTTCCA
CACCTTCAACGCCTTGCTCGAGGGTGGCCAGCAGATGATCCTGACCTCGGACCGTTATCCCAAGGAAATCAGTGGGGTAGAGGAACGTCTGAAGTCGCGC
TTCGGGTGGGGGCTGACCGTGGCGATCGAACCGCCGGAGCTCGAGACGCGGGTTGCCATCCTGATGAAGAAGGCCGATCAGGCCAAGGTCAACCTGCCCC

>Halomonas sabkhae CECT 7246 [NZ_JAUFPM010000008.1:211110..211609], <mnmG oriC2 <COG1126 (glnQ), 583 bp

CCATGCCGCGGATGGAACGCACGACCTGCAGCTGCACATCGAAGGGAAGCGACGTGGAGATGCCGTTGGGATACAGCTCATGGGTCTCCAGGCCCTCCGG
CTCGACGAAGATCTGATGGCTGTTCTTGTCCGCGAAGCGATGCACCTTGTCCTCGATGGACGGGCAATAGCGCGGTCCCACGCTGTCGATGGTGCCGGAA
AACATCGGCGACCGATCCAGGTTAGCGTGAATGATCTCGTGGGTGCGCTCGTTGGTGTGCGCGATGTGGCAACTCACCTGACGCGGATGCAGCTCCCGAT
CGCCCATGTACGACATGACCGGGGTCGGGTCATCACCCGGTTGTGCATCGAGCTGCGAGAAATCGACGCTCTTGGCATCCAGCCGCGGGGGAGTCCCGGT
CTTGAGTCGATCGACGCTGAACGGCTGGGCCCGCAGACGCTCTGCCAGCCCATTGGAGGGCGGATCGCCGGCTCGACCGCCGCGGCTGTTGTCCATCCCG
ATGTGGATAACTCCGCCGAGGAAGGTGCCGGTACACAAGACCACGCTTTCGCTGTGGAAGCGGATTCCGGTTTCGGTGGTGACGCCGCGTACCTGGTTGC
CGTCGACGATCAGGTCGCCGGCGGCCTGCTGAAAGATCGTCAGATTGGGCTGGTTCTCCAGCATGCCGCGGATGGCAGCCTTGTACAGCACGCGGTCCGC
CTGGGCACGGGTCGCCCGCACGGCTGGGCCCTTGCGCGCATTGAGCACGCGAAACTGGATGCCGGCCATGTCGGTGGCCAGGCCCATGGCACCGCCCAGG
GCATCGATCTCCTTGACCAGATGACTCTTGCCGATCCCGCCGATGGCAGGATTGCACGACATCTGACCGAGGGTCTCGATGTTGTGGGTCAGAAGCAGCG
TACGGCAGCCCGTGCGAGCCGAGGCCAGAGCGGCTTCGGTGCCCGCATGGCCACCACCGATGACGATGACGTCGAAACGGTCGGGGTAATCCAAGGTTCA

CCTCGTTCGGCGCATCGGCGCCGATATTGCGAATCGGGGATGAAATCAGCCAGCCAGTATAAACCTCCTGTGGGTAATCGTCAGGGTTTTCCACACCTGG
ATTTTGGTCCGCGTCCTTTCACACACCCTCAGACAACAATATAAATAATATATATAGATAAGAAGTTCTGTTGTTGTTGTTATTACTGGTGTGGAGCTTTTC
**TGTTGATAA**GCGATCATAAACCTTTTACATCAACAACATGTAGTGTTGACTGATACACGCAAAGCTTCCGGGCGAGCTGCGCACGAGCTGTTGGAGGTCT
GTGGATGAAACCTCGACTTGCCCACAAGCCGGAAAAATCAGCCCTTTTCCACCGACCACTACCCGAGTTGTCACCGCGGCTGTGAACAACCTGCCTCCTT
GTTCAGCATTACCGGCGTGGCAGTGCCTGCGAGCGAATTTCGGGGGTCATCGTCCGGCTGGTTATCCACAGGCAGAAAAAAGGCCCTGTCGATAACGACA

GAGCCTTGAGAAAGTGCCTGATTGGCACTCGATGAGATCCTGGAAGGCGATCAATGCCTCCCGCAAATGCGCGACGGTCAGTGCTTGAGCACCCTCGAAA
GGAAACTCTGGGTGGCGGCCTCGACAGGCTGATCGAAGATCGTCTCGGGCGGGCCTTCCTCGACGATGCAGCCCTTGTCGATGAAGATCACGCGGTCGGC
GACCTCGCGGGCAAAGCCCATTTCGTGGGTGACGATCATCATGGTCATGCCGTCATTGGCCAGCTCGCGCATGGCATCGAGCACTTCGCCGATCATCTCG
GGATCCAGCGCCGATGTGGGCTCATCGAAGAGCATCAGGCGGGGTTCCATGGCCAGGGCACGGGCCAGGGCAACACGCTGCTGCTGACCGCCGGAGAGCT
GGGTCGGGTACTTGTAGATCTGGTCGCTGATGCCGACACGCTCGAGCAGGCGTTCACCGGTCGCCACCGCCGTTTCGCGGTCCTGCCCGCGTACCTTCAT
GGGCGCCAGAATCACGTTGTCGAGCACGCTGAGGTGCGGAAAGAGGTTGAATTGCTGGAACACCATGCCGACTTCGGTGCGGATCTTCTGCAGGGCCTGA
GTGGCCTTGCCGTAGGGCAGCAAGGGACTGTCGTCGACATCCAGTTCGCCCTGCTGGAATTCTTCCAGGCCGTTGACGCAGCGGATCAGTGTCGACTTGC
CGGAGCCACTGGCGCCGATGACCACGACGACCTCGCCGGTGGCGACGCTCAGGTCGATATCCTTGAGCACGTGAAAGTCGCCGAAGTGCTTGTTGAGCTT
GTTCATGCGCACGATGGGCTCGGCGCCGGCCGTGGACTTCTGCTGGTTCATGAGGTGCCTTCCTTTATTGTCCGACGAGGCCCTTGCGCTCGAGCATGCG
CAGCGCAAACGACAGGGTCAGGGTGATCGCCAGGTACAGCAGGGCAACCATCAGGTAGACCTCCAGGGCGGTGAAGGTGGTGGCGATGTAGATCTGTCCC

>Halomonas sabkhae CECT 7246 [NZ_JAUFPM010000008.1:232498..232710], dnaA> ig dnaN>, 213 bp

< page 1 | #006 | Halomonas almeriensis CECT 7050

WebSkew GCmin pos. 199020 (GCmin<>oriC1 20.6 kb; GCmin<>oriC2 1.3 kb; oriC2<>oriC1 19.3 kb)

>Halomonas almeriensis CECT 7050 [NZ_JAUFPO010000005.1:c178198..178697], <rpmH oriC1 dnaA>, 509 bp

CCCGGCATCTGGTTGCCCATGTCGACATCGGCGCTCGTGTTCGCAGGCACCGAGAGATCGCTGTCGGCTTCTGACTGCCCGGACGCCGGCGAGCTCTGGT
TCGCAATGCCCGGGGCTTCCGGCGCCTGTGTGGCCTGGCCATAGTCCTGATTCCACTGCACTACGATCAAATAGGCAAGCACTGCCAGGGGAATCAGCAG
GAGGAGTCGTTTTACGTCCATGAGGGGTCTGCCCGATGGGTGGTGAAGATGCCGATGGCGTGGCGAGAGTCGCGCCGCCTGCAAACACGAAAGCCCGCCG
ATGGCAGGCCGGTTCTCGTCAAGGGAGCGTCCGCTTTCGGCGGGGCGACGTCAAGCCTGGTGGCAAGAAGAGATGTTGCGCTCATGCAGGCGCCAGGCGT
CGAGCGCATCGTCAGGCCCGGGCCTTCATCGTTGGCTCGTCGGAGCAGCCGAGGCGTGATGCGTGCGCTTGTTGGCTTCACGCTCGAGCCGACGCCACAT
GCCATGCAATTGGCGGTGGACAACTTCATTGTCCAGCTCCTGTACACCGCGTCTCGCCAGTAACACGATATCCACTGCCGGGAGTCGATGCTGGCGATGC
CGAATGGACTCACGCGTGATGCGCTTGAGGCGGTTGCGGTCCACGGCCCGACGGACATTTTTCTTGCTGAAGATCAGGCCGATGCGAGGGCGGCCCAGAG
CATTGGGTGTGGCGAGCGCCATCATTCCCTTGCCGTGCAGCTTGAAGGCTGCCGTGTCGAAAACGTGTCGATAATCCTCGGCGGTCAGCAGTCTCAACCG
CCGAGGAAAGCCCTGACAGGACACGTCTTGCCGATCAAGCGCTTAGACGCTTGCGGCCCTTGGCGCGGCGACGCGCCAGGACGGCACGGCCATTCTTTGT
GGCCATGCGAGCACGGAAGCCGTGCGCGCGCTTGCGCTTGAGAACGCTGGGTTGAAAGGTGCGTTTCATAACTGATTATCCCACGTAGTGTTGGTACGGA

ATCGAAATGCGTTGAGGTGCCCGATGCTTGCCGAAACCACGGGCAAATCCGGCTCAAGACCGGACATTGTAGTGATTTCACCGATCAAGCGCAATTCCTG
AGCGCCCCTGCGATCGTGGGAATCGAACGCCGGCTTTATCGTATCAATATAATAAGAGGGTTGAATGACTGATGTTGTTATTATTGTTGGGGAAACAATT
**TGTGGATAA**AAGGTTTATTCCATAAACAACAGAAGGTTATGCTTTGGTTGAGGCTGTGTAAAAACCGGTAGGAAAGCGGGGGCCGGTTGTGCATGTCCT
GGGCACGACTGTCGGGCTTATCCACATCGGCTGATAGTCCCCCGTTTGTGCCCAGCATGCAGCGGACGATGACAACAGGGTTTTCCACACTGGGAAATCA
GGGCAGTCGCCATGTGGATAACCTGAAGGTTGGGCGCTACAATGTTTGGTCGTTCAAGAATCTACCATGTGAGGTCGTGTGTCACTCGCACTCTGGCAAC

AGTGTCTGGCTACCCTCCAGGATGAGCTGAGCTCTCAGCAGTTCAATACCTGGATTCGCCCCCTGCAGGCGGAAGAGGGAGAGTCCAATCAGCTGCGTCT
GCTGGCGCCCAACCGCTTCGTGCGCGATTGGGTCAGTGACAAGTATGCCAAGCGCATCAGTGAGTTGATGCGTGAGCTGGCGCCTGCTAAGCCGCCCAAG
GTGGTGTTGGTGGTGGGTAGCCGACGTGCTGCGCCGGCCCCTCAGCCGAGAGAGCTGGGTGATCCGGTATCCGCTGCGCCCCAGGCACCCAATGTGCATA
CCCCGCCGCGTGGTGATGTGGCCGATGAGTCGGAGATCGACCGCACGCGCGATGATGAGGAGACCGAGAAGGAGACCCCGCGCCGACCCTCGGGCGAACG
CCAGGTCCAGGTCGAGGGCAGCCTCAAGCACAACAGTGGTCTGAATCAGAACTTTACCTTCGACACTTTCGTCGAGGGCAAGTCGAACCAGCTCGCCCGG
GCCGCCTCCCGCCAGGTCTCCGAGAACCCCGGGGGCGCCTACAATCCGCTCTTTCTCTATGGCGGCGTCGGCCTGGGCAAGACGCACCTGATGCATGCCG
TGGGCAACGCCCTGGACCTGCGTCGCGAGAATGCCCGCGTGGTGTATCTGCATTCCGAACGCTTCGTGGCCGATATGGTCAAGGCGCTGCAGCTTAATGC
CATCAATGATTTCAAGCGCTTCTATCGCAGTGTGGATGCTTTGCTGATTGACGATATCCAGTTCTTTGCCGGCAAGGAACGCTCCCAGGAAGAGTTCTTC
CATACGTTCAATGCCTTGCTGGAAGGTGGTCAGCAGATGATCCTGACCTCGGATCGATATCCCAAGGAAATCAGTGGCGTGGAAGAACGCTTGAAGTCAC
GCTTCGGGTGGGGACTGACCGTGGCAATCGAGCCGCCCGAGCTCGAAACCCGCGTCGCCATTCTGATGAAGAAGGCCGACCAGGCTAAGGTCAACCTGCC

>Halomonas almeriensis CECT 7050 [NZ_JAUFPO010000005.1:c197479..197978], <mnmG oriC2 <COG1126 (glnQ), 537 bp

CCATGCCGTGGATAGAGCGCACCACCTGTAGCTGGACATCGAATGGCAGGGAGGTGGAAATGCCATTGGGGTAGAGCTCGTGGGTGTTCAGGCCTTCGGG
CTCGACAAAGATCTGATGGCTGTTCTTGTCGGCGAAGCGATGCACCTTGTCCTCGATGGACGGGCAATAGCGCGGGCCGACACTATCGATGGTGCCCGAG
AACATCGGTGAGCGTTCCAGGTTGGCATGGATGATCTCGTGGGTGCGCTCATTGGTATGGGCGATATGGCAACTCATCTGCTGCGGATGCAGCCGACGCT
CGCCCAGGTAGGACATCACCGGTGTGGGGCTGTCGCCGGGTTGTTCTTCCAGCTTCGAGAAATCCACGCTCTTGGCATCGATGCGCGGCGGAGTTCCGGT
CTTGAGTCGATCGACGCGGAATGGCTGGGCACGCAGTCGCTCGGCAAGCGCGTTGGAGGGCGGATCTCCGGCACGGCCGCCGCGACTGTTGTCCAGCCCG
ATGTGGATAACGCCCCCCAGGAAGGTACCGGTGCATAGGACCACGCTTTCGCCCATGAAGCGAATGCCGGTCTCGGTGGTCACGCCGCGGACCTGGTGGT
CGTCGACAATCAGATCACCGGCGGCCTGCTGAAAAAGCGTCAGATTGGGCTGGTTTTCCAGCATGCTGCGAATGGCAGCCTTATACAGCACGCGATCGGC
CTGGGCACGTGTGGCGCGTACAGCGGGTCCCTTGCGTGCATTGAGGACACGAAACTGGATGCCGGCCTTGTCGGTGGCCAGTCCCATGGCCCCGCCCAGG
GCGTCGATTTCCTTGACCAGATGGCTTTTGCCGATCCCGCCGATAGCGGGATTGCAGGACATCTGCCCCAGGGTCTCGATGTTGTGGGTCAGCAAGAGGG
TTCGGCAGCCCGTACGAGCCGAGGCCAGAGCGGCTTCGGTGCCCGCATGGCCGCCACCGATGACGATGACGTCGAAACGGTCGGGGTACTCCAAGGTTCA

CCTCGTTCGGCGCATCGGCGCCGATATTGCGAATCGGGGATGAAATCAGCCAGCCAGTATAAACCTCCTGTGGGTAATCGTCAGGGTTTTCCACACCTGG
ATTTTCGGATCGTTCGCTGTTGGGCTCTTCAGACAATAAAATAAATAATATATAGAAAAGAAGTTCTGTTGTTGTTGTTACTACTGGTGTGCACCTTTTC
**TGTTGATAA**AGGTAGAAAATCCCCGATAATCAGATGGTTGTAATGTTGACTTGTCTTCGCAAAAAGGGCTGGTGACCTGCGGACCCCGTGTGTGAGCCCT
GTGGATGAAAGGCCGAGTTAACCACAAGCCACAGCAACCACGCCTTTTCCACCGCTCTCTACCCGGTTTGTCACCGCGGCTGTGAACAACCCGGCCGGTT
ATCCAGAGGCGCCCGGGTGCCGGACGCTGCAGCCTGATTCGCGCTGATGTCACGGCGAAGTTTTCCACAGCCCAAAAAAAGGCCCTGTCGCGGAGAACAG

GGCCTGGTGAATCGCCGATTCGCGGTTCTTGTCAGTGCTTGAGGACGCGTGCCAGAAAGCTCTGGGTGGGCGCCTCGGTAGGCTGATCGAAGATCGTTTC
GGGCGGCCCTTCTTCGACGATTCGCCCCTTGTCGATGAAGATCACGCGGTCGGCCACTTCGCGGGCAAAGCCCATTTCATGGGTCACGATCATCATGGTC
ATGCCGTCGTTGGCCAGCTCGCGCATGGCGTCCAGGACTTCGCCGATCATCTCCGGGTCGAGGGCCGAGGTGGGCTCGTCGAAGAGCATCAAGCGGGGCT
CCATGGCCAGGGCTCGAGCAAGGGCGACGCGCTGCTGCTGTCCCCCGGAAAGCTGGGTGGGGTACTTGTGCATCTGGTCGCTGATGCCCACGCGTTCGAG
CAGACGCTCGCCGGTTGCCACCGCCGTCTCGCGGTCCTGACCGCGGACCTTCATGGGGGCCAGAATCACGTTGTCGAGGACGCTGAGGTGCGGGAAGAGG
TTGAACTGCTGGAAGACCATACCGACCTCGGTGCGGATCTTCTGCAGGGCCTGAGTGGCTTTGCCGTCGGGCAGCAGTGGGTTGTCGTCGACATCCAGTT
CGCCCTGCTGAAATTCTTCCAGGCCATTGACGCAGCGAATCAGGGTCGACTTGCCGGAGCCACTGGCGCCGATGACCACGACCACCTCGCCGGTGGCAAC
GCTCAGATCGATGTCCTTGAGCACGTGAAAGTCGCCGAAATGCTTGTTGAGCTTGTTCATGCGCACGATGGGCTCGGCGCCGGCCTGGGGCTTTTGCTGG
GTCATTGGGTGCCTTCCTTTATTGTCCGACGAGGCCCTTGCGCTCGAGCATGCGCAGTGCAAACGAGAGGGTCAGTGTGATGGCCAGGTACAGTAGTGCC
ACCATCAGATAGACCTCGAGAGCGGTGAAGGTCGTGGCGATGTAGATCTGTCCCTGGCGAACCAGCTCGCCGACGCCGATGACCGAGAATAGCGAGGTAT

>Halomonas almeriensis CECT 7050 [NZ_JAUFPO010000005.1:c176682..176746], dnaA> ig dnaN>, 65 bp

WebSkew GCmin pos. 19500 (GCmin<>oriC1 34.6 kb; GCmin<>oriC2 8.8 kb; oriC2<>oriC1 25.7 kb)

>Halomonas halophila NBRC 102604 [NZ_BJUS01000022.1:c54358..53859], <rpmH oriC1 dnaA>, 513 bp

GTTGCCGGCCATGTCGTCGGGTGCCGTGTTCCGGGTCGGTACCGAGAGGTCGTCACCGCCGGCGTCCGGGGAAGAGGACGTCGGATTCTCGCTGGTGATG
GTCGGGGCCTGGGGGGCGGTGGCCACCTGACCGTAATCCTGATTCCACTGCACGACGATCAGGTAGGCGAGGACCGCCAGGGGAATCAGCAGTAGTAGTC
GTTTTACATCCATGAGGGGTCTGCTCGATGGGTGGTGAACAATCCGGTGGCGCGATGCCGCTCGCCGGCCCGACATACGCGAAAGCCTGCCGGGGGCAGG
CTAGCGTTGGCTCGCCGAGCGTCCCTTGGGAACGCGACGAGGTCAAGCGGGATGGCGTCTGGCGTGATCACGCGCATGCAGGCTCCGGGCGGTGCCAACA
GCGACGGGCAGTCGGCGTCATCGACGCTCTGCCTCAGGTGAGTCGCTCGTGGCCGCCTTGTTCGCTTCGCGTTCCAGACGTTTCCACATGCCGTGCAGCT
GACGGTGGACGGTGGCGTTGTCCAGATCCTGAACGCCGCGTCTGGCGAGCAGCACGATATCCACAGGGGGAAGCCGGTGTTGGCGAAGACGAATGGAGTC
GCGCGCGATGCGCTTGAGGCGGTTGCGGTCGACGGCCCGACGCACGTTCTTCTTGCTGAAGATCAGTCCGAGGCGGGGATGCCCCAGATCATTGGGGGAG
GCCAGGGCCATCAATCCCTTGCCATGGACCTTGAAGGCAGCGGTGTCGAAGACGCGTCGGTAGTCCCCGGCAGTCAGCAGGCGCAGCTGCCGGGGAAAGC
CCTGACTGGACACGCGAGGTCCGCGATCAGGCGCTCAGGCGCTTGCGGCCCTTGGCGCGACGACGCGCCAGGACGGCACGACCGTTCTTGGTGGCCATGC
GAGCACGGAAACCATGCGCGCGCTTGCGCTTGAGAACGCTAGGTTGAAATGTGCGTTTCATAACTCGTTATTCCCACGTGGTTTTTGGAACAAATCGGCG

ATTCAGAGGCGCCCGGGTTCCGGTGGGGAATCCGGGTAGATTCGGCTCAGGACCGGAAATTGTAGTGATATCGGGCCTCGGGCGCAATTTTTCCTTGCGA
CTTTTCGACGGGTCGTTCTCTCGCGGGCCCTGGTGGATCCACTATTATTACTAGTATGGGTTTTCTATTACACCCTGTGATTATTGGTGGGGAAAAGATT
**TGTGGATAA**CCGATATATTTCCTTTTTCCTCAATAAGCTGTGGATATCATAATCGTGTGGACACCCCGTTGGCGATCGGCGAGCAACCGGTGGATGAAGG
GTGGATGGAATCTCGAGTCATCCACAGGGCCACTCACGTACTCAGATTCCTCGCAGGATCTTCCTACCTCCCCGGGCAGGTTTTCCACAGCGATGTCCGC
AGGCGTCATCCATGTGGATAACCTGCGGGTGGCACGATACAATATCCGGTCGTTCTTCAACCGAATGGTGAGGTCGCGTGTCGCTCGCTCTCTGGCAACA

ATGTCTGGATACCCTGCAGGACGAACTGAGTTCCCAGCAGTTCAACACCTGGATTCGCCCCTTGCAGGCGGAGGAAGGGGACGCCAACCAGCTGCGGCTG
CTGGCCCCCAATCGCTTCGTGCGCGACTGGGTCAGTGACAAGTACGCCAAGCGGATCAGCGAACTCATGCGCGAACTGTCGCCGGCCAAGCCGCCCAAGG
TGGTGGTGACCGTGGGCAGTCGTCGTGTGGCGTCGCCGGCCCCGCAGACCCGCGAACTGGGCGACCCGGTCTCCGCCGCGCCTCGCGCCCCGGCGGTGCA
GACGCCACCGCGTGGCGACGTGGCCGACGAGAGCGAGATCGACCAGCTGCGCGACGAGGGAGCGGAGGCGCCCCGCCGGCCTGGCAGCGAGCGCCAGGTA
CAGGTGGAAGGCAGCCTCAAGCACAACAGCGGCCTGAATCCGAGCTTCACCTTCGAGACCTTCGTCGAGGGCAAGTCCAACCAGCTGGCCCGGGCCGCCT
CCCGCCAGGTCTCGGAGAATCCCGGCGGCGCCTACAACCCGCTGTTCCTCTATGGCGGCGTCGGCCTCGGCAAGACCCACCTGATGCATGCCGTCGGCAA
TGCCCTGGCCGGTCGACGCGAGAACGCGCGGGTGGTCTATCTGCACTCCGAGCGCTTCGTCGCCGACATGGTCAAGGCGCTGCAGCTCAATGCGATCAAC
GACTTCAAGCGCTTCTATCGCAGCGTCGATGCGCTGCTGATCGATGACATCCAGTTCTTCGCCGGCAAGGAGCGCTCCCAGGAGGAGTTCTTCCACACCT
TCAACGCCCTGCTGGAGGGCGGCCAGCAGATGATCCTGACCTCGGATCGCTATCCCAAGGAGATCAGCGGCGTCGAGGAGCGCCTCAAATCGCGCTTCGG
CTGGGGTCTGACGGTGGCCATCGAGCCGCCGGAGCTCGAGACCCGGGTGGCGATCCTGATGAAGAAGGCCGACCAGGCCAAGGTCGATCTGCCCCATGAC

>Halomonas halophila NBRC 102604 [NZ_BJUS01000022.1:c28103..28602], <mnmG oriC2 <COG1126 (glnQ), 537

CCAGGCCCTTCATCGAGCGCACGACCTGCAGCTGAACGTCGAACGGCAGCGAGGTCGAGATGCCGTTGGGATACAGCTCGTGGGTATCCAGCCCTTCCGG
CTCGATGAAGATCTGGTGGCTCGCCTTGTCGGCGAAGCGATGCACCTTGTCCTCGATGGACGGGCAGTAGCGCGGCCCGACGCTCTCGATGGTGCCGGAG
AACATCGGCGAGCGATCCAGGTTGGCGCGGATGATCTCGTGGCTGCGCTCGTTGGTCTGGGCGATGTGGCAGCTGATCTGGCGCGGGTGCATCTCGCGGC
TGCCGAGATAGGACATCACCGGCGTTGGGGTATCGCCCGGCTGCTCCTCGAGCGCCGAGAAATCGACGCTCTTGGCATCGATGCGCGGTGGCGTGCCGGT
CTTCAGGCGATCGACGCGGAACGGCAGCGCACGCAGGCGCTCGGCCAGGGCGTTGGAGGGCGGATCGCCGGCGCGACCGCCACGGCTGTTGTCCAGCCCG
ATGTGGATCACCCCGCCGAGGAAGGTACCGGTGCACAGCACCACGGTCTCACCCATGAACCGCACGCCGGTCTCGGTGACCACGCCACGCACGGTGTCGC
CGTCGACGATCAGGTCGCCGGCGGCCTGCTGGAAGATCGTCAGGTTGGCCTGGTTCTCCAGCATGCCGCGGATCGCCGCCTTGTAGCGCACGCGGTCTGC
CTGGGCGCGTGTCGCCCGCACCGCGGGGCCCTTGCGGGAATTGAGTACCCGGAACTGGATGCCGCCCATGTCGGTGGCGAGTCCCATGGCGCCGCCCAGC
GCATCGATCTCCTTGACCAGATGGCTCTTGCCGATCCCGCCGATGGCGGGATTGCAGGACATCTGACCGAGGGTCTCGATGTTGTGGGTCAGCAGCAGGG
TCCGGCAGCCCATGCGAGCCGAGGCCAGGGCGGCTTCGGTCCCGGCATGGCCACCGCCGATGACGATGACGTCGAAGCGGTCGGGGTACTCCAAGGTTCA

CCTCGATCGGCGCCGTCGCGCCGAATGTTGCGGTTCAGGGGTCGAATCAGCCCGCCAGTATAACCCCCTTGTGGGTAATCGTCAGGGTTTTCCACACCCG
TTCAGGCCGATCCGCCTCCCTTCCGCGGAGACTTAAATAAAAAAACATATAGATAAAAAGAAGTTCTGTTGTGGTAGTGATTACTGGTGTGGACATTTCT
**CGTGGATAA**CTGATTTTACCACTTACTTATCAATACCTTGTAATGTTGACCGAAGCAAGGAAAAGGCGCGGACCGCCTGCGGAGGGCATGTTCGGGGCCT
GTGGGTAACACCCCGGGTTACCCACAAGCCAGCGCGAACACGGCTTGTCCACCGAGTTCTTCCGGGCTTGTCACCGCGGCTGTGAACAACCACCTCGTTC
TGTCCCCAGCCCCGCCATGGCCCGCGTGAGCGACGAGCGGCGAAAAATCTTGTCCACAGGCCGAAAAACGCCCAAAAAAAGGGCCCTGTCGCTGACGACA

GGGCCCTTTGAGGCTGGAGGCATGGCGCTTGTCAGTGCTTGAGCACCCGGGACAGGAACTGCCGGGTGGGCTCGGCCTGAGGCGAGTCGAAGATCGTCTC
CGGTGGCCCTTCCTCGGCGATCTCGCCCTTGTGAATGAAGATCACCCGGTCGGCCACCTCACGGGCGAAGCCCATCTCGTGGGTGACGATCAGCATGGTC
ATGCCCTCGTTGGCGAGCTCGCGCATGGCGTCCAGCACCTCGCCGATCATCTCCGGGTCCAGCGCCGAGGTCGGCTCGTCGAACAGCATCAGCCGCGGTT
CCATGGCCAGCGCCCGGGCCAGTGCCACGCGCTGCTGCTGGCCGCCGGAGAGCTGGGTCGGATACTTGGCGGCCTGATCGGCGATGCCGACCCGGTCGAG
CAGGCGCTCGGCGGTATCGGTGGCCTCATGGCGGCCCAGCCCCCGCACCTTCATCGGCGCCAGGGTGACGTTGTCCAGCACGCTCAGATGCGGAAACAGG
TTGAACTGCTGGAACACCATGCCAACCTCGGTGCGGATCTTCTGCAGCGCCTGGGTGGCCTTGCCGTGGGGCTTCAGGGGATTGCCATCGACCTCCAGCT
CGCCCTGCTGGAACTCCTCCAGGCCGTTGATGCAGCGGATCAGGGTCGACTTGCCGGAGCCGCTGGCACCGATGATCACCACCACCTCGCCGGGAGTGAC
CTCGAGATCGATGTTCTTCAGCACGTGCAGGTCGCCGAAGGACTTGTTGAGCTTCTCCATGCGGACGATGCGCTCGCCGCCGGAGGGGGACGTCGGTTGG
TTCATGTGCGTGTCTTCCTTCATTGTCCGACCAGGCCCTTGCGCTCGAGCAGGCGCAGCGCCGCCGAGAGCGACAGAGTGATGACCAGGTACAGCATGGC
CACCATCAGATAGACCTCCAGGGCGTTGAAGGTGGTGGCGATATAGACCTGGCCCTGGCGGACCAGCTCGCCCACGCCGATCACCGAGAACAGTGAGGTG

>Halomonas halophila NBRC 102604 [NZ_BJUS01000022.1:c55803..55866], dnaA> ig dnaN>, 64 bp

WebSkew GCmin pos. 241800 (GCmin<>oriC1 34.3 kb; GCmin<>oriC2 8.6 kb; oriC2<>oriC1 25.7 kb)

>Halomonas salina DSM 5928 [NZ_JASFAS010000003.1:c207217..207716], <rpmH oriC1 dnaA>, 516 bp

GTTGCCGGCCATGTCGTCGGGTGCCGTGTTCCGGGTCGGTACCGAGAGGTCGTCACCGCCGGCGTCCGGGGAAGAGGACGTCGGATTCTCGCTGGTGATG
GTCGGGGCCTGGGGGGCGGTGGCCACCTGACCGTAATCCTGATTCCACTGCACGACGATCAGGTAGGCGAGGACCGCCAGGGGAATCAGCAGTAGTAGTC
GTTTTACATCCATGAGGGGTCTGCTCGATGGGTGGTGAACAATCCGGTGGCGCGATGCCGCTCGCCGGCCCGACATACGCGAAAGCCTGCCGGGGGCAGG
CTAGCGTTGGCTCGCCGAGCGTCCCTTGGGAACGCGACGAGGTCAAGCGGGATGGCGTCTGGCGTGATCACGCGCATGCAGGCTCCGGGCGGTGCCAACA
GCGACGGGCAGTCGGCGTCATCGACGCTCTGCCTCAGGTGAGTCGCTCGTGGCCGCCTTGTTCGCTTCGCGTTCCAGACGTTTCCACATGCCGTGCAGCT
GACGGTGGACGGTGGCGTTGTCCAGATCCTGAACGCCGCGTCTGGCGAGCAGCACGATATCCACAGGGGGAAGCCGGTGTTGGCGAAGACGAATGGAGTC
GCGCGCGATGCGCTTGAGGCGGTTGCGGTCGACGGCCCGACGCACGTTCTTCTTGCTGAAGATCAGTCCGAGGCGGGGATGCCCCAGATCATTGGGGGAG
GCCAGGGCCATCAATCCCTTGCCATGGACCTTGAAGGCAGCGGTGTCGAAGACGCGTCGGTAGTCCCCGGCAGTCAGCAGGCGCAGCTGCCGGGGAAAGC
CCTGACTGGACACGCGAGGTCCGCGATCAGGCGCTCAGGCGCTTGCGGCCCTTGGCGCGACGACGCGCCAGGACGGCACGACCGTTCTTGGTGGCCATGC
GAGCACGGAAACCATGCGCGCGCTTGCGCTTGAGAACGCTAGGTTGAAATGTGCGTTTCATAACTCGTTATTCCCACGTGGTTTTTGGAACAAATCGGCG

ATTCAGAGGCGCCCGGGTTCCGGTGGGGAATCCGGGTAGATTCGGCTCAGGACCGGAAATTGTAGTGATATCGGGCCTCGGGCGCAATTTTTCCTTGCGA
CTTTTCGACGGGTCGTTCTCTCGCGGGCCCTGGTGGATCCACTATTATTACTAGTATGGGTTTTCTATTACACCCTGTGATTATTGGTGGGGAAAAGATT
**TGTGGATAA**CCGATATATTTCCTTTTTCCTCAATAAGCTGTGGATATCATAATCGTGTGGACACCCCGTTGGCGATCGGCGAGCAACCGGTGGATGAAGG
GTGGATGGAATCTCGAGTCATCCACAGGGCCACTCACGTACTCAGATTCCTCGCAGGATCTTCCTACCTCCCCGGGCAGGTTTTCCACAGCGATGTCCGC
AGGCGTCATCCATGTGGATAACCTGCGGGTGGCACGATACAATATCCGGTCGTTCTTCAACCGAATGGTGAGGTCGCGTGTCGCTCGCTCTCTGGCAACA

ATGTCTGGATACCCTGCAGGACGAACTGAGTTCCCAGCAGTTCAACACCTGGATTCGCCCCTTGCAGGCGGAGGAAGGGGACGCCAACCAGCTGCGGCTG
CTGGCCCCCAATCGCTTCGTGCGCGACTGGGTCAGTGACAAGTACGCCAAGCGGATCAGCGAACTCATGCGCGAACTGTCGCCGGCCAAGCCGCCCAAGG
TGGTGGTGACCGTGGGCAGTCGTCGTGTGGCGTCGCCGGCCCCGCAGACCCGCGAACTGGGCGACCCGGTCTCCGCCGCGCCTCGCGCCCCGGCGGTGCA
GACGCCACCGCGTGGCGACGTGGCCGACGAGAGCGAGATCGACCAGCTGCGCGACGAGGGAGCGGAGGCGCCCCGCCGGCCTGGCAGCGAGCGCCAGGTA
CAGGTGGAAGGCAGCCTCAAGCACAACAGCGGCCTGAATCCGAGCTTCACCTTCGAGACCTTCGTCGAGGGCAAGTCCAACCAGCTGGCCCGGGCCGCCT
CCCGCCAGGTCTCGGAGAATCCCGGCGGCGCCTACAACCCGCTGTTCCTCTATGGCGGCGTCGGCCTCGGCAAGACCCACCTGATGCATGCCGTCGGCAA
TGCCCTGGCCGGTCGACGCGAGAACGCGCGGGTGGTCTATCTGCACTCCGAGCGCTTCGTCGCCGACATGGTCAAGGCGCTGCAGCTCAATGCGATCAAC
GACTTCAAGCGCTTCTATCGCAGCGTCGATGCGCTGCTGATCGATGACATCCAGTTCTTCGCCGGCAAGGAGCGCTCCCAGGAGGAGTTCTTCCACACCT
TCAACGCCCTGCTGGAGGGCGGCCAGCAGATGATCCTGACCTCGGATCGCTATCCCAAGGAGATCAGCGGCGTCGAGGAGCGCCTCAAATCGCGCTTCGG
CTGGGGTCTGACGGTGGCCATCGAGCCGCCGGAGCTCGAGACCCGGGTGGCGATCCTGATGAAGAAGGCCGACCAGGCCAAGGTCGATCTGCCCCATGAC

>Halomonas salina DSM 5928 [NZ_JASFAS010000003.1:c232963..233462], <mnmG oriC2 <COG1126 (glnQ), 537 bp

CCAGGCCCTTCATCGAGCGCACGACCTGCAGCTGAACGTCGAACGGCAGCGAGGTCGAGATGCCGTTGGGATACAGCTCGTGGGTATCCAGCCCTTCCGG
CTCGATGAAGATCTGGTGGCTCGCCTTGTCGGCGAAGCGATGCACCTTGTCCTCGATGGACGGGCAGTAGCGCGGCCCGACGCTCTCGATGGTGCCGGAG
AACATCGGCGAGCGATCCAGGTTGGCGCGGATGATCTCGTGGCTGCGCTCGTTGGTCTGGGCGATGTGGCAGCTGATCTGGCGCGGGTGCATCTCGCGGC
TGCCGAGATAGGACATCACCGGCGTTGGGGTATCGCCCGGCTGCTCCTCGAGCGCCGAGAAATCGACGCTCTTGGCATCGATGCGCGGTGGCGTGCCGGT
CTTCAGGCGATCGACGCGGAACGGCAGCGCACGCAGGCGCTCGGCCAGGGCGTTGGAGGGCGGATCGCCGGCGCGACCGCCACGGCTGTTGTCCAGCCCG
ATGTGGATCACCCCGCCGAGGAAGGTACCGGTGCACAGCACCACGGTCTCACCCATGAACCGCACGCCGGTCTCGGTGACCACGCCACGCACGGTGTCGC
CGTCGACGATCAGGTCGCCGGCGGCCTGCTGGAAGATCGTCAGGTTGGCCTGGTTCTCCAGCATGCCGCGGATCGCCGCCTTGTAGCGCACGCGGTCTGC
CTGGGCGCGTGTCGCCCGCACCGCGGGGCCCTTGCGGGAATTGAGTACCCGGAACTGGATGCCGCCCATGTCGGTGGCGAGTCCCATGGCGCCGCCCAGC
GCATCGATCTCCTTGACCAGATGGCTCTTGCCGATCCCGCCGATGGCGGGATTGCAGGACATCTGACCGAGGGTCTCGATGTTGTGGGTCAGCAGCAGGG
TCCGGCAGCCCATGCGAGCCGAGGCCAGGGCGGCTTCGGTCCCGGCATGGCCACCGCCGATGACGATGACGTCGAAGCGGTCGGGGTACTCCAAGGTTCA

CCTCGATCGGCGCCGTCGCGCCGAATGTTGCGGTTCAGGGGTCGAATCAGCCCGCCAGTATAACCCCCTTGTGGGTAATCGTCAGGGTTTTCCACACCCG
TTCAGGCCGATCCGCCTCCCTTCCGCGGAGACTTAAATAAAAAAACATATAGATAAAAAGAAGTTCTGTTGTGGTAGTGATTACTGGTGTGGACATTTCT
**CGTGGATAA**CTGATTTTACCACTTACTTATCAATACCTTGTAATGTTGACCGAAGCAAGGAAAAGGCGCGGACCGCCTGCGGAGGGCATGTTCGGGGCCT
GTGGGTAACACCCCGGGTTACCCACAAGCCAGCGCGAACACGGCTTGTCCACCGAGTTCTTCCGGGCTTGTCACCGCGGCTGTGAACAACCACCTCGTTC
TGTCCCCAGCCCCGCCATGGCCCGCGTGAGCGACGAGCGGCGAAAAATCTTGTCCACAGGCCGAAAAACGCCCAAAAAAAGGGCCCTGTCGCTGACGACA

GGGCCCTTTGAGGCTGGAGGCATGGCGCTTGTCAGTGCTTGAGCACCCGGGACAGGAACTGCCGGGTGGGCTCGGCCTGAGGCGAGTCGAAGATCGTCTC
CGGTGGCCCTTCCTCGGCGATCTCGCCCTTGTGAATGAAGATCACCCGGTCGGCCACCTCACGGGCGAAGCCCATCTCGTGGGTGACGATCAGCATGGTC
ATGCCCTCGTTGGCGAGCTCGCGCATGGCGTCCAGCACCTCGCCGATCATCTCCGGGTCCAGCGCCGAGGTCGGCTCGTCGAACAGCATCAGCCGCGGTT
CCATGGCCAGCGCCCGGGCCAGTGCCACGCGCTGCTGCTGGCCGCCGGAGAGCTGGGTCGGATACGATCGGCGATGCCGACCCGGTCGAGCAGGCGCTCG
GCGGTATCGGTGGCCTCATGGCGGCCCAGCCCCCGCACCTTCATCGGCGCCAGGGTGACGTTGTCCAGCACGCTCAGATGCGGAAACAGGTTGAACTGCT
GGAACACCATGCCAACCTCGGTGCGGATCTTCTGCAGCGCCTGGGTGGCCTTGCCGTGGGGCTTCAGGGGATTGCCATCGACCTCCAGCTCGCCCTGCTG
GAACTCCTCCAGGCCGTTGATGCAGCGGATCAGGGTCGACTTGCCGGAGCCGCTGGCACCGATGATCACCACCACCTCGCCGGGAGTGACCTCGAGATCG
ATGTTCTTCAGCACGTGCAGGTCGCCGAAGGACTTGTTGAGCTTCTCCATGCGGACGATGCGCTCGCCGCCGGAGGGGGACGTCGGTTGGTTCATGTGCG
TGTCTTCCTTCATTGTCCGACCAGGCCCTTGCGCTCGAGCAGGCGCAGCGCCGCCGAGAGCGACAGAGTGATGACCAGGTACAGCATGGCCACCATCAGA
TAGACCTCCAGGGCGTTGAAGGTGGTGGCGATATAGACCTGGCCCTGGCGGACCAGCTCGCCCACGCCGATCACCGAGAACAGTGAGGTGTCCTTGATGC

>Halomonas salina DSM 5928 [NZ_JASFAS010000003.1:c205709..205772], dnaA> ig dnaN>, 64 bp

WebSkew GCmin pos. 424837 (GCmin<>oriC1 34.2 kb; GCmin<>oriC2 8.6 kb; oriC2<>oriC1 25.7 kb)

>Halomonas smyrnensis AAD6 [NZ_AJKS02000002.1:c390339..390838], <rpmH oriC1 dnaA>, 515 bp

TCTCGCTGGCCATGTCGTCCGCCGAGCGGTTCTGGGTCGGCACCGAGAGGTCGTCACCGCCGCTGTCCCCGGAGGAGGACGTCTGGCTCTCGTTGGAGAT
GGTCGGGGCCTGGGGGGCGGTGGCCACCTGACCGTAATCCTGATTCCACTGCACGACGATCAGGTAGGCGAGGACCGCCAGGGGAATCAGCAGTAGTAGT
CGTTTTACATCCATGAGGGGTCTGCTCGATGGGTGGTGAACAATCCGGTGGCGCGATGCCGCTCGCCGGCCCGACATACGCGAAAGCCTGCCGGGGGCAG
GCTAGCGTTGGCTCGCAGAGCGTCCCTTGGGAACGCGACGAGGTCAAGCGGGATGGCGTCTGGCGTGATCACGCGGATGCAGGCTCCGGGCGGTGCCAAC
AGCGACGGGCAGGGAGCGTCATCGACGCTCTGCCGCAGATGCGTCGCCGGCGGTCGCCTTGTTCGCTTCGCGTTCCAGACGCTTCCACATGCCGTGCAGC
TGGCGGTGGACGGTAGCGTTGTCCAGATCCTGAACGCCGCGTCGGGCGAGCAGCACGATATCCACAGGGGGAAGTCGGTGTTGGCGAAGACGAATGGAGT
CGCGCGCGATGCGCTTGAGGCGGTTGCGGTCGACGGCCCGACGCACGTTCTTCTTGCTGAAGATCAGTCCGAGGCGGGGATGCCCCAGATCATTGGGGGA
GGCCAAGGCCATCAATCCCTTGCCATGGACCTTGAAGGCAGCGCTGTCGAAGACGCGTCGGTAGTCCCCGGCAGTCAGCAGGCGCAGCTGCCGGGGAAAG
CCCTGACTGGACACGGGAGGTCCGCGATCAGGCGCTCAGGCGCTTGCGGCCCTTGGCGCGACGACGCGCCAGGACGGCACGACCGTTTTTGGTGGCCATG
CGAGCACGGAAGCCATGCGCGCGCTTGCGCTTGAGAACGCTGGGTTGAAATGTGCGTTTCATAACTCGTTATTCCCACGTGGTTTTTGGAACAAATCGGC

GAATAGAGGCGCCCGGGTTCCGGTGGGGAATCCGGGTAGATTCGGCTCAGGACCGGAAATTGTAGTGATATCGGGCCTCGGGCGCAATTTTTCCTTGCGA
CTTTTCGCCGGGGCGTTCTCTCCCCGTCGATGGCGAATCCACTACCATTACTAGTATTGGTTTTCTATTACACCCTGTGATTATTGATGGGGAAGGTTTC
**TGTGGATAA**CCGGGACAACCTTTTCCACATCAGATTCTTGGCGTTATCACAATCGTGTGGATAGCCTCTTGGGAAGGTCGAGACAGGCGGTGGACGAGCG
GTGCATGGATATGCCACTCATCCACAGGACCGGGATGGTGCCCAGATTCCGCCCAGCTTCCTCCCGTTTCCCCGGGCAGGTTTTCCACAGCGTTGTCCGC
AGCGGTCATCCATGTGGATAACCCGGCGGTGGCGCGATACAATGTCCGGTCGTTCTTCAACCGAATGGTGAGGTCGCGTGTCGCTCGCTCTCTGGCAACA

ATGTCTGGATACCCTGCAGGACGAACTGAGTTCTCAGCAGTTCAACACCTGGATTCGCCCCTTGCAGGCGGAGGAAGGTGAGGCCAACCAGCTGCGTCTG
CTGGCACCCAACCGCTTCGTGCGCGACTGGGTCAGCGACAAGTTCGCCAAGCGCATCAGCGAGCTCATGCGTGAACTGTCGCCGGCCAAGCCGCCCAAGG
TGGTGGTGACCGTGGGCAGTCGGCGCGTGGCGTCGCCGGCCCCGCAGACCCGCGAACTGGGCGACCCCGTCTCCGCCGCGCCTCGGGCCCCGGCGGTACA
GACCCCGCCGCGCGGCGACGTGGCCGACGAGAGCGAGATCGACCAGCTGCGTGACGAGGGCGGCGACGTGCCCCGCCGGCCCGGCAGCGAGCGCCAGGTG
CAGGTGGAAGGCAGCCTCAAGCACAACAGCGGCCTGAACCCGAGCTTTACCTTCGAGACCTTCGTCGAGGGCAAGTCCAACCAGCTGGCCCGGGCCGCTT
CCCGTCAGGTCTCGGAGAATCCCGGTGGCGCCTACAACCCGCTGTTCCTCTATGGCGGCGTCGGCCTCGGTAAGACCCACCTGATGCATGCCGTCGGCAA
CGCCCTGGCCGGGCGGCGCGAGAACGCGCGGGTCGTCTACCTGCACTCCGAGCGCTTCGTCGCCGACATGGTCAAGGCGCTGCAGCTCAACGCCATCAAC
GACTTCAAGCGCTTCTACCGCAGCGTCGATGCGCTGCTGATCGATGACATCCAGTTCTTCGCCGGCAAGGAGCGCTCCCAGGAGGAGTTCTTCCACACCT
TCAATGCCCTGCTCGAGGGTGGCCAGCAGATGATCCTGACCTCGGATCGCTATCCCAAGGAGATCAGCGGGGTGGAGGAGCGCCTCAAGTCGCGCTTCGG
CTGGGGGCTGACGGTGGCCATCGAGCCGCCGGAGCTCGAGACCCGGGTGGCGATCCTGATGAAGAAGGCCGACCAGGCCAAGGTCGATCTGCCCCATGAC

>Halomonas smyrnensis AAD6 [NZ_AJKS02000002.1:c416029..416528], <mnmG oriC2 <COG1126 (glnQ), 537 bp

CCAGGCCCTTCATCGAGCGCACGACCTGCAGCTGGACGTCGAATGGCAGCGAGGTCGAGATGCCGTTGGGATACAGCTCGTGGGTATCCAGGCCTTCCGG
CTCGATGAAGATCTGGTGGCTCGCCTTGTCGGCGAAGCGATGCACCTTGTCCTCGATGGACGGGCAGTAGCGCGGCCCGACGCTCTCGATGCTGCCGGAG
AACATCGGCGAACGATCGAGGTTGGCGCGGATGATCTCGTGGCTGCGCTCGTTGGTCTGGGCGATATGGCAGCTGATCTGGCGCGGGTGCATCTCGCGGC
TGCCGAGATAAGACATCACCGGCGTCGGGGTATCGCCCGGCTGCTCCTCGAGCATCGAGAAGTCCACGCTCTTGGCATCGATGCGCGGCGGGGTGCCGGT
CTTCAGACGATCGACGCGGAACGGCAGCGCGCGCAGGCGCTCGGCCAGGGCGTTGGAGGGCGGATCGCCGGCCCGGCCACCGCGGCTGTTGTCCAGCCCG
ATGTGGATAACCCCGCCGAGGAAGGTGCCCGTGCACAGCACCACGGTCTCGCCCATGAAGCGCACGCCGGTCTCGGTGACCACGCCGCGCACGGTGTCGC
CGTCGACGATCAGGTCGCCGGCGGCCTGCTGGAAGATCGTCAGGTTGGCCTGGTTTTCCAGCATGCCGCGGATCGCCGCCTTGTAGCGCACACGATCGGC
CTGGGCGCGGGTGGCGCGAACGGCGGGACCCTTGCGGGAATTGAGTACCCGGAACTGGATGCCGCCCATGTCGGTGGCGAGTCCCATGGCGCCGCCCAGC
GCATCGATCTCCTTGACCAGATGGCTCTTGCCGATCCCGCCGATGGCGGGATTGCAGGACATCTGGCCGAGGGTCTCGATGTTGTGGGTCAGCAGCAGGG
TCCGGCAGCCCATGCGAGCCGAGGCCAGGGCGGCTTCGGTCCCGGCATGGCCACCGCCGATGACGATGACGTCGAAGCGGTCGGGGTACTCCAAGGTTCA

CCTCGATCGGCGCCGTCGCGCCGAATGTTGCGGTTCAGGGGTCGAATCAGCCCGCCAGTATAACCCTCTTGTGGGTAATCGTCAGGGTTTTCCACACCCT
CCAGGCTAATCCGCTTTCCGTCCGCCGAGACTTAAATAAAAAAACATATAGATAAAGAAGAAGTTCTGTTGTGGTAGTGATTACTGGTGTGGACATTTTC
**CGTGGATAA**CCGGCTTTACAACTTATCTATCAACAACTTGTAATGTTGACTGAAACAAGGACAAGGCGCGGACCGCCTGCGGAGGGCATGTTCGGGGGCT
GTGGGTAACCCCTCGGGTTACCCACAAGCCGGCTCGGACACGGGTTGTCCACGGAGTTCTTCCGGGCTTGTCACCGCGGCTGTGAACAACCACCCCGATC
TGTCCCCAGCCCCGCCATGGCCCGGCTGGACCGCCCATCGCGGAAAATCTTGTCCACAGGCCGAAAACGGCCAAAAAAAGGCCCCTGTCGCGGACGACAG

GGGCCGTTGAGGCCGAGATGAACGGCGCTCGTCAGTGCTTGAGCACCCGGGACAGGAACTGCCGGGTGGGCTCGGCCTGAGGCGAATCGAAGATCGTCTC
GGGCGGGCCTTCCTCGGCGATCTCGCCCTTGTGAATGAAGATCACCCGGTCGGCCACCTCGCGGGCGAAGCCCATCTCGTGGGTGACGATCAGCATGGTC
ATGCCCTCGTTGGCGAGCTCGCGCATGGCGTCGAGCACCTCGCCGATCATCTCCGGGTCCAGCGCCGAGGTCGGCTCGTCGAACAGCATCAGCCGTGGCT
CCATGGCCAGCGCCCGGGCGATGGCCACGCGCTGCTGCTGGCCGCCGGAGAGCTGGGTCGGATACTTGGCGGCCTGGTCGGCGATGCCGACCCGGTCGAG
CAGGCGCTCGGCGGTGTCGGTGGCCTCACGGTGGTCCATGCCGCGCACCTTCATCGGCGCCAGGGTGACGTTGTCCAGCACGCTCAGATGCGGGAACAGG
TTGAACTGCTGGAACACCATGCCGACCTCGGTGCGGATCTTCTGCAGCGCCTGGGTGGCCTTGCCGTAGGGCTTGAGGGGATTGCCATCGACCTCCAGCT
CGCCCTGCTGGAACTCCTCCAGGCCGTTGATGCAGCGGATCAGGGTCGACTTGCCGGAGCCGCTGGCACCGATGATCACCACCACCTCGCCGGGGGTGAC
CTCGAGGTCGATGTTCTTCAGCACGTGCAGGTCGCCGAAGGACTTGTTGAGCTTCTCCATGCGGACGATGCGCTCGCCGCCGGAGGGGGACGTCGGTTGG
TTCATGTGCGTGTCTTCCTTCATTGTCCGACCAGGCCCTTGCGCTCGAGCAGGCGCAGCGCCGCGGAGAGCGACAGGGTGATGACCAGGTACAGCATGGC
CACCATCAGATAGACCTCCAGGGCGTTGAAGGTGGTGGCGATGTAGACCTGGCCCTGGCGGACCAGCTCGCCCACGCCGATCACCGAGAACAGCGAGGTG

>Halomonas smyrnensis AAD6 [NZ_AJKS02000002.1:c388825..388894], dnaA> ig dnaN>, 70 bp

WebSkew GCmin pos. 236000 (2nd min.) (GCmin<>oriC1 29.8 kb; GCmin<>oriC2 4.2 kb; oriC2<>oriC1 25.6 kb)

>Halomonas borealis ATF 5.2 [NZ_SDMO01000002.1:c205922..206421], <rpmH oriC1 dnaA>, 515 bp

ATGCCGTCGGGGGCCCGGTTCTGGGTCGGCACCGAGAGGTCTCCGCCGTCGGCATCATCGGAGGCCGAGGAGGTCGACGTCTGGGACTCGCTCGAGATGG
CCGGGGCCTGGGGGGCGGTGGCCACCTGACCGTAATCCTGATTCCACTGCACGACGATCAGGTAGGCGAGGACCGCCAGGGGAATCAGTAGTAGTAGTCG
TTTTACATCCATGAGGGGTCTGCTCGATGGGTGGTGAACAATCCGGTGGCGCGATGCCGTTCGCCGGCCCGACATACGCGAAAGCCTGCCAGGGGCAGGC
TAGCGTCAGCTCGCCGAGCGTCCCTTGGGAGGACGACGACGTCAAGCGGGGTGGCGTCTGGCGTGATCACGCGCATGCAGGCTCCGGGCGGTGCTAACAG
CGACCGGCAGGTCGCGTCATGAGCGATCTGCCGTTTGTGCATCGCTCGTCGCCGCCTTGTCCGCTTCGCGTGCCAGGCGTTTCCACATGCCGTGTAGCTG
GCGGTGGACGGTGGCGTTGTCCAGCTCGTGAACGCCGCGTCTGGCGAGCAGTACGATATCCACAGCGGGAAGTCGATGTTGGCGAAGGCGAATGGAGTCG
CGCGCGAGGCGCTTGAGGCGGTTGCGGTCGACGGCCCGACGTACGTTCTTCTTGCTGAAGATCAGCCCGAGGCGGGGATGCCCCAGATCATTCGGGGAGG
CCAGGACCATCAATCCCTTGCCATGGACCTTGAAGGCAGCGCTGTCGAAGACGCGTCGATAATCCCCGGCAGTCAGCAGACGCAGCTGCCGGGGAAAGCC
CTGACTGGACATGTGATGTCCGCGGTTCAGGCGCTCAGGCGCTTGCGGCCCTTGGCGCGACGACGCGCCAGGACGGCACGACCATTCTTGGTGGCCATGC
GAGCACGGAAGCCATGCGCGCGCTTGCGCTTGAGAACGCTGGGTTGAAATGTGCGTTTCATAACTCGTTATTCCCACGTGGTTTTTGGAACAAATCGGCG

ATTCAGAGGTTCCCGGTTTCCGGTGGGGAATCCGGGTAGATTCGGCTCAGGACCGGAAATTGTAGTGATATCGGGCCCTGGGCGCAATTTTTCCTTGCGA
TTTTTCGCTGGGGGCGATCTCGCGAGCCAACATCCCTACAACTGTTATTACTAGTATTGGGGTTCTATTACACCCTGTTATTACTAGTGGGGAGGGTTTT
**TGTGGATAA**CCTAGCACTTCTTTTCCATGACAGTCGCTTATGAGCGTAAGAATCGTGTGGATGGACGCTTGAAACAGGGTGCAAGGAAGGTGGATGAGCG
GTGGATGGATGGGCGAACTATCCACAGCCCGTGAGGCATGCCCAGAGACTCCTCAGGATACCTACGACCTCCAGGCAGGTTGTCCACAGAGTTTTTCACG
CCCCTCATCCATGTGGATAACCCGGCGATGGCACGATACAATGCCCGGTCGTTCTTCAACCGAATGGTGAGGTCGCGTGTCGCTCGCTCTCTGGCAACAA

TGTCTGGATACCCTGCAGGACGAACTGAGTTCCCAGCAGTTCAACACCTGGATTCGCCCCTTGCAGGCGGAGGAAGGGGAGGTCAACCAGCTGTGTCTGC
TGGCGCCGAACCGTTTCGTGCGTGACTGGGTCAGCGACAAGTACGCCAAGCGGATCAGCGAGCTCATGCGCGAGCTGTCGCCGACCAAGCCGCCGAAGGT
GGTGCTCACCGTGGGCAGTCGCCGCACCGTGTCACCGGCACCACAGACCCGCGAGTTGGGTGATCCGGTCTCCGCGGCGCCGCGCGCCCCTGCGGTGCAG
ACGCCGCCGCGCGGTGACGTGGCCGACGAGAGCGAGATCGATCAGCTGCGCGACGAGGGCGAGGCGCCGCGTCGCCCCGTCAACGAACGTCAGGTACAGG
TAGAGGGCAGCCTCAAGCACAACAGTGGCCTGAACCCCAACTTCACCTTCGAGACGTTCGTCGAGGGCAAGTCCAACCAGCTGGCCCGGGCCGCCTCCCG
TCAGGTCTCCGAGAATCCCGGTGGTGCCTATAACCCGCTGTTTCTATACGGTGGTGTCGGCCTGGGTAAGACCCACCTGATGCACGCCGTCGGCAATGCG
CTGGCCGGCCGGCGCGAAAACGCACGGGTCGTCTATCTGCACTCCGAGCGCTTCGTCGCCGACATGGTCAAGGCGCTGCAGCTCAATGCCATCAACGACT
TCAAGCGCTTCTACCGTAGCGTCGATGCCTTGCTGATCGATGACATCCAGTTCTTCGCCGGCAAGGAGCGTTCCCAGGAGGAGTTCTTCCACACCTTCAA
CGCCTTGCTCGAGGGTGGCCAGCAGATGATCCTGACCTCGGACCGCTACCCGAAGGAGATCAGCGGCGTCGAGGAGCGCCTCAAGTCGCGTTTCGGCTGG
GGCCTGACGGTGGCCATCGAACCGCCTGAGCTGGAGACACGCGTGGCGATCCTGATGAAGAAGGCCGACCAGTCCAAGGTCGATCTGCCCCATGACGCGG

>Halomonas borealis ATF 5.2 [NZ_SDMO01000002.1:c232070..231571], <mnmG oriC2 <COG1126 (glnQ), 525 bp

CATGCCTTTCATCGAGCGCACGACCTGCAGCTGGATGTCGAACGGCAGCGAGGTCGAGATGCCGTTGGGATACAGCTCGTGGGTATCCAGCCCCTCCGGT
TCGATGAAGACCTGGTGGCTCGACTTGTCGGCGAAGCGGTGCACCTTGTCCTCGATCGAGGGGCAATAACGGGGCCCGACGCTATCGATCGTGCCCGAGA
ACATCGGCGAACGCTCGAGATTGGCGCGGATGATCTCGTGGCTGCGCTCGTTGGTCTGAGCGATGTGGCAGCTGACCTGGCGTGGGTGCATGGCGCGGTT
GCCGCGATAGGACATCACCGGTGTCGGGGTATCTCCGGGCTGCTCCTCGAGCATCGAGAAATCCACGCTCTTGGCGTCGATACGCGGTGGCGTGCCGGTC
TTAAGCCGATCGACGCGGAATGGCAACGCACGCAGGCGTTCCGCCAAGGCATTGGAAGGCGGATCGCCGGCCCGGCCGCCACGGCTATTGTCCAGCCCGA
TGTGGATAACCCCGCCAAGGAAGGTGCCTGTGCACAACACCACGGTCTCGGACATGAAGCGCACGCCGGTCTCGGTGGTCACGCCGCGCACGGTGTCACC
GTCGACGATCAGGTCGCCGGCGGCCTGCTGGAAGATCGTCAGGTTGGCTTGGTTCTCCAGCATTCCGCGGATCGCCGCCTTGTAGCGAACGCGATCGGCC
TGGGCGCGAGTCGCCCGTACGGCGGGGCCTTTGCGGGAGTTGAGTACCCGAAACTGAATTCCGCCCCGGTCGGTGGCGAGTCCCATGGCACCGCCGAGCG
CATCGATCTCCTTGACCAGATGACTCTTGCCGATCCCGCCGATGGCGGGATTGCAGGACATCTGACCCAGGGTCTCGATGTTGTGGGTCAGCAGCAGGGT
CCGGCGGCCCATGCGGGCCGAGGCCAGGGCGGCTTCGGTCCCGGCATGGCCACCGCCGATGACGATGACGTCGAAGCGGTCGGGGTACTCCAAGGTTCAC

CTCGATCGGCGCCGTCGCGCCGAATGTTGCGGATACAGGGGTGTAATCAGCCGGCCAGTATAACGTCCTTGTGGGTAATCGTCAGGGTTTTCCACACCGG
GGAAGCCAAGCTGGGTTCGTCATTCGATTGACCTAAATATAAAAGCATATAAATAAAAAGGGGTTCTGTTGTTGTAGTTATTATTGGTGTGGACATTTCC
**CGTGGATAA**CCGAAACAAAACAACAAATATCAAGTTCTTACAGTGTTGACTGATGAGCGGAAAAGGCGCGGATCTCCTGCGGAGGGGATGTTGGAGGCCT
GTGGGTAAATGCCCGGGTTAACCACAAGCCTCGCTGAACACGGCTTATCCACGCGATTCTACCGGGCTTGTCACCGTGGGTGTGAACAACCGCGCCGATC
CATCCCCAGCCCGTGATGGTGGGACGCAGCGGCGAACCGCCAAGAATCTTGTCCACAGGCCAAAAAAAGGCCCTGTCGATGACGACAGGGCCTGGTACAG

CGAGAAAAATCGTCGCTATCAGTGCTTGAGTACCCGCGACAGGAAGTGACGAGTCGGCTCGGCCTGGGGGGCATCGAAGATCGTCTTCGGCGGTCCTTCC
TCGGCGATCTCACCCTTGTGAATGAAGATCACTCGATCGGCGACCTCGCGGGCGAAGCCCATCTCATGGGTGACGATCAGCATGGTCATGCCCTCGTTGG
CGAGCTCGCGCATGGCATCCAGCACCTCGCCGATCATCTCCGGGTCCAGCGCCGACGTCGGCTCGTCGAACAGCATCAGTCGGGGTTCCATGGCCAGTGC
TCGGGCCAGTGCCACACGCTGCTGCTGGCCGCCGGAAAGCTGGGTAGGCAGCTTTTCGGCCTGGTCGCTGATGCCCACGCGTTCGAGCAGGCGTTCGGCC
GCCTCGGTGGCTTTCTGGCGGGATAGATTGCGCACCTTGATGGGCGCCAGGGTCACGTTGTCGCGCACGCTCAGGTGGGGGAACAGATTGAACTGCTGGA
AGACCATGCCGACCTCGGTACGGATCTTCTGTAGCGCCCTGGTGGCCTTTCCGTGAGGTTTAAGGGGGTTGCCATCTACCTCGAGCTCGCCTTGCTGGAA
CTCTTCCAGGCCATTGATGCAGCGGATCAGCGTCGACTTGCCGGAACCGCTGGCGCCGATGATCACCACCACTTCGCCGGGGGTGACCTCAAGGTCGATG
TCCTTGAGCACGTGCAGGGTCCCGAAGGACTTGTTGAGCTTGTTCAGGCGCACGATGGGCTCGCCGCCGGGTTGTGACGTCTGTTGAGTCATCTCGTGCA
TTCCTTATTGTCCGACCAGGCCCTGGCGCTCCAGCAGGCGCAATGCCACCGACAGCGAGAGGGTGACCACCAGGTAGAGCATGGCGACCATCAGGTAGAC
CTCGAGGGCATTGAAGGTGGTGGCGATATAGACCTGGCCCTGGCGGACCAGCTCGCCGACGCCGATCACCGAGAACAGCGAGGTGTCCTTGATGCTGATG

>Halomonas borealis ATF 5.2 [NZ_SDMO01000002.1:c204412..204481], dnaA> ig dnaN>, 70 bp

WebSkew GCmin pos. 55300 (2nd min.) (GCmin<>oriC1 34.8 kb; GCmin<>oriC2 8.9 kb; oriC2<>oriC1 25.9 kb)

>Halomonas organivorans CECT 5995 [NZ_JACHXM010000012.1:89838..90337], <rpmH oriC1 dnaA>, 516 bp

GCCGGGAATGCTGCCGGCCATGTCTGCCGTCGCCTGCTGGTTGGCAGGCACCGAAAGATCATCGCCGGAAGACGCCGAGGGCGCCTGGCCCGCGCTTGAA
ATGGCCGGGGCCTCGGGGGCGGTAGCCACCTGACCGTAATCCTGATTCCACTGCACGACGATCAGGTAGGCGAGGACCGCCAGGGGAATCAGCAGTAAGA
GTCGTTTTACATCCATGAGGGGTCTGCTCGATGGGTGGTGAACAATCCGATGGCACGATGCCACTCGACGGTCCGACATACGCAAAAGCCTGCCGAGGGC
AGGCTATCGTCGGCTCGCCGAGCGTCCCTTGGAAGCGCGGCGGCGTCAAGCAGGGGGGCGTCTGGCATGATCATGCGCATGTTGGCTCCTGGCGGTGCCG
ACAGCAGTGTGCAGGCGCATTATCACTGACTTGCCGTAGGGGCATCATGTTTCGCCGCCTTGGCAGCCTCGCGTTCCAGCCGTCGCCACATGCCATGAAG
CTGGCGATGGACGGTGGCGTTATCGAGTTCATGAACGCCGCGTCTGGCAAGCAGCACGATATCCACAGGAGGGAGTCGATGCTGGCTCAGGCGGATGGAC
TCGCGCGCCAGGCGCTTGAGACGGTTGCGGTCGACGGCTCGACGCACGTTCTTCTTGCTGAAGATCAGCCCGAGGCGAGGATGTCCCATGGGGTTGGGCG
AGGCCAGGGCCATTAATCCCTTGCCATGGACCTTGAAGGCCGCCGTCTCGAAGACACGTCGATAGTCCCCGGCGGTCAGCAATCGCAGCCGCCGGGGAAA
GCCCTGACAGGACACGAGGTCCGCGCTCAGGCGCTCAGGCGCTTGCGGCCCTTGGCACGACGACGTGCCAGGACGGCACGACCGTTCTTGGTGGCCATAC
GAGCACGGAAGCCGTGCGCGCGCTTGCGCTTGAGAACGCTGGGTTGAAAGGTGCGTTTCATAACTCGTTATTCCCACGTGGTTTTTGGAACAAATCGGCG

ATTCAGAGGCGCCCGGGTTCCGGTGGGGAATCCGGGTAGATTCGGCTCAGGACCGGAAATTGTAGTGATTTTGTGCCCGCGGCGCAATTTTTCCTTGGCG
ACGCCAGCGGCGGCACCGGCCATGATCGGCCCATATGGCTACTACAGTTACTAAATAAGTCTTTTTATTAAACCCTGTTATTATTAATGGGGAAGGATTT
**TGTGGATAA**GCATGAATTTCAATTAAAATACAAATAATTAAGCGATTCAATCTGGTGTGGATAGGCTATTGGAAAAAGGTGGAGGCCCCGTGGAGGAGCG
GTGTACGTCTGCGGACATTATCCACAGGCCGGAGGTTGTTCCCAGGTGTCACGCATTCTCTCCCGCGCTTCCAGGCCATATTATCCACAGGCGTGGTGGA
ATCTTGGGTTCATGTGGATAACCTGGCAATCGCACGATACAATGCCCGGTCGTTCTTCAACCCAATGGTGAGGTCGCGTGTCGCTCGCTCTCTGGCAACA

GTGTCTCGATACCCTGCAGGACGAACTGAGTTCCCAGCAGTTCAATACCTGGATCCGTCCGCTGCAGGCGGAGGAAGGGGAAGCCAACCAGCTGCGGCTG
CTGGCTCCCAACCGTTTCGTGCGCGATTGGGTGAGTGACAAGTATGCCAAGCGGATCAGCGAGCTCATGCGCGAGCTCGCCCCTGCCAAGCCCCCCAAGG
TGGTGTTGACCGTGGGGAGTCGCCGAGCCGCTGCGGCGCCTGCCCCGCAGCCCCGCGAGCTGGGCGATCCGGTGTCCGCCGCGCCGCGTGCTCCGGCGGT
GCGTACGCCTCCGCGCGGGGATGTCGCCGACGAGAGTGAGATCGAACGGCTTCGTGACGAAGGGGCGGAGCCCGCTCGACGGCCTGGTGGTGAGCGCCAG
GTGCAGGTCGAGGGTAGTCTCAAGCACAATAGCGGCCTCAATCCCAACTTCACTTTCGAGACCTTCGTCGAGGGCAAGTCCAACCAGCTGGCTCGGGCCG
CCTCGCGCCAGGTGTCGGAGAATCCCGGCGGCGCCTACAACCCGCTGTTCCTCTATGGCGGTGTCGGTCTCGGCAAGACCCACTTGATGCATGCGGTTGG
CAATGCCCTGGCAGGGCGGCGGGAGAATGCCCGGGTCGTCTACCTGCACTCGGAGCGATTCGTCGCCGACATGGTCAAGGCGCTGCAGCTTAATGCCATC
AACGATTTCAAGCGCTTCTATCGCAGTGTCGATGCGCTGCTGATCGATGATATCCAGTTCTTCGCCGGCAAGGAGCGTTCGCAGGAGGAGTTCTTCCACA
CCTTCAACGCCTTGCTCGAGGGGGGGCAGCAGATGATCCTGACCTCGGACCGCTATCCCAAGGAGATCAGCGGGGTCGAGGAGCGGCTCAAGTCCCGCTT
CGGTTGGGGGCTCACGGTCGCCATCGAGCCGCCGGAGCTGGAGACCCGGGTGGCGATCCTGATGAAGAAGGCCGATCAGGCCAAGGTCGACCTGCCTCAT

>Halomonas organivorans CECT 5995 [NZ_JACHXM010000012.1:63962..64461], <mnmG oriC2 <COG1126 (glnQ), 529 bp

CAGCCCCTTGATCGAGCGCACCACCTGCAGCTGCACGTCGAACGGCAGCGAGGTGGAGATGCCGTTGGGATACAGCTCGTGGGTATCCAGCCCCTCGGGT
TCGATGAAGATCTGGTGGCTCGCCTTGTCGGCAAAGCGGTGCACCTTGTCCTCGATGGACGGGCAGTAGCGCGGCCCCACGCCCTCGATCGTGCCGGAGA
ACATCGGCGAACGATCCAGGTTGGCGCGGATGATCTCGTGGCTGCGCTCGCTGGTGTGAGCGATATGGCAACTGACCTGGCGAGGATGCATGTCACGGCT
ACCGAGGTAGGACATCACCGGCGTCGGGGCGTCCCCCGGCTGCTCCTCGAGTACCGAGAAATCCACGCTCTTGGCGTCGATACGGGGTGGCGTGCCGGTC
TTCAGGCGGTCGACGCGAAACGGCAGCGCACGCAGGCGTTCGGCCAGGGCGTTGGAAGGCGGATCGCCGGCCCGGCCGCCGCGGCTGTTGTCGAGCCCGA
TGTGGATAACGCCGCCGAGGAAGGTGCCTGTGCATAACACCACGGTATCGCTCATGAAGCGCACGCCGGTCTCGGTGACCACCCCGCGTACGGTGTCACC
GTCGACGATCAGGTCGCCGGCCGCCTGCTGGAAGATCTCAAGGTTGGGCTGGTTCTCCAGCAGACCTCGAATCGCTGCCTTGTAGCGCACGCGATCGGCC
TGGGCACGGGTCGCCCGCACGGCCGGCCCCTTGCGGGCATTGAGTACCCGGAACTGGATGCCGCCTAGGTCGGTGGCCAGCCCCATGGCACCGCCGAGCG
CATCGATCTCCTTGACCAGATGGCTCTTGCCGATCCCGCCGATGGCGGGATTGCAGGACATCTGGCCCAGGGTCTCGATGTTGTGGGTCAGCAGCAGGGT
CCGGCAGCCCATGCGAGCCGAGGCCAGGGCGGCTTCGGTCCCGGCATGACCACCGCCGATGACGATGACGTCGAAGCGGTCGGGATACTCCAAGGTTCAC

CTCGATCGGCGCCGTCGCGCCGAATGTTGCGGATCAGGGTATGAAATCAGCCCGCCAGTATAACTCCCTTGTGGGTAATCGTCAGGGTTTTCCACACCCG
GGAAGCGCCGCGCCCGCGGTCCTTCCTCTCACTTAAATCAATAAACAAATAGATAAAGAGAAGTTCTGTTGTGGTAGTGACTACTGGTGTGGACATTTAT
**CGTGGATAA**CCCATTTTATACTTTACTTATCAACATGTTGAAGTGTTGACCACTGCGCGGAAAAGCGGCGGGTGACCTGCGGACGGGATGTTCGGGCCCT
GTGGAGGAATGCGCGGTTTACCCACAGCCCGTCTCGAACACGGCTTGTCCACGTGATTCTACCGGGCTTGTTACCGCGCCTGTGAACAACCTCGCTGTTT
CCTCCTGTGCCCCGGGATGGCAGGGCTCAGCGGCCATTACCGGAAAATCTTGTCCACAGGCCAAAAAAAGGCCCTGTCGATAACGACAGGGCCCTGTGAG

GGACGCAGAAGACGCGACGCTGTCAGTGCTTGAGCACCCGCGACAGGAAATTTCGGGTGGGTTCGGCCTGGGGCGAGTCGAAGATCGTTTCCGGTGGCCC
CTCCTCGGCGATCTCGCCCTTGTGGATGAAGATCACCCGATCGGCGACTTCGCGGGCGAAGCCCATCTCATGGGTGACGATCAGCATGGTCATGCCCTCG
TTGGCGAGTTCGCGCATGGCATCCAGCACCTCGCCGATCATCTCCGGGTCCAGCGCCGATGTCGGCTCGTCGAACAGCATCAAACGGGGTTCCATGGCCA
GCGCCCGGGCCAGCGCCACGCGCTGCTGCTGGCCGCCGGAGAGCTGAGTAGGGAACTTGGCGGCCTGGTCGCTGATGCCCACTCGCTCCAGCAGCCGTTC
GGCGGTCTCGGTGGCATCCTCACGGCTCCAGCCGCGAACCTTCATCGGGGCCAGGGTGACGTTGTCCAGCACGCTGAGATGCGGAAAGAGGTTGAACTGC
TGGAACACCATGCCGACCTCGGTGCGAATCTTCTGCAGCGCCTTGGTGGCCTTGCCATGGGGTTTCAGTGGGTTGCCGTCGACTTCCAGCTCACCCTGCT
GGAATTCCTCGAGGCCGTTGATGCAGCGGATCAAGGTCGACTTTCCGGAGCCGCTGGCACCGATGATCACCACCACCTCGCCGGGTGTGACCGACAGGTC
GATGTCCTTGAGCACGTGGAGGGTGCCGAAGGACTTGTTGAGCTTGCTCAGGCGCACGATGGGCTCGCCGCCGGATTGGGACGTCTGTTGGTTCATCTCG
TGCATTCCTTATTGTCCGACCAGGCCCTTGCGCTCCAGCAGGCGCAATGCCACTGACAGGGTCAGGGTGACCACCAGGTAGAGCATGGCGACCATCAGGT
AGACCTCGAGGGCATTGAAGGTGGTGGCAATGTAGACCTGGCCCTGACGCACCAGCTCACCTACGCCGATCACCGAGAACAGCGAGGTGTCCTTGATGCT

>Halomonas organivorans CECT 5995 [NZ_JACHXM010000012.1:91785..91855], dnaA> ig dnaN>, 71 bp

WebSkew GCmin pos. 222708 (2nd min.) (GCmin<>oriC1 47.6 kb; GCmin<>oriC2 22.6 kb; oriC2<>oriC1 25.0 kb)

>Halomonas koreensis Lim DSM 23530 [NZ_JARWAK010000004.1:270038..270537], <rpmH oriC1 dnaA>, 557 bp

CGGGCACCGAGAGATCGTCGGCGCCGCCGTCGCCGGGGGCCGGCTGCGTCTCGCTCGTGATGCCCGGAGCCTGGGGCGCGGTGGCCACCTGGCCGTAATC
CTGGTTCCACTGCACGACGATCAGGTAGGCGAGGATCGCCAGGGGAACCAGTAGGAGAAGTCGTTTTACGTCCATGAGGGATCTGCCCGATGGGTGGTGA
ACAATCCGATGGCGCGATGCCTGACGCGCACCCGACATACGCGAAAGCCTGCCGGGGGCAGGCTAGCGTGCGTGTGTCGAGCGTCGGGGCTGGATGCCGC
GACGTCAAGCGGGGAGGCGTCTGGCCGGGTCATGCGCATGCAGGCTCCGGTCGCTGCCGGCAGTGCCTCATCGGCGATCTGCCGCGGGGGTGTCGGAGGC
GCGGGCCTTTGCCGCCCTGGTCGCTTCGCGTTCCAGGCGCCGCCACATGCCATGCAGCTGGCGATGGACGGTCGCGTTGTCCAGCTCGTGCACGCCGCGC
CTGGCGAGCAGCACGATATCCACAGCGGGTAACCGGTGCTGGCGATGACGGATGGACTCGCGCGCGAGGCGCTTGAGGCGGTTGCGGTCGACGGCTCGCC
GCACGTTCTTCTTGCTGAAGATCAGGCCGAGGCGGGGATATCCCAGGGTATTGGGAGAGGCCAGGGCCATCAGCCCCTTGCCGTGGACCTTGTAGGCGGC
CGTTTCGAAGACGCGCCGGTAGTCCCCGGCAGTCAGCAGTCGCAGCTGCCGGGGAAAGCCCTGACTGGACACGCGCGGTCCGCGATCAGGCGCTCAGGCG
CTTGCGGCCCTTGGCGCGACGACGCGCCAGGACGGCACGACCGTTCTTGGTGGCCATGCGAGCACGGAAGCCGTGAGCGCGCTTGCGCTTGAGGACGCTG
GGTTGAAAGGTGCGTTTCATAACTCGTTATTCCCACGTGGTTTTTGGAACAAATCGGCGATTCAGAGGCGCCCGGGTTCCGGTGGGGAATCCGGGTAGAT

TCGGCTCAGGACCGGAAATTGTAGTGATTGCGGGGCCTCGGCGCAATTTTTCCGGGGACGCAATCGTGCGCCGAGCGCTTGCGGAGGGCTCGGCGCGACC
TTCGAGCGCGGGCAGGCGTCCACCGGTGGAAAGCGGTCCAATCGTTATTACGAATTTTATAGGTTTGTAGTGCCTTGTTATTACTATTGGGGACGAATGG
**TGTGGATAA**TTCGATATAGTATCCACAGATCAGGATGTTGCGATTTTTATATCGCTGTGGGCAAGTGGCGGACGAGCGTCGGAGGAGCGGTGGACGGACG
GTGCATGAATGAGGCGCTTGTCCACATGCCGAGGGGTATCCCCGAAATCCCGGCAGCTTGTCCGCACCCTGGCCGCCCGGTTTTCCACAGCGCATCCGGG
GGCAGGGGTTCATGTGGATAACCCGGCGGCAGGACGATACAATGCCCGGTCGTTCTTCAATCGAATGGTGAGGTCGCGTGTCGCTCGCTCTCTGGCAACA

GTGTCTGGATACCCTGCAGGACGAACTGAGCTCCCAGCAGTTCAACACCTGGATTCGTCCGCTTCAGGCGGAGGAGGGGGAGTCCAACCAGTTGCGGCTG
CTGGCTCCCAACCGCTTCGTGCGCGACTGGGTCAGCGACAAGTACGCCAAGCGCATCAGCGAGCTCATGCGCGAGCTGGCCCCCGCCAAGCCGCCCAAGG
TGGTGCTGACGGTGGGCAGCCGCCGGGCCGCGGCCCCCGCCCCCCAGCCCCGCGAGCTGGGGGACCCGGTCTCCGCGGCGCCGCGAGCCCCCGCCGTGCA
CACGCCGCCGCGGGGCGATGTCGCCGACGAGAGCGAGATCGAGCAGGCCGTCGAGGAGCCCGAGCCGCCGCGCCGTCCCGGCGGCGAGCGCCAGGTCCAG
GTGGAGGGCAGCCTCAAGCACAACAGCGGCCTCAATCCCAACTTCACCTTCGAGACCTTCGTCGAGGGCAAGTCCAACCAGCTGGCTCGGGCCGCCTCGC
GCCAGGTCTCGGAGAATCCCGGCGGCGCCTATAACCCGCTGTTCCTCTACGGGGGCGTGGGCCTGGGCAAGACCCACCTGATGCACGCCGTGGGCAACGC
CCTGGCCGGACGGCGCGAGAATGCCCGGGTGGTCTACCTGCACTCCGAACGCTTCGTCGCCGACATGGTCAAGGCGTTGCAGCTCAACGCCATCAACGAC
TTCAAGCGCTTCTACCGCAGCGTCGATGCGCTGCTGATCGACGACATCCAGTTCTTCGCCGGCAAGGAGCGCTCCCAGGAGGAGTTCTTCCACACCTTCA
ACGCCCTGCTCGAGGGCGGCCAGCAGATGATCCTGACCTCGGACCGTTATCCCAAGGAAATCAGCGGGGTGGAGGAGCGGCTAAAGTCCCGGTTCGGCTG
GGGGCTGACGGTGGCCATCGAGCCCCCGGAGCTCGAGACCCGGGTGGCGATCCTGATGAAGAAGGCCGACCAGGCCAAGGTCGACCTGCCTCACGATGCC

>Halomonas koreensis Lim DSM 23530 [NZ_JARWAK010000004.1:245051..245550], <mnmG oriC2 <COG1126 (glnQ), 527 bp

AGGCCGCGGATCGAACGCACCACCTGCAGCTGCACGTCGAAGGGCAACGAGGTGGAGATACCGTTGGGATACAGCTCGTGGGTGTCCAGGCCCTCGGGCT
CGACGAAGATCTGGTGGCTCGCCTTGTCCGCGAAGCGGTGCACCTTGTCCTCGATGGACGGGCAGTAGCGCGGTCCCACGCCCTCGATGACCCCGGAGAA
CATCGGCGAGCGCTCCAGGTTGGCGTGGATGATCTCGTGGGTGCGCTCGTTGGTATGGGCGATATGGCAGCTGAGCTGTCGGGGGTGCATGCCGCGATCG
CCCAGGTAGGACATCACCGGGGTCGGGGTGTCCCCCGGCTGCTCCTCGAGCACCGAGAAGTCCACGCTCTTGGCGTCCAGTCGCGGCGGGGTCCCGGTCT
TCAGCCGATCGACGCGGAACGGCAGGGCCCGGAGCCGCTCGGACAGGGCATTGGACGGCGGGTCGCCGGCGCGTCCGCCGCGGCTGTTGTCCAGCCCGAT
GTGGATAACCCCGCCCAGGAAGGTGCCTGTGCACAACACCACCGTTTCCCCCAGGAAGCGCACCCCGGTCTCGGTGACCACGCCGCGCACGGTGTCGCCG
TCGACGATCAGGTCGCCGGCGGCCTGCTGGAAGAGCGTCAGGTTGGGCTGGTTCTCCAGCATGCCGCGGATGGCCGCCTTGTAGCGCACCCGGTCGGCCT
GGGCTCGCGTGGCCCGGACGGCGGGGCCCTTGCGGGCGTTGAGCACGCGGAACTGGATGCCGCCCAGGTCGGTGGCCAGCCCCATGGCGCCGCCGAGCGC
ATCGATCTCCTTGACCAGATGGCTCTTGCCGATCCCGCCGATGGCGGGATTGCAGGACATCTGGCCCAGGGTCTCGATGTTGTGGGTCAGCAGCAGGGTC
CGGCAGCCCATGCGGGCCGAGGCCAGGGCGGCTTCGGTCCCCGCATGGCCTCCGCCGATGACGATCACGTCGAAGCGGTCGGGGTACTCCAAGGTTCACC

TCGTTCGGCGGCGCCGCGCCGAGTGTTGCGGATCAGGGAAGTGGAATCAGCCCGCCAGTATAACCCTCTTGTGGATCATCGTCAGGGTTTTCCACACCGG
GAGGCCGCGGCCTTGCCGTCCTTCCTCCCTGACCTAATAAATAAACAGATAGATAAAGAGAAGTTCTGTTGTGGTAGTGACTACTGGTGTGGACATTTTC
**TGTGGAAAA**GCCATTTTAATGCTTTTTTATCAACAGCCTGGAATGTTGAGGGATACGGGGACGGCGCGCGGGGGAGCTGCGGACGGCATGTTCGGGGCCT
GTGGATGACTGCGCCGGTTACCCACAGCTCGAGATCGTCACCGCTTTTCCACCGACCCCTACCGGCCTTGTCACCGTGGCTGTGAACAACCCGGGAAGGT
GCCGCTCGCCCCCGTCATCAAGGGGCTGGCGGGGGGTTCGGCCGATTTATTGTCCACAGGCAAAAAAAGGCCCTGTCGATGACGACAGGGCCCCTTTTCG

CGGTCTGGCGACGAGAGCGTCAGTGCTTGAGCACCCGGGACAGGAACTGACGGGTGGGCTCTTCCCGGGGCGCGTCGAAGATCGTCTCCGGGGGGCCCTC
CTCGGCGATCTCGCCCTTGTGGATGAAGATGACCCGATCGGCCACCTCCCGGGCGAAGCCCATCTCGTGGGTGACCACCAGCATGGTCATGCCTTCGTTG
GCGAGCTCGCGCATGGCGTCCAGCACCTCGCCGATCATCTCCGGATCCAGGGCCGAGGTAGGCTCGTCGAACAGCATCAGCCGGGGTTCCATGGCCAGGG
CTCGCGCCAGGGCCACGCGCTGCTGCTGTCCGCCCGACAGCTGGGTCGGATACTTGTGGGCCTGGTCGGCGATGCCGACGCGCTCGAGCAGGCGCTCGGC
GGTCTCGACGGCCGTCGCACGGTTCAGCCCGCGTACCCGCATGGGCGCCAGGGTGACGTTGTCCAGCACGCTCAGATGGGGGAAGAGGTTGAACTGCTGG
AAGACCATGCCGACCTCGGTGCGGATCTTCTGCAGCGCCTGGGTGGCCTTGCCGTGGGGCTTGAGGGGGTTGCCGTCGACGACGAGCTCGCCCCTGCTGGA
ACTCCTCGAGGCCGTTGATGCAGCGGATCAGGGTCGACTTGCCCGAGCCGCTGGCGCCGATGATCACCACCACCTCGCCGGGCGTGACCTCGAGGTCGAT
GTCCTTCAGCACGTGCAGGCTGCCGAAGTGCTTGTTGACCTTGTCCAGGCGCACGATGGGGTCGCCGCCGGTCTTGGACGTCTGTGCGTTCATGTCATGC
CTTCCTTATTGTCCGACCAGGCCCTTGCGCTCCAGCAGGCGCAGCGCGAAGGACAGCGACAGGGTGATCACCAGGTAGAGGGCGGCGACCATCAGGTACA
CCTCCAGGGCGTTGAAGGTGGTGGCGATATAGACCTGGCCCTGGCGCACCAGCTCGCCGACGCCGATGACCGAGAACAGCGAGGTGTCCTTGATGCTGAT

>Halomonas koreensis Lim DSM 23530 [NZ_JARWAK010000004.1:271979..272051], dnaA> ig dnaN>, 99 bp

WebSkew GCmin pos. 185258 (GCmin<>oriC1 34.4 kb; GCmin<>oriC2 8.5 kb; oriC2<>oriC1 25.9 kb)

>Halomonas stenophila CECT 7744 [NZ_JACHXR010000003.1:c150588..151087], <rpmH oriC1 dnaA>, 525 bp

ATGGAGCCGGCCATGTCCGACGCCTCGGCGTCGCTGGGGACCGAGAGGTCATCGCTGCCGGTGGTCGTCGGTGACGGGGCGTCGCCGGCGTCACTGGAGA
TGGCCGGGGCCTGCGGTGCCGTGGCCACCTGGCCGTAATCCTGATTCCACTGCACGACGATCAGGTAGGCGAGGACCGCCAGGGGAATCAGCAATAAGAG
TCGTTTTACGTCCATGAGGGTTCTGCCCGATGGGTGGTGAACATTCCGATGGCGCAACGCAGGTCGCCCACCCGACAAACGCGAAAGCCTGCCGAGGGCA
GGCTAGCGTTGGTCAGCAGAGCATCTGCTTTCGGCGTGACGGCGTCAAGAAGCGTGGTCATGCGCATGCAGGCTCCGCAGGGTGCCGACAGCGGTTGGCT
GGGCGTGTCATCGACGCTCGGTCGCGAGGGCGCCGGTCGATTCTCTCCCTTCCCCCTTCCTCGCGTCCCGCTCCAGTCGTCGCCACATGCCGTGCAACTG
GCGATGCACGGTCTCGTTGTCCAGCTCCTGCACCCCGCGTCGGGCGAGCAGCACGATATCCACGGCGGGGAGGCGATGCTGGCGAAGGCGAATGGATTCA
CGCGCGAGGCGCTTGAGGCGGTTGCGGTCGACGGCGCGCCGGACATTCTTCTTGCTGAAGATGAGCCCGAGGCGAGGATGTCCCAGGGAGTTGGGAGTGG
CCAGGGCCATCAACCCCTTGCCGTGGACCTTGAAGGCAGCCGTATCAAAGACACGCCGATAGTCCCCGGCGTTGAGCAGCCTCAGCCGCCGGGGAAAGCC
CTGACAGGACACGCGAGGTCCGCGCGTCAGGCGCTCAGGCGCTTGCGGCCCTTGGCGCGACGACGCGCCAGGACGGCACGACCGTTCTTGGTGGCCATGC
GAGCACGGAAACCATGCGCGCGCTTGCGCTTGAGAACGCTGGGTTGAAAGGTGCGTTTCATAACTCGTTATTCCCACGTGGTTTTTGGAACAAATCGGCG

ATTCAGAGGCGCCCGGGTTCCGATGGGGAATCCGGGTAGATTCGGCTCAAGACCGGAAATTGTAGTGATTTCGAGCCGCCAGCGCAATTTTTCCGTGCGC
CCCAGGATAGCTGTCGCTGCGCCGGCTCCGCCGCGTAGGCACTATCATTACTAATTATGAAGTTCTATTGAGCCCTGTTATTACTAGTGGGGAAAGTTAT
**TGTGGACAA**GTCTGAATATCGTGCTCGATTCAATCATCTATGGAAAAGACAGGGCTGTGGACAGTGGGGGATGCCCGGTCCGCAACCTGTTGATGAGCG
GTGGGTCGATCCGTGAGCTGTCCACAGTCACAGGATTCTTCACGTTCGGCACCCAGATTGTCCCTGAGGCAACGGCAGGTTTTCCACAGCACTTCCCGCC
ACCGTGCCTCATGTGGATAACCCGGCGGCAGGACGATACAATGTGCGGTCGTTCTTCAACCGAATGGTGAGGTCGCGTGTCGCTCGCTCTCTGGCAACAA

TGTCTGGATACCCTGCAGGACGAACTGAACTCCCAGCAGTTCAACACCTGGATCCGTCCGCTGCAGGCGGAGGAAGGGGAATCCAACCAGCTGCGCCTGC
TGGCGCCCAATCGCTTCGTGCGCGACTGGGTCAGCGACAAGTATGCCAAGCGCATCAGCGAGCTGATGCGGGAGCTGTCGCCGGCCAAGCCGCCCAAGGT
CGTCCTCACCGTGGGCAGTCGCCGGGCGGCCGCGCCTGCGCCCCAGCCTCGCGAGCTGGGCGATCCGATCTCCGCCGCGCCCCGGGCGCCGGCCGTGCAG
ACGCCTCCGCGGGGAGACGTGGCCGATGAGCGCGAGATCGATCAGCTCCGGGGGGAGGGGGAGGGCGCCCGCCGTGCCACGGGAGAGCGCCAGGTCCAGG
TGGAGGGCAGCCTCAAGCACAACAGTGGCCTGAATCCCAACTTCACCTTCGAGACCTTCGTCGAGGGCAAGTCCAACCAGCTGGCACGCGCCGCGTCCCG
CCAGGTGTCGGAGAACCCGGGAGGCGCCTACAATCCCCTCTTTCTCTACGGCGGGGTCGGCTTGGGCAAGACCCACCTGATGCATGCCGTCGGCAATGCG
CTGGCGGGCCGACGCGAGAACGCCCGGGTGGTCTACCTGCACTCCGAGCGCTTCGTCGCCGACATGGTCAAGGCCCTCCAGCTCAACGCCATCAACGACT
TCAAGCGCTTCTATCGCAGCGTCGATGCCCTGTTGATCGACGATATCCAGTTCTTCGCCGGCAAGGAACGCTCCCAGGAGGAGTTCTTCCATACCTTCAA
TGCCCTGCTCGAGGGCGGTCAGCAGATGATCCTGACCTCGGATCGCTACCCCAAGGAAATCAGTGGGGTGGAAGAGCGGCTCAAGTCCCGCTTCGGCTGG
GGGCTGACGGTGGCGATCGAGCCGCCGGAGCTCGAGACTCGGGTGGCGATCCTGATGAAGAAGGCCGATCAGGCCAAGGTCGATCTGCCCCATGACGCGG

>Halomonas stenophila CECT 7744 [NZ_JACHXR010000003.1:c176497..176996], <mnmG oriC2 <COG1126 (glnQ), 537 bp

AGTCCCCGGATCGAGCGTACCACCTGCAGCTGGACGTCGAAGGGCAGCGAGGTGGAGATGCCGTTGGGGTAGAGCTCATGGGTATCCAGTCCCTCGGGTT
CGATGAAGATCTGGTGGCTGGCCTTGTCGGCGAAGCGGTGCACCTTGTCCTCGATGGACGGGCAATAGCGCGGCCCCACGCCTTCGATCGTCCCCGAGAA
CATCGGCGAGCGATCCAGGTTGGCCTGGATGATCTCGTGGCTGCGTTCGTTGGTATGCGCGATGTGACAGCTCACCTGACGCGGATGCATCTCCCGGCGA
CCCAGGTAGGACATCACCGGCGTGGGGCTGTCGCCGGGCTGTTCATCGAGGACCGAGAAATCCACGCTCTTGGCATCGAGGCGCGGCGGTGTCCCGGTCT
TCAGCCGATCGACCCGGAACGGCAGGGCCCGCAGCCGCTCGGCCAGCGCATTGGAGGGCGGATCGCCGGCTCGGCCGCCGCGGCTGTGGTCGAGACCGAT
GTGGATAACCCCGCCGAGGAAGGTGCCTGTGCACAACACCACGCCCTCACCGAGGAAGCGCACGCCGGTCTCGGTCACCACACCCCGCACGGTGTCGCCG
TCGACGATCAGGTCCCCGGCGGCCTGCTGGAAGAGTGTCAGGTTGGGCTGGTTCTCCAGCATGCCGCGGATCGCCGCCTTGTAGCGGACTCGATCGGCCT
GGGCGCGGGTGGCGCGCACGGCGGGGCCCTTGCGGGCATTGAGCACCCGGAACTGGATGCCGCCCATGTCGGTGGCCAGGCCCATGGCCCCACCGAGGGC
GTCGATCTCCTTGACCAGGTGACTCTTGCCGATCCCGCCGATGGCGGGATTGCAGGACATCTGGCCCAGGGTCTCGATGTTGTGGCTCAGCAGCAGGGTC
CGGCAGCCCATGCGGGCCGAGGCCAGGGCGGCTTCGGTCCCCGCATGACCGCCACCGATGACGATGACGTCGAAGCGGTCGGGGTACTCCAAGGTTCACC

TCGTTCGGCGCGACGGCGCCGATATTGCGAATCGGGGGTATGAAATCAGCCAGCCAGTATAACCCTCCTGTGGGTAATCGTCAGGGTTTTCCACACCCGG
ATGACCAGGCGTCCGTCATCCTCGGGTAGCAGATAAACAAATACATAGAATAAGAAAGAGAAGTTCTGTTGTTGTAGTAATTACTGGTGTGGATACTTTC
**CGTGGAAAA**GCAGGCTTATCTTTTATTTATCAACAGCTTGTACTGTTGACCAAAACAAGGAAAAGCGGCGTGCTGCCTGCGGAGGAGATGTTCGGGAGCT
GTGGACGAAAGGCCGAGTTGCCCACAGCCGCGACCGGACACCGCTTGTCCACCGTCCTCTACCGCGCTTGTCACCGCGGCTGTGAACAACCACCGTGTTT
CATCCCCAGGCCCGCCATCATGGGCACTGGCGCCGGTTCGACGCGGTTCGGCGGAGAATCTTGTCCACAGGCCAAAAAAAGGCCCTGTCGCGAACGACAG

GGCCTTTGTCCAGGGCTCGAGGCGTGGCCTCAGTGCTTCAACACCCGGGCCAGGAAGTTCCGGGTGGCTTCCTCCCGGGGCGAATCGAAGATCGTCTCGG
GAGGACCCTCCTCGGCGATTTCCCCCTTGTGGATATAGATCACCCGGTCGGCGACCTCCCGGGCGAAGCCCATCTCGTGGGTCACGATCATCATGGTCAT
GCCCTCGTTGGCGAGCTCGCGCATGGCATCCAGGACCTCGCCGATCATCTCCGGGTCCAGCGCCGAGGTCGGCTCGTCGAAGAGCATCAGCCGGGGCTCC
ATGGCCAGGGCCCGGGCCAGGGCCACCCGCTGCTGCTGGCCGCCGGACAACTGGCCGGGATACTTGGCGGCCTGGTCGCTGATGTCGACCCGCTCGAGCA
GCCGCTCGGCGATCTCGGTGGCATCGGCCCGACTCCAGCCGCGGACCTTCATGGGCGCCAGGGTCACGTTGTCGAGCACGCTGAGGTGGGGGAAGAGGTT
GAACTGCTGGAACACCATGCCCACCTCGGTGCGGATCTTCTGCAGCGCCTTGGTGGCCTTGCCGTAGGGCAGCAGCGGTTTGCCGTCCACCTCCAGGCTG
CCCTTCTGGAACTCCTCGAGGCCATTGATGCAGCGGATCAGGGTGGACTTGCCCGAGCCGCTGGCGCCGATGACCACCACCACCTCGCCGGGGCTGACCT
CGAGGTCGATGTCCTTGAGGACATGCAGGCTGCCGAAGTGCTTGTTGACCTTGTCCAGGCGCACGATGGGCTCGCCGCCGGACTTGGACGTCTCTTGGTT
CATGGCTCGCATTCCTTATTGTCCGACCAGGCCCTTGCGCTCCAGGAAACGCAGCGCCACCGACAGTGATAGGGTGATCACCAGGTAGAGCATGGCGACC
ATCAGGTACACCTCCAGGGCGTTGAAGGTGGTGGCGATGTAGACCTGGCCCTGGCGGACCAGCTCGCCGACACCGATGACCGAGAACAGCGAGGTGTCCT

>Halomonas stenophila CECT 7744 [NZ_JACHXR010000003.1:c149078..149147], dnaA> ig dnaN>, 70 bp

WebSkew GCmin pos. 246000 (2nd min.) (GCmin<>oriC1 30.6 kb; GCmin<>oriC2 10.2 kb; oriC2<>oriC1 20.4 kb)

>Halomonas maura CECT 5298 [NZ_JAUFPQ010000013.1:276369..276868], <rpmH oriC1 dnaA>, 514 bp

CATGTCCGAGGCCCCGGCATCGCTGGGGACCGAGAGATCATCGCTGCCGGCGGTCGTCGGTGACGGGGCATCGCCGGCGTTACTGGAGATGGCCGGGGCC
TGCGGTGCCGTGGCCACCTGGCCGTAATCCTGATTCCACTGCACGACGATCAGGTAGGCGAGGACCGCCAGGGGAATCAGCAATAAGAGTCGTTTTACGT
CCATGAGGGTTCTGCCCGATGGGTGGTGAACATTCCGATGGCGCAACGCAGGTCGCCCACCCGACATACACGAAAGCCTGCCGAGGGCAGGCTAGTGTTC
GTCAGCAGAGCGTCTGCTTTCGGCGTGACGGCGTCAAGGGTGGCGGCGAGAAGCGTGGTCATGCGTATGCTGGCTCCGCGGGGTGCCGACAGCGGTTGGC
GGGACGTGTCATCGTCGCTCGGTCACCAGGGTGCCGGCCGATTCTCTCCCTTCCCTCTGACTCGCCTCTCGCTCCAGTCGCCGCCACATGCCGTGCAACT
GGCGATGCACGGTCGCGTTGTCCAGATCCTGTACCCCGCGTCTGGCGAGCAGCACGATATCCACGGCGGGGAGACGATGCTGGCGAAGGCGAATGGATTC
GCGTGCGAGACGCTTGAGGCGGTTGCGGTCGACGGCGCGCCGGACGTTCTTCTTGCTGAAGATGAGCCCGAGGCGAGGATATCCCAGGGAGTTGGGGGTG
GCCAGGGCCATCAACCCCTTGCCGTGGACCTTGAAGGCAGCCGTATCGAAGACACGTCGGTAGTCCCCGGCGTTGAGCAGCCTCAGCCGCCGGGGGAAGT
CCTGACAGGACACGCGAGGTCCGCGGGTCAGGCGCTCAGGCGCTTGCGGCCCTTGGCGCGACGACGCGCCAGGACGGCACGACCGTTCTTGGTGGCCATG
CGAGCACGGAAACCATGCGCGCGCTTGCGCTTGAGAACGCTGGGTTGAAAGGTGCGTTTCATAACTCGTTATTCCCACGTGGTTTTTGGAACAAATCGGC

GATTCAGAGGCGCCCGGGTTCCGCTGGGGAATCCGGGTAGATTCGGCTCAAGACCGGAAATTGTAGTGATTTCGAGCCGCCAGCGCAATTTTTCCGTGTC
CTCCAGCTCCCTGTCGCCGTCCCGGCCTGGCTGCGGAGGCGCTACCATAACTAATAATGAAGTTCTATTGAGCCCTGTTATTATTAATGGGGAAAGTTAT
**TGTGGATAA**GTGCAAATGGCATCGGCGATTCAAGTGCCTATGACAGCCACAGGGTTGTGGACGGCGCGGGGATGTCCGGCCCGCAAGCTGTTGACAACGG
GTGGATCGATCCTTGAACTGTCCACAGCCACAGGGCATTCCACCTGCGGCGACCAGGCTATCCCTCGGACCACAGCAGGTTTTCCACAGCGCTTTCCGGC
TCCGTGCTTCATGTGGATAACCTGGCCGCAGGGCGATACAATGTGCGGTCGTTCTTCAACCGAATGGTGAGGTCGCGTGTCGCTCGCTCTCTGGCAACAA

TGTCTGGATACCCTGCAGGACGAACTGAACTCCCAGCAGTTCAACACCTGGATCCGTCCGCTGCAGGCGGAGGAAGGGGAATCCAATCAGCTGCGCCTGC
TGGCGCCCAATCGCTTCGTGCGCGACTGGGTCAGCGACAAGTATGCCAAGCGCATCAGTGAGCTGATGCGCGAGCTTTCGCCGGCCAAGCCCCCCAAGGT
GGTCCTGAGCGTGGGCAGTCGCCGGGCGGCCGCCCCGGCGCCCCAGCCCCGCGAGTTGGGGGACCCGGTCTCCGCCGCGCCCCGGGCGCCGGCCGTGCAG
ACGCCTCCCCGGGGGGACGTGGCCGATGAACGCGAGATCGACCAGCTGCGAGAGGAGGGGGAGGGCGCACGCCGTGCCGGGGGAGAGCGCCAGGTGCAGG
TGGAAGGCAGTCTCAAGCACAACAGTGGCCTGAACCCCAACTTCACCTTCGAGACCTTCGTCGAGGGCAAGTCCAACCAGCTGGCACGCGCCGCGTCACG
CCAGGTCTCGGAGAACCCCGGGGGGGCCTACAACCCGCTGTTCCTGTATGGCGGGGTCGGCCTGGGCAAGACCCACCTGATGCATGCGGTCGGCAATGCG
TTGGCAGGCCGACGCGAGAATGCCCGGGTGGTCTATCTGCACTCCGAGCGCTTCGTCGCCGACATGGTCAAGGCCCTCCAGCTCAACGCCATCAACGACT
TCAAGCGCTTCTATCGCAGCGTCGATGCCCTGTTGATCGACGATATCCAGTTCTTCGCCGGCAAGGAGCGCTCCCAGGAGGAGTTCTTCCATACCTTCAA
TGCCCTGCTCGAGGGCGGTCAGCAGATGATCCTGACCTCGGATCGGTATCCCAAGGAAATCAGTGGGGTGGAAGAACGGCTCAAGTCCCGCTTCGGCTGG
GGACTGACGGTGGCGATCGAGCCGCCGGAGCTCGAGACCCGGGTGGCGATCCTGATGAAGAAGGCCGATCAGGCCAAGGTCGATTTGCCCCATGATGCCG

>Halomonas maura CECT 5298 [NZ_JAUFPQ010000013.1:255997..256496], <mnmG oriC2 <COG1126 (glnQ), 536 bp

CAGTCCGCGGATCGAGCGTACCACCCGCAGCTGGACGTCGAAGGGCAGCGAGGTGGAGATGCCATTGGGATAGAGCTCGTGGGTGTCCAGTCCCTCGGGT
TCGATGAAGATCTGGTGGCTGGCCTTGTCGGCGAAGCGGTGCACCTTGTCCTCGATCGACGGGCAATAGCGCGGCCCCACGCCTTCGATGGTGCCCGAGA
ACATCGGCGAGCGATCCAGGTTGGCGTGGATGATCTCGTGGCTGCGCTCGTTGGTGTGGGCGATGTGGCAGCTGACCTGGCGCGGGTGCATGTCGCGGTT
GCCGAGGTAGGACATCACCGGGGTGGGGCTGTCGCCGGGCTGCTCCTCGAGCACCGAGAAATCCACGCTCTTGGCGTCCAGTCGCGGCGGCGTGCCGGTC
TTCAGGCGATCGACCCGGAACGGCAGGGCCCGCAGTCGCTCGGCCAGCGCATTGGAGGGCGGGTCGCCGGCCCGGCCGCCGCGACTGTGATCGAGGCCGA
TGTGGATAACCCCGCCGAGGAAGGTACCTGTGCACAACACCACGCTCTCGCCGAGGAAGCGCACCCCGGTCTCGGTGACCACGCCGCGTACCGTGTCGCC
GTCGACGATCAGGTCGCCCGCGGCCTGCTGGAAGAGCGTCAGGTTGGGCTGGTTCTCCAGCATGCCGCGGATCGCCGCCTTGTAGCGGACCCGATCGGCC
TGGGCGCGGGTCGCGCGCACCGCAGGCCCCTTGCGGGCGTTGAGCACCCGGAACTGGATGCCACCCAGGTCGGTGGCCAGGCCCATGGCCCCGCCGAGGG
CATCGATCTCCTTGACCAGATGGCTCTTGCCGATCCCGCCGATGGCGGATTGCAGGACATCTGTCCCAGGGTCTCGATGTTGTGGCTGAGCAGCAGGGT
CCGGCAGCCCATGCGGGCCGAGGCCAGGGCGGCTTCGGTCCCCGCATGACCGCCACCGATGACGATGACGTCGAAGCGGTCGGGGTACTCCAAGGTTCAC

CTCGTTCGGCGCGCCCGCGCCGATATTGCGAATCGGGGGTATGAAATCAGCCAGCCAGTATAACCCTCCTGTGGGTAATCGTCAGGGTTTTCCACACCCG
GAAGTCCGAGCCGGCGTCATCCCGAGACCCCATTAGACAAATACATAGAATAAGAAAGAGAAGTTCTGTTGTTGTAGTGACTATTGGTGTGGATACTTTC
**CGTGGATAA**GCCGTTTTATTCTATTTACAACAACAGCTTGAATTGTTGACCAAGACAAGGAAAAGCGGCGTGCCGCCTGCGGAGGAGATGTTCGGGAGCT
GTGGACGAAAGGCCGGGTTGCCCACAGGCGTGGCGAGTCACCGCTTGTCCACCGCCGCCTACCGCGCTTGTCACCGTGGCTGTGAACAACTGCCGTGGTT
CATCCCCAGCCCCGGCAGCATGGGCGCTGGGGACGGATTGCCCCGTCCCGGCCGGCGATCTTGTCCACAGGCCAAAAAAAGGCCCTGTCGGGGTCGACAG

GGCCTTTGTGCCGGGCGCGAGGCGTGACCTCAGTGCTTCAACACCCGGGCCAGGAAGTTCCGGGTGGCTTCCTCCCGGGGCGAATCGAAGATCGTCTCGG
GGGGGCCCTCCTCGGCGATTTCCCCCTTGTGGATATAGATCACCCGGTCGGCGACCTCCCGGGCGAAGCCCATCTCGTGGGTCACGATCATCATGGTCAT
GCCCTCGTTGGCGAGCTCGCGCATGGCATCCAGGACCTCGCCGATCATCTCCGGGTCCAGCGCCGAGGTGGGCTCGTCGAAGAGCATCAGCCGGGGCTCC
ATGGCCAGGGCCCGGGCCAGGGCCACCCGCTGCTGCTGGCCGCCGGACAGCTGGCCGGGATACTTGGCGGCCTGGTCGCTGATGTCGACCCGCTCGAGCA
GCCGCTCGGCGATCTCGGTGGCATCGGCCCGACTCCAGCCGCGGACCTTCATGGGGGCCAGGGTGACGTTGTCGAGCACGCTGAGGTGGGGGAAGAGGTT
GAACTGCTGGAACACCATGCCCACTTCGGTGCGGATCTTCTGCAGCGCCTTGGTGGCCTTGCCGTGCGGCAGCAGGGGTTTGCCGTCCACCTCCAGGCTG
CCCTGCTGGAACTCCTCGAGGCCATTGATGCAGCGAATCAGGGTGGACTTGCCCGAGCCGCTGGCGCCGATGATCACCACCACCTCGCCGGGGGTGACCT
CGAGGTCGATGTCCTTGAGGACGTGCAGGCTGCCGAAGTGCTTGTTGACCTTGTCCAGGCGCACGATGGGCTCGCCGCCGGAGGTGGACGTCTCTTGGTT
CATGTCTCGCATTCCTTATTGTCCGACCAGGCCCTTGCGCTCCAGGAAGCGCAGCGCCACCGACAGCGACAGGGTGATCACCAGGTAGAGCATGGCGACC
ATCAGGTACACCTCCAGGGCATTGAAGGTGGTGGCGATGTAGACCTGGCCCTGGCGGACCAGCTCGCCGACACCGATGACCGAGAACAGCGAGGTGTCCT

>Halomonas maura CECT 5298 [NZ_JAUFPQ010000013.1:278309..278378], dnaA> ig dnaN>, 70 bp

WebSkew GCmin pos. 248336 (GCmin<>oriC1 51.3 kb; GCmin<>oriC2 25.5 kb; oriC2<>oriC1 25.7 kb)

>Halomonas nitroreducens 11S [NZ_RXNS01000004.1:c196831..197330], <rpmH oriC1 dnaA>, 515 bp

ATGTCCGACGCGGCGGCGTCGCTGGGCACAGCGAGATCATCGCTGCCGGAGGCCGTCGGAGACGGGGCGTCGCCGGCGTCACTGGAGATGGCCGGGGCCT
GCGGTGCCGTGGCCACCTGGCCGTAATCCTGATTCCACTGCACGACGATCAGGTAGGCGAGGACCGCCAGGGGAATCAGCAATAAGAGTCGTTTTACGTC
CATGAGGGTTCTGCCCGATGGGTGGTGAACATTCCGATGGCGCAGCGCAGGTCGCACACCCGACATACACGAAAGCCTGCCGAGGGCAGGCTAGCGTTCG
TCAGCAGAGCGTCTGCTTTCGGCGTGACGGCGTCAAGGATGGCGGCGAGTAGCGTGGTCATGCGCATGCTGGCTCCGCAACGGTGCCGACAGCGGTTGGC
TGGGCGTGTCATCGACGCTCGGTCATGAGGTCACCGGCCGATTGTCTCTCTTCCACCTTCCTCGCTTCCCGCTCCAGTCGTCGCCACATGCCGTGCAACT
GGCGATGTACGGTCGCATTGTCCAGTTCCTGAACGCCGCGTCTGGCGAGCAGCACGATATCCACGGCGGGCAGGCGATGCTGGCGGAGGCGAATGGATTC
GCGCGCGAGGCGTTTGAGGCGGTTGCGGTCGACGGCACGCCGGACGTTCTTCTTGCTGAAGATGAGCCCGAGGCGGGGATGCCCCAGAGAATTGGGGGTG
GCCAGGACCATCAAGCCCTTGCCGTGGACCTTGAAGGCAGCCGTATCGAAGACACGTCGATAGTCCCCGGCGTTGAGTAGTCTCAGTCGCCGGGGAAAGC
CCTGACAGGACACGCGAGGTCCGCGGTCAGGCGCTCAGGCGCTTGCGACCCTTGGCGCGACGACGCGCCAGGACGGCACGACCATTCTTGGTGGCCATGC
GAGCACGGAAGCCATGCGCGCGCTTGCGCTTGAGAACGCTGGGTTGAAAGGTGCGTTTCATAACTCGTTATTCCCACGTGGTTTTTGGAACAAATCGGCG

ATTCAGAGGCGCCCGGGTTCCGGTGGGGAATCCGGGTAGATTCGGCTCAAGACCGGAAATTGTAGTGATTTCGCGGCCCCAACGCAATTTTTCCGTGCCC
TCCGGGAGAGGTGCCGCGACGGCATGTCCCCGTTGCCGGCTCTTCTATTACTATTCAGGAAGTTCTATTGAACCTTGTTACTATTGATGGGGACGTTTGT
**TGTGGATAA**GTCGATTTCCCCTTTTTGAGACATCCGCTTATGAACGCCATAGGGCTGTGGGCAAGCGGCGAATGGCCGGCCTATAAGCTGTGGAAGGCCA
GTGGGTGAATGCTCGAGCTGTCCACACCCCCAAGGTTGCTCGCGTTCGACGCCCAGGGAGTCCCTGACTCGACAGCAGGTTTTCCACAGCGGCGGCGGAA
GCCGGCTGTCATGTGGATAACCCGGCGGCAGGACGATACAATGTGCGGTCGTTCTTCAACCGAATGGTGAGGTCGCGTGTCGCTCGCTCTCTGGCAACAA

TGTCTGGATACCCTGCAGGACGAACTGAACTCCCAGCAGTTCAATACCTGGATTCGTCCGCTGCAGGCGGAGGAAGGGGAATCCAATCAGCTGCGCCTGC
TGGCGCCCAATCGCTTTGTACGCGACTGGGTCAGCGACAAGTATGCCAAGCGCATCAGTGAGTTGATGCGGGAGCTGTCGCCGGCCAAGCCGCCCAAGGT
GGTGCTGACCGTGGGCAGTCGCCGGGCGGCTGCCCCGGCACCCCAGCCCCGCGAGCTGGGCGATCCGGTTTCTGCAGCGCCCCGGGCGCCGAGCGTGCAG
ACGCCTCCCCGGGGGGACGTGGCCGATGAGCGCGAGATCGATCAGCTCCGCGAGGAAGGGGAGGGCGCCCGCCGTCCCGGCGGAGAGCGGCAGGTCCAGG
TCGAGGGCAGCCTCAAGCACAACAGTGGCCTGAACCCCAACTTCACCTTCGAGACCTTCGTCGAGGGCAAGTCCAACCAACTGGCGCGCGCCGCTTCCCG
CCAGGTATCGGAGAACCCGGGTGGCGCCTACAATCCCCTGTTTCTGTATGGCGGGGTCGGCCTGGGCAAGACCCACCTGATGCATGCCGTCGGCAATGCG
CTGGCCGGCCGGCGCGAGAACGCCCGGGTGGTCTACCTGCACTCCGAGCGCTTCGTCGCCGACATGGTCAAGGCGCTCCAGCTCAATGCCATCAATGATT
TCAAACGCTTCTACCGCAGCGTCGATGCCCTGCTGATCGACGACATCCAGTTCTTCGCCGGCAAGGAACGCTCTCAGGAGGAGTTCTTCCATACTTTCAA
TGCCCTGCTCGAGGGCGGTCAGCAAATGATCCTGACCTCGGACCGTTATCCCAAGGAAATCAGTGGCGTGGAGGAGCGGCTCAAGTCCCGCTTCGGCTGG
GGGCTGACGGTGGCGATCGAGCCGCCGGAGCTCGAGACCCGGGTGGCGATCCTGATGAAGAAGGCCGATCAGGCCAAGGTCGATCTGCCGCATGACGCGG

>Halomonas nitroreducens 11S [NZ_RXNS01000004.1:c222545..223044], <mnmG oriC2 <COG1126 (glnQ), 536 bp

CCAGTCCGTGGATGGAGCGCACCACCTGCAGCTGAACATCGAAGGGCAGCGACGTCGAGATGCCGTTGGGGTAGAGCTCGTGGGTATCGAGTCCCTCGGG
CTCGATGAAGATCTGGTGGCTGGCCTTGTCCGCGAAGCGATGCACCTTGTCCTCGATGGACGGGCAATAGCGCGGTCCCACGCCTTCGATGGTCCCCGAG
AACATCGGCGAGCGATCCAGGTTGGCCTGGATGATCTCGTGGCTGCGCTCGTTGGTGTGGGCGATGTGGCAACTCACCTGTCGCGGATGCAGCTCCCGCC
GGCCCAGATAGGACATCACCGGCGTGGGGCTGTCGCCGGGCTGTTCCTCGAGGACCGAGAAATCCACGCTCTTGGCGTCCAGTCTCGGCGGCGTGCCGGT
CTTCAGCCGATCCACCCGGAACGGCAGGGCCCGCAGTCGCTCGGCCAGCGCATTCGAGGGCGGGTCACCGGCCCGGCCACCGCGACTGTGATCGAGGCCG
ATGTGGATAACCCCGCCGAGGAAGGTGCCTGTGCACAACACCACGCTCTCGCCCAGGAAGCGTACGCCGGTCTCGGTCACTACCCCGCGGACGGTGTCGC
CATCGACGATCAGGTCGCCGGCAGCCTGCTGGAAGAGCGTCAGGTTGGGCTGGTTCTCCAGCATGCCGCGGATCGCCGCCTTGTAGCGGACCCGATCGGC
CTGGGCGCGGGTGGCGCGCACTGCCGGCCCCTTGCGGGCGTTGAGCACCCGGAACTGGATGCCGCCCAGGTCGGTGGCCAGGCCCATGGCCCCACCGAGG
GCATCGATCTCCTTGACCAGGTGGCTCTTGCCGATCCCGCCGATGGCGGGATTGCAGGACATCTGCCCCAGGGTCTCGATGTTGTGGCTCAGCAGCAGGG
TCCGGCAGCCCATGCGGGCCGAGGCCAGGGCGGCTTCGGTCCCCGCATGACCGCCACCGATGACGATGACGTCGAAGCGGTCGGGGTACTCCAAGGTTCA

CCTCGTTCGGCGCGCCAGCGCCGATATTGCGAATCGGGGGTATGAAATCAGCCGCCCAGTATAACTGGCCTGTGGGTAATCGTCAGGGTTTTCCACACCC
GGACAGTTCGCGTGCGATTTCCCCGAAGACAGATAGACAAATACATAGAATAAGAAAGAGAAGTTCTGTTGTTGTAGTGACTATTGGTGTGGACAATTTC
**GGTGGATAA**GCATGTTTAGCCTTTTTTTATCAACAGCTTGAATTGTTGACCAGAACAAGGAAAAGCGGCGCGCGACCTGCGGAGGAGGTGTTCGGGGGCT
GTGGACGAAAGCCGGGGTTGCCCACAGCCGCGACGAGTCACCGCTTGTCCACCGTCCTCTCCCGCGCTTGTCACCGTGGCTGTGAACAACCGCCGCGTTT
GATCCCCAGGCCCGGCAGCATGGGCGCTGTGGGCGAATCGCCCGGGGCCTGTCGCAGCTCTTGTCCACAGGCAAAAAAAAGCCCTGTCGCGTACGACAGG

GCCTTGATTCAGGACTCGTGGGGCGGCCTCAGTGCTTCAATACCCGGGCCAGGAAGTTCCGGGTGGGCTCTTCCCGGGGTGAATCGAAGATCGTCTCGGG
AGGGCCCTCCTCGGCAATTTCCCCCTTGTGGATGAAGATCACCCGATCGGCCACCTCGCGGGCAAAGCCCATCTCGTGGGTCACGATCATCATGGTCATG
CCTTCGTTGGCGAGCTCGCGCATGGCATCCAGTACCTCGCCGATCATCTCCGGATCCAGTGCCGAGGTCGGCTCGTCGAAGAGCATCAGCCGGGGTTCCA
TGGCCAGCGCCCGGGCCAGGGCCACCCGCTGCTGCTGGCCGCCGGACAGCTGGCCGGGATACTTGACGGCCTGGTCGCTGATGCCGACCCGCTCGAGCAG
TCGCTCGGCGATCTCGGTGGCATCGGCCCGGCTCCAGCCGCGGACCTTCATGGGGGCCAGGGTCACATTGTCGAGCACGCTGAGGTGGGGGAAGAGGTTG
AACTGCTGAAACACCATGCCCACCTCGGTGCGGATCTTCTGCAGGGCCTTGGTGGCCTTGCCGTGGGGCAGCAGCGGGTTGCCATCCACCTCGAGGCTGC
CCTTCTGGAATTCCTCCAGGCCATTGATGCAGCGGATCAGCGTGGACTTGCCCGAGCCACTGGCGCCGATGACCACCACCACCTCGCCGGGGGTGACCTC
GAGGTCGATGTCCTTGAGGACATGCAGACTGCCGAAGTGCTTGTTGACCTTGTCCAGGCGCACGATGGGCTCGCCGCCGGACTTGGACGTCTCTTGGTTC
ATGTCTTGCATTCCTTATTGTCCGACCAGGCCCTTGCGCTCCAGGAAGCGAAGCGCCACCGACAGCGACAGGGTGATCACCAGGTAGAGCATGGCGACCA
TCAGGTACACCTCCAGGGCGTTGAAGGTGGTGGCGATGTAGACCTGGCCCTGGCGAACCAGCTCGCCGACACCGATGACCGAGAACAGCGAGGTGTCCTT

>Halomonas nitroreducens 11S [NZ_RXNS01000004.1:c195321..195390], dnaA> ig dnaN>, 70 bp

WebSkew GCmin pos. 177356 (GCmin<>oriC1 19.1 kb; GCmin<>oriC2 1.8 kb; oriC2<>oriC1 17.3 kb)

>Halomonas halocynthiae DSM 14573 [NZ_AUDZ01000013.1:196199..196698], <rpmH oriC1 dnaA>, 521 bp

GGCGGCCTGCACAGGAACCTGCCCAGGAAGAGAGCCTGCAAGACCATCGTTGCTGGCCGGCTGACTGGGGGCTGATAACGCGCTATCAGCTGACCCCGAG
GGCTCACTGTTTTGCTGGTTGGTTACCGTCGGTGGCGCCTGATGCGCGGTGACTGGCTGGCCGTAGTCCTTGTTCCACTGCACGATGATCAGGTAGGCAA
GAAGCGCCAGTGGAATCAGCAGAAATAGTCGTTTTATGTCCATGAAGGGTCTGCCCGATGGGGGAGTGAACATTCCGATGGCGAAACATAGCTCGCACAC
CTGAGATGAACAAAAGCCTGCCAGGGGCAGGCCAATGTCAGGTACTGAGCGTCGGTGCTTAACACGGCTTCGTCAAGCAGTACATCAACGTGAATGACAA
AACGTGAACGGCGAAGGCTTGTGGTCATGGCACTCAGATAGTCAACGGCTGTTTTCGAGTTTCTGTAATTCTCGATCTAAGCGTCGCCACATGCCATGAA
GCTGTCGATGAACCGTTTCGTTATCCAGCTCAACAAGTCCACGCCGTGCCAGCAGCACAATATCCACAGGTGGGAGCTGGTGCTGGCGAAGCCGGATGGA
CTCTCGTGCGATACGTTTGAAGCGATTTCGGTCGACGGCGCGGCGCACATTTTTCTTGCTGAAAATCAAACCAAGGCGCGGATGCCCCAGGTTGCTTGGT
GCTGCCAGTGCCATCAAGCCTTTGCCATGTACCTTGCAGGTTGCCGTATCAAATACGTGACGATAATCACCGGCGGTCAGCAATCGCAGCCGCCGGGGGT
AGCCCTGACAGGACACGCGCAATCAGCAATCAGGCGCTGAGACGCTTACGTCCTTTAGCGCGACGACGCGCCAGGACAGCACGGCCGGCTTTGGTGGCCA
TGCGAGCGCGGAAGCCATGGTTGCGCTTACGCTTGATAACGCTGGGTTGAAATGTGCGTTTCATAACAGGTCATTCCCACAAGTAGGTTTCTGAAAAATC

GGGTATTGCATTGGTGTTGGGTTCCTTTCGGAATCCAAACAGATTCGGTCAAGGACCGGAGATTGTAGAGACTTGCGCGGCGCGGCGCAATTCTTCTGAT
GAGATGAGAGGGGTTTCTGTTGTTGTCCAGAATTACCCTTAATAATAACTATTATATAGAAGTTCTGTAATACCTTGTTATTATGTATGGGGATAAAATC
**TGTTGATAA**TCTATTTAATGTTTTTTAAAACAAAAGGTTGAGCTTGTGATTTCTCTGTGAAAAAGCGTGTGGCAAGCCGCTGGAAAGCTGTGGATCATCG
GTGAATAACATGCGATGTTACCCACAGCGCTAACAATGCTGACAGACCATCACCATAAAATACCTGTCTCTATTGCCACTTTTCCACAAGTTGTCGCCGG
TTCGCTGGGTGTGGATAACTGTGCTTTTGAGCGATACAATAACCGCCGTTCTTAAATTCGCTTTTTAACCCAGGGGTGCGGCCGCGTGTCGATTGTACTT

TGGCAACAATGCCTGGAAACCCTGCAGGATGAGCTCAGCTCACAGCATTTCAATACTTGGATTCGTCCTCTTCAAGCAGAGGAAGAAGAAGGGGCTAATC
AGTTGCGACTGTTGGCTCCCAATCGTTTCGTGCGCGACTGGGTGTGTGACAAATATGTGCAGCGCATCACGGAGCTAATGCGTGATTTTTCATCAGGTCA
GCCACCTAAATTGGTCGTCGTGGTTGGTAGCCGCCGTGCTGCTGCTCCGCCATCTGCCCCGATGCCACGCGAGCTGGGTCAGCCTGTATCTGCTACACCT
AGCGCTTCTTCTATACACACTCCAGCAGTGGGTGGTGGTTCTGATAGCAGCCGTGATGCAGGTAGCCGGCCCGCGAGGGAGGGCACGAGCAGTGAGTTCC
AGCGCCCTGCTGAGCGGCAAGTGCAGGTAGAGGGCAGCATCAAACATAACAGTGGGCTGAATCCTAACTTTACTTTCGATACTTTTGTCGAAGGTAAGTC
GAACCAATTAGCAAAAGCCGCTTCACGCCAGGTGTCAGAGAATCCTGGTGGTGCGTATAACCCGTTGTTTTTGTATGGTGGTGTTGGTTTGGGTAAGACC
CACCTGATGCATGCTGTAGGTAATGCACTGGCCACTCGGCGCAGCAATGCTCGGGTTGTCTATCTGCATTCTGAGCGTTTTGTGGCTGATATGGTCAAGG
CGCTGCAGCTGAATGCCATTAACGACTTCAAGCGCTTTTATCGTAGTGTCGATGCGCTGTTGATAGATGATATTCAGTTTTTTGCTGGCAAAGAGCGGTC
TCAGGAAGAGTTTTTCCACACCTTTAACGCGCTGCTTGAGGGCGGCCAGCAAATGATTCTTACCTCTGACCGCTATCCCAAAGAGATCAGTGGAGTAGAA
GAACGTCTGAAGTCGCGGTTTGGCTGGGGGTTGACGGTGGCAATTGAGCCTCCTGAGCTTGAAACCAGAGTCGCAATCCTGATGAAAAAGGCCGACCAGG

>Halomonas halocynthiae DSM 14573 [NZ_AUDZ01000013.1:178897..179396], <mnmG oriC2 <COG1126 (glnQ), 542 bp

TTTTCCAAACCACGGATAGAGCGAACGACTTGCAGCTGCACATCAAAAGGAAGTGAGGTGGAAATACCATTGGGGTACAGCTCGTGGGTGTTGAGACCTT
CGGGTTCAAGAAATATCTGATGACTGTCTTTATCAGCAAAACGATGAACCTTGTCTTCGATCGATGGGCAGTAACGTGGCCCAACGCCCTCAATAACCCC
ACTAAACATAGGTGAGCGGTCAAGATTATCACGAATCAGCTGATGACTATGGGCATTGGTATGCGTGATATGGCAGCTGACTTGACGTGGGTGCATCTGT
TGGCTACCCATGTACGACATTACTGGCGTTGGCGTATCACCAGGTTGTTCGTCAAGTTGTGAAAAATCTACTGACTTGGCATCAATACGTGGAGGCGTGC
CAGTTTTTAATCGATCAACCCGTAAAGGCATTGCACGTAATCGTTTTGCCAGTGCATTGGAGGGCGGATCACCAGCACGACCAGCATGAGAGGTATCTAA
CCCTATGTGGATAACACCACCAAGGAAAGTCCCTGTGCATAACACAACCGCATGGGCATGAAAGCGAATACCTGTTTCGGTGACAACTCCGGTCACCCTG
TCTTGATCAACTATCAGGTCGCCAGCAGCTTGTTGAAAAATCGTTAAATTCTGTTGGTTTTCCAAGGTATTACGGATAGCAGCTTTATACAGTGCACGAT
CAGCCTGGGCCCGGGTCGCACGCACTGCTGGCCCTTTACGCGCATTGAGTACACGAAACTGAATACCACCCGCATCGGTAGCCAGCCCCATAGCACCACC
AAGTGCATCAATTTCTTTGACCAAATGGGTTTTGCCAATGCCACCAATAGCGGGATTACAGGACATTTGCCCGAGTGTTTCGATGTTGTGTGTCAGTAAC
AGTGTCTGACTACCAGCTCGGGCGGCAGCTAAAGCTGCCTCTGTACCTGCATGACCACCACCAATGACGATGACGTCGAAGCGGTCGGGATAATCCAAGA

AACACCTCGTTTGGCATCAAAGCCAATGCCTAGGAAAAATGAAATTTAGCTCGGCAGTATAACTGCTCCTGTGGGTAAACGTCAGGGTTTTCCACACAGG
GCGACTCATGCTGTTAGCCTGTGTTTTTGATTACAATATAAAAATAAATATATAAATAAGAAGTTCTATTGATGTTGTAATTATTAGACATGCAAAAATC
**TGTTGAAAA**CTCAATATATTTCTTCTATATCAAAAAGATAGCTTGTTGTTTTATAGGTTTGAAAGGTGCTTGAATCTTGCGCGTAGCTTGTGTGGTGGCT
GTGGGTAAAAAAGAAGGTTATGCACAGCGATGACTGTCCCGCAGTTGCCCACAGAGCTTGAGATGCTTTTTCCTTACACCTGTGAACAATTAGCTCGTCC
GTTCGTAAAATAGCTAGAGCAGGCGGTCTGCCTGACGTGAGCTGAGTTCATTCGGTCATCATAAAAAATATTATCCACAGGCATAAAAAAGCCCTGTCGT

AACCGACAGGGCTCAATATGGTGCGTGAACAACCAGAAAATTAGTGCGTCAGTACTCGAGCAAGAAAGCTTTGTGTGCGTTCATTCTGAGGGTTATCGAA
CACGTCTTCTGGTGTGCCTTGTTCAACAATTTCGCCTTTGTGGATGTAGATCACACGATCGGCAACCTCACGGGCAAAGTTCATTTCATGGGTTACAATC
ACCATGGTCATCCCCTCGCGGGCTAACTCACGCATAGCATCGAGCACTTCACCAATCATTTCAGGATCAAGTGCTGACGTGGGCTCATCAAACAACATGA
GCCTAGGTTCCATTGCTAAAGCTCTGGCTAGTGCCACACGTTGCTGCTGGCCGCCTGACAACTGGGTTGGGTACTTGTCGGATTGATCAGCAATTCCCAC
TCGTTCGAGTAAGCGGTTGGCTGTTTGGGTCGCTTCGGCACGGGGAGTATTGCGCACTTTCATCGGTGACAGGGTAACGTTGTCCAGTACGCTGAGGTGA
GGGAAGAGATTGAATTGTTGGAATACCATGCCAACTTCAGTACGAATTTGTTGTAGGGCTCGATTGGCTTTTCCATCAGGGGAAAGACTGTTTCCGTCTA
CAGTAAGCTGCCCTTTCTGAAACTCTTCTAGCCCATTGATGCAGCGAATCATGGTGGACTTACCTGATCCGCTGGCACCAATAATCACAACGACTTCACC
AGGAGTCACTTCAAGATCAACATCTTTCAGAACATGCAGTTTGCCGAAGTGTTTGTTGAGCTTCTGCATGGATACGATAGGTGTAGCGTTAGTATCAACC
GAGGATCTGCTTAGTGTGGTGTCTTTATTCATTGTCCAACTAGCCCTTTGCGTTCAAGCATGCGTAGTGCGATTGAGAGTGTCAGAGTAATGATCAGGTA
CATGATGGCAACCATCATATACACCTCGAGTGCATTGAACGTTGTTGCAATATAGATCTGTCCCTGACGTACCAGCTCACCAACGCCTATCACGGCAAAA

>Halomonas halocynthiae DSM 14573 [NZ_AUDZ01000013.1:198166..198247], dnaA> ig dnaN>, 82 bp

WebSkew GCmin pos. 60264 (GCmin<>oriC1 29.8 kb; GCmin<>oriC2 9.4 kb; oriC2<>oriC1 20.4 kb)

>Halomonas cerina CECT 7282 [NZ_JACHXP010000005.1:89802..90301], <rpmH oriC1 dnaA>, 505 bp

GGCGCTCGGGGCCGCCGGTACCGTCAAGTCGTTGTCGCCGGGTGCCGCTGCATCGTCCCGAGCGGCATCACCGGTGATGCCGGGGGCCTGCGGTGCCGTG
GTCACCTGACCGTAGTCCTGATTCCACTGCACCACGATCAAGTAGGCGAGAATCGCCAGGGGAATCAGAAGGATGAGTCGTTTTACGTCCATGAAGGGTC
TGCCCGATGGGTGGTGAACATTCCGATGGCACAGGCGGGATAAGAAGCGCGGGTCCGACATACACGAAAGCCTGCCGGCGGCAGGCTAAATGGCGTCGTT
GGCGTAACGGCGTTCGGCATGACAGCATCCAGGGAGCTAGTAGCGAATGTCGTGATCATGCCCTTCCGGTCTCCGCTATCCGCCGACAGTCGGGAATGCC
GATCATCGCTGTCCCGATGAGGCTTGCGCCGGAGGATCGACCACGTCGGATCGACCGGCCGGCTGCCTGTGTGCTTCGCGATCCAGGCGGCGCCACATGC
CATGCAACTGGCGATGCACGGCCTCGTTGTCCAGTTCATGAAGGCCACGTCTGGCCAGCACCACGATGTCCACGGATGGGAGCCGATGCTGCTGCAGCCG
TACCGATTCACGCACGAGGCGCTTGAGACGATTGCGGTCGACCGCCCGGCGAACGTTCTTCTTGCTGAAGATCAGGCCAAGGCGGGGGTGTCCCAGTGAA
TTGGGGGTGGCCAGGGCCATCAGCCGCTTGCCATGCACCTTGTAGACCGCCGTATCGAAGACGTGCCGATAGTCCCCGGCCGTCAGCAGTCGCAGAGACC
GGGGGTAGCCGTGACTGGACACGCGCGGAACGCGCTCAAGCGCTCAGGCGCTTGCGGCCCTTGGCACGACGGCGTGCCAGCACGGCACGACCGTTCTTGG
TGGCCATGCGGGCGCGAAAGCCGTGGACACGCTTGCGTTTCAGAACGCTAGGTTGAAATGTACGCTTCATAACTCGTTATTCCCACGTGGTTTTTGGAAC

AGTTGGATAATCTCAGAGGCGCCCTGGCCGCGACAGGAGGCCAGGCTGGAGTCGGCTCAAGACCCGAAATTGTAGTGATTTCGGGGCCTCGGCGCAATTT
TTCTTGTGCATCGTCCGTTGCGCCGGGCAGCCATCGTGTTTCCCGGGACGCTGTCTTCATCATAACCAACGGTTTTTCTGTGACAGGGTGTTATTAATCA
**TGGGGATAA**TGGCTGTTGATAAGTGAGCTAT**C**TTCTTTTTCTCCAATGAGTTGTCTTCGCGAAAAGGTTGTGCATAAACAGGGGGCCCCGACCGGCTTCC
TGTCAGTGATCGGTGAATAGCGGGCCTTTTTATCCGGTTCTTCCCGATTATCCCCAAGTTTCCGTACGGGTTTGTCAGGTTGTCCACAGAAAGCCAGCCA
TCATGCCAGGTGTGGATAACCTGCTGGTGGGGCGATACAATGCTCGGCCGTTCTTCAGCCGATTGGTGAGGTCGCGTGTCGCTTGCTCTCTGGCAACAGT

GTCTGGATACCCTGCAGGATGAACTGAGCTCTCAGCAGTTCAATACCTGGATCCGCCCCTTGCAGGCGGAGGAAGGGGAGTCCAACCAGCTGCGCCTGCT
GGCGCCCAACCGGTTCGTGAGAGACTGGGTCAGCGACAAGTACGCCAAGCGCATCGGTGAGCTGCTGCGCGAACTGGCGCCGGCAAAGCCGCCCAAGGTC
GTGCTGACGGTGGGAAGCCGCCGGGCAGCGCCCAACCCCCAGCCGCGTGAGCTCGGTCATCCGGTATCAGCCACTCCGCGCCGGCCGGTCACACCCGCCG
TCCAGACGCCGCCGCGTGGTGACCTGGCCGACGAGCGGGAGGTCGACCGCCTCCGTGAGGAGGGCACGACGAGTGCCCGTAGCGGCGAGCGACAGGTCCA
GGTGGAAGGCAGCCTTAAGCACAGCAGCGGGCTCAACCCCAACTTCACCTTCGAAACCTTCGTCGAAGGCAAGTCAAACCAGCTGGCCCGTGCCGCTTCC
CGCCAGGTCTCCGAGAACCCTGGTGGCGCCTACAATCCCCTATTCCTGTATGGCGGGGTCGGTTTGGGCAAGACCCACCTGATGCATGCCGTCGGCAATG
CGCTGGCGACCCGTCGCGAAAATGCTCGGGTGGTCTACCTGCACTCCGAACGCTTCGTGGCCGACATGGTCAAGGCGCTGCAGCTCAACGCCATCAATGA
CTTCAAGCGCTTCTATCGCAGCGTCGATGCCCTGCTGATCGACGACATTCAGTTCTTCGCGGGCAAGGAGCGCTCCCAGGAAGAGTTTTTCCATACCTTC
AATGCCCTGCTCGAGGGCGGCCAGCAGATGATCCTGACCTCGGACCGCTATCCCAAGGAGATCAGCGGCGTGGAGGAGCGCCTGAAGTCGCGCTTCGGCT
GGGGGCTTACCGTGGCGATCGAGCCACCCGAGCTCGAGACCCGGGTGGCGATCCTGATGAAGAAGGCCGACCAGGCCAAGGTGGACCTGCCGCATGATGC

>Halomonas cerina CECT 7282 [NZ_JACHXP010000005.1:69431..69930], <mnmG oriC2 <COG1126 (glnQ), 531 bp

TCTCGAGGCCCTCGATCGAGCGCACCACCTGCAGCTGGACGTCGAAAGGCAGCGAGGTGGAGATGCCGTTGGGATAGAGCTCGTGGGTGTCCAGGCCTTC
GGGCTCGATGAAGATCTGGTGACTCGCTTTATCCGCGAAGCGGTGCACCTTGTCCTCGATGGACGGGCAATAGCGCGGTCCCACGCCCTCGATCACCCCG
GAGAACATCGGCGAGCGATCCAGGTTGGCGTGGATGATCTCGTGAGTCCGCTCGTTGGTGTGAGCGATGTGACAACTCACCTGGCGGGGATGCATCTCCC
GGTCGCCGAGATAGGACATCACCGGGGTTGGAACATCACCGGGCTGCTCCTCGAGCCTCGCGAAGTCAATGTTCTTGGCATCGAGGCGCGGCGGGGTACC
AGTCTTCAGGCGATCGACCCGGAACGGCAGGGCCCGCAGGCGTTGGGCCAAGGCGTTGGAGGGTGGATCCCCGGCACGTCCGCCGTGACTGTTGTCCAGG
CCGATGTGGATAACCCCGCCAAGGAAGGTGCCTGTGCACAACACTACCGTCTCGCCGAGGAAGCGCACGCCGGTCTCGGTGACCACGCCCCGCACGCTAT
CCCCTTCGACGATCAGGTCGCCGGCGGCCTGTTGGAAGAGCGTCAGGTTGGGCTGGCTCTCTAGCATGCCGCGGATCGCGGCCTTGTAGCGGACGCGATC
GGCCTGGGCCCGAGTCGCTCGGACGGCAGGGCCCTTGCGGGCGTTGAGCACCCGGAACTGGATGCCGCCCCGGTCGGTGGCCAGGCCCATGGCCCCGCCG
AGCGCATCGATTTCTTTGACCAGATGGCTCTTGCCGATCCCGCCGATGGCGGGATTGCACGACATCTGGCCCAGGGTCTCGATGTTGTGGGTCAGTAGCA
GGGTCCGGCAGCCCATGCGGGCCGAGGCCAGGGCGGCTTCGGTGCCGGCATGGCCGCCGCCGATGACGATGACGTCAAAGCGGTCGGGATACTCCAAGGT

TCACCTCGTTCGGCGTCGCCGCGCCGATGTTGCGGTCGGGGATGGAATCGGGCGCCAGTATAAACCTCCTGTGGGTAATCGTCAGGGTTTTCCACACCCG
ATTCCTACCTGCTACGACAGCTTCTTGCTTCGTATAATCAATAATAAAGAATATAAAGAGAAGTTCTGTTGTGGTTGTAACTACTGGTGTGGACACTTTC
**TGTGGATAA**ACATGGCTAATGGCCTGACAGCAGGCAGTTGTATTGTTGATTAGCGTGGGGACAAACAGCGGATGATCGGCGGACACCGTGTCCATGAGCT
GTGGATAGAATGGCACCCTGTCCACAGCCGTCCGAGCTCACCGGTTGTCCACCGCTCAATACCTCACTTGTTACCGCGGCTGTGAACAACCATCATGATG
CATTCTGCGTCCCGCCCTCATGGGCCCTGACGCTGGATCGGTGCGACTCGTCGGACAATCTTGTCCACAAGCTAAAAAAAAGCCCTGTCGAGATCGACAG

GGCTTTTTCAAGGGCGCTAAGCGGGGCCTCAGTGCTGCAACACCCGGGCCAGGAAGTTTCGGGTGGGGGCTTCCTTAGGGGAATCGAAGATCGTCTCGGG
GGGCCCCTCCTCGGCGATTTCCCCCTTGTGGATATAGATGACCCGATCGGCCACTTCCCGGGCGAAGCCCATCTCGTGAGTCACGATCATCATCGTCATG
CCTTCCTTCGCGAGTTCCCGCATGGCATCCAAGACTTCACCGATCATCTCCGGGTCCAATGCCGAGGTCGGCTCGTCGAAGAGCATCAGTCGAGGCTCCA
TGGCCAGCGCCCGGGCCAGGGCCACCCGTTGCTGCTGGCCGCCGGAGAGTTGGGTCGGGTACTTGGCCGCTTGATCGCTGATGCCCACGCGCTCGAGCAG
CCGCTCGGCGATCTCGGTGGCATCGCTGCGACTCCAACCACGCACCTTCATCGGCGCCAGGGTCACGTTGTCCAGCACGCTAAGATGGGGGAAGAGGTTG
AACTGTTGGAAGACCATGCCCACCTCGGTGCGAATGCGTTGCAGCGCCTTGGTCGCCTTGCCGTAGGGCAGCAAGGGGCTGCCATCGACCTCGAGATGGC
CCTGCTGGAACTCCTCGAGGCCGTTGATGCAACGGATCAGGGTCGACTTGCCCGAGCCACTGGCCCCGATGATCACGACCACCTCGCCGGGGGTCACCTC
GAGGTCGATGTCCTTCAGGACATGCAGGCTGCCAAAGTGCTTGTTGACCTTCTCCAAGCGCACGATGGGGTGGGCGCCGGAGGGAGACGTATCTGGGTTC
ATGTCGCGCTTTCCTTATTGTCCGGCCAGGCCCTTGCGTTCGATGATGCGCAGCACCAACGACAGGGAGAGCGTGATCGCCAGGTAGAGCATAGCGACCA
TCAGGTAGACCTCCAGGGCGGTGAAGGTGGTGGCGATGTAGACTTGACCCTGACGGACCAGCTCGCCGACCCCGATCACCGAGAACAGCGAAGTGTCCTT

>Halomonas cerina CECT 7282 [NZ_JACHXP010000005.1:91744..91815], dnaA> ig dnaN>, 72 bp

WebSkew GCmin pos. 4757486/0 (GCmin<>oriC1 0 kb; GCmin<>oriC2 20.2 kb; oriC2<>oriC1 20.2 kb)

>Halomonas huangheensis BJGMM-B45 [NZ_CP013106.1:4757240..4757486;1..253], <rpmH oriC1 dnaA>, 616 bp

ATAATCCTGGTTCCACTGCACCATGATCAGGTAAGCGAGCACCGCCAGGGGGATCAGCAAAAGTAGTCGTTTTACGTCCATGAGGGGTCTGCCCGATGGG
AAGTGAACAATCCAGTGGCGTCCGTCTGGGCGCCGACGCCTGGGAGAAATACACCAAAGCCTGCCAGAGGCAGGCCAGGGTCCGTACGCAAACATTCTGC
GATGAGTGAGGTGACGTCAATGCTATCGACGTTGGGCTCGTACATGCGCATAGATGGTGTCATCGCATCGATGGGATGCCGCACAGCTTTATCCGCGGCG
CTGGTGAGCGGCGGAGGAGTGGGAGGCCGTCGAGGAGCGAGAGGGTCGGGTCGACTTCTCGGCCTCGCGCTCGAGGCGGCGCCACATGCCGAACAGCTGA
CGATGCAAGGTGTCGTTGTCGACCTCGTTGACGCCACGTCTTGCCAGCAGAACGATATCCACACCGGGAAGTCGATGCTGAGAATGGCGTACGGATTCGC
GGGCCAGTCGCTTGAGACGATTACGATCAACAGCCCTGCGTACGTTCTTCTTGCTGAACACCAGGCCAAGGCGCGGGTGCCCCAGGTCATTAGGGGTAGC
CAGCGCCATCAGGCCCTTGCCATGGACCTTGTATACTGCAGTGTCAAAGACACGGCGGTAATCCCCGGCAGTCAGCAAGCGATAGTGCCGGGGAAAGGTC
AGACAGGTCATATTTGACCTGCGATCAGGCACTCAGGCGCTTGCGGCCCTTGGCACGGCGACGTGCCAGAACGGCGCGGCCGTTTTTGGTGGCCATGCGA
GCACGGAAGCCGTGCGCGCGCTTGCGCTTGAGAACGCTCGGTTGGAATGTGCGCTTCATAACTCGTTAATCCTGAATGGTTTTTTGGATCGGGTAGACAA
CCAAAGGAGCCCGAATTGGCAACGGGCTCCTCGGGGGTATTCGGTTCAAGACCGGACATTCTAGTGATTTCAAGCCTTACGCGCAATTGCTGAGCGCATT

TTTACCTCGACACCCTCTTTCCGAGAGCAGTTTGACGATGAGTCGAGGGGTATAGCTCTCGCCCCTGGCAACGACATTGGGGAAGCCGCTCAAGCTAGCG
GACCTGGAGCGCGGAGTGGTTGGTGATGTATTCGTATGACCATAGTAATTACTAATCTAAAGTATTGAATATACTTGTGATTATTACTGGGGATAGTTTT
**TGTGGATAA**CCGTAAAAGTTTCTTTTAATTCAACATGCTGCATTGCACATAAAAGTGTGGGAAAAACGTTGAACACAGGTGGGAGGACGGTTGATGAGCT
GTGCGGAAAGCGCATCATTATCCACAGGCCTGAAATCATCCTTGGTTTGTCCCCAGTCGTTGTGGGTAGTTTCCGACTGGTTTTCCACAGTACTCAGACA
GCCCTTCTGTCGTGTGGATAACTCGTCTCTGGGGCGATACAATATGCGGTCGCTCTTCCAATCAAGGTAGCTCGTGTGTCGCTCGCTCTTTGGCAACAAT

GTCTGGAAACCCTTCAGGATGAACTGAGTTCGCAACAGTTCAACACCTGGATCCGACCGCTTCAGGCGGAAGAGGAGGCCAATCAGCTGCGGCTGTTGGC
GCCAAACCGGTTTGTGCGTGACTGGGTCAACGACAAGTACGCCAAGCGTATCAGTGAGCTGATGCGTGAGCTTGCGCCTGCCAAACCGCCCAAGGTTCTG
CTGGTGGTAGGCAGTCGGCGTGCTGCGGCCCCAGCCCCTCAGACTCGTGATCTCGGAAACCCCGTGTCGGCTGCACCGACGCGGGCTCCTGCTGTGCGCA
CGCCTCCCCGAGGGGATGTGGCGGATGAGCGCGAGATTGATCGCCTGCGTGAGGAAGGAGAAGGCCCTGTACCCACTCGTCCGCCCAGTGGCGAGCGTCA
GGTGCAGGTGGAGGGCAGTCTCAAGCACTCCAGTGGTCTCAATCCCAATTTTACCTTCGAAACTTTCGTCGAAGGTAAGTCAAACCAGTTGGCACGTGCT
GCATCTCGCCAGGTGTCGGAAAACCCCGGTGGAGCCTACAATCCTCTGTTCCTCTATGGTGGTGTGGGCCTCGGTAAGACCCACCTGATGCATGCTGTGG
GTAATGCGTTGGCCGGGCGCCGAGAGAACGCGAGGGTGGTGTATCTGCACTCCGAGCGTTTTGTAGCCGATATGGTCAAGGCGTTACAGCTCAATGCCAT
CAATGACTTCAAGCGCTTCTACCGTAGTGTTGATGCGCTTTTGATCGATGACATTCAGTTCTTTGCTGGCAAGGAGCGCTCCCAGGAGGAGTTCTTCCAC
ACCTTCAATGCCTTGCTTGAAGGGGGCCAGCAGATGATTCTGACCTCGGATCGTTACCCTAAGGAAATCAGTGGTGTCGAGGAGCGTCTCAAGTCGCGCT
TCGGTTGGGGACTGACAGTTGCCATTGAGCCTCCCGAGCTGGAGACACGTGTCGCGATCCTTATGAAGAAGGCCGACCAGGCCAGGGTCGACCTGCCGCA

>Halomonas huangheensis BJGMM-B45 [NZ_CP013106.1:4737019..4737518], <mnmG oriC2 <COG1126 (glnQ), 643 bp

TCCAGTCCTACCATCGAGCGCACTACCTGCAACTGCACGTCGAAGGGCAGTGAGGTAGAAATACCATTTGGGTATAGCTCGTGAGTATCGAGTCCTTCCG
GCTCGATAAACACCTGATGACTCGATTTGTCGGCGAAGCGATGCACCTTATCCTCGATGGACGGGCAATAACGCGGGCCAACACCCTCGATGACACCGGA
GAACATCGGCGAACGATCCAGGTTGGCGAAGATGATCTCGTGACTGCGCTCGTTGGTATGCGCAATGTGGCAGCTGACCTGACGCGGATGCATCTCGCGG
GAGCCGAGGTAGGACATCACCGGTGTTGGGGTATCGCCCGGCTGCTCCTCGAGGCGTGAGAAATCCACACTACGCGCATCGATACGCGGCGGCGTACCGG
TCTTGAGCCGGTCGACGCGAAATGGCAGTGCGCGAAGCCGGTCTGCCAGGGCATTGGAAGGCGGATCTCCGGCACGCCCACCACGACTCGTTTCGAGACC
AATGTGAATGATTCCACCGAGGAAGGTGCCCGTGCACAGCACCACCGTTTCGGCCAGGAAACGCACCCCGGTCTCTGTCACCACACCGCGGACGGTGTCT
TGATCAACAATCAAGTCACCGGCGGCCTGCTGGAAGATGGTCAGATTGGGTTGGTTTTCCAGTATCGAGCGAATTGCGGCCTTGTAGCGAACGCGATCAG
CCTGAGCCCGTGTTGCACGTACCGCAGGGCCCTTACGGGCATTGAGCACACGAAACTGGATGCCACCCAGGTCTGTCGCCAGACCCATGGCACCGCCCAG
GGCATCGATCTCCTTGACCAGATGACTCTTGCCGATCCCGCCAATGGCCGGATTGCAGGACATCTGACCCAGGGTCTCGATGTTGTGTGTCAGCAGCAGG
GTTTGGCAACCCATTCGGGCAGAGGCCAGAGCAGCTTCCGTTCCGGCATGGCCACCGCCGATGACGATAACGTCAAAGCGGTCGGGATACTCCAAGGTGC

ACCTCGTTTGGCGTTCAACCGCCAGTGTGGAGAAAGTGTCAGGAAAACAAGCCCGCCAGTATAACGTCCCTGTGGGTAAACGTCAGGGTTTTCCACACCC
AGGCCAATCACACGTCGGACAACGACCTTTCTTTATAAATAAACATAGGTAAATAAAGAGAAGTTCTGTTGTGGTTGTTACTAACTGGTGTGGGCATTTTC
**TGTGTACAA**CGCTGATATCCTTTTATAAAACAGCAAATTAAAATGTTGATAGAAATTTGGACAGCGCGGTGATGGGGTGCGTATGAGTTGTCTGGAGCCT
GTGGATGAAAAGCGGACTTGCCCACAAGCGTATCGGTCCACCACTTATCCACCTGGCTCATGAACATTTGTCACCGCACCGGTGAACAAGTGACGAAAAT
GCCCACAGCGCCCGGATTCATGGGGCTGCAGCGTACTGAACATTTTTTTATCCACAGCCGGAAAATGACACAAAATGGCCTGTCGATAACGTAATGCGC

CTCGATATGGAGCAGAAAAAGTGGCTGGTTGGCGCTGACCTGACATCATCCAGCGCGTAGCTGATGGCCGATGCCCAGAAAAAAACCCCGCCGTGAGACA
CGACAGGGCTTCGAGGACAAGAGCGGTGAGAGAGTTCGTCAGTGCTTGAGAACCCGTGCCAGAAAGTTCTGTGTACGTTCGTTGCGAGGGTTGTCGAACA
CGTCTTCAGGCACGCCCTGTTCAACGATCTCCCCCTGGTGAATATAGATAACCCGGTCAGCCACTTCCCGAGCAAAGCCCATCTCATGGGTAACGATGAT
CATGGTCATGCCATCTTTTGCCAGCTCTCGCATGGCATCCAACACTTCACCAATCATCTCGGGATCCAGTGCTGAAGTAGGCTCATCGAACAGCATCAGC
CGTGGTTCCATGGCCAATGCCCGTGCCAGTGCAACGCGCTGCTGCTGACCACCAGACAGCTGAGAAGGGTACTTGTCGGCTTGATCACGGATGCCGACAC
GATCCAGCAGGCGTTCGGCGATGGTGACAGCATCTGTTCTGGCGAGACCACGTACTTTCATCGGCGCAAGGGTGACGTTGTCACGCACGCTGAGATGCGG
GAAGAGGTTGAATTGCTGGAAGACCATGCCAACCTCGGTACGAATGGTCTGGAGGGCGCGGGATGCTTTCCCGTGGGGCGCCAGGTCGTGGCCGTCTACA
CGCAACTCCCCGCTCTGGAACTCCTCCAGTCCGTTGATGCAGCGAATCATGGTTGACTTGCCGGAACCGCTGGCTCCGATCACTACCACCACTTCGCCCT
GGGCGACTTCGAGGTCGATATTCTGGAGCACATGCAGGCTGCCGAAGTGCTTGTTCAGGCTGCGCATTTGTACGATAGGTTCGCCACTCGCCTGTTGTGC
GCTCGTGTCGAGTTTGATGTCGTTCATATCCAATCCCATGAAGTATCAGCACACAGAAGGGCTATTCACCCACCAGACCCTTGCGTTCGAGCATGCGCAG

>Halomonas huangheensis BJGMM-B45 [NZ_CP013106.1:1702..1775], dnaA> ig dnaN>, 74 bp

WebSkew GCmin pos. 301804 (GCmin<>oriC1 22.4 kb; GCmin<>oriC2 1.6 kb; oriC2<>oriC1 20.8 kb)

>Halomonas cupida DSM 4740 [NZ_FRCA01000005.1:323996..324495], <rpmH oriC1 dnaA>, 632 bp

GCACCATGATCAGGTAAGCGAGCACCGCCAGGGGGATCAGCAAAAGTAGTCGTTTTACGTCCATGAGGGGTCTGCCCGATGGGAAGTGAACAATCCAGTG
GCGTCCGTCTGGGCGCCGACGCCTGGGAGAAATACACCAAAGCCCGCCAGAGGCAGGCCAGGATCCGTACGCAGACATTCTGCGATGAGAGAGGTGACGT
CAATGTTATCGACGACGTTGGGCTCGATCATGCGCATATACAGTGTCATCGCATCGAAGGAGCCGGACGGCGCTATCCGCGGCGCTGGTGAGCGGCGGGG
GAGTGGGGGGCCGTCGAGGAGCGAGAGGGTCGGGTCGACTTCTCGGCCTCGCGCTCGAGGCGGCGCCACATGCCGAACAACTGACGATGCACGGTATTGT
TGTCGACCTCATTGACGCCACGTCTCGCCAGTAGAACGATATCCACACCGGGAAGTCGATACTGGGAATGGCGTACAGATTCGCGGGCCAATCGCTTGAG
ACGGTTGCGATCAACGGCCCTGCGTACGTTCTTCTTGCTGAACACCAGGCCAAGGCGCGGGTGCCCCAGATCATTGGGGGTAGCCAGCGCCATCAGGCCC
TTGCCGTGGACCTTGTATACCGCACTGTCAAAGACGTGGCGGTAATCCCCAGCAGTCAGCAAGCGGTAGTGCCGGGGAAAGGTCAGACAGGTCACGCTCG
ACCTGCGATCAGGCGCTCAGGCGCTTGCGGCCCTTGGCACGACGACGTGCCAGAACGGCACGGCCGTTCTTGGTAGCCATGCGAGCACGGAAGCCGTGCG
CGCGCTTGCGCTTGAGAACGCTCGGTTGGAATGTGCGCTTCATAACTTGTTAATCCTGAATGGTTTTTTGGATCGGATAGACAACCAAAGGAGCCCGGAA
TCGGGTCGGGCTCCTCGGGGGTATTCGGTTCAAGACCGGACATTCTAGTGATTTCAAGTCCTCTGCGCAATTGTTGTGCGCATTTTTAGCCCAGCAGTGC

TGGTTACCGCCTTGAATACCGCCTGCCACTGCCAGCTGCTCGGAAGTTCCCGGTGGCTGTCATTCGATGGCATTGAGATGGCGAGGTCGGGAGGAGCCAG
GTACCCACAGCCCGGCTTACCTTCCTTCGAATGTTCAAACCTTAATCACTATAGAACTAAGTACCTGTATATACTTGTTATTATTACTGGGGATAATGTT
**TGTGGATAA**CTGGCAAAGTTTCTTTAAAGTCAATATGCTGCGCCGCACACAAAAGTGTGGGAAAAGCGTTGAACACCGGTGGGCGAGACGTTGATGAGCT
GTGCAGAAAGGTTGGCATTATCCACAGGAGGGAATCCGTCCCTGTTCCATCCCCAGTTCCCTGGGGTAGCAAACCTCCGGTTTTCCACAGAGCTCTCTTG
ATACTTCTGGCGTGTGGATAACTCATTCCCGGAACGATACAATGCGCGGTCGTTCTTCCAATCAAGGTAGCTCGTGTGTCGCTCGCTCTTTGGCAACAAT

GTCTGGAAACCCTCCAGGACGAACTGAGTTCACAGCAGTTCAACACCTGGATTCGACCGCTTCAGGCGGAGGAGGAGGCCAATCAGCTTCGGCTGTTGGC
GCCCAACCGGTTTGTGCGTGATTGGGTCAACGACAAGTACGCCAAGCGTATCAGTGAGCTGATGCGTGAGCTTGCGCCGGCCAAACCACCCAAGGTGATG
CTGGCAGTGGGAAGCCGACGCGCTGCGGCTCAAGCTCCCCTGACTCGTGACCTGGGTGATCCGGTATCGGCCACCCCGCCACAGGCCCCTGCAGTACGTA
CCCCTCCCCGCGGAGACGTAGCAGACGAGCGCGAGATTGATCGTCTGCGCGAGACTGGTGAGGGCGGTGGCACTCGTACGCCCAGCGGCGAACGCCAGGT
GCAGGTGGAGGGTAGCCTCAAGCACTCCAGTGGCCTCAATCCCAACTTCACCTTCGAAACCTTTGTCGAGGGTAAGTCGAACCAGTTGGCCCGTGCCGCC
TCGCGCCAGGTGTCGGAGAACCCGGGGGGCGCCTACAACCCACTGTTCCTGTATGGCGGTGTGGGGCTTGGTAAAACCCACCTGATGCATGCCGTGGGCA
ATGCGTTGGCAGGGCGTCGCGAGAATGCGCGGGTGGTGTATCTGCACTCCGAGCGTTTTGTGGCCGATATGGTCAAGGCGTTGCAGCTCAACGCCATCAA
TGACTTCAAGCGCTTCTATCGCAGCGTTGATGCGCTGCTGATCGATGACATCCAGTTCTTTGCCGGCAAGGAGCGTTCCCAGGAGGAGTTCTTCCATACC
TTCAATGCGTTGCTTGAAGGTGGCCAGCAGATGATTCTGACCTCGGATCGCTATCCGAAGGAGATCAGTGGTGTCGAGGAGCGTCTCAAGTCACGTTTTG
GCTGGGGCCTGACGGTTGCCATCGAGCCGCCCGAGTTGGAGACGCGTGTCGCAATCCTGATGAAGAAGGCTGATCAGGCCAGGGTCGACCTGCCTCACGA

>Halomonas cupida DSM 4740 [NZ_FRCA01000005.1:303189..303688], <mnmG oriC2 <COG1126 (glnQ), 546 bp

CGAACGTACCACCTGCAGCTGCACGTCAAAGGGCAGCGAGGTCGAAATACCATTTGGGTATAGCTCGTGAGTGTCGAGGCCTTCGGGCTCGATGAACACC
TGATGGCTCGACTTGTCGGCAAAGCGATGCACCTTGTCCTCGATGGACGGGCAGTAACGCGGGCCGACCCCCTCGATGACACCGGAAAACATCGGTGACC
GATCGAGGTTGGCGAAGATGATCTCGTGACTACGCTCATTGGTATGCGCGATATGGCAACTCACCTGACGGGGGTGCATATCACGTGAGCCCAGGTAGGA
CATTACCGGTGTTGGTGTATCACCGGGTTGCTCCTCGAGGCGGGAAAAATCCACGCTCCGGGCATCGATACGCGGTGGCGTCCCGGTCTTCAGCCGATCC
ACGCGGAATGGCAGAGCACGCAGGCGTTCGGCAAGTGCATTGGAGGGAGGGTCTCCAGCCCGCCCGCCGCGGCTCTTTTCGAGACCAATATGAATTACCC
CGCCGAGGAAAGTACCGGTACACAACACCACGGTTTCCGCCAGAAAGCGTACTCCGGTTTCCGTCACTACGCCGCGAACCGTGTCCTGATCAACGATCAG
ATCCCCGGCAGCCTGTTGGAAAATGGTCAGGTTGGTCTGGTTTTCCAGCATCGAGCGGATTGCCGCCTTGTAGCGGACACGGTCGGCCTGAGCACGTGTG
GCACGTACCGCAGGGCCCTTGCGTGCATTGAGCACGCGAAACTGGATGCCACCCAGGTCTGTCGCCAGACCCATGGCACCGCCCAGGGCGTCGATTTCCT
TGACCAGATGACTCTTGCCGATCCCACCGATGGCGGGATTGCAGGACATCTGTCCCAGGGTCTCGATGTTGTGTGTCAGCAGCAGGGTTTGGCAACCCAT
TCGGGCAGAGGCCAGAGCAGCTTCCGTTCCGGCATGGCCACCGCCGATGACGATGACGTCAAAGCGGTCGGGATACTCCAAGGTTCACCTCGTTTTGGCG

GTCAATCGCCAGCGTGGAAAAAGTGTCGAGTGAAAACGGTCTTGAAAACAAGCCCGCCAGTATAACGCCCCTGTGGGTAAACGTCAGGGTTTTCCACATA
GGGTTCAGCAGAGGTCCCGCTTCTGGTCTTTCTTTATAAAAAGAATAAGTAAATAAAGAGAAGTTCTGTAATGGTAGTAACTACTGGTGTGGGTAATTTC
**TGTGGATAA**GTTATTTTAATCTTTTAAAATCAATTTCTTGTTGAGTTAATAAAAGTTTGGAAAGCAGGCTTGCGAGTTGCGTATGAGTTGTCTGCAGCCT
GTGGATGAAATGCCGGATTGCCCACAGGCAGAATCCTCCACTCGTTATCCACCGATCTCACGAACACTTGTCACCGCGATGGTGAACAACCCAGTAGAAT
CGCCAGGAGTCCCGTATCCTCCGACTATCGAGGCAATCATCATTTTTTTTATCCACAGGTGAAAAAAAGGCCTGTTGAGAAGCTCAACAGGCCTTGCTTG

AGAAAATCCGGGGTTGGTGAAGCGTGATCAGTGCTTGAGGACGCGTGCCAGAAAGTTCTGTGTGCGCTCGTTGCGAGGGTTATCGAACACCTCTTCGGGA
GCGCCCTGTTCGACGATCTCACCTTGATGGATGTAGATCACTCGGTCGGCCACTTCGCGGGCGAAGCCCATCTCGTGAGTGACGATGATCATGGTCATGC
CTTCCCTCGCCAACTCGCGCATGGCATCAAGGACCTCGCCGATCATCTCGGGATCCAGCGCCGAGGTGGGCTCGTCGAACAGCATCAATCGAGGTTCCAT
GGCCAGAGCGCGTGCCAGTGCCACACGCTGCTGTTGACCGCCGGAGAGCTGGGATGGGTACTTGTCGGCCTGGTCACGAATACCGACGCGATCCAGCAGG
CGCTCAGCAGTGGCGATGGCATCCGACTGCGGGACACCACGTACTTTCATTGGTGCCAGAGTAACGTTGTCGCGCACGCAGAGGTGCGGAAAGAGGTTGA
ACTGCTGGAAGACCATGCCCACCTCAGTGCGGATGGTCTGGAGGGCACGAGAGGCCTTGCCATGGGGCAGTAGGTCGTGGCTATCGACTCGCAGCTCACC
GCTCTGGAATTCCTCCAGCCCATTGATGCAGCGAATCATGGTCGATTTTCCGGAGCCGCTGGCACCGATGACCACTACAACTTCGCCCTGGGAGACCTCG
AGGTCGATATTGTTGAGCACATGCAGGCTGCCGAAGTGCTTGTTCAGGCCATGCATTCGTACGATGGGCTCGTCAGCCTGCTGTGCGCTCTTGTCGAGAT
TGATGTCGTTCATATCCATTTCCATGAAGTATCAGCACACAGCGGGTTATTCACCCACGAGGCCCTTGCGTTCGAGCATGCGCAGCGCAAGGGACAGGGA
GAGCGTGATGGCCAGGTAGAGCAGCGCCACCATCAGGTAGACCTCGAGGGCCGTGAAGGTTGTGGCGATATAGATCTGTCCCTGGCGTACCAGCTCTCCC

>Halomonas cupida DSM 4740 [NZ_FRCA01000005.1:325941..326063], dnaA> ig dnaN>, 123 bp

WebSkew GCmin pos. 84960 (GCmin<>oriC1 29.4 kb; GCmin<>oriC2 8.5 kb; oriC2<>oriC1 20.9 kb)

>Halomonas litopenaei SYSU ZJ2214 [NZ_PXNS01000004.1:c55323..55822], <rpmH oriC1 dnaA>, 601 bp

CCTGGTTCCACTGCACCATGATCAGGTAGGCGAGCACCGCCAGGGGGATCAGCAGGAGTAGTCGTTTTACGTCCATGAAGGGTCTGCCCGAAGGGAAGTG
AACATGCCGATGGCGCCACGCGAGACGCCTACATGAGATACACGAAAGCCTGCCGGAGGCAGGCCAGTATTCGTACGGAAACGTTCCGCGTCGCGAGAGG
TGACGTCAAGGGGCCGGAAGAGGGACTCGTTCATGCTCGGACGAAGTGTGTTCGGCATTGAAGAGCCTCTTGCTGCTAGTTACGGCTTTGCTTCGGCTGT
GACCGAGTGTCACCCCGGGTGTCACTCGCGTGGCGAGGTGTCGTCGATGTCCGCGAGGGACGAGTCGACTTCTCGGCCTCGCGCTCGAGACGACGCCACA
TGCCGAACAGCTGGCGCAGCACGGTGTCGTTGTCGACCTCGTTGACGCCACGCCTGGCCAGCAAGACGATATCCACGGCGGGAAGTCGATGCTGTGCATG
CCGAACGGACTCACGAGCCAGTCGTTTCAGTCGATTGCGATCGACGGCCCGGCGTACGTTCTTCTTGCTGAAGACCAGGCCCAGGCGAGGGTGTCCCAAG
TCATTGGGAATCGCCAACGCCATCAGGCCCTTGCCGTGAACCTTGTAGACCGCCCCATCGAACACCTGGCGGTAGTCCCCGGCAGTCAGCAAGCGGCATT
GCCGGGGAAAGGTCAGACAGGCCACGAAGGACCTGCGCTTAGGCGCTCAGGCGCTTGCGGCCCTTGGCACGGCGACGTGCCAGAACGGCGCGGCCGTTTT
TGGTGGCCATGCGAGCACGGAAGCCGTGCGCGCGCTTGCGCTTGAGAACGCTCGGTTGAAATGTGCGCTTCATAACTCGTTAATCCCACAGTGGTTTTTT
GGAACGGATAGACATCCAGAGGCTCCCGGAATCCGCTCGGGCCCCCGGGGGTATTCGGCTCAAGACCGGATATTTTAGTGATTTCAAGCCCTGGGTGCAA

TCTTTTCGCGGGCCAGAAAGCAGGTTTCAGGGTGTTCGCCCCTGCCAGTCCGGACCAGACGGCTGGGGATAACTCTTGCTCGAGCCGAACCGGTGGGTGA
ACAAGCCACCTGGGACCGGCCTCGCGACATGACCCACTGCCTACATAACAATAAAGAGAAGTTCTTGACTGATGTTGTTATTACTACTGGGGATAGTTTT
**TGTGGATAA**GCTTGTTCTCGTATGAAATTTCAACATGTTGCGTTGCATATAAAAGTGTGGAGAAAACGTTGAGACCGCGGGGAGGAGCGGTGTGGTTCCG
GTGGATCAATGATGAGATTGTCCACAGGGGCTAGTTCGCCAGCAAGTTGTTCCAGCACAACTCCGAATGGCGTACAAGGGTTTTCCACAGCTAGGTTTCG
GTGCCGTTCTCGTGTGGATAACTCGTGGTGACAAGGCTACAATGTGCGGTCGTTCTTCAACCAAGGTAGCTCGCGTGTCGCTCGCTCTCTGGCAACAGTG

TCTACAAACCCTTCAGGATGAACTGAGCTCGCAGCAGTTCAACACCTGGATTCGCCCTCTGCAGGCGGAAGAAGGGGAGTCCAATCAGCTCAGGCTGCTG
GCACCCAACCGCTTCGTGCGCGATTGGGTCAGTGACAAGTACGCCAAGCGTATCAGTGAACTGATGCGGGAGTTGGCGCCGGCCAAGCCGCCCAAGGTGA
TGCTGACGGTGGGCAGTCGTCGTGCTGCGCCGGCCCCACCCCAGACTCGCGAACTCGGCGATCCTGTCTCCGCCGCTCCGGCCCGCTCGGCGGCCGTGCA
CACCAGCTCCCGCGGCGATGTGGGGGATGAGCGCGAGGTCGACCGCCTGCGCGACGAAGGCGGTGAGGCGCCGGTCACGCGTCCACCCGGTGGTGAGCGT
CAGGTTCAGGTCGAGGGCAGTCTCAAGCACTCCAGTGGCCTGAATCCCAACTTCACCTTCGAGACGTTTGTCGAGGGCAAGTCCAACCAGCTTGCCAGGG
CGGCATCCCGCCAGGTGTCGGAGAACCCTGGGGGAGCCTACAACCCACTGTTTCTGTATGGTGGCGTCGGCCTCGGTAAGACCCACCTGATGCATGCGGT
GGGCAATGCCCTGGCGGGGCGTCGCGAGAACGCTCGGGTGGTGTATCTTCACTCGGAGCGCTTCGTGGCCGATATGGTCAAGGCCCTTCAGCTCAACGCC
ATCAATGACTTCAAGCGCTTCTATCGCAGCGTCGACGCGCTGTTGATCGATGACATCCAGTTCTTCGCCGGCAAGGAGCGCTCCCAGGAGGAGTTCTTCC
ACACCTTCAATGCCTTGCTTGAGGGTGGCCAGCAGATGATCCTGACCTCGGATCGCTATCCGAAGGAAATCAGCGGCGTCGAGGAGCGCCTCAAGTCCCG
TTTTGGCTGGGGGCTGACGGTGGCGATAGAGCCGCCTGAGCTCGAGACTCGGGTGGCGATCCTGATGAAGAAGGCCGACCAGGCCAAGGTGGATCTACCC

>Halomonas litopenaei SYSU ZJ2214 [NZ_PXNS01000004.1:c76249..76748], <mnmG oriC2 <COG1126 (glnQ), 540 bp

ACCTTCGATGGAGCGCACCACCTTGAGCTGCACGTCGAAGGGCAAGGAGGTCGAAATACCATTTGGGTATAGCTCATGGGTATCCAGACCCTCCGGCTCG
ACGAACACCTGGTGGCTGGTCTTGTCGGCGAAGCGGTGGACCTTGTCTTCGATGGACGGGCAGTAGCGCGGCCCGACCCCTTCGATCACCCCGGAGAACA
TCGGCGAGCGATCGAGATTGTCGAAGATGATCTCGTGGGTGCGCTCGTTGGTATGGGCGATATGGCAGCTGACCTGACGCGGATGCATGTCCCGTGATCC
CAGGTAGGACATCACCGGAGTCGGACTGTCGCCCGGCTGTTCCTCGAGGCGCGAGAAGTCGACGCTCTTGGCATCGATGCGCGGCGGGGTGCCGGTCTTC
AGGCGATCGACCCGAAACGGCAAGGCGCGCAGACGCTCGGCCAGGGCATTGGAGGGCGGATCCCCGGCACGACCACCACGGCTGTTGTCCAGCCCGATGT
GGATCACCCCGCCGAGGAAGGTACCGGTGCACAGCACCACGGTTTCGGCCAGAAATCGCAGTCCAGTCTCGGTGGCCACGCCGCGCACGGTGTCCTGGTC
GACGATCAGGTCGCCGGCGGCCTGCTGGAACAGCGTCAGGTTCGGTTGGTTTTCCAGCATGCCACGAATCGCGGCCTTGTAGCGCACACGATCGGCCTGG
GCACGGGTCGCCCGTACCGCGGGGCCCTTGCGGGCGTTGAGCACGCGAAACTGGATCCCGCCCAGGTCCGTCGCCAATCCCATGGCGCCGCCGAGGGCGT
CGATTTCCTTGACCAGATGACTCTTGCCGATCCCGCCGATGGCGGGATTGCAGGACATCTGGCCCAGGGTCTCGATGTTGTGGGTCAACAGCAGGGTCTG
GCAACCCATGCGGGCAGAGGCCAGGGCAGCTTCCGTTCCAGCATGGCCACCGCCGATGACGATGACGTCAAAGCGGTCGGGATACTCCAAGGTTCACCTC

GTTCGGCGCAGCGCGCCAGCGGTCTGGATATTGGGTTCGTACGAAAAATAGCCGGCCAGTATAAACCCTCCTGTGGGTAAACGTCAGGGTTTTCCACACC
CCGTCTGTCAGGCAGGAGCTTCACTTACCTACCTATAAAAAAATAAAAGAATAAGAAAGGAAGTTCTGTAGTGGTTGTAACTAACTGGTGTGGACACTTTC
**TGTGGAAAA**ACCATTTAAATCTACAAAAAACAAATAGGTACAATGTTGATCCTTTGTTGGACAAGGGACGTACCAGCGTTTGATGGCGTGTTATCTGCCT
GTGGACACAATCACGACTTGCCCACAGGGCTCGGCTTCCACCAGTTATCCACCGAGCTCGGGGGCGGTTGTCACCGCGACGGTGAACAACTCGCCGACTT
GCCCACAGCCCCCGTGGCAGTGGGCTTCGCAGCCAAATGTGAAAAATCTTGTCCACAGCCCCAAAAAGCCAAAAAAACGCCCTGTCGATCACTCGACAGG

GCGTTTGAGAGGCTCGAACGGGCGGTTTTTTCAGTGCTTGAGCACTCGCGCCAGGAAGCTCTGGGTACGTTCATTCTGCGGATTGTCGAAGACTTCTTCG
GGCTTGCCCTGCTCGACGATCTCACCCTTGTGGATGTAGATCACGCGGTCGGCGACCTCCCGGGCGAAGCCCATCTCGTGGGTCACGATCATCATCGTCA
TGCCCTCTCTGGCGAGCTCACGCATGGCGTCGAGCACCTCACCGATCATCTCCGGGTCCAGCGCCGAGGTGGGCTCGTCGAACAGCATCAGACGGGGCTC
CATGGCCAGTGCGCGGGCCAGCGCCACGCGCTGCTGCTGACCGCCGGAGAGCTGGGTCGGGTACTTGTCGGCCTGGTCCTTGATGCCGACCCGATCGAGC
AGCCGTTCCGCGGTGGCCTCGGCCTTGTCACGGCTTTCGCCGCGCACCTTCATCGGGGCCAGGGTGACGTTGTCACGCACGCTGAGGTGAGGGAACAGGT
TGAACTGCTGGAACACCATGCCGACCTCGGTGCGGATGGTCTGCAGCGCCTGGCTGGCCTTGCCGTCGGGCTTGAGGTCGTTGCCGTCGACCCTGAGGGC
GCCCTGCTGGAATTCTTCCAGGCCATTGACGCAGCGGATGAGGGTCGACTTGCCGGAGCCACTGGCGCCGATGACGACCACCACTTCGCCCTGGGCCACC
TCAAGGTCGATGTTCTTGAGCACGTGCAGCGAGCCGAAGTGCTTGTTGAGTTTGTCGATACGGACGATGGGCTCGCCCTGGGCGTGAGAGGCTGGGTTCA
TGTCGGAATTTTCCATGGGATGTTGCACCTCACTGGCCCACCAGGCCCTTGCGCTCGAGCATGCGCAGGGCGAAGGACAAGGAAAGGGTGATGGCAAGAT
AGAGCAGCGCCACCATCAGATAGACCTCCAGGGCGGTGAAGGTGGTGGCGATGTAGATCTGGCCCTGGCGCACCAGCTCGCCAACGCCGATGACGGCGAA

>Halomonas litopenaei SYSU ZJ2214 [NZ_PXNS01000004.1:c53813..53872], dnaA> ig dnaN>, 60 bp

WebSkew GCmin pos. 229755 (GCmin<>oriC1 45.3 kb; GCmin<>oriC2 25.9 kb; oriC2<>oriC1 19.4 kb)

>Halomonas urmiana TBZ3 [NZ_VBUI01000001.1:184233..184732], <rpmH ig dnaA>, 563 bp

AACGAGCGCCGTCCGACGCGCTGCTGGTCATGGCGGGGGCTTGCGGCTCAGGGGAAACCTGGCCGTAGTCCTGATTCCACTGGACCATCAACAGGTAGGC
GAGTACCGCCAGGGGAATCAGCAGGAGTAGTCGTTTTACGTCCATGAGGGGTCTGCCCGATGGGTAGTGAACATTCCGATGGCGCAACGCAGGTCGCACA
CCCGAGATACACGAAAGCCTGCCGGAGGCAGGCTAGGAAACGCACGCTGGTCGGCGGCATCCGACGCGTCGGCGCGGAGAGGCGCGGCAAAGAGCGAGGT
GATCATGCTCATGCAGGCTCCGGGATGCGCTGGCGCTTGGGGGGTCGGTCATCATATGCGGTGCGTGTCTTCGCTCCGCGTCCCGTGTCGGTCCGGCTCG
CGGGCGGACTTGCTCGCCTCGCGCTCGAGGCGGCGCCACATACCCTGGAGCTGGCGATGTACGGTGGCGTTGTCCAGCTCGTGGACTCCCCGTCTGGCCA
GTACCACGATATCCACGGCGGGAAGGCGCTGCTGCTGGAGCCGGATGGAGTCACGCACGATGCGCTTGAGGCGGTTGCGATCGACGGCCCGGCGCACGTT
CTTCTTGCTGAAGATCAGGCCGAGTCGCGGGTGGCCGAGTGCGTTGGGCGAGGCCAGAGCCATCAGCCCCCTGCCATGGACCTTGAAAGCCGCCGTTTCG
AAGACACGACGATAGTCCCCAGCAGTCAGCAGTCTCAGCTGCCGGGGATAGCCCTGACGTGACACGCGAGGGCCGCGATCAGGCGCTCAGGCGCTTGCGG
CCCTTGGTGCGGCGGCGAGCCAGCACGGCACGACCGTTCTTGGTGGCCATGCGAGCACGGAAGCCATGTGCGCGCTTGCGCTTGAGGACGCTGGGTTGAA
AAGTACGTTTCATAACTCGTTATTCCCACGTGGTTTTCTGAATGACTGGATCACTCGTAGAGATCCGGATGCCGCAGGCGGATCCGGGGGTATTCGACTC

AAGACCGGAAATTGTAGTGATTTCAGCCCTGCGGTGCAATTCTTACCTGCGCTTTGCGCCCCCTGCTGACGCGGCTTTCCGCGTGGTCACTCGGCCGCCA
TCCACCACCCGTTCGCCCGCTCGGCATACCGTAACGCTCCAGGCCCTGTGGTCACTCCTGGGGATGGGTATTGTGAATAAGTCGTTTTATTCCATGTTTA
TCATCGTATTGAGGCGTTATTCCTGCTGTGGAAAAAGTGGGGATTAGGTGGGGTGAGGTGGTGAGGCCCACCGCGATGCCGCCCTGTCGGGCGTGGCCCG
AAGCTCGGCGCTGTCTGAGAGGGTTGTCGCCGTTCCCTGCTGCCAGCGTCCGGGCGGAAGTCAGCGGACATGGCGGCCGGGTTTTCCACAGGACGAAGCG
ATAGGCCGGTCATGTGGATAACCCGTGGCCCGAACGATACAATAGGCGGTCGTTTTCTACGCAAAGGTGAGGTCGCGTGTCACTGGCTCTCTGGCAACAA

TGTCTGGATACCCTGCAGGATGAACTGAATTCGCAGCAGTTCAATACCTGGATCCGTCCGCTGCAGGCGGAGGAAGGGGAGTCCAACCAGTTGCGCCTGT
TGGCGCCGAATCGCTTCGTGCGCGACTGGGTCAGCGACAAGTATGCCAAGCGGATCAGTGAGCTGATGCGCGAACTCTCGCCGGCCAAGCCGCCCAAGGT
GGTCTTGACCGTGGGCAGCCGGCGCAGCGCGGCGCCGAACCCGCAGCCCCGAGAACTGGGCAACCCGGTCTCGGCCGCCCGCCGGGCGCCGGCGGCGCCC
GCCGTGCAGGTGCCCCCCCGGGGAGACGTGGGCGATGAGCGTGAGATCGATCAGCTGCGCGAGGAGGGGGCCGGCCAGCGCCGTGGCGGTGAGCGCCAGG
TGCAGGTAGAGGGCAGTCTCAAGCACAATAGCGGCCTCAATCCCAACTTCACCTTCGAGACCTTCGTCGAGGGCAAGTCCAACCAGCTGGCGCGCGCCGC
CTCCCGTCAGGTCTCCGAGAATCCCGGCGGGGCGTACAATCCGCTGTTCCTCTATGGCGGCGTCGGCCTGGGTAAGACTCACTTGATGCATGCCGTCGGC
AATGCCCTGGCGACCCGGCGCGAGAACGCCCGTGTCGTCTACCTGCATTCCGAACGCTTCGTGGCCGATATGGTGAAGGCGCTGCAGCTCAACGCGATCA
ACGACTTCAAGCGTTTCTACCGCAGCGTCGATGCCCTGTTGATCGATGATATCCAGTTCTTCGCCGGCAAGGAGCGCTCCCAGGAGGAGTTCTTCCACAC
CTTCAATGCGCTGCTCGAGGGCGGCCAGCAGATGATCCTGACATCCGACCGATATCCCAAGGAAATCAGTGGGGTAGAGGAGCGATTGAAGTCTCGTTTC
GGCTGGGGGCTGACGGTGGCCATCGAGCCGCCGGAGCTGGAAACGCGCGTGGCCATCCTGATGAAGAAGGCCGATCAGGCCAAGGTCGACCTGCCTCATG

>Halomonas urmiana TBZ3 [NZ_VBUI01000001.1:203587..204086], <mnmG oriC2 <COG1126 (glnQ), 565 bp

GTTCTCGAAGCCGTTGATCGAGCGCACCACCTGCAGCTGCACGTCGAAGGGCAGCGAGGTGGAGATGCCGTTGGGATAGAGCTCATGGGTATCGAGGCCC
TCGGGCTCGACGAACACCTGGTGGCTGGCCTTGTCGGCGAAGCGGTGCACCTTGTCCTCGATGGACGGGCAGTAGCGCGGCCCGACGCCCTCGATCACCC
CCGAGTACATCGGCGAGCGATCGAGATTGGCGTGGATGATCTCGTGGGTGCGCTCGTTGGTGTGCGCGATGTGGCAGCTCACCTGGCGGGGATGCATCTC
GCGGCTGCCCTGGTAGGACATCACCGGGGTCGGCGTGTCGCCGGGCTGCTCCTCGAGCGAGGCCAGATCGATGCTCTTGGCATCGAGCCGCGGCGGTGTG
CCGGTCTTGAGGCGATCGACCCGGAACGGCAGTGCCCGCAGCCGCTCGGCCAGGCCGTTGGAGGCGGGATCCCCGGCGCGACCGCCGCGGCTGGTGTCGA
GCCCGATGTGGATCACGCCGCCGAGGAAGGTGCCCGTGCACAGCACCACCGTCTCGGCGAGGAAGCGGATGCCGGTCTCGGTGACCACGCCCCGCACGGT
GTCGCCGTCGACGATCAGGTCGCCCGCCGCCTGCTGGAAGAGGGTCAGGTTGGCCTGGTTCTCCAGCATGGTGCGGATCGCGGCCTTGTAGCGGACCCGG
TCGGCCTGGGCGCGGGTGGCCCGGACCGCCGGCCCCTTGCGGGCATTGAGCACGCGAAACTGGATGCCGCCGTGGTCGGTTGCCAGGCCCATGGCACCGC
CCAGGGCATCGATCTCCTTGACCAGATGACTCTTGCCGATCCCGCCGATGGCGGGATTGCAGGACATCTGGCCGAGCGTCTCGATGTTATGGGTCAGCAG
CAAGGTCCGGCAGCCCATGCGGGCCGAGGCCAGGGCGGCTTCGGTTCCGGCATGGCCGCCGCCGATGACGATGACGTCAAAGCGGTCGGGGTACTCCAAG

GGTCACCTCTTGGCGCGGTGCGCCGATGTGGCTGTCAGGGGTGGAAATCAGCGGGCCAGTATAAACCCCCTGTGGGTAATCGTCAGGGTTTTCCACACCC
GAATTCCTTCCCGGTGGCCGTCATCTCTCTTCATATACAATAAAACAGAATATAAAAGAGAAGTTCTGTTGTAGTTGTAACTACTGGTGTGGACATTTTT
**AGTGGATAA**GCTAATAAATCTTACTTTTATCAACAGCTTGCATTGTTGATCCCTCGTCGGAGAAAGGCGTGACAGCCTGGCGACGAGGTGTCCGGAGGCT
GTGGACGGAATGGCGACTTGTCCACACGGGAAAAAGATCACCGCTTGTCCACCGTTCTCTACCGGGCTTGTTACCGTGGCTGTGAACACTCGAGAGCCAG
GCGTCCATCTCCGCGTATGACGGGGCCTGCAGGCGGATCTAGGCAGGTTCCAGAAAACTTCTGTCCACAGGCAAAAAAAGGCCCTGTCGATAACGACAGG

GCCTTTTTTGGAGGGATGCTGAAGGGCTGGTTAGAGGCTCCCGAGGGGATCGGAGGAGGCAAGATCAGTGCTTGAGGACGCGCGAGAGGAAGCTCTGGGT
GCGTTCGTTCCGAGGTGAATCGAAGACCTCCTCGGGCGGCCCCTCCTCGGTGATCTCACCCTTGTGAATGAAGATCACTCGGTCGGCGACCTCACGGGCG
AAGCCCATCTCGTGGGTCACGATCAGCATGGTCATGCCTTCCTTTGCCAGTTCCCGCATGGCGTCCAGGACCTCGCCGATCATCTCCGGATCGAGTGCCG
AGGTGGGCTCGTCGAACAGCATCAGGCGCGGCTCCATGGCCAGCGCACGGGCCAGGGCTACGCGCTGCTGCTGGCCACCGGAGAGCTGACCAGGATGCTT
GGCGGCCTGGTCGCTGATGCCGACCCGCTCGAGCAGGCGCTCGGCGGTCTCGATAGCGTCCTCCCGGCTCCAGCCGCGCACCTTCATCGGCGCCAGGGTC
ACGTTGTCCAGCACGCTCAGATGGGGGAAGAGGTTGAACTGCTGGAAGACCATGCCGACCTCGGTACGGATCTTCTGCAGCGCCGCCGAGGCCTTGCCGT
GGGGCACCAGCTGGCTGCCATCGACCTCCAGGTAGCCCTGCTGGAACTCCTCGAGGCCGTTGATGCAGCGGATCAGCGTCGACTTGCCCGAGCCGCTGGC
ACCGATGATCACCACGACCTCGCCGGGGGAGACCGACAGGTCGATGTCCTTCAGCACATGCAGGCTGCCGAAATGCTTGTTGACCTTGTCCATGCGCACG
ATGGGATCGCTGCCGGACGTGGACTTCTCGTGTGTCATGTCGCGCGTTCCTTATTGTCCGACCAGGCCTCTGCGCTCGAGCTGGCGCAGCACGAACGAGA
GGGTCAGGGTGATCGCCAGATAGAGCAGGGCGACCATCAGGTAGACCTCGAGGGCATTGAAGGTGGTCGCGATGTAGACCTGGCCCTGGCGGACCAGTTC

>Halomonas urmiana TBZ3 [NZ_VBUI01000001.1:182726..182786], dnaA> ig dnaN>, 61 bp

WebSkew GCmin pos. 16300 (GCmin<>oriC1 28.2 kb; GCmin<>oriC2 8.8 kb; oriC2<>oriC1 19.4 kb)

>Halomonas saccharevitans CGMCC 1.6493 [NZ_FPAQ01000016.1:44290..44789], <rpmH ig dnaA>, 573 bp

TTGCTGTCCGCTGCGTTGCTGGTCATGGCGGGGGCTTGCGGCTCAGGGGGAACCTGGCCGTAGTCCTGATTCCACTGGACCATCAACAGGTAGGCGAGTA
CCGCCAGGGGAATCAGCAGGAGTAGTCGTTTTACGTCCATGAGGGGTCTGCCCGATGGGTAGTGAACATTCCGATGGCGCAACGCAGGTCGCACACCCGA
GATACACGAAAGCCTGCCGGAGGCAGGCTAGGAAACGCACGCTGTCCGGCGGCATCTGACGCGTCGGCGTGGAGGGGCGCGGCAAAGAGCGAGGTGGTCA
TGCTCATGCAGGCTCCGGGATGCGCTGGCGCTCAGGGGGGTCGGTCATCATGTGCGCTGCGTGTCTTCGCCCCGTTCTAGCTCTGGCTCGCGGGCGGACT
TGTTCGCCTCACGCTCCAGGCGCCGCCACATGCCGTGGAGCTGGCGATGCACGGTGGCGTTGTCCAGCTCGTGGACGCCTCGCCTGGCCAGTACCACGAT
ATCCACGGCGGGAAGGCGCTGTTGCCGGAGCCGGATGGAGTCGCGCACGATGCGCTTGAGCCGGTTGCGATCGACGGCCCGGCGCACGTTCTTCTTGCTG
AAGATCAGGCCGAGTCGCGGGTGGCCGAGCGCGTTGGGCGAGGCCAGAGCCATCAGCCCTTTGCCATGGACCTTGAAGGCTGCCGTTTCGAAGACACGAC
GATAGTCCCCGGCAGTCAGCAGTCTCAGCTGCCGGGGATAGCCCTGATCTGACACGCGAGGGCCGCGATCAGGCGCTCAGGCGCTTGCGGCCCTTGGCGC
GACGGCGAGCCAGCACGGCACGACCGTTCTTGGTGGCCATGCGAGCACGGAAGCCATGTGCGCGCTTGCGCTTGAGGACGCTGGGTTGAAAGTACGTTT
CATAACTCGTTATTCCCACGTGGTTTTCTGAATGATTTGATCACTCGTAGAGATCCGGAAGCCGCAGGCGGATCCGGGGGTATTCGACTCAAGACCGGAA

ATTGTAGTGATTCCAGCCCTGCGGTGCAATTCTTACCTGCGTTTCGCGGTCGCTCGAGCGCCCTGCTGGCGCGGCCTCCGGCGTGGTCGTTCGACCGCTA
TCCACCATTCGGTTGCCCTCGCGGCATGTCGTAACGTGCCGGGCCCTGTGGTCACTCCTGGGGACGGGTAGTGTGAATAAGATGATTTAATTTATATATA
TCAGCGTATTGAGAGGGTGTCCTTGCTGTGGAAAAAGTGGGGATGGGGTGGGGAGCGAGGTAACGGTGCCTGCGTGGTGTCGTCGTATCGGGTCCGGCTC
GAAGGCTTGCGGCGTCTGAGGTGTCGTCGCGTTTGCCTGCTTGTCGTGTCTTGTCGAAAAACAGCGGACACGGCGGCCAGGTTTTCCACAGCACGAGGCG
TTAGGCCGGTCATGTGGATAACCCAGGGCCTGAACGATACAATAGGCGGTCGTTTTCTACGCAAAGGTGAGTTCGCGTGTCACTGGCTCTCTGGCAACAA

TGTCTGGATACCCTGCAGGATGAACTGAATTCGCAGCAGTTCAATACCTGGATCCGCCCGCTGCAGGCGGAGGAAGGGGAGTCCAATCAGTTGCGTCTGC
TGGCGCCGAATCGCTTCGTGCGCGACTGGGTCAGCGACAAGTATGCCAAGCGGATCAGCGAGCTGATGCGCGAGCTCTCGCCGGCCAAGCCGCCCAAGGT
GGTGCTGACCGTGGGCAGCCGGCGCAGCGCGGCGCCGAACCCGCAGCCCCGTGAGCTGGGCAACCCGGTCTCGGCCGCGCCCCGAGCACCGGCGGCGCCC
GCCGTGCAGGTGCTCCCTCGGGGAGATGTGGGTGACGAGCGCGAGATCGATCAGATGCGCGAGGAGGGGGCCGGCCAGCGGCGAGGCGGTGAGCGACAGG
TGCAGGTCGAGGGCAGCCTCAAGCACAACAGCGGCCTCAATCCCAACTTCACCTTCGAAACCTTCGTCGAGGGCAAGTCCAACCAGCTGGCTCGTGCCGC
CTCCCGCCAGGTCTCCGAGAACCCCGGCGGGGCGTACAACCCGCTGTTCCTCTACGGCGGCGTCGGCCTGGGTAAGACGCACCTGATGCATGCCGTCGGC
AATGCCCTGGCGACACGGCGCGAGAATGCCCGCGTCGTCTATCTCCACTCGGAACGCTTCGTGGCCGACATGGTGAAGGCGCTGCAGCTCAATGCGATCA
ATGACTTCAAGCGTTTCTACCGGAGCGTCGATGCCCTGCTGATCGATGATATCCAGTTCTTCGCCGGCAAGGAGCGCTCCCAGGAGGAGTTCTTCCACAC
CTTCAATGCGCTGCTCGAGGGCGGCCAGCAGATGATCCTGACGTCCGACCGATATCCCAAGGAAATCAGTGGGGTAGAGGAGCGTTTGAAGTCTCGCTTC
GGCTGGGGGCTGACGGTGGCCATCGAACCGCCGGAGCTGGAGACACGCGTGGCCATCCTGATGAAGAAGGCCGATCAGGCCAAGGTCGACCTGCCCCACG

>Halomonas saccharevitans CGMCC 1.6493 [NZ_FPAQ01000016.1:24836..25335], <mnmG oriC2 <COG1126 (glnQ), 576 bp

GTTCTCGAAGCCCTTGATCGAGCGCACGACCTCAAGCTGCACGTCGAAGGGCAGCGAGGTGGAGATTCCGTTGGGATAGAGCTCGTGGGTATCGAGGCCT
TCGGGCTCGACGAACACCTGGTGGCTGGCCTTGTCCGCGAAGCGGTGGACCTTGTCTTCGATCGACGGGCAGTAACGCGGCCCGACGCCCTCGATCACCC
CCGAGTACATCGGCGAGCGATCAAGATTGGCATGGATGATCTCGTGGGTGCGCTCGTTGGTGTGGGCGATGTGGCAGCTCACCTGACGGGGGTGCATCTC
GCGGTTGCCCAGGTAGGACATCACCGGCGTCGGCGTGTCGCCGGGCTGTTCCTCGAGCGAGGCGAGATCGACGCTCTTGGCATCGAGACGCGGCGGCGTG
CCAGTCTTCAGGCGATCGACCCGGAACGGCAACGCCCGCAGCCGGTCGGCCAGGCCGTTGGAGGCGGGATCCCCGGCGCGTCCGCCGCGGCTGGTATCGA
GCCCGATGTGGATCACCCCGCCGAGGAAGGTCCCCGTGCACAGCACCACCGTCTCGGCGAGGAAACGAATGCCGGTCTCGGTGACCACGCCGCGCACCGT
ATCGCCGTCGACGATCAGGTCGCCCGCGGCCTGCTGGAAGAGGGTCAGGTTGGCCTGGTTCTCCAGCATGGTGCGGATCGCAGCCTTGTAGCGGACCCGA
TCGGCCTGGGCCCGTGTGGCTCGCACGGCTGGCCCCTTGCGGGCGTTGAGCACCCGGAACTGGATGCCCCCGAGGTCGGTTGCCAGGCCCATGGCACCGC
CCAGGGCATCGATCTCCTTGACCAGATGGCTCTTGCCGATCCCGCCGATGGCGGGATTGCAGGACATCTGGCCGAGGGTCTCGATGTTATGGGTCAGCAG
CAAGGTCCGGCAGCCCATGCGGGCCGAGGCCAGGGCGGCTTCGGTTCCGGCATGGCCGCCGCCGATGACGATGACGTCAAAGCGGTCGGGATACTCCAAG

GGTCACCTCTTGGCGCGGTGCGCCGATGTGGCAGTCAGGGGTGGAAATCAGCGGGGCAGTATAAACCCCCTGTGGGTAATCGTCAGGGTTTTCCACACCC
GAATTTCCTCACCGTGGCCGTCGTCACTCTCCATATACAATAAAACAGAATATAAAAGAGAAGTTCTGTTGTTGTTGTAACTACTGGTGTGGACATTTTT
**GGTGGATAA**GATAATAAAGCCTTTTTTTATCAACAGTTTGCATTGTTGACCCCTCGCGGGAGAAAGCCGTGACTGCCTGCGGACGAGGTGTCCGGAGACT
GTGGATGAAATGGCGCCTTGTCCACACGCGAAAACGATCACGCCTTGTCCACCGCTTTCTACCGGGCTTGTTACCGCGACTGTGAACAGTTGCCTGTGAG
GTCGCCATACCCGCGTATGACGGGGCCTGCAGACGGGCAGAGGCGGGTATCAAATTTTTTCTATCCACAGGCAAAAAAAGGCCCTGTCGATAACGACAGG

GCCTTGTCTGAAGGGCTTCAGAAGAGCTTTCAAAGTGCCTAAGAGTGGCTCACGAGGTGAGCGGAGTAGGCGAGATCAGTGCTTGAGGACGCGCGAGAGG
AAGCTCTGAGTGCGCTCGTTCTGCGGCGAATCGAAGACCTCTTCGGGTGGCCCCTCCTCGGTAATCTCGCCCTTGTGAATGAAAATCACCCGGTCGGCGA
CCTCGCGGGCGAAGCCCATCTCGTGGGTCACGATCAGCATGGTCATGCCTTCCCGAGCGAGCTCGCGCATGGCGTCCAGGACCTCGCCGATCATCTCTGG
ATCGAGCGCCGAGGTGGGCTCATCGAACAGCATCAGGCGCGGCTCCATGGCCAGCGCACGGGCCAGGGCCACGCGCTGCTGCTGGCCGCCGGAGAGCTGG
CCGGGATGCTTGGCAGCCTGGTCGCTGATGCCTACCCGCTCGAGCAGGCGCTCGGCGGTCTCGGTGGCATCCTCGCGGCTCCAGCCGCGCACCTTCATCG
GCGCCAGGGTCACGTTGTCCATCACGCTCAGATGGGGGAAGAGGTTGAACTGCTGAAAGACCATGCCGACTTCGGTGCGGATCTTCTGCAGCGCCTGGGT
GGCCTTGCCATGGGGCACCAGCGGGCTGCCATCGACCTCCAGATAGCCCTGCTGGAACTCCTCGAGGCCGTTGATGCAACGGATCAGCGTCGACTTGCCC
GAGCCGCTGGCGCCGATGATCACCACCACCTCGCCGGGGGAGACCGACAGGTCGATATCCTTCAGCACGTGAAGGCTGCCGAAATGCTTGTTGACCTTGT
CCATGCGCACGATGGGATCGCTGCCGGACGTGGACTTCTGGTGTGTCATGTCGCGCGTTCCTTATTGTCCGACCAGGCCTCTGCGCTCGAGCTGGCGGAG
CACGAGCGAGAGGGTCAGGGTGATCGCCAGGTAGAGCAGGGCGACCATCAGGTAGACCTCGAGGGCATTGAAGGTGGTGGCGATGTAGACCTGGCCCTGG

>Halomonas saccharevitans CGMCC 1.6493 [NZ_FPAQ01000016.1:46236..46296], dnaA> ig dnaN>, 61 bp

WebSkew GCmin pos. 242526 (GCmin<>oriC1 47.2 kb; GCmin<>oriC2 27.7 kb; oriC2<>oriC1 19.6 kb)

>Halomonas denitrificans DSM 18045 [NZ_PYVX01000003.1:c195069..195569], <rpmH ig dnaA>, 553 bp

CGGTCGGCGCGGAAGCACCGTTCGACGCGCTGCTCGTCATGGCGGGGGCTTGCGGCTCAGGAGAAACCTGGCCGTAGTCCTGATTCCACTGGACCATCAA
CAGGTAGGCGAGTACCGCCAGGGGAATCAGCAGGAGTAGTCGTTTTACGTCCATGAGGGGTCTGCCCGATGGGTAGTGAACATTCCGATGGCGCAACGCA
GGTCGCACACCCGAGATACACGAAAGCCTGCCGGAGGCAGGCTGGGAAACGCACGCTGTCCGGCGGCATCCGAGGCGTCGACGCGATGGGGCGCGGCAAA
GAGCGGGGTGATGAGGCTCATGCAGGCTCCGGGATGCGCTGGGACCGAAGGGGGCCGGTCATCATTTGCGGTGCGTGTCTTCGCTCCGCGTGGTGCCTCG
CTCCGGCTCGCGGGCGGACTTGCTCGCCTCGCGCTCGAGGCGGCGCCACATCCCGTGGAGTTGGCGATGCACGGTAGCGTTGTCCAGCTCGTGGACTCCT
CGCCGGGCCAGCACCACGATATCGACCGCGGGAAGGCGCTGCTGCCGGAGCCGGATGGAGTCACGCACGATGCGCTTGAGGCGGTTGCGATCGACGGCCC
GGCGTACGTTCTTCTTGCTGAAGATCAGGCCGAGTCGCGGGTGGCCGAGCGCGTTGGGTGAGGCCAGAGCCATCAGCCCCTTGCCATGGACCTTGAAAGC
CGCCGTTTCGAAGACACGACGATAGTCCCCGGCAGTCAGCAGTCTCAGCTGCCGGGGATAGCCCTGACGAGACACGCGAGGGCCGCGATCAGGCGCTCAG
GCGCTTGCGGCCCTTGGCGCGACGACGAGCCAGCACGGCACGGCCGTTCTTGGTGGCCATGCGAGCACGGAAGCCATGTGCGCGCTTGCGCTTGAGGACG
CTGGGTTGAAAAGTACGTTTCATAACTCGTTATTCCCACGTGGTTTTCTGAATGATTGGATCACTCGTAGAGATCCGGATGCCGCTGGCGGATCCGGGGG

TATTCGACTCAAGACCGGAAATTGTAGTGATTCCAGCCCTGCGGTGCAATTCTTACCTGTGTTTTGCGGCGCCGGGAACCCCTGACGGCGCGGCTTTCGG
CGCGATCGTGGACCGCGTGCCCCTGCCCGTTCGCCCCCATGACAGGCCGGGGTGTCTCGTACCCTGTGGTCACTCCTGGGGACGGGCTGTGTGAACAAG
TCGTTTTATTTTATATATATCAATGCATTGTTGATGTGTATCTGCTGTGGACAAAGTGGGGATGGGGTGAAGGGGGAGTGCACTCACCGCGATGCGCCCG
TGTCGGGCAAGGCTCGAGGGGCCGGCGGCGTCCGAGGGTCGCTGACCGCTGCGTCCCGCAAGCCGCCGAGGGGTGGCGGGGTTTTCCACAGGACCCGGCG
TTGGCTCGGTCATGTGGATAACCCGAAGCCCGAGCGATACAATGGGCGGTCGTTTTCTACGCAAAGGTGAGGTCGCGTGTCACTGGCTCTCTGGCAACAA

TGTCTGGATACCCTGCAGGATGAACTGAATTCGCAGCAGTTCAATACCTGGATCCGTCCGCTGCAGGCGGAGGAAGGGGAGTCCAACCAGTTGCGCCTGC
TGGCGCCGAATCGCTTCGTGCGCGACTGGGTCAGCGACAAGTATGCCAAGCGGATCAGCGAGCTGATGCGTGAGCTCTCGCCGGCCAAGCCGCCCAAGGT
GGTGCTGACCGTGGGCAGCCGCCGCAGCGCGGCGCCGAACCCGCAGCCCCGTGAGCTGGGCAACCCGGTCTCCGCCGCGCCCCGGGCACCGGCGGCGCCC
GCCGTGCAGGTCCCGCCCCGGGGCGACGTGGGCGACGAGCGCGAGATCGACCAGCTGCGCGAGGAGGGGGCCGGCCAGCGGCGGGGCGGTGAGCGCGAGG
TGCAGGTGGAGGGCAGCCTCAAGCACAATAGCGGCCTCAATCCCAACTTCACCTTCGAGACCTTCGTCGAGGGCAAGTCCAACCAGCTGGCGCGCGCCGC
CTCCCGCCAGGTCTCCGAGAACCCCGGCGGGGCCTACAACCCGCTATTCCTCTACGGCGGCGTCGGCCTGGGTAAGACCCACCTGATGCATGCGGTCGGC
AATGCCCTGGCGACCCGGCGCGAGAACGCCCGCGTGGTCTACCTCCATTCCGAGCGCTTCGTCGCCGACATGGTGAAGGCGCTGCAGCTCAATGCGATCA
ACGACTTCAAGCGTTTCTACCGCAGCGTCGACGCCCTGCTGATCGACGATATCCAGTTCTTCGCCGGCAAGGAGCGCTCCCAGGAGGAGTTCTTCCACAC
CTTCAATGCGCTGCTCGAGGGCGGCCAGCAGATGATCCTCACGTCCGACCGTTATCCCAAGGAAATCAGTGGGGTAGAGGAGCGCTTGAAGTCGCGCTTC
GGCTGGGGGCTGACGGTGGCCATCGAGCCGCCGGAGCTGGAGACGCGCGTGGCCATCCTGATGAAGAAGGCCGATCAGGCCAAGGTCGACCTGCCCCACG

>Halomonas denitrificans DSM 18045 NZ_PYVX01000003.1:c214606..215105], <mnmG oriC2 <COG1126 [(glnQ), 553 bp

GTTCTCGAAGCCCTTGATCGAGCGCACCACCTGCAGCTGCACGTCGAAGGGCAGCGAGGTGGAGATCCCGTTCGGGTAGAGCTCATGGGTATCGAGGCCC
TCGGGCTCGACGAACACCTGGTGGCTGGCCTTGTCGGCGAAGCGGTGCACCTTGTCCTCGATCGAGGGGCAGTAGCGCGGCCCGACCCCCTCGATTACCC
CCGAGTACATCGGCGAGCGATCGAGATTGGCATGGATGATCTCGTGGGTGCGCTCGTTGGTGTGCGCGATGTGGCAGCTCACCTGGCGCGGATGCATCTC
GCGGCTGCCCAGGTAGGACATTACCGGGGTCGGGGTGTCGCCGGGCTGCTCCTCGAGGGAGGCGAGATCGATGCTCTTGGCATCGAGGCGCGGCGGCGTG
CCGGTCTTCAGGCGATCGACCCGGAACGGCAGCGAGCGCAGGCGCCCGGCCAGGCCGTTGGAGGCGGGATCCCCGGCGCGACCGCCGCGGCTGGTGTCCA
GGCCGATGTGGATCACCCCGCCGAGGAAGGTTCCGGTGCACAGCACCACCGTCTCGGCGAGGAAGCGGATGCCGGTCTCGGTGACCACGCCGCGCACGGT
ATCGCCGTCGACGATCAGGTCGCCCGCCGCCTGCTGGAACAGGGTCAGGTTGGCCTGGTTCTCCAGCATGGTGCGGATCGCGGCCTTGTAGCGGACCCGG
TCGGCCTGAGCCCGGGTGGCCCGCACCGCCGGTCCCTTGCGGGCATTGAGCACGCGGAACTGGATGCCGCCGAGGTCGGTTGCCAGGCCCATGGCACCGC
CCAGGGCATCGATCTCCTTGACCAGGTGACTCTTGCCGATCCCGCCGATGGCGGGATTGCAGGACATCTGGCCGAGCGTCTCGATGTTATGGGTCAGCAG
CAAGGTCCGGCAGCCCATGCGGGCCGAGGCCAGGGCGGCTTCGGTTCCGGCATGGCCGCCGCCGATGACGATGACGTCAAAGCGGTCGGGGTACTCCAAG

GGTCACCTCTTGGCGCGGCGCGCCGATGTGGCAGTCAGGGGTGGAAATCAGCGGGCCAGTATAAACCCCCTGTGGGTAATCGTCAGGGTTTTCCACACCC
GAATTCCCTCCCCGCGGTCGTCAACGCACTTCATATACAATAAAACAGAATATAAAAGAGAAGTTCTGTTGTGGTTGTAACTAACTGGTGTGGACATTTTT
**GGTGGATAA**GGGAATAAACCATAATCTTATCAACAGCTTGTATTGTTGAACCCTCGCGGGAGAAAGGCGTGACTGCCTGCGGACGACGTGCGCGGAGGCT
GTGGATGGAATGCCGCCTTGTCCACACGGCGAAGCGATCACCGCTTGTCCACCGCTCCCTACCGGGCTTGTTACCGCGACTGTGAACATTCACCGGCCTG
GCTGCCATCGCCCGGCGTGGCGGGCCTTCCAGGCCGATCGTCGCGACCCGCCGCGATGTTCTGTCCACAGGCAAAAAAAGGCCCTGTCGATAACGACAGG

GCCTTTTTGGATGGGCTGCTGAGGGGCTCCAAAAGGAGCTAGGCAGACGTGGTCAGTGCTTGAGGACCCGCGAGAGGAAGCTCTGGGTGCGCTCGTTCTC
GGGCGAATCGAAGACCTTCTCGGGCGGTCCCTCCTCGGTGATCTCGCCCTTGTGAATGAAGATCACCCGATCGGCGACCTCGCGGGCGAAGCCCATCTCG
TGGGTGACGATCAGCATGGTCATGCCTTCCTTCGCCAGTTCCCGCATGGCGTCCAGCACCTCGCCGATCATCTCCGGATCCAGCGCCGAGGTGGGCTCGT
CGAACAGCATCAGGCGCGGCTCCATGGCCAGCGCACGGGCCAGGGCCACGCGCTGCTGCTGGCCCCCGGAGAGCTGCCCGGGATGCTTGGCGGCCTGGTC
GCTGATGCCGACCCGCTCGAGCAGGCGCTCGGCGGTCTCGGTGGCGTCCTCCTGGCTCCAGCCGCGCACCTTCATCGGCGCCAGGGTGACGTTGTCCAGC
ACGCTGAGGTGGGGGAAGAGGTTGAACTGCTGGAAGACCATGCCGACCTCGGTGCGGATCTTCTGCAGCGCCGCCGAGGCCTTGCCGTGGGGCACCAGCT
GGCTGCCGTCGACCTCCAGGTAGCCCTGCTGGAACTCCTCGAGGCCGTTGATGCAGCGGATCAGCGTCGACTTGCCCGAGCCGCTGGCACCGATGATCAC
CACCACCTCGCCCGGCGAGACCGAGAGGTTGATGTCCTTCAGCACGTGCAGGCTGCCGAAATGCTTGTTGACCTTGTCCATGCGCACGATGGGATCGCTG
CCGGACGTGGACTTCTCGTGTGTCATGTCGCGCGTTCCTTATTGTCCGACCAGGCCTCGGCGCTCGAGCTGGCGCAGCACGAACGAGAGGGTCAGGGTGA
TCGCCAGATAGAGCAGGGCGACCATCAGGTAGACCTCGAGGGCGTTGAAGGTGGTGGCGATATAGACCTGGCCCTGGCGGACCAGTTCGCCGACCCCGAT

>Halomonas denitrificans DSM 18045 [NZ_PYVX01000003.1:c193562..193622], dnaA> ig dnaN>, 61 bp

WebSkew GCmin pos. 1723753 (2nd min.) (GCmin<>oriC1 21.4 kb; GCmin<>oriC2 1.9 kb; oriC2<>oriC1 19.4 kb)

>Halomonas aestuarii Hb3 [NZ_CP018139.1:1744892..1745391], <rpmH ig dnaA>, 530 bp

GACCGAGAGGTCATCGCCGCCGGAGGCGGCCTGGCGGTTGCCGTCCGGGGCGCTGCTGGTCACGGCGGGAGCTTGCGGCTCAGGGGATACCTGTCCGTAG
TCCTGATTCCACTGGACCATCAAGAGGTAGGCGAGTACCGCCAGGGGAATCAGCAGGAGTAGTCGTTTTACGTCCATGAGGAGTCTGCCCGATGGGTAGT
GAACATTCCGATGGCGCAACGCAGGTCGCACGCCCGAGATACACGAAAGCCTGCCGTTGGCAGGCTGGATAACGCACACAGGCCGGTGGCATCCGACGCG
TCGGCGTGAAGGGTCGCGGCGAGATGGGCGGTGCTCATGCTCATGCAGGCTCCGGGATACGCTGGCGCTCAGGGGGGCCGGTCATCATTCGCGGTGCGCG
GCTGCGCTCCGCTTGCCGTGCATGTCCGACTCGCGGGTGGACTTGCCCACCTCGCGCTCAAGGCGACGCCACATCCCCTGGAGCTGGCGATGCACGGTGT
CGTTGTCCAGTTCATGGACTCCCCGTCTTGCCAGTACCACGATATCCACGGCGGGAAGCTGGTGCTGTCGGAGCCGGATGGAGTCACGCACGAGGCGCTT
GAGGCGATTGCGGTCGACGGCCCTGCGCACATTCTTCTTGCTGAAGATCAGGCCGAGACGCGGGTGGCCGAGCGCATTGGGCGAGGCCAGAGCCATCAGC
CCCTTGCCATGGACCTTGTATGCAGCCGTTTCGAAGACACGTCGATAGTCCCCGGCAGTCAGCAGTCTCAGCTGCCGGGGATAGCCCTGACTGGACACGC
GAGGGCCGCTATCAGGCGCTCAGGCGCTTGCGGCCCTTGGCACGGCGACGGGCCAGCACGGCACGACCGTTCTTGGTGGCCATGCGCGCACGGAAACCAT
GCACGCGCTTGCGCTTGAGGACGCTCGGTTGAAAAGTACGTTTCATAACTGGCTATTCCCACGTGGTTTTCTGAATGACTTGATCACTCGTAGACATCCG

GATGCCGCAATCGGACCCGGGGTATTCGATTCAAGACCGGAAATTGTAGTGATTTGCGCCCCGGGGTGCAATTCTTACGTGCGCTCCGGCCCTCTTTGCG
CCGCCGGCAAGGCGCGTGATCCTGGAGGCCAATGCCTCGAGGTCGTCACGGTCGCTCCCTTCCTGTGGTCACTATTGGGGATGAAGATTGTGGGCAACTC
TTATTAATTGTTTATTATCAATGTTCTGAAGGTTAAATCTTGCTGTGGAAAAGGTGGGGGCGATGCGGAAAAGGGCCGGGTTCCCGTGGTGGTGTCCCTT
CCGTTGCCCGACGGGCTCACCGCTTCGGGTCCACGCCGATGCCTCTCGGGTCATGACCAAGCGACGACTGATCAGGACAGGTTTTCCACAGGGTGAGGCG
GCGGGGTGGTCATGTGGATAACCCGGAGCCCGAACGATACAATAGGCGGTCGTTTTCAATGCAAAGGTGAGGTCGCGTGTCACTGGCTCTATGGCAACAA

TGTCTGGATACCCTGCAGGATGAACTGAATTCACAGCAGTTCAATACCTGGATCCGTCCGCTGCAGGCGGAGGAAGGGGAATCCAACCAGTTGCGCCTGC
TGGCGCCGAACCGCTTCGTGCGCGACTGGGTCAGTGACAAGTACGCCAAGCGGATCAGTGAACTGATGCGCGAGCTCTCGCCGGCCAAGCCGCCCAAGGT
GGTGCTGACGGTGGGCAGTCGACGCGCCGCAGCGCCGAATCCGCAGCCCCGTGAGCTGGGCAACCCTGTCTCGGCCACCAAGCGGGCGCCAGCGGCACCG
GCCGTCCAAGTGCCGCCCAGGGGCGACGTCGGGGACGAGCGCGAGATCGACCAGCTGCGGGAGGAGGGGGCCGACCAGCGCCGTGGCGAGCGCCAGGTGC
AGGTGGAGGGCAGCCTCAAGCACAACAGCGGCCTCAATCCCAACTTCACCTTCGAGACCTTCGTCGAGGGCAAGTCCAACCAGCTGGCGCGCGCCGCCTC
CCGCCAGGTCTCCGAGAACCCAGGCGGGGCATACAACCCGCTGTTTCTCTATGGCGGCGTCGGCCTGGGTAAGACCCACCTGATGCATGCCGTAGGCAAC
GCCCTGGCGACCCGTCGCGAGAATGCCCGCGTCGTCTACCTTCACTCGGAGCGCTTCGTGGCCGACATGGTCAAGGCGCTGCAGCTCAACGCGATCAACG
ATTTCAAGCGCTTCTATCGCAGCGTCGATGCCCTGCTGATCGACGACATCCAGTTCTTTGCCGGCAAGGAGCGCTCCCAGGAGGAGTTCTTCCACACCTT
CAATGCGCTGCTCGAAGGCGGCCAGCAGATGATCCTGACGTCCGACCGATATCCCAAGGAAATCAGTGGGGTAGAGGAGCGATTGAAGTCCCGTTTCGGC
TGGGGGTTGACGGTGGCCATCGAGCCGCCGGAGCTGGAGACCCGGGTGGCCATCCTGATGAAGAAGGCCGACCAGGCCAAGGTGGACCTGCCTCATGATG

>Halomonas aestuarii Hb3 [NZ_CP018139.1:1725455..1725954], <mnmG oriC2 <COG1126 (glnQ), 532 bp

GTTCTCGAAGCCCTTGATCGAGCGCACGACCTGCAGCTGCACGTCAAAGGGCAGCGAGGTGGAGATCCCGTTGGGATAGAGCTCGTGGGTATCCAGTCCC
TCCGGCTCGACGAACACCTGGTGGCTGGCCTTGTCGGCGAAGCGGTGGACCTTGTCCTCGATGGACGGGCAGTAGCGCGGGCCAACCCCCTCGATCACGC
CCGAATACATCGGCGAGCGATCGAGGTTGGCGTGGATGATCTCGTGGGTCCGCTCGTTGGTGTGCGCGATATGGCAACTCACCTGCCGGGGATGCATCTC
GCGGCTGCCCAGGTAGGACATCACTGGGGTCGGGGTGTCACCGGGCTGCTCCTCGAGCGCCGCGAGGTCGATGCTGCGGGCATCGAGGCGAGGCGGGGTG
CCGGTCTTGAGGCGATCGACCCGGAACGGCAGGGCCCGCAGCCGCTCGGCGAGCCCGTTGGAGGGCGGATCCCCGGCGCGTCCGCCACGGCTGGTGTCGA
GGCCGATATGGATCACCCCGCCGAGGAAGGTGCCGGTGCACAAGACCACGGTCTCGCCGAGGAAGCGGATGCCCGTCTCGGTGACCACGCCGCGCACGGT
ATCGCCATCGACGATCAGATCGCCCGCCGCCTGTTGGAACAGCGTCAGATTGGCCTGGTTTTCCAGCATGCCGCGGATCGCCGCCTTGTAGCGGACCCGG
TCGGCCTGAGCCCGTGTGGCACGCACAGCCGGACCCTTGCGGGCGTTGAGCACGCGGAACTGGATGCCACCGAGGTCGGTTGCCAGGCCCATGGCCCCCC
CCAGGGCATCGATCTCCTTGACCAGATGGCTCTTGCCGATCCCGCCGATGGCGGGATTGCAGGACATCTGGCCCAGCGTCTCGATGTTGTGGGTCAGCAG
CAGGGTCTGGCAGCCCATGCGGGCCGAGGCCAGGGCGGCCTCGGTGCCGGCATGGCCGCCGCCGATGACGATGACGTCAAAGCGGTCGGGATACTCCAAG

GGTCACCTCTTGGCGCGGTGCGCCGATGTTGCAGTCAGGGGTGGAAATCAGCGGGCCAGTATAAACCCCCTGTGGGTAATCGTCAGGGTTTTCCACACCG
GATTTTCCTTCCCGCTGTCCTTGGTTCCCTACCTAAACAATAAAACAGAATATAAAAGAGAAGTTCTGTTGTGGTTGTAACTAACTGGTGTGGACATTTTC
**GGTGGATAA**GGAAATTAACCTTATATCTTTCAGAATCTTGGGTTGTTGATCAATGGAAGGAGAAAGGCGTGACGTCCTGCGGACGAGGTGTCCGGAGGCT
GTGGATGGAATGTCGGCTTGTCCACACGTTAAAACGGACACCGCTTTTCCACCGCTCCCTACCTGGCTTGTTACCGCGACTGTGAACAATCGGCCTAAGG
ACGTCCATCCCCCCGTGTGACGCAGGCTTCAGGCCGATCGGGGCCATGCCGCCGGGAATTCTGTCCACAGGAAAAAAAAGGCCCTGTCGATAACGACAGG

GCCTTTTCCGAGCGGAGGAAGGAGGCGGGATTCAGTGCTTGAGGACCCGCGAGAGGAAGCTCTGGGTCCGTTCGTTCTGCGGAGAATCGAAGACCTTCTC
GGGCGGGCCCTCCTCGGTGATCTCGCCCTTGTGAATGAAGATCACCCGGTCGGCGACCTCACGGGCGAAGCCCATCTCGTGGGTCACGATGATCATGGTC
ATGCCCTCCCGGGCCAGCTCGCGCATGGCGTCCAGGACCTCGCCGATCATCTCCGGGTCCAGGGCCGAGGTGGGCTCGTCGAACAGCATCAGACGCGGTT
CCATGGCCAGTGCACGCGCCAGGGCCACCCGCTGCTGCTGGCCGCCGGAGAGCTGCCCGGGGTGCTTGGCGGCCTGGTCGCTGATGCCGACCCGCTCGAG
CAGCCGTTCGGCGGTCTCGGTCGCATCGTCGCGGCTCCAGCCGCGCACCTTCATCGGGGCGAGGGTCACGTTGTCCAGCACGCTCAGGTGCGGAAAGAGG
TTGAACTGCTGGAAAACCATGCCGACCTCGGTGCGGATCTTCTGCAGCGCCGCCGAGGCCTTGCCGTGGGGCACCAGCTGGCTGCCATCGACCTCCAGGT
AGCCCTGCTGGAACTCCTCGAGGCCGTTGATGCAGCGGATCAGCGTCGACTTTCCCGAGCCGCTGGCGCCGATGATCACCACCACCTCGCCGGGCGAGAC
CGTCATTTCGATGTCCTTGAGGACGTGCAGGCTGCCGAAATGCTTGTTGACCTTGTCCATGCGCACGATGGGATCGACGCCGGACGTGGACTTGTCGTCT
TTCATGTCGCGTGTTCCTTATTGTCCGACCAGGCCCTTGCGCTCCAGCTGGCGAAGCACGAACGAGAGGGTCAGGGTGATCGCCAGATAGAGCAGGGCGA
CCATCAGGTAGACCTCCAGGGCGGTGAAGGTGGTGGCGATGTAGACCTGCCCCTGGCGGACCAGCTCACCGACCCCGATCACCGAGAACAGCGAGGTGTC

>Halomonas aestuarii Hb3 [NZ_CP018139.1:1746835..1746895], dnaA> ig dnaN>, 61 bp

WebSkew GCmin pos. 35000 (GCmin<>oriC1 27.7 kb; GCmin<>oriC2 8.6 kb; oriC2<>oriC1 19.1 kb)

>Halomonas ventosae CECT 5797 [NZ_SNZJ01000018.1:c7069..7568], <rpmH ig dnaA>, 510 bp

CCGGCAGTCGCCGGGACCGAGAGGTCGTCGCTGCCCGTGGATGCGGGGTCCTGGCGGCTCCCTTCCGGGGCGCTGCTGGTGATGGCGGGAGCTTGCGGCT
CAGGGGTCACCTGTCCGTAGTCCTGATTCCACTGCACGATCAGCAGGTACGCGAGTACCGCCAGGGGAATCACGAGGAGAAGTCGTTTTACGTCCATGAG
GGGTCTGCCCGATGGGTAGTGAACATTCCGATGGCGCAACGCAGGTCGCCCACCCGAGATACACGAAAGCCTGCCGAAGGCAGGCTGGACAACGCGTGCA
GGCTGGCTGCATCCGAGGCGTCAGCGTGGAGGGGCGCAACGAGGCAGGCAGTGGTCATGTCCATGCCGGCTCCGGGGTGCACCGGCACGCTGGGGTGCCG
GTCACCGTGTGCGGCCCGCGTTACCGCTCTCCGGGCTGGGCGAGTCCATCCCGGGGGACGACCTGCGCGCCTCTCGCTCCAGGCGACGCCACATCCCGTG
GAGCTGGCGATGCACGGTGGCATTGTCCAGTTCATGGACTCCTCGTCTTGCCAGCACCACGATATCCACCGAGGGCAGGCGGTGCTGCTGAAGCCGGATG
GATTCACGCACGATGCGCTTGAGGCGGTTTCGGTCGACGGCTCTGCGGACGTTCTTCTTGCTGAAGATCAGGCCGAGTCGCGGATGTCCTCGCTCGTTGG
GAGAGGCGAGGGCCATCAGCCCCTTGCCGTGGACCTTGAATGCAGCCGTTTCGAAGACACGTCGATAATCCCCGGCGGTCAGCAGTCGCAGCTGCCGGGG
ATAGCCCTGACTGGACACGCGAGGGCCGCGGTCAGGCGCTCAGGCGCTTGCGGCCCTTGGCACGGCGACGGGCCAGTACGGCACGACCATTCTTGGTGGC
CATGCGTGCACGGAAGCCATGCACGCGCTTGCGCTTGAGGACGCTGGGTTGAAAAGTACGTTTCATAACTCGTTATTCCCACGTGGTTTTCGGAACGAAT

CGATCATTCGGAGACATCCGGATGCCGCTGGCGGACCCGGGGGTATTCGGTTCAAGACCGGAAATTGTAGTGATTTCAGGCCCGCGGTGCAATTCTTCCG
TGCGTCCCGATGCTGCCGGCCGGGCATGGCACCCCGCTGTCCTCTCGTCGCGCCCTCAGGCAATCACCGGCAGGGGGCAAGCCCTGTTATCACTATTGGG
GATATGTATTGTGACCAAGCCCGTATAATCTCTTTTAATCAATATGTTAGAGATAAAGGATTGCTGTGGACAAGGTGGGGATGCGGCGGGGATTGCCTGC
CCTGGCGCCCGGCAGGTAGCGGCGACCTGCGCGGGTCGCCTCTACCTGTCGAATACGCACTCGGTGCGGGTCGCGCAGGGGTTTTCCACAGACCGCGACT
CGGCGCGGGGCATGTGGATAACCCGTCGTCTGAACGATACAATAGGCGGTCGTTTTCAATGCAATGGTGAGGTCGCGTGTCACTGGCTCTCTGGCAACAA

TGTCTGGATACCCTGCAGGATGAACTGAATTCCCAGCAGTTCAACACCTGGATCCGCCCGCTGCAGGCGGAGGAAGGGGAGTCCAACCAGTTGCGCCTCC
TGGCGCCGAACCGCTTCGTGCGTGACTGGGTAAGCGACAAGTATGCCAAGCGGATCAGCGAGCTGATGCGTGAGCTCTCGCCCGCCAAGCCGCCGAAGGT
GGTGCTGACCGTCGGCAGTCGCCGCGCGGCGCCGAGCCCTCAACCTCGCGAGCTGGGCAATCCCGTCTCGGCGACTCGAGCGCCAGCCGCACCGGCAGTC
CAGGTGCCTCCGCGTGGTGATGTGGGGGATGAGCGCGAGATCGATCAGCTGCGCGAGGAGGGCGCCGGCCAGCGGCGTGGCGGCGAGCGCCAGGTGCAGG
TGGAGGGGAGCCTCAAGCACAGCAGTGGCCTGAATCCCAACTTCACCTTCGAGACCTTCGTCGAGGGCAAATCGAACCAGCTGGCCAGGGCCGCTTCCCG
CCAGGTGTCCGAGAATCCCGGCGGCGCCTACAACCCGCTGTTCCTCTATGGCGGGGTCGGCCTCGGCAAGACCCACCTGATGCATGCCGTGGGTAACGCC
CTGGCAACGCGACGCGAGAATGCCCGCGTCGTTTACCTGCATTCCGAGCGCTTCGTGGCCGACATGGTCAAGGCGCTGCAGCTCAACGCCATCAATGACT
TCAAGCGTTTCTATCGCAGCGTCGATGCCCTGCTGATCGACGATATCCAGTTCTTCGCGGGCAAGGAGCGCTCCCAGGAGGAGTTCTTCCATACCTTCAA
TGCCCTGCTCGAGGGCGGCCAGCAGATGATCCTGACATCTGATCGATACCCCAAGGAAATCAGTGGGGTAGAGGAGCGCCTGAAGTCGCGTTTCGGGTGG
GGGCTGACGGTGGCCATCGAGCCGCCGGAGCTGGAGACCCGGGTGGCGATCCTGATGAAGAAGGCCGACCAGGCCAAGGTCGACCTGCCCCACGACGCCG

>Halomonas ventosae CECT 5797 [NZ_SNZJ01000018.1:c26145..26644], <mnmG oriC2 <COG1126 (glnQ), 531 bp

GTTCTCCAGCCCCTCGATGGAGCGCACCACCTGCAGCTGCACGTCAAAGGGCAGCGAGGTGGAGATCCCGTTGGGGTAGAGCTCGTGGGTGTCGAGGCCT
TCCGGCTCGATGAACACCTGGTGGCTCGCCTTGTCGGCAAAGCGATGCACCTTGTCCTCGATGGAGGGGCAGTAGCGTGGCCCGACCCCCTCGATCACCC
CGGAGTACATCGGTGAACGGTCGAGGTTGGCGAGGATGATCTCGTGGGTGCGCTCGTTGGTGTGCGCGATATGGCAGCTCACCTGCCGGGGATGCATCCC
GCGGTTGCCGAGGTAGGACATCACCGGCGTCGGGCTGTCGCCGGGCTGCTCCTCGAGACACGAGTAATCGACGCTCTTGGCGTCGAGCCGGGGCGGCGTT
CCCGTCTTCAGGCGGTCGACCCGGAACGGCAACGCCCGCAGGCGTTCCGCCAGGGCGTTGGAGGGCGGGTCTCCGGCGCGGCCACCACGGCTGGTGTCGA
GGCCGATATGGATCACCCCGCCGAGGAAGGTTCCGGTGCACAGCACCACGGTCTCGGCCATGAAGCGAATACCGGTCTCGGTGACCACGCCGCGCACGGT
ATCACCCTCGACGATCAGGTCGCCCGCCGCCTGCTGGAAGAGCGTCAGGTTGGCCTGGGTCTCCAGCATGGTGCGAATCGCCGCCTTGTAGCGGACCCGA
TCGGCCTGGGCCCGGGTGGCACGTACTGCCGGTCCCTTGCGGGCATTGAGCACGCGGAACTGGATGCCACCCTGGTCGGTTGCCAGGCCCATGGCACCAC
CCAGGGCGTCGATCTCCTTGACCAGATGGCTCTTGCCGATCCCGCCGATGGCGGGATTGCAGGACATCTGGCCGAGGGTCTCGATGTTATGGGTCAGCAG
CAGGGTCCGGCAGCCCATGCGGGCCGAGGCCAGGGCGGCTTCGGTTCCGGCATGGCCGCCGCCGATGACGATGACGTCAAAGCGGTCGGGATACTCCAAG

GGTCACCTCTTGGCGCATGGCGCCGATGTTGCAGTCAGGAGTGGAAATCAGCGGGGCAGTATAAACCTCCTGTGGGTAATCGTCAGGGTTTTCCACATCC
GATTGCCCAGGCCGATGGTCCTCTCTTCACTCCTAAACAATAAAACAGAATATAAAAGAGAAGTTCTGTTGTGGTAGTAACTAACTGGTGTGGACATTTTC
**CGTGGATAA**CCGTATCAGTTTCATTTTTTTCAACAGCTTGCGTTGTTGACTCCTGCCGGGACAGGAGGGTGACTGGCTGCGGATGAGGTGTCGGGAGCCT
GTGGACGGAAAGCTCGCTTGTCCACAGCGCGACTCGGACACCGCTTGTCCACCGCTGTCTACCCGGCTTGTCACCGTGATGGTGAACAACCTGCGGTGAT
CCATCCGATGCCGCGCATGGCAGGGGCTGCAGGCCGTTGGAGCCCGTTTTTTCGGATTTTCTGTCCACAGGCAAAAAAAGGCCCTGTCGATAACGACAGG

GCCTTTGCGGGAGGGTGCCTGTGAGCGAGATCAGTGCTTGAGAACCCTCGACAGGAACTGCTGGGTCGGCTTGTTGCGGGGATGGTCCAGGACCTCTTCG
GGCGGCCCCTCCTCGGCGATCTCGCCCTGGTGGATGAAGATCACCCGGTCGGCGACCTCCCGGGCGAAGCCCATCTCGTGGGTCACGATCATCATGGTCA
TGCCCTCTTTCGCCAGCTCACGCATGGCATCGAGGACCTCGCCGATCATCTCCGGGTCGAGCGCCGAGGTCGGCTCATCGAACAGCATCAGGCGCGGCTC
CATCGCCAGGGCGCGCGCCAGGGCCACACGCTGCTGCTGACCGCCGGAGAGCTGCCCGGGGTGCTTGGCCGCCTGGTCGCTGATGCCCACCCGCTCGAGC
AGCCGTTCGGCCGTCTCGATGGCATCCGCACGGCTCCAGCCGCGCACCTTCATGGGTGCCAAGGTCACGTTGTCGAGCACGCTCAGGTGCGGGAAGAGGT
TGAACTGCTGGAAGACCATGCCGACCTCGGTACGGATCTTCTGCAGTGCCTGGCCGGCCTTGCCATGGGGCAGCAGCTCGTTGCCATCGACCTCGATATG
GCCCTGCTGGAACTCCTCGAGGCCATTGATGCAGCGGATCAGCGTCGACTTGCCCGAGCCGCTGGCACCGACGATCACCACCACTTCCCCGGGCGAGACC
TCCAGATCGATGTCCTTCAGGACGTGCAGGTCGCCGAAGAACTTGTTGACCTTGTCGAGGCGCACGATGGGGTCGGCGCCGGAAGAGGACTTGTCTTGGT
TCATGTCGCGCTTTCCTTATTGTCCGGCCAGGCCCTTGCGTTCGATGATGCGCAGCACCAGCGACAGGGAGAGCGTGATCGCCAGATAGAGCATGGCGAC
CATCAGGTAGACCTCCAGGGCAGTGAAGGTGGTGGCGATATAGACCTGGCCCTGACGGACCAGCTCGCCGACCCCGATCACCGAGAACAGCGAGGTGTCC
TTGATGCTGATGATGCCCTGGTTGCCCAGCGCCGGGATCATGCGCCGGAAGGCCTGGGGCCAGATGATGTAGCGGAAGGTCTGGCGTCGAGAGAGCCCCA

>Halomonas ventosae CECT 5797 [NZ_SNZJ01000018.1:c5628..5568], dnaA> ig dnaN>, 61 bp

WebSkew GCmin pos. 174020 (GCmin<>oriC1 27.4 kb; GCmin<>oriC2 8.7 kb; oriC2<>oriC1 18.7 kb)

>Halomonas fontilapidosi CECT 7341 [NZ_JACHXQ010000006.1:c146362..146861], <rpmH oriC1 dnaA>, 508 bp

GGGACCGAGAGGTCATCGCTACCCGAGGATGCCGGGTCCTGTCGAGCTCCTTCCGGGGCATTGCTGGTGATGCCGGGAGCTTGCGGCTCAGGGGTTACCT
GCCCGTAGTCCTGATTCCACTGGACGATCAGCAGATAGGCGAGAACCGCCAGAGGAATCACGAGGAGAAGTCGTTTTACGTCCATGAGGAGTCTGCCCGA
TGGGTAGTGAACATTCCGATGGCGCAACGCAGGTCGCACGCCCGAGACATACGAAAGCCTGCCGGTGGCAGGCTGTACAACGCGTGCTGGTTGGCTGCAT
CCGAGGCAACAGCAAGGAGGGGCCCTGCGGGGAGGGCGGTGGTGGGACCCATGCTGGCACCGGTTCGCACCGATACGCTGAAGCGCTGGTCATCGTGCGC
GGCCCGCGTCACCGCTATCCATGCTGGACGCGTCTCCTCCGCAGGACGCTCTGCGCGTCTCTCGCTCCAGGCGACGCCACATCCCGTGGAGCTGGCGATG
CACGGTGGCATTGTCCAGTTCATGGACTCCTCGTCTTGCCAGTACCACGATATCCACCGAAGGCAGGCGATGTTGCTGAAGACGGATGGACTCTCGCACG
ATGCGCTTGAGGCGGTTTCGGTCGACGGCTCTGCGGACATTCTTCTTGCTGAAGATCAGGCCGAGGCGTGGATGATCTCGCTCATTGGGCGAGGCGAGGG
CCATCAGCCCTTTGCCGTGGATCTTGAATGCAGCCGTTTCGAAGACACGTCGATAATCCCCGGCAGTCAGCAGTCGCAGTTGCCGGGGATAGCCCTGATT
GACCACGTAAGAGCCGCGGTCAGGCGCTCAGGCGCTTGCGGCCCTTGGCACGACGACGTGCCAGTACAGCTCGACCATTCTTGGTGGCCATGCGCGCACG
AAAGCCATGCACGCGCTTGCGCTTGAGTACGCTGGGTTGAAAAGTACGTTTCATAACTCGTTATTCCCACATGGTTTTTGGAACGAATCGATCATTCGGA

GGCATCCGGATGCCGCTGACGGACCCGGGGGTATTCGGTTCAAGACCGGAAATTGTAGTGATTTCAGCGTCGCGGTGCAATTCTTCCCTGCGTCTCGATG
CGGTCTGCAAGGTATGTGGCCCAGCTTCTCAGTCCACTGTCGTCGCGGCAGGTTCGCTGCCATGGATTGGGATCCTGTTATCACTATTGGGGACAGATAT
**TGTGGAAAA**GTCTTTTATATAATTTAATATCAATGATTTGTGGCTATGTCCTTGCTGTGTACAAGGTGGGGATGTGGTGGGGATGGCCGGCAACGGGGTC
AGTCACAATGGAGGCATGGTCTGGCTTGGATCTTGAGGCCTGTGCCCACTTCAGGGTTGCCGAAAGGTTTTCCACAGGTAGTGTGGCTGCCGGCGCCATG
TGGATAACCCTGCCACTGAACGATACAATAGGCGGCCGTTTTCGATGCAATGGTGAGGTCGCGTGTCACTGGCTCTATGGCAACAATGTCTGGATACCCT

TCAGGATGAACTGAATTCCCAACAGTTCAATACCTGGATCCGCCCGTTGCAGGCGGAAGAAGGGGACTCCAACCAGTTGCGCCTGCTGGCGCCTAATCGC
TTTGTGCGTGACTGGGTCAGCGACAAGTATGCCAAGCGAATCAATGAGCTGATGCGTGAACTCTCGCCTGCCAAGCCGCCCAAGGTGGTGTTGGCCGTCG
GCAGTCGCCGCGCTGCTCCAAGCCCGCAGCCTCGAGAACTGGGCAATCCCGTCTCGGCAGCACGGAGCCCTGCCGCTCCGGCGGTCCAGGTACCTCCTCG
GGGAGACGTGGCAGACGAACGCGAGATCGATCAACTTCGCGAGGAAGGCCCTAACCAACGGCGGGGCGGCGAGCGCCAGGTGCAGGTAGAGGGAAGCCTC
AAGCACAGCAGTGGCCTCAACCCCAACTTCACCTTCGAAACCTTTGTCGAGGGCAAGTCGAATCAGCTGGCCAGAGCGGCTTCCCGCCAGGTGTCCGAGA
ATCCCGGGGGTGCCTACAATCCGCTGTTCCTCTACGGTGGGGTGGGGCTGGGCAAGACCCACCTGATGCATGCGGTAGGCAATGCCCTGGCAACGCGCCG
CGAGAATGCCCAAGTGGTCTACCTGCATTCCGAACGTTTCGTGGCCGACATGGTCAAGGCCCTTCAGCTCAATGCCATCAACGACTTCAAGCGCTTCTAT
CGCAGCGTCGATGCGCTACTGATCGACGATATCCAGTTCTTCGCCGGCAAGGAACGCTCCCAGGAGGAGTTTTTCCATACCTTCAATGCGCTGCTTGAGG
GCGGCCAGCAGATGATCCTGACGTCTGATCGATATCCCAAGGAAATCAGCGGAGTAGAAGAGCGTCTGAAGTCACGTTTTGGCTGGGGGCTGACGGTAGC
GATCGAGCCGCCCGAATTGGAGACCCGAGTCGCAATCCTGATGAAGAAGGCGGATCAGGCCAAGGTCGATCTTCCTCACGATGCTGTCCTCACGATGCTG

>Halomonas fontilapidosi CECT 7341 [NZ_JACHXQ010000006.1:c165367..165866], <mnmG oriC2 <COG1126 (glnQ), 535 bp

TCTCCAACCCGGTGATGGAGCGCACCACCTGCAGCTGCACATCGAAGGGCAGTGAGGTGGAGATGCCGTTAGGGTAGAGCTCGTGGGTATCGAGGCCTTC
CGGTTCGATGAACACCTGGTGGCTCGCCTTGTCGGCAAAGCGATGCACCTTGTCCTCGATGGAGGGGCAGTAGCGCGGTCCGACCCCTTCGATCACCCCG
GAGTACATCGGCGAACGGTCGAGGTTGGCGAGGATGATCTCATGGGTGCGCTCGTTGGTGTGCGCGATATGGCAGCTCACCTGACGAGGGTGCATCTCGC
GATTACCCAGGTAGGACATCACCGGCGTCGGGCTGTCACCGGGCTGCTCCTCGAGATGCGAGAAATCGACGCTCTTGGCATCGAGCCGGGGTGGTGTTCC
CGTTTTCAGGCGATCAACCCTGAACGGCAATGCCCGCAGCCGTTCGGCCAGGGCGTTGGAGGGCGGGTCACCGGCACGGCCGCCGCGGCTGGTGTCGAGG
CCGATATGGATCACCCCGCCGAGGAAGGTCCCGGTGCACAGCACCACGGTCTCGGCCTTGAAGCGGATACCGGTCTCGGTGACCACCCCGCGCACGGTAT
CGCCCTCGACGATCAGATCACCGGCCGCCTGCTGGAAGAGGGTCAGGTTGGCCTGGGTCTCCAGCATGCGGCGAATCGCTGCCTTGTAGCGGACTCGGTC
GGCCTGGGCGCGGGTAGCGCGTACCGCCGGGCCCTTGCGGGCATTGAGCACACGGAACTGGATGCCGCCGATATCGGTTGCCAGGCCCATGGCACCGCCC
AGAGCATCGATCTCCTTGACCAGATGGCTTTTGCCGATCCCGCCGATGGCGGGATTGCAGGACATCTGGCCAAGGGTCTCGATGTTGTGGGTCAGCAGCA
AGGTCCGGCAGCCCGTGCGGGCCGAAGCCAGCGCAGCTTCGGTTCCGGCATGGCCGCCGCCGATGACGATGACGTCAAAGCGGTCGGGATACTCCAAGGG

TCACCTCTTGGCGCAGGGCGCCGATGTTGCAATCACAAGAGTAGAAATCAGCGGGGCAGTATAAACCTCCTGTGGGTAATCGTCAGGGTTTTCCACACAT
GGATTTCGCATCCTGTTGCAGCGCTCGCTACTCCTAAACAATAAAACAGAATATAAAGAGAAGTTCTGTTGTGGTAGTTACTACTGGTGTGGACATTTTT
**CGTGGATAA**CCAGGGAAAAGTGCTTTTTTACATCAACTTAGAGTGTTGACGCCTGGGGGGAAAGCCAAGTGACCACCTGCGGATGAGGTGTTGGGAGGCT
GTGGATGTAAAGGCGTCTTGTCCACAGCGTGACTCGGACACGTCTTGTCCACCGTCAGCTACCGGGCTTGTCACCGTGGCTGTGAACAACCCGCGAGAGA
TCCTTCCGCGCCGCCAACGACGGGGCTTGCAGGCGGATTTCGCGCAGCTCCACGAACTTTCTGTCCACAGGCAAAAAAAGGCCCTGTCGATAACGACAGG

GCCTTTTTCGTGACGGTGCTGTAGGGGCTGGATCAGTGCTTGAGCACCCTCGACAGGAACTGCTGGGTCGGCTTGTTGCGGGGATGATCCAGGACCTCCT
CGGGTGGCCCCTGCTCGGCGATTTCCCCTTGATGAATGAAGATCACCCGGTCGGCGACATCTCGGGCGAAACCCATCTCGTGCGTCACGATCATCATGGT
CATGCCCTCTTTCGCCAGTTCGCGCATGGCGTCGAGGACTTCGCCGATCATTTCCGGGTCGAGTGCCGATGTTGGTTCATCGAACAGCATGAGGCGTGGC
TCCATCGCCAGGGCTCGCGCCAAGGCGACACGCTGCTGCTGACCGCCGGAGAGCTGCCCTGGATACTTCGCCGCCTGGTCGCTGATACCCACGCGCTCAA
GCAGCCGTTCGGCCGTCTCCATGGCATCCAGACGACTCCAGCCACGCACCTTCATGGGCGCCAGCGTCACGTTGTCGAGCACGCTCAGGTGAGGGAAGAG
GTTAAACTGCTGAAAGACCATGCCGACCTCGGTACGGATCTTCTGCAGTGCCTGGGCGGCCTTGCCATGCGGCAGCAGCTCGTTGCCATCGACCTCGATA
TGCCCCTGCTGGAACTCCTCGAGGCCATTGATGCAGCGGATCAGCGTCGACTTTCCCGAGCCGCTGGCACCAACGATCACCACCACTTCCCCGGGCGAGA
CCTCCAGGTCGATGTCCTTCAGTACGTGCAGGTCGCCAAAGAACTTGTTGACCTTATCGAGGCGCACGATGGGGTTGGCGCTGGAAGAGGACTTTTCTTG
GTTCATGTCGCGCTTTCCTTATTGTCCGGCCAGGCCCTTGCGTTCGATGATGCGTAGCACCAACGACAGGGAGAGCGTGATCGCCAGATAGAGCATGGCG
ACCATCAGGTAGACCTCTAGGGCAGTGAAGGTGGTGGCGATGTAGACCTGGCCCTGACGGACCAGCTCGCCGACCCCGATTACCGAGAACAGCGAGGTGT

>Halomonas fontilapidosi CECT 7341 [NZ_JACHXQ010000006.1:c144875..144935], dnaA> ig dnaN>, 61 bp

< page 1 | #027 | Halomonas daqiaonensis CGMCC 1.9150

WebSkew GCmin pos. 89052 (GCmin<>oriC1 27.6 kb; GCmin<>oriC2 8.4 kb; oriC2<>oriC1 19.0 kb)

>Halomonas daqiaonensis CGMCC 1.9150 [NZ_FOBC01000008.1:c61329..61828], <rpmH oriC1 dnaA>, 508 bp

TCGCGGGTACCGAGAGGTCGTCGCTACCCGACGACGCCGGATCCTGGCTAGCCCCTTCCGAGGCACTGCTCGAGATGGCGGGAGCTTGCGGCTCAGGGGT
CACCTGTCCGTAGTCCTGATTCCACTGCACGATCAACAGGTAGGCGAGTACCGCCAGGGGAATCACGAGGAGAAGTCGTTTTACGTCCATGAGGGGTCTG
CCCGATGGGGAGTGAACATTCCGATGGCGCAACGCAGGTCGCACACCCGAGATACACGAAAGCCTGCCGAGGGCAGGCTGGATAACGCGTGCAGGCTGGC
TGCATTCGAAGCGTCAGCATGGTGGATCGCAGCGAAGATGGCAGTGATGCTCATGCAGGCTCCGGGATGCACTGGCACGCTGAGGTACCGGTCATCGTTC
GCGGCCCGCGTCGCCGGATTCCTGGCTGGGCAAGTCCATCCCAGGGGACGACCTGCGTGCTTCTTTTTCCAGGCGGCGCCACATTCCGTGGAGCTGTCGA
TGCACGGTGGGATTGTCCAGTTCATGGACTCCTCGTCTTGCCAGCACCACGATATCCACCGAGGGCAGACGATGCTGCTGAAGCCGGATGGATTCGCGCA
CGATGCGCTTGAGGCGGTTCCGGTCGACGGCTCTGCGAACATTCTTCTTGCTGAAGATCAGGCCAAGACGTGGATAGCCTCGTCCATTGGGAGAGGCCAG
GGCCATCAGCCCCTTGCCGTGAACCTTGAATACAGCTGTTTCGAAGACACGCCGATAATCCCCGGCGGTCAGCAGTCGCAACTGCCGGGGATAGCCCTGA
CTAGGCACGCGAGGGCCGCGTTCAGGCGCTCAGGCGCTTGCGGCCCTTGGCACGACGGCGGGCCAGTACGGCACGACCGTTCTTGGTGGCCATGCGTGCA
CGAAAGCCATGCATGCGCTTGCGCTTGAGGACGCTGGGTTGAAAAGTACGTTTCATAACTCGTTATTCCCACGTGGTTTTCGGAACAATTGGATCATTAG

GAGGCATCCGGATGCCGCTGGCGGACCCGGGTGTATTCGGTTCAAGACCGGAAATTGTAGTGATTTCAGCCTCGCGGTGCAATTCTTCCTTGCGATCCCA
TGCTGCCGGCAGGGCGAGAGGTGAGGGTGTCCGGTTACCGGCGCTCCTGGTGATCATCGCGGGAGATGTGACCCCTGTTATCACTAATGGGGATATGTAT
**TGTGAACAA**TCCACTATAAGCTTTTGGTATCAGTCACCTACTAGCCTCGTTTTACTGTGGACAAGATGGGGATGTTGTGAATATCTTCTGCCTCGACTGT
TGACACAACAGGCTTGACCCGTAGGTTCCGGGCTCAGCAGCTCCACAGTCTTTGCGGGTCGCAGTGGGGTTTTCCACAGAACGGCGTGTGGTGCTGGCAT
GTGGATAACCCGTTGTCTGAACGATACAATGGGCGGTCGTTTTCAATGCAATGGTGAGGTCGCGTGTCACTGGCTCTCTGGCAACAATGTCTGGATACCC

TGCAGGACGAACTGAACTCCCAGCAGTTCAATACCTGGATCCGCCCGCTGCAAGCGGAGGAAGGGGAGTCCAACCAGTTATGCCTGCTTGCGCCGAATCG
CTTCGTGCGTGACTGGGTCAGCGACAAGTATGCCAAGCGAATCAGCGAGTTGATGCGCGAGCTTTCGCCCGCCAAGCCACCAAAGGTGGTGCTGACCGTC
GGCAGTCGCCGCGCAGCACCTAACCCCCAGCCTCGTGAGCTGGGTAATCCGGTCTCGGCGGCGCGCACCCCCGCAGCACCGGCAGTTCAAGTGCCTTCTT
CCCGCGGGGAAGTCGGTGACGAGCGCGAGATCGATCAGTTGCGCGAGGAGAGTGGTGGTCAGCGACGCGGCGGCGAGCGCCAGGTGCAGGTGGAAGGGAG
CCTCAAGCACAATAGTGGCCTCAACCCCAATTTTACCTTCGAGACCTTCGTTGAGGGCAAGTCCAACCAGCTGGCCAGGGCCGCCTCCCGCCAGGTATCC
GAAAATCCCGGCGGCGCCTATAATCCGCTGTTCCTCTACGGGGGAGTCGGCCTCGGCAAGACCCACCTGATGCATGCCGTGGGTAATGCCTTGGCGACAC
GACGCGAGAATGCACGGGTGGTCTATCTGCATTCCGAGCGCTTTGTGGCCGACATGGTAAAGGCACTTCAGCTCAATGCCATCAACGATTTCAAGCGCTT
CTATCGCAGCGTCGATGCCCTGCTGATCGATGATATCCAGTTCTTCGCCGGGAAGGAGCGCTCCCAGGAGGAGTTCTTCCATACCTTCAATGCGCTGCTC
GAGGGCGGCCAGCAGATGATCCTCACGTCGGATCGATATCCCAAGGAAATTAGTGGGGTTGAGGAGCGCCTGAAGTCGCGTTTCGGTTGGGGACTGACGG
TGGCCATCGAGCCGCCGGAGCTGGAGACTCGGGTGGCGATCCTGATGAAGAAGGCCGATCAGGCCAATGTCGATCTGCCTCACGACGCCGCCTTCTTCAT

>Halomonas daqiaonensis CGMCC 1.9150 [NZ_FOBC01000008.1:c80343..80842], <mnmG oriC2 <COG1126 (glnQ), 533 bp

GTTCTCAAGCCCCGGGATGGAACGAACCACCTGAAGCTGCACGTCGAAGGGCAGCGAGGTGGAAATCCCGTTGGGGTACAACTCATGGGTATCGAGCCCT
TCAGGCTCGATGAACACCTGGTGGCTCGCCTTGTCGGCGAAGCGATGCACCTTGTCCTCGATGGACGGGCAGTAGCGTGGCCCGACCCCCTCGATCACCC
CCGAGTACATCGGCGAACGATCCAGGTTGGCGAGGATGATCTCGTGGGTACGCTCATTGGTATGCGCGATATGGCAGCTCACCTGCCGAGGGTGCATCCC
GCGATTGCCGAGGTAGGACATCACCGGCGTCGGGTTGTCGCCGGGCTGCTCCTCGAGACGCGAGAAATTGACGCTCTTGGCATCCAGCCGTGGCGGCGTT
CCCGTCTTCAGGCGGTCGACCCTGAACGGCAGTGCCCGTAGCCGCTCGGCCAGGGCATTGGAGGGCGGATCGCCTGCACGCCCGCCACGGCTGGTGTCGA
GGCCGATATGGATCACCCCGCCGAGGAAGGTTCCGGTGCACAACACCACGCTTTCGGCCAGGAAGCGGATGCCGGTCTCGGTGACCACGCCACGCACGGT
GTCCCCCTCGACGATCAGATCGCCAGCCGCCTGCTGGAAGAGCGTCAGGTTGGCCTGGGACTCCAGCATGCCACGAATTGCCGCCTTGTAGCGTACTCGG
TCGGCCTGGGCCCGGGTGGCGCGTACTGCCGGCCCCTTGCGGGCATTGAGAACGCGAAACTGGATGCCGCCTAGGTCGGTGGCCAGTCCCATGGCACCAC
CCAGGGCATCAATCTCCTTGACCAGGTGACTCTTGCCGATCCCGCCGATAGCGGGATTGCATGACATCTGGCCGAGAGTCTCGATGTTGTGGGTCAGCAG
CAGGGTCCGGCAGCCCATGCGGGCCGAGGCCAGGGCGGCTTCGGTTCCGGCATGGCCGCCGCCGATGACGATGACGTCAAAGCGGTCGGGATACTCCAAG

GGTCACCTCTTGGCGCACTGCGCCGATGTTGCAGTCAGGAGTGGAAATCAGCGGGCCAGTATAAACCTCCTGTGGGTAATCGTCAGGGTTTTCCACACGG
GATTTCCTGCACCATCGCCTAACTCACCACTCCTAAACAATAAAATAGAATATATAAAAGAAGTTCTGTTGTGGTAGTAACTAACTGGTGTGGACATTTTC
**CGTGGGTAA**GTGGTTTAAACTCAATGTTTTCAAGGTATTGCCGTGTTGATGCCTCGTGGGACATGAGGCTAACTGCCTGCGGATGGGGTGTCGGGAGCCT
GTGGATGGAAAGCCCGCTTGTCCACAGCACGACTGAGACACCGCTTGTCCACCGTTGCCTACCTGGCTTGTCACCGCGACTGTGAACAACCCGACATGAT
TCGCCCATTGCCCCACATGACGGGCACTGCAGGCCGATTCGATCCGGTTCCCCAGGTTTTTTGTCCACAGGCAAAAAAATGGCCCTGTCGATAACGACAG

GGCCTGTGAAGGGCGGTGGAAGGGCGTCGGATCAGTGCTTGAGGACTCTCGACAGGAACTGCTGGGTCGGCTTGTTGCGGGGGCGATCCAGGACCTCTT
CGGGCGGGCCCTCTTCGGCGATCTCGCCCTGGTGTATGAAGATCACCCGGTCGGCGACCTCCCGGGCGAAGCCCATTTCATGGGTCACGATCATCATCGT
CATGCCCTCTCGTGCCAGCTCGCGCATGGCATCGAGGACTTCCCCTATCATCTCGGGATCGAGTGCCGAGGTCGGCTCATCGAACAGCATCAGGCGCGGC
TCCATGGCGAGTGCTCGAGCCAGGGCCACACGCTGCTGCTGACCGCCGGAGAGCTGTCCAGGATGCTTGGCGGCCTGGTCGCTGATGCCCACTCGCTCGA
GCAGGCGTTCGGCCGTCTCGATGGCATCCGCACGACTCCAGCCCCGCACCTTCATTGGTGCCAGGCTTACGTTGTCGAGCACACTCAGGTGCGGGAATAG
GTTGAACTGCTGGAAGACCATGCCGACCTCGGTGCGGATCTTCTGCAGTGCCTGGCCGGCCTTGCCGTGGGGCAGCAACTCGTTGCCATCGACCTCGATA
TGGCCCTGCTGGAACTCCTCGAGGCCATTGATGCAGCGGATCAACGTCGACTTGCCTGAGCCACTGGCACCGACGATCACTACCACTTCCCCGGGCGAGA
CCTTCAGGTCGATGTCCTTCAGCACGTGCAGGTCGCCGAAGAACTTGTTGACCTTCTCCAGGCACACGATGGGGTCGGCGGCGGAAGGAGACTTGTCTTG
GTTCATGTCGCGCTTTCCTTATTGTCCGACCAGGCCTCTGCGCTCCAGCTGTCGCAGCGCAAACGAGAGGCTAAGGGTGATCGCCAGGTAGAGCAGGGCG
ACCATCAGATAGACCTCCAGGGCGTTGAAGGTGGTGGCGATATAGACTTGGCCTTGGCGGACCAGCTCACCGACGCCAATCACCGAGAACAGCGAAGTGT

>Halomonas daqiaonensis CGMCC 1.9150 [NZ_FOBC01000008.1:c59838..59898], dnaA> ig dnaN>, 61 bp

WebSkew GCmin pos. 95128 (GCmin<>oriC1 26.6 kb; GCmin<>oriC2 8.6 kb; oriC2<>oriC1 18.0 kb)

>Halomonas lysinitropha 3(2) [NZ_CABVOU010000015.1:c68300..68799], <rpmH oriC1 dnaA>, 508 bp

CGGGTACCGAGAGGTCGTCGCTTCCCGAGGACGCCGGGTCCTGGCGAGCCCCTTCCGGGGCCTTGCTCGAGATGGCGGGAGCTTGCGGCTCAGGGGTCAC
CTGGCCGTAGTCCTGATTCCACTGGACGATCAACAGGTAGGCGAGTACCGCCAGAGGAATCACGAGGAGAAGTCGTTTTACGTCCATGAGGGGTCTGCCC
GATGGGGAGTGAACATTCCGATGGCGCAACGCAGGTCGCACACCCGAGATACACGAAAGCCTGCCGGAGGCAGGCTAGATAATGCACGCAGGCTGGCTGC
ATTCGATGCATCAGCGTGGAGGATTGCAGCGAAGATGGCAGTAACGATGTTCATGCAGGCTCCGGGATAAACGGGCACGCTGGGGTACCTGTCATCGTTC
GTGGCCTGCTTCACCGGTCTCCTGGCTGGACAAGTCTGCCCCAAGGGGCGACTTTTGTGCTTCTCTCTCCAGTCGCCGCCACATCCCGTGGAGTTGACGA
TGCACGGTGGCATTGTCCAGTTCATGGACTCCTCGTCTTGCCAGCACCACGATATCCACCGAGGGCAGGCGATGCTGCTGAAGCCGGATGGATTCGCGCA
CGATGCGCTTGAGGCGGTTTCGGTCCACGGCCCTACGGACATTCTTCTTGCTGAAGATCAGGCCGAGCCGCGGATAGCCTCGTGCGTTGGGAGAGGCCAG
GGCCATCAGCCCTTTGCCGTGGACCTTGAATGCAGCCGTTTCGAAGACATGTCGATAATCCCCGGCGGTCAGCAGTCGCAACTGCCGGGGAAAGCCCTGA
CGTGACACGCGAGGGCCGCGTTCAGGCGCTCAGGCGCTTGCGACCCTTGGCACGACGACGGGCCAGTACGGCACGACCATTCTTGGTGGCCATGCGCGCA
CGGAAACCATGCATGCGCTTGCGCTTGAGGACGCTGGGTTGAAAAGTGCGTTTCATAACTCGTTATTCCCACGTGGTTTTCGGAACGAATCGATCATTCG

AAGGTATCCGGATACCGCTGGCGGACCCGGGGGTATTCGGTTCAAGACCGGAAATTGTAGTGATTTCAGCCCCGCGGTGCAATTCTTCCTTGCTTCCCTA
TGCCATAAGCAGGGTTGGGGAACGGGTTGCCCGGGGTCTGGCGCTCCAGAAGAGCATTCATATAAAGTGTGACCTTGTTATCACTATGGGGGATAATATT
**TGTGGATAA**CATGCTATATACGTTTAATATCAGCGAGAAGGGAAATTTTTTTATCTGTGGATAAGGTGGGGATGTCAGGGGGACAACTCTGCTCGGCTAG
TGACAGGTCAGGGCGAATTCGTGGCTTCCGGATGTTCCTGCGTCGGAGTCGCTGAGGATCGCAGATAGGTTTTCCACAGCACGGTACGCGCCGTGAGGCA
TGTGGATAACATGCTGCCTGAACGATACAATAGGCGGTCGTTTTCAATGCAATGGTGAGGTCGCGTGTCACTGGCTCTCTGGCAACAATGTCTGGATACC

CTGCAGGATGAACTGAATTCCCAGCAGTTCAATACCTGGATCCGTCCGCTGCAGGCGGAGGAAGGGGAGTCCAACCAGTTGCGCCTGCTGGCGCCGAACC
GCTTCGTGCGGGACTGGGTAAGCGACAAGTATGCCAAGCGGATCAGCGAACTGATGCGTGAGCTCTCGCCCGCCAAGCCGCCGAAGGTGGTGCTGACCGT
CGGCAGTCGGCGCGCAGCGCCCAACCCTCAGCCTCGTGAGTTGGGCAATCCCGTTTCGGCGGGGCGTACTCCCGCCGCACCGGCGGTTCAAGTCTCCCCG
CGGGTGGAGGTGGGGGACGAGCGCGAGATCGATCAGTTGCGCGAGGAGAAGAGTGGCAGCGACGCGGTGGCGAGCGTCAGGTGCAGGTTGAGGGTAGCCT
GAAGCACAACAGTGGCCTCAATCCCAACTTTACCTTTGAGACCTTCGTTGAAGGCAAGTCCAACCAGCTGGCCAGGGCCGCGTCACGTCAGGTATCCGAG
AATCCCGGTGGCGCCTATAATCCGCTGTTTCTCTATGGCGGGGTCGGCCTCGGCAAGACCCACCTTATGCATGCCGTCGGCAATGCCCTGGCGACGCGAC
GCGAGAATGCACAGGTGGTCTACCTGCATTCCGAGCGCTTCGTGGCCGACATGGTCAAGGCGCTGCAACTCAATGCCATCAACGACTTCAAGCGTTTCTA
TCGCAGCGTCGATGCTCTGCTGATCGATGACATCCAGTTTTTCGCCGGCAAGGAACGCTCCCAGGAGGAGTTCTTCCACACCTTCAATGCGCTGCTCGAG
GGTGGCCAGCAGATGATCCTGACGTCTGACCGGTATCCCAAGGAAATCAGTGGAGTAGAGGAACGGCTGAAGTCCCGTTTCGGATGGGGGCTGACGGTGG
CCATCGAGCCACCGGAGCTTGAGACTCGGGTGGCGATCCTGATGAAGAAGGCCGATCAGGCCAAGGTCGATCTCCCCCACGATGCCGCTTTCTTTATTGC

>Halomonas lysinitropha 3(2) [NZ_CABVOU010000015.1:c86316..86815], <mnmG oriC2 <COG1126 (glnQ), 530 bp

CGTTCTCCAGCCCCGCGATGGAGCGTACCACCTGCAACTGCACGTCGAAGGGCAGCGAGGTGGAGATGCCATTGGGATAGAGCTCATGGGTGTCGAGGCC
TTCCGGCTCGATGAATATCTGGTGGCTCGCCTTGTCGGCAAAGCGGTGCACCTTGTCCTCGATGGAAGGGCAGTAGCGTGGCCCGACTCCCTCGATCACC
CCGGAGTACATCGGCGAACGGTCCAGGTTGGCGAGGATGATCTCGTGGGTGCGCTGGTTGGTGTGCGCGATATGGCAGCTTACCTGCCGAGGGTGCAACT
CGCGGTTGCCGAGGTAGGACATCACCGGCGTCGGGCTGTCACCGGGCTGCTCCTCGAGACGGGAGAAATCGACGCTCTTGGCATCGAGCCGGGGCGGCGT
CCCGGTCTTCAGGCGGTCGACCCGGAACGGCAGCGCTCGCAGCCGCTCGGCCAGAGCATTGGAGGGGGGATCACCGGCGCGGCCGCCACGGCTGGTATCG
AGGCCGATATGGATCACCCCGCCGAGGAAGGTGCCGGTGCACAGCACCACGGTCTCGGCCAGGAAACGGATGCCGGTCTCGGTGACCACGCCGCGCACGG
TATCACCTTCGACGACCAGGTCGCCGGCCGCCTGCTGGAAGAGCGTCAGGTTCTCCTGGGTTTCCAGCATGGTGCGAATGGCCGCCTTGTAGCGGACCCG
ATCGGCCTGGGCCCGGGTGGCACGGACCGCCGGCCCCTTGCGGGCATTGAGCACACGGAACTGGATGCCACCGAGATCGGTTGCTAGACCCATGGCACCA
CCGAGGGCATCAATTTCCTTGACCAGATGACTCTTGCCGATCCCGCCGATGGCGGGATTGCAGGACATCTGGCCGAGGGTCTCGATGTTGTGGGTCAGCA
GCAGGGTCCGGCAGCCCATGCGGGCCGAGGCCAGGGCGGCTTCGGTTCCGGCATGGCCGCCGCCGATGACGATGACGTCAAAGCGGTCGGGATACTCCAA

GGGTCACCTCTCGGCGCAATGCACCGAAATCACAGTCATGAGTGGAAATCAGCGGCCCAGTATAAACCTCCTGTGGGTAATCGTCAGGGTTTTCCACATC
GGATCGCATGGATCAACGCTCGCTCTCCACTCCTAAACAATAAAATAGAATATATAAAAGAAGTTCTGTTGTGGTAGTAACTAACTGGTGTGGACATTTTC
**CGTGGATAA**GTAAATAAACTACCATTTTTTCAATGCATTACGGTGTTGACTCCTGAACGGACAAGAGGGTGACTGCCTGCGAACGAGGTGTCGGGAGGCT
GTGGACGGAAAGCTTCCTTATCCACAGCTCACTTCGAACACGGCTTGTCCACCGTCTGCTACCCGGCTTGTTACCGCAACTGTGAACAACCCGACATGAA
GCCTCCATTTCCCCGCATGACGGGGGCTCCAGGCTGCCCGGGCCTGGTTTACCATTTTTTTTGTCCACAGCCAAAAAAAGGCCCTGTCGATAACGACAGG

GCCTTGGCGAGGTACGCGATAAGAGCGGGATCAGTGCTTGAGGACCCTCGACAGGAACTGCTGGGTCGGCTTGTTGCGGGGGCGATCCAGGACTTCTTCG
GGCGGCCCCTCCTCGGCAATCTCGCCCTGATGGATGAAGATCACCCGGTCGGCAACTTCCCGGGCGAAGCCCATCTCGTGGGTCACGATCATCATGGTCA
TGCCTTCTCCTGCCAGCTTACGCATGGCATCGAGGACTTCACCGATCATTTCGGGATCGAGCGCCGATGTCGGCTCATCGAACAGCATCAGGCGTGGCTC
CATCGCCAGGGCGCGGGCCAGGGCCACCCGCTGCTGCTGACCGCCCGAGAGCTGCCCGGGAAACTTAACCGCCTGGTCACTGATGCCCACGCGCTCGAGC
AGTCGCTCTGCCGTTTCGATGGCGTCCAGGCGGCTCCAGCCACGTACCTTCATGGGCGCCAGGGTCACGTTGTCGAGCACGCTCAAGTGCGGGAAGAGGT
TGAACTGCTGGAAGACCATGCCCACCTCGGTACGGATCTTCTGCAGTGCCTTGCCCGCCTTGCCATGCGGCAGCAATTCGTTGCCATCGACCTCGATATG
GCCCTGCTGGAACTCCTCGAGGCCATTGATGCAGCGGATCAGCGTTGACTTGCCTGAACCACTGGCACCGACGATCACGACTACCTCCCCGGGTGAGACC
TCCAGGTCGATATCTTTCAGCACATGCAGGTCGCCGAAGAACTTGTTGACCTTGTCGAGGTGCACGATGGGGTCGGCGGCGGAAGAGGACTTGTCTTGGT
TCATGTCGCGCTTTCCTTATTGTCCGGCCAGGCCCTTGCGTTCGATGATGCGCAGCACCAACGACAGGGAGAGCGTAATCGCCAGATAGAGCATGGCGAC
CATCAGATAGACTTCCAGGGCCGTGAAGGTGGTGGCGATGTAGACTTGGCCCTGACGGACCAGCTCACCGACACCGATCACCGAGAACAGTGAGGTGTCC

>Halomonas lysinitropha 3(2) [NZ_CABVOU010000015.1:c66811..66873], dnaA> ig dnaN>, 61 bp

WebSkew GCmin pos. 14637 (GCmin<>oriC1 36.2 kb; GCmin<>oriC2 8.9 kb; oriC2<>oriC1 27.3 kb)

>Halomonas marinisediminis 204 [NZ_SLTR01000012.1:50575..51074], <rpmH oriC1 dnaA>, 578 bp

TTCATCAGTATTCTCGACTTGGGAGCCGTTGTTTGCGGACTGGCTGCTGGTGAAGGAGGGAGCCTGCGGCTCAGCGGCCATCTGGCCGTAGTCCTGATTC
CACTGTACGACGATCAGATAGGCGAGCACCGCCAGTGGAATCAGCAATAGGAGTCGTTTTACGTCCATGAGGGTTCTGCCCAGTGGGTAGTGAACTTGCC
AGGGGCGGAAGGCGATTCAGCCGCCCGAGATAGACAAAAGCCTGCCAGGGGCAGGCTGTATTCCGAACACGGACTTTCTAGGGTAGCCCCGGCAGCGTCA
AGTCCTGTCTTGCGAAGGAGTGCGCTCATGATCCCGCGAGCTCCGATCCGATGAGGGTGGCTCGATGTTCGGTGGTAGCTGTGTTGCGAACGATTTCTTG
GCGTCGCGAGTCAGCCTTTTCCACATCCCGAACAGCTGCCGGTGCAGGGTCTCATTGTCCAGTTCACCGACTCCGCGTCGTGACAATATCACGATGTCGA
CGGCAGGTAGCCGATGTTGCTGAAATCGGATCGATTCGCGTACCAGCCGTTTGACACGATTACGGTCCACGGCGCGGCGCATGCTCTTCTTGCTGATCAC
AAGGCCGATACGGGGGTGTCCCAGATCGTTGGGCGAGGCGAGTGCCATCAGCCCTTTGCCATGTACTTTAAAGGCAGCGTTCTCGAAGACCTGACGGTAG
TCCCCTGCCGTCAGCAGGCGCAGACGGCGGGGAAACGTTTGACAGGGCACGCAGGGTCCGTGATCAGGCGCTCAGGCGCTTGCGGCCCTTGGCGCGACGA
CGCGCCAGGACGGCACGACCGTTCTTGGTGGCCATCCGGGCACGGAAACCATGCATGCGCTTGCGCTTCAGGACACTGGGCTGAAAGGTGCGTTTCATAA
CTCGTTATTCCCACGTGGTTTTCAGGACGATTCGTCGTTTTCGAGGCGCCTGGATTCCCGCTGGGAACCCGGACGTATTCGGCTCAAGACCGGAAATTGT

AGTGTTTTCCCCCGAGCGGTGCAATTTTTCCGTGTGCTTGTTGCACTTCTCGGCAAGGTAGGGGTGATGGCGATGGCACCGGGAGGTGTGAGATTTCGGC
AGCCAGAGGGAGATTCAGGTACCTAGCAGGCTTGCGCCGCATCAATAACAATAATGTTTATATCTTGTAATGTCTTGTTATTAATGCTGGGGAAGTTGTT
**TGTGGATAA**CTGGTTTTATATCAATTACGACAACTAGTTGGGATGCTATAGAATATGTGGATAGTGACGGGAAAAACGGTGCCCTTCACGCGGGAATCCT
GTGGATGAAAATGGGCCTTCTCCACAACACTGGATTGCCCACAACTCAATACAGAAGGCGTCATCACACCATACGCCTGTTTTCCACAGGGTGTGTCAAG
GTGCCTCGGGGTGTGGATAACCGAGTGTTGTGGCGATACAATGCCCGGCCGTTTTACGACAAGGTGGTGACGTCGCGTGTCGCTGGCTCTCTGGCAACAG

TGTCTGGATACCCTGCAGGATGAGTTGAGCTCGCAGCAGTTCAACACCTGGATTCGCCCGCTTCAGGCAGAGGAGGGCGAAGGGGGAGAGCTGTGCCTGC
TGGCCCCCAATCGCTTCGTGCGCGACTGGGTCAGCGACAAGTTTGCCAAGCGGATCACCGAACTGCTGCGCGAGCTCTCCCCGGCGCGGCCGCCCAAGGT
GGTCCTGAGCGTGGGAAGCCGGCGTGCCGCGCCTCGGCCACAACCGCGGGAGCTCGGCACTCCCGTGTCGGCGACGCCACGGCAGCCGGCAGCTCCGGCG
GTCCATACGCGACCCCGGGGTGAGGTGGCCGACGAGGCCGAGATTCAGGCATCCCGAGACGACGCGCCATCGGCCCGACGACCAGGAGGGGAGCGTCAGG
TCCAGGTGGAGGGGAGTCTCAAGCACACCAGTGGCCTCAATCCCAACTTCACCTTCGAGACCTTTGTCGAGGGCAAGTCGAACCAGCTGGCTCGGGCCGC
CTCGCGCCAGGTATCGGAGAATCCCGGTGGGGCCTATAACCCGCTGTTTCTCTATGGGGGCGTGGGCCTGGGCAAGACTCACCTGATGCATGCCGTAGGC
AATGCCCTGGCGGGCCGGCGCGAGAATGCGCGGGTCGTCTATCTGCATTCCGAGCGCTTCGTGGCCGATATGGTCAAGGCGCTGCAGCTCAACGCCATCA
ACGATTTCAAGCGTTTCTACCGTAGCGTCGACGCCCTGCTGATCGATGACATCCAGTTCTTCGCCGGCAAGGAGCGCTCCCAGGAGGAGTTCTTCCATAC
CTTCAACGCCCTGCTCGAGGGCGGGCAGCAGATGATCCTGACCTCCGATCGTTATCCCAAGGAGATCAGTGGTGTTGAGGAGCGCCTCAAGTCCCGCTTC
GGCTGGGGGCTGACGGTCGCGATCGAGCCCCCGGAGCTCGAGACGCGTGTCGCCATCCTGATGAAGAAGGCCGATCAGGCCAGGGTCGACCTGCCGCACG

>Halomonas marinisediminis 204 [NZ_SLTR01000012.1:23267..23766], <mnmG oriC2 <COG1126 (glnQ), 569 bp

CTCGAGCCCCTTCATGGAGCGTACGACCTGCAGCTGCACATCGAAGGGCAACGAGGTCGAGATGCCATTGGGGTAGAGCTCATGGGTATCGAGCCCCTCG
GGCTCGATAAAGACCTGATGGCTTGCCTTGTCGGCGAAACGGTGGACCTTGTCCTCGATGGAGGGACAGTAACGTGGACCGACCCCCTCGATCGTCCCGG
AGTACATCGGCGAGCGATCGAGGTTGGCATGGATGATCTCGTGGCTGCGCTCGTTGGTATGGGCGATGTGACAGCTTACCTGCCGGGGGTGCTGGTCACG
ACTGCCCATGAACGACATTACCGGGGTGGGTGAATCGCCTGGTTGCTCCTCCAGCTGGGAGAAGTCCAGGCTACGCGCATCCAGGCGCGGCGGTGTGCCG
GTCTTGAGACGATCGACCCGGAATGGCAAGGCACGCAGACGGTCGGCCAGGCGGTTCGAGGGCGGATCACCGGCACGGCCTCCATGAGTGCTGTCGAGCC
CGATATGGATGACCCCACCGAGAAAGGTACCGGTGCACAGCACCACGGTTTCGCCGAGAAAGCGAATACCGGTCTCGGTGACCACGCCCCGTACCTGGTC
ATTTTCGACGATCAGATCACCGGCGGCCTGCTGGAACAGCGTCAGATTCGGCTGATTCTCCAGCAGGCTTCGAATGGCTGCCTTGTAGCGCACCCGGTCG
GCCTGGGCGCGAGTCGCACGCACTGCAGGCCCCTTGCGGGCATTCAGCACGCGGAACTGGATGCCTCCCAGGTCGGTGGCCAGTGCCATGGCTCCACCCA
GGGCATCGATCTCCTTGACCAGATGACTCTTGCCGATGCCCCCGATGGCGGGATTGCAGGACATCTGTCCGAGGGTCTCGATGTTGTGGGTCAGCAACAG
GGTCCGAGAGCCCATGCGCGCGGCGGCCAGGGCGGCTTCGGTTCCCGCATGGCCGCCGCCGATGACGATGACGTCGAAGCGGTCGGGGTAATCCAAGGTG

CACCTCATTGGCGCTATGGCGCCTGTCTTGCGAGCTAGGGATGAAATCAGCCCGCCAGTATAAACCTCCTGTGGGTAATCGTCAGGGTTTTCCACACCCG
CGTCGGCAGAGAGGTCTTTCCTCCGCCCGCCTATCAATTAATATCAGTATATAAATAAAGAAGTTCTGTAATGGTTGTAACTACTGGTGTGGACAATATC
**TGTGGATAA**CCATTTCCTCCCCTTTTTGAT**T**AGTGACTTGCGTTGTTGACCACTAAGAGGAGAAGAGGCTGACAGCCTGCGGACTCAGGGTCCGGAACCT
GTGGATGAAAGGAGGACCTGTCCACAACGGCAACCCTACACCGCTTGTCCACCGTCGGGTACCTGGCTTGTGCCCGTGGCTGTGAACAACCGCCGTAGTT
ATCCACGAACCCCTTGGGAGTGGGCCCCGTAGACGAAATGCAGGACATCAGAAGAAATTCTGTCCACAGTCAAAAAAATGGCCCAGCCGCGATGCGGCTG

GGCCATTTTTCCTTTCCTCAAGGGCAGGGTTCATGAACATGTCTCATGAATAGGACATGGCAATCTCAGTGCTTCAGTACCCGCGAGAGGAAGCTGCGGG
TACGTTCATCGTGCGGGCTATCGAAGATCTTCTCGGGTGGCGCCACCTCGGCGATCTCGCCCTTATGGACATAGATGACTCGATCGGCAACTTCCCGAGC
GAACCCCATCTCATGAGTGACGATGATCATGGTCATGCCTTCCCGGGCCAGCTCACGCATGGCGTCGAGCACCTCGCCGATCATCTCGGGGTCCAGTGCG
GAGGTAGGCTCGTCGAAAAGCATCAGCCGTGGTTCCATGGCCAGCGCCCTGGCCAGGGCAACACGCTGTTGCTGGCCGCCGGAGAGCTGGGCCGGATACT
TCTCGGACTGATCGGCGATCCCGACTCGCTGAAGCAGGCGTTCGGCGGTTTCCACCGCGTCCTCCTGGTTCCAGCCGCGCACCTTCATGGGGGCCAGGGT
GACGTTGTCACGCACGCTCAGATGAGGGAAAAGGTTGAACTGCTGGAACACCATACCCACTTCGGTGCGGATCCTCTGCAGCGCCTTGCCACTCTTGCCG
TAGGGGGTGAGCTCGTTGCCGTCCACATCGATATGGCCGCTCTGAAACTCCTCCAGGCCATTGATGCAGCGAATAAGAGTCGACTTTCCGGAGCCACTGG
CGCCGATGATCACCACCACCTCTCCGGGCATGACCTCCAGGTCGATGTCCTTGAGAACGTGCAGGTTGCCGAAATGCTTGTTGACCTTCTCCATCCGTAC
GATGGCAGAGGTCGTCTTCTCTTTCTGGGTCATGTGTTGGGCTCCCTTATTGGCCGACCAGGCCGCGACGCTCGAGCTGGCGTAGCAGGATCGACAGGGT
CCAGGTGATGGCCAGATAGAGCAGTGCCACCATCAGGTAGACCTCCATGGCGGTGAAGGTGGTGGCGATATAGATCTGGCCCTGTCGCACCAGCTCGCCG

>Halomonas marinisediminis 204 [NZ_SLTR01000012.1:52521..52810], dnaA> ig dnaN>, 289 bp insertion locus tag E0702_RS10090 (183 bp orf, overlapping dnaN)

TGCGTCTGCTGACCAGC**TGA**CCTAGGGCTATCGCTATCGCAGTTATCGGAATTAAGGGGCGCTGGGGACAAGCCGAAGAGGAGGTCCGAGGACCAGGGAT
GGCCGAGGTAGCGTACAGGGAGGTAT**TTA**CAGCGCCTCCTCGTAAGCTTGTCGCCAGATCAGCCCCGGGCTGTCAAGCTCTTTGCGAATGTCCTGCCAGC
TGATCGCTCCGCCCTCCGAGATACTGCCGGCTTGTGGTTTAATGGTCGTCATGCCATCTGGCCGGGCCCCTGCCGGCTGCAACCCCTTCAGGACAAGACG
TGGAGCCGA**CATG**AAATTTTCCATCTCCCGCGAAGCGCTGCTGCGACCGTTGACGCTGGTGGCAGGCGTGGTCGAACGCCGCCAGACCCTGCCGGTACTT

WebSkew GCmin pos. 2546223(2nd min.) (GCmin<>oriC1 35.6 kb; GCmin<>oriC2 8.3 kb; oriC2<>oriC1 27.3 kb)

>Halomonas icarae D1-1 [NZ_WUTS01000001.1:2581558..2582057], <rpmH oriC1 dnaA>, 578 bp

TTCATCAGTATTCTCGACTTGGGAGCCGTTGTTTGCGGACTGGCTGCTGGTGAAGGAGGGAGCCTGCGGCTCAGCGGCCATCTGGCCGTAGTCCTGATTC
CACTGTACGACGATCAGATAGGCGAGCACCGCCAGTGGAATCAGCAATAGGAGTCGTTTTACGTCCATGAGGGTTCTGCCCAGTGGGTAGTGAACTTGCC
AGGGGCGGAAGGCGATTCAGCCGCCCGAGATAGACAAAAGCCTGCCAGGGGCAGGCTGTATTCCGAACACGGACTTTCTAGGGTAGCCCCGGCAGCGTCA
AGTCCTGTCTTGCGAAGGAGTGCGCTCATGATCCCGCGAGCTCCGATCCGATGAGGGTGGCTCGATGTTCGGTGGTAGCTGTGTTGCGAACGATTTCTTG
GCGTCGCGAGTCAGCCTTTTCCACATTCCGAACAGCTGCCGGTGCAGGGTCTCATTGTCCAGTTCACCGACTCCGCGTCGTGACAATATCACGATGTCGA
CGGCAGGTAGCCGATGTTGCTGAAATCGGATCGATTCGCGTACCAGCCGTTTGACACGATTACGGTCCACGGCGCGGCGCATGCTCTTCTTGCTGATCAC
AAGGCCGATACGGGGGTGTCCCAGATCGTTGGGCGAGGCGAGTGCCATCAGCCCTTTGCCATGTACTTTAAAGGCAGCGTTCTCGAAGACCTGACGGTAG
TCCCCTGCCGTCAGCAGGCGCAGACGGCGGGGAAACGTTTGACAGGGCACGCAGGGTCCGTGATCAGGCGCTCAGGCGCTTGCGGCCCTTGGCGCGACGA
CGCGCCAGGACGGCACGACCGTTCTTGGTGGCCATCCGGGCACGGAAACCATGCATGCGCTTGCGCTTCAGGACACTGGGCTGAAAGGTGCGTTTCATAA
CTCGTTATTCCCACGTGGTTTTCAGGACGATTCGTCGTTTTCGAGGCGCCTGGATTCCCGCTGGGAACCCGGACGTATTCGGCTCAAGACCGGAAATTGT

AGTGTTTTCCCCCGAGCGGTGCAATTTTTCCGTGTGCTTGTTGCACTTCTCGGCAAGGTAGGGGTGATGGCGATGGCACCGGGAGGTGTGAGATTTCGGC
AGCCAGAGGGAGATTCAGGTACCTAGCAGGCTTGCGCCGTATCAATATCAATAATGTTCATATCTTGTAATGCCTTGTTATTAATGCTGGGGAAGTTGTT
**TGTGGATAA**CTGGTTTTATATCAATTACGACAACTAGTTGGGATGCTATAAAATATGTGGATAGTGATGGGAAAAACGGTGCCCTTCACGCGGGAATCCT
GTGGATGAAAATGGGCCTTCTCCACAACACTGGATTGCCCACAACTCAATACAGAAGGCGTCATCACACCATACGCCTGTTTTCCACAGGGTGTGTCAAG
GTGCCTCGGGGTGTGGATAACCGAGTGTTGTGGCGATACAATGCCCGGCCGTTTTACGACAAGGTGGTGACGTCGCGTGTCGCTGGCTCTCTGGCAACAG

TGTCTGGATACCCTGCAGGATGAGTTGAGCTCGCAGCAGTTCAACACCTGGATTCGCCCGCTTCAGGCAGAGGAGGGCGAAGGGGGAGAGCTATGCCTGC
TGGCCCCCAATCGCTTCGTGCGCGACTGGGTCAGCGACAAGTTTGCCAAGCGGATCACCGAATTGCTGCGCGAGCTCTCCCCCGCGCGGCCACCCAAGGT
AGTCCTGAGCGTGGGAAGCCGGCGTGCCGCACCTCGGCCACAACCGCGGGAGCTCGGCACTCCCGTATCGGCGATGCCACGGCAGCCGGCAGCTCCTGCG
GTCCATACGCGACCCCGGGGTGAGGTGGCCGACGAGGCCGAGATTCAGGCATCCCGAGACGAAGCGCCATCGGCCCGACGACCAGGAGGGGAGCGTCAGG
TCCAGGTAGAGGGGAGTCTCAAGCACACCAGTGGCCTCAATCCCAACTTCACCTTCGAGACCTTTGTCGAGGGCAAGTCGAATCAGTTGGCTCGGCCGC
CTCGCGCCAGGTGTCGGAGAATCCCGGTGGGGCCTATAACCCGCTGTTTCTCTATGGGGGCGTGGGCCTGGGCAAGACTCACCTGATGCATGCCGTAGGC
AATGCCCTGGCGGGCCGCCGCGAGAATGCGCGAGTCGTCTATCTGCACTCCGAGCGCTTCGTGGCCGATATGGTCAAGGCGCTGCAGCTCAACGCCATCA
ATGATTTCAAGCGTTTCTACCGTAGCGTCGACGCTCTGCTGATCGATGATATTCAGTTCTTCGCCGGCAAGGAGCGTTCTCAGGAGGAGTTCTTCCATAC
CTTCAACGCCCTGCTCGAGGGCGGTCAGCAGATGATCCTTACCTCCGATCGCTACCCCAAGGAGATCAGTGGTGTGGAGGAGCGCCTCAAGTCCCGCTTC
GGCTGGGGACTGACGGTCGCGATCGAACCTCCGGAGCTCGAGACACGAGTCGCCATCCTGATGAAGAAGGCCGATCAAGCCAGGGTCGACCTGCCGCACG

>Halomonas icarae D1-1 [NZ_WUTS01000001.1:2554334..2554833], <mnmG oriC2 <COG1126 (glnQ), 569 bp

CTCGAGTCCCTTCATGGAGCGTACGACCTGTAGCTGCACATCGAAGGGCAACGAGGTCGAGATGCCATTGGGGTAGAGCTCATGGGTATCGAGCCCCTCG
GGCTCGATAAAGACCTGATGGCTTGCCTTGTCGGCGAAACGGTGGACCTTGTCCTCGATGGAGGGACAGTAACGTGGACCGACCCCCTCGATCGTCCCGG
AGTACATCGGCGAGCGATCGAGGTTGGCATGGATGATCTCGTGGCTGCGCTCGTTGGTATGGGCGATGTGACAGCTTACCTGCCGGGGGTGCTGGTCACG
ACTGCCCATGAACGACATTACCGGGGTGGGTGAATCGCCTGGTTGCTCCTCCAGCTGGGAGAAGTCCAGGCTACGCGCATCCAGGCGCGGCGGTGTGCCG
GTCTTGAGACGATCGACCCGGAATGGCAAGGCACGCAGACGGTCGGTCAGGCGGTTCGAGGGCGGATCACCGGCACGGCCTCCATGAGTGCTGTCGAGCC
CGATATGGATGACCCCACCGAGAAAGGTACCGGTGCACAGCACCACGGTTTCGCCGAGAAAGCGAATACCGGTCTCGGTGACCACGCCCCGTACCTGGTC
ATTTTCGACGATCAGATCACCGGCGGCCTGCTGGAACAGCGTCAGATTCGGCTGATTCTCCAGCAGGCTTCGAATGGCTGCCTTGTAGCGCACCCGGTCG
GCCTGGGCGCGAGTCGCACGCACTGCAGGCCCCTTGCGGGCATTCAGCACGCGGAACTGGATGCCTCCCAGGTCGGTGGCCAGTGCCATGGCTCCACCCA
GGGCATCGATCTCCTTGACCAGATGACTCTTGCCGATGCCCCCGATGGCGGGATTGCAGGACATCTGTCCGAGGGTCTCGATGTTGTGGGTCAGCAACAG
GGTCCGAGAGCCCATGCGCGCGGCGGCCAGGGCGGCTTCGGTTCCCGCATGGCCGCCGCCGATGACGATGACGTCGAAGCGGTCGGGGTAATCCAAGGTG

CACCTCATTGGCGCTATGGCGCCTGTCTTGCGAGCTAGGGATGAAATCAGCCCGCCAGTATAAACCTCCTGTGGGTAATCGTCAGGGTTTTCCACACCCG
CGTCGGCAGAGAGGTCTTTCCTCCGCCCGCCTATTAATTAATATCAGTATATAAATAAAGAAGTTCTGTAATGGTTGTAACTAACTGGTGTGGACAATATC
**TGTGGATAA**CCATTTCCTCCCCTTTTTGAT**T**AGTGACTTGCGTTGTTGACCACTAAGAGGAGAAGAGGCTGACAGCCTGCGGACTCAGGGTCCGGAACCT
GTGGATGAAAGGAGGACCTGTCCACAGTGGCAACCCTACACCGCTTGTCCACCGTCGGGTACCTGGCTTGTGCCCGTGGCTGTGAACAACCGCCGTAGTT
ATCCACGAACCCCTTGGGAGTGGGCCCTGTAGACGAAATGCAGGACATCAGAAGAAATTCTGTCCACAGTCAAAAAAATGGCCCAGCCGCGATGCGGCTG

GGCCATTTTTCCTTTCCTCACGGGCAGGGTTCATGAACATGTCTCATGAATAGGACATGGCAATCTCAGTGCTTCAGTACCCGTGAGAGGAAGCTGCGGG
TACGTTCATCGTGCGGGCTATCGAAGATCTTCTCGGGTGGCGCCACCTCGGCGATCTCGCCCTTATGGACATAGATGACTCGATCGGCAACTTCCCGAGC
GAACCCCATCTCATGAGTGACGATGATCATGGTCATGCCTTCCCGGGCCAGCTCACGCATGGCGTCGAGCACCTCGCCGATCATCTCGGGGTCCAGTGCG
GAGGTAGGCTCGTCGAAAAGCATCAGCCGTGGTTCCATGGCCAGCGCCCTGGCCAGGGCAACACGCTGTTGCTGGCCGCCGGAGAGCTGGGTCGGATACT
TCTCGGACTGATCGGCGATCCCGACTCGCTGAAGCAGGCGTTCGGCGGTTTCCACCGCGTCCTCCTGGTTCCAGCCGCGCACCTTCATGGGGGCCAGGGT
GACGTTGTCACGCACGCTCAGATGAGGGAAAAGGTTGAACTGCTGGAACACCATACCCACTTCGGTGCGGATCCTCTGCAGCGCCTTGCCACTCTTGCCG
TAGGGGGTGAGCTCGTTGCCGTCCACATCGATATGGCCGCTCTGAAACTCCTCCAGGCCATTGATGCAGCGAATAAGAGTCGACTTTCCGGAGCCACTGG
CGCCGATGATCACCACCACCTCTCCGGGCATGACCTCCAGGTCGATGTCCTTGAGAACGTGCAGGTTGCCGAAATGCTTGTTGACCTTCTCCATCCGTAC
GATGGCAGAGGTCGTCTTCTCTTTCTGGGTCATGTGTTGGGCTCCCTTATTGGCCGACCAGGCCGCGACGCTCGAGCTGGCGTAGCAGGATCGACAGGGT
CCAGGTGATGGCCAGATAGAGCAGTGCCACCATCAGGTAGACCTCCATGGCGGTGAAGGTGGTGGCGATATAGATCTGGCCCTGTCGCACCAGCTCGCCG

>Halomonas icarae D1-1 [NZ_WUTS01000001.1:2583504..2583610], dnaA> ig dnaN>, 107 bp

WebSkew GCmin pos. 9646 (GCmin<>oriC1 9.2 kb; GCmin<>oriC 29.2 kb; oriC2<>oriC 20.0 kb)

>Halomonas halodenitrificans DSM 735 [NZ_JHVH01000002.1:38583..39082], <rpmH oriC1 dnaA>, 568 bp

```
CCGCCAGGCCTTCTTCGGCGGTTTCGAGCTCGGCGCTACCGTTTCCGGACTGGTTGCTGGTAATGGAGGGAGCCTGCGGCTCGACGGCCGTCTGGCCATA
ATCCTGGTTCCACTGCACGACGATCAGATAGGCGAGCACCGCCAGGGGAATCAGTAGTAGGAGTCGTTTTACGTCCATGAGGGTTCTGCCCAGTGGGTAG
TGAACTTGCCAGGGGCGGAAGGCGATTTGGCCGCCCGAGATAGACAAAAGCCTGCCAAGGGCAGGCTGTATTCCGAACACGGACTTTCGAAGGTTGTCCC
GGCAGCGTCAAGCCTTGCCTTGCGAGGGGGTGCGCTCATGATTCCGTGGGCTCCGATCGGATGAAGATGGCTCGGCGCTTGGTGACGGTTGTCTTGCGAG
CGACTTCTGTGCCTCGCGCGTCAGCCTTTTCCACATCCCGAAAAGCTGCCGATGCAGGGTCTCGTTGTCCAGCTCGCCGACACCGCGTCGCGAGAGTATC
ACGATGTCGACGGCCGGTAGCCGGTGCTGCTGGAATCGGATCGATTCGCGCACGAGACGCTTGATGCGGTTGCGGTCCACGGCACGGCGCATGCTCTTCT
TGCTGATGACAAGGCCAATACGGGGGTGACCCAGATTGTTGGGCGAGGCGAGTGCCATCAGCCCCTTGCCATGTATCTTGAAGGCAGCGTTATCGAAGAC
CTGACGGTAGTCCCCGGCCGTCAGCAGGCGCAGACGACGGGGAAACGACTGACGGGACACGTAGGGTCCGTGATCAGGCGCTCAGGCGCTTGCGACCCTT
GGCGCGACGGCGCGCCAGGACGGCACGTCCGTTCTTGGTAGCCATCCGGGCACGGAAACCGTGCATACGCTTACGCTTCAGAACGCTGGGCTGAAAGGTG
CGTTTCATAAGTCGTTATTCCCACGTGGTTTTCTGGACGATTCGTCGTTTTCGAGGCGCCTGGATTCCCACCGGGAACCCGGGCGTATTCGGTTCAAGAC
```

```
CGGAAATTGTAGTGTTTTCCCTTGGGCGGTGCAATTTTTCCGTACATGAATTGGGCTTTTGATGGCTGGGAGGGGGCGCCTCGGCATGTCGAGATAATGC
GTCGGATCCATTCCGAAGCAGCATTCGAAGGCCGTACCTCGAACTACAATAACAAGGGTATATCTTGTAATGCCTTGTTATTAATACTGGGAAACTTTC
TGTGGATAAGCTTTCATATTTCATCAATTACAAGGCATTGGATGAATGAAACTTGTGTGGACAGCCTCTGCAAGGGTGGTTTGCAAGCCTGGGATAACCT
GTGTATGAAGATGGCACTTCTCCACAGCCCAGCATTACGCACAGTTCGGTCGCGGATGCGTCACCATGTCGTCACCCCGTTTTCCACAAGATGTGTGGAG
GCGTTACGGGGTGTGGATAACCGCGTGGTGGGGCGATACAATGCACGGCCGTTTTTCGACAATGTAGTGAGGTCGCGTGTCGCTGGCTCTCTGGCAACAG
```

```
TGTCTAGATACCCTTCAGGATGAGTTGAGCTCGCAGCAGTTCAACACCTGGATTCGCCCGCTCCAGGCAGAGGAGGGCGAAGGGGGAGAGCTGTGCCTGT
TGGCACCCAATCGCTTCGTACGCGACTGGGTCAGCGACAAGTTTGCCAAGCGGATCACCGAACTGCTGCGTGAGCTGTCGCCGGCACGTCCGCCCAAGGT
CGTGCTGAGCGTGGGGAGTCGACGCACTGCTCCTCGGCCTCAGCCACGCGAGCTCGGCACCCCGGTGTCGGCAATGCCGCGACAACCCGCGGCCCCGGCG
GTACATACACGCTCCCGTGGTGATGTAGCCGATGAAGTCGAGATCGAGTCATCCCGGGAGGAGGCGCCGCCGGCCCGGCGACCGGGAGGCGAGCGTCAGG
TCCAGGTCGAGGGCAGCCTCAAGCATACCAGTGGCCTCAACCCTAACTTCACCTTCGAGACCTTCGTCGAGGGCAAGTCGAACCAGCTGGCCCGGGCCGC
GTCCCGACAGGTGTCGGAGAACCCCGGCGGTGCCTATAATCCGCTGTTTCTCTATGGTGGAGTAGGGCTGGGCAAGACCCACCTGATGCATGCCGTGGGC
AATGCGCTGGCGGGTCGCCGGGAGAATGCGCGGGTGGTCTATCTTCACTCCGAGCGTTTCGTGGCCGATATGGTCAAGGCGCTGCAGCTCAACGCCATCA
ATGACTTCAAGCGGTTCTATCGTAGCGTCGATGCCCTGCTGATCGATGACATCCAGTTCTTTGCCGGGAAGGAGCGCTCCCAGGAGGAGTTCTTCCATAC
CTTCAACGCCCTGCTCGAGGGCGGACAGCAGATGATCCTGACCTCCGACCGCTATCCCAAGGAGATCAGCGGTGTAGAGGAGCGGCTCAAGTCCCGCTTC
GGCTGGGGGCTGACGGTCGCGATCGAGCCGCCCGAGCTCGAGACACGGGTTGCCATCCTGATGAAGAAGGCCGATCAGGCCAAGGTCGAGCTTCCCCATG
```

>Halomonas halodenitrificans DSM 735 [NZ_JHVH01000002.1:18624..19123], <mnmG oriC2 <COG1126 (glnQ), 554 bp

```
TTTCCAGCCCATGCATGGAACGCACCACCTGGAGCTGGACATCGAAGGGCAGCGACGTGGAGATGCCGTTGGGGTAGAGCTCATGCGTATCCAGCCCCTC
GGGCTCGATAAAGACCTGGTGGCTTGCCTTGTCGGCGAAGCGATGGACCTTGTCCTCGATGGAGGGACAGTAGCGAGGACCCGTTCCTTCGATCGTTCCG
GAATACATCGGCGAGCGATCGAGGTTGGCATGAATGATGTCGTGGCTGCGCTCGTTGGTATGGGCGATATGACAGCTCATCTGGCGGGGATGCTGATCGC
GGCTGCCCATGAACGACATTATCGGCGTCGGTGTGTCACCCGGCTGTTCCTCCAGCTTCGAGAAATCCAGGGTCCGCGCATCCAGACGGGGAGGGGTGCC
GGTCTTGAGGCGGTCGACGCGAAACGGGAGAGCACGCAAGCGTTCGGCCAGACGGTTGGAGGGCGGGTCGCCGGCGCGGCCGCCGCGAGTGTTGTCGAGC
CCGATATGGATCACGCCACCGAGGAAGGTGCCGGTGCACAACACGACGGTTTCGCCGAGAAAGCGGATGCCGGTTTCGGTGACCACCCCTCGCACCCGAT
CGTTATCGACGATCAGGTCCCCGGCGGCCTGCTGAAACAGGGTCAGGTTGGGCTGATTTTCCAGCAATCCACGGATGGCCGCCTTGTAGCGTACCCGGTC
GGCCTGGGCGCGAGTGGCTCGCACGGCAGGCCCCTTGCGGGCGTTCAGGACGCGAAACTGGATTCCCCCGAGATCGGTGGCCATCGCCATGGCACCCCCC
AGGGCATCGATTTCCTTGACGAGGTGACTCTTGCCGATGCCGCCGATTGCGGGATTGCAGGACATCTGGCCGAGGGTCTCGATGTTATGGGTCAGCAGTA
GGGTTCGAGAGCCCATACGAGCAGCGGCCAGGGCGGCTTCGGTTCCTGCATGGCCACCGCCAATGACAATGACGTCGAAGCGGTCGGGATAATCCAAGGT
```

```
GCACCTCATTGGCGCCGGGGCGCCTGTCTTGCGGTCAAGGGATGAAATCAGCCCGCCAGTATAAACCTCCTGTGGGTAATCGTCAGGGTTTTCCACACCC
GCCCCGACAACGTCATGACCCCATCCCTCCCTATTAATTAATATCAGTATATAAATAAAGAAGTTCTGTTGTTGTTGTAACTACTGGTGTGGAGGAAATC
TGTTGATAAGCCTTCTAATGTTAGCAATATCAAAAGGATAGACTGGTGACTCTCTTCCGGAGAAACGGCGGGCAGCCTGCGGACTCGCCGCCGAAAAGCT
GTGGATGAAATACCGTTCTGTCCACAGCGCCAGCGCTGCACCGCTTGTCCACCGTCGGGTACCTGGCTTGTGCCCGTGGGTGTGAACAACTGAGGTGGTT
ATCCACGAACCCCTCCATGATGAGGGTTACAGCGGAAATTCAGGCGTTTGCGAAAAATTCTGTCCACAGGCGAAAAAAGGGCCCTGGCGAGCTTCGCCAG
```

```
GACCCTTCCTATCGATATTTCCTTGTTATGACAGAGCGGCAAAGGCCGAGATCAGTGCTGGAGCACCCGCGCCAGGAAGTGACGAGTACGCTCGTCGGTG
GGGCTATCGAAGATCTTCTCGGGAGCCGCCTCTTCGGCGATCTCGCCCTTGTGGACATAGATGACGCGGTCGGCGACTTCGCGGGCAAACCCCATCTCGT
GGGTGACGATGATCATGGTCATGCCTTCCCGGGCCAGCTCGCGCATGGCGTCGAGAACCTCGCCGATCATCTCCGGATCCAGCGCGGAGGTGGGCTCATC
GAACAGCATCAGCCGTGGCTCCATGGCCAGGGCTCGCGCCAGGGCGACGCGCTGCTGCTGGCCACCGGAAAGCTGGGTCGGATACTTATCGGACTGGTCG
GCGATGCCGACGCGGTCGAGCAGGCGGTCGGCGGTCTCCCGCGCGTCTTCCCGGCTCCAGCCGCGTACTCTCAAGGGGGCCAGGGTGACGTTATCGCGCA
CGCTGAGGTGGGGAAACAGATTGAACTGCTGAAACACCATGCCCACCTCGGTACGGATCTTCTGCAGCGCACGGCCGCTCTTGCCGTGGGGCGTCAGCTC
ATTACCGTCCACATCGATATGGCCCTCCTGGAATTCCTCCAGGCCGTTGATGCAACGAATCAGGGTCGATTTACCGGAGCCGCTGGCGCCGATGATCACC
ACCACCTCGCCGGGCATGATCTCCAGGTCGATGTGCTTGAGAACATGCAGACTGCCGAAGTGCTTGTTGACCTTCTCCATGCGTACGATGGCGGTGGTGG
TCGCGTCTTTCTGGGTCATGTGTTTGGCCTCCCTTGCGCGTTGTTAACCCGTTCACTGGCCGACCAGGCCGCGGCGCTCGAGCTGGCGCAGCAGGATCGA
CAGGGTCCAGGTGATGGCCAGGTAGAGCATTGCCACCATCAGGTAGACTTCCATGGCGGTGAAGGTGGTGGCGATATAGATCTGCCCCTGTCGCACCAGT
```

>Halomonas halodenitrificans DSM 735 [NZ_JHVH01000002.1:40529..40634], dnaA> ig dnaN>, 106 bp

WebSkew GCmin pos. 2000000 (2nd min.) (GCmin<>oriC1 30.9 kb; GCmin<>oriC2 12.7 kb; oriC2<>oriC1 18.1 kb)

>Halomonas alimentaria DSM 15356 [NZ_WUTT01000001.1:2030628..2031127], <rpmH oriC1 dnaA>, 569 bp

CCGACAGACCATCCTCGGCGCCTTCGGACCGGGAGTCGCTGTCCCCGGACTGACTGCTGGTAAAGGAGGGAGCCTGTGGCTCGACGGCCGTCTGGCCATA
GTCCTGATTCCACTGCACGACGATCAGATAGGCGAGCACCGCCAGTGGAATCAGCAATAGGAGTCGTTTTACGTCCATGAGGGTTCTGCCCAGTGGGTAG
TGAACATGCCAGGGGCGGAAGGCGTTTCAGCCGCCCGAGATAGACAAAAGCCTGCCACTGGCAGGCTGTATTCCGAACACGGACTCTCGAGGGTAGCCCC
GGCAGCGTCAAGCCTTGCCTTGTGAGGGGGTGCGCTCATGCTTCGGCGGGCTCCGGTCGGATGAAGACGGCTCGACGCTTGATTAGGGATCTCTCGCGGG
CGTCTTCAATGCCTCGCGTACCAGCCTTCGCCACATCCCGAACAGCTGCCGATGCAGGGTCTCGTTATCCAGTTCGCCTACGCCGCGCCGCGAGAGTATC
ACGATGTCGACGGCGGGGAGGCGATGCTGCTGGAAGCGGATCGATTCACGCACCAGTCGCTTGACGCGGTTGCGGTCTACGGCACGGCGCATGCTCTTCT
TGCTGATGACAAGGCCGATGCGGGGATGGCCCAGATCGTTGGGCGAGGCGAGTGCCATCAGCCCCTTGCCATGCACCTTGAAGGTCGCGTTCTCGAAGAC
CTGGCGATAGTCCCCGGCCGTCAGCAGGCGCAGACGGCGGGGAAACGACTGACGGGGCACGCAGGATCCGTGATCAGGCGCTCAGGCGCTTGCGACCCTT
GGCGCGACGACGCGCCAGGACGGCACGTCCGTTCTTGGTGGCCATACGGGCACGGAAACCGTGCGCACGCTTACGCTTCAGGACGCTGGGTTGAAAGGTG
CGTTTCATAACTCGTTATTCCCACGTGGTTTTCTGGACGATTCGTCGTTTGCGAGGCGCCTGGATTCCCGCGGGGAACCCGGACGTATTCGGTTCAAGAC

CGGAAATTGTAGTGATTTCCCCCGGGCGGTGCAATTTTTCTGTGCGCCGATCCGACCGATTGTGCGGCAGCCGTGGCCAGCGAACATCCTTCTGGCATG
GCGATCAGGTTACCGATGTCGCTCGCCAGTCATCCAGGGCTTCCATAACAACAGTGTTTATAACCTGTAATACCTTGTTATTAATACTGGGGAGGCATTC
**TGGGGATAA**GTTATATTAGTCCTTTCAATACAGTATGATGAAGCTATGCTGTTGCTGTGGATAGGCCTGGGATGAGCGGTGTGCGATGCTGGGGTATCCT
GTGGAAGAAAAGCGAGCTCCTCCACAGCCTCTCGTTGCGCACAGTTCGGTCCCGCAGTAGTCACCATGCCAGGGACCCGTTTTTCCACAGGCTGTATAGG
GGTGGTCGGGGGTGTGGATAACCGTGCGGTGGGGCGATACAATGCCCGGCCGTTTTTCGACAAGGTGGTGAGATCGCGTGTCGCTGGCTCTCTGGCAACA

GTGTCTGGATACCCTGCAGGACGAATTGAGCTCTCAGCAGTTCAACACCTGGATTCGCCCGCTCCAGGCCGAGGAGGGCGAAGGGGGAGAGCTGTGCCTG
CTGGCCCCCAACCGCTTCGTCCGTGACTGGGTCAGCGACAAGTTCGCCAAGCGGATCACCGAGCTGCTGCGTGAGCTGTCGCCGGCACGGCCGCCCAAGG
TGGTGCTGAGTGTTGGTAGTCGCCGTGCCGCCCCTCGCCCCCAGCCGCGAGAGCTTGGCACACCGGTGTCGGCGACCCCGCGGCGGCCGGCGGCCCCGGC
GGTGCATACTCCCCCCCGCGGCGCTGCGGCCGACGAGGTCGAGATCGAGGACAGCCGCGAGGAGACGGCGGCTGTCCGGCGCCCCGGTGGTGAGCGCCAG
GTCCAGGTGGAGGGCAGCCTCAAGCACACCAGCGGTCTCAATCCCAACTTCACCTTCGAGACCTTCGTCGAGGGCAAGTCGAACCAGCTGGCTCGGGCCG
CTTCCCGCCAGGTGTCGGAGAACCCCGGCGGTGCCTATAACCCGCTGTTCCTGTATGGCGGGGTGGGCCTGGGCAAGACCCACCTGATGCATGCCGTGGG
CAATGCCCTGGCCGGTCGCCGCGAGAATGCGCGGGTGGTCTACCTGCACTCCGAGCGTTTCGTGGCCGACATGGTCAAGGCGCTGCAGCTCAACGCCATC
AACGACTTCAAGCGTTTCTATCGCAGCGTCGACGCCCTGCTGATCGACGATATCCAGTTCTTCGCCGGCAAGGAGCGCTCCCAGGAGGAGTTCTTCCACA
CCTTCAATGCCCTGCTCGAGGGCGGTCAGCAGATGATCCTGACCTCCGACCGCTATCCCAAGGAGATCAGCGGCGTGGAGGAGCGCCTCAAGTCGCGCTT
CGGTTGGGGCCTGACGGTGGCGATCGAGCCACCGGAGCTCGAGACACGGGTCGCTATCCTGATGAAGAAGGCCGACCAGGCCAAGGTCGACCTGCCCCAC

>Halomonas alimentaria DSM 15356 [NZ_WUTT01000001.1:2012494..2012993], <mnmG oriC2 <COG1126 (glnQ), 543 bp

TTCTCCAGTCCGCGCATGGAGCGCACTACCTGCAACTGCACATCGAAGGGCAGCGAGGTAGAGATGCCGTTGGGATAGAGCTCGTGGGTGTCGAGCCCCT
CGGGCTCGATGAAGACCTGGTGGCTCGTCTTGTCGGCGAAGCGGTGCACCTTGTCCTCGATGGAGGGGCAGTAGCGCGGCCCCACCCCCTCGATCATCCC
GGAGTACATCGGCGAGCGGTCGAGATTGGCGTGGATGATCTCGTGGGTACGCTCGTTGGTGTGGGCGATGTGACAGCTCACCTGGCGGGGATGCTGAGCG
CGATTGCCCATGAACGACATCACCGGGGTGGGGGTATCTCCCGGCTGCTCCTCCAGCTGCGAGAAGTCCAGGCTCCTGGCATCGAGGCGAGGCGGAGTAC
CGGTCTTGAGGCGATCGACACGGAACGGCAGGGCGCGCAGGCGTTCGGCCAGGCGGTTCGAGGGCGGGTCGCCGGCGCGCCCGCCGCGGGTGCTGTCGAG
CCCGATATGGATCACCCCACCGAGGAAGGTTCCGGTACACAGCACCACGGTCTCGCCTAGGAAGTGGATGCCGGTCTCGGTAACCACGCCCCGTACCTCG
TCGTTGTCGACGATCAGGTCGCCGGCGGCCTGCTGGAACAGCGTCAGGTTGGGCTGGTTCTCCAGCAGGCCGCGGATCGCGGCCTTGTAGCGAACCCGGT
CGGCCTGGGCGCGGGTGGCCCTAACCGCGGGCCCCTTGCGGGCATTGAGAACACGGAACTGGATCCCGCCGAGATCGGTGGCCAGGCCCATGGCACCACC
CAGGGCATCGATCTCCTTCACGAGGTGGCTCTTGCCGATGCCGCCGATGGCCGGATTGCAGGACATCTGGCCGAGGGTCTCGATGTTGTGGGTCAGAAGC
AGGGTACGTGAGCCCATGCGAGCGGCGGCCAGGGCGGCTTCGGTTCCCGCATGGCCGCCGCCGATGACGATGACGTCGAAGCGGTCGGGGTAATCCAAGG

TGCACCTCGTTGGCGCCGCGGCGCCTGTCTTTCTGTCAGAGGGTGAAATCAGCCCGGCAGTATAAACCTCCTGTGGGTAATCGTCAGGGTTTTCCACACG
TGTTTCCAACCAGGCTCGTTCCCTCCTGACCTATCAATTAATATCAGTATATAAATAAAGAAGTTCTGTTGTGGTTGTAACTAACTGGTGTGGACAAAAAC
**CGTGGATAA**GTCATTTCCTGCCATTTTCTTCAGTAGCTTACGTTGTTGACCAAGCACAGGAGAAAGGGACGACAGCCTGCGGACTAGCGGTCGCGAACCT
GTGGATGGAATGCCACCTTGTCCACAGTGCCGGCGCTGCACCGCTTGTCCACCGATGGGTACCAGGCTTGTGCCCGCGGCTGTGAACAACTGGGGTATTT
ATCCACGAGCCCCTCCAGCATGGGCCCCGTAGCCAGAATCCGGGGTATCGAAAAAAATTCTGTCCACAGACGAAAAAAGGGCCCTGCCGCAACGCGACAG

GGCCCTTTTTCATCCTGATGGAGTCGATCCTGGTGGTTGAGCTAGTGCTTGAGTACGCGAGAGAGGAAGCTTCGGGTGCGTTCATCATTCGGGCTATCGA
AGATCGTCTCCGGCGGGGCCTCCTCGGCGATCTCGCCCTGATGGACATAGATGACGCGATCGGCCACTTCGCGGGCGAAGCCCATCTCGTGGGTGACAAT
GATCATGGTCATGCCTTCCCGGGCCAGCTCGCGCATGGCGTCGAGCACTTCGCCGATCATCTCCGGATCCAGCGCGGAGGTAGGCTCGTCGAACAGCATC
AGCCGCGGCTCCATGGCCAGGGCGCGGGCCAGGGCGACGCGCTGCTGCTGGCCACCGGAGAGCTGGGCCGGGTACTTGTCGGCCTGGTCGGAAATGCCGA
CGCGCTCCAGCAGGCGTTCGGCGGTCTCCATTGCATCCTCCTGGCTCCAGCCACGTACCTTCATGGGGGCCAGGGTGACGTTGTCGCGCACGCTCAGGTG
GGGGAAGAGGTTGAACTGCTGGAACACCATGCCCACCTCGGTGCGGATCTTCTGCAGGGCGCGGCCGCTCTTGCCGTGGGGGGTGAGTTCATTGCCGTCC
ACCTCGATGTGCCCGGACTGGAACTCCTCCAGGCCGTTGATGCAGCGGATCAGGGTCGACTTGCCCGAGCCGCTGGCGCCGATGACCACCACCACCTCGC
CGGGCATCACCTCGAGGTCGATATCCTTGAGTACGTGCAGCTTGCCGAAGTGCTTGTTGACCTTCTCCATGCGGACGATGGCCGTGGTCGGGTTCTCTTT
CTGTGTCATGTGCTTGGCCTCCCTCGCTCTTTTTTTAACCCCATCACTGGCCGACCAGGCCACGCCGCTCGAGCTGGCGCAGCAGGATCGACAGGGTCCA
GGTGATGGCCAGGTAGAGCAGTGCCACCATCAGGTAGACTTCCATGGCGGTGAAGGTGGTGGCGATATAGATCTGCCCCTGGCGTACCAGCTCGCCCACA

>Halomonas alimentaria DSM 15356 [NZ_WUTT01000001.1:2032575..2032680], dnaA> ig dnaN>, 106 bp

WebSkew GCmin pos. 51788 (GCmin<>oriC1 44.7 kb; GCmin<>oriC2 26.5 kb; oriC2<>oriC1 18.2 kb)

>Halomonas shengliensis CGMCC 1.6444 [NZ_FNIV01000006.1:c6786..7285], <rpmH oriC1 dnaA>, 580 bp

TTCTCGGGAGCCTGGGGTGCGGACTGGCTGGCGGAGAAGGAGGGAGCCTGGGGCTCGACGGCGGTCTGGCCATAGTCCTGATTCCACTGCACGACGATCA
GGTAGGCGAGCACCGCCAGGGGAATCAGCAATAGGAGTCGTTTTACGTCCATGAGGGTTCTGCCCGATGGGTAGTGAACATGCCGGTGGCGGAAGGCGTT
TCATCCGCCCGAGATAGACAAAAGCCTGCCGGGGGCAGGCTGACTTCCGAACACGGACTTTCGAGGCTCGCCCCGACGACGTCAAGCGTGACCTGGCGAA
CCTTGGCACTGCGAAGGGGGGCGCTCATGCTCTCGCGGGCTCCGGTCGGAGGTTGGCGGGGCATCGCCGTTCTCGGCGCGACGGGAGGCAGCCTTGTGTA
CCTCGCGTTCCAGGCGTCGCCACATCCCGACGAGTTGCCGCTGCAGGGTGGCATTGTCCAGCTCGCCGATACCTCGGCGCGACAGCAGCACGATATCCAC
AGCGGGAAGCCGCTGCTGCTGGAGCCTGACCGACTCGCGCGCCAGTCGCTTGATGCGGTTGCGGTCGACGGCGCGGCGCAGGCTCTTCTTGCTGACCACC
AGCCCCAGGCGAGGATGTCCCAGGCCATTGGGTGCCGCCAGTGCCATCAGGCCCTTGCCGTGGACCTTGAAGGCGGCATTGTCGAAGACCTGCCGGTAAT
CCCCGGCCGTCAGCAGGCGCAGACGGCGGGGAAACGACTGACGGGACACGCAGGGTCCGTGATCAGGCGCTCAGGCGCTTGCGGCCCTTGGCACGACGAC
GAGCCAGCACGGCGCGGCCGTTCTTGGTGGCCATCCGGGCACGGAAACCGTGTGTACGCTTGCGCTTGAGGACGCTGGGCTGAAAGGTACGTTTCATAAC
TCGTTATTCCCACGTGGTTTTCTGGACGAATCGTCGATTGCGAGGTGCCCGGAATCCACGGGGGAGTCCGAGCCGGTTCTGTTCAAGACCGGAAATTGTA

GTGATTTCCCGCCGGCGGTGCAATTTTTCCCTGTCTCGACCGCATCGCCAATCGCCGCCTTTGGGTATTGCCAGGGATTCGGGGGAGATGACCGGTCGGG
ACCAGGGGATCACCCGCCTGTAGTGTCCCGTCTGGGTCTGTCTTCCTACTATTACTAGGGTTATTTGTTACACGCAGTGATTACTGGGGAGAAGCTTATC
**TGTGGATAA**CGCCCATATCTCTCACCATCTCAGTCGCCTGGCCTTGGTACCGATAGGTGGGCAAGCTCGGGATGAGCGGTGGCCCGAGGTGGGATAACCG
GTGGATAACCCCTGCTTTCGTCCACAGCCGGGGAAGGCTCCCCATTCGTTCCCTAGGCTGTCGCCAGCCAGTCGGCCTGTTTTCCACAGCCTGGTGCAGG
GGTTGGCGGGGTGTGGATAACAGTGGCGCTGGGCGATACAATGCACGGTCCTTTCTCCACAACCATGGTGAGGTCGCGTGTCGCTGGCTCTCTGGCAACA

GTGTCTGGATACCCTGCAGGACGAGTTGAGCTCCCAGCAGTTCAATACCTGGGTTCGCCCCCTCCAGGCCGAGGAGGGCGAGGGGGGTGAGCTGTGCCTG
CTGGCGCCCAACCGCTTCGTGCGCGACTGGGTCAGCGACAAGTACGCCAAGCGAATCGCCGAGCTGCTGCGCGAGCTGTCGCCGGCCCGTCCTCCCAAGG
TGGTGCTGAGCGTCGGCAGTCGTCGCAGCGCACCGCGCCCGCAGCCGCGGGAGCTGGGCAACCCGGTATCGGCCGCGCCACGGCGTCCGGCGGCGCCGGC
GGTACACACGCCGCCACGGGGTGACGTGGCCGACGAGAAGGAGATCACTCGCCAGGAGGAGGACGCGCCCCGGCGCCCTGGCGGCGAGCGCCAGGTCCAG
GTGGAGGGGAGCCTCAAGCACAACAGCGGCCTCAACCCCAACTTCACCTTCGAGACCTTCGTGGAGGGCAAGTCCAACCAGCTGGCCCGGGCCGCCTCGC
GGCAGGTGTCGGAGAATCCCGGCGGCGCCTACAACCCGCTGTTCCTCTACGGTGGGGTCGGCCTGGGCAAGACCCACCTGATGCACGCGGTGGGCAATGC
CCTGGCCGGACGGCGCGAGAACGCCCGGGTGGTCTACCTGCACTCCGAGCGCTTCGTGGCCGACATGGTTAAGGCGCTGCAGCTCAACGCTATCAACGAC
TTCAAGCGCTTCTACCGTAGCGTCGATGCCCTGCTGATCGACGATATCCAGTTCTTCGCCGGCAAGGAGCGCTCCCAGGAGGAGTTCTTCCACACCTTCA
ATGCCCTGCTCGAGGGCGGCCAGCAGATGATCCTGACCTCCGACCGCTATCCCAAGGAGATCAGCGGCGTAGAGGAGCGACTCAAGTCGCGCTTCGGCTG
GGGCCTGACGGTGGCCATCGAGCCGCCCGAGCTGGAGACCCGTGTGGCGATCCTGATGAAGAAGGCCGACCAGGCCAAGGTCGACCTGCCCCACGACGCC

>Halomonas shengliensis CGMCC 1.6444 [NZ_FNIV01000006.1:c25063..25562], <mnmG **oriC2** <COG1126 (glnQ), 532 bp

TTCTCGAGCCCCTGCATGGAGCGCACCACCTGCAGCTGCACGTCGAAGGGCAGCGAGGTGGAGATGCCGTTGGGATAGAGCTCGTGGGTGTCGAGGCCCT
CCGGCTCGATGAAGACCTGGTGGCTCGTCTTGTCGGCGAAGCGATGCACCTTGTCCTCGATGGAGGGGCAGTAGCGCGGGCCGACCCCCTCGATGGTTCC
CGAGTACATCGGCGACCGGTCGAGGTTGGCGTGGATGATCTCGTGGGTGCGCTCGTTGGTGTGGGCGATATGGCAGCTGACCTGGCGCGGGTGCTGGTCG
CGGCGTCCCATGAACGACATCACCGGGGTGGGGGTGTCGCCGGGTTGCTCCTCGAGCCGCGAGAAGTCCAGGCTCCTGGCATCCAGGCGCGGCGGGGTGC
CGGTCTTGAGGCGGTCGACCCGGAACGGCAGGGCGCGCAGGCGCTCGGCCAAGCGGTTGGAGGGCGGGTCGCCGGCGCGCCCGCCGCGGCTGCTGTCCAG
GCCGATATGGATCACCCCGCCGAGGAAGGTGCCGGTGCACAGCACCACCGTCTCGCCGAGGAAGCGGATACCGGTCTCGGTGACCACGCCACGCACGGTG
TCCTGCTCCACGATCAGGTCGCCGGCCGCCTGCTGGAAGAGCGTCAGGTTGGGCTGGTTCTCCAGCATGCCGCGGATGGCCGCCTTGTAGCGGATACGGT
CGGCCTGGGCGCGGGTGGCGCGCACCGCCGGCCCCTTGCGGGCGTTGAGCACCCGGAACTGGATTCCCCCGAGGTCGGTGGCCAGGCCCATGGCCCCGCC
CAGGGCATCGATCTCCTTGACCAGGTGACTCTTGCCGATGCCACCGATGGCCGGGTTGCAGGACATCTGGCCAGGGTCTCGATGTTGTGGGTCAGCAGCA
GGGTCCGGGAGCCCATGCGCGCGGCGGCCAGGGCGGCTTCGGTTCCCGCATGGCCGCCGCCGATGACGATGACGTCGAAGCGGTCGGGGTAATCCAAGGT

GCACCTCGTTGGCGCCGCGGCGCCTCTCTTCGGTCGGGGGAATGGAATCAGCCCGCCAGTATAAACCTCCTGTGGGTAATCGTCAGGGTTTTCCACACCG
TGAACCCAGCCGTTCGCTTCCGGGTTCTCCTAGTAATCAATATCCAATATATAAAGAGAGAAGTTCTGTTGTGGTTGTGACTACTAGTGTGGACAAAAAC
**CGTGGATAA**GCGCCCTTATTCATTGTTAACCAAAGGCTTGCGTTGTTAACCCTTCTTTGGGAAAACGGCTGACTGCCTGCGCACGCCCGGTCGAGAGCCT
GTGGATGGAGTGCCCGCTTGTCCACAGCGTCGTCGCTGCACCGCTTGTCCACCGCCGGCTACCGGGCTTGTGCCCGCGGCTGTGAACAACCGGGCCGGTT
ATCCACGAGGCCCGCGGTGGCAGCCTCTCCGGCGGTTCATGGCGGTCTCGGCCTCAATCTTTTCCACAGGCGAAAAAAGGGCCCTGTCGCCGCGCGACAG

GGCCCTTCTCGGCGCCGATGAGGCTGTTCTCAGTGCTTGAGCACCCGCGACAGGAAGCTGCGTGTACGTTCGTCGGCGGGGCTGTCGAAGATCGTCTCGG
GGGGCGCCTCCTCGGCGATCTCGCCCTTGTGGACGTAGATGACCCGGTCGGCCACCTCGCGGGCGAAGCCCATCTCGTGGGTGACGATGATCATGGTCAT
GCCCTCCCGTGCCAGCTCGCGCATGGCGTCGAGCACCTCGCCGATCATCTCCGGGTCGAGCGCCGAGGTGGGCTCGTCGAACAGCATCAGGCGCGGCTCC
ATGGCCAGGGCGCGGGCGAGGGCCACGCGCTGCTGCTGGCCCCCGGAGAGCTGGGTGGGGTACTTGTCGGCCTGGTCGGCGATGCCGACCCGCTCCAGCA
GGCGTTCCGCGGTCTCCACCGCGTCCTCGTTGCTCCAGCCGCGCACCTTCATCGGCGCGAGGGTGACGTTGTCACGCACGCTCAAGTGGGGGAAGAGGTT
GAACTGCTGGAACACCATGCCCACCTCGGTGCGGATCTTCTGCAGGGCCTTGCCGCTCTTGCCGTGGGGGGTGAGTTCGTTGCCGTCCACCTCGATGTGT
CCGGACTGGAACTCCTCCAGGCCGTTGATGCAGCGGATCAGGGTCGACTTGCCCGAGCCGCTGGCGCCGATGATCACCACCACCTCCCCCGGCATCACCT
CGAGGTCGATCTCCTTGAGCACGTGCAGGCTGCCGAAGTGCTTGTTGACCTTTTCCATCCGCACGATGGGAGCGGTCGCGTTCTCGTTCTGTGTCATGCG
TGAGCTCCCCTTATTGTCCGACCAGGCCGCGGCGTTCGAGCTGGCGCAGCACGATCGAGAGGGTCCAGGTGATGGCGAGGTAGAGCACCGCCACCATCAG
GTAGACCTCCATGGCGGTGAAGGTGGTGGCGATGTAGATCTGTCCCTGGCGCACCAGCTCACCGACGCCGATCACCGAGAACAGCGAGGTATCCTTGATG

>Halomonas shengliensis CGMCC 1.6444 [NZ_FNIV01000006.1:c5242..5344], dnaA> ig dnaN>, 103 bp

WebSkew GCmin pos. 249480 (GCmin<>oriC1 28.4 kb; GCmin<>oriC2 9.4 kb; oriC2<>oriC1 19.0 kb)

>Halomonas ramblicola CECT 7896 [NZ_JAUFPL010000032.1:277653..278152], <rpmH oriC1 dnaA>, 521 bp

GTCGTCGCCCTGGGCACCGGCCGTGGGTACCGAGAGGCCATCGTCGTCGTCGGCGGAGGGCTGCGTTGCCCCGTCCGAACTGCTGGTGATGGAGGGAGCC
TCCGCTTGCGGGGTCGTCTGGCCATAGTCCTGATTCCACTGCACGACGATCAGGTAGGCGAGCACCGCCAGTGGGATCAGCAATAGGAGTCGTTTTACGT
CCATGAGGGTTCTGCCCGATGGGTGGTGAACATGCCAAGGGTGGAAGGGGAATCATCCGCCCGAGATAGACAAAAGCCTGCCGACGGCAGGCTGTGTTCC
GGACACGGACTTTCGACGTCGGCCCCGGCGGCGTCAAGCATCGCCTTGAGGAGAGGTGCGGACATGGTGTCGCGGGCTCCTGGCTGGCCGGCGGTCCTTG
GCGTCGGGCGCGTCGGCGGTCCCCGGCGTGGCGATGGGCTGGCCGTCTCGCTTGCGCGCCTCGCGCGCCAGGCGCTTCCACATGCCGTGCAGCTGGCGAT
GCAGGACGGCGTTGTCCAGTTCCCCTACGCCGCGTCGGGCGAGGACCACGATATCGACGTCCGGAAGCTGCTCCTGGCGCAGCCGCACCGATTCCCGAAC
GAGTCGCTTGAGACGATTGCGATCGACGGCACGGCGCAGGCTCTTCTTGCTGACGACCAGGCCGAGGCGCGGATGCCCGAGCCGGTTGGGTACCGCCAGT
GCCAGCAGCCCCTTGCCATGGACCTTGAAGACGGCGGTCTCGAAGACCTGTCGGTAATCCCCGGCCGTCAGCAGGCGCAGACGGCGGGGAACGCCTGTC
GGGACACGCAGGGTCCGTGATCAGGCGCTCAGGCGCTTGCGTCCCTTGGCGCGACGACGCGCCAGGATGGCACGACCGCTCTTGGTGGCCATACGGGCAC
GGAAGCCGTGCGTGCGCTTGCGCTTGAGAACGCTGGGCTGGAAGGTGCGTTTCATAACTCGTTATTCCCACGTGGTTTTTCAGGACAAATCGACGATTTC

TCGGCGCCCGGGCTCCACAGGGAAGCCGGGCGTATTCGGTTCAAGACCGGAAATTGTAGTGATTTCGCGGGCCCGGTGCAATTTTTCCTTGCGCCGGAAC
GAGGCGGCGGGATTCACCAGGGACGGCCACAGGCGTGACGATATAACAACAGAATAATGTTCTTCTTGAACCCGTTGTGATTAATGGTGGGGATGCTTTC
**TGTGGAAAA**TTCACAATAAGTCTTCCATATCAATCACAACTGCACATGTTAATACTGTGGACAAGCCTTGTGTCGAGGCGGGATAAAGGCTGCATGGGCG
GTGGATGGCTACCCGACCCATCCACAGGAGCGAACCATGCGGCTCGGTGTCCCCTGACCATCCCCCGGCTTTCGCTATTTTTTCCACAGCCTGCAGGTGG
CGCGCCCGGGGTGTGGATAACCATCCTGCGGGGCGATACAATGCCCGGTCGTTCCACGAACAGATGGTGAGGTCGCGTGTCGCTGGCTCTCTGGCAACAG

TGTCTGGATACCCTGCAGGACGAGCTGAGCTCGCAGCAGTTCAATACCTGGATCCGTCCACTCCAGGCGGAGGAGGGGGATGCCGGCGAACTGTGCCTGC
TGGCACCCAATCGCTTCGTCCGGGACTGGGTCAGTGACAAGTACGCCAAGCGCATCGGCGAGCTGATGCGCGAGCTGTCGCCGGCCAGGCCACCCAAGGT
GGTGCTCTCGGTCGGCAGCCGCCGTGCCGCCCCCACGCCTCAGCCCCGCGAGCTGGGCCACCCGGTCTCCGCCGGGCATCGCCGGCCGATGGCGCCGGCG
GTCCATGCCCCACCGCGCGGCGACGTCGCCGACGAGGGGGAGATCGAGTCGTCGGGTGAGGAAGTCGCGGCCCCCCGTCGTCCCGCCGGAGAGCGTCAGG
TCCAGGTCGAGGGCAGCCTCAAGCACAACAGCGGCCTCAATCCCAACTTCACCTTCGAGACCTTCGTCGAGGGCAAGTCCAACCAGTTGGCCCGGGCGGC
CTCGCGGCAGGTATCCGAGAATCCCGGCGGTGCCTACAACCCGCTGTTCCTGTATGGCGGCGTGGGCCTCGGCAAGACCCACCTGATGCATGCTGTGGGC
AACGCCCTGGCCGGGCGCCGCGAGAATGCTCGGGTGGTCTACCTGCACTCGGAGCGCTTCGTGGCCGACATGGTCAAGGCACTCCAGCTCAATGCCATCA
ACGACTTCAAGCGCTTCTATCGCAGCGTCGATGCACTGCTGATCGACGATATCCAGTTCTTCGCCGGCAAGGAGCGCTCCCAGGAGGAGTTTTTCCACAC
CTTCAACGCCCTGCTCGAGGGCGGCCAGCAGATGATCCTCACTTCGGATCGCTACCCCAAGGAGATCAGCGGTGTGGAGGAGCGCCTCAAGTCGCGTTTC
GGCTGGGGGCTGACGGTGGCCATCGAACCGCCGGAACTGGAGACCCGGGTGGCGATCCTGATGAAGAAGGCCGATCAGGCGAAGGTGGACTTGCCCCACG

>Halomonas ramblicola CECT 7896 [NZ_JAUFPL010000032.1:258613..259112], <mnmG oriC2 <COG1126 (glnQ), 535 bp

CTCCAGGCCGGCGATGGAGCGTACCACCTCGATCTGGACGTCGAAGGGCAGCGAGGTGGAGATGCCGTTGGGGTAGAGCTCATGGGTGTCCAGACCCTCC
GGCTCGATGAACACCTGGTGGCTCGCCTTGTCGGCGAAGCGGTGCACCTTGTCCTCGATCGACGGGCAGTAGCGCGGTCCGACCCCCTCGATCGTCCCGG
AGTACATGGGCGAGCGGTCGAGGTTGGCGAAGATGATCTCGTGGGTGCGCGCATTGGTGTGGGCGATATGGCAGCTCACCTGGTGCGGATGCCGCTCACG
ATCGCCCAGGAAGGACATCACCGGTGTCGGGGTATCGCCCGGCTGCTCCTCGAGCTTGGCGAAGTCGAGGCTCCTGGCATCCAGCCGCGGCGGCGTCCCG
GTCTTCAACCGGTCGACCCGGAACGGCAGGGCGCGCAGCCGCTCGGCCAGGGCGTTGGAGGGCGGGTCGCCGGCACGGCCGCCGCGGCTGCGGTCGAGGC
CGATATGGATGACCCCGCCGAGGAAGGTGCCGGTGCACAGCACCACGCTCTCGGCGTGGAAGCGGATACCGGTCTCGGTGACCACGCCGCGCACGGTGTC
GCCGTCGACGATCAGGTCGCCGGCGGCCTGCTGGAAGAGCGTCAGGTTGGGCTGGTTCTCCAGCAGGCCGCGGATCGCCGCCTTGTAGCGGACCCGGTCG
GCCTGGGCCCGGGTCGCGCGTACCGCGGGGCCCTTGCGGGCGTTGAGGACACGGAACTGGATACCGCCGAGGTCGGTGGCCCTGGCCATGGCACCGCCCA
GGGCATCGATCTCCTTGACCAGGTGGCTCTTGCCGATCCCGCCGATGGCGGGATTGCAGGACATCTGGCCGAGGGTCTCGATGTTGTGGGTCAGCAGCAG
GGTCTGACAGCCCATGCGGGCGGAGGCCAGGGCGGCTTCGGTTCCCGCGTGGCCACCGCCGATGACGATGACGTCGAAGCGGTCGGGGTAATCCAAGGTG

CACCTCGATGGCGCCGTGGCGCCTGTCTTGCGGTCAAGGGGGATGAAATCAGCCCGCCAGTATAAACCCCCTGTGGGTAATCGTCAGGGTTTTCCACACC
GGGATCGCCATCTTCGCGCCTCTCCATCTCCTATCAATAAAAAACAGAGTATTAGAATAGAAGTTCTGTTGTGGTAGTAATTACTGGTGTGGACAAAACC
**CGGGGATAA**GTGGAAAAGTCTCTTTCCTATCAACAGGATCCGCTGTTAACCAAAACCGGGACAAGGGGCTGACGACCTGCGGACGAGGTGTCGCGAACCT
GTGGACGGAACCGCGACTTGCCCACAGCCCCGGGACTGCACCGCTTTTCCACCGCCGGCTACCGGGCTTGTGCCCGTGACTGTGAACAACCCTCGCGATA
CGTCGCCAGGCCCCTGACCATGGGCCTCGGACGGCGAATGGGGAGTCCGAGGCGAATTTTCTGTCCACAGGCAGAAAAAGGGCCCTGTCGATAACGACAG

GGCCCGTCCATGCGCGGCGACGCACGGCGATTCAGTGCTGGAGCACCCGCGACAGGAAGCTGCGAGTCCGCTCGTCGGCGGGAGCGTCGAAGATCGTCTC
CGGTGGCGCCTCCTCGGCGATCTCACCCTTGTGGACGTAGATCACCCGGTCGGCCACCTCGCGGGCGAAGCCCATCTCGTGGGAGACGATGATCATGGTC
ATGCCCTCCCTGGCCAGCTCGCGCATGGCGTCGAGCACCTCGCCGATCATCTCCGGATCCAGCGCCGAGGTGGGCTCGTCGAAGAGCATCAGGCGTGGCT
CCATGGCCAGCGCCCGGGCCAGGGCCACACGCTGCTGCTGGCCGCCGGAGAGCTGGGTCGGTGACTTGGCGGCCTGGTCGGCGATGCCGACGCGCTCGAG
CAGCCGCTCGGCGATCTCGACGGCGTCCTCGCGGCTCCAGCCGCGTACCTTCATGGGCGCCAGGATGACGTTGTCCAGCACGCTCAGGTGGGGAAAGAGG
TTGAACTGCTGGAACACCATGCCCACCTCGGTGCGGATCTTCTGCAGCGCCTTGCCGCTCTTGCCATGGGGCAGCAGCTCGTTGCCGTCCACCACGATAT
GGCCGTCCTGAAACTCCTCCAGGCCGTTGATGCAACGGATCAGCGTCGACTTGCCCGAGCCGCTGGCGCCGATGATCACCACCACCTCGCCGGGATTGAC
CTCCAGGTCGATGTCCTTGAGGACGTGCAGGCTGCCGAAGTGCTTGTTGACCTTCTCCATGTTCACGATGGGCTCGTTGCCGCCCGGCGACTGTTCTGAG
TTCATGTCGCGCTCCCTTATTGTCCGACCAGGCCGCGGCGCTCGAGCTGGCTGAGCAGCAGCGACAGGGTCCAGGTGATCGCCAGGTAGAGCAGGGCCAC
CATCAGGTAGACCTCCATGGCGGTGAAGGTGGTGGCGATGTAGATCTGGCCCTGGCGGACCAGCTCGCCGACCCCGATCACCGAGAACAGCGAGGTGTCC

>Halomonas ramblicola CECT 7896 [NZ_JAUFPL010000032.1:279599..279709], dnaA> ig dnaN>, 111 bp

WebSkew GCmin pos. 3989759/0 (GCmin<>oriC1 0.5 kb; GCmin<>oriC2 19.6 kb; oriC2<>oriC1 19.1 kb)

>Halomonas alkalicola M2 [NZ_CP130143.1:3989049..3989548], <rpmH oriC1 dnaA>, 545 bp

```
GCGGGCACGGCCAGGCCATCCGGGTCGGCCTGCTGGGCCTCGCCGTTGGCCGGGCTGCTGGTGAAGGAGGGGGCCTCGGGCTCAGGGGAGACCTGGCCAT
AGTCCTGATTCCACTGCACGACGATCAGGTAGGCGAGCACCGCCAGGGGGATCAGCAAGAGGAGTCGTTTTACGTCCATGAGGTTTCTGCCCGATGGGTG
GTGAACTTGCCGCGGGCGAGATGGAAAACATCTGCCCGAGATAGACAAAAGCCTGCCAGAGGCAGGCTGGATGCCGAACACGGACTCTCGGCGGGGATGC
CGGCAGGGTCGGGGAACACCATGGAAAGGGGCCTCGACATGGTGACGACTCCTGGTTGGCCGGTCATCCGCCCTGACTGGCGTTCGCGCTGGGTGTCCCG
GTACCTGGCTGGTCGTCGACGGCGGCCTTGCTGGCCTCGCGCTCCAGACGTCGCCACATCCCATGCAGCTGGCGATGCAGGACCTCGTTATCGAGCTCCT
TCACGCCTCGCCTGGCCATCAGCACGATGTCCACGGCGGGCAGACGCTGCTGCTGAAGGCGTATGGATTCGCGGGCGAGTCGCTTGAGGCGGTTGCGGTC
GACGGCACGGCGAAGGTTCTTCTTGCTGACGATCAGGCCAAGGCGTGGATGCCCAAGGTCGTTGGGAATCGCCAGGGCCATGAGGCCCTTGCCATGAACC
TTGTAGACCGCGTTGTCGAAGACCTGCCGATAGTCCCCGGCCGTCAGCAGACGCAGACGGCGGGGGAAGTCCTGACGGGACACCTGGGGTCCGTGATCAG
GCGCTCAGGCGCTTGCGGCCCTTGGCACGACGGCGGGCCAGGACGGCACGGCCGTTCTTGGTGGCCATACGGGCACGGAAACCGTGCACGCGCTTACGCT
TCAGGACGCTGGGCTGAAAGGTGCGTTTCATAACTCGTTATTCCCACGTGGTTTTCAGGAAGAATCGACAATTTCGAGGCGTCCGGTTTCCACAGAGGAC
```

```
CCCGGGGGTATTCGGCTCAAGACCGGAAATTGTAGTGATATCGCGGCCCGGGTGCAATTTTTCCGTACGCGATCGCCAGTGCGATGACGGCACAGGCGCC
TCACGAGCCCTGCCGTCGGGTCTGCCATCGACCACTACAACTACTATTACTATTTTCTATCCCTTTTAAAGAACCTGTTATTAATAGTGGGATAGTATC
TGTGGATAAGATGGCTTTCCCTAACGTAAACAGTCTTCTACCGCAGGGCCTAGCCTGTGGACAGGCAGTGGGGTGGCACGGGATGTCCTGTGCAGAAGCG
GTGGATGCAATCGCCAGTCGTCCACAGTGAGATATTGTCCCCAGCGTTCGGCGGGACCCATCCCCGGGGTCCGGGACAAGTTTTCCACAGCCGCGGCGCG
GCGCCCGCGGGGTGTGGATAACCATCCTCGGGGGCGATACAATATGCGACCGTTTTCCAACCGATGGTGAGGCCGCGTGGCGCTAGCTCTCTGGCAACAG
```

```
TGTCTGGACACCCTTCAGGACGAGCTGAACTCCCAGCAGTTCAACACCTGGATTCGCCCGCTGCAGGCGGAAGAGGGGGAGTCGGGTGAGCTGTGTCTGC
TGGCCCCCAACCGCTTCGTGCGCGACTGGGTCAGCGACAAGTATGCCAAGCGCATCGGTGAGCTGATGCGCGAGCTCTCCCCGGCCAAGCCTCCCAAGGT
GGTGCTGACGGTAGGCAGTCGGCGCACCGCGCCCCAGCCCCAGCCGCGCGAACTGGGCAACCCGGTATCGGCCGCGCCACGACGCCCGGCGGCACCCGCC
GTGCATACTCCCCCGCGTGGTGAGGTGGCCGATGAGCGTGAGCTCGACGCCCAGCGCGAGGAGGGCGCGGGCCGCCGGGCGGCCGGGGAGCGCCAGGTGC
AGGTCGAGGGCAGCCTCAAGCACAGCAGTGGCCTGAACCCCAACTTCACCTTCGAGACCTTCGTGGAGGGCAAGTCCAACCAGCTGGCCCGGGCCGCCTC
GCGGCAGGTCTCGGAGAACCCGGGAGGGGCCTACAATCCGCTGTTCCTCTACGGCGGGGTCGGCCTCGGCAAGACCCACCTGATGCACGCCGTTGGCAAT
GCCCTGGCCGGGCGGCGGGAGAATGCCCGGGTGGTCTACCTGCACTCCGAGCGTTTCGTGGCCGACATGGTCAAGGCGTTGCAGCTCAACGCCATCAACG
ACTTCAAGCGCTTCTACCGCAGCGTGGACGCGCTGCTGATCGATGACATCCAGTTCTTCGCCGGCAAGGAGCGCTCCCAGGAGGAGTTCTTCCACACCTT
CAATGCCCTGCTCGAAGGCGGCCAGCAGATGATCCTCACCTCCGACCGCTATCCCAAGGAGATCAGCGGGGTGGAGGAGCGCCTGAAGTCGCGCTTCGGC
TGGGGGCTGACGGTGGCCATCGAGCCGCCGGAGCTGGAGACCCGGGTGGCGATCCTGATGAAGAAGGCCGACCAGGCCAAGGTGGATCTGCCCCACGATG
```

>Halomonas alkalicola M2 [NZ_CP130143.1:3969919..3970418], <mnmG oriC2 <COG1126 (glnQ), 530 bp

```
TTCTCCAGCCCCCTGATCGAGCGCACCACCTCGAGCTGCACGTCGAAGGGTAGCGAGGTGGAGATGCCGTTGGGGTAGAGCTCGTGGGTATCCAGCCCCT
CCGGCTCGATAAAGACCTGGTGGCTCGCCTTGTCGGCGAAGCGGTGCACCTTGTCCTCGATGGAGGGGCAGTAGCGCGGCCCGACGCCCTCGATGACGCC
GGAGTACATCGGCGAGCGGTCGAGGTTGGCGAAGATGATCTCGTGGGTGCGCGCATTGGTGTGGGCGATATGGCAGCTCACCTGGCGCGGGTGCTGAGCC
CGACTGCCCAGGAAGGACATCACCGGGGTGGGGGTGTCGCCGGGCTGCTCCTCGAGCTGCGAGAAGTCGAGGCTGCGTGCATCCAGCCTGGGCGGGGTGC
CGGTCTTGAGGCGGTCGACCCGGAACGGCAGGGCGCGCAGGCGTTCGGCGAGACGATTGGAGGGCGGGTCGCCGGCGCGGCCGCCGCGGCTCTGCTCGAG
GCCGATATGGATCACCCCGCCGAGGAAGGTGCCGGTACACAGCACCACGGTCTCGCCGAGGAAGCGGATCCCGGTCTCGGTGACCACGCCGCGGACAGTG
TCGCCCTCGACGATCAGGTCGCCGGCGGCCTGCTGGAAGAGCGTCAGGTTGGGCTGGTTCTCCAGCATGCTGCGGATCGCCGCCTTGTAGCGCACCCGAT
CCGCCTGGGCCCGGGTGGCCCGTACCGCGGACCCTTGCGGGCGTTGAGCACGCGGAACTGGATACCGCCGAGATCGGTGGCCATGGCCATGGCGCCACC
CAGTGCATCGATCTCCTTGACCAGATGGCTCTTGCCGATCCCGCCGATGGCGGGATTGCAGGACATCTGCCCGAGGGTCTCGATGTTGTGGGTCAGCAGC
AGGGTCTGACAGCCCATGCGGGCGGAGGCCAGGGCGGCTTCGGTTCCAGCATGGCCTCCGCCGATGACGATGACGTCAAAGCGGTCGGGGTAATCCAAGG
```

```
TGCACCTCGTTGGCGCCGCGGCGCCGTCTTCCGGTCGGGGGATGAAATCAGCCCGGCAGTATAAACCCCCTGTGGGTAATCGTCAGGGTTTTCCACACCG
CCTTCGCCCTTCCCTGTGGTCTCCCTCTCCCTAACATTAAATATCAATATATAAAGAGAGAAGTTCTGCTGTGGTAGTAACTAACTGGTGTGGACAAAAAC
CGTGGATAACCCACAAGAACCTTTTGATATCAATGCTCTGTATTGTTGACCAAAGGCGGGACAAGAGGGTGACTGCCTGCGCACATTCCGCCCTGAGCCT
GTGGATGGAATGGCGACTTGCCCACAGCGGGACTCCTGCACGGCTTGTCCACCGCCGGGTACCTGTCTTGTGCCCGCGGCTGTGAACAACCGGCCTTCTT
ATCCACGACCCCCTCTGGCATGGGCCCTGCAGCGCAGATCGGCCTCGCCGTAGAAAATTCTATCCACAGGCGAAAAAAAGCCCCTGCCGATGCCGACAGG
```

```
GGCTTTTCGGTCGCTCCGAGGGGCGATGTCAGTGCTTGAGGACCCGGGCCAGGAAGCTCCGGGTGCGCTCGTCGGCGGGGCGATCGAAGATCTGCTCCGG
CGGTGCCTGCTCGGCGATCTCTCCCTTGTGGATGTAGATCACCCGGTCGGCGACCTCCCGGGCGAAGCCCATCTCGTGGGTGACGATGATCATGGTCATG
CCTTCTCTCGCCAGCTCGCGCATGGCATCGAGCACCTCGCCGATCATCTCCGGGTCCAGCGCCGAGGTGGGCTCGTCGAAGAGCATCAGGCGCGGCTCCA
TGGCCAGGGCGCGGGCCAGGGCCACGCGCTGCTGCTGGCCGCCGGAGAGCTGGGTGGGGTGCTTGGCGGCCTGGTCGCTGATGCCGACCCGGGCCAGCAG
CCGCTCGGCGGTCTCCACGGCGTCGGCGCGGCTCCAGCCGCGCACCTTCATGGGGGCGAGGATGACGTTGTCCAGCACGCTGAGGTGGGGAAAGAGGTTG
AACTGCTGGAACACCATGCCGACCTCGGTGCGGATCCGCTGCAGCGCACGGCCGCTCTTGCCGTGGGGAGTCAGCTCGTTGCCGTCCACCTCGATGTGGC
CCGCCTGGAACTCCTCCAGGCCGTTGATGCAGCGGATCAGCGTGGATTTCCCCGAGCCGCTGGCGCCGATGATCACGACGACCTCCCCCGGGGTCACCTC
GAGGTCGATCCCCTTGAGCACGTGAAGGTCGCCGAAGTGCTTGTTGACCTTCTCCATGCGCACGATGGGAGCGGTCGCCTTGTCGTTCTGTGTCATGGGT
GGGCTCCCTCTATTGTCCGACCAGGCCGCGCTTTTCCAGCTGGCGCAGCAGCAGCGACAGGCTCCAGGTGATGGCCAGATACATCAGGGCCACCATGAAG
TAGACCTCCAGGGCGGTGAAGGTCGTGGCGATGTAGATCTGCCCCTGGCGGACCAGCTCGCCCACGCCGATCACCGAGAACAGCGAGGTGTCCTTGATGC
```

>Halomonas alkalicola M2 [NZ_CP130143.1:1233..1413], dnaA> ig dnaN>, 181 bp

WebSkew GCmin pos. 100000 (2nd min.) (GCmin<>riC1 26.6 kb; GCmin<>oriC2 7.7 kb; oriC2<>oriC1 19.0 kb)

>Halomonas mongoliensis DSM 17332 [NZ_JARWAL010000011.1:c73128..73627], <rpmH oriC1 dnaA>, 556 bp

CGGCAAGGCCATCCGGGTCGGCGGGCTGGACCTGGCCATTGTCTGAGCTGCTGGTAAAGGAGGGGGCCTCAGGCTCAGGAGCCACCTGGCCATAGTCCTG
ATTCCACTGCACGACGATCAGGTAAGCGAGCACCGCCAGGGGGATCAGCAATAGGAGTCGTTTTACGTCCATGAGGGTTCTGCCCGATGGGTGGTGAACT
TGCCAGGGGCGGGATGGAAAGTACCCGCCCGAGATAGACAAAAGCCTGCCAAAGGCAGGCTCGATGCCGAACACGGACTGTCGGCGGTGGAATCGGCCGG
GGCGAGTGTCGCCATGGCAAGCAGGTGCAGCATGGCAATACCGCCGGTCCGCTCATCCGCTCTGACTGGCGTTCGCACTGGTCGTCCCGTTCTGCGGCTG
ATCGTCACGCTTCCACCTGGCAGCATCTCGCGCCAGGCGTCGCCACATGCCGTGCAGCTGGCGATGCAGGGTGGCGTTATCCAACTCTCCTACGCCTCGC
CTGGCCAGGACCACGATGTCCAGGGCAGGGATCTGCTGTTGCTGGAGACGGATGGACTCACGAACGAGCCGCTTGAGGCGGTTACGGTCGACGGCACGGC
GAAGGTTCTTCTTGCTGACCACCAGGCCAAGACGCGGATGTCCCAGCTCATTGGGTATGGCCAGGGCCATGAGCCCCTTGCCGTGAACCTTGAACGCCGC
GGCCTCGAAGACCTGACGGTAGTCCCCGGCCGTCAGCAGGCGCAGACGGCGGGGAAATGCCTGACGGGACACCAGGAGTCCGTGATCAGGCGCTCAGGCG
CTTGCGACCCTTGGCGCGACGACGGGCCAGGACGGCACGGCCGTTCTTGGTGGCCATGCGAGCACGGAAGCCGTGCACACGCTTACGCTTCAGAACGCTG
GGCTGAAAGGTACGTTTCATAACTCATTATTCCCACGTGGTTTTTAGGACGATTGGACAGCTTGGAGGCGTCCGGATCCCGCTCGGGGCCCGGTGATATT

CGGCTCAAGACCGGAAATTGTAGTGATATCGCGCTCCGGGTGCAATTTTTCCGTTGCCGGATCCGGGCAACGGCCTCGGTGCCCCCGAGCCTCGCTGCCG
GGACAGCGGGAGACTCTGGCCTGACTTCTACCCTCAAGACGCAACAGTAACTATTTATAGTTCTTTTATAAAACCTGTTATTAATGCTGGGGACAGTTCC
**TGTGGATAA**CTTGCCTTTCCATTTCCAGGACAAGGGGATATACAGGTTGAAAGTGTGTGGACGAGCGCTGGAGGCAGCATGGACAGCCTGTGCAGTGAGG
GTGGACAAATGTTGCTCTGGTCCACAGGCTCCGGGTATCCCCAGAACTCAAGGGCAGCAGTCCCGAGGGCCCGCCCAGGGTTTTCCACAGCCCTGCGGCC
TGGAGGCCGGGGTGTGGATAACCGGCCTTGAGGGAGCTACAATACCCGCCCGTTTTCCAACCGATGGTGAGGTCGCGTGCCACTAGCTCTCTGGCAGCAG

TGCCTGGACACCCTTCAGGACGAACTGAACTCGCAGCAGTTCAATACCTGGATCCGCCCGCTGCAGGCGGAAGAGGGAGAGTCCGGCGAGCTGTGTCTGC
TGGCGCCCAATCGCTTCGTGCGGGACTGGGTCAGTGACAAGTACGCCAAGCGTATCGGCGAGCTGATGCGTGAGCTTTCGCCTGCCAAGCCTCCCAAGGT
GGTGCTGAGCGTGGGCAGTCGGCGCAGCGCGCCTCGGCCGCAGCCTCGCGAGCTGGGCAACCCCGTCTCCGCCGCTCCTCGCCGGCCGGCGGCGGCTGCC
GTGCATACCCCTTCTCGAGGCAGTGCTGCCGACGAGCGCGAGGTCGAGGCTCTGCGCGAGGAAGGGGAGGTGCGCCGACCCGGCGGGGAGCGGCAGGTCC
AGGTGGAGGGCAGTCTCAAGCACAGCAGCGGCCTCAACCCCAATTTCACTTTCGAGACCTTCGTGGAGGGCAAGTCCAACCAGCTGGCCCGGGCGGCCTC
GCGGCAGGTCTCCGAGAACCCCGGCGGCGCCTACAACCCGCTGTTCCTATATGGCGGGGTGGGCCTGGGCAAGACTCACCTGATGCATGCGGTGGGCAAC
GCCCTGGCCGGGCGCCGCGAGAACGCCCGGGTCGTCTACCTGCACTCCGAGCGTTTCGTGGCCGATATGGTCAAGGCGCTGCAGCTCAATGCCATCAACG
ACTTCAAGCGCTTCTACCGCAGCGTCGATGCCCTGCTGATCGATGACATCCAGTTCTTCGCGGGCAAGGAGCGCTCCCAGGAGGAGTTCTTCCATACCTT
CAACGCCCTGCTCGAGGGCGGGCAGCAGATGATCCTGACCTCCGATCGCTACCCCAAGGAGATCAGCGGCGTGGAGGAGCGTCTCAAGTCGCGCTTCGGC
TGGGGATTGACCGTGGCGATTGAGCCGCCGGAGCTGGAGACGCGGGTGGCGATCCTGATGAAGAAGGCGGACCAGGCCAAGGTGGACCTGCCCCACGATG

>Halomonas mongoliensis DSM 17332 [NZ_JARWAL010000011.1:c92096..92595], <mnmG oriC2 <COG1126 (glnQ), 532 bp

TTCTCCAGCCCCTTGATAGAGCGCACCACCTGGAGCTGGACATCGAAGGGCAGCGAGGTGGAGATGCCGTTGGGGTAGAGCTCATGGGTTTCCAGCCCTT
CCGGCTCGATAAAGACCTGGTGGCTCGACTTGTCCGCGAAGCGGTGCACCTTGTCCTCGATGGAGGGGCAGTAGCGCGGCCCGACGCCCTCGATGACGCC
GGAGTACATCGGCGAGCGGTCGAGGTTGGCCAGGATGATCTCGTGGGTGCGCTCGTTGGTATGGGCGATATGGCAGCTCACCTGCCGAGGGTGCTGCTCC
CGACGCCCCATGAACGACATCACCGGGGTCGGGGTATCGCCGGGCTGCTCCTCGAGCTGCGAGAAGTCGAGGCTCCTGGCATCCAGCCGCGGGGGGGTGC
CGGTCTTGAGGCGGTCGACCCGGAACGGCAGGGCGCGCAGGCGCTCGGCGAGGCGATTGGAGGGTGGGTCTCCGGCGCGGCCGCCGCGGCTCTGCTCCAG
GCCGATATGGATCACCCCGCCGAGGAAGGTCCCGGTGCAGAGCACCACGGTTTCGGCAAGGAAGCGAATACCGGTCTCGGTGACCACGCCGCGGACGGTG
TCGTTCTCCACGATCAGGTCGCCGGCGGCCTGCTGAAACAGCGTCAGGTTGGGCTGGTTCTCAAGCATGCCGCGAATGGCGGCCTTGTAGCGCACGCGGT
CGGCCTGGGCTCGGGTGGCGCGCACCGCCGGGCCCTTGCGGGCGTTGAGTACCCGGAACTGGATGCCACCGAGGTCCGTGGCCACGGCCATGGCACCGCC
CAGAGCATCGATCTCCTTGACCAGATGGCTCTTGCCGATCCCGCCGATGGCGGGATTGCAGGACATCTGCCCCAGGGTCTCGATGTTGTGGGTCAGCAGC
AGGGTCTGACAGCCCATGCGGGCGGAGGCCAGGGCGGCTTCGGTTCCCGCGTGGCCACCGCCGATGACGATGACGTCAAAGCGGTCGGGATAATCCAAGG

TGCACCTCGTTGGCGCCACGGCGCCGTGTTGCAGACAGGGGGTGAAATCAGCCCGGCAGTATAACCCCCCTGTGGGTAATCGTCAGGGTTTTCCACATGC
CTGTCGGCCGGCGAGGGAGCTTCTCCATGCCTATTAATATAAATAGAGAAATATAAAGAGAAGTTCTGTTGTGGTAGTAACTACTGGTGTGGACAAAAAC
**CGTGGATAA**GTCAATAAACTCATTTTTTATCAACAGGATGGATTGTTGACCAAGGCGGGGAAAGAGAGCCGGCAGGCTGCGGACAGCCGGGGCTGAGGCT
GTGGATGGAATCTCGACTTGCCCACAGCGCCCTTCGCACACGGCTTGTCCACCGCTGGCTACCGCCCTTGTGCCCGTGGCTGTGAACAACCGGCCTTCTT
ATCCACGACCGCCTCTGGCATGGGCCCTGCAGCGCGGATTGTCCCTTCCGGAGAAAATTCTATCCACAGGCGAAAAAAAGCCCCTGTCGATGCCGACAGG

GGCTCCCTTGCTCGCCTTGCAGGGGGATTGTCAGTGCTTGAGAACCCTGGCCAGGAAGCTACGGGTACGCTCATCGGTGGGCTGGTCGAAGATCTGCTCC
GGCGGCGCCTGCTCGGCGATCTCCCCCTTGTGGATGTAGATCACCCGGTCGGCGACCTCCCGGGCGAAGCCCATCTCATGGGTGACGATGATCATGGTCA
TACCCTCTCGCGCCAGCTCGCGCATGGCATCGAGGACCTCGCCGATCATCTCTGGGTCCAGCGCCGAGGTGGGCTCGTCGAAGAGCATCAGGCGCGGCTC
CATGGCCAGGGCTCGAGCCAGGGCCACACGCTGCTGCTGGCCGCCGGAGAGCTGGCTGGGGTACTTGCGGGCCTGGTCGCTGATGCCGACCCGGGCCAGC
AGCCGTTCGGCGGTCTCCAGGGCGTCTGCGCGGCTCCAGCCGCGGACCTTCATGGGCGCCAGGATGACGTTGTCCAGCACGCTGAGGTGAGGAAAGAGGT
TGAACTGCTGGAACACCATGCCGACCTCGGTGCGGATGCGCTGGAGGGCGCGTCCGCTCTTGCCGTTGGGGAGCAGTTCATTGCCGTCCACTTCTATCTG
CCCGCCCTGGAACTCCTCCAGGCCGTTGATGCAGCGAATCAGCGTCGACTTGCCGGAGCCGCTGGCCCCGATGATCACCACCACCTCGCCGGGCGTGACC
GTGAGGTCGATCTCCTTGAGGACATGGAGGCTGCCGAAGTGTTTGTTGACCCTCTCCATGCGCACGATGGGAGCGGTCGAGTTCGCGTTCTGTGTCATGG
TGGGGCTCCCTTATTGACCGACCAGGCCGCGCTTCTCGAGCATGCGGAGCAGCAGCGACAGGGTCCAGGTGATGGCCAGGTACATCAGGGCCACCATGAA
GTAGACCTCCAGGGCGGTGAAGGTGGTGGCGATATAGATCTGCCCCTGGCGGACCAGCTCGCCCACGCCGATCACCGAGAAGAGCGAGGTGTCCTTGATG

>Halomonas mongoliensis DSM 17332 [NZ_JARWAL010000011.1:c71563..71684], dnaA> ig dnaN>, 122 bp

WebSkew GCmin pos. 91802 (GCmin<>oriC1 27.8 kb; GCmin<>oriC2 8.6 kb; oriC2<>oriC1 19.2 kb)

>Halomonas campaniensis 5AG [NZ_JACHZF010000004.1:c63787..64286], <rpmH oriC1 dnaA>, 520 bp

CGCGGGGCGCTGGGGACCGCCAGGCCATCACCGCTGTCGGCCTCGGCAGCGCCCTCGGCTTGTGGAGCGCTGCTGGTGAAGGAGGGGGCTTCTGGTTCAG
GGGCTACCTGACCGTAGTCCTGATTCCACTGAACGACGATCAGGTAGGCGAGCACCGCCAGTGGAATCAGCAATAGGAGTCGTTTTACGTCCATGAGGGT
TCTGCCCGATGGGTAGTGAAGATGCCAATGGCGCGAAAGGGATCGCACGCCCGATATAGACAAAAGCCTGCCGAAGGCAGGCTGGATTCCGGACACGGAC
TGTCGGCGGTCGTTGCGGCGGCGTCAATGATCACCCGGAGAGGAAGTCGTGGCATCGTGCTGACTCCTTGGCTGGCCGGTCACCCGCCGTGGCCGACGGT
CCCGCAGGTGCTTTCGGCTGGCGCACTCCGTTCCGGCGTCGTGGCCAGGACCTTCCCGGCATCTCGAGTGAGACGTCGCCACATGCCTTGCAGCTGGCGG
TGCAGGGTGGGGTTGTCCAGCTCACCCACACCACGCCTGGCCAGCAACACGATATCCACAGGGGGAAGCTGCTGCTGATGCAGGCGGACGGACTCTCGTG
CGAGACGCTTGAGACGGTTGCGATCGACGGCGCGGCGGAGATTCTTCTTGCTGACCACCAGCCCCAGACGGGGGTGCCCCAGATCGTTGGGCACCGCCAG
GACCAGGAGTCCCTCGCCGTGCACCTTGAAGGCCGCCGTCTCGAAGACGCGCCGATAGTCCCCCGCCGTCAGCAGGCGCAGACGGCGGGGAAAAGCCTGA
CGGGACACGTGCAGTCCGTGATCAGGCGCTCAGGCGCTTGCGACCCTTGGCGCGACGACGCGCCAGGACGGCACGACCGTTCTTGGTGGCCATGCGGGCA
CGGAAGCCGTGCACGCGCTTGCGCTTGAGAACGCTGGGCTGAAAAGTGCGTTTCATAACTCGTTATTCCCACGTGGTTTTTAGGACGAATCGACAATTTC

GAGGCGTCCGGATTCCGCTGGGGAACCCGGTGGTATTCGGCTCAAGACCGGAAATTGTAGTGATTTCGGGCCTCCGGTGCAATTTTTCCGTTCCGTCATC
GCCGCGAAGCCTGACGAGGGTTCGCGACCAGGGAGGGACTACCATCACTATTGGTTAAGGTCTTCCTGTAGATGCTGTTATTAATACTGGGGACAGGTTC
**TGTGGATAA**CATGATTTTTTAAATGTGAAACAACCGGTTGTATTCAGTGAAAATCTGTGGGGAAGTGGCGTGAAAGGCGTGTCGAAGCTGTTGGCGACCG
GTGGACCCTATCGGCAGCGATCCACAGGCCGCAGATATCCCCAGGGCCGGGGGGTGGCTGTCCTGCCGGAAGACGCGGAGTTTTCCACAGCCACCAAGGT
CGTGGGCAAGGGTGTGGATAACCCGTGCCGTCAACGATACAATATGCGGTCGTTTTCCAACCGATGGTGAGGTCGCGTGTCGCTAGCTCTCTGGCAACAG

TGTCTGGACACCCTGCAGGATGAATTGAGCTCCCAGCAGTTCAACACCTGGATTCGCCCGCTGCAGGCGGAGGAGGGGGAGTCGAACGAGCTGTGTCTGC
TGGCACCCAACCGCTTCGTGCGGGACTGGGTCAGCGACAAGTATGCCAAGCGCATCGGGGAGCTGATGCGTGAGCTCTCGCCGGCCAAGCCCCCCAAGGT
GGTGCTGTCGGTGGGCAGCCGCCGCACGGCTCCCAGGCCCCAGCCACGGGAGCTGGGCAACCCGGTATCGGCGAGCCCCAGGCGCCCTGCAGCCCCGGCG
GTGCAGGTGCCGCCGCGAGGTGACGTGGGGGACGAACGCGAGATCGAGGCGCTGCGCGAGGAGGGGGCAGCCAGCCGGCGCGCCGGCGGCGAGCGTCAGG
TGCAGGTCGAGGGCAGCCTCAAGCACAACAGTGGCCTCAACCCCAACTTCACCTTCGAGACCTTCGTCGAGGGCAAGTCCAACCAGCTGGCCCGGGCGGC
CTCTCGGCAGGTCTCCGAGAACCCCGGAGGGGCCTACAATCCGCTGTTCCTCTATGGCGGCGTCGGCCTCGGCAAGACCCACCTCATGCACGCCGTGGGT
AACGCCCTGGCCGGGCGCCGCGAGAACGCCCGAGTCGTCTACCTGCACTCCGAACGCTTCGTGGCCGACATGGTCAAGGCGCTGCAGCTCAATGCCATCA
ACGACTTCAAGCGCTTCTACCGTAGCGTCGATGCGCTGCTGATCGATGACATCCAGTTCTTCGCCGGCAAGGAGCGCTCCCAGGAGGAGTTCTTCCACAC
CTTCAATGCCCTGCTGGAGGGTGGCCAGCAGATGATCCTCACCTCGGATCGCTATCCGAAGGAGATCAGCGGCGTCGAGGAGCGGTTGAAGTCGAGGTTC
GGCTGGGGGCTGACCGTGGCCATCGAGCCACCCGAACTCGAGACGCGGGTGGCGATCCTGATGAAGAAGGCCGATCAGGCCAAGGTCGAGCTGCCCCATG

>Halomonas campaniensis 5AG [NZ_JACHZF010000004.1:c82945..83444], <mnmG oriC2 <COG1126 (glnQ), 528 bp

GCGTTCTCCAGCCCCTTGATCGAGCGCACCACCTGCAGCTGCACGTCGAAGGGCAGCGAGGTGGAGATGCCGTTGGGGTAGAGCTCGTGGGTGTCCAGCC
CCTCGGGCTCGACGAACACCTGGTGGCTCGCCTTGTCGGCGAAGCGGTGCACCTTGTCCTCGATGGAGGGGCAGTATCGCGGCCCGACCCCCTCGATCAC
GCCGGAGTACATGGGCGAACGGTCGAGGTTGGCGAGGATGATCTCGTGGGTGCGCTCGTTGGTGTGGGCGATATGGCAGCTCACCTGCCGGGGATGCTGC
GCACGGTTGCCCAGGAAGGACATCACAGGGGTCGGGGTATCGCCGGGCTGCTCCTCGAGGAGCGAGAAGTCGAGGCTCCGGGCATCCAGCCGCGGCGGAG
TGCCGGTCTTGAGCCGGTCGACCCGGAACGGGAGGGCCCGCAGGCGCTCGGCCAGGCGATTGGAGGGCGGGTCACCGGCACGGCCTCCGCGGCTCTGGTC
GAGCCCGATGTGGATCACCCCGCCGAGGAAGGTGCCGGTGCACAGCACCACGGTCTCGCCGAGGAAGCGGATGCCGGTCTCGGTGACCACGCCGCGGACG
GTGTCGCCCTCGACGACCAGGTCACCGGCGGCCTGCTGGAACAGCGTCAGGTTGGGCTGGTTCTCCAGGATGCCGCGGATGGCCGCCTTGTAGCGAACGC
GATCGGCCTGGGCCCGGGTCGCTCTCACCGCCGGCCCCTTGCGGGCATTGAGCACGCGGAACTGGATGCCACCCAGGTCGGTGGCAACGCCCATGGCACC
GCCCAGCGCATCGATCTCCTTGACCAGATGGCTCTTGCCGATGCCGCCGATGGCGGGATTGCAGGACATCTGGCCCAGGGTCTCGATGTTGTGGGTCAGC
AGCAGGGTCTGACAGCCCATGCGGGCGGAGGCCAGGGCGGCTTCGGTTCCCGCATGGCCACCGCCGATGACGATGACGTCAAAGCGGTCGGGGTAATCCA

AGGTGCACCTCGTTGGCGCCGGGCGCCTGTCTCGCGGTCAGGATGAAATCAGCCCGCCAGTATAAACCCCCTGTGGGTAATCGTCAGGGTTTTCCACATG
CCGATTTCGGGTCTCCACCACGCCGGCTTCCCTATCAGTTAAAACAGAATATAAAGAGAGAAGTTCTGTTGTGGTTGTTACTACTGGTGTGGACAGTTTC
**CGTGGATAA**CTAAATAATCCAGATTATTATCAACAGGATGCATTGTTGACCAAAGCCAGGACAAGCGGGTGACTGCCTGCGGAGGGAGCCGCCGGAGCCT
GTGGATGGAATCGCGACCTGTCCACAGCGTGAGGCCTGCAAGGCTTGTCCACCGGCGGCTACAAGGCTTGTGCCCGCGGCTGTGAACAACCGCCCCTGGC
TGTCCACAACGCCTCTGGCATGGGGCCCCTGGGGCATTATCGCGATATCTGGCCGATCTCTTGTCCACAGGCAAAAAAAAAGGCCCTGTCGCAGTCGACA

GGGCCCGATCTGGCGGCGCCGAGCGGGGGTCAGTGCTTGAGGACGCGCGCGAGGAAGCTGCGGGTGCGCTCGTCCTCGGGGGTGTCGAAGATCTTCTCGG
GCGGCGCCTCCTCGGCGATCTCCCCCTTGTGGATGTAGATGACCCGGTCGGCCACCTCCCGGGCGAAGCCCATCTCGTGGGTGACGATGATCATGGTCAT
CCCCTCCCGCGCCAGCTCGCGCATGGCATCCAGCACCTCGCCGATCATCTCCGGGTCGAGCGCGGAGGTGGGCTCGTCGAAGAGCATCAGCCGCGGCTCC
ATGGCCAGCGCCCTGGCCAGCGCCACCCGCTGCTGCTGGCCGCCGGAGAGCTGGCTGGGGTACTTGGCGGCCTGGTCGCTGATGCCGACCCGTTCCAGCA
GCCGCTGGGCCGTCTCCACGGCGTCGGCACGGCTCCAGCCGCGTACCTTCATGGGCGCCAGGATGACGTTGTCGAGCACGCTGAGGTGCGGGAAGAGGTT
GAACTGCTGGAACACCATGCCCACCTCGGTGCGGATCTTCTGCAGCGCCTTGCCGCTCTTGCCGTGGGGGGTGAGCTCGTTGCCGTCCACCTCGATGTGG
CCGTCCTGGAATTCCTCCAGGCCGTTGATGCAGCGGATCAGCGTCGACTTGCCGGAGCCGCTGGCGCCGATGACCACGACCACCTCGCCGGTAGCTACCT
CGAGGTCGATGTCCTGGAGCACGTGCAGGCTGCCGAAGTGCTTGTTGACCTTCTCCATGCGCACGATGGGAGCGGTCGAGTTCTCGTTCTGTGTCATGCG
TGGGCTCCCTCTCATTGGCCGACCAGGCCGCGCTTCTCGAGCTGGCGCAGCAGCAGCGACAGGCTCCAGGTGATGGCCAGGTACATCAGTGCCACCATGA
AGTAGACCTCCAGGGCGGTGAAGGTGGTGGCGATGTAGATCTGCCCCTGGCGGACCAGCTCGCCGACGCCGATCACCGAGAACAGCGAGGTGTCCTTGAT

>Halomonas campaniensis 5AG [NZ_JACHZF010000004.1:c62225..62340], dnaA> ig dnaN>, 116 bp

WebSkew GCmin pos. 1381086 (GCmin<>oriC1 26.7 kb; GCmin<>oriC2 5.1 kb; oriC2<>oriC1 21.5 kb)

>Halomonas campaniensis LS21 [NZ_CP007757.1:1407504..1408003], <rpmH oriC1 dnaA>, 534 bp

AGAACGTCGGTCGTGACAGCAATAAAGTCTCGGCTAGACGAGTTACTTTCTGCTTCTCCAGGCATGCCTTCACTAATGGCATTTTGGGGAGCCGATGAAC
TAGGAACCGATAAATCATCACTGCCAACATTATTGGCGGATGAGCCGTTGCTGTTTTGGGTAATTTCAGGCGTCGAATCGACGCTTATTTGCCCATAATC
CTGATTCCACTGTACAACCAATAGATAGGCAAGTACTGCCAGTGGAATCAGTAATAAAAGTCGTTTTACGTCCATGAGGGTTCTGCCCGATGGGTGGTGA
ACATGCTAATGGCGCAGTGTCATTTACTCGACCCATGACACACGAAAGCCTGCCAGGCGGCAGGCCGAGGTCGAGCGGCGAGGTGTGTCGTCAAGCGTCA
CGTGTTACGCCATTAGGCGGGAGTGGTACAGACACCGCCTGTGCGTCACGCTGAAGTCGTTTCCACATGCCATTTAGCTGGCGATTAAGCGTCTCGTTAT
CAATATCTTGAACACCACGCTTTGCCAGTATCACGATATCCACAGAGGGTAAGTGATCTTGGCGGAGGCGAACAGATTCTCTCACGAGTCGCTTAAAGCG
GTTGCGATCAACAGCGAGCTTCACATTCTTTTTGCTGACCACCAGTCCAATGCGGGGATGTCCCCGGGTACTCAAACGCGCTAGCGCCATCATACCTTTG
CCATGAACCTTCAGGTCTGCTTGTTCAAACACGTGACTAAAATCCCCGGCGTTTAGCAAACGTAAGCACCGGGGAAAGCCTTGATGCGGCACACGCAATC
CGAGATCAGGCGCTCAGACGCTTGCGGCCTTTGGCGCGGCGGCGTGCGATAACGGCACGGCCGTTTTTGGTTGCCATACGAGCACGGAAGCCATGCGCGC
GCTTGCGCTTGAGAACGCTGGGTTGAAAAGTGCGTTTCATAACTCGTTATTCCCACGTGGTTTAGGACGGAAAGTGAATTCCAAAAGACCCGGTCCAGTG

AGGAACTAGCCGGGGAGTTTCAATTCAAGACCGGAAATTGTAGAGACTTCCCGCGTCAGGTGCAATTTTTCCTTGCATAAAAAGCGTTAAAAAGGCAATG
GAGGCTCTTCGTACATCGCCATCGTGGCTTCTTGCCATGACACTACTATTACTAATGAAATTATTTAAAACCTATTGTTATTAATAGTGGGGATACTAAT
**TGTGGATAA**GCTATTTTAGCTTACTTTTATCAACAGCTTAAAAACATAAAAAATCTGTGCATAACCTTCGGATAAGCCTTAAATAAACTGTTAATTAGGT
GTGTTCAACAGGGTATAGAAGACATCAAAACAGCTTTTTCCTTGATCCATTACCAGCTTTTCCCTATGCCTCATGAATAGTTGTCCACAGGGCAAAAAGA
GGCGTTAGTGCGTGTGGATAACCGTCTCGTTGGGCGCTACAATGGTCGGCCGTTTTCAACCGATGGTGAGATGAGTGTCACTCGCGCTTTGGCAACAGTG

CCTGGACTACCTGCAGGACGAACTGAATTCACAGCAGTTCAATACCTGGATACGTCCGCTGCAGGCAGAAGAAGGAGAGACCAATGAGCTGCGCTTGTTG
GCGCCGAATCGCTTTGTGCGCGACTGGGTGAGTGATAAGTATTCAAAGCGTATCAGCGAGCTGATGCGAGAACTTGCGCCATCCAAGCCTCCCAAAGTCA
GTTTGACGGTAGGTAGCCGCCGTGTGGCTGCTCCTGCCCCGCAGCCAAGAGATTTGGGTAATCCGGTCTCAGCATCTCCCTCAAGGCAAGCATCGCCTGC
GGTTCATACGCCGCCAAGGGGAGACCTTGCTGATGAGCGAGAAATTGATCAACTGCGTGAAGAGTCACCGAATCGGCGAGGAAATGAGCGCCAGGTACAG
GTTGAAGGCAGCCTAAAACATGGTAGTGGTCTTAATCCTAATTTTACCTTCGATACCTTTGTTGAAGGTAAATCGAACCAGCTCGCTCGTGCGGCATCCC
GTCAGGTATCTGAAAATCCTGGCGGTGCCTATAACCCGCTGTTTTTATACGGTGGTGTTGGTTTAGGTAAAACACACTTGATGCATGCTGTCGGTAACGC
CCTGGCTGGCCGCCGCGAAAATGCCCGTGTGGTGTATCTACACTCTGAACGCTTTGTCGCGGATATGGTAAAAGCACTGCAGTTAAATGCGATTAACGAC
TTCAAGCGCTTTTATCGCAGCGTTGATGCGCTACTTATTGATGATATTCAGTTTTTTGCTGGAAAAGAGCGTTCCCAGGAAGAGTTCTTCCATACCTTTA
ATGCGTTGTTAGAGGGTGGGCAGCAAATGATACTTACCTCTGATCGCTATCCAAAAGAGATTAGTGGTGTGGAGGAGCGTCTGAAATCGCGCTTTGGCTG
GGGCCTGACAGTGGCTATTGAGCCGCCAGAGCTTGAAACCCGTGTAGCGATTTTGATGAAAAAGGCAGATCAGGCGAAAGTTGATCTTCCACACGACGCT

>Halomonas campaniensis LS21 [NZ_CP007757.1:1385956..1386455], <mnmG oriC2 <COG1126 (glnQ), 565 bp

TAACCCTTTAATTGAGCGAACCACTTGTAGCTGAACATCGAAAGGCAACGACGTCGAGATTCCGTTGGGATAGAGCTCATGGGTATCCAACCCTTCAGGC
TCGATAAATACCTGATGGCTTGATTTGTCGGCAAAGCGGTGAACTTTGTCTTCAATCGATGGGCAGTATCGCGGACCAACACCTTCGATCACACCGGAAT
ACATAGGCGAGCGATCAAGATTCGCCATGATGATTTCATGGGTGCGCTCATTGGTGTGCGCTATGTGGCAACTCACCTGCTGCGGATGCATCTCCCGTGA
ACCAAGGTATGACATGACCGGTGTTGGGGTGTCACCTGGCTGTTCTTCCAACGATGTGAAGTCAACGGTTTTGGCATCAATTCGTGGAGGCGTCCCGGTT
TTTAGCCGATCAACGCGAAACGGTAGTGCGCGTAAGCGTTCAGCTAGCGCGTTGGAAGGCGGGTCACCGGCGCGGCCACCGCGGCTTTGATCTAACCCAA
TGTGGATAACGCCACCCAAAAACGTACCGGTTGACAGCACAACGGACTCCGCATGGAAGCGGATGCCTGTTTCGGTCACAACGCCTCGAACAGTGTTGTT
ATCCACAACCAGATCGCCAGCGGCTTGCTGAAAAATCGTTAAGTTCGGCTGGTTTTCCAGCATTCCCCGAATAGCAGCTTTGTAGCGAATTCGGTCTGCT
TGAGCACGTGTTGCTCGAACGGCTGGACCTTTACGAGCGTTCAATACCCGAAACTGGATGCCGCCTAAGTCCGTGGCTAGCCCCATTGCACCACCGAGTG
CATCAATTTCTTTTACCAGGTGGCTTTTGCCAATCCCACCAATCGCTGGATTGCACGACATTTGGCCGAGCGTTTCGATGTTGTGGGTTAGCAATAGGGT
TTGACAACCCATACGAGCAGAGGCCAATGCGGCTTCGGTTCCCGCATGGCCACCGCCGATGACAATCACGTCAAAGCGGTCGGGATAGTTCAAGAGGCAC

CTCGTCTTGCGCCGTTTGGCACGGATGATGTGAGTCAGTATTGGTTCACTGATTGAGTAAGCCCGCCAGTATAAACCTCCTGTGGGTAACCGTCAGGTTT
TTCCACACCCCAAGCTAAGCATCAATCTATTATTAATAACAAAATAAAAATATATAAGAGAAGTTCTGTTGATGTTGTAACTAACTGGTGTGGACAAAATC
**TGTTGATAA**CCCAAATAAGCTAATATATTTCAGTTGCTTAAATTGTTAGTTTTTACATGATTAAACGGAGGAGTACCTGCGTAGAAGCGGTGCTACTCCT
GTGGATGAAAGTGCGACTTACACACAGGCATGACAGACAACGGGTTATCCACCGACCCATACCGAGTTTGTTACCGCACCTGTGAACAAGTGCGATTGGT
ATCTTATAAAGACCTGGTTAGCGCTCATGCTAGGCATCGTAGAGCGATATGATGCAGCGTGTAAAAAATAGTATCCACAGGCAAAAAAAATGGCCTGTCG

CGATGACAGGCCATTTTACGGGGCAAAAATGGGCAAAAAAGTCAGATTAATTCCGACATTAATGCTTCAAAACACGTGCAAGGAAGGATTGTGTTCGCTC
ATGTTGTGGCGTATCGAACAGTTGCTCGGGAGGGCCTTGCTCGGCAATTTCGCCTTTATGGATAAAAATAACGCGATCTGCCACTTCACGGGCAAAGCCC
ATTTCATGGGTCACTATCACCATGGTCATGCCTTCTTTGGCAAGTTCACGCATGGCATCCAGCACTTCACCAATCATTTCAGGGTCAAGCGCTGATGTAG
GCTCGTCAAACAGCATCAGGCGGGGTTCCATGGCTAACGCCCGGGCTAACGCCACACGCTGCTGCTGCCCACCCGAGAGCTGGCTTGGGTACTTATCTGC
TTGGTCGGCAATACCAACGCGCTCTAACAGTCGTTCAGCGGTCTCTTCGGCATCTTGGCGGCTCCAGCCCCGCACTTTCATGGGCGCAAGGGTAACGTTG
TCGCGCACGCTCAAGTGGGGAAAAAGGTTGAACTGCTGAAACACCATGCCGACTTCAGTACGGATAGTTTGTAGCGCTTTACTGCTTTTACCGTTGGGCA
GTAATTGGTTGCCATCAACATCAAGAGAGCCGGCTTGAAACTCTTCTAAGCCGTTGATACAGCGGATGAGCGTTGATTTACCAGAGCCACTGGCACCGAT
GATAACGACGACTTCACCAGGCACAATTTCAAGATCAACATTATTGAGTACGTGCAGGCTGCCAAAATGCTTATTGAGCTGCTGCATACGCACAATTGGC
GATTGGGTAGAGGTCATTGCACGTTCCTTATTGTCCTGCGAGGCCTCTGCGCTCAATTTGGCGCAGAATCAAAGAGAGAGTCCAGGTGATGGCAAGGTAC
ATCAGCGCCACCATGAAATAGACCTCAAGTGCGGTAAAGGTAGTGGCAATATAAATTTGCCCTTGGCGGACTAATTCACCGACCCCGATAACAGAAAATA

>Halomonas campaniensis LS21 [NZ_CP007757.1:1409445..1409499], dnaA> ig dnaN>, 55 bp

WebSkew GCmin pos. 333234 (GCmin<>oriC1 27.9 kb; GCmin<>oriC2 3.0 kb; oriC2<>oriC1 24.9 kb)

>Halomonas urumqiensis BZ-SZ-XJ27 [NZ_PYUD01000002.1:360855..361354], <rpmH oriC1 dnaA>, 516 bp

TCGGGTCGCTTGCGGTCGCTGGCATGGACAGGGCGTCCTCACCATCGCCTTCCGTCTCGACCTGATTGAGCGGGGCGCTGCTGCTGATAGGGGGTGCGTC
AGGCGGCGTGACTGCCTGGCCATAATCCTGATTCCACTGTACGACGATCAGATAGGCGAGCACCGCCAGTGGAATCACCAAGAGGAGTCGTTTTACGTCC
ATGAAGGTTCTGCCCGATGGGAAGTGAACATGCCAATGGCGTGATAATGAGGAGCGCACGCCCGATACAGACGAAAGCCTGCCAAAGGCAGGCCGAATTC
CGAACAGGGACTCTCGACGATTGTCTCGGCGGCGTCAACGATCACCCGATTGGGAAGCCGTATCATCATGCTGACTCCAGGGCTTGTCGTTGGCTAGCCA
TTGCTGCCGTTGGCGGGATCAAGTGACACATGATTGTCCTGCCTTGCCGATTTGCGAACGTCTTTCGCCAGACGACGCCACATACCTTGTAGCTGCCGAA
ACAGGGTGTCGTTGTCCAGTTCGCCGACGCCACGTCGGGCCAGCAAGACGATATCGACGTCAGGCAGTTGATGTTGCCGGTGCCGGACGGACTCGCGAGC
GATTCGTTTGAGACGGTTACGATCGACAGCGCGGCGCAGGTTCTTCTTGCTGATGACGAGCCCCAGGCGTGGATGACCCATTGCGTTCGGACTTGCCAGG
ACCAACAGACCCTTGCCATGGACCTTGAAGGCCGCTGTATCGAAAACGTGACGATAATCCCCGGCCGTCAGCAGTCGCAGACGGCGGGGAAAGCCTGAG
GGGACAACTGCGTGTCCCGGATCAGGCGCTCAGGCGCTTGCGCCCCTTGGCGCGACGACGCGCCAGAACGGCACGACCATTCTTGGTGGCCATACGGGCA
CGGAAGCCGTGCACGCGCTTGCGCTTGAGGACGCTTGGCTGAAATGTGCGTTTCATAACTCGTTAATCCCACGTGGTTTTTAGGGCAAATTGACAACATA

GAGGCGTCCGGATGACGTGTTGGCCATCCGGGGGTGTTCGGCTCAAGACCGGAAATTGTAGTGATTTCGCGGCCAAGGTGCAATTTTTCCGTTCCGCCCC
CGGAACAGGTGCGACCTGAGACCTGCGGGATGTACCGGTGGTAAACCATCACGACCATTGTTTAGCGATTGATGTTGTTATCAATACTGGGGATAGTATC
**TGTGCATAA**CAAACCATATGCATTTAAAATCAATTACTTGCTGCATAGCAATAACAGTGGATAAATGTAGCGTGACGCGAGCCGCCGCCGGAGATTCCCT
GTGGAAGCCGCGGGCGCTAAATCCACCTCCCTGATCCAGGCATCACCTGGCATGGCATTCCCCCTGGCCCCGGATGTGTTTTCCACAACATGGAGGAAAG
CTGGCGGGGTGTGGATAACCCTGCCCCCGGGAGATACAATATGCGGTCATTTTCCATCTATGGTGAGGTCGCGTGTCGCTCGCTCTCTGGCAACAGTGTC

TTGAAACCCTGCAGGATGAACTGAGTTCCCAACAGTTCAATACCTGGATCCGCCCCTTGCAGGCGGAGGAGGGTCAGGCCAGTGAGCTATGCCTGCTGGC
ACCGAATCGATTCGTGCGTGACTGGGTAAGTGACAAGTATGCCAAGCGCATCGGTGAATTGATGCGTGAGCTTGCGCCTGCCAAGCCTCCCAAGGTAGTG
CTCACGGTGGGCAGCCGGCGGACCGCGCCACGACCACAGCCTCGCGAGTTGGGGAATCCCGTCTCGGCTAGCCCTCGTCGACCCGGGGCACCCGCCGTGC
ATGTGGCGCCGCGTGGAGATGTGGCCGACGAGCGCGAGATCGACTCACAACGCGAAGCTAATCAGTCTACCGGCAATATCGACCCGTCGCCTGCACGGCG
TCCCGGTAGTGAGCGGCAAGTCCAGGTAGAGGGTAGCCTCAAGCATAACAGCGGCTTGAACCCCAATTTCACTTTCGAGACCTTCGTCGAGGGCAAGTCG
AACCAGTTGGCCAGGGCAGCCTCGCGCCAGGTGTCCGAGAACCCTGGTGGTGCCTACAATCCGCTGTTCCTCTATGGTGGCGTCGGCCTCGGCAAGACCC
ACCTGATGCATGCCGTGGGCAACGCATTGGCGACTCGTCGTGAGAATGCGCGCGTGGTCTACCTTCACTCAGAGCGGTTCGTGGCCGACATGGTCAAGGC
GCTGCAACTCAACGCTATCAATGATTTCAAGCGCTTCTATCGAAGCGTCGATGCCTTGCTGATCGATGACATCCAGTTCTTCGCCGGCAAGGAGCGCTCA
CAGGAGGAGTTCTTCCACACCTTCAATGCGTTACTCGAAGGTGGACAGCAGATGATCCTGACATCCGACAGATATCCCAAGGAAATCAGTGGGGTGGAGG
AGCGTCTCAAGTCCCGCTTCGGATGGGGGCTGACGGTTGCCATCGAGCCTCCCGAGCTCGAGACACGTGTGGCGATCCTCATGAAGAAGGCCGACCAGGC

>Halomonas urumqiensis BZ-SZ-XJ27 [NZ_PYUD01000002.1:335989..336488], <mnmG oriC2 <COG1126 (glnQ), 574 bp

ACATCGAACGGCAGTGACGTGGAGATCCCATTGGGATAGAGCTCGTGGGTGTCGAGGCCTTCGGGTTCGATGAAAATCTGGTGGCTCTGCTTGTCGGCAA
AGCGATGCACCTTGTCCTCGATCGATGGGCAGTAACGTGGCCCGGTGCCCTCGATTGCACCGGAATACATCGGTGAACGATCGAGATTCGCGAAGATGAT
CTCGTGGGTGCGCTGGTTGGTATGCGCGATGTGACAGCTCATCTGACGCGGGTGCTGTTCGGGCTTGCCCATGAACGACATCACCGGCGTCGGGGAATCG
CCCGGCTGCTCCTCGAGCATCGAAAAATCGACGCTCTTGGCATCGAGGCGTGGTGGAGTACCGGTCTTCAGGCGGTCGACCCGGAATGGCAACGAACGCA
GACGCTCGGCCAGGGCGTTGGAAGGCGGGTCACCGGCACGGCCACCCAGGCTCTTGTCCAGCCCGATATGGATCACCCCGCCGAGGAAGGTGCCGGTGCA
CAGTACGACCGTCTCGGCAAGAAAACGGATCCCGGTCTCGGTGACCACGCCGCGCACGGTGTCGCCATCAACTACCAGGTCTCCGGCCGCCTGCTGGAAG
ATCGTCAGGTTGGCCTGGTTTTCCAGCATGCCCCGAATCGCCGCCTTGTAACGGACTCGGTCGGCCTGGGCGCGAGTTGCACGCACAGCGGGCCCCTTGC
GGGAATTGAGCACGCGAAACTGGATGCCTCCCAGGTCGGTAGCCAGCGCCATGGCCCCACCCAACGCATCGATCTCCTTGACCAGATGACTCTTGCCAAT
GCCGCCGATGGCGGGATTGCAGGACATCTGGCCCAGGGTCTCGATGTTGTGGGTCAAAAGCAGGGTCTGACAGCCCATGCGAGCGGAGGCCAGCGCGGCT
TCGGTTCCCGCATGGCCACCGCCGATGACGATCACGTCAAAGCGGTCTGGGTAATCCAAGGTGCACCTCGTTGGCGCCGATAGCGCCTGCGTTGTCGTCA

GGTAGTTGCGTTGCCCCGCAAGCTCCGGAAACGGAGCCGCTGTGAACAAGCCGGCCAGTATAAACCCCTTGTGGGTAATCGTCAGGGGTTTTCCACACCT
GGTTTTCAGGCCCGTGGTGGTATTGGGCTATCTATCAGTTAAATCAAAATATATAGAGAGAAGTTCTGTTGTGGTAGTAACTAACTGGTGTGGACAAAAAC
**CGTGGATAA**GTGAATTAATCAACATAAAAACAAAATGATACGCTGTTGACTCAAAGACGCACAAGCGGGTGACTGCCTGGGAGGGACCTGTCTGGAGCTT
GTGGACGGAATGCCACTCTATCCACAGTGGAGAGAATCTACCGCTTGTCCACCGTCGGGTACATGCCTTGTGCCCGTGGCTGTGAACAACCTCGATGATG
TGTGGTGAAGGCCTTGGGAGTGGGCCTTGGCGGCGTATATGCCAGGGCGCCAGGAAAATTCTATCCACAGGCATGAAAAAGGCCCTGTCGATAACGACAG

GGCCCTCTTTCGCTCGTCTTCTACGCCTGTTTTTCAGTGCTTTAGCACGCGGGAGAGGAAACTGCGTGTTCGCTCGTCCTTGGGTTGATCGAAGATCGTC
TCGGGCGGTGCGGCCTCGGCGATCTCGCCCTTGTGGATGTAGATGACGCGATCCGCCACTTCCCTGGCAAAGCCCATCTCATGAGTGACGATCACCATGG
TCATGCCTTCGCGGGCGAGCTCACGCATGGCATCCAGTACTTCGCCGATCATTTCAGGATCGAGTGCCGAGGTGGGCTCGTCGAACAGCATCAGCCGAGG
TTCCATGGCCAGTGCCCGGGCCAGCGCCACACGCTGCTGCTGACCTCCGGAGAGCTGCGTGGGATGCTTGGCGGCCTGGTCGCTGATGCCGACCCTTGCA
AGGAGACGCTCGGCGGTCTCCACCGCGTCCGCTCGGCTCCAGCCACGTACCTTCATGGGTGCCAGGATGACGTTGTCGAGCACGCTCAGGTGGGGAAACA
GGTTGAACTGCTGGAACACCATGCCGACTTCGGTGCGGATCTTTTGAAGTGCCTTGCCGCTCTTGCCGTGGGGGGTGAGTTGGTTGCCATCGACCATGAT
ATGGCCGTCCTGGAACTCCTCGAGGCCATTGATGCAGCGAATCAGCGTCGACTTGCCGGAACCACTGGCACCGATCACCACCACTACCTCGCCGGGTACC
ACGTTCAGGTCGATGTCCTTCAGCACATGCAGGTCGCCGAAGTGCTTGTTGACCTTCTCCATCTGGACGATGGGAGCGGTCGCTGTGTCGTTATCGGTCA
TGCGGTGACTCTCCTCATTGTCCGGCCAGGCCGCGCTGTTCAAGGAAACGCAACACCAGCGATAGCGTCCAGGTGATGGCGAGGTAGAGCAGCGCCACCA
TGAAGTAGACCTCAAGCGCCGTGAAGGTGGTGGCGATATAGATCTGGCCTTGACGCACCAGCTCACCGACCCCAATCACCGAGAACAGAGAGGTATCCTT

>Halomonas urumqiensis BZ-SZ-XJ27 [NZ_PYUD01000002.1:362821..362935], dnaA> ig dnaN>, 115 bp

WebSkew GCmin pos. 83834 (GCmin<>oriC1 27.6 kb; GCmin<>oriC2 8.5 kb; oriC2<>oriC1 19.0 kb)

>Halomonas heilongjiangensis DSM 26881 [NZ_PNRE01000019.1:c56014..56513], <rpmH oriC1 dnaA>, 519 bp

CTTCGGCCCGCGGCGTCGTCGGGACGGCGAGGTCGTCGTCTCCATCCGGGGCGGCGGTGTCCTCGCCAGGGGCTCCAGTGGTGAAGGAGGGGGCTGTCGG
TTCAGGCGCTACCTGACCGTAATCCTGGTTCCACTGCACGACGATCAGGTAGGCGAGCACCGCCAGGGGAATCAGCAAGAGGAGTCGTTTTACGTCCATG
AGGGGTCTGCCCGATGGGTGGTGAACATTCCGATGGCGTGGTGCGGGTCGCACGCCCGATATACACGAAAGCCTGCCGAAGGCAGGCTAGGCTCAGTACA
CGGATTCTCGGCACTCGGCATGCTGGTAGCGATGATCGTCCGGCAAGAGGTGGTGGCTCTCATGCAGACTCCGTGACTGGCCGCGCCGCGCCGATGCCGA
TACGCGGCGACGCAGCTGGCAAGTTGCCCTGGGCCCATCAGCCGGTCTTCGACTTCTGCACGTCTCGCGTCAGACGCTTCCACATGCCGTGAAGCTGGCG
ATGCAGGCTGGCGTTGTCGAGCTCGTGAACGCCACGCCTGGCCAGAATCACGATGTCCACGGAGGGCAGGCGCTGTTGCTGCAGCCGGATCGACTCGCGA
GCGAGTCGCTTGAGGCGGTTGCGGTCGACGGCTCGCCGGACGTTCTTCTTGCTGAAGACCAGGCCGAGTCGGGGGTGTCCCAGGTCGTTGGGAGAGGCCA
GTGCCATCAACCCCTTGCCATGGACCTTGTATGCAGCCGTATCGAAGACGCGCCGAAAATCCCCGGCCGTCAGAAGACGCAGACGCCGGGGAAAATCCTG
ACGGGACACGTGCGGTCCGCGTTCAGGCGCTCAGGCGCTTGCGACCCTTGGCGCGACGACGCGCCAGAACGGCACGACCATTCTTGGTGGCCATGCGGGC
ACGGAAGCCGTGCACGCGCTTGCGCTTGAGAACGCTGGGCTGAAAGGTGCGTTTCATAACTCGTTATTCCCACGTGGTTTTCAGGAAGAATCGACAATGT

CGAGGCGTCCGGATCCCGTATCGGAACCCGGGGGGATTCGGCTCAAGACCGGAAATTGTAGTGATTTCGGTGCTCGGGTGCAATTTTTCCGTATCGATGC
CGGTGGATGGCAGGGGAAGGGGAACGCCGCCCTGCCACGAATCAGTCATTATAATTAAAGATCTTTAGATGATGTTGTTATTAATACTGGGGACAGCTTC
**TGTGGATAT**GTGCAATTAACCGCATTAGATCAAAATGTTATTTGATAGACTTGGCTGTGGGCAAGTCTCGCATGACCGGAGGATGGCCTGTGCATCGACG
GTGTACCCCGCCGGGCATCCATCCACAGGGCCGGGCCGCCCCCAGGGGAATCCCGCGGCTGTCCCCGCCGATCATCCGGGTTGTCCACAGCATGCCGCGA
GGGGGTGGCTCATGTGGATAACCTTGGCCCTGAACGATACAATGTCGGTCGTTTTTTCGATGAATGGTGAGGTCGCGTGTCGCTCGCTCTCTGGCAACAG

TGTCTGGACTACCTGCAGGACGAACTGAATTCCCAGCAGTTCAATACCTGGATTCGCCCCTTGCAGGCGGAGGAGGGGGAGTCCAACGAGCTTTGTCTGC
TGGCACCCAACCGGTTCGTCCGTGACTGGGTCAGCGACAAGTATGCCAAGCGCATCAGTGAGTTGATGCGGGAGCTGTCGCCCGCCAAGCCACCCAAGGT
GGTGCTGTCGGTGGGCAGTCGTCGCACGGCGACCGCTCCCCAGCCGCGGGAACTCGGCAATCCGGTGTCGGCCAGCCCCCGACAACCGGTGGCACCGGCG
GTCCAGGTGCCGTCGCGTGGCCGCGGAGAGGTCGGGGACGAGCGCGAGATCGAAGCGCTGCGCGATGAGGGGGGCGCAGCCGGTCGGCGTCCCGGCGAGC
GCCAGGTCCAGGTAGAGGGCAGCCTCAAGCACAACAGCGGACTGAATCCGAACTTCACCTTCGAGACCTTCGTCGAGGGCAAGTCCAACCAGTTGGCGAG
GGCGGCGTCGAGGCAGGTCTCCGAGAATCCTGGAGGGGCCTACAACCCGCTCTTCCTCTATGGGGGCGTGGGTCTCGGCAAGACCCACCTGATGCATGCC
GTGGGCAATGCCCTGGCCGGACGTCGCGAGAATGCGCGGGTGGTCTACCTGCATTCCGAGCGTTTCGTGGCCGACATGGTCAAGGCGCTCCAGCTCAATG
CCATCAACGACTTCAAGCGCTTCTATCGCAGCGTGGATGCCCTGCTGATCGACGATATCCAGTTCTTTGCCGGCAAGGAGCGTTCCCAGGAGGAGTTCTT
CCACACCTTCAATGCGTTGCTCGAGGGTGGCCAGCAGATGATCCTCACGTCGGACCGTTATCCCAAGGAAATCAGTGGGGTAGAGGAGCGGTTGAAGTCA
CGTTTCGGCTGGGGGCTGACGGTGGCGATCGAGCCGCCGGAGCTCGAGACCCGGGTGGCGATCCTGATGAAGAAGGCCGATCAGGCCAAGGTCGAGCTGC

>Halomonas heilongjiangensis DSM 26881 [NZ_PNRE01000019.1:c75036..75535], <mnmG oriC2 <COG1126 (glnQ), 563 bp

CAGTTGCACGTCGAAGGGCAACGAGGTGGAGATGCCGTTGGGGTAGAGCTCGTGGGTGTCGAGGCCCTCGGGCTCGATGAACACCTGGTGGCTCGCCTTG
TCGGCGAAGCGGTGCACCTTGTCCTCGATGGAGGGGCAATAGCGCGGCCCGACCCCCTCGATCACCCCGGAATGCATCGGCGAGCGGTCGAGGTTGGCCA
GGATGATCTCGTGGCTACGCTCGTTGGTGTGGGCGATGTGGCAGCTCACCTGGCGCGGATGCTGATCGCGCCGGCCCATGAACGACATCACCGGCGTCGG
GGTATCACCGGGCTGCTCCTCGAGCACCGAGAAATCGAGACTCCTGGCATCGAGGCGCGGCGGCGTCCCGGTCTTCAGGCGATCGACCCGGAACGGCAGT
GCCCGCAGCCGCTCGGCCAGGGCATTGGACGGTGGATCCCCGGCGCGGCCGCCGCGACTCCGGTCGAGCCCGATATGGATCACCCCGCCGAGGAAGGTGC
CGGTGGCGAGCACGACGGTCTCGCCGCGGAAGCGGATGCCGGTCTCGGTGACCACGCCACGCACGGTGTCGCCATCGACGATGAGATCGCCGGCGGCCTG
CTGGAACAGCGTCAGGTTGGGCTGGTTCTCCAGCATCCCGCGGATGGCCGCCTTGTAGCGAACCCGGTCGGCCTGGGCGCGAGTGGCGCGTACGGCGGGC
CCCTTGCGGGCGTTGAGTACCCGGAACTGGATCCCGCCGAGGTCGGTGGCCGACGCCATGGCACCGCCCAGGGCATCGATCTCCTTGACCAGGTGGCTTT
TGCCGATCCCGCCGATGGCGGGGTTGCAGGACATCTGGCCGAGCGTCTCGATGTTGTGGGTCAACAGCAGGGTCTGACAGCCCATGCGAGCGGAGGCCAG
AGCGGCTTCGGTTCCCGCATGGCCACCGCCGATGACGATGACGTCAAAGCGGTCGGGATACTCCAAGGTGCACCTCGTTTGGCGCCGTGGCGCCGATGTC

GTTGTTTCGGGTCGTCGATAGGGACCGTCGGAAAGTGAGTCCGAATCAGCCCGGCAGTATAAACCTCCTGTGGGTAATCGTCAGGGTTTTCCACACCGGG
TCTCGCCGAAGGCCGCCTGCGGGCTTGCCAGCTAATCAGATAAAACAGAATAAAAGAGAGAAGTTCTGTTGTGGTAGTGACTATAGGTGTGGACAAAATC
**CGGGGATAA**CCATGTTTTCTCTTTTATTATCCACTGGTTGCATTGTTGACCAAGAGCTGGACAAGGGGGCGGGTGGCTGCGTACAGCTGGTCGAGAGGCT
GTGGACGGAATGGCGACTTGCCCACAGCGACGCGGGCGCACCGCTTGTCCACCGCCGGCTACCGGGCTTGTGCCCGTAGCTGTGAACATCCCTTCGCGGG
TGCAACGAAGCCGACGGTGGCGGGGGGTTCAGCGCCTTGGAGAGGACCTGTGCAGGAATTCTGTCCACAGGCAGAAAAAAGGCCCTGTCGTTGACGACAG

GGCCGCGAAGGTGGCCGGAAAGGCAGGATTCAGTGCTTGAGCACCCGTGACAGGAAGTTCCGGGTGCGTTCGTTCTGCGGTGAGTCGAAGAGTCGTTCGG
GGGGGGCCTGCTCGACGATCTCCCCCTGGTGGATGAAGATCACCCGGTCGGCCACCTCGCGGGCGAAGCCCATCTCGTGGGTGACGATCACCATGGTCAT
GCCCTCCTTCGCCAGTTCACGCATGGCGTCGAGCACCTCGCCGATCATCTCCGGGTCGAGCGCCGAGGTGGGTTCGTCGAACAGCATCAGCCGCGGTTCC
ATGGCCAGCGCCCGGGCCAGGGCCACCCGCTGCTGCTGGCCGCCGGAGAGCTGGCCGGGATGCTTGGCACTCTGGTCGGCGATGCCGACGCGCTCGAGCA
GCCGTTCGGCGGTCTCGACGGCATCCGCCCGGCTCCAGCCGCGCACCTTCATGGGGGCCAGGATGACGTTGTCGAGCACGCTGAGATGGGGAAAGAGGTT
GAACTGCTGGAACACCATGCCGACCTCGGTGCGGATCTTCTGCAGCGCCCTGGCACTCTTGCCGTCGGGCAGCAACTCCTTGGCGTCGACCTCGATATGG
CCCTGCTGGAACTCCTCGAGGCCGTTGATGCAGCGGATCAGGGTCGACTTGCCGGAGCCGCTGGCGCCGATCACCACTACCACCTCGCCGGGCTGGACCT
CGAGGTCGATATCCTTGAGCACGTGCAGGTTACCGAAGTGCTTGTTGACCCTCGCCATGCGCACGATGGGCGTCGTGGTCGTCTTGCTCTGTGTCATGCG
GTGGTTCCCCTAGCAGTCTCGTGGACCTTCACTGGCCGACCAGGCCGCGGCGCTCGAGCAGGCGCAGCGCGATGGAGAGGCTCCAGGTGATGGCCAGGTA
CATCAGGGCCACCATGAAGTAGACCTCGAGGGCGGTGAAGGTGGTGGCGATGTAGATCTGCCCCTGGCGCACCAGTTCGCCGACGCCGATCACCGAGAAC

>Halomonas heilongjiangensis DSM 26881 [NZ_PNRE01000019.1:c54456..54561], dnaA> ig dnaN>, 106 bp

WebSkew GCmin pos. 137067 (GCmin<>oriC1 30.1 kb; GCmin<>oriC2 6.7 kb; oriC2<>oriC1 23.4 kb)

>Halomonas korlensis CGMCC 1.6981 [NZ_FPBP01000003.1:166934..167433], <rpmH oriC1 dnaA>, 565 bp

CGGCACGCTCACGCTGCTCGCTACCGGTGATGGCCGGGGCATCGGGCACGGTGGTGGCTTGCCCGTAATCCTGATTCCATTGCACTACCAGCAAGTAAGC
GAGCAGCGCCAGGGGAACCAGCAATAGGAGTCGTTTTAGGTCCATGAAGAATCTGCCCGATGGGAGGTTAACATTTCGATAGCGGCATAGCGGCTGCCTG
CCTGATAGACGCAAAAGCCTGCCGAGGGCAGGCTGTGTGTCGGTCGCGACGATGCTTCACCTGGCGCGCTAGCAAACAGGCGGTTCCGGACGAGGGCCGT
CATACAGACTCCATGAGGCTGTCGAGGGGCGTGACTGAGACGTATCGGTTGCGGTGGCAGCACGAGGCATTTCGCTTCAGGTCGTTATCGATGACGCATT
GCGTACATCTCGCTCAAGGCGTCGCCACATGCCGTGGATCTGACGGTGCAGGGTCTCATTGTCCAGCTCCTGGACCCCTCGCTTGGCCAATACGACGATA
TCCACGGCGGGCAAACGGTCTTGTTGCAAACGCACGGATTCGCGAACCAACCGCTTGAAGCGGTTGCGATCGACTGCGCGGCGAACGTTCTTCTTGCTCA
CAATCAATCCGAGACGAGGATGGCCCAACGCATTGGGCATGGCCAAGGCCATCAACCCCTTTCCATGAATCTTGTAAACAGCCTTGTCGAAAACAGTCCG
ATACGCCTCGGCGTTCAACAAGCGAAGACACCGAGGAGAGCTCTGATGAGGCACTAGTGAGCCGCGTTCAGGCGCTTAGACGCTTGCGACCCTTGGCGCG
ACGGCGTGCCAGAACGGCACGACCATTCTTGGTGGCCATGCGAGCACGAAAACCGTGCACACGCTTGCGCTTGAGAACGCTAGGTTGAAAAGTGCGTTTC
ATAACTGGTTATTCCCACGTGATTTTTTGGACAAATAGGCAAATTCGAGACATCTGAACCCCCATCGGGGAACCCGGGGTGTTTGACTCAAGACCGGAAA

TTGTAGTGATTTCGCCGCTCCGGTGCAATTCTTCTTGTCGATAGACTCAAGGGAAGCCGGAGGAAACAGTGACTAATGCTGACGGTGGATGGCTGGCTCG
GCATCCGGATGAGGGGGTGTCGTGGGCAGTGTTGGGGATAGAATCTGTTGATAAGCTTAAAAAGTATAGTTTTATCAATAAATTAATGTGTTAATCGTTT
**TGTGAAGAA**AGGTGTGAGTATAAAGCGGCTATTGCCATGATGGCGCGGAACGATGAAATGGCTTCTTTATCGAGTTTAGTGCACCTCTATGAATGTGCTT
ATCGGTTGCGTTAGCGTGCGTTGGCTAGAAGAGTCCGAATGTCGCGGGTAACTGATATGTCGAGGATGGCGTTTTCCACAGTTCGTCGCACAGGCCGAGC
TCGTGTGGATAACTTTGCTGCTGGACGATACAATACCCGGCCGTTTTCCGATCAATGGTGAGGTCGCGTGTCGCTTGCGCTTTGGCAACAATGTCTGGAC

TCCCTACAGGATGAATTGAGCTCACAACAGTTCAATACCTGGATCCGCCCCCTGCAGGCGGAAGAAGGCGAGGCCAATGAGCTGTGTCTGCTGGCGCCGA
ACCGGTTTGTCCGTGACTGGGTTAGCGATAAGTATGCCAAGCGTATCAGCGAGTTAATGCGCAATCTGTCGCCTGCCAAACCCCCCAAGGTGGTGTTGAC
AGTGGGCAGTCGCCGTCGAGCCACGACGCCCCAGCCCCGAGAACTCGGCAACCCGGTGTCCGCGACCCCACCAGCCCAGCAACAGCCCGGCGCGCCCTCG
GTGCGAACACCGAAACGGGGCTCAAGCGACGACGAGCGTGAGATAAACGTATTGCGTGAAGAGCAGGATGGGGCGCGTCGACCCGGAGAACGGCAGGTGC
AGGTGGAAGGAGGCCTCAAGCACCACAGTGGGCTCAATCCCAATTTCACGTTCGAGACTTTCGTCGAGGGTAAATCGAATCAGCTTGCCAGAGCGGCTTC
CCGTCAGGTTTCCGAGAACCCCGGCGGTGCTTATAACCCCCTGTTCCTTTACGGGGGCGTGGGCCTGGGCAAGACGCACCTCATGCATGCCGTGGGCAAC
GCCCTGTCCACCCGGCGTGAGAACGCGCAAGTGATCTATCTGCATTCCGAGCGTTTCGTGGCCGATATGGTCAAGGCACTCCAGCTCAACGCCATCAATG
ATTTCAAGCGCTTCTATCGTAGTGTCGATGCCCTGCTGATCGACGATATCCAGTTCTTTGCCGGCAAGGAGCGGTCCCAGGAAGAGTTCTTCCACACCTT
CAATGCGTTGCTCGAAGGTGGCCAGCAAATGATCCTGACATCGGATCGTTACCCGAAGGAAATCAGCGGTGTTGAAGAGCGTCTCAAGTCGCGCTTCGGA
TGGGGCCTGACGGTGGCCATCGAGCCTCCCGAGCTCGAGACGCGCGTGGCGATCCTGATGAAGAAAGCGGATCAGGCGAAGGTCGACCTGCCCAACGATG

>Halomonas korlensis CGMCC 1.6981 [NZ_FPBP01000003.1:143509..144008], <mnmG oriC2 <COG1126 (glnQ), 546 bp

CATGCCCTCGATGGAGCGCACCACCTGAAGCTGGACATCGAAGGGCAGCGAGGTCGAGATGCCGTTGGGGTAGAGCTCATGCGTCTCCAGCCCCTCGGGT
TCGACGAAGATCTGGTGGCTCGTCTTGTCGGCAAAACGATGGACCTTGTCCTCGATCGACGGGCAGTAGCGGGGCCCGACGCCCTCGATGACGCCCGAGT
ACATGGGTGAGCGCTCGAGGTTGGCCAGGATGATCTCGTGGGTGCGCTCATTGGTATGGGCGATATGGCAGCTCACCTGGCGAGGGTGCATGGCCTGCGA
ACCGCGGTAGGACATCACCGGCGTGGGGCTGTCGCCCGGCTGCTCCTCGAGTCGCGACAGGTCGAGGCTACGGGCATCCAGGCGCGGCGGCGTGCCGGTC
TTGAGGCGATCGACCCGGAAGGGCAGGGCGCGCAGCCGCTCGGCAAGAGCGTTGGAGGGCGCATCACCGGCGCGTCCACCGCGACTGCGATCGAGCCCGA
TATGGATGACCCCGCCGAGGAAGGTTCCGGTACACAGCACCACGCTTTCGCCATGAAAGCGGATCCCGGTTTCGGTGATTACGCCCCGCACGGTATCGCC
GTCGACGATCAGGTCCCCGGCTGCTTGCTGAAAGAGCGTGAGGTTGGGCTGGTTTTCCAGCATGGCGCGAATTGCTGCCTTGTAGCGGACGCGATCGGCC
TGGGCCCGTGTCGCCCGAACCGCGGGACCCTTGCGGGCATTGAGCACGCGAAACTGGATGCCGCCCTGGTCGGTGGCAAGCGCCATGGCCCCTCCCAGCG
CATCGATCTCCTTGACCAGATGGCTCTTGCCGATCCCGCCGATAGCGGATTGCACGACATCTGACCCAGCGTCTCGATGTTGTGCGTCAGCAGCAGGGT
CTGACAGCCCATACGAGCGGAGGCCAGCGCGGCTTCGGTTCCCGCATGGCCGCCGCCAATGACGATGACGTCGAAGCGGTCGGGGTAATCCAAGGTACAC

CTCGTCGGGCGCAGCGGCGCCCGTGTTGCGATAAGTCAGTGATGAAATCAGCCCGGCAGTATAAACCTCCTGTGGGTAAACGTCAGGGTTTTCCACATCG
CGTTTCGAAGGTTCCTCAAGAGACGCTGCTCTATTATTAAATTAAACAATATATAAAGAGAAGTTCTGTTGTTGTTGTTACTACTGGTGTGGACAAAAAC
**CGTGGATAA**GCAACTAAATCCATTATTTAACCAGCAGGTTGCATTGTTGACTTCTGGGCGGATAAGCAGCGAATGAGCTGCGTACGGCCTGCCGGGAAGCT
GTGGATGAAAAGCCGCTTTGTCCACCGCAGCAAAACAGCACCGCTTGTCCACCGTCGAGTACCTGGCTTGTCACCGCAGTCGTGAACACCCGCTGATCGA
ATGAAGGAATCCGCGCCTCATCACGGTTGCAGAGCGATAAGTCGACGAAAGACGATTAAGTTATCCACAGCGCTAAAAAGCATGAAAAAGGCCCTGTCGA

TAACGGCAGGGCCTTGGTCAGCGGAGAAATAACGTGGTGTCAGTGTTTCAGGACCCGGGAAAGAAAGCTTCGGGTACGCTCGTTACGCGGCGCATCGAAG
ACCTGTTCGGGCGGCCCTTCCTCGGCAATCTCGCCCTGGTGGATATACACGACGCGATCGGCGACTTCACGGGCGAAGCCCATTTCGTGGGTGACGATCA
TCATGGTCATGCCTTCCTTGGCAAGTTCGCGCATGGCATCGAGTACCTCGCCGATCATCTCCGGGTCCAGGGCCGAGGTCGGCTCGTCGAAGAGCATCAG
GCGGGGTTCCATGGCCAGCGCCCTGGCCAAGGCCACACGCTGCTGCTGGCCCCCGGAGAGCTGGGTGGGGTATTTGGCCGCCTGATCGCTGATATCGACG
CGCTCCAGCAGACGCTCGGCAGTCTCGGTGGCGTCCACGCGATTCCAGCCTCGCACCTTCATCGGTGCCAGCGTGACATTGTCGAGCACGCTCAAGTGAG
GGAAGAGATTGAACTGCTGGAAGACCATGCCGACTTCGGTACGGATCTTCTGCAATGCCTTGGTCGCCTTGCCATGGGGCACCAGATCGTTGCCATCGAC
CTGAATATGCCCTTGCTGGAACTCTTCCAGGCCGTTGATGCAGCGAATCAGCGTCGACTTGCCCGAACCGCTGGCGCCGATGATGACCACAACCTCGCCG
GAGGAAACGGCAAGATCGATGTCCTTCAGCACGTGGAGGCTGCCGAAATGCTTGTTGACCTTGTCCAGGCGCACGATGGGGTCGGCGCCGGAAAGGGACT
CGTCATGGTTCATGTCGCGCATTCCTTATTGTCCGACCAGGCCTTTGCGCTCGAGCATGCGCAGCGCAAACGAGAGGGTGAGGGTGATCGCCAGATAGAG
CATGGCGACCATCAGGTAGACTTCCAGGGCGTTGAAGGTGGTGGCGATGTAGACCTGGCCCTGGCGCACCAACTCACCGACTCCGATCACCGAGAACAGC

>Halomonas korlensis CGMCC 1.6981 [NZ_FPBP01000003.1:168877..168950], dnaA> ig dnaN>, 74 bp

< page 1 | #041 | Halomonas sulfidivorans MCCC 1A13718

WebSkew GCmin pos. 4535424 (GCmin<>oriC1 37.5 kb; GCmin<>oriC2 10.7 kb; oriC2<>oriC1 26.7 kb)

>Halomonas sulfidivorans MCCC 1A13718 [NZ_CP053383.1:4572639..4572976], <rpmH oriC1 dnaA>, 472 bp

TGGGGCAGGGCAGCGTAAACGATGTCACCTCCCTGGGGATCGATGCGGACATCGAGCACATCGGTCACGACGGCGATCAGATCGCGAGAGCCACTCGTCT
GCTCGGTCTCGGTCGGCATGTCGCCGGCCATCTGCCGCTCCGCCTGAGGCGCGGCGGGCACGGCCAGGCCATCCTCGTCGGCTGGCTGCTGTTCATCGGG
TGCCGGTGCGGTGATCGCGGGAGCATCGGGCTCAGGAGCAACCTGGCCATAATCCTGATTCCACTGCACGACAAGCAAATAGGCGAGCACGGCCAGTGGG
ATCAGCAATAGGAGTCGTTTTACGTCCATGAGGGTTCTGCCCGGTGGGTAGTGAACATGCCGCTGGAACGAAATCCGCTGTCATGTAGGAAAGGTCGATA
GCCTGACGAGACAGGCTATCGAGGCGGAGAAAGACTCCGTATCCGGTCAGGAGGCCATCGACTTTCGGGCGTCACGCTCCAGGCGACGCCACATGCCGTA
GAACTGGCGATGCAGCGTTGCTCTGTCCAGCTCGTTCGCGCCACGCCGAGCCAGCAGGACGATGTCGACGGCGGGTAGCGAGCACTGCTGCAGGCGAATG
GATTCTCGAGCGAACCGCTTGAGGCGATTGCGATCCACAGCCTTGCGCACGTTCTTCTTGCTGATGACCAGGCCAAGGCGGGGGTGGTCCAGTCCATTGG
GGCACGCCAGGGCCATCAGCCCCTTGCCATGTACCTTGAATGAAGCGTGATCGAAAACGTATCGATAATCCCCGGCAGTCAACAAGCGCAGCTGCTTGGG
GAAGCCATGACCGGACACGCGCAATCCGCTATCAGGCGCTCAGACGCTTGCGGCCCTTGGCGCGGCGGCGAGCCAGAACGGCGCGACCATTCTTGGTGGC
CATGCGCGCACGGAAACCGTGGACACGCTTGCGCTTCAGAACGCTGGGTTGAAAGGTACGTTTCATAACACTCGATTCCCACGTGGTTTATTGACGGAAC

GTCTTACAGGAATGCCCTGGCTCCCATCAGGAACCAGGGGCGGATTCGGTTCAAGACCGGAAATTGTAGAGACTTCGGCTGTCCGACGCAATTGTTCCGC
TCGATCCGAGGGGCTGGCGCGGTGGATAGGCGGGCAAGATCGCGGGTGCTGGCGCCATGCCGGCTCGGCATACGCTGTGGTCAATACTGGGGACGGTCAC
**TGTTGATAA**GTCCTGACTTTAATGCCTAATCAACGAGGTAGGCAGTGCTCAGGAGTGTGGAAAAACCGTTCGCAAGTGCTTGCTGCCAGAACTTGCCCCC
AATTTGGCGACATGATGCGATGTGTCCTTCTTCGCCGGCGTTTTCCACAGGGGTGGTGGGTGTCGAACCGCATGTGGATAACCCGTCGGGCGAAGGATAC
AATGTCCGGTCATGCATCAACGGGTATGGTGAGGTCGAGTGTCTCTTGCTCTCTGGCAACAGTGTCTGGATTACCTGCAGGACGAGCTGAACGCACAGCA

GTTCAATACCTGGATTCGCCCGCTTCAAGCCGAAGAGGGGGAAGCCAGCGAGTTGCGTCTGCTGGCGCCCAATCGCTTCGTCCGCGACTGGGTTGGTGAC
AAGTATGCCAAGCGCATCAATGAGCTGCTCAGGGACCTGGCGCCTGCCAAGCCGCCGCGGGTAGTGCTGACAGTCGGCAGCCGCAAGGCGGCGGCGATCA
AGCCACAGCCGCGCGACCTGGGGACGCCGGTGTCGGCCAGCCCCCATTCCCCCCTGGCCCCGGCGGTGCAGGTACCGCCGCGAGGTGATATCGGCGACGA
GCGCGAAATCGACCAGCTGCGCGAGGAAGGAGCTACGGCGCGGCGTTCCGGCGGCGATCGCGAAGTGCAGGTCGAGGGCAGCCTGAAGCACAACAGCGGG
CTCAACCCCGGCTTTACCTTTGAAACCTTCGTCGAGGGCAAGTCCAACCAGCTTGCCCGCGCCGCGTCGCGGCAGGTGTCGGAAAATCCCGGTGGCGCCT
ACAACCCGCTGTTCCTCTACGGCGGCGTGGGCCTCGGCAAGACTCACTTGATGCACGCCGTGGGCAATGCCTTGGCGGCCCGGCGCGAAAACGCCCGTGT
CGTCTATCTGCACTCCGAGCGCTTCGTGGCCGACATGGTCAAGGCGCTACAGCTCAACGCGATCAACGACTTCAAGCGTTTCTATCGCAGCGTCGATGCG
CTGCTGATCGACGACATCCAGTTCTTTGCCGGCAAGGAGCGCTCCCAGGAGGAGTTCTTCCATACGTTCAATGCCTTGCTCGAAGGCGGTCAGCAGATGA
TCCTCACCTCGGACCGCTATCCAAAGGAGATCAGCGGCGTCGAGGAGCGGCTCAAGTCGCGCTTCGGCTGGGGGCTGACCGTGGCCATCGAGCCTCCCGA
GCTGGAGACCCGGGTCGCCATCCTGATGAAGAAGGCAGACCAGGCCAAGGTGGATCTGCCCCACGATGCCGCCTT

>Halomonas sulfidivorans MCCC 1A13718 [NZ_CP053383.1:4545925..4546424], <mnmG oriC2 <COG1126 (glnQ), 536 bp

CGTTCTCCAGCCCCTGGATCGAGCGCACCACCTGCAGCTGCACGTCGAAGGGCAGCGAGGTGGAAATGCCGTTGGGGTAGAGCTCATGAGTATCGAGGCC
CTCGGGTTCGATGAAGATCTGATGGCTCGCCTTGTCGGCGAAGCGGTGCACCTTGTCCTCGATCGACGGACAGTAGCGCGGCCCCACTCCCTCGATCACG
CCGGAGTACATCGGCGAGCGATCGAGATTGGCGAAGATGATCTCGTGGGTGCGCTCGTTGGTGTGAGCGATGTGGCAGCTCACCTGGCGCGGATGCATGT
CGCGGCGGCCCAGGTAGGACATGACCGGAGTCGGGCTGTCGCCGGGCTGCTCCTCGAGCACGCTGAAATCCACGCTCTTGGCATCCAGCCGCGGCGGCGT
GCCGGTCTTGAGCCGGTCGACCCGGAACGGCAACGCACGCAGGCGCTGGGCGAGGGCATTGGAGGGGGGATCCCCGGCGCGGCCGCCGCGGCTCTGCTCC
AGACCGATATGGATCACCCCGCCGAGGAAGGTGCCGGTACACAGCACCACCGTTTCGGCACGGAAGCGAATACCGGTCTCGGTGACCACCCCGCGTACCG
TGTCGTTTTCCACCTGCAGATCTCCCGCCGCCTGCTGAAAGATCGTCAGGTTCGGCTGGTTCTCCAGCATGCCACGGACGGCGGCCTTGTAACGCACGCG
GTCGGCCTGAGCGCGGGTGGCACGCACGGCGGGCCCCTTGCGCGCGTTGAGCACGCGGAACTGGATGCCGCCGAGATCGGTGGCCAGGCCCATGGCGCCG
CCCAGCGCATCGATTTCCTTGACCAGGTGGCTCTTGCCGATGCCGCCGATGGCGGGATTGCACGACATCTGCCCGAGGGTCTCGATGTTGTGGGTCAGCA
GCAGGGTCTGACAGCCCATGCGGGCGGAGGCCAGAGCGGCTTCGGTTCCCGCATGGCCGCCGCCGATGACGATGACGTCAAAGCGGTCGGGATACTCCAA

GGTGCACCTCGTTGCGCCAGCGCGCGGATGTTGCCAGTGACGAAAAACAGCCTGCCAGTATAAACGTCCTGTGGGTAAGCGTCAGGGTTTTCCACACCGT
CCTTGTCCAAGCCTCCCACTTTCCCTCCAGATAACCAAAACAGAATATCTAAAAAGAAAGAAGTTCTGTTGTGGTTGTGACTACTGGTGTGGACAATATC
**TGTGGATAA**GCCATAGAAGATATTTAATTCCAATGGCTTGCTTTGTTAACTTCTATGCGGAGAAGAGGGTGACGAGCTGAGGGTTGCCGGTGCGGGGGCT
GTGGATGAATGCCGGCCTTTTCCACAGCCCGAACGCTACACCCTTTTTCCACCGTTGCCTACCTGGCTTGTCACCGCGACTGTGAACAGTCACGCGAAGT
GAACGAAAAGGCGGTAACCATGCCGCTTGCAGCGACGAATCGTGGGTGGAGAAAAAATATTTATCCACAGGGCGAAAAAATGCCCTGTCGAGTTCGACAG

GGCTTGCCAGGGCGGAGGGCAGCGGCTTTCTGGAGCTCAGTGCTTGAGCACCCGCGACAGGAAGCTCTGGGTGCGCTCCTGCCGGGGGGAGTCGAAAAGC
GTGTCCGGCGCCGCCTGCTCGACGATCTCGCCGCGATCGATGAAAACCACGCGATCGGCGACTTCGCGGGCGAAGCCCATCTCGTGGGTGACGATGATCA
TGGTCATACCCTCACGGGCAAGTTCGCGCATGGCATCGAGCACTTCGCCGATCATCTCCGGATCGAGTGCCGAAGTGGGTTCGTCGAAGAGCATCAGGCG
AGGCTCCATGGCCAGTGCCCTGGCCAGGGCGACGCGCTGCTGCTGCCCTCCGGAGAGTTGCGACGGGTACTTGTCGGCCTGGTCGGCGATACCGACGCGT
TCGAGCAGTCTATAGGCGGTCTCGGTGGCATCGGCCCGGCTCCAGCCGCGTACCTTCATGGGGGCCAGCGTGACGTTGTCGAGTACGCTGAGATGCGGGA
ACAGGTTGAACTGCTGGAACACCATGCCGACTTCGGTGCGGATCTTCTGCAGTGCGCGGCCGCTCTTGCCGTCCGGGAGCAGCTCGTTGCCATCGACCTC
GATGTGGCCGGCCTGGAACTCCTCGAGGCCGTTGATGCAGCGAATCAGGGTGGACTTGCCCGAGCCGCTGGCGCCGATCACCACCACCACCTCGCCGGAG
GCGATGGCCAGATCGATATCCTTGAGCACGTGCAACTCGCCGAAGTGCTTGTTGACCTTGTCCATGTGAACGATGGGGACAGCGTCGGCGTGTGTATCGT
CTCGGCTCATGTCGCGCTCCTTATTGTCCGACCAGGCCCTTGCGCTCGAGCAGGCGCAGGGCAAATGACAGGCTCAGGGTTATCGCCAGATAGAGGAGGG
CGACCATCAAGTAGACTTCCAGGGCGTTGAAGGTGGTGGCGATGTAGACCTGGCCCTGACGTACCAGCTCGCCAACTCCGATCACCGAGAACAGCGAGGT

>Halomonas sulfidivorans MCCC 1A13718 [NZ_CP053383.1:1574..1636], dnaA> ig dnaN>, 63 bp

WebSkew GCmin pos. nd (GCmin<>oriC1 nd; GCmin<>oriC2 nd; oriC2<>oriC1 26.7 kb)

>Halomonas antri Y3S6 [NZ_JAHYCA010000007.1:c44361..44860], <rpmH oriC1 dnaA>, 473 bp

GGGGCAGGGCAGCGTAAACGATGTCACCTCCCTGGGGATCGATGCGGACATCGAGCACGTCGGTCACGACGGCGATCAGATCGCGAGAGCCACTCGTCTG
CTCGGTCTCGGTCGGCATGTCGCCGGCCATCTGCCGTTCCGCCTGAGGAGCGGCGGGCACGGCCAGGCCATCCTCGTTGGCTGGCTGCTGTTCATCGGCT
GCCGGTGCGGTGATCGCGGGAGCATCGGGCTCAGGAGCAACCTGGCCATAATCCTGATTCCACTGCACGACAAGCAAATAGGCGAGCACGGCCAGTGGGA
TCAGCAATAGGAGTCGTTTTACGTCCATGAGGGTTCTGCCCGGTGGGTAGTGAACATGCCGCTGGAACGAAATCCGCTGTCATGTAGGAAAGGTCGATAG
CCTGACGAGACAGGCTATCGAGGCGGAGAAAGACTCCGTATCCGGTCAGGAGGCCATCGACTTTCGGGCGTCACGCTCCAGGCGACGCCACATGCCGTAG
AACTGGCGATGCAGCGTTGCTCTGTCCAGCTCGTTCGCGCCACGCCGAGCCAGCAGGACAATGTCGACGGCGGGTAGCGAGCACTGCTGCAGGCGAATGG
ATTCTCGAGCGAACCGCTTGAGGCGATTGCGATCCACAGCCTTGCGCACGTTCTTCTTGCTGATGACCAGGCCAAGGCGGGGGTGGTCCAGTCCATTGGG
GCACGCCAGGGCCATCAGCCCCTTGCCATGTACCTTGAATGAAGCGTGATCGAAAACGTATCGATAATCCCCGGCAGTCAACAAGCGCAGCTGCTTGGGG
AAGCCATGACCGGACACGCGCAATCCGCTATCAGGCGCTCAGACGCTTGCGGCCCTTGGCGCGGCGGCGAGCCAGAACGGCGCGACCATTCTTGGTGGCC
ATGCGCGCACGGAAACCGTGGACACGCTTGCGCTTCAGAACGCTGGGTTGAAAGGTACGTTTCATAACACTCGATTCCCACGTGGTTTATTGACGGAACG

TCTTACAGGAATGCCCTGGCTCCCATCAGGAACCAGGGGCAGATTCGGTTCAAGACCGGAAATTGTAGAGACTTCGGCTGTCCGACGCAATTGTTCCGCT
CGATCCGAGGGGCTGGCGCGGTGGATAGGCGGGGCAAGATCGCGGGTGCTGGCGCCATGTCGGTTCGGCATACGCTGTGGTCAATATTGGGGACGGGCAC
**TGTTGATAA**GTCCTAACTTTGATGCCTAATCAACTAGGTAGGCAGTGCCCAGGAGTGTGGAAAAACCGTTCGCAAATGCTTTTCTGCCGAACTTGGCTCC
CCATTTGGCGACATGATGCGATGCGTCTCTCTTGCCGGCGTTTTCCACAGGGACGGCAGGTGACGAACCGCATGTGGATAACCCGCCGGGCGAAGGATAC
AATGTCCGGTCATGCATCAACGGGTATGGTGAGGTCGAGTGTCTCTTGCTCTCTGGCAACAGTGTCTGGATTACCTGCAGGACGAGCTGAACGCACAGCA

GTTCAATACCTGGATTCGCCCGCTTCAAGCCGAAGAGGGGGAAGCCAGCGAGTTGCGTCTGCTGGCGCCCAATCGCTTCGTCCGCGACTGGGTTGGTGAC
AAGTATGCCAAGCGCATCAATGAGCTGCTCAGGGACCTGGCTCCTGCCAAGCCGCCGCGGGTAGTGCTGACAGTCGGCAGCCGCAAGGCGGCGGCGGTCA
AGCCGCAGCCGCGCGACCTGGGGACGCCGGTGTCGGCCAGCCCCCATTCCCCCCTGGCCCCGGCCGTGCAGGTACCGCCGCGAGGTGATATCGGCGACGA
GCGCGAAATCGACCAGATGCGCGAGGAGGGAGCTACGGCGCGGCGTTCCGGCGGCGATCGCGAAGTGCAGGTCGAGGGCAGCCTGAAGCACAGCAGTGGG
CTCAATCCCGGCTTTACCTTCGAAACCTTCGTTGAGGGCAAGTCCAACCAGCTTGCCCGCGCCGCGTCGAGGCAGGTGTCGGAAAATCCCGGTGGCGCCT
ACAACCCGCTGTTCCTCTACGGCGGCGTGGGCCTCGGCAAGACTCACTTGATGCACGCCGTGGGCAACGCCTTGGCGGCCCGGCGCGAAAACGCCCGTGT
CGTCTATCTGCACTCCGAGCGCTTCGTGGCCGACATGGTGAAGGCGCTACAGCTCAACGCGATCAACGACTTCAAGCGTTTCTATCGCAGCGTCGATGCG
CTGCTGATCGATGACATCCAGTTCTTTGCCGGCAAGGAACGCTCCCAGGAGGAGTTCTTCCATACGTTCAATGCCTTGCTCGAAGGCGGTCAGCAGATGA
TCCTCACCTCGGATCGCTATCCAAAGGAGATCAGCGGCGTCGAGGAGCGGCTCAAGTCGCGCTTCGGCTGGGGGTTGACCGTGGCCATCGAGCCTCCCGA
GCTGGAGACGCGGGTCGCCATCCTGATGAAGAAGGCAGACCAGGCCAAGGTGGATCTGCCCCATGATGCCGCATTCTTCATTGCCCAGAAGATCCGCTCC

>Halomonas antri Y3S6 [NZ_JAHYCA010000007.1:c71079..71578], <mnmG oriC2 <COG1126 (glnQ), 534 bp

CGTTCTCCAGCCCCTGGATCGAGCGTACCACCTGCAGCTGCACGTCGAAGGGCAGCGAGGTGGAAATGCCGTTGGGGTAGAGCTCGTGAGTATCGAGGCC
CTCGGGTTCGATGAAGATCTGATGGCTCGCCTTGTCGGCGAAGCGGTGCATCTTGTCCTCGATCGACGGACAGTAGCGCGGCCCCACTCCCTCGATCACG
CCGGAGTACATCGGCGAGCGATCGAGATTGGCGAAGATGATCTCGTGGGTGCGCTCGTTGGTGTGAGCGATGTGGCAGCTCACCTGGCGCGGATGCATGT
CGCGGCGGCCCAGGTAGGACATGACCGGCGTCGGGCTGTCGCCGGGCTGCTCCTCGAGCACGCTGAAATCCACGCTCTTGGCATCCAGCCGCGGCGGCGT
GCCGGTCTTGAGCCGGTCGACCCGGAACGGCAACGCACGCAGGCGCTGGGCGAGGGCATTGGAGGGGGGGTCACCGGCACGGCCGCCGCGACTCTGCTCG
AGGCCGATGTGGATCACCCCACCGAGGAAGGTGCCGGTACACAGCACCACCGTTTCGGCATGGAAGCGAATACCAGTCTCGGTGACCACCCCGCGCACCG
TGTCGTTTTCCACCAACAGATCTCCCGCCGCCTGCTGAAAGATCGTCAGGTTCGGCTGGTTCTCCAGCATGCCGCGGACGGCGGCCTTGTAGCGCACACG
GTCGGCCTGGGCGCGGGTGGCACGCACGGCGGGCCCCTTGCGCGCGTTGAGCACGCGGAACTGGATGCCGCCGAGATCGGTGGCCAAGCCCATGGCGCCG
CCCAGCGCATCGATTTCCTTGACCAGGTGGCTCTTGCCGATGCCGCCGATGGCGGGATTGCACGACATCTGCCCGAGGGTCTCGATGTTGTGGGTCAGCA
GCAGGGTCTGACAGCCCATGCGGGCGGAGGCCAGAGCGGCTTCGGTTCCCGCATGGCCGCCGCCGATGACGATGACGTCAAAGCGGTCGGGATACTCCAA

GGTGCACCTCGTTGCGCCAGCGCGCGGATGTTGCCAGTGACGAAAAACAGCCTGCCAGTATAAACGTCCTGTGGGTAAGCGTCAGGGTTTTCCACACCGC
CTTCGTCCAAGCCTCCCACTTTCCCTCCAGATAACCAAAACAGAATATCTAAAAAGAAAGAAGTTCTGCTGTGGTTGTGATTACTGGTGTGGACAATATC
**TGTGGATAA**GCCGTAGAAGGTATTTAATTTCAATGCTTTGTTTTGTTAACTTCTATGCGGAGAAGAGGGTGACGAGCTGAGGGTAGCCGGTGTGGGGGCT
GTGGATGAATGCTGGCCTTTTCCACAGCGCGAACGCTACACCTTTTTTCCACCGTTGCCTACCGGGCTTGTCACCGTGACTGTGAACAGTCGCGGTAAGC
GAACGAGAAGTCGACAACCACGCATCTGAGGCGGCAGATTCGGGTGAACAGAATTTTTTTTATCCACAGGCCAAAAAAATAGGCCCTGTCGTATTCGACA

GGACCTGTCAGACGGAAAAGGCGCTCTACGAGCTTCAGTGCTTGAGCACACGGGACAGGAAGCTCCTGGTGCGTTCTTGCTGCGGAGAGTCGAACAGCGT
GACCGGCGCCGCCTGCTCGACGATCTCGCCGCGATCGATGAAAACCACGCGATCGGCGACCTCGCGGGCGAAGCCCATCTCGTGGGTGACGATGATCATG
GTCATGCCCTCACGGGCAAGTTCGCGCATGGCATCGAGCACTTCGCCGATCATCTCCGGATCGAGTGCCGAGGTGGGTTCGTCGAAGAGCATCAGGCGGG
GCTCCATGGCCAGTGCCCTGGCCAGGGCGACGCGCTGTTGCTGCCCTCCGGAGAGTTGCGACGGATACTTGTCGGCCTGGTCGGCGATACCGACGCGTTC
GAGCAGCCTATAGGCGGTCTCGGTGGCATCGGCCCGGCTCCAGCCGCGTACCTTCATGGGGGCCAGCGTGACATTGTCGAGTACGCTGAGATGCGGGAAC
AAGTTGAACTGCTGGAACACCATGCCGACTTCTGTGCGGATCTTCTGCAGTGCGCGGCCGCTCTTGCCGTCCGGGAGCAGCTCGTTGCCGTCGACCTCGA
TGTGGCCGGCCTGAAACTCCTCGAGGCCGTTGATGCAGCGAATCAGGGTGGACTTGCCCGAGCCGCTGGCGCCGATCACCACCACCTCGCCGGAGGC
GACGGCCAGATCGATGTCCTTGAGCACGTGCAACTCGCCAAAGTGCTTGTTGACCTTGTCCATGTGCACGATGGGGTCAGCGTCGGCGTGTGTATCGTCT
CGGCTCATGTCGCGCTCCTTATTGTCCGACCAGGCCCTTGCGCTCTAGCAGGCGCAGGGCAAATGACAGGCTCAGGGTTATCGCCAGATAGAGGAGGGCG
ACCATCAAGTAGACTTCCAGGGCGTTGAAAGTGGTGGCGATGTAGACCTGGCCCTGACGTACCAGCTCACCGACCCCGATCACCGAGAACAGCGAGGTGT

>Halomonas antri Y3S6 [NZ_JAHYCA010000007.1:c42887..42949], dnaA> ig dnaN>, 63 bp

< page 1 | #043 | Halomonas sulfidoxydans MCCC 1A11059

WebSkew GCmin pos. 4491948/0 (GCmin<>oriC1 0.0 kb; GCmin<>oriC2 27.1 kb; oriC2<>oriC1 27.1 kb)

>Halomonas sulfidoxydans MCCC 1A11059 [NZ_CP053381.1:4491624..175], <rpmH oriC1 dnaA>, 462 bp

GCAGGGCGGCGTAGACGATGTCACCCCCCTGGGGGTCGATGCGAACGTCGAGCACGTCGGTCACCACGGCGACCAGATCGCGCGTGCCGCTCGACGGCTC
CGCCTCGGCCGGCATGTCGCCGGCCATCTCCTGTTCCGCCCGAGGCGCGGCGGGTACCGCCAGGCCATCCTCGTCAGGTGCCTGCTGCTCGTCGGCGGCC
GGTGCGGTGATGGCGGGAGCGTCAGGCTCAGGAGCAACCTGGCCATAATCCTGATTCCACTGCACGACCAGCAGGTAGGCGAGCACGGCCAGTGGAATCA
GCAATAGGAGTCGTTTTACGTCCATGAGGGTTCTGCCCGGTGGGTAGTGAACATGCCGCTGGAACGAAATCCGCTGCCATGTAGGAAAGGTCGATAGCCT
GCCGAGACAGGCTATCGAAGCGGAGTAAAGACTCCGTATCCGGTCAGGAGGTCATCGACTTACGGACGTCACGCTCCAGGCGGCGCCACATGCCGAAGAA
CTGGCGATGCAGCGTGGCCCGGTCCAGCTCGTTCGCGCCACGCCGAGCCAGCAAGACGATGTCGACGGCGGGTAGCGAGCGCTGCTGCAGGCGGATGGAT
TCACGAGCGAACCGCTTGAGGCGATTGCGATCCACGGCCCTGCGGACGTTCTTCTTGCTGATGACCAGGCCAAGGCGGGGGTGGTCCAGTCCATTGGGGC
ACGCCAGGGCCATCAGCCCCTTGCCATGTACCTTGAATGCAGCGTGATCGAAAACGTATCGATAATCCCCGGCAGTCAGCAGACGCAGCTGCTTGGGGAA
ACCATGACCGGACACGCGCAATCCGAGATCAGGCGCTCAGACGCTTGCGGCCCTTGGCGCGGCGGCGAGCCAGAACGGCGCGACCATTCTTGGTAGCCAT
GCGCGCACGGAAACCGTGGACACGCTTGCGCTTCAGAACGCTGGGTTGAAAGGTACGTTTCATAACACTGGATTCCCACGTGGTTTATTGACGGAACGTC

TTACAGGAATGCCCTGGCTCCCGCGATAGGAACCAGGGGCGGATTCGGTTCAAGACCGGAAATTGTAGAGACTTCGGCTGCCCGACGCAATTGTTCCGCT
CGAAGCCGTGGGGCTTGTCGTCGCCGCCGGCGGGGCGGGGGCGTGGCCTCGATACCGCGATGGCCAGTCGTCCGCTGTGGTCAATACTGGGGACGAGCAC
**TGTTGATAA**GTCGAAGCTTTAATTAATAAACAATGGTATAGGTCGTGTTCAGGGCTGTGGAAAAGCGTTTCGTAAATGCGCAGCGCTGGCGGCCTGCCGG
CCTGGCCTGCGCTTCCTGCGGCGCCGCCGTTTTCCACAGCACCCCGCAGGCGAGCCGCATGTGGATAACCCGTCGGGCGGAGGATACAATGGCCGGTCAT
GCATCAACGGGTATGGTGAGGTCGAGTGTCTCTTGCTCTCTGGCAACAGTGTCTGGATTACCTGCAGGACGAGCTGAACGCACAGCAGTTCAATACCTGG

ATTCGCCCGCTTCAAGCCGAAGAGGGGGAGGCCAACGAGCTGCGCCTGCTGGCACCCAATCGCTTCGTTCGCGACTGGGTTGGCGACAAGTATGCCAAGC
GCATCAACGAGCTGCTCCGGGACCTGGCGCCTGCCAAGCCGCCACGGGTGATACTGACGGTCGGCAGCCGCAGGGCGGCCGCCGTCAAGCCGCAGCCCCG
TGACCTGGGTGCCCCGGTGTCGGCCAACCCTCACTCGCCCTTGGCGCCGGCCGTGCAGGTGCCGCCGCGAGGCGACATCGGCGACGAGCGCGAAATCGAC
CAGATGCGCGAGGAGGGGGCCGCCGCCCGCCGTTCCGGTGGCGATCGCGAAGTGCAGGTCGAGGGCAGCCTGAAGCACAGCAGCGGCCTCAATCCCGGCT
TCACCTTCGAGACCTTCGTCGAGGGCAAGTCCAACCAGCTGGCCCGCGCCGCCTCGCGCCAGGTTTCCGAGAACCCGGGCGGTGCCTACAACCCGCTGTT
CCTGTACGGCGGCGTGGGCCTGGGCAAGACTCACCTGATGCACGCCGTCGGCAACGCGCTGGCGGCCCGGCGCGAGAACGCCCGGGTCGTCTACCTGCAC
TCCGAGCGCTTCGTGGCCGACATGGTCAAGGCACTGCAGCTCAACGCCATCAACGATTTCAAGCGATTCTACCGCAGCGTCGATGCGCTGCTGATCGACG
ACATCCAGTTCTTCGCCGGCAAGGAGCGCTCCCAGGAGGAGTTCTTCCACACTTTCAACGCCTTGCTCGAGGGCGGCCAGCAGATGATTCTCACCTCGGA
CCGCTACCCCAAGGAGATCAGCGGCGTCGAGGAGCGCCTCAAGTCGCGCTTCGGCTGGGGGCTGACCGTGGCCATCGAGCCGCCCGAGCTGGAAACCCGG
GTCGCCATCCTGATGAAGAAGGCGGACCAGGCCAAGGTGGACCTGCCGCACGACGCCGCCTTCTTCATTGCCCAGAAGATTCGCTCCAACGTGCGCGAGC

>Halomonas sulfidoxydans MCCC 1A11059 [NZ_CP053381.1:4464564..4465063], <mnmG oriC2 <COG1126 (glnQ), 537 bp

GTTTTCGAGTCCCTTGATCGAGCGCACCACCGCGAGCTGCACGTCGAAGGGCAGCGAGGTGGAGATGCCGTTGGGGTAGAGCTCGTGGGTATCGAGCCCT
TCCGGCTCGATGAAGATTTGGTGGCTCGCCTTGTCGGCGAAGCGGTGCACCTTGTCCTCGATCGACGGGCAGTAACGCGGGCCGACGCCCTCGATCACGC
CCGAATACATCGGCGAACGGTCGAGGTTGGCGAAGATGATCTCGTGGCTGCGCTCATTGGTGTGAGCGATGTGGCAGCTCACCTGGCGCGGATGCATGTC
GCGACGACCCAGGTAGGACATCACCGGCGTCGGGCTGTCGCCCGGCTGCTCCTCGAGCACGGCGAAATTCACGCTCTTGGCGTCCAGCCGTGGCGGGGTG
CCGGTCTTGAGCCGGTCGACCCGGAACGGCAGCGCGCGCAGGCGCTGCGCCAGGGCATTGGAGGGCGGGTCGCCGGCGCGGCCACCGCGGCTCTGATCCA
GGCCGATATGGATCACGCCGCCGAGGAAGGTGCCGGTACACAGCACCACCGTCTCGGCCCGGAAGCGAATGCCGGTCTCGGTGACCACCCCGCGCACCGT
GTCGTTTTCCACCAGCAGGTCGCCCGCCGCCTGCTGGAAGATCGTCAGGTTCGGCTGGTTCTCCAGCATGCCGCGGATGGCGGCCTTGTAGCGCACGCGG
TCGGCCTGGGCGCGGGTGGCGCGCACAGCCGGGCCCTTGCGGGCGTTGAGCACGCGGAACTGGATACCGGCGAGATCGGTGGCCAGGCCCATGGCGCCGC
CCAGTGCATCGATCTCCTTGACCAGGTGGCTCTTGCCGATCCCGCCGATGGCGGGATTGCACGACATCTGGCCGAGAGTCTCGATGTTGTGGGTCAACAG
CAGGGTCTGACAGCCCATGCGGGCGGAGGCCAGGGCGGCTTCGGTTCCCGCATGGCCGCCGCCGATGACGATGACGTCAAAGCGGTCGGGATACTCCAAG

GTACACCTCGTTGCACGTGCGTGCGGATATTGCCAAAGGGGGTGAAAAACAGCCTGCCAGTATAAACCGCCTGTGGGTAAGCGTCAGGGGTTTTCCACAT
CACGTTGGTCAGTACCGAGTGGAGCAGCTCACTAATCAATACAGAATGATTAAAAGAGAGAAGTTCTGTAGTGGTTGTGACTACTGGTGTGGACAAAAAC
**CGTGGATAA**GCGGAATTAATCCTTTGGTATCATCTGCTTGCGCTGTTGACTCTTGCGGTGAGAAAAGGGTGACGAGCTGAGGGCGGCCGGTGGTGAGCCT
GTGGATGACCGCGGCGCTTTTCCACAGGGCGGACACTACACCCTTTTTCCACCGCCGGCTACCCGGCTTGTCACCGCGACTGTGAACAGCCGCGTGAGAC
GCCCAGGAAGTCGACAACCATGCTGCCTGGAGCGGCAAATCGAGGATGAGGAAAAAATTCTTATCCACAGGCCAAAAAAATGCCCTGCCGAGTGCGACAG

GGCCTGCCGGAGTGGAGATGAGGCGACTTTTTGGGTCTCAGTGCTTGAGTACCCGGGACAGGAAGCTCTGGGTGCGCTCGTGCCGTGGAGAGTCGAAAAG
CGTGTCCGGTGCTGCCTGCTCGACGATCTCACCGCGATCGATGAAGACGACGCGATCGGCGACCTCGCGGGCGAAGCCCATCTCGTGGGTGACGATGATC
ATGGTCATGCCTTCCCGGGCCAGCTCGCGCATGGCGTCGAGGACCTCGCCGATCATCTCCGGATCGAGCGCCGAGGTGGGTTCGTCGAACAGCATCAGGC
GCGGCTCCATGGCCAGCGCCCGGGCCAGGGCGACGCGCTGCTGCTGCCCGCCGGAGAGCTGGCCGGGGTACTTGCCCGCCTGGTTGGCGATGCCCACGCG
CGTGAGCAGACGCTCGGCGGTCTCGATCGCGTCGGCGCGGCTCCAGCCGCGCACCTTCATCGGCGCCAGGGTGACGTTGTCGAGCACCCGGAGATGGGGA
AACAGGTTGAACTGCTGGAACACCATGCCGACCTCGGTGCGGATCCTCTGCAGCGCACGGCTGCTCTTGCCGTCGGGCAGCAGCTCGTTGCCATCGACCT
CGATGTGGCCGGCCTGGAACTCCTCGAGGCCGTTGATGCAGCGAATCAGGGTGGACTTGCCCGAACCACTGGCGCCGATCACCACCACCACCTCGCCGGG
GGCGACCGTCAGGTCGATGTCCTTGAGCACGTGCAGCTCGCCGAAATGCTTGTTGACCCGCTCCATGCGCACGATGGGCTGGGCCGCGGCGTGGGAAGTG
TCTCGGTTCATGTCGCGCTCCTTATTGTCCGACCAGGCCCTTGCGCTCGAGCAGGCGCAGTGCGAATGACAGGCTCAGGGTTATCGCCAGATAGAGGAGG
GCGACCATCAGGTAGACTTCGAGGGCATTGAAGGTGGTGGCGATGTAGACCTGGCCCTGGCGCACCAGCTCGCCGACGCCGATCACCGAGAACAACGAGG

>Halomonas sulfidoxydans MCCC 1A11059 [NZ_CP053381.1:1574..1637], dnaA> ig dnaN>, 64 bp

WebSkew GCmin pos. nd (GCmin<>oriC1 nd; GCmin<>oriC2 nd; oriC2<>oriC1 nd)

>Halomonas lactosivorans KCTC 52281 [NZ_QHGY01000015.1:15832..16331], <rpmH oriC1 dnaA>, 473 bp

TCTCGACGGCGATCAGGTCGCGGGAACTGTCCGACTGTTCCGACTCGGCCGGCATGTCGCCGTCCATCTCCTGCTCCGCCCGAGGCGTGGCGGGTACCGC
CAAGTCGTCGTCGTCTGGCGCTTCCTGCTCATCGGCTTCTCTTGCGGTGATGGCGGGAGCGTCGGGCTCAGGAGCAACCTGGCCATAATCCTGATTCCAC
TGCACGACCAGCAGGTAGGCGAGCACGGCCAGTGGAATCAGCAATAGGAGTCGTTTTACGTCCATGAGGGTTCTGCCCGGTGGGTAGTGAACATGCCGCT
GGAACGGGATCCGCTGCCATGTGAAGGGGGACGACAGCCTGCCGGGCAGGCCGTCGAGGCGGCGTTCGGTAGCGCTGTGGCCGTGTTGGCGCTGCTTGCT
CGCGCGCGCCGCTTCGGTCAGCGCTTCCCGAGGCATCGCATCCGGTCAGGAGGTCATCGACTTTCGTGCGTCACGCTCCAGGCGGCGCCACATGCCGGAA
AACTGGCGATGCAGCGTGGCCCTGTCCAGCTCGTTCGCGCCGCGTCGAGCCAGCAGGACGATGTCGACGGCGGGTAGCGAGCACTGCTGCAGGCGAATGG
ATTCACGAGCGAACCGCTTGAGACGGTTGCGATCCACGGCCTTGCGGACGTTCTTCTTGCTGATGACCAGGCCAAGGCGGGGGTGGTCCAGTCCATTGGG
GCACGCCAGGGCCATCAGCCCCTTGCCATGTACCTTGTATGTCGCGTGATCGAAAACGTATCGATAATCCCTGGCAGTCAGCAGGCGCAGCTGCTTGGGG
AAACCATGACCGGACACGCGCAATCCGCTATCAGGCGCTCAGACGCTTGCGGCCCTTGGCGCGGCGGCGAGCCAGAACGGCGCGACCATTCTTGGTGGCC
ATGCGAGCACGGAAACCGTGGACACGCTTGCGCTTGAGAACGCTGGGTTGAAAGGTACGTTTCATAACACTTGATTCCCACGTGGTTAATTGACGGAACG

TCTTACAGGAATGCCCTGGCTCCCGCGAGGAACCAGGGGCGTATTCGGTTCAAGACCGGAAATTGTAGAGACTTCGGCTGCCCGACGCAATTGTTCCGCA
CGCTGCCGTGCGCCCTGACCCGGCGGCCGCCGGGCGGAGCCGCGCCGAGGGCATCGCCAGGCGGCGTATGCCTGCTGTGGTCAGTATTGGGGATGGCTTC
**TGTTGATAA**GTTGGAGCTTTAATTATTAATCAAGGAGATAAAGGGTGCCCATAGGTGTGGAAAAACGTTTCGCAACCGGGCCGCCCGGAGGGCGAGCGCG
TGGCCGGTCGACGGCAGGCGAGCCTCCTTCCCGCAGGCGTTTTTCCACAGGCGTTTGCGCTGTCGAACCGCGTGTGGATAACCCGCCGGGCGAAGGATAC
AATGGCCGGTCATGCATCAACGGGTATGGTGAGGTCGAGTGTCTCTTGCTCTCTGGCAACAGTGTCTGGATTACCTGCAGGACGAGCTGAACGCACAGCA

GTTCAACACCTGGATTCGCCCCCTTCAAGCCGAAGAGGGTGAAGCCAACGAGCTGCGCCTGCTGGCGCCCAATCGCTTCGTTCGCGACTGGGTCGGCGAC
AAGTATGCCAAGCGCATCAATGAGCTGCTTCGGGACCTGGCGCCAGCCAAGCCGCCGCGGGTGGTGTTGACGGTGGGTAGCCGCAAGGCGGCGGTCAAGC
CGCAGCCCCGCGACCTGGGCGCCCCGGTCTCGGCCAACCCCCACTCTCCCTTGGCGCCGGCCGTGCAGGTGCCGCCGCGAGGCGACATCGGCGACGAACG
CGAGATCGACCAGCTGCGCGAGGAAGGCGGTGCGCCGCGTCGCTCCGGCGGCGACCGTGAGGTGCAGGTCGAAGGCAGCCTCAAGCACAACAGCGGGCTC
AACCCCAGCTTCACCTTCGAGACCTTCGTCGAGGGCAAGTCCAACCAGCTGGCACGCGCCGCCTCGCGGCAGGTCTCCGAGAACCCGGGAGGCGCCTACA
ACCCGCTGTTCCTGTATGGCGGCGTAGGGCTGGGCAAGACCCACCTGATGCATGCGGTCGGCAACGCCCTGGCGGCTCGGCGCGAGAACGCCCGCGTCGT
CTACCTGCACTCCGAGCGTTTCGTGGCCGATATGGTCAAGGCGCTGCAGCTCAACGCCATCAACGACTTCAAGCGCTTCTATCGCAGCGTCGACGCGCTG
CTGATCGACGATATCCAGTTCTTCGCCGGCAAGGAGCGCTCCCAGGAGGAGTTCTTCCATACCTTCAACGCCCTGCTCGAGGGTGGCCAGCAGATGATTC
TCACCTCGGATCGCTATCCCAAGGAGATCAGCGGCGTGGAGGAGCGCCTCAAGTCCCGCTTCGGCTGGGGGCTGACCGTGGCCATCGAGCCTCCTGAGCT
CGAGACCCGGGTGGCGATCCTGATGAAGAAGGCCGACCAGGCCAAGGTGGACCTGCCCCATGATGCCGCCTTCTTCATTGCCCAGAAGATCCGCTCCAAC

>Halomonas lactosivorans KCTC 52281 [NZ_QHGY01000013.1:12567..13066], <mnmG oriC2 <COG1126 (glnQ), 529 bp

TTCTCCAGCCCCTGGATCGAGCGCACCACCGCGAGCTGCACGTCGAAGGGCAGCGAGGTGGAGATGCCGTTGGGGTAGAGCTCGTGGGTGTCGAGCCCCT
CCGGCTCGATGAATACCTGATGGCTCGCCTTGTCGGCGAAGCGGTGCACCTTGTCCTCGATCGACGGGCAGTAGCGCGGCCCGACACCCTCGATCACGCC
GGAATACATCGGCGAGCGGTCGAGGTTGGCGAAGATGATCTCGTGGGTGCGCTCGTTGGTATGGGCGATATGGCAGCTCACCTGGCGCGGGTGCATGTCG
CGGCTGCCCAGGTAGGACATCACCGGCGTCGGCGTATCGCCCGGCTGCTCCTCGAGCTGCGCGTAGTCGACGCTTTTGGCGTCCAGCCGTGGCGGGGTAC
CGGTCTTGAGCCGGTCGACCCGGAACGGCAGCGCCCTCAGGCGACCGGCCAGCGCGTTGGAGGGCGGATCGCCGGCGCGACCGCCGCGGCTCTGCTCGAG
ACCGATGTGGATCACGCCACCGAGGAAGGTGCCGGTGCACAGCACCACCGTCTCCGCCAGCAGGCGAATGCCGGTCTCGGTGACCACCCCGCGCACGGTG
TCGCCCTCGACGATCAGGTCGCCGGCGGCCTGCTGGAAGAGCGTCAGGTTGGCCTGGTTCTCCAGCATGCCGCGGATAGCGGCCTTGTAGCGCACCCGGT
CGGCCTGGGCGCGGGTGGCGCGCACCGCCGGCCCCTTGCGTGCGTTGAGCACGCGAAACTGGATACCGCCGAGGTCGGTGGCCAGGCCCATGGCCCCGCC
CAGGGCATCGATCTCCTTGACCAGGTGGCTCTTGCCGATCCCGCCGATGGCGGGATTGCAGGACATCTGGCCGAGCGTCTCGATGTTGTGGGTCAGTAGC
AGGGTCTGACAGCCCATGCGGGCGGAGGCCAGAGCGGCTTCGGTTCCCGCATGGCCACCGCCGATGACGATGACGTCAAAGCGGTCGGGATACTCCAATG

TGCACCTCGTTGCGCTTGCGCGTGGAAATTGGCAAAGGGGCGAAAAACAAGCCTGCCAGTATAAACCGCCTGTGGGTAAGCGTCAGGGGTTTTCCACATC
GGGTTCGCCAGCGTGTCTGCCCGGTACCCTCACTAATCAAAACAGAATGATTTAAGAGAGAAGTTCTGTTGTGGTAGTGACTATTGGTGTGGACAAATTC
**TGTGGGTAA**GCGGAAAGATGTGATATTTATCAACAAGTTGAATTGTTGACTTTCGTGGTGAGAAGAGGGTGACGAGCTGAGGGCGGCCGGTGTCGAGCCT
GTGGATGAAACGGGGGCTTTTCCACAGAGCGAACGCTACACCGTTTTTCCACCGCCGGCTACCCCGCTTGTCACCGTGGCTGTGAACAATCCCCCGTAAA
GCTGGCGCATGCCCCAACCATGCGGTCTGTGGCCCGGAATGGGGGCGCTGGCGGATAATGTTATCCACAGGCAGAAAAAAGCCCTGGGGGAGTCCCGCCA

GGGCGTCGACCGAGAAAGGGAAGGGGGTCAGTGCTTGAGCACGCGGGACAGGAAGTTCTGCGTGCGTTCGTGACGCGGAGCGTCGAACAGCGCGTCCGGG
GCGGCCTGCTCGACGATCTCGCCCTGGTGGATGAAGACGACGCGGTCGGCGACCTCGCGGGCGAAGCCCATCTCGTGAGTGACGATGATCATGGTCATCC
CCTCGCGGGCCAGCTCGCGCATGGCGTCGAGGACCTCGCCGATCATCTCGGGATCGAGCGCCGAGGTCGGCTCGTCGAACAGCATCAGGCGTGGCTCCAT
GGCCAGCGCCCGGGCCAGGGCCACGCGCTGCTGCTGGCCACCGGAGAGCTGGCTGGGGTACTTGTCGGCCTGGTCGGCGATGCCCACCCGCGCCAGCAGG
CGCTCGGCGGTCTCGATCGCGTCGTCCCGGCTCCAGCCGCGCACCTTCATCGGTGCCAGCGTGACGTTGTCGCGCACCTTGAGGTGGGGGAAGAGATTGA
ACTGCTGGAACACCATGCCGACCTCGGTGCGGATCTTCTGCAGCGCCCGGCTGCTGCGGCCGTTGGGCAGCAGCTCGTTGCCGTCGACCTCGATATGACC
GGCCTGGAATTCCTCGAGACCGTTGATGCAGCGAATCAGGGTGGACTTGCCGGAGCCGCTGGCGCCGATCACTACCACCACCTCGCCGGGGGCGATCGTC
AGGTCGATGTCCTTGAGCACGTGCAGGTCGCCGAAGTGCTTGTTGACCTTGTCCATGCTTACGATGGGGGCGGCCGGAGAGGGGGAATTGCCTCGGTTCA
TGTCGCGCTCCTTATTGTCCGACCAGGCCCTTGCGCTCGAGCAGGCGCAGTGCGAATGACAGGCTCAGGGTGATCACCAGGTAGAGGAAAGCGACCATCA
GGTAGACTTCCAGGGCGTTGAAGGTGGTGGCGATGTAGACCTGGCCTTGGCGCACCAGCTCGCCGACGCCGATCACCGAGAACAGCGAGGTGTCCTTGAT

>Halomonas lactosivorans KCTC 52281 [NZ_QHGY01000015.1:17740..17807], dnaA> ig dnaN>, 68 bp

< page 1 | #045 | Halomonas bachuensis DX6

WebSkew GCmin pos. 163590 (GCmin<>oriC1 33.9 kb; GCmin<>oriC2 8.8 kb; oriC2<>oriC1 22.1 kb)

>Halomonas bachuensis DX6 [NZ_JAAQTO010000018.1:197244..197743], <rpmH oriC1 dnaA>, 485 bp

TGGGGCAGGGCGGCATAGACGATGTCCCCCCCCTGGGGATCGATGCGAACATCGAGCACATCGGTCACGACGGCAACCAGGTCGCGAGAACCGCTCGGCT
GCTCCGTCTCGCCCGGCATTTCGCTGGCCATCTCCTGCTCAGCACGGGGCGCAGCGGGAACCGTCAGGCCATCGTCGTCGGGTGCCTGCTCTTCACCGGG
GGGCGATGCGGTGATGGCCGGAGCGTCAGGCTCAGGAGCAACCTGGCCGTAGTCCTGATTCCACTGCACGACAAGCAGGTAGGCGAGCACGGCCAGTGGA
ATCAGCAATAGGAGTCGTTTTACGTCCATGAGGGTTCTGCCCGGTGGGTAGTGAACATGCCGCTGGAACGAGATCCGCTGTCATGTAGATGAAGACGATA
GCCTGCTAGGCAGGCTATCGAGGCGGAGAAAGGCTCCGCATCCGGTCAGGAGGTCATCGTTTTTCGGGCGTCACGCTCCAGGCGGCGCCACATGCCGAAT
AACTGGCGATGCAGCGTAGCCTTGTCCAGCTCGTTCGCGCCGCGTCGAGCCAGCAAGACGATATCGACGGCGGGTAGCGAGCACTGCTGCAGGCGTATGG
ATTCACGAGCGAACCGCTTGAGACGGTTGCGATCCACGGCCTTGCGGACGTTCTTCTTGCTGATGACCAGGCCAAGGCGGGGGTGGTCCAGTCCATTGGG
GCACGCCAGGGCCATCAGCCCCTTGCCATGTACCTTGAATGAAGCGTGATCGAAAACGTATCGATAATCCCCGGCAGTCAACAGACGCAACTGTTTGGGG
AAACCATGACCGGACACGCGCAATCCGCTGTCAGGCGCTCAGACGCTTGCGGCCCTTGGCGCGGCGACGAGCCAGAACGGCGCGACCATTCTTGGTAGCC
ATGCGAGCACGGAAACCGTGGACACGCTTGCGCTTCAGAACGCTGGGTTGAAAGGTACGTTTCATAACACTTGATTCCCACGTGGATTATTGACGGAACG

TCTTACAGGAATGCCCTGATTCCCGACAGGAACCAGGGACAGATTCGGTTCAAGACCGGAAATTGTAGAGACTTCGGCTGCCCGACGCAATTGTTCCGCA
TGCTGCAGCGAGTCGAGCCGTCGCCGCTGGTGGGCTGAACCGCAGTCTCGACCCCTTGGAGCCACGCTTCCCAACTGTGATTAACCTTGGGGACTCTCCC
**TGTTGATAA**GTTTTTTATTTGATTAATAAACAATATAGTAGGTGGCACCATGCTTTGTGGAAAAGCGTTTGGTAAGTCCTGCTCATGAGGGGTCAATGGC
GTAGCGAAGCCAAGGCACTCGCCAGGCTGTCGAGAGTCCAACCTCGAGCCGCTTTTCCACAGGCCTGTCGGGTGTCGAGCTGCATGTGGATAACCCGCCA
GCCGAAGGATACAATGGCCGGTCATGCATCAACGGGTATGGTGAGGTCGAGTGTCTCTTGCTCTCTGGCAACAGTGTCTGGATTACCTGCAGGATGAGCT

GAACGCACAGCAGTTCAATACCTGGATTCGCCCGCTTCAAGCCGAAGAGGGGGGAGCCAACGAGCTGCGCCTGCTGGCGCCCAACCGCTTCGTTCGCGAT
TGGGTTGGCGACAAGTATGCCAAGCGCATCAACGAACTGCTCCGGGACCTGGCGCCTGCCAAACCGCCACGGGTGGTACTGACAGTCGGTAGCCGCAGAG
CGGCGGCGGTCAAGCCGCAGCCGCGAGACCTGGGTGCGCCGGTATCGGCCAACCCTCACTCTCCGTTGGCGCCGGCCGTGCAGGTGCCACCGCGAGGAGA
TATCGGCGACGAACGTGAAATCGACCAGCTACGCGAAGAAGGGGCCGGCTCGGGGCGCCGCTCCAGCGGTGATCGCGAAGTGCAGGTCGAAGGCAGCCTG
AAGCACAACAGCGGCCTCAATCCCAGCTTCACCTTCGAAACCTTCGTCGAGGGCAAGTCCAACCAGCTGGCACGGGCCGCCTCGCGGCAGGTTTCCGAGA
ACCCCGGAGGCGCCTACAATCCTCTGTTCCTGTATGGCGGTGTCGGGTTGGGCAAGACCCACCTGATGCACGCCGTAGGCAACGCCTTGGCGTCGCGTCG
CGAGAACGCGCGGGTCGTCTACCTCCACTCCGAGCGCTTCGTGGCCGACATGGTCAAGGCACTTCAGCTCAACGCCATCAACGACTTCAAGCGTTTCTAC
CGCAGCGTAGACGCGCTATTGATCGACGACATCCAGTTCTTTGCCGGCAAGGAGCGTTCCCAGGAGGAGTTCTTTCATACCTTCAATGCGCTACTCGAAG
GTGGCCAGCAGATGATCCTCACCTCCGATCGCTATCCCAAGGAGATCAGCGGCGTCGAGGAGCGACTCAAGTCGCGCTTCGGCTGGGGATTGACGGTGGC
CATCGAGCCGCCGGAGCTGGAAACCCGTGTGGCTATCCTGATGAAGAAGGCCGACCAGGCCAAGGTCGACCTGCCCCACGATGCTGCCTTCTTCATCGCC

>Halomonas bachuensis DX6 [NZ_JAAQTO010000018.1:172126..172625], <mnmG oriC2 <COG1126 (glnQ), 530 bp

TTTTCCAGCCCCTTGATCGAGCGCACCACCTGCAGTTGCACGTCGAAGGGCAGCGAGGTGGAAATCCCGTTGGGGTAGAGCTCATGGGTCTCCAGCCCCT
CGGGCTCGATGAAGATCTGATGGCTCGCCTTGTCGGCGAAGCGGTGCACCTTGTCCTCGATCGACGGACAGTAGCGCGGCCCAACGCCTTCGATCACGCC
GGAGTACATCGGTGAACGATCGAGGTTGGCGAAGATGATCTCGTGGGTACGCTCGTTGGTATGGGCGATATGACAGCTTACCTGGTGCGGATGCATGTCG
CGCCGGCCAAGGTAGGACATGACGGGGGTCGGTGTATCGCCCGGTTGTTCCTCGAGCTGCGAGAAATCCACGCTTTTGGCGTCCAGCCGCGGTGGCGTAC
CGGTCTTCAGCCGGTCGACCCGGAACGGCAGCGCGCGCAGGCGCTCGGCCAGGGCATTGGAGGGCAGATCGCCGGCGCGACCGCCGCGGCTCTTGTCGAG
CCCGATGTGGATCACGCCACCGAGGAAGGTGCCCGTGCACAGCACCACCGTCTCGGCCAGGAAGCGAATCCCGGTCTCGGTGACCACGCCACGCACGGTG
TCGTCGTTCACCACCAGATCGCCAGCGGCCTGCTGAAAGATCGTCAGATTGGCCTGGTTCTCCAGCAAACCACGGATGGCTGCCTTGTAGCGCACGCGGT
CGGCCTGGGCCCGGGTGGCACGAACTGCGGGCCCCTTGCGTGCGTTGAGCACACGGAACTGGATACCGCCGAGATCGGTGGCCAGGCCCATGGCGCCGCC
CAGGGCATCGATTTCCTTGACCAGATGGCTCTTGCCGATCCCGCCAATGGCGGGATTGCAGGACATCTGTCCGAGAGTCTCGATGTTATGGGTCAGTAGC
AGGGTCTGACAGCCCATGCGAGCGGAGGCCAGAGCGGCTTCGGTTCCCGCATGGCCGCCGCCGATGACGATGACGTCAAAGCGGTCGGGATACTCCAAGG

TGCACCTCGTTGCGCTTGTGCGCAGATATTGCCTAAGGACGAAAATCAGCCTGGCAGTATAAACCTCCTGTGGGTAAGCGTCAGGGGTTTTCCACACCCG
CTTTCACGGAGCCCCCTTTCCCCTCACCTGGTTAAACAAAACAGAATGATTAAAAGAGAGAAGTTCTATTGTGGTTGTAACTACTAGTGTGGATAAAAAC
**CGTGGATAA**CCAAGAATATCATATACAAAACATGATCTTGAATTGTTGACTTATACGGTGAAAAAAGGATGACGAGCTGAGGGCGGCCGGTGACGAGCCT
GTGGATGAAAGGCATGTTTGTCCACAGAGCGAACGCTGCACCTCTTTTCCACCGCCGCCTACCGTACTTGTCACCGCGACTGTGAACAACCGGCCTAGGA
CATGACGTAGGCACCAACCATGCGCTCTGGGAGCAAGGGAAGGGTGTTTTCGAGAAAATCTTATCCACAGGCAAAAAAAAGCCCTGACGATGACGTCAGG

GCTTCGCAGGGGCGTGTCCGGGAAGCTGCTAGTGCTTGAGTACGCGAGAGAGAAAGCTCTGGGTGCGCTCGTGTCGAGGGGAGTCGAACAGCACGTCGGG
AGCGGCCTGTTCGACGATCTCTCCCCGATGAATGAACACCACCCGATCGGCGACTTCACGGGCGAAGCCCATCTCGTGGGTAACGATGATCATGGTCATG
CCCTCGCGAGCGAGCTCGCGCATGGCGTCGAGCACTTCACCGATCATTTCGGGATCGAGTGCCGAGGTGGGTTCGTCGAACAGCATCAGCCGCGGCTCCA
TGGCAAGCGCCCGGGCCAGGGCGACGCGCTGCTGTTGACCTCCGGAGAGCTGGGAAGGGTACTTGTCTGCCTGGTCGGCGATCCCGACACGTTCGAGTAG
ACGATTGGCGGTCTCGGTGGCATCGGCGAGGTTCCAGCCGCGCACTTTCATCGGTGCCAGTGTGACGTTGTCGCGCACGCTGAGATGGGGAAAAAGATTG
AACTGCTGGAATACCATGCCGACCTCGGTGCGGATCGTCTGCAGCGCACGGCTGCTCTTGCCATTGGGTAGCAATTCGTTGCCGTCGACCTCGATATGGC
CGGCTTGGAACTCCTCGAGGCCGTTGATGCAGCGAATCAGGGTCGATTTTCCGGAGCCGCTGGCGCCGATGACCACCACCACCTCACCAGGCCGGATCGT
AAGGTCGATATCCTTGAGCACGTGCAACTCACCGAAGTGCTTGTTGACCTTCTCCATTTGCACGATGGGGGCGGCATCGGAGGGAGTATTGTCTCGGTTC
ATTTCGCGCTCCTTCATTGTCCAACCAGGCCCCTGCGCTCGAGCAGGCGCAGTGCGAATGACAGGCTCAGGGTTATCGCCAGATAGAGGAGGGCGACCAT
GAGATAGACTTCCAGGGCGTTGAAGGTAGTGGCGATATAGACTTGGCCCTGGCGCACCAGCTCGCCGACACCGATGACCGAGAACAGCGAAGTGTCCTTG

>Halomonas bachuensis DX6 [NZ_JAAQTO010000018.1:199170..199236], dnaA> ig dnaN>, 68 bp

< page 1 | #046 | Halomonas diversa MCCC 1A13316

WebSkew GCmin pos. 4454080 (GCmin<>oriC1 36.8 kb; GCmin<>oriC2 11.7 kb; oriC2<>oriC1 25.2 kb)

>Halomonas diversa MCCC 1A13316 [NZ_CP053382.1:4490725..293], <rpmH oriC1 dnaA>, 486 bp

TGGGGCAGGGCGGCATAGACGATGTCCCCCCCCTGCGGGTCGATGCGAACATCGAGTACGTCGGTGACGACGGCAACCAGGTCGCGAGAACCGCTCGGCT
GCTCCGTCTCGTCCGGCATGCTGCCGGCCATCTCCTGCTCAGCACGGGATGCAGCGGGGACCGTCAGGCCATCGTCGTCGGGTGCCTGCTCTGCACCCGG
TGCCGGTGCGGTGATAGCGGGAGCGTCAGGCTCAGGAGCAACCTGGCCATAGTCCTGATTCCACTGCACGACGAGCAGATAGGCGAGCACGGCCAGTGGA
ATCAGCAATAGGAGTCGTTTTACGTCCATGAGGGTTCTGCCCGGTGGGTAGTGAACATGCCGCTGGAACGAGATCCGCTGTCATGTAGATAAGGACGATA
GCCTGCGCGGCAGGCTATCGAGGCGGAGAAAGGCTCCGCATCCGGTCAGGAGGTCATCGTTTTTCGGGCGTCACGCTCCAGGCGACGCCACATGCCGAAG
AACTGGCGATGCAGTGTAGCCTTGTCCAGCTCATTCGCGCCGCGCCGAGCCAGCAATACGATATCGACGGCGGGTAGCGAGCACTGCTGAAGGCGTATGG
ATTCACGAGCGAACCGCTTGATACGGTTGCGATCCACGGCCTTGCGGACGTTCTTCTTGCTGATGACCAGGCCTAGGCGGGGGTGGTCCAGTCCATTGGG
GCACGCCAGGGCCATCAGCCCCTTGCCATGTACCTTGAATGAAGCGTGATCGAAAACGTATCGATAATCCCCGGCAGTCAACAGACGCAGCTGCTTGGGG
AAACCATGACCGGACACGCGCAATCCGCTGTCAGGCGCTCAGACGCTTGCGGCCCTTGGCGCGACGGCGAGCCAGGACGGCACGACCATTCTTGGTAGCC
ATGCGAGCACGGAAACCGTGGACACGCTTGCGCTTCAGAACGCTGGGTTGAAAGGTACGTTTCATAACACTTGATTCCCACGTGGTTTATTGACGGAACG

TCTTACAGAAATGCCCTGATTCCCGACAGGAACCAGGGGCAGATTCGGTTCAAGACCGGAAATTGTAGAGACTTCGGCTGCCCGACGCAATTGTTCCGCT
TGCTGCAGCGGGTCGAGCAGCCGCTGGCGGTTGGCCGAATCGCAGACTTGCTGTCATGGCTTCCCGGCCCACGGCTGTGGTTAACATTGGGGACTCTATC
**TGTTGATAA**GTTATTTGATTAATGAAAGATCAAAGACATAGGTTGGGGCATGATCTGTGGAAAAGCGTTTGGTAAGTCCTGCGCATCAGGGGCGGTTGAA
GAAGCGAATCCAAGGCACGCAGCCAGACTGTCGAGAGCCTGCTGCAGAGCCGCTTTTCCACAGGGTTGTCTGGTGTCGAACCGCATGTGGATAACCCGCC
AGGCGAAGGATACAATGGCCGGTCATGCATCAACGGGTATGGTGAGGTCGAGTGTCTCTTGCTCTCTGGCAACAGTGTCTGGATTACCTGCAGGACGAGC

TGAATGCACAGCAGTTCAATACCTGGATTCGCCCGCTTCAAGCCGAAGAGGGGGAAGCCAACGAGCTGCGCCTGCTGGCACCCAATCGCTTCGTTCGTGA
TTGGGTTGGCGACAAGTATGCCAAGCGCATCAACGAACTGCTCCGGGATCTGGCCCCAGCCAAACCGCCAAGGGTAGTGCTGACAGTCGGAAGCCGCAGG
GCGGCAGCGGTCAAGCCACAGCCACGAGACCTGGGAGCCCCGGTATCGGCCAACCCTCACTCTCCCCTGGCGCCGGCCGTGCAAGTTCCACCGCGAGGAG
ATATCGGCGACGAGCGTGAAATTGACCAGCTGCGTGAAGAGGGGCTCGGCTCGGGGCGTCGTCCGAGCGGGGATCGCGAAGTACAGGTCGAAGGCAGCCT
GAAACACAACAGCGGCCTCAATCCTACGTTTACCTTCGAGACCTTTGTCGAGGGCAAGTCGAACCAGTTGGCGCGGGCCGCCTCGCGGCAGGTTTCCGAG
AACCCGGGAGGCGCCTACAACCCACTGTTCCTGTATGGCGGCGTCGGTCTGGGCAAGACCCACCTGATGCACGCGGTGGGCAACGCGCTGGCAACCCGGC
GCGAGAATGCACGCGTCGTCTACCTGCACTCCGAACGTTTCGTGGCCGACATGGTCAAGGCGCTGCAGCTCAACGCCATCAACGACTTCAAGCGTTTCTA
CCGCAGCGTGGACGCGCTGTTGATCGACGATATCCAGTTCTTTGCCGGCAAGGAGCGCTCCCAGGAGGAGTTCTTTCATACCTTCAATGCACTGCTCGAG
GGTGGCCAGCAGATGATCCTCACCTCTGATCGCTATCCCAAGGAGATCAGCGGTGTCGAGGAGCGTCTCAAGTCGCGCTTCGGCTGGGGGTTGACCGTTG
CCATCGAGCCGCCCGAGCTGGAAACCCGTGTCGCCATTCTGATGAAGAAGGCAGACCAGGCCAAGGTGGACCTGCCGCACGATGCCGCCTTCTTCATTGC

>Halomonas diversa MCCC 1A13316 [NZ_CP053382.1:4465560..4466059], <mnmG oriC2 <COG1126 (glnQ), 529 bp

GTTTTCCAGCCCCTTGATCGAGCGCACCACCTGCAATTGCACGTCGAAGGGCAGCGAGGTGGAAATCCCGTTGGGGTAGAGCTCATGGGTATCCAGCCCC
TCGGGCTCGATGAATATCTGGTGGCTCGCCTTGTCGGCAAAGCGGTGCACCTTGTCCTCGATCGACGGGCAGTAGCGCGGCCCAACGCCTTCGATCACAC
CGGAGTACATCGGTGAACGATCGAGGTTGGCGAAGATGATCTCGTGGGTACGCTCGTTGGTATGGGCGATATGACAGCTTACCTGACGCGGATGCATGTC
ACGCCGTCCCAGGTAGGACATCACCGGAGTCGGGGTATCGCCGGGTTGCTCCTCGAGCTGCGAGAAATCCACGCTTTTGGCGTCCAGCCTTGGCGGCGTA
CCGGTCTTGAGCCGATCGACACGGAACGGCAGCGCACGCAGACGCTCGGCCAGGGCGTTGGAAGGCGGGTCCCCGGCGCGCCCCCCGCGGCTCTGATCGA
GCCCGATATGGATTACGCCACCGAGGAAGGTGCCCGTGCACAGCACCACCGCCTCGGCCAGGAAGCGAATCCCGGTCTCGGTGACCACGCCTCGTACGGT
GTCGTCGTCCACCAGCAGGTCGCCGGCGGCCTGCTGGAAGATCGTCAGATTGGCCTGGTTCTCGAGCAAGCCGCGAACGGCTGCTTTGTAGCGTACGCGG
TCGGCCTGGGCCCGGGTGGCACGAACCGCAGGGCCCTTGCGTGCGTTGAGCACACGGAACTGAATACCACCGAGATCGGTAGCCAGGCCCATGGCGCCGC
CCAGGGCATCGATTTCCTTGACCAGATGGCTCTTGCCGATCCCACCAATGGCGGGATTGCACGACATCTGCCCGAGAGTCTCGATGTTATGGGTCAGTAG
CAGGGTCTGACAGCCCATGCGAGCGGAGGCCAGAGCGGCTTCGGTTCCCGCATGGCCGCCGCCGATGACGATGACGTCAAAGCGGTCGGGATACTCCAAG

GTGCACCTCGTTGCGCTTGTGCGCAGATATTGCCTAGGGACGAAAATCAGCCTGGCAGTATAAACCTCCTGTGGGTAAGCGTCAGGGGTTTTCCACACCC
ACTCGCAAGGAAGCCTCTTTCCCTCTTCTGGATAAACAAAACAGAATGATTAAAAGAGAGAAGTTCTGTTGTGGTTGTTACTACTAGTGTGGACAAAAAC
**CGTGGATAA**CCCTGGTTAGTCTTTTGGCATCAAAAGCTTGCATTGTTGACTTCTGCGCTGAGAAAAGGGTGACGAGCTGAGGGCGGCCGGTGACGAGCCT
GTGGATGAAAAGCATGCTTTTCCACAGCGCGAACGCTGCACCTCTTTTCCACCGCCGCCTACCGGACTTGTTACCGCGATTGTGAACAACCGGCCGCGAA
CATGACGTAGGCACCAACCATGCACCCTGGAAGCAAGAAAGAGGCGTTTTCAAAAAAAATTTATCCACAGGCAAAAAAAAGCCCTGACGATGGCGTCAGG

GCTTCGCAAGAGCGTATCGGGGAAGCTGCTAGTGCTTGAGCACGCGAGAGAGAAAGCTCTGGGTACGCTCGTGTTGGGGGGAGTCGAACAGCACGTCGGG
AGCGGCCTGTTCGACGATCTCCCCACGATGTATGAACACGACACGGTCGGCGACTTCACGGGCGAACCCCATCTCGTGGGTAACGATGATCATGGTCATG
CCCTCGCGAGCGAGCTCGCGCATGGCGTCGAGCACTTCACCGATCATTTCGGGATCGAGTGCCGAAGTGGGTTCGTCGAACAGCATCAGCCGCGGCTCCA
TGGCAAGTGCCCGCGCCAGGGCGACGCGCTGTTGCTGGCCTCCGGAGAGCTGGGAAGGGTACTTGTCGGCCTGGTCGGCGATCCCCACACGTTCGAGTAG
GCGATTGGCTGTCTCGGTGGCATCTGCGAGGTTCCAGCCGCGCACTTTCATCGGCGCCAGTGTGACGTTGTCGCGTACGCTGAGATGGGGAAAAAGATTG
AACTGCTGGAATACCATGCCGACCTCGGTGCGGATCGTCTGCAGCGCACGGCTGCTCTTGCCATAGGGCAGCAATTCGTTGCCGTCGACCTCGATATGGC
CGGCTTGGAACTCCTCAAGGCCGTTGATGCAGCGAATCAGGGTCGATTTTCCGGAGCCGCTGGCACCGATGACCACCACCACCTCACCAGGCCGGATCGT
AAGGTCGATGTCCTTGAGCACGTGCAACTCACCGAAGTGCTTGTTGACCTTCTCCATTTGCACGATGAGGGCGGCATCGGAGGGAGTATTGTCTCGGTTC
ATGTCGCGCTCCTTCATTGTCCAACCAGGCCCCTGCGCTCGAGCAGGCGCAGTGCCAATGACAGGCTCAGGGTTATCGCCAGATAGAGGAGGGCGACCAT
GAGATAGACTTCCAGGGCGTTGAAGGTAGTGGCGATATAGACCTGGCCCTGGCGCACCAGCTCGCCGACCCCGATGACCGAGAACAGCGAAGTGTCCTTG

>Halomonas diversa MCCC 1A13316 [NZ_CP053382.1:1721..1789], dnaA> ig dnaN>, 69 bp

WebSkew GCmin pos. 5015118/0 (GCmin<>oriC1 0.0 kb; GCmin<>oriC2 38.4 kb; oriC2<>oriC1 38.4 kb)

>Halomonas tianxiuensis BC-M4-5 [NZ_CP035042.1:5014740..121], <rpmH oriC1 dnaA>, 499 bp

GCGTCATGCGGTGCCGCGGCAGGGCCGCATAGACGATGTCGCCGCCTTGGGGATCGATGCGAACGTCGAGCACGTCGGTCATGACCGCGATCAGGTCGCG
CGTTCCGCTCGCCTGCTCCGTCTCGGCTGGCATTTCGCTGGCCACGTCGCGCTCCGCTTGCGGAGCGGCGGGAACCGCCAGGCCGTCATCGTCGGGCGCT
TGCGCTTCACCGGGTGCCGGTGCCGTGATGGTGGGAGCTTCGGGCTCAGGAGCAACCTGGCCATAGTCCTGGTTCCACTGCACGACTAGCAGATAGGTGA
GCACGGCCAGTGGAATCAGCAATAGGAGTCGTTTTACGTCCATGAGGGTTCTGCCCGGTGGGTAGTGAACATGCCGCTGGAACGGAATCCGCTGCCATGT
GAAAGAGGGCGATAGCCTGCCGAGCAGGCTATCGGGGCGGAGACGAGTACCGCATCCGGTCAGGAGGTCATCGCTTTTCGTGCATCACGCTCCAGGCGCC
GCCACATGCCGAAAAGCTGGCGATGCAGCGTGGCCTTGTCCAGCTCGCTCGCGCCGCGCCGCGCCAGCAGGACGATATCGACGGCGGGTAGCGAGCATTG
CTGAAGACGAATGGATTCACGAGTCAACCGCTTGAGGCGGTTGCGGTCCACGGCCCTGCGGACGTTCTTCTTGCTGATGACCAGGCCAAGGCGGGGGTGG
TCCAGTCCATTGGGGCACGCCAGGGCCATCAGCCCCTTGCCATGTACCTTGAATGTAGCGTGATCGAAAACGTATCGATAATCCCCGGCAGTCAGCAAGC
GCAGCTGCTTGGGGAAACCATGACCGGACACGCGCAATCCGCTATCAGGCGCTCAGACGCTTGCGGCCCTTGGCGCGGCGGCGGGCCAGAACGGCACGAC
CATTCTTGGTAGCCATGCGTGCACGGAAGCCGTGGACACGCTTGCGCTTCAGAACGCTGGGTTGAAAAGTACGTTTCATAACACTCGATTCCCACGTGGT

TTTTGACGGTACGTCGTTACAGGAACGCCCTGGCTCCCGATAGGAACCAGGGACAGATTCGGTTCAAGACCGGAAATTGTAGAGACTTCGGCTGCCCGAC
GCAATTGTTACGCTCCATGGCGCGACTGGAAGCGGGGCCGGCGAACGGCGTAATTTCCTGATTTTTGTGGGTTGTTGTGATTAATACTGTGGGCGACGTC
**TGTTGATAA**GTAATAAAATCATATTCGAAACAATAGGTTAGGTTTTCTTTGGGTGTGTGGAAAAGCTCTTTGTAAACAGTGGCTAGACAGTGGGTGAGGC
TGGCAGGGTGACGGTCTGGGAAATCGGCTTCCCCCATGCGTAGCGAAGCCAAGGGGAGCCCGTAGCCAGACGCGATGACGTTTTCCACAGGAGCGTTCGG
CATTGCCTGGCATGTGGATAACCCGCCGGCCGAAGGATACAATGGCCGGTCATGCATCAACGGGTATGGTGAGGTCGAGTGTCTCTAGCTCTCTGGCAAC

AGTGTCTGGAATACCTGCAGGACGAGCTGAACGCCCAGCAGTTCAATACTTGGATTCGTCCGCTCCAAGCCGAGGAGGGGGACGCCAACGAGCTGCGTCT
GCTGGCGCCCAATCGCTTCGTTCGCGACTGGGTTGGCGACAAGTATGCCAAGCGTATCAACGAGCTGCTCAGGGATCTGTCACCTGCCAAGCCGCCGCGT
GTGGTGCTGACCGTGGGCAGCCGCCAGGCGGCGACAGTCAAGCCGCAGCCGCGCGACTTGGGGGCTCCTGTCTCGGCCAACCCCCACTCGCCCTTGGCGC
CGGCGGTTCAGGTGCCACCGCGTGGCGATATCGGTGACGAGCGCGAAATTGACCAACTGCGCGAGGAGGGGGCCACGTCACGCCGCTCCGGCGCGGAACG
TGAGGTGCAAGTCGAAGGCAGTTTGAAGCACAGCAGTGGCCTCAACCCCAGCTTCACCTTCGAAACCTTCGTCGAGGGCAAGTCCAACCAGTTGGCTCGG
GCCGCCTCGCGACAGGTTTCCGAGAACCCCGGCGGCGCCTATAACCCGCTCTTTCTCTATGGTGGCGTCGGCCTGGGCAAGACTCACCTGATGCACGCCG
TGGGCAACGCCCTGGCGGCTCGGCGCGAGAACGCTCGTGTCGTCTACCTGCATTCCGAGCGCTTCGTGGCGGACATGGTCAAGGCGTTGCAGCTCAACGC
CATCAACGACTTCAAGCGCTTCTACCGCAGTGTCGATGCGCTGCTGATCGACGACATCCAGTTCTTTGCCGGCAAGGAGCGCTCCCAGGAGGAGTTCTTC
CATACTTTCAATGCCCTGCTCGAGGGTGGCCAGCAGATGATCCTCACCTCGGACCGCTACCCCAAGGAGATCAGCGGTGTCGAGGAGCGTCTCAAGTCGC
GCTTCGGCTGGGGGCTGACCGTGGCCATCGAACCGCCCGAACTGGAAACCCGGGTCGCCATCCTGATGAAGAAGGCGGACCAGGCCAAGGTGGATCTGCC

>Halomonas tianxiuensis BC-M4-5 [NZ_CP035042.1:4976356..4976855], <mnmG oriC2 <COG1126 (glnQ), 539 bp

GCGTTCTCCAATCCCTTGATCGAGCGCACCACCTGCAGCTGCACGTCGAAGGGCAGCGAAGTGGAGATGCCGTTGGGGTAGAGCTCGTGAGTATCGAGCC
CTTCCGGTTCGATGAACACCTGGTGACTCGCCTTGTCGGCGAAGCGGTGCACCTTGTCCTCGATCGACGGGCAGTAACGCGGCCCGACGCCCTCGATCAC
GCCCGAATACATCGGCGAGCGGTCGAGGTTGGCGAAGATGATCTCGTGGGTGCGCTCATTGGTATGAGCGATGTGGCAGCTCACCTGGCGCGGATGCATG
TCGCGACGTCCCAGGTAGGACATCACCGGCGTCGGGCTGTCGCCTGGCTGCTCCTCGAGCACGGTGAAATTCACGCTCTTGGCATCCAGCCGTGGCGGGG
TGCCGGTCTTGAGCCGGTCGACCCGGAACGGCAGCGCCCGCAGACGCTCCGCCAGGGCATTGGAGGGAGGATCGCCGGCACGCCCGCCGCGGCTCCGATC
CAGGCCGATATGGATCACGCCGCCGAGGAAGGTGCCGGTGCACAGCACCACGGTTTCGGCACGGAAGCGGATGCCGGTCTCGGTGACTACCCCGCGCACC
GTGTCGTTTTCCACCAGCAGATCTCCTGCAGCCTGCTGGAAGATCGTCAGGTTCGGCTGGTTCTCCAGCATGCCGCGGATGGCCGCCTTGTAGCGTACGC
GGTCGGCCTGAGCACGGGTGGCCCGCACGGCCGGTCCCTTGCGAGCGTTGAGCACGCGGAACTGGATGCCGCCGAGATCGGTGGCCAGGCCCATGGCTCC
CCCCAGCGCGTCGATCTCCTTGACCAGGTGGCTCTTGCCGATGCCGCCGATGGCGGGATTGCACGACATCTGCCCGAGGGTCTCGATGTTGTGGGTCAGC
AGCAGGGTCTGACAGCCCATGCGGGCGGAGGCCAGAGCGGCTTCGGTTCCCGCATGGCCGCCGCCGATGACGATGACGTCAAAGCGGTCGGGATACTCCA

AGGTGCACCTCGTTGCGCTAGTGCGCGGATGTTGCCAGTGACGAAAAACAGCCTGCCAGTATAAACCTCCTGTGGGTAAGCGTCAGGGTTTTCCACACCG
CCTCTGCCCGCAGGCTCGTTTTCCTCCCAGATAACCAAAACAGAATATCTAAAAAGAAAGAAGTTCTGCTGTGGTTGTGACTACTGGTGTGGACAATATT
**TGTGGATAA**GCAGAAAAATTCCAGCCAAGT**T**AACGGTTTGCTTTGTTAACTTCTATGCGGAGAAGAGGGTGACGAGCTGAGAGTTGCCGGTGCGGGGGCT
GTGGATGAACGCCAAGCTTTTCCACAGCCCGGACGCTACACCTTTTTTCCACCGTTGCCTACCTGGCTTGTCACCGCGACTGTGAACAGTCACGCGAGGC
GAACGAGAAGGCGGCAACCATGCCGCTTGGAGCGATGAATCGAGGGTAGAGCAAAAATTCTTATCCACAGGCCGAAAAAAAGCCCTGCCGAGTGCGACAG

GGCCTGCCAGGGTGGAGAGCAGCGGCTATCTGTGGGCCCCTCAGTGCTTGAGCACCCGCGACAGGAAGCTCTGGGTGCGCTCGTGCCGTGGAGAGTCGAA
AAGAGTGTCCGGTGCCGCTTGCTCGACGATCTCTCCGCGATCGATGAAGACCACTCGATCGGCGACCTCGCGGGCGAATCCCATCTCATGGGTAACGATG
ACCATGGTCATACCCTCGCGGGCAAGCTCTCGCATGGCATCGAGCACTTCACCGATCATTTCCGGATCGAGCGCCGAGGTGGGTTCGTCGAACAGCATCA
GGCGCGGTTCCATGGCCAGCGCTCTGGCCAGGGCGACGCGCTGCTGCTGCCCGCCGGAGAGCTGGGATGGGTACTTGTCCGCCTGGTCGGCGATGCTGAC
GCGTTCGAGCAGGCGGTAGGCAGTCTCGGTGGCATCGGCTCGGCTCCAGCCGCGCACCTTCATGGGCGCCAGCGTGACGTTATCGAGCACGCTGAGATGT
GGAAAGAGGTTGAACTGCTGGAACACCATGCCGACTTCGGTGCGGATTCTCTGCAGCGCGCGACCGCTCTTGCCGTTCGGTAGCAGTTCGTTGCCATCGA
CTTCGATGTGGCCGGCCTGAAACTCCTCGAGGCCATTGATGCAGCGGATCAGGGTGGACTTGCCCGAGCCGCTGGCACCGATAACCACCACCACCTCGCC
AGGCTGGATCTCGAGCACGATGTCCTTGAGCACGTGCAGTTGTCCGAAGTGCTTGTTCACCCTGTCAAGGCGCACGATGGGCTGGGTGTCGGCCAGGGAT
TCGTCTCGGCTCATGTCGCGCTCCTTATTGTCCGACCAGGCCCTTGCGCTCGAGCAGGCGCAGGGCGAATGACAGGCTCAGGGTTATCGCCAGATAGAGG
AGGGCGACCATCAAGTAGACTTCCAGGGCGTTGAAGGTGGTGGCGATGTAGACCTGGCCCTGGCGGACCAGCTCGCCAACCCCGATCACCGAGAATAGCG

>Halomonas tianxiuensis BC-M4-5 [NZ_CP035042.1:1573..1639], dnaA> ig dnaN>, 67 bp

WebSkew GCmin pos. 178350 (GCmin<>oriC1 40.3 kb; GCmin<>oriC2 8.0 kb; oriC2<>oriC1 32.2 kb);

>Halomonas zhangzhouensis MCCC 1A11036 [NZ_JABFTT010000001.1:137809..138308], <rpmH oriC1 dnaA>, 499 bp

GCGTCATGCGGTGCTGCGGCAGGGCCGCATAGACGATGTCGCCGCCCTGGGGATCGATGCGAACGTCGAGTACGTCGGTCACGACGGCGATCAGGTCGCG
CGTTCCGCTCGGCTGCTCCGTCTCGGCTGGCATTTCGCTTGCCACGTCGCGCTCTGCTTGCGGAGCGGCGGGAACCGCCAGGCCATCATCGTCGGGCGCT
TGCGCTTCACCAGGTGCCGGTGCCGTGATGGTGGGAGCTTCGGGCTCAGGAGCAACCTGGCCATAGTCCTGGTTCCACTGCACGACCAACAGATAGGTGA
GCACGGCCAGTGGAATCAGCAATAGGAGTCGTTTTACGTCCATGAGGGTTCTGCCCGGTGGGTAGTGAACATGCCGCTGGAACGAAATCCGCTGCCATGT
GAAAGAGGGCGATAGCCTGCCGAGCAGGCTATCAGGGCGGAGACGAGTACCGCATCCGGTCAGGAGGTCATCGCTTTTCGTGCATCACGCTCCAGGCGCC
GCCACATGCCGAAAAGCTGGCGATGCAGCATGGCCTTGTCCAGCTCGCTCGCGCCTCGCCGCGCCAGCAGGACGATATCGACAGCGGGTAGCGAGCATTG
CTGAAGACGAATGGATTCACGAGTCAACCGCTTGAGGCGGTTGCGGTCCACGGCCCTGCGGACGTTCTTCTTGCTGATGACCAGGCCAAGGCGGGGGTGG
TCCAGTCCATTGGGGCACGCCAGGGCCATCAGCCCCTTGCCATGTACCTTGAATGTAGCGTGATCGAAAACGTATCGATAATTCCCGGCAGTCAGCAAGC
GCAACTGCTTGGGGAAACCATGACCGGACACGCGCAATCCGCTATCAGGCGCTCAGACGCTTGCGGCCCTTGGCGCGGCGGCGGGCCAGAACGGCACGAC
CATTCTTGGTAGCCATGCGTGCACGGAAGCCGTGGACACGCTTGCGCTTCAGAACGCTGGGTTGAAAAGTACGTTTCATAACACTCGATTCCCACGTGGT

TTTTGACGGTACGTCGTTACAGGAACGCCCTGGCTCCCGACAGGAACCAAGGACAGATTCGGTTCAAGACCGGAAATTGTAGAGACTTCGGCAGCCCGAC
GCAATTGTTACGCTCCATGCCGCGAGTGGAAGCGGGGCCGGTGAACGGCGTAATTTCCTGTTTTTTGTGGGTTATTGTGGTTAATACTGTGGGCGATAAC
**TGTTGATAA**GTAATAAAACCATATTTGAAACAATAGGATAGATTGGATTTTGGTATGTGGAAAAGCCTTTTGTAAACAGTGGCCAGGCAGTGGGTGGGGC
TGGCTGGGTGGTGGTTTGAGAAATCGGCTTCCCCCATGTGTAGCGAAGCCAAGGGGAACCCGTAGCCAGTCGTGATGACGTTTTCCACAGGAGCGTTCGG
CATTGCCTGGCATGTGGATAACCCGCCGGCCGAAGGATACAATGGCCGGTCATGCATCAACGGGTATGGTGAGGTCGAGTGTCTCTTGCTCTCTGGCAAC

AGTGTCTGGATTACCTGCAGGACGAGCTGAACGCCCAGCAGTTCAATACTTGGATTCGTCCGCTCCAAGCCGAGGAGGGGGCGCCAACGAGCTGCGTCT
GCTGGCACCCAATCGCTTCGTTCGCGACTGGGTTGGCGACAAGTATGCCAAGCGTATCAACGAGCTGCTCAGGGATCTGTCACCTGCCAAGCCGCCGCGT
GTGGTGCTGACCGTGGGTAGCCGCCAGGCGGCGGCAGTCAAGCCGCAGCCGCGCGACCTGGGAGCTCCTGTCTCGGCCAACCCCCACTCGCCCCTGTCGC
CAGCGGTTCAGGTGCCGCCGCGTGGCGATATCGGTGACGAGCGCGAAATCGACCAGCTGCGCGAGGAGGGGGCCTCGTCACGCCGCTCCGGCGCGGAACG
TGAGGTGCAAGTCGAAGGCAGCTTGAAGCACAACAGCGGCCTCAACCCCAGCTTCACCTTCGAAACCTTTGTCGAGGGCAAGTCCAACCAGTTGGCCCGG
GCTGCCTCGCGACAGGTCTCCGAGAACCCCGGCGGCGCCTACAACCCGCTCTTTCTCTATGGTGGCGTCGGCCTGGGCAAGACCCACCTGATGCACGCCG
TGGGCAACGCTCTGGCGGCGCGGCGCGAGAATGCTCGTGTCGTTTACCTGCATTCCGAGCGTTTCGTGGCGGACATGGTCAAGGCGCTGCAGCTCAACGC
CATCAACGACTTCAAGCGCTTCTACCGTAGTGTCGATGCGCTGCTGATCGACGACATCCAGTTCTTTGCCGGCAAGGAGCGCTCCCAGGAGGAGTTCTTC
CATACTTTCAATGCCCTGCTCGAGGGTGGCCAGCAGATGATTCTCACCTCGGACCGCTACCCCAAGGAGATCAGCGGTGTCGAGGAGCGTCTCAAGTCGC
GCTTCGGCTGGGGGCTGACCGTGGCCATCGAACCGCCTGAACTGGAAACCCGGGTCGCCATCCTGATGAAGAAGGCGGACCAGGCCAAGGTGGATCTGCC

>Halomonas zhangzhouensis MCCC 1A11036 [NZ_JABFTT010000001.1:170039..170538], <mnmG oriC2 <COG1126 (glnQ), 537 bp

GCATTCTCCAATCCCTTGATCGAGCGCACCACCTGCAGCTGCACGTCAAAGGGCAGCGAAGTCGAGATGCCGTTGGGGTAGAGCTCATGGGTATCGAGCC
CTTCCGGTTCGATGAACACCTGGTGGCTCGCCTTGTCGGCGAAGCGATGCACCTTGTCCTCGATCGACGGACAGTAACGCGGTCCGACGCCCTCGATCAC
GCCCGAATACATCGGTGAGCGGTCGAGGTTGGCGAAGATGGTCTCGTGGGTGCGCTCGTTGGTGTGAGCAATGTGGCAGCTCACCTGGCGTGGGTGCATG
TCGCGACGCCCCAGGTAGGACATTACCGGTGTCGGGCTATCGCCCGGCTGCTCCTCGAGCACGGCGAAATTCACGCTCTTGGCATCCAGCCGTGGCGGAG
TGCCGGTCTTGAGCCGGTCGACCCGGAACGGCAGCGCCCGCAGACGCTCCGCCAGGGCATTGGAGGGAGGATCGCCGGCACGACCGCCGCGGCTCCGATC
CAGGCCGATATGGATCACCCCACCGAGGAAAGTGCCGGTACACAGCACCACGGTTTCAGCATGGAAGCGAATGCCGGTCTCGGTGACTACCCCACGCACC
GTGTCGTTTTCCACCAGCAGATCTCCCGCCGCCTGCTGGAAGATCGTCAGGTTCGGCTGGTTCTCCAGCATGCCGCGGATAGCGGCCTTGTAGCGCACGC
GATCGGCCTGAGCACGGGTAGCCCGCACGGCCGGTCCCTTGCGAGCGTTGAGTACGCGGAACTGGATGCCGCCGAGATCGGTGGCCAGGCCCATGGCTCC
CCCCAGCGCATCGATCTCCTTGACGAGATGGCTCTTGCCGATGCCACCGATGGCGGGATTGCACGACATCTGCCCGAGGGTCTCGATGTTGTGGGTCAGC
AGCAGGGTCTGACAGCCCATGCGGGCGGAGGCCAGAGCGGCTTCGGTTCCCGCATGGCCGCCGCCGATGACGATGACGTCAAAGCGGTCGGGATACTCCA

AGGTGCACCTCGTTGCGCTAGTGCGCGGATGTTGCCAGTGACGAAAAACAGCCTGCCAGTATAAACCTCCTGTGGGTAAGCGTCAGGGTTTTCCACACCG
TCTCTGCCAGCAGGATCACTTTCCTACCAAATAACCAAAACAAAGTATCTAAAAAGAAAGAAGTTCTGCTGTGGTTGTGACTACTGGTGTGGACAATATT
**TGTGGATAA**GAAGAAAAAATCAAGCAGGGT**T**AACGATTTGCTTTGTTAACTTCTATGCGGAGAAGAGGGTGACGAGCTGAGAGTTGCCAGTGTGGGGGCT
GTGGATGAACCCTAACGTTTTCCACAGCCAGGACGCTACACCCTTTTTCCACCGTTGCCTACCTGGCTTGTCACCGTGACTGTGAACAGTCACGCGAAGC
CAACGAAAAGGCGGCAACCATGCCGCTTGGAGCAACGAATCGAGGGTAGAGGAAAAATTCTTATCCACAGGCCGAAAAAACCCTGTCGAGTGCGACAGGG

CCTGCCAGGGCGGACAGCAGCGGCCTTCTGTGGATCCCTCAGTGCTTGAGCACCCGCGACAGAAAGCTCTGGGTGCGCTCGTGCTGCGGGGAGTCGAAAA
GGGTGTCTGGGGCAGCCTGCTCGACAATCTCGCCGCGATCGATGAAGACCACGCGATCGGCGACCTCCCGAGCGAATCCCATCTCGTGGGTAACGATGAC
CATGGTCATACCCTCGCGGGCGAGCTCACGCATGGCATCGAGCACTTCACCGATCATTTCCGGATCGAGCGCCGAGGTGGGTTCGTCGAACAGCATCAGG
CGCGGTTCCATGGCCAGCGCTCTAGCCAGGGCGACGCGCTGCTGCTGCCCGCCGGAGAGCTGGGATGGGTACTTGTCCGCCTGGTCGGCGATGCCGACGC
GTTCGAGCAGGCGGTAGGCAGTCTCGGTGGCATCGGCTCGGCTCCAGCCGCGCACCTTCATGGGCGCCAGCGTGACGTTATCGAGCACGCTGAGATGTGG
AAACAGGTTGAACTGCTGGAACACCATGCCGACTTCGGTGCGGATTCTCTGCAGCGCGCGACCGCTCTTGCCGTCCGGTAGCAGCTCGTTGCCATCGACT
TCGATGTGGCCAGCCTGGAACTCCTCAAGGCCATTGATGCAGCGGATCAGGGTGGATTTACCCGAGCCGCTGGCACCGATCACCACCACCACCTCGCCAG
GCTGGATCTCGATCACGATGTCCTTGAGCACGTGCAGTTGTCCGAAGTGCTTGTTGACCCTGTCAAGGCGCACGATGGGGACAGCGTCGGCGTGTGTATC
GTCTCGGCTCATGTCGCGCTCCTTATTGTCCGACCAGGCCCTTGCGCTCGAGCAGGCGCAGGGCGAATGACAGGCTCAGGGTTATCGCCAGATAGAGGAG
GGCGACCATCAAGTAGACTTCCAGGGCGTTGAAGGTGGTGGCGATGTAGACCTGGCCCTGGCGGACCAGCTCGCCGACCCCGATCACCGAGAATAGCGAG

>Halomonas zhangzhouensis MCCC 1A11036 [NZ_JABFTT010000001.1:136314..136357], dnaA> ig dnaN>, 45 bp

WebSkew GCmin pos. nd (GCmin<>oriC1 nd; GCmin<>oriC2 nd; oriC2<>oriC1 nd)

>Halomonas desiderata FB2 [NZ_JAAQTN010000031.1:15032..15531; <rpmH oriC1 dnaA>, 513 bp

GGGCAGTGCCGCATAGACGATGTCTCCCCCCTGTGGGTCGATGCGCACGTCGAGCACGTCGGTCACCACGGCGACCAGGTCGCGGGTGCTGGTGGGCTCG
GCAGGCTCCGCCGGCATGTCGCCGGTCATCTCCTGCTCCGCCCTGGGGGCGGCCGGCACGGCCAGCGCGTCGTCATCGGGCGCCTGCGCCTCGCCGGGTG
CCGGGGCGGTGATGGTGGGAGCATCTGGCTCAGGAGCAACCTGGCCATAATCCTGGTTCCACTGTACCACCAGCAGGTAGGTGAGCACGGCCAGTGGAAT
CAGCAATAGGAGTCGTTTTACGTCCATGAGGGGTCTGCCCGGTGGGTAGTGAACATGCCGCTGGAACGAGATCCGCTGCCATGTGAAAGAGGGCGACAGC
CTGCCGGGGCAGGCCATCGAAGCGGAGACTAATACCGCATCCGGTCAGGAGGTCATCGACTTCTGCGCGTCACGCTCCAGGCGCCGCCACATGCCGAAAA
GCTGGCGATGCAGCGTGGCCTTGTCCAGTTCGCTCGCGCCGCGACGTGCCAGCAGGACGATATCCACGGCGGGTAATGAGCGCTGGCGCAGACGTACGGA
TTCGCGCACCACACGTTTCAGGCAGTTGCGATCCACGGCTCTGCGGACGTTCTTCTTGCTGATGACCAGGCCAAGGCGGGGGTGGTCCAGTCCATTGGGG
CACGCCAGGGCCATCAGCCCCTTGCCATGTACCTTGAGTGCAGCGTGATCGAAAACGTATCGATAATCCCCGGCAGTCAGCAGTCGCAGCTGTTTGGGGA
AACCATGACCGGACACGCGCAATCCGCGATCAGGCGCTCAGACGCTTGCGGCCCTTGGCGCGGCGGCGAGCCAGAACGGCACGACCATTCTTGGTAGCCA
TGCGAGCACGGAATCCGTGGACACGCTTGCGCTTCAGAACGCTGGGTTGAAAGGTGCGTTTCATAACTCTTGATTCCCACGTGGTTTGTTGACGGATCGT

CTTACAGGAATGCCCTGGCTCCCGACAGGAACCAGGGGCAGATTCGGTTCAAGACCGGAAATTGTAGAGACTTCGGCTGCCCGACGCAATTGTTCCTCAC
GCTCCCGCATCACCACAGGGCGTCCGCAGCGATCGCCACTGTAGCAAACATCATAGGATGTTTTCTTGAACCTGCTGTGACTAATAATGGGGATCGCTTC
**TGTTGATAA**AGTGATATTCACCTTTCCTATCAGAGGCCTGACCGTTGGCTGAGTCTGTGGAAAAGCGTTGGGGAAGGCGTGCTCATGCTGTGATTCGACT
GTTGATGGCTACCTGACTCATCCACAGGTCGAGGTGGTCCGCGGTTGATGGACAGAATGATCACGGGTGCCGCGAGGGCTTTTCCACAGCCATGTTCCGG
CAGTTTCCGCTGTGGATAACCCGGCGGGCGAAGGATACAATGTCCGGTCACGCATCAACGGGTAATGGTGAGGTCGAGTGTCTCTTGCTCTCTGGCAACA

GTGTCTGGATTACCTGCAGGACGAGCTGAATGCCCAGCAGTTCAATACCTGGATTCGCCCGCTTCAAGCCGAGGAGGGCGAAGCCAACGAGCTGCGACTG
CTGGCGCCCAACCGCTTCGTGCGCGACTGGGTCGGCGACAAGTATGCCAAGCGTATCAATGAGCTGCTCAGGGAGCTGGCGCCTGCCAAGCCGCCGCGGG
TGGTGCTGGCCGTGGGCAGTCGCACGGTGGCGGCGGTCAAGCCGCAGACGCGCGAGCTCGGCGCTCCGGTCTCGGCCAATCCACACTCTCCCCTGGCGCC
TGCCGTTCAGGTGCCGCCTCGTGGCGACATCGGCGATGAGCGCGAGATCGACCAGCTGCGCGAGGAGGGCGCCACGGCACGTCGCGGCGGCAGCGAGCGC
GAAGTGCAGGTCGAGGGCAGCCTCAAGCACAGCAGCGGCCTCAACCCCAGCTTCACCTTCGAAACCTTCGTCGAGGGCAAGTCCAACCAGCTGGCTCGAG
CCGCCTCGCGGCAGGTCTCGGAGAACCCCGGTGGCGCCTACAACCCGCTGTTCCTCTATGGCGGCGTTGGCCTGGGCAAGACCCACCTGATGCACGCCGT
AGGCAATGCCCTGGCCTCGAGGCGCGAGAACGCCCGGGTGGTCTATCTGCACTCCGAGCGCTTCGTGGCGGACATGGTCAAGGCACTGCAGCTCAACGCC
ATCAACGATTTCAAGCGTTTCTATCGCAGCGTCGATGCGCTGCTGATCGACGACATCCAGTTCTTCGCCGGCAAGGAGCGCTCCCAGGAGGAGTTCTTCC
ACACTTTCAATGCCTTGCTCGAGGGCGGTCAGCAGATGATCCTCACCTCGGACCGCTACCCCAAGGAGATCAGCGGGGTCGAGGAGCGCCTCAAGTCGCG
CTTCGGCTGGGGGCTGACCGTGGCCATCGAGCCGCCGGAGCTGGAAACCCGGGTCGCCATCCTGATGAAGAAGGCCGATCAGGCCAAGGTGGATCTGCCC

>Halomonas desiderata FB2[NZ_JAAQTN010000055.1:c7945..8444], <mnmG oriC2 <PRK13623 (ErpA), 536 bp

CGTTCTCCAGCCCCTTGATCGAGCGCACCACCTGCAGCTGCACGTCGAACGGCAGCGAAGTCGAAATCCCGTTGGGGTAGAGCTCGTGGGTATCGAGGCC
TTCGGGCTCGATGAACACCTGGTGGCTCGCCTTGTCGGCGAAGCGGTGCACCTTGTCCTCGATCGACGGGCAGTAGCGCGGCCCGACCCCCTCGATCACG
CCGGAATACATCGGCGAGCGGTCGAGGTTGGCGAAGATGATCTCGTGGGTGCGCTCGTTGGTGTGCGCAATGTGGCAGCTCACCTGGCGCGGGTGCATCT
CGCGGCTACCGAGGAAGGACATCACCGGCGTCGGCGTGTCGCCGGGCTGCTCCTCGAGCACCGAGAAATCGACGCTCTTGGCATCCAGCCGCGGCGGGGT
GCCGGTCTTGAGGCGATCGACCCGAAACGGCAGCTCGCGCAGGCGGCTGGCCAGGGCGTTGGAGGGCGGGTCGCCGGCGCGGCCGCCGCGGCTCTGGTCG
AGCCCGATGTGGATCACCCCACCGAGGAAGGTGCCGGTACACAGCACCACGGTTTCGGCACGGAAGCGGATGCCGGTCTCGGTGACCACGCCACGCACTG
TGTCGCCATCGACGATCAGATCCCCGGCCGCCTGCTGGAAGATCGTCAGGTTAGGGGCGTTCTCCAGCATGCCGCGAATGGCCGCCTTGTAGCGTACACG
GTCGGCCTGGGCGCGGGTGGCACGCACCGCCGGTCCCTTGCGTGCGTTGAGCACACGGAACTGAATACCGCCGAGATCGGTGGCCAGGCCCATGGCGCCG
CCCAGCGCATCGATCTCCTTGACCAGATGGCTCTTGCCGATCCCGCCGATGGCGGGATTGCAGGACATCTGACCGAGGGTCTCGATGTTGTGGGTCAGCA
GCAGGGTCTGACAGCCCATGCGGGCGGAGGCCAGGGCGGCTTCGGTTCCCGCATGGCCGCCGCCGATGACGATGACGTCAAAGCGGTCGGGATACTCCAA

GGTTCACCTCGTTGCGCTTGCGCGCGGATATTGCCTAGGGATGAAAAACAGCCTGCCAGTATAAACCGACTGTGGGTAACCGTCAGGGTTTTCCACACCG
CGTTTTGTCAGTCGGGCCGATCTACCATCCACTAAACAAAACAGAATGATTAAAAGAGAGAAGTTCTGCTGTGGTAGTGACTATTGGTGTGGACAAATTC
**TGTGGGTAA**CCGTGAATAGTGTAAAAATTTCAAAATCTTGCATTGTTGACTTTCATGCTGAGAAGAGGGTGACGAGCTGAGGGCGAGCGGTGTCGAGACT
GTGCATAAGAGAGCGGCTTTTCCACAATGCCAACGCTACACCGCTTTTCCACCGCTGCCTACCGGGCTTGTCACCGTGGCTGTGAACAATCGTGCTGGCG
AGTGCAAGAGCCACGCCTCATGCGGCTTGTCGGCCAATGGGGAGCGTGTTGCGAGGAAATTTATCCACAGGCGAAAAAAAGCCCTGTCGAAAACGACAGG

GCTTGCGATCGAGCTCAAAAGTGGGGGCAGCCGGGATCAGACCATGAAGGAGGCACCGCAGCCGCAGGTGGTTGTCGCGTTGGGGTTCTGCACCAGGAAG
CGAGCGCCCGCGAGTCCCTCTTCGTAGTCCACGGTGGACCCGACGAGGTATTGATAGGAGAGTTCGTCGACCACCAGGGCCACCTCACCGAATTCGATGA
CGGTATCGTCCTCGCCTACGGTGTCGGCAAAGTCGAAGCCGTACTGGAAACCCGAGCAGCCGCCGCCCGTCACGTAGACGCGCAGCTTGAGTTCGGGCTT
GCCCTCCTCCGCAATCAGCGCCTGGAGCCGCTTCCTGGCGGCTGCGCTGAGCATCATGGGAGTGGGGACGAAAGATTCTGCGCCGCTCATGGGCATCCTC
CCGTGAGCCGAATGAATGGATAACAGTCGATTATCTTCAATTCCTAGCAAAAGGGTCAACTATTCAGGAAAAGGACTCGCCGATGACGCCATCCCTGGCG
TCGAACTTCCCTGCGGTCTGACCCCGGAGCCCCAATATCTCCGGGTGAAATCATCAGGGCATCAGGGCGGGGGCGGCGAGTCCGGCATGCTCGTCGAAGC
CGAACATCAGGTTGAGGTTCTGCACGGCCTGCCCGGAAGCGCCCTTGACCAGATTGTCGATCACCGAGAGCACTACTACGGTGTCGCCGTTACCCGGCCG
ATGCACGGCCAGACGGCAGACGTTGGCGCCCTTGACGCTGCGCGTCTCGGGGTGGCTGCCGGCAGGCATCACGTCGACGAAGGGCTCGTCGGCGAAGCGC
TGCTCGAACAGTGCCTGCAGATCGCCGGGCTCGCCGGTCAGGCGCCCGTACAGGGTGGCATGAATGCCGCGGATCATCGGCGTCAGGTGGGGCACGAAGG
TCAGGCCCACCGGGCCGTTGGCAGCATCGCGGAGGCCCTGGCTGATTTCCGGCAGGTGGCGGTGGCCGGAAGCGCCGTAGGCCTTCATCGATTCGCTGGC

>Halomonas desiderata FB2 [NZ_JAAQTN010000031.1:16981..17050], dnaA> ig dnaN>, 69 bp

WebSkew GCmin pos. 84960 (GCmin<>oriC1 49.5 kb; GCmin<>oriC2 8.8 kb; oriC2<>oriC1 40.7 kb)

>Halomonas daqingensis CGMCC 1.6443 [NZ_FNVC01000006.1:134250..134749], <rpmH oriC1 dnaA>, 513 bp

GGGCAGTGCCGCATAGACGATGTCTCCCCCCTGGGGGTCGATGCGCACGTCGAGCACGTCGGTCACCACGGCGACCAGGTCGCGGGTGCTGGTCGGCTCG
GCAGGCTCCGCCGGCATGTCGCCGGTCATCTCCTGCTCCGCCCTGGGGGCGGCCGGCACGGCCAGCGCGTCGTCATCGGGCGCCTGCGCCTCGCCGGGTG
CCGGGGCGGTGATGGTGGGAGCATCAGGCTCAGGAGCAACCTGGCCATAATCCTGGTTCCACTGTACCACCAGCAGGTAGGTGAGCACGGCCAGTGGAAT
CAGCAATAGGAGTCGTTTTACGTCCATGAGGGGTCTGCCCGGTGGGTAGTGAACATGCCGCTGGAACGAGATCCGCTGCCATGTGAAAGAGGGCGACAGC
CTGCCGGGGCAGGCCATCGAAGCGGAGACTAATACCGCATCCGGTCAGGAGGTCATCGACTTCTGCGCGTCACGCTCCAGGCGCCGCCACATGCCGAAAA
GCTGGCGATGCAGCGTGGCCTTGTCCAGTTCGCTCGCGCCGCGACGTGCCAGCAGGACGATATCCACGGCGGGTAATGAGCGCTGGCGCAGACGTACGGA
TTCGCGCACCACACGTTTCAGGCAGTTGCGATCCACGGCTCTGCGGACGTTCTTCTTGCTGATGACCAGGCCAAGGCGGGGGTGGTCCAGTCCATTGGGG
CACGCCAGGGCCATTAGCCCCTTGCCATGTACCTTGAGTGCAGCGTGATCGAAAACGTATCGATAATCCCCGGCAGTCAGCAGTCGCAGCTGTTTGGGGA
AACCATGACCGGACACGCGCAATCCGCGATCAGGCGCTCAGACGCTTGCGGCCCTTGGCGCGGCGGCGAGCCAGAACGGCACGACCATTCTTGGTAGCCA
TGCGAGCACGGAAGCCGTGGACACGCTTGCGCTTCAGAACGCTGGGTTGAAAGGTGCGTTTCATAACTCTTGATTCCCACGTGGTTTGTTGACGGATCGT

CTTACAGGAATGCCCTGGCTCCCGACAGGAACCAGGGGCAGATTCGGTTCAAGACCGGAAATTGTAGAGACTTCGGCTGCCCGACGCAATTGTTCCTCAC
GCTCCCGCATCACCACAGGGCGTCCGCAGCGATCGCCGCTGTAGCAAACATCATAGGATGTTTTCTTGAACCTACTGTGACTAATAATGGGGATCGCTTC
**TGTTGATAA**AGTGATATTCACCTTTCCTATCAGAGGCCTGCCCGTTGGCTGAGTCTGTGGAAAAGCGTTGGGGAAGGCGTGCTCATGCTGTGATTCGACT
GCTGATGACTACCTGACTCATCCACAGGTCGAGGTGGTCCGCGGTTGATGAACAGAATGATCACGGGTGCCGCGAGGGCTTTTCCACAGCCATGTTCCGG
CAGTTTCCGCTGTGGATAACCCGGCGGGCGAAGGATACAATGTCCGGTCACGCATCAACGGGTAATGGTGAGGTCGAGTGTCTCTTGCTCTCTGGCAACA

GTGTCTGGATTACCTGCAGGACGAGCTGAATGCCCAGCAGTTCAATACCTGGATTCGCCCGCTTCAAGCCGAGGAGGGCGAAGCCAACGAGCTGCGACTG
CTGGCGCCCAACCGCTTCGTGCGCGACTGGGTCGGCGACAAGTATGCCAAGCGTATCAATGAGCTGCTCAGGGAGCTGGCGCCTGCCAAGCCGCCGCGGG
TGGTGCTGGCCGTGGGCAGTCGCACGGTGGCGGCGGTCAAGCCGCAGACGCGCGAGCTCGGCGCTCCGGTCTCGGCCAATCCACACTCTCCCCTGGCGCC
TGCCGTTCAGGTGCCGCCTCGTGGCGACATCGGCGATGAGCGCGAGATCGACCAGCTGCGCGAGGAGGGCGCCACGGCACGTCGCGGCGGCAGCGAGCGC
GAAGTGCAGGTCGAGGGCAGCCTCAAGCACAGCAGCGGCCTCAACCCCAGCTTCACCTTCGAAACCTTCGTCGAGGGCAAGTCCAACCAGCTGGCTCGAG
CCGCCTCGCGGCAGGTCTCGGAGAACCCCGGTGGCGCCTACAACCCGCTGTTCCTCTATGGCGGCGTCGGCCTGGGCAAGACCCACCTGATGCACGCCGT
AGGCAATGCCCTGGCCTCGAGGCGCGAGAACGCCCGGGTGGTCTATCTGCACTCCGAGCGCTTCGTGGCGGACATGGTCAAGGCACTGCAGCTCAACGCC
ATCAACGATTTCAAGCGTTTCTATCGCAGCGTCGATGCGCTGCTGATCGACGACATCCAGTTCTTCGCCGGCAAGGAGCGCTCCCAGGAGGAGTTCTTCC
ACACTTTCAATGCCTTGCTCGAGGGCGGTCAGCAGATGATCCTCACCTCGGACCGCTACCCCAAGGAGATCAGCGGGGTCGAGGAGCGCCTCAAGTCGCG
CTTCGGCTGGGGGCTGACCGTGGCCATCGAGCCGCCGGAGCTGGAAACCCGGGTCGCCATCCTGATGAAGAAGGCCGATCAGGCCAAGGTGGATCTGCCC

>Halomonas daqingensis CGMCC 1.6443 [NZ_FNVC01000006.1:93564..94063], <mnmG oriC2 <PRK13623 (ErpA), 536 bp

GCTTGCGATCGAGCTCAAAAGTGGGGGCAGCCGGGATCAGACCATGAAGGAGGCACCGCAGCCGCAGGTGGTTGTCGCGTTGGGGTTCTGCACCAGGAAG
CGTTCTCCAGTCCCTTGATCGAGCGCACCACCTGCAGCTGCACGTCGAACGGCAGCGAAGTCGAAATCCCGTTGGGGTAGAGCTCGTGGGTATCGAGGCC
TTCGGGCTCGATGAACACCTGGTGGCTCGCCTTGTCGGCGAAGCGGTGCACCTTGTCCTCGATCGACGGGCAGTAGCGCGGCCCGACCCCCTCGATCACG
CCGGAATACATCGGCGAGCGGTCGAGGTTGGCGAAGATGATCTCGTGGGTGCGCTCGTTGGTGTGCGCAATGTGGCAGCTCACCTGGCGCGGGTGCATCT
CGCGGCTACCGAGGAAGGACATCACCGGCGTCGGCGTGTCGCCGGGCTGCTCCTCGAGCACCGAGAAATCGACGCTCTTGGCATCCAGCCGCGGCGGGGT
GCCGGTCTTGAGGCGATCGACCCGAAACGGCAGCTCGCGCAGGCGGCTGGCCAGGGCGTTGGAGGGCGGGTCGCCGGCGCGGCCGCCGCGGCTCTGGTCG
AGCCCGATGTGGATCACCCCACCGAGGAAGGTGCCGGTACACAGCACCACGGTTTCGGCACGGAAGCGGATGCCGGTCTCGGTGACCACGCCACGCACGG
TGTCGCCATCGACGATCAGATCCCCGGCCGCCTGCTGGAAGATCGTCAGGTTAGGGGCGTTCTCCAGCATGCCGCGAATGGCCGCCTTGTAGCGTACACG
GTCGGCCTGGGCGCGGGTGGCACGCACCGCCGGTCCCTTGCGTGCGTTGAGCACACGGAACTGAATACCGCCGAGATCGGTGGCCAGGCCCATGGCGCCG
CCCAGCGCATCGATCTCCTTGACCAGATGGCTCTTGCCGATCCCGCCGATGGCGGGATTGCAGGACATCTGACCGAGGGTCTCGATGTTGTGGGTCAGCA
GCAGGGTCTGACAGCCCATGCGGGCGGAGGCCAGGGCGGCTTCGGTTCCCGCATGGCCGCCGCCGATGACGATGACGTCAAAGCGGTCGGGATACTCCAA

GGTTCACCTCGTTGCGCTTGCGCGCGGATATTGCCTAGGGATGAAAAACAGCCTGCCAGTATAAACCGACTGTGGGTAACCGTCAGGGTTTTCCACACCG
CGTTTTGTCAGTCAGGTCGATCTACCATCCACTAAACAAAACAAAATGATTAAAAGAGAGAAGTTCTGCTGTGGTAGTGACTATTGGTGTGGACAAATTC
**TGTGGGTAA**CCGTGGATAGTGTATAAATTTCAAAATCTTGCATTGTTGACTTTCATGCTGAGAAGAGGGTGACGAGCTGAGGGCGAGCGGTGTCGAGGCT
GTGCATAAGAGAGCGGCTTTTCCACAATGCCAACGCTACACCGCTTTTCCACCGCTGCCTACCGGGCTTGTCACCGTGGCTGTGAACAATCGTGCTGGCG
AGTGCAAGAGCCACGCCTCATGCGGCTTGTCGGCCAATAGGGAGCATGTTGCAAGAAAATTTATCCACAGGCGAAAAAAAGCCCTGTCGACAACGACAGG

GCTTGCGATCGAGCTCAAAAGTGGGGGCAGCCGGGATCAGACCATGAAGGAGGCACCGCAGCCGCAGGTGGTTGTCGCGTTGGGGTTCTGCACCAGGAAG
CGAGCGCCCGCGAGTCCCTCTTCGTAGTCCACGGTGGACCCGACGAGGTATTGATAGGAGAGTTCGTCGACCACCAGGGCCACCTCACCGAATTCGATGA
CGGTATCGTCCTCGCCTACGGTGTCGGCAAAGTCGAAGCCGTACTGGAAACCCGAGCAGCCGCCGCCCGTCACGTAGACGCGCAGCTTGAGTTCGGGCTT
GCCCTCCTCCGCAATCAGCGCCTGGAGCCGCTTCCTGGCGGCTGCGCTGAGCATCATGGGAGTGGGGACGAAAGATTCTGCGCCGCTCATGGGCATCCTC
CCGTGAGCCGAATGAATGGATAACAGTCGATTATCTTCAATTCCTAGCAAAAGGGTCAACTATTCCGGAAAAGGACTCGCCGATGATGCCATCCCTGGCG
TCGAACTGCCCCTGCGGTCTGACCCCGGAGCCCCAATACCTCCGGGTGACATCATCAGGGCATCAGGGCGGGGGCGGCGAGTCCGGCATGCTCGTCGAAG
CCGAACATCAGGTTGAGGTTCTGCACGGCCTGCCCGGAAGCGCCCTTGACCAGATTGTCGATCACCGAGAGCACTACTACGGTGTCGCCGTTACCCGGCC
GATGCACGGCCAGACGGCAGACGTTGGCGCCCTTGACGCTGCGCGTCTCGGGGTGGCTGCCGGCAGGCATCACGTCGACGAAGGGCTCGTCGGCGAAGCG
CTGCTCGAACAGTGCCTGCAGATCGCCGGGCTCGCCGGTCAGGCGCCCGTACAGGGTGGCATGAATGCCGCGGATCATCGGCGTCAGGTGGGGCACGAAG
GTCAGGCCCACCGGGCCGTTGGCAGCATCGCGGAGGCCCTGGCTGATTTCCGGCAGGTGGCGGTGGCCGGAAGCGCCGTAGGCCTTCATCGATTCGCTGG

>Halomonas daqingensis CGMCC 1.6443 [NZ_FNVC01000006.1:136200..136268], dnaA> ig dnaN>, 69 bp

WebSkew GCmin pos. 420979 (GCmin<>oriC1 42.0 kb; GCmin<>oriC2 6.1 kb; oriC2<>oriC1 35.9 kb)

>Halomonas ethanolica MCCC 1A11081 [NZ_JABFTX010000004.1:378701..379200], <rpmH oriC1 dnaA>, 513 bp

GGGCAGTGCCGCATAAACGATGTCTCCCCCCTGGGGATCGATGCGCACGTCGAGCACGTCGGTCACCACGGCGACCAGGTCGCGGGCGCTGGTCGGCTCG
GCAGATTCCGCCGGCATGTCGCCGGTCATCTCCTGCTCCGCCTGGGGGGCGGCAGGCACGGCCAGCGCGTCGTCATCGGGCGCTTGCGTGTCCCCAGGGG
CCGGGGCGGTGATGGTGGGAGCATCTGGCTCAGGAGCAACCTGGCCATAATCCTGGTTCCACTGTACCACCAGCAGGTAGGTGAGCACGGCCAGTGGAAT
CAGCAATAGGAGTCGTTTTACGTCCATGAGGGTTCTGCCCGGTGGGTAGTGAACATGCCGCTGGAACGAGATCCGCTGCCATGTGAAAGAGGGCGACAGC
CTGCCGAGGCAGGCCATCGAAGCGGAGACTAACACCGCATCCGGTCAGGAGGTCATCGACTTCTGCGCGTCACGCTCCAGGCGCCGCCACATGCCGAAAA
GCTGGCGATGCAGCGTGGCCTTGTCCAGTTCGCTCGCGCCGCGACGTGCCAGCAGGACGATATCCACGGCGGGTAATGAGCCCTGGCGCAGGCGTAAGGA
CTCGCGCGCCACACGCTTCAGGCAGTTGCGATCCACGGCTCTGCGGACGTTCTTCTTGCTGATGACCAGGCCAAGGCGGGGGTGGTCCAGTCCATTTGGG
CACGCCAGGGCCATCAGCCCCTTGCCATGTACCTTGAGTGCCGCGTGATCGAAAACGTTTCGATAATCCCCGGCTGTCAGCAATCGCAGCTGTTTGGGGA
AACCATGACCGGACACGCGCAATCCGCGATCAGGCGCTCAGACGCTTGCGGCCCTTGGCGCGGCGGCGAGCCAGAACGGCACGACCATTCTTGGTAGCCA
TGCGAGCACGGAAGCCGTGGACACGCTTGCGCTTCAGAACGCTGGGTTGAAAGGTGCGTTTCATAACTCTTGATTCCCACGTGGTTTGTTGACGGATCGT

CTTACAGGAATGCCCTGATTCCCGACTGGAACCAGGGGCAGATTCGGTTCAAGACCGGAAATTGTAGAGACTTCGGCTGCCCGACGCAATTGTTCCTCAC
CTTCCCGCATCATCGCAGGGTATCCGCAGAGAACGGCGCTAGAGCAATCATCATAAGAGGTTTTCTTGAAGCTATTGTGACTAATGGTGGGGATCAATTC
**TGTTGATAA**GGTGGTATTTCCCATTCTGATCAGAGGATTGTCTGATGCCCGGTTCTGTGGAAAAGCGTGGGGAAAGGCGTGTTCATGCTGTGACTTGGCT
GTTGATGGTTGCCCGACTCATCCACAGCCTGGGGTGGTCCGCCATTGATGGACAGGATGATCACCGGTGCCACCTGAGCTTTTCCACAGCTGTTTTGCGG
GGTTTTCCGCTGTGGATAACCCGGCGGGCGAAGGATACAATGTCCGGTCACGCATCAACGGGTCATGGTGAGGTCGAGTGTCTCTTGCTCTCTGGCAACA

GTGTCTGGATTATCTGCAGGACGAGCTGAACGCCCAGCAGTTCAATACATGGATTCGCCCTCTGCAGGCCGAGGAGGGCGAAGCCAACGAGCTGCGGCTG
CTGGCGCCCAACCGTTTCGTACGTGATTGGGTCGGCGACAAGTATGCCAAGCGCATCAATGAGCTGCTCAGGGAGCTGGCGCCTGCCAAACCGCCGCGGG
TGGTACTGGCCGTGGGCAGTCGCAAGGTGGCGGCGGTCAAGCCCCAGACGCGCGAACTCGGGGCTCCTGTCTCGGCCAATCCTCACTCCCCCTTGGCGCC
TGCCGTTCAGGTGCCGCCTCGCGGCGACATCGGCGATGAGCGTGAGATCGATCAGCTGCGCGAGGAGGGCGGCACGGCGCGGCGTTCCGGCAGCGAGCGC
GAAGTGCAGGTCGAGGGTAGCCTCAAGCACAGCAGCGGGCTCAACCCCAGCTTCACCTTCGAAACCTTCGTCGAGGGCAAGTCCAACCAGCTGGCTCGGG
CCGCCTCGCGGCAGGTCTCGGAGAACCCCGGTGGTGCCTACAACCCACTGTTCCTCTACGGTGGCGTCGGTCTGGGCAAGACCCACCTGATGCATGCCGT
GGGTAATGCCCTGGCCGCGCGGCGCGAGAATGCGCGGGTGGTCTATCTGCACTCCGAGCGCTTCGTGGCGGACATGGTCAAGGCGCTGCAGCTCAATGCC
ATCAACGATTTCAAGCGTTTCTATCGCAGCGTCGATGCGCTGCTGATCGACGACATTCAGTTCTTCGCCGGCAAGGAGCGCTCCCAGGAGGAGTTCTTTC
ACACTTTCAATGCCTTGCTCGAGGGCGGTCAGCAGATGATCCTCACCTCGGACCGCTATCCCAAGGAGATCAGCGGCGTCGAGGAGCGGCTCAAGTCGCG
CTTCGGCTGGGGCCTTACCGTGGCCATCGAGCCGCCGGAGCTGGAAACTCGGGTCGCCATCCTGATGAAGAAGGCGGATCAGGCCAAGGTGGATCTGCCT

>Halomonas ethanolica MCCC 1A11081 [NZ_JABFTX010000004.1:414629..415128], <mnmG oriC2 <PRK13623 (ErpA), 534 bp

CGTTCTCCAGTCCCTTGATCGAGCGCACCACCTGCAGCTGCACGTCGAACGGCAGCGAAGTCGAAATCCCGTTGGGGTAGAGCTCGTGAGTGTCGAGGCC
CTCGGGCTCGATGAACACCTGGTGGCTCGCCTTGTCGGCGAAGCGGTGCACCTTGTCCTCGATCGATGGACAGTAGCGCGGCCCGACACCCTCGATCACG
CCGGAATACATCGGCGAGCGGTCGAGGTTGGCAAAGATGATCTCGTGGGTGCGCTCGTTGGTGTGCGCAATGTGACAGCTCACCTGACGCGGGTGCATCT
CGCGATTACCGAGGAAGGACATCACCGGGGTCGGCGTGTCGCCGGGCTGCTCCTCGAGCACCGAGAAATCGACGCTCTTGGCATCCAGCCGCGGCGGGGT
GCCGGTCTTGAGGCGATCGACCCGAAACGGCAGCTCGCGCAGGCGGTTGGCCAGGGCGTTGGAGGGCGGGTCGCCGGCGCGGCCGCCGCGGCTCTGTTCG
AGCCCGATGTGGATCACCCCACCGAGGAAGGTGCCGGTACACAGCACCACGGTTTCGGCACGAAAACGGATGCCGGTCTCGGTGACCACGCCACGCACGG
TGTCGCCATCGACGATCAGATCCCCGGCCGCCTGCTGAAAGATCGTCAGGTTGGGGGCGTTCTCCAGCATGCCGCGAATGGCCGCCTTGTAGCGCACACG
GTCGGCCTGGGCGCGAGTAGCACGCACCGCCGGACCCTTGCGGGCGTTGAGTACGCGGAACTGAATGCCACCGAGATCGGTGGCCAGGCCCATGGCACCG
CCCAGCGCATCGATCTCCTTGACCAGATGGCTCTTGCCGATCCCGCCGATGGCGGGATTGCAGGACATCTGACCGAGTGTCTCGATGTTGTGGGTCAGCA
GCAGGGTCTGACAGCCCATGCGGGCGGAGGCCAGAGCGGCTTCGGTTCCCGCATGGCCGCCGCCGATGACGATGACGTCAAAGCGGTCGGGATACTCCAA

GGTGCACCTCGTTGCGCTTGCGCGCGGATATTGCCTAGGGATGAAAAACAGCCTGCCAGTATAAACCGACTGTGGGTAACCGTCAGGGTTTTCCACACCG
GGTTTGCTCGTCAGACGCGATCTACCACTCACTAAACAAAACAGAATGATTAAAAGAGAGAAGTTCTATTGTGGTTGTGACTACTGGTGTGGACAAAATC
**TGTGGATAA**CAGTTGATACCACATAGATTTCATGTTCTTGCATTGTTGACTTTCATGGTGAGAAGAGGATGACGAGCTGAGGGCGAACGGTGGCGAGGCT
GTGTATAAAAGAGGGGCTTTTCCACAGCGCCAACGCTACACCGCTTTTCCACCGTTGCCTACCGTGCTTGTCACCGCAGCTGTGAACAGTCGAGCTGGAA
ACGGCAAAAGCCCGGGCTCATGCGGCTTGCCAGCCCCTGGGGAGCGCCATGGGAGAAAATTTATCCACAGGCGAAAAAAAAGCCCTGTCGGGAACGACAG

GGCTTACGATCGGGCTTGAAGAGAGGGAGTAGGCTCAGACCATGAAGGAAGCGCCGCAGCCGCAGGTGGTCGTGGCGTTGGGGTTCTGCACCAGGAAGCG
AGCGCCCGCGAGTCCCTCTTCGTAGTCGACGGTGGACCCTACGAGATACTGATAAGAGAGTTCGTCGACCACCAGGGCCACCTCACCGAATTCGATGACG
GTATCGTCCTCGCCTGTGGTGTCGGCAAAGTCGAAGCCGTACTGGAAACCCGAGCAGCCGCCGCCCGTCACGTAGACGCGCAGCTTGAGTTCGGGCTTGC
CCTCCTCCGCAATCAGCGTCTGGAGCCGCTTCCTGGCGGCTGCGCTGAGCATCATGGGAGTGGGGACGAAAGATTCTGCGCCGCTCATGGGCATCCTCCC
GTGAGCCGAATGAATGGATAACAGTCGATTATCTTCAATTCCTAGCAAAAGGGTCAACTATTGCCGAGGGCCACCTGGCGGTGAGCCTCGGCAGTAAGGG
TATATGCAAAGGGTCAGGGCATCAGGGCGGGGGCGGCGAGTCCGGCATGCTCGTCGAAGCCGAACATCAGATTGAGGTTCTGCACCGCCTGGCCGGAAGC
GCCCTTGACCAGGTTGTCGATCACCGAGAGTACCACCACGGTGTCGCCGTTGCCCGGCCGATGCACGGCCAGACGGCAGACGTTGGCGCCCTTGACGCTG
CGCGTCTCGGGGTGGCTGCCGGCGGGCATCACGTCGACGAAGGGCTCGTCGGCGAAGCGCCGCTCGAACAGCGCCTGCAGATCGCCGGGCTCACCGGTCA
GGCGTCCGTACAGGGTGGCGTGAATGCCGCGGATCATTGGCGTCAGGTGGGGCACGAAGGTCAGACCCACCGGGCCGTTGGCGGCATCGGCGAGGCCCTG
GCGGATTTCCGGCAGGTGGCGGTGGCCGGAAGCGCCGTAGGCCTTCATCGATTCGCTGGCTTCGGCCAGCAGTGAGGGCACCTTGGCGCCGCGGCCGGCG

>Halomonas ethanolica MCCC 1A11081 [NZ_JABFTX010000004.1:377182..377250], dnaA> ig dnaN>, 69 bp

WebSkew GCmin pos. 362043 (GCmin<>oriC1 49.0 kb; GCmin<>oriC2 8.3 kb; oriC2<>oriC1 40.7 kb)

>Halomonas aerodenitrificans MCCC 1A11058 [NZ_JABFTV010000003.1:c312723..313222], <rpmH oriC1 dnaA>, 513 bp

GGGCAGTGCCGCATAGACGATGTCTCCCCCCTGGGGGTCGATGCGCACGTCGAGCACGTCGGTCACCACGGCGACCAGGTCGCGGGTGCTGGTCGGCTCG
GCAGGTTCCGCCGGCATGTCGCCGGTCATCTCCTGCTCCGCCCTGGGGGCGGCCGGCACGGCCAACGCGTCGTCATCGGGGGCCTGCGCGTCACCGGGTG
CCGGGGCGGTGATGGTGGGAGCATCTGGCTCAGGAGCAACCTGGCCATAATCCTGGTTCCACTGTACCACCAGCAGGTAGGTGAGCACGGCCAGTGGAAT
CAGCAATAGGAGTCGTTTTACGTCCATGAGGGTTCTGCCCGGTGGGTAGTGAACATGCCGCTGGAACGAGATCCGCTGCCATGTGAAAGAGGGCGACAGC
CTGCCGAGGCAGGCCATCGAAGCGGAGACAAATACCGCATCCGGTCAGGAGGTCATCGACTTCTGCGCGTCACGCTCCAGGCGCCGCCACATGCCGAAAA
GCTGGCGATGCAGCGTGGCCTTGTCCAGTTCGCTCGCGCCGCGACGTGCCAGCAGGACGATATCCACGGCGGGTAATGAGCACTGGCGCAGACGTAAGGA
TTCGCGCACCACACGCTTCAGGCAGTTGCGATCCACGGCTCTGCGGACGTTCTTCTTGCTGATGACCAGGCCAAGGCGGGGGTGGTCCAGTCCATTGGGG
CACGCCAGGGCCATCAGCCCCTTGCCATGTACCTTGAGTGCAGCGTGATCGAAAACGTATCGATAATCCCCGGCTGTCAGCAGTCGCAGCTGTTTGGGGA
AACCATGACCGGACACGCGCAATCCGCGATCAGGCGCTCAGACGCTTGCGGCCCTTGGCGCGGCGGCGAGCCAGAACGGCACGACCATTCTTGGTAGCCA
TGCGAGCACGGAAGCCGTGGACACGCTTGCGCTTCAGAACGCTGGGTTGAAAGGTGCGTTTCATAACTCTTGATTCCCACGTGGTTTGTTGACGGATCGT

CTTACAGGAATGCCCTGATTCCCGACAGGAACCAGGGGCAGATTCGGTTCAAGACCGGAAATTGTAGAGACTTCGGCTGCCCGACGCAATTGTTCCTCAC
CTTCCCGCATCACCTCAGGGCATCCGCAGCGAACGCCACTGGGGTAATCATCATAAGAGGTTTTCTTGAATCTACTGTGACTAATAATGGGGATCGATTC
**TGTTGATAA**AGTAGTATTTGCTTTTCTCATCAGCTGGTTGGCTGATGTCTGGGTCTGTGGAAAAGCGTTGGGAAAGGAGTGCTCATGCTGTGACTGGTCG
GTTGATGATTGTCCGACTCATCCACACGCCGATGTGGTTCGCCATGGATGAACACGATGATCACTGGCGCCACCCGGGCTTTTCCACAGCCATTTTCCGG
CGTTTTCCGCTGTGGATAACCCGGCGGTCGAAGGATACAATGGCCGGTCACGCATCAACGGGTCATGGTGAGGTCGAGTGTCTCTTGCTCTCTGGCAACA

GTGTCTGGATTACCTGCAGGACGAGCTGAACGCCCAGCAGTTCAATACGTGGATTCGCCCGCTTCAGGCCGAGGAGGGCGAAGCCAACGAGCTGCGACTG
CTGGCGCCCAACCGCTTCGTGCGTGACTGGGTCGGCGACAAGTATGCCAAGCGCATCAATGAGCTGCTCAGGGAGCTGGCGCCTGCCAAACCACCGCGGG
TGGTACTGGCTGTGGGCAGTCGCAAGGTGGCTGCGGTCAAACCCCAGACACGCGAACTCGGGGCTCCGGTCTCGGCCAATCCTCACTCCCCCTTGGCGCC
TGCCGTTCAGGTACCGCCGCGCGGCGACATCGGCGATGAGCGCGAGATCGACCAACTGCGCGAAGAGGGCGGCTCGGCGCGGCGTTCCGGCAGCGAGCGC
GAAGTGCAGGTCGAGGGTAGTCTCAAGCACAGCAGCGGTCTCAACCCCAGCTTCACCTTCGAAACCTTCGTCGAGGGCAAGTCCAACCAACTGGCTCGGG
CCGCCTCGCGGCAGGTCTCGGAGAACCCCGGCGGTGCCTACAACCCACTGTTTCTCTACGGTGGGGTCGGTCTGGGCAAGACCCACCTGATGCATGCCGT
GGGCAATGCCCTGGCAACACGGCGCGAGAACGCACGGGTCGTCTATCTGCACTCCGAGCGCTTCGTGGCGGACATGGTCAAGGCGCTGCAGCTCAATGCC
ATCAATGATTTCAAGCGCTTCTATAGAAGCGTCGATGCGCTGCTGATCGACGATATCCAGTTCTTCGCCGGCAAGGAGCGCTCTCAGGAAGAATTCTTCC
ATACTTTCAATGCCTTGCTTGAGGGCGGTCAGCAGATGATCCTGACCTCGGACCGCTATCCCAAGGAGATCAGCGGCGTCGAGGAGCGTCTCAAGTCACG
TTTCGGCTGGGGCCTTACCGTGGCCATCGAGCCACCGGAGCTGGAAACCCGGGTCGCCATCCTGATGAAGAAAGCGGATCAGGCCAAGGTGGACTTGCCT

>Halomonas aerodenitrificans MCCC 1A11058 [NZ_JABFTV010000003.1:c353465..353964], <mnmG oriC2 <PRK13623 (ErpA), 533 bp

CGTTCTCCAGCCCCTTCATCGAGCGCACCACCTTGAGCTGCACGTCGAAGGGCAGCGAGGTGGAGATGCCGTTGGGGTAGAGCTCGTGGGTGTCGAGGCC
TTCAGGCTCGATGAACACCTGGTGGCTCGCCTTGTCGGCGAAGCGGTGCACCTTGTCCTCGATCGATGGACAGTAGCGCGGCCCGACGCCCTCGATCACG
CCGGAATACATCGGCGAGCGGTCGAGGTTGGCGAAGATGATCTCGTGGGTGCGCTCGTTGGTGTGCGCAATGTGACAGCTCACCTGACGCGGGTGCATCT
CGCGATTACCGAGGAAGGACATCACCGGGGTCGGCGTGTCGCCCGGCTGCTCCTCGAGCACCGAGAAGTCGACGCTCTTGGCGTCCAGCCGCGGCGGGGT
ACCGGTCTTGAGCCGATCGACCCGGAACGGCAGTTCGCGCAGACGGCTGGCCAGTGCGTTGGAGGGAGGATCGCCGGCGCGTCCGCCACGGCTCTGATCG
AGACCGATATGGATCACCCCACCGAGGAAGGTGCCGGTGCACAGCACCACCGTTTCGGCATGGAATTGAATACCGGTCTCGGTGATCACGCCACGCACGG
TGTCGCCATCGACGATCAGATCCCCCGCCGCCTGTTGGAAGATCGTCAGGTTGGGGGCGTTCTCCAGCATGCCGCGAATGGCTGCCTTGTAGCGCACTCG
GTCGGCCTGGGCGCGGGTGGCACGCACCGCCGGTCCCTTACGTGCGTTGAGCACGCGAAACTGGATGCCGCCGAGATCGGTGGCCAGACCCATGGCACCG
CCCAGCGCATCGATCTCCTTGACCAGATGGCTCTTGCCGATCCCGCCGATGGCGGGATTGCAGGACATCTGACCGAGGGTCTCGATGTTATGGGTCAGCA
GCAGGGTCTGACAGCCCATGCGGGCGGAGGCCAGAGCGGCTTCGGTTCCCGCATGGCCGCCGCCGATGACGATGACGTCAAAGCGGTCGGGATACTCCAA

GGTTCACCTCGTTGCGCTTGTGCGCGGATATTGCCTAGGGATGAAAAACAGCCTGCCAGTATAAACCGACTGTGGGTAACCGTCAGGGTTTTCCACACCG
GGTTTGCCTGCCAGCCGCGAGCTACCACTCACTAAACAAAACAGAATGATTAAAAGAGAGAAGTTCTGTTGTGGTTGTGACTACTGGTGTGGACAATTTC
**TGTGGATAA**CACAATGTCCTACAGATATATCAGTAAGTTGCATTGTTGACTTTTGTGGTGAGAAGAGGATGACGAGCTGAGGGCGAGCGGTGACGAGGCTT
GTGTATAAAAGAGGGTCTTTTCCACAGCGGCAACGCTACACCTCTTTTCCACCGTTGGCTACCGGGCTTGTCACCGCGACTGTGAACAGTTGCGCTGATG
ACGACAAGATCCCGGCCTCATGCGGTTTGCAGCGCGATCGGGAGCTTCAAGAGAAGAATGTTATCCACAGGCGAAAAAAAGCCCTGTCGGGAACGACAGG

GCTTACGATCGGGCTCGAAGAGAGGGAGAAGGATCAGACCATGAAGGAAGCGCCGCAGCCGCAGGTAGTCGTCGCGTTGGGGTTCTGCACCAGGAAGCGA
GCGCCTGCGAGTCCCTCTTCGTAGTCGACGGTGGACCCTACGAGGTACTGGTAGGAGAGCTCGTCGACCACCAGGGCCACCTCTCCGAATTCGATGACGG
TATCGTCCTCGCCGACGGTATCGGCAAAGTCGAAGCCGTACTGGAAACCCGAGCAGCCGCCGCCCGTCACGTAGACGCGCAGCTTGAGTTCGGGCTTGCC
CTCCTCCGCAATCAGTGTCTGGAGCCGCTTCCTGGCGGCTGCGCTGAGCATCATGGGAGTAGGGACGAAAGATTCTGCGCCGCTCATGGGCATCCTCCCG
TGAGCCGAATGAATGGATAACAATCGATTATCTTAAATTCCTAGCAAAAGGGTCAACTATTCAGGAAAAGGACTCACCAGTGACGGCATCGCTAGGGGCT
GAGCCGTCGGCAGGTCTACGAGGAGGCGCTGTAAACCCATCCCTGGGCGCTATCGACGCCATCCCTGGCGTAGAACCTCCTCTACGACCTGCCCCCGGCT
CCCAATACCTCTGGTGAAATCATCAGGGCATCAGGGCGGGGGCGGCGAGTCCGGCATGCTCGTCGAAGCCGAACATCAGGTTGAGGTTCTGCACCGCCTG
GCCGGAAGCGCCCTTGACCAGGTTGTCTATCACCGAGAGTACCACCACGGTGTCGCCGTTGCCCGGCCGATGCACGGCCAGGCGGCAGACATTGGCGCCC
TTGACGCTGCGCGTCTCGGGGTGGCTGCCGGCGGGCATCACGTCGACGAAGGGCTCGGCGGCGAAGCGCTGCTCGAACAGTGCCTGCAGATCGCCGGGCT
CGCCGGTCAGGCGCCCGTACAGGGTGGCATGAATGCCGCGGATCATCGGCGTCAGGTGGGGCACGAAGGTCAGGCCCACCGAGCCCTTGGCAGCATCGCG

>Halomonas aerodenitrificans MCCC 1A11058 [NZ_JABFTV010000003.1:c311204..311272], dnaA> ig dnaN>, 69 bp

WebSkew GCmin pos. 74520 (GCmin<>oriC1 38.3 kb; GCmin<>oriC2 9 kb; oriC2<>oriC1 29.3 kb)

>Halomonas kenyensis DSM 17331 [NZ_JACEFT010000002.1:112576..113075], <rpmH oriC1 dnaA>, 481 bp

TCATGCGATGCTGGGGCAGGGCAGCATAGACGATGTCGCCCCCTTGAGGGTCGATGCGCAGGTCGAGCACATCGGTGACTACCGCGATCAGGTCGCGAGT
CCCCGTCGGCTGCTCCGGCTCGGCCGGCATCTCACCAGCCATCTCTTGTTCCGCCTGGGGTGCGGCGGGCACCGCCAGGTCATCGTCATCAGGCGCCTGG
GCGTCGTCGACGGCCGGTGCGGTGATGGTCGGAGCCTCTGGCTCAGGAGCAACCTGGCCATAATCCTGGTTCCACTGCACGACCAGCAGATAGGTCAGCA
CGGCCAGTGGAATCAGCAATAGGAGTCGTTTTACGTCCATGAGGGTTCTGCCCGGTGGGTAGTGAACATGCCGCTGGAACGAAATCCGCTGCCATGTGAA
AGAGGGCGTCAGCCTGTCGGGACAGGCTGACGAGGCGGCGACAGGCACCGTATCCGGTCAGGAGGTCATCGTTTTTCGTACATCACGCTCAAGGCGACGC
CACATGCCGTGAAGCTGACGATGTAGCGTGGCCTTGTCCAGCTCGCTCGCGCCTCGCCTTGCCAGCAGGACGATGTCGACGGCAGGAAGCGAGCATTGCT
GGAGACGAATGGATTCACGAGCCAACCGCTTGAGGCGGTTGCGGTCCACGGCCTTGCGGACGTTCTTCTTGCTGATGACCAGGCCAAGGCGGGGGTGGTC
CAGTCCATTGGGGCACGCCAGGGCCATCAGCCCCTTGCCATGTACCTTGAATGTAGCGTGATCGAAAACGTATCGATAATCTCCCGCAGCCAACAAGCGC
AACTGCTTGGAGAAACCATGACCGGACACGCGCGATCCGCTATCAGGCGCTCAGGCGCTTGCGGCCCTTGGCACGGCGGCGAGCCAGAACGGCGCGACCA
TTCTTGGTGGCCATGCGCGCACGGAAGCCGTGAACACGCTTGCGCTTCAGAACGCTGGGTTGAAAAGTACGTTTCATAACACTCGATTCCCACGTGGTTT

GTTGACGGTGCGTCTTACAGGAATGCCTTGGCTCCCGGGAGGAACCAAGGGCAGATTCGGTTCAAGACCGGAAATTGTAGAGACTTCGTCTTCCCGACGC
AATTATTACGCGCGTCGTTCGTGGCCGGGTTCGTGAGGGTCCTACTCCAACAAGGCTTTCTTTACTAGCCCTACTTGTGATCAATGCTGGGGATGGTGTC
**TGTGCATAA**CTTCCATAAATACATATAAAACAACAGGGTAATAGAATATCTGATTTGTGGAAAAACCGTTTGCAAGCAGTGCCCTTCCGCTGAGCCACCG
GCTATCCCGGCGTTGGGAGGGAAAGCCGGTAACGCGGGCCGGCTTGCTGGTGTCCGTACGTTTTCCACAGGATGGCGGAGTGGTGAAGGGCGTGTGGATA
ACCCGCCTGTCGAAGGATACAATGGCCGGTCACGCATCAACGGGTATGGTGAGGTCGAGTGTCTCTTGCTCTCTGGCAACAGTGTCTGGATTACCTGCAG

GACGAGTTGAACGCCCAGCAGTTCAACACCTGGATTCGTCCGCTCCAAGCCGAGGAAGGGGAAGCCAACGAGCTGCGCTTGCTGGCGCCCAATCGCTTCG
TGCGCGACTGGGTGGGTGACAAGTATGCCAAACGTATCAACGAGCTGCTTAGAGAGCTCTCGCCGACAAAGCCTCCCCGGGTGGTATTGACGGTCGGAAG
TCGCCAGGCGGCAGCGGTCAAGCCACAGCCGCGTGAGCTGGGCGCCCCGGTGTCAGCGAACCCTCACTCCCCGCTGGCCCCCGCCGTGCAGGTGCCGCCG
CGGGGCGATATCGGTGACGAGCGCGAAATCGACCAGATGCGCGAAGAGGGCGCCTCTCCACGCCGCAGTGATCGCGAGGTGCAGGTTGAAGGCAGTCTCA
AGCACAACAGCGGTCTCAACCCAAGCTTCACCTTCGAAACCTTTGTCGAGGGTAAGTCCAACCAGCTTGCCAGGGCGGCTTCTCGACAGGTCTCCGAGAA
TCCCGGTGGCGCCTATAATCCGCTTTTCCTCTATGGCGGGGTAGGGCTTGGCAAGACTCACCTGATGCACGCGGTGGGCAACGCGCTGGCGTCACGTCGC
GAAAACGCACGCGTCGTCTATCTGCACTCGGAGCGCTTCGTGGCAGACATGGTGAAGGCGCTGCAGCTCAACGCCATCAATGACTTCAAGCGTTTCTATC
GCAGCGTGGATGCCCTGCTGATCGATGATATCCAGTTCTTTGCCGGCAAGGAGCGCTCCCAGGAGGAGTTCTTCCATACTTTCAATGCCCTGCTCGAGGG
CGGCCAGCAGATGATCCTTACCTCGGACCGCTATCCGAAGGAAATCAGCGGGGTCGAGGAGCGTCTCAAGTCCCGCTTCGGCTGGGGACTGACCGTGGCC
ATCGAGCCCCCCGAACTCGAAACGCGTGTCGCCATCCTGATGAAGAAGGCAGACCAGGCCAAGGTGGACCTGCCTCACGATGCGGCCTTTTTCATTGCCC

>Halomonas kenyensis DSM 17331 [NZ_JACEFT010000002.1:83276..83775], <mnmG oriC2 <PRK13623 (ErpA), 541 bp

TTTTCCAGCCCCTTGATCGAGCGCACCACCTCGAGCTGCACGTCGAAGGGGAGCGAAGTCGAGATCCCGTTGGGGTAGAGCTCATGAGTCTCCAGCCCCT
CGGGTTCGATGAAGATCTGGTGACTTGCCTTGTCGGCGAAGCGGTGCACCTTGTCCTCGATCGACGGGCAATAGCGAGGCCCGACGCCCTCGATCACCCC
GGAGTACATCGGCGAGCGGTCCAGGTTGGCGAAGATGATCTCGTGGGTACGCTCGTTGGTGTGGGCGATGTGGCAGCTCACCTGGCGTGGATGCATGTCA
CGACTGCCCAGGTACGACATCACCGGCGTCGGCGAATCGCCGGGCTGCTCCTCGAGCACGCTGAAATCGACGCTCTTGGCGTCCAGTCGTGGCGGTGTAC
CGGTCTTGAGTCGGTCGACCCGGAAGGGCAACGCGCGCAGCCGCTCGGCGAGGGCATTGGAAGGCGGGTCACCGGCGCGCCCACCGCGGCTCCGGTCGAG
GCCGATGTGAATCACCCCGCCGAGGAAGGTGCCGGTACACAGCACCACGGTTTCGCTCAGGAAGCGAATTCCGGTTTCAGTCACTACCCCGTGCACGCTG
TCACCATCGACGATCAGATCGCCGGCCGCCTGCTGGAAGATCGTCAGGTTCGGCTGGTTTTCCAGCATGCTGCGGATGGCGGCCTTGTAGCGCACGCGAT
CGGCCTGCGCTCGGGTGGCGCGAACCGCCGGCCCCTTGCGGGCATTGAGTACGCGAAACTGAATGCCTCCCAGATCGGTGGCCAGGCCCATGGCACCGCC
CAGTGCATCGATCTCCTTGACCAGATGGCTCTTGCCGATCCCGCCAATGGCGGGATTGCAGGACATCTGGCCCAGGGTCTCGATGTTGTGGGTCAGCAGC
AGGGTCTGACAGCCCATGCGGGCGGAGGCCAAGGCGGCTTCGGTTCCCGCATGGCCGCCGCCGATGACGATGACGTCAAAGCGGTCGGGATACTCCAAGG

TACACCTCGTTGCGCTTGCGCGCGGATATTGCCTAAGGGATGAAAAACAGCCTGGCAGTATAAACCGCCTGTGGGTAAGCGTCAGGGGTTTTCCACACCG
TGTCCGCCACGAGCCGCATCTTCATTGGTCACTAAACAAAACAGAATGATTAAAAGAGAGAAGTTCTGTTGTGGTTGTGACTATTGGTGTGGACAAAAAC
**AGTGGATAA**GCCAGGAATTAGCTTTAATTCCAATGGGATGGATTGTTGATGCCTGTGGTAAGAAACGGCGGGCGAGCTGAGGGCGAACGGTGTCGAGGCT
GTGGATGAAACCAGCCTTTTTCCACAGAGCAAACGCTACACCGTTTTTCCACTGCCGGCTACCCGGCTTGTCACCATGGCTGTGAACAGCCGTGGTGAGC
GAGCTCAGAGGCGCTCACTGCGCGGCCGGGAAGGGATTATAGGCACTAAGTGGAAAAATGTTATCCACAAGCAAAAAAAAGCCCTGTCGTGGCCGACAGG

GCTTGGAACAAGGCTGGTTGCGGTGGGGCAAGCCTGAAATCACACCATGAAGGAGGCGCCGCAACCGCAGGTGGTCGTGGCGTTGGGGTTCTGGACCAGG
AAGCGAGCTCCGGCCAAGCCTTCCTCATAATCGACGGTAGAGCCGACCAGGTACTGGTAGGAGAGTTCGTCGATTACCAGACCTACATCACCGAATTCGA
TGACGGTATCGTCCTCGCCGATGTCATCGGCGAAGTCGAAGCCGTACTGGAACCCGGAGCAGCCACCGCCCGTCACGTAGACGCGCAGCTTGAGGTCGGG
TTTGTTCTCCTCCGCAATCAGCGCCTGGAGCCGCTTCCTGGCGGCATCAGAGAGCATCATGGGAGTGGGAACGAAAGATTCTGCGCCGCTCATGGGCATC
CTCCCGTGAGCCGAATGAATGAGTAACAGTCGATTATCTTCAATTCCTAGCAAAAGGGTCAACTATTCAGGCGCCCCAATACTCCCGGGTGAAATCATCA
GGGCATCAGTGCAGGGGCGGCGAGTCCAAAATGCTCGTCGAATCCGAACATCAGGTTGAGGTTCTGCACCGCCTGGCCGGAAGCGCCCTTGACCAGGTTG
TCGATCACCGAGAGCACCACCACGGTGTCGCCGTTGCCGGGGCGATGCACGGCCAGGCGGCAGACGTTGGCGCCCTTGACGCTGCGCGTCTCGGGGTGGC
TGCCGGCGGGCATCACGTCGACGAAGGGCTCGTCGGCGAAGCGTTGCTCGAACAGCGCCTGTAGATCGCCGGGCTCGCCGGTCAAGCGCCCGTAAAGAGT
GGCGTGAATGCCGCGAATCATCGGTGTCAGGTGCGGCACGAAGGTCAGGCCCACCGGCCCACGGGCCGCATCGCGAAGGCCCTGACTGATTTCCGGCAGG
TGGCGGTGTCCGCTGGCGCCGTAGGCCTTCATCGATTCGCTGGCTTCGGCCAGCAGCGAGGGCACCTTGGCGCCGCGGCCGGCGCCGGTGACGCCCGACT

>Halomonas kenyensis DSM 17331 [NZ_JACEFT010000002.1:114501..114563], dnaA> ig dnaN>, 63 bp

< page 1 | #054 | Halomonas saliphila LCB169

WebSkew GCmin pos. 186880 (GCmin<>oriC1 35.1 kb; GCmin<>oriC2 9.1 kb; oriC2<>oriC1 26.0 kb)

>Halomonas saliphila LCB169 [NZ_PJRN01000005.1:221778..222277], <rpmH oriC1 dnaA>, 509 bp

GGGCAGTGCGGCATAGACGATGTCGCCGCCCTGAGGATCGATGCGCACATCGAGCACGTCGGTGGCCACGGCAACCAGGTCGCGCGTACCGGAGGATTGC
TCCGTCTCGCTCGGCATGTCGCCCGTCACTTCGCTATCCGCTCGGGGGGAGGCGGGAACGGCCAACCCGTCCTCGTCCGGAACCGTCGCTTCGTCGCCTG
GGGCCGGAGCGGTGAAAGAAGGTGCTTCAGGTTCAGGAGCAACCTGACCGTAGTCCTGATTCCACTGCACGACCAGCAGGTAAGCGAGTACGGCCAGTGG
AATCAGCAATAGGAGTCGTTTTACGTCCATGAGGGATCTGCCCGGTGGGTAGTGAACATGTCGCTGGAACGATATCCGCTGCCATGTGGAAAGGGCGATA
GCCTGCTCGAGGCAGGCTATCGTGGCGGCGGCAGGCACCACATCCGGTCAGGATGCCATCGACTTGCGTACGTCACGCTCCAAGCGACGCCACATGCCGA
ACAGCTGGCGATGAAGCGTGGCGTTGTCCAGCCCGTCCGCGCCGCGCCGCGCCAGGAGCACGATATCGACGGCGGGGAGCGAGTGTTGCTGAAGACGAAT
GGATTCGCGCGTCAGTCGCTTGAGACGATTGCGATCCACAGCCTTGCGGACGTTTTTCTTGCTGATGACCAGGCCAAGGCGGGGGTGGTCCAGTCCGTTG
GGGCACGCCAGGGCCATTAGCCCTTTGCCATGTACCTTGAATGCAGCGTGATCGAAGACGTATCGATAATCCCCGGCCGACAGCAGTCGTAGTCGCTTGG
GGAAGTCATGACCGGACACGCGCGATCCGCGATCAGGCGCTCAGGCGCTTGCGGCCCTTGGCACGACGGCGAGCCAGCACGGCGCGACCGTTCTTGGTGG
CCATGCGCGCACGGAAGCCATGCACACGTTTGCGCTTGAGAACGCTGGGTTGAAAAGTGCGTTTCATAACTCTTGATTCCCACGTAATGAGTTGGACGGA

ACGTTGACAGGAACGCCCTGACTCGTACGGGGAGCCAGGGGGAGATTCGATTCAAGACCGGAAATTGTAGAGACTTCGGCAGCGCGACGCAATCGTTCCG
CTCCTGCCCTTGGTTTTCGGCTAGCCGTGCGCCGGGAACGGTCGCTGGCACGAGTGAGTGTTAACCTTTCGATACTGTGATTAAGGTTGGGGACGTTGCC
**TGTTGATAA**GATTTAATTTTTCCTATTCACCAATAAGTTGAAGTAATGTTCCTATTGTGGAAAAGCGGTTAGAAGCCGGTGATCAGCCTGTGGGAGGATC
AAGAATGTGAGCCTGATGATTCGCGGTGCCTCGCTGAGCCTTGGTCACGGACGAGCTCTCCAAAGCGAACGGGTGGGCTTTTCCACAACCTTTTCCCGGG
GCGTGCCGCATGTGGATAACCCGGCGGGCGAAGGATACAATGACCGGTCATGCATCAACGGGGATGGTGAGGTCGAGTGTCACTTGCTCTCTGGCAACAA

TGTCTGGATTACCTGCAGGACGAGTTGAACGCTCAGCAGTTCAATACCTGGATTCGCCCGCTTCAGGCCGAGGAAGGGGAGGGGAACGAGCTGCGTCTGC
TGGCGCCCAATCGCTTCGTGCGCGACTGGGTGGGTGACAAGTATGCCAAGCGCATCAATGAACTGCTCAGAGATCTGGCGCCAGCCAAGCCGCCCCGGGT
GATGCTGACCGTCGGCAGCCGCCAGTCGGCTGCGCCCAAGCCGCAGCCGCGTGAACTCGGCGCGCCGGTGTCGGCCAGCCCCCACTCCCCGCTGGCACCG
GCCGTGCAGGTGCCGCCCAGGGGCGACATCGGCGACGAGCGCGAGATCGATCAGCTGCGCGAAGAGGGCGCTTCCGCACGCCGGCCCGGAGAGCGTGAAG
TTCAGGTAGAAGGCAGTCTCAAGCACAACAGCGGACTCAATCCCAACTTCACCTTTGAAACCTTCGTCGAGGGCAAGTCCAACCAGCTCGCCCGGGCGGC
CTCGCGACAGGTATCGGAGAACCCGGGCGGCGCCTACAATCCATTGTTTCTATATGGCGGAGTGGGCCTGGGCAAGACTCACCTGATGCATGCGGTGGGC
AATGCACTGGCCATACGGCGCGAGAACGCGCGAGTCGTCTATCTCCACTCCGAGCGCTTCGTGGCCGATATGGTCAAGGCGCTGCAGTTGAACGCCATCA
ACGACTTCAAGCGTTTCTATCGCAGCGTCGACGCGCTGCTGATCGACGATATCCAGTTCTTCGCCGGAAAGGAACGCTCACAGGAAGAATTCTTCCATAC
TTTCAATGCCTTGCTGGAAGGCGGCCAGCAGATGATTCTGACGTCGGATCGCTATCCCAAGGAGATCAGCGGTGTCGAGGAGCGTCTGAAGTCGCGCTTC
GGCTGGGGCCTGACGGTGGCCATCGAGCCGCCGGAGCTGGAAACCCGGGTCGCCATCCTGATGAAGAAGGCCGACCAGGCCAAGGTCGACCTGCCTCATG

>Halomonas saliphila LCB169 [NZ_PJRN01000005.1:195781..196280], <mnmG oriC2 <COG1126 (glnQ), 555 bp

TTTCCAGCCCCTTGATCGAGCGCACCACCTGGAGCTGCACGTCGAAGGGCAGCGAGGTGGAGATGCCGTTGGGGTAGAGCTCGTGGGTATCGAGTCCTTC
CGGTTCGACGAACACCTGATGGCTCGCCTTGTCGGCGAAACGGTGCACCTTGTCCTCGATCGACGGGCAGTAACGCGGCCCGACGCCTCCGATCACGCCG
GAGTACATGGGCGAGCGGTCGAGGTTGGCGAAAATGATCTCGTGGGTGCGCTCGTTGGTGTGGGCAATGTGGCAGCTCACCTGGCGCGGATGCATGTCGC
GCCGGCCCAGATAGGACATCACCGGCGTCGGCGTGTCGCCGGGCTGCTCCTCGAGCACCGAGAAATCGACGCTCTTGGCATCCAGCCGCGGCGGGGTGCC
GGTCTTGAGCCGGTCGACCCGAAACGGCAGCGCGCGCAGGCGGGTCGCCAGAGCGTTCGAGGGCGGGTCGCCGGCGCGGCCGCCGCGACTCTGCTCAAGA
CCGATATGAATCACTCCACCGAGGAAGGTGCCGGTGCACAGCACCACCGTTTCGGCACGCAAGCGAATGCCGGCTTCGGTTACCACGCCGCGGACCGTGT
CGCCGTCGACGATCAGGTCGCCCGCAGCCTGCTGGAACAGCGTCAGATTGGGCTGGCTTTCCAGCATACCGCGTATGGCCGCCTTGTAGCGTACGCGGTC
GGCCTGAGCGCGGGTGGCGCGTACCGCCGGCCCCTTGCGGGCGTTGAGCACACGGAACTGAATACCGCCGAGATCGGTGGCCAGCCCCATGGCGCCGCCC
AACGCATCGATCTCCTTGACCAGATGGCTCTTGCCGATCCCGCCGATGGCGGGGTTGCAGGACATCTGGCCGAGAGTCTCGATGTTGTGGGTCAACAACA
GGGTCTGACAGCCCATGCGGGCGGAGGCCAGAGCGGCTTCGGTACCCGCATGGCCGCCGCCGATGACGATGACGTCAAAGCGGTCGGGATACTCCAAGTT

ACACCTCGTTGCGCTTGCGCGCGGATATTTCCATAGGGGGTGAAAAACAGCCTGGCAGTATAAACCGCCTGTGGGTAAGCGTCAGGGGTTTTCCACATTC
GGTTCGCCGGCGAGCTTTCAATCTCACCAGACTAAACAAAACAGAATGATTAAAAGAGAGAAGTTCTGTTGTGGTTGTAACTACTGGTGTGGACAAAAAC
**GGTGGATAA**CCTTAAAAATCACTTATGCAACATTAGGATGCATTGTTGACGGCTGCGGGGGAAAGCGGCTGACGCCCTGAGGGTGAACGGTGTCGAGCCT
GTGGATGAATGGAGCGGTTGCCCACAGGCGAGACGCTACACGCTTTTTCCACCGCCGGCTACCGGGTTGTCACCGCAACTGTGAACAACCGGATGTTGC
AGTCGGCAATGGCGCAACCGTGCGGCTGCGAGAGCAAAACAGGGCAATAAGCGTGAAAAGATATCCACAGGCATGAACAGGCATAAAAAAGGCCCTGTCG

ATCGCGACAGGGCCTGCACGGCAATTTCCGCAAGCGCTTGCGAATAGAGTTATCAGTGCTTGAGCACCCGCGACAGGAAGTTCTGGGTACGCTCGTGCTG
TGGTGAATCGAAGAGGATGTCGGGAGCCGCCTGCTCGACGATTTCGCCCTTGTGAATAAAGACCACCCGATCGGCGACCTCGCGGGCGAAGCCCATCTCG
TGTGTGACCACGACCATGGTCATGCCCTCTCGTGCCAGCTCGCGCATGGCGTCGAGGACCTCGCCGATCATCTCCGGGTCGAGCGCCGAGGTGGGCTCGT
CGAACAGCATCAGGCGTGGCTCCATGGCGAGTGCCCGCGCCAGAGCGACGCGCTGCTGCTGGCCGCCCGAGAGCTGGCTGGGGTATTTGCTTGCCTGATC
GGCGATACCGACCCGCTCGAGCAGACGTTCGGCGGTCTCGACCGCGTCGGCGCGGCTCCAGCCGCGCACTTTCATCGGCGCCAGCGACACGTTGTCGAGT
ACGCTGAGGTGGGGAAAGAGATTGAACTGCTGAAACACCATGCCGACTTCGGTGCGGATCCGCTGCAGTGCCCGGTTGCTCTTGCCGTTGGGCAGCAGCT
CGTTGCCGTCGACCTCGATGTGCCCATGCTGGAACTCCTCGAGGCCGTTGATGCAGCGAATCAAGGTGGATTTGCCGGAACCGCTGGCGCCGATCACCAC
GACCACTTCGCCGGGTTGGATCGCGAGGTCGATGTCCTTGAGCACGTGCAGGTCGCCGAAGTACTTGTTGACCTTCGCCATGTGCACGATGGGTTCGGCG
CTGGAAGGGGAGTTGTCTTGGGTCATGTCGCGCTCCTTATTGTCCGACCAGGCCCTTGCGCTCGAGCAGGCGCAATGCGAAAGACAGGCTCAGCGTTATC
GCCAAGTAGAGGAGGGCGACCATCAGGTAGACTTCCAGGGCGTTGAAGGTGGTGGCGATGTAGACCTGCCCCTGGCGCACCAGCTCGCCCACGCCGATCA

>Halomonas saliphila LCB169 [NZ_PJRN01000005.1:223724..223785], dnaA> ig dnaN>, 62 bp

< page 1 | #055 | Halomonas pellis L5

WebSkew GCmin pos. 84832 (GCmin<>oriC1 33.5 kb; GCmin<>oriC2 8.5 kb; oriC2<>oriC1 25.0 kb)

>Halomonas pellis L5 [NZ_VTPY01000006.1:c51035..51534], <rpmH oriC1 dnaA>, 508 bp

TGCTGAGGCAGCGCGGCATAGACGATGTCGCCTCCCTGGGGATCGATGCGCACATCGAGGACGTCGGTGGCGACGGCGATCAGGTCGCGAGTCGCGGAGG
GTTGTTCCGCTTCACCCGGCATGTCGCCCGCCACGCTATCCGCTCGGGGGGAGGCGGGAACGGCCAATCCGTCCTCGTCGGGAGGCGTCGCTTCGTCGCC
AGGGGCCGGAGCGGAGAAAGAAGGTGCTTCAGGTTCAGGAGCAACCTGACCGTAGTCCTGATTCCACTGCACGACCAGCAGGTAGGCAAGTACGGCCAGT
GGAATCAGCAATAGGAGTCGTTTTACGTCCATGAGGGGTCTGCCCGGTGGGTAGTGAACATGTCGCTGGAACGATATCCGCTGCCATGTGGAAAGGGCGA
TAGCCTGCCCGAGGCAGGCTATCGTGGCGGCGGCAGGCACCGCATCCGGTCAGGATGCCATCGATTTGCGTACGTCACGCTCCAAACGACGCCACATGCC
GAACAGCTGGCGATGAAGCGTGGCGTTGTCCAGTCCGTCCGCGCCGCGCCGTGCCAGGAGCACAATATCGACGGCAGGGAGCGAGCGCTGCTGAAGACGA
ATGGATTCGCGCGTCAATCGCTTGAGACGGTTGCGATCCACAGCCTTGCGGACGTTTTTCTTGCTGATGACCAGGCCAAGGCGGGGGTGGTCCAGTCCAT
TGGGGCACGCCAGGGCCATCAGCCCTTTGCCATGTACCTTGAATGCAGCGTGATCGAAGACGTATCGATAATCCCCGGCCGACAGCAGTCGCAGTCGCTT
GGGGAAGCCATGACCGGACACGCGCGATCCGCGATCAGGCGCTCAGGCGCTTACGGCCCTTGGCACGACGGCGAGCCAGCACGGCGCGACCGTTCTTGGT
GGCCATGCGCGCACGGAAGCCATGCACACGCTTGCGCTTGAGAACGCTGGGTTGAAAAGTGCGTTTCATAACTGATTCCCACGTGATGAGTTGGACGGAA

CGTTTGACAGGAACGCCCTGGCTCGCACGGGGAGCCAGGGGGAGATTCGATTCAAGACCGGAAATTGTAGAGACTTCGGCAGCGCGACGCAATCGTTCCG
CTCCTGGCCTTGGTTTTCGATGAAGCGCTTGCCGGGAGAGGCCACTGGCAAGAGTGTGTGTTCATTCTTCGATACTGTGATTAAGACTGGGGAACCTGCC
**TGTTGATAA**GAACGTATATGTCTTATTTGACAGTGGGTTGTAGTAGAAACTGTGCTGTGGAAAACCGGTTCGGAGCCGGTGGTCAGGCTGTGGAGGGAAT
TAAGGCATGCCGAGCCATCCTCGGCCCCCTTGAGCCCGGCTGCTGGTAGGTTTTTTTGGTGGCATGCAAGCGAACGGGCTTTTCCACAACCTTTTTCCGC
CACCTGTCGCGTGTGGATAACCCGGCGAGCGAAGGATACAATGACCGGTCATGCATCAACGGGGATGGTGAGGTCGAGTGTCACTTGCTCTCTGGCAACA

ATGTCTGGATTACCTGCAGGACGAGTTGAACGCTCAGCAGTTCAATACCTGGATTCGCCCGCTTCAGGCCGAGGAAGGGGAGGGGAATGAGCTGCGTCTG
CTGGCGCCCAATCGCTTCGTACGCGACTGGGTGGGCGACAAATATGCCAAGCGCATCAATGAGTTGCTAAGAGATTTGGCACCAGCCAAGCCGCCCAGAG
TTATGCTGACCGTCGGCAGCCGGCAATCTGCTGCGCTCAAGCCGCAGCCGCGCGATTTGGGAGCACCGGTTTCGGCCAATCCCCACTCTCCACTGGCGCC
GGCCGTGCAGGTGCCGCCCAGGGGCGACATTGGTGATGAGCGCGAGATCGATCAACTGCGTGAAGAGGGGGTCTCCTCGCGCCGGTCCGGAGAACGTGAA
GTACAGGTAGAGGGCAGCCTCAAGCATAACAGCGGCCTCAACCCCAACTTCACTTTCGAAACCTTTGTCGAGGGCAAATCCAACCAGCTCGCTCGTGCCG
CCTCGCGGCAGGTATCGGAAAACCCGGGCGGCGCTTACAACCCGCTGTTCCTGTACGGCGGGGTAGGCCTGGGCAAGACCCACCTGATGCACGCGGTTGG
CAACGCGCTGGCGACGCGACGCGAGAACGCACGGGTCGTTTACCTCCACTCCGAGCGCTTCGTGGCCGATATGGTCAAGGCGCTGCAGCTGAACGCCATC
AACGACTTCAAGCGCTTCTATCGCAGCGTCGATGCGTTGCTGATCGACGATATCCAGTTCTTTGCCGGCAAGGAGCGTTCACAGGAGGAATTTTTCCATA
CTTTCAATGCCTTACTTGAAGGCGGTCAGCAAATGATCCTGACGTCGGATCGCTATCCCAAGGAGATCAGCGGCGTCGAGGAGCGTCTTAAGTCGCGTTT
CGGCTGGGGGCTGACCGTGGCGATCGAGCCGCCGGAGCTGGAAACCCGAGTGGCGATCCTGATGAAGAAGGCTGACCAGGCCAAGGTCGACCTGCCTCAT

>Halomonas pellis L5 [NZ_VTPY01000006.1:c76102..76601], <mnmG oriC2 <COG1126 (glnQ), 521 bp

TTCTCCAGCCCTTCGATCGAGCGCACCACCTGGAGCTGCACGTCGAAAGGCAGCGAAGTGGAAATGCCGTTGGGGTAGAGCTCGTGGGTATCGAGCCCTT
CCGGTTCAATGAACACCTGGTGGCTCGCTTTGTCGGCGAAGCGGTGCACTTTGTCCTCGATCGAGGGACAGTAACGCGGCCCGACGCCTTCGATCATGCC
GGAGTACATGGGCGAGCGGTCGAGGTTGGCGAAGATGATCTCGTGGGTGCGCTCGTTGGTGTGGGCGATATGACAGCTCACCTGGCGCGGATGCATGTCG
CGCCTTCCCAGATAGGACATCACCGGCGTCGGCATATCGCCGGGCTGCTCCTCGAGGGCGGAAAAATCGACGCTCTTGGCATCCAGCCGGGGCGGCGTCC
CGGTCTTGAGCCGGTCGACCCGGAACGGCAGCGCACGCAGCCGAGATGCCAGGGCATTCGAGGGCGGATCGCCGGCGCGACCGCCGCGGCTTTGCTCGAG
ACCGATATGAATCACGCCACCGAGGAAGGTACCGGTGCACAGCACGATCGTTTCGGCATGGAAGCGAATGCCGGTCTCGGTCACCACACCGCGCACCGTA
TCGCCGTCGACGATCAAATCGCCCGCGGCCTGCTGGAACAGCATTAGATTGGGCTGGTTCTCCAGCATGCCGCGAATGGCCGCCTTGTAGCGTACGCGAT
CCGCCTGAGCGCGGGTCGCCCGCACCGCCGGCCCCTTACGGGCGTTGAGCACGCGAAACTGGATGCCGCCGAGGTCGGTGGCCAGCCCCATGGCACCGCC
CAGCGCATCGATCTCCTTGACCAGATGGCTCTTGCCGATCCCGCCGATGGCGGGATTGCAGGACATCTGCCCGAGAGTCTCGATGTTGTGGGTCAGCAGC
AGGGTCTGACAGCCCATGCGGGCGGAGGCCAAAGCGGCTTCGGTTCCCGCATGGCCGCCGCCGATGACGATGACGTCAAAGCGGTCGGGATACTCCACGT

TACACCTCGTTGCGCTTGAGCGCGGATATTCCAAAGGGGGCGAAAAACAGCCTGCCAGTATAAACCGCCTGTGGGTAAGCGTCAGGGGTTTTCCACATCG
CGTTTGCCAACGGACTTGCCGATTCACCAGACTAAACAAAACAGAATGATTAAAAGAGAGAAGTTCTGTTGTGGTTGTAACTACTGGTGTGGACAAAAAC
**GGTGGATAA**CCTGAAAAACTACTTTCGCAACATTAGGATGCATTGTTGACGGATGCGGGGAAAGCGGCTGATGTCCTGAGAGTGGAGGGTGACGAGCCT
GTGGATGAATGGGCTGGTTGCCCACAGGCGAGACGCTACACGCTTTTTCCACTGTCGGCTACCGCGGTTGTCACCGCAGCTGTGAACAACGGCATGCTGC
CGACGGCAATGGCGTAGCTGCGCGGCTCCGGAGGGAAAAAGGGGAGGAAGCGAAAAAAGATATCCACAACCAAAAAAAAGCCCTGTCGGTGCCGACAGGG

CCTGCTTGCAGGAACGCTATCAGTGCTTGAGCACCCGCGACAGGAAGTTCTGGGTACGTTCGTGTTGTGGCGAGTCGAACAACACGTCGGGAGCGGCCTG
CTCGACGATTTCCCCCTTGTGGATATAGACCACTCGATCGGCGACCTCGCGAGCGAAGCCCATCTCATGGGTAACGACCACCATGGTCATGCCCTCCCGT
GCCAGCTCACGCATGGCATCGAGGACCTCACCGATCATCTCCGGGTCGAGCGCCGAGGTGGGTTCGTCGAACAGCATCAGGCGTGGCTCCATGGCGAGCG
CCCGCGCCAGAGCGACGCGCTGTTGCTGGCCGCCCGAGAGCTGGCTGGGGTATTTGCGCGCCTGATCGGCGATACCGACCCGTTCGAGCAAACGTTCGGC
AGTCTCGACCGCGTCGGTGCGGCTCCAGCCTCGCACCTTCATCGGCGCCAGCGAGACATTGTCGAGCACGCTGAGGTGGGGAAAGAGATTGAATTGCTGA
AACACCATGCCGACCTCGGTACGAATCCGCTGCAGTGCTCGGCTGCTCCTGCCGTTGGGCAGCAGTTCGTTGCCATCGACCTCGATATGGCCGTGCTGAA
ACTCCTCGAGGCCGTTGATGCAGCGAATCAGGGTGGACTTGCCGGAACCGCTGGCACCGATCACCACGACCACCTCGCCGGGTTGAATCGTAAGGTCGAT
ATCCTTGAGTACATGCAGGTCGCCGAAAGACTTGTTGACCTTCGCCATGTGCACGATGGGCTCGGCGCTGGAAGGTGAGTTGTCCTGGGTCATGTCGCGC
TCCTTATTGTCCGACCAAGCCCTTGCGCTCGAGCAGCCGCAAGGCGAAAGACAGGCTCAGCGTTATCGCCAGATAGAGGAGGGCGACCATCAGGTAGACT
TCCAGGGCATTGAAGGTGGTGGCAATGTAGACCTGGCCCTGGCGCACCAGCTCGCCCACGCCGATCACCGAGAACAGTGAGGTATCCTTGATACTGATGA

>Halomonas pellis L5 [NZ_VTPY01000006.1:c49524..49587], dnaA> ig dnaN>, 64 bp

< page 1 | #056 | Halomonas chromatireducens AGD 8-3

WebSkew GCmin pos. 3909432 (GCmin<>oriC1 >1 Mb; GCmin<>oriC2 62.0 kb; oriC2<>oriC1 (>1.2 Mb))

>Halomonas chromatireducens AGD 8-3 [NZ_CP014226.1:1152618..1152618], <rpmH oriC1 dnaA>, 501 bp

TTCGGTGCTGGGGCAGCGCGGCGTAAACGATATCGCCCCCCTGGGGATCGATACGCAGGTCCAGGACATCAGTGACCACGGCAATCAGGTCGCGGGTACC
CGTCGCACGTGCCGGTTCGCCAGGCATGTCCCCCGGCACCTCTTCTTCGGCGCGGGGCGCGGCAGGGACGGCCAAGTCATCGTCCGGATCCACGCTCTCT
TCCCCGGGCGCAGGCGCGGTGAAGGTGGGTGCTTCAGGCTCAGGAGCAACCTGGCCGTAGTCCTGGTTCCACTGAATGACGAGCAAGTAGGCAAGCACGG
CCAGTGGAATCAACAAAAGGAGTCGTTTTACGTCCATTGAGAGTTCTGCCCGGTGGGTAGTGAACATGCCGCTGGAACGGTATGCGCTGCCACGTTGATA
TGCGAAAGCCTGCCCAAGGCAGGCTATCGTGGCGGCGACAGGCACCGCATCCGGTCAGGATGCCATCGACTTGCGTACGTCTCGCTCCAGCCGACGCCAC
ATGCTGGAGAGCAGGCGGTGCAGTGTCGCATTATCCAGCTCATTGGCGCCCCGTCGAGCCAGTACCACGATGTCGACGGCGGGAAGCTGATGCTGCTGAA
GGCGAATGGACTCGCGAGTCAGCCGCTTCAGGCGATTGCGATCCACGGCTCTACGAACGTTCTTCTTGCTGAAGACCAGGCCGAGACGGGGGTGGCCCAG
TCCGTTGGGACACGCCAGGGCCATTAGCCCATTGCCATGTACCTTGAATGCAGCGTGATCGAAGACAAATCGATAGTCCCCGGCAGTCAGTAAACGCAGC
TGCGGGGGGAAGTCATGACCGGACACGCGCAATCCGCGATCAGGCGCTCAGGCGCTTGCGGCCCTTGGCGCGACGGCGCGACAGTACGGCACGACCGTTC
TTGGTGGCCATGCGAGCACGGAAGCCGTGCACACGCTTGCGCTTGAGAACGCTGGGTTGAAAGGTGCGTTTCATAACTCTTGATTCCCACGTGGTGTGTT

TGGACGGAATGTCTTACAGGAAATGCCCGGGCTCCTTGCGGGAACCTGGGGAGATTCGGTTCAAGACCGGAAATTGTAGTGACTTCGGGGCTCCGACGCA
ATTCCTCAGCCGCCCAAATTTCAATTGGCATGCCGCCAGCGAATCTATAGTTACTAAATGAATGTTTACTGAAACTGTTATTAATACTGGGGATGCTTTC
**TGTTGATAA**GCCACAAAAACGCTTGATAAACAATAGTGTGCAGTGCAGCATGGCTTGTGGAAAAGCTCTGTGCTTCCGGTAGGGTTTCTGTTGATTCCTT
GTGTGAAATTGTCGTTTTTGTCCACAGCTCGGGCCGCCTTCCTGGCTATACAAGGGTTATCCTGTTGATCCATGTGTGATTTTCCACAGCGCTTTTCGAC
GGCTGCTGCGTGTGGATAACCCGACGGGTGAAGGATACAATGGCCGGTCATGCATCAACCGGATGGTGAGGTCGAGTGTCGCTTGCTCTTTGGCAAAAGT

GTTTGGATTTCCTGCAGGATGAGCTGAACGCCCAGCAGTTCAACACCTGGATCCGTCCGCTTCAGGCGGAAGAGGGGGAGTCCAATGAGCTGCTTCTTCT
GGCACCCAATCGCTTTGTTCGCGATTGGGTTGGGGATAAGTATGCCAAGCGGATCAACGAGCTGCTGCGCGACCTGGCCCCAGCCAAGCCTCCCCGCGTG
GTATTGACGGTCGGCAGCCGCCGCGCCGCTGCTCCCAGGCCGCGAGAGCTCGGAGCGCCGGTCTCGGCCAGTCCGCGATCGCCCCTCGGGCCTGCCGTAC
ATACCCCCCACGGGGAGATATCGGCGACGAGCGTGAGATCGATCGACAGCGAGAAGAGGGGCGTTCGGCACGGCGCGCCACTGGTGAACGAGAGGTACA
GGTCGAGGGTAGCCTCAAGCACAACAGTGGCCTCAACCCCAATTTCACTTTCGAGACCTTCGTCGAGGGCAAGTCGAACCAGCTGGCCAGGGCGGCTTCC
CGTCAGGTCTCGGAAAACCCGGGTGGCGCCTACAACCCGCTATTTCTCTATGGCGGTGTCGGCCTGGGCAAGACACACCTGATGCACGCCGTGGGTAACG
CACTGGCCACCCGCCGGGAGAATGCCACGGTGGTCTATCTCCATTCGGAGCGTTTCGTCGCCGACATGGTCAAGGCGCTTCAGTTGAACGCGATCAACGA
TTTCAAGCGTTTCTATCGTAGCGTCGATGCGCTGCTGATCGACGATATCCAGTTCTTTGCCGGCAAGGAGCGCTCCCAGGAGGAGTTCTTCCACACCTTC
AATGCCTTGCTCGAGGGTGGCCAGCAGATGATCCTGACCTCCGATCGCTATCCCAAGGAGATCAGCGGCGTCGAGGAGCGACTCAAGTCGCGTTTCGGCT
GGGGATTGACGGTGGCGATCGAGCCGCCGGAGCTGGAGACGCGGGTGGCTATCCTGATGAAGAAGGCCGATCAGGCCAAGGTCGACCTGCCCCACGATGC

>Halomonas chromatireducens AGD 8-3 [NZ_CP014226.1:3971200..3971699], <mnmG oriC2 <COG1126 (glnQ), 542 bp

TTTTCCAGTCCGGGGATCGAGCGTACCACCTGCAGCTGCACGTCGAAGGGCAGCGAGGTGGAGATGCCGTTGGGATAGAGCTCGTTGGTGTCGAGCCCTT
CCGGCTCGATGAATATCTGGTGGCTCGCCTTGTCGGCAAAGCGGTGCACCTTGTCCTCGATCGAGGGGCAGTAGCGCGGACCGACCCCCTCGATGACGCC
GGAATACATCGGTGAACGATCCAGGTTGGCCAGGATGATCTCGTGGGTGCGCTCGTTGGTATGCGCGATATGGCAGCTCACCTGCCGCGGATGCATCTCG
CGACTCCCCAGGTAGGACATCACCGGAGTGGGGGTATCACCGGGCTGCTCTTCGAGCACCGAGAAATCGACGTTCTTGGCATCGAGTCGCGGCGGCGTGC
CGGTCTTGAGCCGATCGACCCGGAACGGCAGTGCTCGCAGACGCCTGGCCAGGTCATTGGAGGGCGGATCCCCGGCACGACCGCCACGACTCTGGTCGAG
GCCGATATGGATCACGCCGCCGAGGAAGGTTCCGGTGCAGAGCACCACCGTTTCGCCGAGGAAACGGATACCGGTCTCGGTCACCACGCCACGCACGGTG
TCGCTGTCGACGATCAGGTCACCGGCGGCCTGCTGGAACAGCGTCAGATTGGGCTGATTTTCCAGCATGCCGCGAATGGCTGCCTTGTAGCGGACCCGGT
CGGCCTGGGCTCGTGTCGCCCTCACTGCCGGTCCCTTGCGTGCGTTTAGTACACGGAACTGGATACCGCCCAGGTCAGTTGCCAGGCCCATGGCACCACC
CAGGGCATCGATTTCCTTGACCAAATGACTCTTGCCGATCCCGCCGATGGCGGGATTGCAGGACATCTGTCCCAGGGTCTCGATGTTGTGGGTCAGCAAC
AGGTCTGACAGCCCATGCGAGCGGAGGCCAGGGCGGCTTCGGTTCCCGCATGGCCGCCGCCGATGACGATGACGTCAAAGCGGTCGGGGTAATCCAAGG

TGCACCTCATTGCGCTTCTGCGCGGATATTGCGTAGGGGATGAAAATCAGCCTGGCAGTATAAACCTCCTGTGGGTAAGCGTCAGGGGTTTTCCACACCA
CGTCATCCGGGAGACCTTGATCTTCCAGGTTTCTATAAAATAAATAGAATGATTTAAAGGTAGTTCTGTTGATGTTGTTACTACTAGTGTGGACAAAATC
**TGTGGGTAA**CTGACTAACCCCTTTATTTTTCAGTACTCTGGATTGTTGACTAGTGGCGTGGAAAGGGGCGTGCCTCCTGAGGGCTACCGGTGACGAGCCT
GTGGATGAAATGCAAAGTTACGCACAGGTGGAATCCTGCACCGCTTGTCCACCGCCGGCTACCTGCCTTGTCACCATGGCTGTGAACAAGCTGCTTGATG
ATTAGACAATGCCCCTGCGGCGGGGCTTGCAGAGCGAAAAAGCAGCCTGCGGGAAAAATTTTTTCCCCAGCCCGCAAAAAGCCCCCTCGATAACCCCACA

GGGCCTTTTCAACGCTTTCATCAAGCAAAGCTGTGCTGTTTCAGTGCTTGAGCACCCGCGACAGGAAGCTCTGAGTGCGCTGATTTGCCGGAGAGTCGAA
CACCTGCTTCGGCGGGCCTTCCTCGACGATTTCGCCTCGATGAATGAAGATCACCCGGTCGGCGACCTCCCTGGCGAAGCCCATCTCGTGGGTCACGATA
ATCATGGTCATGCCCTCGCGGGCCAGCTCGCGCATGGCATCGAGAACCTCGCCGATCATCTCGGGATCGAGTGCTGAAGTAGGCTCGTCGAACAGCATCA
GTCGCGGCTCCATGGCCAGTGCCCGGGCCAGGGCAACGCGCTGCTGCTGTCCCCCAGACAGCTGGCTGGGGTACTTGGAGGCCTGGTCGCTGATACCGAC
CCTGTTGAGCAGCCGCTCGGCCGTCTCGGTGGCATCCGCCCGGTTCCAGCCGCGCACCTTCATGGGTGCCAGGGTCACGTTGTCCAGCACGCTGAGATGC
GGGAAGAGATTGAACTGCTGGAAGACCATACCCACCTCGGTACGAATTCGCTGCAGTGCCTTGCTGCTCTTTCCATCGGGTACCAGTTCATTGCCGTCGA
CCTCGATGTGGCCCTGCTGGAACTCTTCCAGGCCGTTGATACAGCGGATCAGTGTAGACTTGCCGGAACCGCTGGCGCCGATGACGACGACTTCCCC
GGGCTTCACCGCCAGTTCGATGTCCTTCAGCACGTGCAGGTCGCCAAAGTGCTTGTTGACTCGCTCGATGCGTACGATGGGATCGGATGTGGACGCTGTC
ATGCAAAGCTCCCTTGCTTCTCTTTATTGACCGACCAGGCCCCTGCGCTCAAGCATGCGCAGCGCGATGGACAGGCTCCAGGTAATGGCCAGATACATCA
GCGCCACCATGAAATAGACCTCCAGGGCGGTAAAGGTCGTGGCAATGTAGATCTGGCCCTGGCGCACTAGCTCACCGACGCCGATCACCGAGAACAGTGA

>Halomonas chromatireducens AGD 8-3 [NZ_CP014226.1:1151107..1151175], dnaA> ig dnaN>, 69 bp

WebSkew GCmin pos. 252255 (GCmin<>oriC1 38.4 kb; GCmin<>oriC2 8.9 kb; oriC2<>oriC1 29.5 kb)

>Halomonas montanilacus PYC7W [NZ_QPII01000003.1:290415..290914], <rpmH oriC1 dnaA>, 502 bp

TCCGATGCTGGGGCAGAGCGGCGTAGACGATATCGCCGCCCTGGGGGTCGATACGCAGGTCGAGCACATCGGTGACTACGGCGATGAGATCGCGGGTGCC
CGTGGCGCTCGCCGGTTCGCCGGGCATGTCGCCCGTTACATCCTCTTCGGCGCGGGGTGCGGCAGGGACGGCCAGGTCGTCGTCCGAATCTACGCGCTGT
TCGCCGGGAGCGGGGGCGGTGAAGGTGGGTGCTTCAGGCTCAGGAGCAACCTGGCCGTAGTCCTGATTCCACTGAATCACGAGCAAGTAGGCAAGCACGG
CCAGTGGAATCAGCAAAAGAAGTCGTTTTACGTCCATTGAGAGTTCTGCCCGGTGGGTAGTGAACATGCCGCTGGAACGGTATGCGCTGCCATGTTGATA
TGCGAAAGCCTGCCCAAGGCAGGCTATCGTGGCGGCGACAAGCACCGCGTCCGGTCAGGATACCATCGACTTGCGTGCGTCTCGCTCCAGCCGACGCCAC
ATGCCGAAGAGCTGGCGATGCAGTGTCGCATTGTCCAGCTCATTGGCACCCCGTCGAGCCAGTACCACGATGTCGACGCCGGGAAGCTGATGCTGCTGGA
GGCGGATGGATTCGCGAGTCAGCCGTTTCAGGCGGTTGCGATCCACGGCCCTGCGGACATTCTTCTTGCTGAAGACCAGGCCGAGACGGGGGTGGCCCAG
TCCGTTGGGGCACGCCAGGGCCATTAGCCCATTGCCATGAACCTTGAATGCAGCGTGATCGAAGACAAATCGATAATCCCCGGCAGTCAGCAGACGCAGC
TGCGGGGGAAAGTCATGACCGGACACGCGCAATCCGCAATCAGGCGCTCAGGCGCTTGCGGCCCTTGGCACGACGGCGCGACAGTACGGCACGACCGTTC
TTGGTGGCCATGCGAGCACGAAAGCCGTGCACACGCTTGCGCTTCAGAACGCTGGGTTGAAAGGTGCGTTTCATAACTCTTGATTCCCACGTGGTGTGTT

TGGACGGAATGTCTTACAGGAAATGCCCTGGCTCCTTGCTGGAACCTGGGGAGATTCGGTTCAAGACCGGAAATTGTAGTGACTTCGGGCCCCCGACGCA
ATTCCTCCGGCGATGCAATTTTGAAACGCCATCCCCGGAGGCTGTAACAGTAATAAAGAGATATTTTACTGATACTGTTATTAATGTTGGGGATTCTTTC
**TGTTGATAA**ACCGGGAAAGCCTTTGTTTACAGATGTGTGCAGCGCAGCATGATCTGTGGAAAAGCACTGCCCTTCCGGTGTCCTCCCTGTTGAGTCCCT
GTGCGAAATTGTCGCTTTTGCCCACAGCTCGTAGACGCTACCCCGACCATACCGGGGTTATCCTGTTGATCCACGTGTAGTTTTCCACAGCGTTCCTCGG
TAACGGCTGCGTGTGGATAACCCGGCGGGTGAAGGATACAATGGCCGGTCATGCATCAACCGGATGGTGAGGTCGAGTGTCGCTTGCTCTTTGGCAACAG

TGTTTGGATTTCCTGCAGGATGAGCTGAACGCCCAGCAGTTCAATACCTGGATCCGTCCTCTTCAGGCGGAAGAGGGGGAGTCCAATGAACTGCGTCTTC
TGGCGCCCAATCGCTTTGTCCGCGACTGGGTGGGGGACAAGTATGCCAAGCGAATCAACGAGCTGCTGCGCGACCTGGCGCCGGCCAAGCCACCCAGGGT
GGTGTTGACCGTCGGTAGCCGAAGCTCCGCTACGGCCCCCAGGCCGCGGGAGCTGGGAGCGCCGGTTTCGGCCAGCCCGCGCTCCCCCCTGGGGCCGGCC
GTACATACCCCGCCGAGGGGAGACATCGGCGACGAGCGTGAGATCGACCGCCAGCGTGAGGAGGGGCGGTCGGCACGACGTGCCGCAGGTGAGCGTGAAG
TGCAGGTCGAGGGTAGCCTCAAGCACAACAGTGGCCTCAATCCAAACTTCACTTTCGAGACATTCGTCGAGGGCAAGTCGAACCAACTGGCCAGGGCAGC
CTCTCGGCAGGTCTCGGAGAATCCGGGTGGCGCCTATAACCCGCTGTTTCTTTACGGTGGTGTCGGCCTGGGCAAGACACACCTGATGCACGCCGTGGGC
AATGCCTTGGCCACCAGGCGGGAGAATGCCACGGTAGTCTATCTCCATTCCGAGCGTTTCGTCGCCGATATGGTCAAGGCACTTCAGCTTAACGCGATCA
ACGATTTCAAGCGTTTCTATCGTAGCGTCGATGCGTTGCTGATCGACGATATCCAGTTCTTTGCCGGCAAGGAGCGCTCCCAGGAGGAGTTCTTTCACAC
CTTCAATGCCTTGCTCGAGGGTGGCCAGCAGATGATCCTGACGTCCGATCGCTACCCCAAGGAGATCAGTGGCGTCGAGGAGCGACTCAAGTCGCGTTTC
GGCTGGGGGTTGACGGTGGCGATCGAGCCGCCGGAGCTGGAGACGCGGGTGGCGATCCTGATGAAGAAGGCCGACCAGGCCAAGGTCGACCTGCCCCACG

>Halomonas montanilacus PYC7W [NZ_QPII01000003.1:260895..261394], <mnmG oriC2 <COG1126 (glnQ), 542 bp

TTTTCCAGCCCGTGAATCGAGCGCACGACCTGCAGTTGCACGTCGAAGGGCAGCGAGGTGGAGATCCCGTTGGGATAGAGCTCATTGGTGTCGAGCCCTT
CCGGCTCGATGAATATCTGGTGACTCGCCTTGTCGGCAAAGCGGTGAAACTTGTCCTCGATCGAGGGGCAATAGCGCGGCCCGACCCCTTCGATCACCCC
GGAATACATCGGCGAACGATCGAGGTTGGCCAGGATGATCTCGTGGGTGCGTTCGTTGGTATGGGCGATGTGGCAGCTCACCTGCCGCGGATGCATCTCG
CGACTTCCCAGGTAGGACATCACCGGGGTCGGTGTATCACCGGGCTGCTCCTCGAGTATCGAGAAATCGACGCTCTTGGCATCGAGTCGCGGCGGCGTTC
CGGTCTTGAGACGATCAACCCGGAAAGGAAGTGCCCGCAGCCGCTGGGCCAGGGCATTGGAGGGTGGATCTCCGGCGCGACCCCCGCGGCTCTGGTCGAG
ACCGATATGGATTACGCCGCCGAGAAAGGTCCCTGTGCAGAGCACCACCGTTTCGCCGAGGAAACGGATGCCGGTTTCTGTTACCACGCCACGTACGGTG
TCGTTGTCGACGATCAGGTCACCGGCGGCCTGCTGGAACAGCGTCAGATTGGGCTGATTTTCCAGCATGCCACGAATGGCGGCCTTGTAGCGCACCCGAT
CAGCCTGGGCTCTTGTCGCCCTCACCGCGGGGCCCTTTCGCGCGTTGAGCACACGAAACTGGATACCGCCCCGATCGGTAGCCAAGCCCATGGCGCCGCC
CAGGGCATCGATTTCCTTGACCAGATGACTCTTGCCGATCCCGCCGATGGCGGGATTACAGGACATCTGCCCCAGGGTTTCGATGTTGTGGGTCAGCAAC
AGGGTCTGACAGCCCATGCGAGCGGAGGCCAGAGCGGCTTCGGTTCCCGCATGGCCGCCGCCGATGACGATGACGTCAAAGCGGTCGGGGTAATCCAAGG

TACACCTCGTTGCGCTTATGCGCGGATATTGCGTAAGGGATGAAAATCAGCCTGGCAGTATAAACCTCCTGTGGGTAAGCGTCAGGGGTTTTCCACACGG
GGTCCGTCGAGAGAGCTCGTTCTTCGGTATTTCTATAAAATAAATAGAATGATTTAAAGGTAGTTCTGTTGATGTTGTTACTACTAGTGTGGACAAAATC
**TGTGGGTAA**CTAAATAAAGTCACCATTTTTCAACACATTGAATTGTTGACTAGTGCCGTGAGAAGCGGCGCGTCTCCTGAGGGTTACCGGTGGCGAGGCT
GTGGATGAAATGGCAGGTTACGCACAAGCGAGATCCTGCACCGCTTGTCCACCGGCGGCTACCCGCCTTGTCACCATGGCTGTGAACAAGCTCATCGATG
ATGAGTCTATGCCCCTGCGATGGGGCTTGCAGAGCGAAAAGTCAGCCTGCGGAAAAAATCTTATCCACAGCCCGAAAAAGGCCCTGTCGATAACCCGACA

GGGCCTTTTTCACCGCGTTCATCAAGCAAGCTGTGCAGATTCAGTGCTTGAGCACCCGCGACAGAAAGCTCTGGGTGCGCTCATTGCGAGGGGAATCGAA
GACCTGCTGTGGCGGTCCCTCCTCGACGATCTCGCCTCGATCAATGAAGATCACCCGGTCGGCGACTTCTCTGGCGAAACCCATTTCGTGGGTCACGATG
ATCATGGTCATCCCCTCGCGGGCCAGTTCACGCATGGCATCGAGAACCTCGCCGATCATCTCGGGGTCGAGTGCCGAGGTGGGTTCGTCGAACAGCATGA
GCCGGGGTTCCATGGCCAGTGCCCGGGCCAGAGCAACCCGCTGCTGCTGGCCACCGGACAGCTGGCTGGGGTATTTGGACGCCTGGTCGCTGATACCGAC
CCTGTTGAGCAGCCTTTCCGCCGTCTCGGTGGCATCCGCCCGGTTCCAGCCGCGCACCTTCATGGGGGCCAGGGTCACGTTGTCGAGCACGCTGAGATGC
GGAAAGAGGTTGAATTGCTGGAAGACCATACCCACTTCGGTACGAATCCGCTGCAGCGCCTTGCTACTCTTTCCATCGGGTACCAGCTCGTTGCCGTCGA
CCTCGATATGACCCTGCTGGAATTCTTCCAGGCCGTTGATACAGCGAATCAGGGTGGACTTGCCGGAGCCACTGGCGCCGATGACGACGACGACTTCGCC
GGGCTTTATCGCCAGTTTGATGTCCTTCAGCACATGCAGGTCGCCGAAGTGCTTGTTGACCCGTTCCATGCGCACGATGGGATCGGATGTGGACGCTGTC
ATGCAAAGCTCCCTTGCTTCTCTTTATTGACCGACCAGGCCCTTGCGCTCGAGCATGCGCAACGCGATGGACAGGCTCCAGGTGATGGCCAGATACATCA
ATGCGACCATGAAGTAGACCTCCAGGGCGGTGAAGGTCGTGGCGATGTAGATCTGGCCCTGGCGGACCAGTTCGCCGACACCGATCACCGAGAACAGCGA

>Halomonas montanilacus PYC7W [NZ_QPII01000003.1:292361..292428], dnaA> ig dnaN>, 68 bp

< page 1 | #058 | Halomonas endophytica MC28

WebSkew GCmin pos. 151875 (GCmin<>oriC1 30.1 kb; GCmin<>oriC2 8.3 kb; oriC2<>oriC1 21.8 kb)

>Halomonas endophytica MC28 [NZ_PNRF01000027.1:121516..122015], <rpmH oriC1 dnaA>, 500 bp

TGCGGTGCTGTGGCAGCGCGGCATAGACGATGTCACCACCCTGCGGGTCGATGCGCACGTCCAGTACGTCAGTAACGACGGCGATCAGGTCACGTGTACC
GGTCGGGCGCTCGGGCTCGCCGGGCATGTCACCCGTCGTGGCGTCCTCGGCCCGCGGAGCGGCGGGGACAGCCAGGTCGTCATCATCGGGGGCCGCCGTT
TCCTCGCCGGGAGCGGGCGCCGTGAAGGTGGGTGCTTCCGGCTCAGGAGCAACCTGGCCGTAGTCCTGGTTCCACTGAATGACCAGCAAATAGGCCAGCA
CTGCCAGTGGAATCAGCAATAGGAGTCGTTTTACGTCCATGAGGGTTCTGCCCGGTGGGTAGTGAACATGCCGTTAACGGTATGCGCTGCCAGAGTGATA
TATGCGAAAGCCTGCCCAAGGCAGGCTATCGTGGCGGCGACAGGCATCGCATCCGGTCAGGATGCCATCGACTTGCGTACGTCTTTTTCCAGGCGACGCC
ACATGCCAAGAAGCTGACGATGCAGTGTCGCGTTGTCCAGCTCGTTTGCGCCCCGCCGGGCCAATACTACGATATCGACGGCGGGCAGCTGATGCTGCTG
AAGACGGATGGATTCGCGAGTAAGCCGTTTCAGGCGATTGCGATCGACGGCTCTACGGACGTTTCTCTTGCTGAAGACCAGGCCGAGGCGGGGGTGGCCT
AGCCCGTTGGGGCATGCCAGGGCCAACAGCCCCTTGCCATGTACCTTGAATGCAGCGCGATCAAAGACGCATCGATAGTCCCCGGCAGTCAGCAAGCGCA
GCTGCGGGGGAAACTCATGACCGGACACGCGCAATCCGCGATCAGGCGCTCAGACGCTTGCGGCCCTTGGCGCGACGACGTGCCAGTACGGCACGACCAT
TCTTGGTGGCCATGCGAGCACGGAAGCCGTGCACACGCTTGCGCTTGAGAACGCTGGGTTGAAAGGTGCGTTTCATAACTCTTGATTCCCACGTGGTGTG

TTTGGACGGAACGTCTTCCAGATAACGCCCAGGCACCTGATGGTGCCCGGGGAGATTCGGTTCAAGACCGGAAATTGTAGTGACTTCGGCAGCCCGACGC
AATTCTTCCTGGCATCCATTTTTGCCGGGTGTGAGGTAAGCTGCCTTTCATTACCAAAAGGTATCCTGATACTGCTGTTATCAATACTGGGGACGCAATT
**TGTTGATAA**GTATTAAATTCATAGCTTTATCAGAATGTTGCATTGCGACATCATCTGTGGAAAAGCCTTTCGGATCCAGTGTCTGAACTGTGTTCATCCT
GTGAAGAAATGTCGGCTTTATCCAGGCCTCAGCCCCGGTCCCATGCTGATCCCACGGGTATCCTCCTGTTTCGATGCCACTTTTCCACAGGCTGGCGGAG
CATCTCGTGCGTGTGGATAACCCTGCGGGTGAAGGATACAATGGCCGGTCATGCATCAACCGGATGGTGAGGTCGAGTGTCGCTTGCTCTATGGCAACAG

TGTTTGGATTACCTGCAGGACGAACTGAACGCCCAGCAGTTCAATACCTGGATCCGTCCGTTACAGGCGGAGGAGGGGGAGTCCAACGAGCTGCGTCTCC
TGGCGCCCAACCGTTTCGTGCGAGATTGGGTGGGAGACAAGTATGCCAAGCGAATCAATGAGCTTCTGCGGGACCTGGCGCCCGCCAAGCCCCCCAGGGT
CATATTGACCGTCGGTAGTCGGCGAACGGCAACGGCACCCAAGCCCCGGGACCTGGGAGCACCTGTCTCGGCCAGCCCCCGTTCACCGCTTGGCCCTGCG
GTTCACACCCCACCCAGAGGTGATATTGGCGACGAACGGGAAATCGATCGGCTTCGTGATGAGGGCGGCTCGACGCGTCGTGGTGGAGGGGAGCGTGCAG
TGCAGGTCGAAGGCAGTCTCAAGCACAATAGCGGCCTCAATCCCAATTTCACCTTCGAAACCTTCGTCGAGGGCAAGTCGAACCAGCTGGCCAGAGCGGC
GTCACGGCAGGTGTCCGAGAACCCCGGCGGCGCCTACAATCCCCTGTTTCTCTACGGCGGCGTCGGCCTGGGCAAGACTCACCTGATGCATGCCGTCGGT
AATGCTCTCGCCACGCGGCGAGAAAACGCCACGGTCGTCTATCTGCATTCCGAGCGTTTCGTTGCCGACATGGTCAAGGCACTGCAGCTCAATGCCATCA
ATGATTTCAAGCGATTCTATCGCAGTGTCGATGCGCTGCTCATCGATGACATCCAGTTCTTTGCCGGCAAGGAGCGTTCTCAGGAAGAGTTCTTTCACAC
CTTCAACGCACTCCTTGAGGGTGGCCAGCAGATGATCCTCACTTCCGATCGCTATCCCAAGGAGATCAGTGGTGTCGAGGAGCGCCTCAAGTCGCGCTTT
GGCTGGGGTCTCACCGTGGCCATCGAGCCACCGGAACTTGAAACCCGCGTCGCTATTCTGATGAAAAAGGCCGACCAGGCAAAGGTGGACCTGCCGCATG

>Halomonas endophytica MC28 [NZ_PNRF01000027.1:143300..143799], <mnmG oriC2 <COG1126 (glnQ), 528 bp

GTTCTCCAGGCCCTTGACCGAGCGAACCACCCGCAGCTGGACGTCGAAAGGCAGCGAGGTGGAGATCCCGTTGGGGTAGAGCTCGCGGGTATCCAGCCCT
TCCGGTTCGATGAAAATCTGGTGGCTCGACTTGTCGGCGAACCGGTGCACCTTGTCCTCGATGGACGGGCAATAGCGGGGCCCGACGCCTTCGATCACAC
CGGAGTACATCGGCGAACGATCCAGGTTGGACAGGATGATCTCATGGGTGCGCTCGTTGGTATGCGCGATGTGACAGCTCACCTGCCGTGGGTGCATCTC
GCGACGCCCCAGGTAGGACATGACCGGGGTCGGGGTATCGCCGGGCTGCTCCTCGAGCACGGAAAAATCGATGCTCTTGGCGTCGAGCCGCGGAGGGGTA
CCGGTCTTGAGCCGATCGACCCGGAACGGCAGTGCCCGCAGGCGTTCCGCCAGGGCATTGGAAGGCGGATCGCCTGCACGCCCCCCCGACTCTGGTCGA
GGCCGATGTGTATCACACCGCCCAGGAAGGTCCCGGTACACAGCACCACGCTACTCCCGTGAAAGTGGATGCCCGTCTCGGTCACGACGCCACGGACCGT
ATCGTTCTCGACAATCAGATCGCCTGCCGCCTGCTGGAACAACGTCAGATTGGGCTGACTCTCCAGCATTCCCCGAATGGCGGCCTTGTAACGAACCCGA
TCGGCCTGAGCCCGGGTCGCTCGTACTGCCGGCCCCTTGCGGGCATTCAGCACGCGAAACTGGATGCCACCCATGTCGGTGGCCAGGCCCATGGCGCCGC
CCAGGGCATCGATCTCCTTGACCAGATGGCTCTTGCCGATCCCGCCGATGGCGGGATTGCAGGACATCTGGCCCAGGGTCTCGATGTTGTGGGTCAGCAG
CAGGGTCTGACAGCCCATGCGGGCGGAGGCCAAGGCGGCTTCGGTTCCCGCATGGCCGCCGCCGATGACGATGACGTCAAAGCGGTCGGGGTAATCCAAG

ATGCACCTCATTGCGCTTGTGCGCGGATATTGCGTCGGGGTGAAAAATCAGCCTGGCAGTATAAACCTCCTGTGGGTAAGCGTCAGGGGTTTTCCACATG
GGGTCAACCAACCGGAGGGGTCTCTCGGTTTTACATAAATAAAATAGAATGATTTAAAGGAAGTTCTGTTGTTGTTGTTACTACTAGTGTGGACAGAATC
**CGTGGATAA**CCTTATTTTTCCGTTATTTTTCATGCAATTGAATTGTTGATGAACGACAGGGGAAAGCGCTATGTAGCTGAGGGCGACTGGTGCGGAGCCT
GTGGATGAAATGCCGAGTTGCACACAGGTGCGATTCTGCACCGCTTGTCCACCGGCGGCTACCTGGCTTGTCACCGTAGCTGTGAACAATGTCATGGAAG
TGAGCAAAGGCCGCTATGACGCCGGGTAGGACCGGAATGTCGGCAGGTGAGAAAAAAGTTGTCCACAGCCAAAAAAAGGCCCTGTCACTACCGACAAGGC

CTTTTTCGGGTGCGCGGGGGAGGTATTTCAGTGCTTCAGTACCCGAGACAGGAAGCTTTGAGTGCGTTCGTGACGCGGGTTGTCGAACAGGTCGGCGGGG
GGAGCTTCCTCAAGGATCTGCCCCTGATGAATGAAGATCACCCTGTCAGCGACCTCCCGGGCAAAGCCCATTTCGTGGGTCACGATGATCATGGTCATGC
CTTCCCGGGCCAGTTCGCGCATGGCATCGAGTACTTCGCCAATCATCTCGGGGTCGAGAGCCGAAGTGGGCTCATCGAACAGCATGAGGCGCGGTTCCAT
GGCCAGCGCCCGGGCCAGGGCAACACGCTGCTGCTGGCCGCCCGATAGCTGGCTGGGATACTTGATGGCCTGATCGCTGATGCCGACACGGTCGAGCAGC
CGCTCCCCTGTCTTCACGGCATCGGCACGGTTCCAGCCGCGTACCTTCATCGGCGCCAGCGTCACGTTGTCGAGTACGCTCAAATGCGGGAACAGGTTGA
ACTGCTGGAAGACCATGCCGACCTCGGTGCGGATCCGCTGGAGTGCCTTGCTGCTTCTGCCGTCGGGGAGCAGCTCGTTGCCGTCGACCTCGATATGACC
CTCCTGGAACTCCTCCAGGCCGTTGATACAGCGGATCAGGGTGGATTTTCCCGAGCCGCTGGCGCCGATGACGACGACCACTTCACCGGGTTTTACCTGC
AGGTCGATATCCTTGAGGACATGCAGGCTGCCGAAGTGCTTGTTGACCTTTGCCATGCGCACAATGGGATTGGATACGGACTCTGTCATGCGGTGGTTCC
CCTTTTCCTGCCCCTTTACTGGCCAACGAGGCCCTTGCGCTCGAGCAGGCGCAAGGCGATGGAAAGACTCCAGGTAATGGCCAGATACATCAGGGCCACC
ATGAAGTAGACCTCCAGCGCGGTGAACGTCGTGGCGATGTAGATCTGCCCCTGACGCACCAGCTCACCGACGCCGATCACCGAGAACAGCGAGGTATCCT

>Halomonas endophytica MC28 [NZ_PNRF01000027.1:120000..120069], dnaA> ig dnaN>, 70 bp

WebSkew GCmin pos. 139946 (GCmin<>oriC1 26.1 kb; GCmin<>oriC2 9.1 kb; oriC2<>oriC1 17.0 kb)

>Halomonas campisalis A4 [NZ_JABFUC010000002.1:165780..166279], <rpmH oriC1 dnaA>, 506 bp

TGCTGGGGCAGGGCGGCGTAGATGATATCGCCGCCCTGGGGGTCGATGCGCAGGTCCAGCACGTCGGTGACCACGGCCACCAGGTCACGGGTGCCGTCGG
GACGTACCGGCTCGCTCGGCATCTCGGCAGCCACGTCATCTTCTGCCCGGGGAGCCGCGGGAACCGCCAGATCGTCCGCGTCAGGCCGCGCTGCTTCCTC
ATCGGGGGCGGTGGTGGTAAAGGTGGGTGCTTCCGGCTCAGGGGCTACCTGGCCGTAGTCCTGGTTCCACTGCACGACGATCAGATAGGCGAGCACGGCC
AGTGGAATCAGCAATAGGAGTCGTTTTACGTCCATGAGGGTTCTGCCCGGTGGGTAGTGAACATGCCGCTGGAACGGTACGCATTGCAGGCCAATACGCG
AAAGCCTGCCCAGGGCAGGCTATCGTGACGGCGGCCGGCGGATCCGGTCAGGATGCCATCGACTTGTGAACGTCCCGTTCAAGGCGTCGCCACATGCCGT
GCAGCTGGCGGTGAAGCGTCGCGTTGTCCAGCTCATGAGCGCCCCGTCGGGCCAGCAGTACGATATCGACGGCGGGTAGCCGGTGCTGCTGAAGACGGAC
GGATTCGCGAGCGAGCCGCTTGAGGCGGTTGCGATCGACGGCTCGGCGGACGTTTCGCTTGCTGAAGACCAGGCCGAGGCGGGGGTGGCCCAGCCCGTTT
GGGCATGCCAGGGCCATCAGCCCCTTGCCATGAACCTTGAATGCAGCGTGATCGAAGACCTGTCGGTAATCCCCAGCGGTCAGCAGCCGCAGCCGCGGGG
GAAAGCCATGATGACCGGACACGCGCAATCCGCGATCAGGCGCTTAGACGCTTGCGGCCCTTGGCGCGACGACGAGCCAGAACGGCACGGCCATTCTTGG
TGGCCATGCGCGCACGAAAGCCGTGCACACGCTTGCGCTTCAGGACGCTGGGCTGAAAGGTACGTTTCATAACTCTTGATTCCCACGTGGTGTGTTTGGA

CGGTACGTCTTAGTCTTACAGGAGACGTCCGGGTTCCTGCCGGAACCCGGGGAGATTCGGTTCAAGACCGGAAATTGTAGTGATTTCGGCCGCCGGGTGC
AATTTTTCCGTTGGGTCTCATGGCAGGCGGCAAGCCTTGCGCTTCACTGGTTACCCCGCATGAGTTTCTTCATGCTGTGGTTAATATTGGGGACGCATCT
**TGTGAATAA**CCAGTAAAAACCCTTTTTGTTCAGTGTTTTATGGATGTTGTTTGGCAGTGGAAAAACTGCGATCAAGCGGTTGGCTGGCGGCCAGGCGGCA
GCGCCTGGCGGGAAAAGTTATCCACAGCCGACAATTTATTCCGCTCTCCCACGCCGTTCGCCTCGGCCCGGCGCTGGCGTTTTCCACAGGCTGTCATCGT
GGCGTCGGGCGTGTGGATAACCTGTCTTGGCAAGGCTACAATGTGCGGTCATTCATCAACCGGACGGTGAGGTCGCGTGTCGCTCGCTCTTTGGCAACAG

TGCTTGGATTACCTGCAGGATGAGCTGAATGCCCAGCAGTTCAATACCTGGATCCGTCCGCTCCAGGCGGAGGAGGGGGAGTCCAATGAGCTTCGGCTGC
TGGCACCGAATCGCTTCGTGCGCGACTGGGTCGGTGACAAGTACGCCAAGCGTATCAATGAACTGCTTCGCGAGCTGGCCCCTGCCAAACCGCCCAGGGT
GGTGCTGACCGTCGGGAGTCGCCGCGCGGCGGCGCCCATGCCGCAGCCTCGTGAGCTGGGCACCCCGGTATCGGCCAGTCCGCGCTCTCCGCTGGGGCCC
GCCGTGCAGACCGCGCCTCGCGGGGATGTCGGCGATGAGCGCGAGATCGACCAGATGCGGGAAGATGCCGCCGGTGCGCGGCGTTCCGGCGGTGAGCGCG
AGGTGCAGGTCGAGGGCAGCCTCAAGCACAGCAGCGGGCTCAATCCCAATTTCACCTTCGAGACCTTCGTCGAAGGCAAGTCCAACCAGCTGGCCCGCGC
CGCCTCACGGCAGGTCTCCGAGAATCCCGGCGGGGCCTACAACCCGCTGTTCCTGTATGGCGGCGTGGGCCTGGGCAAGACCCACCTGATGCATGCGGTA
GGCAATGCCCTGGCGGTGCGTCGCGAGAACGCCCGGGTGGTCTACCTGCACTCCGAGCGTTTCGTCGCCGACATGGTCAAGGCGCTGCAGCTCAATGCCA
TCAACGACTTCAAACGCTTCTACCGCAGCGTCGATGCGCTGCTGATCGATGACATCCAGTTCTTCGCCGGCAAGGAGCGCTCCCAGGAGGAGTTTTTCCA
TACCTTCAACGCCCTGCTCGAGGGCGGTCAGCAGATGATCCTTACCTCCGACCGCTATCCCAAGGAGATCAGCGGTGTCGAGGAGCGCCTCAAGTCGCGC
TTCGGCTGGGGGTTGACCGTGGCCATCGAGCCGCCGGAGCTGGAAACGCGAGTGGCGATCCTGATGAAGAAGGCCGACCAGGCCAAGGTCGACCTGCCCC

>Halomonas campisalis A4 [NZ_JABFUC010000002.1:148778..149277], <mnmG oriC2 <PRK13623 (ErpA), 733 bp

TTTTCCAGGCCCTGGATCGAGCGCACCACCTGCAGCTGCACATCGAAGGGCAGCGAGGTGGAGATGCCGTTGGGGTAGAGCTCATGGGTGTCCAGGCCTT
CCGGCTCGATGAACACCTGGTGGCTCGCCTTGTCGGCGAAGCGGTGCACCTTGTCCTCGATGGAGGGGCAGTAGCGCGGCCCCACGCCCTCGATCACCCC
GGAATACATGGGCGAACGGTCCAGGTTGGCCAGGATCAACTCATGGGTCTGGGCGTTGGTATGGGCGATATGGCAGCTCACCTGACGGGGATGCATCTCG
CGGCGGCCCAGAAACGACATCACCGGCGTCGGGCTGTCGCCGGGCTGCTCCTCGAGCACCGAGAAGTCCACGCTCTTGGCGTCGAGCCGCGGCGGCGTAC
CGGTCTTGAGCCGGTCGACGCGAAACGGCAGGGCGCGCAGCCGCTCGGCCAGGGCATTGGAGGACGGGTCCCCGGCGCGTCCACCGCGACTCTGGTCGAG
GCCGATATGGATCACCCCGCCGAGGAAGGTGCCGGTGCACAGCACCACGGTCTCGCCGAGGAAACGGATGCCGGTCTCGGTGACCACCCCGCGCACGCTG
TCGCCGTCGACGATCAGGTCGCCGGCGGCCTGCTGGAAGAGCGTCAGGTTGGGCTGGTTTTCCAGCATCCCCCGAATCGCCGCCTTGTAGCGGACCCGAT
CGGCCTGGGCACGCGTCGCTCGAACCGCCGGGCCCTTGCGGGCATTGAGCATGCGGAACTGGATACCGCCCAGGTCGGTGGCCAGTCCCATGGCGCCACC
CAGCGCATCGATCTCCTTGACCAGATGACTCTTGCCGATCCCGCCGATGGCGGGATTGCAGGACATTTGACCGAGGGTCTCGATGTTGTGGGTCAGCAGC
AGGGTCTGACAGCCCATGCGGGCGGAGGCCAGGGCGGCTTCGGTTCCCGCATGGCCGCCGCCGATGACGATGACGTCAAAGCGGTCGGGGTAATCCAAGG

TGCACCTCGTTGGCGCCGCGGCGCCGTCTTCCAGTCAGGGAATGAAATCAGCCCGGCAGTATAAACCTCCTGTGGGTAATCGTCAGGGTTTTCCACACCC
GGGATCGCCTCCGAGCCGAACCGTCGTCTCACTAAGTAAAAACAGAATATATATAAAGAGAAGTTCTGCTGTGGTTGTAACTACTGGTGTGGACAAAACC
**CGTGGATAA**GTGGATATCCCGCAAATCTATCAACAAGATGGCTTGTTGCCAAAAACCGTGGAAAAGGGCTGACTGGCTGCGGACGAGCGGTGGTGAGCCT
GTGGGCAAAAGCGCTGCTTGTCCACAGCGCCAGTTGGACACCGCTTGTCCACCGTTGGGTACCGGGCTTGTTACCGTGGCTGTGAACAACCGCGCGCATT
TTGATCCAAGCCGCCAGCGGCGGGGCCTGGCGGCCGATTGGAGGGGGCTAGGGAAAAATCTTGTCCACAGGCCAAAAAACGGCCCTGTCGCAAGCGACAG

GGCCGTTTTTTGGCAGGATCAGGCGGGAGGTGAGAGCCGGTTGGCAACGTTCTCGGATGACTCAGTGCTTGAGCACCCGGGAGAGGAAGCTGCGCGTCCG
GTCGTCGTCGAAGAGCATCAATCGCGGCTCCAGCCGCGTACCTGCATGGGGGCAAGGGTCACGTTGTCGAGCACGCTCAGGTGGGGGAACCGGCGGCGGT
CAACGCCTGGGCTTAGACCATGAACCAAGGCTTAGACCATGAAGGAGGCGCCGCAGCCACAGGTAGTGGTGGCGTTGGGGTTCTGGACCAGGAAGCGAGC
GCCGGCCAGCCCCTCCTCGTAGTCCACGGTGGAGCCGACCAGATACTGGTAGGAGAGCTCATCGATGACCAGCGCCACCTCGCCGAACTCGATCAGGGCG
TCGTCGTCGCCCACGTCCTCGGCAAAGTCGAAGCCGTACTGAAAGCCGGAGCAGCCCCCGCCGGTCACATAGACCCGCAGCTTGAGCTGGCTGTTGCCCT
CTTCCTCGATCAGCGACTGGAGCCGCTTGCGTGCGCTGTCGGACAGCAGCAGGGGAGTGGGAACAAAGGATTCGGCACCGCTCATGGAGAGCTCCCTTGG
GGGAAATGGGAAATTCGACCGGCCATTATCCCCAATCCCCAGCAAAGGGGTCAACTTTTGCTGGGGGGAGGGCTATCACAGATAGCTTGCATTCCTCGCT
TTTCGTAACTGCGATAGCCCTGGGGGTGTGAGATCAGGGCATCAGCGCCGGGGCGGCCAGGCCGGCCCGTTCATCGAAGCCGAACATCAGGTTCAGGTTG
TGGATGGCCTGGCCGGAGGCGCCCTTGACCAGGTTGTCGATCACCGCCAGCACCACCACGGTGTCACCGTTGCCCGGCCGGTGCACCGCCAGCCGGCAGA
GGTTGGCGCCCTTGACGCTGCGGGTCTCCGGGTGGCTGCCGGCGGGCATGACGTCGACGAAGGGCTCACTGGCGAAGCGCGTCTCGAACAGCGCCTGGAG

>Halomonas campisalis A4 [NZ_JABFUC010000002.1:167729..167799], dnaA> ig dnaN>, 69 bp

< page 1 | #060 | Halomonas azerbaijanica TBZ202

WebSkew GCmin pos. 112965 (GCmin<>oriC1 33.0 kb; GCmin<>oriC2 9.3 kb; oriC2<>oriC1 23.7 kb)

>Halomonas azerbaijanica TBZ202 [SBLD01000003.1:145766..146265], <rpmH ig dnaA>, 388 bp

GAAACTGATGCGTCCGGAGTGGCCGTCGATCTGCAGGCCCGAACGGGCGACGTAGCTGCGCGACTCGCTGTCGGAGAGCAGCACGTAGGGGTGCTCGCTG
TCCCGCGACAGTCGATGCTGCGGCAGAGCCGCGTAGACGACGTCGCCGCCCTGCGGGTCGATGCGCAGGTCGAGCACGTCCGTCTCGACCACGATCAGTT
CGCGGCTGGTATCGGGGCGCCCGGCGGGGGCGGGCATCTCGCCGGCCAGCTCGCCCTCGCTGCGGGGCGCCGCCGGCACGGCCAGGCCGTCCTCGTCGGG
CGTCTCGCCCTGCTGATCGGGGGCGGGCGTGGTGACGCTCGGGGTGTCCGGCTCAGGGGTGACCTGGCCATAGTCTTGGTTCCACTGCACCATGATCAGG
TAGGCGAGCACGGCCAGTGGAATCAACAGAAGGAGTCGTTTTACGTCCATGAGGGTTCTGCCCGGTGGGTAGTGTACATGCCGCTGGAACGGGGCCTGCT
GGCAGACCGATATGCGCGAAAGCCTGCCCGAGGCAGGCTATCGTGGCGGAAAGCGGCACCGCATCCGATCAGGATGCCCTCGACTTGTGAACGTCGCGTT
CCAGCCGGCGCCACATGCCGTACAGCTGGCGGTGGAGCGTGGCGTTGTCCAGTTCATGGGCGCCTCGACGGGCCAGCACCACGATGTCGACGGCGGGCAG
CCGATGCTGCTGGAGTCGGATGGACTCGCGGACGAGCCGTTTCAGGCGGTTGCGATCGACGGCTCGGCGGACGTTTCTCTTGCTGAACACCAGGCCGAGT
CGTACGTGCTGCCGCGCGTTGGGACACGCCAGGGCCAGGAGCCCCTTGCCATGAACCTTGAATGCAGCGTGATCGAAGACGTAGCGGTAGTCCCCGGCGG
TCAGCAGGCGCAGCCGTGGGGGAAAACCGTGACCGGACACGGGCAATCCGCGGTCAGGCGCTCAGACGCTTGCGGCCCTTGGCGCGGCGACGTGCCAGCA

CGGCACGGCCGTTCTTGGTGGCCATGCGCGCACGAAAGCCGTGCACACGCTTGCGCTTGAGAACGCTGGGTTGAAAGGTGCGTTTCATAACTCTTGATTC
CCACGTGGTTTAATTGGACGGAACGGCTTCAAGGCAACACCCGGACGCCGCACGGCCCCCGGGAGGATTCGATTCAAGAGCGGAAATTGTAGAGACTTCG
CTGGCCCGACGCAATTGTTGCGCCCCTCTCGCGAGGACGGACGCGGCGGCGAGGAGCGCCGCCGCGCGGCAGTGGCCGGTCCGGGCTTGTGGGCAGGTTT
GGGGATCTGTCTTGTGAATAAATCGCGTAGTGGCTGTTGAATCAATGAGCTATGCCATGAATAGCGAAGTGGACAAGGGGCGACATCCACAACAGATGGA
GCGGCACGGCATGTGGATAACCCGCCCGGGGAAAGATACAATGTCCGGTCATTCATCAACCGGATGGTGGGGTCGCGTGTCGCTTGCTCTCTGGCAACAG

TGTCTGGATTACCTGCAGGACGAGCTGAACGCTCAGCAGTTCAACACCTGGATCCGCCCGCTGCAGGCGGAGGAGGGGGAGTCCAACGAGCTGCGCCTGC
TGGCGCCCAATCGCTTCGTGCGCGACTGGGTGGGCGACAAGTACGCCAAGCGCATCAACGAACTGCTGCGCGACCTGGCGCCGGCCAAGCCGCCGCGTGT
GGTGCTGACGGTGGGCAGTCGGCGGGCCGCCGTGGCCAAGCCGCAGCCGCGCGAGCTGGGCTCGCCGGTCTCGGCCAGCCCCCGTTCTCCGCTGGGGCCG
GCGGTGCACACACCGCCACGCGGCGATATCGGCGACGAGCGTGAGCTCGAGCCGCTCAGCGAGGAGGGGGAGGCGCTCGCGGGCCGGCGCAGCGGGGCCG
AGCGCGAGGTGCAGGTCGAGGGTAGCCTCAAGCACAGCAGTGGGCTCAACCCCAGCTTCACCTTCGAGACCTTCGTCGAGGGCAAGTCCAACCAGCTGGC
CCGGGCGGCCTCGCGCCAGGTCTCGGAGAACCCCGGCGGTGCCTACAACCCGCTGTTCCTCTACGGCGGGGTGGGGCTCGGCAAGACCCACCTGATGCAC
GCCGTGGGCAATGCGCTGGCCGGTCGGCGCGAGAACGCGCGCGTTGTCTACCTGCACTCCGAGCGCTTCGTGGCCGACATGGTCAAGGCGCTGCAGCTCA
ACGCCATCAACGACTTCAAGCGCTTCTACCGTAGCGTCGATGCGCTGCTCATCGACGACATCCAGTTCTTCGCCGGCAAGGAGCGCTCCCAGGAGGAGTT
CTTCCACACCTTCAACGCCCTGCTCGAGGGCGGCCAGCAGATGATCCTCACCTCGGACCGCTACCCCAAGGAGATCAGCGGGGTCGAGGAGCGCCTCAAG
TCGCGCTTCGGCTGGGGCCTGACCGTGGCCATCGAACCGCCGGAACTCGAGACGCGGGTGGCCATTCTCATGAAGAAGGCCGACCAGGCCAAGGTCGACC

>Halomonas azerbaijaica TBZ202 [SBLD01000003.1:122020..122519], <mnmG oriC2 <COG1126 (glnQ), 525 bp

CTCGAGGCCCTGGATCGAGCGCACCACCGCGAGCTGCACGTCGAAGGGCAGCGAGGTGGAGATGCCGTTGGGGTAGAGCTCGTGGGTATCCAGCCCCTCC
GGCTCGATGAACACCTGGTGGCTCGCCTTGTCGGCGAAGCGGTGCACCTTGTCCTCGATCGACGGGCAGTAGCGCGGTCCCACGCCCTCGATGACGCCGG
AGTACATCGGCGAGCGGTCGAGGTTGGCAAGGATGATCTCGTGAGTGTGCGCGTTGGTGTGGGCGATGTGGCAGCTCACCTGGCGCGGGTGCATCTCGCG
CGTTCCCAGGTAGGACATCACCGGGGTGGGGGTGTCGCCCGGCTGCGCCTCGAGTACGGAGAAGTCGACGCTCCTGGCGTCGAGGCGCGGGGGCGTGCCG
GTCTTGAGGCGGTCGACGCGGAACGGCAGCGCGCGCAGGCGCTCGGCCAGCGCATTGGAGGGCGGATCGCCGGCGCGGCCACCACGGCTGTTGTCGAGGC
CGATGTGGATGACACCGCCGAGGAAGGTGCCGGTGCACAGCACCACCGTCTCGGCCAGGAAGCGGATGCCCGTTTCGGTGACCACCCCGCGGACGGTGTC
GTTATCCACAACCAGGTCGCCGGCGGCCTGCTGGAAGAGCGTCAGGTTGGGCTGGTTTTCCAGCATGCCGCGGATGGCAGCCTTGTAGCGGACGCGGTCG
GCCTGGGCACGGGTGGCACGCACCGCGGGTCCCTTGCGGGCATTGAGCACGCGGAACTGGATGCCGCCGCGGTCGGTGGCCAATCCCATGGCGCCCCCCA
GTGCATCGATCTCCTTGACCAGGTGGCTCTTGCCGATGCCGCCGATGGCGGGATTGCAGGACATCTGGCCCAGCGTCTCGATGTTGTGGGTGAGCAGCAG
GGTCTGACAGCCCATGCGGGCGGCAGCCAGGGCGGCTTCGGTTCCCGCATGGCCGCCGCCGATGACGATGACGTCAAAGCGGTCGGGATACTCCAAGGTG

CACCTCGTTGCGCGTCACGCGCGAATCGTCCGAAGAGGATGAAAATCAGCCTGCCAGTATAAACCTCCTGTGTGTAAGCGTCAGGGGTTTTCCACATGCC
CTTGACCAGGGATCTCACTACACTGCACAGACCAACAATAAAATAGAAAGATTAAAAGAGAAGTTCTGTTGTGGTTGTTACTACTGGTGTGGACAAAAAC
**TGTGGATAA**GGCAGATTAATGCTATTGCATCAGAGAGATGCATTGTTGACCAAAACCGGGACAAGCGGATGACGACCTGCGCGGGGCCGGTGAGGAGGCT
GTGGATGAAAAGGGCGTTGATCCACAGCCCCGTGAGCCCACGTCTTTTCCACCGTTCCCTACCGACGTTGTCACCGCGGCTGTGAACAACCGGCTCGATA
GGCCGCGAGACGCGCCATGGCGGGCCTGGTGGGCTGCGGCGCCATAATCTTGTCCACAGGCAAAAAAAAAAGGCCCCGTCGTCGGTGGCGGGGCCCGGTC

GAAGCGGTATGACGTCGCGCTTCAGTGCTTGAGCACGCGCGACAGGAAGTTGCGGGTGGGTTCGTTGCGGGGGTTGTCGAAGATCTGTCCGGGAGGCCCC
TCCTCGGCGATCTCGCCCCGATGGATGTAGATGACCCGGTCGGCCACCTCGCGGGCGAAGCCCATCTCGTGGGTGACGATCAGCATCGTCATGCCTTCGC
GGGCGAGTTCGCGCATGGCGTCGAGGACCTCGCCGATCATCTCCGGGTCGAGCGCCGAGGTGGGCTCGTCGAAGAGCATCAGGCGCGGTTCCATGGCCAG
CGCCCGGGCCAGGGCGACGCGCTGCTGCTGGCCGCCGGAGAGCTGGCTGGGGTACTTGGCGGCCTGGTCGGCGATGTCCACGCGCTCCAGCAGCCGCTCG
GCCGTCTCGGTGGCATCGGCCCGGCTCCAGCCGCGCACCTTCATCGGCGCCAGGGTCACGTTGTCGAGCACGCTCAGGTGTGGAAACAAATTGAACTGCT
GGAAGACCATGCCCACTTCGGTGCGGATCTTCTGCAGCGCCTTGCCGGCCTTGCCGTGGGGCACCAGGTCGTTGCCGTCGACCTCGATATGGCCGCTCTG
GAACTCCTCCAGTCCGTTGATGCAGCGGATCAGGGTCGACTTGCCGGAACCGCTGGCGCCGATGATGACCACCACCTCGCCGGGGGAGACCGCCAGGTCG
ATGTCCTTGAGGACGTGCAGGCTGCCGAAGTGCTTGTTGACCTTGTCCAGGCGCACGATCGGGTCGGCGCCGGAGGGGGACGTCTCTGGGTTCATGTCGC
GCGTTCCTTATTGTCCGACCAGGCCCCTGCGCTCCAGCTGGCGCAGTGCAAACGACAGGCTCAGGGTGATCACCAGGTAGAGCAGGGCGACCATCAGGTA
GACCTCCAGGGCATTGAAGGTCGTGGCGATGTACACCTGGCCCTGGCGGACCAGCTCGCCGACGCCGATCACCGAGAACAGTGAGGTGTCCTTGATGCTG

>Halomonas azerbaijanica TBZ202 [SBLD01000003.1:147721..147805], dnaA> ig dnaN>, 85 bp

< page 1 | #061 | Halomonas gudaonensis CGMCC 1.6133

WebSkew GCmin pos. nd (GCmin<>oriC1 nd; GCmin<>oriC2 nd; oriC2<>oriC1 40.0 kb)

>Halomonas gudaonensis CGMCC 1.6133 [NZ_FNES01000002.1:97497..97996], <rpmH ig dnaA>, 377 bp

GTATAGCTGATGCGGCCCGAGTGACCATCGAGCTGTAGCCCGGAACGCGCCACGTAGCTGCGCGTTTCACTGTCGGACAGCATCACGTAGGGGTGCTCGC
TGTCTCGCGTGAGCCGGTGCTGCGGCAGTGCTGCGTAGACGACGTCGCCGCCCTGGGGATCGATGCGCAAGTCGAGGACGTCCGTCTCCACCACGATCAG
ATCCCGGCTGGTGCCGGGACGTGCCGTGTCGCTAGGCATGTCGCCGGCCATTTCGTCCTGTTCGCGCGGGGCGCTGGGGACGGCAAGGCCTTCTTCATCA
GGCGTTTCGGCCGTCTCGGCTTCCTGTTCGCTGGCCGTGGTGGTGACGCTGGGAGTGTTCGGCTCAGGCGTGTTCTGGCCATAGTCCTGGTTCCACTGCA
CCATGATCATGTAGGCGAGCACGGCCAGAGGAATCAACAGTAGGAGTCGTTTTACGTCCATGAGGGTTCTGCCCGGTGGGTAGTGTACATGCCGCTGGAA
CGGGGCCTGCTGCCAGGCCGATATACGCGAAAGCCTGCCCGAGGCAGGCTATCGTGGCGGAAACCGGCACCGCATCCGCTCAGGATGCCTTCGACTGGCG
AACGTCCCGTTCCAGCCGGCGCCACATGCCATGCAGCTGGCGATGGAGCGTAGCGTTATCCAGTTCATGGGCGCCACGACGGCCCAGCACCACGATATCG
ACGGCGGGTAGCCTATGTTGCTGGAGTCTGATGGACTCGCGGGCAAGCCGCTTCAGGCGATTGCGATCGACGGCGCGGCGAACGTTTCTCTTGCTGAACA
CCAGGCCGAGTCGGACGTGTCCTTGCTGGTTGGGGCACGCCAGGGCCAGCAGCCCTTTGCCGTGAACCTTGAAAGCAGCGTGATCGAAGACGTGGCGGTA
GTCCCCGGCAGTCAGCAAGCGCAACCGAGGGGGAAAACCATGACCGGACACGGGCAATCCGCGATCAGGCGCTCAGACGCTTGCGGCCCTTGGCGCGACG

ACGTGCCAGTACGGCACGGCCGTTCTTGGTGGCCATGCGCGCACGAAAGCCATGCACACGCTTGCGCTTGAGAACGCTGGGTTGGAAAGTGCGTTTCATA
ACTCTTGATTCCCACGTGGTTAAATTGGACGGAACGTCTTCAAGACAGCACCCTGAGGCCACGTGCGGCCCCGGGAGGATTCGGTTCAAGACCGGAAATT
GTAGTGACTTCGCCGCTTCGACGCAATTGTTGCGCCCCTGGTTTCTGCCGTCGGAATGGGCTCTGCCCGGAGCGTGGCGGTTCGGCCTGTGGGCAGGAGC
GCGGTGCCGAGTTGTGGATAATTGCGCTTTATTCTTATTAAACAATCGCCTGTTTATTGTATAAAGGTGTGGATATGGGTTGTCGGTCGTAACCAGGAGG
CGGAGTGCACGCATGTGGATAACCTGCCGGGGGAAGGATACAATGATCGGTCATTCATCAACCGGGGTGAGGTCGCGTGTCGCTGGCTCTCTGGCAACAG

TGTCTGGATTTCCTGCAGGATGAGCTGAACGCTCAGCAGTTCAACACCTGGATTCGCCCGTTGCAGGCGGAAGAGGGGGAGTCCGACGAGCTGCGCCTGC
TGGCGCCCAATCGCTTCGTGCGTGACTGGGTGGGCGACAAGTACGCCAAGCGCATCAACGAGCTGCTGCGCGAACTGGCACAAGGCAAGCCGCCGCGGGT
AGTGCTTACCGTGGGTAGCCGACGGGCGGCGGTGGCCAAACCACAGCCCCGTGAACTGGGTGCACCGGTTTCCGCAAGCCCTCGCGCGCCGGCCGTGCAG
ACACCGCCACGCGGCGATATCGGCGATGAGCGCGAGATCGAGTCGTCGAACGAGGAAGGCTCGGCATCCGTCGGTCGCCGTGGAGAACGCGAAGTTCAGG
TCGAGGGTAGCCTCAAGCACAGTAGCGGCCTCAATCCCAATTTCACGTTCGATACCTTCGTCGAGGGCAAGTCGAACCAGTTGGCTCGTGCGGCCTCGCG
TCAGGTCTCCGAGAATCCCGGGGGGGCCTACAATCCTCTGTTCCTCTATGGGGGAGTGGGGCTGGGCAAGACGCACCTGATGCACGCCGTGGGCAATGCT
CTGGCCATTCGGCGCGAGAATGCGCGTGTCGTCTACTTGCATTCGGAGCGCTTCGTCGCCGACATGGTCAAGGCGCTTCAACTCAACGCTATCAATGACT
TCAAGCGTTTCTATCGCAGCGTCGATGCGCTGCTCATCGACGACATTCAGTTCTTCGCCGGCAAGGAGCGCTCTCAGGAGGAGTTCTTCCACACTTTCAA
TGCCCTGCTCGAGGGCGGTCAGCAGATGATCCTCACTTCCGACCGCTATCCCAAAGAGATCAGTGGCGTCGAAGAGCGTCTCAAGTCACGCTTCGGTTGG
GGGCTCACGGTGGCCATCGAGCCGCCGGAACTGGAGACGCGGGTCGCCATCCTGATGAAGAAGGCCGACCAGGCCAAGGTCGACCTGCCACACGACGCCG

>Halomonas gudaonensis CGMCC 1.6133 [NZ_FNES01000002.1:57496..57995], <mnmG oriC2 <COG1126 (glnQ), 523 bp

CATTCTCCAGGCCCTGGATGGAGCGCACCACCTGTAGCTGGACGTCGAAGGGCAGCGAGGTGGAGATACCGTTGGGGTAGAGTTCGTGGGTGTCGAGCCC
CTCGGGCTCGATGAACACCTGGTGGCTCGCCTTGTCGGCGAAACGATGCACCTTGTCCTCGATGGACGGGCAATAGCGCGGGCCTACGCCTTCGATCACC
CCGGAGTACATCGGTGAGCGGTCGAGGTTGGCGTGGATGATCTCGTGGGTACGCGCATTGGTGTGAGCGATGTGGCAGCTGACCTGATGGGGGTGCATGT
CGCGCGTTCCCAGGTAGGACATCACCGGAGTCGGAGTGTCGCCGGGCTGCTCCTCGAGGGCCGAGAAATCCACGCTCTTGGCGTCGAGGCGTGGCGGTGT
GCCGGTCTTGAGCCGATCCACGCGGAAGGGCAGCGCACGCAGCCGTTCGGCCAGGCGGTTCGACGGTGGGTCACCAGCGCGCCCCCCGCGGCTGTTGTCG
AGGCCGATGTGGATGACCCCGCCGAGAAAGGTGCCGGTGCACAGCACCACCGTTTCGGCATGGAAGCGAATGCCGGTCTCGGTGACCACGCCGCGTACGG
TGTCATTTTCCACCACCAGATCGCCAGCGGCCTGCTGGAAGAGCGTCAGGTTGGGCTGATTTTCCAGCAGCCCGCGAATCGCCGCCTTGTAGCGAATCCG
GTCCGCCTGGGCGCGGGTGGCGCGCACCGCCGGTCCCTTGCGGGCATTGAGAACCCGGAACTGGATACCGCCCCGATCGGTGGCCAAACCCATGGCGCCA
CCCAGCGCATCGATCTCCTTGACCAGATGACTCTTGCCGATGCCGCCGATGGCGGGATTGCAGGACATCTGTCCCAGCGTCTCGATGTTGTGGGTCAGCA
GCAGTGTCCGACAGCCCGTGCGTGCAGCGGCCAAGGCGGCTTCGGTTCCTGCATGGCCGCCGCCGATGACGATGACGTCGAAGCGGTCGGGGTAATCCAA

GGTGCACCTCGCTGGCGCCGTTGCGCCTGTCCTGGAATCAGGGGAAATCAGCCCGCCAGTATAACTCCCCTGTGGGTAAACGTCAGGGGTTTTCCACATG
GCCGAAGCCAGCATGCACAGGGGGTCTCTCCATCAATTACTATCTATATGAAATAAAGAGAAGTTCTGTTGTGGTTGTAATTACTGGTGTGGACATTATT
**TGTGGATAA**GGTTGATTACTCTTTTAAAGACAGTCATTTGCTGTGTTGACTGAAACTCGGAAAGACGGTCGACGTCCTGGGCGCGAACGGTGAGGACGCT
GTGGATGGAATCGCCAGTCGTCCACAGCCTGACACATCAAGGCGTTTTCCACCGATGGATACCGGAGTTGTCATCGTGGCTGTGAACAACCGGGCCAAAC
CGGGTTCGGCGGCGCAACCGCGGGGCTGGGTGCCATTCGGTGAAAAATCTTGTCCACAGGCCGAAAAAAGGCCCTGACGGTATCGACAGGGCCTGCGGAG

GATTGCCAGAGAGGCTGGGGGAATCAGTGCTTGAGCACACGCGCCAAGAAGTTGCGGGTAGGCTCGTTCTTCGGCGAATCGAAGATCTCCTCCGGTGGGC
CTTCCTCGGCAATTTCTCCCTGATGGATGTAGATGACGCGATCGGCAACCTCGCGGGCGAAGCCCATTTCGTGGGTGACGATCATCATGGTCATGCCTTC
GTGGGCCAATTCACGCATCGCATCCAGCACTTCACCGATCATCTCGGGGTCGAGTGCCGATGTCGGTTCGTCGAAGAGCATCAGGCGAGGTTCCATGGCC
AGGGCGCGTGCCAGTGCGACACGCTGCTGCTGGCCGCCGGAGAGCTGACTCGGGTACTTGCTGGCCTGGTCGGCGATATCGACGCGTTCCAGGAGGCGCT
CGGCCGTCTCCGTTGCATCCTTGCGGCTCCAGCCGCGCACTTTCATCGGGGCCAGGGTCACGTTGTCCAGCACGCTGAGGTGGGGAAAGAGGTTGAACTG
CTGGAATACCATGCCCACTTCGGTACGAATCTTCTGTAGTGCCTTGGCGGCCTTGCCGTAGGGCACCAGATCGTTGCCATCGACCTGGATGTGACCCTGC
TGAAATTCCTCGAGCCCGTTGATGCAACGTATCAGGGTCGATTTTCCGGAGCCGCTGGCACCGATGACGACCACCACCTCGCCAGCAGCGACCTCGAGGT
CGATGTCCTTGAGTACGTGCAGGCTGCCGAAGTGCTTGTTCACCTTCTCCAGTTGCACGATGGGAGCGTCAGTGGACGGAGACTTGTCGTGATTCATGTC
GCGCGTTCCTTATTGTCCGACCAGGCCTTTGCGCTCCAGCTGGCGCAGCGCAAACGACAGGCTCAGGGTGATCGCCAGATAGAGCAGAGCCACCATCAGG
TAGACTTCCAGGGCGGTGAAGGTGGTCGCGATGTAGACCTGACCCTGACGAACCAGCTCGCCGACCCCGATGACCGAGAACAGTGACGTGTCCTTGATGC

>Halomonas gudaonensis CGMCC 1.6133 [NZ_FNES01000002.1:99437..99494], dnaA> ig dnaN>, 58 bp

WebSkew GCmin pos. nd (GCmin<>oriC1 nd; GCmin<>oriC2 nd; oriC2<>oriC1 nd)

>Halomonas aquamarina 558 [NZ_FODB01000017.1:18821..19320], <rpmH oriC1 dnaA>, 534 bp

TCTAGAACATCCGTGGTGACCGCAATGAAGTCACGGCTGGCGGTGTCGCTTTGCTCATCGCCAACGCCTTCGCTAATAGCGTTGTTAGGCGCAGAAGTAC
TGGGGACGGAGAGATCGTCGCTACCGCTTTCGTTCGCAGGAGCCCCATTGCTGCTTTGGGTAATCTCTGGCGTCGAATCGTAGGTGGTCTGCCCGTAATC
CTGATTCCACTGTATAACTAGAAGGTAGGCTAGTACGGCCAGTGGAATCAGTAATAAAAGTCGTTTTACGTCCATGAGGGTTCTGCCCGATGGGTGGTGA
ACATGCTAATGGCGCAGTGTCATTTACTCGACCCATGACACACGAAAGCCTGCCAAGCGGCAGGCCAAGGTCGAGCGGCGAGGTGCGTCGTCAAGCGTCA
CGTGTTACGCCATTAGGCGGGGGAGATGCAGACACCGCCTGTGCGTCGCGCTGAAGCCGTTTCCACATGCCGTGTAGCTGGCGAGTGAGCGTTGCGTTAT
CGATATCCTGAGCGCCGCGCTTTGCCAATATCACAATATCCACAGAGGGTAAGTGATCTTGACGCAGACGAACGGATTCCCGAACGAGTCGCTTGAAGCG
GTTACGATCAACAGCGAGCTTCACATTTTTTTTGCTGACCACCAGTCCAATGCGGGGATGTCCCCGGGTACTCATGCGCGCGAGCGCCATCATACCTTTG
CCATGAACTTTCAGGTCTGCCTGTTCAAACACGTGACTAAAATCCCCGGCATTTAGCAAACGTAAGCACCGGGGAAAGCCTTGTTGCGACACACGCAATC
CGAGATCAGGCGCTCAGACGCTTGCGGCCTTTGGCGCGGCGGCGTGCGATAACGGCACGGCCGTTTTTGGTTGCCATACGAGCACGGAAGCCATGCGCGC
GCTTGCGCTTGAGAACGCTGGGTTGAAAAGTGCGTTTCATAACTCGTTATTCCCACGTGGTTTAGGACGGAAAGTGAATTCCAAAAGACCCGGTCCAGTG

AGGAACTAGCCGGGGAGTTTCAATTCAAGACCGGAAATTGTAGAGACTTCCCGCGCCGGGTGCAATTTTTCCTTGCATAAAAAGCTCGAAAAAGGCAATG
GCGGCGCGTCTCCGCTGTGGAGTCGATTCAACTTGGGGCACACTACTATTACTAATGAAGTTATTTAAAACCTACTGTTATTAATAGTGGGGATACTATT
**TGTGGATAA**GTGGTTTTAGCACAATTTTATCAACAGCTTAAAATAATAAAAAATCTGTGTATAAGCATCGAATAACGAGCGAATAAACTGTGAGTTTGAT
GTGGTCAACCGAAGACAAAACAGCCCAAAATAGAGTTGTACCCATGGCATCACTGGGTTTTCCTGAACCCTTACCTAAAGTTGTCCACAGGTGAAAAAGA
GCCGTTAGTGCGTGTGGATAACCACTTCGTTGGGCGCTACAATGGTCGGCCGTTTTCAACCGATGGTGAGATGAGTGTCACTCGCGCTTTGGCAACAGTG

CCTGGACTACCTGCAGGACGAACTGAATTCACAGCAGTTCAATACCTGGATACGTCCGCTGCAGGCAGAAGAAGGAGAGTCCAATGAGCTACGCTTGTTG
GCGCCGAATCGTTTTGTGCGCGACTGGGTGAGTGATAAGTACGCCAAGCGTATTAGTGAGCTAATGCGCGAGCTTGCCCCTTCTAAGCCTCCCAAAGTGA
GCCTGACCGTGGGCAGCCGACGAGCTGCAGCCCCTGCGCCACAGCCTAGAGAGCTTGGCAATCCGGTCTCGGCAGCTCCCGCTCAACCGGCTTCACCTGC
GGTCCATACGCCCCCCCGGGGCGATATTGGCGATGAGCGTGAAATCGACCAGCTACGTGAAGAGAGTGGCCCTAGCCGTCGTGGTAACGAGCGCCAGGTG
CAGGTAGAAGGCAGCTTGAAGCACAGTAGTGGCCTAAACCCTAATTTTACCTTCGACACCTTTGTGGAAGGTAAGTCTAACCAGCTGGCGCGTGCGGCTT
CTCGTCAGGTCTCCGAAAATCCTGGTGGCGCTTATAACCCGCTGTTTTTATACGGTGGTGTTGGCTTAGGTAAAACGCACCTAATGCATGCAGTGGGTAA
TGCGCTGGCGGGCCGACGTGAAAATGCCCGCGTGGTCTACCTGCACTCCGAGCGTTTTGTTGCTGATATGGTGAAGGCGCTGCAGCTTAATGCGATTAAC
GACTTTAAGCGTTTCTATCGCAGCGTCGATGCCCTGCTGATTGACGATATTCAATTTTTTGCCGGTAAAGAGCGTTCTCAGGAAGAGTTTTTCCACACGT
TCAACGCGCTGTTAGAGGGTGGTCAGCAGATGATCCTTACCTCTGACCGCTACCCCAAAGAGATCAGTGGCGTTGAAGAGCGGCTGAAGTCGCGCTTTGG
GTGGGGGCTGACCGTCGCCATTGAGCCGCCGGAGCTTGAGACCCGAGTGGCGATTCTGATGAAGAAAGCCGATCAAGCCAAAGTGGATCTACCCCACGAT

>Halomonas aquamarina [NZ_FODB01000057.1:22528..23027], <mnmG oriC2 |, 522 bp

CTAACCCTTTGATCGAGCGCACCACCTGCAGCTGCACATCAAACGGCAGCGAGGTCGAGATGCCGTTGGGATAGAGCTCGTGGGTATCCAGGCCTTCAGG
CTCGATGAACACCTGATGGCTCGATTTGTCGGCAAAGCGGTGCACTTTATCTTCAATCGAGGGGCAGTAGCGCGGGCCGATCCCTTCAATCACGCCAGAG
TACATTGGCGAGCGATCCAAGTTACCCAGAATAATCTCGTGCGTGCGCTCGTTGGTGTGGGCGATATGGCAGCTCACCTGCTGGGGATGCATCTCCCGGG
AGCCGAGGTAAGACATCACCGGCGTTGGGGTATCGCCCGGCTGCTCTTCCAACTGCGAAAAATCGACGGTTTTGGCATCGATACGCGGCGGCGTGCCAGT
TTTGAGCCGGTCAACATGAAACGGCAACGCACGCAGTCGTTCTGCCAGCGCGTTGGAGGGCGGATCACCCGCGCGGCCTCCGCGGCTTTGATCCAACCCG
ATGTGGATAACGCCACCCAGGAAAGTACCGGTTGAGAGCACGACCGATTCCGCGTGAAAACGAATGCCGGTTTCAGTTACGACGCCGCGCACAGTGTCGT
TATCCACAATGAGATCACCTGCCGCCTGCTGAAAAATCGTCAGGTTGGGCTGGTTTTCCAGCATGCCGCGAATGGCGGCTTTGTAGCGGATACGGTCGGC
TTGGGCACGGGTTGCACGTACCGCTGGCCCTTTGCGTGCGTTGAGCACGCGAAACTGGATGCCGCCTAAATCCGTGGCTAGCCCCATTGCACCGCCGAGC
GCATCAATTTCTTTCACCAAATGGCTTTTGCCGATGCCGCCAATGGCGGGATTGCACGACATTTGGCCAAGCGTCTCGATGTTGTGAGTTAGCAATAGGG
TTTGACAGCCCATACGAGCAGAGGCCAGTGCGGCTTCAGTTCCCGCATGGCCACCGCCGATGACAATCACGTCAAAGCGGTCGGGGTAGTTCAAGAGACA

CCTCGTCTTGCGCCTGTTGGCACGGATGATGTGGTCAGCATATCGGTGCTGTTGGAATAAGCCCGCCAGTATAAACCTCCTGTGGGTAACCGTCAGGTTT
TTCCACACCCCAACCAAAGGGTTCAGTCTTTATTAATAACAAAATAAAAATAAATAAGAGAAGTTCTGTTGATGTAGTAACTACTGGTGTGGACAAAATC
**TGTTGATAA**GTCTACTAACGCTAGCTTTATCAGTATTTTACATTGTTGACCTTTAGGTGATTAAGCAGCGGAGTGCCTGCGTAGAAGCGGTGTTGGAGCT
GTGGATGAAAGTGCGACTTACCCACAAGGCTGACTGACAACGGGTTATCCACCGCTGGGTACCTGGCTTGTTACCGCACCTGTGAACAACCTGCTTCACA
TACTCAAATCATGCTGATAAGCGCTCATGGTGGGCTTCGTGGCGAAGTAGCGAGGAGCGTGTAAAAAATAGTATCCACAGGATCGAGTAGGAGGGGGAGT

CGCCTCCCCCGTCCTCTCACACCACCGGGCAAACGGATCCGTACCACGGCGGTTCCTGTCCCCCATTATCGGGCTGAGCTTCGTTCCGCTGAATCCGCTC
GCCTGTGTGCTCTGAGCGCTTCGACAGTCTTTGGGCTTCCGGCCCGCCATACAGTTATCGAGAGCCCTCCGGGGAGGCTTCTGCTCGTACACATCAGAGA
GTGGTCAACTATCCACTCGTGACAGGTTCAGCCCTTCAGCTCTTGGTTAGAGAGCCTAATATGGCCTCTGCTGACTCCTGTTCGCCCATCCCGACACCTC
ACGGTGTCG

>Halomonas aquamarina 558 [NZ_FODB01000017.1:20765..20822], dnaA> ig dnaN>, 58 bp

WebSkew GCmin pos. 1827760 (GCmin<>oriC1 55.5 kb; GCmin<>oriC2 5.5 kb; oriC2<>oriC1 49.9 kb)

>Halomonas meridiana ACAM 246 [NZ_FSQY01000001.1:1882976..1883475], <rpmH oriC1 dnaA>, 534 bp

TCTAGAACATCCGTGGTGACCGCAATGAAATCACGGCTGGCGGTGTCGCTTTGCTCATCGCCAACGCCTTCGCTAATAGCGTTGTTAGGCGCAGAAGTAC
TGGGGACGGAGAGATCGTCGTTACCGCTTTCGTTCGCAGGAGCCCCATTGCTGCTTTGGGTGATCTCTGGCGTCGAATCGTAGGTGGTCTGCCCGTAATC
CTGATTCCACTGTATAACCAGAAGGTAGGCGAGTACGGCCAGTGGAATCAGTAATAAAAGTCGTTTTACGTCCATGAGGGTTCTGCCCGATGGGTGGTGA
ACATGCTAATGGCGCAGTGTCATTTACTCGACCCATGACACACGAAAGCCTGCCAAGCGGCAGGCCAAGGTCGAGCGGCGAGGTGCGTCGTCAAGCGTCA
CGTGTTACGCCATTAGGCGGGGGAGATGTAGACACCGCCTGTGCGTCGCGCTGAAGCCGTTTCCACATGCCGTGTAGCTGGCGAGTGAGCGTTTCGTTAT
CGATATCCTGAGCGCCGCGCTTTGCCAATATCACAATATCCACAGAGGGTAAGTGATCTTGACGCAGACGAACAGATTCCCGAACGAGTCGCTTGAAGCG
GTTACGATCAACAGCGAGCTTCACATTTTTTTTGCTGACCACCAGTCCAATGCGGGGATGTCCCCGGGTACTCATGCGCGCGAGCGCCATCATACCTTTG
CCATGAACTTTCAGGTCTGCCTGTTCAAACACGTGACTAAAATCCCCGGCATTTAGCAAACGTAAGCACCGGGGAAAGCCTTGTTGCGACACACGCAATC
CGAGATCAGGCGCTCAGACGCTTGCGGCCTTTGGCGCGGCGGCGTGCGATAACGGCACGGCCGTTTTTGGTTGCCATACGAGCACGGAAGCCATGCGCGC
GCTTGCGCTTGAGAACGCTGGGTTGAAAAGTGCGTTTCATAACTCGTTATTCCCACGTGGTTTAGGACGGAAAGTGAATTCCAAAAGACCCGGTCCAGTG

AGGAACTAGCCGGGGAGTTTCAATTCAAGACCGGAAATTGTAGAGACTTCCCGCGCCGGGTGCAATTTTTCCTTGCATAAAAAGCTCGAAAAAGGCAATG
GCGGCGCGTCTCCGCTGTGGAGTCGATTCAACTTGGGGCACACTACTATTACTAATGAAGTTATTTAAAACCTACTGTTATTAATAGTGGGGATACTATT
**TGTGGATAA**GTGGTTTTAGCACAATTTTATCAACAGCTTAAAATAATAAAAAATCTGTGTATAAGCATCGAATAACGAGCGAATAAACTGTGAGTTTGAT
GTGGTCAACCGAAGACAAAACAGCCCAAAATAGAGTTGTACCCATGGCATCACTGGGTTTTCCTGAACCCTTACCTAAAGTTGTCCACAGGTGAAAAAGA
GCCGTTAGTGCGTGTGGATAACCACTTCGTTGGGCGCTACAATGGTCGGCCGTTTTCAACCGATGGTGAGATGAGTGTCACTCGCGCTTTGGCAACAGTG

CCTGGACTACCTGCAGGACGAACTGAATTCACAGCAGTTCAATACCTGGATACGTCCGCTGCAGGCAGAAGAAGGAGAGTCCAATGAGCTACGCTTGTTG
GCGCCGAATCGTTTTGTGCGCGACTGGGTGAGTGATAAGTACGCCAAGCGTATTAGTGAGCTAATGCGCGAGCTTGCCCCTTCTAAGCCTCCCAAAGTGA
GCCTGACCGTGGGCAGCCGACGAGCTGCAGCCCCTGCGCCACAGCCTAGAGAGCTTGGCAATCCGGTCTCGGCAGCTCCCGCTCAACCGGCTTCACCTGC
GGTCCATACGCCCCCCCGGGGCGATATTGGCGATGAGCGTGAAATCGACCAGCTACGTGAAGAGAGTGGCCCTAGCCGTCGTGGTAACGAGCGCCAGGTG
CAGGTAGAAGGCAGCTTGAAGCACAGTAGTGGCCTAAACCCTAATTTTACCTTCGACACCTTTGTGGAAGGTAAGTCTAACCAGCTGGCGCGTGCGGCTT
CTCGTCAGGTCTCCGAAAATCCTGGTGGCGCTTATAACCCGCTGTTTTTATACGGTGGTGTTGGCTTAGGTAAAACGCACCTAATGCATGCAGTGGGTAA
TGCGCTGGCGGGCCGACGTGAAAATGCCCGCGTGGTCTACCTGCACTCCGAGCGTTTTGTTGCTGATATGGTGAAGGCGCTGCAGCTTAATGCGATTAAC
GACTTTAAGCGTTTCTATCGCAGCGTCGATGCCCTGCTGATTGACGATATTCAATTTTTTGCCGGTAAAGAGCGTTCTCAGGAAGAGTTTTTCCACACGT
TCAACGCGCTGTTAGAGGGTGGTCAGCAGATGATCCTTACCTCTGACCGCTACCCCAAAGAGATCAGTGGCGTTGAAGAGCGGCTGAAGTCGCGCTTTGG
GTGGGGGCTGACCGTCGCCATTGAGCCTCCGGAGCTTGAGACCCGAGTGGCGATTCTGATGAAGAAAGCCGATCAAGCCAAAGTGGATCTACCCCACGAT

>Halomonas meridiana ACAM 246 [NZ_FSQY01000001.1:1833057..1833556], <mnmG oriC2 <COG1126 (glnQ), 579 bp

CTAACCCTTTGATCGAGCGCACCACCTGCAGCTGCACATCAAACGGCAGCGAGGTCGAGATGCCGTTTGGATAGAGCTCGTGGGTATCCAGGCCTTCAGG
CTCGATGAACACCTGATGGCTCGATTTGTCGGCAAAGCGGTGCACTTTATCTTCAATCGAGGGGCAGTAGCGCGGGCCGATCCCTTCAATCACGCCAGAG
TACATTGGCGAGCGATCCAAGTTACCCAGAATAATCTCGTGGGTGCGCTCGTTGGTGTGCGCGATATGGCAGCTCACCTGCTGGGGATGCATCTCCCGCG
AGCCGAGGTAAGACATCACCGGCGTTGGGGTATCGCCCGGCTGCTCTTCCAACTGCGAAAAATCGACGGTTTTGGCATCGATACGCGGCGGCGTGCCGGT
TTTGAGCCGGTCAACGTGAAACGGCAACGCACGTAGTCGTTCTGCCAGCGCGTTGGAGGGCGGATCACCCGCGCGGCCTCCGCGGCTTTGATCCAACCCG
ATGTGGATAACGCCACCGAGGAAAGTACCGGTTGAGAGCACGACCGATTCCGCGTGAAAGCGAATGCCGGTTTCAGTTACGACGCCGCGCACGGTGTCGT
TATCCACAATGAGATCACCTGCTGCCTGCTGAAAAATTGTCAGGTTGGGTTGGTTTTCCAGCATCCCGCGAATGGCTGCTTTGTAGCGGATGCGGTCGGC
CTGGGCACGGGTTGCGCGCACCGCTGGCCCTTTGCGTGCGTTGAGCACGCGAAACTGGATGCCGCCTAAATCTGTGGCTAGCCCCATTGCACCGCCGAGC
GCATCAATTTCTTTCACCAAATGGCTTTTGCCGATGCCGCCAATGGCGGGATTGCACGACATTTGGCCAAGTGTCTCGATGTTGTGGGTTAGCAATAGGG
TTTGACAGCCCATACGAGCAGAGGCCAGTGCGGCTTCAGTTCCCGCATGGCCACCGCCGATGACAATCACGTCAAAGCGGTCGGGGTAGTTCAAGTGACA

CCTCGTCTTGTGCCTGTTGGCACGGATGATGTGGTCAGCATATCGGCGCTGATGGAATAAGCCCGCCAGTATAAACCTCCTGTGGGTAACCGTCAGGTTT
TTCCACACCCCGGTTTAAGGGTTCTGTCTTTATTAATAACAAAATAAAAATAAATAAGAGAAGTTCTGTTGATGTAGTAACTACTGGTGTGGACAAAATC
**TGTTGATAA**GTCTACTAACGCTAGCTTTATCAGTATTTTACATTGTTGACCTTTAGGTGATTAAGCAGCGGAGTGCCTGCGTAGAAGCGGTGTTGGAGCT
GTGGATGAAAGTGCGACTTACCCACAAGGCTGACTGACAACGGGTTATCCACCGCTGGGTACCTGGCTTGTTACCGCACCTGTGAACAACCTGCTTCACA
TACTCAAATCATGCTGATAAGCGCTCATGGTGGGCTTCGTGGCGAAGTAGCGAGGAGCGTGTAAAAAATAGTATCCACAGGCAAAAAAAAATGGCCTGTC

ATGAAGACAGGCCATTAGATAGGACGTTTTAAGCGCAATATTGGCGATTTAAGAGCGCATGGTTAGCACGACATTAGTGCTTCAGTACCCGTGCCAGGAA
CGACTGAGTGCGTTCGTGTTGAGGGGAATCGAAGAGCTGCTCAGGGGTGCCTTGTTCGGCAATTTCGCCTTTATGGATAAAGATGATGCGGTCGGCTACC
TCACGTGCAAAGCCCATCTCGTGAGTAACAATCACCATGGTCATACCCTCTTTGGCAAGCTCCCGCATGGCGTCTAGCACTTCGCCAATCATCTCGGGGT
CAAGGGCTGAGGTTGGCTCATCAAACAGCATGAGGCGCGGCTCCATGGCCAACGCACGCGCCAGCGCCACACGCTGCTGCTGGCCACCGGAGAGCTGGCT
GGGGTACTTATTTGCCTGGTCGGCAATACCTACGCGATCAAGTAGCCGCTTGGCAGTCTCTTCGGCATCCTGACGGCTCCAGCCACGCACCTTCATCGGT
GCGAGAGTGACGTTATCGAGCACGCTAAGATGCGGGAACAGATTGAACTGCTGAAACACCATGCCCACTTCGGTGCGGATGGTTTGTAGCGCCTGACTGC
TTTTACCGTTCGGTAGCAGCGTATTGCCGTCCACATCCAAGCTGCCTGACTGAAACTCTTCTAAGCCGTTGATACAGCGAATGAGTGTGGATTTACCCGA
GCCGCTGGCCCCGATGACCACGACGACTTCACCAGGCACAATTTCCAGGTCGATATCTTTGAGTACGTGCAGGCTGCCAAAGTGCTTGTGGAGCTTTTGC
ATACGCACAATGGGCGTTTGAGTAGAAGTCATTGTGCGCTCCTTATTGTCCGGCGAGGCCGCGGCGCTCGATTTGGCGCAGCACCAGTGAGAGGGTCCAG
GTAATGGCGAGATACATCAGTGCCACCATGAAGTAGACCTCCAGCGCGGTGAAGGTGGTGGCGATATAGATCTGCCCTTGGCGTACCAGCTCACTGACCC

>Halomonas meridiana ACAM 246 [NZ_FSQY01000001.1:1884920..1884977], dnaA> ig dnaN>, 58 bp

< page 1 | #064 | Halomonas axialensis FXH-124

WebSkew GCmin pos. nd (GCmin<>oriC1 nd; GCmin<>oriC2 nd; oriC2<>oriC1 nd)

>Halomonas axialensis FXH-124 [JAJGND010000025.1:40907..41406], <rpmH oriC1 dnaA>, 534 bp

TCTAGAACATCCGTGGTGACCGCAATGAAATCACGGCTGGCGGTGTCGCTTTGCTCATCGCCAACGCCTTCGCTAATAGCGTTGTTAGGCGCAGAAGTAC
TGGGGACGGAGAGATCGTCGCTACCGCTTTCGTTCGCAGGAGCCCCATTGCTGCTTTGGGTGATCTCTGGCGTCGAATCGTAGGTGGTCTGCCCGTAATC
CTGATTCCACTGTATAACCAGAAGGTAGGCTAGTACGGCCAGTGGAATCAGTAATAAAAGTCGTTTTACGTCCATGAGGGTTCTGCCCGATGGGTGGTGA
ACATGCTAATGGCGCAGTGTCATTTACTCGACCCATGACACACGAAAGCCTGCCAAGCGGCAGGCCAAGGTCGAGCGGCGAGGTGCGTCGTCAAGCGTCA
CGTGTTACGCCATTAGGCGGGGGAGATGCAGACACCGCCTGTGCGTCGCGCTGAAGCCGTTTCCACATGCCGTGTAGCTGGCGAGTGAGCGTTTCGTTAT
CGATATCCTGAGCGCCGCGCTTTGCCAATATCACAATATCCACAGAGGGTAAGTGATCTTGACGCAGACGAACGGATTCCCGAACGAGTCGCTTGAAGCG
GTTACGATCAACAGCGAGCTTCACATTTTTTTTGCTGACCACCAGTCCAATGCGGGGATGTCCCCGGGTACTCATGCGCGCGAGCGCCATCATACCTTTG
CCATGAACTTTCAGGTCTGCCTGTTCAAACACGTGACTAAAATCCCCAGCGTTTAGCAAACGTAAGCACCGGGGAAAGCCTTGAGGCGACACACGCAATC
CGAGATCAGGCGCTCAGACGCTTGCGGCCTTTGGCGCGGCGGCGTGCGATCACGGCACGGCCGTTTTTGGTTGCCATACGAGCACGGAAGCCATGCGCGC
GCTTGCGCTTGAGAACGCTAGGTTGAAAAGTGCGTTTCATAACTCGTTATTCCCACGTGGTTTAGGACGGAAAGTGAATTCCAAAAGACCCGGTCCAGTG

AGGAACTAGCCGGGGAGTTTCAATTCAAGACCGGAAATTGTAGAGACTTCCCGCGCCGGGTGCAATTTTTCCTTGCATAAAAAGCTCGAAAAAGGCAATA
GCGGCGCGGTTTGGCGCCCACTACTCGTTTTCTTGCATCGCACTACTATTACTAATGAAGTTATTTAAAACCTACTGTTATTAATAGTGGGGATACTATT
**TGTGGATAA**GTCATTTTAGGAGATTATTATCAACAGCTTGAAGAAATAAGAAATCTGTGTATAAGCATCTAATAACGCGCGAATAAACTGTGAATTTGGT
GTGGTCAGGCGGTGACAACTTGCTCGAAAGGAGGGTTGTACCCATGGCATCACCGGGTTTTCCTAAGCCCTTACGGAAAGTTGTCCACAGGCGAAAAAGA
GCTGCTAACGCGTGTGGATAACCGTGTCGTTGGGCGCTACAATGGTCGGCCGTTTTCAACCGATGGTGAGATGAGTGTCACTCGCGCTTTGGCAACAGTG

CCTGGACTACCTGCAGGACGAACTGAATTCACAGCAGTTCAATACCTGGATACGTCCGCTGCAGGCAGAAGAAGGAGAGTCCAATGAGCTACGCTTGTTG
GCGCCGAATCGTTTTGTGCGCGACTGGGTGAGTGATAAGTACGCCAAGCGTATTAGTGAGCTAATGCGCGAGCTTGCCCCTTCTAAGCCTCCCAAAGTGA
GCCTGACCGTGGGCAGCCGACGAGCTGCAGCCCCTGCGCCACAGCCTAGAGAGCTTGGCAATCCGGTCTCGGCAGCTCCCGCTCAACCGGCTTCACCTGC
GGTCCATACGCCCCCCCGGGGCGATATTGGCGATGAGCGTGAAATCGACCAGCTACGTGAAGAGAGTGGCCCTAGCCGTCGTGGTAACGAGCGCCAGGTG
CAGGTAGAAGGCAGCTTGAAGCACAGTAGTGGCCTAAACCCTAATTTTACCTTCGACACCTTTGTGGAAGGTAAGTCTAACCAGCTGGCGCGTGCGGCTT
CTCGTCAGGTCTCCGAAAATCCTGGTGGCGCTTATAACCCGCTGTTTTTATACGGTGGTGTTGGCTTAGGTAAAACGCACCTAATGCATGCAGTGGGTAA
TGCGCTGGCGGGCCGACGTGAAAATGCCCGCGTGGTCTACCTGCATTCCGAACGCTTTGTGGCCGACATGGTAAAAGCGCTGCAGCTTAATGCGATTAAC
GACTTTAAGCGTTTCTACCGCAGCGTTGATGCGCTTCTGATTGACGATATTCAGTTTTTTGCCGGTAAAGAGCGCTCTCAAGAAGAGTTCTTTCATACGT
TTAATGCGCTGTTGGAAGGCGGTCAGCAGATGATTTTGACCTCTGACCGTTATCCCAAAGAGATCAGTGGCGTTGAAGAGCGGCTGAAGTCGCGCTTTGG
GTGGGGGCTGACCGTCGCCATTGAGCCGCCGGAGCTTGAGACCCGAGTGGCGATTCTGATGAAGAAAGCCGATCAAGCCAAAGTGGATCTACCCCACGAT

>Halomonas axialensis FXH-124 [JAJGND010000012.1:11308..11807], <mnmG oriC2 <COG1126 (glnQ), 579 bp

CTAACCCTTTGATCGAGCGCACCACCTGCAGCTGCACATCAAACGGCAGCGAGGTCGAGATGCCGTTGGGATAGAGCTCGTGGGTATCCAGGCCTTCAGG
CTCGATGAACACCTGATGGCTCGATTTGTCGGCAAAGCGGTGCACTTTATCTTCAATCGAGGGGCAGTAGCGCGGGCCGATCCCTTCAATCACGCCAGAG
TACATTGGCGAGCGATCCAAGTTACCCAGAATAATCTCGTGGGTACGCTCGTTGGTGTGCGCGATATGGCAACTTACCTGCTGGGGATGCATCTCCCGCG
AGCCGAGGTAAGACATCACCGGCGTTGGGGTATCGCCCGGCTGCTCTTCCAACTGCGAAAAATCGACGGTTGTGGCATCGATACGCGGCGGCGTGCCGGT
TTTGAGCCGGTCAACGTGAAACGGCAACGCACGCAGTCGTTCTGCCAGCGCGTTGGAGGGCGGATCACCCGCGCGGCCTCCGCGGCTTTGATCCAACCCG
ATGTGGATAACGCCACCTAGGAAGGTACCGGTTGAGAGCACAACGGATTCTGCGTGAAAGCGAATGCCGGTTTCAGTCACGACGCCGCGTACGGTGTCGT
TATCCACAATGAGATCACCTGCCGCCTGCTGAAAAATCGTCAGGTTGGGCTGGTTTTCCAGCATGCCGCGAATGGCGGCTTTGTAGCGGATACGGTCGGC
TTGGGCACGGGTTGCACGTACCGCTGGCCCTTTGCGTGCGTTGAGCACGCGAAACTGGATGCCGCCTAAATCCGTGGCTAGCCCCATTGCACCGCCGAGC
GCATCAATTTCTTTCACCAAATGGCTTTTGCCGATGCCGCCAATGGCGGGATTGCACGACATTTGGCCAAGCGTCTCGATGTTGTGAGTTAGCAATAGGG
TTTGACAGCCCATACGAGCAGAGGCCAGTGCGGCTTCAGTTCCCGCATGGCCACCGCCGATGACAATCACGTCAAAGCGGTCGGGGTAGTTCAAGAGACA

CCTCGTCTTGCGCCTGTTGGCACGGATGATGTGGTCAGCGATTCAGCGCTGTTGGAATAAGCCCGCCAGTATAAACCTCCTGTGGGTAACCGTCAGGTTT
TTCCACACCCCAACCAAAGGGTTCAGTCTTTATTAATAACAAAATAAAAATAAATAAGAGAAGTTCTGTTGATGTAGTAACTAACTGGTGTGGACAAAATC
**TGTTGATAA**GTCTACTAACGCTAGCTTTATCAGTATTTTACATTGTTGACCTTTAGGTGATTAAGCAGCGGAGTGCCTGCGTAGAAGCGGTGTTGGAGCT
GTGGATGAAAGTGCGACTTACCCACAAGACTGACTGACAACGGGTTATCCACCGCTGGGTACCTGGCTTGTTACCGCACCTGTGAACAACCTGCTTCACA
TACTCAAATCATGCTGATAAGCGCTCATGGTGGGCTTCGTGGCGAAGTAGCGAGGAGCGTGTAAAAAATAGTATCCACAGGCAAAAAAAAATGGCCTGTC

ATGAAGACAGGCCATTAGATAGGACGTTTTAAGCGCAATATTGGCGATTTAAGAGCGCATGGTTAGCACGACATTAGTGCTTCAGTACCCGTGCCAGGAA
CGACTGAGTGCGTTCGTGTTGAGGGGAATCGAAGAGCTGCTCAGGGGTGCCTTGTTCGGCAATTTCGCCTTTATGGATAAAGATGATGCGGTCGGCTACC
TCACGTGCAAAGCCCATCTCGTGAGTAACAATCACCATGGTCATACCCTCTTTGGCAAGCTCCCGCATGGCGTCTAGCACTTCGCCAATCATCTCGGGGT
CAAGGGCTGAGGTTGGCTCATCAAACAGCATGAGGCGCGGCTCCATGGCCAACGCACGCGCCAGCGCCACACGCTGCTGCTGGCCACCGGAGAGCTGGCT
GGGGTACTTATTTGCCTGGTCGGCAATACCTACGCGATCAAGTAGCCGCTTGGCAGTCTCTTCGGCATCCTGACGGCTCCAGCCACGTACCTTCATCGGT
GCGAGAGTGACGTTATCGAGCACGCTAAGATGCGGGAACAGATTGAACTGCTGAAACACCATGCCCACTTCGGTGCGGATGGTTTGCAGCGCCTGACTGC
TTTTACCGTTCGGTAGCAGCGTATTGCCGTCCACATCCAGGCTGCCTGACTGAAACTCTTCTAAGCCGTTGATACAGCGAATGAGTGTGGATTTACCCGA
GCCGCTGGCCCCGATGACCACGACGACTTCACCAGGCACAATTTCCAGGTCGATATCTTTGAGTACGTGCAGGCTGCCAAAGTGCTTGTGGAGCTTTTGC
ATACGCACAATGGGCGTTTGAGTAGAAGTCATTGTGCGCTCCTTATTGTCCGGCGAGGCCGCGGCGCTCGATTTGGCGCAGCACCAGTGAGAGGGTCCAG
GTAATGGCGAGATACATCAGCGCCACCATGAAGTAGACCTCCAGCGCGGTGAAGGTAGTGGCGATATAGATCTGCCCTTGGCGTACCAGCTCACTGACCC

>Halomonas axialensis FXH-124 [JAJGND010000025.1:39402..39462], dnaA> ig dnaN>, 61 bp

WebSkew GCmin pos. nd (GCmin<>oriC1 nd; GCmin<>oriC2 nd; oriC2<>oriC1 nd)

>Halomonas lionensis RHS90 [NZ_MWVI01000087.1:40606..41105], <rpmH oriC1 dnaA>, 534 bp

AAGTACGTCGGTGGGGACGGCAATAAAGTCGCGGCTGGCGGTGTCGCTTTCAGCATCACCGACGCATTCGCTGATGGCGTTTTGAGGAGCGGAAGTGCTA
GGTACAGAGAGATCATCGCTACCGTTCTCGCTCGCAGGAGCCCCGTTTCCTGCTTTGGGTTATCTCAGGCGTTGAGTCGTAGGTGGTCTGCCCGTAATCC
TGATTCCACTGTACAACCAAAAGGTAGGCAAGCACTGCCAGTGGAATCAGTAATAAAAGTCGTTTTACGTCCATGAGGGTTCTGCCCGATGGGTGGTGAA
CATGCTAATGGCGCAGTGTCATTTACTCGACCCATGACACACGAAAGCCTGCCAAGCGGCAGGCCGAGGTCGAGCGGCGAGGTATGTCGTCAAGCGTCAC
GTGTTACGCCATGAGGCGGGGGAGGTGCAGACACCGCCTGTGCGTCGCGATGAAGTCGTTTCCACATGCCGTGTAGCTGGCGAGTAAGCGTTTCGTTATC
GATATCCTGCACGCCGCGCTTCGCCAATATCACGATATCCACAGAGGGTAAGTGATCCTGACGTAGACGAACGGATTCCCGGACGAGTCGCTTGAAGCGG
TTACGATCAACAGCGAGCTTCACATTTTTTTTGCCTGACCACCAGTCCAATACGGGGATGCCCCCGGGTACTCATGCGCGCTAGCGCCATCATACCTTTG
CCATGAACTTTCAGGTCGGCCTGTTCAAACACGTGACTAAAATCCCCGGCATTTAGCAAACGTAAGCACCGGGGAAAGCCTTGTTGCGACACACGCAATC
CGAGATCAGGCGCTCAGACGCTTGCGGCCTTTGGCGCGGCGGCGTGCGATAACGGCACGGCCGTTTTTGGTTGGCATACGAGCACGGAAGCCATGCGCGC
GCTTGCGCTTGAGAACGCTGGGTTGAAAAGTGCGTTTCATAACTCGTTATTCCCACGTGGTTTAGGACGGAAAGTGAATTCCAAAAGACCCGGTCCAGTG

AGGAACTAGCCGGGGAGTTTCAATTCAAGACCGGAAATTGTAGAGACTTCCCGCGCCGGGTGCAATTTTTCCTTGCATAAAAAGAGTGAAAAAGGCAATA
GCGCTGCCGCTTCCATCAGGTTTTGATTCAACCAAGGCCACACTACTATTACTAATGAAGTTATTTAAAACCTACTGTTATTAATAGTGGGGATACTATT
**TGTGGATAA**GTGGTTTTAGCATAATTTTATCAACAGGTTAAAATAATAAAAAATCTGTGTATAAGTATCAAATAACCGGCCAATAAACTGTGAATTAGAT
GTGTTCAACGTAAGACAACTTACTCAAAAATACCCTTGTACCCATTTCATTACCGGGTTTTCCTTAGCCCTCACGCACAGTTGTCCACAGGCAAAACAGA
GCCGCTAGCGCGAGTGGATAACCGCTTCGTTGGGCGCTACAATGGTCGGCCGTTTTCAACCGATGGTGAGATGAGTGTCGCTCGCGCTTTGGCAACAGTG

CCTGGACTACCTGCAGGACGAACTGAATTCACAGCAGTTCAACACCTGGCTACGTCCGCTGCAGGCAGAAGAAGGAGACACCAATGAGCTACGCTTATTG
GCGCCGAATCGCTTTGTGCGCGACTGGGTGAGTGATACGTCCGCTAAGCGTATTAGTGAGCTAATGCGTGAGCTCGCCCCTTCTAAACCTCCCAAGGTCA
GCTTAACCGTCGGCAGCCGGCGGGCTGCAGCGGCTGCGCCGCAACCCAGAGAGGTGGGTAATCCGGTCTCGGCCGCACCAGCACAGCCGTCATCCCCCGC
CGTTCATACTCCGCCGCGTGGTGATATTGGCGATGAGCGTGAGATTGACCAACTGCGTGAAGAGAATGCCCCTAGCCGTCGTGGCAATGAGCGCCAGGTT
CAAGTCGACGGCAGCTTGAAGCACAGCAGTGGCTTAAACCCTAATTTCACCTTCGATACCTTTGTAGAAGGTAAATCGAACCAACTTGCCCGTGCAGCTT
CCCGCCAGGTGTCGGAAAATCCTGGTGGCGCTTATAACCCGCTGTTTTTATACGGTGGTGTTGGCTTGGGCAAAACGCACTTGATGCATGCTGTGGGCAA
TGCATTGGCAGGCCGTCGTGAAAATGCACGCGTGGTCTACCTGCACTCCGAGCGTTTTGTTGCTGATATGGTGAAGGCGCTGCAGCTTAATGCGATTAAC
GACTTTAAGCGTTTCTATCGCAGCGTCGATGCCCTGCTGATTGACGATATTCAATTTTTTGCCGGTAAAGAGCGTTCTCAGGAAGAGTTTTTCCACACGT
TCAACGCGCTGTTAGAGGGTGGTCAGCAGATGATTCTCACCTCTGACCGCTACCCTAAAGAGATTAGTGGCGTTGAAGAGCGTCTTAAGTCGCGCTTTGG
CTGGGGGCTTACCGTGGCGATCGAGCCGCCGGAGCTTGAGACACGAGTGGCGATTTTGATGAAAAAAGCCGATCAAGCAAAAGTGGATTTGCCCCACGAT

>Halomonas lionensis RHS90 [NZ_MWVI01000003.1:c1..377], <mnmG oriC2 (3'-part from H. axaliensis), 579 bp

TAACCCTTTGATCGAGCGCACCACCTGCAGCTGCACATCAAACGGCAGCGAGGTCGAGATGCCGTTGGGATAGAGCTCGTGGGTATCCAGGCCTTCAGGC
TCGATGAACACCTGATGGCTCGATTTGTCGGCAAAGCGGTGCACTTTATCTTCAATCGAGGGGCAGTAGCGCGGGCCGATCCCTTCAATCACGCCAGAGT
ACATTGGCGAGCGATCCAAGTTACCCAGAATAATCTCGTGGGTGCGCTCGTTGGTGTGCGCGATATGGCAGCTCACCTGCTGGGGATGCATCTCCCGCGA
GCCGAGGTAAGACATCACCGGCGTTGGGGTATCGCCCGGCTGCTCTTCCAACTGCGAAAAATCGACGGTTTTGGCATCGATACGCGGCGGCGTGCCGGTT
TTGAGCCGGTCAACGTGAAACGGCAACGCACGCAGTCGTTCTGCCAGCGCGTTGGAGGGCGGATCACCCGCGCGGCCTCCGCGGCTTTGATCCAACCCGA
TGTGGATAACGCCACCCAGGAAGGTACCGGTTGAGAGCACAACGGATTCCGCGTGAAAGTGAATGCCGGTTTCAGTCACGACGCCGCGTACGGTGTTGTT
ATCCAACAATGAGATCACCTGCCGCCTGCTGAAAAATCGTCAGGTTGGGCTGGTTTTCCAGCATGCCGCGAATGGCGGCTTTGTAGCGGATGCGGTCGGC
CTGGGCACGGGTTGCACGTACCGCTGGCCCTTTGCGTGCGTTGAGCACGCGAAACTGGATGCCGCCTAAATCCGTGGCTAGCCCCATTGCACCGCCGAGC
GCATCAATTTCTTTCACCAAATGGCTTTTGCCGATGCCGCCAATGGCGGGATTGCACGACATTTGGCCAAGCGTCTCGATGTTGTGAGTTAGCAATAGGG
TTTGACAGCCCATACGAGCAGAGGCCAGTGCGGCTTCAGTTCCCGCATGGCCACCGCCGATGACAATCACGTCAAAGCGGTCGGGGTAGTTCAAGAGACA

CCTCGTCTTGCGCCTGTTGGCACGGATGATGTGGTCAGCGATTCAGCGCTGTTGGAATAAGCCCGCCAGTATAAACCTCCTGTGGGTAACCGTCAGGTTT
TTCCACACCCCAACCTAAGGGTTCAGTCTTTATTAATAACAAAATAAAAATAAATAAGAGAAGTTCTGTTGATGTAGTAACTACTGGTGTGGACAAAATC
**TGTTGATAA**GTCTATTAAACCTATTTTTATCAATATTTTAAATTGTTGACCTTTAGGTGATTAAGCAGCGGAGTGCCTGCGTAGAAGCGGTGTTAGAGCT
GTGGATGAAAGTGCGACTTACCCACAAGACTGACTGACAACGGGTTATCCACCGCTGGGTACCTGGCTTGTTACCGCaccTGTGAACAAcctgcttcaca
tactcaaatcaTGCTGATAAgcgctcatggtgggcttcgTGGCGAAGTagcgaggagcgTGTAAAAAAtaGTATCCACAggcaaaaaaaaatggcctgtc

atgaagacaggccattagataggacgttttaagcgcaatattggcgatttaagagcgcatggTTAGCACGAcattagtgcttcagtacccgtgccaggaa
cgactgagtgcgttcgtgttgaggggaatcgaagagctgctcaggggtgccttgttcggcaatttcgcctttatggataaagatgatgcggtcggctacc
tcacgtgcaaagcccatctcgtgagtaacaatcaccatggtcataccctctttggcaagctcccgcatggcgtctagcacttcgccaatcatctcggggt
caagggctgaggttggctcatcaaacagcatgaggcgcggctccatggccaacgcacgcgccagcgccacacgctgctgctggccaccggagagctggct
ggggtacttatttgcctggtcggcaatacctacgcgatcaagtagccgcttggcagtctcttcggcatcctgacggctccagccacgtaccttcatcggt
gcgagagtgacgttatcgagcacgctaagatgcgggaacagattgaactgctgaaacaccatgcccacttcggtgcggatggtttgcagcgcctgactgc
ttttaccgttcggtagcagcgtattgccgtccacatccaggctgcctgactgaaactcttctaagccgttgatacagcgaatgagtgtggatttacccga
gccgctggccccgatgaccacgacgacttcaccaggcacaatttccaggtcgatatctttgagtacgtgcaggctgccaaagtgcttgtggagcttttgc
atacgcacaatgggcgtttgagtagaagtcattgtgcgctccttattgtccggcgaggccgcggcgctcgatttggcgcagcaccagtgagagggtccag
gtaatggcgagatacatcagcgccaccatgaagtagacctccagcgcggtgaaggtagtggcgatatagatctgcccttggcgtaccagctcactgaccc

>Halomonas lionensis RHS90 [NZ_MWVI01000087.1:39104..39161], dnaA> ig dnaN>, 58 bp

< page 1 | #066 | Halomonas piezotolerans NBT06E8

WebSkew GCmin pos. 3945801/0 (GCmin<>oriC1 0 kb; GCmin<>oriC2 21.1 kb; oriC2<>oriC1 21.1 kb)

>Halomonas piezotolerans NBT06E8 [NZ_CP048602.1:3945328..3945801], <rpmH oriC1 dnaA>, 534 bp

TCCAGAACGTCCGTGGTGACCGCAATGAAATCACGGCTGGCGGTGTCGCTTTGCTCATCGCCAACGCCTTCGCTAATAGCGTTGTTAGGCGCCGAAGTAC
TGGGCACGGAGAGATCGTCGCTACCGCTTTCGTTCGCAGGAGCCCCATTGCTGCTTTGGGTGATCTCTGGCGTCGAATCGTAGGTGGTCTGCCCGTAATC
CTGATTCCACTGTACAACCAGAAGGTAGGCTAGTACGGCCAGTGGAATCAGTAATAAAAGTCGTTTTACGTCCATGAGGGTTCTGCCCGATGGGTGGTGA
ACATGCTAATGGCGCAGTGTCATTTACTCGACCCATGACACACGAAAGCCTGCCAAGCGGCAGGCCGAGGTCGAGCGGCGAGGTGTGTCGTCAAGCGTCA
CGTGTTACGCCATGAGGCGGGAAAGGTGTTGATGCCGCCTGTGCGTCGCGCTGAAGCCGTTTCCACATGCCGTATAGTTGACGGCTGAGCGTTTCGTTAT
CGACATCCTGCACGCCTCGCTTTGCCAATATCACGATATCCACAGAGGGTAAGTGATCTTGACGCAGACGAACGGATTCGCGAACGAGTCGCTTGAAGCG
GTTACGATCGACAGCAAGCTTTACATTTTTTTTGCTAACGACCAGTCCGATGCGGGGGTATCCCCGGGTAGTCATGCGCGCTAGCGCCATCATACCTTTG
CCATGGACTTTGAGGTCAGCCTGCTCAAACACGTGACTGAAATCCCCGGCAGTTAGCAACCGCAAGCACCGGGGAAACCCTTGAGGCGACACACGCAATC
CGAGATCAGGCGCTCAGACGCTTGCGGCCTTTGGCACGGCGGCGTGCGATCACGGCACGGCCGTTTTTGGTTGCCATACGAGCACGGAAGCCATGCGCGC
GCTTGCGCTTGAGAACGCTGGGTTGAAAAGTGCGTTTCATAACTCGTTATTCCCACGTGGTTTAGGACGGAAAGTGAATTCCAAAAGACCCGGTCCAGTG

AGGAACTAGCCGGGGAGTTTCAATTCAAGACCGGAAATTGTAGAGACTTCCCGCGCTGGGTGCAATTTTTCCTTGCATAAAAAGCTCGAAAAAGGCAATG
GCGGCGCGGCTTGGCGCCCACCACTCGTTTTCTTGCATCGCACTACTATTACTAATGAAGTTATTTAAAACATACTGTTATTAATAGTGGGGATACTATT
**TGTGGATAA**GTCATTTTAGGATATTGTTATCAAAAGCTTGGAGTAATAAAAAATCTGTGAATAAGCATCGAATAACGCGCGAATAAACTGTGAATTTGGT
GTGGTCAGGCGGTGACAACTTGCTCGAAAGGAGGGTTGTACCCATGGCGTCACCGGGTTTTCCTAAGCCCTTACGGAAAGTTGTCCACAGGCGAAAAAGA
GCCGTTAGCTCGTGTGGATAACCGTGTCGTTGGGCGCTACAATGGTCGGCCGTTTTCAACCGATGGTGAGATGAGTGTCACTCGCGCTTTGGCAACAGTG

CCTGGACTACCTGCAGGACGAACTGAATTCACAGCAGTTCAATACCTGGATACGTCCGCTGCAGGCAGAAGAAGGAGACACCAATGAGCTGCGCTTGTTG
GCGCCGAATCGCTTTGTGCGCGACTGGGTGAGTGATAAGTACGCCAAGCGTATTAGTGAATTAATGCGCGAGCTTGCACCCTCCAAGCCCCCTAAGGTCA
GTTTGACGGTAGGTAGCCGCCGGACCGCTGCTCCAGCTCCCCAGCCTCGCGATCTGGGTAATCCGGTGTCAGCCGCCCCGGCTCAGCCGACGTCGCCCGC
CGTACATACCCCGCCCCGGGGTGATATTGCCGACGAGCGTGAAATCGATCAGCTGCGTGAAGAGAGTGCCTCTCATCGTCGTGGCAACGAGCGTCAGGTG
CAGGTAGAAGGCAGCTTGAAACACAATAGCGGCCTAAACCCTAATTTCACGTTCGATACCTTTGTGGAAGGTAAATCTAACCAGTTGGCACGTGCGGCGT
CTCGCCAAGTGTCTGAAAATCCTGGCGGTGCTTATAATCCACTGTTCTTATACGGTGGCGTGGGCTTGGGTAAAACCCACTTGATGCATGCCGTAGGCAA
TGCGTTGGCAGGTCGTCGGGAAAATGCCCGCGTGGTTTATCTTCACTCTGAGCGCTTCGTGGCTGACATGGTGAAGGCGCTGCAGCTCAATGCGATCAAC
GATTTCAAACGTTTCTACCGCAGTGTCGATGCCCTCCTGATCGATGACATTCAGTTCTTTGCCGGTAAAGAGCGCTCCCAAGAAGAGTTCTTCCATACGT
TCAATGCCTTGCTAGAGGGCGGTCAGCAGATGATCTTGACCTCTGATCGCTATCCAAAAGAGATCAGTGGTGTCGAAGAGCGGCTGAAGTCGCGCTTTGG
CTGGGGTCTCACCGTCGCCATCGAACCGCCAGAGCTCGAGACGCGCGTGGCGATTCTGATGAAGAAAGCCGATCAGGCCAAGGTGGACCTGCCCCACGAT

>Halomonas piezotolerans NBT06E8 [NZ_CP048602.1:3923823..3924322], <mnmG oriC2 <COG1126 (glnQ), 578 bp

CGAGCCCTTTGATCGAGCGCACCACCTGCAACTGCACGTCGAAAGGCAGCGAGGTCGAGATGCCGTTGGGATAGAGCTCGTGGGTATCCAGACCTTCCGG
TTCGATGAACACCTGATGGCTCGATTTGTCGGCAAAGCGGTGCACTTTGTCTTCAATCGACGGGCAGTAGCGCGGGCCGATCCCTTCGATTACGCCAGAG
TACATTGGCGAGCGATCCAGGTTACCCAGGATGATCTCGTGGGTACGCTCGTTGGTGTGAGCAATATGGCAGCTCACCTGCTGGGGGTGCATTTCCCGCG
AGCCGAGATAAGACATTACTGGCGTTGGCGTATCGCCCGGCTGCTCTTCCAACTGCGAAAAATCGACGGTCTTGGCGTCAATACGTGGCGGCGTACCGGT
TTTGAGTCGGTCAACGTGAAACGGCAACGCACGCAGACGTTCCGCTAGCGCGTTGGAGGGCGGATCACCGGCGCGGCCGCCGCGGCTTTGATCGAGTCCA
ATGTGAATAACCCCACCCAGGAAGGTGCCGGTCGAGAGCACGACGGATTCTGCGTGGAAGCGAATGCCCGTCTCGGTCACGACGCCACGCACGGTGTCGT
TATCCACAATGAGATCGCCAGCCGCCTGCTGAAAAATCGTCAGGTTGGGTTGGTTTTCCAACATGCCGCGAATGGCGGCTTTGTAGCGGACGCGATCAGC
CTGGGCACGAGTTGCGCGTACCGCGGGCCCTTTGCGTGCATTGAGCACGCGAAACTGGATGCCGCCTAAATCCGTGGCTAGCCCCATTGCACCGCCGAGC
GCATCGATTTCTTTGACCAAATGACTTTTGCCAATGCCACCGATGGCCGGGTTGCACGACATTTGGCCGAGCGTCTCGATGTTGTGGGTCAGCAATAGGG
TTTGACAGCCCATACGAGCAGAGGCCAGTGCGGCTTCGGTTCCCGCATGGCCACCGCCGATGACAATCACGTCAAAGCGGTCGGGGTAATTCAAGAGACA

CCTCGTCTTGCGCCTATTGGCACGGATGATGTGGTCAGCGATTCGCCGCTGATGAAATTAGCCCGCCAGTATAAACCTCCTGTGGGTAACCGTCAGGTTT
TTCCACACCCCGTATGCAACGCAGTGTCGTTATTAATAACAAAATAAAAATATATAAGAGAAGTTCTGTTGATGTTGTAACTATTGGTGTGGACAAAATC
**TGTTGATAA**GTCAATATTCTCTATATTTTT**T**ATAACCTTATATTGTTGACCTTTACGTGATTAAAGGGCGGAGTGCCTGCGTAGAAGCGGTGGCGGAGCT
GTGGATGAAACATCGACTTACCCACAACGCTGCCTGACAACGGGTTATCCACCTATGGATACCGTGCTTGTTACCGTACCTGTGAACAAGCCGGTCGATA
CGGCCGTATCGCTTCAAGAGGTGCCCATGCTGGGGGCTGTGGCGATGTAAGAGTGTGCTTGTAAAAAATAGTATCCACAGGCAAAAAAAATGGCCTGTCA

GCAAGACAGGCCATTGGGTGGGTCGCTTTTTGTGCTTTGGTCGCCTTTAACAGAGCGGAGATCAAGTACGTATCAGTGCTTGAGTACGCGCGCCAAAAAC
GATTGGGTGCGCTCGTGTTGAGGCGCATCAAAGAGTTGCTCGGGTGGGCCTTGTTCGGCAATTTCGCCCTTATGGATAAAGATGATGCGGTCGGCTACCT
CACGGGCAAAGCCCATTTCGTGGGTCACGATCACCATGGTCATGCCCTCCTTAGCTAGCTCTCGCATGGCATCCAGCACTTCACCAATCATCTCGGGATC
GAGCGCCGACGTTGGCTCGTCGAACAGCATGAGGCGCGGCTCCATCGCCAACGCACGCGCCAGCGCCACACGCTGCTGCTGGCCGCCGGAGAGCTGGCTG
GGATACTTGTTGGCCTGATCGGCAATGCCGACGCGCTCCAGCAGCCGCTTGGCGGTCTCTTCGGCGTCTTGGCGGCTCCAGCCGCGGACCTTCATGGGTG
CCAGCGTAATGTTGTCGAGCACGCTCAGATGGGGAAAGAGATTGAACTGCTGGAACACCATGCCCACTTCGGTACGGATGGTTTGCAGCGCCTGGCTGCT
TTTACCGTTGGGCAACAGCGTGTTGCCGTCGACCTCTAAACTACCTGACTGAAACTCTTCCAGACCGTTGATACAGCGGATCAACGTGGACTTGCCGGAA
CCACTGGCTCCGATGACCACGACGACCTCACCCGGGACGATGTCCAGGTCGATATCCTTGAGAACGTGCAGACTGCCAAAGTGCTTGTGGAGCTTTTGCA
TACGCACAATGGGCGTTTGAGTTGAAGTCATCGTGCGCTCCTTATTGTCCGGCGAGGCCACGGCGCTCTAGTTGACGCAGCACCAGGGAAAGGGTCCAGG
TAATGGCGAGATACATCAGCGCCACCATGAAGTAGACCTCCAACGCGGTGAAGGTGGTCGCGATATAGATTTGCCCTTGGCGTACTAGCTCACTCACCCC

>Halomonas piezotolerans NBT06E8 [NZ_CP048602.1:1471..1530], dnaA> ig dnaN>, 60 bp

< page 1 | #067 | Halomonas lutescens CGMCC 1.15122

WebSkew GCmin pos. 78523 (GCmin<>oriC1 23.0 kb; GCmin<>oriC2 1.1 kb; oriC2<>oriC1 21.9 kb)

>Halomonas lutescens CGMCC 1.15122 [NZ_BMHM01000004.1:55244..55743], <rpmH oriC1 dnaA>, 534 bp

TCCAGGACATCAGTGGTCACCGCAATAAAGTCGCGGCTGGCAGTATCGCTTTCAGCATCACCCACGCCTTCGCTGATGGCGCTTTGGGGTGCGGAAGTGC
TAGGTACGGAGAGATCGTCGTTAGCGTTCTCGCTGGCGGAGGCCCCATTACTGCTTTGGGTAATTTCCGGCGTCGAGTCGTAAGTGGTCTGCCCGTAATC
CTGATTCCACTGTACAACCAAAAGATAGGCAAGCAATGCCAGTGGAATCAGTAATAAAAGTCGTTTTACGTCCATGAGGGTTCTGCCCGATGGGTGGTGA
ACATGCTAATGGCGCAGTGTCATTTACTCGACCCATGACACACGAAAGCCTGCCAAGCGGCAGGCCAAGGTCGAGCGGCGAGGTGTGTCGTCAAGCGTCA
CGTGTTACGCCATTAGGCGGGGGAGGTGCAGACACCGCCTGTGCGTCGCGCTGAAGTCGTTTCCACATGCCGTGTAGCTGGCGAGTAAGCGTTTCGTTGT
CGATATCCTGCACGCCTCGTTTTGCCAATATCACGATATCCACAGAGGGTAAGTGATCTTGACGAAGACGCACGGATTCTCGAACGAGTCGCTTGAAGCG
GTTGCGATCCACAGCGAGCTTCACATTTTTTTTGCTGACCACCAGTCCAACACGGGGATACCCCCGGGTACTCATGCGCGCTAACGCCATCATACCTTTG
CCATGAACTTTGAGGTCTGCCTGTTCAAACACGTGACTAAAATCCCCGGCATTTAGCAAACGTAAGCACCGGGGAAAGCCTTGATGCGACACACGCAATC
CGAGATCAGGCGCTCAGACGCTTGCGGCCTTTGGCGCGGCGGCGTGCGATAACGGCACGGCCGTTTTTGGTTGCCATACGAGCACGGAAGCCATGCGCGC
GCTTGCGCTTGAGAACGCTGGGTTGAAAAGTGCGTTTCATAACTCGTTATTCCCACGTGGTTTAGGACGGAAAGTGAATTCCAAAAGACCCGGTCCAGTG

AGGAACTAGCCGGGGAGTTTCAATTCAAGACCGGAAATTGTAGAGACTTCCCGCGCCGGGTGCAATTTTTCCTTGCATAAAAAGCGTGAAAAAGGCAATA
GCGCTGCGGCGTCGGCGTACTGCTTGGGTACTTTGTATCAGACTATTATTACTAATGAAGTTATTTAAAACTTACTGTTATTAATAGTGGGGATACTAAT
**TGTGGATAA**GCTATTTTATGATAATTTTATCAACAGCTTAAAATAATAAAAAATCTGTGCATAAGTCCCGAATAAGCGATAAATAAACTGTGAATTATGT
GTGTGCAACGTGGCACAACTTGGTCGGAAATGAGGTTGTGCCCACTCCGCTACCGGGTTTTCCTTAGCCCTCACATATAGTTGTCCACAGGCAAAAAAGA
GCCGTTAGCGCGTGTGGATAACCGCTTCGTTGGGCGCTACAATGGTCGGCCGTTTTCAACCGATGGTGAGATGAGTGTCGCTCGCGCTTTGGCAACAGTG

CCTGGACTACCTGCAGGACGAACTGAATTCACAGCAGTTCAATACCTGGATACGTCCGCTGCAGGCAGAAGAAGGAGACACCAATGAGCTACGCTTATTG
GCGCCGAATCGCTTTGTGCGTGACTGGGTGAGTGATAAGTATGCAAAACGCATTAGTGAATTGATGCGCGAACTTGCCCCTTCTAAGCCTCCCAAAGTCA
GTTTGACGGTGGGCAGCCGCCGGGCCGCAGCCCCTGCACCGCAGCCCAGAGACCTTGGCAATCCAGTCTCGGCAGCCCCTGCACAGCCGGCATCACCTGC
TGTGCATACGCCACCGCGAGGTGATATTGGCGATGAGCGTGAAATCGACCAACTTCGTGAAGAGAGTGGCCCTAGCCGCCGTGGTAACGAACGCCAAGTG
CAAGTAGAAGGCAGTTTGAAGCACAGCAGTGGCCTAAATCCTAATTTTACCTTCGACACTTTTGTAGAAGGTAAGTCTAACCAGCTGGCACGCGCGGCTT
CTCGTCAGGTCTCCGAGAATCCTGGTGGTGCTTATAACCCTCTGTTTTTATACGGTGGTGTGGGTTTAGGTAAAACGCACTTAATGCATGCGGTGGGTAA
TGCACTGGCTGGCCGACGTGAAAATGCTCGTGTGGTTTACCTGCATTCTGAGCGCTTTGTCGCCGACATGGTGAAGGCGCTTCAGCTTAATGCGATTAAT
GACTTTAAGCGTTTTTACCGCAGCGTCGATGCCCTGCTGATTGATGACATTCAATTTTTTGCTGGTAAAGAGCGTTCTCAAGAAGAGTTTTTTCATACCT
TTAATGCGCTGTTAGAGGGCGGTCAGCAGATGATCCTCACCTCTGACCGCTATCCTAAAGAGATCAGTGGCGTTGAAGAGCGGCTAAAGTCGCGCTTTGG
CTGGGGGCTGACCGTGGCGATCGAGCCGCCAGAGCTTGAAACTCGAGTGGCGATTCTGATGAAGAAAGCCGACCAAGCTAAAGTAGATTTACCCCATGAT

>Halomonas lutescens CGMCC 1.15122 [NZ_BMHM01000004.1:77160..76661], <mnmG oriC2 <COG1126 (glnQ), 576 bp

CTAACCCTTTGATCGAGCGCACCACCTGCAGCTGCACATCAAACGGCAGCGAAGTCGAGATGCCGTTGGGATAGAGCTCGTGGGTATCCAGGCCTTCTGG
CTCGATGAACACCTGATGGCTCGATTTATCGGCAAAGCGGTGCACTTTGTCTTCAATCGACGGGCAGTAGCGCGGGCCGATGCCTTCAATGACGCCAGAG
TACATCGGCGAGCGATCCAAATTACCCAGAATAATCTCGTGGGTGCGCTCGTTGGTGTGCGCGATATGGCAGCTCACCTGCTGGGGGTGCATTTCCCGCG
AGCCAAGGTAAGACATTACCGGCGTTGGCGTATCGCCTGGCTGCTCTTCCAGCTGCGAAAAATCGACGGTTTTGGCATCGATACGCGGCGGCGTGCCGGT
TTTGAGTCGGTCAACGTGAAACGGCAACGCACGCAGTCGTTCGGCTAGCGCGTTGGAGGGCGGATCACCCGCGCGGCCTCCTCGGCTTTGATCCAACCCA
ATGTGGATAACACCGCCCAGGAAGGTACCCGTTGAGAGCACGACAGATTCCGCATGAAAGCGAATGCCGGTTTCGGTCACGACACCGCGCACGGTGTGGT
TATCCACAATGAGATCACCTGCCGCCTGCTGAAAAATCGTCAGGTTCGGCTGGTTTTCCAACATACCGCGTATGGCGGCTTTGTAGCGAATGCGGTCGGC
CTGGGCACGGGTTGCGCGTACCGCTGGCCCTTTACGTGCGTTGAGCACACGAAACTGGATGCCACCTAAGTCCGTCGCTAACCCCATTGCACCGCCGAGC
GCATCAATTTCTTTCACCAAATGGCTCTTGCCGATGCCGCCAATGGCGGGATTGCACGACATTTGGCCAAGCGTCTCGATGTTGTGAGTTAGCAATAGGG
TTTGACAGCCCATACGAGCAGAGGCCAGTGCGGCTTCAGTTCCCGCATGGCCACCGCCGATGACAATCACGTCAAAGCGGTCGGGGTAGTTCAAGAGACA

CCTCGTCTTGTGCCTGTTGGCACGGATGATGTGGTCAGCATGTCAGCGCTGACGGAATAAGCCCGCCAGTATAAACCTCCTGTGGGTAACCGTCAGGTTT
TTCCACACCCCAACTGAGGGGTTCTGTCTTTATTAATAACAAAATAAAAATAAATAAGAGAAGTTCTGTTGATGTAGTAACTACTGGTGTGGACAAAATC
**TGTTGATAA**GTCTATTAACGTTATTTTTATCAACATTTTACATTGTTGACCTTTAGGTGATTAAGCAACGGAGTGCCTGCGTAGAAGCGGTGTTGAAGCT
GTGGATGAAAATACGACTTACCCACAGCGCTGGCTAGCAACGGGTTATCCACCGCTGGATACCTGGCTTGTTACCGCACCTGTGAACAACCTCCATAGGA
GCGCGTTGACACGTTTAAAAGCGCTCATGGCGGGGGGTGTAGCGGAGGTGAGAGAACGTTGTAAAAAATAGTATCCACAGGCAAAAAAAATGGCCTGTCA

TGAAGACAGGCCATTTTAGGTGGCGTTATTGGCGTTAAACGCGCCGTGTGAAACCGTTATTTAGCGTGGTTTAGTGCTTCAATACCCGTGCCAAAAACGA
TTGGGTGCGTTCGTGTTGAGGGGCATCAAACAGCTGCTCGGGTGGCCCTTGTTCGGCAATTTCGCCTTTATGGATAAAGATGATGCGGTCAGCCACTTCT
CGGGCAAAGCCCATCTCGTGGGTAACAATCACCATGGTCATGCCCTCTTTGGCAAGCTCCCGCATGGCGTCTAGCACTTCACCAATCATCTCGGGGTCAA
GGGCTGACGTTGGCTCGTCAAACAGCATGAGGCGCGGCTCCATGGCCAGCGCACGGGCCAGCGCCACACGCTGCTGCTGGCCGCCGGAGAGCTGGCTTGG
GTACTTATTTGCTTGGTCGGCAATGCCTACGCGATCAAGCAGTCGCTTAGCGGTCTCTTCGGCATCCTGGCGGCTCCAGCCACGCACTTTCATCGGGGCG
AGAGTGACGTTATCGAGCACGCTAAGATGGGGGAACAGATTGAACTGCTGAAACACCATGCCCACTTCGGTGCGGATGGTTTGCAGCGCTTTGGTGCTCT
TGCCCAGCGGTAGTAGCTCATTGTTGTCGACCACCAATGTGCCTGACTGGAACTCTTCCAACCCATTAATGCAGCGAATTAGGGTGGATTTCCCCGAGCC
GCTGGCACCGATAATCACCACCACTTCGCCGGGGATAATTTCTAAATCAATGCCGTTAAGTACATGAAGACTGCCAAAGTGCTTATTCAGCTTTTGCATA
CGCACAATGGGTTGCGCTGTGTCGTTCTTGATCGCGTCGTTCTTGATTGCATCGTTCATTGCGCGTGTCCTTATTGTCCGACCAGGCCTCTGCGCTCAAG
CTGACGCAGTAAGATTGACAGTGTCCAGGTGATGGCCAGATACATCAGCGCCACCATGAAGTAGACCTCAAGGGCGGTGAACGTGGTGGAGATATAAATT

>Halomonas lutescens CGMCC 1.15122 [NZ_BMHM01000004.1:53740..53799], dnaA> ig dnaN>, 60 bp

WebSkew GCmin pos. 89261 (2nd min.) (GCmin<>oriC1 21.2 kb; GCmin<>oriC2 2.1 kb; oriC2<>oriC1 19.1 kb)

>Halomonas stevensii S18214 [NZ_AJTS01000041.1:110218..110717], <rpmH oriC1 dnaA>, 533 bp

GTCCAGCACATCGGTAGTGACGGCGATGAAGTCACGACTGGCGGTCGAACTTGGCGAGTCGCTCATGCCTTCATCGATGGCGGCCTGAGGTGCGGAAGTA
CTGGGAACGGAAAGATCGTCGCTGCCGCCTTCACTCGCAGGTGCCCCATTGCTTGTCTGGGTCATTTCGGGAGTGGAGTCGTAGCTGGTCTGCCCGTAAT
CCTGATTCCACTGTACAACCAGTAGGTAGGCAAGCACTGCCAGTGGAATCAGTAATAAAAGTCGTTTTACGTCCATGAGGGTTCTGCCCGATGGGTGGTG
AACATGCTAATGGCGCAGTGTCATTTGCTCGGCCCATGACACACGAAAGCCTGCCAAGCGGCAGGCCGAGGTCGAGCGGCGAGGTGTGTCGTCAAGTGTC
ACGTGTCAGGCCATGTGATGGCGGAGATGTCGATACCGCCGGAGCCTCGCGCTGAAGCCGTTTCCACATGCCATGCAATTGGCGAGTCAGCGTCTCGTTA
TCGATCTCCTGAACGCCTCGCTTGGCCATGATCACGATATCCACAGCGGGCAAGCGATCCTGACGAAGGCGAACGGATTCTCTAACCAGGCGCTTGAAGC
GGTTACGATCCACAGCAAGCTTCACATTTTTCTTGCTGACTATCAGACCAATGCGGGGATGTCCCAGGCTGTTCAAGCGCGCTAGCGCCATCATGCCCTT
GCCATGGACTTTCAAGTCGGCTTGTTCGAACACGTGACTGAAATCCCCAGCGTTCAGCAATCGCAAGCACCGGGGAAAGCCTTGAGGCGACACACGCAAT
CCGAGATCAGGCGCTCAGACGCTTGCGGCCTTTGGCACGGCGGCGTGCGATGACGGCACGGCCGTTCTTGGTTGCCATACGAGCACGGAAGCCATGCGCG
CGCTTGCGCTTGAGAACGCTGGGTTGAAAAGTGCGTTTCATAACTCGTTATTCCCACGTGGTTTAGGACGGAAATTGAATTCCAGATGACCCGGTCCAGT

GAGGAACCAGCCGGGGAGTTTCAATTCAAGACCGGAAATTGTAGAGACTTCCTGGCCTCGGTGCAATTTTTCCTTGCATGAAAAGCGTGAAAAGGGCAAT
GGCGACCCGCGATGCCCCGTCAGTCTGACGAACGTCGATGACACTACTATTACTAATGGGTTTACTTAAACCTGTTGTTATTAATAGTGGGGATACTAAT
**TGTGGATAA**CTCATTTATCCCTATTTTTATCAAACTCTTAAAATGATAAAAAATCTGTGGGTAACTGGCTGTAAAGTGGCGTAAAAGCTGTGATTTATGT
GTGATTAACGCAGCACAACTAAGCAGTATGCGGTTTTGTACCCATGCCGTCCGCTGCTTTTCCTAAGCCCTCATGGACATTTGTCCACAGGGGAAAAAGA
GCCCCGAGTGCATGTGGATAACCGTTTCGTTGGGCGCTACAATGGTCGGCCGTTTTCAATCGATGGTGAGATGAGTGTCACTCGCGCTTTGGCAACAGTG

CCTGGACTACCTGCAGGACGAACTGAATTCGCAACAGTTCAATACCTGGATACGTCCGCTGCAGGCAGAAGAAGGAGAGTCCAACGAGCTTCGCTTGTTG
GCGCCGAACCGCTTCGTGCGCGATTGGGTGAGCGATAAATATGCCAAGCGAATCAGTGAACTGATGCGCGAGCTTGCCCCTTCCAAACCTCCCAAGGTCA
GCCTTACCGTGGGTAGTCGTCGCACCTCGGCAGCTGCGCCACAGCCTAGAGAGCTGGGTAATCCCGTCTCCGCATCGCCTGCCCAGGCACCTGCTCCCGC
CGTCCATACTCCGCCCCGCGGTGAAGTGGGCGATGAGCGTGAGATCGAGCAGTTGCGCGAAGAGCCGGGCGCTGGCCGACGCGGCAACGAGCGCCAGGTA
CAGGTAGAAGGTAGCCTGAAGCACAGCAGCGGCCTGAATCCCAACTTCACCTTTGATACATTCGTGGAAGGTAAATCGAACCAGTTGGCGCGTGCGGCCT
CTCGGCAGGTATCCGAAAATCCGGGTGGGGCCTATAACCCGCTGTTCCTCTACGGGGGTGTAGGCCTGGGTAAAACCCACTTGATGCATGCCGTGGGCAA
TGCCTTGGCAGGTAGGCGGGAAAACGCCCGAGTGGTGTATCTGCACTCCGAGCGCTTCGTGGCGGATATGGTGAAGGCGCTTCAGCTCAACGCCATCAAT
GACTTCAAGCGTTTTTATCGCAGTGTCGATGCCTTGTTGATCGATGATATTCAATTCTTTGCTGGCAAGGAGCGCTCCCAGGAGGAGTTTTTCCATACCT
TCAACGCCTTGTTGGAAGGTGGCCAGCAGATGATCCTGACCTCGGATCGCTACCCCAAAGAGATCAGTGGCGTCGAAGAGCGTCTGAAGTCGCGGTTCGG
TTGGGGCTTGACGGTGGCCATCGAGCCGCCTGAGCTGGAGACGCGGGTCGCGATCCTGATGAAGAAAGCCGACCAGGCCAAGGTCGAGTTACCGCATGAT

>Halomonas stevensii S18214 [NZ_AJTS01000041.1:91093..91592], <mnmG oriC2 <COG1126 (glnQ), 542 bp

TCCAACCCTTTGATGGAGCGCACGACCTGCAGCTGCACATCGAAGGGCAGTGAGGTGGAGATCCCGTTGGGGTAGAGCTCGTGGGTATCCAGACCTTCCG
GTTCGATGAACACCTGATGGCTCGATTTGTCGGCGAAGCGATGCACCTTGTCTTCGATCGACGGGCAGTAGCGGGGGCCGATGCCTTCGATGACCCCAGA
GTACATGGGGGAACGATCGAGGTTGTTCAGGATGATCTCGTGGGTGCGCTCGTTGGTGTGCGCAATATGGCAGCTCACTTGCTGAGGATGCATGCCCCTA
TGCCCCAGGTAGGACATGATCGGCGTGGGCGTGTCGCCTGGCTGCTCTTCCAGCTGTGAAAAGTCGACGGATTTGGCATCGATACGCGGAGGGGTACCGG
TCTTCAACCTGTCGACGTGAAAAGGCAGTGCCCGCAACCGCTCGGCCAGCGCATTGGAAGGCGGGTCGCCTGCGCGGCCACCACGGCTTTGATCCAACCC
GATGTGGATAACCCCGCCAAGGAACGTGCCCGTGCTCAGCACTACCGACTCGGCATGAAAACGAATGCCTGTCTCGGTCACTACACCGCGTACGGTGTCG
TTATCCACAATGAGGTCGCCAGCCGCCTGCTGAAAAATCGTCAGGTTCGGCTGGTTTTCCAGCATGCCGCGAATGGCGGCCTTGTAGCGGATCCTGTCTG
CCTGGGCGCGGGTCGCGCGTACGGCAGGCCCCTTACGTGCGTTGAGTACTCGAAACTGGATGCCGCCCAAATCCGTGGCTAGCCCCATGGCACCACCCAA
CGCATCGATCTCCTTGACCAGATGGCTCTTGCCGATCCCACCAATGGCGGGATTGCAGGACATTTGGCCCAGCGTCTCGATGTTGTGTGTCAGCAATAGG
GTTTGACAGCCCATGCGAGCAGAGGCCAGTGCGGCTTCGGTTCCCGCATGGCCACCGCCGATGACAATCACGTCAAAGCGGTCGGGGTAGTTCAAGAGAC

ACCTCGTCTTGTGCCCAAGGCACGGATGATGTGGTCAGCGGATCACCGCTGTTTGAATAAGCCCGCCAGTATAAACCTCCTGTGGGTAACCGTCAGGTTT
TTCCACACCCCTGCTATCAGCGAGTTTATAGATTAATAACAAAATAAAAATATATAAGAGAAGTTCTGTAAATGTAGTAACTAATGGTGTGGACAAAATC
**TGTTGATAA**ATGGTTTTTTCCCATATATATCATGGATTTGTATTGTTGACCAAGCAGTGGCGAACCGGCGGAGTGCCTGCGTAGAAGCGGTGACAGAGCT
GTGGATGAAAAGCTTACTTACCCACAGCCCAGGTTAACGACGGCTTATCCACCGCTGGCTACCGCGCTTGTTACCGCACCTGTGAACAAGTGATCTGGTG
ACCATCATATGCGTTGAAAGATGCCCATGGCGCGGGTGTTAGCCCGATAGTGGCAGCCGAGAAAAAAATATTATCCACAGGCAAAAAAAAGGCCTGTCGC

AAGGACAGGCCGGTCGCGGAGGTACCCTGATGGGCTGCTAGTGCTTCAATACGCGCGCCAAGAAGGACTGGGTGCGTTCATGCTGAGGTGCATCGAACAG
CTGTTCCGGTGGCCCTTGCTCGGTGATTTCCCCTTTGTGAATGAAGATGACGCGGTCGGCTACTTCCCGCGCAAAGCCCATTTCATGGGTCACGATGACC
ATGGTCATGCCTTCCTTGGCCAGCTCACGCATGGCGTCCAGCACTTCACCGATCATCTCCGGGTCTAGCGCCGAGGTGGGCTCGTCGAAGAGCATGAGCC
GGGGTTCCATGGCCAGCGCGCGGGCCAGGGCGACCCGCTGCTGCTGCCCTCCAGAGAGCTGGCTGGGGTACTTCTGTGCCTGATCGGCAATGCCGACGCG
TTCCAACAGCCGCTGAGCCGTCTCTTCGGCATCCTTACGGCTCCAGCCACGCACTTTCATGGGGGCCAGGGTAATGTTGTCGATGACGCTCAGATGGGGG
AAGAGGTTGAACTGCTGGAACACCATGCCGACTTCGGTGCGGATGGTTTGCAGGGCCTTGCTGCTTTTGCCATCGGGCAGCAGCGTATTGCCATCGACAT
CGAGGCTGCCGGACTGAAACTCTTCCAGCCCGTTGATGCAGCGGATCAGCGTGGATTTACCAGAGCCGCTGGCGCCGATTACGACGACGACTTCCCCAGG
GACGATTTCCAGATCGATGTTCTTGAGCACATGCAGGCTGCCAAAGTGCTTGTTGAGCTTCTGCATTTGCACAATGGGCGATGAGGTCGAAGTCATTGTG
CGCTCCTTATTGGCCAACGACGCCGCGCCGTTCCAACTGGCGCAGGATGATAGAGAGGGTCCAGGTAATGGCCAGGTACATCAGGGCGACCATGAAATAG
ACTTCCAGCGCCGTGAAGGTGGTGGCAATGTAGATCTGTCCTTGGCGCACCAGCTCACCCACGCCAATGACGGAAAACAGCGAGGTGTCCTTGATGCTGA

>Halomonas stevensii S18214 [NZ_AJTS01000041.1:112216..112162], dnaA> ig dnaN>, 55 bp

WebSkew GCmin pos. 64400 (GCmin<>oriC1 22.4 kb; GCmin<>oriC2 1.1 kb; oriC2<>oriC1 21.3 kb)

>Halomonas johnsoniae KCTC 22157 [NZ_BMXO01000019.1:c41770..42269], <rpmH oriC1 dnaA>, 533 bp

GTCCAGCACATCGGTAGTGACGGCGATGAAGTCACGACTGGCGGTCGAACTTGGCGAGTCGCTCATTCCTTCATCGATGGCGGCCTGAGGTGCGGAAGTA
CTGGGAACGGAAAGATCGTTGCTGCCGCCTTCACTCGCCGGTGCCCCATTGCTTGTCTGGGTCATTTCGGGAGTGGAGTCGTAGCTGGTCTGCCCGTAAT
CCTGATTCCACTGTACAACCAGTAGGTAGGCAAGCACTGCCAGTGGAATCAGTAATAAAAGTCGTTTTACGTCCATGAGGGTTCTGCCCGATGGGTGGTG
AACATGCTAATGGCGCAGTGTCATTTGCTCGGCCCATGACACACGAAAGCCTGCCAAGCGGCAGGCCGAGGTCGAGCGGCGAGGTGTGTCGTCAAGTGTC
ACGTGTCAGGCCATGTGATGGCGGAGATGTCGATACCGCCGGAACCTCGCGCTGAAGTCGTTTCCACATGCCATGCAATTGGCGAGTCAGCGTCTCGTTG
TCGATCTCCTGAACGCCCCGCTTGGCCATGATCACGATATCCACAGCGGGCAAGCGATCCTGACGAAGGCGAACGGATTCTCTAACCAGGCGCTTGAAGC
GGTTACGTTCCACAGCAAGCTTCACATTTTTCTTGCTGACTATCAGGCCAATGCGGGGGTGCCCCAGAGTATTCAAGCGCGCTAGCGCCATCATGCCCTT
GCCATGGACTTTCAAGTCGGCTTGTTCGAACACGTGACTGAAATCCCCAGCGTTCAGCAATCGCAAGCACCGGGGAAAGCCTTGAGGCGACACACGCAAT
CCGAGATCAGGCGCTCAGACGCTTGCGGCCTTTGGCACGGCGGCGTGCGATGACGGCACGGCCGTTCTTGGTTGCCATACGAGCACGGAAGCCATGCGCG
CGCTTGCGCTTGAGAACGCTGGGTTGAAAAGTGCGTTTCATAACTCGTTATTCCCACGTGGTTTAGGACGGAAATTGAATTCCAGATGACCCGGTCCAGT

GAGGAACCAGCCGGGGAGTTTCAATTCAAGACCGGAAATTGTAGAGACTTCCTGGCCCCGGTGCAATTTTTCCTTGCATGAAAAGCGTGAAAAGGGCAAT
GGCAGCCCACGATGCCCTGTCAGTCTGACGAACGTCGATGACACTACTATTACTAATAGGTTTATTTAAACCTGTTGTTATTAATAGTGGGGATACTAAT
**TGTGGATAA**CTCATTTATCCCTATCTTTATCAAAACCTTAAAATGATAAAAAATCTGTGGGTAACTGGCTGTGAAGTAGCGTAAAAGCTGTGATTTATGT
GTGATTAACGCAGCACAACTCAGCAATTTGCGGTTTTGTACCCATGCCGTCCGCTGTTTTTCCTAAGCTCTCATGGTCATTTGTCCACAGGCGGAAAAGA
GCCCCGAGTGCGTGTGGATAACCGTTTCGTTGGGCGCTACAATGGTCGGCCGTTTTCAACCGATGGTGAGATGAGTGTCACTCGCGCTTTGGCAACAGTG

CCTGGACTACCTGCAGGACGAACTGAATTCGCAACAGTTCAATACCTGGATACGTCCACTGCAGGCAGAAGAAGGAGAGTCCAACGAGCTTCGCTTGTTG
GCGCCGAACCGCTTTGTGCGCGATTGGGTGAGCGATAAATACGCGAAGCGAATCAGTGAACTGATGCGCGAGCTTGCTCCTTCCAAACCTCCCAAGGTCA
GCTTGACCGTGGGTAGTCGTCGCACCTCGGCACCTGCGCCACAGCCTAGAGAGCTGGGTAATCCCGTCTCCGCATCGCCTGCTCAGGCGCCTGCTCCCGC
CGTCCATACTCCGCCCCCGCGGTGAAGTGGGCGATGACCGTGAAATCGAGCAGTTGCGCGAAGAGCCGGGCGCCGGCCGACGCGGCAACGAGCGCCAGGTA
CAGGTAGAAGGTAGCCTGAAGCACAGCAGCGGCCTGAATCCCAACTTCACCTTTGATACCTTCGTGGAAGGTAAATCGAACCAGTTGGCGCGTGCGGCCT
CTCGGCAGGTATCCGAAAATCCGGGTGGGGCCTATAACCCGCTGTTCCTCTACGGCGGGGTAGGCCTGGGTAAAACCCACTTGATGCATGCCGTGGGCAA
TGCCCTGGCAGGTAGGCGGGAAAACGCCCGAGTGGTGTATCTGCACTCCGAGCGCTTCGTGGCGGATATGGTGAAGGCGCTTCAGCTCAACGCCATCAAT
GACTTCAAGCGTTTTTATCGCAGTGTCGACGCCTTGTTGATCGATGATATTCAATTCTTTGCTGGCAAGGAGCGCTCCCAGGAGGAGTTTTTCCATACCT
TCAACGCCCTGTTGGAAGGTGGCCAGCAGATGATCCTGACCTCGGATCGCTACCCCAAAGAGATCAGTGGCGTCGAAGAACGTTTGAAGTCGCGGTTCGG
TTGGGGCTTGACGGTAGCCATCGAGCCGCCTGAGCTGGAGACGCGGGTCGCGATCCTGATGAAGAAAGCCGACCAGGCCAAGGTCGAGTTACCGCATGAT

>Halomonas johnsoniae KCTC 22157 [NZ_BMXO01000019.1:c63041..63540], <mnmG oriC2 <COG1126 (glnQ), 542 bp

TCCAACCCTTTGATGGAGCGCACGACCTGCAGCTGCACATCGAAGGGCAGTGAGGTGGAGATCCCGTTGGGGTAGAGCTCGTGGGTATCCAGGCCTTCCG
GTTCGATGAACACCTGATGGCTCGATTTGTCGGCGAAGCGATGCACCTTGTCTTCGATCGACGGGCAGTAGCGGGGGCCGATGCCTTCGATTACCCCAGA
GTACATGGGGGAACGATCGAGGTTGTTCAGGATGATCTCGTGGGTGCGCTCGTTGGTGTGCGCAATATGGCAGCTCACCTGCTGAGGATGCATGCCCCGA
TGCCCCAGGTAGGACATGATCGGCGTGGGCGTGTCGCCTGGCTGCTCTTCCAGCTGTGAAAAGTCGACGGATTTGGCATCGATACGCGGAGGGGTACCGG
TCTTCAACCTGTCGACGTGAAAAGGCAGTGCCCGCAGTCGCTCAGCCAGCGCATTGGAAGGCGGGTCGCCTGCGCGGCCACCACGGCTTTGATCCAACCC
GATGTGGATAACCCCGCCAAGGAACGTGCCCGTGCTCAGCACCACCGACTCGGCATGAAAACGAATGCCTGTCTCGGTCACTACACCGCGTACGGTGTCG
TTATCCACAATGAGGTCGCCAGCCGCCTGCTGAAAAATCGTCAGGTTCGGCTGGTTTTCCAGCATGCCGCGAATGGCGGCCTTGTAGCGGATTCTGTCTG
CCTGGGCGCGGGTCGCGCGTACGGCAGGCCCCTTACGTGCGTTGAGTACTCGAAACTGGATGCCGCCCAAATCCGTGGCTAGCCCCATGGCACCACCCAA
CGCATCGATCTCCTTGACCAGATGGCTCTTGCCGATCCCACCAATGGCGGGATTGCAGGACATTTGGCCCAACGTCTCGATGTTGTGGGTCAGCAATAGG
GTTTGACAGCCCATGCGAGCAGAGGCCAATGCGGCTTCGGTTCCCGCATGGCCACCGCCGATGACAATCACGTCAAAGCGGTCGGGGTAGTTCAAGAGAC

ACCTCGTCTTGTGCCCAAGGCACGGATGATGTGGTCAGCGGATCACCGCTGTTGGAATAAGCCCGCCAGTATAAACCTCCTGTGGGTAACCGTCAGGTTT
TTCCACACCCCTGCTATCAGTGGTTTTATAAATTAATAACAAAATAAAAATATATAAGAGAAGTTCTGTAAATGTAGTTACTAATGGTGTGGACAAAATC
**TGTTGATAA**ATGGTTTTTTCCTATATGAATCATTATTTTGTATTGTTGACCAAGCAGTGGCGAACCGGCGGAGTGCCTGCGTAGAAGCGGTGACGGAGCT
GTGGATGAAAAGCTTACTTACCCACAGGCCAGGTTAACGACGGCTTATCCACCGCTGGCTACCGCGCTTGTTACCGCACCTGTGAACAAGTGATCTGGTG
ACCATCATATGCGTTGAAAGATGCTCATGGTGCGAGTGTTAGCCAGATAGGGGCAGCCGAGGAAAAAATATTATCCACAGGCAAAAAAAAGGCCTGTCGC

AAGGACAGGCCGGTCACTGATGTGCCCTGATGGGCTGCTAGTGTTTCAATACGCGCGCCAAAAAGGACTGGGTGCGTTCGTGCCGAGGTGCATCGAACAG
CTGTTCCGGTGACCCTTGCTCGGTGATTTCCCCTTTGTGAATGAAGATGACGCGGTCGGCCACTTCCCGTGCAAAGCCCATTTCATGGGTCACGATGACC
ATGGTCATGCCTTCCTTGGCCAGCTCACGCATGGCGTCCAGCACTTCACCGATCATCTCCGGGTCTAGCGCCGAGGTGGGCTCGTCGAAGAGCATGAGCC
GGGGTTCCATGGCCAGCGCGCGGGCCAGGGCGACCCGCTGCTGCTGCCCTCCAGAGAGCTGGCTGGGGTACTTCTGTGCCTGATCGGCAATGCCGACGCG
TTCCAACAGCCGCTGAGCCGTCTCTTCGGCATCCTTACGGCTCCAGCCACGCACTTTCATGGGGGCCAGGGTAATGTTGTCGATGACGCTCAGATGGGGG
AACAGATTGAACTGCTGGAACACCATGCCGACTTCGGTGCGGATGGTTTGCAGGGCCTTGCTGCTTTTGCCATCGGGCAGCAGCGTATTGCCATCGACAT
CGAGGCTGCCGGACTGAAACTCTTCCAGCCCGTTGATGCAGCGGATCAGCGTGGATTTGCCAGAACCGCTGGCGCCGATCACGACGACGACTTCTCCAGG
GACGATTTCCAGATCGATGTTCTTGAGCACATGCAGGCTGCCAAAGTGCTTGTTGAGCTTCTGCATTTGCACAATGGGCGATGAGGTCGAAGTCATTGTG
CGCTCCTTATTGGCCAACGACGCCGCGCCGTTCCAACTGGCGCAGGATGATAGAGAGGGTCCAGGTAATGGCCAGGTACATCAGGGCGACCATGAAATAG
ACTTCCAGCGCCGTGAAGGTGGTGGCAATGTAGATCTGTCCTTGGCGCACCAGCTCACCCACCCCGATGACGGAAAACAGCGAGGTGTCCTTGATGCTGA

>Halomonas johnsoniae KCTC 22157 [NZ_BMXO01000019.1:c40271..40326], dnaA> ig dnaN>, 56 bp

WebSkew GCmin pos. 78921 (GCmin<>oriC1 23.4 kb; GCmin<>oriC2 1.6 kb; oriC2<>oriC1 21.8 kb)

>Halomonas hamiltonii KCTC 22154 [NZ_BMXN01000011.1:55252..55751], <rpmH oriC1 dnaA>, 533 bp

GTCCAGCACATCGGTAGTGACGGCGATGAAGTCACGACTGGCGGTCGAACTTGGCGAGTCGCTCATGCCTTCATCGATGGCGGCCTGAGGTGCGGAAGTA
CTGGGAACGGAAAGATCGTCGCTGCCGCCTTCACTCGCAGGTGCCCCATTGCTTGTCTGGGTCATTTCGGGAGTGGAGTCGTAGCTGGTCTGCCCGTAAT
CCTGATTCCACTGTACAACCAGTAGGTAGGCAAGCACTGCCAGTGGAATCAGTAATAAAAGTCGTTTTACGTCCATGAGGGTTCTGCCCGATGGGTGGTG
AACATGCTAATGGCGCAGTGTCATTTGCTCGGCCCATGACACACGAAAGCCTGCCAAGCGGCAGGCCGAGGTCGAGCGGCGAGGTGTGTCGTCAAGTGTC
ACGTGTCAGGCCATGTGATGGCGGAGATGTCGATACCGCCGGAGCCTCGCGCTGGAGTCGTTTCCACATGCCATGCAATTGGCGAGTCAGCGTCTCGTTG
TCGATCTCCTGAACGCCTCGCTTGGCCATGATCACGATATCCACAGCGGGCAAGCGATCCTGACGAAGGCGAACGGATTCTCTAACCAGGCGCTTGAAGC
GGTTACGTTCCACAGCAAGCTTCACATTTTTCTTGCTGACTATCAGGCCAATGCGGGGGTGCCCCAGAGTATTCAAGCGCGCTAGCGCCATCATGCCCTT
GCCATGGACTTTCAAGTCGGCTTGTTCGAACACGTGACTGAAATCCCCAGCGTTCAGCAATCGCAAGCACCGGGGAAAGCCTTGAGGCGACACACGCAAT
CCGAGATCAGGCGCTCAGACGCTTGCGGCCTTTGGCACGGCGGCGTGCGATGACGGCACGGCCGTTCTTGGTTGCCATACGAGCACGGAAGCCATGCGCG
CGCTTGCGCTTGAGAACGCTGGGTTGAAAAGTGCGTTTCATAACTCGTTATTCCCACGTGGTTTAGGACGGAAATTGAATTCCAGATGACCCGGTCCAGT

GAGGAACCAGCCGGGGAGTTTCAATTCAAGACCGGAAATTGTAGAGACTTCCTGGCCTCGGTGCAATTTTTCCTTGCATGAAAAGCGTGAAAAGGGCAAT
GGCGGCCCACGATGCCCTGTCAGTCTGACGAACGTCGATGACACTACTATTACTAATGGGTTTACTTAAACCTGTTGTTATTAATAGTGGGGATACTAAT
**TGTGGATAA**CTCATTTATCCCTATTTTTATCAAAACCTTAAAATGATAAAAAATCTGTGGGTAACTGACTGTGAAGCAGCGTAAAAGCTGTGATTTATGT
GTGATTAACGCAGCACAACTCAGCAATTTGCGGTTTTTTACCCATGCCGTCCGCTGTTTTTCCTAAGCTCTCATGGTCATTTGTCCACAGGCGGAAAAGA
GCCCCGAGTGCGTGTGGATAACCGTTTCGTTGGGCGCTACAATGGTCGGCCGTTTTCAACCGATGGTGAGATGAGTGTCACTCGCGCTTTGGCAACAGTG

CCTGGACTACCTGCAGGACGAACTGAATTCGCAACAGTTCAATACCTGGATACGTCCACTGCAGGCAGAAGAAGGAGAGTCCAACGAGCTTCGCTTGTTG
GCGCCGAACCGCTTTGTGCGCGATTGGGTGAGCGATAAATACGCGAAGCGAATCAGTGAACTGATGCGCGAGCTTGCTCCTTCCAAACCTCCCAAGGTCA
GCTTGACCGTGGGTAGTCGTCGCACCTCGGCACCTGCGCCACAGCCTAGAGAGCTGGGTAATCCCGTCTCCGCATCGCCTGCTCAGGCGCCTGCTCCCGC
CGTCCATACTCCGCCCCGCGGTGAAGTGGGCGATGAGCGTGAAATCGAGCAGTTGCGCGAAGAGTCGGGCGCCGGCCGACGCGGCAACGAGCGCCAGGTA
CAGGTAGAAGGTAGCCTGAAGCACAGCAGCGGCCTGAATCCCAACTTCACCTTTGATACCTTCGTGGAAGGTAAATCGAACCAGTTGGCGCGTGCGGCCT
CTCGGCAGGTATCCGAAAATCCGGGTGGGGCCTATAACCCGCTGTTCCTCTACGGCGGGGTAGGCCTGGGTAAAACCCACTTGATGCATGCCGTGGGCAA
TGCCCTGGCAGGTAGGCGGGAAAACGCCCGAGTGGTGTATCTGCACTCCGAGCGCTTCGTGGCGGATATGGTGAAGGCGCTTCAGCTCAACGCCATCAAT
GACTTCAAGCGTTTTTATCGCAGTGTCGACGCCTTGTTGATCGATGATATTCAATTCTTTGCTGGCAAGGAGCGCTCCCAGGAGGAGTTTTTCCATACCT
TCAACGCCCTGTTGGAAGGTGGCCAGCAGATGATCCTGACCTCGGATCGCTACCCCAAAGAGATCAGTGGCGTCGAAGAACGTTTGAAGTCGCGGTTCGG
TTGGGGCTTGACGGTAGCCATCGAGCCGCCTGAGCTGGAGACGCGGGTCGCGATCCTGATGAAGAAAGCCGACCAGGCCAAGGTCGAGTTACCGCATGAT

>Halomonas hamiltonii KCTC 22154 [NZ_BMXN01000011.1:77026..76527], <mnmG oriC2 <COG1126 (glnQ), 542 bp

TCCAACCCTTTGATGGAGCGCACGACCTGCAGCTGCACATCGAAGGGCAGTGAGGTGGAGATCCCATTGGGGTAGAGCTCGTGGGTATCCAGGCCTTCCG
GTTCGATGAACACCTGATGGCTCGATTTGTCGGCGAAGCGATGCACCTTGTCTTCGATCGACGGGCAGTAGCGGGGGCCGATGCCTTCGATTACCCCAGA
GTACATGGGGGAACGATCGAGGTTGTTCAGGATGATCTCGTGGGTGCGCTCGTTGGTGTGCGCAATATGGCAGCTCACCTGCTGAGGATGCATGCCCCGA
TGCCCCAAGTAGGACATGATCGGCGTGGGCGTGTCGCCTGGCTGCTCTTCCAGCTGTGAAAAGTCGACGGATTTGGCATCGATACGCGGAGGGGTACCGG
TCTTCAACCTGTCGACGTGAAAAGGCAGTGCCCGCAGTCGCTCAGCCAGCGCATTGGAAGGCGGGTCGCCTGCGCGGCCACCACGGCTTTGGTCCAACCC
GATGTGGATAACCCCGCCAAGGAACGTGCCCGTGCTCAGCACCACCGACTCGGCATGAAAACGAATGCCTGTCTCGGTCACTACACCACGTACGGTGTCG
TTATCCACAATGAGGTCGCCAGCCGCCTGCTGGAAAATCGTCAGGTTTGGCTGGTTTTCCAACATGCCGCGAATGGCGGCCTTGTAGCGGATCCTGTCTG
CCTGGGCGCGGGTCGCGCGTACGGCAGGCCCCTTACGTGCGTTGAGTACTCGAAACTGGATGCCGCCCAAATCCGTGGCTAGCCCCATGGCACCACCCAA
CGCATCGATCTCCTTGACCAGATGGCTCTTGCCGATCCCACCAATGGCGGGATTGCAAGACATTTGGCCCAGCGTCTCGATGTTGTGGGTCAGCAATAGG
GTTTGACAGCCCATGCGAGCAGAGGCCAATGCGGCTTCGGTTCCCGCATGGCCACCGCCGATGACAATCACGTCAAAGCGGTCGGGGTAGTTCAAGAGAC

ACCTCGTCTTGTGCCCAAGGCACGGATGATGTGGTCAGCGGATCACCGCTGTTGGAATAAGCCCGCCAGTATAAACCTCCTGTGGGTAACCGTCAGGTTT
TTCCACACCCCTGCTATCAGTGGTTTTATAAATTAATAACAAAATAAAAATATATAAGAGAAGTTCTGTAAATGTAGTTACTAATGGTGTGGACAAAATC
**TGTTGATAA**ATGATTTTTTCCTATATGAATCATTATTTTGTATTGTTGACCAAGTAGTGGCGAACCGGCGGAGTGCCTGCGTAGAAGCGGTGACGGAGCT
GTGGATGAAAAGCTTACTTACCCACAGGCCAGGTTAACGACGGCTTATCCACCGCTGGCTACCGCGCTTGTTACCGCACCTGTGAACAAGTGATCTGGTG
ACCATCATATGCGTTGAAAGATGCCCATGGCGCGGGTGTTAGCCAGATAGTGACCGCCGAGGAAAAAATATTATCCACAGGCAAAAAAAAGGCCTGTCGC

AAGGACAGGCCGGTCTCTGAGGTGCCCTGATGGGCTGCTAGTGTTTCAATACGCGCGCCAAGAAGGACTGGGTGCGTTCGTGCCGAGGTGCGTCGAACAG
CTGTTCCGGTGGCCCTTGCTCGGTGATTTCCCCTTTGTGAATGAAGATGACGCGGTCGGCTACTTCCCGTGCAAAGCCCATTTCATGGGTCACGATAACC
ATGGTCATGCCTTCCTTGGCCAGCTCACGCATGGCGTCCAGCACTTCACCGATCATCTCCGGGTCTAGCGCCGAGGTGGGCTCGTCGAAGAGCATGAGCC
GGGGTTCCATGGCCAGCGCACGCGCCAGTGCAACCCGCTGCTGCTGCCCTCCAGAGAGCTGGCTGGGGTACTTCTGTGCCTGATCGGCAATGCCGACGCG
TTCCAACAGCCGCTGAGCCGTCTCTTCGGCATCCGAACGGCTCCAGCCACGCACTTTCATGGGGGCCAGGGTAATGTTGTCGATGACGCTCAGATGAGGG
AAGAGGTTGAACTGCTGGAACACCATGCCGACTTCGGTGCGGATGGTTTGCAGGGCCTTGCTGCTTTTGCCATCCGGCAGCAGCGTATTGCCATCGACAT
CGAGGCTGCCGGACTGAAACTCTTCCAGCCCGTTGATGCAGCGGATCAGCGTGGATTTACCGGAGCCGCTGGCGCCGATCACGACGACGACTTCTCCGGG
GACGATTTCCAGATCGATGTTCTTGAGCACATGCAGGCTGCCAAAGTGCTTGTTGAGCTTCTGCATTTGCACAATGGGCGATGAGGTCGAAGTCATTGTG
CGCTCCTTATTGGCCAACGACGCCGCGCCGTTCCAACTGGCGCAGGATGATAGAGAGGGTCCAGGTAATGGCCAGGTACATCAGGGCGACCATGAAATAG
ACTTCCAGCGCCGTGAAGGTGGTGGCAATGTAGATCTGCCCTTGGCGCACCAGCTCGCCCACCCCGATGACGGAAAACAGCGAGGTGTCCTTGATGCTGA

>Halomonas hamiltonii KCTC 22154 [NZ_BMXN01000011.1:53753..53807], dnaA> ig dnaN>, 56 bp

< page 1 | #071 | Halomonas humidisoli WN018

WebSkew GCmin pos. 64685 (GCmin<>oriC1 22.0 kb; GCmin<>oriC2 0.6 kb; oriC2<>oriC1 21.4 kb)

>Halomonas humidisoli WN018 [NZ_NSKA01000003.1:42372..42871], <rpmH oriC1 dnaA>, 534 bp

AGCACGTCGGTCGTGACAGCAATAAAGTCTCGGCTAGACGAGTTACTTTCTGCTTCTCCAGGCATGCCTTCACTAATGGCATTTTGGGGAGCCGACGAAC
TAGGAACCGATAAATCATCACTGCCAACATTATTGGCGGATGAGCCGTTGCTGTTTTGGGTAATTTCAGGCGTCGAATCGACGCTTATTTGCCCATAATC
CTGATTCCACTGTACAACCAATAGGTAGGCAAGTACTGCCAGTGGAATCAGTAATAAAAGTCGTTTTACGTCCATGAGGGTTCTGCCCGATGGGTGGTGA
ACATGCTAATGGCGCAGTGTCATTTACTCGACCCATGACACACGAAAGCCTGCCAGGCGGCAGGCCGAGGTCGAGCGGCGAGGTGTGTCGTCAAGCGTCA
CGTGTTACGCCATTAGGCGGGAGTGATACAGACACCGCCTGTGCGTCACGCTGAAGTCGTTTCCACATGCCATTTAGCTGGCGAGTAAGCGTCTCGTTAT
CAATATCTTGAACACCACGCTTTGCCAATATCACGATATCCACAGAGGGTAAGTGATCTTGGCGGAGGCGAACAGATTCTCTCACGAGTCGCTTAAAGCG
GTTGCGATCAACAGCGAGCTTCACATTCTTTTTGCTGACCACCAGTCCAATGCGGGGATGTCCCCGGATACTCAAACGCGCTAGCGCCATCATACCTTTG
CCATGAACCTTCAGGTCTGCTTGTTCAAACACGTGACTAAAATCCCCGGCGTTTAGCAAACGTAAGCACCGGGGAAAGCCTTGATGCGGCACACGCAATC
CGAGATCAGGCGCTCAGACGCTTGCGGCCTTTGGCGCGGCGGCGTGCGATAACGGCACGGCCGTTTTTGGTTGCCATACGAGCACGGAAGCCATGCGCGC
GCTTGCGCTTGAGAACGCTGGGTTGAAAAGTGCGTTTCATAACTCGTTATTCCCACGTGGTTTAGGACGGAAAGTGAATTCCAAAAGACCCGGTCCAGTG

AGGAACTAGCCGGGGAGTTTCAATTCAAGACCGGAAATTGTAGAGACTTCCCGCGTCAGGTGCAATTTTTCCTTGCATAAAAAGCGTGAAAAAGGCAATG
GAGGCTCTTCGTACATCGCCATCGTGGCTTCTTGCCATGACATTACTATTACTAATGAAGTTATTTAAAACTTACTGTTATTAATAGTGGGGATACTAAT
**TGTGGATAA**GCTATTTTAGCTCATTTTTATCAACAGCTTAAAAACATAAAAAATCTGTGCATAACCTTCGGATAAGCCTTAAATAAACTGTTAATTAGGT
GTGTTCAACAGGGTATAGAAGACATCAAAACAGCTTTTTCCTTGATCAGTTACCAGCTTTTCCCTATGTCTTGTGAATAGTTGTCCACAGGGCGAAAAGA
GGCGCTAGTGCGTGTGGATAACCGTCCCGTTGGGCGCTACAATGGTCGGCCGTTTTCAACCGATGGTGAGATGAGTGTCACTCGCGCTTTGGCAACAGTG

CCTGGACTACCTGCAGGACGAACTGAATTCACAGCAGTTCAATACCTGGATACGTCCGCTGCAGGCAGAAGAAGGAGAGACCAATGAGCTGCGCTTGTTG
GCGCCGAATCGCTTTGTGCGCGACTGGGTGAGTGATAAGTATTCAAAGCGTATCAGTGAGCTGATGCGAGAACTTGCGCCATCCAAGCCTCCCAAAGTCA
GTTTGACGGTAGGTAGCCGCCGTGTGGCTGCTCCTGCCCCACAGCCAAGAGATTTGGGTAATCCGGTCTCAGCATCTCCCTCTAGGCAAGCATCGCCTGC
GGTTCATACGCCGCCAAGGGGAGACCTTGCTGATGAGCGAGAAATTGATCAACTGCGGGAAGAGTCACCGAATCGGCGAGGAAATGAGCGCCAGGTACAG
GTTGAAGGCAGCCTAAAACATGGTAGTGGTCTTAATCCTAATTTTACCTTCGATACCTTTGTTGAAGGTAAATCGAACCAGCTCGCTCGTGCGGCATCTC
GTCAGGTATCTGAAAATCCTGGCGGTGCCTATAACCCGCTGTTTTTATACGGTGGCGTTGGTTTAGGTAAAACACACTTGATGCACGCTGTCGGTAACGC
CCTGGCTGGCCGACGTGAAAATGCCCGTGTGGTGTATCTACACTCTGAACGCTTTGTCGCGGATATGGTGAAAGCGCTGCAGTTAAATGCGATTAACGAC
TTCAAGCGCTTTTATCGCAGCGTTGATGCGTTACTTATTGATGATATTCAGTTTTTTGCTGGAAAAGAGCGTTCCCAGGAAGAGTTCTTCCATACCTTTA
ATGCATTATTAGAGGGTGGGCAGCAAATGATACTTACCTCTGACCGCTATCCAAAAGAGATTAGTGGTGTGGAGGAGCGTCTGAAATCGCGCTTTGGCTG
GGGCCTGACAGTGGCTATTGAGCCGCCAGAGCTTGAAACCCGCGTAGCAATATTAATGAAAAAAGCAGATCAGGCGAAAGTTGATCTTCCACACGACGCT

>Halomonas humidisoli WN018 [NZ_NSKA01000003.1:63807..64306], <mnmG oriC2 <COG1126 (glnQ), 565 bp

TAACCCTTTAATTGAGCGAACCACTTGTAACTGAACATCGAAAGGCAACGACGTCGAGATTCCGTTGGGATAGAGCTCATGGGTATCCAAACCTTCAGGC
TCGATAAATACCTGATGGCTTGATTTGTCGGCAAAGCGGTGAACTTTGTCTTCAATCGATGGGCAGTATCGTGGTCCAACGCCTTCGATCACACCGGAAT
ACATAGGTGAGCGATCAAGATTTGCCATGATGATTTCATGGGTGCGCTCATTAGTGTGCGCTATGTGGCAACTCACCTGCTGTGGATGCATCTCCCGTGA
ACCAAGGTATGACATGACTGGCGTTGGGGTGTCGCCTGGCTGTTCTTCCAACGATGTGAAGTCAACGGTTTTGGCATCAATTCGTGGAGGCGTCCCGGTC
TTTAGCCGATCAACGCGAAACGGTAGTGCGCGTAAGCGTTCAGCTAGCGCGTTGGAAGGCGGGTCACCGGCGCGGCCACCGCGGCTTTGATCTAACCCAA
TGTGGATAACGCCACCCAAAAACGTACCGGTTGACAGCACAACGGACTCCGCATGGAAGCGGATGCCTGTTTCGGTCACAACGCCTCGAACAGTGTTGTT
ATCCACAACCAGATCGCCAGCGGCTTGCTGAAAAATCGTTAAGTTCGGCTGGTTTTCCAACATTCCCCGAATAGCGGCTTTGTAACGAATTCGGTCTGCT
TGAGCACGTGTTGCTCGAACGGCTGGACCTTTACGAGCGTTCAATACCCGAAACTGGATGCCGCCTAAGTCCGTGGCTAGCCCCATTGCACCACCGAGTG
CATCAATTTCTTTTACCAGGTGGCTTTTGCCAATCCCACCAATCGCTGGATTACACGACATTTGGCCGAGCGTTTCGATGTTGTGGGTTAGCAATAGGGT
TTGACAACCCATACGAGCAGAGGCCAATGCGGCTTCAGTTCCCGCATGGCCACCGCCGATGACAATCACGTCAAAGCGGTCGGGATAGTTCAAGAGGCAC

CTCGTCTTGCGCCGTTTGGCACGGATGATGTGAGTCAGTATTGGTTCACTGATTGAGTAAGCCCGCCAGTATAAACCTCCTGTGGGTAACCGTCAGGTTT
TTCCACACCCCAAGCTAAGCATTCATCTATTATTAATAACAAAATAAAAATATATAAGAGAAGTTCTGTTGATGTTGTAACTACTAGTGTGGACAAAATC
**TGTTGATAA**CCCAAATAAACTAATATATTTCAGTTGCTTAAATTGTTAGTTTTACATGATTAAACGGAGGAGTAGCTGCGTAGAAGCGGTGCTACTCCT
GTGGATGAAAGTGCGACTTACACACAGGCATGACAGACAACGGGTTATCCACCGGCCCATACCGAGTTTGTTACCGCACCTGTGAACAAGTGCGATTGGT
ATCTTAAAAAGATCTGATTAGCGCTCATGGTAGGCGTCGTAGCGCGATATGACGCAGTGTGTAAAAAATAGTATCCACAGGCAAAAAAAATGGCCTGTCG

CGATGACAGGCCATTTTACGGGGTAAAAACGGGCAAAAACGTCAGATTAATTCCGACATTAATGCTTCAAAACACGTGCAAGGAAGGATTGTGTTCGCTC
ATGTTGTGGCGTATCAAACAGTTGCTCGGGAGGGCCTTGCTCGGCAATTTCGCCTTTATGGATAAAAATAACGCGGTCTGCCACTTCACGGGCAAAACCC
ATTTCATGGGTCACTATCACCATGGTCATGCCTTCTTTGGCAAGTTCACGCATGGCATCCAGCACTTCACCAATCATTTCAGGGTCAAGCGCTGATGTAG
GCTCGTCAAACAGCATCAGGCGGGGTTCCATGGCTAACGCCCGGGCTAACGCCACACGCTGCTGCTGTCCACCCGAGAGCTGGCTTGGATACTTATCTGC
TTGGTCGGCAATACCAACGCGCTCTAACAGTCGTTCAGCGGTCTCTTCGGCATCTTGGCGGCTCCACCCCCGCACTTTCATGGGGGCAAGGGTAACGTTG
TCTCGCACGCTCAAGTGGGGAAAAAGGTTGAATTGCTGAAATACCATACCGACTTCGGTACGGATAGTTTGCAGCGCTTTACTGCTTTTACCGTTGGGCA
GTAATTGGTTGCCATCAACATCAAGAGATCCGGATTGAAACTCTTCTAAGCCGTTGATACAGCGGATGAGCGTTGATTTACCAGAGCCACTGGCACCGAT
GATAACGACGACTTCACCAGGCACAATTTCAAGATCAACATTATTGAGTACGTGCAGGCTGCCAAAATGCTTATTGAGCTGCTGCATACGCACAATTGGC
GATTGGGTAGAGGTCATTGCACGTTCCTTATTGTCCTGCGAGGCCTCTGCGCTCAATTTGGCGCAGAATCAAAGAGAGAGTCCAGGTGATGGCAAGGTAC
ATCAGCGCCACCATGAAATAGACCTCTAGTGCGGTAAAGGTAGTGGCAATATAAATTTGCCCTTGGCGGACTAATTCACCGACCCCGATAACAGAAAATA

>Halomonas humidisoli WN018 [NZ_NSKA01000003.1:40876..40930], dnaA> ig dnaN>, 55 bp

< page 1 | #072 | Halomonas alkaliphila DSM 16354

WebSkew GCmin pos. 64610 (GCmin<>oriC1 22.6 kb; GCmin<>oriC2 1.0 kb; oriC2<>oriC1 21.6 kb)

>Halomonas alkaliphila DSM 16354 [NZ_JABASV010000002.1:41797..42296], <rpmH oriC1 dnaA>, 534 bp

AGAACGTCGGTCGTGACAGCAATAAAGTCTCGGCTAGACGAGTTACTTTCTGCTTCTCCAGGCATGCCTTCACTAATGGCATTTTGGGGAGCCGATGAAC
TAGGAACCGATAAATCATCACTGCCAACATTATTGGCGGATGAGCCGTTGCTGTTTTGGGTAATTTCAGGCGTCGAATCGACGCTTATTTGCCCATAATC
CTGATTCCACTGTACAACCAATAGATAGGCAAGTACTGCCAGTGGAATCAGTAATAAAAGTCGTTTTACGTCCATGAGGGTTCTGCCCGATGGGTGGTGA
ACATGCTAATGGCGCAGTGTCATTTACTCGACCCATGACACACGAAAGCCTGCCAGGCGGCAGGCCGAGGTCGAGCGGCGAGGTGTGTCGTCAAGCGTCA
CGTGTTACGCCATTAGGCGGGAGTGGTACAGACACCGCCTGTGCGTCACGCTGAAGTCGTTTCCACATGCCATTTAGCTGGCGATTAAGCGTCTCGTTAT
CAATATCTTGAACACCACGCTTTGCCAGTATCACGATATCCACAGAGGGTAAGTGATCTTGGCGGAGGCGAACAGATTCTCTCACGAGTCGCTTAAAGCG
GTTGCGATCAACAGCGAGCTTCACATTCTTTTTGCTGACCACCAGTCCAATGCGGGGATGTCCCCGGGTACTCAAACGCGCTAGCGCCATCATACCTTTG
CCATGAACCTTCAGGTCTGCTTGTTCAAACACGTGACTAAAATCCCCGGCGTTTAGCAAACGTAAGCACCGGGGAAAGCCTTGATGCGGCACACGCAATC
CGAGATCAGGCGCTCAGACGCTTGCGGCCTTTGGCGCGGCGGCGTGCGATAACGGCACGGCCGTTTTTGGTTGCCATACGAGCACGGAAGCCATGCGCGC
GCTTGCGCTTGAGAACGCTGGGTTGAAAAGTGCGTTTCATAACTCGTTATTCCCACGTGGTTTAGGACGGAAAGTGAATTCCAAAAGACCCGGTCCAGTG

AGGAACTAGCCGGGGAGTTTCAATTCAAGACCGGAAATTGTAGAGACTTCCCGCGTCAGGTGCAATTTTTCCTTGCATAAAAAGCGTTAAAAAGGCAATG
GAGGCTCTTCGTACATCGCCATCGTGGCTTCTTGCCATGACACTACTATTACTAATGGAATTATTTAAAACCTATTGTTATTAATAGTGGGGATACTAAT
**TGTGGATAA**GCTATTTTAGCTTACTTTTATCAACAGCTTAAAAACATAAAAAATCTGTGCATAACCTTCGGATAAGCCTTAAATAAACTGTTAATTAGGT
GTGTTCAACAGGGTATAGAAGACATCAAAACAGCTTTTTCCTTGATCCATTACCAGCTTTTCCCTATGCCTCATGAATAGTTGTCCACAGGGCAAAAAGA
GGCGTTAGTGCGTGTGGATAACCGTCTCGTTGGGCGCTACAATGGTCGGCCGTTTTCAACCGATGGTGAGATGAGTGTCACTCGCGCTTTGGCAACAGTG

CCTGGACTACCTGCAGGACGAACTGAATTCACAGCAGTTCAATACCTGGATACGTCCGCTGCAGGCAGAAGAAGGAGAGACCAATGAGCTGCGCTTGTTG
GCGCCGAATCGCTTTGTGCGCGACTGGGTGAGTGATAAGTATTCAAAGCGTATCAGCGAGCTGATGCGAGAACTTGCGCCATCCAAGCCTCCCAAAGTCA
GTTTGACGGTAGGTAGCCGCCGTGTGGCTGCTCCTGCCCCGCAGCCAAGAGATTTGGGTAATCCGGTCTCAGCATCTCCCTCAAGGCAAGCATCGCCTGC
GGTTCATACGCCGCCAAGGGGAGACCTTGCTGATGAGCGAGAAATTGATCAACTGCGTGAAGAGTCACCGAATCGGCGAGGAAATGAGCGCCAGGTACAG
GTTGAAGGCAGCCTAAAACATGGTAGTGGTCTTAATCCTAATTTTACCTTCGATACCTTTGTTGAAGGTAAATCGAACCAGCTCGCTCGTGCGGCATCCC
GTCAGGTATCTGAAAATCCTGGCGGTGCCTATAACCCGCTGTTTTTATACGGTGGTGTTGGTTTAGGTAAAACACACTTGATGCACGCTGTCGGTAACGC
CCTGGCTGGCCGCCGCGAAAATGCCCGTGTGGTGTATCTACACTCTGAACGCTTTGTCGCGGATATGGTAAAAGCACTGCAGTTAAATGCGATTAACGAC
TTCAAGCGCTTTTATCGCAGCGTTGATGCGCTACTTATTGATGATATTCAGTTTTTTGCTGGAAAAGAGCGTTCCCAGGAAGAGTTCTTCCATACCTTTA
ATGCGTTGTTAGAGGGTGGGCAGCAAATGATACTTACCTCTGATCGCTATCCAAAAGAGATTAGTGGTGTGGAGGAGCGTCTGAAATCGCGCTTTGGCTG
GGGCCTGACAGTGGCTATTGAGCCGCCAGAGCTTGAAACCCGTGTAGCGATTTTGATGAAAAAGGCAGATCAGGCGAAAGTTGATCTTCCACACGACGCT

>Halomonas alkaliphila DSM 16354 [NZ_JABASV010000002.1:63347..63846], <mnmG oriC2 <COG1126 (glnQ), 565 bp

TAACCCTTTAATTGAGCGAACCACTTGTAGCTGAACATCGAAAGGCAACGACGTCGAGATTCCGTTGGGATAGAGCTCATGGGTATCCAACCCTTCAGGC
TCGATAAATACCTGATGGCTTGATTTGTCGGCAAAGCGGTGAACTTTGTCTTCAATCGATGGGCAGTATCGCGGACCAACACCTTCGATCACACCGGAAT
ACATAGGCGAGCGATCAAGATTCGCCATGATGATTTCATGGGTGCGCTCATTGGTGTGCGCTATGTGGCAACTCACCTGCTGCGGATGCATCTCCCGTGA
ACCAAGGTATGACATGACCGGTGTTGGGGTGTCACCTGGCTGTTCTTCCAACGATGTGAAGTCAACGGTTTTGGCATCAATTCGTGGAGGCGTCCCGGTT
TTTAGCCGATCAACGCGAAACGGTAGTGCGCGTAAGCGTTCAGCTAGCGCGTTGGAAGGCGGGTCACCGGCGCGGCCACCGCGGCTTTGATCTAACCCAA
TGTGGATAACGCCACCCAAAAACGTACCGGTTGACAGCACAACGGACTCCGCATGGAAGCGGATGCCTGTTTCGGTCACAACGCCTCGAACAGTGTTGTT
ATCCACAACCAGATCGCCAGCGGCTTGCTGAAAAATCGTTAAGTTCGGCTGGTTTTCCAGCATTCCCCGAATAGCAGCTTTGTAGCGAATTCGGTCTGCT
TGAGCACGTGTTGCTCGAACGGCTGGACCTTTACGAGCGTTCAATACCCGAAACTGGATGCCGCCTAAGTCCGTGGCTAGCCCCATTGCACCACCGAGTG
CATCAATTTCTTTTACCAGGTGGCTTTTGCCAATCCCACCAATCGCTGGATTGCACGACATTTGGCCGAGCGTTTCGATGTTGTGGGTTAGCAATAGGGT
TTGACAACCCATACGAGCAGAGGCCAATGCGGCTTCGGTTCCCGCATGGCCACCGCCGATGACAATCACGTCAAAGCGGTCGGGATAGTTCAAGAGGCAC

CTCGTCTTGCGCCGTTTGGCACGGATGATGTGAGTCAGTATTGGTTCACTGATTGAGTAAGCCCGCCAGTATAAACCTCCTGTGGGTAACCGTCAGGTTT
TTCCACACCCCAAGCTAAGCATCAATCTATTATTAATAACAAAATAAAAATATATAAGAGAAGTTCTGTTGATGTTGTAACTACTGGTGTGGACAAAATC
**TGTTGATAA**CCCAAATAAGCTAATATATTTCAGTTGCTTAAATTGTTAGTTTTTACATGATTAAACGGAGGAGTACCTGCGTAGAAGCGGTGCTACTCCT
GTGGATGAAAGTGCGACTTACACACAGGCATGACAGACAACGGGTTATCCACCGACCCATACCGAGTTTGTTACCGCACCTGTGAACAAGTGCGATTGGT
ATCTTATAAAGACCTGGTTAGCGCTCATGCTAGGCATCGTAGAGCGATATGATGCAGCGTGTAAAAAATAGTATCCACAGGCAAAAAAAATGGCCTGTCG

CGATGACAGGCCATTTTACGGGGCAAAAATGGGCAAAAAAGTCAGATTAATTCCGACATTAATGCTTCAAAACACGTGCAAGGAAGGATTGTGTTCGCTC
ATGTTGTGGCGTATCGAACAGTTGCTCGGGAGGGCCTTGCTCGGCAATTTCGCCTTTATGGATAAAAATAACGCGATCTGCCACTTCACGGGCAAAGCCC
ATTTCATGGGTCACTATCACCATGGTCATGCCTTCTTTGGCAAGTTCACGCATGGCATCCAGCACTTCACCAATCATTTCAGGGTCAAGCGCTGATGTAG
GCTCGTCAAACAGCATCAGGCGGGGTTCCATGGCTAACGCCCGGGCTAACGCCACACGCTGCTGCTGCCCACCCGAGAGCTGGCTTGGGTACTTATCTGC
TTGGTCGGCAATACCAACACGCTCTAACAGTCGTTCAGCGGTCTCTTCGGCATCTTGGCGGCTCCAGCCCCGCACTTTCATGGGCGCAAGGGTAACGTTG
TCGCGCACGCTCAAGTGGGGAAAAAGGTTGAACTGCTGAAACACCATGCCGACTTCAGTACGGATAGTTTGTAGCGCTTTACTGCTTTTACCGTTGGGCA
GTAATTGGTTGCCATCAACATCAAGAGAGCCGGCTTGAAACTCTTCTAAGCCGTTGATACAGCGGATGAGCGTTGATTTACCAGAGCCACTGGCACCGAT
GATAACGACGACTTCACCAGGCACAATTTCAAGATCAACATTATTGAGTACGTGCAGGCTGCCAAAATGCTTATTGAGCTGCTGCATACGCACAATTGGC
GATTGGGTAGAGGTCATTGCACGTTCCTTATTGTCCTGCGAGGCCTCTGCGCTCAATTTGGCGCAGAATCAAAGAGAGAGTCCAGGTGATGGCAAGGTAC
ATCAGCGCCACCATGAAATAGACCTCAAGTGCGGTAAAGGTAGTGGCAATATAAATTTGCCCTTGGCGGACTAATTCACCGACCCCGATAACAGAAAATA

>Halomonas alkaliphila DSM 16354 [NZ_JABASV010000002.1:40301..40355], dnaA> ig dnaN>, 55 bp

WebSkew GCmin pos. 62436 (GCmin<>oriC1 21.6 kb; GCmin<>oriC2 0.0 kb; oriC2<>oriC1 21.5 kb)

>Halomonas venusta NBRC 102221 [NZ_BJUL01000002.1:c40612..41111], <rpmH oriC1 dnaA>, 535 bp

GAACGTCGGTAGTGACAGCAATAAAGTCTCGGCTAGACGAGTTACTTTCTGCTTCTCCCGGCATGCCTTCACTAATGGCATTTTGGGGAGCCGACGAACT
GGGAACCGATAAATCATCACTGCCAACATTGTTGGCGGATGAGTCGTTGCTGTTTTGGGTAATTTCAGGCGTCGAATCGACGCTTATTTGCCCATAATCC
TGATTCCACTGTACAACCAATAGGTAGGCAAGTACTGCCAGTGGAATCAGTAATAAAAGTCGTTTTACGTCCATGAGGGTTCTGCCCGATGGGTGGTGAA
CATGCTAATGGCGCAGTGTCATTTACTCGACCCATGACACACGAAAGCCTGCCAGACGGCAGGCCGAGGTCGAGCGGCGAGGTGTGTCGTCAAGCGTCAC
GTGTTACGCCATTAGGCGGGAGTGATACAGACACCGCCTGTGCGTCGCGCTGAAGCCGTTTCCACATGCCATTTAGCTGGCGATTCAGCGTCTCGTTATC
AATATCTTGAACACCGCGCTTTGCCAGTATCACGATATCCACAGAGGGTAAGTGATCTTGGCGGAGGCGAACAGATTCTCTCACGAGTCGCTTAAAGCGG
TTGCGATCAACAGCGAGCTTCACATTCTTTTTGCTGACCACCAGTCCAATGCGGGGATGCCCCCGGGTACTCAAACGCGCTAACGCCATCATACCTTTGC
CATGAACCTTCAGGTCTGCTTGTTCGAACACGTGACTAAAATCCCCAGCGTTTAGCAAACGTAAGCACCGGGGAAAGCCTTGATGCGGCACACGCAATCC
GAGATCAGGCGCTCAGACGCTTGCGGCCTTTGGCGCGGCGGCGTGCGATAACGGCACGGCCGTTTTTGGTTGCCATACGAGCACGGAAGCCATGCGCGCG
CTTGCGCTTGAGAACGCTGGGTTGAAAAGTGCGTTTCATAACTCGTTATTCCCACGTGGTTTAGGACGGAAAGTGAATTCCAAAAGACCCGGTCCAGTGA

GGAAACTAGCCGGGGAGTTTCAATTCAAGACCGGAAATTGTAGAGACTTCCCGCGTCCGGTGCAATTTTTCCTTGCACAAAAAGCGTTAAAAAGGCAATG
GAGGCCCTGTGCACATCGCCATCGTGGCTTCTTGTCATGACACTATTATTACTAATGAGGTTATTTAAAACCTACTGTTATTAATAGTGGGGATACTTAT
**TGTGGATAA**GTTGTTTTAGATGATTTTTATCAACAGCTTAAAAAAATAAAAAATCTGTGCATAACCTTCGGATAAGCCTTAAATAAACTGTTAATTAGGT
GTGTTCAACAGGTTATAGAATACATCAAAACAGCTTTGTCCCTGATCGATTCCCAGCTTTTCCCTATGCCTTATGAATAGTTGTCCACAAGGCAAAAAGA
GGCATTAGTGCGTGTGGATAACCGGTCCGTTGGGCGCTACAATGGTCGGCCGTTTTCAACCGATGGTGAGATGAGTGTCACTCGCGCTTTGGCAACAGTG

CCTGGACTACCTGCAGGACGAACTGAATTCACAGCAGTTCAATACCTGGATACGTCCGCTGCAGGCAGAAGAAGGAGAGACCAATGAGCTACGCTTGTTG
GCGCCGAATCGCTTTGTGCGCGACTGGGTGAGTGATAAGTATTCAAAGCGTATCAGCGAACTGATGCGCGAGCTTGCGCCATCCAAGCCTCCTAAAGTCA
GTTTGACAGTAGGTAGCCGCCGTGTGGCAGCACCTGCCCCGCAGCCAAGAGACTTGGGTAATCCCGTCTCAGCATCTCCCTCAAGGCAAGCATCGCCTGC
GGTTCATACGCCGCCAAGAGGGGATCTTGCCGATGAGCGAGAAATTGATCAGCTACGGGAGGAGTCACCAAGTCGGCGAGGTAATGAGCGCCAGGTACAG
GTTGAAGGCAGCCTAAAACATGGCAGCGGGCTAAACCCTAATTTTACCTTCGATACTTTTGTTGAAGGTAAATCGAACCAGCTCGCTCGTGCGGCATCTC
GTCAGGTATCTGAAAATCCTGGCGGAGCCTATAACCCACTGTTTTTATACGGTGGCGTTGGTTTAGGTAAAACACACTTGATGCACGCTGTCGGTAATGC
CCTGGCTGGCCGGCGTGAAAATGCCCGAGTGGTGTATCTACACTCTGAGCGCTTTGTCGCGGATATGGTGAAAGCACTGCAGTTAAATGCGATTAACGAT
TTCAAGCGTTTCTATCGCAGCGTTGATGCACTACTTATTGATGATATTCAGTTTTTTGCTGGAAAAGAGCGTTCCCAGGAAGAGTTCTTCCACACCTTTA
ATGCATTATTAGAGGGTGGACAGCAAATGATACTTACCTCTGACCGCTACCCAAAAGAGATTAGTGGTGTAGAGGAGCGTCTGAAATCTCGCTTTGGCTG
GGGGCTGACTGTGGCTATTGAGCCGCCAGAGCTTGAAACCCGTGTAGCGATTTTAATGAAAAAAGCCGATCAGGCGAAAGTTGATCTCCCGCACGACTCT

>Halomonas venusta NBRC 102221 [NZ_BJUL01000002.1:c62148..62647], <mnmG oriC2 <COG1126 (glnQ), 566 bp

TAAGCCTTTAATTGAGCGAACCACTTGTAGCTGAACATCAAAAGGCAACGACGTCGAGATTCCGTTGGGATAGAGCTCATGGGTATCCAATCCTTCAGGC
TCGATAAACACCTGATGGCTTGATTTGTCGGCAAAGCGGTGAACTTTGTCCTCAATCGATGGGCAGTATCGCGGACCAACGCCTTCGATCACACCGGAAT
ACATAGGCGAGCGATCAAGATTCGCCATGATGATTTCATGGGTGCGCTCATTGGTGTGCGCTATGTGGCAACTCACCTGCTGTGGATGCATCTCCCGTGA
ACCAAGGTACGACATGACCGGCGTGGGGGTGTCACCTGGCTGTTCTTCCAACGCTGTAAAATCAACAGTTTTGGCATCAATTCGTGGAGGCGTTCCGGTT
TTTAGTCGATCAACGCGAAACGGCAGTGCGCGTAAGCGTTCAGCTAGCGCGTTGGAAGGCGGGTCACCGGCGCGGCCACCGCGGCTTTGATCTAACCCAA
TGTGGATAACGCCACCCAAAAATGTACCGGTTGACAGCACAACGGACTCCGCATGGAAGCGGATGCCTGTTTCGGTCACAACGCCTCGAACAGTGTTGTT
ATCCACAATCAGATCGCCAGCGGCTTGCTGAAAAATGGTTAAGTTCGGCTGGTTTTCCAACATTCCCCGAATAGCAGCTTTATAGCGAACTCGGTCTGCC
TGAGCACGTGTTGCTCGAACGGCTGGGCCTTTACGAGCGTTCAATACCCGAAACTGGATGCCGCCTAAGTCCGTGGCTAGCCCCATTGCACCACCGAGTG
CATCAATTTCTTTTACCAGGTGGCTTTTGCCAATCCCACCAATCGCTGGATTGCACGACATTTGGCCGAGCGTTTCGATGTTGTGGGTTAGCAATAGGGT
TTGACAACCCATACGAGCAGAGGCCAATGCGGCTTCAGTTCCCGCATGGCCACCGCCGATGACAATCACGTCAAAGCGGTCGGGGTAGTTCAAGAGGCAC

CTCGTCTTGTGCCATTCGGCACGGATGATGTGAGTCAGTATTGGTTCACTGATTGAGTAAGCCCGCCAGTATAAACCTCCTGTGGGTAACCGTCAGGTTT
TTCCACACCCTAAACGAAGCATTCATCTATTATTAATAACAAAATAAAAATATATAAGAGAAGTTCTGTTGATGTTGTAACTAACTGGTGTGGACAAAATC
**TGTTGATAA**CTGAAATAAATCATTATTTTTCAGGTTTTTACATTGTTAGTTTTTACATGATTAAACGGCGGAGTACCTGCGTAGAAACGGTGCTACTCCT
GTGGATGAAAGTGCAACTTACACACAAGCATAGCAAACAACGGGTTATCCACCGGCCCATACCGAGTTTGTTACCGCACCTGTGAACAAGTTCGATTGGT
ATCTTCAAAAGACCTTATTAGCGCTCATGGTAGGCATCGTAGAGCGATATGCCCCAGCGTGTAAAAAATAGTATCCACAGGCAAAAAAAATGGCCTGTCG

CGATGACAGGCCATTTAGCGGGACAAAAATGGGCAAAAAAAGTCAGATTAATTGCGACATTAATGCTTCAAAACACGTGCAAGGAAGGATTGTGTTCGCT
CATGTTGTGGCGTATCGAACAGTTGCTCGGGGGGACCTTGTTCGGTAATTTCGCCTTTATGGATGAAAATGACACGGTCTGCCACTTCGCGGGCAAAGCC
CATTTCATGGGTCACAATCACCATGGTCATGCCTTCTTCGGCAAGTTCACGCATGGCGTCCAGTACTTCGCCGATCATTTCAGGGTCAAGCGCTGATGTA
GGCTCGTCAAACAGCATAAGACGAGGTTCCATGGCCAATGCCCGGGCTAACGCCACACGCTGCTGTTGCCCACCTGAAAGTTGGCTTGGGTACTTAGCTG
CTTGGTCGGCAATACCAACGCGCTCTAACAGTCGTTCAGCGGTCTCTTCGGCATCCTGACGGTTCCATCCCCGCACTTTCATGGGGGCAAGAGTAACGTT
GTCGCGCACGCTCAAGTGGGGAAAAAGGTTGAACTGCTGAAACACCATGCCGACTTCAGTACGGATAGTTTGCAGCGCCTTACTGCTTTTACCGTTGGGC
AGTAATTGGTTGCCATCAACATCAAGAGAGCCAGCTTGAAACTCTTCTAAGCCATTGATACAGCGGATGAGCGTTGATTTACCTGAGCCACTGGCACCGA
TGATAACGACGACTTCACCAGGCACAATTTCGAGATCAATATTTTTGAGTACGTGCAGGCTGCCAAAATGCTTATTGAGCTGCTGCATACGCACAATTGG
CGATTGGGTAGAGGTCATTGCACGTTCCTTATTGTCCTGCGAGGCCTCTGCGCTCAATTTGGCGCAGAATCAAAGAGAGGGTCCAGGTGATGGCAAGGTA
CATCAGCGCCACCATGAAATAGACCTCAAGTGCGGTAAAGGTAGTCGCAATATAAATTTGCCCTTGGCGGACTAATTCACCGACCCCGATGACCGAAAAT

>Halomonas venusta NBRC 102221 [NZ_BJUL01000002.1:c39116..39170], dnaA> ig dnaN>, 55 bp

WebSkew GCmin pos. 4095280 (GCmin<>oriC1 25.5 kb; GCmin<>oriC2 3.7 kb; oriC2<>oriC1 21.8 kb)

>Halomonas hydrothermalis Slthf2 [NZ_AP022843.1:4120513..189], <rpmH oriC1 dnaA>, 535 bp

GAACGTCGGTCGTGACAGCAATAAAGTCTCGGCTAGACGAGTTACTTTCTGCTTCTCCCGGCATGCCTCCACTAATGGCATTTTGGGGAGCCGACGAACT
AGGAACCGATAAATCATCGCTGCCAACATTATTGGCGGATGAGCCGTTGCTGTTTTGGGTAATTTCAGGCGTCGAGTCGACGCTTATTTGCCCATAATCC
TGATTCCACTGTACAACCAATAGGTAGGCAAGTACTGCCAGTGGAATCAGTAATAAAAGTCGTTTTACGTCCATGAGGGTTCTGCCCGATGGGTGGTGAA
CATGCTAATGGCGCAGTGTCATTTACTCGACCCATGACACACGAAAGCCTGCCAGACGGCAGGCCGAGGTCGAGCGGCGAGGTGTGTCGTCAAGCGTCAC
GTGTTACGCCATTAGGCGGGAGTGATACAGGCACCGCCTGTGCGTCGCGCTGAAGCCGTTTCCACATGCCGTGTAGCTGGCGATTTAGCGTCTCGTTATC
AATATCTTGAACACCACGCTTTGCCAATATCACGATATCCACAGAGGGTAAGTGATCCTGGCGGAGGCGAACAGATTCTCTCACGAGTCGCTTAAAGCGG
TTGCGATCAACAGCGAGCTTCACATTCTTTTTGCTGACCACCAGTCCAATGCGGGGGTGCCCCCGGGTACTCAAACGCGCTAACGCCATCATACCTTTGC
CATGAACCTTCAGGTCTGCTTGTTCGAACACGTGACTAAAATCCCCAGCGTTTAGCAAACGCAAGCACCGGGGAAAGCCTTGATGCGGCACACGCAATCC
GAGATCAGGCGCTCAGACGCTTGCGGCCTTTGGCACGGCGGCGTGCGATAACGGCACGGCCGTTTTTGGTTGCCATACGAGCACGGAAGCCATGCGCGCG
CTTGCGCTTGAGAACGCTGGGTTGAAAAGTGCGTTTCATAACTCGTTATTCCCACGTGGTTTAGGACGGAAAGTGAATTCCAAAAGACCCGGTCCAGTGA

GGAAACTAGCCGGGGAGTTTCAATTCAAGACCGGAAATTGTAGAGACTTCCCGCGCCCGGTGCAATTTTTCCTTGCATAAAAAGCGTTAAAAAGGCAATG
GAAGCCCTGTCGACATCACCATCGTGGCTTTTTGTCATGACACTACTATTACTAATGAGGTTATTTAAAACCTACTGTTATTAATAGTGGGGATACTTAT
**TGTGGATAA**GTTGTTTTAGACGATTTTTATCAACAGCTTAAAAAAATAAAAAATCTGTGCATAACCTTCGGATAAGCCTTAAATAAACTGTTAATTAGGT
GTGTTCAATAGGTTATAGAACACATCAAAACAGCTTTGTCCCTGATCGATTCCCAGCTTTTCCCTATGCCTTATGAATAGTTGTCCACAAGGCAAAAAGA
GGCATTAGTGCGTGTGGATAACCGGTCCGTTGGGCGCTACAATGGTCGGCCGTTTTCAACCGATGGTGAGATGTGTGTCACTCGCGCTTTGGCAACAGTG

CCTGGACTACCTGCAGGACGAACTGAATTCACAGCAGTTCAATACCTGGATACGTCCGCTGCAGGCAGAAGAAGGAGAGACCAATGAGCTACGCTTGTTG
GCGCCGAATCGCTTTGTGCGCGACTGGGTGAGTGATAAGTATTCAAAGCGTATCAGCGAACTGATGCGCGAACTTGCGCCATCCAAGCCTCCCAAAGTCA
GTTTGACGGTAGGTAGCCGCCGTGTGGCAGCACCTGCCCCGCAGCCAAGAGACTTGGGTAATCCCGTCTCAGCATCTCCCTCAAGGCAAGCGTCGCCTGC
GGTTCATACGCCGCCAAGAGGGGATCTTGCCGATGAGCGAGAAATTGATCAGCTACGGGAGGAGTCACCGAGTCGGCGAGGTAATGAGCGCCAGGTACAG
GTTGAAGGCAGCCTAAAACATGGCAGTGGTCTAAACCCTAATTTTACCTTCGATACTTTTGTTGAAGGTAAATCGAACCAGCTCGCTCGTGCGGCATCTC
GTCAGGTATCTGAAAATCCTGGCGGAGCCTATAACCCACTGTTTTTATACGGTGGCGTTGGTTTAGGTAAAACACACTTGATGCACGCTGTCGGTAATGC
CCTGGCTGGCCGACGTGAAAATGCCCGAGTGGTGTATCTACACTCTGAGCGCTTTGTCGCGGATATGGTGAAAGCACTGCAGTTAAATGCGATTAATGAT
TTCAAGCGTTTCTATCGCAGCGTTGATGCACTACTTATTGATGATATTCAGTTTTTCGCTGGAAAAGAGCGTTCCCAGGAAGAGTTCTTCCACACCTTTA
ATGCATTATTAGAGGGTGGACAGCAAATGATACTTACCTCTGACCGCTACCCAAAAGAGATTAGCGGTGTAGAGGAGCGTCTGAAATCGCGCTTTGGTTG
GGGGCTGACAGTGGCTATTGAGCCGCCAGAGCTTGAAACCCGTGTAGCGATTTTAATGAAAAAAGCCGATCAGGCGAAAGTTGATCTCCCGCACGACGCT

>Halomonas hydrothermalis Slthf2 [NZ_AP022843.1:4098735..4099234], <mnmG oriC2 <COG1126 (glnQ), 565 bp

TAAGCCTTTAATTGAGCGGACCACTTGTAGCTGAACATCAAAAGGCAACGACGTCGAGATTCCGTTGGGATAGAGCTCATGGGTATCCAATCCTTCAGGC
TCGATAAATACCTGATGGCTTGATTTGTCGGCAAAGCGGTGAACTTTGTCCTCAATCGATGGGCAGTATCGCGGACCAACGCCTTCGATCACACCGGAAT
ACATAGGCGAGCGATCAAGATTCGCCATAATGATTTCATGGGTGCGCTCATTGGTGTGCGCTATGTGGCAACTCACCTGCTGTGGATGCATCTCCCGTGA
ACCAAGGTACGACATGACCGGTGTGGGGGTGTCACCTGGCTGTTCTTCCAACGCTGTGAAATCAACGGTTTTGGCATCAATTCGTGGAGGCGTTCCGGTT
TTTAGTCGATCAACGCGAAACGGCAGTGCGCGTAAGCGTTCAGCTAGCGCGTTAGAAGGCGGATCACCGGCGCGGCCACCGCGGCTTTGATCTAACCCAA
TGTGGATAACGCCACCCAAAAACGTACCGGTTGACAGCACAACGGACTCCGCATGGAAGCGGATGCCTGTTTCGGTCACAACACCTCGAACAGTGTTGTT
ATCCACAATCAGATCGCCAGCGGCTTGCTGAAAAATGGTTAAGTTCGGCTGGTTTTCCAACATTCCCCGAATAGCAGCTTTATAGCGAACTCGGTCTGCC
TGAGCACGTGTTGCTCGAACGGCTGGACCTTTGCGAGCGTTCAATACCCGAAACTGGATGCCGCCTAAGTCCGTGGCTAGCCCCATTGCACCACCGAGTG
CATCAATTTCTTTTACCAGGTGGCTTTTGCCAATCCCACCAATCGCTGGATTGCACGACATTTGGCCGAGCGTTTCGATGTTGTGGGTTAGCAATAGGGT
TTGACAACCCATACGAGCAGAGGCCAATGCGGCTTCAGTTCCCGCATGGCCACCGCCGATGACAATCACGTCAAAGCGGTCGGGGTAGTTCAAGAGGCAC

CTCGTCTTGTGCCATTTGGCACGGATGATGTGGGTCAGTATTGGTTCACTGATTGAGTAAGCCCGCCAGTATAAACCTCCTGTGGGTAACCGTCAGGTTT
TTCCACACCCTAAGCTAAGCATTCATCTATTATTAATAACAAAATTAAAATATATAAGAGAAGTTCTGTTGATGTTGTAACTAACTGGTGTGGACAAAATC
**TGTTGATAA**CTGGAATAAATCATTATATTTCAGTTTTTTACATTGTTAGTTTTTACATGATTAAACGGCGGAGTAGCTGCGTAGAATCGGTGCTACTCCT
GTGGATGAAAATGCAACTTACACACAAGCATGGCAGACAACGGGTTATCCACCGGCTCATACCGAGTTTGTTACCGCACCTGTGAACAAGTGCGATTGGT
ATCGTTAAAAGACCTAATTAGCGCTCATGGTCGGCATCGTAGAGCGATATGCCCCAGCGTGTAAAAAATAGTATCCACAGGCAAAAAAATGGCCTGTCGC

GGTGACAGGCCATTTAGCGGGGCAAAAATGGGCAAAAAAAGTCAGATTAATTGCGACATTAATGCTTCAAAACACGTGCAAGGAAGGATTGTGTTCGCTC
ATGTTGTGGCGTATCGAACAGTTGCTCGGGGGGACCTTGTTCGGTAATTTCGCCTTTATGGATAAAAATGACACGGTCTGCCACTTCGCGGGCAAAGCCC
ATTTCATGGGTCACAATCACCATGGTCATGCCTTCTTCGGCAAGTTCACGCATGGCGTCCAGTACTTCGCCAATCATTTCAGGGTCAAGCGCTGATGTAG
GCTCGTCAAACAGCATAAGACGAGGTTCCATGGCTAATGCCCGGGCTAATGCCACACGCTGCTGTTGTCCACCTGAAAGCTGGCTTGGGTACTTAGCTGC
TTGGTCGGCAATACCAACGCGCTCTAACAGTCGTTCAGCGGTCTCTTCGGCATCCTGACGGTTCCATCCCCGCACTTTCATGGGGGCAAGAGTAACGTTG
TCGCGCACGCTCAAGTGGGGAAAAAGGTTGAACTGCTGAAACACCATGCCGACTTCAGTACGGATAGTTTGCAGCGCCTTACTACTTTTACCGTTGGGCA
GTAATTGGTTGCCATCAACATCAAGAGAGCCAGCTTGAAACTCTTCTAAGCCATTGATACAGCGGATGAGCGTTGATTTACCTGAGCCACTGGCACCGAT
GATAACGACGACTTCACCAGGCACAATTTCGAGATCAATATTTTTGAGTACGTGCAGGCTGCCAAAATGCTTATTGAGCTGCTGCATACGCACAATTGGC
GATTGGGTAGAGGTCATTGCACGTTCCTTATTGTCCTGCGAGGCCTCTGCGCTCAATTTGGCGCAGAATCAAAGAGAGAGTCCAGGTGATGGCAAGGTAC
ATCAGCGCCACCATGAAATAGACCTCAAGAGCGGTAAAGGTAGTCGCAATATAAATTTGCCCTTGGCGGACTAATTCACCGACCCCGATGACTGAAAATA

>Halomonas hydrothermalis Slthf2 [NZ_AP022843.1:1631..1685], dnaA> ig dnaN>, 55bp

WebSkew GCmin pos. 63168 (GCmin<>oriC1 22.3 kb; GCmin<>oriC2 0.8 kb; oriC2<>oriC1 21.5 kb)

>Halomonas andesensis DSM 19434 [NZ_RZHG01000019.1:c40635..41134], <rpmH oriC1 dnaA>, 533 bp

ACAGCAATGAAGTCTCGGCTTGACGAGTTGCTTTCCGCTTCCCCGGGCATGCCCTCACTGATGGCATTTTGAGGAGCTGAAGAGCTAGGAACCGAAAAAT
CATCACTGCCACTATTGTTGGCGGATGGGCCGTTGCTGCTTTGGGTAATATCAGGCGTCGAATCAACGCTAATTTGCCCATAATCCTGATTCCACTGCAC
AACCAATAAGTAGGCAAGTACTGCCAGTGGAATCAGTAATAAAAGTCGTTTTACGTCCATGAGGGTTCTGCCCGATGGGTGGTGAACATGCTAATGGCGC
AGTGTCATTTTGCTTGACCTCATTTGCTCGACCCATGACACACGAAAGCCTGCCAGACGGCAGGCCGAGGTCGAGCGGCGAGGTGTGTCGTCAAGCGTCA
CGTGTTACGCCGTTAGGCGGGGAAGATGCAGGCACCGCCTGTGCGTCGCGCTGAAGCCGTTTCCACATGCCATTTAGTTGGCGAGTAAACGTTTCGTTGT
CGATATCGTGTGCACCACGTTTTGCCAGTACCACGATATCCACAGAGGGAAAGTGATCTTGACGGAGGCGAACGGATTCTCTAACGAGTCGCTTGAAGCG
GTTACGATCAACAGCGAGCTTCACATTCTTTTTGCTGACCACCAGTCCAACGCGGGGATGCCCCCGGGTACTCAGACGCGCTAACGCCATCATACCTTTG
CCATGAACTTTGAGGTCTGCTTGTTCAAACACGTGACTAAAGTCCCCGGCGTTCAGCAAACGTAAGCACCGGGGAAAGCCTTGATGCGGCACACGCAATC
CGAGATCAGGCGCTCAGACGCTTGCGGCCTTTGGCGCGACGGCGTGAAATAACGGCACGGCCGTTTTTGGTTGCCATACGAGCACGGAAGCCATGCGCGC
GCTTGCGCTTGAGAACGCTGGGTTGAAAAGTGCGTTTCATAACTCGTTATTCCCACGTGGTTTAGGACGGAAAGTGAATTCCAAAAGACCCGGTCCAGTG

AGGAACCAGCCGGGGAGTTTCAATTCAAGACCGGAAATTGTAGAGACTTCCCGCGTCAGGTGCAATTTTTCCTTGCATAAAAAGCATGAAAAAGGCAATG
GCGTCTCTTCGTAGATTGCGATGTTGGCTTCTTGTCGTGACACTACTATTACTAATAGAGTTATTTAAAATCTATTGTTATTAATAGTGGGGATACTAAT
**TGTGGATAA**GCCATTTTATCTGATTTATATCAATATCTTAAAATCATAAAAAATCTGTGCATAACCTTCGTATAAGCCTTAAATAAACTGTTAATTAGGT
GTGTTCAACAGGTTATAAAACATAGCTAATCCGTTTTGTCCTCGATCCATTACCAGTTTTCCTTATCCCTTACGAATAGTTGTCCACAGGGCAAAAAGAG
GTGGTAGTGCGTGTGGATAACCACTCCGTTGGGCGCTACAATGGTCGGCCGTTTTCAACCGATGGTGAGATGAGTGTCACTCGCGCTTTGGCAACAGTGC

CTGGACTACCTGCAGGACGAACTGAATTCACAGCAGTTCAATACCTGGATACGTCCGCTGCAGGCAGAAGAAGGAGAAACCAATGAGCTGCGCTTGTTGG
CGCCGAATCGCTTTGTGCGCGACTGGGTGAGTGATAAGTATTCAAAGCGTATCAGTGAGCTGATGCGTGAACTCGCGCCATCCAAACCTCCCAAAGTCAG
TTTGACAGTAGGTAGCCGCCGTGTGGCAGCGCCTGCCCCTCAGCCAAGAGATTTAGGTAATCCGGTCTCTGCGTCTCCTTCTAGGCAGGCATCGCCCGCG
GTTCATACACCGCCAAGGGGAGACCTTGCTGATGAGCGTGAAATTGATCAACTGCGGGAAGCGTCATCGACTCGGCGCAGTAATGAGCGCCAAGTACAGG
TTGAAGGTAGTCTGAAACATGGTAGCGGCCTTAATCCTAATTTTACCTTTGATACATTTGTTGAGGGTAAGTCGAACCAACTTGCCCGAGCGGCATCGCG
TCAGGTTTCTGAAAATCCTGGTGGTGCTTACAACCCACTGTTTTTGTACGGTGGCGTTGGCTTAGGTAAAACACACTTAATGCACGCGGTAGGTAACGCG
TTGGCAACTCGCCGTGAGAATGCCCGTGTGGTTTATCTACACTCTGAGCGTTTTGTGGCAGATATGGTGAAAGCCCTTCAGCTAAACGCGATTAACGACT
TCAAGCGTTTTTATCGCAGCGTTGATGCGTTACTTATTGATGATATTCAATTTTTTGCTGGTAAAGAGCGTTCGCAAGAAGAGTTTTTCCACACCTTTAA
TGCGTTGTTAGAGGGTGGGCAGCAAATGATTCTTACCTCTGACCGCTACCCAAAAGAAATCAGCGGTGTAGAAGAACGTCTAAAATCACGCTTTGGCTGG
GGTCTGACAGTAGCTATTGAGCCACCGGAGCTTGAAACCCGCGTAGCTATTTTGATGAAAAAAGCAGATCAGGCGAAAGTTGATCTTCCACATGACGCTG

>Halomonas andesensis DSM 19434 [NZ_RZHG01000019.1:c62130..62629], <mnmG oriC2 <COG1126 (glnQ), 578 bp

AGACCGTTAATTGAACGAACCACTTGCAGCTGAACATCGAAAGGCAACGACGTTGAGATTCCGTTGGGATACAGCTCATGGGTATCCAGCCCTTCAGGTT
CAATAAACACCTGGTGGCTCGATTTGTCGGCAAAGCGGTGAACTTTGTCCTCAATCGATGGGCAGTATCGTGGACCAACGCCTTCGATAACACCGGAATA
CATCGGTGAACGGTCAAGGTTTGCCATGATGATTTCATGGGTGCGCTCGTTGGTATGCGCAATGTGACAGCTTACCTGCTGAGGATGCATGTCCCGTGAG
CCCAGGTATGACATCACCGGTGTTGGCGTATCACCTGGCTGCTCTTCTAACACTGAGAAATCAACGGTTTTAGCATCAATCCGCGGGGGGGTGCCGGTTT
TTAACCGGTCGACACGAAACGGTAATGCACGTAAGCGTTCGGCTAAGGCGTTGGAAGGTGGATCGCCTGCGCGTCCGCCGCGGCTTTGATCTAATCCGAT
GTGGATAACGCCACCCAAAAACGTACCGGTTGATAGCACAACGGATTCCGCATGGAAGCGTATGCCTGTTTCGGTCACGACGCCTCGAACGGTGTTGTTA
TCCACAATCAGGTCGCCAGCCGCTTGCTGAAATATCGTCAAGTTCGGCTGGTTTTCTAACATGCTACGAATAGCCGATTTGTAACGAATTCTATCTGCTT
GAGCACGTGTTGCTCGAACGGCTGGGCCCTTACGAGCGTTCAAAACTCGAAACTGAATGCCGCCTAGGTCCGTGGCTAGCCCCATTGCACCACCGAGTGC
ATCAATTTCTTTTACCAAGTGGCTTTTGCCAATCCCACCAATTGCTGGATTGCACGACATTTGGCCGAGCGTTTCAAAATTGTGGGTTAGCAACAGGGTT
TGACAACCCATACGAGCAGAGGCCAATGCGGCTTCAGTTCCCGCATGGCCACCGCCGATGACAATCACGTCAAAGCGGTCGGGATAGTTCAAGAGGCACC

TCGTCTTGCGCCGTTTGGCACGGATGAAAATGGGTCAGCGTGACTTCACTGATTGAGTAAGCCGGCCAGTATAAACCTCCTGTGGGTAACCGTCAGGTTT
TTCCACACCCCAAACTTATTACTCATGCATTATTAATAAAAAAATAAAAATATATAAGAGAAGTTCTATTGATGTTGTAATTATTAGAGTGGACAAAAAC
**TGTTGATAA**CCCTATTAATACTAATTTTATCAATGATTTAAATTGTTAGCTTTACATGATTAAACGGCGGAGTACCTGCGTAGAAGCGGTGTTGCTCCT
GTGGATGAAAGTACGACTTACACACAGGCACGATAAACACCGGGTTATCCACCGACCCATACCGAGTTTGTTACCGCACCTGTGAACAAGTGAGATTGGA
ATCTCAAAAAGTCCTTATTAGCGCTTATGGCAGGTGTCTTAGAACGATAGTGTGCTGCGTGTAAAAAATAGTATCCACAGGCAAAAAAAATGGCCAGTCG

CGATGACAGGCCATTTAATGAGGTAAAAATTGGCAAAAAAGCCAGATTTAAAACCAGATTTACGCAGACATTAGTGTTTCAAAACGCGTGCAAGGAAGGA
TTGTGTACGCTCATGTTGAGGTGTATCGAACAGTTGTTCAGGAGGTCCTTGTTCAGCAATTTCGCCTTGATGAATGAAAATAACGCGATCTGCCACTTCA
CGGGCAAAGCCCATTTCATGCGTCACAATCACCATGGTCATGCCTTCTTTAGCAAGCTCACGCATGGCGTCCAGCACTTCACCAATCATTTCAGGGTCAA
GCGCCGAGGTAGGCTCGTCAAACAGCATAAGGCGGGGTTCCATAGCCAACGCCCGTGCTAACGCTACTCGTTGCTGCTGCCCGCCCGAGAGCTGACTCGG
ATACTTGTCTGCTTGGTCGGAAATACCAACACGCTCTAGCAGTTTGTCAGCCGTTTCCTCGGCATCCTGGCGGCTCCATCCCCGTACTTTCATAGGCGCG
AGAGTGACGTTGTCGCGCACGCTTAAATGGGGAAAAAGGTTGAACTGCTGAAACACCATACCAACTTCCGTACGGATGGTTTGCAGCGCTTTACTGCTTT
TACCGTTAGGCAATAATTCGTTGCCATCAACGTCAAGGGAACCTGATTGAAACTCTTCTAAGCCATTGATACAGCGTATAAGGGTTGATTTACCAGAGCC
GCTGGCACCGATGACGACGACGACTTCACCGGGGATAATCTCAAGATCAATATTATTGAGCACGTGCAGGCTGCCAAAATGCTTATTGAGCTGCTGCATA
CGCACAATTGGCGATTGTGTAGAGGTCATTGCACGTTCCTTATTGTCCTGCGAGGCCTCTGCGCTCGATTTGGCGCAGAATCAAAGAGAGGGTCCAGGTA
ATGACAAGATACATAAGTGCCACCATGAAATAGACCTCTAGTGCGGTGAAGGTGGTCGCAATATAAATTTGCCCTTGGCGAACCAATTCACCGACCCCGA

>Halomonas andesensis DSM 19434 [NZ_RZHG01000019.1:c39141..39194], dnaA> ig dnaN>, 54 bp

WebSkew GCmin pos. 609738 (GCmin<>oriC1 21.2 kb; GCmin<>oriC2 0.3 kb; oriC2<>oriC1 21.5 kb)

>Halomonas zhaodongensis NEAU-ST10-25 [NZ_JACCDD010000002.1:630709..631208], <rpmH oriC1 dnaA>, 534 bp

AGGACATCCGTGGTGACCGCAATAAAATCTCGGCTTGCGGTGTCACTCTCGGACGCATCTGGAATGCCTTCGCTGATGGCATTTTGTGGTGCTGATGCAC
TGGGTACAGCGAAGTCGCCGTCATTGCTAGTATCGCTGACGGGGGCTCCATTGTTACGTTGAGTGATCTCAGGCGTCGAATCATAGGTTGTCTGCCCGTA
ATCCTGATTCCACTGCACAACCAATAAGTAGGCAAGTAATGCTAGTGGAATCAGTAATAAAAGTCGTTTTACGTCCATGAGGGTTCTGCCCGATGGGTGG
TGAACATGCTAATGGCGCAGTGTCATTTACTCGACCCAAGACACGCGAAAGCCTGCCAAACGGCAGGCCGAGGTCGAGCGGCGAGGTGTGTCGTCAAGCG
TCACGTGTTACGCCATTAGGCGGAGAAACGGCGACCGCCTTTGCGTCGTGCTGAAGTCGTTTCCACATGCCGTGTAACTGGCGAAAAAGCGTTTCGTTGT
CGATATCCTGTACGCCACGTTTAGCCAATATCACGATATCCACAGAGGGTAGGTGATCCTGGCGGTGACGAACGGATTCCCGAACAAGTCGCTTAAAGCG
GTTGCGATCAACAGCAAGCTTCACATTTTTTTTGCTGACCACCAGTCCAACGCGAGGGTGCCCGCGGGTAGTCAGACGCGCTAGCGCCATCATACCTTTA
CCATGAACTTTCAGGTCTGCCTGTTCAAACACGTGACTATAGTCCCCGGCATTTAACAAACGTAAGCACCGGGGAAAGCCTTGATGCAGCACACGCAATC
CGAGATCAGGCGCTCAGACGCTTGCGGCCTTTGGCACGGCGGCGTGCGATAACGGCACGGCCGTTTTTGGTTGCCATACGAGCACGGAAGCCATGCGCGC
GCTTGCGCTTGAGAACGCTAGGTTGAAAAGTGCGTTTCATAACTCGTTATTCCCACGTGGTTTAGGACGGAAATTGAATTCCAAAAGGCCCGGTCCAGTG

AGGAACTAGCCGGGGAGTTTCAATTCAAGACCGGAAATTGTAGAGACTTCCGGCGCTAGGTGCAATTTTTCCTTGCATAAAAAGCGTGATCAAGGCAATG
CCGGAGGGCAATGGGTCGTTTTCACGTTAACCAACCATATCACTACTATTACTAATTAAGTTATTTAAAACCAACTGTTATTAATAGTGGGGATACTAAT
**TGTGGATAA**GCGATTTTAAATGATTAATATCAAGATATTGAAGATATAAAAAATCTGTGGATAAGCAACGAATAAGCCTTAAATAAGCTGTTGATTAGGT
GTGATCAACAGGCAAGAACTTGGCTCGGAGTAGAGTTGTCCTCAAGCTACCAGCGATTTTTCCTTAGCCCTTACGAGCAGTTGTCCACAGGGTAAAAAGA
GGCCTGAGTGCGTGTGGATAACCGGCTCGTTGGGCGCTACAATGGTCGGCCGTTTTCAACTGATGGTGAGATGAGTGTCACTCGCGCTTTGGCAACAGTG

CCTGGACTACCTGCAGGACGAACTGAATTCACAGCAGTTCAATACCTGGATACGTCCGCTGCAGGCAGAAGAAGGAGAGACCAATGAGCTTCGCTTATTG
GCGCCGAACCGCTTTGTGCGCGACTGGGTGAGTGATAAGTATGCAAAGCGTATTAGTGAACTAATGCGTGAGCTTGCGCCGTCTAAACCTCCTAAAGTTA
GTTTAACAGTGGGGAGTCGCCAGACGGCTGCGCCCGCCCCGCAGCCAAGAGAATTAGGGAACCCCATTTCAGCGACTCCGGCTCGACCGTCAGCGCCAGC
GGTTCATACATCATCACGTCGTGATAATAGCGACGAGCGTGATATAGATAGGCTGCGGGAAGATAATTCTGCAAACCGTCGTGGCAATGAACGCCAAGTT
CAAGTTGAAGGTAGCTTGAAACACAACAGTGGTCTCAATCCTAATTTTACCTTCGATACCTTTGTTGAAGGTAAATCGAACCAACTTGCTCGTGCGGCAT
CACGCCAGGTATCTGAAAACCCTGGTGGCGCATATAACCCACTGTTTTTATACGGTGGTGTTGGCTTGGGTAAAACCCACTTGATGCACGCAGTAGGCAA
CGCGTTGGCTGGCCGCCGTGAGAATGCCCGTGTGGTGTATCTACACTCTGAGCGTTTTGTCGCAGACATGGTGAAAGCACTCCAGCTAAACGCGATTAAC
GACTTCAAGCGCTTTTACCGCAGTGTGGATGCTCTACTGATTGATGATATTCAGTTTTTTGCTGGTAAAGAGCGCTCCCAGGAGGAGTTTTTCCACACCT
TTAATGCACTGTTAGAGGGCGGTCAGCAGATGATCCTGACATCTGATCGTTATCCCAAAGAGATCAGTGGTGTTGAGGAGCGTTTGAAGTCAAGGTTTGG
CTGGGGTCTGACCGTGGCTATTGAGCCACCGGAGCTTGAAACCCGTGTCGCTATTTTGATGAAAAAAGCAGACCAGGCGAAAGTAGATCTCCCACATGAC

>Halomonas zhaodongensis NEAU-ST10-25 [NZ_JACCDD010000002.1:609156..609655], <mnmG oriC2 <COG1126 (glnQ), 564 bp

AAGCCCTTTAATTGAGCGAACTACCTGCAGTTGTATATCAAAAGGCAGTGAGGTTGAAATCCCGTTGGGATACAGCTCATGGGTATCTAGCCCTTCGGGT
TCGATAAACACTTGATGGCTGGCTTTATCGGCAAAGCGGTGGACTTTATCTTCAATCGATGGGCAATAGCGCGGGCCAACCCCTTCGATAACACCAGAAT
ACATGGGTGAACGGTCAAGGTTCGCCATGATAATTTCATGGGTGCGCTCATTGGTGTGAGCAATATGACAGCTCACCTGCTGAGGATGCATATTCCGTGA
ACCAAGATACGACATGACTGGAGTCGGCGTATCGCCAGGCTGTTCTTCCAACGGTGCAAAATCAACCGTTTTGGCATCAATGCGTGGTGGAGTCCCCGTT
TTTAATCGATCCACGCGAAACGGCAGTGCACGAAGCCGATCTGCTAGCGCATTGGATGGCGGATCACCGGCACGGCCACCGCGGCTCTGATCCAGCCCAA
TATGGATAACCCCACCTAAAAATGTGCCTGTGGACAAGACCACAGAGTCCGCATGAAAGCGAATACCAGTTTCCGTTACAACACCGCGAACAGTGTCGTT
ATCCACAATCAGGTCACCCGCCGCCTGCTGAAAAATGGTCAGGTTAGGTTGATTTTCCAACATTCCGCGAATGGCGGCTTTGTAACGAATGCGGTCAGCC
TGAGCACGTGTGGCACGAACTGCTGGGCCTTTGCGTGCATTGAGTACACGAAACTGGATGCCGCCTAAGTCCGTGGCTAGCCCCATTGCACCACCGAGTG
CATCGATTTCTTTCACCAAGTGGCTTTTGCCGATCCCACCAATCGCTGGATTACACGACATTTGGCCGAGTGTTTCGATGTTGTGGGTCAATAACAGGGT
TTGACAGCCCATACGAGCAGAGGCCAATGCGGCTTCGGTTCCCGCATGGCCACCGCCGATGACAATAACGTCAAAGCGGTCGGGATAGTTCAAGAGACAC

CTCGTCTTGCGCCTTTTGGCACGGATGATGTGAGTCAGCGTATCGGCGCTGATTTCGTAAGCGAGCCAGTATAAACCTCCTGTGGGTAACCGTCAGGTTT
TTCCACACCCAAAATAACCTCGTCACTTATTATTAATAACAAAATAAAAATATATAAGAGAAGTTCTATTGATGTTGTTACTACTAGTGTGGACAAATTC
**TGTTGATAA**CTGAAAAAAACTCAAATTCTTCATGATGTTAAATTGTTAGCTTTTACATGATTAAACGGCGGAGTGCCTGCGTAGAAGTCGTGTTGAGCCT
GTGGATGAAAGTGGTCTTTATCCACAGGCGCAGCTATTGACGTCTTATCCACCGGCTCATACCGAGTTTGTTACCGCACCTGTGAACATTTCTGCTGACT
TAGTATAAAAGCCATTAACAGTGCTCATAGCAGGGATCACAGAGGAGTACCTGGTGACAGAAAAAAATAGTATCCACAGGCAAAAAAAATGACCTGTCAC

TACGACAGGCCATTTTATGGGCTAAAAAATGAAGAAAAAGAGGCTGGATAACCTACATCAGTGCTTTAAAACACGCGCTAGGAAGGATTGGGTTCGCTCA
TGTTGAGGCGTATCAAACAGCTGCTCAGGTGGTCCTTGTTCGGTAATTTCACCTTTATGGATAAAGATAACGCGGTCAGCCACTTCTCTGGCAAAGCCCA
TTTCGTGGGTGACAATCACCATGGTCATACCTTCTTTGACGAGTTCACGCATAGCATCTAGCACTTCGCCGATCATTTCGGGATCAAGTGCTGACGTAGG
CTCGTCGAACAGCATAAGGCGGGGTTCCATCGCCAGTGCGCGCGCCAGCGCTACGCGCTGTTGTTGTCCACCTGAGAGTTGGCTTGGGTACTTATCCGCC
TGGTCGGAAATCCCAACACGCTTTAATAGGCGCTTGGCAGTTTCTTCAGCATCCTGACGGTTCCATCCACGCACTTTCATTGGGGCAAGTGTGATGTTGT
CGAGCACGCTTAAGTGGGGAAACAGGTTGAACTGCTGAAACACCATGCCCACTTCTGTCCGAATAGTTTGCAGTGCTTTGCTGCTTTTACCATTAGGCAA
TAGCTCGTTGCCATCGACATCTAACGTGCCTGACTGGAACTCTTCCAGGCCGTTAATGCAACGGATAAGGGTCGACTTACCGGAGCCACTGGCACCAATC
ACCACCACCACTTCGCCAGGGACAATTTCAAGGTCAATATTATTCAGCACGTGCAGGCTACCAAAGTGCTTATTGAGCTGCTGCATACGCACAATGGGTT
GTGTGACATTGTTCATTGCACTTGTCCTTATTGTCCAACGAGGCCTCTGCGCTCTAGTTGGCGCAGTAGGAGAGAGAGGGTCCAGGTGATCGCAAGGTAC
ATCAGCGCTACCATGAAATAGACCTCTAGGGCTGTAAACGTGGTGGCAATATAAATCTGCCCCTGGCGCACTAACTCACCCACCCCGATGACGGAAAAGA

>Halomonas zhaodongensis NEAU-ST10-25 [NZ_JACCDD010000002.1:632653..632708], dnaA> ig dnaN>, 56 bp

< page 1 | #077 | Halomonas nigrificans MBT G8648

WebSkew GCmin pos. 65250 (GCmin<>oriC1 22.1 kb; GCmin<>oriC2 0.0 kb; oriC2<>oriC1 22.1 kb)

>Halomonas nigrificans MBT G8648 [NZ_NWUX01000004.1:c42893..43392], <rpmH oriC1 dnaA>, 534 bp

ACGTCAGTGGTAACGGCAATGAAGTCACGGCTCGCTGTTTCGCTTTCCGCGGCATCCTGCAGGCCCTCGCCAACAGCGCTCTGCGGAGCGGATGTGCTTG
GAACTGCAAAGTCGTCGCCATTGGAGACGTCGCTTGCTGGGGTTCCATTGCTGCTTTGAGTGATTTCAGGCGTCGAATCGTAGGTTGTCTGCCCGTAATC
CTGATTCCACTGCACAACCAGCAGATAGGCAAGTACTGCCAGTGGAACCAGTAATAGAAGTCGTTTTACGTCCATGAGGGTTCTGCCCGATGGGTGGTGA
ACATGCTAATGGCGCAGTGTCATTTACTCGACCCATGACACACGAAAGCCTGCCAGACGGCAGGCCGAGGTCGAGCGGCGATGTGTGTCGTCAAGCGTCA
CGTGTTACGCCATCAAGCGGTGAAGGTGGCGATATCGCCTGTGCATCTCGCTGGAGTCGCTTCCACATGCCGTTTAACTGGCGGTAAAGCGTTTCGTTGT
CGATATCCTGAACACCTCTTTTAGCCAGAATCACAATATCCACAGGAGGGAGGTGATCTTGGCGAAGCCGAACAGATTCTCTAACAAGACGCTTAAAGCG
ATTGCGATCAACAGCGCGCTTCACATTCTTCTTGCTGACTATCAGTCCGACGCGAGGGTGGCCCTGAGTACTTAAGCGCGCTAGCGCCATCATACCTTTG
CCATGGACTTTCAAGTCGGCCTGTTCAAACACGTGACTAAAATCCCCAGCGTTTAGCAAACGTAAGCACCGGGGAAAGCCTTGATGCGGCACACGCAATC
CGAGATCAGGCGCTCAGACGCTTGCGGCCTTTGGCACGGCGGCGTGCGATAACGGCACGGCCGTTTTTGGTTGCCATACGAGCACGGAAGCCATGCGCGC
GCTTGCGCTTGAGAACGCTGGGTTGAAAAGTGCGTTTCATAACTCGTTATTCCCACGTGGTTTAGGACGGAAAGTGAATTCCAAAAGACCCGGACCAGTG

AGGAACTAGCCGGGTAGATTCAATTCAAGACCGGAAATTGTAGAGACTTCCTGCGTCGGGTGCAATTTTTCCTTGCACAAAAAGCTTCAAAAAGGCAATG
CAGATTTTGCAATTGGAGCTACAGAGTGTCAGAAACCTACCACTATTATTACTAATAGTATTACTTAAAACCTATTGTTATTAATAGTGGGATACTAAT
**TGTGGATAA**GTGGTTTATGCTTTTTATTAACCAATAGCTTAAAGATATAAAAAATCTGTGTGTAAGTCCCGAATAAAGGCCAAATAAGCTGTTAATTAGGT
GTGTTCAATACCCCCGAAAGTTGCCCTTTCTTAAGTTGTCCTCTCCCCCTCCCATACTTTTCCTAGTAGCTTACGATAAGTTGTCCACAGGGGTAAAAGG
GCGTCTAGTGCGTGTGGATAACCGCTCCATTGGGCGCTACAATGGTCGGCCCTTTTCAACTGATGGTGAGATGAGTGTCACTCGCGCTTTGGCAACAGTG

CCTGGACTACCTGCAGGACGAACTGAATTCACAGCAGTTCAATACCTGGATACGTCCGTTGCAGGCAGAAGAAGGAGAGTCCAACGAGCTACGCTTGTTG
GCGCCGAACCGCTTTGTGCGCGACTGGGTGAGCGATAAGTACGCCAAGCGGATCAGCGAGTTGATTCGTGAGCTATCACCCGCAAAACCTCCCAAAGTTA
GCTTAACGGTAGGTAGCCGTCGTATTGCTACCCCAGCTCCACAGCCACGAGAATTAGGCAACCCGGTATCGGCAGCTCCTTCTCAGAGAATGGCCCCTGC
TGTGCATACCCCCCCACGAGGTGATATTGCTGATGAGCGGGAAATTGGCCAGTTAAGAGATAGAGAAGAAGCACCTAGTCGTCGTAATAACGAGCGACAA
GTACAGGTTGAAGGTAGCCTGAAGCATGGTAGCGGCCTTAATCCTAATTTTACCTTCGATACTTTTGTTGAAGGTAAATCGAACCAACTTGCTCGCGCCG
CATCACGCCAGGTGTCAGAAAATCCCGGTGGCGCTTATAACCCACTGTTCTTATACGGTGGAGTTGGCTTGGGTAAAACCCACTTAATGCATGCAGTAGG
TAACGCTCTGGCGGGCCGCCGAGAAAATGCTCGAGTAGTTTATTTACACTCTGAGCGTTTTGTGGCTGACATGGTAAAAGCGCTGCAGCTAAACGCAATT
AATGATTTTAAGCGGTTTTACCGTAGTGTTGATGCACTGCTTATTGATGACATACAGTTTTTTGCGGGTAAAGAGCGCTCACAGGAAGAGTTTTTCCATA
CGTTTAACGCGCTGTTAGAGGGAGGGCAGCAAATGATCCTCACCTCTGATCGTTATCCTAAAGAGATTAGTGGTGTTGAAGAGCGTTTGAAGTCACGCTT
TGGGTGGGGACTTACCGTCGCTATTGAACCACCCGAGCTTGAAACTCGTGTGGCGATTTTGATGAAAAAAGCTGATCAGGCGAAAGTTGATTTGCCTCAT

>Halomonas nigrificans MBT G8648 [NZ_NWUX01000004.1:c64971..65470], <mnmG oriC2 <COG1126 (glnQ), 554 bp

CAAGTCCTTTAATCGAGCGCACCACCTGCAACTGCACATCAAAGGGCAGTGAGGTTGAAATACCATTAGGGTACAACTCGTGGGTATCTAGCCCTTCTGG
CTCGATAAATACCTGATGGCTTGATTTATCAGCAAAGCGATGCACTTTGTCTTCGATAGATGGGCAGTAGCGCGGGCCAACCCCTTCAATCACCCCTGAA
TACATTGGTGAACGATCTAGATTACCAAGAATAATTTCATGGGTGTGCTCATTGGTATGGGCAATATGGCAGCTCACCTGTTGAGGGTGCATATCCCGTG
AGCCAAGGTAGGACATCACTGGCGTTGGCGTATCACCAGGCTGCTCTTCCAACTGGGAAAAATCGACAGTCTTGGCATCAATACGTGGTGGTGTGCCAGT
TTTAAGGCGATCAACGCGAAATGGAAGTGCCCTTAGGCGTTCTGCTAGTGCATTTGAGGGCGGGTCACCGGCACGACCACCGCGGCTTTGATCCAACCCT
ATGTGGATAACACCACCTAAAAATGTACCTGTCGAAAGTACCACAGCTTCTGCATGAAAACGGATACCAGTTTCTGTAACAACACCACAAACAGCGTTGT
TATCCACAATCAAATCACCAGCTGCCTGCTGAAAGATTGTCAAATTTGGCTGGTTTTCAAGCATCCCGCGAATTGCTGCTTTGTAGCGAATGCGATCAGC
TTGGGCACGTGTTGCCCGAACGGCTGGGCCTTTGCGGGCATTCAACACGCGAAACTGGATGCCGCCTAAATCCGTGGCTAACCCCATTGCACCACCAAGT
GCATCAATTTCTTTCACTAGGTGGCTTTTGCCAATCCCACCAATCGCTGGATTGCACGACATTTGGCCGAGCGTCTCGATGTTGTGAGTTAGCAATAGAG
TTTGACAGCCCATACGAGCAGAGGCCAGTGCGGCTTCAGTTCCCGCATGGCCACCGCCGATGACAATCACGTCAAAGCGGTCGGGGTAGTTCAAGAGACA

CCTCGTCTTGCGCCTTTTGGCACGGATGATGTGGTCAGTGTTTAGCCACTGATTGAATAGGTCGGCAAGTATAAACCTCCTGTGGGTAACCGTCAGGTTT
TTCCACACCCCAAGCAATCAGCTCACCTATTATTAATAACAAAATAAAAATAAATAAGAGAAGTTCTGTTAATGCTGTAACTAATGGTGTGGACAAAATC
**TGGGGATAA**CCCTATAAATAAATTTAAAATCAAATGATTACGTTGTTGACCTTTCAGTGATCAAGCAGCGTAGTGCCTGAGTAGAAGCGGTGTTAGAGCT
GTGGATGAAAAGGCGACTTATGCACAAGGCTATCGAGTAACGGGTTATCCACCGGCTCATACCGACTTTGTTACCGCATCTGTGAACATCTTGCCATAAC
GCCTCTAGAAGTGCTTAAAGGTCGGCAGAGTAGGAGCTACAGAGCACATGAGAGGTCCCGTAAAAAAATAGTATCCACAGGCAAAAAAAATGGCCTGTCG

CTTTGACAGGCCATATAACAGACATTAATTAGCAGTTTTATGCGTGTTTTAATGTTTTAACACGCGAGCCAAGAATGAACGGGTGCGCTCATGTTGAGGC
GCATCGAACAGCTGTTCGGGGGGACCCTGTTCAGTGATTTCTCCTTTGTGGATGAAGATAACGCGGTCAGCCACTTCACGGGCAAATCCCATTTCATGGG
TAACAATCACCATGGTCATGCCTTCTTTAGCAAGTTCGCGCATGGCATCAAGCACTTCACCGATCATTTCGGGATCGAGAGCTGAGGTAGGCTCATCAAA
CAGCATCAATCGTGGTTCCATTGCCAGTGCTCGTGCCAATGCAACCCGCTGCTGCTGGCCACCAGAAAGCTGACTTGGATACTTATATGCTTGGTCGGAG
ATACCCACCCTCGTTAGTAGGCGTTGAGCGGTATCTTCAGAGTCTTTACGGTTCCAGCCGCGCACTTTCATAGGTGCAAGGGTGATATTGTCGATTACGC
TTAGGTGAGGAAACAGGTTGAATTGCTGGAACACCATACCTACTTCAGTGCGAATAGTTTGTAGGGCTTTGCCGCTTTTTCCATTAGGTAGCAGTTCATT
GCCATCCACATCCAGTGAACCTGATTGAAATTCTTCAAGGCCGTTAATACAGCGAATCAGCGTCGATTTGCCAGAGCCACTGGCACCAATCACCACCACT
ACTTCGCCTGGGATAATTTCAAGGTCAATATTATTGAGTACGTGCAGGCTGCCAAAATGCTTGTTGAGCTTCTGCATACGCACAATAGGTTGCGTTGCGT
TATTCATTGCACTTGTCCTTATTGTCCGACGAGGCCTCTGCGCTCTAATTGACGCAGTAAGATGGAGAGAGTCCAGGTAATGGCGAGATACATAAGTGCG
ACCATGAAGTAGACCTCTAGGGCAGTGAAGGTGGTGGCGATATAGATCTGACCTTGGCGTACCAGCTCACCTACTCCTATGACGGAAAACAGCGATGTGT

>Halomonas nigrificans MBT G8648 [NZ_NWUX01000004.1:c41389..41445], dnaA> ig dnaN>, 57 bp

WebSkew GCmin pos. 40992 (GCmin<>oriC1 30.5 kb; GCmin<>oriC2 8.9 kb; oriC2<>oriC1 21.6 kb)

>Halomonas malpeensis YU-PRIM-29 [NZ_WHVL01000008.1:71261..71760], <rpmH oriC1 dnaA>, 532 bp

AGCACGTCGGTGGTGACGGCAATGAAGTCACGGCTGGCAGATTCGCTGCCGTTGTCCGTGGGAATCGCCTCGTTGGCGCTGCCCTGGGGGGCATTGGAAG
TGGGAACGGCCAAGTCGTTACCGCCGCCGTCGTCGACGCCTTGCTCGCTGCTCTGGCGCTGGGTGATCTCCGGCGTCGAGTCGAAGCTGGTCTGCCCGTA
GTCCTGATTCCACTGAACAACCAGAAGATAGGCCAGTACGGCCAGTGGAATCAGCAATAAAAGTCGTTTTACGTCCATGAGGGTTCTGCCCGATGGGTGG
TGAACATGCTAATGGCGCAGTGTCATTTCTCGCCCCGAGACACACGAAAGCCTGCCAGGAGGCAGGCCGAGGCCGAGCGGTAAGGCGTGCGATCAAGCGC
CGTGCGTAACATTAGCAAGCGAGGGAGGCGGCGAGAGTGCAAGCGCATCGCGCGAAAGTCGCTTCCACATGCCGTGCAGCTGGCGGGCGAGCGTTTCATT
GTCAATATCCTGCACACCGCGCTTGGCCAGGATCACAATGTCCATGGAGGGCAAACGGTGCTGGCGAAGCCGGACGGATTCTCTGACAAGACGCTTGAAG
CGATTACGATCGACGGCGAGCTTGACATTCTTCTTGCTGACCACCAGTCCAACGCGAGGATGCCCCCGGGTACTGAGACGCGCGAGCGCCATCATACCTT
TGCCGTGGACCTTCAGGTCTGCCTGCTCGAATACGTAGCTGAAATCCCCGGCATTCAGTAAGCGCAAATCCCGGGGAAAGCCTTGGTGCGGCACACGCAA
TCTGAAATCAGGCGCTCAGACGCTTGCGGCCCTTGGCGCGACGACGGTTGATGACGGCGCGGCCATTCTTGGTGGCCATGCGAGCACGGAAGCCATGCGC
GCGCTTGCGCTTGAGAACGCTGGGTTGAAAAGTGCGTTTCATAACTGGTTATTCCCACGTGGTTTAGGACGAAAGTGAATTCCAAACGACCCGGTCCAGT

GGAGGAACTAGCCGGGGAGGTTCAATTCAAGACCGGAAATTGTAGAGACTTCGCGGTCTGGGTGCAATTTTTCCATGCATAAAAAGCGCTAAATATGCTC
TAACGCCCCTTAACGTGACCTCAAATCGAGGTTTTGACGCTACTATCATTACTAGTGAAGTTATTTAAAACCCATTGTTATTAATAGTGGGATACTAAT
**TGTGGATAA**GTGGTTTTTGAATTTTATTAACAACTGCTTACAAGCATAAAAAATGTGTGCATAAGCCGTTGGAAAACGCTTTGGAAGATGTGCGTGGGGT
GTGCATGGATCCTTGAAAACGACGATTTTGCCTTTCTGTCCTCATGTCGTGCCCGGCTTCTCCCAATGGCTCATGAGGAGTTGTCCACAGCCCGTAATGA
GCCCATTTCGCGTGTGGATAACCGGCGCCATGGGCGGTACAATAGTCGGCCGTTTTCAACTGATGGTGAGATGAGTGTCACTCGCGCTTTGGCAACAGTG

CCTGGACTATCTGCAGGACGAACTAAGCTCCCAGCAGTTCAACACCTGGATTCGCCCCTTGCAGGCTGAAGAGGGGGAGGCCAACGAGCTCTGCTTGCTG
GCGCCCAACCGCTTCGTGCGCGACTGGGTGAGCGACAAGTACGCCAAGCGCATCAGCGAGCTGATTCGTGAGCTCGCGCCCTCCAAACCCCCCAAGGTGA
GCCTGAGCGTGGGCAGCCGCCGCACGGTGGCCGCCACGCCCCAGCCGCGCGATCTGGGTCATCCGGTGTCCGCCGCGCCGGAGCGGCCCGCCGCTCCCGC
CGTGCATACCCCGCCTCGCGGCGATATCGCCGATGAGCGCGAGATCGACCGGCTGCGCGAGGAACCCGCCGGGCGACGCGGCAGCGAGCGCCAGGTACAG
GTCGAGGGCAGTCTTAAGCACGTCAGTGGCCTGAACCCCAATTTCACCTTCGACAAGTTCGTCGAGGGCAAGTCGAACCAGCTGGCCCGCGCCGCTTCGC
GCCAGGTATCGGAAAATCCTGGCGGTGCCTACAACCCGCTGTTTCTGTATGGCGGCGTGGGGCTTGGCAAGACCCACCTGATGCACGCCGTGGGCAACGC
TCTGGCCGGACGGCGCGAGAACGCCCGGGTGGTCTACCTGCACTCCGAGCGCTTCGTCGCCGACATGGTCAAGGCGCTGCAGCTCAACGCCATCAACGAT
TTCAAGCGCTTTTATCGCAGCGTCGATGCGCTATTGATCGACGACATTCAGTTCTTCGCCGGCAAGGAGCGCTCCCAGGAGGAGTTCTTCCATACCTTCA
ATGCCTTGCTGGAAGGCGGTCAGCAGATGATTCTCACCTCGGATCGTTATCCGAAAGAGATCAACGGGGTGGAGGAGCGTCTCAAGTCGCGCTTCGGCTG
GGGGCTGACGGTGGCCATCGAGCCGCCGGAGCTCGAGACCCGGGTGGCGATTCTGATGAAGAAGGCCGATCAGGCGAGCGTCGATCTGCCCCACGATGCG

>Halomonas malpeensis YU-PRIM-29 [NZ_WHVL01000008.1:49657..50156], <mnmG oriC2 <COG1126 (glnQ), 565 bp

GCCCCTTGATCGTGCGCACCACCTGCAGTTGCACGTCAAAAGGCAGCGAGGTGGAGATGCCGTTGGGGTAGAGCTCATGCGTGTCGAGACCTTCGGGCTC
GATGAACACCTGGTGGCTCGACTTGTCGGCGAAGCGGTGGACCTTGTCCTCGATCGACGGACAGTAGCGCGGCCCCACGCCTTCGATGACACCGGAGTAC
ATCGGCGAGCGATCCAGGTTCGCCATGATGATCTCGTGGGTGCGCTCGTTGGTGTGCGCGATGTGGCAGCTCACCTGCTGCGGGTGCATTTCCCGGGTAC
CGAGATAGGACATCACTGGTGTCGGGGTGTCGCCGGGCTGCTCCTCGAGCTGGCTGAAATCGACGCTCTTGGCGTCGATGCGCGGCGGCGTGCCGGTCTT
GAGTCGGTCGACGCGAAACGGCAGCGCCCGTAGGCGTTCGGCCAGGGCGTTCGAGGGTGGATCCCCCGCGCGGCCACCCCGGCTCTTGTCGAGCCCGATG
TGGATCACGCCGCCCAGGAAAGTGCCGGTCGACAGCACCACGGTGTCGCTCAGAAAGCGGATGCCAGTTTCGGTGACGACCCCACGCACGGTGTCGTTCT
CCACGATAAGATCGCCCGCCGCCTGCTGGAACAGCGTCAGGTTCGGCTGGTTTTCCAGCATGCCACGAATGGCGGCCTTGTAGCGAATGCGATCCGCCTG
GGCACGGGTGGCGCGCACTGCCGGCCCCTTGCGGGCGTTCAACACGCGAAACTGAATACCGCCCAGGTCAGTGGCAAGCCCCATGGCGCCGCCCAGCGCA
TCGATCTCCTTGACCAGGTGGCTCTTGCCGATGCCACCGATGGCGGGATTGCAGGACATCTGCCCCAGGGTCTCGATGTTGTGGGTGAGCAGCAGGGTTT
GACAACCCATGCGAGCAGAGGCCAGCGCGGCTTCGGTTCCCGCATGGCCACCGCCGATGACAATGACGTCAAAGCGGTCGGGGTAATTCAATTGACACCT

CAACTTGGCCCTTCAATGAGCGCCGAATTCGTCGAATTAACGTGAGCCTGGGCTCGATAAGCCCGCCAGTATAAACCGCCTGTGCATAACAGTCAGGTTT
TTCCACACCGCTTTCAGACGGGTCATCACTTACTAATAACAAAATAAAAAATATAGAAAGAAGTTCTGTTGATGTTGTAACTAACTGGTGTGGATAAAATC
**TGTGTATAA**CCAACTAAAACTATATTTATCAGGGTAATATATTGTTGACCGATCCGTGGTTAAACGGCGGAGTGCCTGCGTGCAAGCGGTGGCGAGGCT
GTGGATCGAAAGGCCAATTATGCACAGCGTCGCGAAAGCACTGGTTATCCACTGCTGCATACCGGTACTGTCCGCCGGGCTGTGAACAAGCGTGGTGATG
AATGCGAATTTCACCAGAGGCCGCTGTAGCGGGCCTCGTGGAGGGGCGTGAATAAATTCGAAAAGAAAATATCCACAGGCACAAAAAAGGCCTGTCATGA

CGACAGGCCTGGAAAATTAACGGTTTCGAGCGCACACACGCTTGGCTAGACGTGGCTCAATGCTTGAGTACCCGCGCCAGAAACGCCTGGGTACGCTCGT
GCTGGGGAGCGTCGAACAACTGCTCTGGCGCACCCTGCTCGGCGATCTCGCCCTTGTGGATGAAGATCACCCGGTCCGCCACCTCGCGGGCGAAGCCCAT
CTCGTGAGTGACGATCACCATGGTCATGCCCTCCTTGGCGAGTTCGCGCATGGCGTCGAGCACCTCGCCGATCATCTCCGGGTCGAGCGCCGAGGTGGGC
TCGTCGAACAGCATCAGCCGCGGCTCCATGGCGAGTGCCCGGGCCAGCGCCACGCGCTGCTGCTGGCCGCCGGAGAGCTGGCTCGGGTACTTGTTCGCCT
GGTCGGCGATGCCCACTCTGGCCAGTAGACGCTTGGCGGTTTCGATGGCCTGATCGCGGCGCTGTCCGCGCACCTTGATCGGCGCGAGCGTGACGTTGTC
GATCACGCTCAGGTGCGGGAACAGGTTGAACTGCTGAAACACCATGCCGACTTCGGTGCGGATGGTCTGCAGCGCCTTGGTGCTCTTGCCCATGGGCTTC
AAAAGGTTGCCGTCCACCTCGAGCAGGCCGGACTGGAACTCCTCGAGCCCATTGATGCAGCGAATCAGCGTGGACTTGCCCGAACCGCTGGCGCCGATGA
TCACCACCACTTCGCCGGGGGTGATTTCCAGATCGATGTCATTGAGTACGTGCAGGCTGCCGAAGTGCTTGTGGAGCTTTTGCATGCGCACGATGGGGGG
CTTGTTCGTCGTCATGGCACGCTCCTTATTGGTCGGCGAGGTTGGAGCGCTCGAAGTAGCGCAGCAGTAGGGAGAGGGTCCAGGTGATGGCCAGGTACAT
CAGTGCCACCACGAAATAGACCTCGAGAGCAGTAAAGGTGGTAGCGATGTAGATCTGTCCCTGGCGGACCAGCTCGCCCACGCCGATCACCGAGAAGAGC

>Halomonas malpeensis YU-PRIM-29 [NZ_WHVL01000008.1:73202..73266], dnaA> ig dnaN>, 65 bp

WebSkew GCmin pos. 607948 (GCmin<>oriC1 22.5 kb; GCmin<>oriC2 2.8 kb; oriC2<>oriC1 19.7 kb)

>Halomonas populi MC [NZ_RZHE01000015.1:630223..630722], <rpmH oriC1 dnaA>, 534 bp

TACGTCGGTGGTGACCGCGATAAAGTCACGACTTGAGGTGTCACTTTCCGTGTCGCCAGCCAGCCCTTCGCCCACTGCATTGTTCGGTGCAGACGTTGAC
GGGACCGCCAGGTCGTTACCGCTTTCAGCATTGGGTAAGGGGGCATTGCTACTGCTTTGGGTAATTTCAGGCGTCGAGTCGTAATCTGTCGGCCCATAAT
CCTGATTCCACTGAATGACCAGTAGATAGGCAAGTACTGCCAGTGGAATCAGTAATAGAAGTCTTTTTACGTCCATGAGGGTTCTGCCCGCTGGGTGATG
AACATGCTAATGGCGCAGTGTCATTTACTCGACCCATGACACACGAAAGCCTGCCAAGCGGCAGGCCGAGGTCGAGCGGCGAGGTGCGTCGTCAAGTGTC
ACGTGTTACGCCATCAATCGGTGGTGCAACTGATACCGTCTGGGCGTCACGCTGAAGGCGTTTCCACATGCCATGTAGCTGGCGGTTAAGCGTTTCGTTA
TCGATATCCTGCACACCGCGCTTGGCTAGTATCACAATATCCATGGAGGGAAGGTGATGCTGACGAAGTCTGGCGGATTCTCGGACAAGACGCTTAAAGT
GGTTGCGGTCTACAGCGCGCTTTACATTTTTCTTGCTGACCACCAGTCCAATGCGGGGGTATCCCCGGGTACTCATGCGCGCTAGCGCCATCATACCTTT
GCCATGAACTTTAAGATCGGCCTGTTCAAACACGTGACTAAAGTCCCCGGCGTTTAAAAGACGTAAACACCGGGGAAAGCCTTGAGGCGGCACACGCAAT
CTGAAATCAGGCGCTCAGACGCTTGCGGCCTTTCGCACGGCGGCGCGAGATAACGGCACGGCCGTTTTTGGTTGCCATGCGAGCACGGAAGCCATGCGTG
CGCTTGCGTTTGAGAACGCTGGGTTGAAAAGTGCGTTTCATAACTCGTAATTCCCACGTGGTTTAGGACGGATGTGAATCCAAAAGACCCGGTCCAGTGA

GGAAACTAGCCGGGGAGTTTCAATTCAAGACCGGAAATTGTAGAGACTTCCCGCGTCGGGTGCAATTTTTCCTTGCATAAAAACCAACAAAAAGGCAATG
GGCATCGACCCGGTTATATTTCATCGTTCCCAACAGCGACTACTACTATTACTAGTTAAGGTTTTAAATGATGTTGTTATTAATAGTGGGGATACTATT
**TGTGGATAA**GTGATTATTCCTTATTTTTATCAAATAACTAAAAGCCAAAAAACAATGTGCATAAGTGCCGAATAACCGGCCGATAAACTGTTAATTTAAT
GTGAGTAGCTGTATGGAAAAACCCCAATAATGGATCTGTCCCCATTTCTTCAGGAGGTTTTCCCGTGTGCTTATGGAAAGTTGTCCACAGGCACAAATTG
ATAGCCAAACCCGTGTGGATAACCAAGCCCTAGAGCGCTACAATGGTCGGCCGTTTTCAACTGATGGTGAGATGAGTGTCACTCGCGCTTTGGCAACAGT

GCCTGGACTACCTGCAGGACGAATTGAATTCACAACAATTCAATACCTGGATACGTCCCTTGCAGGCAGAAGAAGGAGAGTCCAATGAGCTGCGCTTACT
GGCGCCTAATCGCTTTGTGCGCGACTGGGTGAGTGACAAGTATGCCAAGCGGATCAGTGAGCTGATTCGCGAGCTTGCACCGTCTAAACCACCCAAGGTA
AGCCTGACCGTGGGTAGCCGCCGCCCGGCGGCACCCGCGCCCCAACCACGTGATTTGGGCAACCCGGTTTCCGCCAACACCTCCGCGCCTTCCGTGCATA
CACCGCCGCGTGGTGATATTGCCGATGAGCGTGAGATCGACCAGCTGCGTGAGCGTGAGGAAGCGCCGCGTCGGCGTACCAATGAGCGTCAGGTTCAGGT
CGAAGGCAGCCTCAAGCACGGCAGCGGGCTTAACCCGAATTTTACGTTTGATACTTTTGTTGAAGGTAAATCCAACCAATTGGCTCGGGCTGCTTCACGG
CAGGTTTCGGAGAACCCAGGCGGCGCCTATAACCCCCTGTTTCTCTACGGTGGGGTAGGTCTGGGTAAAACGCACTTGATGCACGCAGTGGGTAATGCGC
TGGCTGGCCGCCGGGAAAATGCCCGGGTGGTTTACCTGCATTCAGAGCGTTTTGTTGCGGATATGGTCAAGGCACTTCAACTTAACGCGATCAACGATTT
CAAACGTTTCTATCGCAGCGTTGACGCCTTACTGATTGATGATATTCAGTTTTTTGCTGGCAAAGAGCGCTCTCAGGAAGAGTTTTTTCACACCTTCAAT
GCGTTATTGGAAGGTGGCCAGCAGATGATTCTCACCTCTGATCGTTACCCCAAGGAAATCAGCGGTGTCGAGGAGCGCCTGAAGTCGCGCTTTGGATGGG
GGTTGACGGTGGCGATCGAGCCGCCGGAGCTTGAGACCCGAGTGGCCATCTTGATGAAGAAAGCCGACCAGGCCAAGGTCGATTTACCCCATGATGCTGC

>Halomonas populi MC [NZ_RZHE01000015.1:610482..610981], <mnmG oriC2 <COG1126 (glnQ), 555 bp

ATCCCTTGATCGAGCGGACTACCTGCAGCTGTACATCGAAAGGCAGCGAGGTGGAGATACCGTTAGGGTAGAGCTCGTGGGTGTCGAGGCCTTCAGGTTC
AATAAATACCTGATGGCTCGATTTATCCGCAAAGCGGTGCACCTTGTCTTCAATTGACGGGCAGTAGCGCGGGCCAATGCCTTCGATGACGCCGGAGTAC
ATCGGCGAGCGGTCAAGGTTGGCCAGAATCAGCTCGTGGGTGCGCTCATTGGTGTGCGCAATATGACAGCTGACTTGACGCGGGTGCATGTCCCGCGAAC
CAAGGTATGACATCACCGGTGTTGGGGTATCGCCTGGTTGTTCTTCAAGCTGGGAAAAATCCACGCTTTTGGCATCGATACGTGGGGGAGTACCGGTCTT
TAACCGGTCGACCCGAAACGGCAATGCACGCAAACGCTCAGCCAGAGCGTTGGAAGGCGGGTCACCCGCGCGGCCACCACGGCTCTGATCCAGGCCGATA
TGGATAACGCCACCTAAAAACGTGCCTGTCGAAAGCACCACGGATTCGGCATAAAAACGAATCCCGGTTTCCGTCACGACACCTCGTACAGTGTCGTTAT
CCACAACCAGATCGCCTGCAGCCTGCTGGAAAATGGTCAGGTTAGGCTGGTTTTCAAGCATGCCTCGGATCGCCGATTTATAGCGGATCCGGTCAGCCTG
GGCACGTGTGGCACGTACTGCCGGGCCCTTGCGAGCGTTCAGCACACGAAACTGAATGCCGCCTAAATCAGTGGCAAGCCCCATCGCGCCGCCTAACGCA
TCGATTTCCTTGACCAGATGACTCTTGCCAATCCCGCCAATCGCGGGATTGCACGACATCTGGCCCAGAGTCTCGATGTTATGGGTGAGCAGCAGGGTTT
GACAGCCCATACGAGCAGAGGCCAATGCGGCTTCAGTTCCCGCATGGCCACCGCCGATGACAATCACGTCAAAGCGGTCGGGGTAATTCAAGAGACACCT

CTTCTTGCGCCAGTTGGCACGGATGAGGGGAGCCGGTGGATAGGCACCCAGATTCAATAAGCCCGCCAGTATAAACCCCCTGTGGGTAACCGTCAGGTTT
TTCCACACCTCAATTCATCAGCTTGTCTCTATTAATAACAATATAAAAATAAAATAAGAGAAGTTCTGTTAATGTTGTAACTACTAGTGTGGACAGTTTC
**TGTGCATAA**CACATTTATTATTATAAATTTCAGAAGCTTAAATTGTTTACAATTATGTGATTAAGCAGCGCAGTGCCTGCGTAGAAGCAGTGCGGCCCCT
GTGGATGAAAAAGCGACTTACCCACAACACTCAAAAATCACCACTTATCCACCTTCCCATACCGGGCTTGTTACCGCACCTGTGAACAAGTCATTGGGTT
AACATCTTATTGGCTTAAAGACCCCTTTGACGGGGGGGCGGGCTTGGTGGAGTAGAAAAAACAAAAAATATTATCCACAGGCAAAAAAATGGCCTGTCAC

AAGGACAGGCCATAAAAATTCAGGGTAAAGCGAGTGTTTTTATCGATTAATGCTTTAACACACGAGCCAAAAATGAACGGGTTTGCTCATTTTGAGGCGC
GCTAAAAATCTGTTCAGGCGTACCCTGTTCGGCAATTACTCCCTTGTGAATAAAGATAACGCGGTCGGCCACTTCGCGGGCAAAGCCCATTTCATGGGTA
ACGATCACCATGGTCATGCCTTCTTTGGCAAGCTCACGCATTGCATCCAGCACTTCACCGATCATCTCGGGGTCAAGCGCCGAGGTAGGCTCGTCAAACA
GCATCAGGCGTGGCTCCATGGCCAAGGCACGTGCCAGCGCGACCCGCTGCTGTTGGCCACCGGAGAGCTGGCTGGGGTACTTATCGGCCTGGTCGGCAAT
ACCTACCCGGTCAAGCAGACGTTTTGCCGTATTGACTGCATCTTCCCGGTTCCAGCCGCGTACTTTCATAGGGGCCAAGGTAATGTTGTCGATAACGCTC
AGGTGAGGAAACAGGTTGAACTGCTGAAATACCATGCCGACTTCGGTACGGATGGTTTGTAGTGCCTTGCTGCTCTTGCCATTGGGTTTTAGCGTATTGC
CATCCACGCTCAGCGTGCCTGATTGAAACTCCTCAAGCCCGTTAATGCAGCGAATAAGGGTTGATTTGCCCGAACCGCTTGCGCCAATGACGACGACGAC
TTCCCCAGGGACAATCTCAAGGTCGATATCATTGAGTACATGCAGGCTGCCAAAGTGCTTATTGAGCTTTTGCATACGCACAATGGGGGGCGTTGTAGGT
GACACAGTATCGGTCATTGTACGTGTCCTTATTGGCCGGCAATGCCGCGGCGCTCAAGCTGGCGCAGAAGAATGGAAAGGGTCCAGGTGATCGCTAGGTA
CATCAGCGCCACCATGAAATAGACCTCAAGCGCGGTAAACGTGGTGGCGATGTAAATCTGTCCCTGGCGTACCAGTTCACCCACGCCAATCACGGAAAAT

>Halomonas populi MC [NZ_RZHE01000015.1:632162..632218], dnaA> ig dnaN>, 57 bp

WebSkew GCmin pos. 61758 (GCmin<>oriC1 17.8 kb; GCmin<>oriC2 1.8 kb; oriC2<>oriC1 19.6 kb)

>Halomonas nanhaiensis JCM 18142 [NZ_RZHE01000015.1:c43725..44224], <rpmH oriC1 dnaA>, 534 bp

TACATTGGTGGTAACCGCGATAAAGTCACGGCTTGAGGTTTCGCTTTCCGTGTCGCCAGCCAGTCCTTCGCCCACTGCATTGTTCGGTGCAGACGTTGAC
GGAACCGCCAGGTCGTTACCGCTTTCTGCATTGGCTAAGGGTGCATTGCTACTGCTTTGGGTAATTTCAGGCGTCGAGTCGTAATCTGTCGGCCCATAAT
CCTGATTCCACTGAATAACCAGTAGATAGGCAAGTACTGCCAGTGGAATCAGTAATAGAAGTCTTTTTACGTCCATGAGGGTTCTGCCCGCTGGGTGATG
AACATGCTAATGGCGCAGTGTCATTTACTCGACCCATGACACACGAAAGCCTGCCAAGCGGCAGGCCGAGGTCGAGCGGCGAGATGTGTCGTCAAGTGTC
ACGTGTTACGCCATCAAGCGGCGGTGCAACTGATACCGTCTGGGCGTCACGCTGAAGGCGTTTCCACATGCCATGTAGCTGGCGGTTAAGCGTTTCGTTA
TCGATATCCTGCACACCGCGCTTGGCTAGTATCACAATATCCATAGAGGGAAGGTGATGCTGCCGAAGCCTGGCGGATTCTCGGACAAGACGCTTAAAGC
GGTTGCGATCTACAGCGCGCTTTACATTTTTCTTGCTGACCACCAGTCCAATGCGGGGATAGCCCCGGGTACTCATGCGCGCTAGCGCCATCATACCTTT
ACCATGCACTTTGAGGTCGGCCTGTTCAAACACGTGACTAAAATCCCCGGCGTTTAAAAGACGTAAACACCGGGGAAAGCCTTGAGACGGCACACGCAAT
CTGAAATCAGGCGCTCAGACGCTTGCGGCCTTTCGCACGGCGGCGCGAGATAACGGCACGGCCGTTTTTGGTTGCCATGCGAGCACGGAAGCCATGCGTG
CGCTTGCGTTTGAGAACGCTGGGTTGAAAAGTGCGTTTCATAACTCGTAATTCCCACGTGGTTTAGGACGGATGTGAATCCAAAAGACCCGGTCCAGTGA

GGAAACTAGCCGGGGAGTTTCAATTCAAGACCGGAAATTGTAGAGACTTCCCGCGTCGGGTGCAATTTTTCCTTGCATAAAAACCAACAAAAAGGCAATG
GGCATCGGCCCGGTTATATTTCATCGTTCATGGCTTAGACAACTACTATTACTAGTTAAGGTATTTAAATGATGTTGTTATTAATAGTGGGGATACTATT
**TGTGGATAA**GTAGTTATTTCTTATGTTTATCAAATTGCTAAAATTATAAAAACAGTGTGCATAAGTATAGAATAAACGGCCGATAAGCTGTTAGTTTGGT
GTGAGTAGCAATATGAAAAAACCCCAAAAATGGGTCTGTCCCCATGTCTTCAGAAGGTTGTCCCACGTGCTCATGGGAAGTTGTCCACAGGCACAAATTG
ATAGCCAAACCCATGTGGATAACCAAGCTCTAGAGCGCTACAATGGTCGGCCGTTTTCAACTGATGGTGAGATGAGTGTCACTCGCGCTTTGGCAACAGT

GCCTGGACTACCTGCAGGACGAATTGAATTCACAACAATTCAATACCTGGATACGTCCCTTGCAGGCAGAAGAAGGAGAGTCCAATGAGCTGCGCTTACT
GGCGCCTAACCGCTTTGTGCGCGACTGGGTGAGTGACAAGTACGCCAAGCGGATCAGTGAGCTGATTCGTGAGCTTGCGCCGTCTAAACCACCCAAGGTA
AGTCTGACCGTTGGTAGCCGCCGCCCGGCGGCACCTGCGCCTCAACCACGTGATTTGGGCAATCCGGTTTCCGCCGCCACCTCTGCGCCTTCTGTGCATA
CACCTCCGCGTGGAGATATTGCCGATGAGCGTGAGATCGACCAGCTGCGTGAGCGTGAAGAAGCGCCACGTCGGCGTACCAATGAGCGTCAGGTGCAAGT
CGAAGGTAGCCTCAAGCACGGCAGTGGACTTAATCCGAATTTTACGTTTGATACCTTTGTTGAAGGTAAATCCAACCAACTGGCCCGGGCGGCTTCTCGG
CAGGTCTCGGAGAACCCGGGCGGTGCCTATAACCCCTTGTTTCTCTACGGCGGGGTAGGTCTGGGTAAAACGCACTTAATGCATGCGGTGGGTAATGCGC
TGGCTGGCCGCCGGGAAAATGCCCGGGTGGTTTACCTGCATTCAGAGCGTTTTGTTGCGGATATGGTCAAGGCGCTTCAGCTTAACGCGATCAACGATTT
CAAGCGCTTCTATCGAAGCGTTGATGCTTTACTTATTGATGATATTCAGTTTTTTGCTGGCAAGGAGCGCTCTCAAGAAGAGTTTTTCCACACGTTCAAT
GCATTATTGGAAGGTGGCCAGCAGATGATCCTTACCTCTGATCGTTATCCAAAGGAGATCAGTGGTGTCGAGGAGCGCCTGAAGTCGCGCTTTGGATGGG
GTTTGACGGTGGCGATCGAGCCGCCGGAGCTTGAGACCCGAGTGGCCATCTTGATGAAGAAAGCCGACCAGGCCAAGGTCGATCTGCCCCATGATGCTGC

>Halomonas nanhaiensis JCM 18142 [NZ_RZHE01000015.1:c63307..63806], <mnmG oriC2 <COG1126 ((glnQ), 554 bp

AGCCCCTTGATCGAGCGAACCACCTGTAGCTGTATGTCAAAGGGCAGCGAGGTGGAAATACCGTTGGGATAGAGCTCGTGGGTATCGAGACCTTCCGGTT
CGATAAACACCTGGTGGCTCGACTTATCCGCAAAGCGGTGCACTTTGTCTTCAATCGACGGGCAGTAGCGGGGGCCAATGCCTTCGATCACTCCTGAATA
CATTGGCGAGCGATCAAGATTGGCCAGAATTAGCTCATGTGTACGTTCATTGGTATGCGCAATGTGGCAGCTGACCTGTCGAGGATGCATTTCCCGCGAA
CCAAGGTACGACATCACAGGCGTTGGGGTATCGCCAGGTTGCTCTTCAAGCTGGGAAAAATCCACGCTTTTGGCATCGATGCGCGGTGGCGTGCCAGTTT
TTAACCGGTCAACCCGAAAAGGTAGCGCGCGCAAACGCTCGGCCAGGGCGTTGGAAGGTGGATCGCCTGCTCGGCCACCCCGGCTTTGATCCAGGCCAAT
ATGGATAACGCCGCCTAAAAACGTGCCTGTCGAAAGCACGACGGATTCGGCATAAAAACGAATGCCGGTTTCCGTGACCACGCCACGAACAGTATCGTTA
TCCACAACCAGATCGCCTGCGGCCTGCTGGAAAATGGTCAGGTTGGGCTGGTTTTCAAGCATGCCCCGGATTGCCGCTTTATAGCGGATCCGGTCGGCCT
GGGCACGTGTAGCACGTACTGCAGGACCCTTACGAGCATTTAGCACACGAAACTGAATGCCGCCCAAATCAGTGGCAAGCCCCATTGCGCCACCCAGTGC
ATCGATTTCCTTAACCAGATGACTTTTGCCAATGCCACCGATGGCCGGATTGCACGACATCTGGCCTAACGTCTCGATGTTATGGGTGAGCAGCAGGGTT
TGACAGCCCATACGAGCAGAGGCCAATGCGGCTTCAGTTCCCGCATGGCCACCGCCGATGACAATCACGTCAAAGCGGTCGGGGTAATTCAAGAGACACC

TCTTCTTGTGCCAATTGGCACGGATGATGGGAGCCGATGGATAGGCCCCAGATTCAATAAGCCCGCCAGTATAAACCCCCTGTGGGTAACCGTCAGGTTT
TTCCACACCTCAATTCATCTATCTGTCTCTATTAATAACAACATAAAAATAAAATAAGAGAAGTTCTGTTAATGTTGTAACTACTAGTGTGGACAGTTTC
**TGTGCATAA**CACGATTATTAGCATAAATTTCAGCTGTTTAAATTGTTCACATTTATGTGATTAAGCAGCGTAGTGCCTGCGCAGAAGCGGTGTGGCCCCT
GTGGATGAAAAAGCGACTTACCCACACCACTCAAAAATAACCGCTTATCCACCTTCCCATACCGGGTTTGTTACCGCACCTGTGAACAAGTCAGTCCGTC
AATTTCTTATTGTCTTGAAGGTCCTGTTAATGGGGGGTAGGGCCTGGTGAAGAAGAAAAATCAAAAAACATTATCCACAGGCAAAAAAATGGCCTGTCAC

AAGGACAGGCCATTAAAATTCAGGGTAAAGCGAGCGTTTTTATCGATTAATGCTTTAACACGCGGGCCAGAAATGAACGGGTTTGCTCATTTTGAGGCGT
GCTAAAAATCTGTTCAGGCGTTCCCTGTTCGGCAATTACCCCCTTGTGAATAAAGATAACGCGGTCAGCCACTTCACGGGCAAAGCCCATTTCATGGGTA
ACGATCACCATAGTCATGCCTTCTTTGGCAAGCTCACGCATTGCATCCAGCACTTCACCTATCATCTCCGGGTCCAGGGCCGAGGTAGGCTCGTCAAACA
GCATCAAACGTGGCTCCATGGCCAAAGCGCGTGCCAGTGCCACCCGCTGCTGCTGGCCACCGGAGAGCTGGCTTGGGTACTTGTTGGCCTGGTCGGCAAT
GCCTACCCGATCAAGCAGACGTTTAGCCGTGGCGACCGCATCTTCCCGGTTCCAGCCACGTACTTTCATGGGGGCCAAGGTAATGTTGTCGATAACGCTG
AGATGGGGAAACAGGTTGAACTGCTGGAATACCATGCCGACTTCGGTACGGATAGTCTGCAGCGCCTTGCTGCTCTTGCCATCGGGTTTCAGGGTATTGC
CATCAACGGCCAGCGTGCCCGACTGAAACTCTTCCAGGCCGTTGATGCAGCGAATCAGGGTCGATTTACCCGAGCCGCTGGCACCGATAATCACGACGAC
TTCGCCTGGCACAATCTCAAGGTCGATATCATCGAGTACATGCAGGCTGCCAAAGTGCTTGTTGAGCTTCTGCATACGCACAATAGGCGGCGTTGTAGGT
GACACAGTATCGGTCATTGTACGTGTCCTTATTGGCCGGCAATGCCTCTGCGCTCAAGCTGGCGCAGAAGAATGGAAAGGGTCCAGGTGATCGCCAGGTA
CATCAGCGCCACCATGAAATAGACCTCAAGGGCGGTAAACGTGGTGGCGATGTAAATCTGCCCCTGGCGCACCAGCTCACCCACGCCAATCACGGAAAAC

>Halomonas nanhaiensis JCM 18142 [NZ_RZHE01000015.1:c42229..42285], dnaA> ig dnaN>, 57 bp

WebSkew GCmin pos. 68252 (GCmin<>oriC1 24.0 kb; GCmin<>oriC2 1.1 kb; oriC2<>oriC1 23.0 kb)

>Halomonas songnenensis CGMCC 1.12152 [NZ_PVTK01000004.1:c43937..44436], <rpmH oriC1 dnaA>, 534 bp

CATCGGTAGTCACCGCGATAAAGTCACGGCTCGAGGTTTCGCTTTCGGTATCGCCAGCCAGGCCTTCGCCTACTGCACTGTCCGGTGCCGACGTTGACGG
AACCGCCAGGTCGTCATTACCACTTTGTGCATTGGGCAAAGGCGCGCTGTTGCTGCTTTGCGTAATCTCGGGCGTCGAGTCGTAATCTGTCGGCCCATAA
TCCTGATTCCACTGGATGACCAGCAGATAGGCAAGTACTGCCAGTGGAATCAGTAAGAGAAGTCGTTTTACGTCCATGAGGGGTCTGCCCGCTGGGTGAT
GAACATGCTATGGCGCAGTGTCATTTACTCGACCCATGACACACGAAAGCCTGCCAGGCGGCAGGCCGAGGTCGAGCGGCGAGATGCATCGTCAAGCGTC
ACGTGTTACGCCATCAGGCGGTGGTGGTGCTGGTACCGTCTGGGCGTCACGCTGAAGGCGTTTCCACATGCCGTGTAGCTGGCGGTTAAGCGTTTCGTTA
TCGATATCCTGCGCACCGCGCTTGGCCAATATCACGATATCCATGGAGGGTAGGTGATGCTGACGAAGTCTGGCGGACTCACGAACAAGACGCTTGAAGC
GGTTGCGATCTACAGCGCGCTTTACATTCTTCTTGCCTACCACCAGTCCAAAGCGGGGGTAGCCCCGGGTACTCATGCGCGCTAGCGCCATCATACCTTT
GCCATGCACTTTGAGGTCGGCTTGCTCAAACACGTGACTGAAATCCCCGGCGTTTAAAAGACGTAAACACCGGGGAAAGCCTTGATGCGGCACACGCAAT
CCGAGATCAGGCGCTCAGACGCTTGCGGCCTTTCGCACGACGGCGCGAGATAACGGCGCGGCCGTTCTTGGTTGCCATGCGAGCACGGAAGCCATGCGTG
CGCTTGCGTTTGAGAACGCTGGGTTGAAAAGTGCGTTTCATAACTCGTAATTCCCACGTGGTTTAGGACGGATGTGAATCCAAAAGACCCGGTCCAGTGA

GGAAACTAGCCGGGGAGTTTCAATTCAAGACCGGAAATTGTAGAGACTTCCCGCGTCGGGTGCAATTTTTCCTTGCATAAAAACCAACAAAAAGGCAAAG
GGCCAAGCCCCGATGACCCATCGTCGTTTGACACACCGACAACTACTATTACTAGTTAAGGTTTTAAATGATGTTGTTATTAATAGTGGGGATACTATT
**TGTGGATAA**GCACTTATTCCTTTTATTTATCAAATTACTAAAATCATAAAAACAGTGTGCATAAGTGCCGAATAACCGGTCGGTAAACTGTTGGTTTGGT
GTGAGTAGAGGTGTGGAAAACCCTAAAAAGCAGATCTGTCCCCATCTCTTCAGGAGGTTGTCCCACGTGCTCATGACAAGTTGTCCACAGGCGAAAAATC
GCCTCCAAACCCGTGTGGATAACCGGGGCTTAGGGCGCTACAATGGTCGGCCGTTTTCAACTGATGGTGAGATGAGTGTCACTCGCGCTTTGGCAACAGT

GCCTGGACTACCTGCAGGACGAATTGAATTCCCAGCAATTCAATACCTGGATACGTCCGCTGCAGGCAGAAGAAGGAGAGTCCAACGAGCTGCGCCTACT
GGCGCCTAATCGCTTTGTGCGCGACTGGGTGAGTGACAAGTACGCCAAGCGAATCAGCGAACTCATCCGCGAACTCGCACCGTCCAAGCCCCCCAAGGTA
AGCCTGACCGTAGGCAGCCGTCGGCCGGCTACCCCGGCCACGGCGCCGCGAGACCTTGGCAATCCGGTGTCGGCCTCCCCGGAGGCGCCCGCCGTGCATA
CGCCGGCGCGCGGCGATATCGGCGATGAGCGTGAAATCGACCAGCTGCGCGAGCGCGAAGAGGCCCCGCGGCGACGCAACAACGAGCGCCAGGTCCAGGT
AGAAGGCAGCCTCAAGCACGGCAGCGGCCTCAACCCCAATTTCACCTTCGATACCTTCGTGGAAGGTAAATCCAACCAGCTGGCCCGTGCGGCCTCGCGC
CAGGTATCAGAAAACCCGGGCGGCGCCTATAACCCATTGTTTCTGTACGGCGGCGTGGGTCTGGGTAAGACTCACCTGATGCACGCAGTGGGCAACGCGC
TGGCGGGCCGCCGGGATAACGCTCGGGTGGTGTATCTGCACTCCGAGCGTTTCGTGGCCGACATGGTCAAGGCGCTTCAGCTGAACGCGATCAACGACTT
CAAGCGTTTCTACCGCAGTGTCGATGCGTTACTGATTGATGACATTCAGTTTTTTGCCGGAAAGGAGCGCTCTCAGGAAGAGTTTTTCCACACCTTCAAT
GCGCTTCTGGAAGGCGGTCAGCAGATGATTCTGACCTCCGACCGCTATCCAAAAGAGATCAGCGGCGTGGAGGAGCGCCTCAAATCGCGTTTCGGCTGGG
GGCTGACGGTGGCCATCGAGCCGCCGGAGCTGGAAACCCGGGTGGCTATCTTGATGAAAAAAGCCGACCAAGCTAAGGTCGAGCTGCCACATGATGCGGC

>Halomonas songnenensis CGMCC 1.12152 [NZ_PVTK01000004.1:c66894..67393], <mnmG oriC2 <COG1126 (glnQ), 556 bp

GTCCCTTGATCGAGCGTACTACCTGTAGCTGAACGTCAAACGGCAGCGAGGTGGAGATACCGTTGGGGTAGAGCTCGTGAGTGTCGAGTCCTTCGGGTTC
GATGAACACCTGATGGCTCGACTTGTCCGCGAAGCGGTGCACCTTGTCCTCGATCGACGGGCAGTAGCGCGGGCCAATGCCTTCGATGACACCAGAGTAC
ATCGGCGAGCGGTCAAGATTCGCCATGATGATCTCGTGGGTGCGCTCATTGGTGTGCGCAATATGGCAGCTGACCTGCCGCGGGTGCATGTCCCGCGAGC
CAAGATAGGACATCACCGGCGTTGGCGTGTCGCCCGGCTGCTCTTCAAGCTGGGAAAAATCCACGCTTTTGGCATCGATGCGCGGTGGCGTACCGGTTTT
TAACCGGTCGACCCGAAACGGCAGGGCGCGCAGACGTTCTGCAAGGGTATTGGAAGGCGGGTCGCCTGCCCGGCCACCGCGGCTTTGATCCAGACCGATA
TGAATGACGCCGCCCAGGAACGTGCCGGTCGAAAGCACCACGGATTCGGCATGAAAACGAATCCCGGTTTCCGTTACCACGCCACGCACGGTGTCGTTAT
CCACAATCAGATCGCCCGCTGCCTGCTGAAAGATGGTCAGGTTGGGCTGGTTTTCCAGCATGCCGCGAATCGCTGCCTTGTAGCGGATTCGGTCGGCCTG
GGCACGCGTGGCACGCACCGCCGGGCCCTTGCGGGCGTTCAGCACACGAAACTGGATGCCGCCCAAATCAGTGGCAAGCCCCATCGCGCCGCCCAGTGCA
TCGATTTCCTTGACCAGATGGCTCTTGCCGATACCGCCAATAGCGGGATTACACGACATCTGGCCCAGGGTTTCGATGTTGTGGGTGAGCAGCAGGGTTT
GACAGCCCATGCGAGCAGAGGCCAATGCGGCTTCAGTTCCCGCATGGCCACCGCCGATGACAATCACGTCAAAGCGGTCGGGGTAATTCAAGAGACACCT

CATCTTGCGCCAGTCGGCACGGATCATGTGAGTCGGTGGATAACCACCCGGTTTCAATAAGCCCGCCAGTATAAACCCCCTGTGGGTAACCGTCAGGTTT
TTCCACACCTCGATTCGTTAACCAGTCTCTATTAATAACAACATAAAAATAAAATAAGAGAAGTTCTATTGATGTAGTAACTATTAGTGTGGACAGTTTC
**TGTGCATAA**CTGGATTATTGTTTTAAATTTCATACTTTTATATTGTTGACCTTTACGTGGTTAAAGCGCGGAGTGCCTGCGCAGAAGCGGTGCGGGCCCT
GTGGATGAAACGAAGACTTACCCACAGCGCCTGGAAAACACCGTTTATCCACCTGCCCATACCGGGCTTGTTACCGCACCTGTGAACAACGTTGCTGACG
CCTTTTCAATCCGCTTGAAAGGCAGCACGGTGGGCAGCGTAGCGTCATGGGCCGTTAAACAGGAAAAAAATTATCCACAGGCAAAAAAATGGCCTGTCAC

GGGGACAGGCCATTAAAAAATGCCGTTGAACGCGTTTTTTACGTCGTTTAATGCTTCAGAACGCGAGCAAGGAAAGAGCGCGTATGGTCATTTTTAGGCG
CGCTGAAGAACTCTTCAGGTGTGCCCTGTTCGGCAATTTCACCTTTATGGATAAAGATGATGCGATCAGCCACCTCGCGGGCAAAGCCCATTTCGTGGGT
GACAATCACCATGGTCATGCCCTCCTTGGCAAGCTCACGCATGGCGTCCAGCACTTCACCGATCATCTCCGGATCAAGCGCTGAGGTTGGCTCGTCAAAC
AGCATCAGGCGTGGCTCCATGGCGAGCGCCCGCGCCAGTGCCACCCGCTGCTGTTGGCCGCCGGAGAGCTGGCTTGGGTACTTGTCGGCCTGGTCGGCGA
TGCCTACCCGATCCAGCAGGCGCTTTGCCGTGGCCACTGCATCACTGCGGCTCCAGCCGCGTACTTTCATGGGGGCGAGAGTAATGTTGTCGATGACGCT
GAGGTGGGGAAACAGGTTGAACTGCTGGAAGACCATGCCGACTTCAGTACGAATGGTCTGCAGCGCCTTGCTGCTCTTGCCGTCGGGTTTCAGGGTATTG
CCATCGACGGTCAGGGTGCCCGACTGAAACTCTTCCAGGCCGTTGATACAGCGAATCAGCGTCGATTTACCCGAGCCGCTGGCACCAATGATCACGACAA
CTTCGCCAGGGGTAATTTCAAGGTCGATATCATTGAGTACATGCAGGCTGCCAAAATGCTTGTTGAGCTTCTGCATACGCACAATTGAGGGTTCTGATGA
CGTAGTAACGGTCATTGTACGTGTCCTTATTGGCCGGCGATGCCGCGGCGCTCAAGCTGGCGCAGCAGGATGGAGAGCGTCCAGGTGATCGCCAGGTACA
TCAGCGCGACCATGAAGTAGACCTCAAGGGCGGTAAAGGTAGTGGCGATATAGATCTGCCCCTGACGAACCAGCTCGCCCACGCCAATCACGGAAAACAG

>Halomonas songnenensis CGMCC 1.12152 [NZ_PVTK01000004.1:c42441..42497], dnaA> ig dnaN>, 58 bp

WebSkew GCmin pos. 67419 (GCmin<>oriC1 25.1 kb; GCmin<>oriC2 1.2 kb; oriC2<>oriC1 26.4 kb)

>Halomonas zhanjiangensis DSM 21076 [NZ_KB898653.1:c42024..42523], <rpmH oriC1 dnaA>, 534 bp

TACGTCTGTGGCCACTGCGATCAGGTCACGGCTGGACGTTTCACTTTCCGCGTCGCCAGGCATGCCTTCGTCTATCGCCGTATTGGGCGTTGACGTAGAT
GGAACGGCTAAACCGTTTTCGTCTTCGCTACCGCTTAAGGGCGCATTGTTCGTGCTTTGTGTAATCTCAGGCGTCGAGTCGTAATCTGTCGGCCCATAAT
CCTGATTCCACTGAATAACCAGTAGATAGGCAAGTACTGCCAGTGGAATCAGTAATAGAAGTCTTTTTACGTCCATGAGGGTTCTGCCCGCTGGGTGATG
AACATGCTAATGGCGCAGTATCATTTACTCGACCCATGACACACGAAAGCCTGCCAAGCGGCAGGCCAAGGTCGAGCGGCGAGATGCGTCGTCAAGAGTC
ACGTGTTACGCCATCAGACGGTGGTGCTACTGATACCGTTTGGGCGTCACGCTGAAGCCGTTTCCACATGCCATGTAACTGGCGAAAAAGCGTTTCGTTA
TCGATGTACTGTACGCCGCGCTTAGCGAGTATCACGATATCCATTGAGGGTAAGTGATCCTGGCGAAGGCGGGCGGATTCTCGAACAAGACGCTTAAAGC
GGTTGCGATCGACAGCACGCTTTACATTTTTTTTGCTGACCACCAGTCCTACGCGGGGGTGTCCCCGGGTACTCATGCGCGCTAACGCCATCATACCTTT
GCCATGAACTTTGAGGTCGGCCTGTTCAAACACGTGACTAAAATCCCCGGCATTTAGCAGACGTAAACACCGGGGATAGCCTTGATGCGGCACACGCAAT
CCGAGATCAGGCGCTCAGACGCTTGCGGCCTTTGGCGCGGCGGCGTGCGAGTACGGCACGGCCGTTTTTGGTTGCCATACGAGCACGGAAGCCATGCGCG
CGCTTGCGCTTGAGAACGCTGGGTTGAAAAGTGCGTTTCATAACTCGTTATTCCCACGTGGTTTAGGACGGATGTGAATCCAAAAGACCCGGTCCAGTGA

GGAAACTAGCCGGGGAGTTTCAAATCAAGACCGGAAATTGTAGAGACTTCCCGCGCCGGGTGCAATTTTTCCTTGCATAAAAAGCGACAAAAAGGCAATG
GGCATTTGTTGGATAGGTTATCTTCACCCATTACCTTGGCAACTACTATTACTAGTTAAGTATTTTAAAACCTATTGTTATTAATAGTGGGGATAAAATT
**TGTGGATAA**GTGGTTAATCTTTCTATTTATCATTGCCTTAAAATCATAAAAACAATGTGCATAAGTATCGTATAACCGGCAAATAAGCTGTTGGCTTGGT
GTGATTAAGCCCCTTGGAAAACCGGAAAATCCAATCTATCCCCACGCCGTTAAGGGGGTTGTCCCGAGACCTTAAAGAAAGTTGTCCACAGGCGAAATTG
AAGGCTAAACCCGTGTGGATAACCGAGCCTTTGGGCGCTACAATGGTCGGCCGTTTTCAACCGATGGTGAGATGAGTGTCACTCGCGCTTTGGCAGCAGT

GCCTGGACTACCTGCAGGACGAATTGAATTCACAGCAATTCAATACCTGGATACGTCCACTCCAGGCAGAAGAAGGAGAGTCCAACGAGCTGCGCTTACT
GGCGCCTAATCGCTTTGTGCGCGATTGGGTGAGTGACAAGTATGCTAAGCGGATCAGCGAGCTGATTCGTGAGCTTGCGCCCTCTAAACCGCCTAAGGTA
AGCCTTACGGTGGGTAGCCGCCGTCCAGCGGCACCGGCTCCTCAACCTCGCGATTTGGGCAACCCCGTTTCAGCGGCCACTGACCGTTCGTCGGCGCCTG
CTGTACATACGCCGCCGCGCGGAGATATTGCCGATGAGCGTGAAATCGACCAGATGCGCGAACGCGAAGAAGCGCCGCGTCGGCGCAACAATGAGCGCCA
GGTTCAGGTGGAGGGAAGCCTTAAGCACGGCAGTGGGCTTAATCCTAATTTTACTTTTGATACCTTTGTTGAAGGTAAGTCGAACCAGCTGGCCCGCGCT
GCCTCCCGACAAGTGTCGGAAAATCCGGGCGGTGCTTATAACCCACTATTTCTATACGGTGGGGTAGGCTTAGGTAAAACGCACTTGATGCATGCTGTAG
GTAATGCATTAGCCGGCCGCCGGGATAATGCCCGAGTGGTTTACCTGCATTCAGAGCGTTTTGTCGCGGACATGGTTAAAGCGCTTCAGCTTAATGCTAT
TAACGACTTCAAGCGGTTTTACCGCAGTGTTGATGCACTGTTAATTGACGATATTCAGTTTTTCGCTGGCAAAGAGCGCTCTCAGGAAGAGTTTTTCCAC
ACCTTTAATGCGCTATTAGAAGGTGGCCAGCAAATGATTCTTACCTCTGACCGCTATCCCAAAGAGATAAGCGGCGTGGAAGAGCGCTTGAAATCCCGGT
TTGGCTGGGGGTTGACGGTGGCAATTGAGCCGCCTGAACTTGAGACCCGGGTCGCTATCTTGATGAAAAAGGCCGACCAGGCCAAGGTCGATTTACCCCA

>Halomonas zhanjiangensis DSM 21076 [NZ_KB898653.1:c68382..68881], <mnmG oriC2 <COG1126 (glnQ), 574 bp

AACCACCTGCAGCTGCACATCAAACGGCAGCGAGGTCGAGATACCGTTGGGGTAGAGCTCGTGAGTATCCAGGCCTTCCGGCTCAATAAATACCTGATGG
CTGGATTTATCGGCAAACCGGTGAACCTTGTCTTCAATCGACGGGCAGTAGCGCGGGCCAACCCCTTCGATCACGCCGGAATACATGGGTGAGCGGTCAA
GGTTTGCCAAAATGATTTCATGAGTGCGCTCATTGGTGTGCGCAATATGGCAACTGACCTGCTGAGGATGCATGTCCCGCGAGCCAAGATACGACATTAT
CGGCGTTGGCGTATCACCTGGCTGTTCTTCAAGCTGGGAAAAATCCACGCTCTTGGCGTCAATTCGCGGTGGCGTGCCGGTTTTCAACCGGTCAACGCGA
AACGGTAAAGCACGTAGACGCTCGGCCAGAGCGTTCGAGGGCGGGTCACCTGCCCGGCCGCCGCGACTCTGATCCAAACCAATATGGATCACACCGCCTA
AAAACGTCCCGGTGGATAGCACTACCGATTCGGCATAAAAACGAATGCCGGTTTCGGTAACCACGCCACGTACCGTGTCGTTATCCACAACCAGATCGCC
AGCCGCTTGCTGAAAAATCGTTAAGTTTGGCTGGTTTTCCAACATACCGCGGATCGCTGCTTTGTAGCGTACACGATCTGCTTGAGCACGTGTGGCACGC
ACTGCAGGCCCTTTTCGAGCGTTAAGCACCCGAAACTGAATACCGCCCAAATCCGTGGCAAGCCCCATCGCCCCGCCTAGCGCATCGATTTCTTTGACCA
AATGGCTTTTGCCAATGCCGCCAATGGCAGGGTTACACGACATCTGACCCAAGGTCTCGATGTTGTGAGTGAGCAGTAGGGTCTGACAGCCCATACGAGC
AGAGGCCAATGCGGCTTCGGTTCCCGCATGGCCACCGCCGATGACAATCACGTCAAAGCGGTCGGGGTAATTCAAGAGACACCTCGTCTTGCGCCTGTTG

GCACGAATGATGTGACCAATGTCGTCAGTTTTTTCCGACATAGCCACGGTAGTTCAATAAGCCGGGCAGTATAAACCCCCTGTGGGTAACCGTCAGGTTT
TTCCACACCTCAATTCATTAACTGACCTCTATTAATAACAATATAAAAATAAAATAAGAGAAGTTCTGTTAATGTTGTAACTACTAGTGTGGACAGTTTC
**TGTGCATAA**CTAACTTATAGTTATTAGTATCATGTACTTAAAATGTTGACCTTTACGTGGTTAAGCAGCGTATTGCCTGAGTAGAAACGGTGTCTCGCCT
GTGGATGAAATAGCTACTTACCCACAGCCTGCCAAAAACACGGGTTATCCACCTACCCATACCGGGCTTGTTACCGTACCTGTGAACATGTCACTCAACC
ACATGGTAAAGCGATGAGAAACGCTGTTTATCAGCGTTACAGCGTCATAGGGACAATTAAAATAAAAAGTTTATCCACAGGGCAAAAAAATGGCCTGTCG

CAAGGACAGGCCATTAAAAAGCACGTTTGAGTCGATAATTTATTCAGATTTAGTGCTTGAGCACCCGGGCTAGAAACGATTGGGTGCGTTCATGCTGAGG
GGTATCAAAGAGCTGTTCAGGCGTTCCCTGCTCTGCAATAACGCCTTTATGAATAAAGATAATCCGGTCGGCCACTTCGCGTGCGAAGCCCATTTCATGG
GTAACAATCACCATGGTCATACCTTCTTTGGCAAGCTCGCGCATGGCATCGAGTACCTCACCAATCATCTCGGGGTCAAGCGCCGAGGTAGGCTCGTCAA
ACAGCATTAGCCGGGGTTCCATGGCCAGAGCTCGAGCAAGCGCTACGCGCTGTTGCTGGCCGCCTGAGAGCTGGCTGGGATACTTATCCGCCTGGTCGGC
AATACCCACCCGATCAAGCAGGCGGTTTGCGGTAGCCACCGCATCGCTTCGGCTCCAGCCACGCACTTTCATCGGCGCCAGGGTGATATTGTCGATAACG
CTTAGATGAGGAAACAGATTGAACTGCTGGAACACCATGCCGACTTCCGTACGGATGGTTTGCAGCGCTTTACTGCTTTTACCGTCAGGTTTCAGCGTAT
TGCCATCAACGCTTAAGGTGCCTGATTGAAACTCTTCCAGGCCATTAATACAGCGAATCAGCGTTGATTTACCTGACCCGCTGGCGCCAATCACCACAAC
GACTTCGCCAGGAACAATCTCAAGGTCAATATCATTGAGTACATGCAGGCTGCCAAAGTGTTTGTTCAGCTTCTGCATACGCACAATGGGTGGCACTGTA
GGTGACATTGTATCGGTCATTGTACGTGTCCTTATTGGCCGGCAATGCCGCGGCTCTCAAGCTGGCGCAGAATAATGGAAAGGGTCCAGGTGATCGCCAG
ATACATCAGCGCGACCATAAAGTAGACCTCAAGGGCGGTAAACGTCGTGGCAATATAGATCTGCCCCTGACGCACCAGCTCGCCTACCCCGATTACGGAA

>Halomonas zhanjiangensis DSM 21076 [NZ_KB898653.1:c40531..40575], dnaA> ig dnaN>, 45 bp

< page 1 | #083 | Halomonas titanicae BH1

WebSkew GCmin pos. 948689 (GCmin<>oriC1 24.6 kb; GCmin<>oriC2 2.9 kb; oriC2<>oriC1 21.7 kb)

>Halomonas titanicae BH1 [NZ_AOPO01000001.1:973036..973535], <rpmH oriC1 dnaA>, 533 bp

CCAGAACATCGGTGGTCACGGCAATGAAGTCGCGGCTTGATGTCTCGCTTTCGGTATCAGGAATACCTTCACCTACCGCACTTTGCTGTGAAGACGTGCT
CGGCACCGCTAAGCCGCTTTCTCCATCTTCAGCATTGCTAGGAGACGTGGAATTGCTGCGTTGCGTCGTTTCAGGCACTGAATCGTAATTCGGCTGCCCG
TAATCCTGATTCCACTGGACGACTAGCAAGTAAGCGAGTACGGCTAGTGGAATCAGTAATAAAAGTCGTTTTACGTCCATGAGGGTTCTGCCCGCTGGGT
GGTGAACATGCTAATGGCGCAGTGTCATTTACTCGACTCTTGACACACGAAAGCCTGCCAGTTGGCAGGCCGAGGTCGAGCGGCGAGGGGTGCACCAAAG
GTCACGTGTCACGCTTAGGCGTTGGTGCTGCGATCACCGCCTGTGCGTCACGCTGTAGTCGTTTCCACATGCCGTGTAACTGGCGATGCAAGGTGTCGTT
ATCGAGATCCTGAACGCCACGTCGCGCAAGTACCACTATATCCACAGAGGGTAGGTGATCTTGACGAAGGCGAACGGATTCTCGAACCAGACGCTTGAAA
CGGTTGCGATCTACGGCGCGCTTAACATTTTTTTTGCTGACCACCAGTCCAAGGCGGGGGTGTCCCCGAGTACTAAAGCGAACTAACGCCATCATACCTT
TGCCATGCACTTTCAGGTCGGCTTGCTCAAACACGTGACTAAAGTCCCCAGCATTTAGTAACCGTAATTGCCGGGGAAAGCCTTGCTGCGGTACACGCAA
TCTGAGATCAGGCGCTCAGACGCTTGCGGCCCTTGGCACGGCGACGTGCGAGGACGGCGCGACCGTTTTTGGTTGCCATACGAGCACGGAAGCCATGCGC
GCGCTTGCGCTTGAGAACGCTAGGTTGAAAAGTGCGTTTCATAACTCGTTATTCCCACGTGGTTTAGGACGGAAAGTGAATTCCAAAAAGCCCGGTCCAG

TGAGGAACTAGCCGGGGAGATTCAATTCAAGACCGGAAATTGTAGAGACTTCCTCTGCTGGGTGCAATTTTTCCTTGCATAAAAGCGTTATCACGCCTCA
ACCAGCGTGCTTGATACGGAAAGAGTAGCCATTAATAAAGTCACTATTATTACTAATAGGTAGTTCTAAAACTGTTGTTATTAATAGTGGGGATACTATT
**TGTGGATAA**GTGGTTTTAAACGATATTTTTCAATTGCCTATAACAATAAAAAATCTGTGTGTAAGTCCCGTATAAGCCGTCTACAAGGTGTTGGCGAGAT
GTGATTAACCCGTGTGTAAGACACGTATTTAACGATTATCCCCGTCCCTCCACAAGGTTTTCCCATGGGCTCATCGCTAGTTGTCCACAGGCCAATATTG
AGGGCTAAACCCGTGTGGATAACCGAGGCTTTGGGCGCTACAATGGTCGGCCGTTTTCAACCGATGGTGAGATGAGTGTCACTGACGCTTTGGCAACAGT

GCCTGGACAACCTGCAGGATGAGCTGAATTCACAGCAGTTCAATACCTGGATACGCCCGCTGCAGGCAGAAGAAGGAGAGTCCAACGAGCTGCGCTTATT
GGCGCCGAATCGTTTTGTGCGCGATTGGGTGAGCGATAAGTATGCCAAGCGGATTAGTGAGCTGATGCGTGAACTCGCACCGGCCAAGCCGCCAAAGGTC
AGCCTGACGGTGGGTAGTCGCCGTTTGGCAGCCCAGGCCCCTCAGCATCGTGATTTAGGTAATCCTGTATCGGCTTCGCCGTCGCGGCCAGCATCGCCAG
CGGTACATACGCCGCCACGCGGCGATATTGCTGATGAGCGTGAAATCGATAGGCTGTGCGATGAAGCGCCGAGTCGGCGTAATAATGAACGTCAGGTACA
GGTAGAAGGCAGTCTGAAACATGGCAGCGGGCTCAATCCGAATTTTACTTTCGATACGTTCGTGGAAGGTAAGTCTAACCAGTTGGCTCGTGCTGCCTCC
CGCCAGGTGTCTGAAAACCCTGGTGGCGCTTATAACCCCTTGTTTTTATACGGCGGCGTAGGCTTGGGTAAAACCCACTTAATGCATGCAGTGGGGAACG
CATTGGCAACCCGGCGTGAAAACGCCCGGGTGGTATACCTGCACTCAGAGCGGTTTGTCGCCGATATGGTTAAAGCGCTCCAGCTTAATGCGATTAATGA
TTTTAAGCGTTTTTATCGCAGTGTTGATGCGTTACTTATTGATGACATTCAGTTTTTTGCTGGAAAAGAGCGCTCTCAAGAAGAGTTTTTTCATACCTTC
AATGCGTTATTAGAAGGTGGTCAGCAAATGATCCTGACCTCTGATCGCTATCCTAAAGAGATCAGTGGGGTAGAGGAGCGCCTTAAATCACGCTTTGGTT
GGGGGCTTACGGTCGCGATAGAGCCGCCTGAGCTTGAAACCCGTGTGGCCATCTTGATGAAAAAGGCCGACCAGGCTAAGGTCGATTTGCCTCACGATGC

>Halomonas titanicae BH1 [NZ_AOPO01000001.1:951350..951849], <mnmG oriC2 <COG1126 (glnQ), 617 bp

ACGCACCACCTGAATCTGCACGTCAAAGGGCAGCGAAGTCGAGATACCGTTGGGGTAGAGCTCATGGGTATCCAAGCCTTCGGGTTCGATAAACACCTGG
TGGCTGGATTTATCGGCAAAGCGATGAACCTTGTCTTCTATCGATGGGCAGTAACGTGGCCCAATCCCTTCGATGACGCCGGAATACATCGGCGAGCGAT
CAAGATTGGCCAGAATCAGATCGTGGGTGTGCTGATTGGTGTGAGCAATGTGGCAACTCACCTGCTGAGGGTGCATCTCCCGCGATCCCAGGTAGGACAT
AACCGGCGTTGGGCTATCACCGGGCTGCTCTTCCAACTGGGAAAAATCCACGCTTTTGGCATCGATGCGCGGCGGCGTACCGGTTTTTAGGCGGTCAACC
CGAAACGGGAGGGCGCGCAGACGTTCAGCCAGCGCGTTGGAAGGCGGGTCGCCCGCTCGGCCGCCTTTGCTTTGATCCAAACCAATATGAATCACGCCAC
CCAAAAATGTCCCGGTACACAGCACGACGCTGTCGGCATGAAAACGAATGCCGGTTTCCGTGACGACGCCTCGTACGGTGTCGTTATCCACAATCAGATC
ACCAGCGGCTTGCTGGAAAATCGTTAAGTTGGGCTGGTTTTCCAGCATTCCGCGGATCGCTGACTTGTAACGGATGCGGTCAGCCTGGGCACGGGTGGCC
CGAACGGCGGGCCCTTTGCTGGCATTGAGCACACGAAACTGGATACCACCCAAGTCCGTGGCAAGCCCCATGGCGCCGCCCAGTGCATCAATTTCCTTGA
CCAAGTGACTTTTGCCGATGCCGCCAATCGCGGGATTACACGACATTTGGCCCAGAGTTTCGATGTTATGGGTGAGCAGCAGGGTCTGACAGCCCATGCG
AGCAGAGGCCAGTGCGGCTTCAGTTCCCGCATGGCCACCGCCGATGACAATCACGTCAAAGCGGTCGGGGTAATTCAAGAGACACCTCGTCTTGTGCCTG

TCGGCACGGATGATGTGGCCAATGCAGTCGATTCTAGACGCAGCCACCTTAGATCAATGAGTCGGCAAGTATAAACCCCCTGTGGGTAACCGTCAGGTTT
TTCCACACCTCAAATCGCTAAACCGCCTCTATTAATAACAACATAAATATAAAATAAGAGAAGTTCTGTAAATGTTGTAACTACTAGTGTGGACAATTTC
**TGTGTATAA**GCGATTAATTATTAATTATTTCATCAACTTATATTGTTGACTTCTACATGTTTGAAGGCGGGAGTGCCTGGGTAGAAAGGGTGCTAAGCCT
GTGGATGAAATAGCGCCTTATCCACAGATGCAATTGGCCACGGCTTATCCACCGGCCAGTACCTGGGTTGTTACCGCAGCTGTGAACAAGCAACGTTGGG
TGTGGAGCTAACGCTGAGAGACGCCCATGGTAGGCCTTGCGGAGATAAAAACGATTAATGTAAAAAAAGTTTATCCACAGGCAAAAAAATGGCCTGTCGT

GTCGACAGGCCGGTAAAAAAGCGTCATTTCGCATTAGCGAACGCGTTATTTAGGTGCCAAGCGCAGGTCTACGCGAGGGCATTGAGTGTTCACATCTAGT
GTTTAAGCACCCGCGCAAGGAAGGATTGAGTACGTTCATGTTGCGGTGAATCGAACAGTTTTTCCGGGTGTCCCTGCTCGGTAATCTCGCCTTTATGGAT
AAAAATCACCCGGTCGGCAACCTCACGGGCAAAGCCCATTTCATGGGTAACAATCACCATGGTCATGCCCTCTTTGGCCAGTTCACGCATGGCATCGAGC
ACTTCACCGATCATTTCCGGGTCAAGCGCCGATGTGGGCTCGTCAAATAGCATCAGCCGGGGTTCCATGGCCAAGGCACGAGCGAGCGCCACACGCTGCT
GCTGGCCGCCTGAGAGCTGGCTGGGGTACTTATCGGCCTGGTCGGCGATACCCACCCGGTCAAGCAAACGTTTGGCATTGCTGACCGCGTCGTCACGGCT
TAAACCGCGCACTTTCATCGGCGCCAGTGTGACATTATCGATGACGCTGAGATGGGGAAACAGGTTGAATTGCTGAAAAACCATGCCCACTTCGGTGCGG
ATGGTTTGCAGTGCTTGGGTGCTCTTTCCATAGGGCAGCAGTTCATTGTCATCGACCACCAGACTGCCTGATTGGAACTCTTCCAGGCCGTTAATACAGC
GAATCAGCGTCGATTTACCTGAGCCACTGGCGCCGATGATCACGACAACTTCACCGGGCACAATCGTAAGGTCAATATCGTTGAGTACGTGCAGGCTGCC
AAAGTGCTTGTTGAGCTGCTGCATACGCACAATTGGTTGCGTTGTGTTGTTCAATGTCTCGCTCATTGCGCGTGTCCTTATTGTCCGGCCAGGCCTCTGC
GCTCAAGTTGGCGCAGGATGATGGAGAGCGTCCAGGTGATCGCCAGATACATCAGGGCGACCATAAAGTAGACCTCAAGGGCCGTGAAGGTGGTGGCGAT

>Halomonas titanicae BH1 [NZ_AOPO01000001.1:974978..975037], dnaA> ig dnaN>, 60 bp

WebSkew GCmin pos. 69441 (GCmin<>oriC1 22.8 kb; GCmin<>oriC2 1.1 kb; oriC2<>oriC1 21.7 kb)

>Halomonas sedimenti QX-2 [NZ_JACCGK010000004.1:c46416..46915], <rpmH oriC1 dnaA>, 533 bp

CAAGAACATCTGTGGTCACGGCAATGAAGTCGCGGCTTGATGTCTCTCTTTCGGTATCAGGAATACCTTCACCTACCGTACTTTGCTGTGAAGACGCGCT
CGGCACCGATAAGCCGCTTTCTCCATCTTCAGCATTGCTAGGAGACGTGGAATTGCTGCGTTGCGTCGTTTCAGGCACTGAATCGTAATTCGGCTGCCCG
TAATCCTGATTCCACTGGACGACTAGCAAGTAAGCGAGTACGGCTAGTGGAATCAGTAATAAAAGTCGTTTTACGTCCATGAGGGTTCTGCCCGCTGGGT
GGTGAACATGCTAATGGCGCAGTGTCATTTACTCGACTCTTGACACACGAAAGCCTGCCAGTTGGCAGGCCGAGGTCGAGCGGCGAGGGGTGCACCAAAG
GTCACGTGTCACGCTTAGGCGTTGGTGCTGCGATCACCGCCTGTGCGTCACGCTGTAGTCGTTTCCACATGCCGTGTAACTGGCGATGCAAGGTGTCGTT
ATCGATATCCTGAACGCCACGTCGCGCAAGTACCACTATATCCACAGAGGGTAGGTAATCTTGACGAAGACGAACGGATTCTCGAACCAGACGCTTGAAA
CGGTTGCGATCTACGGCGCGTTTAACATTTTTTTTGCTGACCACCAGTCCAAGGCGGGGGTGTCCCCGAGTACTTAAGCGAACTAACGCCATCATACCTT
TGCCATGCACTTTTAGGTCGGCTTGCTCAAACACGTGACTAAAGTCCCCAGCATTTAGTAACCGTAATTGCCGGGGAAAGCCTTGCTGCGGTACACGCAA
TCTGAGATCAGGCGCTCAGACGCTTGCGGCCCTTGGCACGGCGACGTGCGAGGACGGCGCGACCGTTTTTGGTTGCCATACGAGCACGGAAGCCATGCGC
GCGCTTGCGCTTGAGAACGCTAGGTTGAAAAGTGCGTTTCATAACTCGTTATTCCCACGTGGTTTAGGACGGAAAGTGAATTCCAAAAAGCCCGGTCCAG

TGAGGAACTAGCCGGGGAGATTCAATTCAAGACCGGAAATTGTAGAGACTTCCTCTGCTGGGTGCAATTTTTCCTTGCATAAAAGCGTTATCACAGCTCA
ACCAGCGTGCTTGATACGGTAAGAGTAGCCATTGATAAAGTCACTATTATTACTAATAGGTAGTTCTAAACCTGTTGTTATTAATAGTGGGGATACTATT
**TGTGGATAA**GTAGTTTTAAGCTATGTTTTTCAATTGCCTATAACAATAAAAAATCTGTGTGTAAGTCCCGTATAAGCCGTCTACAAGGTGTTGGCGAGAT
GTGATTAACCCGTGTGTAAGACGCATATTTAACAATTATCCCCGTCCCTCCACAAGGTTTTCCCATGGGCTCATCGTTAGTTGTCCACAGGGCAATATTG
AGGCCTAAACCCGTGTGGATAACCGAGGCTTTGGGCGCTACAATGGTCGGCCGTTTTCAACCGATGGTGAGATGAGTGTCACTGACGCTTTGGCAACAGT

GCCTGGACAACCTGCAGGATGAGCTGAATTCACAACAGTTCAATACCTGGATACGCCCGCTGCAGGCAGAAGAAGGAGAGTCCAACGAGCTGCGCTTATT
GGCGCCGAATCGTTTTGTGCGCGATTGGGTGAGCGATAAGTATGCCAAGCGGATTAGTGAGCTGATGCGTGAACTCGCACCGGCCAAACCGCCAAAGGTC
AGCCTGACGGTGGGTAGTCGCCGTTTGGCAACCCAGGCCCCTCAGCATCGTGATTTAGGTAATCCTGTATCGGCTTCGCCGTCGCGGCCAGCGTCGCCAG
CGGTACATACGCCGCCACGCGGCGATATTGCTGATGAGCGCGAGATCGATAGGCTACGCGATGAAGCGCCGAGTCGGCGTAATAATGAACGTCAGGTACA
GGTAGAAGGCAGTCTGAAACATGGCAGCGGGCTCAATCCGAATTTTACTTTCGATACGTTCGTGGAAGGTAAGTCTAACCAGTTGGCTCGTGCTGCCTCC
CGCCAGGTGTCTGAAAACCCCGGTGGCGCTTATAACCCTTTGTTTTTATACGGCGGCGTAGGCTTGGGTAAAACCCACTTAATGCATGCAGTGGGGAACG
CATTGGCAACCCGGCGTGAAAACGCCCGGGTGGTATACCTGCACTCAGAGCGGTTCGTCGCCGATATGGTTAAAGCGCTCCAGCTTAATGCGATTAATGA
TTTTAAGCGTTTTTATCGCAGTGTTGATGCGTTACTTATTGATGACATTCAGTTTTTTGCTGGAAAAGAACGCTCTCAAGAAGAGTTTTTTCATACCTTC
AATGCGTTATTAGAAGGTGGTCAGCAAATGATCCTGACCTCTGACCGCTATCCTAAAGAGATCAGTGGTGTAGAGGAACGCCTTAAATCACGCTTTGGTT
GGGGGCTTACGGTCGCGATAGAGCCGCCAGAGCTTGAAACCCGTGTGGCCATCTTGATGAAAAAGGCCGACCAAGCTAAGGTCGATTTGCCTCACGATGC

>Halomonas sedimenti QX-2 [NZ_JACCGK010000004.1:c68091..68590], <mnmG oriC2 <COG1126 (glnQ), 617 bp

GCGCACGACTTGAATTTGCACATCAAAAGGCAACGAGGTCGAGATGCCGTTGGGGTAGAGCTCATGGGTATCCAAGCCTTCGGGTTCGATAAACACCTGG
TGGCTGGATTTATCGGCAAAGCGATGAACCTTGTCTTCAATCGATGGGCAGTAACGCGGCCCGATCCCTTCAATTACGCCGGAATACATCGGCGAACGAT
CCAGGTTGGCCAGAATCAGGTCGTGGGTGTGCTGATTGGTGTGAGCAATGTGGCAACTCACCTGCTGAGGGTGCATCTCCCGCGATCCCAGGTAGGACAT
AACTGGTGTTGGGCTATCACCGGGCTGCTCTTCCAACTGGGAAAAATCCACGCTTTTGGCATCGATGCGCGGCGGCGTACCGGTTTTTAGACGATCAACC
CGAAACGGCAGGGCGCGCAGACGTTCAGCCAGCGCGTTGGAGGGAGGATCACCCGCTCGGCCGCCTTTGCTTTGATCCAAACCAATATGAATCACGCCAC
CCAAAAATGTACCGGTACACAGCACTACGCTGTCGGCATGAAAGCGAATCCCAGTCTCCGTGACAACGCCTCGTACGGTGTCGTTATCCACAATCAGATC
ACCAGCGGCTTGCTGGAAAATCGTTAAGTTGGGCTGGTTTTCCAGCATTCCGCGGATTGCTGCCTTATAACGGATGCGGTCAGCCTGGGCACGGGTGGCC
CGAACAGCGGGCCCTTTGCTGGCATTGAGCACGCGAAACTGGATGCCACCCAAATCCGTGGCAAGCCCCATGGCGCCACCCAGTGCATCAATTTCCTTGA
CCAAGTGACTTTTGCCGATGCCGCCAATCGCGGGATTACACGACATTTGGCCCAGAGTTTCGATGTTATGGGTGAGCAGCAGGGTCTGACAGCCCATACG
AGCAGAGGCCAGTGCGGCTTCAGTTCCCGCATGGCCACCGCCGATGACAATCACGTCAAAGCGGTCGGGGTAATTCAAGAAACACCTCGTCTTGTGCCTG

TCGGCACGGATGATGTGGCCAATGCAGTCGATTCTAGACGCAGCCACCTTAGATCAATGAGTCGGCAAGTATAAACCCCCTGTGGGTAACCGTCAGGTTT
TTCCACACCTCAAATCGCTAAACCGCCTCTATTAATAACAATATAAATATAAAATAAGAGAAGTTCTGTAAATGTTGTAATTACTAGTGTGGACAATTTC
**TGTGTATAA**GCGATTAATCATTAATTATTTCATTAACTTATATTGTTGACTTCTACATGTTTGAAGACGGGAGTGCCTGGGTAGAAAGGGTGCTAAGCCT
GTGGATGAAATAGCACCTTATCCACAGATGCAATTAGCCACGGCTTATCCACCGGCCAGTACCTGGGTTGTTACCGCAGCTGTGAACAACCATCGTTGGG
TGTGGAGCTAACGCTGAGAGACTCCCATGGTAGGCCTTACGGAGATAAAAACGATCAATGTAAAAAAAGTTTATCCACAGGCAAAAAAATGGCCTGTCGT

GTCGACAGGCCGGTAAAAAAGCGTCATTTCGCATCAGCGAACGAGTTATTTAGGTGCCAAACGCAGGTCTATGCTAGGGCATTGAATGGCTACATCTAGT
GTTTTAGCACCCGCGCAAGGAAGGATTGAGTACGTTCATGCTGCGGTGAATCGAACAGTTTTTCTGGATGGCCCTGCTCGGTAATCTCGCCTTTATGGAT
AAAAATCACCCGGTCGGCAACCTCACGGGCAAAGCCCATTTCATGGGTAACAATCACCATAGTCATGCCCTCTTTGGCCAGTTCACGCATGGCATCGAGC
ACTTCACCGATCATTTCCGGGTCAAGCGCCGATGTGGGTTCGTCAAATAGCATCAGCCGGGGTTCCATAGCCAAGGCACGAGCGAGCGCCACACGCTGCT
GTTGGCCGCCTGAGAGCTGGCTGGGGTACTTATCGGCCTGGTCGGCGATACCCACCCGGTCAAGCAAACGTTTGGCATTACTGACCGCGTCGTCACGGCT
TAAACCGCGCACTTTCATCGGCGCGAGGGTGACGTTATCAATGACGCTGAGATGAGGAAACAGGTTGAACTGCTGAAAGACCATGCCTACTTCGGTGCGG
ATGGTTTGCAGTGCTTGGGTGCTCTTTCCATGGGGCAGCAGTTCGTTGTTATCGACCACCAAACTGCCTGATTGGAACTCTTCCAGGCCGTTAATACAGC
GAATCAGCGTCGATTTACCTGAGCCACTGGCACCGATGATCACGACAACTTCACCGGGCACTATCGTAAGGTCAATATCGTTGAGTACGTGCAGGCTGCC
AAAGTGTTTGTTGAGCTGCTGCATACGCACAATTGGTTGCGTTGTGTTTTTCAATGTCTCGCTCATTGCGCGTGTCCTTATTGTCCGGCCAGACCTCTGC
GCTCAAGTTGGCGCAGGATGATGGAGAGCGTCCAGGTGATCGCGAGATACATAAGGGCGACCATAAAGTAGACCTCAAGGGCCGTGAAGGTGGTGGCGAT

>Halomonas sedimenti QX-2 [NZ_JACCGK010000004.1:c44914..44973], dnaA> ig dnaN>, 60 bp

WebSkew GCmin pos. 69422 (GCmin<>oriC1 24.8 kb; GCmin<>oriC2 1.3 kb; oriC2<>oriC1 23.5 kb)

>Halomonas alkaliantarctica CRSS [NZ_JACCGK010000004.1:44350..44849], <rpmH oriC1 dnaA>, 533 bp

```
CGAGAACATCGGTGGTCACGGCAATGAAGTCGCGGCTTGATGTATCGCTTTCGGTATCAGGAATACCTTCACCTACCACACTTTGCTGTGAAGACGCGCT
TGGCACTGCTAAGCCGCCTTCACCATCTTCAGCATTGCTAGGAGACGTGGAATTGCTGCGTTGCGTCGTTTCAGGCACTGAATCGTAATTCGGCTGCCCG
TAATCCTGATTCCACTGGACGACTAGCAAGTAAGCGAGTACGGCTAGTGGAATCAGTAATAAAAGTCGTTTTACGTCCATGAGGGTTCTGCCCGCTGGGT
GGTGAACATGCTAATGGCGCAGTGTCATTTACTCGACTCTTGACACACGAAAGCCTGCCAGTTGGCAGGCCGAGGTCGAGCGGCGAGGGGTGCACCAAAG
GTCACGTGTCACGCTTAGGCGTTGGTGCTGCGATCACCGCCTGTGCGTCACGCTGTAGTCGTTTCCACATGCCGTGTAACTGGCGATGCAAAGTCTCGTT
ATCGATATCCTGAACACCGCGTCGCGCAAGCACCACTATATCCACTGAGGGCAGGTGATCTTGACGAAGACGAACGGATTCTCGAACCAGACGCTTGAAA
CGGTTGCGATCTACGGCGCGTTTAACATTTTTTTTGCTGACCACCAGTCCAAGGCGGGGATGACCCCGAGTACTTAAGCGAACTAACGCCATCATACCTT
TGCCATGCACTTTTAGGTCGGCTTGCTCAAACACGTGACTAAAATCCCCAGCATTTAGTAACCGTAATTGCCGGGGAAAGCCTTGCTGCGGCACACGCAA
TCTGAGATCAGGCGCTCAGACGCTTGCGGCCCTTGGCACGGCGACGTGCGAGGACGGCGCGACCGTTTTTGGTTGCCATACGAGCACGGAAGCCATGCGC
GCGCTTGCGCTTGAGAACGCTAGGTTGAAAAGTGCGTTTCATAACTCGTTATTCCCACGTGGTTTAGGACGGAAAGTGAATTCCAAAAAGCCCGGTCCAG

TGAGGAACTAGCCGGGGAGATTCAATTCAAGACCGGAAATTGTAGAGACTTCCTCTGCTCGGTGCAATTTTTCCTTGCATAAAAGCGTTAACACACCTCA
GCCAGCGTGCTTAATACGCTGAGAGTAACGATTAATGAAGTCACTATTATTACTAATAGGTAGTTCTAAACCTGTTGTTATTAATAGTGGGGATACTATT
TGTGGATAAGTGTTTTTAACTATTACTTTTCAATGGCCTATGATAATAAAAAATCTGTGTGTAAGTCCCGTATAAACCGTCCACAAGGTGTTGGCTAGAT
GTGATTAACCCGTTTATAAGGCACCTTTTTAACAATTGTCCCCGTCCCTCCACAAGGTTTTCCCATAGGCTCACTGTAAGTTGTCCACAGGCCAAGATTG
AGGGGTAAACCCGTGTGGATAACCGAGGCTTTGGGCGCTACAATGGTCGGCCGTTTTCAACCGATGGTGAGATGAGTGTCACTGACGCTTTGGCAACAGT

GCCTGGACAACCTGCAGGATGAGCTGAATTCACAGCAGTTCAATACCTGGATACGCCCGCTGCAGGCAGAAGAAGGAGAGTCCAACGAGCTGCGCTTATT
GGCGCCGAATCGTTTTGTGCGCGATTGGGTGAGCGATAAGTATGCCAAGCGGATTAGTGAGCTGATGCGAGAACTTGCACCGGCCAAACCGCCAAAGGTA
AGCCTGACGGTGGGTAGTCGCCGCTTGGCCACCCAGGCCCCTCAGCATCGTGATTTAGGTAATCCTGTATCGGCTTCACCGTCGCGGCCAGCATCGCCTG
CGGTACATACGCCGCCCCGCGGCGATATCGCCGATGAGCGCGAAATCGATAGACTGCGCGATGAAGCGCCGAGTCGACGTAATAATGAACGTCAGGTACA
GGTAGAAGGCAGTCTGAAACATGGCAGCGGGCTTAATCCGAATTTTACTTTCGATACGTTTGTAGAAGGTAAATCGAACCAATTGGCTCGCGCCGCCTCC
CGCCAGGTGTCTGAAAATCCTGGCGGTGCTTATAACCCATTGTTTTTATACGGTGGTGTAGGCCTGGGTAAAACCCACTTAATGCATGCGGTGGGGAACG
CATTGGCAACCCGGCGTGAGAATGCCCGGGTGGTATACCTGCACTCAGAGCGGTTTGTCGCCGATATGGTTAAAGCGCTCCAGCTTAATGCGATTAATGA
TTTTAAGCGTTTTTATCGCAGTGTTGATGCGTTACTTATTGATGATATTCAGTTTTTTGCTGGAAAAGAGCGGTCTCAAGAAGAGTTTTTTCATACCTTC
AATGCGTTATTAGAAGGTGGTCAGCAAATGATTCTCACCTCTGATCGCTATCCTAAAGAGATCAGTGGGGTAGAGGAGCGCCTGAAATCACGCTTTGGTT
GGGGCCTTACGGTCGCGATAGAGCCGCCAGAGCTTGAAACCCGTGTGGCCATCTTGATGAAAAAGGCCGACCAAGCTAAGGTCGATTTGCCTCATGATGC
```

>Halomonas alkaliantarctica CRSS [NZ_JACCGK010000004.1:67874..68373], <mnmG oriC2 <COG1126 (glnQ), 616 bp

```
AACGCACGACCTGAATCTGCACGTCAAAGGGTAGCGAGGTCGAGATGCCGTTGGGGTAGAGCTCATGGGTATCCAGGCCTTCGGGTTCGATAAATACCTG
GTGGCTGGATTTATCGGCAAAGCGATGAACCTTGTCTTCGATCGATGGGCAGTAACGCGGTCCGATACCTTCAATGACGCCGGAATACATCGGCGAGCGA
TCAAGGTTGGCCAGAATCAGATCATGAGTCTGCTGATTGGTGTGAGCAATATGGCAACTCACTTGCTGAGGATGCATTTCCCGTGATCCCAGATAGGACA
TAACCGGCGTTGGGCTATCGCCGGGCTGCTCTTCCAACTGAGAAAAATCCACGCTTTTGGCATCAATCCGCGGCGGTGTGCCGGTTTTTAGGCGATCAAC
CCGAAACGGCAGTGCCCGAAGACGTTCGGCCAGCGCGTTGGAGGGCGGATCGCCTGCTCGGCCACCTTTGCTTTGATCCAAACCAATGTGTATCACACCA
CCCAAAAATGTCCCGGTACACAGCACGACGCTGTCGGCATGAAAGCGGATGCCGGTTTCGGTGACGACGCCTCGTACTGTGTCGTTATCCACAATCAGAT
CGCCAGCGGCTTGCTGGAAGATCGTTAAGTTGGGCTGGTTTTCCAGCATTCCGCGGATCGCTGCCTTGTAACGGATGCGGTCAGCCTGGGCACGAGTGGC
CCGAACGGCGGGCCCTTTGCTGGCATTGAGCACGCGAAACTGGATGCCACCCAAGTCCGTGGCAAGCCCCATGGCGCCGCCCAGTGCATCAATTTCCTTG
ACCAAGTGACTTTTGCCGATGCCGCCAATGGCGGGATTACACGACATTTGGCCCAGAGTTTCGATGTTATGGGTGAGCAGTAGGGTCTGACAGCCCATGC
GAGCAGAGGCCAGTGCGGCTTCGGTTCCCGCATGGCCACCGCCGATGACTATCACGTCAAAGCGGTCGGGGTAATTCAAGAGACACCTCGTCTTGTGCCA

ATTGGCACGGATGATGTGGCCAGGCAGTCGATTCTAGACGCAGCCACCTTAGATCAATGAGGCGGCAAGTATAAACCCCCTGTGGGTAACCGTCAGGTTT
TTCCACACCCCGAATCGTTAAACAGGCTCTATTAATAACAACATAAATATAAAATAAGAGAAGTTCTGTAAATGTTGTAACTACTAGTGTGGACAATTTC
TGTGCATAAGCGATTAATCATTAATTATTTCATCAACTTATATTGTTGACTTCTACATGTTTGAAGGCGGGAGTGCCTGGGCAGAAAAGGTGTTAAGCCT
GTGGATGAAATAGCACCTTATCCACAGATACAATTAGCCACGCCTTATCCACCGGCCAGTACCTGGGTTGTTACCGCACCTGTGAACAACCACCGTTGGG
TGTGGAGCTAATGCTAAGAGGCTCCCATGGTAGGCCCTACGGAGATAAAAACGATCAATGTAAAAAAAGTTTATCCACAGGCAAAAAAATGGCCTGTCGT

GTCGACAGGCCGGTAAAAAAGCGTCATTTCGCATTAGCGAACGGGTTATTTAGGTGCCAAGCGCAGGTCTATGCGAAGGCATGAAGTGATCACATTTAGT
GTTTAAGCACCCGGGCAAGGAAGGATTGAGTACGTTCATGCTTCGGTGAATCGAACAGTTTTTCCGGGTGTCCCTGTTCGGTAATCTCGCCTTTATGGAT
AAAAATCACCCGGTCGGCAACCTCACGGGCAAAGCCCATTTCATGGGTCACAATCACCATGGTCATGCCCTCTTTGGCCAGTTCGCGCATGGCATCCAAC
ACTTCACCGATCATTTCCGGGTCAAGCGCCGATGTGGGCTCGTCAAATAGCATCAGCCGGGGCTCCATGGCCAAGGCGCGAGCAAGTGCCACACGCTGCT
GCTGGCCGCCTGAGAGCTGGCTGGGGTACTTATCGGCCTGGTCGGCGATACCCACCCGGTCAAGCAGCCGCTTGGCATTGCTGACCGCGTCGTCACGGCT
TAAACCGCGCACTTTCATTGGCGCCAGGGTGACGTTATCAATGACGCTGAGATGGGGAAACAGGTTGAACTGCTGAAAGACCATGCCCACTTCGGTGCGG
ATGGTCTGCAGTGCTTGGGTACTTTTACCATGGGGTAGCAGTTCGTTGTTATCGACCACCAGATTGCCTGATTGGAACTCTTCCAGGCCGTTAATACAGC
GAATCAGCGTCGATTTGCCTGAGCCACTGGCGCCGATGATCACGACTACTTCACCGGGCACAATCGTAAGGTCAATATCGTTGAGTACGTGCAGGCTGCC
AAAGTGCTTGTTGAGCTGCTGCATACGCACAATTGGTTGCGTTGTGTTGTTCAATGTCTCGCTCATTGCGCGTGTCCTTATTGTCCGGCCAGGCCTCTGC
GCTCAAGTTGGCGCAGGATAATGGAGAGCGTCCAGGTGATCGCCAGATACATCAGAGCGACCATGAAGTAGACCTCAAGGGCCGTGAAGGTGGTGGCGAT
```

>Halomonas alkaliantarctica CRSS [NZ_JACCGK010000004.1:42848..42907], dnaA> ig dnaN>, 60 bp

WebSkew GCmin pos. nd (GCmin<>oriC1 nd; GCmin<>oriC2 nd; oriC2<>oriC1 nd)

>Halomonas neptunia CECT 5815 [NZ_JAUFPP010000021.1:c44390..44889], <rpmH oriC1 dnaA>, 533 bp

CGAGAACATCGGTGGTCACGGCAATGAAGTCGCGGCTTGATGTATCGCTTTCGGTATCAGGAATACCTTCACCTACCGCACTTTGCTGTGAAGACGCGCT
TGGCACTGCTAAGCCGCCTTCACCATCTTCAGCATTGCTAGGAGACGTGGAATTGCTGCGTTGCGTCGTTTCAGGCACTGAATCGTAATTCGGCTGCCCG
TAATCCTGATTCCACTGGACGACTAGCAAGTAAGCGAGTACGGCTAGTGGAATCAGTAATAAAAGTCGTTTTACGTCCATGAGGGTTCTGCCCGCTGGGT
GGTGAACATGCTAATGGCGCAGTGTCATTTACTCGACTCTTGACACACGAAAGCCTGCCAGTTGGCAGGCCGAGGTCGAGCGGCGAGGGGTGCACCAAAG
GTCACGTGTCACGCTTAGGCGTTGGTGCTGCGATCACCGCCTGTGCGTCACGCTGTAGTCGTTTCCACATGCCGTGTAACTGGCGATGCAAAGTCTCGTT
ATCGATATCCTGAACACCGCGTCGCGCAAGCACCACTATATCCACTGAGGGCAGGCGATCTTGACGAAGACGAACGGATTCTCGAACCAGACGCTTGAAA
CGGTTGCGATCTACGGCGCGTTTAACATTTTTTTTGCTGACCACCAGTCCAAGGCGGGGATGACCCCGAGTACTTAAGCGAACTAACGCCATCATACCTT
TGCCATGCACTTTTAGGTCGGCTTGCTCAAACACGTGACTAAAATCCCCAGCATTTAGTAACCGTAATTGCCGGGGAAAGCCTTGCTGCGGCACACGCAA
TCTGAGATCAGGCGCTCAGACGCTTGCGGCCCTTGGCACGGCGACGTGCGAGGACGGCGCGACCGTTTTTGGTTGCCATACGAGCACGGAAGCCATGCGC
GCGCTTGCGCTTGAGAACGCTAGGTTGAAAAGTGCGTTTCATAACTCGTTATTCCCACGTGGTTTAGGACGGAAAGTGAATTCCAAAAAGCCCGGTCCAG

TGAGGAACTAGCCGGGGAGATTCAATTCAAGACCGGAAATTGTAGAGACTTCCTCTGCTCGGTGCAATTTTTCCTTGCATAAAAGCGTTAACACACCTTA
GCCAGCGTGCTTAATACGCTGAGAGTAACGATTAATGAAGTCACTATTATTACTAATAGGTAGTTCTAAACCTGTTGTTATTAATAGTGGGATACTATT
**TGTGGATAA**GTGTGTTTAACTATTACTTTTCAATGGCCTATGATAATAAAAAATCTGTGTGTAAGTCCCGTATAAGCCGTCCACAAGGTGTTGGCTAGAT
GTGATTAACCCGTTTATAAGACACCTTTTTAACAATTGTCCCCGTCCCTCCACAAGGTTTTCCCATAGGCTCACTGTAAGTTGTCCACAGGCCAAGATTG
AGGGGTAAACCCGTGTGGATAACCGAGGCTTTGGGCGCTACAATGGTCGGCCGTTTTCAACCGATGGTGAGATGAGTGTCACTGACGCTTTGGCAACAGT

GCCTGGACAACCTGCAGGATGAGCTGAATTCACAGCAGTTCAATACCTGGATACGCCCGCTGCAGGCAGAAGAAGGAGAGTCCAACGAGCTGCGCTTATT
GGCGCCAAATCGTTTTGTGCGCGATTGGGTGAGCGATAAGTATGCCAAGCGGATTAGTGAGCTGATGCGAGAACTTGCACCGGCCAAACCGCCAAAGGTA
AGCCTGACGGTGGGTAGTCGCCGCTTGGCCACCCAGGCCCCTCAGCATCGTGATTTAGGTAATCCTGTATCGGCTTCACCGTCGCGGCCAGCATCGCCTG
CGGTACATACGCCGCCCCGCGGCGATATCGCCGATGAGCGCGAAATCGATAGACTGCGCGATGAAGCGCCGAGTCGACGTAATAATGAACGTCAGGTACA
GGTAGAAGGCAGTCTGAAACATGGCAGCGGGCTTAATCCGAATTTTACTTTCGATACGTTTGTAGAAGGTAAATCGAACCAATTGGCTCGCGCCGCCTCC
CGCCAGGTGTCTGAAAATCCTGGCGGTGCTTATAACCCATTGTTTTTATACGGTGGTGTAGGCCTGGGTAAAACCCACTTAATGCATGCGGTGGGGAACG
CATTGGCAACCCGGCGTGAGAACGCCCGGGTGGTATACCTGCACTCAGAGCGGTTTGTCGCCGATATGGTTAAAGCGCTCCAGCTTAATGCGATTAATGA
TTTTAAGCGTTTTTATCGCAGTGTTGATGCGTTACTTATTGATGATATTCAGTTTTTTGCTGGAAAAGAGCGGTCTCAAGAAGAGTTTTTTCATACCTTC
AATGCGTTATTAGAAGGTGGTCAGCAAATGATTCTCACCTCTGATCGCTATCCTAAAGAGATCAGTGGGGTAGAGGAGCGCCTGAAATCACGCTTTGGTT
GGGGCCTTACGGTCGCGATAGAGCCGCCAGAGCTTGAAACCCGTGTGGCCATCTTGATGAAAAAGGCCGACCAAGCTAAGGTCGATTTGCCTCATGATGC

>Halomonas neptunia CECT 5815 [NZ_JAUFPP010000005.1:5293..5792], <mnmG oriC2 <COG1126 (glnQ), 616 bp

AACGCACGACCTGAATCTGCACGTCAAAGGGTAGCGAGGTCGAGATGCCGTTGGGGTAGAGCTCATGGGTATCCAGGCCTTCGGGTTCGATAAATACCTG
GTGGCTGGATTTATCGGCAAAGCGATGAACCTTGTCTTCGATCGATGGGCAGTAACGCGGTCCGATCCCTTCAATGACGCCGGAATACATCGGCGAGCGA
TCAAGGTTGGCCAGAATCAGATCATGAGTCTGCTGATTGGTGTGAGCAATATGGCAACTCACTTGCTGAGGATGCATTTCCCGTGATCCCAGATAGGACA
TAACCGGCGTTGGGCTATCGCCGGGCTGCTCTTCCAACTGGGAAAAATCTACCGTTTTGGCATCAATCCGCGGTGGCGTACCGGTTTTTAGGCGATCAAC
CCGAAACGGCAGTGCGCGCAGACGTTCGGCCAGCGCATTGGAGGGTGGATCGCCTGCTCGGCCACCTTTGCTTTGATCCAAACCAATGTGAATCACGCCA
CCCAAAAATGTCCCGGTACACAGCACGACGCTGTCCGCATGAAAGCGGATACCGGTTTCGGTGACGACGCCTCGTACGGTGTCGTTATCCACAATCAGAT
CGCCAGCGGCTTGCTGGAAGATCGTTAAGTTGGGCTGGTTTTCCAGCATTCCGCGGATCGCTGCCTTGTAACGGATGCGGTCAGCCTGGGCACGAGTGGC
CCGAACGGCGGGCCCTTTGCTGGCATTGAGCACGCGAAACTGGATGCCACCCAAGTCCGTGGCAAGCCCCATGGCGCCGCCCAGTGCATCAATTTCCTTG
ACCAAGTGACTTTTGCCGATGCCGCCAATGGCGGGATTACACGACATTTGGCCCAGAGTTTCGATGTTATGGGTGAGCAGTAGGGTCTGACAGCCCATGC
GAGCAGAGGCCAGTGCGGCTTCGGTTCCCGCATGGCCACCGCCGATGACTATCACGTCAAAGCGGTCGGGGTAATTCAAGAGACACCTCGTCTTGTGCCA

ATTGGCACGGATGATGTGGCCAGGCAGTCGATTCTAGACGCAGCCACCTTAGATCAATGAGGCGGCAAGTATAAACCCCCTGTGGGTAACCGTCAGGTTT
TTCCACACCCCTAATCGTTAAACAGCCTCTATTAATAACAACATAAATATAAAATAAGAGAAGTTCTGTAAATGTTGTAACTACTAGTGTGGACAATTTC
**TGTGCATAA**GCGATTAATCATTAATTATTTCATCAACTTATATTGTTGACTTCTACATGTTTGAAGGCGGGAGTGCCTGGGCAGAAAGGGTGTTAAGCCT
GTGGATGAAATAGCACCTTATCCACAGATGCAATTAGCCACGCCTTATCCACCGGCCAGTACCTGGGTTGTTACCGCACCTGTGAACAACCACCGTTGGG
TGTGGAGCTAATGCTAAGAGGCTCCCATGGTAGGCCCTACGGAGATAAAAACGATCAATGTAAAAAAAGTTTATCCACAGGCAAAAAAATGGCCTGTCGT

GTCGACAGGCCGGTAAAAAAGCGTCATTTCGCATTAGCGAACGGGTTATTTAGGTGCCAAGCGCAGGTCTATGCGAAGGCATGAAGTGATCACATTTAGT
GTTTAAGCACCCGGGCAAGGAAGGATTGAGTACGTTCATGCTGCGGTGAATCGAACAGTTTTTCCGGGTGTCCCTGCTCGGTAATCTCGCCTTTATGGAT
AAAAATCACCCGGTCGGCAACCTCACGGGCAAAGCCCATTTCATGGGTCACAATCACCATGGTCATGCCCTCTTTGGCCAGTTCGCGCATGGCATCCAAC
ACTTCACCGATCATTTCCGGGTCAAGCGCCGATGTGGGCTCGTCAAATAGCATCAGCCGGGGCTCCATGGCCAAGGCGCGAGCAAGTGCCACACGCTGCT
GCTGGCCGCCTGAGAGCTGGCTGGGGTACTTATCGGCCTGGTCGGCGATACCCACCCGGTCAAGCAGCCGCTTGGCATTGCTGACCGCGTCGTCACGGCT
TAAACCGCGCACTTTCATTGGCGCCAGGGTGACGTTATCAATGACACTGAGATGGGGAAACAGGTTGAATTGCTGAAAGACCATGCCCACTTCGGTGCGG
ATGGTCTGCAGTGCTTGGGTACTTTTACCATGGGGTAGCAGTTCGTTGTTATCGACCACCAGATTGCCTGATTGGAACTCTTCCAGGCCGTTAATACAGC
GAATCAGCGTCGATTTGCCTGAGCCACTGGCGCCGATGATCACGACTACTTCACCGGGCACAATCGTAAGGTCAATATCGTTGAGTACGTGCAGGCTGCC
AAAGTGCTTGTTGAGCTGCTGCATACGCACAATTGGTTGCGTTGTGTTGTTCAATGTCTCGCTCATTGCGCGTGTCCTTATTGTCCGGCCAGGCCTCTGC
GCTCAAGTTGGCGCAGGATAATGGAGAGCGTCCAGGTGATCGCCAGATACATCAGAGCGACCATGAAGTAGACCTCAAGGGCCGTGAAGGTGGTGGCGAT

>Halomonas neptunia CECT 5815 [NZ_JAUFPP010000021.1:c42888..42947], dnaA> ig dnaN>, 60 bp

WebSkew GCmin pos. 171680 (GCmin<>oriC1 17.3 kb; GCmin<>oriC2 0.3 kb; oriC2<>oriC1 17.0 kb)

>Halomonas olivaria TYRC17 [NZ_JALPZO010000066.1:188734..189233], <rpmH oriC1 dnaA>, 534 bp

ACGTCGGTAGAGACAGCAATAAAGTCACGGCTTGCTGTTTCACTCTCTGTCTCGCCCGGCATGTCCTCGCTAATGGCACTCTGAGGAGTCGACACACTGG
GAACAGCCAACTCATCACCATTACCGGCGTTGCCGTTCTGTGCCCCATTACTGCTTTGGGTAATGTCAGGCGTCGAATCGTAGGTTGTCTGCCCGTAATC
CTGATTCCACTGTACAACCAGTAAATAGGCGAGTACCGCCAGTGGTATCAGTAATAAAAGTCGTTTTACGTCCATGAGGGTTCTGCCCGATGGGTGGTGA
ACATGCTAATGGCGCAGTGTCATTTACTCGACCCATGACACACGAAAGCCTGCCAGGCGGCAGGCCGAGGTCGAGCGGCGAGGTGTGTCGTCAAGTGTCA
CGTGTTATGCCTGCTAATAGCGGAGGCGTAGCTACTGCCAGGGCATCACGCTGCAGTCGTTTCCACATACCGTCTAACTGACGGTATAGCGTTTCGTTAT
CAATATCCTGCACACCACGCTTAGCCAGAACCACTATATCCACAGAGGGTAAGTGATCTTGACGCAGCCGGACGGACTCTCTAACGAGGCGTTTGAAGCG
GTTGCGATCAACAGCGAGCTTCACATTTTTTTTGCTGACTACCAGTCCAACGCGAGGGTGACCCCGAGTATTCAAGCGTGCTAGCGCCATCATACCTTTG
CCATGAACCTTTAAGTCTGCTTGTTCAAACACGTGACTAAAATCCCCAGCGTTTAGCAAACGTAAGCACCGGGGAAAGCCTTGATGCGGCACACGCAATC
CGAGATCAGGCGCTCAGACGCTTGCGGCCTTTGGCGCGGCGGCGTGCGATAACGGCACGGCCGTTTTTGGTTGCCATACGAGCACGGAAGCCATGCGCGC
GCTTGCGCTTGAGAACGCTGGGTTGAAAAGTGCGTTTCATAACTCGTTATTCCCACGTGGTTTAGGACGGAAAGTGAATTCCAAAGACCCGGTCCAGTG

AGGAGCTAGCCGGGTAGATTTAATTCAAGACCGGAAATTGTAGAGACTTCCCGCGTCGGGTGCAATTTTTCCTTGCACAAAAAGCATCAAAAAGGCAATG
GCGATTTCGCTGACCGCTAGTCCATTGCTTGAACAAGACCCACTATTATTACTAATTAAGTAGTTTAAAACCTACTGTTATTAATAGTGGGGATACAAAT
**TGTGGATAA**GCCATTTATTATCAAATTTATCAACAGATTAAAATCATAAAAAATCTGTGCATAAACATAGAATAAGAACTAAATAAGCTGTGAATTAGGT
GTGATCAATGGATATGAACCCATCCATTTATGGCTTTATCCTCATCCCCTCCCAGGCTTTTCCCAATAGCTTACGAAAAGTTGTCCACAGGCCAAAAAGA
GCCTTGAGTCCGTGTGGATAACCGCTTCGTTGAGCGCTACAATGGTCGGCCGTTTTCAACCGATGGTGAGATGAGTGTCACTCGCGCTTTGGCAACAGTG

CCTGGACTACTTGCAGGACGAACTGAATTCACAGCAGTTCAATACCTGGATACGTCCTTTGCAGGCAGAAGAAGGAGGGTCCAATGAGCTGCGCTTATTG
GCGCCGAACCGCTTTGTGCGCGACTGGGTGAGTGATAAGTACGCTAAACGGATAAGTGAATTGATCCGTGAGCTTGCGCCCTCCAAGCCTCCCAAAGTCA
GTTTAACGGTGGGTAGCCGACGCCCCGCGTCACCAGCGCCACAACCCAGAGAGCTAGGCAACCCTGTCTCGGCTGCACCTTCTCAGCCAACGGCACCAGC
TATACATACGCCACCACGTGGTGATATCGCGGATGAACGGGAGATAGATCAACTGCGTGAAGAGATACCTGGTCGGCGTGCTAATGAGCGTCAGGTACAG
GTCGAGGGCAGTCTGAAGCATAATAGCGGCCTCAATCCTAATTTTACTTTTGATACCTTTGTTGAAGGTAAATCGAACCAGTTGGCCCGTGCGGCCTCCC
GTCAGGTTTCAGAGAATCCTGGTGGTGCCTATAACCCGTTGTTTTTATATGGTGGCGTTGGTTTAGGTAAAACCCACTTAATGCATGCGGTCGGTAATGC
GTTAGCTGGTCGTCGTGAAAACGCCCGGGTGGTTTATCTGCACTCAGAACGCTTTGTTGCTGATATGGTAAAAGCATTGCAGCTAAACGCGATTAATGAT
TTCAAGCGCTTTTATCGCAGTGTCGATGCCTTATTAATTGATGATATTCAATTTTTTGCAGGGAAAGAGCGCTCACAGGAAGAGTTTTTCCACACCTTCA
ATGCTCTTTTGGAGGGGGGGCAGCAGATGATCCTCACCTCGGATCGTTATCCTAAAGAGATCAGTGGTGTTGAAGAGCGTTTGAAATCGCGCTTTGGATG
GGGGCTCACCGTTGCAATTGAACCGCCGGAGCTTGAAACCCGGGTGGCTATTTTGATGAAAAAAGCCGATCAGGCTAACGTTGATTTACCCCATGATGCC

>Halomonas olivaria TYRC17 [NZ_JALPZO010000066.1:171740..172239], <mnmG oriC2 <COG1126 (glnQ), 552 bp

TCGAGCCCTTTAATTGAGCGTACTACCTGAAGTTGCACGTCAAAAGGCAGCGAGGTGGAGATACCGTTGGGGTAAAGCTCGTGCGTATCCAGGCCTTCCG
GTTCGATAAACACCTGATGACTCGATTTGTCGGCAAAACGGTGCACTTTGTCTTCAATCGACGGGCAGTAGCGCGGACCAATACCTTCAATCACCCCTGA
ATACATGGGCGAGCGATCAAGATTGCCAAGAATAATCTCGTGGGTACGCTCATTGGTGTGCGCAATATGGCAGCTCACCTGCTGGGGATGCATGTCACGC
GACCCAAGGTAAGACATCACGGGCGTTGGTGTGTCACCGGGCTGCTCTTCAAGCTGAGAGAAGTCAACCGTTTTGGCATCAATACGTGGCGGTGTACCGG
TTTTTAGCCGGTCGACGCGAAATGGTAAAGCACGCAAGCGTTCCGCCAATGCGTAGGAAGGCGGATCGCCTGCCCGACCACCACGACTTTGATCCAGCCC
GATATGGATGACACCACCAAGGAACGTACCTGTGGATAACACGACGGACTGTGCATGAAAACGAATGCCTGTTTCGGTGATAACACCACGAACAGTATCG
TTATCCACTATCAGGTCACCCGCTGCCTGCTGAAAGATCGTCAAGTTGGGTTGGTTTTCCAGCATTCCACGAATCGCCGCTTTGTAGCGAATGCGATCAG
CTTGAGCACGGGTGGCACGCACAGCGGGCCCTTTACGGGCGTTGAGGACACGAAACTGAATGCCACCTGAATCTGTGGCAAGCCCCATTGCACCACCAAG
TGCATCAATTTCTTTAACCAGGTGGCTTTTGCCAATTCCACCAATTGCCGGGTTGCACGACATTTGGCCGAGCGTCTCGATATTGTGGGTTAGCAATAGG
GTTTGACAACCCATACGAGCAGAGGCCAGTGCGGCTTCAGTTCCCGCATGGCCACCGCCGATGACAATCACGTCAAAGCGGTCGGGATAATTCAAGAGAC

ACCTCTTCTTGTGCCATTCGGCACGAATGTTGTGAGTCAGTACATCGGTACTGATCGTAAGCCCGCCAGTATAAACCTCCTGTGGGTAACCGTCAGGTTT
TTCCACACCCCATCCTAACCAGTCAGCTACTATTAATAACAAAATAAAAATATATAAGAGAAGTTCTGTTAATGTTGTAACTAATGGTGTGGACAAATTC
**TGTGTATAA**CTCAATTTATTCCATATATATCAATTGATTAAATTGTTAACCTATGCGTGGTTAAAGGGCGGAGTGCCTGCGTAGAAGCGGTGTCTGACCT
GTGGATGGAATGCTGCCTTATCCACAGGGCTGGATGATAAGAGGTTATCCACCGCTCCATACCTTGTTTGTTACCGCACCTGTGAACAAGACTGCATTGA
GGTGATTAAACCCTCTGAAGGCCTAGCGGGAGGGAGTTACAAGGCAAACGGGAGGGGAAAATAAAAAATAGTATCCACAGGCGAAAAAAATGGCCTGTCG

CTCTGACAGGCCATTAAATGAGCAAAAATCAGGGGTATGGCGTCGCTTTAATGCTTCAATACCCGTGCTAAAAATGACCTGGTTCGTTCATGCTGAGGGG
TATCAAATAGCTGTTCTGGGGTTCCTTGCTCGGTTATCTCTCCTTTATGGATAAAGATGATGCGATCGGCTACTTCACGTGCAAAGCCCATTTCATGGGT
GACAATCACCATCGTCATTCCCTCTTTAGCCAGCTCACGCATGGCATCAAGCACTTCCCCAATCATTTCAGGGTCAAGAGCGGAGGTTGGCTCATCAAAT
AGCATGAGCCTGGGTTCCATGGCCAGTGCACGGGCCAGCGCCACGCGTTGCTGCTGGCCGCCAGAGAGCTGGCTGGGATATTTGTCTGCCTGGTCGGCAA
TGCCTACCCGCTCTAACAGACGTTTTGCCGTCTCCTCGGCATCCTGGCGGTTCCAGCCGCGTACTTTCATGGGGGCGAGGGTGATGTTATCGAGCACGCT
AAGGTGGGGAAACAGGTTGAACTGCTGAAACACCATACCCACTTCGGTACGAATGGTTTGCAGGGCTTTACTGCTTTTGCCGTTGGGCAACAGCTCATTG
CCATCCACATCCAGTGAGCCGGACTGAAACTCTTCCAGGCCATTAATGCAGCGAATCAGTGTTGATTTGCCTGAACCGCTGGCGCCAATGATGACCACGA
CTTCGCCAGGGACAATCTCAAGGTCAATAGTATTGAGCACGTGCAGGCTGCCAAAATGCTTATTGAGCTTCTGCATACGCACAATGGGTTGTGTAACGTT
GTTCATTGCGCTTGTCCTTATTGTCCGACGAGGCCTCTACGCTCCAACTGGCGCAGGATGATGGAGAGGGTCCAGGTGATGGCGAGATACATCAGTGCCA
CCATGAAATAGACCTCAAGGGCGGTGAAGGTGGTCGCGATGTAAATTTGCCCCTGGCGTACCAGTTCACCCACGCCAATCACGGAGAAGAGTGATGTATC

>Halomonas olivaria TYRC17 [NZ_JALPZO010000066.1:190675..190724], dnaA> ig dnaN>, 50 bp

WebSkew GCmin pos. nd (GCmin<>oriC1 nd; GCmin<>oriC2 nd; oriC2<>oriC1 21.8 kb)

>Halomonas boliviensis LC1 [NZ_JH393257.1:c1297567..1298066], <rpmH oriC1 dnaA>, 534 bp

AAGAACATCGGTAGTAACCGCAATGAAGTCGCGGCTTGATGTCTCGCTTTGGGTATCCGGGATACCTTCACCAACGGCGCTTTGCGGTGAAGAAGTACTC
GGAACAGCTAAGCTGCCCTCTTGATCTTCACCGTTGCTAGGAGACGCCGAATTGCTGCGTTGGGTTGTTTCAGGCACTGAATCATAATTCGGTTGCCCGT
AATCCTGATTCCACTGGACGACTAGCAAGTACGCTAGTACGGCTAGTGGAATCAGTAATAAAAGTCGTTTTACGTCCATGAGGGTTCTGCCCGCTGGGTG
GTGAACATGCTAATGGCGCAGTGTTATTTACTCGACTCTTGACACACGAAAGCCTGCCAGTTGGCAGGCCAAGGTCGAGCGGCGAGAGGTGCACCAAAGG
TCACGTGTCACGCTTAGGCGTTGGTACTGCGATCATCGCCTGTGCGTCACGCTGAAGTCGCTTCCACATGCCGTGTAACTGGCGATGCAAGGTTTCGTTA
TCTATATCCTGAACACCGCGTCGTGCAAGCACCACTATATCCACAGAGGGTAGGTGATCTTGTCGAAGGCGAACGGATTCTCTAACCAGACGCTTGAAAC
GGTTGCGATCTACGGCGCGTTTAACATTCTTTTTGCTGACCACCAGTCCAAGGCGGGGGTGTCCCCGGGTACTCAAGCGGACTAACGCCATCATACCTTT
GCCATGTACTTTCAGGTCGGCTTGTTCAAACACGTGACTAAAATCCCCAGCATTTAATAACCGTAATTGCCGGGGAAAGCCTTGCTGCGGCACACGCAAT
CTGAGATCAGGCGCTCAGACGCTTGCGGCCCTTGGCACGACGACGTGCGAGGACGGCGCGACCGTTTTTGGTAGCCATACGAGCACGGAAGCCATGCGCG
CGCTTGCGCTTGAGAACGCTAGGTTGAAAAGTGCGTTTCATAACTCGTTATTCCCACGTGGTTTAGGACGGAAAGTAAATTCCAAAAAGCCCGGTCCAGT

GAGGAACTAGCCGGGGAGATTCAATTCAAGACCGGAAATTGTAAAGACTTCCTCTGCTGGGTGCAATTTTTCCTTGCATAAAAAGCCTTATCACGCCCCA
GCCAGCTTGCTTGATGCTGAAAGAGTAGCAGCCAATAAGGTCACTATTATTACTAATAGGTAGTTCTAAAACTATTGTTATTAATAGTGGGGATACTATT
**TGTGGATAA**CTGCTTTTAATTGTTATTTTTCAATAGTATGTAGTCATAAAAAATCTGTGTGTAAGTCTCGTATAAGCAGTCCATAAGGTGTTGGCTAGAT
GTGATTAACCATTGCGTAATGAGCATTTCTATCGATTGTCCTCGTTCCTCCACATGGTTTTCCCATAGGCTCATCGGTAGTTGTCCACAGGTCAACATTG
AGGACTAAACCCGTGTGGATAACCGAGGCTTTGGGCGCTACAATGGTCGGCCGTTTTCAACCGATGGTGAGATGAGTGTCACTGACGCTTTGGCAACAGT

GCCTGGACAACCTGCAGGATGAGCTGAATTCACAGCAGTTCAATACCTGGATACGCCCGCTGCAGGCAGAGGAAGGAGAGTCCAACGAGCTGCGCTTATT
AGCGCCGAATCGCTTTGTGCGCGATTGGGTGAGCGATAAGTATGCCAAGCGGATTAGTGAGCTGATGCGTGAACTCGCACCGGCCAAACCGCCAAAGGTA
AGCCTGACGGTGGGTAGCCGCCGCTTGGCAACCCAGGCCCCTCAGCATCGTGATTTAGGTAATCCTGTATCCGCTTCACCATCGCGGCCCGTATCGCCAG
CGGTTCATACGCCGCCACGCGGCGATATTGCCGATGAACGCGAAATCGATAGGCTGCGCGATGAAGCGCCGAGTCGGCGCAATAATGAACGTCAGGTACA
GGTAGAAGGCAGCCTGAAACATGGCAGTGGGCTTAATCCAAATTTTACTTTCGATACATTTGTGGAAGGTAAATCGAACCAACTGGCTCGCGCTGCCTCC
CGCCAAGTATCTGAAAACCCTGGCGGCGCTTATAACCCTTTGTTTTTATACGGTGGCGTGGGCCTTGGTAAAACCCACTTAATGCATGCGGTGGGGAACG
CACTGGCAACCCGGCGTGAAAACGCCCGGGTGGTGTACCTGCACTCAGAGCGGTTCGTCGCCGATATGGTTAAAGCGCTTCAGCTTAACGCGATTAATGA
TTTTAAACGCTTTTACCGCAGTGTTGATGCGCTACTTATTGATGACATTCAGTTTTTTGCTGGAAAAGAGCGCTCGCAAGAAGAATTCTTTCATACCTTC
AATGCACTTTTGGAAGGTGGGCAGCAAATGATCCTCACCTCTGATCGTTATCCTAAAGAGATCAGTGGTGTAGAGGAGCGCCTTAAATCACGCTTTGGTT
GGGGCCTAACGGTAGCGATTGAACCGCCGGAGCTTGAAACCCGCGTTGCTATTTTGATGAAAAAGGCCGACCAAGCTAAGGTCGATTTGCCTCACGATGC

>Halomonas boliviensis LC1 [NZ_JH393257.1:c1319364..1319863], <mnmG oriC2 <COG1126 (glnQ), 595 bp

GCGCACCACCTGAATCTGCACGTCAAAGGGCAGCGACGTCGAGATGCCGTTGGGGTAGAGCTCATGGGTATCCAAGCCTTCGGGTTCAATAAACACCTGA
TGGCTGGATTTATCGGCAAAGCGATGAACCTTGTCTTCGATTGACGGGCAATAGCGCGGCCCAATCCCTTCGATCACGCCCGAATACATTGGCGAGCGAT
CAAGGTTTGCCAGGATGAGATCGTGGGTCTGCTGATTGGTATGCGCTATGTGACAACTGACCTGCTCCGGGTGCATTTCCCGTGACCCCAGGTAGGACAT
AACGGGCGTCGGGCTATCGCCGGCTTGCTCTTCCAGCTGAGAAAAATCTACCGTTTTAGCATCAATCCGCGGTGGTGTGCCGGTTTTCAGACGATCAACG
CGAAACGGCAGTGCGCGTAGTCGTTCGGCCAGCGCATTGGAAGGCGGATCACCGGCGCGACCACCTTTGCTTTGATCCAGACCAATATGAATCACGCCAC
CCAAAAACGTCCCGGTGCACAGCACGACGCTTTCGGCGTTAAAACGAATGCCGGTTTCCGTGACGACACCGCGAACGGTGTCGTTATCCACAATCAGATC
GCCAGCGGCCTGTTGAAAAATCGTCAAGTTGGGCTGGTTTTCCAGCATTCCGCGGATAGCGGATTTATAGCGGATACGGTCAGCCTGGGCACGGGTGGCC
CGAACCGCAGGCCCTTTGCTGGCATTGAGCACACGAAACTGGATGCCACCTAAATCCGTGGCAAGCCCCATGGCGCCGCCCAATGCATCAATTTCCTTGA
CCAAATGACTTTTACCGATGCCGCCAATGGCGGATTACACGACATTTGGCCCAGGGTTTCGATGTTGTGGGTGAGCAGCAGGGTCTGACAGCCCATGCG
AGCAGAGGCCAGTGCGGCTTCGGTTCCCGCATGGCCACCGCCGATGACAATCACGTCAAAGCGGTCGGGGTAATTCAAGAGACACCTCGTCTTGTGCCCA

TCGGCACGGATGATGTGACCTGCGCAGTCGGTTTAAGACGCGGCCACCTTAGATCAATGAGGCGGCAAGTATAAACCCCCTGTGGGTAACCGTCAGGTTT
TTCCACACCCACAATCCCTAAACAGCCTCTATTAATAACAACATAAATATAAAATAAGAGAAGTTCTGTAAATGTTGTAACTACTAGTGTGGACAATTTC
**TGTGCATAA**GCCACTTATATTCAATAATTTCAGGTGTTTATATTGTTGACTTCTACATGTTTGAAAGCGGGAGTGCCTGGGCAGAAAGGGTGTTAAGCCT
GTGGATGGAATAGCGCTTTATCCACAGATGTAATTAGGCACGGCTTATCCACCGGCCAGTACCTGGGTTGTTACCGCAGCTGTGAACAACCTCCGTTGGG
TGTGGATCTAACGCTAAGAGACTCCCATGATAGGCCTTACGAGCAAAAAAACGATCGATGAAAAAAAGGTTATCCACAGGCAAAAAAATGGCCTGTCGT

GTCGACAGGCCAGCAAAAAAGCGTAATTCAGCGCGAATAAGCGGTTATTTTAAAGCCTCAAAGGGTGTTTAATTTAATGTTTAAGCACGCGCGCAAGGAA
GGATTGGGTGCGTTCGTGTTGCGGAGAATCGAACAGTTTCTCAGGGTGGCCCTCTTCAGTAATCTCGCCTTTATGGATAAAAATAACCCGGTCGGCGACC
TCGCGGGCAAAGCCCATTTCATGGGTGACAATCACCATGGTCATGCCTTCTTTGGCCAGCTCGCGCATGGCATCCAGCACTTCACCGATCATTTCCGGGT
CAAGCGCCGATGTGGGCTCGTCAAACAGCATCAGACGCGGCTCCATGGCCAAGGCGCGAGCCAGCGCCACACGCTGCTGCTGGCCGCCTGAGAGCTGGCT
GGGGTACTTATCGGCCTGGTCGGCGATACCCACCCGATCAAGCAATCGTTTGGCATTGCTGACCGCCTCGTCACGGCTTAAACCGCGTACTTTCATCGGC
GCCAGGGTGACGTTATCGATGACGCTGAGGTGGGGAAAGAGATTGAACTGCTGAAAGACCATGCCGACTTCGGTGCGAATGGTTTGCAGCGCTTTGGTGC
TCTTGCCATGGGGCAGTAGCTCATTGCTATCGACTACTAGACTGCCGGACTGGAACTCTTCCAAGCCATTAATACAGCGAATTAGCGTCGATTTGCCTGA
GCCACTGGCACCGATAATCACGACCACCTCACCAGGCACAATGGTAAGATCAATATCGTTGAGTACGTGCAGGCTGCCAAAGTGTTTATTGAGCTTCTGC
ATACGCACAATGGGGTGCGTTGCGCTGTTCATTGTATTGCTCATTGCGCGTGTCCTTATTGTCCGGCCAGGCCTCTGCGCTCAAGTTGGCGCAGGATGAT
GGAGAGCGTCCAGGTGATCGCCAGATACATCAGGGCGACCATGAAGTAGACCTCAAGGGCCGTAAAGGTGGTGGCGATATAGATTTGCCCCTGGCGCACC

>Halomonas boliviensis LC1 [NZ_JH393257.1:c1296065..1296124], dnaA> ig dnaN>, 60 bp

WebSkew GCmin pos. 84587 (GCmin<>oriC1 24.8 kb; GCmin<>oriC2 1.3 kb; oriC2<>oriC1 23.5 kb)

>Halomonas maris QX-1 [NZ_JABWCV010000006.1:c59558..60057], <rpmH oriC1 dnaA>, 535 bp

ATGAAATCACGGCTAGACGTTTCGCTTTCAGTATCGCCAGGAATACCTTCACCAACCGCATTTGGCTGCGATGTGGTACTCGGAACGGCTAAGCCGCCCT
CGCCATCGTCAACATTGCTGGATGGTACTGAATTGCTAGATGGTGCTGAATTACTGCGTTGAGTCGTTTCAGGCACTGAATCGTAATTCGGCTGCCCGTA
ATCCTGATTCCACTGGACGACTAGCAAGTACGCCAGTACGGCTAGTGGAATCAGTAATAAAAGTCGTTTTACGTCCATGAGGGTTCTGCCCGCTGGGTGG
TGAACATGCTAATGGCGCAGTGTCATTTACTCGACCCTTGACACACGAAAGCCTGCCAGGCGGCAGGCCGAGGTCGAGCGGCGAGGTGTGCACCAAAGGT
CACGTGTCACGCTCTGGCGTTGGTGCTGCGATCACCGCCTGTGCGTCACGCTGAAGTCGCTTCCACATGCCGTGTAACTGGCGATGCAAGGTGTCGTTGT
CGATATCCTGAACGCCGCGTCGGGCAAGCACCACGATATCCACAGAGGGTAAGTGATCTTGACGAAGACGGACAGATTCTCGAACGAGACGCTTGAAACG
GTTGCGATCTACGGCGCGTTTGACATTTTTTTTGCTGACTACCAGTCCAAGGCGGGGGTGCCCCCGAGTACTCAAGCGAACTAACGCCATCATACCTTTG
CCATGCACTTTCAGGTCGGCTTGCTCAAACACGTGACTGAAATCCCCAGCATTTAGTAACCGTAACTGCCGGGGAAAGCCTTTCTGCGGCACACGCAATC
CGAGATCAGGCGCTCAGACGCTTGCGGCCCTTGGCACGGCGGCGTGCGAGGACGGCGCGACCGTTTTTGGTTGCCATACGAGCACGGAAGCCATGCGCGC
GCTTGCGCTTGAGAACGCTAGGTTGAAAAGTGCGTTTCATAACTCGTTATTCCCACGTGGTTTAGGACGGAAAGTGAATTCCAAAAAGCCCGGTCCAGTG

AGGAACTAGCCGGGGAGATTCAATTCAAGACCGGAAATTGTAGAGACTTCCTTTGCTGGGTGCAATTTTTCCTTGCATAAAAAGCGTTATGGCGTCATTG
CGGCCATGCTGGCAGTGACGAATATAACAATCAACACTCTCGCTATTATTACTAATTAGGTAGTTCTAAACCTGTTGTTATTAATAGTGGGGATACTATT
**TGTGGATAA**GTGGTTTTAATTATTATTTATCAATCACCTATAATAATAAAAAATCTGTGTGTAAGTTTCGTATAAACAGTCTACAAGGTGTTAAGTAGGT
GTGTTTAACCCAAACGTAAGTCGCTTATTTGTTAATTGTCCCCGTTCCTCCACACGGTTTTCCCGTAGGCTCACTACAAGTTGTCCACAGGACAATATTC
AATCTTAAACCCGTGTGGATAACCGAAGCTTTGGGCGCTACAATGGTCGGCCGTTTTCAACCGATGGTGAGATGAGTGTCACTGACGCTTTGGCAACAGT

GCCTGGACAACCTGCAGGATGAGCTGAATTCACAGCAGTTCAATACCTGGATACGCCCGCTGCAGGCAGAAGAAGGAGAGTCCAACGAGCTGCGCTTATT
GGCGCCGAATCGCTTTGTGCGCGATTGGGTGAGCGATAAGTATGCCAAGCGGATTAGTGAACTAATGCGTGAGCTCGCGCCGGCCAAGCCGCCGAAAGTG
AGTTTGACGGTAGGTAGCCGCCGCCCGGCAACGCAGGCTCCTCAGCATCGCGACTTAGGCAACCCCGTATCGGCTTCTCCATCGCGTCCTTCGGCACCGG
CTGTTCATACGCCTCCGCGCGGCGATATTGCCGATGAGCGTGAAATAGACAGGCTGCGTGATGAAGCTCCCAGTCGGCGTAACAATGAGCGTCAGGTACA
GGTAGAGGGCAGCCTGAAACATGGCAGTGGGCTAAATCCTAATTTTACTTTCGATACTTTTGTTGAAGGTAAATCGAACCAACTGGCTCGCGCTGCCTCA
CGCCAGGTGTCTGAAAACCCCGGCGGCGCCTATAACCCGCTGTTTTTATATGGCGGCGTGGGCCTTGGTAAAACTCACTTGATGCACGCCGTGGGTAATG
CACTGGCAACGCGGCGCGAAAATGCCCGCGTGGTTTATCTGCACTCAGAGCGTTTTGTCGCCGATATGGTCAAGGCGTTACAGCTCAATGCGATTAACGA
CTTTAAGCGCTTTTATCGCAGTGTTGATGCATTACTTATTGATGATATTCAGTTTTTCGCCGGCAAAGAGCGTTCCCAGGAAGAGTTTTTTCATACCTTC
AACGCACTGTTGGAAGGTGGCCAGCAAATGATCCTTACCTCTGATCGCTATCCGAAAGAGATCAGTGGGGTAGAGGAGCGCCTGAAATCACGCTTTGGTT
GGGGGCTCACGGTGGCGATTGAACCGCCGGAACTTGAAACCCGCGTAGCCATCTTGATGAAAAAGGCCGACCAAGCTAAGGTCGATTTGCCTCACGATGC

>Halomonas maris QX-1 [NZ_JABWCV010000006.1:c83015..83514], <mnmG oriC2 <COG1126 (glnQ), 591 bp

CGTACGCACGACTTGAATCTGCACGTCAAACGGCAGCGAGGTCGAGATACCGTTGGGGTAAAGCTCGTGGGTATCCAGGCCTTCGGGTTCGATGAACACC
TGATGGCTGGATTTATCGGCAAAGCGATGAACCTTGTCTTCGATCGATGGGCAATAACGCGGCCCAATTCCTTCGATCACACCGGAGTACATCGGCGAGC
GATCAAGGTTCGCCAGAATCAGATCGTGGGTTTGCTGATTGGTGTGAGCAATGTGGCAACTCACCTGCCGAGGGTGCATTTCCCGTGACCCCATGTAGGA
CATGACCGGTGTAGGGCTGTCACCGGGTTGTTCTTCCAACTGGGAAAAATCCACGGTTTTGGCATTAATACGCGGCGGTGTGCCGGTTTTCAGCCGATCG
ACACGAAATGGCAGCGCACGCAGCCGTTCGGCCAGTGCGTTGGAAGGCGGATCACCTGCTCGACCACCCTTGCTTTGATCCAAGCCAATGTGAATCACAC
CACCCAAAAATGTGCCGGTACACAGCACGACACTTTCAGCATGGAAGCGAATACCGGTTTCCGTGACAACGCCACGCACGGTGTCGTTATCCACAATGAG
ATCACCAGCAGCCTGCTGGAAGATCGTCAAGTTGGGCTGGTTTTCCAGCATACCGCGGATAGCGGATTTATAGCGGATGCGGTCAGCCTGGGCACGGGTT
GCCCGAACGGCAGGCCCTTTGCGGGCATTGAGCACGCGAAACTGGATGCCACCTAAATCCGTGGCAAGCCCCATGGCGCCGCCCAGCGCATCGATTTCCT
TAACCAAATGGCTTTTGCCGATGCCGCCAATCGCGGGGTTACACGACATTTGGCCCAGGGTTTCGATGTTGTGGGTGAGCAGCAGGGTCTGACAGCCCAT
ACGAGCAGAGGCCAATGCGGCTTCAGTTCCCGCATGGCCACCGCCGATGACAATCACGTCAAAGCGGTCGGGATAATTCAAGAGGCACCTCGTCTTGCGC

CTTTCGGCACGGATGATGTGGCCAGTGCAGTTCTCAAACATGGCCACCTTAGATCAATGAGCCGGCAAGTATAAACCCCCTGTGGGTAACCGTCAGGTTT
TTCCACATCCCAAATCACCAACCCTCCGTTATTAATAACAACATAAATATAAAATAAGAGAAGTTCTGTAAATGTTGTAATTACTAGTGTGGACAATTTC
**TGTGCATAA**GCGATTTATTCTTAATAATTTCATGCTCTTATATTGTTGACTTCTACATGTTTGAAGGCGGGAGTGCCTGGGTAGAAGGGGTGCTAAGCCT
GTGGATGGAATAGTGCTTTATCCACAGTTTCAATAAAGCACGGCTTATCCACCGGCCAGTACCTGGGTTGTTACCGCACCTGTGAACAACCACCGATCAA
TGTGGAGCTAACGCTAAGCGACGTACATGGTAGGCCTTACGGAGATAAAACCGATCAATGTAAAAAAAGTTTATCCACAGGCAAAAAAATGGCCTGTCGT

GTCGACAGGCCAGTAAAAAAGCGTAATTCAGAGCAAATTAGCGGTTATTTTAACGCGTAAAAAGGGCTCAATTTAATGTTTGAGTACGCGAGCAAGGAAG
GATTGAGTACGTTCGTGTTGAGGAGAATCGAACAGTTTTTCAGGGTGGCCCTGCTCTGTAATCTCGCCTTTATGGATAAAAATCACCCGGTCGGCGACCT
CGCGGGCAAAGCCCATTTCATGGGTGACAATCACCATGGTCATGCCCTCTTTGGCCAGCTCGCGCATGGCATCCAGCACTTCGCCGATCATTTCCGGGTC
AAGCGCCGATGTGGGCTCGTCAAACAGCATCAGGCGGGGTTCCATCGCCAGGGCGCGGGCGAGTGCGACGCGCTGCTGTTGACCGCCTGAGAGCTGGCTG
GGGTACTTGTCGGCCTGGTCGGCGATACCTACCCGGTCAAGCAAGCGTTTGGCGTTGCTGACTGCGTCGTCACGGCTTAAACCACGCACTTTCATCGGCG
CCAGAGTGACATTATCGATCACGCTGAGATGGGGAAAGAGATTGAATTGCTGAAACACCATACCCACTTCGGTACGGATGGTTTGCAGCGCTTTAGTGCT
CTTGCCGTGCGGTAATAGTTCGTTGCTGTCGACCACCAGACTGCCGGACTGAAACTCTTCCAAGCCGTTAATACAGCGAATTAGTGTGGATTTGCCTGAG
CCACTGGCACCGATAATCACCACCACTTCACCAGGCACAATCGTCAGATCAATATCGTTGAGTACGTGCAGACTGCCAAAGTGTTTATTGAGCTGCTGCA
TACGCACAATTGGTTGCGTTGTGTTGCTCATTGTCTCGCTCATTGCGCGTGTCCTTATTGTCCGGCCAGGCCTCTGCGCTCGAGTTGACGCAGGATGATG
GAGAGAGTCCAGGTGATTGCCAGGTACATCAGGGCGACCATGAAGTAGACCTCAAGCGCCGTAAAGGTGGTGGCGATATAGATCTGCCCCTGACGCACTA

>Halomonas maris QX-1 [NZ_JABWCV010000006.1:c58056..58115], dnaA> ig dnaN>, 60 bp

< page 1 | #090 | Halomonas glaciei DD 39

WebSkew GCmin pos. 66204 (GCmin<>oriC1 18.9 kb; GCmin<>oriC2 1.5 kb; oriC2<>oriC1 17.4 kb)

>Halomonas glaciei DD 39 [NZ_JACCDE010000017.1:c47059..47558], <rpmH oriC1 dnaA>, 534 bp

AATGAAATCACGGCTAGACGTTTCGCTTTCAGTATCGCCAGGAATACCTTCACCAACCGCATTTGGCTGCGATGTGGTACTCGGAACGGCTAAGCCGCCC
TCGCCATCGTCAACATTGCGGGATGGTACTGAATTGCTAGATGGTACTGAATTACTGCGTTGAGTCGTTTCAGGCACTGAATCGTAATTCGGCTGCCCGT
AATCCTGATTCCACTGGACGACTAGCAAGTACGCCAGTACGGCTAGTGGAATCAGTAATAAAAGTCGTTTTACGTCCATGAGGGTTCTGCCCGCTGGGTG
GTGAACATGCTAATGGCGCAGTGTCATTTACTCGACCCTTGACACACGAAAGCCTGCCAGACGGCAGGCCGAGGTCGAGCGGCGAGGTGTGCACCAAAGG
TCACGTGTCACGCTCTGGCGTTGGTGCTGCGATCACCGCCTGTGCGTCACGCTGAAGTCGCTTCCACATGCCGTGTAACTGGCGATGCAAGGTGTCGTTG
TCGATATCCTGAACGCCGCGTCGGGCAAGCACCACGATATCCACAGAGGGTAAGTGATCTTGACGAAGACGGACAGATTCTCGAACGAGACGCTTGAAAC
GGTTGCGATCTACGGCGCGTTTGACATTTTTTTTGCTGACTACCAGTCCAAGGCGGGGGTATCCCCGAGTACTTAAGCGAACTAACGCCATCATACCTTT
GCCATGCACTTTCAGGTCGGCTTGCTCAAACACGTGACTGAAATCCCCAGCATTTAGTAACCGTAACTGCCGGGGAAAGCCTTTCTGTGGCACACGCAAT
CCGAGATCAGGCGCTCAGACGCTTGCGGCCCTTGGCACGGCGACGTGCGAGGACGGCGCGACCGTTTTTGGTTGCCATACGAGCACGGAAGCCATGCGCG
CGCTTGCGCTTGAGAACGCTAGGTTGAAAAGTGCGTTTCATAACTCGTTATTCCCACGTGGTTTAGGACGGAAAGTGAATTCCAAAAAGCCCGGTCCAGT

GAGGAACTAGCCGGGGAGATTCAATTCAAGACCGGAAATTGTAGAGACTTCCTTTGCTGGGTGCAATTTTTCCTTGCATAAAAAGCGTTGTGGCGTCATT
GCGGCCATGCTGGCAGTGAGAATTATAGCAATCTATGTTCTGGCTATTATTACTAATAAGTAGTTCTAAACCTGTTGTTATTAATAGTGGGGATACTATT
**TGTGGATAA**GTGGTTTAATTATTATTTATCAATTGCCTATAGACATAAAAAATCTGTGTGTAAGTTCCGTATAAACAGTCTACAAGATGTTAAGTAGAT
GTGTTTAACCTAAACGTAAGTCGCTTTTTTGTTAATTATCCCCGTTCCTCCACATGGTTTTCCCGTAGGCTCACTGCAAGTTGTCCACAGGGCAATATTC
AAGCTTAAACCCGTGTGGATAACCGAGGCTTTGGGCGCTACAATGGTCGGCCGTTTTCAACCGATGGTGAGATGAGTGTCACTGACGCTTTGGCAACAGT

GCCTGGACAACCTGCAGGATGAGCTGAATTCACAGCAGTTCAATACCTGGATACGCCCGCTGCAGGCAGAAGAAGGAGAGTCCAACGAGCTGCGCTTATT
GGCGCCGAATCGCTTTGTGCGCGATTGGGTGAGCGATAAGTATGCCAAGCGGATTAGTGAACTAATGCGTGAGCTCGCGCCGGCCAAGCCGCCGAAAGTG
AGTTTGACGGTAGGTAGTCGCCGTCCGGCCACGCAGGCACCTCAGCATCGCGACTTGGGCAACCCCGTATCGGCTTCTCCATCGCGTCCTTCGGCACCGG
CTGTTCATACGCCTCCGCGCGGCGATATTGCCGATGAGCGTGAAATAGGCAGGCTGCGTGATGAAGCTCCCAGTCGGCGTAATAATGAGCGCCAGGTACA
GGTAGAGGGCAGCCTGAAACATGGTAGCGGACTAAACCCGAATTTTACTTTCGATACTTTTGTTGAAGGTAAATCAAACCAACTGGCTCGCGCTGCCTCC
CGCCAGGTGTCTGAAAACCCCGGCGGCGCTTATAACCCGCTGTTTTTATACGGCGGCGTGGGTCTTGGTAAAACTCACTTGATGCACGCCGTGGGTAATG
CGCTGGCAACGCGGCGCGAAAATGCCCGCGTGGTCTATCTGCACTCAGAGCGTTTTGTCGCCGATATGGTCAAGGCGTTACAGCTCAATGCGATTAACGA
TTTCAAGCGCTTTTATCGCAGTGTTGATGCATTACTTATTGATGATATTCAGTTTTTCGCTGGCAAAGAGCGTTCCCAGGAAGAGTTTTTTCATACCTTC
AATGCGTTATTAGAAGGTGGTCAGCAAATGATCCTGACCTCTGACCGCTATCCTAAAGAGATCAGTGGGGTAGAGGAGCGCCTTAAATCACGCTTTGGTT
GGGGGCTCACGGTCGCGATTGAACCGCCGGAACTTGAAACCCGCGTAGCCATCTTGATGAAAAAGGCCGACCAAGCTAAGGTCGATTTACCTCACGATGC

>Halomonas glaciei DD 39 [NZ_JACCDE010000017.1:c64481..64980], <mnmG oriC2 <COG1126 (glnQ), 594 bp

ACGCACGACTTGAATCTGAACGTCAAACGGCAGCGAGGTCGAGATGCCGTTGGGGTAGAGCTCATGGGTATCCAAACCTTCGGGCTCGATAAACACCTGA
TGGCTCGATTTATCGGCAAAGCGATGAACCTTGTCTTCAATCGATGGGCAGTAACGCGGCCCGATGCCTTCGATCACACCGGAATACATCGGCGAGCGAT
CAAGGTTGGCCAGAATCAGATCGTGAGTCTGCTGATTGGTGTGAGCAATATGGCAACTCACCTGCTGAGGATGCATTTCCCGTGATCCCAGGTAGGACAT
AACCGGCGTTGGGCTATCGCCGGGCTGCTCTTCCAACTGGGAAAAATCCACGCTTTTGGCATCAATACGCGGCGGTGTGCCGGTTTTTAGGCGATCAACG
CGAAACGGCAGCGCGCGCAGACGTTCGGCCAACGCGTTGGAGGGCGGATCGCCTGCTCGGCCACCTTTGCTTTGATCCAAACCAATGTGAATCACGCCAC
CCAAAAATGTCCCGGTACACAGCACGACGCTGTCGGCATGAAAGCGGATGCCGGTTTCCGTGACGACGCCTCGTACGGTGTCGTTATCCACAATCAGATC
GCCTGCGGCTTGCTGGAAGATCGTTAAGTTGGGCTGGTTTTCCAGCATTCCGCGGATCGCTGCCTTGTAACGGATGCGGTCAGCCTGGGCACGAGTGGCC
CGAACGGCGGGCCCTTTGCTGGCATTGAGCACGCGAAACTGGATGCCACCCAAGTCCGTGGCAAGCCCCATGGCGCCGCCCAGTGCATCAATTTCCTTGA
CCAAGTGACTTTTGCCGATGCCACCGATGGCGGGATTACACGACATTTGGCCCAGAGTTTCGATGTTATGGGTGAGCAGCAGGGTCTGACAGCCCATGCG
AGCAGAGGCCAGTGCGGCTTCAGTTCCCGCATGGCCACCGCCGATGACAATCACGTCAAAGCGGTCGGGGTAATTCAAGAGACACCTCGTCTTGTGCCTA

TCGGCACGGATGATGTGGCCAGTGCAGTCGATTCTAGACGCAGCCACCTTAGATCAATGAGGCGGCAAGTATAAACCCCCTGTGGGTAACCGTCAGGTTT
TTCCACACCCCAAATTGTTAAACAGTCTCTATTAATAACAACATAAATATAAAATAAGAGAAGTTCTGTAAATGTTGTAATTACTAGTGTGGACAATTTC
**TGTGTGTAA**GCGATTTATTCTTAATAATTTCATGCGCTTATATTGTTGACTTCTACATGTTTGAAGGCGGGAGTTCCTGGGCAGAAAGGGTGTTAAGCCT
GTGGATGGAATAGTGCTTTATCCACAGCTGTAATTAGCCACGGCTTATCCACCGTCCAGTACCTGGGTTGTTACCGCACCTGTGAACAATCAACGATTGG
TGTGGAGCTAACGCTAAGCGATGTACATGGTGGGCCTTACGGAGATAAAACCGATCAATGTAAAAAAAGTTTTTCCACAGGCAAAAAAATGGCCTGTCGT

GTCGACAGGCCAGTAAAAAAGCGTTATTCAGCGTGAATTGGCGGTTATTTTAACGCGTAGAAGGGGCTCAATTTAATGTTTGAGCACGCGAGCAAGGAAG
GATTGAGTACGTTCGTGTTGAGGAGAATCGAACAGTTTTTCAGGGTGGCCCTGCTCGGTAATCTCGCCTTTATGGATAAAAATCACCCGGTCGGCGACCT
CGCGGGCAAAGCCCATTTCATGGGTAACAATCACCATGGTCATGCCCTCTTTGGCCAGCTCGCGCATGGCATCCAGCACTTCACCGATCATTTCCGGATC
CAGCGCCGATGTGGGCTCGTCAAACAGCATCAGGCGGGGTTCCATCGCCAAGGCACGAGCCAGTGCCACACGTTGCTGTTGACCGCCTGAGAGCTGGCTG
GGGTACTTGTCAGCCTGATCAGCGATACCTACCCGGTCAAGCAAGCGTTTGGCGTTGCTGACCGCGTCGTCACGGCTTAAACCGCGCACTTTCATTGGCG
CCAGGGTGACGTTATCGATCACGCTGAGATGAGGAAAGAGATTAAATTGCTGAAAGACCATGCCCACTTCGGTACGGATGGTTTGCAGCGCTTGGGTACT
CTTCCCATGGGGCAATAGTTCGTTGCTGTCGACCACCAGGCTGCCCGATTGGAACTCTTCCAAACCGTTAATACAGCGAATTAGCGTGGATTTGCCAGAG
CCACTGGCACCGATGATCACGACGACTTCACCAGGCACAATCGTTAGGTCAATATCGTTAAGTACGTGCAGGCTGCCAAAGTGTTTATTAAGCTGCTGCA
TACGCACAATGGGTTGCGTTGTGTTGCTCATTGTCTCGCTCATTGCGCGTGTCCTTATTGTCCTGCCAGGCCCCTGATCTCAAGCTGGCGGAGGATGATT
GATAGGGTCCAGGTGATTGCCAGGTACATCAGGGCGACCATGAAGTAGACCTCAAGCGCCGTAAAGGTGGTGGCGATATAGATCTGCCCCTGACGCACTA

>Halomonas glaciei DD 39 [NZ_JACCDE010000017.1:c45557..45616], dnaA> ig dnaN>, 60 bp

WebSkew GCmin pos. 138225 (GCmin<>oriC1 22.6 kb; GCmin<>oriC2 1.6 kb; oriC2<>oriC1 20.9 kb)

>Halomonas sulfidaeris SST4 [NZ_QNTU01000008.1:160542..161041], <rpmH oriC1 dnaA>, 533 bp

GGACATCAGTCGTCACGGCAATGAAATCGCGACTTGATGTCTCGCTTTCAGTATCGTCAGGAATACCTTCTCCAACGGCATTTTGCGGTGAAGAGGTACT
TGGAACGGCCAAGTCACCGTCCCCATCTTCAATATCGCGGGAAGGAGCCGAGCTGCTGCGTTGAGTTGTCTCGGGCACTGAATCGTAATTCGGCTGCCCG
TAATCCTGATTCCACTGGACGACTAGCAAGTACGCAAGCACGGCTAGTGGAATCAGTAATAAAAGTCGTTTTACGTCCATGAGGGTTCTGCCCGCTGGGT
GGTGAACATGCTAATGGCGCAGTGCCTTTTGCTCGACCCTTGACACACGAAAGCCTGCCAGGCGGCAGGCCGAGGTCGAGCGGTGAGGTGTGCACCAAAG
GTCACGTGTCACGCGCTGACGTTGCTACTGCGATCACCGCCGGTGCGTCACGCTGCAGTCGCTTCCACATACCGTGTAACTGGCGATGCAAGGTGTCGTT
ATCGATGTCCTGAACACCGCGCTTGGCAAGCACCACGATATCCACAGAGGGTAAGTGATCTTGACGAAGACGGACGGATTCCCGAACGAGACGCTTGAAG
CGGTTGCGATCTACGGCGCGTTTGACATTTTTCTTGCTGACTATTAATCCAAGGCGGGGATGCCCCCGTGTACTCAAGCGAACTAACGCCATCATACCTT
TGCCATGCACTTTTAGATCAGCTTGCTCAAATACGTGACTAAAATCCCCAGCATTTAGTAACCGTAATTGCCGGGGAAAGCCTTGCTGCGGCACACGCAA
TCCGAGATCAGGCGCTCAGACGCTTGCGGCCCTTGGCACGGCGACGTGCGAGGACGGCGCGACCGTTTTTGGTTGCCATACGAGCACGGAAGCCATGCGC
GCGCTTGCGCTTGAGAACGCTAGGTTGAAAAGTGCGTTTCATAACTCGTTATTCCCACGTGGTTTAGGACGGAAAATGAATTCCAAAAAGCCCGGTCCAG

TGAGGAACTAGCCGGGGAGATTCAATTCAAGACCGGAAATTGTAGAGACTTCCTCTGCTGGGTGCAATTTTTCCTTGCATAAAAAGCTTCCCCGCGCCGT
AGCGATCCGTTTGGCGTACGGTAAGTACCGCTCTTGGCTACGACTATTATTACTAATAGGTATTTCTAAACCTGTTGTTATTAATAGTGGGGATACTATT
**TGTGGATAA**GCTGATTTATTTTCTAATTTTCAATTGCCTATGTATATAAAAAATCTGTGTGTAAGTCTCGTATAAGCGGTCCAAAAGGTGTTGGTTAGGT
GTGGTTAAGCTTCTAGAAAAGGGCATTGTGAAAAATTGTCCCCTTTCCTCCACACGGTTTTCCCATAGGCTCATGTCAAGTTGTCCACAGGCCAGTATTG
AGGGCTAAACCCGTGTGGATAACCGAGGCTTTGGGCGCTACAATGGTCGACCGTTTTCAACCGATGGTGAGAAGAGTGTCACTGACGCTTTGGCAACAGT

GCCTGGACAACCTGCAGGATGAGCTGAATTCACAGCAGTTCAATACCTGGATACGCCCGCTGCAGGCAGAAGAAGGAGAGTCCAACGAGTTGCGCTTATT
GGCGCCGAATCGTTTTGTGCGCGATTGGGTGAGCGATAAGTATGCCAAGCGAATCAGTGAGCTGATGCGTGAACTCGCGCCAGCAAAACCGCCGAAAGTA
AGTTTAACGGTGGGTAGCCGTCGCTTGGCAACACAAGCACCTCAGCATCGAGATTTAGGAAATCCTGTCTCGGCTTCTCCATCGCGACCTTCGGCACCCG
CAGTGCATACCCCTCCGCGCGGTGATATCGCCGATGAGCGCGAAATCGACAAGTTGCGCGATGAAGCGCCAGGGCGACGTAATAATGAGCGTCAGGTACA
GGTAGAAGGCAGTCTGAAACATGGTAGTGGGCTAAATCCAAATTTTACTTTCGATACCTTTGTTGAAGGTAAATCGAACCAACTGGCCCGTGCTGCCTCG
CGACAAGTCTCAGAAAACCCTGGTGGTGCTTATAATCCATTGTTTTTATATGGTGGCGTGGGGCTTGGTAAAACCCACTTAATGCATGCCGTGGGCAACG
CATTGGCAACCCGGCGTGAAAATGCCCGTGTGGTATATCTGCACTCAGAGCGTTTCGTCGCCGACATGGTCAAGGCGTTACAGCTCAATGCGATTAATGA
TTTCAAACGCTTTTATCGCAGTGTTGATGCATTACTGATTGATGACATTCAGTTTTTTGCTGGCAAAGAGCGTTCGCAAGAAGAATTTTTTCATACCTTC
AACGCACTTTTGGAAGGTGGCCAGCAAATGATACTTACCTCTGATCGCTATCCTAAAGAGATTAGCGGTGTAGAGGAGCGTCTGAAATCACGCTTTGGTT
GGGGCCTCACGGTGGCGATTGAACCGCCTGAGCTTGAAACCCGTGTCGCGATCTTGATGAAAAAAGCTGACCAAGCTAAGGTCGATTTGCCTCACGACGC

>Halomonas sulfidaeris SST4 [NZ_QNTU01000008.1:139636..140135], <mnmG oriC2 <COG1126 (glnQ), 593 bp

ACGCACCACCTGAATCTGCACGTCAAAGGGCAGCGAGGTCGAGATGCCGTTGGGATAGAGCTCATGGGTATCCAAGCCTTCGGGTTCGATAAACACCTGG
TGGCTGGATTTATCGGCAAAACGATGCACTTTGTCTTCAATCGATGGGCAGTAACGCGGCCCGATGCCTTCGATGACACCGGAGTACATCGGTGAGCGAT
CAAGGTTGGCCAGAATCAGATCGTGGGTCTGTTGATTGGTGTGAGCAATATGACAACTCACCTGCCGAGGGTGCATCTCCCGCGACCCCAGGTAGGACAT
CACCGGCGTTGGGCTATCGCCTGGCTGCTCTTCCAACTGAGAAAAATCAACGGTTTTGGCATCGATACGCGGCGGCGTACCGGTTTTTAAGCGATCAACG
CGAAACGGCAGTGCGCGCAGACGTTCGGCCAGCGCGTTGGAGGGCGGATCGCCTGCGCGGCCACCTTTGCTTTGATCCAAACCAATGTGAATCACGCCAC
CCAAAAATGTCCCGGTACACAGAACGACGCTTTCAGCATGAAAGCGAATACCGGTTTCCGTCACCACGCCACGTACGGTGTCGTTATCCACAATGAGATC
ACCAGCGGCCTGCTGGAAAATCGTCAAATTTGGTTGGTTTTCCAGCATTCCGCGGATCGCTGACTTGTAACGAATGCGGTCAGCCTGGGCACGGGTGGCC
CGAACGGCGGGCCCTTTGCTGGCATTGAGCACGCGAAACTGAATCCCACCTAAATCCGTGGCAAGCCCCATAGCGCCGCCGAGTGCATCAATTTCCTTGA
CCAAATGACTTTTGCCGATGCCGCCAATGGCGGGGTTACACGACATTTGGCCGAGGGTTTCGATGTTGTGGGTGAGCAGCAGGGTCTGACAGCCCATGCG
AGCAGAGGCCAATGCGGCTTCGGTTCCCGCATGGCCACCGCCGATGACAATCACGTCAAAGCGGTCGGGATAATTCAAGAGACACCTCGTCTTGTGCCTA

TCGGCACGGATGATGTGGCCAGCGTCGGCAATTTCTGACGGCGCTACCTTAGATCAATGAGCCGGCAAGTATAAACCCCCTGTGGGTAACCGTCAGGTTT
TTCCACACCCGGATTACTCAGAACAACTCTATTAATAACAACATAAATATAAAATAAGAGAAGTTCTGTAAATGTTGTAATTACTAGTGTGGACAATTTC
**TGTGTATAA**GCAATTTATCAATAACAATTTCAATAGTTTATATTGTTGACTTCTACGTGTTTGAAGGAGGGATTCCCTGGGTAGAAGAAGTGCTAAGCCT
GTGGATGAAATAGTGCCTTATCCACCGATACAACTAGGCACCGCTTATCCACCGGCCAGTACCTGGGTTGTTACCGCACCTGTGAACAACGCTAGACTGG
TGTGGAGCTAACGCTAAGAGATGCCCATGGTAGGCCTTACAGAGATAAAGTCGCACAATGTAAAAAACTTTATCCACAGGCAAAAAAAAGCCTGTCGTG

TCGACAGGCCAGTAAAAAAGCGTCATTCAGCGAAAATTAATGCTTATTTTAAAGCGCTAACAGGGCTCAATTTAGTGTTTTAGCACGCGTGCAAGGAAGG
ATTGAGTACGTTCATGTTGAGGCGAATCGAACAGTTGTTCAGGCGGCCCCTGCTCGGTAATTTCACCTTTATGGATAAAAATAACCCGGTCGGCGACTTC
GCGGGCGAAGCCCATTTCATGGGTAACAATCACCATGGTCATGCCTTCTTTAGCCAGCTCGCGCATGGCATCCAGTACTTCCCCGATCATTTCCGGGTCA
AGCGCCGAGGTGGGCTCGTCAAACAGCATTAAGCGCGGTTCCATCGCTAGGGCACGGGCCAGTGCTACCCGTTGCTGTTGGCCGCCTGAGAGCTGGCTGG
GGTACTTATCGGCTTGGTCGGCGATGCCCACTCGGTCTAGTAGGCGCTTAGCGTTACTAATGGCGTCATCACGGCTTAAACCACGGACTTTCATGGGGGC
CAGAGTGACGTTGTCAATCACGCTGAGATGAGGAAATAGATTGAACTGCTGAAACACCATGCCCACTTCGGTACGGATGGTTTGCAGCGCTTGTGTGCTC
TTTCCGTGTGGTAACAGTTCATTGTTATCGACGACCAAGTTGCCCGTTTGGAACTCTTCCAAGCCGTTAATACAGCGAATCAGCGTGGATTTACCTGAGC
CACTGGCACCGATAATCACTACCACCTCGCCAGGTACAATTGTAAGGTCAATGTCGTTGAGTACGTGCAGGCTGCCAAAGTGCTTATTGAGCTGCTGCAT
ACGCACAATAGGTTGCGTTGTGCTGTTCATTGTATCGCTCATTGCGCGTGTCCTTATTGTCCTGCCAGGCCTCTGCGCTCAAGTTGGCGCAGGATGATGG
AGAGCGTCCAGGTAATGGCTAGATACATCAAGGCGACCATGAAGTAGACCTCAAGCGCCGTGAAGGTGGTGGCGATATAGATCTGCCCTTGACGAACCAG

>Halomonas sulfidaeris SST4 [NZ_QNTU01000008.1:162484..162547], dnaA> ig dnaN>, 64 bp

WebSkew GCmin pos. 64860 (GCmin<>oriC1 19.0 kb; GCmin<>oriC2 1.4 kb; oriC2<>oriC1 17.7 kb)

>Halomonas utahensis NBRC 102410 [NZ_BJXV01000011.1:c45550..46049], <rpmH oriC1 dnaA>, 534 bp

GAGGACATCGGTAGTCACCGCAATGAAATCGCGGCTAGACGTCTCGCTTTCGGTATCGCCGGGGATACCTTCACCCACGTTGCCTTGTGGCGCCGAGGTG
CTTGGCACCGCTAAGCCCTCGCTATCCTCAACGTTGCTGGGAGGCGCCGAGCTGCTGCGTTGCGTCGTTTCAGGTACCGAATCGTAATTCGGCTGCCCGT
AATCCTGATTCCACTCGACGACTAGCAGGTACGCCAGTACGGCTAGTGGAATCAGTAATAAAAGTCGTTTAACGTCCATGAGGGTTCTGCCCGCTGGGTG
GTGAACATGCTAATGGCGCAGTGTCATTTACTTGACTCTTGACACACGAAAGCCTGCCAGACGGCAGGCCGAGGTCGAGCGGCAAGGGGTGCACCTAATG
TCACGTGTTACGCTCCGGCGTTGGTGCTGCGATCATCGCCTGTGCGTCACGCTGAAGCCGCTTCCACATACCATGTAACTGGCGATGCAAGGTGTCGTTG
TCTATATCCTGAACGCCGCGCCGGGCAAGCACCACAATATCCACAGAGGGTAAGTGATCTTGACGAAGACGTAAGGATTCTCGAACGAGACGCTTAAAAC
GGTTGCGGTCGACGGCGCGCTTAACATTTTTTTTGCTGACCACCAGTCCAAGGCGAGGGTGTCCCCGGGTACTTAGGCGAACTAATGCCATCATACCTTT
GCCATGTACTTTTAGATCGGCTTGCTCAAACACGTGACTGAAGTCCCCAGCATTTAGTAACCGTAATTGCCGGGGAAAGCCTTGCTGCGGCACACGCAAT
CCTAGATCAAGCGCTCAGACGCTTGCGGCCTTTGGCACGACGACGTGCCAGGATGGCGCGACCATTTTTGGTAGCCATACGAGCACGGAAGCCATGCGCG
CGCTTGCGCTTAAGAACGCTGGGTTGAAAAGTGCGTTTCATAACTCGTTATTCCCACGTGGTTTAGGACGGAAAGTGAATTCCAAAAAGCCCGGTCCAGT

GAGGAACTAGCCGGGGAGATTCAATTCAAGACCGGAAATTGTAGAGACTTCCTCCGCTGCGTGCAATTTTTCCTTGCACAAAAAGCGTTGTCACCTCCCA
GCCAGCTTGCGCGGCCGGTCTAGAGTAGTCGCCCATGTTGTCACTATTATTACTAATAGGTAGTTCTAAACCTGTTGTTATTAATAGTGGGGATACTATT
**TGTGGATAA**GTGGTTTTAGTGTCTTTATTTCAACAACCTATAGGTATAAAAAATCTGTGTGTAAGTCGCGTATAAGCAGGAGATAAGCTGTTATTTTGAT
GTGATTAACCGCTGTGTAAGCAGTGATATTGAGGTTTATCCCTATCTATTCACAACGTTTTCCCCGTGGCTTATGACAAGTTGTCCACAGGGCGATATTG
ATGTCTAAACCTGTGTGGATAACCGCGGCTTTGGGCGCTACAATGGTCGGCCGTTTTCAACCGATGGTGAGATGAGTGTCACTGACGCTTTGGCAACAGT

GCCTGGACAACCTGCAGGATGAACTGAATTCACAGCAGTTCAACACCTGGATACGCCCACTGCAGGCAGAAGAAGGAGAGTCCAACGAGCTGCGCTTATT
AGCGCCGAATCGCTTTGTGCGCGATTGGGTGAGCGATAAGTATGCCAAGCGGATTAGTGAGCTGATGCGTGAGCTGGCACCGGCCAAACCACCTAAAGTC
AGTTTGACGGTAGGTAGTCGTCGCCCAGCTAACCACGCACCTCAGCATCGTGATTTGGGCAACCCCGTATCGGCGTCACCATCGCGGCCTTCAGCGCCGG
CGGTGCATACGCCGCCGCGTGGCGATATACCCGATGAGCGCGAATTAGATAAGCAACGCGAAGCGGCTTCGGGTCGGCGTACCAATGAGCGTCAGGTACA
AGTCGAAGGCAGCTTGAAACACGGCAGCGGTCTAAATCCAAACTTTACCTTTGACACCTTTGTGGAAGGTAAGTCGAATCAGCTCGCCCGCGCGGCGTCG
CGCCAAGTGTCAGAAAACCCCGGTGGTGCTTACAACCCCTTGTTTTTATACGGGGGAGTGGGCCTGGGTAAAACCCACCTAATGCATGCGGTGGGTAATG
CGTTAGCCACCCGGCGCGAAAATGCGCGGGTTGTCTATCTGCATTCAGAGCGTTTTGTGGCGGATATGGTCAAAGCACTACAGCTAAATGCGATTAATGA
TTTTAAGCGCTTTTACCGTAGCGTCGACGCCTTATTAATCGATGATATTCAGTTTTTCGCTGGCAAAGAGCGGTCGCAGGAGGAGTTCTTTCACACCTTC
AATGCGCTTTTGGAAGGCGGCCAACAGATGATTTTGACCTCTGATCGCTATCCGAAAGAGATCAGTGGTGTTGAAGAGCGGCTTAAATCACGTTTTGGTT
GGGGCCTGACGGTAGCGATTGAGCCGCCGGAGCTTGAGACCCGCGTGGCTATCCTGATGAAAAAGGCTGACCAGGCGAAGGTCGATTTGCCCCACGATGC

>Halomonas utahensis NBRC 102410 [NZ_BJXV01000011.1:c63206..63705], <mnmG oriC2 <COG1126 (glnQ), 617 bp

CGCACCAGCTGAATTTGCACGTCAAACGGCAGCGAGGTCGAGATCCCGTTGGGATACAGCTCGTGCGTATTTAAGCCTTCGGGTTCAATAAACACCTGAT
GGCTGGATTTATCGGCAAAGCGATGAACTTTGTCTTCGATTGACGGGCAATAGCGCGGCCCAACCCCTTCGATCACGCCCGAATACATCGGTGAGCGATC
AAGGTTTGCCAGAATCAGATCGTGGGTCTGCTGATTGGTATGCGCTATGTGGCAGCTGACCTGCTGGGGGTGCCGATCTCGCGAGCCCAGGTAGGACATC
ACCGGCGTTGGCGTATCGCCTGGCTGCTCTTCCATCTGGGAAAAGTCAACGCTTTTGGCATCAATGCGCGGCGGCGTGCCGGTTTTTAACCGGTCAACGC
GAAACGGTAATGCCCGCAGCCGTTCGGCCAGCGCATTGGAAGGCGGGTCGCCCGCTCGGCCGCCTTTGCTTTGGTCCATGCCAATGTGGATCACGCCACC
TAAAAACGTCCCGGTACTCAACACAACGCTGTCGGCGTGAAAGCGAATGCCGGTTTCCGTAACAACGCCACGCACGGTGTCGTTATCCACAATGAGATCG
CCAGCGGCCTGTTGGAAAAGCGTTAAATTTGGCTGGTTTTCCAACATGCCACGGATAGCGGCTTTGTAAAGAATACGGTCAGCCTGAGCACGGGTTGCCC
GAACGGCTGGTCCCTTGCGGGCATTGAGCACGCGAAATTGGATACCGGCTAAATCCGTGGCAAGCCCCATGGCGCCGCCCAACGCATCAATTTCCTTGAC
CAAATGGCTTTTGCCGATGCCGCCAATCGCGGGGTTACACGACATTTGACCCAGGGTTTCGATGTTGTGGGTGAGCAGCAGGGTCTGACAGCCCATGCGA
GCAGAGGCCAGTGCGGCTTCAGTTCCCGCATGGCCACCGCCGATGACAATCACGTCAAAGCGGTCGGGGTAATTCAAGAGACACCTCGTCTTGTGCCTAT

CGGCACGGATGATGTGAGCCGGTGCAGTCAGCTCCAGACATGAGCACCGTAGATCAATGAGCGGGGCAGTATAAACCCCCTGTGGGTAACCGTCAGGTTT
TTCCACACCCGGATTCACCAGAGAGGCTCTATTAATAACAACATAAATATAAATTAAAAGAAGTTCTGTAAATGTTGTAATTACTAGGGTGGACAATTTC
**TGTGTATAA**GCGATTTATTGCTATTTTTTTCAACAGTTTACTTTGTTGACTTCTACATGTTTGAAAGCGGGACTGCCTGGGTAGAAGGGGGGTTAAGCCT
GTGGATGAAATAGCGCTTTATCCACAGAGATAGTTAAGCACCTTTTATCCACCGGCCAGTACCTGGGTTGTTACCGCACCTGTGAACAACCGTGGCGGGA
TGTGGAGCTAACGCTAAGAGACTCGCAGGATAGGCCTTACAGAGGTATAGAGGAGCAAAGTAAAAAAAGTTTATCCACAGGCAAAAAAATGGCCTGTCAC

GTGGACAGGCCGGTAAAAAACGCCATTTAGCAGGGTCAAACATATTATTTTGGTGCCGAGTGCAGGTCTACGCGAGGCGATTAAGTGATGACATTTAGTG
TTTAAGTACCCGTGCAAGGAAGGATTGGGTGCGCTCATGTTGCGGCGTATCGAACAGTTTTTCAGGGGGGCCCTGTTCAGCAATTTCGCCTTTATGAATA
AAAATAACCCGGTCGGCGACTTCGCGGGCAAAGCCCATTTCATGGGTAACAATCACCATGGTCATGCCCTCTTTGGCCAGCTCGCGCATGGCATCGAGCA
CTTCACCGATCATTTCCGGGTCAAGCGCCGATGTGGGCTCGTCAAACAGCATAAGGCGCGGTTCCATGGCCAATGCGCGGGCTAGCGCCACACGCTGCTG
CTGGCCGCCTGAGAGATGGCTGGGGTACTTGTCGGCTTGGTCGGCGATACCCACACGGTCAAGCAAGCGTTTGCCGCTACTGATGGCGTCATCACGACTC
AAACCGCGTACTTTCATGGGGGCTAGGGTAACGTTGTCGAGCACACTGAGATGGGGGAAAAGATTGAACTGCTGGAACACCATACCCACTTCGGTGGAAT
CTTTTGCAGCGCTTGGGTGCTCTTTCCATAGGGCAGCAGTTCGCTGCTATCGACGACCAGATTGCCCGACTGAAACTCTTCCAAGCCGTTTATACAGCGG
ATCAATGTCGATTTACCCGAGCCACTGGCACCGATAATTACGACCACTTCGCCAGGCACAATGTCGAGGTCGATGTTGTTGAGTACGTGCAGGCTGCCAA
AATGCTTATTGAGCTGCTGCATACGCACAATTGGCTGCGTTGTATCGTTCATTACGCGTGTCCTTATTGTCCAGCCAGGCCTCTGCGCTCTAGCTGGCGC
AGCAGGATTGAGAGCGTCCAGGTGATCGCTAGATACATCAGCGCGACCATGAAATAGACCTCAAGGGCGGTGAAGGTAGTGGCGATATAAATTTGTCCCT

>Halomonas utahensis NBRC 102410 [NZ_BJXV01000011.1:c44048..44107], dnaA> ig dnaN>, 60 bp

WebSkew GCmin pos. 62682 (GCmin<>oriC1 20.7 kb; GCmin<>oriC2 0.9 kb; oriC2<>oriC1 21.6 kb)

>Halomonas profundi MT13 [NZ_JAHRFY010000004.1:c41757..42256], <rpmH oriC1 dnaA>, 534 bp

AAGAACGTCGGTCGTCACGGCAATAAAATCGCGGCTTGTTGACTCGCTTTCAGTGCTGCTTGCTATGCCTTCTCCAACCGGATCTTGCTGCGGAGAACTG
CTGGGAACGGCTAAGCCATCGTCAGTTTCGACATTGCTAGGCACTGCCGAGTTGCTGCGTTGCGTCATTTCAGGAGCCGAATCAAAATTTGGCTGCCCGT
AATCCTGATTCCACTGGACGACTAACAAATAGGCTAGTACGGCTAGTGGAATCAGTAATAAAAGTCGTTTTACGTCCATGAGGGTTCTGCCCGCTGGGTG
GTGAACATGCTAGTGGCGCAGTGTCATTTACTCGTCCCTTGACACACGAAAGCCTGCCGGGGGGCAGGCCGAGGTCGAGCGGCGAGGTGTGCACCAAATG
TCACGTGTTACGCTCTGGCGTTGGTGCTGCTGTTACCGCCTGTGCGTCACGCTGGAGTCGCTTCCACATGCCATATAACTGGCGATGTAAGGTGTCGTTA
TCGATATCCTGAACACCCCGTCGAGCAAGAACCACGATATCCACAGAGGGTAAGTGGTCTTGGCGAAGGCGCACGGATTCTCGTACGAGACGCTTGAAAC
GGTTGCGGTCTACGGCGCGTTTAACATTTTTTTTGCTGACTATCAGTCCAAGGCGAGGGTAACCCCGAGTACTTAGGCGAACTAACGCCATCATACCTTT
GCCATGCACTTTTAAGTCGGCTTGTTCAAACACGTGACTGAAGTCCCCAGCATTTAGTAACCGTAATTGCCGGGGAAAGCCTTGCTGCGGCACACGCAAT
CCGAGATCAGGCGCTCAGACGCTTGCGGCCTTTGGCACGGCGGCGTGCTAGGACGGCGCGACCGTTTTTGGTTGCCATACGAGCACGGAAGCCATGCGCG
CGCTTGCGCTTGAGAACGCTAGGTTGAAAAGTGCGTTTCATAACTCGTTATTCCCACGTGGTTTAGGACGGAAAGTGAATTCCAAAAAGCCCGGTCCAGT

GAGGAACTAGCCGGGGAGATTCAATTCAAGACCGGAAATTGTAGAGACTTCCTACGCTGGGTGCAATTTTTCCTTGCATAAAAACCCTTGCAGCGGCATC
GCTGCGCTGCTTTTGGGGCATTGTTAGGCAATCGACCCTACCACTATTATTACTAGTTAGTAGTTCTAAACCTGTTGTTATTAATAGTGGGGATACTATT
**TGTGGATAA**GTGATTTTAATAGCCATATTTCAATAGCCTATAATGATAAAAAATCTGTGTGTAAGTGTCGTATAAGCGGTCTAAAAGGTGTTGGTTAGAT
GTTTATAACCTTTAAGAAAGCGTTATTATTAATATTTATCCCCCTTTCCTCCCGAGCTTTTCCCGTAGGCTCATCGCGAGTTGTCCACAGGCCAATATGG
GTCGCTAAACCCATGTGGATAACCGGGGCTTTGAGCGCTACAATGGTCGGCCGTTTTCAACCGATGGTGAGATGAGTGTCACTGACGCTTTGGCAACAGT

GCCTGGACAACCTGCAGGATGAACTGAATTCACAGCAGTTCAATACCTGGATACGCCCGCTGCAGGCAGAAGAAGGAGAGTCCAACGAGCTGCGCTTATT
GGCGCCGAATCGCTTTGTGCGCGATTGGGTGAGCGATAAGTACGCCAAGCGGATAAGTGAGCTGATGCGTGAACTCGCACCCGCGAAACCGCCTAAGGTC
AGCCTGACGGTAGGCAGTCGGCGCCCGGCAACACAGGCACCTCAGCATCGTGATTTAGGTAACCCTGTATCGGCTTCGCCGTCGCGCCCTGCCTCAGCAG
CGGTGCATACGTCATCGCGTGCAGATACCGCCGACGAGCGCGAAATCGATAAATTGCGTGAGCAGCAGGCGTCTGCGCGGCGCCCCAATGAGCGCCAGGT
ACAGGTTGAGGGCAGCCTGAAACATGGCAGCGGCCTTAATCCTAATTTTACCTTTGATACTTTTGTTGAAGGTAAATCGAACCAGCTGGCACGCGCGGCG
TCGCGTCAGGTGTCTGAAAATCCTGGTGGTGCCTATAACCCGCTGTTTTTATACGGTGGAGTGGGGCTTGGTAAAACTCACTTGATGCATGCTGTTGGCA
ATGCGTTAGCGACCCGGCGTGAAAATGCTCGCGTTGTCTATCTGCATTCCGAGCGATTTGTTGCCGATATGGTGAAGGCATTACAGCTTAATGCGATTAA
TGATTTTAAGCGCTTTTACCGCAGCGTCGATGCCTTACTAATTGATGATATTCAGTTTTTTGCGGGCAAAGAGCGCTCACAAGAAGAGTTCTTCCATACG
TTCAATGCGCTCTTGGAAGGCGGCCAACAGATGATCCTGACCTCTGATCGCTATCCGAAAGAGATCAGCGGTGTAGAAGAGCGTCTGAAATCACGCTTTG
GCTGGGGGCTTACGGTAGCGATTGAGCCGCCGGAGCTTGAGACACGCGTGGCCATTTTGATGAAGAAAGCAGATCAAGCTAAGGTCGATTTACCTCATGA

>Halomonas profundi MT13 [NZ_JAHRFY010000004.1:c63348..63847], <mnmG oriC2 <COG1126 (glnQ), 592 bp

AGCCTTTGATAGAGCGCACAACTTGAATCTGCACGTCAAAGGGCAGCGAGGTCGAGATTCCATTGGGATACAGCTCGTGAGTGTCCAGGCCTTCGGGCTC
GATAAACACTTGATGGCTCGACTTGTCGGCAAACCGATGGACTTTATCTTCAATCGATGGGCAGTAGCGCGGGCCAACGCCTTCAATCACGCCAGAATAC
ATGGGTGATCGATCAAGGTTAGCCAGAATCAGATCGTGAGTCTGCTGGTTGGTGTGAGCAATGTGACAGCTCACTTGCTGAGGGTGCATTTCCCGCGACC
CCAGATAGGACATCACCGGCGTTGGTGTATCGCCTGGCTGTTCTTCCATCTGTGAAAAATCAACCGTTTTAGCATTAATGCGCGGCGGGGTGCCTGTTTT
TAAACGATCAACCCGAAACGGCAACGCCCTTAATCGCTCGGCGAGAGCGTTGGAAGGCGGATCACCTGCCCGGCCGCCCTTGCTTTGATCCAAGCCAATA
TGAATCACACCACCCAAAAATGTCCCGGTACACAGCACAACGCTTTGAGCATGAAAACGAATACCTGTTTCGGTCACAACGCCACGTACGGTGTCATTTT
CGACGATTAAATCACCGGCCGCCTGCTGGAAAAGCGTCAAATTCGGTTGGTTTTCCAACATTCCGCGAATAGCCGCCTTGTAGCGGACACGGTCTGCCTG
AGCCCGAGTAGCTCTAACGGCAGGCCCTTTGCGTGAATTCAGCACACGAAACTGGATGCCACCTAAATCCGTGGCAAGCCCCATAGCGCCGCCCATCGCA
TCGATTTCTTTCACCAAATGGCTTTTGCCAATGCCGCCAATCGCAGGATTACACGACATTTGGCCCAGTGTTTCGATGTTGTGAGTGAGCAGCAGGGTCT
GACAGCCCATACGAGCAGAGGCCAGTGCGGCTTCAGTTCCCGCATGGCCACCGCCGATGACAATCACGTCAAAGCGGTCGGGATAATTCAAGAGACACCT

CGTCTTGTGCCATTGGGCACGGATGATGTGAGCCAGTGCTTATGCACCCTAGTTCAATACGCCGGCAAGTATAAACCCCCTGTGGGTAACCGTCAGGTTT
TTCCACACCCACTCAAATTGAGCAGTCTTTATTAATAACAATATAAAGATAAGATAAGAGAAGTTCTGTAAATGTAATAACTATTAGTGTGGACAATTTC
**TGTGCATAA**GCAATTAATCCATTTATCTTTCATTAGGTTACATTGTTGACTTCTACATGTTTGAACAGCGTATTGCCTGGGCAGAAGCGGTGCTACTCCT
GTGGATGAAATACCCCTTTATCCACAGCATCAATTCTACACGCCTTATCCACCGTCCTATACCTGGGTTGTTACCGCACCTGTGAACATCCAGAGCATGC
CTGTTAACAAATGCTAATAGATCCCTATGGCAGGCCCTATAGACGCGAAGTGGATAAAGTAAAAAACTTTATCCACAGGCAAAAAAAAGCCTGTCACG

AGGACAGGCCGGTAAAAAAAGCAATTTAACGCGCGTTTAAGTCACCATTAATGTCAAATCAGCGCTGAAAAAGTAGGGTCGATCTAGTGTTTTAGTACCC
GTGCAAGGAAGGATTGGGTGCGCTCATTTTGTGGCGTATCAAACAGTTTTTCAGGCGCTCCCTGCTCAGCAATTTCACCTTTATGGATAAAAATAACGCG
ATCGGCGACTTCACGGGCAAAGCCCATTTCATGAGTGACGATTACCATCGTCATCCCTTCTTTGGCGAGTTCGCGCATAGCGTCCAGTACCTCACCAATC
ATTTCGGGATCAAGTGCCGATGTAGGCTCGTCAAACAGCATCAGGCGCGGTTCCATTGCCAAAGCGCGTGCCAGTGCGACACGTTGCTGTTGGCCGCCAG
AGAGCTGGCTTGGGTACTTCTCGGCCTGATCAGCGATACCCACACGGTCTAGCAGACGTTTGGCACTGCTGACAGCATCAGCGCGGCTAGCGCCGCGTAC
TTTCATCGGTGCTAACGTAACGTTGTCTTGTACGCTGAGGTGGGGAAATAGATTGAATTGCTGAAAGACCATACCGACTTCAGTACGAATGGTTTGTAAC
GCTTTAGTGCTTTTTCCATTCGGCAGAAGTTCGTTGCTATCAACCATAAGGCTGCCTGCCTGAAATTCCTCAAGGCCGTTGATACAGCGAATCAGAGTCG
ACTTACCTGAGCCGCTGGCACCGATAATCACGACCACTTCACCAGGAATAATGTCGAGATCAATGTTGTTGAGTACGTGCAGGCTGCCAAAATGTTTATT
GAGCTTCTGCATACGTACAATTGGCTGCGTTGTGTTGTTCAATGTCTCGCTCATTGCGCGTGTCCTTATTGTCCGGCCAGACCTCTACGCTCAAGCTGGC
GTAGTAAGATTGAAAGGGTCCAGGTGATCGCTAAATACATCAGTGCGACCATGAAGTAGACCTCAAGGGCGGTGAACGTGGTGGAAATGTAGATTTGTCC

>Halomonas profundi MT13 [NZ_JAHRFY010000004.1:c40252..40311], dnaA> ig dnaN>, 60 bp

WebSkew GCmin pos. 555262 (GCmin<>oriC1 25.8 kb; GCmin<>oriC2 3.0 kb; oriC2<>oriC1 22.8 kb)

>Halomonas subterranea CGMCC 1.6495 [NZ_FOGS01000001.1:581060..581559], <rpmH oriC1 dnaA>, 532 bp

GGTGGTCACCGCGATGAAGTCACGGCTGTCATTGCTGCTCTCGCCGCTGTTCAGTGCGTCCTGCGTAGCGGCAGCGCCATCACCGGGAACCGACAGGTTA
TCCGCGTCGCTGCCGCCAGACGTACCGCCACCTTCCGCCGTTGCGTTGTCATTGCTGCTTTGGCGTTGCGATTGTTCGGACGACGATTGATAAGACGTCT
GCCCGTAATCCTGATTCCACTGTACAACCAGTAGATAGGCCAGAATGGCAAGTGGAATCAGTAATATAAGTCGTTTTACGTCCATGAGGGTTCTGCCCGC
TGGGTGGTGAACATGCTAATGGCGCAGTATCAATTGCTCGACCCATGACACACGAAAGCCTGCCAGGCGGCAGGCCGAGGTCGAGCGCTGAGGTGCCCAT
CAAGTAGTCTGCTTTGAGCTTGCAGGTAGCGCGTTGGCTGCCTGGCTGTCGCGCCGAAGCCGTTTCCACATGCCATGGAGCTGACGATATAAAACGTCAT
TGTCCATGTCCTGAACGCCACGCCGAGCAAGTACCACAATATCCACTGCAGGAAGCTGGTCTTGACGTAGCCGAACCGACTCTCGAACGAGTCTTTTGAA
ACGATTTCGATCAACGGCGAGCTTCACATTTTTTTTGCTGACCACCAGGCCAAGGCGGGGATGCCCCAGGGTATTCCGGCGTGCTAACGCCATTATCCCT
TTGCCATGTACCTTGAGGTCTGCTTGCTCAAAAACGTAGCGGTAGTCCCCGGCGTTAAGTAACCGACAACACCGGGGAAACTCTTGATGTGGCACACGCA
ATCCGAGATCAGGCGCTCAGACGCTTACGGCCTTTGGCGCGGCGACGCGAAATGATAGCGCGGCCGTTTTTGGTTGCCATACGAGCACGGAAGCCATGCG
CGCGCTTGCGCTTGAGAACGCTAGGTTGAAAAGTGCGTTTCATAACTCGTTATTCCCACGTGGTTAGGACGGAAAATAAATTCCAAAAGACCCGGTCCAG

TGAGGAACTAGCCGGGAGATTCAATTCAAGACCGGAAATTGTAGAGGCTTCCCAGGTTGCGTGCAATTTTTTCCTTGTAAAAAATGCGTTCTGTGCAAGC
AGGTGTCTCTCTCGAGCCACATCATGCGCCTGCCGGGAAACATATCAGTCTCCTACAATGGGTTTTCTGCTGCCTTGTGGCTAATAGTGGGGATACATTT
**TGTTAACAA**CCTTATTTTTTTGTTTAAAATCAAGCATATAAAAGCTATTTGAGGTTGTGTATAAGTTGCGGAAAAGGCGATGATAAGTCGTTGATTAGCT
GTGAATTAATGACCGGAAATAATGAGCGCTAGCGGTTGTTCAGCTCTTGGACGGAGGGTCACTTTTTATCCTTCTGAAAGTTTTCCACAGCTTATTAAAA
CAGCCTGCTCCTGTGTGGATAACCACGCCGTTGGACGCTACAATGGTCGGCCTTTTTCAACTGATGGTGAGATGTGTGTCACTCGCGCTTTGGCAACAGT

GCTTGGATTACCTGCAGGACGAGCTGAGTTCGCAGCAGTTCAACACGTGGATACGTCCACTGCAGGCTGAAGATAGTGAGGCTAATGAGCTGCGCTTGTT
GGCACCTAATCGCTTTGTGCGCGATTGGGTGAGTGATAAGTACGCCAAGCGGATTAACGAGCTGATGCGCGAGCTCGCCCCTTCGAAACCTCCCAAGGTT
AGCCTCACGGTAGGTAGCCGTCGATCGGCCTCTCCTGCTCCCCAGCCCCGTGATCTGGGTAATCCGGTCTCAGCCTCGCCGGCACCCCCGCGCGCGTCGG
CAGCTGTTCATACGCCGCCACGTGGTGATATTGCTGACGAGCGTGAAATCGACCAGCTACGCGAGGAGGGAAGTGGCTCAGACCGACGGACGAACGAGCG
CCAGGTTCAGGTCGAGGGTGGTCTCAAGCATACCAGCGGGCTAAACCCTAATTTTACGTTTGACACCTTTGTTGAAGGTAAATCCAACCAGCTTGCGCGT
GCGGCTTCCCGCCAGGTTTCCGAAAACCCCGGGGGGGCCTATAACCCGCTATTCCTGTATGGCGGCGTCGGGCTGGGTAAAACCCACTTGATGCATGCGG
TGGGTAATGGTCTGGCGTTGCGCCGTGAAAATGCCCGTGTGGTTTATCTCCACTCAGAGCGTTTCGTGGCCGATATGGTGAAGGCGCTTCAGCTTAATGC
CATCAATGACTTTAAGCGCTTTTACCGTAGTGTCGATGCGTTACTTATTGATGATATTCAATTTTTTGCTGGAAAAGAGCGCTCTCAGGAAGAGTTCTTC
CATACGTTTAATGCGTTATTGGAAGGCGGTCAGCAAATGATTCTGACGTCGGATCGCTACCCTAAGGAAATCAGTGGGGTTGAAGAGCGTCTGAAATCCC
GCTTTGGGTGGGGCCTTACGGTTGCCATCGAGCCGCCTGAGCTGGAAACCCGGGTGGCTATCTTGATGAAGAAAGCTGATCAGGCCAAAGTCGACTTGCC

>Halomonas subterranea CGMCC 1.6495 [NZ_FOGS01000001.1:558031..558530], <mnmG oriC2 <COG1126 (glnQ), 557 bp

TGCCCTTGATCGAGCGCACAACCTGAAGCTGTATATCGAACGGCAGGGAAGTGGAAATCCCGTTGGGATACAGCTCATTGGTATCCAGACCTTCGGGTTC
GATAAATATCTGGTGGCTTGATTTATCAGCAAAGCGATGGACTTTATCTTCTATCGAAGGGCAATAGCGCGGCCCGATGCCCTCGATTACGCCCGAATAC
ATCGGCGAACGTTCGAGATTCGCCATGATGATTTCATGAGTGCGCTCGTTGGTGTGAGTAATATGACAGCTCAGCTGCTTAGGATGCATTTCCCGTGAGC
CGAGGTAGGACATTACCGGTGTAGGCGTGTCACCGGGTTGCTCTTCAAATTGAGAGAAATCAACGCTTTTGGCATCAATGCGCGGCGGCGTACCGGTTTT
CAGCCGATCAACCCGGAACGGCAGAGCACGCAGTCGTTCAGCCAGGGCGTTTGACGGCGCATCGCCCGCTCGGCCACCGCGACTTTGGTCGAGCCCGATG
TGAATTACCCCGCCGAGGAAAGTACCGGTCGAGAGCACCACGGATTCACTGTGAAATCGGATACCGGTTTCCGTTACCACGCCACGAACCGTGTTGTTGT
CGACCATGAGGTCGCCCGCAGCCTGCTGGAAAATCGTCAGGTTTGACTGATTCTCAAGCATGCCACGGATGGCTGCTTTGTAGCGTACGCGGTCAGCCTG
GGCGCGGGTTGCTCGAACGGCAGGCCCTTTGCGTGCGTTCAAGACACGAAACTGAATGCCACCTGCGTCTGTGGCAAGCCCCATGGCGCCGCCCAGCGCA
TCAATTTCCTTGACGAGATGGCTTTTGCCAATCCCGCCAATGGCCGGGTTACACGACATTTGGCCGAGTGTTTCGATGTTGTGCGTGAGCAGCAAGGTTT
GACAGCCCATGCGAGCAGAGGCCAGCGCGGCTTCAGTTCCCGCATGGCCACCGCCGATGACAATGACATCAAAGCGGTCAGGGTAATTCAAGAGACACCT

CGTCTTGTGCCATCAGGCACGGATGAGTTGAGTCGGTGCATGAGCACCTTAATTCAATAAGCCCGCCAGTATAAACCCCTTGTGGGTAACCGTCAGTTTT
TTCCACACCCCCATTGGCCCAGCAGCCCTTATTAATAACAACATAAATATTATATAAGAGAAGTTCTATTGATGTTGTTACTACTGGTGTGGACAAAATC
**TGTTGATAA**GCAAATTAAATTTATAATTTTCAATGGTTTATATTGTTGACTTTTGCATGATTAAACGGCGGACTAACTGAGTAGAAACGGTGTTGGGCCT
GTGGATGAAAGCTCGTTTTATCCACCGATGTGATAAGGCACGGGTTTTCCACTCGTGTATACCGGAGTTGTTACCGTGACTGTGAACAAGTGAAGGAAAT
GCGATCAATCCCGCGTTAAGGCGGCATGATGGTGGCTAGCGAGGGATTGAGGCCATTCAGAAAATTAGTTTATCCACAGGCAAAAAAAATGGCCTGTGGT

GATCACAGGCCATTGAAGGTCGTTTATCAAGCGTATTTTTTGGCTAACTCAGTGCTTGAGTACCCGGGCTAAAAATGACTGGGTACGTTCGTGTCGAGGC
TTGTCGAACAGCTGTTCAGGAGGTCCCTCTTCGGTGATTTCCCCCTGGTGAATGAACACCACACGGTCCGCAACTTCACGCGCGAAACCCATTTCATGGG
TAACAATGACCATGGTCATGCCTTCTTTAGCAAGCTCTCGCATGGCATCCAGCACCTCACCGATCATTTCAGGGTCCAGCGCCGAGGTAGGCTCGTCAAA
CAGCATTAACCGCGGCTCCATGGCAAGTGCACGGGCGAGCGCCACACGCTGCTGCTGACCGCCGGAAAGCTGACTGGGATACTTGTCGGCCTGGTCGGCG
ATGCCTACCCGGTCGAGGAGCCGCTTGGCAGCTTCCAGGGCATCTGAACGGTTCCAGCCGCGCACTTTCATGGGTGCAAGGGTGACGTTGTCGATAACGC
TTAAGTGAGGAAAGAGATTGAACTGCTGGAACACCATACCGACTTCAGTGCGGATGGTTTGCAGTGCCTGGCCACTTTTGCCGTTGGGCAGTAACTCGTT
ACCGTCGACCTCCAGCGAACCGGACTGGAATTCCTCCAGTCCGTTGATGCAGCGGATCAGTGTCGATTTACCCGACCCGCTGGCACCGATCACCACGACC
ACTTCACCAGGGGTGATGTGCATATCAATGTTTTTGAGTACGTGCAGGCTACCGAAGTGTTTATTGAGCTGCTGCATACGCACAATCTGCTGAGCATTGT
CGTTCATTGTGCGTCTCCTTATTGGCCTACCAAGCCTTTGTTTTCAAGCTGGCGCAAGATGATTGAAAGGGACCAGGTGATAGCGAGGTACATGATAGCC
ACCATCAGGTAGACCTCCATCGCGGTAAAGGTGCTGGCAATGTAAACCTGGCCCTGACGAACCAGCTCACCTACCCCGATCACCGAAAATAAAGAGGTAT

>Halomonas subterranea CGMCC 1.6495 [NZ_FOGS01000001.1:583011..583064], dnaA> ig dnaN>, 54 bp

WebSkew GCmin pos. 68043 (GCmin<>oriC1 23.6 kb; GCmin<>oriC2 0.8 kb; oriC2<>oriC1 22.7 kb)

>Halomonas janggokensis DSM 18043 [NZ_JARWAJ010000002.1:c44223..44722], <rpmH oriC1 dnaA>, 532 bp

GTGGTCACCGCGATGAAGTCACGGCTGTCGCTATCACTTTCGTCGCTGTTTAGCGCGTCCTGCGAAGCGGCAGTGCTATCTCCGGGAACCGACAAGTTTT
CCGCGTCGTCATTGCCAGACGTACCGCCACTTCCCGACGATGCGTTGTTATCGCGTGTTTGATCTTGCGATACCTCGGACGACGGTTGATAAGACGTCTG
TCCGTAATCCTGATTCCACTGTACAACCAGTAGATAGGCCAGAATGGCAAGTGGAATCAGTAATATAAGTCGTTTTACGTCCATGAGGGTTCTGCCCGCT
GGGTGGTGAACATGCTAATGACGCAGTATCAATTACTCGACCCATGACACACGAAAGCCTGCCAGGCGGCAGGCCGAGGTCGAGCGATGAGGTGCCCATC
AAGTAGTGTGCTTTGAGCTTGCAGGTAGCGCCTTGTCTGCCTGGCTGTCGCGCCGAAGCCGTTTCCACATGCCATGGAGCTGACGATATAAAACGTCATT
GTCCATGTCCTGAACGCCACGCCGAGCAAGTACCACAATATCCACTGCGGGAAGCTGGTCTTGACGTAGCCTGACTGACTCTCGAACGAGTCTTTTGAAG
CGATTTCGATCAACGGCGAGCTTCACATTTTTTTTGCTGACCACGAGGCCAAGGCGGGGATGCCCGAGGGTATTCCGGCGTGCTAACGCCATAATCCCTT
TACCATGTACCTTGAGATCTGCTTGCTCAAAAACGTAGCGGTAATCCCCGGCGTTAAGTAACCGAAAACACCGGGGAAACCCTTGATGTGGCACACGCAA
TCCGAGATCAGGCGCTCAGACGCTTACGGCCTTTGGCGCGGCGACGCGAAATAATGGCGCGGCCATTTTTGGTTGCCATACGAGCACGGAAGCCATGCGC
GCGCTTGCGCTTGAGAACGCTAGGTTGAAAAGTGCGTTTCATAACTCGTTATTCCCACGTGGTTAGGACGGAAAATAAATTCCAAAAGACCCGGTCCAGT

GAGGAACTAGCCGGGAGATTCAATTCAAGACCGGAAATTGTAGAGGCTTCCCGGGTTACGTGCAATTTTTTCCTTGTAAAAAATACCGCCTGTCCACGCA
GGTGCCTGTCAGGGAGCACGCAATGGTGCTTGCCGGGAAGTGCACCGGCGTCCAGCCATTGGTCTTCTGCTGCCTTGTGGCTAATAGTGGGGATACATTT
**TGTTAACAA**CTGTATATTTTTCTTTAAATTCAACTATATAAAAGCTAGATGAGGTTGTGTATAAGTTGCGGAAAAGACGATGATAAGTGGTTGATTAACT
GTGAATTAATGCCCAGCGATAATGAACAGTAAGCTGTCGATCAGCACTTGGAATATCTATCAATTCGACACCTGAAAAGTTTTCCACAGCTTATTAAAAC
GCCCCAGTCCTGTGTGGATAACCACGCCGTTGGACGCTACAATGGTCGGCCTTTTTCAACCGATGGTGAGATGTGTGTCACTCGCGCTTTGGCAACAGTG

CCTGGATTACCTGCAGGACGAGCTGAGTTCGCAGCAGTTCAACACGTGGATACGTCCACTGCAGGCTGAAGATAGTGAGGCTAACGAGCTGCGCTTGTTG
GCGCCTAACCGCTTTGTGCGCGATTGGGTGAGTGACAAGTACGCCAAGCGGATTAGCGAACTGATGCGCGAGCTTGCCCCTTCGAAACCTCCCAAGGTTA
GCCTCACGGTAGGTAGTCGACGGCCAGCAGCACCTGCCCCCCAACCCCGTGATCTGGGTAATCCGGTCTCGGCCTCTCCGGCACCGCGTGCGTCGGCGGC
TGTTCATACGCCGCCACGGGGCGACATTGCTGACGAGCGTGAAATCGATCAGCTACGCGAGGAGGGAAGTGGTTCAGACCGGCGTTCGAACGAGCGCCAG
GTTCAGGTTGAGGGTGGGCTCAAGCATACCAGCGGGCTAAATCCTAACTTTACTTTTGATACTTTCGTGGAAGGTAAATCCAACCAGCTCGCGCGCGCGG
CTTCTCGCCAGGTTTCCGAAAATCCGGGTGGGGCCTATAACCCGCTATTCTTGTATGGCGGTGTTGGTCTGGGTAAAACCCACTTGATGCATGCCGTAGG
TAATGGTCTGGCGTTGCGCCGTGAAAATGCGCGTGTGGTCTACCTTCACTCGGAGCGTTTCGTGGCCGATATGGTGAAGGCGCTACAGCTCAATGCCATC
AATGACTTCAAGCGTTTTTACCGTAGTGTTGATGCGTTACTCATTGACGATATTCAATTTTTTGCCGGTAAAGAGCGCTCTCAGGAAGAGTTCTTCCATA
CGTTTAATGCGTTATTGGAAGGCGGTCAGCAAATGATTCTGACGTCGGATCGCTACCCGAAGGAAATCAGTGGTGTCGAAGAACGCCTGAAATCACGCTT
TGGGTGGGGCCTTACTGTCGCTATTGAGCCGCCTGAGTTGGAAACCCGTGTGGCTATTTTGATGAAAAAAGCGGATCAGGCCAAAGTCGACTTGCCCCAT

>Halomonas janggokensis DSM 18043 [NZ_JARWAJ010000002.1:c66958..67457], <mnmG oriC2 <COG1126 (glnQ), 557 bp

ATTCCCTTGATCGAGCGTACGACCTGAAGCTGTATATCAAACGGCAATGAAGTGGAAATTCCGTTGGGATACAGCTCATTGGTATCCAGGCCTTCGGGTT
CGATAAATATCTGGTGGCTCGATTTATCGGCAAAGCGATGGACCTTGTCTTCAATCGAAGGGCAATAGCGTGGCCCGATGCCCTCGATCACGCCCGAATA
CATTGGTGAACGTTCGAGGTTCGCCATGATGATCTCATGGGTGCGCTCGTTGGTGTGCCCAATGTGACAGCTCATCTGTTCGGGGTGCATTTCCCTTGAG
CCCAGATAGGACATCACCGGCGTTGGCGTGTCACCAGGTTGTTCTTCAAGTTGAGAGAAATCAACGGTTTTGGCATCAATGCGTGGGGGTGTACCGGTTT
TCAGACGATCAACCCGGAACGGCAGGGCACGCAGTCGCTCAGCCAATGCATTGGAGGGCGCATCGCCTGCTCGGCCACCGCGAGTTTGATCAAGTCCGAT
GTGGATCACACCGCCGAGAAAAGTACCGGTCGAGAGCACGACGGATTCACTGTGAAACCGGATACCGGTTTCCGTCACCACGCCCCGCACGGTGTCGTTA
TCCACAATCAGATCGCCTGCAGCCTGCTGGAAAATCGTCAGATTTGGCTGGTTTTCAAGCATGCCACGGATGGCAGCTTTGTAGCGTACGCGGTCAGCCT
GGGCGCGGGTCGCTCGAACAGCAGGCCCCTTACGTGCGTTCAAGACACGAAACTGGATGCCACCTGCGTCTGTGGCAAGACCCATGGCGCCGCCTAACGC
GTCAATTTCCTTGACGAGATGGCTCTTGCCGATCCCGCCAATTGCCGGATTACACGACATTTGGCCGAGCGTTTCGATGTTGTGCGTAAGCAACAGGGTT
TGACAGCCCATGCGAGCAGAGGCCAATGCGGCTTCAGTTCCCGCATGGCCACCGCCGATGACAATGACATCAAAGCGGTCAGGGTAATTCAAGAGACACC

TCGTCTTGTGCCATCAGGCACGGATGAGTTGAGTCGGTGCATGCGCACCTTAATTCAATAAGCCCGCCAGTATAAACCCCTTGTGGGTAACCGTCAGTTT
TTTCCACACCCCGTTTGTTCAGCAGCGCTTACTAATAACAACATAAATATTATATAAGAGAAGTTCTATTGATGTTGTTACTACTGGTGTGGACAAAATC
**TGTTGATAA**GCGATTTTATCTCATTTTTATCAATACTTTATATTGTTGACTTCTACATGATTAAACGGCGGACTAACTGAGTAGAAACCGTGTTGGGCCT
GTGGATGAAAGCTTGTTTTATCCACCGCTGTGATGAGGCACGGGTTTTCCACTCGTGTATACCAGGGTTGTTACCGTGACTGTGAACAAGGGAAGGAAAT
ACGCTGGTATCGAGTTCGAAGGCGGCGTGATGCAGGTTAAAAGCAGGTTGTGACAACCTATAGAAATAAATTATCCACAGGCAAAAAAAATGGCCTGTGG

TGATCACAGGCCATCAAATGTCGGTTACCAAGCGTATTTTTTGGCTAAATCAGTGCTTGAGTACCCGGGCTAAAAACGATTGCGTACGTTCGTGTCGAGG
CGCATCGAATAGCTGTTCGGGAGGCCCCTCTTCGGTGATTTCCCCCTGGTGAATGAACACCACACGGTCGGCGACTTCACGCGCAAAGCCCATTTCATGG
GTGACGATCACCATGGTCATGCCCTCTTTAGCGAGCTCTCGCATGGCATCCAGTACCTCGCCAATCATTTCGGGGTCCAGCGCCGACGTAGGCTCGTCAA
ACAGCATTAATCGCGGTTCCATGGCAAGTGCACGCGCGAGTGCCACACGCTGCTGCTGACCGCCAGACAGCTGGCTGGGATACTTATCCGCCTGGTCGGA
GATCCCAACGCGGTCAAGCAGTCGCTTGGCAGCTTCCAGGGCGTCTGAACGGTTCCAGCCACGTACTTTCATGGGAGCGAGGGTGATATTGTCGATAACG
CTTAAATGAGGAAACAGGTTGAATTGCTGAAACACCATGCCGACTTCGGTACGGATGGTTTGTAGTGCCTGGCTACTCTTGCCGTTAGGTAGCAACTCGT
TACCGTCAACTTCCAGCGAACCGGACTGGAATTCTTCCAGTCCGTTGATGCAGCGGATCAGGGTGGATTTACCCGACCCACTTGCACCGATCACCACGAC
CACTTCGCCAGGTGTGATGTGCATATCTATGTCTTTGAGTACGTGCAGGCTACCAAAGTGTTTATTGAGCTGCTGCATACGCACAATCTGCTGAGCTTTG
TCGTTCATTGTGCGTCTCCTTATTGGCCTACCAAGCCTTTGTTTTCAAGCTGTCGCAAGATGATTGAAAGGGTCCAGGTGATAGCCAGGTACATGATGGC
CACCATCAGATAGACCTCCATCGCGGTAAAGGTGCTGGCGATATAAACCTGGCCTTGACGAACCAGCTCACCCACCCCGATTACCGAAAATAAAGAGGTA

>Halomonas janggokensis DSM 18043 [NZ_JARWAJ010000002.1:c42721..42775], dnaA> ig dnaN>, 55 bp

WebSkew GCmin pos. 296664 (GCmin<>oriC1 24.1 kb; GCmin<>oriC2 2.8 kb; oriC2<>oriC1 21.3 kb)

>Halomonas gomseomensis DSM 18042 [NZ_JARWAI010000003.1:320536..321035], <rpmH oriC1 dnaA>, 532 bp

GGTGGTGACTGCGATGAAGTCACGGCTGTCGCTATCGCTTGCGTCGCTGTCGAGCGTCTCTTGAGGTGAGGCGTCGCCGTCGCTGGGAACAGAGAGATCA
TTCGCGTCGTCGCCACGAGATGCATCGCCGCCGCCAGACCCTGCATTATTGCTGCTGGTTTGGCTTTGCGATATCTCCGACGACGATTGATAAGACGTCT
GTCCGTAATCCTGATTCCACTGGACAACCAGCAGATAGGCCAGAATGGCTAGCGGAATCAGTAATATAAGTCGTTTTACGTCCATGAGGGTTCTGCCCGC
TGGGTGGTGAACATGCTAATGGCGCAGTGTCAATTACTCGACCCTTGACACACGAAAGCCTGCCAGGCGGCAGGCCGAGGTCGAGTGGTGAGGTGCCCAT
CAAGCGTTGCGTATTGAGCCTTCAGGCGGCGCACTGGCCGACTGGGCATCGCGCCGAAGCCGTTTCCACATGCCATGAAGCTGACGGTAGACAACATCAT
TTTCCATGTCTTGGACGCCACGTCGAGCAAGCACCACGATATCCACAGCGGGAAGCTGGTCTTGACGTAGGCGGGCTGACTCGCGAACGAGCCTTTTGAA
GCAGTTACGATCAACGGCGAGCTTCACATTTTTTTTGCTGACTATCAGGCCCAGGCGCGGATGCCCGAGGGTATTCCAGCGCGCAAGCGCCATCATCCCT
TTACCATGAACCTTGAAGTCTGCCTGCTCAAAAACGAAGCGGTAATCCCCAGCGTTCAATAACCGACAACACCGGGGAAAACCTTGATGTGGCACACGCA
AGCCGAAATCAGGCGCTCAGACGCTTACGGCCCTTGGCGCGGCGACGGGCGATGATGGCGCGGCCATTTTTGGTTGCCATACGAGCACGGAAACCATGCG
CGCGCTTGCGCTTGAGAACGCTAGGTTGAAAAGTGCGTTTCATAACTCGTTATTCCCACGTGGTTTAGGACGGAAAATGAATTCCAAAAGGCCCGGTCCA

GTGAGGAACTAGCCGGGAGATTCAATTCAAGACCCGAAATTGTAGAGGCTTCCCGCGTTGCGTGCAATTTTTCCTTGCAAAAAATGCCTTCTGCAGAGGC
GCTTGCCTGTCGCATCGACATCGCCAAGCGGGCAATATATCGTCTAGTGATGCACAGGCGATGATACCGCCTTGCTGTGGCTAATAGTGGGGATACTTTT
**TGTTAGTAA**CCCACTATTATTGTTTTTATTCAAATGCTTACTGTTGGTTCAGTGTTGTGGACAAGTGTCCTTAAAGTGGATGATAAGCAGTTGATTAGCT
GTGAATTAAAGCGTCGTCAGGGTTGACTTTCGGGATGCTCCCGTCAGTTCGGGAGGCGACTTAAACGGCGAGGGTTAAAGTTTTCCACAGCTTGTTGAAT
CGCTTCTTGCCTGTGTGGATAACCACGCCGTTGGGCGCTACAATGGTCGGCCTTTTTCAATCGATGGTGAGATGAGTGTCACTCGCGCTTTGGCAACAGT

GCCTGGATTACCTGCAGGACGAGCTGAGTTCCCAACAGTTCAATACATGGATACGTCCATTGCAGGCCGAAGATAGTGAGTCCAATGAGCTTCGCTTGTT
GGCGCCCAACCGGTTTGTGCGTGATTGGGTGAGTGACAAGTATGCCAAGCGAATTAGCGAACTGGTGCGTGAGCTATCTCCTTCGAAGCCTCCCAAGGTC
AGTCTTACGGTCGGCAGCCGGCGGCCGGCCACACCAGCCCCCCAACCTCGTGATTTGGGCAATCCGGTATCTGCCGCACCGGCGCCGCGTTCATCACCGG
CGGTTCATACGCCGCCGCGTGGTGACATTGCCGATGAGCGTGAGATAGACCAGCTACGTGAAGAGAACGAAGAAAATAACGGCCGTCGCAATAATGATCG
CCAGGTTCAGGTTGAAGGCAGTCTCAAGCATACCAGCGGGCTGAACCCCAATTTCACCTTCGACACCTTCGTTGAGGGTAAATCTAACCAGCTTGCCCGC
GCGGCCTCGCGGCAGGTCTCGGAAAACCCCGGTGGCGCCTATAACCCGCTTTTCCTGTATGGCGGCGTTGGCCTGGGTAAAACCCACTTGATGCATGCGG
TCGGGAATGGGCTGGCCGTTCGCCGCGAAAATGCGCGCGTCGTGTACCTTCACTCTGAGCGTTTCGTTGCCGACATGGTGAAGGCACTTCAGCTCAACGC
GATCAACGATTTCAAGCGTTTCTACCGTAGCGTAGACGCGCTGCTCATAGATGACATTCAGTTTTTTGCTGGTAAAGAACGGTCGCAGGAAGAGTTCTTC
CATACGTTCAATGCCTTGTTGGAAGGCGGGCAGCAAATGATCCTGACATCCGACCGTTACCCCAAGGAAATCAGCGGTGTAGAAGAGCGTCTGAAATCCC
GCTTCGGGTGGGGGCTTACTGTCGCCATCGAGCCACCCGAACTGGAAACCCGCGTTGCGATCCTGATGAAGAAAGCCGATCAGGCCAAGGTCGACCTGCC

>Halomonas gomseomensis DSM 18042 [NZ_JARWAI010000003.1:299176..299675], <mnmG oriC2 <COG1126 (glnQ), 558 bp

GGCCCTTTATCGAGCGCACGACCTGCAACTGGATATCGAACGGCAGCGACGTTGAAATGCCATTCGGGTACAGCTCGTTGGTGTCCAAGCCTTCGGGCTC
GACGAATATCTGGTGGCTCGCTTTATCGGCAAAGCGATGCACCTTGTCTTCGATGGATGGGCAATAACGCGGCCCGATGCCTTCGATGACGCCTGAGTAC
ATCGGCGAACGCTCAAGGTTCGCCATGATGATCTCGTGGGTATGCGGGTTGGTCTGCGCGATATGGCAGCTCATCTGCCGGGGATGCATATCCCGCGAGC
CCAGATAAGACATCACCGGCGTGGGGGTATCACCGGGCTGCTCCTCCATTTGCGAGAAGTCCACCGTCTTGGCATCGATGCGCGGTGGCGTACCGGTTTT
AAGCCGGTCAACCCGAAACGGTAATGCACGCAGTCGCTCTGCCAGGGCATTCGAAGGCGCATCGCCGGCCCGGCCACCGCGGCTTTGATCGAGGCCGATA
TGAATGATCCCGCCGAGGAAGGTGCCTGTCGAGAGAACGACCGTCTCGCTATGAAACCGGATGCCGGTTTCCGTGACCACGCCGCGTACGGTGTTGTTGT
CCACCATCAGGTCGCCGGCGGCCTGCTGGAAAATCGTCAGGTTGGGCTGGTGTTCCAGCATGCCACGGATAGCCGCTTTGTAACGAACTCGGTCGGCCTG
GGCACGAGTGGCGCGTACCGCCGGCCCCTTGCGGGCGTTCAACACACGAAACTGAATACCGCCCATGTCGGTGGCGAGCCCCATGGCACCGCCCAGGGCA
TCAATTTCCTTGACGAGATGGCTCTTGCCAATCCCGCCAATCGCCGGATTACACGACATTTGACCGAGGGTTTCGATGTTATGGGTCAGCAGCAGGGTTT
GACAGCCCATGCGAGCAGAGGCCAGTGCGGCTTCGGTTCCCGCATGGCCACCGCCGATGACAATCACGTCAAAGCGGTCGGGGTAATTCAAGAGACACCT

CGTCTTGTGCCCTGAGGCACGGATGAGTTGAGCCGGTGCACAGGCACCTTAATTCACTAGGTCGGCAAGTATAAACCCCCTGTGGGTAACCGTCAGGTTT
TTCCACACCCCGATTTGTTCCTTGGCCTCTATTAATAACAACATAAATATAAAATAAGAGAAGTTCTGTTAATGCTGTAATTACTGGTGTGGACAAAATC
**TGTTGATAA**GTCTATAAAGCACATATTTATCAGAAAATTGCATTGTTGATCTTTACGTGATTAAACGGCGTACTAGCTGCGTAGAAACGGTGCTGGGCCT
GTGGATGAAATGCAAGTTTATCCACCGGTGAAATAAAACACGCCTTGTCCACCGCTGCATACCGGAGTTGTTACCGCGACTGTGAACAAGAGGCAAGCCA
CCTGGTGTATCGCGCCTAACGCCAGCGTGCCGCTGGATGGCAGCGGATTGGCATGGATCGTAAAAAAATACTATCCACAGGCAAAAAAAATGGCCTGTGA

AGATCACAGGCCATGAAACGCGGTCTTTAGAGCGGGATAATTGGCTGAATCAGTGCTTGAGTACCCGCGCTAAAAACGATTGGGTGCGTTCGTGCTGAGG
CGTATCGAAGAGCTGTTCAGGAGGGCCTTGTTCGGTAATTTCGCCTTGGTGAATGAATATCACCCGGTCAGCAACCTCGCGGGCAAAACCCATTTCGTGG
GTAACGATCACCATGGTCATACCTTCTTTTGCCAGCTCGCGCATGGCATCCAGCACCTCACCGATCATTTCCGGATCAAGGGCCGAGGTAGGCTCGTCAA
ACAGCATCAGCCGGGGTTCCATGGCCAACGCGCGGGCCAGTGCCACGCGTTGCTGTTGGCCGCCTGACAGCTGGCTTGGGTACTTGTCCGCCTGATCCGA
GATGCCGACCCGCTCGAGCAGTCGCTTGGCAGTCTCCAGGGCGTCGGACGCGCTCCAGCCGCGTACTTTCATCGGGGCCAGGGTAATGTTGTCGATAACG
CTCAAGTGTGGAAACAGATTGAATTGCTGGAACACCATGCCGACTTCGGTGCGGATGGTCTGCAGCGCCTTGCCACTTTTGCCATGGGGTAGCAATTCGT
TGCTATCGACCTCCAGCTTACCGGTCTGGAATTCTTCCAACCCGTTGATACAGCGGATCAAGGTGGACTTACCCGAGCCGCTGGCGCCAATCACCACGAC
GACTTCGCCGGGGGTGATCTCCATGTCGATATCCTTGAGTACGTGCAGGCTGCCAAAGTGTTTATTGAGCTGCTGCATACGCACAATTCGCTGGGGTGTG
TCGTTCATTGTGCGTCTCCTTATTGGCCGACCAGGCCTTTGCTTTCAAGCTGGCGCAGGACAATCGATAGGGTCCAGGTAATGGCGAGGTACATGAGGGC
CACCATCAGGTAAACCTCCATGGCCGTGAAGGTGCTGGCGATATATACCTGCCCTTGGCGAACCAGTTCACCCACGCCGATCACTGAAAATAGGGACGTA

>Halomonas gomseomensis DSM 18042 [NZ_JARWAI010000003.1:322487..322542], dnaA> ig dnaN>, 56 bp

WebSkew GCmin pos. 22626 (GCmin<>oriC1 nd; GCmin<>oriC2 0.8 kb; oriC2<>oriC1 nd)

>Halomonas arcis CGMCC 1.6494 [NZ_FNII01000021.1:c44909..45408], <rpmH oriC1 dnaA>, 532 bp

CCGTGGTGACGGCAATAAAGTCACGGCTGTCGCTTTCGCTGCTATCCAGTGCCTCTTGAGACGACGAGCCACCTTCGTTGGGCACAGCCAAGCTCTCAGC
GTTATCGTCGCCGGTTCCTGCCCCGCTACTCGACGATGGAACGGCGCTGCTGCTCGTTTGGTTTTGCGATATCTCAGACGACGATTGGTAAGACGTCTGT
CCGTAATCCTGATTCCACTGTACAACCAGTAGATAAGCCAGGATGGCAAGTGGAATCAGTAATATTAGTCGTTTTACGTCCATGAGGGTTCTGCCCGCTG
GGTAGTGAACATGCTATGGCGCAGTGTCAATTACTCGACCTATGACACACGAAAGCCTGCCAGGCGGCAGGCCGAGGTAGAGCGATGAGGTGCCGATCAA
GTATCGCGCATTGAGCCGCTAGCAAGAGGCGCATTAGCTGCTTGTGCATCGCGCCGAAGTCGCTTCCACATGCCGTGGAGTTGACGGAATAACACGTCGT
TGTCCACTTCCTGAACGCCACGCCGAGCAAGCACCACAATATCCACAGCGGGGAGCTGGTCTTGACGTAGTCGAACCGACTCGCGAACGAGGCGTTTGAA
GCGATTTCGATCAACGGCGAGTTTCACATTTTTTTTGCTGACCACTAAGCCAAGGCGAGGATGCCCTAAGGTATTCCAGCGTGCTAGCGCCATCATCCCT
TTGCCATGCACCTTGAGGTCTGCTTGCTCAAACACGAAACGGTAATCCCCAGCGTTAAGTAACCTAAAGCACCGGGGAAACCCTTGATGTGGCACACGCA
ATCCGAGATCAGGCGCTCAGACGCTTACGGCCCTTGGCGCGGCGACGTGCGATAATGGCACGGCCGTTTTTGGTTGCCATACGAGCACGGAAGCCATGCG
CGCGCTTGCGCTTAAGAACGCTAGGTTGAAAAGTGCGTTTCATAACTCGTTATTCCCACGTGGTTTAGGACGGAAAATGAATTCCAGAATGCCCGGTCCA

GTGAGGAACTAGCCGGGAGATTTAATTCAAGACCGGAAATTGTAGAGGCTTCCTACGTCACGTGCAATTTTTCCTTGCAAAAAGACCGGATGTTCAGAG
ATGACGCCCCCGATGTGATGCGCTGCGTTTCGCTATCACGTCATTTATCGTATCCGTACAATGTTGATAAGACGCTGTGACTAATAGTGGGATACATTT
**TGTTAACAA**CTCGATATTAACTATCAAATTCAGTTGCTTATATGTTGTTTTATTATGTGTATAAGTTGCGCAAAAGACGATGATAAGCCGTTGATTAGCT
GTGAATTAAAGTCTCGCTTTGATGAGCTTTTAGTCACCGTTAGCTAGCTAAACAGCCAAGCCTTATCGTGAAAAAGAAAGTTTTCCACAGGTTATTAAAT
CAGCCAATACCTGTGTGGATAACCACGCCGTTGGGCGCTACAATGGTCGGCCGTTTTCAACTGATGGTGAGATGAGTGTCACTCGCGCTTTGGCAACAGT

GCCTGGATTACCTGCAGGACGAGCTGAGTTCGCAGCAGTTCAATACATGGATACGTCCATTACAGGCCGAAGATAGTGAGGCTAATGAGCTGCGCTTGTT
GGCCCCCAACCGCTTTGTGCGTGATTGGGTCAGTGACAAGTATGCCAAGCGGATTAGTGAGTTAATGCGTGAGCTTGCCCCTTCCAAGCCGCCTAAAGTC
AGCCTGACCGTTGGCAGCCGTCGTCCTGCGACACCTGCTCCGCAGCCTCGCGACTTGGGTAACCCGGTGTCCGCCTCCCCGGCACCCCGGGCGTCGGCGG
CGGTTCACACGCCGCCGCGTGGTGATATTGCGGATGAGCGTGAAATTGATCGATTAAGAGATGAAAACGGTTCGGGAGGCGGGCGACGCAACAATGAGCG
TCAAGTACAGGTAGAGGGCAGTCTCAAGCACCATAGCGGACTGAACCCTAATTTTACTTTCGACACCTTTGTTGAAGGTAAATCTAACCAACTCGCCCGT
GCGGCATCTCGGCAGGTTTCCGAGAATCCGGGTGGGGCCTATAACCCGCTATTTTTATACGGCGGTGTGGGGCTTGGTAAAACCCACTTGATGCACGCGG
TGGGTAATGCCCTGGCGGTGCGCCGTGAAAACGCGCAGGTGATATATCTTCACTCAGAACGTTTCGTCGCCGATATGGTCAAGGCGCTACAGCTAAACGC
TATCAATGACTTCAAGCGCTTTTATCGCAGTGTCGATGCGTTACTTATTGATGATATTCAGTTTTTTGCTGGAAAAGAGCGCTCGCAGGAAGAGTTCTTC
CATACGTTCAATGCGTTATTGGAAGGCGGTCAGCAGATGATCTTGACGTCAGACCGCTACCCCAAAGAAATCAGTGGCGTTGAAGAGCGTCTGAAATCAC
GTTTTGGCTGGGGCCTAACGGTGGCGATTGAGCCGCCTGAGCTGGAAACTCGCGTTGCCATTTTGATGAAAAAAGCCGAGCAAGCCAAAGTTGACTTGCC

>Halomonas arcis CGMCC 1.6494 [NZ_FNII01000001.1:21555..22054], <mnmG oriC2 (BLR97_RS19495> ignored) <COG1126 (glnQ), 557 bp

ACCCTTTGATTGAGCGAACCACCTGCAGTTGTATATCAAAAGGCAGGGAGGTGGAGATGCCATTGGGATATAGCTCATGGGTATCCAACCCTTCGGGTTC
GATAAACACCTGATGGCTCGTTTTATCGGCGAAGCGATGCACTTTGTCTTCGATGGACGGGCAGTAGCGTGGCCCGATGCCCTCGATAACGCCCGAATAC
ATCGGCGAACGCTCAAGGTTTGCCATGATGATTTCGTGGGTACGCTCGTTGGTGTGCGCAATGTGACAGCTCATCTGCTCGGGGTGCATGTCCCGCGAGC
CGAGGTAAGACATCACCGGTGTTGGCGTGTCACCGGGTTGCTCTTCGAGCTGTGAGAAATCAACGCTTTTAGCATCAATGCGCGGCGGTGTGCCGGTTTT
TAGCCGATCAACGCGGAACGGCAGTGCGCGCAGCCGCTGGGCCAGGGCGTTTGAAGGTGGATCGCCGGCACGGCCACCGTGGCTTTGATCCAGGCCAATA
TGAATGACGCCGCCTAGGAAAGTCCCGGTCGAGAGCACGACCGATTCGGCATGAAAGCGAATGCCCGTCTCAGTCACCACGCCCCGAACGGTGTCGTTAT
CCACAATCAGATCCCCAGCCGCTTGCTGGAAGATCGTCAAATTCGCCTGGTTTTCCAACATGCTGCGAATAGCGGATTTGTAACGCACACGATCTGCTTG
AGCACGAGTGGCTCGAACAGCGGGCCCTTTGCGTGCGTTCAGAACACGAAACTGGATGCCACCGACGTCGGTGGCTAGCCCCATCGCGCCGCCTAACGCA
TCGATTTCCTTGACGAGATGGCTTTTGCCAATCCCGCCGATCGCCGGGTTACACGACATCTGGCCGAGGGTTTCGATGTTGTGGGTTAGCAACAGGGTCT
GACAGCCCATACGAGCAGAGGCCAATGCGGCTTCAGTTCCCGCATGGCCACCGCCGATGACAATCACGTCAAAGCGGTCGGGGTAATTCAAGAGACACCT

CGTCTTGTGCCTTGGGCACGGATGAGTTGAGCGCGGTGCGCAAGCACCTTAATTCAATAAGCCCGCCAGTATAAACCCCCTGTGGGTAACCGTCAGGTTT
TTCCACATGCCAATTTGTTCATCAGCTGCTATTAATAACTACATATAAATAAAATAAGAGAAGTTCTGTTAATGTTGTAATTACTGGTGTGGACAAAATC
**TGTTGATAA**GCTACTTATTTAACTATATTTCATGATGTTACGATGTTGACCTTTACATGATTAAACGGCGGACTAGCTGCGCACAAGCGGTGTTAGTCCT
GTGGATGAAACCTTCACTTATCCACCGATACAAGCCCGCACGGCTTTTCCACCGCTGCATACCAGGGTTGTTACCGCAACGGTGAACAATTGGCTATCAA
TGGTTGGCATGCTCGTTAATCCCTTCACCGCAGGGGCTAGAGGCCTGTTTAACAAGACAACAAAAAAAGTTTATCCACAGGCAAAAAAATGGCCTGTGGT

AATCACAGGCCATCAAAAATCATCAATTAATCGGGTTGTGGGCTAAATTTAGTGCTTGAGTACGCGTGCCAAAAACGATTGAGTACGTTCGTGTTGTGGG
GTATCAAACAGTTTTTCAGGAGGCCCTTCTTCAGTAATTTCTCCTTGATGGATAAATACCACGCGGTCGGCAACTTCCCGGGCAAACCCCATTTCATGGG
TAACGATCACCATGGTCATGCCTTCTTTAGCGAGCTCTCGCATGGCGTCTAGCACTTCCCCGATCATTTCGGGGTCAAGGGCCGAGGTAGGTTCATCAAA
CAGCATCAGCCGCGGCTCCATTGCCAGTGCGCGTGCTAAGGCAACCCGCTGCTGTTGGCCGCCTGAAAGATGACTGGGGTATTTATCCGCTTGGTCAGCA
ATACCCACACGCTCTAGCAGGCGCTTAGCGGCTTCTACGGCATCAGTTCGACTCCAACCACGTACTTTCATGGGGGCGAGGGTGATATTGTCGATAACGC
TCAAATGGGGAAACAGATTGAATTGCTGAAACACCATGCCGACTTCGGTTCGGATGGTTTGCAGCGCCTCACTGCTCTTCCCATAGGGCCGCAGTTCATT
GCTATCGACCTCCAGGAATCCGCTTTGGAATTCCTCCAACCCATTGATGCAGCGAATTAAGGTGGATTTACCCGAGCCGCTTGCGCCGATAACCACGACG
ACCTCACCAGGGGTAATCTGCATATCAATATGTTTTAGTACGTGCAGACTGCCAAAATGTTTATTGAGCTGCTGCATACGCACAATTGGCTGCGTTGTGT
TGTTCATTGCGCGTGTCCTTATTGTCCTGCCAGGCCTCTGAGCTCAAGTTGGCGCAGTACGATGGAGAGGGTCCAGGTAATTGCTAGATACATCAGGGCG
ACCATGAAATAGACCTCAAGGGCCGTGAAGGTGGTCGCGATATAGATCTGCCCCTGACGCACCAATTCACCGACGCCAATCACTGAGAATAGCGATGTGT

>Halomonas arcis CGMCC 1.6494 [NZ_FNII01000021.1:c43401..43457], dnaA> ig dnaN>, 57 bp

WebSkew GCmin pos. nd (GCmin<>oriC1 nd; GCmin<>oriC2 nd; oriC2<>oriC1 nd)

>Halomonas salicampi BH103 [NZ_JACCDF010000018.1:20780..21279], <rpmH oriC1 dnaA>, 524 bp

ATAGAGGATATCGCCACCGTGAGGATCGATACGCACATCGAGTACGTCGGTAGTGACGGCAATAAAGTCGCGGCTGTCTGTTTCGCTTTCTTCAGCCGGC
ATGCTGTCGTCGATATCGCCCTGCTGAGTGCCACTTGAGGGCACGCTAAGGCCACCATCATCTTGCGTTTCTTCTGGGGTCGAGCCGTTACGCTGTACGA
TTTCGGTGGAAGAAGACGGCGTAGCAACCGGCTGACTATAGTCCTGACTCCATTGAACAACTAGGAGATAGGCCAAAATAGCCAATGGAATCAGTAGGAT
AAGTCGTTTTACGTCCATGAGGGTTCTGCCCGCTGGGTGGTGAACATGCTAATGGCGCAGTGTGATTTTACTCGACATATGACACACAAAAGCCTGCCAA
GCGGCAGGCCGAGGTCGAGCGGTAAAACGTAGTCGTCAAGCGTCACGTTGCGCTTCGCGCTGCAGACGTTTCCACATGCCGTGTAGCTGGCGATGTAACG
TTTCGTTGTCGAGATCATGAACGCCTCGTCGTGCCAGAACCACGATATCAACCGCAGGTAGCTGGTGTTGGCGTAAGCGAATAGATTCCCGTACGAGCCG
TTTGAAACGGTTGCGTTGCACGGCAAGCTTCACATTTTTTTTGCTAACCACAAGCCCAAGCCGGGGATGCCCGAGGGTATTCCAGCGAGCTAGCGCCATC
ATACCTTTACCATGAACTTTTAGATCAGCGTGTTCGAAAACGTGACGGAAATCCCCGGGGCTGAGTAGTCGTAAGCACCGAGGATAGCCTTGACTGAGCA
CACGCAATCCGAAATCAGGCGCTCAGACGCTTGCGGCCTTTGGCACGACGGCGTGAGATTACAGCACGGCCGTTTTTGGTTGCCATACGAGCACGGAAGC
CATGCGCGCGCTTGCGCTTTAGAACGCTGGGTTGAAAAGTGCGTTTCATAACTCGTTATTCCCACGTGGTTTTTTGACGGAATGTCGATTTTAAGAGGCC

CAAACCAGCAAGGCACTGGCATAGGGAGATCCAATTCAAGACCGGAAATTGTAGAGCCTTACCGCGATAGGTGCAATTTTTTTGTCGAGCTACCGTGATG
GCCGTACCTGCCGCTATGCTCCATTCTGCTCAAGGTGACACAACATTAACAATAATAGTATTACTTAAATGATGTTGTTATTAATAGTGGGGATACAATT
**TGTGGATAA**GTGTTTTAAATCGTTTATTAACAGGTAGATAAACATTTTAAAAATGTGTGGATAAGCGGTTATTTAGCAGGGGGTTCGCTGTGGGGAAACG
GTGCAAAAGCGAGGATAAATCGCCCATGCTAACAAGCGTGCCATGCTTATCCTCGGGCTGTTCGTAACTTGGTAAGAGGGTTATCCACAGAAAATATTTA
GCCGTACGCTGTGTGGATAACTGGCTTTTGGGGCGCTACAATATCCGGCCGTTTTCAACCGAAGGTGAGTTGAGTGTCTCTCGCGCTATGGCAACAGTGT

CTGGACTATCTGCAGGACGAACTGAATTCACAGCAGTTCAATACCTGGATACGTCCCCTTCAAGCAGAGGAGGGGGAGTCCAACGAGCTGCGCTTGTTAG
CGCCTAATCGCTTTGTGCGCGACTGGGTGAGCGACAAGTACGCCAAGCGTATCAGCGAGCTGATTCGGGAGCTTGCACCGACCAAGCCGCCAAAAGTGAG
CTTGACGGTAGGAAGCCGCCGCGTTGCCGCGGCACCGCCCCAGCCGCGTGAGCTTGGCAACCCGGTTTCCGCATCGCCGGCAAAACCGGCGGCACCGGCA
GTGAGTACGCCCCCCCGTGGCGATATCGCCGATGAGCGCGAGATTGATCAGCTGCGCGAAGAACAGGCGTCAAAGCGGCCTAAGGAGCGTCAGGTTCAGG
TGGAAGGGGGGCTGAAACACAATAGCGGCTTGAACCCTAATTTTACTTTCGACACCTTTGTGGAAGGTAAATCAAACCAACTGGCGAGAGCTGCTTCGCG
CCAAGTGTCAGAAAACCCCGGTGGGGCATACAACCCACTTTTCTTATACGGTGGAGTGGGGCTAGGTAAAACTCACTTAATGCACGCGGTAGGCAATGCG
CTGGCGACCCGGCGTGAGAATGCGCAGGTGGTATATTTGCACTCAGAGCGCTTCGTTGCCGATATGGTGAAAGCCCTCCAATTGAATGCGATTAACGACT
TTAAACGCTTTTATCGTAGCGTGGATGCGTTACTGATTGATGATATTCAGTTTTTCGCCGGCAAAGAGCGCTCTCAAGAGGAATTTTTCCACACCTTCAA
CGCGCTGTTAGAAGGCGGGCAGCAGATGATTCTGACCTCTGACCGCTACCCTAAAGAGATTAGTGGCGTCGAAGAGCGCTTGAAGTCTCGCTTTGGTTGG
GGGTTAACCGTTGCCATTGAACCACCGGAGCTTGAGACGCGCGTGGCCATTTTGATGAAAAAGGCCGAACAGACCAAAGTGGATTTGCCTCATGATGCGG

>Halomonas salicampi BH103[NZ_JACCDF010000008.1:c691..1190], <mnmG oriC2 <COG1126 (glnQ), 551 bp

ATATGCGCGTTTTCCAGTCCCTTGATCGAGCGCACTACCTGAAGCTGAATGTCAAACGGCAGCGAGGTGGAGATTCCGTTGGGATACAGCTCGTTGGTAT
CTAAACCTTCCGGCTCAATGAAAATCTGGTGGCTCGATTTATCCGCAAAGCGATGGACCTTGTCCTCAATCGACGGGCAGTAGCGCGGCCCGATGCCTTC
GATCACGCCGGAATACATCGGCGAGCGATCCAGGTTTTTTAGAATGATGTCATGCGTTTGCGTATTGGTATGGGCAATGTGACAGCTCATCTGGCGCGGA
TGCATTTCGCGGTTGCCGAGATACGACATCACCGGCGTGGGCGAATCCCCCGGCTGCTCTTCAAGGTGAGAGAAATCTACGGTTTTGGCATCAAGACGCG
GCGGCGTGCCGGTTTTTAGCCGGTCAACGTGAAACGGCAGCGCGCGTAATCGTTCGGCTAGGGCGTTGGAGGGTGGATCCCCTGCGCGTCCACCGCGATT
TTGATCAAGTCCGATGTGAATCACGCCACCGAGGAAGGTTCCTGTACACAGCACGACGGATTCGCCGTGAAAACGAATCCCTGTTTCAGTTAACACCCCG
CGAACGGTGTCGTTGTCCACGATCAGATCGCCCGCGGCTTGCTGGAAGATGGTCAGATTCGGCTGGTTTTCCAGCATGCCGCGGATAGCCGCTTTGTAAC
GCACCCGGTCAGCTTGAGCACGGGTAGCGCGCACCGCCGGTCCTTTGCGAGCGTTAAGCACACGAAACTGGATACCACCCAGGTCCGTGGCTAGCCCCAT
CGCACCCCCAAGGGCATCGATTTCTTTGACCAAATGGCTTTTGCCGATGCCGCCGATGGCGGGGTTACACGACATTTGGCCCAGAGTTTCGATGTTGTGG
GTGAGCAGCAGAGTTTGACAGCCCATGCGAGCAGCGGCCAGCGCGGCTTCGGTCCCCGCATGGCCACCGCCGATGACAATTACGTCAAAGCGGTCGGGAT

AATTCATGACACCTCGTCTTGCGCGTTTTAGCGCGACTTTATTCAGCATGCATCAGTAAGCCCGGCAGTATAAACCTCCTGTGGGTAACCGTCAGGTTTT
TCCACATCCGGATTTCATCATTTCACTCACTATTAATAACAACATATATATAAATAAAGGAAGTTCTGTTGTTGTTGTAATTAATTGGTGTGGATAAAATC
**TGTGGAAAA**GAGTATTTATTTATCTACTATCAATTGGTTGCAATGTGGGCAGATATGTTAAGAAAGCGCGTACTAGACGCGTACAAACGGTGTTGAGCCT
GTGGATGGAAACGCGATTTACCCACAGGACAAATTACTGACGGCTTTTCCACTGCTGCGTACCTGACTTGTTACCGTGGCTGTGAACAACCCGGGGGTTT
CTCCACAATGACGATTAAGCACCGGCATGCCAACGTTTGAGGCTGATACATGAAGGCGTGAAAAAAATAGTATCCACAGGCAAAAAAAAGCCTGTCACGA

GGACAGGCAATGAAATAGAGCGATTTTTGGCGCTTTTCTTAGCGCTAATGTGGCGAATCTAGTGTTTTAAAACACGCGCCAAAAATGACTGTGTACGTTC
GTGCTGGGGTTTATCAAACAGCTGCTCGGGTGGGCCTTGTTCGGTAATTTCCCCCTGGTGGATGAAGATTATCCGGTCGGCGACTTCGCGAGCAAAGCCC
ATTTCGTGGGTAACAATCACCATCGTCATGCCCTCTTTCGCTAACTCGCGCATGGCGTCGAGCACTTCACCGATCATCTCCGGATCAAGCGCTGACGTAG
GCTCGTCAAAGAGCATTAACCGGGGTTCCATGGCTAAGGCGCGGGCAAGGGCGACACGCTGCTGCTGACCGCCTGAAAGCTGTGCCGGGTACTTATCCGC
TTGATCGGAAATGCCGACACGCTCAAGTAGGCGCTTGGAGGTTTCGACGGCATCAGCCCGGCTCCAACCGCGCACCTTCATCGGTGCAAGCGAAATATTG
TCGATCACGCTGAGATGGGGGAACAGGTTGAACTGCTGAAATACCATGCCCACTTCAGTGCGGATTTTTTGGGCTCTTCGTCGTTGGGTGTGGAATTCAC
CCCCTGATCTGGGCCAGGATATGACCTCCACGACGACAGGAGGCATGGCATGGACGGCACACAGATTTTGACCCTCGGCCTGGGCTTGGA

>Halomonas salicampi BH103 [NZ_JACCDF010000018.1:22720..22773], dnaA> ig dnaN>, 54 bp

WebSkew GCmin pos. 23754 (GCmin<>oriC1 23.8 kb; GCmin<>oriC2 0.2 kb; oriC2<>oriC1 23.6 kb)

>Halomonas vilamensis DSM 21020 [NZ_JARWAN010000010.1:47329..47828], <rpmH oriC1 dnaA>, 534 bp

ATATCGCCACCGTAAGGATCGATACGTACATCAAGCACATCGGTCGTAACGGCAATGAAGTCGCGGCTATCAGTTTCGCTTGCGTCAGCCGCCATGCTGT
TGTCGATGTCGCCCTGTTGAGTGCCGCTAGAGGGCACGCTCAGGCCACCATCATCTTGCGTCTCTGCTGGGGTCGAGCCGTTACGCTGTACCACTTCGGT
AGAAGACGGCGTATCAATCGGCTGACTGTAGTCCTGACTCCATTGAACAACCAATAGATAGGCCAAAATAGCCAATGGAATCAGTAGTATAAGTCGTTTT
ACGTCCATGAGGGTTCTGCCCGCTGGGTAGTGAACATGCTAATGGCGCAGTGTGATTTTACTCGACCTATGACACACAAAAGCCTGCCAAGCGGCAGGCC
GAGGTCGAGCGGTAAAGCGTAGTCGTCAAGCGTCACGTTGCGCTTCGCGCTGCAAACGTTTCCACATGCCGTGTAGTTGGCGATGTAACGTTTCGTTGTC
GAGATCATGTACGCCTCGTCGTGCCAGTACTACGATATCTACTGCAGGTAGCTGGTGTTGGCGTAAGCGAACAGATTCCCGTACGAGCCGTTTGAAACGA
TTACGTTGCACGGCAAGCTTCACATTTTTTTTGCTAACCACAAGCCCAAGGCGAGGATGCCCAAGAGTATTCCAGCGTGCTAGCGCCATCATACCTTTGC
CATGAACTTTAAGATCAGCATGTTCGAAAACGTGACGGAAATCCCCGGCGCTGAGTAGTCGTAAGCTCCGAGGATAGTCTTGACTGAGCACACGCAATCC
GAAATCAGGCGCTCAGACGCTTGCGGCCTTTGGCACGACGGCGTGAGATGACGGCACGGCCGTTTTTGGTTGCCATACGAGCACGGAAGCCATGCGCGCG
CTTGCGCTTAAGAACGCTGGGTTGAAAAGTGCGTTTCATAACTCGTTATTCCCACGTGGTTTTTGGACGGAATGTCGATTTTAAGAGACCTAAACCAGCG

AGGCACTGGCATAGGGAGATCCAATTCAAGACCGGAAATTGTAGAGCCTTACCGCGATAGGTGCAATTTTTTTGTCGAGTTCCCGTCGTGATGCCATGCG
CCTTATGCGCCGTTCTCGTCAAAACGAATCAACAGCCACTACAACAATTATTATAAAATATTATTTAAATTCTGCTGTTATTAATACTGGGGATACAATT
**TGTGGATAA**GTGTTTTAAAGTGTTTATAAACAGCTAGGTAAGCGTTTTAAAAATATGTGGATAAGCGGTTATTTAGCAGGGGTATAGCTGTGGGCAAACG
GTGTAAAAACGAAGATAGTTCGTCGTTATTAAGCAGTGTGCCCTGCTTATCCGCGGGGTATTGTTGATTTGGTCAGAGGGTTATCCACAGAGAATATTTA
ACCGAACGCTGTGTGGATAACTGGCTTTTGGGGCGCTACAATATCCGGCCGTTTTCAACCGAAGGTGAGTTGAGTGTCTCTCGCGCTATGGCAACAGTGT

CTGGACTATCTGCAGGACGAACTGAATTCACAGCAGTTCAATACCTGGATACGTCCCCTTCAAGCAGAGGAGGGGGAGTCCAACGAGCTGCGCTTGTTAG
CGCCTAATCGCTTTGTACGCGACTGGGTGAGTGATAAGTACGCCAAGCGTATTAGCGAGTTGATACGAGAACTCGCGCCGACTAAGCCGCCAAAAGTGAG
CTTAACGGTGGGAAGCCGCCGTGTCGCCGCGGCAGCACCTCAGCCACGTGACCTTGGCAACCCAGTTTCTGCGTCGCCAGCGAGACCGTCGTCCCCGGCA
GTGAGCACGCCTCCCCGTGGTGATATTGCTGACGAGCGCGAGATTGATCAGCTGCGCGAAGAGCAGGCGGCAAAGCGTCCTAAAGAACGCCAGGTTCAGG
TAGAAGGGGGCTTGAAACACAATAGCGGTTTGAATCCCAATTTTACCTTCGACACCTTTGTGGAAGGTAAGTCCAACCAGCTGGCGAGAGCCGCCTCGCG
TCAAGTGTCAGAAAATCCCGGCGGGGCTTACAATCCGCTTTTTTTATACGGTGGTGTAGGGCTAGGTAAAACCCACTTAATGCACGCGGTAGGCAATGCG
CTGGCGACGCGACGTGAGAATGCGCAGGTGGTGTACTTACACTCTGAACGCTTTGTCGCGGATATGGTGAAAGCCCTGCAGCTGAATGCGATTAACGATT
TCAAACGCTTCTATCGTAGTGTGGACGCGTTACTGATTGACGATATTCAGTTTTTCGCCGGCAAAGAGCGCTCTCAGGAGGAGTTTTTTCATACCTTCAA
CGCGCTGTTAGAGGGCGGACAGCAAATGATTCTGACGTCTGACCGCTACCCGAAAGAGATTAGTGGCGTCGAAGAGCGCTTGAAGTCTCGCTTTGGCTGG
GGGTTAACCGTTGCCATTGAACCACCGGAGCTTGAGACGCGGGTGGCCATTTTGATGAAAAAAGCCGAGCAGACCAAAGTGGATTTGCCTCACGATGCGG

>Halomonas vilamensis DSM 21020 [NZ_JARWAN010000010.1:23710..24209], <mnmG oriC2 <COG1126 (glnQ), 550 bp

ATATGCGCGTTTTCCAGCCCCTTGATTGAGCGTACCACCTGAAGCTGAATATCAAACGGCAGCGACGTAGAAATCCCGTTGGGATACAGCTCGTTTGTAT
CTAAGCCTTCCGGCTCGATAAACACTTGGTGACTCGATTTATCCGCAAAACGGTGAACTTTGTCTTCAATCGACGGGCAGTAGCGCGGTCCAATGCCTTC
GATCACGCCAGAGTACATCGGCGAACGGTCCAGATTTTGTAGAATAATATCGTGGGTTTTCTCATTGGTATGGGCAATGTAGCAGCTCATCTGCTGCGGA
TGCATCGCACGATTACCCAGATACGACATCACCGGCGTGGGTGAATCCCCCGGCTGTTCTTCGAGTTGGGAGAAATCCACGGTTTTGGCGTCAATACGCG
GCGGCGTGCCGGTTTTGAGTCGATCAACACGAAACGGCAGTGCACGTAACCGTTCTGCCAAGGCGTTGGAAGGCGGATCCCCCGCGCGTCCACCGCGACT
TTGATCAAGCCCGATATGAATCACACCCCCGAGGAAGGTTCCTGTGCAAAGCACCACGGATTCGCCGTGAAAGCGAATGCCGGTCTCGGTTACCACACCG
CGAACGGTGTCATTGTCGACAATCAAATCGCCCGCTGCTTGCTGGAAAATCGTCAAATTCGGTTGGTTTTCCAACATACCGCGGATAGCGGCTTTGTAAC
GTACTCGGTCGGCTTGCGCCCGAGTGGCGCGTACTGCCGGCCCTTTGCGAGCGTTAAGCACACGAAATTGAATGCCGCCTAGGTCCGTGGCTAGCCCCAT
TGCACCGCCAAGTGCATCAATTTCTTTGACCAAATGGCTTTTGCCAATACCGCCAATCGCGGGATTACACGACATCTGGCCCAAGGTTTCGATGTTATGG
GTGAGCAGCAGGGTTTGACAGCCCATACGAGCAGAGGCCAGCGCGGCTTCGGTACCCGCATGGCCACCGCCGATGACAATCACATCAAAGCGGTCGGGAT

AGTTCATGACACCTCGTCTTGCGCGTTTCACCGCGACTTTATTCAGCATGCATCAGTAAGCCCGGCAGTATAAACCTCCTGTGGGTAACCGTCAGGTTTT
TCCACATGGGAATCTTGTCACTTCAGCTACTATTAATAACAACATATATATAAATAAAGGAAGTTCTGTTGTTGTTGTAATTATTGGTGTGGATAAAATC
**TGTGGAAAA**GCATAATTGTCCGATAACTATCAATGAGTTGCTATGTGGGCGAATGTGTTAAGAAGACGCGTACGAGACGCGTACAAACGGTGCTAAGCCT
GTGGATGGAAATGCGATTTACCCACAAGACAAATCATCGACGGGTTTTCCACTGCTGCATACCTGACTTGTTACCGTGGCTGTGAACAACCGTATATTTT
CTCCACAGGGTCGATTAACCCCCGGCATATGCAGACTTAAGAGCAATAAAGAAAATCGTGGAAAAAATATTATCCACAGGCAAAAAAAGCCTGTCACGAG

GACAGGCGATAAAATAATGGGGTTTTTGGCGCTTTTCTTAGCGCTAATGTTGCTCAGTCAGTGCTTTAAAACACGTGCTAGGAAAGACTGAGTGCGTTCG
TGTTGGGGTTTATCAAACAGCTGCTCGGGAGGACCTTGCTCGGTAATTTCTCCCTGGTGAATGAAGATCACTCGGTCGGCGACTTCGCGGGCAAAGCCCA
TTTCGTGCGTCACAATCACCATGGTCATGCCCTCTTTCGCCAATTCACGCATGGCATCGAGTACTTCGCCGATCATCTCCGGGTCCAGCGCTGACGTGGG
CTCGTCAAACAACATTAGCCGTGGCTCCATGGCCAAGGCGCGTGCCAGTGCGACACGTTGCTGTTGGCCGCCTGAGAGCTGGCTGGGGTACTTTTGCGCC
TGATCGGAAATCCCCACGCGCTCGAGCAAGCGGTTCGCGCTATCGATAGCATCAGCGCGGCTCCAACCGCGCACCTTCCTTGGCGCAAGCGAAATGTTGT
CGATCACGCTAAGGTGAGGAAAAAGGTTGAACTGCTGAAATACCATGCCGACTTCAGTACGGATGGCCTGCAAGGCTTTAGTACTTTTGCCATGAGGTAA
CAGCGGTTTGCTATCCACCGTAAGGCTGCCTTGCTGGAACTCTTCTAGCCCGTTAATGCAGCGAATGAGCGTGGATTTCCCCGACCCGCTGGCACCGATG
ACCACCACGACCTCGCCGGGTACGATTTCAAGGTCGATATCTTTAAGTACGTGCAGACTGCCAAAATGCTTATTTAACGCCTGCATCTGCACAATTGGTT
GCGTTGTATCGTTCATTGCGCGTGTCCTTATTGTCCAGCCAGGCCTCTGCGTTCAAGTTGACGCAGGATGATAGAGAGGGTCCAAGTGATCGCAAGGTAC
ATCAGCGCGACCATAAAGTAGACCTCAAGCGCGGTGAAGGTGGTCGCAATATAGATTTGCCCTTGACGCACCAGCTCACCGACGCCGATCACGGAAAATA

>Halomonas vilamensis DSM 21020 [NZ_JARWAN010000010.1:49269..49346], dnaA> ig dnaN>, 78 bp

WebSkew GCmin pos. 686730 (GCmin<>oriC1 31.9 kb; GCmin<>oriC2 5.2 kb; oriC2<>oriC1 26.7 kb)

>Halomonas massiliensis Marseille-P2426 [NZ_LT699745.1:718399..718898], <rpmH oriC1 dnaA>, 522 bp

GCGGCATAAAGAATATCGCCACCGAAAGGATCGATACGCACGTCAAGCACGTCAGTGGTGACGGCGATAAAGTCGCGGCTGTCGGTCTCGCTTTCGCCAG
CCGCCATGCCGTCGTCGATATTGCCTTGCGAAGAGGCGCTCGAAGGCACGCTAAGCCCGCCGTTATCTTGCGTCTCGTCAGGGGTCGAGCCGTTTTGCTG
AGATACCTCCGCGGAGGAGGGCATATCAACCGGTTGACCGTAGTCCTGATTCCATTGAACAACCAGGAGATAGGCCAAAACGGCCAATGGAATCAGTAGA
ATAAGTCGTTTTACGTCCATGAGGGTTCTGCCCGCTGGGTGGTGAACATGCTGATGGCGCAGGGTGATTTTACTCGACCTATGACACACAAAAGCCTGCC
AAGCGGCAGGCCGAGGTCGAGCGGTAAGGCGTGTGCGTCAAGCGTCACGTTGCGCTTCGCGCTGCAGTCGTTTCCACATGCCGTGTAGCTGGCGATGTAA
CGTCTCGTTGTCGAGTTCATGCACGCCCCGCCGTGCCAATACCACGATATCAACCGCGGGTAACTGGTGCTGGCGTAAGCGAATCGATTCTCGTACAAGC
CGTTTGAAACGGTTGCGTTGCACGGCAAGCTTCACATTTTTTTTGCTAACGACAAGTCCAAGGCGGGGATGCCCGTGAGTATTCCAGCGCGCTAGCGCCA
TCATACCTTTGCCATGAACTTTAAGATCAGCGTGTTCGAAAACATGACGGAAATCCCCGGCGCTAAGCAGACGCAGGCACCGAGGATAGCCTTGACGGAG
CACACGCAATCCGAAATCAGGCGCTCAGACGCTTGCGGCCTTTGGCACGGCGGCGTGAGATCACGGCACGGCCGTTTTTGGTCGCCATACGAGCACGGAA
GCCATGCGCGCGCTTGCGCTTTAGAACGCTGGGTTGAAAAGTGCGTTTCATAACTCGTTATTCCCACGTGGTTTTTGGACGGAATGTCGATTTAAAAGAC

CTAAACCAGCAAGGCACTGGCCTAGGGGGATCCAATTCAAGACCGGAAATTGTAGAGCCTTACTAGGTTAGGTGCAATTTTTTTGTGGCGTGACCGCGTG
GGTATCGACCGTTATCAGCGCCACGCTACGCGACAAGACTCTTTAATAACAATAATAGTATTACTTAAATGATGTTGTTATTAATAGTGGGGATAAAAAT
**TGTGGATAA**GTGATTTAAGTATTTTATTAACAACACCCTATGCGTTTAAAAAAGGTGGGGATAAGCGGTGGTTTAGACGGGGTATAGCTGTGAGTAAACG
GTGCAGTCGCGTGGATAAATAGGCGGTGGTAAGAGTTGTACCCCGTTTATCCGCCGCGCGTTGAGCATTCGATAAGAGGGTTATCCACAGCTGATATTTA
GCCGCGCCCTGTGTGGATAACTGGCTTTTGGGGCGCTACAATACCCGGCCGTTTTCAACCGAAGGTGAGTTGAGTGTCTCTCGCGCTATGGCAACAGTGT

CTGGACTATCTACAGGACGAGCTGAATTCACAGCAGTTCAATACCTGGATACGTCCCCTCCAAGCAGAGGAGGGGGAGTCCAACGAGCTGCGCTTGTTGG
CGCCCAATCGCTTTGTACGCGACTGGGTGAGCGATAAGTACGCCAAGCGTATTAGCGAACTGATGCGCGAGCTTGCACCGACCAAACCCCCTAAAGTGAG
CTTGACGGTGGGGAGTCGCCGCGCGGCCGCGCCGTCTCCCCAGCCAAGGGAGTTGGGCAACCCAGTGTCGGCATCGCCCGCCAAACCTTCGGCCCCGGCC
GTTAGCACGCCGCCACGCGGCGACATTGCCGATGAGCGTGAGATCGATCAGCTGCGCGAGGAGACGCCAGCAAAACGCCCCAATGAGCGTCAGGTTCAGG
TCGAAGGCAGCCTTAAGCACAACAGCGGCTTGAACCCCAATTTCACCTTCGAAACCTTCGTGGAGGGTAAATCAAACCAGCTGGCCAGGGCCGCCTCACG
CCAGGTATCGGAGAACCCCGGCGGGGCCTATAATCCGCTTTTCCTATACGGCGGCGTGGGGCTGGGTAAAACCCACCTGATGCACGCGGTGGGTAATGCG
CTGGCGACCCGGCGGGAGAACGCGCGCGTGGTGTATCTGCACTCTGAGCGCTTTGTCGCGGATATGGTGAAAGCCCTTCAGCTCAATGCGATTAACGATT
TTAAGCGCTTTTACCGCAGTGTTGATGCGCTACTGATTGATGACATTCAGTTTTTCGCCGGCAAGGAGCGCTCTCAGGAAGAGTTTTTCCATACTTTCAA
CGCGCTGCTTGAGGGTGGTCAGCAGATGATACTGACCTCGGACCGCTACCCAAAAGAGATTAGCGGCGTTGAGGAGCGGTTAAAGTCTCGCTTCGGGTGG
GGGCTCACTGTCGCCATTGAGCCGCCCGAGCTTGAGACCCGCGTGGCCATTTTGATGAAAAAGGCCGATCAGTCCAAAGTCGACCTGCCCCACGATGCGG

>Halomonas massiliensis Marseille-P2426 [NZ_LT699745.1:691666..692165], <mnmG oriC2 <COG1126 (glnQ), 551 bp

TATGCGCGTTTTCAAGCCCCTTGATTGAGCGGACGACCTGCAGCTGAACGTCAAACGGTAGCGAAGTGGAGATACCGTTGGGGTACAGCTCGTGGGTATC
CAAGCCTTCGGGTTCGATGAAAATCTGATGGCTCGATTTATCCGCGAAGCGATGCACCTTGTCCTCGATCGACGGGCAGTAGCGCGGGCCAATGCCTTCA
ATCACGCCGGAATACATGGGCGATCGATCGAGGTTTTCCATGATGATCTCATGGGTTTGCGCGTTGGTATGGGCAATATGGCAGCTTACCTGGTGCGGAT
GCATATCGCGGTTACCCATGTACGACATCACCGGGGTCGGCGTATCCCCCGGCTGCTCTTCGAGCTGGGTGAAATCCAGCGTTTTGGCATCCAGACGTGG
CGGCGTGCCGGTTTTGAGCCGGTCAACGCGAAACGGCAGCGCGCGCAGGCGTTCGGCCAAGGCGTTGGAGGGGGGATCCCCCGCGCGGCCACCCCGGCTC
TGATCGAGGCCGATATGAATCACGCCGCCGAGAAAGGTGCCCGTGCAGAGCACGACGGATTCGGCGAAAAAGCGAATCCCGGTCTCGGTCACCGCACCGC
GAACGGTGTCGTTATCCACCACCAGATCCCCCACCGCTTGCTGGAAGATCGTCAGGTTGGGTTGGTTTTCCAGCATGCCGCGAATGGCCGCTTTGTACCG
CACTCGGTCCGCTTGAGCGCGGGTGGCGCGTACCGCCGGCCCTTTGCGCGCATTGAGGACACGAAACTGAATGCCCCCCAGGTCCGTGGCTAGCCCCATC
GCCCCGCCGAGCGCATCAATCTCTTTGACCAAGTGGCTTTTGCCGATCCCCCCAATCGCGGGATTACACGACATTTGGCCAAGCGTCTCGATGTTGTGGG
TGAGCAGCAGGGTTTGACAGCCCATACGGGCGGAGGCCAGCGCGGCTTCGGTCCCCGCATGGCCACCGCCGATGACAATCACGTCAAAGCGGTCGGGATA

GTTCATGACACCTCGTCTTGCGCGTTATCAACGCGACTTTGTTCAGCATGCAGCGTTAAGCCCGGCAGTATAAACCTCCTGTGGGTAACCGTCAGGTTTT
TCCACACGCGAATCGCACCACTTCACCGACTATTAATAACAACATAAATATAAATAAAGGAAGTTCTGTTGTTGTTGTTACTACTAGTGTGGATAAAATC
**TGTGGAAAA**GCATCTTTAGTTTTAAAAATCATGCATTTGCTATGTGGGCGGCTATGTTGAGAAAGCGCGGATGAGGCGCGGACAAGCGGTGTTGGGCCT
GTGGATGGAAAAGCGAGTTACCCACAGCGTGAAATAGCGACGGCTTTTCCACCGCTGAGTACCCGACTTGTTACCGCAGCTGTGAACAACCCGGCGTTTT
CTCCACATAGGGTCTTGAACGCGGGCATGACAGCGTTTAAGGCGACTCATCAAGGGCGTAAAAAAATAGTATCCACAGGCAAAAAAAAGCCTGTCACGAG

GACAGGCTGTGAAATAGCGGGGTTTTTGGCGCTTATCTTAGCGCTAACATTGCCAAGCTAGTGCTTTAGAACACGCGCGAGGAAAGACTGTGTGCGTTCG
TGTTGGGGTTGATCGAACAGCTGCTCGGGCGGGCCTTCTTCGGTGATTTCCCCCTGGTGGATATAGATCACCCGGTCGGCGACTTCGCGGGCAAAGCCCA
TTTCGTGGGTGACAATCACCATGGTCATGCCCTCTTTGGCTAGCTCGCGCATGGCATCGAGCACTTCGCCGATCATCTCCGGGTCGAGCGCCGAGGTGGG
CTCGTCAAAGAGCATCAACCGCGGCTCCATGGCCAACGCGCGGGCGAGCGCCACACGCTGCTGCTGGCCGCCGGAGAGCTGCGAGGGGTACTTATCCGCC
TGGTCGGCAATGCCGACGCGCTCAAGCAGGCGTTTGGCGGTTTCCACCGCATCCTCACGGCTCCAGCCACGCACTTTCATCGGCGCCAGCGTGATATTGT
CGATCACGCTGAGGTGGGGAAAGAGATTGAACTGCTGAAATACCATACCCACTTCGGTGCGGATTTTTGCAGCGCTTTGCTGCTTTTACCGTGGGGCTG
AAGGGGGTTGCCATCGACCATCAGGCTGCCCTGCTGAAACTCTTCTAGCCCGTTGATACAGCGAATCAGCGTGGATTTGCCCGAACCGCTGGCACCAATC
ACCACCACCACTTCGCCGGGTACGATTTCAAGGTCGATATTTTTGAGAACATGCAGGCTGCCAAAGTGTTTATTCAGCGCCTGCATGGCCACGATAGGCT
GTGTCGTCTCGTTCATGTCCGCGTCCTTTATTGTCCGACCAGGCCTCTACGCTCTAACTGGCGTAGGAGAATGGAGAGGGTCCAGGTGATCGCCAAATAA
AGCAGGGCCACCATCAAATAGACCTCCATCGCGGTAAACGTCGTGGCGATATAAATCTGCCCTTGGCGGACCAGCTCGCCTACCCCGATCACCGAAAACA

>Halomonas massiliensis Marseille-P2426 [NZ_LT699745.1:720339..720381], dnaA> ig dnaN>, 43 bp

WebSkew GCmin pos. 20125 (GCmin<>oriC1 18.3 kb; GCmin<>oriC2 1.4 kb; oriC2<>oriC1 16,9 kb)

>Halomonas azerica TBZ9 [NZ_JABFHI010000006.1:c1590..2089], <rpmH oriC1 dnaA>, 577 bp

AGTCTCGGCTATCGGTTGTGCTCTCATCTGCGGACATCCCTTCATCGATGTCCGCATTCTGAGAAGTACTCGAAGGTACAGCCAAACCGCCGTCGCCGTC
CTGTGTGTCAGCAGAAGTCGTATCATTGGCACTTCGCTGCACAACGTCACTGGAGGACGGACTGTCAATCGGTTGACCGTAGTCCTGATTCCATTGCACA
ACCAGTAAATAAGCCAGTACGGCTAATGGAATCACGAGGATAAGTCGTTTTACGTCCATGAGGGGTCTGCCCGCTGGGTGGTGAACCTGCTAATGGTGCA
GTGATTTTTTACTCGACCTATGACACACAAAAGCCTGCCAAACGGCAGGCCGAGGTCGAGCGGCGCTATATAATCGTCAAGCGTCATGCTGAGTGCTTGC
TGGTGATGCCAATTTTGTGCACTCATGCTGTCTCGCTTCACGCTGCAGCCGTTTCCACATGCCATGTAGCTGACGATGTAACGTATCGTTGTCAAGATCA
TGTACACCGCGCCTAGCCAGAACAACAATGTCAAGCATTGGTAACTGATGTTGGCGTAATCGTACGGATTCTCTAACGAGACGTTTGAAACGATTGCGCT
GTACGGCAAGTTTCACATTTTTTTTACTGACCACTAGCCCGATTCGTGGATGACCGAGCATGTTCCAACGTGCAAGTGCCATCATACCTTTACCGTGGAC
TTTCAGGTCAGCTTGTTCGAAAACGTGACTGAAATCCCCAGCATTAAGCAAACGTAAACACCGGGGATAGCCTTGACGGAGCACACGCAATCCGGATTCA
GGCGCTCAGACGCTTACGGCCTTTGGCACGACGGCGCGTGATAATGGCGCGGCCACTTTTGGTTGCCATGCGTGCGCGGAAGCCATGCGTACGCTTGCGC
TTAAGAACGCTAGGTTGAAAGGTGCGTTTCATAACTCGTTATTCCCACGTGGTTTTAGGACGGAACGTCGATTTCAGGAACCCGATTTAGTCAATAACCA

TCTCGGGTACTGCTAATTCAAGACCGGAAATTGTAGAGCTTTAGCTCGCCAGGTGCAATTTTTTTGTCTGAAGAACCCAGTGATGTCGCATTGGCAGTC
CAGTGATGGGTCGTTCAATGCCACTCAGGGTTATTGCCAGGCCTTATCTACCATTATCTATATCTTTTTAAATATTGTTATTAATAGTGGGGATAGAAAT
**TGTGGATAA**CACCATATAAGTTATTATATTCAGATATTTGAAATGATAAAAAAACTGTGGATAAGCAGTGCTTAAGCGGTAGTTTAGCTGTGACGAAAGC
GTGGATAAATGAAGGTTTAGCATTATTTCAGGTTTTTTCCCCAGCTCATCCAAAGGGCTCAAGCCATAGCAGTAAGAAGGTTATCCACAGAAGTGTCGGA
GATGATACTTGTGTGGATAACTTGGTTCTGGGGCGCTACAATATCCGGCCGTTTTCAACCGATGGTGAGTTGAGTGTCTCTCGCGCTTTGGCAGCAATGC

CTCGATTATCTGCAGGACGAGCTGAATTCGCAGCAGTTCAATACTTGGATACGTCCTCTTCAGGCCGAAGAAGGCGAGTCCAATGAGCTGTGTCTTTTAG
CGCCTAACCGTTTTGTACGTGATTGGGTAAATGATAAATATGCCAAACGAATCAGTGAGTTGGTCAGAGAGCTCTCACCGGCAAAGCCTCCCAAGGTAAG
CCTGACAGTCGGTAGCCGCCGCGTGGCTGCTCCTGCACCGCAGCCACGTCAGCTGGGAACCCCGTTTCAGCCTCAGCCAACAGGTCGCCGTCACCGGCTG
TCAGTACACCTTCACGCACTCATATGAACGATGATCGTGAAATTGATCAACTGCGTGAGGAATCTGCGTCTAAACGCTCCGGTGAGCGTCAGGTGCAGGT
CGAAGGCAGTCTGAAACACACCAGCGGACTTAATCCTAACTTCACTTTTGAAACCTTCGTTGAAGGTAAATCCAACCAATTGGCGCGGGCTGCATCACGT
CAGGTATCAGAAAACCCCGGTGGGGCTTATAACCCGTTGTTTTTATATGGTGGTGTGGGGTTGGGTAAGACCCACTTGATGCATGCGGTGGGCAATGCGC
TGGCAGAGCGACGGGAAAATGCGCGCGTGGTGTATTTGCACTCAGAGCGTTTTGTAGCGGATATGGTAAAAGCCCTGCAATTGAATGCCATCAATGATTT
CAAGCGTTTTTACCGTAGTGTTGATGCCCTGTTGATTGATGATATTCAATTCTTTGCTGGTAAGGAGCGTTCTCAGGAAGAGTTTTTCCATACGTTTAAT
GCCCTGCTAGAAGGTGGGCAGCAGATGATCCTTACCTCTGACCGCTATCCGAAAGAGATCAGTGGCGTTGAAGAGCGTTTGAAATCACGTTTTGGTTGGG
GGCTGACGGTGGCGATTGAACCACCTGAACTTGAAACCCGCGTGGCTATTCTGATGAAGAAAGCCGATCAATCCAAAGTGGATCTTCCCCATGATGCGGC

>Halomonas azerica TBZ9) [NZ_JABFHI010000006.1:c18453..18952], <mnmG oriC2 <COG1126 (glnQ, 584 bp

TTCTCCAACCCCTTGATGGAGCGTACTACCCGCAGCTGGATATCGAAGGGCAGTGAGGTGGAAATGCCGTTGGGATACAGCTCATGGGTGTCCAGCCCCT
CTGGCTCGATAAAGATCTGGTGGCTGGATTTGTCGGCAAAACGATGCACCTTGTCTTCAATCGAAGGGCAGTAACGCGGCCCGATGCCTTCAATCGTGCC
GGAGTACATCGGCGAGCGATCAAGATTAGCCAGAATCAGCTCATGAGTCTGAGGGTTGGTGTGGGCAATATGACAACTCACCTGACGTGGATGCATGGCC
TGATTTCCCAGATAGGACATCACCGGCGTGGGCGTGTCACCAGGCTGTTCTTCAAGCTGGGTAAAATCGACGGTTTTGGCATCCAGGCGTGGCGGTGTAC
CGGTTTTCAGGCGGTCGACCCGTAACGGTAAGGCTCTCAGGCGTTCTGCCAGCGCGTTGGACGGTGGGTCACCTGCGCGGCCACCACGGCTTTGATCCAT
CCCAATATGCATGACGCCACCGAGAAAAGTGCCTGTGCATAACACCACGGATTCGGCATGAAAACGTATCCCGGTTTCGGTGACCACGCCTCTGACCGTG
TCATTTTCAACCATCAGATCGCCTGCAGCCTGCTGAAAAATCGTCAGGTTGGGCTGGTTTTCCAGCAGGCCACGGATAGCTGCCTTGTAGCGAACCCGAT
CTGCCTGGGCACGGGTGGCTCTCACCGCTGGCCCCTTGCGGGCATTGAGAACCCGGAACTGGATGCCACCCAGATCTGTTGCCAGCCCCATTGCTCCGCC
AAGTGCGTCAATTTCCTTGACCAGATGGCTTTTGCCAATCCCGCCAATGGCGGGGTTGCAGGACATTTGTCCCAGGGTTTCAACGTTGTGGGTTAGTAAC
AGGGTTTGACAGCCCATACGGGCAGAGGCCAGTGCGGCTTCGGTTCCCGCATGGCCCCCACCGATGACAATAACGTCAAAGCGGTCGGGATAGTTCATGA

CATCTCATCCTGAATTTTGGGCGGCATTGAATTGAATAACCTATTGCCGCTGTGATCTGGCCGGACAGTATAAACCTCCTGTGGGTAACCGTCAGGTTTT
TCCACATGGGAAATGACAGAGCTGACTGCTTTTAGTAATAATATAAAGATTAAATAAAAGAACTTCTGTAAATGCTGTTACTACTGGTGTGGATAAAAAC
**TGTGGTTAA**GTGATATTTATCTATTACTTTCAGATGTTTATAATGTTTATAAAGATGTTTATGAGTTGCGTAGGGAGCGGGTATAAACCGTCATGAACCT
GTGGATGAAATAGCGACTTACCCACAGGGCAAGTTGACCCCTGCTTATCCACCGACCCGTACATGGGTTGTGAGCCTAGCTGTGAACAAGTGGCTATTCT
AGCGGCGGTACTCTGCAGGGACTGGCATAGAGGCGTATAAGCAGGTTGCCCAGCAGCTATAAAAATTAATTATCCACAGGCAAAAAAGCCTGTCACAGGG

ACAGGCAATAAATTAGCGGGTTTTGACGTGCCTTTTTGGTGCTTAAAACACCTCTCTGACAGCTTGTTCATCTATAGGAACGTCAATGTTTCAGTACCCT
GGCCAGGAAGGATTGGGTACGTTCGTGCTGAGGTTTATCAAACAATTGTTGCGGTGGCCCCTGCTCGGTAATTTCACCCTGATGGATGTAGATAACCCGG
TCAGCGACTTCACGGGCAAAGCCCATTTCATGGGTGACAATCACCATGGTCATGCCTTCCTTGGCCAGTTCACGCATGGCATCCAGTACTTCACCGATCA
TTTCCGGGTCAAGAGCTGAGGTTGGCTCATCAAACAGCATCAGACGCGGTTCCATGGCCAGTGCACGGGCCAGGGCCACACGCTGCTGCTGGCCGCCTGA
GAGCTGGGTAGGCGATTTCTTGGCCTGGTCGGCAATACCTACCCGTTCAAGCAGGCGCATGGCCGTTTCCTCGGCGTCTTCACGGTTCCAGCCGCGTACT
TTCATGGGTGCCAAGGTGATGTTGTCGAGAACGCTAAGGTGCGGAAAGAGGTTGAATTGCTGGAATACCATGCCGACTTCAGTGCGGATCTTTTGTAGCG
CCTTGGTGCTTTTACCAAAGGGCATGAGGTCATGACCGTCAACGGTGAGGTTACCGGACTGAAACTCTTCCAGGCCATTGATACAGCGGATCAGGGTGGA
CTTACCCGAACCGCTGGCGCCGATCACAACGACGACTTCACCTGGCACGATTTCCAGGTTGATATCTTTGAGAACATGCAGGCTGCCAAAGTGCTTATTC
AACTTCTGCATGGTGACGATAGGCTGCGTCGAATCATTCATGTCTGCGTCCTTATTGTCCTGCCAGGCCTCTGCGCTCCAGTTGGCGCAGCAGGATTGAA
AGCGACCAGGTGATTATCAGGTAAAGCATGGCCACCATCAGATAGACCTCCATGGCGGTGAAGGTGGTGGCGATGTAGATTTGCCCCTGGCGGACCAGTT

>Halomonas azerica TBZ9 NZ_JABFHI010000006.1:c161598..161620], dnaA> ig dnaN>, 23 bp

WebSkew GCmin pos. 83732 (GCmin<>oriC1 32.0 kb; GCmin<>oriC2 1.7 kb; oriC2<>oriC1 33.7 kb)

>Halomonas jeotgali Hwa [NZ_AMQY01000003.1:51516..52015], <rpmH oriC1 dnaA>, 728 bp

GGCCGTAGTCCTGGTTCCATTGAACCACCAGCAGATACGCCAGCACTGCCAATGGAATCACTAATATCAGTCGTTTAACGTCCATGAGGGTTCTGCCCGC
TGGGTGATGATCATGCAATGGCGCAGTGTCGTTTTACTCGACCCATGACACACGAAAGCCTGCCGTGCGGCAGGCCGGGGTCGAGCGGCGAAGTGTTGAC
ATCAAGCATCACGTTTTGTGCCGGTCGGCGCGGACGCGGTGTCGAAACGTCGGGCATCGCGCTGCAGCCGTTTCCACATGCCGTGCAGTTGGCGCTGTAG
CGTGGCGTTATCCAGATCATCAACGCCGCGTCGTGCCAATAACACGATATCGAGTGTGGGCAGCTGTTGCTGGCGCAAGCGGAACGATTCGCGTGCCAGA
CGCTTGAAGCGATTACGCTGTACGGCCAGCTTCACGTTTTTTTTACCGGCAACCAGTCCGATCCGGGTATGACCGCAGGAGTTGAAGCGCGCCAGCGCCA
GCATACCCTTGGCATGAACCTTGAGATCGGCCTGTGCGAAAACGTAACGGAAATCCCCGGCGTTCAGCAGACGTAATGACCGGGGAAAACCTTGACGGGA
CACACGCAAGCTGAAATCAGGCGCTCAGACGCTTGCGGCCCTTTGCACGGCGACGTGCGATGACGGCACGGCCGCTTTTGGTAGCCATACGGGCACGGAA
GCCGTGCATGCGCTTGCGCTTGATAACGCTGGGTTGAAAAGTGCGTTTCATAACTTATTCCCACGTGGTTTGAAGACGGGTTTGAGGACTGGTTTCAAGG
ACTGGCTTCACAGGGAGCGTCAATACATGAAAACCCGTGGCGGCACCTTGCCGAACGGGAGGATCCAATTCAAGACCGTGAATTGTAGAGCTTTCATTGG
TCCGGTGCAATTTTTCCATGCCGGAGCCCGGGAAGCGGTTGTGGATAGCGGCCCTGGCAGGCGTTGAAACCCGCCCATGTCGGCACTGAGCGGCTTTTGT

CCTTGATCGGGATCTTGAATCAAGCGTCTTGAATAAGGGCTTTGATAAAGCCCGGGTTTCAGGGCTCGGGTAGGGCTCAGCAAATCGGAAATGACTTAAC
TACGTCGCAGCGGGTCGTTTGCCGGTGACGATGGCGCATGGCGCAACAATTACTATAATTATAACAGATTAATGCTGTTATTATTGATGGGGATAATTTT
**TGTGGATAA**ACGCATATTTATAATTTTATACAAGAGGTTATTGTTATAATAACCCGGTGTAAAACCATCGGAAAACAAGCGTATCAGCTGTGGGGCAGTG
GTGAACAGAAATCGCCAGGGCAAATAAACGCCCGGTTGTGCATAAATTATAACGGGGAGGTCCCTGTGTTGACCATAGGGTTATCCACAGGTGTTTTCGG
CGCTTTTGGTATGTGGATAACAGGGGCTTTGGGCGCTACAATGGCCGGCCTTTTTCAACTTCAACCGATGGTGAAATGAGTGTCGCTCGCGCTTTGGCAT

CAGTGTCTGGACTATTTGCAAGATGAGCTGAGCTCCCAGCAGTTCAACACCTGGATACGTCCGCTGCAGGCGGAGCAGGGAGAGACCAACGAGATTTTCC
TGCTGGCACCCAACCGCTTTGTACGCGACTGGGTAAGCGACAAGTACGCCGGGCGTATCAGCGAGCTGATGCGTGAGCTGGCCCCCACCAGACCGCCGCG
AGTCAGCTTGAGTGTGGGCAGCCGTCGAGCCGCGGCGCCGGCCCCTCAGCCGCGGGATCTGGGGCATCCGGTATCTGCGGTGCCGTCCGGAAAGAGCGCA
CCGTCGGTTCAGCCGCCGCCCCGTGGGGATATCGCCGATGAGCGTGAGATCAATCGGTTGAGTCAGGAAACCACGCATCGCCAGGGCGGCGGCCGTCAGG
TGCAGGTCGAAGGTAGCCTCAAGCACGGAAGCGGACTCAACCCCAACTTCACCTTTGATACCTTTGTAGAGGGTAAGTCCAACCAGCTGGCCCGGGCGGC
GTCGCGTCAGGTATCGGAAAATCCGGGCGGCGCCTATAACCCGCTGTTTCTGTATGGCGGCGTCGGGCTGGGTAAGACCCACCTGATGCACGCTGTGGGC
AACGCCCTGGCCGGGCAGAGAGAAAACGCGCGAGTGGTTTATCTGCATTCCGAACGCTTTGTCGCCGACATGGTCAAGGCGCTGCAGCTCAACGCCATCA
ACGATTTCAAGCGTTTTTACCGCAGCGTGGATGCCCTGCTGATTGATGATATTCAGTTTTTTGCCGGCAAGGAGCGCTCCCAGGAAGAGTTTTTCCACAC
TTTCAACGCTCTGCTAGAGGGCGGTCAGCAGATGATCCTGACGTCCGATCGCTATCCCAAGGAAATCAACGGCGTTGAAGAGCGTTTGAAGTCGCGCTTT
GGCTGGGGGCTGACCGTGGCTATCGAGCCGCCCGAGCTGGAAACGCGGGTCGCGATTCTGATGAAGAAGGCCGACCAGGCAAAAGTGGATCTGCCCCACG

>Halomonas jeotgali Hwa [NZ_AMQY01000003.1:85188..85687], <mnmG oriC2 COG4221> <COG1126 (glnQ), 707 bp

GAGCGCACCAGCTCCAGCTGAACGTCAAACGGCAGCGAGGTGGAAATACCGTTGGGATACAGTTCGTGGGTATCCAGACCCTCGGGCTCAATGAAAATCT
GATGGCTGCTCTTGTCGGCAAAGCGGTCGACCTTGTCCTCGATCGACGGGCAATAGCGTGGACCAATGCCTTCGATGCTGCCCGAATACATCGGCGAGCG
ATCGATATTATCGCGGATAATCGCGTGGGTGCGCTCGTTGGTATGGGCAATGTGGCAGCTCATCTGCCGCGGATGCATATCGCGGCTGCCCAGATACGAC
ATTACCGGTGCGGGATCATCGCCGGGCTGCTCATCCAGCCGGGAAAAATCCACGGTTTTGGCATCCAGCCGCGGCGGCGTGCCGGTTTTCAGCCGATCGA
CGCGAAACGGCAGCGCACGCAGGCGTTCGGCCAGAGCGTTGGAGGGCGCATCGCCGGCGCGGCCCCCACGGGTGGTATCCGTCCCGATGTGAATCACCCC
GCCGAGAAAAGTCCCGGTACAGAGCACGACGGTTTCGCCGTGGAAATGAATACCGGTTTCGGTCACCACGCCTCTTGTGCTGTCGCCGTCCATGATCAGG
TCGCCGGCGGCCTGCTGGAAAATGGTCAGATTGGGTTGATTTTCCAGCACGCGGCGAATAGCGCCCTTGTAGCGAACCCGGTCGGCCTGGGCGCGAGTGG
CACGCACGGCCGGCCCCTTGCGCGCGTTGAGCACGCGAAACTGGATGCCGCCCTGGTCCGTGGCCAGCCCCATGGCGCCGCCCAGCGCATCGATTTCCTT
GACCAGGTGACTTTTGCCGATCCCGCCGATCGCAGGGTTACACGACATCTGGCCCAGAGTTTCGATGTTATGGGTCAGCAGCAGAGTCTGACAGCCGGTG
CGGGCGGAGGCCAGTGCGGCTTCGGTTCCCGCATGGCCACCGCCGATGACGATAACGTCAAAGCGGTCGGGGTAATTCAAGAGACACCTCGTCTTGTGCG

TTGGGCACCGTTGAGCATGAGTATCGAGGAACGGCGCACAGGCACCATGGTTGTTTATGGCCGGGCAGTATAAACCTCCTGTGGGTAACCGTCAGGTTTT
TCCACACCTCGGTTTTGGCATCAGAGCGCCTGTTAACAGTTAATATAATATATATAAAAGAAGTTCTGTTGTTGTTGTAATTACTGGTGTTGATAAAATC
**TGTGGAAAA**CCTAACTTTTAATAATTTTATCAATTGCTTGAGACGTGGATACCATCGTGATGAAAGCGCTGGAAAGACGCGTATAAGCCTTGAATCAACG
GTGGACAACATGCCGTTTTATCCACCGCGCCACCTGGCTACCGCTTTTCCACCGATTCATACCGCCCTTGTTACCGCGGGTGTGAACAACATGGGGTCGG
CTGTCTGAACAGCGCTGAAAGCGCTCAGTGCGCGGTCGGGAAGGGTGGTCGGCCAGCCGTGTAAAGAATCGTATCCACAGGCAGGAATAACGGCTGAAAG

TCTTGCTGAGGGCGAAAAACGCGATTGTTGGACGCCAAAACAGCGAACCATGACGGTCGGAAAGCGGGAGTATTGAACAGAAATCGAGCAGGACTATCCT
GCTGATAGAGAGCGAGACGTTGAGGGCGGGAAATCGGCGGCGTTAACCTGAGCGCCTTATGCCTTTTAGCTGCAACAGGAGTTCAGCATGAACGAGCGCG
TATTGTTGATTACCGGCGCTTCCAGCGGTATCGGCGCGGCAACGGCCCGAGCCGCTGCGGAGAAGGGCTATCGTCTGGTGCTCGCCGCACGCAGCGGTGA
GAAGCTCGATGCCCTCGCCGATGAGATCGGCCGGCAGCAGGTGGTCACCGTGAGCTGCGATGTCACGTCGATGGACGACCAGCGCGCCATGGTCGAACAG
GCGCTAGCTGCTTTTGGCCGCCTGGATGCGGTATTCGCCAACGCCGGGCGCGGCGGTTCACCGGGCGGTTTCAGCGAGGCGGATCACGAGCAGTGGAAGG
ATATGCTGCTGACCAACGTTTACGCTGTTGGCCTGACCCTGCAGGCAACGCTGCCGGCGCTCAGAGAGAGCCGTGGGCAGGTGCTGTTGACCGGCTCAGC
CGCCGGGCGCGCCACCATTCCCGGGTCCATGTACAGCGCCAGCAAATGGGCAGTAACCGGCATCGGCTACAACCTGCGCGAAGAGCTTCGCGGCACCGGT
ATTCGCGTTACCCTGCTTGAGCCGGGCATGGTGGATACCCCCTTCTTCGATGAGCCGCCGACCCATGCCATGAAGGATACCGATATCGCCAATGCGGCGA
TGTATGCCCTCGACCAGCCGGCGCATGTGGACGTCAACGAAATTCTGGTACGCCCCACGCCGGCGGTGGAATAGCACACCCCGGCCCCGTAAGGCCGGGT
AGTCAGTGCTTGAGTATCCGGCCCAGAAAGTTCTGCGTGCGTTCGTGTTGCGGGGCGTCAAACAGCTGCTCCGGTGGCCCCTGTTCGGTAATTTCGCCCT

>Halomonas jeotgali Hwa [NZ_AMQY01000003.1:50009..50069], dnaA> ig dnaN>, 61 bp

WebSkew GCmin pos. 2015280 (GCmin<>oriC1 49 kb; GCmin<>oriC2 11.0 kb; oriC2<>oriC1 38.8 kb)

>Halomonas subglaciescola ACAM 12 [NZ_LT670847.1:2064116..2064615], <rpmH oriC1 dnaA>, 544 bp

TCGCGGCCTTGGCTATTGCCATCGCCGGCACCGTCGGCCAGACCGCTGTCGTTACCGGCATTGGCTGGAGATGTGCTGCTGGGAACGGAGAGGCTATTAC
CGTCCGCGTCGGCAGTTGAAGCCGCGCCGCTATCGCGCTGAGTGCTTTCAACGGCAGGGGAGGGCTCAGCCGCTTGCCCGTAATCCTGGTTCCATTGAAC
CACTAGCAGATACGCCAGTACTGCCAATGGAACCAGTAATATCAGTCGTTTAACGTCCATGAGGGTTCTGCCCGCTGGGTGATGAACATGCAATGGCGCA
GTGTCATTTTACTCGACTCATGACACACGAAAGCCTGCCATACGGCAGGCCGGGGTCGAGCGGCGAGGTGTTGGCATCAAGCGTCACGTCGTGTATCGGC
CGGGGTGGGGGCGGCTTGGCTGTGCCGTGAGGCATCGCGCTGTAGCCGTTTCCACATCCCGTGTAGCTGGCGCTGTAACGTGGCGTTATCCAGCTCGTGA
ACGCCGCGCCGGGCAAGTATGACCACATCAACCGTGGGCAGTTGCTCCTGGCGTAACCGAAAAGACTCACGCGCCTGACGCTTGAAGCGATTGCGCTGAA
CGGCCAGCTTCACGTTTTTTTTACCGACAATCAGTCCGACCCGGGCATGACCACAGGTATTGAAGCGCGCAAGCGCCAGCATACCTTTGGCGTGGATCTT
GAGATCAGCCTGTTCGAAGACGTAACGGAAGTCCCCGGCGCTTAGCAGACGTAGCGGCCGGGGAAAGCCTTGATGGGACACACGCAATCTGAAATCAGGC
GCTCAGACGCTTGCGGCCTTTTGCACGGCGACGCGTAATGACGGCGCGGCCGCTTTTGGTTGCCATACGAGCACGGAAGCCGTGCGAGCGCTTGCGCTTG
AGAACGCTGGGTTGAAAAGTGCGTTTCATAACTCGTTATTCCCACGTGGTTTTAAGACGGAACGTCAATACTAGAAGACCCGTATCGGCAGGTCACCGAA

CGGGGATATCCAATTCAAGACCCGAAATTGTAGAGCCTTCCTGCGCCAGGTGCAATTTTTCCTTGCCGATATTGTTCAGCCGGCAGCTGCCGTGGCGGAA
AAGCGTCCGCTGGATTCAGCCAATATTCACCCGCAATGCCGTGCAATAGTTACTATTATATAAGATAACTAATGCTGTTATTAATGGTGGGATAATAAT
**TGTGGATAA**GCTTATTTTAACAAATTTAATCAAAGGGTTATTGGCATAAAAAATCTGTGTAAAAGCGTTGGAAAAGCAGCGCGAAAACTGTGATGAGGCG
GTGAACGGTTCAGGATAAGTCGAGAAATTGCCAAGTTGTCCGCAGTTTATCACCCAGTAGTGTCACGGTTTCCAAGAGACTTGTCCACAGGCCGGAAAGG
CCGCATTCGCGTGTGGATAACAGGGGCTTTGGGCGCTACAATGGCCGGCCTTTTTCAACCGATGGTGAAATGAGTGTCGCTCGCGCTTTGGCATCAGTGT

CTGGACTATTTGCAAGATGAGCTTAGCTCCCAGCAGTTCAACACCTGGATACGCCCGCTGCAGGCAGAACAGGGCGAGGCCAACGAGCTTTGCCTGCTCG
CGCCGAACCGCTTTGTGCGTGACTGGGTGAGCGACAAATATGCCAGGCGAATCAGCGAGCTCATGCGCGAGCTGGCACCCGCCAAACCGCCCAAGGTAAG
CCTGAGCGTGGGCAGCCGTCGCACCGCCGCCGCCGCTGCCCCGCAGACGCGTAATCTGGGGCACCCGGTCTCGGCCACACCTGCCGGGCAACGCGCGCCG
GCGGTGCATACCTCGCCGCGCGGATCGACCCGGGGTAATCATGGTAATGATCGTGAAATCAATCGCTTAAGCCAGCAAACGGGGCATCGTCAGGGCGAGC
GTCAGGTACAGGTGGAAGGTGGTCTTAAACACGGCAGTGGACTTAACCCGAACTTCACCTTTGATACCTTTGTCGAGGGTAAGTCCAACCAGTTGGCACG
GGCGGCGTCGCGTCAGGTGTCGGAAAACCCCGGCGGGGCATATAATCCGTTGTTTTTGTACGGCGGTGTGGGGCTGGGTAAAACGCACCTGATGCATGCC
GTTGGCAATGCCCTGGCAGAGCGCCGGGAAAATGCGCAGGTGATATATCTGCACTCCGAGCGTTTCGTTGCCGACATGGTCAAGGCCCTGCAGCTCAATG
CCATCAACGATTTCAAGCGTTTTTACCGTAGCGTTGATGCGCTGCTGATCGATGATATTCAGTTTTTTGCCGGTAAGGAGCGTTCGCAGGAAGAGTTCTT
TCATACGTTCAACGCGCTGCTTGAAGGCGGCCAGCAGATGATCCTGACCTCTGATCGCTACCCCAAGGAAATCAACGGCGTTGAAGAGCGCTTGAAATCG
CGTTTTGGCTGGGGGTTAACGGTGGCCATCGAGCCGCCTGAGCTTGAAACTCGGGTAGCTATCTTGATGAAAAAGGCCGATCAGGCCAAAGTAGACCTGC

>Halomonas subglaciescola ACAM 12 [NZ_LT670847.1:2026060..2026559], <mnmG oriC2 COG4221> <COG1126 (glnQ), 652 bp

TTTTCCAGCCCTTCGATGGAGCGCACTACCTCAAGCTGAACGTCAAACGGCAGCGAGGTGGAAATGCCGTTGGGGTACAGCTCGTGGGTATCGAGCCCTT
CGGGTTCGATAAAAATCTGATGGCTGTCTTTATCGGCAAAGCGGTGGACCTTGTCTTCGATCGACGGGCAATAGCGCGGGCCAACGCCTTCGATATCACC
CGAATACATCGGTGAGCGATCCAGATTGGCGCGAATAATTTCGTGGGTACGCGCATTGGTATGGGCAATGTGGCAACTCATCTGGCGCGGATGCATCTCG
CGATTGCCCAGATACGACATCACCGGCGTTGGCGTATCACCGGGCTGTTCATCAAGCCTGGAAAAATCCACCGTTCGCGCATCGATGCGTGGCGGGGTGC
CGGTTTTCAGCCGATCAACGCGAAACGGCAGTGCCCGCAGGCGCTCGGCCAGTGCGTTGGAGGGCGCATCGCCGGCGCGTCCACCAAGGGTTTTATCCAG
CCCGATGTGGATAACGCCGCCCAGGAACGTGCCGGTGCACAGCACCACGGTGTCGCTATGAAAGTGGATGCCGGTTTCGGTGACAACACCGCGCACGGTG
TTGTCATCCACGATCAAATCGCCGGCGGCCTGCTGAAAAATCGTCAAATTCGACTGGTTTTCCAGGGTACTGCGAATGGCCGCTTTGTAGCGAATGCGGT
CGGCCTGGGCACGGGTGGCACGAACGGCGGGGCCTTTGCGGGCGTTGAGCACGCGAAACTGGATGCCGCCCTTATCCGTTGCCAGCCCCATGGCGCCGCC
CAGCGCATCAATTTCCTTGACCAAATGGCTCTTGCCAATGCCGCCAATGGCGGGATTACACGACATTTGTCCCAGGGTTTCGATGTTGTGGGTCAGCAAC
AGGGTTTGACAGCCCATACGAGCGGAGGCCAATGCGGCTTCAGTTCCCGCATGGCCACCGCCGATGACAATCACGTCAAAGCGGTCGGGATAATTCAAAG

AACACCTCGTCTTGTGCGTTGGGCACGGATGTGAAAAGCGCAGCAGCGCTGAATGACTAGGGCCGGGAGTATAAACCTCCTGTGGGTAACCGTCAGGTTT
TTCCACACCGGGTTTTCTTCTGTTCTGCGCTTTTAGTAACTATATATATATAAATAAAAGAAGTTCTATTGTTGTTGTAACTACTGGTGTGGACGAATTC
**TGTGGAAAA**GCCATTATTCCCTTTATATCTCAATCAAATAGAGCGTTGATAGATGCGCGTGAAAGGTGCGCGTGGAGCGCGGACAAGCGCTGGAGGACCT
GTGTATAAGGAGGGTGCTTATCCACAGGCGCAGATAATGACAACTTCTCCACCGCTGCATACCGGTGTTGTTACCGCGGCTGTGAACAACCCGAGCGTGT
CTTGCGTAACGCCGCTAAACGCCCGCGTGACAGGGCCTGGCAGGTAAAAAAAGCGGAAAGGAAAAAAATACTATCCACAGGCAAAAAAAAGCCCGTGAAG

CGGGATTCTTGAGGAAAAAAAGGGCAGATAAGCCGTATTTCTATACCGGGAGAAGACGTGTCGTCTTGAGAGGATCTTGAGAGAGCCTTGAGAGAAAGAG
ACCAGCGGCGTTAATCTGGAGCATCGTCTGTCACTCAACAGGAGCTCAGTATGGATGATCGTGTATTGCTGATTACCGGCGCCTCCAGCGGTATCGGTGC
CGCCACCGCCCGCGCCGCCGCCGAGAAAGGTTACAAACTGGTATTGGCGGCACGCCGCCAGGATAAGCTCGAAGCGCTCGCCGAGGAGATCGGCAAGGAC
CGGGTCGTCACCGTGAGCTGCGACGTGACGTCGATGCAGGAACAGCAGGCCATGGTCGAGCAGGCACTGGAACGCTTTGGTCGCCTGGATGTGGTGTTTG
CCAACGCCGGGCGCGGGGGTTCAGCCGGCGGCTTCAGCGAGGCGGATCATGAGCAATGGAAGGAGATGCTGATGACCAACGTTTATGGCGTCGGCCTGAC
TCTGCAGGCGACGCTGCCCGCTCTTAAGCAAAGCCAGGGGCATGTGCTGCTGACCGGCTCTGCTGCCGGGCGTGCCACCATTCCCGGCTCCATGTATAGC
GCTAGCAAATGGGCGGTGACCGGCGTCGGCTACAACCTGCGGGAAGAACTGCGCGGTACGGGCATTCGCGTCACCCTGATCGAACCCGGCATGGTGGATA
CGCCCTTCTTCGACGAACGCCCCGCCCATGCCATGCAAGATGAGGATATCGCCAACGCGGTGATCTACGCCCTCGACCAGCCCGCCCATGTGGACGTCAA
CGAAATCCTGATACGCCCGACGCCGGCCAGGGAATGAACCCGAAAAACGCCGTTGAGCAGGCGTCAGTGCTTGAGCACCCGGGACAAAAAGCCCTGGGT
GCGCTCATGCTGCGGAGCATCAAACAGCTGTTCAGGCGGGCCCTGCTCGGTGATTTCCCCCTGATGGATGTAAATCACGCGGTCGGCCACTTCACGGGCA

>Halomonas subglaciescola ACAM 12 [NZ_LT670847.1:2066068..2066118], dnaA> ig dnaN>, 51 bp

WebSkew GCmin pos. 17100 (GCmin<>oriC1 23.4 kb; GCmin<>oriC2 1.6 kb; oriC2<>oriC1 21.9 kb)

>Halomonas rituensis TQ8S [NZ_QPIJ01000013.1:40270..40769], <rpmH oriC1 dnaA>, 543 bp

GCGATGAAGTCACGGCTGCTGGTCGAGGTGTCTTCCTCCACCGCTGGCATGCCCTCCTCCCCAGCTTGACGCTCGGGTGCGGTGGGAACGGCAAGCCCGT
TACCGTCAGGGGATTCTTCACTAACGCCATTGGCGGGCTGAGAAATCCCCGGTGCCTGGGCAACAGGTTCAGTAACTGGCTGCCCGTAATCCTGATTCCA
CTGGATTACCAGAAGGTAAGCCAGAATAGCTAAAGGAATCAGCAGAATAAGTCGTTTTACGTCCATAAGGGTTCTGCCCGATGGGTGGTGAACATGCTAA
TGGCGCAATGTCATTTGCTCGACCCTTGACACACGAAAGCCTGCCATGTGGCAGGCCGAGGTCGAGCGGCGAAAGGTATGCCGGTCAAGAACCACTGGTC
TGACGGGACGAAACCGTATGCTTGTCAGCTTGTGCTTCACGTAAGAGCCGTTTCCACATCCCGTGAAGCTGGCGATGCAGAGTGTCGTTGTCCAGCTCAT
GAACGCCGCGTCTGGCGAGTACCACGATATCCACAGCGGGAAGGTGGTGTTGCCTGAGACGAAAGGAATCACGAGCAAGCCGCTTGAAGCGATTACGGGC
AACGGCGTGCCTCACATTTTTCTTGCTGACTATCAAGCCAAGGCGGGGATATCCCAGGGCATTGAAACGCACAAGCGCCATCATGCCTTTGCCATGAACC
TTGATATCTGCATGCTCGAAAACGTGACTGTAATCCCCGGCGTTAAGCAAGCGTAAGCGTCGGGGAAAGTCTTGAGCGGATGCACGCAATCTGAGATCAG
GCGCTCAGACGCTTACGGCCCTTGGCACGACGACGCGCCAGAATGGCACGGCCGTTTTTGGTAGCCATGCGAGCACGGAATCCATGCGCGCGCTTGCGCT
TGAGAACGCTGGGTTGAAAAGTGCGTTTCATAACTCGTTATTCCCACGTGGTTTAGAACGGAAGATGAATTCCAGGGAGCCCCGGATAGAGAGGCTCTGA

CCGGGGGAGATTCAATTCAAGACCGGAAATTGTAGAGGCTTCGTGAGCCCGGTGCAATTCTTCCGTGCACAAAAGCCTTCAGGTGACGTGCACAAACGGT
CATGAATGGACTGGTGCAGGCTGAATGTAAAAAACTACCTCGCTATAATTACTAACTTAATAGAGTTTTTAATGCTGTTATTAATAGTGGGATAGTAAT
**TGTGGATAA**GATTAAAATTTAAATGTATTTCAATGCTATATAGTTATAAAAAATCTGTGCATAAGCTTGGGTAAAAGGTTGAGAGAGTGTTAGAAAGCT
GTGAATGAAACTGGGTGTTGATACTGGCTGACAAGCTATCCCCCGATTATTGGGTATTTTTCCAAAGCCCTCCTTTTAAGTTATCCACAGTCGAAAAGAC
AGCCAAAGGACGTGTGGATAACCTGAGCTTGGGGCGCTACAATGTGCGGCCGTTTCCAACCGATGGTGAGATGTGTGTCGCTCGCGCTCTGGCAGCAGTG

TCTGGATTACCTGCAGGACGAACTGAATTCCCAGCAGTTCAATACCTGGATTCGCCCATTGCAGGCGGAGGAAGGGGAGTCCAATGAACTGTGTTTGCTG
GCACCCAATCGCTTTGTGCGCGATTGGGTGAGTGACAAGTACGCCAAGCGTATCAGTGAGCTGATGCGCGAACTGGCCCCGGCTAAACCTCCTAAGGTGA
GCCTGACGGTAGGCAGTCGTCGTCCCGCTACGCCGGCTCCGCAACCACGAGAGCTCGGCCATCCTGTCTCTGCCGCCCCCCACAGATCAGTGGCACCCAC
GCCGCATACGCAGCGTCGAGGTGATATTGGCGATGAGCGTGAAATCGATCAATTGCGTGAGGAAAATGCCGCTCATCGTCGGCGAGGTAACGCCCGCCAG
GTTCAGGTGGAAGGTAGTCTCAAGCATGGTAGTGGACTGAATCCGAATTTTACCTTTGATACCTTTGTGGAAGGTAAGTCCAACCAGTTGGCGCGTGCGG
CATCACGTCAGGTATCGGAAAACCCCGGTGGGGCTTACAACCCTTTGTTTCTCTATGGTGGGGTAGGTTTGGGTAAAACCCATTTAATGCATGCCGTGGG
CAATGCGTTAGCCACGCGGCGGGAAAACGCTCGTGTGGTGTATCTGCATTCCGAACGCTTTGTCGCGGATATGGTAAAAGCGCTACAGCTTAATGCGATC
AACGATTTCAAGCGTTTCTATCGCAGTGTCGATGCGCTTCTAATTGATGATATTCAGTTTTTTGCCGGCAAAGAGCGCTCCCAGGAAGAATTCTTTCATA
CTTTCAACGCCCTCTTGGAAGGCGGTCAGCAAATGATTCTGACCTCCGACCGTTATCCCAAGGAAATCAGTGGGGTGGAGGAGCGCTTGAAGTCGCGGTT
TGGCTGGGGCTTGACCGTTGCAATCGAACCGCCTGAGCTTGAAACCCGAGTGGCGATTCTGATGAAGAAGGCCGATCAGGCCAAGGTCGATTTGCCGCAT

>Halomonas rituensis TQ8S [NZ_QPIJ01000013.1:18407..18906], <mnmG oriC2 <COG1126 (glnQ), 566 bp

TCGAGCGGACTACCTTGAGCTGAATATCGAACGGCAACGAGGTCGAAATCCCGTTGGGATAAAGCTCGTGGGTATCCAGCCCTTCAGGCTCGATAAACAC
CTGATGGCTATTCTTGTCGGCGAAACGATGCACCTTGTCTTCGATCGAAGGGCAGTAACGCGGCCCCACGCCCTCGATCACCCCGGAATACATCGGTGAG
CGTTCCAGGTTGGCGAGGATGATGTCGTGGGTCTGCTCGTTGGTATGCGCGATATGACAATTGACCTGCGCCGGGTGCATTTCCTGCTTGCCCATGTACG
ACATCACGGGAGTGGGTGTATCTCCTGGCTGAGTTTCGAGCCCCGAGAAATCCACACTCTTGGCATCCAGGCGTGGCGGTGTTCCGGTTTTTAGGCGGTC
GACACGAAATGGCAGTGCGCGCAGACGTTCGGCAAGCGCATTGGAGGGAGGATCGCCGGCCCGGCCGCCGCGACTCTGGTCCAGACCGATGTGGATAACA
CCGCCCAGAAAAGTCCCTGTGGACAACACAACCGTCTTGGAAAGAAAGCGAATGCCGGTTTCGGTCACCACACCGCAAACGCTGTCGTTATCCACAATCA
GATCGCCTGCCGCTTGCTGAAACAACGTCAGGTTGGGCTGGTTTTCCAGCATTCCCCGGATAGCGGATTTGTAGCGGATGCGATCGGCCTGGGCTCTTGT
CGCTCGAACTGCCGGGCCTTTGCGAGCATTGAGCACACGAAACTGGATACCACCGAGGTCGGTTGCAAGCCCCATGGCTCCCCCCAAGGCATCGATTTCC
TTGACCAGATGGCTCTTGCCGATCCCGCCAATCGCTGGATTACAGGACATTTGCCCCAGGGTTTCGATGTTGTGGGTCAGCAGCAGGGTCTGACTGCCCA
TGCGAGCGGAGGCCAGTGCGGCTTCAGTTCCCGCATGGCCTCCGCCGATGACGATGACGTCAAAGCGGTCGGGGTAATTCAAAAGACACCTCGTCTTGCA

CCGATGACGGCGCGATGTGATCAGCCGGCAAGGACGCTGCCTTGGCGGTATTCAAAATGAAGCTGGGCAGTATAAACCTCCTGTGGGTAACCGTCAGGTT
TTTCCACACCCTGTCAGCCAGCGACTCAAGTATTAAATAACAATATAAATATAAATAAAGAAGTTCTGTTGTTGTTGTAATTACTAGTGTGGGCAAAATC
**TGTTATTAA**GCCATTTAATCCTATAAAAATCATATTATTACGTTGTTGATGATCGCGGTTGAAAGCAGCGTACTACCTGCTGGTTAACGGTGAACCTCCT
GTGGATGAAAAGGCGCTTTATCCACAGGGCCAGTAATTCACGGTTTGTCCACCGCTCCATACCCTAGTTGTTACCGCACCTGTGAACAATTTTGCAAGCG
TCCTTATTGTTTCGCTCATGCCTGCCATGCTCTGCCTGGGCAAAGGTTTCTCCACTGGCCAAAAATAAAGTTGTCCACAGGCAAAAAAAAGGCCTATCAC

GAAGATAGGCCGGTAATTTAGGGTGCATTGCATGCCGAAACGTAGGTTTTAGTGAGTCAAAACCCGTGAAAGAAACGACTGGGTACGTTCGTGTTGAGGG
TTATCGAACAGGCTTTCGGGCTCTCCCTGTTCGGTGATTTCACCTTGATGGATAAAGATGACCCGGTCGGCCACTTCACGGGCAAAGCCCATTTCGTGCG
TCACGATCACCATGGTCATGCCTTCCTTGGCGAGCTCACGCATGGCGTCCAATACCTCCCCGATCATTTCCGGGTCCAGGGCCGAGGTGGGCTCATCGAA
CAGCATCAGGCGGGGTTCCATGGCCAAGGCACGCGCCAAGGCGACGCGTTGCTGTTGGCCACCGGAAAGCTGTGCGGGATATTTGTGAGCCTGATCGGTA
ATTCCCACGCGCTCGAGCAGGCGGTTGGCGGTTTCCAGGGCATCAGCGCGGCTCCAGCCGCGTACTTTCATCGGGGCGAGCGTTATGTTGTCGACCACGC
TGAGGTGCGGAAACAGATTGAACTGCTGAAACACCATTCCGACTTCGGTACGAATACGCTGCAAGGCGCGGTTGCTCTTGCCAAAGGGTAGAAGCTGGCT
GCCATCGACTTCCAGCTTGCCCAGCTGGTATTCCTCGAGGCCGTTGATACAGCGAATCAAGGTGGATTTGCCCGAGCCACTGGCCCCGATCACCACCACC
ACTTCACCGGGAACGATCTCCAGATCGATATCCTTCAATACGTGCAGGTTGCCAAAATGCTTGTTCAGCGCCTGCATGCGCACGATGGGTTGGGTTGCAT
CATCATTCATCGCAGGGGTCCTTATTGTCCGGCCAGGCCTCTACGCTCGAGTTGGCGCAGCAGGATTGAAAGCGTCCAGGTGATGACCAGGTACATTAGG
GCCACCATGAAATAGACCTCCAGAGCGGTAAACGTCGTGGCGATGTAGATCTGCCCCTGACGTACCAACTCGCCGACGCCGATCACCGAGAACAGCGAGG

>Halomonas rituensis TQ8S [NZ_QPIJ01000013.1:42217..42286], dnaA> ig dnaN>, 70 bp

WebSkew GCmin pos. 66300 (GCmin<>oriC1 25.0 kb; GCmin<>oriC2 1.3 kb; oriC2<>oriC1 23.7 kb)

>Halomonas zhuhanensis ZH2S [NZ_WTKP01000010.1:c41073..41572], <rpmH oriC1 dnaA>, 544 bp

ACGGCGATGAAGTCACGGCTCGTCGAGGCGTCTTCCTCCACCGCGGGCATGCCCTCCCCTCCAGCTTGACGCTCGGGTGCGGTGGGAACGGCAAGCCCGT
TACCGTCAGGGGATTCTTCACTAACGCCATTGGCGGACTGAGAAATTCCCGGTGCCTGGGCAACAGGTTCAGGAACTGGCTGCCCGTAATCCTGATTCCA
CTGGATTACCAGAAGGTAAGCCAGAATAGCTAAAGGAATCAGCAGAATAAGTCGTTTTACGTCCATAAGGGTTCTGCCCGATGGGTGGTGAACATGCTAA
TGGCGCAATGTCATTTGCTCGACCCTTGACACACGAAAGCCTGCCATGTGGCAGGCCGAGGTCGAGCGGCGAAAGGTATGCCGGTCAAGAACCACTGGTC
TGACGGGACGAAACCGTATGCTTGTCAGCTTGTGCTTCACGTGAGAGCCGTTTCCACATCCCGTGAAGCTGGCGATGCAGAGTGTCGTTGTCCAGCTCAT
GAACGCCGCGTCTGGCGAGTACCACGATATCCACAGCGGGAAGGTGGTGTTGCCTGAGACGAAAGGAATCACGAACAAGCCGCTTGAAGCGATTACGGGC
AACGGCGTGCCTCACATTTTTCTTGCTGACTATCAAGCCAAGGCGGGGATATCCCAGGGCATTGAAACGCACAAGCGCCATCATGCCTTTGCCATGAACC
TTGATATCTGCATGCTCGAAAACGTGACTGTAATCCCCGGCGTTAAGCAAGCGTAAGCGTCGGGGAAAGTCTTGAGCGGATGCACGCAATCCGAGATCAG
GCGCTCAGACGCTTACGGCCCTTGGCACGACGACGCGCCAGAATGGCACGGCCGTTTTGGTAGCCATGCGAGCACGGAAACCATGCGCGCGCTTGCGCT
TGAGAACGCTGGGTTGAAAAGTGCGTTTCATAACTCGTTATTCCCACGTGGTTTAGAACGGAAGATGAATTCCAGGGAGCCCCGGATAGAGAGGCTCTGA

CCGGGGGAGATTCAATTCAAGACCGGAAATTGTAGAGGCTTCGTGAGCCCGGTGCAATTCTTCCGTGCACAAAAGCCTTCAGGCAACGTGCACAAAGGGT
CTTGAATGGACTGGTGCAGGCTGAATGTAAAAAACTACTTCGTTATAATTACTAACTTAATAGGGTTTTAATACTGTTATTAATAGTGGGGATAGTAAT
**TGTGGATAA**GTTTGAAATTTAAATGTATTTCAATGCTATATAGTTATAAAAAATCTGTGCATAAGCTTGGGTAAAAAGGTTGAGAGAGTGTTAGAAAGCT
GTGAATGAAACTGGCTGTTGATATTGGCTGACAAGCTAACCTCCGATTGTTGGGGTATTTTTCCAAAGCCCTCCTTTTAAGTTATCCACAGTCGAAAAGA
CAACCAAAGGACGTGTGGATAACCTGCGCCTGGGGCGCTACAATGTGCGGCCGTTTCCAACCGATGGTGAGATGTGTGTCGCTCGCGCTCTGGCAGCAGT

GCCTGGATTACCTGCAGGACGAACTGAATTCCCAGCAGTTCAATACCTGGATTCGCCCATTGCAGGCGGAAGAAGGGGAGTCCAATGAACTGTGTTTGCT
GGCACCCAATCGCTTTGTACGCGATTGGGTGAGCGACAAGTACGCCAAGCGTATCAGTGAGCTGATGCGCGAGCTGGCACCGGCCAAACCGCCCAAGGTG
AGCCTGACGGTGGGCAGTCGTCGTCCCGCTACGTCGGCACCTCAACTTCGGGAGTTAGGTCATCCTGTCTCTGCTGCACCGCACAGATCAGTGGCGCCCA
CGCCGCATACGCAGCGCCGGGGTGATATTGGCGATGAGCGTGAAATCGATCAATTGCGTGAGGAAAATGCCGCTCATCGTCGACGCGGTAACGCTCGCCA
GGTTCAGGTGGAAGGTAGTCTCAAGCATGGTAGTGGACTGAATCCGAATTTCACCTTCGATACCTTTGTGGAAGGCAAATCCAACCAGTTGGCGCGTGCG
GCGTCACGTCAGGTATCGGAAAACCCCGGTGGGGCTTACAACCCTTTGTTTCTCTACGGTGGGGTAGGCTTGGGTAAAACCCATTTGATGCATGCCGTGG
GCAATGCGTTAGCCACGCGGCGGGAAAACGCCCGTGTGGTGTATCTGCATTCCGAGCGCTTTGTCGCGGATATGGTAAAAGCGCTACAGCTCAATGCGAT
CAACGATTTCAAACGTTTCTATCGCAGTGTCGATGCGCTACTGATTGATGACATTCAGTTTTTTGCCGGCAAAGAGCGCTCTCAGGAAGAATTCTTTCAC
ACATTCAATGCTCTTCTGGAAGGTGGTCAGCAAATGATTCTGACCTCCGATCGTTATCCCAAGGAGATCAGTGGGGTGGAGGAGCGCTTGAAGTCGCGGT
TTGGCTGGGGTTTGACCGTTGCAATCGAACCGCCCGAGCTTGAAACCCGGGTGGCGATTCTGATGAAGAAGGCCGACCAGGCCAAGGTCGATTTGCCGCA

>Halomonas zhuhanensis ZH2S [NZ_WTKP01000010.1:c64724..65223], <mnmG oriC2 <COG1126 (glnQ), 565 bp

TCGAGCGGACTACCTTGAGCTGAATATCGAACGGCAACGAGGTCGAAATGCCGTTGGGATAAAGCTCATGGGTATCCAGCCCCTCAGGTTCGATAAACAC
CTGATGGCTGTTCTTGTCGGCGAAACGATGCACCTTGTCTTCGATCGAAGGACAGTAACGCGGCCCAACCCCCTCGATCACCCCGGAATACATCGGTGAA
CGTTCCAGGTTGGCGAGGATGATGTCGTGGGTCTGCTCGTTGGTATGCGCGATATGACAATTGACCTGCGCTGGGTGCATTTCCTGCTTGCCCATGTACG
ACATCACGGGAGTGGGTGTATCTCCCGGCTGAGTTTCGAGCCCCGAGAAATCCACAGTCTTGGCATCCAGGCGCGGCGGTGTTCCGGTTTTTAGCCGGTC
GACACGAAATGGAAGTGCGCGCAGACGTTCGGCAAGCGCATTGGAAGGTGGGTCGCCGGCCCGGCCGCCGCGGCTCTGGTCCAGACCGATGTGGATAACA
CCACCCAAAAAAGTCCCTGTGGATAACACGACCGTCTTAGCCAGAAAGCGAATGCCAGTCTCTGTAACCACACCGCAAACGCTGTCGTTATCCACAATCA
GATCGCCTGCCGCTTGCTGAAATAACGTCAGGTTGGGCTGGTTTTCCAGCATTCCCCGGATCGCGGATTTGTAGCGGATGCGATCGGCTTGGGCTCTTGT
CGCTCGAACTGCCGGGCCTTTGCGGGCATTGAGCACACGAAACTGGATACCACCGAGGTCGGTTGCAAGTCCCATGGCTCCCCCCAAGGCATCGATTTCC
TTGACCAGATGGCTCTTGCCGATCCCGCCAATTGCCGGATTACAGGACATTTGCCCCAGGGTTTCGATGTTGTGGGTCAGCAGCAGGGTCTGACTGCCCA
TGCGAGCGGAGGCCAGTGCGGCTTCGGTTCCCGCATGGCCACCGCCGATGACGATGACGTCAAAGCGGTCGGGGTAATTCAAAAGACACCTCGTCTTGCA

CCGATGACGGCGCGATGTGATCAACCGGCAAGGATTCTCCCTTGGCGGTATTCAAAATGAAGCTGGGCAGTATAAACCTCCTGTGGGTAACCGTCAGGTT
TTTCCACACCCTGTCAGCCAGTGACTCCAGTATTAAATAACAATATAAGTATAAATAAAGAAGTTCTATTATTGTTGTAATTACTACTGAGGGCAAAATC
**TGTTATTAA**TCCATTTAAATTTATAAAAATCAGGTTGTTACGTTGTTGATGATCGAGAGTAAAAGCAGGGCACTACCTGCTGATTAACGGTGCAACTCCT
GTGGATGAAAAGGCGCTTTATCCACAGGGCCAATAATTCACTGCTTATCCACCGCTCCATACCCTAGTTGCTACCGCACCTGTGAACAATTTTGCAAGTG
CCCTTATTCTTCCCTTCGTGCCTACCATGCTCTGCCTGGGCAAGGTTTTCCCGCTAGCCAAAAATAAAGTTGTCCACAGGCAAAAAAAAGGCCTATCACG

AGGATAGGCCGGTCATTTGGCATGTATTGCATGCCGAAACGAAGGGTTTAGTGAGTCAAAACCCGTGAAAGAAACGACTGGGTACGTTCGTGTTGAGGGT
TATCGAACAGGCTTTCGGGCTCTCCCTGTTCGGTGATCTCACCTTGATGGATAAAGATGACCCGGTCGGCCACTTCACGCGCAAAGCCCATTTCGTGCGT
CACGATCACCATGGTCATGCCTTCCTTGGCGAGCTCACGCATGGCGTCCAATACCTCCCCGATCATTTCTGGGTCCAATGCCGAGGTGGGCTCATCGAAC
AGCATCAGGCGGGGTTCCATGGCCAAGGCACGCGCCAAGGCGACGCGTTGCTGTTGGCCGCCGGAAAGCTGAGCGGGATATTTGTGGGCCTGGTCGGTGA
TTCCCACCCGCTCGAGCAGGCGGTTGGCGGTTTCCAGAGCATCGGCGCGGCTCCAGCCGCGTACTTTCATCGGGGCGAGCGTTATGTTCTCGACCACGCT
GAGATGCGGAAACAGGTTGAACTGCTGAAACACCATCCCGACTTCGGTACGGATACGCTGCAAGGCGCGGTTACTCTTGCCAAAGGGTAGAAGCTGGCTG
CCATCGACTTCCAGTTTGCCCAGCTGGTATTCTTCAAGGCCGTTGATACAGCGAATCAAGGTGGATTTGCCCGAGCCACTGGCCCCGATCACCACCACCA
CTTCACCGGGAACGATCTCCAGATCGATATCCTTCAATACGTGCAGGTCGCCGAAATGCTTGTTCAGCGCCTGCATGCGCACGATGGGTTGGGTTGCATC
ATCATTCATCGCAGCAGTCCTTATTGTCCGGCTAGGCCCCTACGCTCGAGTTGGCGCAGCAGGATTGAAAGCGTCCAGGTGATGACCAGGTACATCAGGG
CCACCATGAAATAGACTTCCAGAGCGGTAAACGTCGTGGCGATGTAAATCTGCCCCTGACGTACCAGCTCGCCGACGCCGATCACCGAGAACAGCGAGGT

>Halomonas zhuhanensis ZH2S [NZ_WTKP01000010.1:c39554..39624], dnaA> ig dnaN>, 71 bp

< page 1 | #106 | Halomonas pacifica NBRC 102220

WebSkew GCmin pos. nd (GCmin<>oriC1 nd; GCmin<>oriC2 nd; oriC2<>oriC1 nd)

>Halomonas pacifica NBRC 102220 [NZ_BJUK01000023.1:c45735..46234], <rpmH oriC1 dnaA>, 556 bp

CAGGTCCAGTACATCGGTGGTCACGGCGATCAGGTCGCGGCGGCTACCCTGGGCGCCTGGTGCTTCCGGCATCTCGCCATCGATGGCGTTCGCCCGAGGC
GCCGTCGGCACGGCCAGGCCATCATCGCCGGGGGAGGCGGCTTCACCGCTCTGGGTGATGGTGGGGGCGGTAGGCTCGGGGGTGGCCTGACCGTAATCCT
GATTCCACTGCACGATCAGCAGATAGGCGAGCACGGCCAATGGAATCAGTAGGAGTAGTCGTTTTACGTCCATGGGGATCCTGCCCGGTGGGTGATGAAG
GTGTGAGCACGCCGTCTTCGAGACACACGAAAGCCTGCCGGAGGCAGGCTGAAGGCGTCACCGTTACGGCGACAACGGCGGTGTCAGGTCAGGACCCTGT
GGCGTCGACCGCCATCTTGCGAGCGTCTTTTTCCAGGCGTCGCCACATGCCGTGAAGCTGGCGGTGCAGGGTGGCATTGTCCAGCTCATGCGCCCCGCGT
CGAGCGAGGACCACGATATCCACCGCGGGCAGGCGCTGCTGCTGCAGTCGGATCGACTCGCGCACCAGCCGTTTGAGGCGGTTGCGATCGACGGCACGAC
GCACATTCTTCTTGCTGAAGACCAGGCCGAGGCGCGCATGACCGAGATCATTGCGCCTTGCCAGGGCCATCAGCCCCTTGCCATGGACCTTGAAGTCCGC
ATGGTCGAAGACATGTCGATAGTCCCCGGCCGTCAGCAGACGCAGCCGCCGGGGAAAGCCCTGACAGGACACGCGTGATCCGCGATCAGGCGCTCAGACG
CTTACGGCCCTTGGCGCGACGGCGCGCCAGGACGGCACGACCGTTCTTGGTGGCCATGCGAGCACGGAAGCCGTGCACACGCTTGCGCTTGAGAACGCTC
GGTTGAAAGGTGCGTTTCATAACTCTCGATTCCCACGTGGTTGGTTGGACTGAACGTCAGAGTAGCGGCCCGGTTTCCCCGACGGATTACGCGGGAGCTT

TCGGGTCAAGACCGGGAATTCTAGAGAATTCGCTGCCCCGGTGCAATTTTTCCGCGCCGCCAGGGCGAAGAAACTGCGCCACGATGGGGGAGCCGGCGTG
CCGGAATGGGAGGGGCGCGCCATTCGGCCAGGCTAGTGCACTCAATTATAATAATTTAGCCCTTTTAAACGATGTTGTTATTAATACTGGGGAAAAGATT
**TGTGGATAA**CTGATATTAGTCATTTCATATCAGTGGGCTAGCCTTTAGAAAAGGGTGTGGAAAAGTGCCGGGAAACGGTGGGGAAGCTGGGAGGAGCCT
GGTGACGAGAGGCGCGTTCGTCCACAGGACCCGGGGCGTCCACCCTGGGTCCTCCGTTTCTCCCAGGGGTTGGGGACAAGTTGTCCACAGCACCGCCGGT
GGTGCCTCTCGTGTGGATAACCCGGGCCTGTGACGATACAATGGCCGGTCATTCGACAACTGCATGGTGAGGTCGCGTGTCCGTCGCTCTTTGGCAACAG

TGTCTGGATTACCTGCAGGACGAGCTGAACTCGCAGCAGTTCAATACCTGGATTCGTCCCCTGCAGGCGGAGGAAGGCGAGTCCAATGAGCTGCGCCTGC
TGGCTCCCAATCGCTTCGTGCGCGATTGGGTCAGCGACAAGTACATCAAGCGCATCAATGAACTGCTGCGTGAGCTGGCCTCGGGCAAACCGCCCAAGGT
GAGTCTCACGGTGGGCAGCCGCGGCAGCGTGCCCAGCCCGCAGCCCCGCAACCTGGGGACCCCGGTCTCGGCGACGCCTCGCCGTCCCAGTGCCCCGGCG
GTGAAGACACCGCCCCGCGGCGACGACTATGCCGACGAGCGGGAGATCGAGGCCCTCGAGGAGAAGGCCCCGCGCCGCGGCGAGCGGCAGGTGCAGGTGG
AGGGTAGCCTCAAGCACTCCAGCGGCCTCAACCCCAACTTCACCTTCGAGACCTTCGTCGAGGGCAAGTCGAACCAACTGGCTCGAGCCGCCTCCCGCCA
GGTCTCCGAGAACCCCGGCGGGGCCTACAACCCGCTGTTTCTGTATGGCGGCGTGGGCCTGGGCAAGACCCACCTGATGCATGCGGTGGGCAATGCCCTG
GCCGGGCGTCGCGAGAACGCCAAGGTGGTCTACCTGCACTCCGAGCGCTTCGTGGCCGATATGGTCAAGGCGCTGCAGCTCAATGCCATCAACGACTTCA
AGCGCTTCTACCGCAGCGTCGACGCCCTGCTGATCGATGATATCCAGTTCTTCGCCGGCAAGGAGCGGTCCCAGGAGGAGTTCTTCCACACCTTCAATGC
CCTGCTCGAGGGCGGCCAGCAGATGATCCTCACCTCGGATCGCTATCCCAAGGAGATCAGCGGGGTAGAGGAGCGCCTCAAGTCGCGCTTCGGCTGGGGG
CTGACGGTGGCCATCGAGCCGCCGGAGCTGGAGACGCGGGTGGCGATCCTGATGAAGAAGGCCGATCAGGCCAAGGTCGATCTGCCCCACGACGCGGCCT

>Halomonas pacifica NBRC 102220 [NZ_BJUK01000013.1:c1550..2049], <mnmG oriC2 <COG1126 (glnQ), 544 bp

CCAGTCCACGGATGGAACGCACCACCTGCAACTGCACGTCGAAGGGCAGCGAGGTGGAGATGCCGTTGGGGTAGAGCTCGTGGGTATCCAGGCCCTCCGG
CTCGACGAACACCTGGTGGCTCGCCTTGTCGGCGAAGCGATGCACCTTGTCCTCGATAGACGGGCAGTAGCGCGGCCCGACGCCCTCGATCACCCCGGAA
TACATCGGCGAGCGATCCAGGTTGGCGAGGATGATCTCGTGGGTGCGCTCGTTAGTATGGGCGATATGGCAATCCACCTGGCGCGGGTGCATCTCGCGAC
TGCCCAGGTAGGACATCACCGGGGTCGGGGTATCGCCGGGCTGGGCCTCGAGCACCGAGAAGTCGACGCTCTTGGCATCGATGCGCGGCGGGGTACCGGT
CTTCAGCCGGGCCACGTTGAAGGGCAGGGCGCGCAGGCGCTGGGCCAGTGCATTGGAGGGCGGATCCCCGGCACGGCCACCGCGACTCTGGTCGAGCCCG
ATATGGATCACCCCGCCGAGGAAGGTCCCGGTGCACAGCACCACGCTCTCGGCGTGGAAGCGAATGCCGGTCTCGGTGACCACCCCACGCACGGTGTCGT
TATCCACAAGCAGATCGCCGGCGGCCTGCTGGAAGAGCGTCAGGTTGGGCTGGTTCTCCAGCAGGCCGCGGATCGCCGCCTTGTAGCGCACCCGGTCCGC
CTGGGCTCGGGTCGCCCGCACCGCCGGGCCCTTGCGGGCGTTGAGCACCCGGAACTGGATGCCGCCCAGGTCCGTGGCCAGGGCCATGGCCCCACCCAGG
GCATCGATCTCCTTGACCAGATGACTCTTGCCGATGCCGCCGATGGCGGGATTGCAGGACATCTGGCCCAGGGTCTCGATGTTATGGGTCAGCAGCAGGG
TCTGACAGCCCATACGGGCGGAGGCCAGGGCGGCTTCGGTTCCCGCATGGCCACCGCCGATGACGATGACGTCAAAGCGGTCGGGGTAATCCACGTTGCA

CCTCGAATGGCGCCGGCGGCGCCGCAGTAGAGCTCGGAGCAGAAAATCAGCCGCACAGTATAAACCCCCTGTGGGTAACCGTCAGGGGTTTTGCACAACC
TCGTCGGCCTTTCCTTGTCTTCACTTCCTAACTAGTTATTAAAGAAGAGAGATAAAAGAGAAGTTCTGTTGCTGTTGTGTCTATTGGTGTGGACAAAATC
**CGGGGACAA**GTGAATAAACTTCTTTTTTAACAACAGGATGCCGTGTTGACGAGTCTCGGGAGAAGCTCGTGGGAGGCTGCGAACAAGCGCACCCGAGGCT
GTGGATGAAAGTCCCACTTATGCACAGCGCGAATGGTGCACCGTTTTTCCACCGTCGCCTACCCGACTTGCGCCCGCGGTGGTCAACAGCCGCCGAATCG
CGCGCCAGGCCCGGCATGGTGGGCCCTGGGGGCATTCGTCGAAAAATTGTATCCACAGGCAAAAAAAGGGCCTGTCGATAAGCCGACAGGCCCTTTTTC

CGCCGAAGGTGGTCGATGATGGCCGTGCAGGGGCGCCCCTAGTGCTTCAACACCCGGGCCAGAAAGCCGCGGGTGCGTTCATCCTGAGGCGAATCGAAGA
TGCGCTCGGGCGTCGCCTCCTCGGCGATCTCGCCCTTGTGGACATAGATGACCCGATCGGCCACCTCCCGGGCGAAGCCCATCTCGTGGGTGACGATGAT
CATCGTCATGCCCTCCCGGGCCAGTTCGCGCATGGCATCCAGCACCTCGCCGATCATCTCCGGATCCAGGGCCGAGGTGGGCTCGTCGAAGAGCATTAGC
CGAGGTTCCATGGCCAGGGCGCGGGCCAGGGCGACCCGCTGCTGCTGGCCCCCGGAGAGCTGGCTGGGGTACTTGTCGGCCTGGTCGCGGATGCCCACCC
GGTCGAGCAGTCGCTCGGCGCTCTCGCCGGCCTGGGCGGCGGTGACACCGCGTACCTTGAGGGGCGCCAGGGTGACGTTGTCGCGCACCGAGAGGTGGGG
AAAGAGGTTGAACTGCTGGAAGACCATGCCCACCTCGGTGCGGATGCGCTGCAGGGCCTTGGCCCCCTTGCCGTGGGGCAGCAGGTCCTTTCCATCCACC
ATGATATGACCGTCCTGGAACTCCTCCAGGCCGTTGATGCAGCGGATCAGGGTCGACTTGCCGGAGCCGCTGGCGCCGATGATCACCACCACCTCGCCGG
GCATGACCTCCAGATCGATATTCTTGAGCACATGCAGGCTGCCGAAATGCTTGTTGAGCTTGCGCATGCGCACGATGGCTTGGGACTCGTGTTGGCTCTC
TGTCATGGATGGGGTCCCTTATTGTCCGACCAGGCCGCGGCGTTCCAGCACACGCAGCAGCAGGGAAAGGCTCCAGGTGATGGCCAGGTAGAGCAGGGCC
ACCATCAGATAGACCTCCAGGGCGGTGAAGGTCGTGGCGATATAGATCTGCCCCTGGCGCACCAGCTCGCCGACGCCGATCACCGAGAACAGCGAGGTGT

>Halomonas pacifica NBRC 102220 [NZ_BJUK01000023.1:c44234..44297], dnaA> ig dnaN>, 64 bp

WebSkew GCmin pos. 281480 (GCmin<>oriC1 35.8 kb; GCmin<>oriC2 9.5 kb; oriC2<>oriC1 26.3 kb)

>Halomonas salipaludis WRN001 [NZ_NSKB01000003.1:316987..317486], <rpmH oriC1 dnaA>, 501 bp

GCCCTGAGGGTCGATGCGCACGTCGAGAACATCGGTGACCACCGCGATCAGGTCACGGCTGCGGGGCTCGCTTTCGGTGGCCTGCGGAAGCTCGCCCTCC
ACCGTCTCCTGGGGAGCGTGGGGAACGCTCAGGCCGCCATCGTCGTCGGGCTCACCGCTTTCGACCTGCGGGGCACCGCCGTTGGTGAAGGAGGGGGCGT
CCGGCGCAGGGGCTGCCTGGCCGTAGTCCTGGTTCCACTGCACGACCAGCAGATATGCCAGCACGGCCAGCGGAACCAGCAGAAGTAGTCGTTTTACGTC
CATGAGGGTCCTGCCCGATGGGTAATGAACATTCTGGCGGCAACATCGACATCAGCAGTGCCGATACGCACGAAAGCCTGCCAGGGGCAGGCTCGAAGCG
TCACCGCGAAGACGGCGAGGAAGTGTCCTGGCTGACAGCGGCCGTCGACTGCTTGCGCACATCCTTCTCCAGACGCCGCCACATGCCGTAAAGCTGGCGG
TGCAGGGTGTCGTTGTCCAGCTCGTGTACGCCTCGCCGGGAGAGTACCACGATGTCGACAGCGGGAAGTTGGTGCTGCTGAAGGCGGATGGATTCGCGAA
CCAGTCGTTTCAAACGATTGCGATCCACGGCCCGGCGCACGTTCTTCTTGCTGAAGACCAAGCCGACGCGGGGATGACCCAGTGCGTTGGGGCTTGCCAG
TGCCAGCAATCCTTTGCCGTGAATCTTGAACGCCGCGTGATCGAAGACGTGTCGGTACTCCCCGGCATTCAGCAGTCGCAGATGCCGGGGGAAACCCTGA
CAGGACACACGCAAGCCGAGTTCAGGCGCTCAGACGCTTGCGGCCCTTGGCGCGTCGACGCGCCAGCACGGCGCGGCCATTCTTGGTAGCCATACGGGCA
CGAAAACCGTGCACGCGCTTGCGCTTGAGAACGCTGGGTTGAAAAGTGCGTTTCATAACTATCGATTCCCACGTGGTTAATTGGACGTAACGTCAATGGT

GAGGCGCCCTGGCTCCGGCGGGGAACCCGGGGTAATTCGGCTCAAGACCGGGCATTTTAGTGAATTCGGGCCACGGGTGCAATTTTTCCTCACCGTCATC
GCATGGGACTCGGCCGCTGCCCAGCAACCGTTCTAGATATCGGCGTGGGAACGCTTGCCGGTGATTAGAACATCTTGTTATTAAAATTGGGGATGATAAA
**TGTGGATAA**GTATTTTTTCATCATGTATTACAGTTGGATAGCTCTATCTCATGGGTGTGGGAAACACTTTGATAACCGGCTATGACCTGACGCCTGGCCG
GTTACGCAACCGCGCCTGGGTAACGCTTCGGCGGGCGCTCGACGAGCGCGGCGTCGGCGAGTTTTCCACAGCGGCGAGGCACGACCGGTGCGTGTGGATA
ACTCGCTGCTTGGGCGATACAATGGCCGGTCATGCATCAACGGGATGGTGAGGTCGAGTGTCGGTCGCTCTTTGGCAACAATGTCTGGATTTCCTGCAGG

ACGAGCTCAGCTCGCAGCAGTACAACACCTGGATTCGCCCCCTCCAGGCGGAACAGGGGGAGTCCAACGAACTCTGCCTGTTGGCGCCCAATCGCTTCGT
GCGTGACTGGGTAAGCGACAAGTACATCAAGCGAATCAACGAATTGCTGCGTGAACTGGCGCCAACCAAGCCACCCAAGGTGACGCTGGCAGTCGGCAGC
CGCCGGGCCGTACCAAGCCCGCAGCCACGCGACCTGGGCAATCCTGTGTCTGCTGCGCCGCGCCAGCCTGCATCGCCGGCGGTACACACGCCGCCGCGTG
GCGACCTCGCCGACGAACGCGAGATCGAGGCGCCCATCGATCCACCGTCGCCACGTGGTGGGTCCCGTGCATCATCACGGGGGGGGGCGAACGGCAGGT
ACAGGTGGAGGGCAGCCTCAAGCACCAGAGTGGCCTCAACCCCAATTTCACCTTCGAAACCTTCGTCGAGGGCAAATCGAACCAGCTGGCTCGTGCTGCT
TCCCGCCAGGTATCGGAAAACCCCGGTGGCGCCTATAATCCGCTGTTTCTTTACGGTGGAGTGGGTCTGGGTAAGACTCACTTGATGCATGCGGTCGGTA
ATGCACTCGCCACTCGCCGCGAGAATGCCACGGTGGTGTACCTGCACTCCGAGCGTTTCGTGGCCGACATGGTCAAGGCGCTGCAGCTAAATGCCATCAA
CGACTTCAAGCGTTTCTATCGCAGCGTCGACGCCCTGCTGATCGACGATATCCAGTTCTTCGCCGGCAAGGAGCGTTCCCAGGAAGAGTTCTTCCACACC
TTCAATGCGTTGCTCGAGGGCGGTCAGCAGATGATCCTGACGTCCGACAGATATCCAAAAGAAATCAATGGTGTAGAAGAGAGGCTCAAGTCTCGCTTCG
GATGGGGGCTAACCGTTGCCATCGAGCCACCGGAGCTTGAGACGCGAGTGGCGATTCTGATGAAGAAGGCAGATCAGGCCAAGGTTGACCTGCCTCACGA

>Halomonas salipaludis WRN001 [NZ_NSKB01000003.1:290732..291231], <mnmG oriC2 <COG1126 (glnQ), 544 bp

CCAGTCCCTTGATCGAGCGCACTACCTGCAGCTGCACATCGAAGGGCAGCGAGGTGGAGATCCCGTTGGGATAGAGCTCGTGGGTATCCAGGCCCTCGGG
TTCGACGAACACCTGATGGCTGGCCTTGTCGGCGAAGCGATGGACCTTGTCCTCGATCGACGGGCAGTAGCGCGGCCCCACGCCTTCGATGACGCCGGAG
TACATTGGCGAGCGATCGAGATTGGCGAGGATGATCTCGTGCGTTCGCGCGTTGGTGTGGGCGATATGACAGCTGACCTGGCGCGGGTGCATCCCGCGTT
CACCAAGGTAGGACATCACCGGGGTGGGACTGTCGCCGGGCTGCTCCTCGAGCTGGCTGAAATCGACGCTCTTGGCGTCCAACCGCGGCGGTGTGCCGGT
TTTCAGCCGTGCCACGTTGAACGGCAGCGCTCGCAGTCGCTCGGCGAGGGCGTTGGACGGCGGGTCACCGGCACGGCCTCCGCGGCTGCGATCGAGGCCG
ATATGGATCACCCCGCCGAGGAAGGTTCCGGTACACAGCACCACGCTCTCGGCGTGAAAACGGATGCCGGTTTCAGTGACCACGCCGCGTACGCTGTCGC
CATCGACGATCAGGTCGCCGGCGGCTTGCTGAAACAGCGTCAGGTTGGGCTGCTGCTCGAGCATTCCGCGAATCGCAGCCTTGTAGCGCACCCGGTCAGC
CTGAGCGCGAGTGGCCCGTACCGCCGGCCCCTTGCGTGCATTGAGCACCCGGAACTGGATCCCTCCCCGGTCGGTGGCCAACGCCATGGCACCGCCCAAT
GCATCGATCTCCTTGACCAAATGGCTCTTGCCGATCCCGCCGATGGCGGGATTGCAGGACATTTGGCCCAGGGTCTCGATGTTATGGGTCAGCAGCAGGG
TCTGACAGCCCATGCGAGCGGAGGCCAGAGCGGCTTCGGTTCCCGCATGGCCACCGCCGATGACGATCACGTCAAAGCGGTCGGGGTAATCCACGTTGCA

CCTCGAATGGCACCACTGGTGCCGAAAAAATGATCAATGCTGGAAATTGGCTGCCCAGTATAAACCTCCTGTGGGTAAGCGTCAGGGGTTTTCCACACCG
TCTTGGTCAGTCCTGCCTCACCTGACTAAGACAAGAATAAAAATAAGTAGAGTAAGAAAGAAGTTCTGTTGTTGTGGTAACTACTGGTGTGGACACTTTC
**TGTGGGTAA**GTAGATTTTCCTTTTTTACATCAATGGATTGTAGTGTTGAGGGATTGACGGACAAGAGGCGTATCGGCTGCGGACAGATGGCCGTGAGCCT
GTGGACGAAAAGGCGTTTTGTGCACAGCCACAATCGCCCACCGTTTATCCACCGGGCTTGGTGGTGCTTGTCACCGTGACTGCCCACAGTCGGTCAAAAA
ATCACCAGCACCATGGTGGTGGGGGCTGTAGAGCGATTCGATGGCGGTGGGATTTTTTTTGTCCACAGGCAAAAAAATGCCAAAAAAAAGCCAGTCGATA

CACGACTGGCCTCTCTCAGTATCCTGGTTGACGCGATATCAGTGTTTGAGAACCCGCGACAGGAAGTTTTGGGTGCGATCATGCTGGGGGCTATCGAATA
GCGTTGCGGGAGGCCCCTGTTCGACGATTTCACCGCTGTGGATGTAGACCACGCGATCGGCGACTTCGCGGGCGAAGCCCATCTCGTGGGTGACGATCAT
CATCGTCATGCCTTCCCTGGCCAGCTCGCGCATGGCATCGAGGACCTCACCGATCATCTCCGGATCCAGCGCCGATGTCGGCTCGTCGAACAGCATCAGC
CGAGGCTCCATGGCCAATGCTCGGGCCAAGGCGACGCGCTGCTGCTGGCCGCCGGAGAGCTGGCTGGGGTACTTGTCGGCCTGGTCGCTGATACCGACTC
GCTCGAGCAGACGCTCGGCGGTGCTTGCTGCCTGCTGCCTGCTGACGCCGCGAACCTTGAGTGGTGCCAGGGTGATGTTGTCGCGCACGCTGAGGTGCGG
AAACAGGTTGAATTGCTGGAAAACCATGCCGACTTCGGTACGAATGGTCTGTAGTGCTTTGCTGCTCTTGCCGTGCGGGGTCAAGGCGTTGCCGTCTACC
TCGATATGACCGGCCTGAAACTCTTCGAGGCCGTTGATGCAGCGGATCAGCGTCGACTTGCCGGAGCCGCTGGCACCGATGATTACCACCACTTCACCGG
GGGCGACGCTGAGCTCGATGTCCTTGAGGACGTGCAGGCTGCCGAAGTGCTTGTTGACCTGCTGCATATGGACGATGGGCTGGGCGGCGGGGGATTGGGT
TGTCATGGATAGCTCCTTATTGTCCGACCAGGCCGCGGCGCTCCAACTGGCGCAGGATCAGCGATAGCGTCCAGGTAATCGCCAGGTACATCAGCGCGAC
CATGAAGTAGACCTCCAGAGCGGTGAAGGTGGTGGCGATGTAGATCTGTCCCTGGCGCACCAGCTCACCGACGCCGATCACCGAGAATAGCGAGGTGTCC

>Halomonas salipaludis WRN001 [NZ_NSKB01000003.1:318932..318979], dnaA> ig dnaN>, 48 bp

WebSkew GCmin pos. 313236 (GCmin<>oriC1 36.0 kb; GCmin<>oriC2 8.3 kb; oriC2<>oriC1 27.7 kb)

>Halomonas qiaohouensis DSM 26770 [NZ_JARWAM010000001.1:c276962..277461], <rpmH oriC1 dnaA>, 502 bp

TGAGGGTCGATACGCACGTCGAGGACGTCAGTGGTCACCGCGACCAGGTCGCGGCTCCGCGTTTCGGTCTCGGCGGCTGAGGGCAGATCGCTCTCCACCG
CCGCCTGCTGATCGGCGCTGGGAACGCTCAGGCCGCCATCGTCGTCGGTCTCAGCACTGTCGACCTGCGGGGCCCCGCCGTTGGTAAAGGAGGGCGCGTC
CGGCTCAGGCGCTGCCTGGCCGTAGTCCTGGTTCCACTGCACGACCAGCAGATATGCCAGCACGGCCAGCGGAACCAGCAGAAGTAGTCGTTTTACGTCC
ATGAGGGTCCTGCCCGATGGGTAATGAACATTCTGACGGCAACATCGATATCAGCGGTGCCGATACACACGAAAGCCTGCCAGAGGCAGGCTCGAAGCGT
CACCGCGAAGACGGCGAGGAAGTGTCCTGGCTGACAGCGGCCGTCGACTGCTTGCGTACATCCTTCTCCAGACGCCGCCACATGCCGTAAAGCTGGCGGT
GCAGGGTGTCGTTGTCCAGCTCGTGTACGCCTCGCCGGGAGAGTACCACGATGTCGACAGCGGGAAGTTGGTGCTGCTGAAGGCGGATGGATTCGCGAAC
CAGTCGTTTCAAACGATTGCGATCCACGGCCCGGCGCACGTTCTTCTTGCTGAAGACCAAGCCGACGCGGGGGTGACCCAGTGCGTTGGGGCTTGCCAGT
GCCAGCAATCCTTTGCCGTGAATCTTGAACGCCGCGTGATCGAAGACGTGTCGGTACTCCCCGGCATTCAGCAGTCGCAGATGCCGGGGAAACCATGAC
AGGACACACGCAAGCCGAGTTCAGGCGCTCAGACGCTTGCGGCCCTTGGCGCGACGACGCGCCAGCACGGCGCGGCCATTCTTGGTAGCCATGCGGGCAC
GGAAACCGTGCACGCGCTTGCGCTTGAGAACGCTGGGTTGAAAAGTGCGTTTCATAACTATCGATTCCCACGTGGTTAATTGGACGGAACGTCAATGGTG

AGGCGCCCTGGCTCCGGCGGGGAACCCGGGGTAATTCGGCTCAAGACCGGGCATTTTAGTGAATTCGGGCCACGGGTGCAATTTTTCCTCACCGTCATCG
CCTAGGGCCTTGGCCGTCGTCCAGCACGCGTTCTAGACAGCGCCGCGGCTAGGCTTGCCAGTGATTAGAACATCTTGTTATTAAAATTGGGGATGGTATA
**TGTGGATAA**GCCATTTTTCATCATGTATTACAGTTAGATAGCACTCTTCGATGGGTGTGGGAACACTTTGATAACTGACGGCGACCCGCGTCTCGGTGG
GTTGCGCGGTGGCGCCTGGGTAACGCTTCGGCGGGGCTCGATGGAGTGCGGCGCCGGCGAGTTTTCCACAGCGATGAGGCATGGTCGGTGCGTGTGGATA
ACTCGCTGCTTGGGCGATACAATGGCCGGTCATGCATCAACGGGATGGTGAGGTCGAGTGTCGGTCGCTCTTTGGCAACAATGTCTGGATTTCCTGCAGG

ACGAGCTCAGCTCGCAGCAGTACAACACCTGGATTCGCCCCCTCCAGGCGGAACAGGGGGAGTCCAACGAACTCTGCCTGTTGGCGCCCAATCGCTTCGT
GCGTGACTGGGTAAGCGACAAGTACATCAAGCGAATCAACGAGTTGCTGCGTGAGCTGGCACCCACCAAGCCGCCCAAGGTAACGCTGGCAGTCGGCAGT
CGCCGGGCCGTGCCAAGCCCGCAGCCACGCGACCTGGGCAATCCGGTGTCTGCTGCGCCGCGCCAGCCTGCATCGCCGGCGGTACACACGCCGCCGCGTG
GCGACCTCGCCGACGAGCGCGAGATCGAGGCGCCCAGCGATACACCGTCGCCACGCGGCGGGTCCCGCGGGGGAGGCGCATCGCGCGGTGGTGGCGAGCG
CCAGGTACAGGTTGAGGGCAGCCTCAAGCACCAGAGTGGCCTCAACCCCAATTTCACCTTCGAAACCTTCGTCGAGGGCAAGTCGAACCAGCTGGCCCGC
GCCGCCTCGCGGCAAGTCTCGGAAAACCCTGGTGGTGCCTATAACCCGCTGTTCCTGTACGGCGGGGTTGGCCTGGGTAAGACTCACCTGATGCACGCCG
TGGGCAATGCGCTGGCCACCCGCCGCGAGAATGCCACCGTGGTGTACCTGCACTCCGAGCGCTTCGTGGCCGACATGGTCAAGGCGCTGCAGCTCAATGC
CATCAACGACTTCAAGCGCTTCTATCGCAGCGTCGACGCGCTGCTGATCGACGATATCCAGTTCTTCGCCGGTAAGGAGCGTTCCCAGGAAGAGTTTTTC
CACACCTTCAATGCGTTGCTCGAGGGCGGTCAGCAGATGATCCTGACGTCCGACAGGTATCCCAAAGAAATCAATGGTGTAGAAGAGAGGCTCAAGTCTC
GCTTCGGATGGGGGCTAACCGTTGCCATCGAGCCACCGGAGCTTGAGACGCGAGTGGCGATTCTGATGAAGAAGGCAGATCAGGCCAAGGTCGATCTGCC

>Halomonas qiaohouensis DSM 26770 [NZ_JARWAM010000001.1:c304650..305149], <mnmG oriC2 <COG1126 (glnQ), 544 bp

CCAGTCCCTTGATCGAACGCACCACCTGCAGCTGCACATCGAAGGGCAGCGAGGTGGAGATCCCGTTGGGGTAGAGCTCGTGGGTATCCAGGCCCTCGGG
TTCGACGAACACCTGATGGCTGGCCTTGTCGGCGAAGCGATGGACCTTGTCCTCGATCGACGGACAGTAGCGCGGCCCCACGCCTTCGATGACGCCGGAG
TACATTGGCGAGCGATCGAGATTGGCGAGGATGATCTCGTGCGTTCGCGCGTTGGTGTGGGCGATATGACAGCTGACCTGGCGCGGATGCATCCCGAGTT
CACCGAGGTAGGACATCACCGGGGTGGGGCTGTCGCCGGGCTGCTCCTCGAGCTGGCTGAAATCGACGCTCTTGGCGTCCAACCGCGGCGGTGTGCCGGT
TTTCAGCCGTGCCACGTTGAACGGCAGCGCTCGTAGCCGCTCGGCGAGGGCGTTGGACGGCGGGTCGCCGGCACGGCCCCCGTGGCTGCGGTCGAGGCCG
ATATGGATCACCCCGCCGAGGAAGGTTCCGGTACACAGCACCACGCTCTCGGTGTGAAAGCGGATACCGGTTTCAGTGACCACGCCGCGCACGCTGTCGC
CATCGACGATCAGGTCGCCGGCGGCCTGCTGGAACAGCGTCAGGTTGGGCTGCTGCTCGAGCATGCCGCGGATCGCAGCCTTGTAGCGCACCCGGTCGGC
CTGAGCGCGAGTGGCCCGCACCGCCGGCCCCTTGCGTGCGTTGAGCACCCGGAACTGGATCCCTCCCCGGTCGGTGGCCAGCGCCATGGCACCGCCCAGC
GCATCGATCTCCTTGACCAGATGACTCTTGCCGATCCCGCCGATGGCGGGATTGCAGGACATCTGGCCCAGGGTCTCGATGTTATGGGTCAGCAGCAGGG
TCTGACAGCCCATGCGAGCGGAGGCCAGGGCGGCTTCGGTTCCCGCATGGCCACCGCCGATGACGATGACGTCAAAGCGGTCGGGGTAATCCACGTTGCA

CCTCGAATGGCACCAGTGGTGCCGAAAAAATGATCAATGCTGGAAATTGGCTGCCCAGTATAAACCTCCTGTGGGTAACCGTCAGGGGTTTTCCACACCG
TCTTGGTCAGTCCTGCCTCACCTGACTAAGACAAGAATAAAAATAAGTAGAGTAAGAAAGAAGTTCTGTTGTTGTAGTGACTACTAGTGTGGACACTTTC
**TGTGGGTAA**GTGGACTTTACTAACAAATATCAATGAATTGCAGTGTTGAGGAATTGACGGACAAGAGGCGTATCGGCTGCGGACAGATGGGCGTGAGCCT
GTGGATGAAAAGCCGCTTTGTGCACAGCCACAATCGCCCACCGTTTATCCACCGAGCTTGGTGATGCTTGTCACCGCGACCGCCCACAGGCGGTGAAAAA
ATCACCAGCACCATGGTGACGGGGGGTGTAGCGCGATTCGATGGCGGTTGGATTTTTTTTGTCCACAGGCAAAAAAATGCCAAAAAAAGGCCAGTCGCTG

TGCGACTGGCCTTTATCAGCATCCCGCTTGACGCAGTGTCAGTGCTTGAGAACCCGCGCCAGGAAGCTTTGGGTGCGATCATGCTGAGGGCTATCGAATA
GGGTTGCGGGAGGCCCCTGTTCGACGATTTCACCGCCGTGGATGTAGACCACACGATCGGCGACTTCGCGGGCGAAGCCCATCTCGTGGGTGACGATCAT
CATCGTCATGCCCTCCCTGGCCAGCTCGCGCATGGCATCGAGCACTTCGCCGATCATTTCCGGATCCAGCGCCGATGTCGGCTCGTCGAACAGCATCAGC
CGCGGTTCCATGGCCAGCGCCCGGGCCAGTGCCACACGCTGCTGCTGGCCGCCGGAGAGCTGGCTGGGGTACTTGTCGGCCTGATCGCTGATGCCGACTC
GCTCGAGCAGACGCTCGGCGGTGCTTGCTGCTTGCTGCCGGCTGACGCCGCGAACCTTGAGCGGTGCCAGGGTGATGTTGTCGCGCACGCTGAGGTGCGG
AAACAGGTTGAACTGCTGGAAGACCATGCCGACTTCGGTGCGGATGGTCTGCAGCGCCTTGCTGCTCTTGCCGTGCGGGGTCAAGGCGTTGCCGTCGACC
TCGATATGACCGGCCTGAAATTCCTCGAGGCCATTGATGCAGCGGATCAGCGTCGACTTGCCGGAGCCGCTGGCACCGATGATCACCACCACTTCACCGG
GGGCGACGCTGAGTTCGATGTCCTTGAGCACGTGCAGGCTACCGAAGTGCTTGTTGACCTGCTGCATATGGACGATGGGCTGGTCGGCGGGGGTTTGGGT
TGTCATGGGTAGCTCCTTATTGTCCAACCAGGCCACGGCGCTCCAGCTGGCGCAGGATCAGCGATAGCGTCCAGGTAATCGCCAGGTACATCAGCGCGAC
CATGAAGTAGACCTCCAGAGCGGTGAAGGTGGTGGCGATGTAGATCTGTCCCTGGCGTACCAGCTCACCGACACCGATCACCGAGAATAGCGAGGTGTCC

>Halomonas qiaohouensis DSM 26770 [NZ_JARWAM010000001.1:c275463..275510], dnaA> ig dnaN>, 48 bp

WebSkew GCmin pos. 168987 (GCmin<>oriC1 30.2 kb; GCmin<>oriC2 3.9 kb; oriC2<>oriC1 26.3 kb)

>Halomonas pantelleriensis AAP [NZ_FNGH01000005.1:198912..199411], <rpmH oriC1 dnaA>, 501 bp

ACAATATCGCCACCTTGAGGGTCGATGCGTATGTCGTGTACATCGGTAACCACGGCAATCAGGTCGCGGCTACGGGTATCGCTCTCGGTAGCCTGTGGCA
GCTCGCTTTCTGCCGCCTCCTGACTGGCGCTGGGAACGCTCAGGCCTCCATCGTCGTCGCTAGGCTCTTGCTCGACCTGAGGCGCGCCGCCGTTGGTGAA
GGAGGGGGCGTCCGGCTCAGGCGTTGCCTGGCCGTAATCCTGATTCCACTGTACCACCAGCAGATATGCCAGCACGGCCAGCGGAACCAGCAGCAGTAGT
CGTTTTACGTCCATGAGGGTCCTGCCCGATGGGTAATGAACATTCTGACGGCAACATCGATATCAGCGGTGCCGATACACGCGAAAGCCTGCCGGAGGCA
GGCTCGAAGCGTCACCGCGAAGACGGCGAGGAAGTGTCCTGACTGACAGCGGCCGTCGACTGCCTACGCACATCCTTTTCCAGACGCCGCCACATGCCGT
ATAGCTGACGATGCAGGGTGTCGTTGTCCAGCTCGTGTACGCCTCGCCGGGAGAGTACCACGATGTCGACAGCGGGAAGCCGATGCTGCTGGAGACGGAT
GGATTCGCGAACCAGTCGTTTTAACCGGTTGCGATCCACGGCCCGGCGCACGTTCTTCTTGCTGAAGACCAAGCCCACGCGCGGGTGGCCCATTGCGTTG
GGGCTTGCCAGTGCCAGCAATCCTTTGCCGTGAATCTTGAACGCCGCGTGATCGAAGACGTGTCGGTATTCCCCGGCATTCAGCAGTCGCAGACGCCGGG
GGAAGCCCAGACAGGACACACGCAAGCCGAGATCAAGCGCTCAGACGCTTGCGGCCCTTCGCGCGACGCCGCGCCAGAACGGCGCGACCGTTCTTGGTGG
CCATGCGGGCACGGAAACCGTGCACGCGCTTGCGCTTGAGAACGCTGGGTTGAAAAGTGCGTTTCATAACTATCGATTCCCACGTGGTTAATTGGACGGA

ACGTCAATGGTGAGGCGCCCTGGCTCCGGCGGGGAACCCGGGGTAATTCGGCTCAAGACCGGGCATTTTAGTGAATTCGAGCCACGGGTGCAATTTTTCT
CGTGGCTAGGGCACGAAGCGTCAAGCGGACTTGCCATGTGGCATCGCGTGAATCCCGGCCATGGTTACACCATCTTGTTATTAAGACTGGGGACGAAATA
**TGTGGATAA**GAAGATTTTCTGTAGTTATTACATATAGATAGTCGGGAGTTGGCGGTGTGGGAAAAGTATTGACAACTTGTTAGGCGATGGCCAAAAGCAT
CACCGCCTGCTGCTGTGTGATTGCTTGGCGGATCGCTGTGCCATCTGGGTTGCGGCACGCTGGTGTGCTGAGTTTTCCACAGCGCGGCTTCGGTGCTGGT
GCGTGTGGATAACTTGCCTCTTGAGCTATACAATGGCCGGTCATTCATCAACGGGATGGTGAGGTCGCGTGTCGGTCGCTCTTTGGCAACAATGTTTGGA

TTTTTTGCAGGACGAGCTCAGCTCACAGCAGTTCAACACCTGGATTCGCCCTCTTCAGGCGGAGCAGGGAGAATCCAACGAACTCTGCCTGTTGGCCCCC
AACCGTTTCGTACGTGATTGGGTAAGCGACAAGTACATCAAGCGAATCAATGAACTGCTGCGCGAACTGGCGCCTACCAAGCCGCCCAAGGTGACGCTGG
TAGTGGGCAGTCGACGCGCCGTGCCTAGCCCGCAGCCACGAGAATTGGGCAATCCTGTCTCTGCCACTCTTCGTCAGCCCGCGGCACCGGCAGTGCATAC
GCCACCACGCGGCGATCTTGCCGACGAGCGTGAGATCGAGACACCCAGCGAGGCGTCATCGCGCAGCGGGTCATCGCGTGGCGGCGGTGAAAGGCAGGTG
CAGGTTGAAGGCAGTCTGAAGCACCAGAGTGGCCTCAACCCCAATTTTACCTTCGAGACGTTTGTCGAGGGTAAGTCGAACCAGTTGGCACGTGCTGCTT
CGCGCCAGGTCTCTGAGAACCCCGGCGGCGCCTATAATCCGCTATTCCTCTACGGTGGCGTTGGCCTGGGTAAGACTCACTTGATGCACGCGGTGGGCAA
TGCACTGGCCACCCGGCGCGACAATGCCAAGGTGGTCTATCTGCACTCCGAGCGTTTCGTGGCCGACATGGTCAAGGCGCTGCAGCTCAACGCGATCAAC
GACTTCAAGCGCTTCTATCGCAGCGTCGATGCTCTGCTGATCGATGATATCCAGTTCTTCGCCGGCAAGGAGCGCTCCCAGGAAGAGTTCTTCCACACCT
TCAATGCGCTGCTCGAGGGCGGCCAGCAGATGATCCTGACGTCTGACAGATATCCCAAAGAAATCAATGGTGTAGAAGAACGACTCAAGTCTCGCTTCGG
GTGGGGGCTGACCGTGGCCATCGAGCCACCGGAGCTCGAGACGCGGGTGGCGATCCTGATGAAGAAGGCCGACCAGGCCAAGGTCGATCTGCCCCACGAT

>Halomonas pantelleriensis AAP [NZ_FNGH01000005.1:172660..173159], <mnmG oriC2 <COG1126 (glnQ), 547 bp

CATCCCCTTGATCGAGCGTACTACCTGCAGCTGCACGTCAAACGGTAGCGAGGTGGAAATCCCATTGGGATAAAGTTCGTGGGTATCCAGGCCTTCCGGC
TCGACGAACACCTGGTGACTGGCCTTGTCGGCAAAGCGATGAACCTTGTCCTCGATTGACGGGCAATAGCGCGGCCCCACGCCTTCGATAAGCCCGGAAT
ACATCGGTGAACGATCGAGATTGGCGAAGATGATCTCGTGAGTACGCTCGTTGGTATGGGCGATATGGCAGCTGACCTGGCGCGGATGCATGCTGCGATC
GCCAAGATAGGACATCACCGGCGTTGGGCTATCGCCGGGCTGTTCCTCGAGTTGGCTGAAGTCGACGCTCTTGGCATTTAGCCGCGGCGGCGTGCCGGTC
TTCAGCCGTGCGACGTTGAACGGCAGCGCCCGCAGCCGCTCAGCCAATGCATTCGAGGGTGGGTCACCGGCGCGTCCTCCACGACTCTGGTCTAGGCCAA
TATGAATGACCCCACCGAGGAAGGTTCCGGTGCACAGCACCACGCTCTCGGCATGGAAGCGAATGCCGGTCTCGGTGACCACGCCGGTTACTCGATCGTC
GCTGACCATCAGGTCGCCGGCAGCCTGCTGAAATAGCGTCAGGTTGGGCTGCTGCTCCAGCATGCCGCGGATGGCAGCCTTGTAGCGTACCCGATCGGCT
TGGGCACGAGTGGCACGCACCGCCGGTCCCTTGCGCGCGTTGAGCACTCGGAACTGGATTCCTCCCCGGTCAGTGGCCAGTCCCATGCCACCACCCAGCG
CATCGATCTCCTTGACCAGATGACTCTTGCCGATCCCGCCGATAGCGGGATTGCAGGACATCTGACCGAGTGTCTCGATGTTGTGAGTTAGCAGTAGGGT
CTGACAGCCCATGCGAGCGGAGGCCAGGGCGGCTTCGGTTCCCGCATGGCCACCGCCGATGACGATGACGTCAAAGCGGTCGGGGTAATCCACGTTGCAC

CTCGAATGGCACCGTTGGTGCCTAAAAAATGATCAATGCCAAAAATTGGGCGCCCAGTATAAACCTCCTGTGGGTAACCGTCAGGGGTTTTCCACACTGC
CTTGCCCTGCCGGGTGACATCACGCGTCAAACAAGAATAAAAATAAGTAGAGTAAAAAAGAAGTTCTGTTGTTGTTGTGACTACTGGTGTGGATATTTTC
**TGTGGGTAA**GCCACTTTTCCATTTCCTAATCAGTCTCTTGGAGTGTTGACCGTCACGCGGAAAAACAGCGTATCAGCTGCGGACAGCCGGTCATGAGCCT
GTGGATGAAAGCGCGACTTGTGCACAGCCCGAGGCGTGCACCGCTTATCCACCGACCTCGTTGGCAGTTGTTACCGCGCCCGCCCACATGCTGGTGAAAA
TTCCACCAACACCATGGTGGTGGGGGCTGGCGAGCATTCTCGCGGCAAATGAATTTTTTTGTCCACAGGCCAAAAAAGCGCCAAAAAAAGGCCAGTCGT

TGTCCGACTGGCCTTTTTGAGAGGTGCGGGGAGCAAAAAATCAGTGCTTGAGCACCCGCGATAGGAAGCTTTGCGTACGAGCATGCTGTGGGTTGTCGAA
CACGGCTGCAGGGGGGCCTTGCTCGACGATCTGACCGCTGTGGATGTAGATCACGCGATCGGCCACCTCCCGAGCAAAGCCCATCTCGTGGGTGACGATC
ATCATCGTCATGCCTTCTCGGGCCAGCTCACGCATGGCGTCGAGCACTTCGCCGATCATCTCCGGGTCCAGCGCTGAGGTCGGCTCGTCGAACAGCATAA
GACGTGGCTCCATGGCCAGCGCCCTGGCCAATGCCACGCGCTGCTGTTGGCCACCTGAGAGCTGGCTTGGGTACTTGTCGGCCTGGTCGCTGATGCCCAC
CCGTTCCAGGAGGCGTTGGGCAGTACTCGATGCCTGTTGGCGACTGACGCCGCGGACCTTGATGGGGGCCAGGGTGACGTTGTCGCGCACGCTGAGATGG
GGAAACAGGTTGAACTGCTGGAAGACCATACCGACTTCGGTGCGGATGGTCTGCAGTGCCTTGCTGCTCTTGCCGTAGGGGGTCAGCGGGTTGCCGTCGA
CCTCGATGTGACCGGACTGGAACTCTTCGAGGCCGTTGATGCAGCGAATCAGCGTCGATTTTCCGGAGCCGCTGGCGCCGATAATCACGACCACTTCGCC
AGGCTCGACGCTGAGTTCGATATCCTTGAGCACGTGCAGGCTACCAAAATGCTTGTTGACCTGTTGCATGTGCACGATGGGTTGGGCCGCGGTGGGTGTG
TCAGTTGTCATGCGTGGCTCCTTTATTGTCCGACCAGGCCGCGGCGCTCCAGTTGGCGCAGGATCAGCGATAACGTCCAGGTGATCGCCAGGTACATCAG
CGCGACCATGAAGTAGACCTCCAGTGCGGTGAAGGTGGTGGCGATGTAGATCTGGCCCTGGCGCACCAGCTCACCCACGCCGATCACCGAGAACAGCGAG

>Halomonas pantelleriensis AAP [NZ_FNGH01000005.1:200856..200902], dnaA> ig dnaN>, 47 bp

WebSkew GCmin pos. 68448 (GCmin<>oriC1 35.9 kb; GCmin<>oriC2 9.3 kb; oriC2<>oriC1 26.6 kb)

>Halomonas taeanensis BH539 [NZ_FNCI01000005.1:104087..104586], <rpmH oriC1 dnaA>, 568 bp

```
GGCGACCAGGGCACGACTGTTGTCGTTGGAAGCCGCTGGATCACGCATGCTGCCGGTGAGATCCGATGACCCTTGCGGGTTCGAAGAAGTCGTGGGTACC
GAAAGGCCAGATGTCTCCTCGGAGGTATTCTGACCAGTGGGCTGGCCACTGGTGGCAACCGAGGGCGCCTGAGTCTCAGCGGTGACCTGGCCGTAGTCCT
GGTTCCACTGTACGACCAGCAGATAGGCCAGTACGGCGAGAGGAATCAGCAGTAGGAGTCGTTTTACGTCCATGAGGGTCCTGCCCGATGGAAAATGAAC
ATTCTGCAGTGCGGATGTTGCCCCGTTAATACACACGAAAGCCTGCCAGGGGCAGGCTCCGAGGATCGCCGGTGTGTCCGATCAGGACAGCGAGGTCGAC
TGCTTGCGAACGTCTTTTTCCAGACGCCGCCACATGCCATGAAGCTGGCGATGCAGAGTGTCATTATCCAGCTCATGAACGCCGCGCCGGGCCAGGACGA
TGATGTCCACGGCAGGTAGGCGATGCTGCTGGTGACGAATGGATTCACGCACTAGGCGCTTAAGACGGTTGCGATCGACGGCTCGTCGCACGGTCTTCTT
GCTGAATACCATGCCAACTCGAGGGTGGCCAAGCGGGTTATGACAGGCGAGAGCCAGCATCCCCTTACCGTGAACCTTGAGTGATGCATGATCGAAGACG
TGACGGTATTCCCCGGCGTTCAGCAGGCGCAGCGACCGGGGAAAGTCCTGATGGGGCAAGTGCAATCCGAGATCACGCGCTCAGACGCTTGCGACCCTTG
GCACGACGACGTGCGAGGACGGCGCGACCATTCTTGGTGGCCATGCGTGCACGGAAGCCATGGTTACGCTTGCGCTTGAGGACGCTGGGCTGAAAGGTGC
GTTTCATAACTATTGATTCCCACGTGGTTAATTGGACATGACGTCTAATTGGCGAAGCCCTGGCTCCGACGGGGAACCCGGGTAGATTCGGCTCAAGACC
```

```
GGGAATTCTAGTGAATTCGCCGTCTCGGTGCAATTTTTCCGTCGACTTCTGGCGGTCTTTTCTACCTGATCGACGCTGTCGCGGTATCGAGCGGCCTGCT
ACCCGATAGGCACAGCCAGGTGCTCAAAACGCCGTAGCAGCTTTAATAACTATTATTAGATCTTCTTACTGATGTTGTTATTAATAGTGGGGACAATATA
TGTGGATAAGTCAAAATATGTCATCTATTACAGTTTTTATACTGAAGACAAGCCTGTGTGGAAACGGTCGATGCAGAGAGGAGAACCGGGGGACAGGCT
GATGACAAGATCAACGGCTGTCCACAACCCAAGGTTTCCCTCAGCGTCATGCAATGGTTCTTCAGGGAGAAGTTGAGATGTTGTCCACAGGGAGAAATCG
ATGCCGCGTCATGTGGATAACCCAAGCCTCGAACGCTACAATGCGGATCTTTCGTCAACCGGTTGGTGAGGTCGCGTGTCTGTCGCTCTTTGGCAACAAT
```

```
GTCTGGATTACCTGCAGGACGAACTGAGCTCCCAGCAGTTCAATACCTGGATCCGTCCTTTGCAGGCGGAGGAAGGGGAGTCCAACGAGCTTTGCCTGTT
GGCCCCCAACCGCTTCGTGCGTGACTGGGTCAGTGACAAGTACGCCAAGCGCATCAGTGAATTGATGCGTGAACTGTCCCCGGCGAAGCCTCCTAAGGTG
TCGTTGACCGTGGGCAGTCGACGTGCGGCGCCGGCGCCGCAGCCTCGTGAGCTTGGCAACCCGGTTTCCGCCGGGTCGCGCCCGGAACAAGCCGCACCGG
CGGTGCATACGCCGCCGCGCGGGGATTACGCTGATGAGCGTGAGATCGATCAGATGCGTGACGAGGGCGCCCGGCGGCCTGCTGGCAGTACTCGTCAGGT
TCAAGTCGAGGGTAGCCTCAAGCACCAGAGCGGCCTCAACCCCAACTTCACCTTCGAGACCTTCGTTGAGGGTAAGTCGAACCAGTTGGCCCGTGCCGCC
TCCCGTCAGGTCTCCGAGAATCCTGGTGGCGCCTACAATCCGCTGTTTCTGTATGGCGGCGTGGGTCTGGGTAAGACGCACCTGATGCACGCGGTAGGCA
ACCAGTTGGCCGGCCGCAGCGAGAACGCCAAGGTGGTCTATCTGCATTCCGAGCGTTTCGTCGCCGATATGGTCAAGGCGCTGCAGCTCAATGCCATCAA
CGACTTCAAGCGCTTCTACCGCAGCGTGGATGCCCTGCTGATCGACGATATCCAGTTTTTCGCCGGCAAGGAGCGTTCCCAGGAAGAGTTCTTCCATACC
TTCAATGCGTTGCTTGAAGGCGGCCAGCAGATGATCCTGACGTCTGACCGTTATCCCAAGGAAATCAGTGGGGTAGAGGAACGACTGAAGTCGCGGTTTG
GTTGGGGACTGACGGTTGCCATCGAGCCGCCCGAGCTCGAGACCCGGGTGGCGATCCTGATGAAGAAGGCCGATCAGGCCAAGGTCGACCTGCCGCATGA
```

>Halomonas taeanensis BH539 [NZ_FNCI01000005.1:77503..78002], <mnmG oriC2 <COG1126 (glnQ), 625 bp

```
TCGAGCGCACTACCTCGAGCTGCACGTCGAAGGGCAGCGAGGTGGAAATCCCGTTGGGATAGAGCTCGTGGGTATCCAGGCCCTCAGGCTCCAGGAATAT
CTGGTGGCTCGCCTTGTCGGCAAAGCGATGCACCTTGTCCTCGATGGAGGGGCAATAACGCGGGCCAACGCCCTCGATGGTGCCGGAATACATCGGCGAT
CGGTCGAGGTTAGCGAAGATGATCTCGTGAGTACGCTCATTGGTATGCGCGATGTGACAACTGACCTGGTCGGGATGCATCTCCCGGTTACCCAGATACG
ACATCACTGGCACCGGGCTGTCGCCGGGCTGCTCTTCGAGATGGGTGAAATCGACGGTGCGTGCATCGATGCGCGGCGGTGTGCCGGTCTTCAGTCGGGC
GACGTTGAAGGGCAGCGCCCGCAGGCGTTCCGCCAGGGCATTCGATGGCGCATCACCGGCGCGTCCGCCGCGGCTTTTGTCGAGGCCGATATGGATCACG
CCACCGAGGAAGGTACCGGTGCACAGCACCACGCTCTCGCCCAGAAAGCGAATGCCGGTTTCGGTGACGGCGCCACGCACGGTGTCGCCATCGACGATCA
GGTCGCCGGCTGCCTGCTGGAAAATCGTCAGATTGGGCTGGCTTTCCAGCATGCCGCGAATCGCCGCTTTGTAGCGGACTCGGTCGGCCTGGGCGCGAGT
GGCACGCACGGCAGGCCCCTTGCGGGCATTGAGCCGGCGAAATTGGATGCCGCCGACGTCGGTGGCCAGCGCCATGGCGCCACCTAGCGCATCGATCTCC
TTGACCAGATGGCTCTTGCCGATCCCGCCGATGGCCGGATTGCAGGACATCTGGCCCAGGGTCTCGATGTTGTGAGTCAGCAGCAGGGTCTTGGCACCCA
TGCGAGCAGAGGCCAGGGCGGCTTCGGTTCCCGCATGGCCTCCGCCGATGACGATGACGTCAAAACGGTCGGGGTAATCCACGTTGCACCTCGAATGGCG
```

```
CCGATGGCGCCGTGTCCATAACCGTCGTCAACCGACGGGGTGAGAAATCAGCGGCACAGTATAAACCTCCTGTGGGTAACCGTCAGGGGTTTTCCACACC
CAGATCCGAAGCCAGGATTCAAAGCGGGGTTCTATATAAAATACAGATTAAATAAGAAAGAAGTTCTGTTGTTGTAGTAACTACTGGTGTGGACACTTTC
TGTGGAAAACCAGAATATCTCTATTTTTATCAATAGCCTGAAATGTTGACCGATCGTGTGAAAAACGGCGGAGGGCGTGCGTACAGCTTGTCGTGAGCCT
GTGGATGAAAGGGCGCCTTGTCCACAAACCTAAATGACCCAAGGTTATCCACCGAGCCTAGCCCGGGTTGTTAGCCTGGTGGTCAACATGGCAGGCCGAT
ATGCTCGAAGACGCTGATGCCGGTGTTCGTGGAAGGATTTGGGGCCTTTATGAAAAAAACTTGTCCACAGCCAAAAAAAGGGCCTGTCGAGAAGTCGACA
```

```
GGCCCTTTGAAGTGCCAAAATCAGGTTTTTTGCAGCGCTAGCGGAGCGATTTATCGCCTATGAGCCGCGCGCTGCTCCTCTTGCGTTAGTTCTTGGGTCA
GTGCTTGAGAACGCGCCCAAGGAAGCTCTGGGTGCGTTCATTCTGTGGGTTGTCGAACACCTCTTCCGGCTTGCCCTGCTCGACGATCTGCCCCTTGTGG
ATATAGATCACGCGATCAGCCACTTCCCGGGCAAAGCCCATCTCGTGGGTCACGATCATCATCGTCATGCCGGCCTTGGCGAGCTCGCGCATGGCATCCA
GCACCTCGCCTATCATCTCCGGGTCCAGGGCCGAGGTGGGCTCGTCGAAGAGCATCAGCCGCGGTTCCATGGCCAGGGCACGGGCCAGCGCCACGCGCTG
CTGTTGGCCGCCTGAGAGCTGACCTGGCAGCTTATCGGCCTGGTCGGCGATGTTGACCTGCTCGAGCAGCCTGTCGGCGTTCTCGATGGCATCGGACTCG
CTGGTGCCGCGGACCTTGATCGGTGCCAGGGTGACGTTGTCGCGCACGCTAAGATGCGGGAAGAGATTGAACTGCTGGAAGACCATGCCGACTTCCGTGC
GGATGGTCTGCAGCGACTTGCTGCTCTTGCCATTTGGCAGCAGCTGGTTGCCGTCGACTTCGATATGGCCGTGCTGGAATTCCTCGAGGCCGTTGATGCA
GCGGATCAGGGTGGACTTGCCAGAGCCGCTGGCGCCGATGATCACTACTACTTCGCCGTGCGTGACTGAAAGATCGATGTCCTGCAGCACGTGCAGGCTG
CCGAAGAACTTGTTGACCTTCTCCATGCGCACGATCAGTTCGGCGCCGGGTTTTTCTTGATATCTATCATGATCGTTTCTCTCGTTGTATCGATGTGCG
TTTACCGGCGTTATTGACCTACCAGTCCCTTGCGCTCGAGCAGGCGCAGGGCCAGCGAGAGGCTCAGGGTTATCGCGAGATACATCAGGGCGACCATGAG
```

>Halomonas taeanensis BH539 [NZ_FNCI01000005.1:106032..106092], dnaA> ig dnaN>, 61 bp

WebSkew GCmin pos. 82368 (GCmin<>oriC1 28.4 kb; GCmin<>oriC2 1.7 kb; oriC2<>oriC1 26.6 kb)

>Halomonas niordiana ATF 5.4 [NZ_SDSD01000004.1:110480..110979], <rpmH oriC1 dnaA>, 557 bp

ACGTCGGTGGTGACAGCGACCAGCGCGCGGCTGTTGCTTTGCGCGCCCGGGGTGCTCATGTCACCGCCGAGATCCGATGCTGCATTCTGACCGGATGTGG
TGGGTGCCGATAGGCCCGACGTTTCATCACGGGCAGCTTGCTCGGTGGGCTGGCCACTGTTGGCGACCGTAGGCGCCTGGGTTTCAGTGGTGATCTGGCC
ATAGTCTTGGTTCCACTGTACGACCAGCAGATAGGCCAGGACGGCGAGGGGGATCAGCAATAGGAGTCGTTTTACGTCCATGAGGGTCCTGCCCGATGGA
AAATGAACATTCTGCAGTGCAGCGATTGTTGCCCCGTGAATACACACAAAAGCCTGCCAGGGGCAGGCTTTGAGGATCGCCGGTGCGCCCGATCAGGGCG
ACGAGGTCGACTGCTTGCGAACGTCTTTTTCCAGACGCCGCCACATGCCATGAAGCTGGCGATGCAGAGTGTCATTATCCAGCTCATGAACGCCACGCCG
GGCCAGGACAATGATATCCACGGCAGGTAGGCGATGCTGCTGGTGACGAATGGATTCACGCACGAGGCGCTTAAGACGGTTGCGGTCGACGGCTCGACGC
ACGGTCTTCTTGCTGAACACCATGCCGACTCTTGGGTGACCCAGCGGGTTGGTGCAGGCCAGAGCCAGCATCCCCTTGCCGTGAACCTTGAGTGATGCAT
GGTCGAAGACGTGTCGATAATCCCCGGCGTTCAGCAGGCGCAGCGACCGGGGAAAATCCTGATGGGGCAAGTGCAATCTGAGATCACGCGCTCAGACGCT
TGCGACCCTTGGCACGGCGGCGTGCGATAACGGCGCGACCATTCTTGGTGGCCATGCGTGCACGGAAGCCATGGTTACGCTTGCGCTTGAGAACGCTCGG
CTGAAAAGTGCGTTTCATAACTATTGATTCCCACGTGGTTAATTGGACATGACGTCAAATTGGCGAAGCCCTGGCTCCGACGGGGAACCCGGGTAGATTC

GGCTCAAGACCGGGAATTCTAGTGAATTCGCCGCCGCGGTGCAATTTTTCCGGCGATCTCGAGCGGTTTTTTCGCGTCGCCATCGAAGCTGAGGCCAAGC
ATGGTGCGCCAGTCCTGCCTTAGTCATCAACCGCGATCTTCTTTAATAACTATTATTAGGTTCTCTTACTGATGTTGTTATTAATGGTGGGGACAATATT
**TGTGGATAA**GTCATTATAAACCATCTATTACAACTTATTATCGTGACGACAAGCTTGTGGGAAAGCGGTCAACCTCATGCGTGCATTCGGGGGATAAGCT
GCTGACAGAATTACCACCTGTCCACAGCACGATGGTACTCCCCGTTTTGCACAGTGTTTCTTAAGGGAGCAACCGCAAGGTTGTCCACAGGGCGAAATCG
ATACGCCGTCATGTGGATAACCTAGTCCTCGGACGATACAATGCGTGTCTTTCGTCAACCGGTTGGTGAGGTCGCGTGTCTGTCGCTCTTTGGCAACAAT

GTCTGGATTACCTGCAGGACGAACTGAGCTCCCAGCAGTTCAACACCTGGATCCGTCCCTTGCAGGCGGAGGATGGAGAGTCCAACGAGCTTTGCCTGTT
GGCACCCAATCGCTTCGTGCGTGACTGGGTCAGTGACAAATACGCCAAGCGTATTAATGAACTCATGCGTGAGTTGTCACCGGCGAAGCCACCTAAGGTG
TCGCTGACGGTCGGTAGCCGACGTGCGGCGCCGGCCCCCCAGCCGCGCGACCTGGGTACGCCAGTTTCCGCTGGTTCGCGTCAGGAGCCTGCCTCGCCCG
CGGTGCATACACCGCCGCGGGGTGATTTCGCCGACGAGCGTGAGATCGACCGCATGCGAGACGAGGGGCCAAAGCGGCCCGCCGGTAATTCGCGTCAGGT
ACAGGTCGAGGGTGGCCTCAAGCATCAGACTGGCTTGAATCCCAACTTCACCTTCGAGACCTTTGTCGAGGGTAAATCGAACCAGCTGGCCAGAGCCGCT
TCTCGCCAGGTATCGGAAAACCCCGGTGGCGCCTATAACCCACTGTTCCTGTACGGCGGCGTAGGTCTGGGTAAGACGCACTTGATGCACGCGGTGGGTA
ACCAGCTGGCCGGGCGGAGCGAGAATGCCAAGGTGGTCTATCTGCACTCCGAGCGCTTCGTGGCCGACATGGTCAAGGCACTGCAGCTTAATGCCATCAA
TGATTTCAAGCGCTTCTATCGCAGCGTCGATGCGCTGCTGATCGATGATATTCAGTTCTTCGCGGGTAAGGAGCGCTCTCAGGAAGAGTTCTTCCATACC
TTCAATGCGCTGCTCGAAGGCGGTCAGCAGATGATCCTGACGTCGGATCGTTATCCCAAGGAAATCAGTGGGGTAGAGGAGCGGCTGAAGTCACGTTTTG
GCTGGGGACTGACGGTTGCCATCGAACCGCCGGAGCTTGAGACGCGGGTCGCGATCCTGATGAAGAAGGCCGACCAGGCCAAGGTCGACCTGCCTCATGA

>Halomonas niordiana ATF 5.4 [NZ_SDSD01000004.1:83832..84331], <mnmG oriC2 <COG1126 (glnQ), 617 bp

GAGCGCACTACCTCAAGCTGCACGTCGAAAGGCAACGAGGTAGAAATCCCGTTGGGGTAGAGCTCGTGGGTATCTAGCCCTTCGGGTTCGACAAAGATCT
GGTGGCTCGCCTTGTCGGCGAAGCGATGCACCTTGTCTTCGATGGACGGGCAATAGCGCGGACCGACGCTATCGATGGAGCCTGAATACATCGGCGACCG
ATCAAGATTGGCGAAGATAATCTCGTGAGTGCGCTCATTGGTATGCGCGATATGGCAGCTGACTTGCTCAGGATGCATATCACGGCTGCCAAGATACGAC
ATCACCGGCGTCGGGCTGTCACCGGGCTGCTCCTCGAGGCCACTGAAATCGACACTGCGAGCATCGATGCGTGGCGGTGTACCGGTCTTCAATCTTGCCA
CATTGAACGGCAGTGCCCGCAGACGTTTTGCCAGGGCGTTGGACGGTGCGTCACCGGCACGGCCACCCAGGCTCTGGTCGAGACCGATATGAATCACGCC
GCCCAGGAAGGTACCGGTGCATAGCACCACGCTTTCGCCGAGGAAGCGGATACCGGTTTCGGTTACCGCACCACGTACGGTGTCGCCATCGACGATCAAG
TCGCCTGCCGACTGCTGGAAAATCGTCAGGTTGGGCTGATTCTCCAGCATGCCGCGAATCGCCGCCTTGTAGCGGACTCGGTCTGCCTGAGCGCGAGTGG
CGCGAACTGCCGGTCCCTTGCGGGCATTGAGCCGACGGAACTGGATGCCTCCCAGGTCGGTGGCCAGCGCCATGGCGCCGCCGAGCGCATCGATTTCCTT
GACCAGATGACTCTTGCCGATTCCGCCGATGGCGGGATTGCAGGACATCTGGCCGAGCGTCTCGATGTTGTGGGTCAGCAGCAGGGTCTTGGCACCCATG
CGAGCAGAGGCCAGGGCGGCTTCGGTTCCCGCATGGCCTCCGCCGATGACGATGACGTCAAAACGGTCGGGGTAATCCACGTTGCACCTCGAATGGCGCC

GATGGCGCCGTGTCCATAGCCGTCGTCAAACGACGGGGTGAGAAATCAGCCGCAGAGTATAAACCTCCTGTGGGTAACCGTCAGGGGTTTTCCACACCCG
GATTTGTATTCGACCCTCTGTGAACCGGACTTAATATAAAATACAAATTAAATAAAAGAGAAGTTCTGTTGTTGTTGCTACTACTGGTGTGGACACTTTC
**TGTGGAAAA**CCTCGCTATTTGTATATTTATCAATACGCTGTAATGTTGACCAATCGTGTGAAAAACGGCGGAGGAGATGCGTACAGCATGTCATGAGCCT
GTGGATGAAAGAGGCCTTTGTCCACAGACTCAAATAGCCCAAGGTTATCCACCGAGCCCAGGCCGGGTTGTTACCGTGGCGATCAACATCCGAAAGCAAC
CGGCGTGAAGTCGCTGATGCTGCCGGTCATGGGACGATTCGGCGCCTTTCTGAAAAAAACTTGTCCACAGCCAAAAAAAGGGCCTGTCGAGAAGTCGACA

GGCCCTTTGAGGTGCCATTATCCGCAGACGTGATGGCCTGGCGGAGCCATATAGAAACGATTTAGGTCACGATCACCGACCATTCACCGACGCTTCATCA
GTGTTTGAGGACTCGTCCCAGGAAGCTCTTGGTGCGTTCATTCTGAGGGTTGTCGAACACGGCTTCCGGCTTGCCCTGTTCGACGATCTGCCCCTTATGA
ATGTAGATCACGCGATCCGCCACCTCTCGAGCAAAGCCCATCTCATGCGTGACTATCATCATCGTCATGCCTGCATCGGCCAGTTCACGCATGGCATCCA
GCACTTCACCGATCATTTCCGGGTCAAGGGCCGATGTGGGCTCGTCGAAGAGCATTAGCCGGGGTTCCATGGCCAGCGCCCTGGCTAGCGCCACACGCTG
CTGTTGGCCGCCTGAAAGCTGGCCCGGAAGCTTGTCGGCCTGGTCGGCGATATTGACCTGCTCCAGCAGTCGGTTGGCGCTGTCGCTCGCCTGCTTTTCA
CTGAGGCCACGGACCTTGATGGGGGCCAGGGTGACGTTGTCGCGAACGCTGAGATGGGGGAATAGGTTGAACTGCTGGAAGACCATGCCGACCTCGGTGC
GAATGGTCTGCAGGGACTTGCTGCTCTTGCCATTTGGCAGCAGTTCATTGCCATCGACATCGATATGGCCGTGCTGGAATTCCTCTAGTCCATTGATACA
GCGGATTAGTGTCGACTTGCCTGAGCCACTGGCCCCGATGACCACTACGACCTCGCCGTGGCTGACCGAAAGCTCAATGTCTTGCAGCACGTGAAGGCTG
CCGAAGAACTTGTTGACCTTTTCCATACGCACGATGGGCTCGGCGCCGGCCTCTGTCTTGATATCTGTCATGATCGGTTCCCTCGTTGTACCGATGTTCG
TTGTGCCGGCGTTATTGACCTACCAGCCCCTTGCGCTCAAGCAGGCGCAGGGCCAGCGAGAGACTGAGGGTGATGGCGAGATACAGCATGGCGACCATCA

>Halomonas niordiana ATF 5.4 [NZ_SDSD01000004.1:112425..112485], dnaA> ig dnaN>, 61 bp

< page 1 | #112 | Halomonas anticariensis FP35 = DSM 16096

WebSkew GCmin pos. 175582 (GCmin<>oriC1 28.9 kb; GCmin<>oriC2 1.1 kb; oriC2<>oriC1 27.8 kb)

>Halomonas anticariensis FP35 = DSM 16096 [NZ_KE332393.1:146437..146936], <rpmH oriC1 dnaA>, 547 bp

CGCCTTGAGGATCGATTCGCACATTGAGCACGTCAGTGACTACTGCGATCAGGTCGCGACTGTCGGTAGGGGAGGGAGTCGCTCCTGGTAGGTCGCTGTC
GGAAATCGGTTCCGCGTCACTGGGGGTTGCGGGAGCCACTAAACCTTCCTCCCCTGGCGCTGTCTCGCCGTTACGAGGGGCATCATCGGTAGTGATGGCG
GGAGCTTCCGGTTCAGGTGTGACCTGACCGTAGTCCTGATTCCATTGCACGACCAGCAAATAGGCCAGTACGGCAAGTGGAATCAATAGCAGAAGTCGTT
TTACGTCCATGAGGGGCCTGCCCGGTGGGTAATGAACAGTCTGACGGCATAGTGCATGAATAAAATGACACACGAAAGCCTGCCGAGGCAGGCCGCCGGT
GTCCCGTCAGGTGGTATTGGCCGATTGCTTGCGAACGTCCTTTTCCAGACGCCGCCACATGCCGTAGAGCTGGCGATGCAGGGTGTCACTGTCCAGTTCC
CGTACGCCACGCCGGGATAGGACAACGATGTCCACGGCGGGTAGGCGGTGTTGCTGGAGGCGAAAGGACTCGCGAACGAGACGCTTCAAGTGATTGCGAT
CGACTGCGCGTCGCACGTTCTTCTTGCTGAATACCAAACCGACCCGTGGGTGACCCAGATCGTTGTTGCAGGCCAATGCCAGCAACCCCTTGCCGTGAAC
CTTGAAGGCGGCCTGGTCGAAAACGCGTCGGTAGTCCCCGGCGGTCAGCAGTCGCAGCTGCCGGGGAAAGCCCTGACGGGACACACGCAATCCGAGATCA
GGCGCTCAGACGCTTGCGACCCTTGGCGCGGCGGCGGGCCAGGACGGCGCGGCCGTTCTTGGTGGCCATGCGGGCACGGAAGCCATGCACACGCTTGCGC
TTGAGAACGCTGGGTTGAAAGGTGCGTTTCATAACAATTGATTCCCACGTGGTTAATTGGACTTTACGTCAACTTAGAGGCGCCCTGACTTCGGAGGAGG

CCAGGGGGGATTCGGCTCAAGACCGGGAATTCTAGTGAATTCGCCGTCAAGGTGCAATTTTTCCCGCTACATTCGCACCACTGGATGAAGTGTTATCCCT
TGGCCATGATTCATGAGGAACATGGCAGCTGATTTTGAACGATAACTACTATCATTATTGGTTTTTAGATTTCGTTGTTATTAATACTGGGGATAGTTTT
**TGTGGATAA**CCATGATTGATGCTTATATTACAATGATGTATAGAGGATACAACCATGTGAGGAAAAGCAGGACAGCCGTTTGGTTCATTGGGGATAGCCT
GGGGGAAGATCGATGAGTCTTCCACAATGTTGTCGTTGTGCCCAGTTCGAGCAGAGGTTGTTCAAGCGTAGGTAGGCAAGTTTTCCACAACTACACTTCC
TTGGGATCGTCATGTGGATAACCTATGGCTGGAGCGATACAATGGCGGGTCATTTTCACAATCGGGATGGTGAGGTCGAGTGTCGGTCGCTCTTTGGCAA

CAATGTCTGGATTACCTGCAGGACGAACTGAATTCTCAGCAGTTCAATACCTGGATTCGCCCTCTTCAGGCGGAGGAGGGTGAATCCAATGAGCTATGCC
TGCTGGCGCCCAATCGTTTCGTCAGAGACTGGGTCAATGACAAGTATGTGAAGCGTATCAACGAGTTGCTACGCGAGCTGTCCCCTGTCAAGCCGCCCAA
GATATCGTTGACCGTGGGCAGCAGGCGTGCAGCACCTTCGCCTCAGCCACGTGAACTGGGTAATCCCGTTTCCGCCAGTCCCCGTCAGCCTGCATCACCT
GCCGTGCATACGTCGCCACGCGGAAATTACGCTGATGAGCATGAGATCGATCAGGCTCGTGAGGAGGGGAGCCGTACTGGAAGCAGCCGCACCTCAGGAA
GCAGTGAGCGTCAGGTTCAAGTCGAAGGGAGCCTCAAGCATCAGAGTGGCCTTAACCCCAATTTCACCTTCGAAACCTTCGTTGAAGGTAAGTCCAACCA
ATTGGCCAGAGCTGCTTCACGCCAAGTGTCGGAAAATCCCGGTGGCGCTTATAACCCTTTGTTTCTGTATGGCGGTGTCGGTCTCGGCAAGACACACCTG
ATGCATGCCGTGGGCAATCAGCTAGCCAATCGTCGCGAGAATGCCAAGGTGGTCTATCTCCATTCCGAGCGTTTCGTTGCCGATATGGTCAAGGCACTAC
AACTGAATGCCATCAATGACTTCAAACGCTTCTATCGTAGTGTCGATGCCTTGCTGATCGACGATATTCAGTTCTTTGCTGGCAAAGAGCGTTCTCAGGA
AGAATTCTTCCACACCTTCAACGCCTTGCTTGAAGGTGGCCAGCAAATGATCTTGACATCGGACAGATACCCCAAGGAAATCAATGGGGTAGAGGAGCGT
CTGAAGTCCCGTTTCGGATGGGGGCTGACGGTAGCCATTGAGCCTCCAGAGCTGGAAACGCGGGTGGCGATCCTGATGAAAAAGGCTGATCAGGCAAAGG

>Halomonas anticariensis FP35 = DSM 16096 [NZ_KE332393.1:c174290..174789], <mnmG oriC2 <COG1126 (glnQ), 549 bp

CAACCCCTTGATCGAGCGCACTACCTCGAGCTGAACGTCGAAAGGCAACGAGGTCGAGATGCCGTTGGGATAGAGCTCATGGGTATCCAATCCCTCGGGC
TCGATGAATATCTGATGACTCGACTTGTCCGCGAAGCGATGCACCTTGTCCTCGATAGACGGACAGTAGCGAGGACCGATAGTGTCGATCACCCCCGAGT
ACATCGGTGAACGATCGAGATTGGCGAAGATGATCTCGTGGGTGCGTTCGTTAGTATGGGCGATATGGCAGCTCACCTGGCGGGGATGCATTCCTCGTGA
ACCAAGATACGACATCACCGGGGTGGGAGTATCACCGGGTTGCTCCTCGAGCACCGAAAAATCGATGCTCTTGGCATCCAGGCGCGGTGGTGTGCCGGTC
TTGAGCCGCGCTACGTTGAACGGCAGCGCGCGCAAGCGTTGAGCCAGGGCGTTGGAAGGGGGATCGCCAGCCCGGCCACCACGGCTTTGATCAAGACCGA
TGTGGATCACGCCGCCAAGGAATGTCCCGGTGCATAACACTACCGTTTCACTATGGAATCGAATGCCGGTCTCGGTGACGACACCGCGTACGGTGTCACC
ATCGACGATCAAGTCACCGGCAGCCTGCTGAAAGACCGTCAGGTTGATCTGGTTTTCCAGCATACCGCGAATCGCCGCTTTGTAACGCACGCGATCGGCC
TGTGCGCGAGTCGCGCGCACAGCAGGGCCTTTGCGGGCATTGAGCACGCGAAACTGGATACCGCCTTGGTCAGTGGCCAGAGCCATGGCACCACCAAGGG
CATCGATTTCCTTGACCAGATGACTCTTGCCGATCCCGCCGATGGCAGGATTGCATGACATTTGGCCAAGAGTCTCGATGTTGTGGGTCAGCAGTAGGGT
CTGACAGCCCATGCGGGCAGAGGCCAGAGCGGCTTCGGTTCCCGCATGGCCTCCGCCGATGACGATGACGTCAAAGCGGTCCGGGTAATTCACGTAACAC

CTCGAATGGCGCGATTCTGCGCCGTAATGAAAAACAGAGGGTGAAATCAGCCGCCCAGTATAAACCTCCTGTGGGTAACCGTCAGGGGTTTTCCACATCC
GGATGATGTCCGATAGGAAATAGCGACATTCCCAATATCAATATTAAAGAAAGAGAAAGGAAGTTCTGTTGTGGTAATGATTATTGGTGTGGACATTTTT
**TGTGGATAA**GCCATATTTTATGTTAATAATCAATTCGATAAAATGTTAGCCAAAAGTTGGACAAGGCGCGTAGCACTTGCGTACAGCATGTCATGTGCCT
GTGGATGAAATCATGACTTGCCCACATGCGGAACGAATCACCGGTTATGCACTGGTGAGTCCCGTGTTTGTCACCGTGGCTGTGCACATGGTGGGGTAAC
TGGCTTGATGAGGGCAACCATGCGGCTTTGTGGAAAAGAAGGTGCTGTTTGACGCTGAATTTATCCACATGAAGAAAAATGGGCCTACCGGTCGTCGGTA

GGCCCATGGAGATGGCGAATTGAGTGTGGAAGTTGGCACCGTTCAGTGCTTGAGCACACGCGACAGGAAGTTCTGAGTCCGTTCGTGTTGCGGGTTGTCG
AAGACCTCGCGGGGTGAGCCCTGTTCGACGATCTGGCCGCCGTGAATATAGATGACGCGATCGGCAACTTCACGGGCAAATCCCATCTCGTGGGTAACGA
TCATCATGGTCATGCCTTCATTGGCCAGTTCACGCATGGCGTCGAGTACCTCACCGATCATTTCCGGGTCGAGCGCCGAAGTCGGTTCGTCGAATAGCAT
CAAGCGCGGTTCCATGGCCAGGGCGCGGGCCAGGGCCACTCGCTGTTGTTGGCCGCCAGAGAGCTGGCTCGGGTACTTGTCGGCCTGGTCGCTGATCCCG
ACCCGCTCCAGTAGCTTGTTGGCAATCTCGGTGGCCCGGCCACGGTCGAGTCGGCGTACCTTGATCGGTGCCAGGGTGACGTTGTCACGCACGCTCAAGT
GAGGAAAGAGGTTGAATTGCTGAAAGACCATGCCGACTTCGGTGCGGATGGTCTGCAGGGCCTTCGAACTCTTGCCGTTCGGCAGCAGCTCGTTGCCGTC
GACCTCGATATGACCGTTCTGGAATTCCTCCAGGCCGTTGACACAACGGATCAGGGTCGACTTGCCCGAGCCACTGGCACCGATGATGACGACGACTTCC
CCAGGGGTGATCTCGAGGTCGATATCTTGCAGCACATGGAGACTGCCGAAATACTTGTTGACCTTGTCCATATGGACGATGGGCTTGTTCGAAGCTGAGT
TGGAGTGATTCATGGGCGATCCTTATTGTCCGACCAGGCCCTTGCGTTCCAGCGTACGCAGGATGAGAGAAAGGCTCCAGGTGATAGCCAGGTACAACAA
GGCCACCATCAGATAGACCTCGAGGGCGGTGAAAGTGGTGGCAATATACACTTGGCCCTGGCGGACCAGTTCGCCGACGCCGATGACCGAGAACAAGGAC

>Halomonas anticariensis FP35 = DSM 16096 [NZ_KE332393.1:c144894..144975], dnaA> ig dnaN>, 82 bp

WebSkew GCmin pos. 23475 (GCmin<>oriC1 30.1 kb; GCmin<>oriC2 1.8 kb; oriC2<>oriC1 28.3 kb)

>Halomonas qijiaojingensis KCTC 22228 [NZ_BMXS01000003.1:53348..53847], <rpmH oriC1 dnaA>, 565 bp

CGATCCGCACATCGAGTACGTCGGTGGTTACCGCGACCAGATCGCGACTGCTGACAGAAGTGGAGGTTGCCCCGGGCAGGTCGCCGTCGGTGACCGACTC
CGCCGCACTGGGTGTTGCAGGGGCTGCCAATCCTTCCTCGTCAGGCAATGGCTCACCATTGCGAGGTGCGTCGGCGGTGATGGTGGGGACTTCCGGTTCA
GGTGTGAGCTGACCGTAGTCCTGGTTCCATTGCACGACCAGCAGGTATGCCAGTACGGCAAGTGGAATCAATAACAGAAGTCGTTTTACGTCCATTGGGG
GCCTGCCCGGTGGGTAATGAACTGTCTGACGGCATAATGCATGAGCGAAAGGGCACACGAAAGCCTGCCGAGGCAGGCCGCTGCTGTCCCGTCAGGAGGT
GTTGGCCGACTGCTTGCGAACGTCCTTTTCCAGACGCCGCCACATGCCGTGGAGCTGGCGATGCAGGGTGTCGTTGTCCAGTTCTTGTACGCCACGCCGG
GATAGGACAACGATGTCCACGGCGGGTAGGCGATGCTGCTGGAGACGAAAGGATTCGCGAACAAGGCGTTTCAGGCGATTGCGATCGACTGCGTGGCGCA
CGTTCTTCTTGCTGAATACCAATCCGACTCGTGGATGACCCAGCTCATTGCTGCAAGCCAGTGCCAGCAAGCCCTTGCCGTGAACCTTGAAAGCAGCCCG
GTCGAAGACACGTCGGTAGTCCCCGGCGGTCAGCAGCCGCAGACGCCGGGGAAAGCCCTGACAGGACACACGCAATCCGAGATCAGGCGCTCAGACGCTT
GCGACCCTTGGCGCGGCGGCGGGCCAGAACGGCACGGCCATTCTTGGTGGCCATGCGGGCACGGAAGCCGTGCACACGCTTGCGCTTGAGAACGCTGGGT
TGAAAGGTGCGTTTCATAACGGTTGATTCCCACGTGGTTAATTGGACTTTACGTAAATGCTAGAGGCGCCCGGGGCCTGGTAGGACCCGAGGGAATTCGG

CTCAAGACCGGGAATTCTAGTGAATTCGCCAGCCGGGTGCAATTTTTCCCTTGCTACTCCCTTTCTGCCGTCGCTTGCTCGCCAAAGGACGTGGCTGCAT
TCGTATGCGGACAGGAGCGGATTGCTGGCCTCATGTCCATCGTAAGTTATCACTATTGTCATCCTTAGAGCCCGTTGTTATTAACACTGGGGATATTTTT
**TGTGGATAA**CCTAAAATAATTAATTAAAA**C**AATGAGATATATTGATTCGCAGCGTGTGGGAAAGATGCGGGTAATGGCTGGACTCTCTGGGGATAGGCTG
GTGAGAAGTCAGCAATTTTTCCACAGCCTCGAACTCGTCGCGGGGCGACTTCCAACCAGTGGTTATTCATTCCAATGCGTACCAGTTTTCCACAGCAAGG
GCGACGATGCGACGGCATGTGGATAACCCTTGGCTTGAGCTATACAATGTTGGGTCATTCGCAAACGGGATGGTGAGGTCGCGTGTCGGTCGCTCTTTGG

CAACAATGTCTGGATTACCTACAGGACGAGCTGAATTCCCAGCAGTTCAATACCTGGATCCGCCCCTTGCAGGCGGAGGAGGGGGAATCCAATGAACTGT
GCCTGCTGGCGCCCAATCGCTTCGTTAGGGATTGGGTCAATGACAAGTATGTGAAGCGCATCAACGAGTTGTTGCGCGAGCTTTCGCCTGCCAAGCCCCC
CAAGGTTTCGTTGAGCGTGGGCAGTCGCCGTGCGGCACCTTCGCCTCAGCCTCGTGAATTGGGCAATCCCGTTTCCGCCAGCCCACGTCAACCTGCGTCG
CCGGCCGTTCATACTCCGCCGCGCGGAGACTATGCCGACGAGCGAGAGATCGACCAGGCACGAGAGGATGGTGCGCGTGCAGGGGTTGGCCGTTCGGGAA
ATAGCGAGCGCCAGGTCCAGGTCGAGGGCAGCCTGAAGCACCAGAGTGGCTTGAACCCCAATTTCACTTTCGAAACCTTCGTCGAAGGCAAATCCAACCA
ATTGGCCAGGGCAGCCTCTCGCCAGGTGTCGGAGAATCCCGGAGGGGCCTATAACCCACTGTTTCTGTATGGCGGTGTCGGCTTGGGCAAGACGCACCTG
ATGCATGCGGTGGGAAACCAGCTTGCCGGCCGGCGTGAGAATGCCAAGGTGGTGTACCTGCATTCCGAGCGTTTCGTCGCTGATATGGTCAAGGCACTGC
AATTGAACGCGATCAACGACTTCAAGCGCTTCTATCGCAGTGTCGATGCCTTGCTGATCGACGACATACAGTTCTTTGCCGGCAAGGAACGCTCGCAAGA
AGAGTTCTTTCATACGTTCAACGCGCTGCTCGAGGGCGGCCAGCAAATGATCCTGACCTCTGATAGATATCCCAAAGAAATCAATGGGGTAGAAGAGCGT
CTGAAGTCTCGCTTCGGGTGGGGATTGACGGTGGCAATCGAGCCGCCGGAGCTGGAAACGCGGGTAGCGATCCTCATGAAGAAGGCCGATCAGGCCAAAG

>Halomonas qijiaojingensis KCTC 22228 [NZ_BMXS01000003.1:25069..25568], <mnmG oriC2 <COG1126 (glnQ), 549 bp

CAACCCCTTGATCGAGCGCACCACCTCGAGCTGCACGTCGAACGGCAGCGAGGTCGAGATGCCATTGGGATAGAGCTCATGGGTATCCAGCCCTTCGGGC
TCGATGAACACCTGATGGCTGGTCTTGTCGGCGAAACGATGCACTTTGTCCTCGATCGACGGACAGTAGCGAGGACCGATGCTCTCGATCGTGCCGGAGT
ACATCGGCGAGCGATCGAGATTGGCGAAGATGATCTCGTGGGTGCGCTCGTTGGTGTGGGCGATATGGCAGCTTACCTGGCGAGGATGCATATCGCGAGT
ACCCATGTATGACATGACCGGTGTCGGCGTGTCGCCCGGCTGTTCCGCGAGCACCGAGAAATCGACACTCTTGGCGTCCAGGCGTGGCGGTGTGCCGGTC
TTGAGTCGGGCCACATTGAACGGCAAGGCGCGCAATCGTTGAGCCAAGGCATTGGAGGGGGGATCGCCTGCTCGACCGCCGCGGCTCTGGTCAAGACCGA
TGTGCATCACGCCACCGAGGAAAGTCCCGGTACACAGCACCACCGTTTCGGCATGGAAACGGATACCCGTCTCTGTGACCACGCCCCTGACAGTCTCGTT
ATCCACAATCAGATCGCCCGCAGCTTGCTGGAAAATTGTCAGCCCAGGCTGGTTTTCCAGGATGCTACGGATCGCGGCCTTGTAACGCACTCGATCGGCC
TGGGCGCGTGTCGCACGCACCGCCGGGCCCTTGCGTGAATTGAGCACGCGGAACTGGATGCCTCCCCGGTCGGTGGCCAGAGCCATGGCACCGCCGAGAG
CGTCGATCTCCTTGACCAGATGGCTCTTGCCAATGCCCCCGATGGCGGGGTTGCACGACATTTGGCCCAACGTCTCGATGTTGTGAGTCAGCAGCAAGGT
CTGACAGCCCATGCGGGCGGAGGCCAAAGCGGCTTCGGTTCCCGCATGGCCTCCGCCGATGACGATGACGTCAAAGCGGTCCGGGTAATCCACGTTACAC

CTCGAATGGCGCGATTCTGGCGCCGTGATCAAAAACAGTGGGTGAAATCAGCCGCCCAGTATAAACCTCCTGTGGGTAACCGTCAGGGGTTTTCCACATC
CGGATGGCATTTGGTATCACCATGCCAGAGGTTCAATATCAATATAAAGATAAAGAAGAGAAGTTCTGTTGTGGTAGTAACTACTGGTGTGGACATTTTT
**TGTGGATAA**GCTTTCTTCTCCTTTTATAAACATGATGTTGCAATGTTGATCAAGAGTTGGACAAGCAGCGTAGTGGTTGCGTACAGCGTGTCAGGTGGCT
GTGGATGGAATCGCGACTTGCCCACATGCCAAATCGATCACCGGTTGTGCACCGACCACTACCGGTCTTGTCACCGTGGCTGTGCACATCATGGATCGGT
TGTCCCGATAAGGGCAACCATGCGGGTTTACGGGCGATGAGGTGCGGATTGACGTTGAAGTTATCCACAGGCAAAAAAAATGGGCCTGCCGGTCGACGGC

TGGCCCATGGAGGCGCTCACTGAGCGTGAGAATTGGCATCGTTCAGTGCTTGAGCACCCGCGACAGGAAGTTCTGGGTCCGCTCGTTCTGCGGGTTGTCG
AAGACCTCGAGGGGCGAGCCTTGTTCGACGATCTGACCACCATGGATATAGATGACGCGATCGGCGACCTCGCGGGCGAAGCCCATCTCATGGGTGACGA
TTATCATGGTCATGCCCTCGTTGGCCAATTCGCGCATGGCATCGAGTACCTCACCGATCATCTCCGGGTCGAGCGCTGAGGTAGGCTCGTCGAACAGCAT
CAGGCGTGGTTCCATGGCCAGTGCGCGGGCCAGCGCCACACGTTGTTGCTGCCCCCCGGAAAGTTGGCTGGGGTACTTGTCGGCCTGGTCGCTGATCCCG
ACCCGCTCCAGTAGCTTGTTGGCAATCTCGGTGGCCCGGTCACGGTCGAGTTGACGGACCTTGATCGGTGCCAGGGTGACGTTGTCACGCACGCTCAAGT
GAGGAAAGAGGTTGAATTGCTGGAAGACCATGCCGACTTCGGTGCGGATGGTCTGTAATGCCCTGCCGCTCTTGCCATGCGGCAAAAGCTCGTTGCCATC
GACCTCGATATGCCCTTGTTGAAATTCTTCCAGACCATTGACGCAGCGGATCAGCGTCGATTTGCCCGAGCCGCTGGCACCGATGATCACCACCACTTCA
CCGGGAACGATTTCGAGATCGATATCCTGCAACACATGCAGGCTACCGAAATACTTGTTGACCTTGTCCATGCGGACGATAGGTTTGCTCGAATTTGAGT
TGGAGTGGTTCATTGGCGTTCCTTATTGTCCGACCAGGCCCTTGCGTTCAAGCGTACGCAGGACGAGGGAAAGGCTCCACGTGATGGCCAGGTACAACAA
CGCCACCATCAGATAGACCTCCAGGGCGGTGAAGGTGGTGGCGATATATACCTGGCCCTGGCGAACCAGCTCGCCGACGCCGATGACCGAGAACAGGGAG

>Halomonas qijiaojingensis KCTC 22228 [NZ_BMXS01000003.1:55309..55398], dnaA> ig dnaN>, 90 bp

< page 1 | #114 | Halomonas rifensis CECT 7698 (no sequences available at NCBI as of January 2024)

WebSkew GCmin pos. 289083 (GCmin<>oriC1 27.9 kb; GCmin<>oriC2 2.4 kb; oriC2<>oriC1 25.5 kb)

>Halomonas xinjiangensis TRM 0175 [NZ_JPZL01000003.1:316710..317209], <rpmH oriC1 dnaA>, 531 bp

ATATCGCCGCCTTGCGGATCGATCCGGATATCCAGGACATCAGTGGTAACGGCAATCAGATCGCGGGTATTGGCGGCGCTGCTGCCTGTCTCGCCGGGAA
GGTCCTGATCGACGCTGCCGTTGCCGGCTGTCGAGCCCTGAGGAACCGTCAGGCCTTCTTCCTGAGCTCCGTCGGTCTGGGGGCTACTGGAGTCGCTGCG
AGAGGTCAGTGCCGGGGCGTCGGGTTGCGGTGCAACCTGACCGTAATCCTGATTCCATTGCACGACCAGCAGATAAGCCAGCACGGCAAGGGGAATCACT
AGCAAAAGTCGTTTTACGTCCATGAGGAGAGCCTGCCCGGTGGGTAGTGATCTATCTGACGGCGAAGTGGGTGAGCATTAGGCACACGAAAGCCTGCCGA
GGCAGGCTGAGTGGTCCCGTCAGGGGGTGTTGCCCGACTGCTTGCGAACGTCCTTATCCAGACGCCGCCACATGCCGTAGAGCTGGCGATGCAGGGTGTC
GTTGTCGATTTCCTGTACGCCACGCCGGGATAGGACAATGATGTCCACAGCGGGGAGGCGATGCTGCTGGAGGCGGAAGGATTCGCGAACGAGGCGCTTG
ATGCGGTTTCTGTCTACCGCTCGCTTCACGTTCTTCTTGCTGAATACCAATCCGACTCGCGGGTGGTCCAAGCCGTTGCCGCACGCCAAGGCCAGCAACC
CTTTCCCATGAACCTTGAAGGCAGCCTGGTCGAAAACGCGTCGGTATTCCCCAGCGACCAGCAATCGGCGGTCACGGGGGAAGGCCTGACAGGACACACG
CAATCCGAGATCAGGCGCTCAGACGCTTGCGACCCTTGGCGCGGCGGCGAGCCAGAACGGCACGGCCGTTCTTGGTGGCCATGCGAGCACGGAAGCCATG
CACGCGCTTGCGCTTGAGAACGCTAGGTTGAAAAGTGCGTTTCATAACGATTGATTCCCACGTGATTAATTGAACTGGTACTTCGATGTAAGAGGCGCCC

TGGCTCCCTAGGGAACCCGGGGGGATTCGGCTCAAGACCGGGAATTCTAGAGAATTCGTGGGTCGTGTGCAATTTTTCCTCGCGCTATCGTCATCTTCTT
GCGCGGCCAACGCGTGCGCGAGGTTGACCGACTCACAGGCGTTACCTTCAGCTGGAGACACTTTTTATAGATTTGTGTTATTAAACCTGTGGACCCATTT
**TGGGGATAA**GTTACTTATATAGTTATTTTTCAGTAAGTTATATTGATATATTAGCTGTGGGCAACCTGAGGGAAGGTTCTGGGTTCACTGGCCGTTTCGT
CATGGGCCGCTGTCATGCGGACGAGCCTATGGCATTTTTTGACCCGGTTATTCCCATCCTTCAAGCAAGAAAGACACAGCTTTTCCACAGAACGCTCCGG
TGTGCCTACCTATGTGGATAACCCAAGCCATGGGCGATACAATGGCCGGTCACTCGTTAACGGGATGTGAGGTCGCGTGTCGGTCGCTCTTTGGCAACAA

TGTCTGGATTACCTGCAGGACGAGCTGAATTCACAGCAGTTCAATACCTGGATCCGTCCGTTACAAGCGGAGGAGGGTGATGCCAATGAACTCTGCTTGC
TGGCGCCCAATCGCTTCGTCAGGGATTGGGTCAATGACAAGTATGCCAAGCGAATCAATGAGTTGATGCGTGAGTTGTCCCCTGCCAAGCCGCCCAAGGT
CTCCATCAGTGTGGGCAGCCGCCGGGTCGCGTCGCCTCCCCAGCCCCGGGAGTTGGGCGATCCAGTCTCGGCCAGTCCCCGTCAGCCTCCCCAGGCTCCT
GCCGTCAGGACCACGCCACGCAGTGAGTATCGTGATGAGCGGGAAATCGACCAGTTACGCGAAGAAGGCGGGCGCGGAAACACAGCGACCCGGCCCGCGG
GCAATGAGCGACAGGTGCAGGTTGAAGGCAGTCTGAAGCATCAAAGCGGGCTGAACCCCAACTTCACCTTCGAAACGTTCGTCGAGGGTAAATCGAACCA
ACTGGCCCGTGCCGCTTCTCGCCAGGTCTCCGAGAACCCGGGCGGGGCGTACAATCCTCTCTTCCTCTATGGAGGCGTGGGGCTGGGCAAGACACACCTG
ATGCATGCCGTCGGTAATCAATTGGCGGGGCGGCGTGAAAATGCCAAGGTGATCTACCTGCACTCCGAGCGCTTCGTCGCCGACATGGTCAAGGCCCTGC
AGCTCAACGCCATCAATGACTTCAAGCGCTTTTATCGGAGCGTCGATGCGCTGCTGATCGACGATATCCAGTTTTTTGCCGGCAAGGAGCGTTCGCAGGA
AGAATTCTTCCACACGTTCAATGCGTTGCTGGAGGGCGGTCAGCAAATGATCCTGACCTCCGATCGTTACCCCAAGGAAATTAGCGGGGTGGAAGAGCGG
TTGAAGTCGCGATTCGGCTGGGGCCTTACCGTGGCCATCGAGCCTCCTGAGCTCGAAACCCGGGTTGCGATTCTCATGAAGAAAGCCGACCAGGCAAAGG

>Halomonas xinjiangensis TRM 0175 [NZ_JPZL01000003.1:291235..291734], <mnmG oriC2 <COG1126 (glnQ), 537 bp

TCGAGCCCCTTGATGGAACGCACCACCTGCAACTGGACATCGAAGGGCAGCGACGTCGAGATGCCGTTGGGGTAAAGCTCGTGGGTCTCGAGGCCTTCGG
GCTCGATGAAGACCTGATGGCTGGCCTTGTCGGCGAAACGATGAACCTTGTCCTCGATGGAGGGGCAGTAGCGTGGCCCTACCCCCTCGATCACCCCCGA
ATACATCGGGGATCGGTCCAGGTTGGCGAAAATGATTTCGTGGGTTCGCTCGTTGGTATGCGCGATATGGCAGCTCACCTGCGGTGGGTGCTGCTCGCGA
GAGCCAAGATACGACATCACCGGTGTCGGCGTATCCCCGGGCTGCTCCTCGAGCTGGCCGAAATCGATGCTCCGGGCGTCCAGACGCGGAGGCGTACCGG
TCTTCAGGCGTGCCACGTTGAATGGCAGGGCACGTAGACGCTCAGCCAGCGCATTGGAAGGGGGATCGCCGGCACGACCACCGCGGCTCTGATCCAGCCC
AATGTGGATCACCCCACCCAGGAAGGTGCCGGTGCACAGCACGACCGCGTCGGCGAGGAAGCGGATTCCAGTTTCGGTGACGACTCCCCGGACGGTGTCA
GCGTCGACGATCAAATCGCCTGCAGCCTGCTGAAAGATCGTCAGGTTGGCCTGGTTCTCCAGCATGCCGCGTATGGCGGCCTTGTAACGAACGCGATCGG
CCTGGGCTCGCGTGGCCCTGACCGCGGGTCCCTTTCGAGCGTTGAGCACGCGAAATTGAATACCGCCGAGATCGGTGGCCCGCGCCATGGCGCCTCCAAG
CGCATCGATCTCCTTGACCAGATGGCTTTTGCCGATGCCGCCAATGGCGGGATTGCAGGACATCTGACCCAGCGTCTCGATGTTGTGAGTGAGCAGCAGG
GTCTGACAGCCCATGCGGGCAGAGGCCAGGGCGGCTTCGGTTCCGGCATGGCCTCCGCCGATGACGATGACGTCAAAGCGGTCCGGGTAATCCAAATGGC

ACCTCGAATGGCACCGTCGTGGCGCCTTGTAGAGCAATGGGTGAAAACAGCCGCCCAGTATAAACCTCCTGTGGGTAACCGTCAGGGGTTTTCCACACCT
CGAAGGCCAGCGGGATACCTCCACTGCCTATCAATCAATATCAAATAAATAAAAAGAAAGAAGTTCTGTTGTGGTTGTAACTACTGGTGTGGACACTTTT
**AGTGGATAA**CCCAATAAAACCAAGCAATAACATGACCTTGTAATGTTGACCAAGGGTTGGACAAGGGACGTGTGACTTGCGTACAGGTTGTCGCGAGCTT
GTGGATGGATTCTCGAGTTGCCCACAAGCCACGTCAACCACCGCTTATGCACTGCTGCATACCTGGCTTGTTACCGCGGCTGTGCACACACGACCGAGCT
TCTCTGAAAGACCCGAGGTATCAGGTCTGGCTGCCGATTGACGTGCCTTTGACCGGTTGAGTTATCCACACCCAAAAAAAAAGGCCTGTCGCAAACGGAC

AGGCCTTTTTTTGACATCTTCGCACGTGAATATCAGTGCTTGAGTACCCGAGAAAGGAAGCTCTGGGTGCGCTCATGCTGGGGATTATCGAAAACGACCC
CGGGAGGGCCTTGTTCGACGATCTGACCGCCATGGATGTAAATCACACGATCGGCAACCTCACGGGCGAAACCCATTTCGTGGGTGACGATCATCATGGT
CATGCCGTCACGGGCAAGTTCCCGCATGGCATCGAGTACCTCACCGATCATTTCGGGATCCAGCGCGGAGGTCGGTTCGTCGAACAGCATCAGCCTCGGT
TCCATGGTCAGTGCGCGCGCCAGGGCGACACGCTGTTGCTGGCCCCCGGAGAGCTGGGTAGGGGACTTGTCAGCCTGGTCGGCAATACCCACACGCTCAA
GCAGACGTCGTGCGTTGGCGACCGCTTGTTGCCGGGGCGTACCGCGTACCTTGATCGGGGCCAGGGTCACATTGTCGAGAACCGAGAGATGGGGAAACAG
ATTGAACTGCTGAAAGACCATGCCAACTTCGGTGCGTATGGTCTGTAGTGCCTTGGCGCTCTTGCCGTTGGGCAGCAGTTCGTTGCCGTCGACCTCGATA
TGGCCATACTGGAATTCCTCGAGGCCATTGACGCAGCGGATCAGGGTCGATTTTCCCGACCCGCTGGCGCCGATGATTACGACCACCTCGCCGGGGGCAA
TCTCGAGGTCGATATCCTTGAGAACGTGAAGGTCGCCAAAATGCTTGTTTACCTTGTCCATGCGCACGATAGGCGTTTCGGACGTAGGTGTCTGGGATGA
ACTCATGGCTAGGCTTCCTTATTGACCGACCAGACCCTTGCGCTCGAGCAGGCGCAGGATGAGGGAGAGGCTCCAGGTGATCGCCAGATAGAGCAGGGCA
ACCATCAGGTAGACCTCGAGGGCGGTAAAGGTCGTCGCGATATAGACCTGACCCTGGCGTACCAGCTCGCCGACACCGATGACCGAGAACAGCGAGGTAT

>Halomonas xinjiangensis TRM 0175 [NZ_JPZL01000003.1:318671..318755], dnaA> ig dnaN>, 85 bp

WebSkew GCmin pos. 250376 (GCmin<>oriC1 35.1 kb; GCmin<>oriC2 2.0 kb; oriC2<>oriC1 33.1 kb)

>Aidingimonas halophila DSM 19219 [NZ_FNNI01000003.1:285242..285741], <rpmH oriC1 dnaA>, 447 bp

CGGGTCGATGCGGACATCCAGGACGTCCGTGATGGCGGCAATCAACTCGCGTCCGTTCTCTTCCTCGTCTGCGACGTCATCGGCCATCTCGGTGTCGACG
TCATCTTCCGAGCTTGGGGCGGCCAGGCCTTCGTCATCTGCCGAGTCACCCTCGTTACCCGGTGCGTCCGAGTCAGTTCCAGCCTCTTCTGCACTGGAAA
TGGAAGGCGCATCTGTGTCCGGAGGCACTTGACCGTAATCCTGATTCCACTGCACCACCAAAAGGTAGGCAAGCACGGCTAATGGAATCAGTAGCAGGAG
TCGTTTTACATCCATGAGAGTTCTGCCCGATGGAAGTGGAGGTCTGGCGACGTATCAGCCGACGCGATGTTATCAGAGAAGCTGGCGTAACGATCTGCCT
CAGTGCAAGCCACCAATACTATCCGGGTGTCAGTGACATCTCGTCATGTATCGGAAACCGAGGCATGGTGTACCTGCTTTTCCAGACGTCGCCACAACCC
ATGAAGTTGGCGATACAAAGTGGCATTGTCCAGTTCGGTGATACCACGCCGGGCCAGGATGACGATATCAACAGCCGGAAGCCGATGTTGTCGCAGGCGA
AAGGACTCGCGCGTCACACGTTTGACGCGGTTACGATCTACCGCACGGCGAGCGCTTTTCTTGCTGATCACGAGCCCAAGCCGAGGATGCCCCGAATCGT
TATTGCATGCCAGGGCCAGTAATCCTTTGCCGTGTACCTTGTACGTCGCGTTGTCGAAAACGTGTCGATAGTCTCCGGCGCTGAGCAACCGCTGGCGCCG
GGGAAATTCATATTGAGACACGTATTCGCCGTCAGGCGCTCAGCCGCTTGCGACCCTTGGCACGACGACGCGCCAGAACAGCACGACCATTTTTGGTTGC
CATGCGTGCCCGGAAGCCGTGCATGCGCTTGCGCTTGATAACGCTGGGTTGGAAAGTGCGTTTCATAATGCGTGATTCCCACGTGGTTGAATGTGACTTG

ATGACGAATCAATGCGACCCCGTGTTCTATACAAGCGTCGGGGTCTTCGGACCAAGACGGGGATTTTAGTGAATTCGCCAGTCCGGTGCAATTTCCTCGA
CACGTTGTTTGCGCAAGAGGCGACGATAGCGGGGTGATGGGTGTGTGTCGTGTCGGTCCGCCTGTTAGAAGTATTTGTTATTAACCTTGGGGATGGTATT
**TGTGGATAA**CCTTTTAAATCCTTTATATT**C**AACTATTTAAAATGATTCCAAGGTTGTGGCTAACTTGGTGGAAAGAACGTGGGGCTCGTGGGGATCGCTA
ATGGAGCATCATGAGTTTTCCACAGTACTTGATAGTGGGTGACATGTGGATAACTTGGGCTCAGGACGATACAATGTCCGGTCTTCATTGGCCCATGAAT
GGATGAGGTGGCGGTGTCTGTCGCTCTTTGGCAACAGTGTCTTGAGTACCTGCAGGATGAATTGAGTTCACAGCAGTTCAATACGTGGATTCGTCCGTTG

CAAGCAGAGGAAGGTGAATCCAACGAATTGCGCCTGCTGGCGCCCAATCGCTTTGTGCGTGACTGGGTCAGTGACAAGTATGCCAAGCGTATCAATGAAT
TGATACGTGAACTGTCCCCTTCCAAGCCTCCCAAGGTGTCATTGACTGTAGGCAGTCGCCGGGCGGGACCCGCCCCTCAGCCACGGCAATTGGGGAATCC
GGTATCCGCCGATGCGTACAGTAATAGACCGGCGGCCCCGGCCGTGCAGGTTCCGCCTCGTGGGGATCTGCCCGATGAGGGAGAAATCGGTGACGCCGGG
GCCGGTGAAAGCGAAGTTTCCCAACAGGCGGAATCAAACACCAGAGCCGGCGGCGAGCGCCAGGTGCAGGTGGAGGGAAGTCTCAAGCACCAGAGTGGAC
TCAATCCCAATTTCACTTTCGAAACGTTTGTGGAAGGTAAGTCCAACCAACTGGCCAGAGCCGCTTCACGTCAGGTGTCCGAAAACCCGGGTGGGGCTTA
CAATCCCCTGTTTCTTTATGGCGGTGTCGGGTTGGGTAAAACGCACTTGATGCATGCCGTCGGGAATCAATTAGCGATGAAAAGCGACAATGCCAAGGTG
GTTTATCTGCATTCGGAACGCTTCGTCGCTGATATGGTCAAGGCACTCCAGCTGAACGCCATCAATGATTTCAAGCGTTTTTACCGTAGCGTCGATGCGC
TGTTGATCGATGATATTCAGTTCTTTGCCGGTAAGGAGCGCTCGCAGGAGGAGTTCTTCCACACTTTCAATGCCCTGTTGGAAGGTGGACAGCAAATGAT
CTTGACTTCTGATCGCTATCCCAAGGAAATCAATGGAGTAGAAGAACGTCTGAAGTCCCGTTTTGGATGGGGTCTGACCGTTGCCATAGAGCCGCCTGAA
CTTGAAACGCGTGTAGCGATCCTGATGAAAAAGGCGGATCAGGCCAAGGTGGATCTGCCTCATGATGCTGCGTTTTTCATTGCACAGAAAATACGCTCCA

>Aidingimonas halophila DSM 19219 [NZ_FNNI01000003.1:252118..252617], <mnmG oriC2 <COG1126 (glnQ), 550 bp

GACCATCGATCGACCGGACGACCTGTAACTGGATGTCGAAGGGCAGTGAGGTCGATATGCCATTGGGATATAGCTCATGAGTCTCGAGTCCCTCGGGTTC
AACGAAAATCTGGTGACTCGACTTGTCGGCGAATCGATGCACCTTGTCCTCAATCGAGGGGCAGTAACGTGGACCGACGCCTTCAATGGTTCCCGAGTAC
ATGGGGGAACGATCCAGATTGGCCAGGATGATCTCGTGAGTTGTCTCGTTGGTGTGTGCGATATGGCAGCTCACTTGGCGTGGATGCCACTCACGTTCGC
CCATGTATGACATGACTGGTGTCGGCGTATCGCCGGGCTGCTCCTCCAGGCGGGAGAAGTCGATGCTCTTGGCGTCCAGCCGTGGTGGAGTTCCTGTCTT
GAGCCTGTCGACACGGAATGGCAGTTCGCGTAGCCGAGATGACAGTGCATTCGAAGGGGGGTCACCGGCGCGTCCACCACGACTGTTGTCGAGCCCGATA
TGAATGACTCCGCCCAGGAAAGTTCCGGTACACAGGACCACAGTGTCGCTATGAAAACGGATGCCCGTTTCGGTAATGACGCCACGCACGGTATCGCCTT
CAACGATCAGGTCGCCGGCGGCTTGCTGGAAGATCGTCAAATTCGGCTGGGTTTCGAGACGGCGTCGAATGGCAGCCTTGTAAAGAACACGATCAGCTTG
TGCTCGAGTGGCACGAACTGCGGGTCCTTTGCGGGCATTGAGAATGCGGAACTGGATGCCGGCTTCATCCGTGGCTTCCGCCATGGCGCCTCCCAGGGCG
TCGATCTCCTTGATCAGGTGACTCTTGCCGATTCCCCCCATGGCAGGATTACAGGACATCTGACCGAGTGTCTCGATGTTGTGTGTCAGCAACAGGGTTC
GGCTGCCTGAACGGGCAGCGGCCAGCGCAGCCTCTGTTCCTGCATGGCCACCACCGATGACGATGACTTCATAACGGTCGGGATAATCCACGTTGCACCT

CGAGGGATGCCAGGTCGGGCATCGTTGCTAAAGGATGTATAGCGAAATCAGCCAGCCAGTATAAACCTCTTGTGGGTAACCGTCAGGGGTTTTCCACACC
TCGTTCGTTTCTGCCTGCCTGTCCATCCTTTCAGTAATCATTATTGGTTATTAAATAAAGAAGTTCTGTTGTGGTTGTTATTACTGGTGTGGATCATTTT
**TGTGGAAAA**GAAGATTTTTCATTTTAAAAA**C**AATAAGATATTGTGTTCATAAGAATAAGGAATCCACGGGTAAAAGCAGCGCACGAACGGCGATGAAGCT
GTGGATGAAATTGCTGTCTATCCACAGCCCGGTCCATTCAGCAGCTATACACCGATCTGTACCGGATCTGTCACCGTAGCTGTGAACAAGTTGGTTCTTT
ATCAGAGAAGCCTCTGTGATAAGGGTTGTAAGCGGATAAGACTACTGGAAGGAGGAAAATTGTCCACAAGGACGAAAACGGGCCTGTGATGAATCACAGA

CCCGTCAATGGCAGCTATCGATGAAAATCCGTCAGGCAGGATTAGTGTTGGAGCACTCGAGAAAGAAAATTTTGGGTCCTTTCATTCCGGGGATTATCGA
AGACCTCATGAGGCGGCCCCTGTTCGACGATTTCCCCTTGATGGATATAGATCACCCGATCGGCGACTTCGCGGGCAAAGCCCATCTCGTGGGTCACGAC
CATCATGGTCATGCCTTCGCGAGCCAGTTCCCGCATGACATCCAGTACTTCACCGATCATCTCCGGATCGAGAGCCGAGGTCGGTTCATCGAACAGCATC
AGTCGTGGATCCATTGCCAGAGCCCGGGCAATGGCAACGCGCTGTTGCTGCCCCCCGGAGAGTTGTGTCGGGTATTTGTCGGCTTGATCAGCAATCCCGA
CTCGTTCGAGAAGCCGTTCGGCGGTCGTGTCCGCTTCCTGTGAGGTCATCCCCCTGACCTTGAGCGGTGCCAGTGTCACATTGGCCTTGACCGTCAAATG
GGGGAACAGATTGAACTGTTGGAAAACCATACCCACTTCGGTTCGGATGGCCTGCAGGGATTGACCGCTTTTGCCATAGGGCAGGAGTTCCTTGCCGTCG
ACATGCAGCCGTCCTTTTTGATACTCCTCCAGACCATTGACGCAGCGTATCAAGGTGGATTTGCCTGAACCGCTTGCCCCGATGATCACCAGTACTTCGC
CGGGCGTGACGTCCAGATCGATGTCCTTCAGGACGTGCAGGCTGCCGAAGTGTTTGTTCAGCTTCTCAAGAGAGACAATGGAATGCTCTGGCGAGGAAGG
GGTCTGTGAATGTTCATCATTCATCGCAATGTTGTCCCTTATTGTCCGACCAGGCCCTTGCGCTCGAGCATGCGCAAGGCCAGAGAAAGGCTCAAGGTGA
TGACCAGATAGAGGATCGCTACCATGAAATAGACTTCAAGGGCGTTGAAGGTGGTCGCAATGTAGACCTGTCCCTGGCGAACCAACTCGCCGACACCAAT

>Aidingimonas halophila DSM 19219 [NZ_FNNI01000003.1:287152..287205], dnaA> ig dnaN>, 54 bp

WebSkew GCmin pos. 471126 (GCmin<>oriC1 36.9 kb; GCmin<>oriC2 3.3 kb; oriC2<>oriC1 33.5 kb)

>Aidingimonas lacisalsi XHU 5135 [NZ_VRYH01000001.1:507747..508246], <rpmH oriC1 dnaA>, 459 bp

```
AAGTACGTCGGTGGTAACAGCAACCAGCTTGCGACCGTTGTCCGCCGGATCGGCGGCTTCATCGGCCATGTCACTGTCGATGTTGTCATCCCCGCTCGGT
GCGGCCAAGCCTTCGCTGTCGCTGTCTCCCGAGGCGCTCTCGTCGTCGACGCTATTGTCGTTGACGCTGGGGTCGTCGGAACTGGAAATCGCGGGAGCGT
CGGTTATCGGAGGCGCTTGACCATAATCCTGATTCCACTGCACAACCAGCAGATAGGCGAGCACGGCCAATGGAATCAGTAGTAGGAGTCGTTTTACGTC
CATGAGAGTTCTGCCCGATGGAAGTGGAGGTCTGGCGACGTGTCGGCCGACGGAGCGACCCCGTATGGCTGGCGTGCCAGCCGACGGCAATGCAGGCTGG
CCATGTCGCCCGCATGGGGCTCTGGCACCTCGTCATGGACCGGTAGCCTATGCATTGCGCACACTCTTCTCGAGACGCCGCCACATGCCGTGTAGTTGGC
GATGCAGTGTAGCATTGTCCAGTTCGGTGATACCCCGGCGCGCCAGTATGACGATATCGACAGCCGGCAGCCGATGCTGTTGCAGGCGAAAGGACTCGCG
AGTCAAACGTTTGATGCGGTTGCGATCGACAGCCCGGCGTGCGCTTTTCTTGCTGATCACCAAGCCAAGCCTTGGATGTTCCGTACCGTTTCTGCAAGCC
AGGGCCAATAACCCTTTGCCGTGCACCTTGTACGTCGCATTGTCGAAAACGTGTCGATAGTCTCCGGCGCTGAGTAGTCGCAGGCGCCGGGGAAATTCGT
TACGAGACACGCGCTCGCCGTCAGGCGCTCAGATGCTTACGACCCTTGGCGCGGCGGCGCGCCAGTACGGCACGACCATTCTTGGTTGCCATGCGTGCCC
GGAAGCCATGCATGCGCTTGCGCTTGAGAACGCTGGGTTGGAAGGTGCGTTTCATAATGAGTGATTCCCACGTGGTTGAATTGGTCTCAATGTCGAATCG

TTCTGCCCCGACGCTTGTTCCAGAATGTCAGGGTCTTCGGATCAAGACGGGGAATTTTAGTGAATTCGCTAGTCCGGTGCAATTTTTCTGAGATTACGAT
TGCATCACTCTGTCACTGGCGAGCCAGACCCAAGGGCGGTGTAGTGCCGAAAGCGCCGTTCATACTAGAAGGCATTGTTATTAACATTGGGGATCAAGGT
TGTGAATATTTCTAAAAAATACATAAAAATCAACCATTTGAAATATTTTATAAACTGTGATCAATTTAGTGGACAGATCGTTGGGTAATGGAATGTGCGT
GCCGAGGTAACAAACGTTTTCCACAGCACCGAGAGTGCGTCGTCGTGTGGATAACTCAGCCTCAGAAAGCTACAATGGCCGGTCTTCGTTAACGCGTTAG
TGGGTGAGGTTGCGGTGTCGGTCGCTCTTTGGCAACAATGTCTTGATTTTCTGCAGGATGAATTGAACTCTCAGCAGTTCAATACGTGGATTCGTCCCCT

GCAGGCGGAAGAAGGTGAATCCAATGAGCTTTGCCTGCTAGCCCCCAACCGTTTCGTACGTGACTGGGTCAACGACAAGTATGCCAAGCGCATCAATGAA
TTGATGCGTGAGTTGGCGCCGACCAAGCCCCCGAAAGTGTCCTTGACGGTAGGCAGCCGACGTGCTGCGCCTGCTCCGCAACCCCGTGAATTGGGTAATC
CGGTTTCTGCCGGCGCCTATAGCCAACCGGCGGCCCCTGCAGTGCATACGCCACCGCGAGGCGATATCCCCGATGAACGTGAAATCGAAGCCACTGCAGA
CAACGATACGCCGCAGCAGGAAAATGCGGCGCGTAGCGGTAGTGGTGAGCGTCAAGTCCAGGTCGAAGGTAGTCTCAAGCACCAGAGTGGACTTAATCCC
AACTTCACTTTCGAAACTTTCGTGGAAGGGAAATCCAACCAACTGGCGCGTGCAGCGTCGCGCCAGGTCTCGGAAAACCCGGGAGGAGCCTACAACCCGC
TGTTCCTCTATGGTGGCGTTGGATTGGGCAAGACACACTTGATGCATGCTGTAGGAAACCAACTGGCGGTGAAGAGTGAGAAGGCCAAGGTCGTCTATCT
GCATTCAGAACGCTTCGTGGCCGATATGGTCAAAGCATTGCAGCTCAATGCAATCAATGATTTCAAACGCTTCTATCGTAGTGTCGATGCCTTGTTGATC
GATGATATCCAGTTCTTTGCCGGTAAGGAGCGTTCTCAGGAGGAGTTTTTCCATACCTTCAACGCCCTGCTTGAGGGTGGCCAGCAAATGATCTTGACCT
CGGATCGTTACCCCAAGGAAATCAATGGTGTAGAAGAGCGCTTGAAGTCTCGTTTTGGATGGGGGCTCACCGTCGCCATCGAGCCGCCGGAACTCGAGAC
GCGAGTGGCGATCTTGATGAAGAAAGCGGACCAGGCCAAGGTCGATCTACCCCATGATGCCGCTTTCTTCATCGCCCAGAAAATTCGCTCCAATGTGCGT
```

>Aidingimonas lacisalsi XHU 5135 [NZ_VRYH01000001.1:474226..474725], <mnmG oriC2 <COG1126 (glnQ), 559 bp

```
GACCACCGATTGACCGGACTACCTGTAACTGGATGTCGAAGGGCAGTGAGGTCGAAATGCCGTTGGGATAGAGCTCATGGGTCTCGAGCCCTTCAGGCTC
GACAAAAATTTGATGGCTCGACTTGTCGGCGAAACGATGAACCTTGTCTTCGATCGATGGGCAATAACGTGGCCCTACTCCCTCGATGGTTCCCGAATAC
ATCGGCGAACGGTCCAGGTTGGCCAGGATGATCTCGTGTGTTTTCTCGTTGGTATGAGCAATATGGCAACTCATCTGTCGAGGATGCCAGTCGCGTTGCC
CCATGTATGACATGACGGGGGTCGGTGTATCGCCCGGCTGTTCCTCGAGACGTGAGAAATCGACGCTCTTGGCGTCCAATCGAGGCGGTGTTCCAGTCTT
GAGGCGATCGACGCGAAACGGCAACTCGCGCAAACGGGACGAGAGTGCATTGGAAGGAGGATCCCCTGCACGACCACCGCGACTGTTATCGAGTCCGATA
TGAATCACGCCGCCCAGGAAGGTACCTGTGCACAGGACAACGGTATCACCATGAAACCGGATACCGGTTTCAGTGACGACCCCGGTCACGGTATCACCCT
CGACGATCAGATCGCCGGCGGCCTGCTGGAATATCACCAGATTCGATTGAGTTTCCAACTGGCGACGTATGGCGGCCTTGTAGAGTACACGATCGGCTTG
TGCCCGAGTTGCTCTGACGGCTGGGCCCTTGCGGGCATTGAGGACGCGGAACTGAATGCCTGCCAGGTCAGTAGCTTTGGCCATTGCGCCATCCAGAGCA
TCGATTTCCTTGATCAGGTGGCTTTTGCCTATGCCTCCCATGGCGGGGTTACAGGACATCTGGCCGAGTGTCTCGATGTTGTGTGTCAGCAGCAGTGTTC
TGGCGCCAGTGCGGGCAGCGGCCAGGGCGGCCTCGGTTCCCGCATGACCGCCACCGATGACGATGACCTGGTAACGATCGGGATAATCCACGTTGCACCT

CGAGTGATGCTGGCTTGGCATCGTTGCTAAAAGACATATAGCGAAATCAGCCAGCCAGTATAAACCTCTTGTGGGTAACCGTCAGGGGTTTTCCACACCT
CGTGTGTCGTACCCTGGTTGTTCAGGGTCTCTAATAATTATTATTGATTGTTAAGTAAAGAAGTTCTGTTGTTGTTGCTACTACTGGTGTGGATGTTTTT
TGTGGAAAACTATATATAGTCTCCGAAAAACATATAGATACAGTGTTCACAAGTATGCGGAAAGTACGGGCAGTAGCTGCGTACGGACAGCGAGGAAGCT
GTGGACGATAATGGCATTTTTCCACAGGACTCACTTTTCACCACTTGTCCACCGTCGTCTACCGAGCTTGTCACCGTGGCTGTTAACACATCGGTTGTAA
TTCGATGAACGGCTCAAGAGCAGGGGATGAGAGGGAATTGGAATGGATTGGCGGGATAAGTTATCCACAGAGCGAAAACGGGCCCATCATGAACGACGGG

CCCGCGATGGCCTTTCACCAATGAAGGAACTTCAATGCTTCACTGGCCCCTCAATGCTGAAGTACCCTCGACAGGAAGTTCTGAGTACGTTCCTGTTGTG
GATTGTCGAACACCTCATGGGGCGGTCCTTGCTCGATGATTTCGCCCTGGTGAATGTAAATGACACGATCGGCTACCTCTCGAGCGAACCCCATCTCGTG
AGTCACCACCATCATCGTCATGCCTTCCCGGGCAAGTTCACGCATGACATCCAGGACTTCACCGATCATTTCCGGATCCAGAGCGGATGTGGGCTCATCG
AATAGCATCAAGCGGGGTTCCATCGCCAGTGCTCTAGCGATCGCTACACGTTGCTGTTGTCCGCCTGAGAGTTGAGTCGGGTATTTGTCAGCCTGATCGG
CAATTCCGACCCGCTCCAGAAGTCGTTCTGCGGCAGTGCCGGAGTCTTGTTGGGATAGCCCCCGTACCTTGATAGGCGCCAGCGTGACATTTTCCCGGAC
CGTCAGGTGAGGGAAAAGATTGAATTGCTGGAAGACCATACCGACCTCGGTACGGATGGCCTGAAGTGCTTGACCGCTCTTGCCATGTGGTAGGAGCTCC
TTCCCGTCAACGCGCAATTGGCCACTTTGATACTCCTCCAGGCCATTGATACAACGTATCAGGGTGGACTTGCCTGAGCCACTGGCGCCAATGATCACGA
GGACTTCACCTGCGCTCACTTCGAGATCGATGTCGTTGAGGACATGGAGGTTGCCGAAGTGCTTGTTCAGTCTCTCGAGCTCGACGATGGGCACTGGCGT
CGAGGATTCCTGGGAATGTTCGTCACTCATCGCATTTATCCCTTATTGTCCGACCAAGCCCTTGCGCTCGAGCTGGCGCAAGACGATGGAGAGGGTCAAG
GTGATGGCCAGGTAGAGTATTGCGACCATGAAATAGACTTCGAGGGCATTGAAGGTCGTTGCAATATACACTTGACCCTGACGTACCAACTCGCCAACAC
```

>Aidingimonas lacisalsi XHU 5135 [NZ_VRYH01000001.1:509649..509703], dnaA> ig dnaN>, 55 bp

WebSkew GCmin pos. 975357 (GCmin<>oriC1 65.9 kb; GCmin<>oriC2 32.3 kb; oriC2<>oriC1 33.6 kb)

>Modicisalibacter tunisiensis LIT2 [NZ_JAGXFD010000001.1:c909242..909741], <rpmH oriC1 dnaA>, 607 bp

CGTCGGTCTTCACGTCGATCAGGCTGCGCCGAGTGTCGGTGGCGGCGTTGTCCGCCATCGTCGGGCTGGCGCTGCCCCCTGACTTGCCGGAGGGTGCAGC
CAGGCTCTGCTCGTCGGCGGGCTGGCCGCCGCCATCGCCGGCGGCCGGCTGACCGCTGCTGCTGAACGAGGGCGAGGAGGTTTCGGCAGGGGTCTGCCCG
TAGTCCTGATTCCACTGGATGACCAGCAGGTAGGCAAGAACCGCGAGCGGAATCACGAGTAACAGTCGTTTTACGTCCATTAGGGATCTGCCCGATGGGA
AGTTGGATCACTCTGGCGACTCGGCCAGCACGCTGTTCCGGTCAGGAGGGCGAGGCGACCGCCGCCTGCTTGCGAACCTCCTTTTCCAGGCGTCGCCACA
GGCCGTGCAGCTGGCGGTGCAGCGTGGGGTTATCCAGGTCGCTGACGCCGCGACGGGCGAGGACCACGATATCCACGGCGGGCAGGCGATGCTGCCGGAG
TCGGATGGATTCGCGCACGAGGCGCTTGAGACGATTGCGATCAACGGCACGACGGACATTCTTCTTGCTCACCACCAGGCCGACACGGGGATGCCCCAGT
GCATTGGCTCTGGCGAGAACCAGGATGCCCCTGGCGTGAACCTTGTACGTGGCGCGATCGAAGACGTATCGATAGTCCCCGGCATTCAGCAGGCGCAGTT
GCCGGGGAAAGCCCTGACAGGACATTCGCGGCCGATCAAGCGCTCAGACGCTTGCGACCCTTGGCACGACGACGTGCCAGAACCTGACGACCATTCTTGG
TGGCCATGCGAGCACGGAAGCCGTGCGCACGCTTGCGCTTGAGAACGCTGGGTTGAAAGGTGCGTTTCATGACGTTGAATTCCCACGTGGTGGATTGGAA
GTGACAGTTGTCCCCCCTCCCCAGGCGGGGCAGCGGGACGGATTTGGCTCAAGGCGGGAAATTCTAGTGAATTCGCCAGCCACGTGCAATTTCTGCGCGC

GATTATCCACAACAGGTTCTTTCAACGCGCCTGGCGGTTCCGCTGTTCGCCGCCGTGCTGTGGCCGCCTTCCTGCCAAAAATGCCAGCCGTCAGCCGGGC
GAAGACCGTCGTCGGATCAGTGCTTCCGTCGTGATCGACGGTCCTATTACAATTATTGGGTTTCTTGACTGATGTTGTTATTACTATTGGGGATAAAAAT
**TGTGGATAA**CCTGATAGATTCCTTGTGAATCATGCAGGTGGGCGTTTGATGAATGTGTGTGCAGTCGTTTGGAAAACGGTGATTCTCCTGCCGGGAGCCT
GTGGATGAAATGACGGGTTATCCACGGTTCGCATCCTGTCCCCGAAAGATACCCGACGCGTCCACCTTCCCGTACCCGAGTTGTCCACAAAGCCATCGTG
GCCGGGTGGCGTGTGGATAACTCGCCTCGAGAACGCTACAATGCCCGGTCATTCGCAAATCGGGATGGGTCACACGCGTGTCGGTCGCTCTTTGGCAACA

ATGTCTGGACTACCTGCAGGACGAGCTGAACTCGCAGCAGTTCAATACCTGGATTCGCCCCTTGCAGGCAGAAGATACGGCGTCCAACGAGCTCACCTTG
CTGGCGCCCAATCGCTTCGTGCGCGATTGGGTCAGCGACAAGTACCTGAAGCGCATCAACGAGCTGCTGCGTGAACTGGCGCCAGCCAAGCCGCCCAAGG
TGGCGCTGGCGGTGGGCAGCCGGCGTGCCGCGCCCAGCCCCCAGCCCCGGGAACTCGGCAACCCGGTGTCGGCCAGCCGCTTTCGTCAGCCCGCGGCCCC
GGCGGTGCAGACCCCGCCGCGGGGCGACTACGCCGACGAGCAGGAGATCGAGACGCTGCGCGAGGAGTCCGCGCGGCGCGGTGGCTCGAACGGCGAGCGC
CAGGTTCAGGTGGAGGGCAGCCTCAAGCACCAGAGCGGCCTCAACCCCAACTTCACCTTCGACACCTTCGTCGAGGGCAAGTCCAACCAGCTGGCCCGGG
CAGCCTCGCGGCAGGTCTCGGAAAATCCCGGTGGCGCCTACAATCCACTGTTCCTGTACGGCGGGGTGGGGCTGGGCAAGACGCACCTGATGCACGCGGT
GGGCAATCAGCTGGCCATGCAGCGCGACAACGCGCGGGTGGTCTACCTGCATTCCGAGCGCTTCGTCGCCGACATGGTCAAGGCGCTGCAGCTCAACGCC
ATCAACGACTTCAAGCGCTTCTATCGCAGCGTCGACGCCCTGCTGATCGACGATATCCAGTTCTTCGCCGGCAAGGAGCGCTCTCAGGAAGAGTTCTTCC
ATACCTTCAATGCGCTGCTCGAGGGCGGTCAGCAGATGATCCTCACCTCGGATCGCTACCCCAAGGAGATCAGCGGGGTCGAGGAACGGCTCAAGTCGCG
CTTCGGCTGGGGGCTGACGGTGGCGATCGAGCCGCCGGAGCTGGAGACGCGGGTGGCGATCCTGATGAAGAAGGCCGACCAGGCCAAGGTCGACCTGCCG

>Modicisalibacter tunisiensis LIT2 [NZ_JAGXFD010000001.1:c942849..943348], <mnmG oriC2 <COG1126 (glnQ), 554 bp

GTTCTCGAGCCCCTTGATCGAGCGCACCACCTGCAGCTGCACGTCGAAGGGCAGCGAGGTGGAGATGCCATTGGGGTAGAGCTCGTGAGTCTCCAGACCC
TCCGGCTCGATGAAGATCTGATGACTGGCCTTGTCGGCGAAGCGATGAACCTTGTCCTCGATCGACGGGCAGTAGCGCGGGCCGACGCCCTCGATCGTCC
CCGAGTACATCGGCGAGCGGTCGAGGTTGGCGAAGATGATCTCGTGGGTGCGCTCGTTGGTATGCGCGATGCGACAGCTCACCTGCCGCGGGTGCAGGTC
CCGTGATCCCAGGTAGGACATCACCGGGGTCGGGGTGTCGCCGGGCTGTTCCTCGAGCTGGCTGAAGTCGAGGCTCTTGGCATCGAGCCGCGGCGGCGTG
CCGGTCTTGAGGCGATCGACGCGGAACGGCAGGGCGCGGAGCCGTTGCGCGAGGGCGTTGGCGGGAGGATCGCCGGCCCGGCCGCCGGTGTGGCTGTCCA
GCCCGATGTGGATAACCCCGCCGAGGAAGGTGCCGGTGCACAGCACCACGGTGTCGGCATGAAAGCGGATACCGGTCTGGGTGACCGCGCCCCTGACCGT
GTCGCCGTCGACGATCAGGTCATCCACACCCTGCTGGAACAGCGTCAGGTTGGGCTGGTTCTCCAGCATGCCGCGAATCGCCGCCTTGTAGCGCACGCGG
TCGGCCTGGGCACGGGTGGCGCGCACGGCCGGACCCTTGCGGGCGTTGAGCACACGGAACTGGATGCCGGCATGGTCGGTTGCCAGCCCCATGGCCCCGC
CCAGGGCGTCGATCTCCTTGACCAGGTGGCTCTTGCCGATGCCACCGATCGCCGGATTGCAGGACATCTGTCCCAGTGTCTCGATGTTGTGAGTCAGCAG
CAGGGTCTGACAGCCCATGCGGGCAGCGGCCAGTGCGGCTTCTGTTCCCGCATGGCCACCGCCGATGACGATGACGTCAAAGCGGTTGGGATAATCCACG

GATACCTCGACTTGCGCGCCGGGCGCGCGACTTCAGGGCAAGTATCAACAAGCCGCCCAGTATAACGCCCCTGTGGGTAACCGTCAGGGGTTTTCCACAT
CAGGTTCCGGTCATGCTCTCCCCCTTCGAGTCTTAAAGAAACAATATAAGAAGAGAAAAGAAGTTCTGTTGTGGTAGTAACTACTGGTGTGGACATTTTC
**TGTGGATAA**GACGGACTTTTCGTTCGTTGTCAGTCCCCTGAAATGTTGACGATGCATTGCAAAAAAGGTAGACGGCCTGCGGAGACCCGGTTCGGGAGCT
GTGGATGAAAAGGACGGTCATCCACAGCCGTCCCCGTCCCCGGCTTGTCCACCGGCTTGTACCGGGGCTGTGCGTGTAGCGATCAACAGTGTGGCTTGTC
ACCCGTCAGGACACGCCTGGCGCGGGCTGCAGGCAAGACAGCGTGGACACGGCGAAAAACTTGTCCACAGGCGGAAATCGGGGCTGGGGAAGGAATCTGG

AGTGGGGGCGAGCGGAAGAAAGACGCCCCTCGCGGTCGGCGAGGGGCGGTGGGTCAGTGCTTGAGCACGCGGGCGAGGAAGGACTGGGTGCGCTCGTGCT
CGGGGCTGTCGAAGACCGTCTCGGGCGGGCCCTCCTCGACGATGCGCCCCTTGTGGACGAAGATCACCCGGTCGGCGACCTCGCGGGCGAAGCCCATCTC
GTGGGTGACGATCATCATGGTCATGCCGTCGTTGGCGAGCTCGCGCATGGCGTCGAGCACCTCGCCGATCATCTCCGGGTCGAGCGCCGAGGTCGGCTCG
TCGAAGAGCATCAGTCGCGGCTCCATGGCCAGCGCCCGGGCCAGGGCGACGCGCTGCTGCTGGCCACCGGAAAGCTGGCCCGGGTACTTGTCGGCTTGGT
CGGCGATGCCGACCCGGTCGAGCAGCCGGTCGGCGATCTGCACGGCCTGGTCGCGGCTCAGGCCGCGGACCTTGATCGGCGCGAGGGTGACGTTGTCACG
CACCGAGAGGTGCGGAAACAGGTTGAACTGCTGGAAGACCATGCCCACCTCGGTGCGGATGGTCTGCAGGGAGCGCCCGCTCTTGCCGTAGGGCAACAGC
TCGTTGCCGTCCACCGTGATGTGGCCGCTCTGGAACTCCTCGAGGCCGTTGACGCAGCGGATCAGCGTCGACTTGCCCGAGCCGCTGGCGCCGATGATCA
CCACCACCTCGCCGGGGGCGACCTCGAGGTCGATGTCCTGCATCACGTGCAGGCTGCCGAAGTACTTGTTGACCTTGTCCAGGCGCACGATGGGCTGGAC
GGTCTCGCTGTGGGGCTTGTCGTTCATGGCGTTGTCCTCACTGACCGACCAGGCCACGGCGCTCGAGCAGGCGCAGGGCGAAGGAAAGACTCAGGGTGAT
CGCCAGATAGAGCATGGCGACCATGAAGTACACCTCGAGGGCGGTGAAGGTCGTGGCGATGTAGATCTGGCCCTGGCGAACCAGCTCGCCGACACCGATC

>Modicisalibacter tunisiensis LIT2 [NZ_JAGXFD010000001.1:c907721..907791], dnaA> ig dnaN>, 71 bp

WebSkew GCmin pos. 106200 (GCmin<>oriC1 31.2 kb; GCmin<>oriC2 8.8 kb; oriC2<>oriC1 22.4 kb)

>Halomonas coralii 362.1 [NZ_QWBW01000012.1:137172..137671], <rpmH oriC1 dnaA>, 567 bp

TGTCGCCGCCACGGGGGTCGATACGCACATCGAGCACATCGGTACTGACCTTGATCAGGCTGCGCCGATTGTCGGTCGCGGCCTCGTCGGCCATGGCCGG
GCTGGTGCCGTCGGTCGCCGGCTTGGCCTGGTTCGACGGGGCCGCCAGGCCCTGTTCGTCGTCGTTGCTCTGGTCACTCGCGTCAGGGGTGGCGCTGTCG
CTGAAGGACGGTGCCTGGGTCTCGACGGGGCTCTGGCCGTAGTCCTGGTTCCACTGGATGACCAGCAGGTAGGCGAGAACCGCGAGCGGGATCACGAGTA
GCAGTCGTTTTACGTCCATTAGGGATCTGCCCGATGGGATGTTGGATCATTCTGGTGACGTGCGACGCCAAGCAGTGCCGGTCAGGAGGATGAGGCAACG
GCCGAATGCTTGCGAACCTCCTTTTCCAGGCGTCGCCACAAGCCGTGCAGCTGACGATGCAACGTGGGGTTATCCAGGTCGCTGACGCCACGGCGGGCGA
GGACCACGATATCCACGTCGGGCAGACGATGCTGCCGAAGACGGATGGATTCACGCACCAGACGCTTCAAACGGTTGCGATCGACGGCACGACGAACATT
CTTCTTGCTCACGACCAGCCCGACGCGAGGATGGCCCAGCGCGTTGGATCTGGCCAGGGCCAGGATCCCTCTGGCATGGACCTTGTACGTGGCGTGATCG
AAAACGTAGCGATAGTCCCCGGCGTTCAGCAGACGCAATTGCCGGGGAAAGCCCTGACAGGACATTCGGGGCCGATCAAGCGCTCAGACGCTTGCGGCCC
TTGGCACGACGGCGCGCCAGGATCTGACGACCATTCTTGGTAGCCATGCGAGCACGGAAGCCGTGCACACGCTTGCGCTTCAGAACGCTGGGTTGAAAAG
TGCGTTTCATAACGGCGGTTTCCCACGTGATGGATTGGATTGCCAGGTGCCGGGCCCCGCGAAGGGAACGCTGGGCGGATTCGGCTCAAGGCCGGGAATT

CTAGTGAATTCGCGGGCCGCGTGCAATTTTTGCGCGCGATTATCCACACCCGGTTTTCGACGCTCGGCCATCAGCGGTTGCGGGGCCGTGTGTCTCGCGA
CTTCGCCGGTGCCCGTGGCGGGAAGATTCATCCTGGCAGGCAGTCACTACAATAACTTTTCATTTTAAATGATGTTGTTATTACTATTGGGGATACTTTT
**TGTGGATAA**CCAGGTTTTACCCTTTCGTATCAAATGCTTGGTCTATTGACACCTGCGTTGACAAGATGCCCGGAAGCTGTCGGCGGCCTGCGTTGGGCCT
GTGGATGAAAGCGACGGTTGTCCACCGGGTCCGGGTTGTTCAGGCTGTGTACCCCATTCGTCAACCGTCCTGTACGTCAGTTTTCCACAGCATGCGGGCA
ACCCCCGCTCGTGTGGATAACTCGACCTCGGGGCGCTACAATGCCCGGTCATTCGCAAACTGGGATGGATCGCTCGCGTGTCGGTCGCTCTTTGGCAACA

ATGTCTGGATTACCTGCAGGACGAACTGAGTTCACAGCAGTTCAATACCTGGATCCGCCCGCTCCAGGCGGAGGATACGTCGTCCAACGAGCTCAGCCTG
CTGGCACCCAATCGCTTCGTGCGTGACTGGGTCAGCGACAAGTACCTCAAGCGAATCAACGAGCTGCTCCGGGAACTGGCACCCGCCAAGCCACCCAAGG
TGTCGCTGGCGGTGGGCAGTCGGCGTGCCGCGCCCAGCCCGCAGCCGCGGGAGCTGGGCAACCCGGTTTCCGCGAGCCGCTTTCGTCAGCCGCCTGCGCC
GGCGGTACAGACGCCGCCGCGAGGCGATTACGCCGACGAGCAGGAGATCGAGCAACTGCGTGACGAGACCGCCAGGCGTGCCTCGCCGCCGGCGGAGCGC
CAGGTTCAAGTCGAGGGCAGCCTCAAGCACCAGAGCGGGCTCAATCCCAACTTCACCTTCGACACCTTCGTCGAGGGCAAGTCGAACCAGCTGGCCCGGG
CGGCATCCCGGCAGGTCTCCGAGAATCCGGGGGGCGCCTACAATCCGCTGTTTCTGTATGGTGGCGTGGGTCTGGGCAAGACCCACCTGATGCATGCGGT
GGGCAATCAGCTGGCGCTGCAGCGCGACAATGCGCGAGTGGTCTACCTGCACTCCGAGCGCTTCGTCGCCGACATGGTCAAGGCGCTGCAACTCAATGCC
ATCAATGATTTCAAGCGTTTCTATCGCAGCGTCGATGCCTTGCTGATCGATGACATCCAGTTCTTCGCCGGCAAGGAACGTTCCCAGGAGGAGTTCTTCC
ACACCTTCAACGCCCTGCTCGAGGGCGGCCAGCAGATGATTCTCACCTCGGATCGCTACCCGAAGGAGATCAGCGGGGTGGAAGAGCGCCTCAAGTCGCG
TTTCGGCTGGGGGCTGACGGTGGCGATCGAGCCACCCGAGCTCGAGACACGGGTGGCGATCCTGATGAAGAAGGCCGATCAGGCCAAGGTGGACCTGCCG

>Halomonas coralii 362.1 [NZ_QWBW01000012.1:114733..115232], <mnmG oriC2 <COG1126 (glnQ), 564 bp

TCCAGCCCCTCGATCGAGCGCACCACCTGCAGCTGCACGTCGAACGGCAGCGAGGTCGAGATACCGTTGGGGTAGAGCTCGTGGGTGTCGAGGCCTTCCG
GTTCGATGAAGACCTGATGGCTCGACTTGTCGGCGAAGCGATGGACCTTGTCCTCGATCGACGGACAGTAGCGCGGCCCCACGCCGCCGATCACCCCGGA
GTACATCGGCGAGCGATCGAGATTGGCGAAGATGATCTCGTGGGTGCGCGCGTTGGTATGCGCGATGCGGCAACTGACCTGACGCGGATGCATGTCCCGC
GAGCCGAGATACGACATCACCGGCGACGGCGCATCGCCGGGCTGCTCCTCGAGGCGGTCGAAGTCGATGCTCTTGGCATCGAGCCGCGGCGGCGTACCGG
TCTTGAGGCGATCGACCCGGAACGGCAGGGCCCGCAGCCGCTGGGCCAGGGCATTGGCGGGCGGATCGCCGGCCCGTCCGCCAGTGTGGTTGTCCAGCCC
GATGTGGATAACCCCGCCGAGGAAGGTGCCTGTGCACAACACGACCGTCTCGGCGTGAAAGCGGATCCCGGTCTGGGTCACGGCGCCCTTGACGGTGTCG
TTCTCGACGATCAGGTCATCCACACCCTGCTGGAACAGCGTGAGATTGGGCTGGTTCTCCAGCACACCGCGAATCGCCGCCTTGTAGCGCACGCGGTCGG
CCTGGGCGCGGGTGGCGCGCACGGCCGGCCCCTTGCGGGCGTTGAGGACGCGAAACTGGATGCCGGCGCGATCGGTGGCCAGAGCCATGGCGCCACCCAG
GGCATCGATCTCCTTGACCAGATGGCTCTTGCCGATGCCTCCAATCGCCGGATTGCAGGACATCTGGCCCAGTGTCTCGATGTTGTGCGTCAGCAACAGG
GTCTGACAGCCCATGCGGGCGGCGGCCAGTGCGGCTTCCGTTCCCGCATGGCCACCGCCGATGACGATGACGTCAAAGCGGTTGGGATAATCCACGGTGT

ACCTCGACTTGCGCCTGCGGCGCGTCGGTTTCATCAGGCAAGTAGCGACAAGCCGCACAGTATAACGTCCCTGTGGGTAACCGTCAGGGGTTTTCCACAC
CGTGGCCAAGCCGAGCGTTTTGCGCCTCGCTGAATAAATACAATGTAATGAAGAAAAGAGAAGTTCTGTTGTGGTTGTGACTACTGGTGTGGACATTTTT
**TGTGGATAA**GCGGTTTTTCCACCAACTTTTCAATTCGTTGCAATGTTGACGATGCGTTGCAAAAAAGGTCGACGACATGCGGACAGGCTGCGTCGGGCCT
GTGGATGAAAGAGGGCCTTGTCCACAGCGCCGCCGATCCCCGGCTTATCCACCGCTGCATACCGGGGCTGTGCGTGATGTCGTCAACATCGGGTGTCGAG
GCGTTTCAGGCCGACAACCGTGGGGGTTGTGGAAGATTCGAGGTGGATTGGCCGAAACTCTTGTCCACAGGCCCTCGGGCGCGGGTGTGAACCTCCGGGA

GCCGGCCGAAGAGCGGGCGGGGGAGCCGCCCCGCGCCGGGAACGACGCGGGGCAGGGGGTCAGTGCTTGAGGACGCGGGCGAGAAAGGACTGGGTGCGTT
CGTGTTCGGGACTGTCGAAGACGCGTTCGGGCGGGCCCTGTTCGACGATCCGGCCCTTGTGGATGAAGATCACCCGATCGGCGACCTCGCGGGCGAAGCC
CATCTCGTGGGTGACGATCATCATCGTCATGCCATCGTTGGCGAGCTCTCGCATGGCGTCCAGGACCTCGCCGATCATCTCCGGGTCGAGCGCCGAGGTC
GGCTCGTCGAAGAGCATCAGGCGCGGTTCCATGGCCAGCGCGCGGGCCAGGGCGACGCGTTGCTGCTGTCCGCCGGAGAGCTGGCCCGGGTACTTGTCGG
CCTGGTCGGCGATCCCCACCCGCTCGAGCAGCCGTTCGGCGGTTTCCACGGCCTGGCCGCGGCTGATGTTGCGTACCTTGAGCGGCGCGAGGGTGACGTT
GTCGCGTACCGAGAGGTGTGGAAACAGGTTGAACTGCTGAAACACCATGCCGACCTCGGTGCGGATGGTCTGCAGCGACCGGCCGTTCTTGCCGTAGGGC
AAAAGCTGGTTGCCGTCGACCTCGATCCGGCCGCTCTGGAACTCCTCGAGGCCGTTGACGCAACGGATCAGCGTCGACTTGCCCGAGCCGCTGGCCCCGA
TGATCACCACCACCTCCCCGGGGGCGACCGCGAGGTCGATGTCCTGCAACACGTGCAGACTGCCGAAATGCTTGTTGACCTTGTCGAGACGCACGATGGG
GTCATTCTGGCCCGCCAGGGTCTCTTGGGCGCGTTGCTTATCGTTCATCGCCTTGGCTCCTCATTGACCGACCAGGCCGCGACGCTCGAGCAGGCGCAGC
GCGAAGGACAGACTCAGGGTGATCGCCAGGTAGAGCATGGCGACCATGAAATACACCTCGAGGGCGGTGAAGGTGGTGGCGATGTAGATCTGGCCCTGAC

>Halomonas coralii 362.1 [NZ_QWBW01000012.1:139122..139185], dnaA> ig dnaN>, 64 bp

WebSkew GCmin pos. 3237125 (GCmin<>oriC1 185.7 kb; GCmin<>oriC2 96.9 kb; oriC2<>oriC1 88.9 kb)

>Halomonas zincidurans B6 [NZ_JNCK01000001.1:c3051117..3051616], <rpmH oriC1 dnaA>, 569 bp

ATCGCCGCCACGCGGGTCGATACGCACATCGAGGGCATCGGTCGTCACCCTGATCAGGCTGCGGCGATTGTCGTCCTGAGCGCTCTCGGCCATGTCCGGG
CTGTTGCCGGCCTGCTTGGCGTCACTGGAAGGCGCCGCCAGGGCCTGCTCGGTCTCGGTGTTGGTGGTGTTATCCGTATCGCGGACGTTCGACTGGCTGT
AGGAAGGGGCTTCAGTCTCGACGGCGGTCTGGCCATAATCCTCATTCCACTGGATCACCAAGAGATAGGCGAGTACCGCGAGCGGGATGACGAGGAGGAG
TCGTTTTACGTCCATTAGGGATCTGCCCGATGGGATATTGGATCATTCTGCGGTTGGTCGGCACGCTGTTCCGGTTCAGGAGGTTGAAGCCATAGCTGAC
TGCTTGCCAGCCTCCTTTTCAAGACGTCGCCACAAGCCGTGAAGCTGGCGGTGCAATGTGGGATTGTCTAGCTCGTGTACGCCACGGCGGGCCAGGACAA
CGATATCCACAGCGGGTAGGCGATGCTGCCGGAGGCGAATGGATTCGCGCACCAAGCGCTTGAGACGGTTGCGATCTATGGCACGGCGAACGTTCTTCTT
GCTGAAGACCAGGCCCACGCGCGGATGATCCAGGGAGTTGTCTCTGGCCAACACCAGCAACCCTTTGGCGTGTACCTTGTACGAGGCGTGTTCGAAGACG
TACCGATAGTCCCCGGCAGTCAGCAGACGCAGTCGCCGGGGAAAGCCCTGACCGGACATTCGCGAGCCGCGATCAAGCGCTCAGACGCTTGCGGCCCTTG
GCACGGCGGCGAGCCAGGACCTGGCGGCCGTTCTTGGTGGCCATGCGTGCACGGAAGCCGTGGACACGCTTGCGCTTCAGTACGCTGGGTTGAAATGTGC
GTTTCATGACGGCGAATTCCCACGTGGTTGATTTGACATTGACCGATGCCCCGGCTCCCGCATGGAACCCGGGCGGATTTCGTTCAAGGCCGGGAATTCT

AGAGGCTTTGCAAGCCGGGTGCAATTTCTGCGCGCAATTCTCCACACCGCCTTGTCGCTCGCGGCGTTGACGATAGCCGGCGAATGCCTGCGGTGCCGGT
GCATCGCGGATTGGGCACTCGCTATGCGATGCTGCCGCGCCACTTACAATAATTACAGGTGTCATTGGAAGTTGTTGTTATTGATAGTAGGGGTGGTTTC
**TGTTTATAA**CCTTATTTAATTTTTATAAATCAAATAGTTAATATAAAAGCAAGGGTGTGGACAAAGTCCGCGGATCCAGTGGCTGGAGGGCGTCAAGCT
GTGTACCGATTGCTGGAATATCCACAGCCGAGACGCATCCGCACAGTCATACCGTATCGCTCCACCCAGCCATACTGTAGTTTTCCACAGCATGGCTCGC
ATCGTTTCGCGTGTGGATAACTTGCGGTGAGGGCGATACAATGTTCGGTCTTTTCGCAACGGGATGGTGAGGTCGCGTGTCGGTCGCTCTTTGGCAACAA

TGTCTGGCTTATCTGCAGGATGAGCTGAGCTCGCAGCAGTTCAATACCTGGATTCGCCCCCTGCAGGCGGAAGACGCAACGCCTAACGAGCTCAGTCTGC
TGGCGCCCAATCGTTTCGTGCGTGACTGGGTCAGCGACAAGTACCTCAAACGGATCAATGAACTGCTGCGCGAGCTGGCGCCCAACAAGCCGCCCAAGGT
GAGCTTGGCAGTGGGTAGCCGGCGCGTTGCACCGAGTCCGCAACCGCGGGATCTGGGCAACCCGGTCTCGGCCAGTCGTTTTCGTCAGCCTCCCGCACCG
GCAGTGCATACCCCGTCCCGCGGCGACGAGTACGCCGAGGAGCGCGAAATCGACAGGCACAATGGCGAAGGCCCGCGGCGTTCCACTCAGCAGCCCGCGG
GCGAGCGTCAGGTTCAGGTGGAAGGCAGCCTCAAGCACCAGAGCGGACTCAACCCCAACTTCACCTTCGACACCTTCGTCGAGGGCAAGTCCAACCAGCT
GGCCAGGGCCGCTTCGCGGCAGGTGTCGGAGAACCCTGGCGGTGCCTACAATCCACTGTTCCTGTACGGTGGGTAGGGCTGGGCAAGACCCACCTGATG
CACGCCGTGGGCAACCAGCTGGCTATCCTGCGCGACAATGCCAAGGTGGTCTATCTGCACTCCGAGCGCTTCGTCGCCGACATGGTCAAGGCGCTGCAGC
TCAACGCCATCAATGATTTCAAGCGGTTTTACCGAAGCGTCGATGCGCTATTGATCGACGATATTCAGTTCTTCGCCGGCAAGGAGCGCTCTCAGGAAGA
ATTTTTCCATACTTTCAATGCTTTGCTGGAAGGCGGTCAGCAGATGATCCTGACCTCCGACCGCTACCCTAAGGAAATCAGCGGGGTGGAGGAACGGCTC
AAGTCGCGTTTCGGCTGGGGCCTGACGGTGGCAATCGAACCGCCCGAACTGGAAACGCGGGTGGCGATCCTGATGAAGAAGGCCGATCAGGCAAAGGTCG

>Halomonas zincidurans B6 [NZ_JNCK01000001.1:c3139984..3140483], <mnmG oriC2 <COG1126 (glnQ), 587 bp

GTTTTCCAGCCCCCTGATCGAGCGCACCACCTCGAGCTGCACGTCGAACGGCAGCGAGGTGGATATGCCGTTGGGGTAGAGCTCGTTGGTCTCCAGTCCC
TCGGGTTCGATGAATACCTGATGGCTCGCCTTGTCGGCGAAGCGATGGACCTTGTCCTCGATCGACGGGCAATAGCGCGGCCCGACGCCCTCGATCACGC
CGGAATACATCGGCGAGCGGTCGAGGTTGGCGAAGATGATCTCGTGGGTGTGCGCGTTGGTATGGGCGATGTGACAACTCACCTGACGCGGATGCATCGC
GTGCGAACCCATGTAGGACATTAGCGGGCTAGGCGCGTCGCCGGGCTGCTCCTCGAGGACGCTGAAGTCGACGCTCTTGGCATCGATGCGCGGCGGTGTG
CCGGTCTTGAGCCGATCGACACGAAACGGCAGTGCCCGCAGGCGTTGTGCCAGGGCATTGGCCGGCGGGTCGCCGGCACGGCCGCCGGTATGGGTATCCA
GACCGATGTGGATGATGCCGCCGAGGAAGGTGCCGGTACACAGCACCACGGTTTCACCGAGAAAGCGGATTCCGGTCTGGGTCACCGCGCCGCGGACAGT
GTCGCCGTCGACGATCAGGTCATCCACCCCCTGCTGGAACAGCGTCAGGTTGGGCTGGTTCTCCAGAAGGCCGCGAATCGACGCCTTGTAGCGAACACGG
TCGGCCTGGGCACGCGTGGCGCGCACCGCCGGTCCCTTGCGGGCATTGAGTACCCGGAACTGGATACCGGCCTGGTCGGTTGCCAGCCCCATCGCGCCGC
CCAGGGCGTCGATCTCCTTGACCAGGTGGCTCTTGCCGATGCCGCCGATGGCCGGATTGCAGGACATCTGTCCGAGCGTCTCGATGTTGTGCGTCAGCAG
CAGCGTCTGGCAACCCATGCGGGCAGCGGCCAATGCGGCTTCCGTTCCCGCATGACCACCGCCGATGACGATGACATCGAAGCGGTTGGGATAATCCACG

TTTTTACCTCGACTTGCGCGGTGCGCGTACGTTCTGGCGAGTGTGATGAAGCGCGCCAGTATAACGTCCCTGTGGGTAACCGTCAGGGGTTTTCCACACC
GCCTTGGACACAACGCGCTGAGCGTACGACTATCAATAACAACGTAGATATATAAAGAAGAAGTTCTGTTGTGGTTGTTACTACTGGTGTGGACATTTTT
**TGTGGAAAA**GTACTTTTAAAGTAGCAAAAACAAAAATTTGCATTGTTGATGGTGCATTGCAAAACAAGTCGACCGGAACGGTACAGGCTGTCGTGAGACT
GTGGATGGATGGCCCGGATATCCACAGCCGAGATTGTACCCCGTTTATCCACCGATACGTACCGGGGCATTGCCGGTAATGATCAACAGCCCTGTCTCGA
AGTGTGGAAACCGCCTGCGATACGGCTTGCGGCCCGATTCGAGGCGGCTGGCAGAAAATCTTATCCACAAGCCAAAAAAGCCGCTGTGGGCGGCTTCTTG

GTGAGGTGGATTCTGGAAATCGAGGGAGTCGCTGCGTCCAGGCAGGCGAACGATTGCTCATTGCCTGCCTGGGCAAAGTCGAACGGTCAGTGCTTGAGCA
CTCGCGCGAGGAAGGATTGCGTACGTTCGTGGCGGGGGTTGTCGAAGATCTCGTCCGGCGGGCCCTGCTCGACGATGCGGCCGCCATGGACGAAGATCAC
CCGGTCGGCGACTTCCCGGGCGAAGCCCATCTCGTGAGTGACGATCATCATCGTCATACCCTCGTTGGCCAGCTCGCGCATGGCATCGAGCACTTCACCG
ATCATCTCCGGGTCGAGCGCCGAGGTCGGCTCGTCGAACAGCATCAAGCGCGGTTCCATGGCCAGCGCCCGGGCCAGGGCGACGCGCTGCTGCTGGCCGC
CGGAGAGCTGGCTGGGATACTTGTCGGCCTGATCGCTGATGCCGACCCGCTCGAGCAGCCGGTCGGCGATCTCGGTCGAATGCTGGCGATTCCAGCCACG
TACCTTGAGCGGTGCCAGGGTGACGTTGTCGCGCACCGACAGATGCGGAAACAGGTTGAACTGCTGGAAGACCATGCCGACTTCGGTGCGAATGGTCTGT
AGCGACCTGCCGCTCTTGCCGTAGGGCAACAGCTGATTGCCGTCGACTTCGATGGCGCCGCTCTGGAACTCCTCGAGACCGTTGACGCAGCGAATCAGGG
TCGACTTGCCGGAGCCGCTGGCACCGACGATCACCACCACCTCGCCGGGGGCCACCTCGAGATCGATGTCCTGCATGACGTGCAGCGAACCGAAATGCTT
GTTGACCTTGTCCATCTTGACGATGGGCGCCTTGTGGAGGGCCTCGTGACGCTGTGAATCGTTCATGGCGTGTTCCTTATTGGCCGACCAGACCGCGGCG
CTCGAGCAGCCGCAGGGCGAATGACAGGCTCAGAGTGATCGCCAGATACATCAGCGCGACCATGAAGTAGACCTCCAGCGCAGTGAAAGTGGTGGCAATG

>Halomonas zincidurans B6 [NZ_JNCK01000001.1:c3049599..3049655], dnaA> ig dnaN>, 57 bp

WebSkew GCmin pos. 77742 (GCmin<>oriC1 28.4 kb; GCmin<>oriC2 6.0 kb; oriC2<>oriC1 22.4 kb)

>Halomonas radicis EAR18 [NZ_CAAHFN010000048.1:c49066..49565], <rpmH oriC1 dnaA>, 572 bp

TGTCGCCGCCACGTGGGTCGATACGCACATCGAGGGTATCGGTCACCACTTCGATCAGGCTGCGACGATTGCCGTCCTGGTCGCTTTCGGCCATGCCCGG
ATTGTCGCTCGGCGCTTGGCTGCCGCCGGTGGGAGCCGCCAGGCCCTGCTCGGTATCGGCTGAGCCGTCCGCATCGCCGCTGTTCGACTGGCTATAGGTG
GGGACTTCGGTCTCGACGGCGGTCTGGCCGTAATCCTCATTCCACTGGACCACCAGGAGATAGGCGAGTACCGCAAGTGGAATGACGAGGAGGAGTCGTT
TTACGTCCATTAGGGATCTGCCCGATGGGATATTGGATCGTTCTGCGGGTTGGTCGGGTCACGCTGTTCCGAATTCAGGAGGCCGAGATGGCAGGTGACT
GCTTGCCAACCTCCTTTTCCAGCCGTCGCCACAAGCCGTGAAGCTGTCGGTGCAGTGTGGGATTGTCCAGCTCGTGTACGCCACGGCGGGCCAGGACAAC
GATATCCACAGCGGGTAGGCGATGCTGCCGGAGACGGATGGACTCACGCACCAAGCGCTTTAGACGGTTGCGATCTACGGCACGACGAACGTTTTTCTTG
CTGAAGACCAGGCCCACGCGCGGGTGACCCAAGGAGTTGTCCCTGGCCAGTACCAGCAACCCTTTGGCGTGTACCTTGTACGATGCTTGTTCGAAGACGT
ACCCATAGTCCCCGGCAGTCAGCAGACGCAGTTGCCGGGGAAAGCCCTGACAGGACATTCGCGAGCCGCGATCAAGCGCTCAGACGCTTGCGGCCCTTGG
CACGGCGGCGAGCCAGGACCTGGCGGCCGTTCTTGGTGGCCATGCGTGCACGGAAGCCGTGGACACGCTTGCGCTTCAATACGCTGGGTTGAAATGTGCG
TTTCATGACGGCGGATTCCCACGTGGTTGGTTTGACATTGATCGATGCCCCGGCTCCCACATGGAACCCGGGCGGATTTCGTTCAAGGCCGGGAATTCTA

GAGGCTTTGCAAGCCGGGTGCAATTTCTGCGCGCAATTATCCACACCGCTGCCGGCGTGGTGCGGCGATGCGCCGCTGGCGGAGCTTCGCCGCATCGAGG
TGTCGAGCGACCCGGAGCTCGACGGCCAGGGGCTATGGGCTTACTACAATAACTAGAAGGTTTATTTAATGCTGTTGTTATTAATGCTGGGGACCTTTTC
**TGTGGATAA**CCTTGTAAAAACATTACGCATCAAAACACTACATCCAGTATAAAGGCGTGGACAACGTGCGTAGAGGCGGTGATTCAAGGGTCATGAAGCT
GTGCATGGAAAGCCGGGATATCCACCGCCAAGGGTTGTCCACGCAATCATACCGGGCCGGTCCCCCGAGGCGTACCCTAGTTTTCCACAGCACCGTTCGA
ACCGCCTTGCGTGTGGATAACTCGTAGTGAGAACGCTACAATGGCCGGTCTTTTTCGCAAACGGGATGGTGAGGTCGCGTGTCGGTCGCTCTTTGGCAAC

AATGTCTGGCTTATCTGCAGGATGAGCTGAGTTCGCAGCAGTTCAATACCTGGATTCGTCCCCTGCAGGCGGAAGACGCTACCCCCAACGAGCTCAACCT
GTTGGCGCCCAATCGCTTTGTACGCGACTGGGTCAGCGACAAGTACCTCAAGCGGATCAATGAGTTGTTGCGCGAACTGGCGCCCAGCAAGCCGCCCAAG
GTGAGCCTGTCGGTGGGCAGTCGGCGTGCTGCGCCGAGCCCGCAGCCGCGGGACCTGGGTAACCCGGTCTCGGCCAGCCGCTTTCGCCAGCCTTCCGCAC
CCGCGGTGCACACGCCGCCCCGGGGTGACGAATACGCCGATGAGCGGGAGATCGACAAGCTCAGCGGCGAGGGCCAGCGCCGTTCCTCACAGCAGCCCGC
GGGGGAGCGGCAGGTCCAGGTGGAAGGCAGCCTCAAGCACCAGAGCGGTCTCAACCCCAACTTCACGTTCGACACCTTCGTCGAGGGTAAGTCCAACCAG
CTTGCCAGGGCCGCTTCGCGACAAGTCTCCGAGAACCCCGGCGGCGCTTACAATCCACTGTTCCTGTATGGTGGGGTGGGCTTGGGCAAGACCCACCTGA
TGCACGCCGTGGGCAACCAACTGGCTGCCCTGCGCGACAATGCCAAGGTGGTCTATCTGCATTCCGAACGCTTCGTCGCCGACATGGTCAAGGCGCTTCA
GCTCAACGCCATCAACGACTTCAAGCGGTTCTATCGAAGCGTCGATGCGCTGCTGATCGACGATATCCAGTTCTTTGCCGGCAAGGAGCGCTCCCAGGAG
GAGTTCTTTCATACTTTCAACGCCTTGCTGGAAGGCGGTCAGCAGATGATCCTGACCTCGGACCGCTACCCCAAGGAGATCAGCGGGGTGGAAGAGCGAC
TCAAGTCGCGTTTCGGCTGGGGGCTGACGGTCGCCATCGAACCTCCCGAACTGGAAACCCGGGTGGCTATCCTGATGAAGAAGGCCGACCAGGCCAAGGT

>Halomonas radicis EAR18 [NZ_CAAHFN010000048.1:c71463..71962], <mnmG oriC2 <COG1126 (glnQ), 638 bp

GTTTTCCAGCCCCTTGATCGAACGCACCACCTCGAGCTGCACGTCGAACGGCAACGAGGTGGAGATGCCGTTGGGGTAGAGTTCGTTGGTCTCCAGCCCC
TCGGGTTCGATGAATATCTGATGGCTCGCCTTGTCGGCGAAGCGATGAACCTTGTCCTCGATCGACGGACAGTAGCGCGGCCCGACGCCTTCGATCACTC
CGGAATACATCGGCGAGCGGTCGAGATTGGCGAAGATGATCTCGTGGGTACGCTCGTTGGTGTGAGCAATGTGACAACTCACCTGCCGCGGATGCATCTC
GCGGGTGCCCAGGTAGGACATCACCGGGGTGGGGGCGTCACCGTGTTGCTCCTCGAGGGTCGTGAAATCGACGCTCTTGGCATCGATACGCGGCGGCGTC
CCGGTCTTGAGTCGGTCGACACGAAACGGCAGTGCGCGCAGCCGCTGCGCCAGGGCATTGGCCGGCGGATCGCCGGCGCGACCGCCGGTATGGGTGTCGA
GACCGATGTGGATGACCCCGCCGAGGAAGGTCCCGGTGCACAGCACCACGGTCTCGGCATGGAAGCGAATTCCGGTCTGGGTCACCGCGCCGCAGACCGT
ATCGCCGTCGACGATCAAGTCATCCACACCCTGTTGGAACAGCGTCAGGTTGGGCTGGTTCTCGAGCATGCCGCGAATGGCTGCCTTGTAGCGTACACGG
TCGGCCTGAGCCCGGGTCGCGCGAACGGCAGGCCCCTTGCGGGCGTTGAGCACTCGGAACTGGATACCGGCCCGATCGGTTGCCAGCCCCATTGCGCCAC
CCAGGGCATCGATCTCCTTGACCAGGTGGCTCTTGCCGATGCCGCCAATGGCCGGATTGCAGGACATCTGTCCCAGGGTCTCGATGTTGTGCGTCAGCAG
TAGTGTCTGGCACCCCATGCGAGCGGCGGCCAGTGCGGCTTCCGTTCCCGCATGGCCACCGCCGATGACGATGACATCGAAGCGGTTGGGATAATCCACG

TTGTTACCTCGACTTGCGCGATGCGCGTGCGTTTCTGGCGAATGTGATCAAGCGCGCCAGTATAACGTCCCTGTGGGTAACCGTCAGGGGTTTTCCACAC
ACCCCGGCGGACGCTGGGTCACCGAATCACTATCAATAATAAAGTAAGTATATAAAGAAGAAGTTCTGTTGTTGTTATTACTACTAGTGTGGACATTTTT
**TGTGGAAAA**GTGGTTTTAGTTTAATGGAATCATAAATTTGCATTGTTGATGGTGCATTGCAAAACGAGTCGACGGCGTCTGTATGGACGGTCGTCAGGCT
GTGGACGAAAGAGCGTCCTATCCACAGCTGGACTTGTGCCCCGCTTATCCACCGTCGAGTACCTGGGCTTTGCCGGTATCGATCAACAGCCGGTCGTCGA
AGGGTGGAAGGCGCCTGTGACGCGGGCTGTGGAAAGATCGTCAGCGTCGCGGAAAAATAATTATCCACAGGCGAAAAAAGCCGCTGTGGGCGGCTTTTCG

TACGAGGCGGGAGTCGAACCGGGTGCCCGGGAGCGTCTGCCGAGTCGACGCTCCCGGCACGCTTCGGACAAGGCCCGAAGTAAGGACAGGCATACTTGTC
ACCGCCTGGACGGGCGGTAACCAGAACAAGGGCTCCGTCAGTGCTTGAGGACCCGCGCGAGGAAGGATTGGGTGCGCTCGTGGCGGGGATTGTCGAAGAT
CTCGCCCGGCGGGCCCTGTTCGACGATACGCCCGCCGTGAACGAAGATCACCCGATCGGCGACTTCGCGGGCGAAGCCCATCTCGTGGGTGACGATCATC
ATCGTCATGCCCTCGTTGGCCAGCTCGCGCATGGCATCGAGCACTTCGCCGATCATTTCCGGATCGAGTGCCGAGGTCGGCTCGTCGAACAGCATCAGCC
GCGGTTCCATGGCCAGCGCCCGGGCCAGGGCGACCCGCTGCTGCTGGCCGCCGGAGAGCTGATTGGGGTACTTGTCGGCCTGGTCCGTGATGCCGACCCT
TTCGAGCAGCCGGTCGGCGATCTCGGTCGAACGCCGGCGATCCCAGCCGCGCACCTTGAGCGGCGCCAGGGTGACATTGTCGCGCACCGAGAGGTGGGGA
AACAGATTGAACTGCTGAAAGACCATGCCGACTTCGGTGCGGATGGTCTGCAGCGACTTGCCGCTCTTGCCGTAGGGCAGCAACTGATTGCCGTCGACCT
CGATGGCCCCGCTCTGGAACTCCTCGAGGCCGTTGACGCAGCGGATCAGGGTCGATTTGCCCGAGCCGCTGGCACCGACGATCACCACCACCTCGCCGGT
GGCCACCTCGAGGTCGATGTCCTGCATCACATGCAGCGAACCGAAATGCTTGTTGACCTTGTCCATCCTGACCATGGGCGCCTTGTCGGCGGTCCGGGCG
GCGTCGTGACGCTGTGAATCGTTCATGGCGTGCTCCTCATTGACCGACCAGGCCGCGGCGTTCGAGCAACCGCAGGGCAAAAGACAGACTCAGCGTGATC

>Halomonas radicis EAR18 [NZ_CAAHFN010000048.1:c47545..47605], dnaA> ig dnaN>, 61 bp

< page 1 | #122 | Halomonas lutea DSM 23508

WebSkew GCmin pos. 259424 (GCmin<>oriC1 24.8 kb; GCmin<>oriC2 6.0 kb; oriC2<>oriC1 18.8 kb)

>Halomonas lutea DSM 23508 [NZ_KB899996.1:c234336..234835], <rpmH oriC1 dnaA>, 570 bp

AGCACGTCGGTGACGACCTGGACCAGGCTGCGACGATTGTCGGTAGCGTTCTCGCCCGGCATAGTGCTGCTGGCGGGGCTAGCCGTGGCGTCCGATGGTG
CCGCCAGGCCCTCGTCATCCGTCGCGGATGCGGCATTACCGCTACGTGGCGCTTCGCTGACAGTGGGAGCGCTGGGAACATTGACGTCAGGCTGGCCGTA
ATCCTGATTCCATTGGATTACCAGCAGGTAGGCGAGTACCGCTAGCGGTATCACGAGAAGAAGTCGTTTTACATCCATGGGGATCTGCCCGGTGGGGTGA
TGGATCAATCTGGCGGCTTCGTTCAGCACCGATGACCGGAGTGGGCATGACGAGGCCGCGCTGCTGCTATCCGGTCAAGAGAGTGAGGAGGCGACCATCT
GCTTGCGGACGTCCTTCTCCAAGCGACGCCACATGCCATAAAGCTGGCGATGCAGTGTCTCGTTGTCCAGATCGTTAACGCCGCGGCGTGCCAAAACGAC
GATATCCACAGCGGGTAGCCGATGCTGCTGCAGACGAATGGATTCGCGCACGAGGCGTTTCAGGCGGTTGCGGTCGACGGCCCGACGAACGTTCTTCTTA
CTGAATACCAGCCCCACCCGTGGATATCCCAGCGCATTGTCACTGGCCAGTGCCAGCAACCCTTTGCCATGTATCTTGTACGAGGCCTGGTCGAAGACTC
GTCGATATTCCCCGGCGGTCAGCAGACGCAGCTGCCGGGGAAATCCCTGACAGGACATTGGCAACCGGGATCAAGCGCTCAGACGCTTGCGACCCTTGGC
GCGACGACGCGCAATGACCTGGCGGCCGTTCTTGGTGGCCATGCGGGCACGAAAGCCGTGCACGCGCTTGCGCTTCAGAACGCTGGGCTGAAAAGTACGT
TTCATAGCTGCTGCTTCCCACGTGGTTAAGTGAACAAGAGGTCAAACGGAGTGCCCTGGCTCCGATCGGGAACCAGGGCGGATTCGGTTCAAGGTCGGGA

ATTCTAGTGAATTCGCAAGCCACGTGCAATTTTTACGCGAATTTATCCACACGTGGTTTGGGAATGCCCTGCCGTGAGTGTTTTCAACCGCATGTCGAAT
GACTGCATCAGCAATTAACGAAATCAGAAGATGCCTATCGAACCACTATAATAACGTTTGGTTTCTTACTGATGTTGTTATTAATACTGTGGGCAATATG
**TGTGGATAA**GTGATTAAAAGTCAGGCTT**C**TTAAGGGCTTATAGCAATAATATTACTGTGGAGGGCATGCTGATAGCCGGTGTGGGGCGCGGGGATAGCGA
TGTGGATAACTTCCATTCCATCCACAGGGTAGACTTGTCCTCTACTATATACCTCTCCTGTCCACTTGGCTGTGGGCAAGTTTTCCACAGGAAGAAATGC
CTTGTCGTCGTGTGGATAACTCGCCCCGAGAGCGATACAATCCCCGATCATTCGCTAACTGGATGGTGAGGTCGCGTGTCGGTCGCTCTTTGGCAACAAT

GTCTTGGCTTTCTGCAGGACGAGCTGAATTCTCAGCAGTTCAATACCTGGATTCGTCCTTTGCAGGCGGAGGAAGGGGATGCCAACGAGCTATGTCTGTT
GGCGCCCAATCGCTTTGTTCGGGACTGGGTCAGCGACAAGTACCTCAAGCGCATCAATGAGTTGTTGCGTGAGCTATCACCAGCCAAGCCCCCTAAGGTA
ACCCTGGCGGTAGGTAGCCGGCGTGCGGCGCCAAGCCCGCAGCCCCGCGATCTGGGCAGCCCGGTCTCGGCCAGTCGTTTCCCGCAGTCGGCTGCACCGG
CTGTACATACCTCTCCGCGCGGTGATTACGCTGACGAGCAGGAAATAGAGCAGCTACGCGAGGAAGCGCCTCGACGTGGTGCGTCGGGCGAGCGGCAGGT
GCAAGTCGAAGGTAGCCTCAAACATCAGAGCGGCCTCAACCCCAACTTCACTTTCGATACCTTCGTCGAGGGTAAGTCGAACCAGCTGGCGAGGGCCGCC
TCGCGACAGGTCTCGGAGAATCCTGGTGGCGCTTACAATCCATTGTTCCTTTACGGGGGCGTCGGCCTAGGTAAGACGCACCTCATGCATGCCGTTGGCA
ACCAGCTCGCCAAACAACGGGAAAATGCGAAGGTAATCTACCTGCACTCAGAGCGCTTCGTGGCGGATATGGTCAAGGCGCTGCAGCTCAATGCCATTAA
CGATTTCAAGCGTTTCTACCGAAGTGTCGACGCGCTATTGATCGACGACATCCAGTTCTTTGCCGGCAAGGAGCGCTCCCAGGAAGAGTTTTTCCATACC
TTCAATGCCCTGCTTGAGGGCGGCCAGCAGATGATCCTGACTTCCGACCGCTACCCTAAGGAAATCAGTGGAGTGGAAGAACGCCTGAAGTCACGCTTTG
GCTGGGGACTGACGGTAGCGATCGAGCCTCCGGAGTTGGAAACACGCGTGGCGATCCTGATGAAGAAGGCTGACCAGGCCAAAGTCGATCTGCCCCATGA

>Halomonas lutea DSM 23508 [NZ_KB899996.1:c253111..253610], <mnmG oriC2 <COG1126 (glnQ), 573 bp

CGTTTTCGAGTCCCTTGATCGAGCGAACCACCTGCAACTGCACGTCGAACGGCAGCGAGGTGGATATCCCGTTGGGGTAAAGCTCGTGGGTATCCAGGCC
CTCGGGCTCGATGAATACCTGATGGCTGGACTTGTCGGCAAAGCGATGGACCTTGTCCTCGATCGAAGGACAGTAGCGCGGCCCGACGCCTTCGATGACC
CCAGAATACATCGGCGAACGATCGAGGTTAGCGAAGATGATCTCGTGAGTGCGCTCGTTGGTGTGCGCAATATGGCAGCTGACCTGTCGTGGATGCATCT
CGCTCGAGCCGAGGTAGGACATTACCGGCGTTGGTGTATCGCCAGGCTGCTCCTCGAGCTCGGAGAAATCGACGCTCTTGGCATCGATGCGCGGTGGTGT
TCCGGTCTTGAGACGCGCAACACGAAAAGGCAACGCACGCAGCCGCTGAGCCAGCGCATTGGCCGGTGGATCACCGGCACGGCCACCGGCATGTTTGTCC
AACCCGATGTGGATAACTCCACCCAGGAAGGTGCCGGTACACAGCACCACGGTTTCGGCATGAAAGCGGATTCCAGTCTGGGTGATTGCGCCACGCACAG
TGTCGTTTTCGACGATCAGATCATCCACACCCTGTTGGAATAGGGTCAAGTTGGGCTGGTTTTCCAACATGCCGCGAATGGCTGCCTTGTAACGCACACG
ATCGGCTTGGGCACGGGTGGCTCGCACAGCAGGCCCCTTGCGTGCGTTGAGCACGCGGAACTGGATGCCGCCGCGGTCGGTGGCTAATGCCATGGCCCCG
CCGAGGGCATCAATTTCCTTGACGAGGTGGCTCTTGCCGATCCCGCCGATCGCCGGGTTGCAGGACATCTGCCCCAGCGTCTCGATGTTGTGTGTCAGTA
ACAGGGTCTGACAGCCCATGCGGGCAGCGGCCAGTGCGGCTTCTGTTCCCGCATGGCCACCGCCGATGACGATGACGTCAAAGCGGTTGGGGTAATCCAA

GATGCACCTCGACTTGCGCCTGTCGGCGCGGCGTTTCTGCTAGGTGAGGGTAAATAGCCGCCCAGTATAACGCCCCTGTGGGTAACCGTCAGGGGTTTTC
CACATGAGGTTTATCAGTCACGTGTTGTTATCTATTAATAAAAATAGAATTAAAGAGAAGAAGTTCTGTTGTGGTTGTTACTACTAGTGTGGATAGTTTT
**TGTGGATAA**CCTAAAAAAGTAAAAACTTTT**C**ATTAGGTTGCATTGTTAACCGAACGTGGGAAAAGCGGTAAGGTAGCTGCGGAAATCATGCCGTGAGGCT
GTGGATGGAATGACTCTTTATCCACACATGTATTTATGCCCCGTTTATCCACCTGTGCCTACCGTAGCTGTCATCAGGGTGATTAACAAAGTGGGTCAGG
AAGGAGAAAGTCGGTCGTGGCAGGGGGAGCAGGGCGATTAGCGGCCGATAATTCGAAAAATTATCCACAGGCCAGAAAAGTGGCTGTGCATAGCAAAAAA

GCCAGTGAGGCGGGAAATAAAAAAAGAACGCCCGCTTCAAGAGCGGGCGTTCGGTAGTAACAAGAGGTGCAGCTCAGTGCTTGAGCACGCGGGACAGGAA
CGATTGAGTTCGCTCGTGGCGAGGGTGGTCGAAGACCTCGTGGGGCGGCCCCTGTTCGACGATGCGACCACCATGTACGAAGATCACCCGATCGGCGACC
TCTCGGGCAAAGCCCATCTCATGGGTGACGATCATCATGGTCATGCCATCGTTGGCGAGCTCACGCATGGCGTCGAGTACCTCACCGATCATCTCCGGAT
CGAGCGCCGAGGTGGGCTCGTCGAAAAGCATCAGGCGGGGTTCCATGGCCAGGGCGCGTGCCAGCGCCACACGCTGCTGTTGGCCACCGGAGAGCTGACT
GGGATACTTTTCAGCCTGATCCTCGATGCCGACCCGGGCCAGCAGCTTGTGTGCAGTGTCTAGTGCCTTGTCGCGGGGCCAATTGCGGACCTTCATCGGT
GCCAGGGTGACATTGTCGAGTACCGACAGGTGCGGGAACAGGTTAAATTGCTGAAAGACCATGCCAACCTCGGTGCGGATGGTCTGCAGTGTCTTGCCGC
TCTTGCCGTTGGGCAGCAGTTCGTTGCCGTCTACCACGATACGACCACCTTGGAACTCTTCAAGGCCGTTGACACACCGAATCAGAGTAGACTTGCCGGA
GCCGCTGGCTCCGACGATGACGACCACTTCGCCGGGGGCCACCTCGAGGTCGATATCCTGCATAACGTGCAACTTGCCGAAGTGCTTGTTGACCTTGTCG
AGGCGGACGATAGGCTCATGGTCACCGGTCTTCGGCGAGGTATGGGTTACTTCATTCATGCGCTTTCTCCTCACTGTCCGACCAGGCCGCGGCGTTCGAT
CTGACGCAGGGCAAAGGACAGGCTCAGCGTGATTGCCAAGTACATCATGGCGACCATGAAATAAACTTCGAGGGCAGTGAAGGTGGTGGCGATGTAGATT

>Halomonas lutea DSM 23508 [NZ_KB899996.1:c232824..232890], dnaA> ig dnaN>, 67 bp

WebSkew GCmin pos. 7695 (GCmin<>oriC1 48.6 kb; GCmin<>oriC2 6.8 kb; oriC2<>oriC1 41.9 kb)

>Halomonas xianhensis CGMCC 1.6848 [NZ_FOPY01000006.1:56097..56596], <rpmH oriC1 dnaA>, 570 bp

GGACGTCGGTGACGACCTGGACAAGGCTGCGGCGGTTGTCGGCAGCGCCTTCGCCCGGCATGTTGCTGCTGGCCGGCGTCGCCGAGGCATCCGATGGTGC
CGCCAGGCCCTGATCATCCGCAGGGGATGTGTCGTTGCCGCCCTGGGGCGCTTCGCTGACAGTGGGGGCACTGGGGACGTTGACGTCAGGCTGGCCGTAA
TCCTGATTCCACTGGATTACCAGCAGGTAGGCGAGCACCGCTAGCGGTATCACGAGAAGAAGTCGTTTTACATCCATGGGGGATCTGCCCGGTGGGGTGA
TGGATCATTCTGGCGGCTTCGTTCAGCACCGATGGCCGGAGGGGGCATGGCGAGGCCGCGCTGTTGCTATCCGGTCAAGAGAGTGAGGAGGCTGCCGTTT
GCTTGCGGACATCCTTCTCCAGGCGACGCCACATGCCGTAAAGCTGGCGATGCAGTGTCTCGTTGTCCAGATCGTTGACGCCGCGGCGTGCCAAAACGAC
GATGTCCACAGCGGGCAAACGATGCTGCTGCAGACGAATGGATTCGCGCACGAGGCGTTTCAGGCGGTTGCGATCGACGGCACGACGTACGTTCTTCTTG
CTGAATACCAGCCCCACCCGCGGATATCCCAGCGCGTTGTCACTGGCCAGTGCCAGCAACCCCTTGCCATGGATCTTGTAGGTGGCCTGGTCGAAGACGC
GCCGATATTCCCCGGCGGTCAGCAGACGCAGCTGCCGGGGAAATCCCTGACAGGACATTCGCAACCGGGATCAAGCGCTCAGACGCTTGCGACCCTTGGC
GCGACGACGCGCAATGACCTGGCGGCCGTTCTTGGTGGCCATACGAGCACGGAAACCGTGCACGCGCTTGCGCTTCAGTACGCTGGGCTGAAAAGTGCGT
TTCATAGTTGCTGCTTCCCACGTGATTGAGTGAACGAGAGGTAAAACGGAGTGCCCCGATTCCAACCTGGAATCGGGGCGGATTCGGTTCAAGGTCGGGA

ATTCTAGTGAATTCGCAAGCTGCGTGCAATTTTTAAGCGCATTTATCCACACAGGGTTTGTCGCCTTGCTTGATGGCAGTGCCTAGGATCGAAAAAAGGC
AAAAGCCGAAGTACCCCTAGGGGCGTTCGTCGTCTATCGACTCACTACAATAATGTTTGGTTTACCTGATGATGTTGTTATTAATACTGTGGGCAATATT
**TGTGGATAA**GTAACAAAAAAATACTTATAT**G**AATGACTTATAACAATGATATCGATGTGGAGAAATTGCGGACATGCGGTGCGGGGCACGGGGACGGTCG
TGTGGATAACTAGCGATTCCTCCACAGGCAGGCGTTATCCTCACAATCATACCGATTCTGTCCACCTGGCTTTGGGCAAGTTTTCCACAAGGCCAGCAAA
GCGGATGGCGTGTGGATAACTCGTCCGTAGGACGATACAATGCCCGGTCATTCGCTAACTGGATGGTGAGGTCGCGTGTCGGTCGCTCTCTGGCAACAAT

GTCTTGGCTTTTTGCAGGACGAGCTGAATTCGCAGCAGTTCAACACCTGGATTCGTCCCTTGCAGGCGGAAGAAGGTGACGCCAACGAGCTGAGTCTGTT
GGCGCCCAATCGCTTTGTACGAGACTGGGTCAGCGACAAGTATCTCAAGCGCATCAATGAATTGCTGCGTGAGCTATCGCCAGCCAAGCCGCCCAAGGTG
ACCCTGGCGGTGGGTAGTCGTCGTGCGGCGCCCAGCCCTCAACCCCGTGATCTGGGCAGCCCCGTCTCGGCCAGTCGTTTCCAGCAGCAGCCCGCAGCGC
CTGCCGTACAGACTTCTGCGCGCGGCGACTATGCCGACGAGCAGGAAATTGAGCAGTTGCGCGAGGAGACGCCCCGGCGCGGGACGACGGGCGAGCGCCA
GGTCCAGGTGGAAGGCAGCCTCAAGCACCAGAGCGGCCTCAACCCCAACTTCACCTTCGATACCTTCGTGGAAGGTAAGTCGAACCAGCTGGCTCGGGCT
GCGTCACGACAGGTCTCGGAAAATCCCGGTGGCGCCTATAACCCCTTGTTCCTCTACGGGGGAGTTGGCCTGGGCAAGACGCACCTGATGCATGCCGTCG
GCAACCAGCTTGCCGGGCAGCGCGAAAACGCCAAAGTGATCTATCTGCACTCGGAGCGCTTCGTGGCGGATATGGTCAAGGCGCTGCAGCTCAATGCCAT
CAATGACTTCAAGCGTTTCTACCGCAGTGTCGATGCATTGTTGATCGACGATATCCAGTTCTTTGCCGGCAAGGAGCGTTCTCAGGAAGAGTTCTTCCAT
ACGTTCAATGCGCTGCTCGAGGGGGGGCAGCAGATGATCCTCACCTCGGATCGCTACCCCAAGGAGATCAGCGGGGTGGAAGAACGTCTCAAGTCGCGCT
TTGGCTGGGGACTCACGGTGGCGATCGAACCGCCGGAACTGGAAACGCGCGTGGCCATCCTGATGAAGAAGGCCGATCAGGCCAAGGTCGACCTGCCGCA

>Halomonas xianhensis CGMCC 1.6848 [NZ_FOPY01000006.1:14208..14707], <mnmG oriC2 <COG1126 (glnQ), 586 bp

GTTTTCCAGTCCCTTGATCGAGCGTACCACCTGAAGTTGCACATCGAACGGCAGCGAGGTCGAAATCCCGTTGGGGTAGAGCTCGTGAGTGTCCAGGCCT
TCGGGTTCGATGAATACCTGATGGCTGGTCTTGTCGGCAAAGCGATGGACCTTGTCCTCGATCGACGGGCAATAGCGCGGCCCGATGCCTTCGATCACCC
CCGAATACATCGGCGAGCGATCCAGATTGGCGAAGATGATCTCGTGGGTGCGCTCGTTGGTATGCGCTATATGACAACTGGCCTGGGGCGGATGCATCTC
GCGCGAGCCGAGATACGACATCACCGGCGTCGGTGTGTCGCCAGGCTGCTCCTCGAGCACCGAGAAGTCGACGCTCTTGGCATCAATGCGTGGTGGGGTG
CCGGTCTTTAGGCGTGAGACATGAAACGGCAACGCACGCAGCCGCTGGGCCAGGGCATTGGCCGGCGGATCACCGGCACGCCCACCGGCATGGTTGTCCA
TCCCGATGTGGATGACGCCACCCAGGAACGTGCCTGTGCATAACACCACTGTTTCGGCATGAAAGCGAATTCCGGTCTGGGTAACGGCCCCGCGCACGGT
GTCATTCTCGACGATCAAGTCATCCACACCCTGTTGGAACAGGGTCAGGTTGGGCTGGTTTTCCAGCATGCCGCGGATGGCCGCCTTGTAGCGCACGCGA
TCGGCCTGGGCGCGCGTGGCCCGTACTGCCGGCCCCTTGCGGGCGTTGAGTACGCGGAACTGGATGCCGCCGAGATCGGTAGCCAGGGCCATGGCCCCAC
CGAGGGCATCGATCTCCTTGACCAGATGGCTCTTGCCGATCCCGCCGATGGCCGGGTTGCAGGACATCTGGCCCAGCGTCTCGATGTTGTGTGTCAGCAA
TAGGGTCTGACAGCCCATGCGGGCAGCGGCCAGTGCGGCTTCTGTTCCCGCATGGCCACCGCCGATGACGATAACGTCAAAGCGGTTGGGATAATCCAAG

GTGCACCTCGACTTGCGCCTGTCGGCGCGCCGGTTTGGCAGGTGAGGGTAAATAGCCGCCCAGTATAACGTCCCTGTGGGTAACCGTCAGGGGTTTTCCA
CACGCGCTTTTCCACCTATCACCTTTCCATCTTCATTTAAAAAATAGAGTTAAAGAAAAGAAGTTCTGTTGTGGTTATGACTATTGGTGTGGACATTTTT
**TGTGGATAA**CCTAAAAAACAACTTTTCTTTCATTAAATTGCATTGTTAATCGATAGGGGGAAAAGCAGTAGGGGGGCTGCGGAAATCCCGCGGTCAGGCT
GTGGATGATATTGGAGATTATCCACAGCACGATTCATGCCCTGCTTATCCACCTGCCCATACCTCAACTATCAGCAGGCTGGTCAACAATGTTGGAAAAA
CTCAGGCATGCCAGCCATGACGGGGGCTGCCGGGCATAATCGAAGCCTTGAGAATGAAAAATATCCACAGGCAAAAAAGACCTTGTGTGCAGAATATCAG

GGGAAATGAAGCGCAACATCACCATGAGAAACGCCCACTCGATGAGCGGGCGCTGGAAGGAGGATGGGGGAGAGGCGCGGTGCGCTCAGTGCTTGAGCAC
ACGCGACAGGAATGACTGGGTCCGCTCGTGACGGGGATGGTCGAACACTTCTTCGGGTGGCCCCTGCTCGACGATGCGTCCGCCGTGCACGAAGATCACC
CGGTCGGCAACTTCGCGGGCGAACCCCATTTCATGGGTGACGATCATCATGGTCATGCCCTCGTTGGCGAGTTCACGCATGGCATCCAGCACCTCTCCGA
TCATCTCCGGGTCGAGTGCCGAGGTCGGCTCGTCGAACAGCATCAGGCGCGGCCCCATGGCCAGGGCGCGCGCCAGAGCCACGCGTTGCTGCTGCCCGCC
GGAGAGCTGGCCGGGATACTTGTCGGCCTGATCCTCGATGCCGACCCGGGCCAGCAGCTTGCGCGCCGTTTCCAGCGCCTTGTCGCGCGGCCAGTTACGC
ACTTTCATGGGGGCGAGAGTAACGTTGTCGAGTACCGACAGGTGCGGGAACAGATTGAACTGCTGGAAGACCATGCCGACTTCGGTGCGGATGGTCTGCA
GCGACTTGCCACTCTTGCCGTTGGGCAGCAGTTCGTTGCCATCGACCACGATGCGTCCGCCCTGGAATTCCTCGAGACCGTTGACGCAGCGGATCAGGGT
CGACTTGCCGGAGCCACTGGCGCCGATGATCACCACCACTTCACCGGGAGCGACCTCGAGGTCGATGTCCTGCATGACGTGCAGCTTGCCGAAATGCTTG
TTGACCTTGTCCATGCGGACGATAGGCTCGCGTTCGCCGGTCTGGCGTGATGCGGTGTTGGTTGTTTCGTTCATGCGCTTCTCCTCACTGTCCGGCCAGG
CCGCGGCGCTCTATCTGACGCAGGGCGAAGGACAGGCTCAGCGTGATGACCAGGTACATCATGGCGACCATGAAATAGACCTCGAGGGCGGTAAACGTCG

>Halomonas xianhensis CGMCC 1.6848 [NZ_FOPY01000006.1:58045..58112], dnaA> ig dnaN>, 68 bp

< page 1 | #124 | Halomonas ilicicola DSM 19980

WebSkew GCmin pos. 82720 (GCmin<>oriC1 39.2 kb; GCmin<>oriC2 8.2 kb; oriC2<>oriC1 31.0 kb)

>Halomonas ilicicola DSM 19980 [NZ_FQUJ01000004.1:43295..43794], <rpmH oriC1 dnaA>, 571 bp

GACACTGACCAGGCTGCGTCGATTGCCTGCTCCGGTCTCCTCCGGCATGGCATTGCTGGCCGGCGCATCGGCATCGCCCGTGGGCGCCGCCAGCCCCTGT
TCGTCGTCTGCAGTGGCGGATTCACCACGCTGGGTCTCGCTGATGGCTGGCGAGCTGGGCTGCGCGTCGAGCTGACCATAATCCTGGTTCCACTGGACGA
CCAGCAGGTAGGCAAGCACCGCGAGCGGAATCAGGAGGAGAAGTCGTTTTACATCCATGAGGATCTGCCCAGTAGGGTGATGGAACGTTCGAGACTCGTC
TGCCGCGTCGGAATTCGCCGGGAAGAAAACGGATCCAGGCGAACCACGGGCATAGTCCAATCAAGAGTGCGAGGAGGCTGCCGGCGATCGGTGCACGTCC
TTCTCCAGACGGCGCCACATGCCGTGCAGCTGACGATGCAGAGTGGCATTGTCCAGGTCGTTGACCCCCCGGCGCGCCAGCACGACGATATCCACGGCAG
GCAGGCGCTGTTGTCGGAGGCGAATGGATTCCCGCACCAAGCGCTTCAAGCGGTTGCGGTCGACGGCCCGACGAACGTTCTTCTTGCTGAAGACCAGGCC
TACCCGGACATGGCCCAGTGAATTGTCTCTGGCCAATGCCAGAAGACCCTTACCGTGAACCCTGTAATCGGCCTGTTCGAAGACGTGTCGGTACTCCCCG
GCGGTCAGAAGGCGCAGCGTCCGGGGAAAGGTCTGACTAGACACCTGCCGAGGCCGGCGCTCAGGCGCTCAGGCGCTCAGGCGCTTGCGACCTTTGGCAC
GACGACGCGCGAGAACCTGCCGACCATTCTTGGTTGCCATGCGCGCGCGAAAGCCGTGCACACGCTTACGCTTCAGAACGCTCGGTTGAAAAGTGCGTTT
CATATCAACTTCTTCCCACGTGATTGAGTGAACGATCGATCATGGAAAAGCGCGAGCTCCCGGAAAGGGGCTCGAACGGATCCATCTCAAGAGCGGGAAT

TCTAGTGAATTCGTTGCCGGTGTGCAATTTTTGCGTGTTATCCACACCCGGTGTGATCGAGCCCCTCCTCCACGCCTTCCGGGGGACGGGGACGATGACG
ATGACGATCCCCTGTGCCGCCAAGATGCCTGCGAAGCCTCTTACTACAATAATTTTTTGGTCTTTTAAATACTGCTGTTATTAATACTGTGGGTAATGTT
**TGTGGATAA**CCTGCGATAGCATACTTATATCAGTAATCTATAGCATTAAGAAAACTGTGGAATGGATGCGTGAAACCGCTTTGCATCCGGGGGAAAAGAG
TGTGGACAGATCGCCATCTGTCCACAGATGGTGCCTGTGCACAGCCCCGTACACTCTTTTTCACGCTTTTCCTCGCAGGATTGTCCACAGCGAAGTCGAG
CCTTCCTCGTGTGGATAACTAGAGGTTCGAACGCTACAATGATCGGCCATTTGCATGTTGGACGGTGAGGTCGTGTGTCAGTCGCGCTTTGGCAACAGTG

TCTGAATTACCTTCAGGACGAACTGAACTCTCAGCAGTTCAACACCTGGATTCGCCCCTTGCAGGCCGAGGATGGCGAGCCCAACGAGCTGAATCTGCTG
GCGCCCAACCGTTTTGTCCGGGACTGGGTCAGCGATAAATATCTCAAGCGAATCAATGAGTTGATGCGTGAGCTGTCGCCTGCCAAGCCACCCAAGGTGT
CGTTGACCGTGGGGAGTCGGCGGTCGGCCCCCAGCCCTCAGCCACGCGAACTGGGAAACCCCGTGTCCGCTAGCCGTTTCCCCAGGCCCGCGGCGCCTGC
GGTGCATAGCCCGCCGCGCAGCGATTACGCGGACGAGCAGGAAATCGAAAAGCCCAGGAACGACTCGACGTCGCGATCCGAGGGACGTCAGGTGCAGGTC
GAGGGTAGCCTCAAGCACCAGAGCGGCCTCAATCCCAATTTCACCTTCGACACCTTCGTCGAGGGCAAGTCCAACCAGCTGGCGCGCGCAGCCTCCCGTC
AGGTGTCCGAAAACCCCGGTGGCGCCTACAATCCGTTGTTCCTGTACGGTGGTGTCGGTCTGGGCAAGACGCACCTGATGCACGCGGTAGGCAACCAACT
GGCCGGTCAGCGCGAGAATGCCAAGGTCATCTACCTGCACTCGGAGCGTTTCGTGGCGGACATGGTCAAGGCTCTGCAGCTCAACGCGATCAATGACTTC
AAGCGCTTCTATCGCAGTGTCGACGCCCTGTTGATCGACGACATCCAGTTCTTCGCCGGCAAGGAGCGTTCCCAGGAAGAGTTCTTTCACACCTTCAATG
CCTTGCTCGAGGGTGGTCAGCAGATGATCCTGACCTCGGATCGCTACCCCAAGGAAATCAGTGGCGTGGAAGAACGTCTCAAGTCGCGTTTCGGTTGGGG
GCTGACGGTGGCGATCGAGCCCCCGGAACTCGAGACACGGGTGGCCATCCTGATGAAGAAGGCCGATCAGGCCAAGGTCGATCTGCCCCATGATGCCGCC

>Halomonas ilicicola DSM 19980 [NZ_FQUJ01000004.1:74288..74787], <mnmG oriC2 <COG1126 (glnQ), 591 bp

TCTCCATGCCGCGAATCGAGCGCACCACCTCGAGCTGCACGTCGAAGGGCAGCGAGGTGGAAATGCCATTGGGATAGAGCTCGTGGGTGTCGAGCCCCTC
CGGCTCGATGAAGATCTGATGGCTCTGCTTGTCGGCAAACCGGTGCACCTTGTCCTCGATGGAGGGGCAATAGCGCGGTCCCACGCCGCCGATCACCCCG
GAGTACATGGGCGAGCGATCGAGGTTGGCGAGGATGATCTCGTGTGTCCGGGCGTTGGTGTGCGCGATGTGGCAGTTCACCTGATCCGGATGCATCCCCC
GGTTGCCGCGATAGGACATCACCGGTGTCGGCGTATCGCCGGGCTGCTCCTCCAGGCCCGAGAAGTCGAGGCTCCTGGCATCCAGGCGGGGCGGGGTGCC
GGTCTTGAGGCGATCGACCCTGAACGGCAGGGCACGCAGGCGGCTTGCCAGGGCGTTGGCGGGCGGGTCCCCGGCGCGCCCGCCGTTGTGGTGATCGAGC
CCGATGTGGATGACCCCGCCGAGGAAGGTCCCGGTGCACAGCACCACGGTCTCGGCGAAGAAGCGGATGCCGGTCTGGGTCACCGCACCGCGCGCCGTGT
CACCGTCGACGATCAGGTCGTCGACGCCCTGCTGGAAGAGGGTCAGGTTGGGCTGGTTCTCGAGCATGCCACGAATCGCCGCCTTGTAGCGCACGCGATC
CGCCTGGGCCCGGGTCGCCCGCACCGCCGGCCCCTTGCGGGCGTTCAGCACGCGAAACTGGATGCCTGCCCGATCCGTGGCCAGGGCCATGGCGCCCCCC
AGGGCGTCGATCTCCTTGACCAGGTGACTCTTGCCGATGCCGCCGATGGCGGGATTGCAGGACATCTGCCCCAGGGTCTCGATGTTGTGTGTTAGCAGCA
GGGTCTGACAATTCATGCGGGCAGCGGCCAGTGCGGCTTCCGTTCCCGCATGACCACCGCCGATGACGATGACGTCAAAGCGGTTGGGATAATCCAAGGT

TGTACCTCGACTTGCGCCAGGGAGCGCTTGGTTGGTCTGTGAGGAAAATCGAGCCGCACAGTATAACGTCCTTGTGGGTAACCGTCAGGGGTTTTCCACA
CCTGTTCCAACCGCTCTTGTCACCACGACCCTATAAAATAAAAACAGAATAGAAATAAAGAAGTTCTGTTGTTGTTGTATCTATTGGTGTGGACATTTTT
**TGTGGAAAA**GCCGTTTAATACGTTTAATAACAATGGCATGCAATGTTAACCGATGGCTGGAGAAGCGGTAAGGCAACTGCGGACAAGCGGTCCGGAGGCT
GTGGATGGAATCTGCGGTTATGCACAGCGGGTTTCCTCCCCGCCTTATCCACCGACGTCTACCGAGGCTGTGGGCAGGGTCGTCAACAATCGCGCCGGGA
AGGCATCAGCCTGCTGGTAGCGGGGCCTGCAGCGCAGATTCCGCCAAGTGTAGGAAAAAGTTGCCCACAGGGGCCGAAGGCGTCCTGTGGGCGGTTTCGA

GGGGAAAAAGAGACGGGCAAAAGCGGTGGGCGAAAAGCGAGTTCGGCCGCTGGACGAACGAACTCGGAGGTGTGCCAGGCCCTTGCTGTCAGTGCTTGAG
CACCCGGGACAGGAAGGCTTGAGTGCGCTCCTGACGAGGATTGTCCAGCACCTCCTGGGGAGGGCCTTCCTCGACGATCTGGCCGTCGTGCACGAAGATC
ACCCGGTCGGCGACTTCCCGGGCGAAGCCCATCTCGTGGGTGACGATCATCATGGTCATGCCGTCCTTGGCCAGTTCGCGCATGGCATCGAGCACCTCGC
CGATCATCTCGGGGTCGAGGGCCGAGGTGGGCTCGTCGAACAGCATCAGGCGCGGTTCCATGGCCAGGGCCCGCGCCAGAGCCACACGCTGCTGCTGACC
GCCCGAGAGCTGACTGGGGTACTTGTCGGCCTGATCCTCGATGCCGACCCGTTCCAGCAAGCGATGCGCCGTCTCCAGGGCCTTGTCCCGGGACCAGCCG
CGCACCTTCCTGGGCGCCAGGGTCACGTTGTCGAGGACCGACAGATGGGGAAAGAGGTTGAACTGCTGGAACACCATGCCGACCTCGGTGCGCACCGCCT
GAAGCGACTTGCCGCTCTTGCCGTCGGGCAGCAGCTCGTTGCCATCGACCTCGATGCGCCCGCTCTGGAACTCCTCCAGGCCGTTGACGCAACGGATCAG
CGTCGACTTCCCCGAGCCACTGGCCCCGACGATCACCACCACCTCGCCCGGGGCGACTTCGAGATCGATGTCCTGCATCACGTGCAGCTTGCCGAAATAC
TTGTTGATCTTTTCCAGCCGTACCATGGGCTGCGTCGTCGCTTGCGTCGTGTCGTTCATCGTCTGCTTCCTCATTGTCCGACCAGGCCGCGGCGTTCGAG
CTGACGCAGGGCGAGGGAGAGGCTCAAGGTGATGGCCAGATACATCAGGGCGACCATGAAATAGACCTCGAGGGCGGTGAAGGTGGTGGCGATGTAGACC

>Halomonas ilicicola DSM 19980 [NZ_FQUJ01000004.1:41790..41856], dnaA> ig dnaN>, 67 bp

WebSkew GCmin pos. 199732 (GCmin<>oriC1 28.9 kb; GCmin<>oriC2 8.9 kb; oriC2<>oriC1 19.9 kb)

>Halomonas muralis DSM 14789 [NZ_FNGI01000004.1:228403..228902], <rpmH oriC1 dnaA>, 564 bp

CAACCAGGCTACGCCGGCTATCTGTCCGAGGGGCTTCTGGCATGGCATCGGCGGCTGGCGCCGAAGTGGCATTGGAAGGTGCCGCCAGCCCCTGGTCGCT
GCGTGTTTCGTTATCGCCGTCGCTGGGCGCCGCGCTGAAGCCGGGAGCCTCGGGTTCAACGGTGACCTGGCCGTAATCCTGGTTCCACTGGATGACCAGC
AGGTAGGCGAGCACCGCAAGCGGAATCACGAGAAGGAGTCGTTTTACGTCCATGGGGGTTCTGCCCGGTGGGGTAATGGATCATTCTGGCGGCTTGTCTT
CCGACCGCTACTCGCCAGCGGCGAGGCGGATCGGTACGACCTTGCTAATACGAAGCGCGAATCGGGTCAGGAAGACGCCGCCACGGTCGACTGTTTGCGA
ACGTCTTTTTCCAGGCGCCGCCACATGCCGTAGAGCTGGCGGTGCAAGGTCGCGTTATCCAGTTCGTTAGCGCCGCGACGCGCCAGGACGATGATGTCCA
CGGCGGGCAGACGATGTTGCTGGAGACGAATGGACTCGCGGACGAGACGCTTCAGACGATTGCGATCGACGGCACGGCGAACGTTTTTCTTGCTGAAGAC
CAGACCGATCCGTGGGTGGCCCAACGCGTTGTCGCTGGCCAACGCCAGCAACCCCTTGCCATGGACCTTGAACGGGGCATGGTCGAAGACGCGTCGATAT
TCCCCGGCAGTCAAGAGACGCAACTGCCGTGGGAAGTTCTGACATGACATTCGCGGCTCGAGTTTAAGCGCTCAGACGCTTGCGACCCTTGGCGCGGCGG
CGCGCCAGAACCTGGCGACCACCCTTGGTGGCCATGCGCGCACGAAAGCCATGTACACGCTTGCGCTTGAGAACGCTCGGTTGAAAAGTGCGTTTCATAA
CTGCTGATTCCCACGTGGTTAAATTTGGACTGATGACAAGCCGATGTGCCTTGATACCCGCCAGGAGCCGGGCGGATTTGGCTCAAGGTCGGGCATTTTA

GTGAATTCGCCAGCCGCGTGCAATTTTTCCGCGGATTTATCCGCGCGGGGCCAAGATTTTTTTAGTATTTTGCCGTCGTTGGCCAGTAAGTGGCCGAATG
TCTATCCCCAGGGTCGGTGCAGTGACCTGCCAAGGCAATACAAACATTATATTAACTATCGATACCAGATAATGCTGTTATTAACATTGGGAATCAATTT
**TGTGGACAA**GCAGTAAAATAAATTATTTTTCAATAAGTTAACCGAATATTGCTTTTGTGGAAAAACTGCGCGAAACGTTTAGCGGCATTAGGAGCAGCGT
TTGAACGGCGGGATTATATTTGGGGGTTGCCGAGTCCGACGCGATTCGATGGCGGGCTCATTCACCGGTTGTCCACAGTTTTTTCACGGCCCTAATCGTG
TGGATAACTCGCCCCACGAACGATATACTGGCCGGTCATTCGCTAACAGGATGTGAGGTCGCGTGTCGGTCGCTCTCTGGCAACAATGTCTGGATTACCT

GCAGGATGAGCTGAGTTCCCAGCAGTTCAATACCTGGATTCGTCCGTTGCAGGCGGAAGAGGGTGGAGTCGACGAGCTCAGCCTGTTGGCACCCAACCGC
TTCGTGCGGGACTGGGTCAGCGATAAGTATCTAAAGCGTATCAATGAGCTGCTACGTGAACTGGCGCCTGCCAAGCCGCCCAAGGTGGCCCTGGCGGTGG
GCAGTCGGCGTGCCACGCCCAGCCCACAGCCACGTGAATTGGGCAACCCGGTATCTGCCAGCCGCTTCAAACAGCCTGGGCCACCTTCGGTGCATACGCC
ACCACGCAACGATTACGCTGATGAGCACGAGATCGAGCAGTTACGCGAGAAGAGCGCTCGTCGTTCGGGAGATCGCCAGGTTCAGGTCGAGGGCAGCCTC
AAGCACCAAAGTAGCCTCAACCCCAATTTCACCTTCGACACCTTCGTTGAGGGTAAATCCAACCAGTTGGCGCGCGCGGCCTCGCGCCAGGTATCGGAAA
ACCCCGGCGGCGCCTATAACCCGTTGTTTCTGTACGGTGGCGTGGGCCTGGGCAAGACGCACCTGATGCACGCCGTAGGCAATCAACTGGCATTGCGGCG
TGAGAACGCCACCGTGGTTTACCTGCATTCCGAGCGCTTCGTCGCCGATATGGTCAAGGCGCTTCAGCTCAATGCCATCAACGATTTCAAGCGCTTCTAT
CGCAGTGTCGATGCCTTGCTGATCGATGACATCCAGTTCTTCGCCGGCAAGGAGCGCTCGCAGGAGGAGTTCTTTCATACCTTCAACGCCTTGCTTGAAG
GTGGCCAGCAAATGATTCTCACTTCTGACAGATATCCCAAGGAAATCAGTGGTGTAGAGGAGCGTCTCAAGTCACGTTTCGGGTGGGGGCTGACGGTGGC
GATCGAACCGCCGGAGCTCGAGACGCGGGTTGCGATCTTGATGAAGAAGGCCGATCAGGCCAAGGTCGACTTGCCCCACGATGCCGCTTTCTTCATTGCC

>Halomonas muralis DSM 14789 [NZ_FNGI01000004.1:208466..208965], <mnmG oriC2 <COG1126 (glnQ), 611 bp

CTCCAGGCCAGCGATCGAACGCACCACGCGTAGCTGGATATCGAACGGCAGCGAGGTCGAAATGCCGTTGGGATACAGCTCGAAGGTATCGAGGCCTTCC
GGTTCGATGAATATCTGGTGGCTGGTCTTGTCGGCGAAACGGTGGACCTTGTCCTCGATCGATGGGCAATAGCGCGGCCCCACGCCTTCGATCACGCCGG
AATACATCGGCGAACGATCGAGATTGGCGAGAATGATCTCGTGAGTGCGCTCATTGGTATGCGCGATATGACAACTCACCTGACGCGGATGGTCGTTGCG
CGAGCCGAGATAGGACATCACCGGCGTGGGGGAATCGCCGGGCTGTTCCTCGAGGTTCGCAAAATCGATGCTCTTGGCATCGAGGCGCGGCGGCGTGCCG
GTCTTCAGTCGATCGACCCGGAACGGCAATGCACGCAGGCGTTGCGCCAGGGCGTTGGCCGGTGGATCGCCGGCCCGCCCGCCGGCGTGATTATCGAGCC
CGATATGCATCACGCCACCGAGGAAGGTGCCGGTACACAGCACCACCGTTTCGCCATGAAAGCGGATACCGGTCTGGGTGATCGCGCCACGCACGGTGTC
GCCATCGACGATCAGATCGTCGACGCCCTGTTGGAACAGCGTCAGATTGGGCTGGCTTTCCAGCATGGCGCGAATGGCTGCCTTGTAGCGCACGCGATCG
GCCTGGGCACGCGTCGCGCGTACCGCCGGGCCCTTGCGGGCATTGAGCACGCGAAACTGGATGCCGCCCTGGTCGGTGGCCAGCGCCATGGCGCCGCCCA
GCGCATCGATCTCCTTGACCAGATGGCTCTTGCCGATCCCGCCGATCGCGGGATTACAGGACATCTGGCCGAGGGTCTCGATGTTATGCGTCAGCAACAG
CGTTTGGCAGCCCATGCGAGCGGCGGCCAGTGCGGCTTCCGTTCCCGCATGGCCCCCGCCTATGACGATAACGTCAAAGCGGTTGGGGTAATCCACGATG

TACCTCGACTTGCGCCCGTGGCGCGCGGTTATTTCGGTGAGCTGAAATCAAGCCGCCCAGTATAACGTCCCTGTGGGTAACCGTCAGGGGTTTTCCACAT
GCGGGTTTCACCGTCCAAGGGAGTCGTCCCTTAGTTATTAAAATAAGTATAAAAAGAAAGAAGTTCTGTTGTGGTTGTAACTACTAGTGTGGACACTTTC
**AGTGGGTAA**GCGGTTTATTTTTATATTTTTCATGGAGTTGAAATGTTGATAAAACATTGTAAAAGCGGTCGGGTTGCTGCGTACATCCGCTGGGCTTGCT
GTGGACCGAATGACTCGTTGTGCACATGCCGAATCATGCCAAGCTTATCCACCGCTGGCTACCGACTCTGTGAGCAGGTTAGTCAACAGCTCGCCCGATA
TTGCTGAAAAGTCCAGCCCCGGCGGGTTGGCGATGAGATATCGGTGTCTAGCGAGAAATTTTATCCACAGACCCAAAAGACCCACGTGAAGGGTCATTCC

GCGCTCATGCAAAGCCGTGCGGCGAAGAATCGGCGCCTAACAACGCAAGAATCCGCCTCGATGGGCGGATCCTTGCGTTGCGATAGACGGAGAACTCGTA
GCGGAATCTAGTGCTTGAGCACGCGTGACAGGAAGGATTGCGTGCGCTCGTTTTGCGGGTTGTCGAGAACCTCATGGGGTGGGCCCTGCTCGACGATCTG
GCCGCCATGGATGTAGATCACCCGGTCGGCGACCTCGCGAGCGAAGCCCATTTCGTGAGTGACGATCATCATCGTCATGCCGTCATTGGCGAGCGTGCGC
ATGGCATCCAGCACCTCGCCGATCATCTCGGGATCCAGCGCCGAGGTCGGCTCGTCGAATAGCATCAGCCGCGGCTCCATGGCCAGAGCGCGTGCCAGCG
CCACGCGTTGCTGCTGACCGCCCGAGAGCTGGCCGGGATGCTTGGCCGCCTGATCCTGTATGCCGACCCGCTCAAGCAGCTTGTGGGCCGTTTGCAGCGC
CTTGTCGCGTGGCCAGCCGCGTACCTTGATGGGTGCCAGGGTGACGTTGTCGAGCACCGATAAGTGGGGAAACAGGTTGAACTGCTGAAACACCATGCCG
ACTTCGGTGCGAATGGTCTGCAGCGATTTGCCGCTCTTGCCGTAGGGCAATAGCTCGTTGCCGTCGACTTCGATATTACCGCTCTGGAACTCCTCGAGGC
CGTTGACGCAGCGAATCAGCGTCGATTTGCCCGAGCCGCTGGCGCCGACGATCACGACGACCTCGCCGGGCACCACTTCGAGATCGATATCCTGCAGCAC
ATGCAGATTGCCGAAGTGCTTATTGACCTTCGCCAAGCGTACGATGGGTGGCTGCTGAGCGGTTTCTTGCGCGGCTTCGCGCGTTTCGTCGTTCATTGCT
TGGATCCTTTACTGACCGACCAAACCCCTGCTTTCGAGCAGGCGCAGGGCGTAGGAGAGACTCAGCGTAATCACCAGGTACAGCATTGCCACCATGAAAT

>Halomonas muralis DSM 14789 [NZ_FNGI01000004.1:230329..230393], dnaA> ig dnaN>, 65 bp

WebSkew GCmin pos. 227934 (GCmin<>oriC1 29.2 kb; GCmin<>oriC2 0.3 kb; oriC2<>oriC1 28.9 kb)

>Salinicola halimionae CPA60 [NZ_PZJQ01000002.1:c198690..199189], <rpmH oriC1 dnaA>, 594 bp

CTTGGGCGGCGTCGGCGTTGGCCGGCATGTCGTCTTGGATCGGCTTTGCCTTGCCGTCGCTGTCCGTCGGTACAGAAAGAGAACCGACGTTATCACCGTC
GTCACCGTTATGGCTGCCACCCGATGAAGCCGCCATCTCCGGCGCCGGGTTGTTCACCACGTTCTGACCATAGTCCTGATTCCACTGCATGATCAGCAGA
TAGGCCAACACGGCCAGGGGAATCACGAAAGGAGGCGTTTCAAATCCATATTGGGGTCTGCCCGGTAAGGGAATGGGTCAGTGGGCGCCGGGCATCGGC
GATTGACCGGCAACATTCGCCGGTGCTCCCCGGCTTTCGACGCGGGGGTGGCCGATCAAGAAGACGAAACGTCCTGATGCGCCTGACGAGCGGACTTTTC
CAGGCGTCGCCACATGCCATTGAGCTGGCGGGAGACGCTGGCATTATCCTGCTCGTGCAACCCTCGTCGAGCGAGTATGACGATATCCACCGCCGGTAAA
CGATGCTGTTGATGGCGGATGGATTCGCGCACAAGGCGCTTGAGGCGATTGCGGTCGACGGCACGGCGCACGTTCTTCTTGCTGAAGATCAACCCGATCC
GCGGATGGCCCAGCGCGTTGGGACGGGCCAACGCCAGTATTCCTTTACCGTGGACCTTATAGGGAGCGTGGTCGAAAACGTGCTTGTAGTCCCCGGCATT
CAGCAAGCGCAGTTGCCGAGGAAAACGCTGATCGGACACCCGGCGTCCGATCAGGCGCTCAGACGCTTACGGCCTTTGGCGCGACGGCGAGCAAGAACTT
GACGGCCATTCTTGGTGGCCATGCGGGCGCGAAAACCGTGGACGCGCTTGCGCTTCAGAACGCTGGGCTGAAAGGTGCGTTTCATAGCTGCTGATTCCCA
ATTAGTTGATTTCTAAAGAAAACGGGTTGAAGCGGGGTGCCCGGCACGCGCAACGAGAGCGGGCTCACTCGACTCAAGAGCGGGAATTCTAGTGAATCCG

TAGTTCGGGTGCAATTTTTCCGTGAATTATCCACACCCAGGTTGTTCCTGAATGGCGGTCTGCCGAACCGGATTTTGGGGTGCTGTCGAGGCTCGCTGTT
ACCAGGGACCGGGTTTTCTTCGATTGATCGATCGACCGGCGAGATCGGGGTTTCGAGAGGCTGTCAATCACACCATGTGGTTAACAATGGGGATAACAGT
**TGTGAACAA**CCCTATTTTTTTCTTATAAAA**T**AGTGGCTTATTGTTTTAATTGGCTTGTGGGGAAAGTGCTTGTAGGCGGTGTACAGGCGGGGATGGATG
GACGACAGGAACTTTTCCAACACAGGGTTTTATCGATGTGTCCCTCCGCGGTACCGTAAAATCCACAGGGTTGTCATCCTGTTGTCCGCCAGGTTATCCA
NNTGGATAACCTGGCGGGCCGACGCTACAATGTGCGGCCGTTTTTTCGATTCCATTCTCTCAACGTTCGGTGAGGTCGTGTGTCTGTCGCTCTCTGGCAA

CAATGTCTGAACTCGCTGCAGGATGAATTGAGCTCTCAGCAATTCAATACTTGGATTCGTCCTCTGCAGGCAGAGGATGGCGAAATGGACGAACTCTGTC
TTCTTGCCCCCAATCGCTTCGTCCGTGATTGGGTCAGCGACAAGTATTTGAAACGCATCAATGAGCTGCTGCGCGAATCTGCGTCCAGCAGCAAACCCCC
CAAGGTGCAGCTGACGGTCGGCAGTCGCAGGGCAGCGGCAGTGTCGCCGCGCGATCTGGGCGCACCGGTTTCTGCACCCAGCCGCCCGCCCCAGCGGCCG
GCGGCGGCTCCCAGCGTGCAGACTGCCCCGCGTTCGAGTGAAGCGGCGGAAGAGCGCCAACTCGACCGATCTCGTGAAGAGAGCGGGCGGCGAGGCAGAG
AGCGCCAAGTCCAGGTCGAAGGTGGACTCAAGCACGGCAATGGCCTCAATTCTAACTTCACCTTCGAAACTTTCGTCGAGGGTAAGTCGAACCAGTTGGC
GCGTGCCGCCTCGCGCCAGGTCTCGGAAAATCCCGGCGGTGCCTACAATCCGCTATTTCTCTACGGTGGCGTGGGGCTGGGCAAGACTCACCTGATGCAC
GCAGTAGGCAATCATCTGGCAGGGTTACGGGACAACGCCAAGGTCGTCTACCTTCACTCCGAGCGTTTCGTCGCTGACATGGTCAAGGCGTTGCAGCTCA
ATGCGATCAACGATTTCAAGCGTTTTTACCGCAGCGTTGACGCGTTACTGATCGACGACATTCAGTTCTTCGCTGGCAAGGAGCGGTCCCAGGAGGAGTT
TTTCCACACGTTCAACGCCCTGCTGGAAGGCGGGCAGCAAATGATCCTGACCTCGGATCGCTATCCCAAGGAGATCAGCGGCGTCGAAGAGCGCTTGAAA
TCCCGTTTCGGTTGGGGATTGACGGTGGCGATCGAACCGCCGGAGCTCGAAACGCGCGTCGCTATCCTGATGAAAAAGGCCGATCAGGCCAAGGTCGACC

>Salinicola halimionae CPA60 [NZ_PZJQ01000002.1:c227407..227906], <mnmG oriC2 <COG1126 (glnQ), 1026 bp

GTATCCAGCCCTTCGGGTTCGACGAAGACCTGATGGCTGTTCTTGTCGGCGAAACGATGAACCTTGTCCTCGATCGACGGGCAGTAACGCGGCCCGACGC
CCTCGATCACCCCGGAATACATCGGCGAGCGGTCGAGATTGGCGAAGATGATCTCATGAGTGCGTTCGTTGGTGTGGCCGATATGGCAATCGACCTGCCG
CGGATGTTGCTCGCGGCTGCCCAGATACGACATCACCGGCAACGGCGTGTCGCCCGGCTGACGCTCGAGTTTGGAAAAATCGACGCTTTTGGCATCGATC
CGCGGCGGGGTACCGGTCTTCAGCCGATCGACCCGGAACGGCAGGGCGCGCAACCGCTGAGCCAGCGCATTGGCCGGCGGATCTCCGGCCCTGCCGCCGG
CATGATGATCCAGACCGATGTGGATCACGCCACCCAGGAAGGTTCCGGTACACAGCACCACGGTATCCGCCATGAAGCGAATGCCGGTCTGGGTCGTGAC
ACCGACGACGCTGTCGTTGTCGACGATCAGATCGTCGACGCCCTGCTGGAACAGGGTCAGATTGGGCTGGTTTTCCAACATTCCACGGATGGCGGCCTTG
TAACGGACTCGATCGGCCTGAGCCCGTGTCGCTCGCACCGCCGGCCCCTTGCGTGAGTTGAGCACGCGAAACTGAATGCCAGCCAGGTCAGTCGCCTGTG
CCATGGCACCGCCCAAGGCGTCGATTTCCTTGACCAGATGACTCTTGCCGATCCCGCCGATGGCGGGGTTGCAGGACATCTGGCCCAGCGTCTCGATGTT
GTGCGTCAGCAGCAGGGTACGGACACCACTGCGAGCCGCAGCGAGTGCGGCTTCGGTGCCGGCGTGGCCACCGCCGATCACGATGACCTGAAAACGGTCG
GGATACTCCACGATTTACCTCACGAGGTGCTGATACGAACGCATCGAATCGGCACCGACAGCAAATGGATCTCGAGACCTCGCCAAGACGAGGTTCAGCG

CCGTCGCAGCGACGCTTTTCATGGTGGCCGAGAACGGGTGGCCACGAAAGCCGCCCAGTATAAACCTCCTGTGGGTAACCGTCAGGGGTTTTCCACACCC
GATTCGACAGTGAGTCGCTAACTCACAGAGTCATTATTAATTAATGAGAAATATATAAAGAAGTTCTGTTGTGGTAGTAACTAATGGTGTGGACATTTTC
**AGTGGATAA**GTGGAAATACTCTATAAATTACGAGAACTTGCATTGTTATTCGATTGCTTGAAAAGCATGGGGCATGCTGCTGACAGAGGCGTCGATTCCT
GTGGACAGAAGCCTTATTTGTGCACAGCCCGCACTGCACACCTGTTATCCACCGCTGCATACGTTATCTGTTACCGCGGTGGTCAACACCCCCTCGATCA
TTCGCTAGCGTCGCGTGAGTCGGGGCTGCAGCAAGATCGGCAAAAAAACTATCCACAGGCGAAAAAAGGCGGTCTGGATAACCTGAAACGACAAATTTCG

ACCTTCCTGGGCGGATTTTCTGCTTCCGACAAGCGGGAATGAACGTGCGGGTTCGGCCGGACGGTTCAGAATCGGCAAAATCGATGGAGAACGACGAGCC
AGACAACCGCAGAAAAGACGGCGATTCGAACGTCAATGGTTCGAGTCCGAAACAGCCGCCAATGAAACGGAAGGGCGATCAACATGGGAAAAAGCGCTGG
GGGAAATCAATGGGGGGAAATCGATGGGCAAAAACCATAGAAAACAGTGCCGAAGAAAAGCGGCGGGGAAAGGGAGCGAGAAGTGAGGCAAGGAGCACAG
CGGCGCAGACAGCGAGCGTGACTCGGTCTGGAAATAGGTCTCCGCGGCACTCGACAGGAGAGCCGATCCACCGGGCAGCAGATCGCGTGCAAGCGACTTG
AAGGGAGCCCTCAAGCCGCGCACAACCCGGTTATCGACTAGTGCTTGAGCACCCGCGCCAGAAACGCCTGAGTACGCTCATGCTGCGGCTGGTCGAAGAG
CTGATCGGGCGGGCCCTGTTCGACGATTCGACCTTCGTGCACGAACACCACACGATCGGCGACTTCACGGGCAAAGCCCATCTCGTGGGTGACGATCATC
ATGGTCATGCCGTCGTTGACCAGCTCGCGCATGGCGTCGAGTACCTCGCCGATCATCTCCGGATCCAGCGCCGAGGTCGGCTCGTCGAACAGCATCAGTC
GAGGTTCCATTGCCAGCGCACGAGCAAGCGCCACGCGTTGCTGCTGACCGCCCGAAAGCTGGGTCGGATACTTGTCGGCCTGATCGGAGATGCCGACCCG
CTCCAGCAACTTGTCGGCGGTGGTGTTGGCCTGCTGACGCGTCCAGCCACGTACCTTGCGCGGCGCCAGCGTGACGTTGTCGCGGATCGACAGGTGCGGG
AACAGATTGAACTGCTGGAACACCATCCCGACTTCGGTGCGCACGGCCTGCAGTGATTTGCCACTGGCGCCGTTGGGCAGCAACGTCTTGCCATCGACTT

>Salinicola halimionae CPA60 [NZ_PZJQ01000002.1:c197050..197234], dnaA> ig dnaN>, 185 bp

GTCGCCGCGCTGCGAATTCGACCTCGACAGGGATTGCGGGCGGCGTCCGGCGAGGCAAGAATGGGACGGCACGGCCCGACAGCCTTCACCGCCCAGCCCA
TTTCAGGATGGACCCGATCCAGATTCGTCGATCCTGTTGAGGCGTATTGATTTCAGCTCAGATTTATTCCAACTACGGAGCCCCA

WebSkew GCmin pos. 281358 (GCmin<>oriC1 32.2 kb; GCmin<>oriC2 1.9 kb; oriC2<>oriC1 30.3 kb)

>Salinicola halophyticus CR45 [NZ_PZJN01000009.1:313301..313800], <rpmH oriC1 dnaA>, 617 bp

TGCCTTGCGCGGTATCGGCGTCGGCGGGCATGTCGTCCTGAATCGGCTTGGCCTTGCCGTCGCTGCCGGTAGGTGCAGAGAGCGCACCGACATCGCCGTC
TTCACCGCTATTGTTGCCACCCGATGAAGCCGCCATCTCCGGCGCCGGGTTGTTCACGACGTTCTGACCGTAGTCCTGATTCCACTGCATGATCAGCAGA
TAGGCCAGCACGGCCAGAGGAATCACGAAAAGGAGGCGTTTCAAATCCATATAGGGGTCTGCCCGGTAGGGGAATGGGTCAGTGGGCGCCGGGCATCGGC
GGACGACCGGCAGCATTTGCCGGTGCTCCCCGGCCTTTGACGCGGGGATGTCCGATCAAGAAGACGAAACGTCCTGTTGCGTCTGACGAGCGGATTTTTC
CAGACGCCGCCACATGCCATTGAGCTGACGAGAGACGCTGGCGTTATCCTGCTCGTGCAATCCTCGTCGCGCGAGTATGACGATATCCACTGCCGGTAAA
CGATGCTGCTGATGGCGAACGGACTCGCGCACAAGACGCTTGAGGCGATTGCGGTCGACGGCACGGCGCACGTTCTTCTTGCTGAAGATCAACCCGATCC
GCGGATGGCCCAGTGCGTTGGGACGGGCCAACGCCAGTATTCCTTTACCGTGGACCTTATAGGGAGCGTGGTCGAAAACGTGCTTGTAGTCCCCGGCTGT
CAGCAAACGCAGCTGCCGGGGAAAACGCTGATCGGGCACCCGGTGTCCGATCAGGCGCTGAGACGCTTACGGCCTTTGGCGCGACGACGAGCAAGGACTT
GGCGGCCGTTCTTGGTGGCCATGCGGGCGCGAAAACCGTGGACGCGCTTACGCTTCAGAACGCTGGGCTGGAAGGTGCGTTTCATAGCTGCTGATTCCCA
ATTAGTTGATTTCTAAAGAAAACGGGTTGAAGCGGGGTGCCCGGCACGCGCAACGGGAGCGGGCTCACTCGACTCAAGAGCGGGAATTCTAGTGAATCCG

TAGTTCGGGTGCAATTTTTCCGTGAATTATCCACACCCGGGCCAGGCTCGAATGGCGGCCCGCCCTGCTCGATTTCTTCGGCTGGCAGCCCTTGGCGTTG
TCGCGATGAGCGATCGATTTCGATTCCTCGGCCGAGCGGCGAGATCGCGATTTCTCGACTCTGTCAATCACACCATGTGGTTAACAATGGGGATAACAGT
**TGTTAATAA**CTCATTTATAAACAAGTAAAACAATAAGTTATGCTTTATTTTAGGTTGTGGGAAAGTGCGTGAAGGCGGTGTATAGCGCGGGGATGAGTG
TGCTCAGGGAAACTTGGCCGAACGACGCTGCGTCGATGTTTCCCCATCCTATTACCGTAGAATCCACAGGATTGTCACCTTGCTATCCTTCAGTTATCCA
CAGTCGGTCGAGATTTCGACATCGTGTGGATAACCTGACGGGTCGACGCTACAATGTGCGGCCGTTTTTTCGATTCCATTTCCCAACGTTCGGTGAGGTC

GTGTGTCTGTCGCTCTCTGGCAACAATGTCTGAACTCGCTGCAGGACGAATTGAGCTCTCAGCAATTCAATACCTGGATTCGTCCTCTGCAGGCAGAGGA
CGGCGAAATGGACGAGCTCTGTCTTCTCGCCCCCAATCGCTTCGTACGTGACTGGGTCAGTGACAAGTATCTGAAGCGAATCAATGAACTGTTGCGTGAA
TCCACGTCCAGCAGCAAGCCTCCCAAGGTGCAACTGACGGTTGGCAGTCGCCGGGCCGCAGCGGTATCGCCGCGAGATCTGGGGGCACCGGTTTCCGCGC
CCAGCCGTCCGTCCCAGCGACAGGCGGCCGCGCCCAGCGTGCAGACCGAATCGCGTTCGAGCGAGGCGGCGGAGGAGCGTCAGCTCGATCGATCCCGCGA
AGAGAGCGGCCGGCGAGGCAGAGAGCGCCAGGTTCAGGTCGAAGGTGGGCTCAAGCACGGCAATGGTCTCAACTCGAACTTCACCTTCGAAACGTTCGTC
GAGGGCAAGTCGAACCAGCTGGCGCGTGCCGCCTCGCGGCAGGTCTCGGAAAACCCCGGCGGTGCCTACAACCCGCTGTTTCTCTACGGGGGCGTGGGGT
TGGGCAAGACCCACCTGATGCACGCGGTGGGCAATCATCTGGCTGGGCTTCGCGACAATGCCAAGGTCGTCTATCTTCACTCCGAGCGTTTCGTCGCCGA
CATGGTCAAGGCGCTGCAGCTCAACGCGATCAACGACTTCAAGCGTTTCTACCGCAGCGTCGATGCTCTGCTGATCGACGATATCCAGTTTTTTGCCGGC
AAGGAGCGTTCCCAGGAGGAGTTCTTCCATACGTTCAACGCCCTGCTGGAAGGCGGGCAGCAGATGATCCTGACCTCGGACCGTTACCCTAAGGAGATTA
GCGGCGTCGAAGAGCGGCTGAAATCCCGCTTCGGCTGGGGCCTGACGGTGGCGATCGAGCCGCCGGAGCTCGAGACGCGCGTGGCGATCCTGATGAAAAA

>Salinicola halophyticus CR45 [NZ_PZJN01000009.1:283033..283532], <mnmG oriC2 <COG1126 (glnQ), 975 bp

GTATCCAGCCCTTCGGGTTCGACGAAGACCTGATGGCTGGTCTTGTCGGCGAAGCGATGAACCTTGTCCTCGATCGACGGGCAGTAACGCGGCCCGACGC
CCTCGATCACGCCGGAATACATCGGCGAGCGGTCGAGATTGGCGAAGATGATCTCGTGGGTGCGTTCGTTGGTGTGTCCGATATGGCAATCGACCTGCCG
CGGATGCTGATCGCGGCTGCCCAGATACGACATCACCGGCAACGGCGTGTCACCTGGCTGACGCTCGAGTTTGGAAAAATCGACGCTCTTGGCATCGATC
CGCGGCGGGGTACCGGTCTTCAGCCGATCGACCCGGAATGGCAGGGCTCGCAGCCGCTGAGCCAGCGCATTGGCCGGCGGATCTCCGGCTCTGCCGCCGG
CATGATGATCCAGACCGATGTGGATTACGCCACCCAGGAACGTTCCGGTACACAAAACCACGGTATCCGCCATGAAGCGAATGCCGGTCTGGGTCGTGAC
ACCGACGACGCTGTCGTTGTCGACGATCAGATCGTCGACGCCCTGCTGGAACAGGGTCAGATTGGGCTGGTTTTCCAGCATTCCACGGATGGCGGCCTTG
TAACGGACTCGATCGGCCTGAGCCCGTGTCGCTCGCACTGCCGGTCCCTTGCGCGAATTGAGCACGCGAAACTGGATGCCCGCCAAGTCGGTGGCCTGCG
CCATGGCGCCGCCCAGGGCGTCGATCTCCTTGACCAGATGGCTCTTGCCGATCCCGCCAATGGCGGGGTTGCAGGACATCTGGCCCAGCGTCTCGATGTT
GTGCGTCAGCAGCAGGGTGCGGACACCACTGCGAGCCGCAGCGAGTGCGGCTTCGGTGCCGGCGTGGCCACCGCCGATCACGATGACCTGAAAACGGTCG
GGATACTCCACGGTTTACCTCACGAGGTGCTGATGCGAACGCATCGAATCGGCACCGACAGCAAATGGATCTCGAGACCTCGCCAAGACGAGGTTCAGCG

CCGTCGCAGCGACGCTTTTCATGGTGGCCGAGAACGGTTGGCCACGAAAGCCGCCCAGTATAAACCTCCTGTGGGTAACCGTCAGGGGTTTTCCACACCC
GATTCGACGGTGAGTCGCTAACTCACAGAGTCATTATTAATTAATGAGAAATATATAAAGAAGTTCTGTTGTGGTAGTAACTAATGGTGTGGACATTTTC
**AGTGGATAA**GTGGAAATACTTTATATATTACGAGAACTTGCATTGTTATTCGATTGCTTGACAAGCATGGGGCATGCTGCTGACAGAGGCGTTGATTCCT
GTGGACAGAAGCCTTGTTTGTGCACAGCCCGGACTGCACACCTGTTATCCACCGCTGCATACGTCATCTGTTACCGCGGTGGTCAACACCCCCTCGATCG
TTCGCTAGCGTCGCGTGAGTCGGGGCTGCAGCAAGATCGGCAAAAAAACTATCCACAGGCGAAAAAAGCCTATCTGGATAACCTGAAACGGCAAATTTCG

ACCTTTTCGGGCGGATTTTTGATGCCGACAAGCGGGAATGAACGTCGCGGGTTCGGCCGGACGGGTCAGAATCGGCAAAATCGATGGAGAACGACGAGC
CAGACAGTCGCAGGAAAGACGGCGATTCGAACGCCAATGGTTCGAGTCCGAAACAGCCGACAATGAAACGGAAGGGCGATCAACATGGGAAACAGCGCTG
AAGAAAAGCGGCGGAAACAAGGGAGCGAGAAGTGAGGCAAGGAGCGCAGCGGCGCAGACAGCGAGCGTGATTCGGTCTGGAAATAGGTCTCCGCGGCACT
CTACAGAAGAGCCTATCCACCGGGCAGCAGATAGCGTGCAGGCGGCTTGAAGGAAGCCCTCAAGCCGCGCACAACCCGGTTATCGACTAGTGCTTGAGCA
CCCGCGCCAGAAACGCCTGAGTGCGCTCATGCTGCGGCTGGTCGAAGAGTTGATTGGGCGGTCCCTGTTCGACGATCCGGCCTTCGTGCACGAACACCAC
GCGATCGGCGACTTCCCGGGCAAAGCCCATCTCGTGGGTGACGATCATCATGGTCATGCCGTCGTTCACCAGCTCGCGCATGGCGTCGAGAACCTCGCCG
ATCATCTCCGGGTCCAGCGCCGAGGTCGGCTCGTCGAACAGCATCAGCCGAGGTTCCATTGCCAGCGCCCGGGCGAGCGCTACGCGTTGCTGCTGACCGC
CCGAAAGCTGAGTCGGATACTTGTCGGCCTGATCGGAAATGCCGACCCGCTCGAGCAGCTTGTCGGCGGTGGCATCGGCCTGCTGGCGCGTCCAGCCACG
GACCTTGCGTGGTGCCAACGTGACGTTGTCGCGGATCGACAGGTGCGGAAACAGATTGAACTGCTGGAACACCATCCCGACCTCGGTGCGCACGGCCTGC
AGCGATTTGCCGCTGGCCCCGTTGGGAATCAGGGTCTTGCCATCGACTTCGATGCGCCCGGACTGGAAGGGTTCGAGTCCGTTGACGCAGCGAATCAGCG

>Salinicola halophyticus CR45 [NZ_PZJN01000009.1:315279..315462], dnaA> ig dnaN>, 184 bp

GCCACTGCGCCGTGAATTCGAGCCCGTAAGGGATTGCGGGCGGCGTCCGGCGAGGCAAGAATGGGACGGCGCGGTCCGAGAGCTTACATCGGCGATCCCC
ATGCGGGATCGTTCGCGTTCAGGTCCATCGAGCCGGCCGGGAACTATTGCTTCAGCTTAGATTCATTCCAACTACGGAGCCCCA

WebSkew GCmin pos. 19684 (GCmin<>oriC1 30.1 kb; GCmin<>oriC2 1.5 kb; oriC2<>oriC1 28.6 kb)

>Salinicola socius SMB35 = DSM 19940 [NZ_MSDO01000012.1:49563..50062], <rpmH oriC1 dnaA>, 604 bp

CAGATTGCTGCCCTGAGCCGGATCGGCGTCAGCGGGCATGTCGTCCTGAATCGGTTTGGCGTTGCCGTCACTGTCCGTCGGTACGGAAAGCGTTCCGCCT
TCGCTGCTGTCGCTGTTACCCGCAGAGGCCGCCATCTCCGGCGCCGGGTTGTTCACGACGTTCTGACCGTAGTCCTGATTCCACTGCATGATCAGCAGAT
AGGCCAGTACGGCCAGCGGAATCACGAAAAGGAGTCGTTTCAAATCCATATAGAGGTCTGCCCGGTAAGGGAGTGGGTCAGTGGGCGCCGGGCATCGACG
GTCGACCGGCGGAGCTCGCCGATGCGTCCTGGCCTTCGGCGCAGGGAGGTCCGATCAGGATGACGAGGCGTCTTGGCGCGTCTGACGGGTGGACTTTTCC
AGGCGCTTCCACATGCCATTGAGCTGGCGAGAGAAGCTGGCGTTGTCCAACTCGTGTACCCCTCGTCGAGCGAGGATGACGATGTCCACTGCCGGTAAAC
GATGCTGTTGATGGCGAATGGATTCGCGCACGAGGCGTTTGAGACGGTTGCGGTCGACGGCACGGCGCACGTTCTTTCTGCTGAAGATCAACCCGATCCG
CGGATGGCCCAGAGCGTTGGGTCGGGCCAACGCGAGCAATCCTTTACCATGGACCTTGTAAGGCGCGTGGTCGAAAACGTGCTTGTAGTCCCCGGCGTTC
AGCAAGCGAAGCTGCCGGGGAAAACGCTGATCAGACACCCGGTATCCGATCAGGCGCTAAGACGCTTACGACCTTTGGCGCGACGACGAGCAATCACCTG
ACGGCCGTTTTTGGTGGCCATGCGGGCGCGAAAACCGTGGACGCGCTTACGCTTCAGAACGCTAGGCTGGAAAGTACGTTTCATAGCTGCTGATTCCCAA
TTAGTTGATTTCTAAAGAAAACGGGTTGAAACGGGATGCCCGATACGTTTGCGGGAAGCGGGCTCGCTCGACTCAAGAGCGGGAATTCTAGTGAATCCGC

GGTTCCGGTGCAATTTTTCCTTGAATTATCCACACCCTGTTCGGGGGCTGTCGCCGGCCGAAACGAATCTTGTGGGATGCTGAGGGAGTGCCGAATTCCC
GCTGGCTCGCGTGGAACGGACAGCGGTCGAATCCCGAGGTGAGATCGCAGTTCACTGACGGTGTCAATCACACCATGTGGTTAACGATGGGGATAACCTT
**TGTGTATAA**GCCATTTAAATCCTTTATAAACATTGGTTTATAGTAATTCTAAGGTTGTGGGGAAGTTGCGCGTGACCGGTGTACAGACTGGGTACGATGG
GCACGGGAATGGGCCGCTCGATACAGGTTATCCCAGGCTGTCACCAAGAAATCCACAGTTTGATGAGCCTGCTCCGCACTCAGTTATCCACAATTTTTCG
ATTTTATCCCCGTGTGGATAACCTCGTGGGTCGACGCTACACTGTGCGCCCGTTTTTCCATTCTTCTCTCAACGTTCGGTGAGGTCGTGTGTCGGTCGCT

CTCTGGCAACAATGTCTGAACTCGCTGCAGGATGAGTTGAGCTCTCAGCAATTCAATACCTGGATTCGTCCCCTGCAGGCAGAGGACGGCGAAGTGGACG
AACTCTGTCTATTGGCGCCCAACCGTTTTGTCCGCGACTGGGTCAGCGACAAGTATCTGAAACGGATCAACGAACTGTTGCGGGACGTCGCCTCCAGCAA
CAAGCCTCCCAAGGTTCGGCTGACGGTGGGTAGTCGTCGGGCCGCGGCGGTTTCCCCACGAGATCTGGGGGCGCCCGTGTCCGCCCCCAGCCGGCCACCG
CAACGGCCATCGTCGCCCAGTGTGCAGACGTCTTCGCGCTCCAACGATGCAGCCGAGGAAAGGCATATCGATCGATCGCGCGAACGGGAAGAGAGTGGGC
GGCGTGGCGGCCAGCGTCAGGTCCAGGTCGAGGGGAGCCTCAAGCATGGCAGCGGTCTCAATCCCAACTTCACGTTTGAGACCTTCGTCGAGGGTAAGTC
GAACCAGCTGGCGCGCGCCGCCTCCCGTCAGGTATCGGAGAATCCCGGCGGTGCCTACAATCCGCTGTTTCTCTACGGTGGCGTGGGTCTGGGCAAGACC
CACCTGATGCATGCGGTGGGCAATCATCTGGCGGGGCTTCGCGACAACGCCAAGATCGTCTATCTCCACTCCGAGCGTTTCGTCGCCGACATGGTCAAGG
CGCTGCAGATCAATGCCATCAACGACTTCAAGCGCTTCTACCGCAGCGTCGATGCCCTGCTGATCGACGATATCCAGTTCTTTGCCGGCAAGGAGCGTTC
CCAGGAAGAGTTCTTCCACACGTTCAACGCGTTGCTCGAGGGCGGCCAGCAGATGATTCTGACCTCGGATCGCTATCCGAAGGAGATCAGCGGTGTCGAG
GAGCGCCTGAAATCGCGCTTCGGCTGGGGCCTGACGGTGGCGATCGAACCGCCGGAGCTCGAGACCCGCGTGGCGATTCTGATGAAGAAGGCCGACCAGG

>Salinicola socius SMB35 = DSM 19940 [NZ_MSDO01000012.1:20935..21434], <mnmG oriC2 <COG1126 (glnQ), 911 bp

TCGAACGCACCACCTGCAGCTGGACGTCGAACGGCAGCGACGTGGAAATCCCGTTCGGGTAGAGCTCGTGGGTATCCAGGCCCTCCGGCTCGACGAAGAC
CTGATGGCTGGTCTTGTCGGCAAAGCGGTGAACCTTGTCCTCGATCGACGGGCAGTAACGCGGCCCGACGCCTTCGATCACGCCGGAATACATCGGCGAA
CGGTCGAGATTGGCGAAGATGATCTCGTGGGTGCGTTCGTTGGTATGGGCGATATGGCAATCGACCTGACGCGGATGCTGGTCGCGGCTGCCCAGATACG
ACATCACCGGCAAAGGCGTATCGCCAGGCTGGCGTTCGAGCTTCGAGAAATCGACGCTCTTGGCATCGATCCGCGGCGGCGTCCCGGTCTTAAGCCGATC
GACCCGGAACGGCATTGCCCGCAACCGGTTTGCCAGCGCATTGGCCGGCGGATCGCCGGCTCTGCCACCCGCATGATGATCCAGTCCGATGTGGATCACG
CCACCCAGGAAAGTTCCCGTGCACAGCACCACGGTATCCGCCATGAAGCGAATCCCGGTCTGGGTCCTGACGCCAACGACGGTCTCGTTTTCCACAACCA
GATCGTCGACACCCTGCTGGAACAGCGTGAGATTGGGCTGGTTTTCCAGCATGCCGCGAATGGCGGCCTTGTAACGAACCCGATCGGCCTGGGCTCGAGT
GGCACGAACCGCCGGTCCCTTGCGTGAATTGAGCACGCGAAACTGGATACCGGCCAGATCGGTTGCCTGAGCCATGGCGCCACCCAGGGCGTCGATCTCC
TTGACCAGATGGCTCTTGCCGATCCCGCCGATGGCGGGGTTGCAGGACATCTGGCCCAGCGTCTCGATGTTGTGCGTCAACAGCAGGGTGCGGGCGCCGC
CGCGGGCCGCTGCCAGAGCGGCTTCGGTGCCAGCGTGACCACCGCCGATCACGATGACCTGAAAACGGTCTGGATATTCCACGTTTTACCTCACGAGGTG

CTGAAACGATTAGCTCGAACTGGCACCCGACAGGGAATGGATCTCGAAAAGCCGCCCAGTATAAACCTCCTGTGGGTAACCGTCAGGGGTTTTCCACACC
CGATTGGCTGTGAGCCACTGGTCGACAGAGTCATTATTAATAAATGAGAAATATATAAAGAAGTTCTGTTGTGGTAGTAACTAATGGTGTGGACATTTTC
**AGTGGATAA**GTCGAAATACTCTATAGATTCCGAAAGCTTGCATTGTTATTCGATTGCTTGACAAGCATGGGGCATGCTGCTGACAGAGGCGTTGATTCCT
GTGGACAGAAGCTCGCTTTGTGCACAGCCCGGTCTGCACACGTGTTATCCACCGCTGCGTACGTTACCTGTTACCGCGGTGGTCAACACCCCCCCGATCG
TTCGCCAGCGTCGCGTGAGTCGGGGCTGCAGCAAGATCGTCAAAAAAACTATCCACAGGCGGAAAAAGCCGTTCTGGATAACCTGAAACGACAAATATCG

ATCTTTCCGAACGGATTTTCTGGTTCTGGAAAGCGGGAATGAGCATCCCGGGTTGGCTGGAAGGTCCAGAATCGGCCCAATCGATGAAGAACGGCGAGTC
GGACAGTCGAAGGAAAAGACGGAAATGTGAACGGAAAATGGTTCGAGTCCGAATCGGACTGCAAACGGAACGGAAGGGCGATCGACATCAGGGGACGGCG
CTGAGAAAAGCGCCAGGGAAAGGCGAGCGCGAGAAGCGAGACAGGCAGCGTTGCGGCGAGCAGGCAGCGGTCGTGACTCGGTCGTCAAATAGGTATTTGC
GGAACTCGATGGAAGGATCGATCCATCAAACAGCGTACAGCGGGCAGGCGGCTTGAAGAGAACCCTCAAGCCGCGCACGGCCCGGTTAATCGATTAGTGC
TTGAGAACCCGTGAGAGGAACGCTTGAGTGCGTTCATGCTGGGGCTGATCGAACAGTCGATCGGGCGGCCCCTGTTCGACGATCTGGCCTTCATGCACGA
AGACCACGCGGTCGGCGACTTCCCGGGCAAAGCCCATCTCGTGGGTGACGATCATCATGGTCATGCCGTCATTGACCAGTTCGCGCATGGCGTCGAGAAC
CTCGCCGATCATCTCCGGATCCAGCGCCGAGGTCGGCTCGTCGAACAGCATCAGCCGCGGCTCCATCGCCAGCGCCCTGGCGAGCGCCACGCGCTGCTGC
TGGCCCCCGGAGAGCTGCGTCGGATACTTGTCGGCCTGATCGGAGATGCCGACCCTTTCGAGTAGCTTGTCGGCGGTGACCTCGGCCTGCTGACGCGTCA
TGCCACGGACCTTGCGGGGCGCCAGGGTGACGTTGTCGCGGATCGACAGATGCGGGAACAGATTGAACTGCTGGAACACCATGCCGACTTCGGTGCGCAC
GGCCTGCAGCGATTTGCCGCTGGCGCCATTGGGCATCAGCGTCTTGCCGTCGACCTCGATGCGGCCGGACTGGAACGGCTCGAGTCCGTTGACGCAGCGA

>Salinicola socius SMB35 = DSM 19940 [NZ_MSDO01000012.1:51530..51860], dnaA> ig dnaN>, 331 bp

GTCGCTATCGCGTCATTTCCGCAACCACAGGGATTGCGGGCGGCGCCTGCCGAGGCAAGAATGGGATGGTGCGATCCGGCAGCTTTCGTCACCAGTCCGG
CTCGGGTCTATTGATCCGACGATTTTGGTTTATTGACTCAGTTTTATTGACTCAGCTTAGGTGCTGTCTGAAAAGTCGACGAGCGAAGGCCAGGCAAGGC
AAAAATCGGCGAAAGCGGACCGGGCTGGCGCACCAGCCGATTTTTAACGCCGCATGGGCGAGCGTAGTACCTTTCAGACAGTGCCTAGGCTTATTGGTCC
AGCTTAGATTCATTCCAACTACGGAGCCCCA

WebSkew GCmin pos. nd (GCmin<>oriC1 nd; GCmin<>oriC2 nd; oriC2<>oriC1 nd)

>Salinicola salarius DSM 18044 [NZ_PZJT01000012.1:8783..9282], <rpmH oriC1 dnaA>, 606 bp

TCAGGTTGCTGCCTCGGGCCGGATCGGCGCCGGCGGGCATGTCATCCTGGATCGGCTTGGCATTGCCGTTGCTGTCCGTCGGTGCGGACAGGGCGCTGCC
TTCACCATTGTCGCTGTTACCCGATGACGCTGCCATCTCCGGCGCCGGGTTGTTCACGACGTTCTGACCATAGTCTTGATTCCACTGCATGATCAGCAGA
TAGGCCAGCACGGCCAGCGGAATCACGAAAAGGAGTCGTTTCAAATCCATATAGAGGTCTGCCCGGTAAGGGAATGGGTCAGTGGGCGCCGGGCATCGGC
AGTCGACCGGCAAAATTCGCCGGTACGCCCCGGCTTTCGGCGCGGGGGAGTCCGATCAGGATGACGAAGCGTCTTGTCGCGTCTGACGAGCGGACTTTTC
CAGGCGCTTCCACATGCCATTGAGCTGGCGAGAGACGCTGGCGTTGTCCAACTCGTGTACCCCTCGCCGAGCGAGTATCACGATATCCACGGCCGGTAAA
CGATGCTGCTGATGGCGGATGGATTCGCGCACGAGGCGTTTGAGACGGTTGCGGTCGACGGCACGACGCACGTTCTTCTTGCTGAAGATCAACCCGATCC
GCGGATGGCCCAACGCGTTGGGACGAGCCAACGCCAGCAATCCTTTACCATGGACCTTGTAAGGAGCGTGGTCGAAAACGTGCTTGTAGTCCCCGGCATT
CAGCAGACGCAACTGCCGGGGAAAACGCTGATCGGACACTCGGCGTCCGATCAGGCGCTGAGACGCTTGCGGCCTTTGGCGCGGCGACGAGCAATGACCT
GACGGCCGTTCTTGGTGGCCATACGGGCGCGAAAACCGTGGACGCGCTTACGCTTCAGAACGCTGGGCTGGAAGGTGCGTTTCATAGCTGCTAATTCCCA
ATTAGTTGATTTCTAAAGAAAACGGGTTGAAGCGGGGTGCCCGACACGTTTGCAGGAAGCGGGCTCACTCGACTCAAGAGCGGGAATTCTAGTGAATCCG

CGGTGCCGGTGCAATTTTTCCGTGAATTATCCACACCCTGTTACGGGTCAAGCGCGGGCCCTGACGTGTCGGGTCGGATACTCGTGCCGCAACGGATCCG
AACTGGGCGACCCCGCGATGATCGGCCTTGGAGGCGTACCGTGGATGACCGTTTCCGAAGTGGGCTTGTCGCTCATGTGGTTAACAACGGGGATAATGTT
**TGTGGATAA**GCTGTTTTTTCCCCTTTATAACAAAGGCCTATGTGTTGATTCAGGTTGTGGGGAAGTTGCGCATTAGCGGTGGGCGGGATGGATGCCTTCG
AAGATGAAGGCAAGCGCCTGCAACAGGTTATTCGCAGCTATCACCAGAAAATCCACAGATTTCGGCCTGAAGGACGGCAACTTATCCACAATTGCCTCGA
ATTTACCCTCGTGTGGATAACCTGGCAGGTCGACGCTACAATGTGCGCCCGTTTTTTCCATTCTCGTTTTCTCAACGTTCGGTGAGGTCGTGTGTCGGTC

GCTCTCTGGCAACAATGTCTGAACTCGCTGCAGGATGAGTTGAGCTCTCAGCAATTCAATACCTGGATTCGTCCCCTGCAGGCAGAGGACGGCGAAGTGG
ACGAACTCTGTCTATTGGCGCCCAACCGTTTTGTCCGTGACTGGGTCAGCGACAAGTACCTGAAGCGAATCAACGAACTGTTGCGTGAGGTCGCGTCCAG
CAACAAACCGCCCAAGGTGCGCTTGACGGTGGGAAGTCGCCGGGCAGCCGCGGTCTCTCCGCGAGATCTGGGTGCCCCCGTTTCCGCGCCCAGTCGGCCG
CCGCAACGGCCATCCTCGCCCAGTGTGCAGACGGCGACGCGGTCTGACGAGGCGGCCGAGGAAAGACAGGTCGACCGCTCGCGGGAACGGGAAGAGAACG
GCCGGCGCGGAGGGCAGCGTCAGGTCCAGGTCGAGGGCAGCCTCAAGCATGGCAGCGGGCTCAATCCCAATTTCACGTTCGAAACCTTCGTCGAAGGCAA
GTCGAACCAGTTGGCCCGCGCCGCCTCGCGCCAGGTGTCCGAAAACCCTGGCGGTGCCTATAATCCGTTGTTTCTCTACGGTGGCGTGGGTCTGGGCAAG
ACCCACCTGATGCATGCGGTGGGCAATCATCTGGCAGGGCTGCGCGACAACGCCAAGATCGTCTATCTGCACTCCGAGCGCTTCGTCGCCGACATGGTCA
AGGCGCTGCAGCTCAATGCCATCAATGATTTCAAGCGCTTCTACCGGAGCGTTGATGCCCTGCTGATCGACGATATCCAGTTCTTCGCCGGCAAGGAGCG
TTCCCAGGAAGAGTTCTTCCATACGTTCAATGCCCTGCTGGAAGGCGGCCAGCAGATGATCCTGACCTCGGATCGCTATCCCAAGGAGATCAGCGGGGTC
GAGGAGCGGCTGAAATCCCGCTTCGGCTGGGGCCTGACGGTGGCGATCGAGCCGCCGGAGCTCGAGACGCGCGTGGCGATCCTGATGAAGAAGGCGGATC

>Salinicola salarius DSM 18044 [NZ_PZJT01000021.1:39482..39981], <mnmG oriC2 <COG1126 (glnQ), 761 bp

CGAGCGCACGACCTGTAGCTGAACGTCGAACGGCAGGGACGTGGAAATCCCGTTCGGGTAGAGCTCGTGAGTGTCCAGACCTTCGGGTTCGATGAACACC
TGATGACTGGTCTTGTCGGCGAAGCGGTGGACCTTGTCCTCGATCGACGGGCAGTAGCGCGGCCCCACGCCTTCGATCACGCCGGAATACATCGGCGAAC
GGTCGAGATTGGCGAAGATGATCTCGTGGGTTCGCTCGTTGGTATGGGCGATATGACAATCGACCTGGCGCGGATGCTGTTCGCGGCTGCCCAGATACGA
CATTACCGGCAACGGCGTGTCGCCGGGCTGTGGCTCGAGCCTGGAGAAATCGACGCTCTTGGCATCGATCCGCGGTGGTGTCCCGGTTTTCAGCCGATCG
ACCCGGAACGGCAGCGCACGCAGACGTTCCGCCAACGCGTTGGCCGGCGGATCGCCGGCGCGACCGCCGGCATGATGATCCAGCCCGATGTGAATCACGC
CGCCTAGGAAGGTACCGGTACACAGCACCACGGTGTCCGCCATGAAGCGGATGCCGGTCTGGGTCCGGACACCGACGACGGTGTCGTTTTCCACAATCAG
ATCGTCGACACCCTGCTGGAACAGCGTGAGATTCGGCTGGTTCTCCAGGATGCCACGAATGGCGGCCTTGTAGCGCACGCGATCCGCCTGGGCTCGAGTG
GCGCGAACCGCCGGCCCCTTGCGGGAATTGAGCACGCGAAACTGGATGCCGGCCAGGTCGGTTGCCTGTGCCATGGCACCACCCAGGGCATCGATCTCCT
TGACCAGATGGCTCTTGCCGATCCCGCCGATGGCGGGATTGCAGGACATCTGGCCCAGCGTCTCGATGTTATGCGTCAGCAGCAGGGTGCGGGCACCGCC
ACGTGCCGCTGCCAGAGCGGCTTCGGTACCGGCATGGCCACCGCCGATCACGATGACCTGAAAACGGTCGGGATACTCCACGGTTTACCTCACGAGGTGC

TGAAAACGAGTAGCTCGAACAGGCACCCGACAGTGAATGAATCTCGAAAAGCCGCCCAGTATAAACCTCCTGTGGGTAACCGTCAGGGGTTTTCCACACC
CGATTGTCTTCAGCCCACCAGCCAGCAGAGTCATTACTAAATAAAGAAAAATATATAAAGAAGTTCTGTTGTGGTAGTAACTAATGGTGTGGATGCTTTC
**AGTGGATAA**GTCGAATTACTCTATGGATGA**T**AACCACTTGCATTGTTATTCGATTGTTTGAAAAGCACTGGGGAGGCTGCTGACAACGGCGTTGGTTCCT
GTGGACAGAAGATCGCTTTGTACACATCGCCAGCCGCACACGGGTTGTCCACCGCCGCATACGCCAACTGTTACCGCGATGGTCAACAAACCGCGGATTC
CTCGCCAGCGCCGGACGAATCAGGGTTACAGCCAGATAGGCGAAAAAACTATCCACAGGCGGAAAAACGCGCCTGGGATAACCATGAATGACGAAATCGG

CCTTGGCCGACCCGATCTTTCGACCCTGAGGCGAACCGATGGCTCGGCGTTTATCGTTCGAGCCAGGAGACGAAGCGGTAAAACGCGGCGAAGAGGACAG
GCGGAATGGCGATGGAAATGAAGAAGCGCAGTGCAGGAAGGGCTGAGCGTGCTGGAGGCGGGTAGAGCGTACCTCGAGGAAGGCGGACACCGGAAATGCC
TCCGGTGCCCGCGGTTTTCGGAACGCTCGGCCCGCGTTCGGGCTAGTGCTTGAGCACCCGTGAAAGGAACGCTTGCGTGCGTTCGTGGCGGGGATGATCG
AAGAGCTGCTCCGGCGGGCCCTGCTCGACGATCTGGCCGCCGTGGACGAAGACCACACGATCGGCGACCTCGCGAGCGAAGCCCATCTCGTGAGTGACGA
TCATCATGGTCATGCCGTCGTTGGCCAGTTCGCGCATGGCGTCGAGCACCTCGCCGATCATCTCCGGATCCAGCGCGGAGGTGGGCTCGTCGAACAGCAT
CAGCCGCGGCTCCATGGCCAGCGCTCGGGCGAGCGCCACGCGTTGCTGCTGCCCGCCGGAGAGCTGGGTCGGATACTTGTCGGCCTGATCGGCAATGCCG
ACCCGTTCGAGCAGCTTGTCGGCGATGGCGTTGGCCTGGTCGCGCGTGACGCCGCGGACCTTGCGCGGTGCCAGCGTGACGTTGTCGCGGATCGACAGAT
GCGGGAACAGGTTGAACTGCTGGAACACCATGCCGACTTCGGTGCGCACCGCCTGCAGGGACTTGCCGCTGGCGCCGTTGGGCATCAGGGTCTTGCCGTC
GACTTCGATACGCCCGGACTGAAACGGTTCGAGCCCGTTGACGCAGCGGATCAGGGTCGATTTTCCCGAGCCGCTGGCGCCGATGATCACCACGACTTCG
CCGTGGGCCACGCTGAGATCGATCTCCTGCATCACGTGCAGGTTGCCGAAGTGCTTGTCGACCTTGTCGAGCCGGACGATCGGCGACGCATCCGCCGTCC

>Salinicola salarius DSM 18044 [NZ_PZJT01000012.1:10753..10921], dnaA> ig dnaN>, 169 bp

WebSkew GCmin pos. 277632 (GCmin<>oriC1 35.2 kb; GCmin<>oriC2 8.4 kb; oriC2<>oriC1 26.8 kb)

>Salinicola lusitanus CR50 [NZ_PZJM01000009.1:c242229..242728], <rpmH oriC1 dnaA>, 605 bp

CAGGTTGCTGCCTTTGGCCGGATCGGCGTCGGCGGGCATGTCGTCCTGAATCGGCTTGGCTTTACCGTTGCTGCCCGTCGGCGCAGACAGGGCGCTGCCC
TCGCCGTCATTGCCGTTACCCGATGAGGCCGCCATCTCCGGCGCCGGGTTGTTCACGACGTGCTGACCGTAGTCCTGATTCCACTGCATGATCAGCAGGT
AGGCCAGCACGGCCAACGGAATCACGAAAAGGAGTCGTTTCAAATCCATATAGGGGTCTGCCCGGTAAGGGAATGGGTCGGTGGGCGCCGGGCATCGGCG
ATCGACCGGCGAGGTTCGCCGGTGCTCCCCGGCCTTCGGCGCGGGGATGTCCGATCAGGAAGACGAAACGTCTTGACGCATCTGGCGAGCGGACTTCTCC
AGGCGTCGCCACATGCCATTGAGCTGGCGGGCGACGCTGGCGTTGTCCAGCTCATGTACCCCTCGTCGCGCGAGTATGACGATATCCACTGCCGGTAAAC
GATGCTGTTGATGGCGAATGGATTCGCGCACGAGGCGCTTGAGGCGATTGCGGTCGACGGCACGGCGCACGTTCTTCTTGCTGAAGATCAACCCGATCCG
TGGATGGCCCAGAGCGTTGGGTCGGGCCAACGCCAGCAATCCTTTACCATGGACCTTGTAGGGAGCGTGGTCGAAGACGTGCTTGTAGTCCCCGGCATTC
AGCAACCGCAACTGCCGGGGAAAACGCTGATCGGACACCCGGCGTCCGATCAGGCGCTCAGGCGTTTACGGCCTTTGGCACGGCGACGGGCAATGACCTG
ACGGCCGTTCTTGGTGGCCATGCGGGCGCGGAAACCGTGGACGCGCTTACGCTTCAGAACGCTGGGCTGGAAGGTGCGTTTCATAGTTGCTAATTCCCAA
TTAGTTGATTTCTAAAGAAAACTGGTTGAAGCGGGACGCCCGGCGCATTCGCGGGAAGCGGGCTCACTCGACTCAAGAGCGGGAATTCTAGTGAATCCGC

GGTTCGGGTGCAATTTTTCCGTGAATTATCCACACCCCGTTGTGAGTGGATCGCCAGCCGTTACGCCCCGAACGGGAGACCCGTGGCGTGAGTGGCCATC
GACGATTGTCGTGGCGATGCCGGTAATCGGCTCGGGATGCGAGATCGAAGTTGCTCGAGGGTGTCAATCACACCATGTGGTTAACGACGGGGATAACCTT
**TGTGGATAA**CCTTATAAAATCAATATTTATCATTGGTTTAATGGCTTTTTTTGGGTTGTGGGGAAGTTGCGAACGAGCCGTGCATAGGGTGAGGATCGCC
GGCGTGTGGCAAGACCGTCCACACCGCCTTGTCCACGCGGATTACCCTGGACATCCACAGTATTGCCAGCGGTGACGGGCGAGTTATCCACAGTTGTTGG
AAAGTCGACTTCGTGTGGATAACCTGGCAGGCCGACGCTACAATGTGCGGCCGTTTTTTCTATTCTCTCTCAACGTTCGGTGAGGTCGTGTGTCTGTCGC

TCTCTGGCAACAATGTCTGAACTCGCTGCAGGATGAATTGAGCTCTCAGCAATTCAATACCTGGATTCGTCCTCTGCAGGCCGAGGACGGCGAAGTGGAC
GAACTCTGTCTGTTGGCCCCCAACCGATTCGTCCGTGATTGGGTCAGCGACAAGTACCTGAAGCGGATCAACGAACTGTTGCGCGAGGTTGCCTCCAGCA
GCAAGCCTCCCAAGGTCCACCTGACGGTGGGGAGCCGGCGGGCCGCGGCGGTGTCGCCGCGGGATCTTGGCGCGCCGGTTTCCGCCCCGAGTCGTCAGCC
TCAGCGATCCACGGCGCCGGGGATTCAGACGAGCCCGCGCTCCAACGAGGCGGCCGAGGAGAAGCAGCTCGATCGAACCCGTGAACGCGAAGAGAGTGGA
CGCCGGGGCAGCGAACGCCAGGTCCAGGTCGAGGGTAGCCTCAAGCACGGCAGCGGACTCAATCCCAATTTCACCTTCGAAACCTTCGTCGAGGGCAAGT
CGAACCAGCTGGCGCGGGCCGCCTCGCGCCAGGTATCCGAGAACCCCGGCGGGGCCTACAATCCGCTGTTCCTCTACGGCGGCGTGGGTCTGGGCAAGAC
CCACCTGATGCATGCGGTGGGCAACCACCTCGCGGGTCTCAGAGACAACGCCAAGGTGGTCTATCTCCACTCCGAGCGCTTCGTCGCCGACATGGTCAAG
GCGCTGCAGCTCAACGCGATCAATGATTTCAAGCGTTTCTATCGCAGCGTCGATGCGCTGCTGATCGACGATATCCAGTTCTTTGCCGGCAAGGAGCGCT
CCCAGGAGGAGTTCTTCCACACCTTCAACGCCTTGCTGGAAGGCGGACAGCAGATGATCCTGACCTCGGATCGCTATCCCAAGGAGATCAGCGGCGTCGA
GGAGCGCCTGAAATCGCGCTTCGGCTGGGGGTTGACGGTGGCGATCGAGCCGCCGGAACTCGAGACCCGGGTGGCGATCCTGATGAAGAAGGCGGATCAG

>Salinicola lusitanus CR50 [NZ_PZJM01000009.1:c268991..269490], <mnmG oriC2 <COG1126 (glnQ), 749 bp

CCAGGCCCTCCGGTTCGACGAAGACCTGATGGCTGGTCTTGTCGGCGAAGCGGTGGACCTTGTCCTCGATCGACGGGCAGTAACGCGGGCCCACGCCCTC
GATCACGCCGGAGTACATCGGCGAACGATCGAGATTGGCGAAGATGATCTCGTGGGTACGCTCGTTGGTGTGGGCGATGTGGCAGTCGACCTGACGCGGG
TGCTGCTCGCGGTTGCCCAGGTAGGACATCACCGGCAGCGGCGTGTCGCCGGGCTGGCGCTCGAGCCTGGAGAAATCGACGCTCTTGGCATCGATCCGCG
GTGGCGTGCCGGTCTTCAGCCGATCGACCCGGAACGGGAGCGCTCGCAGCCGCTGGGCCAGCGCATTGGCCGGCGGATCGCCGGCGCGGCCACCGGCATG
GTGATCCAGACCGATATGGATCACACCGCCCAGGAAGGTTCCGGTACACAGCACCACGGTATCCGCCATGAAGCGAATGCCGGTCTGGGTGGTGACGCCG
ACGACGGTCTCGTTTTCCACAACCAGATCGTCGACGCCCTGCTGGAACAGCGTGAGATTGGGCTGATTTTCCAGCATGCCGCGAATGGCGGCCTTGTAGC
GGACGCGATCGGCCTGGGCTCGGGTGGCCCGAACCGCCGGGCCCTTGCGTGAGTTGAGCACGCGAAACTGGATGCCGGCCAGGTCGGTAGCCTGGGCCAT
GGCCCCACCCAGGGCATCGATCTCCTTGACCAGATGGCTTTTGCCGATCCCGCCGATGGCGGGATTGCAGGACATCTGGCCCAGGGTCTCGATGTTGTGC
GTCAGCAGCAGCGTACGGGCACCACTTCGTGCCGACGCCAGCGCGGCTTCGGTTCCGGCATGGCCACCGCCGATCACGATGACCTGAAAACGGTCGGGAT
ACTCCACGGTTTACCTCACGAGGTGCCCGCAGCGCTTTGGCTCGAGCGGGCACCCGACGATGAATGGATCACGAAATCGGCCAGGCCGATTTTACCGTCC

TGTTCACGACGGCGATGGCACGGCCCCGGGGGGTTCGATGGCCACGAAAGCCGCCCAGTATAAACCTCCTGTGGGTAACCGTCAGGGGTTTTCCACATCG
GGATCGAGGCGATCCGTCAGTCATCAGATTCATTATTAAAAAAATGAAAAATATATAAAGAAGTTCTGTAGTGATAGTGACTATTGGTGTGGACACTTTT
**CGTGGATAA**GTAAAAAATCACCAATAGAATCATATTCTTGGATTGTTATTCGATTGTTTGAAAAGCTCTGGCGAGCCTGCTGACAAGGGCGCTGATTCCT
GTGGACAGCTTCGCCTCTTGTACACAGGCGCGACCGCGCACGGGTTATCCACCGCTGCATACGGTAGCTGTTACCGCGACGGTCAACAAACCCCCGATCA
CGCCAGAGCGGCTGAGGCGTCAGGGCTGGAGACGAATCGACGAAAAAATTGTCCACAGGCGAAAAATACGCCCCGGGATAACTCACGAGCGATGAAATCC

GGGTCACGCCACCGTTTTTCCGGCTCCGACGATCTCGATCGCACGACGGCCAGAACCGGTCTGGACGAATCAATGAAAGGGAAGAGAGGGAGAGGGTGTA
GGGGGTGGGCGGCACGGAGGAGATCCATCCGTGCCTGCTATTTGAGAAACCGATCGCTAGTGCTTGAGCACGCGGGAGAGGAACGCCTGGGTGCGCTCGT
GGCGGGGCTGATCGAACAGCTGGTCGGGTGGACCCTGTTCGACGATCTGGCCGCCGTGGACGAATACCACGCGATCGGCGACCTCGCGGGCGAAGCCCAT
CTCGTGGGTGACGATCATCATGGTCATGCCGTCGTTGACCAGTTCCCGCATGGCATCCAGCACTTCACCGATCATCTCCGGATCCAGCGCCGAGGTCGGC
TCGTCGAACAGCATCAGACGAGGTTCCATGGCCAACGCCCGGGCCAGCGCCACGCGCTGTTGCTGCCCCCCGGAGAGTTTAGTCGGATGCTTGTCGGCCT
GATCCGAGATGCCGACCCGCTCCAGCAGCCTGTCGGCGATGGCATCGGCCTGCCGCCGGGTCATGCCGCGGACCTTGCGGGGCGCCAGGGTGACGTTGTC
GCGGATCGACAGGTGCGGGAACAGGTTGAACTGCTGGAACACCATCCCGACCTCGGTACGGACTGCCTGGAGCGACTTGCCGCTGGCGCCATTGGGCAGC
AGCGTCTTGCCGTCGACCTCGATACGACCGGACTGGAAGGGCTCGAGGCCATTGACGCAGCGGATCAGCGTCGATTTTCCCGAACCACTGGCACCGATGA
TCACCACGACCTCGCCCTGGGCCACGCTCAGGTCGATGTCCTGCATCACGTGCAGACGGCCGAAGTGCTTGTTGACCCCTTCGAGACGCACGATCGGCGA
CGATCCGGACGATCGAGCGGTCGCTTGCCGGTTTGCGCTATCGTTCATCCCACTAACCCCTTGCGTTCGAGCAGACTCAGAGCCAACGACAGACTCAGGG

>Salinicola lusitanus CR50 [NZ_PZJM01000009.1:c240595..240760], dnaA> ig dnaN>, 166 bp

GGCGGCGCGTGACGAATTCCCTGGCCGCGGGGATTGCTAGCGACGACCGGCACGGCAAGAATGAGACGGCGCGGCCCGACGGTCTGCTCGTCCGTCCCAT
GATGCAAAGCGATCCACCGGTTACCGGATTCAACTTCAGGCTTATTGAACCAACAACGGAGCCCCA

WebSkew GCmin pos. 154200 (GCmin<>oriC1 35.4 kb; GCmin<>oriC2 8.6 kb; oriC2<>oriC1 26.8 kb)

>Salinicola acroporae LMG 28587 [NZ_PZJW01000003.1:118529..119028], <rpmH oriC1 dnaA>, 604 bp

CAGGTTGCTGCCTTTGGCCGGATCGGCGTCGGCAGGCATGTCGTCCTGAATCGGCTTGGCTTTGCCGTTGCTACCCGTCGGTGCGGACAGGGCGCTGCTC
TCGCCGTCATTGCCGTTACCCGATGAAGCCGCCATCTCCGGCGCCGGGTTGTTCACGACGTGCTGACCGTAGTCCTGATTCCACTGCATGATCAGCAGGT
AGGCCAGCACGGCCAACGGAATCACGAAAAGGAGTCGTTTCAAATCCATATAGGGGTCTGCCCGGTAAGGGAATGGGTCGGTGGGCGCCGGGCATCGGCG
ATCGACCGGCGAGGTTCGCCGGTGCTCCCCGGCCTTCGGCGCGGGGATGTCCGATCAGGAAGACGAAACGTCTTGACGCATCTGGCGAGCGGACTTCTCC
AGCCGTCGCCACATGCCATTGAGCTGGCGGGCGACGCTGGCGTTGTCCAGCTCATGTACCCCTCGTCGCGCGAGTATGACGATATCCACTGCCGGTAAAC
GATGCTGTTGATGGCGAATGGATTCGCGCACGAGGCGCTTGAGGCGATTGCGGTCGACGGCACGGCGCACGTTCTTCTTGCTGAAGATCAACCCGATCCG
TGGATGGCCAAGCGCGTTGGGTCGGGCCAACGCCAGCAATCCTTTACCATGGACCTTGTAGGGAGCGTGGTCGAAGACGTGCTTGTAGTCCCCGGCATTC
AGCAATCGCAGTTGCCGGGGAAAACGCTGATCGGACACCCGGCGTCCGATCAGGCGCTCAGGCGTTTACGGCCTTTGGCACGGCGACGGGCAATGACCTG
GCGACCGTTCTTGGTGGCCATGCGGGCGCGAAAACCGTGGACGCGCTTACGCTTCAGAACGCTGGGCTGGAAGGTGCGTTTCATAGTTGCTGATTCCCAA
TTAGTTGATTTCTAAAGAAAACTGGTTGAAGCGGGACGCCCGACGCATTCGCGGGAAGCGGGCTCACTCGACTCAAGAGCGGGAATTCTAGTGAATCCGC

GGTTCGGGTGCAATTTTTCCGTGAATTATCCACACCCCGTTGTGAGTGAATCACCAGCCGTCATGCCTTGGACGGAAGACCGATGACGTGGGCGGCCATC
GACGATTGTGGTGGCGATGCCGGCAACCGGATGGAGATGCGAGATCGCAGTTCCGCGAGAGTGTCAATCACACCATGTGGTTAACGACGGGGATAACCTT
**TGTGGATAA**CCCATAAAATTCATTTTGTATCATGAAGTTGTTAGGTTTTTTGGGTTGTGGGAAGTTGCGCATGAGCCGTGCATAGGGTGAGGATCGCCA
ACTGGCGACAAGACCGCCCACACCGTCTTGTCCACGCGGATTGCCCTGGACATCCACAGCATTGCCAGCGATGAGCGTCGACTTATCCACAGTCGTTGGA
AAATCGACTTCGTGTGGATAACCTGGCAGGCCGACGCTACAATGTGCGGCCGTTTTTTCCATTCTCTCTCAACGTTCGGTGAGGTCGTGTGTCTGTCGCT

CTCTGGCAACAATGTCTGAACTCGCTGCAGGATGAGTTGAGCTCTCAGCAATTCAATACCTGGATTCGTCCTCTGCAGGCGGAGGACGGCGAAGTGGACG
AACTCTGTCTGTTGGCCCCCAATCGATTCGTCCGTGACTGGGTCAGCGACAAGTACCTGAAGCGGATCAACGAACTGTTGCGCGAGGTTGCCTCCAGCAG
CAAGCCTCCCAAGGTCCATCTGACGGTGGGTAGCCGACGGGCCGCGGCGGTGTCGCCGCGGGATCTTGGTGCGCCGGTTTCCGCCCCGAGTCGTCAGCCT
CAGCGATCCGCGGCCCCGGGGATACAGACGAGCCCGCGCTCCAACGAGGCGGCCGAGGAGAAGCAGCTCGATCGAGCCCGTGAACGCGAGGAGAGTGGAC
GGCGGGGCAGCGAGCGCCAGGTCCAGGTCGAGGGCAGCCTCAAGCACGGCAGCGGACTCAACCCCAATTTCACCTTCGAAACCTTCGTCGAGGGCAAGTC
GAACCAACTGGCGCGGGCCGCTTCGCGTCAGGTATCCGAGAACCCAGGCGGGGCCTACAATCCGCTGTTTCTCTACGGTGGCGTGGGTCTGGGCAAGACC
CACCTGATGCATGCGGTGGGCAACCATCTCGCGGGTCTCAGAGACAACGCCAAGGTGGTCTATCTTCACTCCGAGCGTTTCGTCGCCGACATGGTCAAGG
CGCTGCAGCTCAACGCGATCAACGATTTCAAGCGCTTCTATCGCAGCGTCGATGCGCTGCTGATCGACGATATCCAGTTCTTTGCCGGCAAGGAGCGCTC
CCAGGAGGAGTTTTTCCACACGTTCAACGCCTTGCTGGAAGGCGGGCAGCAGATGATCCTGACCTCTGATCGCTATCCCAAGGAGATCAGCGGCGTCGAG
GAGCGCCTGAAGTCGCGTTTCGGCTGGGGGTTGACGGTCGCGATCGAGCCGCCGGAACTCGAAACCCGGGTGGCGATCCTGATGAAGAAGGCGGATCAGG

>Salinicola acroporae LMG 28587 [NZ_PZJW01000003.1:145338..145837], <mnmG oriC2 <COG1126 (glnQ), 748 bp

ATCCAGGCCTTCCGGTTCGACGAATACCTGATGGCTGGTCTTGTCGGCGAAGCGGTGGACCTTGTCCTCGATCGACGGGCAGTAACGCGGGCCCACGCCT
TCGATCACGCCGGAATACATCGGCGAACGATCGAGATTGGCGAAGATGATCTCGTGGGTGCGTTCGTTGGTATGGGCGATATGGCAATCGACCTGACGCG
GATGCTGCTCGCGGCTGCCCAGGTAGGACATCACCGGCAGCGGCGTGTCACCGGGCTGGCGCTCGAGCCTGGAGAAATCGACGCTCTTGGCATCGATCCG
CGGCGGCGTGCCGGTCTTCAGCCGATCGACCCGGAACGGGAGCGCTCGCAGCCGCTGGGCCAGCGCATTGGCGGGCGGATCGCCGGCGCGACCGCCGGCA
TGGTGGTCCAGACCGATATGGATCACGCCGCCCAGGAAGGTCCCGGTACACAGCACCACGGTATCCGCCATGAAGCGAATGCCGGTCTGGGTGGTGACGC
CGACAACGGTATCGTTTTCCACAACCAGATCGTCGACACCCTGCTGGAACAGCGTGAGGTTGGGCTGATTTTCCAGCATGCCGCGAATGGCGGCCTTGTA
GCGGATCCGATCGGCCTGGGCCCGGGTGGCCCGAACCGCCGGGCCCTTGCGCGAGTTGAGCACGCGAAACTGGATGCCGGCCAGATCGGTGGCCTTGGCC
ATGGCCCCACCCAGGGCATCGATCTCCTTGACCAGGTGGCTTTTGCCAATCCCGCCGATGGCGGGATTGCAGGACATCTGGCCGAGGGTCTCGATGTTGT
GTGTCAGCAGCAGCGTACGGGCACCACTTCGCGCCGACGCCAGCGCAGCTTCGGTTCCGGCATGGCCACCGCCGATCACGATGACCTGAAAACGGTCGGG
ATACTCCACGGTTTACCTCACGAGGTGCCCGCGGCGCTCTGGCTCGAGCGGGCACCCGACGATGAATGGATCACGAAATCGGCCAAGCCGATTTTACCGT

CCTGTTCACGACGGCGATGGCATGGCCCCTAGGGTTCGGTGGCCACGAAAGCCGCCCAGTATAAACCTCCTGTGGGTAACCGTCAGGGGTTTTCCACACC
GGGATCGAGGCGATCGGTCAGTCATCAGATTCATTATAAAAAAATGAAAAATATATAAAGAAGTTCTGTTGTGATAGTGACTATTGGTGTGGACACTTTC
**CGTGGATAA**GTAAAAATTCGATAACAAAATCATGCTCTTGGATTGTTATTCGATTGCTTGACAAGCTCTGACGAGCCTGCTGACAAGGCGTTGATTCCT
GTGGACAGCTTTGTCTCTTGTACACAGGCACGACCGCGCACGGGTTATCCACCGCTGCATACGGTAGCTGTTACCGCGACGGTCAACAAACCCCCGATCG
TGCCGAGGTGCCTGAGGCGTCAGGGCTGGAGGCAGATCGGTGAAAAAACTGTCCACAGGCAAAAAATACGCTTCGGGATAACCCACGGGTGATGAAATCC

GGGGCACGCTATCGATTTTTCCGGCCCCGACGAGCTCGATCGCACGACGCCTGGGACCGAATCCGGTCTGGATGAGCCAATAAAAGGGAGGAGA**C**AAAGA
GGGATCGGGGGAGGCGCGGAGGAGACCCATCCGCGCTTTCTGTTTGTGAACCGAGCGCTAGTGCTTGAGCACGCGGGAGAGAAACGCTTGGGTGCGCTCG
TGGCGGGGCTGATCGAACAGCTGATCGGGTGGGCCTTGTTCGACGATCTGGCCGCCGTGGACGAATACCACGCGATCGGCGACCTCGCGGGCGAAGCCCA
TTTCGTGGGTGACGATCATCATGGTCATGCCGTCGTTGACCAGTTCCCGCATGGCATCCAGCACTTCACCGATCATCTCCGGATCCAGCGCGGAGGTCGG
CTCGTCGAACAGCATCAGGCGAGGTTCCATGGCCAACGCCCGGGCCAGCGCCACCCGCTGTTGCTGTCCCCCGGAGAGTTGAGTCGGATGTTTGTCGGCC
TGATCCGAGATGCCGACCCGCTCCAGTAGCCTGTCGGCGATGGCATTGGCCTGCCGCCGGGTCATGCCGCGGACCTTGCGTGGCGCCAGGGTGACGTTGT
CGCGGATCGACAGATGCGGGAACAGGTTGAACTGCTGGAACACCATGCCGACCTCGGTACGGACTGCCTGCAGCGACTTGCCGCTGGCGCCATTGGGCAG
CAGCGTCTTGCCGTCGACCTCGATACGACCGGACTGGAATGGCTCGAGACCATTGACGCAGCGGATCAGCGTCGATTTTCCCGAGCCACTGGCGCCGATG
ATTACCACGACCTCGCCCTGGGCCACGCTCAGGTCGATGTCCTGCATCACGTGCAGGCGGCCAAAGTGCTTGTTGACCCCTTCGAGACGCACGATCGGCG
ATGAGCCGGACGATTGGGCGGTCTCTTGCCGGTTTGCGCTATCGTTCATCCCACTAACCCCTTGCGTTCGAGCAGACTCAGAGCCAACGACAGACTCAGG

>Salinicola acroporae LMG 28587 [NZ_PZJW01000003.1:116896..117061], dnaA> ig dnaN>, 166 bp

GTCGGCGTGTAACGAATTCCCTGGCCTCGGGGATTGCTAGCGACGACCGGCACGGCAAGAATGAGACGGCGCGGCCCGACGGTTTCCTCGTCCGTCCCAT
GATGCAAAGCGATCCACCGGTTACCGGATTCAACTTCAGGCTTATTGAACCAACAACGGAGCCCCA

WebSkew GCmin pos. 193930 (GCmin<>oriC1 28.0 kb; GCmin<>oriC2 1.1 kb; oriC2<>oriC1 26.9 kb)

>Salinicola corii L3 [NZ_VTPX01000004.1:c165640..166139], <rpmH oriC1 dnaA>, 610 bp

GCCCTTGGCCGGATCGGCGCCAGCAGGCATGTCGTCCTGAATCGGCTTGGCATTGCCGTTGCTGTCCGTCGGCGCGGACAGGGCATTGCCCTCGCCGTCA
TCGCCGTTACCCGATGAAGCCGCCATCTCCGGCGCCGGGTTGTTCACGACGTGCTGACCGTAGTCCTGATTCCACTGCATGATCAGCAGGTAGGCCAGCA
CGGCCAACGGAATCACGAAAAGGAGTCGTTTCAAATCCATATAGGGGTCTGCCCGGTAAGGGAATGGGTCAGTGGGCGCCGGGCATCGGCGATCGACCGG
CGAGATTCGCCGGTGCTCCCCGGCCTTCGGCGCGGGGATGTCCGATCAGGAAGACGAAACGTCTTGACGCATCTGGCGCATCTGGCGAGCGGACTTCTCC
AGGCGTCGCCACATGCCATTGAGCTGGCGGGTGACGCTGGCGTTATCCAGCTCATGTACCCCTCGTCGCGCGAGTATGACGATATCCACTGCCGGTAAAC
GATGCTGTTGATGGCGAATGGACTCGCGCACGAGGCGCTTGAGGCGATTGCGGTCGACGGCACGGCGCACGTTCTTCTTGCTGAAGATCAACCCGATCCG
TGGATGGCCCAGCGCGTTGGGTCGGGCCAACGCCAGCAATCCTTTACCATGGACCTTGTAGGGAGCGTGGTCGAAGACGTGCTTGTAGTCCCCGGCATTC
AGCAACCGCAGTTGCCGGGGAAAACGCTGATCGGACACCCGGCGTCCGATCAGGCGCTCAGGCGTTTACGGCCTTTGGCACGGCGACGGGCAATGACCTG
GCGACCGTTCTTGGTGGCCATGCGGGCGCGAAAACCGTGGACGCGCTTACGCTTCAGAACGCTGGGCTGGAAGGTGCGTTTCATAGCTGCTAATTCCCAA
TTAGTTGATTTCTAAAGAAAACTGGTTGAAGCGGGATGCCCGACGCGTTTGCGGGAAGCGGGCTCACTCGACTCAAGAGCGGGAATTCTAGTGAATCCGC

GGTTCGGGTGCAATTTTTCCGTGAATTATCCACACCCCTTTGTGAGTGAATCGTCACCCGTCACGCCTTTGGCTGGAGACCGGTAACGTGGGCAGCCATC
GACGATTTTCGTGGCGATGCCGGCAACCGGATCGAGGAACGAGATCGAAGTTCCTCGAGAGTGTCAATCACACCATGTGGTTAACGATGGGGATAACCTT
**TGTGGATAA**CAAGATTATATTCTTTAATATCATTACGTTAATATATTTTTTTGGTTGTGGTAAGTTGCGCATGACCCGTGCACAGGTGAGGATCGCCT
ACCAGACGGCGAGATCGGTCGTACCACTTTGTTCACGGCAATTACCATGGGCATCCACAGCATCATCAGCCGCGCGGGGTGACTTATCCACAATTGCTCG
AAAGTCGACTTCGTGTGGATAACCTGGCAGGCCGACGCTACAATGTGCGGCCGTTTTTTCTATTCTCTATTCTCTCAACGTTCGGTGAGGTCGTGTGTCT

GTCGCTCTCTGGCAACAATGTCTGAACTCGCTGCAGGATGAGTTGAGCTCTCAGCAATTCAATACCTGGATTCGTCCTCTGCAGGCGGAGGACGGCGAAG
TGGACGAACTCTGTCTATTGGCCCCCAATCGATTCGTCCGTGATTGGGTCAGCGACAAATACCTGAAGCGGATCAACGAACTGTTGCGCGAGGTTGCGTC
CAGCAGCAAGCCTCCCAAGGTTCATCTGACGGTGGGGAGTCGACGTGCCGCGGCGGTGTCGCCGCGAGATCTGGGGGCGCCGGTTTCCGCTCCGAGCCGC
CAGCCTCAGCGATCCACAGCGCCGGGAGTACTGACGAGCCCGCGCACAAACGAGGCGACCAAAGAGAAGCAGCTCGATCGAGCGCGTGAACGCGAGGAGA
GCGGTCGGCGGGGCAGCGAGCGCCAGGTTCAGGTCGAGGGCAGCCTCAAGCATGGCAGCGGACTCAATCCCAATTTCACCTTCGAAACCTTTGTCGAGGG
TAAGTCGAACCAACTGGCACGCGCCGCCTCGCGTCAGGTATCCGAGAACCCTGGCGGGGCCTACAATCCCCTGTTTCTCTACGGTGGCGTGGGTCTGGGC
AAGACTCACCTGATGCACGCCGTGGGCAACCATCTCGCGGGTCTCAGAGACAATGCCAAGGTGGTCTATCTCCACTCGGAGCGCTTCGTCGCCGACATGG
TCAAGGCGCTGCAGCTCAACGCGATCAATGATTTCAAGCGTTTCTATCGTAGTGTCGATGCGCTGCTGATCGACGATATCCAGTTTTTTGCCGGCAAGGA
GCGCTCCCAGGAAGAGTTTTTCCACACGTTCAACGCTCTGCTCGAGGGGGGGCAGCAGATGATTCTGACCTCGGACCGCTATCCCAAGGAGATCAGCGGC
GTCGAGGAACGCCTCAAATCGCGCTTCGGCTGGGGGCTGACGGTAGCGATCGAACCGCCGGAACTGGAAACCCGCGTGGCGATTCTGATGAAGAAGGCGG

>Salinicola corii L3 [NZ_VTPX01000004.1:c192579..193078], <mnmG oriC2 <COG1126 (glnQ), 761 bp

ACGCACGACCTGCAACTGGACATCGAACGGCAGTGACGTAGAAATCCCGTTGGGATAGAGCTCGTGGGTGTCCAGCCCTTCGGGCTCGACGAAGACTTGG
TGACTGGTCTTGTCGGCAAAGCGATGGACCTTGTCTTCGATCGACGGACAGTAGCGCGGGCCAACGCCCTCGATCACGCCGGAATACATCGGCGAGCGAT
CGAGATTGGCAAAGATGATCTCGTGAGTGCGCTCGTTGGTATGGGCGATATGACAATCGACCTGACGCGGATGTTGTTCGCGGCTACCCAGATAGGACAT
CACCGGTAACGGCGTATCGCCAGGCTGGCGCTCGAGCCGGGAAAAATCGACGCTCTTGGCATCGATCCGCGGCGGCGTTCCAGTTTTCAGCCGATCAACC
CGGAACGGTAGCGCCCGCAGCCGCTGAGCCAGTGCATTGGCCGGCGGGTCGCCGGCGCGCCCTCCGGCATGATGGTCCAGACCGATATGGATCACGCCAC
CCAGGAACGTCCCGGTACACAGCACCACGGTATCCGCCATGAAGCGAATGCCGGTCTGGGTCGTGACGCCGACGACGGTATCGTTTCCACAACCAGATC
GTCAACGCCCTGCTGGAACAGCGTGAGATTGGGCTGATTTTCCAGCATCCCGCGAATGGCGGCCTTGTAGCGTACGCGATCCGCCTGGGCTCGGGTGGCA
CGAACCGCCGGTCCCTTGCGCGAATTGAGCACGCGGAACTGGATGCCGGCCAGATCGGTGGCCTGGGCCATGGCCCCGCCGAGGGCATCGATCTCCTTGA
CCAGATGGCTCTTGCCGATCCCGCCGATGGCGGATTGCAGGACATCTGGCCCAGCGTCTCGATGTTGTGCGTCAGTAGCAGGGTGCGGGCACCGCCACG
TGCCGCTGCTAGCGCGGCTTCGGTACCGGCATGGCCACCGCCGATCACGATGACCTGAAAACGGTCGGGATACTCCACGGTTTACCTCACGAGGTGCTGG

AAACGAGTAGCTCGAACAGGCACCCGACAGTGGTGAATGAATCTCGAAAAGCCGCCCAGTATAAACCTCCTGTGGGTAACCGTCAGGGGTTTTCCACACC
CGATTGGCTTCAGCCCATCAGCCAACAGAGTCATTACTAAATAAAGAAAAATATATAAAGAAGTTCTGTTGTGGTAGTAACTAATGGTGTGGACATTTTC
**AGTGGATAA**GTCGAATTACCCTATATCCTACAGCTACTTGCATTGTTATCCGATTGTTTGAAAAGCTTTGGGGAGGCTGCTGACAACGGCGTTGGTTCCT
GTGGACAGAAGATCACTTTGTGCACATCGCTGTTCGAGCACCGGTTGTCCACCGTCGTATACGCCAACTGTTACCGCGGTGGTCAACAAACCGCGAATTC
TCCACCAGTGCCGGACGAATCAGGGTTACAGCCGGATAGGCAAAAAAACTATCCACAGGCGAAAAAAGTGTCTGGGGTAATCGTGAATGACGAAATCGAC

CCTGTCCGACCTGGTTTTCCGACCTCGAGGCGAGCCGATGGCTCGGCTGCTTTTCGCTCGAGCGAAGACAGGAAGCACCAAGGTTCGATGAAGCAGACAG
GCAATGACGGGAATGGAGAAGCGTGGTTCAGGGGAGAGCCGAGAGTACTGTCAGGGAACGAAGTGTCCGTCGAGGCAGGCGGACACCGGGAATGGCACCG
GTGCCCGCGTTTTTTGGAACGCTTTAACCCACGTTCGAATTAGTGCTTGAGCACCCGGGAGAGGAACGCCTGGGTACGCTCGTGGCGGGGGTGATCGAAT
AGCTGGTCGGGTGGGCCCTGTTCGACGATCTGGCCGCCGTGGACGAAGACCACGCGATCGGCCACCTCGCGGGCAAAGCCCATCTCGTGAGTGACGATCA
TCATGGTCATGCCGTCGTTGACCAGTTCCCGCATGGCATCCAGGACTTCGCCGATCATCTCCGGATCCAGCGCGGAGGTAGGCTCGTCGAACAGCATCAG
GCGCGGTTCCATGGCCAGCGCCCGGGCCAGCGCCACGCGCTGCTGCTGCCCACCGGAGAGCTGGGTCGGATGCTTGTCGGCCTGATCCGAGATGCCGACC
CGCTCCAGCAGCCTGTCGGCGGTGGCATCGGCCTGCCGCCGGGTCATGCCGCGGACCTTGCGGGGCGCCAGGGTGACGTTGTCGCGGATCGACAGATGCG
GGAACAGATTGAACTGCTGGAACACCATGCCGACCTCGGTCCGCACCGCCTGCAGCGATTTGCCGCTGGCCCCATTGGGCAGCAGGGTCTTGCCGTCGAC
CTCGATGCGACCGGACTGGAATGGCTCGAGGCCATTGACGCAGCGGATCAGTGTCGACTTTCCTGAGCCACTGGCGCCGATAATCACCACGACCTCGCCC
TGGGCCACGCTGAGATCGATGTCCTGCATGACGTGCAGGCGACCGAAGTGCTTGTTGACCCCGGCGAGTCGCACGATCGGCGAGGTCCCGGACGCTTGGG

>Salinicola corii L3 [NZ_VTPX01000004.1:c164000..164166], dnaA> ig dnaN>, 167 bp

GTCGATGTGTGACGAATTCCCTGGCCGCGGGGATTGCTAGCGACGTCCGGCACGGCAAGAATGGGATAGCGCGGCCCGACGGCTTGTTTCGTCCGTCCCA
TGTCCCAAAGCGATCCACCGGTTACCGGATTCAACTTCAGGCTTATTGAACCAACAACGGAGCCCCA

WebSkew GCmin pos. nd (GCmin<>oriC1 nd; GCmin<>oriC2 nd; oriC2<>oriC1 nd)

>Salinicola peritrichatus JCM 18795 [NZ_PZJU01000083.1:5670..6169], <rpmH oriC1 dnaA>, 618 bp

GGCCATGGCCGGGCTGTCGCCGTCGGCCGATGACTTGGCCTGATTCGCGGGGGCCGCCAGGCCCTGATCGTCGGCGCTTGGGTCACTCGCATCGTTACCG
GCGCTGTCGCTGAAGGAGGGCGCCTGGGTCTCGACGGCGGTCTGGCCGTAGTCCTGATTCCACTGCATGATCAGCAGGTAGGCGAGCACGGCCAGCGGAA
TCACGAAAAGGAGTCGTTTCAAGTCCATAAAAGGTCTGCCCGGTAAGGGAATGGTTCGGTGGGCGCCGGGCATCGGCGGTCATCCGGCGACGAACGCCGG
GACTCCCCCGGCCTGCAGCGCAGGGATGTCCGATCAGGTCGACGACGATGAGTCGTGCCGCGTTTGGCGGGCGGACTTTTCCAGACGTCGCCACATGCCG
TTGAGCTGGCGGAAGACGCTGGCGTTGTCCAACTCATGTACGCCCCGCCGAGCGAGCAAGACGATATCCACTGCCGGTAAACGGTGCTGTTGCTGACGAA
TGGATTCGCGCACGAGGCGTTTGAGGCGATTGCGATCGACGGCACGACGCACGTTCTTCTTGCTGAATATCAACCCGATCCGTGGATGGCCCAGTGTGTT
GGGGCGTGCCAACGCCAGCATTCCTTTACCGTGGACCTTGTAGGGAGCGTGGTCGAAAACGTGCTTGTAATCCCCGGCATTCAGTAACCGCAGTTGCCGG
GGGAAACCCTGATCGGACACCCGATGTCCGATCAGGCGCTGAGACGCTTGCGGCCCTTGGCGCGACGACGGGAGAGGACCTGACGGCCGTTCTTGGTCGC
CATGCGGGCGCGGAAACCGTGGACGCGCTTGCGCTTCAGAACGCTCGGCTGGAAAGTACGTTTCATGGCTGCTGATTCCCAATTGGTTGAGTGGTTAAGC
TATTGAAGTAGCAAGGGAAATTGGTTCGAAACGGGATGCCCGACACGCTCAACGAAACCGGGCTCACTCGGCTCAAGAGCGGGAATTCTAGTGAATCCGC

GGGTCAGGTGCAATTTTTCTCTGAATTATCCACACCCCATTGTGGGCGAAAAATACCGAGGCCGGCGGATGTATCGTGAATTTCGCTTCGCGCCGGCTGC
CGCTCTCGCCTCGATAGGTTTCCCGCAGCGGTCGCCCTGGTGGCCCCCGGCTGCCACGGGTTGTCAATAACACCATGTGGTTAAAGGTGGGGATAAGGAT
**TGTGAATAA**GCCAATTTATCCATTTTAAATCAGAGAGATAAAATTTTTTCAGGGCTGTGGGAAAGATGCTCAAGAGTGGTGCAAAAGTCAGTGGCAGGAA
AAATTTTTGGCCTGCGCTGGTGCAGCCAGCGGTAGATCTCCTCAGAGCTTTTCCCAGGATTGCCAACGCCTTGTCATAAGCTTATCCACAACGTGTCGAT
CGACCGACCTCGTGTGGATAACCTATCCACTCGACGCTACAATGTGGCCGTTTCCTTTTCATCTCTCGAGATTCGTGAGGTCGTGTGTCGGTCGCACTCT

GGCAACAATGTCTCAATTCGCTGCAGGACGAGTTGAGCTCGCAGCAGTTCAACACTTGGATCCGTCCGCTGCAGGCGGAAGACGGCGAGACGGATGAGCT
TTGTCTGCTGGCACCCAATCGCTTCGTGCGTGACTGGGTCAGCGACAAGTACCTCAAGCGCATCAATGAGCTATTGCGTGAAGTGGGCTCCAGCGGCAAG
CCGCCCAAGGTGAGGCTGACCGTGGGCAGCCGGCGCGCTGCCGCCGTTTCGCCACGCGATCTGGGTGCGCCGGTGTCGGCGCCCGGCCGGTCGCCGCAAC
GCCCGGCGGCGCCCAGCGTACAGACGGCCTCGCGTTCGGGCGAGGCGGTCGAGGAACGCCAGCAGGATCAGGCGCGTGAACGTGAGGAGAGTGGTCGACG
CTCCAACGAACGGGGAGGCGAGCGCGGCGGCGAACGTCAGGTTCAGGTCGAAGGCAGTCTCAAACACGGCAGTGGCCTCAATCCCAACTTTACCTTCGAA
ACCTTCGTCGAGGGTAAGTCGAACCAGCTGGCCCGCGCCGCCTCGCGCCAGGTCTCGGAAAACCCCGGCGGGGCCTACAATCCGCTGTTTCTCTACGGTG
GCGTCGGTCTGGGCAAGACCCACCTGATGCACGCCGTGGGCAACCACCTGGCCGGTCTGCGCGACAACGCCAAGGTGGTCTATCTGCACTCCGAGCGTTT
CGTCGCCGACATGGTCAAGGCGCTGCAGCTCAATGCGATCAATGATTTCAAGCGCTTCTATCGCAGCGTCGATGCGCTGCTGATCGACGACATCCAGTTC
TTCGCCGGCAAGGAGCGCTCCCAGGAGGAGTTCTTCCATACCTTCAATGCCTTGCTGGAAGGCGGCCAGCAGATGATCCTCACCTCCGACCGCTATCCGA
AGGAGATCAGCGGCGTCGAGGAGCGTCTCAAGTCACGTTTCGGCTGGGGACTGACGGTGGCGATCGAGCCGCCGGAGCTGGAAACCCGAGTGGCGATCCT

>Salinicola peritrichatus JCM 18795 [NZ_PZJU01000033.1:12099..12598], <mnmG oriC2 <COG1126 (glnQ), 588 bp

CAGCCCGCTGATCGAGCGCACCACCTGCAGCTGAACGTCGAACGGCAGCGAGGTCGAGATACCGTTGGGGTAGAGCTCGTGGGTATCGAGGCCTTCGGGC
TCGATGAATACCTGGTGGCTCGACTTGTCGGCGAAGCGGTGAACCTTGTCCTCGATCGACGGGCAATAGCGCGGGCCGACGCCCTCGATCACCCCGGAGT
ACATCGGAGAACGATCGAGATTGGCGAAGATGATCTCGTGGGTGCGCTCGTTGGTATGCGCGATGTGGCACTCGACCTGGCGCGGGTGCTGCTCGCGACT
GCCCAGATAGGACATCACCGGCAATGGCGTGTCGCCCGGCTGCGGCTCGAGCTTGGCGAAATCGACGCTCTTGGCGTCCAGCCGCGGTGGCGTGCCGGTC
TTCAACCGATCGACCCGGAATGGCAGGGCGCGCAGGCGCTGCGCCAGGGCGTTGGCCGGCGGATCGCCGGCGCGGCCACCGGCATGGTTGTCCAGGCCGA
TGTGGATAACGCCACCCAGGAAGGTACCGGTGCACAGCACCACGGTATCGGCCATGAAACGGATACCGGTCTGGGTCTTGACGCCGACGACGGTGTCGTT
CTCCACGATCAGATCGTCGACCGCCTGCTGGAACAGCGTGAGATTGGGCTGGTTCTCCAGCATTCCGCGAATTGCTGCCTTGTAGCGCACGCGGTCCGCC
TGGGCACGGGTGGCCCGCACCGCTGGCCCCTTGCGCGAATTGAGCACACGAAACTGGATGCCGGCCAGATCGGTGGCGTGGGCCATGGCGCCGCCAAGCG
CATCGATCTCCTTGACCAGATGGCTCTTGCCGATGCCACCGATGGCAGGGTTGCAGGACATTTGACCCAGGGTCTCGATGTTGTGCGTCAGCAGTAGCGT
GCACGCACCGCTACGGGCCGCGGCCAGCGCGGCTTCGGTACCGGCATGGCCACCGCCGATGACGATGACCTGAAAACGCTCGGGGTACTCCACGTTTCAC

CTCGTTGGGTGCTCTGCATGGCACCCGACAGTGAATGAATTCTGGAAAAAGCCGCCCAGTATAAACCTCCTGTGGGTAACCGTCAGGGGTTTTCCACACC
CGTTTTCCAGCCGCCTGTCAGTCATCAGTTTCATTATTAAAAAAGAAAAATATATAGAAGAAGTTCTGTTGTGATAGTGACTACTGGTGTGGACACTTTC
**TGTGGATAA**ACAACTTTTCGGTTTATTTTCCAATGACTTATATTGTTATTGAATGGTTTGAAAAACCCTGAGGAGGCTGCTGACAGCCAGCTTCGTTCCT
GTGGACGAAATCCCTTTTTATACACACGCCACGACGCGCACCGGTTTTCCACCGCTGCATACGCTAGCTGTCACCGCAGCGCTCAACAACCCGTTGATCG
TGCGGCAACGCAGTGTTCACGGGGGGTATGGCCGAATTGGCGGAAAAACTATCCACAGCCGAAAAAAACGGCCTTGGATAACCGGAGGGAAGCGGAATCA

CGCTCGATGGCCACGGCGACAGCGCCATGGTTGTCGCGATGGCAAGGCGGAACGCCGAGGCTGGCCGGCGTTCCCGGTGGGATCAATGTTTGAGAACGCG
GGAGAGAAAGGATTGGGTGCGTTCGTGGCGCGGATGGTCGAACAGTTGGTTCGGCGGGCCCTGTTCGATGATCTGGCCACCATGCACGAACACCACGCGG
TCGGCGACCTCGCGGGCAAAGCCCATCTCGTGGGTGACGATCATCATCGTCATGCCGTCTTTGGCCAGCTCGCGCATGGCATCGAGCACTTCGCCGATCA
TCTCCGGATCGAGCGCCGACGTCGGCTCGTCGAACAGCATCAGCCGCGGTTCCATGGCCAGTGCGCGGGCCAGTGCCACGCGTTGCTGCTGACCGCCGGA
GAGCTGGCTGGGATACTTGTCGGCCTGGTCGGCGATACCCACTCGTTGAAGCAGTCGCTCGGCGGTGGCGTTGGCCTGCTGTCGCGACCAGCCGCGCACC
TTGCGTGGTGCCAGGGTGACGTTGTCGCGAATCGACAGGTGCGGAAACAGGTTGAACTGCTGGAACACCATGCCGACTTCGGTACGTACGGCCCGCAGCG
ACTGGCCGCTGGCACCATTGGGCAACAGTGTCTTGCCATCCACCTCGATTCGGCCGGACTGGAACGGTTCGAGACCGTTGACGCAGCGGATCAGGGTCGA
CTTGCCCGAACCGCTGGCGCCGATGATCACGACGACCTCACCGGGGACGACCGAGAGATCGATGTCCTGCATCACGTGCAGATCGCCGAAGTGCTTATTG
ACCCTGTCGAGACGCACGATCGGCTGGCGATCGTCGTCGGGTGACAGCGTGGTTGCGGTGTTGTCGTTCATCCCACCAGCCCCTTGCGTTCGAGCAGACT
CAGAGCCATCGACAGGCTCAACGTGATCACCAGATAGAGCAGGGCGATCATGAAATAGACCTCCAGCGCGGTGAAGGTATTGGCGATGTAGACCTGGCCC

>Salinicola peritrichatus JCM 18795 [NZ_PZJU01000083.1:7657..7838], dnaA> ig dnaN>, 182 bp

TTCGCTGAACCTCAATTACCGCAAGGCAGGGATTGCGACTGGCGTCCGGCGAGGCAAGAATGGGACGGCGCGCAGTTAGACCGCCTTGAACGATCGATCG
TTTGGCAGGATTGATCCACCGGTTTCTTGATTTTCCGATTTCTTGATTCAATCCAGGTTACTGATTCGACTACGGAGCCCCA

WebSkew GCmin pos. 379916 (GCmin<>oriC1 34.3 kb; GCmin<>oriC2 7.1 kb; oriC2<>oriC1 27.1 kb)

>Salinicola rhizosphaerae KCTC 32998 [NZ_BMZI01000002.1:413962..414461], <rpmH oriC1 dnaA>, 621 bp

GCCGGCTCGGAGCCCGCGGGCATGTCGTCTTCGATCGGTTTGGCATTGCCGTTGCCATCGGTCGGTGCCGAGAGCGCGCCGTTACCGCTGCTGTCACCGC
TACCCGAAGAGGCGGCGACTTCCGGCGCCGGATTGTCCACCACATTCTGATGGTAGTCCTGATTCCACTGCATGATCAGCAGGTAGGCCAGCACGGCCAG
CGGAATCACGAAAAGGAGTCGTTTCAAGTCCATACAGAGGTCTGCCCGTCAAGGGAGTGGGTCGGTGGGCGCCTTGGTATCGGCGATCGTCCGGCGACGA
TCAGCAAGACTCCCCGGCCGTAGGCGCGGTCGTGTCCGGTCAGGTCGATGTGTCGGGCCGAGCCTGTCGAGCGGACTTCTCGAGACGTCGCCACATGCCG
TGGAGCTGACGGCTGACGCTGTCGTTGTCCTGCTCGTGAAGGCCTCGTCGCGCAAGCAGTACGATATCCACAGCCGGTAGACGGTGCTGCTGATGACGGA
TGGATTCGCGGACGAGGCGTTTGAGACGGTTGCGGTCGACGGCACGACGCACGTTCTTCTTGCTGAAGATCAACCCGATCCGCGGATGGCCCAGTGCGTT
GGGACGTGCCAACGCCAGCATTCCCTTACCGTGGACCTTGAGAGGAGCGTGGTCGAATACGTGCTTGTAGTCCCCGGCGGTCAGCAAACGGAGTTGCCGG
GGGAAGCCATGATCGGACACCCGTTGTCCGATCAGGCGCTGAGACGCTTGCGGCCCTTGGCACGACGACGGTTGATGACCTGACGGCCGTTCTTGGTGGC
CATGCGAGCGCGGAAACCGTGGACGCGCTTGCGCTTCAGAACGCTGGGCTGAAAGTACGTTTCATGACTGCTGCTTCCCGAGGTAGTTGAGTGGCTAAG
CTGATGATTCAGCTTATAAAAAAGGTTCGATACGGGACGTCGGACACGCGCGACGGGTCCGGGTCACTCGGCTCAAGAGCGGGAATTCTAGTGAATACGG

CGGTCAGGTGCAATTTTTCCGTGAATTATCCACACCCGGTTGTGAGCGAAACGACGCACTGCCCAATGACAGAATTCGCGCTGCTAGGTTACTCGGAAAG
GTTTTCAGGATATCGCGGCCGGCCGTGCCCGGTTGAAGTCTTGAGCGCGAGCACGGGAGCTTGTCAATCGCACCATGTGGTTAAGGATGGGGATAGTTAT
**TGTGAATAA**CTCTTATTATATAAATGATATCAACGGCCTACGATCTATCTTTGCTTGTGAGGAAAATGCGTATCCCTGGTGCAAAAGCTGCGGACTGTGC
ACCGATGAAGCGGCACGATGGTGCACACGACCGCCGGTTTTTCCGTCTTCTTCCGCAACTTGCCAACCGCTTGTCGGCTGTTTATCCACAGCTTTTAGCG
TGCGAGACTCGGTGTGGATAACCCGGGAGTCGGACGCTACAATGTGCGGCCGTTTCTCATTCATCTCTTGTTATCCGGTGAGGTCGTGTGTCCGTCGCGC

TCTGGCAACAATGCCTCAATTCGCTGCAGGATGAGTTGAGTTCGCAGCAGTTCAATACCTGGATCCGCCCGCTGCAGGCGGAGGACAGTGACTCGGACGA
ACTCTGCCTGCTCGCCCCCAACCGCTTCGTGCGTGATTGGGTGAGCGACAAGTATCTCAAGCGCATCAATGAACTGTTGCGCGAAGTCGGTACCAGCAAC
AAGCCGCCCAAGGTTCGTCTGAGCGTCGGCAGCCGTCGCGCTGCGGCGGTCTCGCCGCGCGATCTGGGGGCTCCGGTATCGGCTCCTGGACGATCGGAAC
AGCGGCCCGCGGCGCCGAGCGTGCAGACGCCGCCGCGCAGCGAGGAAGCGGCAGACGCGCGCGAGGAGGATCGGAGCCGGGAAGGGGGGCGGGAAGAGAG
TGGCCGGCGCGCCAGCGACCGCGGCGATCGCCAGGTACAGGTCGAGGGCAGTCTCAAGCACCAGAGTGGCCTCAATCCCAACTTCACCTTCGAGACGTTC
GTCGAAGGTAAGTCGAACCAGTTGGCGCGTGCCGCATCGCGGCAGGTCTCGGAAAACCCCGGCGGTGCCTACAACCCGCTCTTTCTCTACGGCGGCGTCG
GCCTGGGCAAGACCCACCTCATGCACGCGGTCGGCAATCATCTGGCCGGGCTACGCGACAATGCCAAGGTCGTCTACCTTCACTCCGAGCGTTTCGTGGC
CGACATGGTCAAGGCGCTTCAGCTTAACGCAATTAACGATTTCAAGCGCTTCTATCGCAGTGTCGATGCGCTGTTGATCGACGACATCCAGTTCTTTGCC
GGCAAGGAGCGGTCGCAGGAGGAGTTCTTCCATACTTTCAACGCGCTGCTCGAGGGCGGTCAGCAGATGATTCTGACTTCCGATCGCTATCCGAAGGAGA
TCAGCGGCGTCGAGGAGCGTCTCAAGTCCCGCTTCGGCTGGGGGCTGACGGTGGCGATCGAGCCGCCGGAACTCGAGACACGCGTGGCGATCCTGATGAA

>Salinicola rhizosphaerae KCTC 32998 [NZ_BMZI01000002.1:386819..387318], <mnmG oriC2 <COG1126 (glnQ), 734 bp

GGGATAGAGCTCGTGAGTCTCCAGGCCCTCCGGCTCGATGAAGACCTGATGACTGGTCTTGTCAGCGAAGCGGTGGACCTTGTCCTCGATCGACGGGCAG
TAACGCGGCCCGACGCCCTCGATGACGCCGGAGTACATCGGTGAACGATCGAGATTGGCGAAGATGATCTCGTGAGTTCGCTCGTTGGTGTGCGCGATGT
GGCACTCGACCTGGCGCGGATGCTGATCGCGAGACCCCAGATAGGACATTACCGGTAGCGGCGTGTCACCCGGCTGCGGTTCGAGACGCGAGAAGTCGAC
GCTCTTGGCATCGATTCGTGGCGGTGTACCGGTCTTCAGCCGATCGACGCGAAACGGCAGCGCGCGTAGCCGCTGCGCCAGCGCATTCGCAGGCGGGTCG
CCGGCGCGGCCACCGGCGTGATGGTCGAGCCCAACGTGAATGATCCCGCCGAGAAACGTACCGGTACACAGCACCACGGTATCGGCCATGAACCGGATAC
CCGTCTGCGTGACCACGCCGGTCACGGTGTCGTTATCCACAATCAGATCATCGACGCCCTGCTGGAACAGCGTGAGATTGGCCTGATTCTCCAGCATGCC
GCGAATGGCGGCCTTGTAACGGACCCGGTCCGCCTGGGCACGCGTGGCCCTCACTGCCGGCCCCTTGCGCGAATTGAGCACGCGGAACTGGATACCGGCG
AGATCGGTTGCCTCGGCCATGGCACCGCCCAGCGCATCGATCTCCTTGACCAGATGGCTCTTGCCGATACCGCCGATCGCCGGGTTGCAGGACATCTGGC
CCAGCGTCTCGATGTTGTGCGTCAAAAGCAGCGTGCGGGCACCGCTGCGGGCAGCGGCGAGCGCCGCTTCGGTACCGGCATGGCCACCGCCGATCACGAT
GACCTGAAAACGGTCGGGGTATTCCACGTCTCACCTCGTTGGGTACCCGCCGGCACCCGACAGTGAATGAATCTGGGCAAAGGGCGTCCCGAAAGATTAT

TGACGCAGCGTCAATGAACCCGGAACGGCAGCCGCCGATGGCGACGAAAGCCGCACAGTATAAACCTCCTGTGGGTAACCGTCAGGGGGTTTTCCACACC
GACTTTTCGTCCGGGTCCGCTACCGATACTTCATCAATATAAAAAAAGAAAAATATAAAGAAGTTCTGTTGTGATAGTGACTATTGGTGTGGACATTTTC
**TGTGGATAT**CTGTTTTTTCTTCAACGAAAACGGTAGCTTAAATTGTTATCGAATGGTTTGAAAAACGCTGGGAAGGCTGCTGACGAGCGCTGTTCAAGCT
GTGGATGACTCCGGGGTTTATGCACAGGCCAGCGAGCGCACGACTTTTCCACCGCTGCATACGCTAACTGTCACCGCGACTCTCAACAACCGCCGATATT
GGCTCGGAGGCAGAGGCGGTGCGGGCTATCGTCAGGTCGTGAAAAAAATTGTCCACAGGGGAAAGTCAGCCCTGGGGAGGATGTCGAATGACCGATTCCA

GTCGTGTTGACAGGTTGCGCGGGAAAAGTGTAGCTACCGGGCCGGTGAAGGAGCGTCGCTGGAGAGGAAATGACGCAAGGCGTGGCCAGGTTGGCGACAC
CGTCCCTGCCATGGGCGAGAGTGTCCCGGCAGCCGAAACGGCGTCGCGTTCGAGGGTATGTCTAGTGCTTGAGCACGCGCGAGAGGAATGCCTGAGTGCG
ATCGTGCTGCGGTGTGTCGAAAAGCGCAGCCGGCGGGCCCTGTTCGACGATCTGGCCGTTGTGGACGAACACAACGCGATCGGCGACTTCGCGGGCGAAG
CCCATCTCGTGTGTCACGATCATCATCGTCATGCCGTCATTGGCCAGTTCGCGCATGGCATCCAGCACTTCACCGATCATTTCGGGATCGAGCGCCGAGG
TGGGCTCGTCGAACAGCATCAGGCGAGGTTCCATCGCCAGGGCACGCGCCAGCGCCACGCGCTGTTGCTGACCACCCGAGAGCTGGCCGGGAAACTTGTC
GGCCTGATCGATGATGCCGACCCGTTCGAGCAGGCGTCTGGCGGTATCGTCGGCCTGCTGCCGCGTCCAGCCACGGACCTTGCGCGGGGCCAGCGTCACG
TTGTCGCGAATCGACAGGTGGGGAAACAGATTGAACTGCTGAAACACCATGCCGACTTCGGTACGCACTGCCTGCAGCGACTTGCCGCTGGCACCGTTGG
GCGAGAGCGTGTTGCCGTCGACCTCGATCCGTCCTGACTGGAACGGCTCCAGGCCGTTGACGCAGCGAATCAGCGTCGATTTGCCCGAGCCGCTGGCACC
GATGATCACCACCACTTCGCCAGGGGCGACCGAGAGGTCGATATCCCGCATCACGTGCAGTTCGCCGAAATGCTTGTTGACTCGCTCCATTCGCACGATG
GGCGTCGTGTCAGCCGTCTCGTCTGTCGGCCGGTTGCGTTTCACTTTGTCATTCATCCCACCAGCCCCTTGCGTTCGAGCAGACTCAGAGCCATCGACAG

>Salinicola rhizosphaerae KCTC 32998 [NZ_BMZI01000002.1:415943..416070], dnaA> ig dnaN>, 128 bp

GGCGTCGCCGTCTCGTCTCGCCGCCGACAGGGATTGCGAGCGGCGCCTGGCGAGGCAAGAATGGGACGGCGCTCAGGTTCTGCCGCAGACAGACTCGATC
CAGCCATTGATTCGAATACGGAGCTCCA

WebSkew GCmin pos. 1513704 (GCmin<>oriC1 31.1 kb; GCmin<>oriC2 1.9 kb; oriC2<>oriC1 29.2 kb)

>Salinicola tamaricis F01 [NZ_CP023559.1:c1482361..1482860], <rpmH oriC1 dnaA>, 622 bp

GGTTCACTGCCCGCGGGCATGTCGTCCTCGATCGGCTGGGTCTTGCCTTTACCACCGCTATCGGGGGCGGAGAGGGCGTTACCTTCTTTGCCACCGTCAC
TGGTGGCGGAGGCGACTTCCGGTGCCGGGTTGTTCACGACGTTCTGACCGTAGTCCTGATTCCACTGCATGATCAGCAGATAGGCGAGCACGGCCAGCGG
AATCACGAAAAGGAGTCGTTTGAAATCCATAGAGGGGTCTGCCCGGTAAGGGAGTGGTTCGGTAGGCGCCGGGCAACGGCGATGATTCGGCGATGGACGC
CGAAGCTCCCCGGCCAGCAGCGCGGGGATGTCCGATCAGGTCGACGAGGTGTCGTGGCGCGTCTGACGCGCGGACTTTTCCAGGCGTCGCCACATGCCGT
TGAGCTGGCGCGCGACGCTTTCATTGTCCAGCTCGTGTACGCCTCGCCGAGCGAGAAGAACGATATCCACGGGCGGCAGGCGATGCTGCTGCAGACGGAT
GGATTCGCGCACGAGGCGTTTGAGGCGATTGCGGTCGACGGCACGACGCACGTTCTTCTTGCTGAAGATCAACCCGATCCGCGGATGGCCAAGCGCGTTG
GGGCGTGCCAACACCAGCATGCCTTTACCGTGGACCTTGAAGGGAGCGTGGTCGAAAACGTGCTTGTAGTCCCCGGCGTTGAGCAACCGAAGCTGCCGGG
GGAAACCTTGATCGGACACCCGATGTCCGATCAGGCGCTGAGACGCTTGCGACCTTTGGCACGACGACGATTGATGACCTGACGGCCGTTTTTGGTCGCC
ATGCGGGCACGGAAACCGTGGACGCGCTTGCGCTTCAGGACGCTCGGCTGAAAAGTACGTTTCATGGTGGCTTTTTCCCAATTGGTTGAGTGATTAAGCT
ATTGATTTAGCTCGAAAAACTGGTTCGAAACGGGATGCCCGACACGCGTACGGAAACCGGGCTCACTCGGCTCAAGAGCGGGAATTCTAGTGAATCCGGC

GATCGGGTGCAATTTTTCTATGAATTATCCACACCCGTTTGTGGGTGAAACCCTTGCGTCACATCGCCGCGGCGGTCACGCGCGCACGCCGTTAGGAAGC
GAAGCGATCGGCGGCCGAGGGTGGGCATGATCGTCCGCGGCTTATCCCCAACCTGTCGGCTTGTCAATCGCACCATGTGGTTAAGGATGGGGATAATGTT
**TGTGGGTAA**CTCATTATAATATTTACTTTTCAAAATCTTATGTGATTTTCTGTCCTGTGGGGAAGATGCGGAAGACAGGTGAGAATCCTGCGTGTGGCGA
GAACCCGCGGGACACGATGTGTCTTTGAACGAGAACCGCCGGGCGATCATTTTCCCAAGGCTGTCAACGGGTTCTCCGCCGCTTATCCACAGTGGGATGG
CCTGGCGACCACGTGTGGAACCTGCCTGGCCGGCGCTACACTGTGCGGCCGTTTTCTTTTCATCTGCCGAAATCCCCGTGAGGTCGTGTGTCGGTCGCTC

TCTGGCAACAATGTCTCAATTCGCTTCAGGATGATTGAGCTCGCAGCAGTTCAACACCTGGATCCGCCCTTTGCAGGCGGAGGATGGCGAGACGGACGAG
CTCTGTCTGCTGGCGCCCAACCGCTTTCGTGCGCGACTGGGTCAGCGACAAGTATCTGAAGCGTATCAACGAGCTGCTACGCGAGGTGGCGTCGAGCAGC
AAGCCGCCCAAGGTGCGTCTGGTGGTGGGCAGCCGCCGCGCGGCTGCCGTGTCGCCGCGGATCTGGGCGCGCCGGTATCGGCCCCCAGCCAGCCGCCGCA
GCGCCCAGTGCGCCCGCGTGAAGACGTCGCCGCGCTCGCCCGAGGTGGCCGAGGAGCAGGCACTCGACCAGAACCGTGAGCGCGAGGAGGGCGGCCGGCG
TGGCGGTGAACGCCAGGTGCAGGTCGAGGGTAGCCTCAAGCACCAGAGCGGGCTTAACCCCAACTTCACCTTCGAAACCTTCGTCGAGGGCAAGTCGAAC
CAACTGGCGCGGGCCGCCTCGCGTCAGGTCTCGGAGAATCCCGGTGGCGCCTACAACCCGCTGTTCCTCTACGGCGGCGTGGGGCTGGGCAAGACCCACC
TGATGCATGCGGTGGGCAATCATCTGGCCGGCCTGCGCGATAACGCCAAGGTGGTCTATCTGCACTCCGAGCGCTTCGTCGCCGACATGGTCAAGGCCTT
GCAGCTCAATGCGATCAACGACTTCAAGCGCTTCTACCGTAGCGTCGATGCACTGTTGATCGACGATATCCAGTTCTTCGGTGGCAAGGAGCGCTCCCAG
GAGGAGTTCTTCCACACCTTCAACGCCCTGCTCGAGGGCGGCCAGCAGATGATCCTCACTTCCGACCGCTATCCCAAGGAGATCAGCGGGGTGGAGGAGC
GTCTCAAGTCGCGCTTCGGCTGGGGGCTGACGGTGGCGATCGAGCCGCCGGAGCTGGAGACCCGGGTGGCGATCCTGATGAAGAAGGCCGATCAGGCCAA

>Salinicola tamaricis F01 [NZ_CP023559.1:c1511581..1512080], <mnmG oriC2 <COG1126 (glnQ), 795 bp

GTAGAGCTCGTGGGTGTCGAGTCCCTCGGGCTCAAGAAAGATCTGGTGGCTCGACTTGTCGGCGAAGCGATGGACCTTGTCCTCGATCGACGGGCAGTAG
CGCGGGCCGACGCCTTCGATCACCCCGGAGTACATCGGCGAGCGGTCGAGATTGGCGAAGATGATCTCGTGGGTCCGGGCGTTGGTGTGCGCGATGTAGC
AATCGACCTGACGCGGATGCTGCTCACGCGAGCCGAGATAGGACATCACCGGCAGCGGCGTATCGCCCGGCTGCGGCTCGAGCCGGCTGAAGTCGACGCT
CTTGGCGTCGATCCGCGGCGGCGTGCCGGTCTTCAGCCGATCGACCCGGAACGGCAGCGCGCGCAGCCGCTGGGCCAGGGCGTTGGCCGGCGGATCTCCG
GCGCGACCGCCAGCGTGGTGGTCGAGACCGATATGGATCACACCGCCGAGGAAGGTCCCGGTGCACAGCACCACGGTATCGGCCATGAAGCGGATACCTG
TCTGGGTGACCACGCCAGTGACCTGGTCGTTATCCACAACCAGATCGTCGACCCCTTGCTGGAACAGCGTCAGGTTGGGCTGGTTTTCCAGCAGCGAGCG
AATGGCGGCCTTGTAACGCACACGGTCGGCCTGGGCACGCGTCGCCCGCACCGCCGGACCCTTGCGTGAGTTGAGCACGCGGAATTGGATCCCGGCGAGA
TCGGTCGCGGTCGCCATGGCGCCGCCGAGCGCATCGATCTCCTTGACCAGATGACTCTTGCCGATGCCACCGATGGCCGGGTTGCACGACATCTGCCCCA
GCGTCTCGATGTTGTGTGTCAGCAGCAGCGTGCGTGCGCCGCTGCGGGCCGCTGCCAGCGCGGCTTCGGTGCCGGCATGGCCGCCACCGATGACGATGAC
CTGAAAGCGGTCGGGGTACTCCACAGTCCACCTCGTTGAGTGCTCGAACCGGCACCCGACAGTGAATGAATCTCGAAGGATGGCTGCCCGGTGGGATTGT

TGACTTGGAGTCAAGGATCCCATCGGCAGCGGCGACCGGAGTCGCGAAAGCCGCCCAGTATAAACGTCCTGTGGGTAACCGTCAGGGGTTTTCCACACCT
CTCGAGCCGCGTCAGCCAGGTCACAAGGTCATTACTAAAAAATAAAGAAATATAGAAAAGAAGTTCTGTAGTGATGGTTACTACTGGTGTGGACATATTT
**TGTGGAAAA**GCGAATTTTCCCTAAAGAAAACATCCACTTGGATTGTTATTGGATGTATTGAAAAGGGCTTAGGAGGCTGCTGACAGACAGCGCTCAAGCT
GTGGATGAAAAGCGTCGTTATACACAGCGTCAGCTACCCAGCGCTTGTCCACCGCTGCATACGCCTGCTGTCACCGTGACGGTCCCCAACGCGGGCATCG
CGCTGGAGAACACGGCTGAGGCGGGCTGCGTCGAGATAGGCTGAAAAACTATCCACAGGCGAAAAAAACCGCTCGGGATAACCTTTCTGGCGCTATCGCG

GGCAGATCTCAAGACGATTCGAAGGTGATTCGGGGGTGATACCGAGAGGAATCCGGCGCCCATCGTCAGAGGGTTTGGAAATGGGCTTCAAGGGGGCATG
ACGCGAAGACAGAGCGGCGACAGGAACCGCAGCCAGCCAGGGGCTCGTGAGAGACGGTCGAAGCTAGCGAGAACGGGCCACCGTCTAGGCGGCCGCGTTC
GCGCCGAGTGGGGGCGGGCTCAGTGCTTGAGCACGCGAGAAAGGAAGGAGCGGGTACGCTCGTTCTGGGGATCATCGAAGACGGCCTCCGGCGTTCCCTG
CTCGACGATCTCGCCGCATGGACGAATACCACACGGTCGGCGACCTCGCGCGCGAAGCCCATTTCGTGGGTCACGATCATCATGGTCATGCCGTCCTTGG
CCAGCTCGCGCATGGCATCCAGCACTTCGCCGATCATCTCCGGATCCAGCGCCGAGGTTGGCTCGTCGAACAGCATCAGCCGTGGTTCCATGGCCAGTGC
CCGGGCCAGCGCCACGCGTTGTTGCTGGCCGCCGGAGAGCTGGCTGGGGTACTTGTCGGCCTGATCGGCGATCCCGACGCGTTCGAGCAGGCGCTCGGCG
GTGGTGTTGGCCTGCCGGCGTGACCAGCCACGTACCTTGCGCGGGGCCAGGGTCACGTTGTCGCGAATCGACAGGTGAGGAAACAGGTTGAACTGCTGGA
ACACCATGCCGACTTCGGTGCGTACCGCCTGCAACGACTTGCCGCTGGCGCCGTGGGGGGCCAGGGTCTTGCCGTCGACCTCGATGCGCCCGGATTGGAA
CGGCTCCAGCCCGTTGACGCAGCGTATCAGGGTCGACTTGCCCGAGCCGCTGGCGCCGATGATCACCACCACTTCGCCCGGGTCCACGCTGAGGTCGATG
TCACGCATCACGTGCAGTTCGCCGAAGTGCTTGTTGACCTTGTCGAGACGCACGATCGGCGCCGCGTCGCCCTGGATCTGCTTGTCGTTGTTTTTGTCGA

>Salinicola tamaricis F01, dnaA> ig dnaN> not resolved, annotation issues

WebSkew GCmin pos. 327522 (GCmin<>oriC1 36.2 kb; GCmin<>oriC2 7.1 kb; oriC2<>oriC1 29.2 kb)

>Salinicola endophyticus CPA92 [NZ_PZJO01000003.1:363494..363993], <rpmH oriC1 dnaA>, 624 bp

GGAGCACTACCCGCGGGCATGTCGTCCTCGATCGGCTGGGTCTTGCCACTGCCGCCGTCGGGGGCGGAGAGGGCATTGCCTTCTTTGCCACCGTCGCTGG
TGGCCGAAGCGACTTCCGGTGCCGGGTTGTTCACGACGTTCTGACCGTAGTCCTGATTCCACTGCATGATCAGCAGATAGGCGAGTACGGCCAGCGGAAT
CACGAAAAGGAGTCGTTTGAAATCCATAGAGGGGTCTGCCCGGTAAGGGAGTGGTTCGGTAGGCGCCGGGCAACGGCGATCATTCGGCGATGGACGCCGA
AGCTCCCCGGCCTGCAGCGCGGGGATGTCCGATCAGGTCGACGACGAAGTGTCGTGGCGCGTCTGGCGCGCGGACTTTTCCAGGCGTCGCCACATGCCGT
TGAGCTGGCGCGCGACGCTGGCATTGTCCAGCTCGTGTACGCCTCGCCGAGCGAGAAGAACGATATCCACGGGCGGCAGGCGATGCTGCTGCAGACGGAT
GGATTCGCGCACGAGGCGTTTGAGGCGATTGCGGTCGACGGCACGACGCACGTTCTTCTTGCTGAAGATCAACCCGATCCGCGGATGGCCAAGCGCGTTT
GGGCGTGCCAACACCAGCATGCCTTTACCGTGGACCTTGAAGGGAGCGTGGTCGAAAACGTGCTTGTAGTCCCCGGCGTTGAGCAACCGTAGCTGCCGGG
GGAAACCCTGATCGGACACCCGATGTCCGATCAGGCGCTGAGGCGCTTGCGGCCTTTGGCGCGACGACGGGCAATGACCTGACGGCCGTTCTTGGTCGCC
ATGCGGGCACGGAAACCGTGGACGCGCTTGCGCTTCAGGACGCTCGGCTGAAAAGTACGTTTCATGGTGGCTTTTTCCCAATTGGTTGAGTGATTAAGCT
ATTGATTTAGCTCGAAAAACTGGTTCGAAACGGGATGCCCGACACGCGTACGGAAACCGGGCTCACTCGGCTCAAGAGCGGGAATTCTAGTGAATCCGGC

GATCGGGTGCAATTTTTCTCTGAATTATCCACACCCGTTTGTGGGTGAAACCCTTGCGTCAGATCGCCGCGGCGGTCACGCGCGTACGCCGTTAGGAAGG
GGCGCCATCGGCGGTCGAAGGTGGAAATGGTCCTCCTCGGCTTATCCCCACCCTGGACGCTTGTCAATCACACCATGTGGTTAAGGATGGGGATAATGTT
**TGTGAATAA**CTCACAATAACCCTTATTATTCATGTATCTATGCATTTTTCTGGCCTGTGGGGAAGATGCGGAAGAGAGGTGAAGATCCTGCGTGCAGTGA
CAAGGTTCGAGGTCTGGCGCGGCGCTGAATGAGCACGGCCGGGTCATGATTTTCCCAAGGCTGTCAACGGGTTCTCCGCCGTTTATCCACAGAGGGATGA
CCTGGCGACCACGTGTGGATAACCTGTCTGGCCGGCGCTACACTGTGCGGCCGTTTTCTTTTCATCTGCCGAAATTCCCGTGAGGTCGTGTGTCGGTCGC

TCTCTGGCAACAATGTCTCAATTCGCTTCAGGATGAGTTGAGCTCGCAGCAGTTCAACACCTGGATCCGCCCCCTGCAGGCGGAGGATGGCGAGACGGAC
GAGCTCTGTCTGCTGGCGCCCAACCGCTTCGTGCGCGACTGGGTCAGCGACAAGTACCTGAAGCGCATCAACGAGCTGCTGCGCGAAGTGGCATCGAGCA
GCAAGCCGCCGAAGGTGCGTCTGGTGGTGGGCAGCCGTCGCGCGGCCGCCGTGTCGCCGCGGGATCTGGGCGCGCCGGTGTCGGCGCCCAGCCAGCCGCC
GCAGCGCCCCAGTGCGCCCAGTGTCCAGACACCGTCGCGCTCCCCCGAGGTGGCCGAGGAGCAGGCACTCGACCAGAGCCGCGAGCGTGAAGAGAGCGGC
CGGCGTGGCGGTGAACGCCAGGTGCAGGTGGAGGGCAGTCTCAAGCACCAGAGCGGCCTCAACCCCAACTTCACCTTCGAAACCTTCGTCGAGGGCAAGT
CGAACCAGCTGGCGCGGGCCGCTTCGCGCCAGGTCTCGGAGAATCCCGGTGGCGCCTACAACCCGCTGTTCCTCTACGGCGGGGTGGGCCTGGGCAAGAC
CCACCTGATGCATGCGGTGGGCAATCATCTGGCCGGCCTGCGCGATAACGCCAAGGTGGTCTATCTGCACTCCGAACGCTTCGTCGCCGACATGGTCAAG
GCGCTGCAGCTCAACGCGATCAACGACTTCAAGCGCTTCTACCGCAGCGTCGATGCGCTGCTGATCGACGATATCCAGTTCTTTGGTGGCAAGGAGCGCT
CCCAGGAGGAGTTCTTCCACACCTTCAACGCCCTGCTCGAGGGCGGCCAGCAGATGATCCTCACCTCCGACCGTTACCCCAAGGAGATCAGCGGGGTGGA
GGAGCGTCTCAAGTCGCGCTTCGGCTGGGGGCTGACGGTGGCGATCGAGCCGCCGGAGCTGGAGACCCGGGTGGCGATCCTGATGAAGAAGGCCGATCAG

>Salinicola endophyticus CPA92 [NZ_PZJO01000003.1:334339..334838], <mnmG oriC2 <COG1126 (glnQ),778 bp

GGTAGAGCTCGTGGGTGTCCAGCCCCTCGGGCTCGATGAACACCTGGTGGCTCGACTTGTCGGCGAAGCGGTGGACCTTGTCCTCGATCGACGGGCAGTA
GCGCGGCCCAACGCCGCCGATCACGCCGGAGTACATTGGCGAGCGGTCGAGATTGGCGAAGATGATCTCGTGGGTCCGGGTGTTGGTGTGCGCGATATGG
CAATCGACCTGGCGCGGATGCTGCTCGCGGCTGCCCAGGTAGGACATCACCGGCAGCGGCGTATCGCCGGGCTGGCGCTCGAGCCTGGAGAAATCGACGC
TCTTGGCATCGATTCGCGGCGGCGTGCCGGTTTTCAGCCGATCGACCCGGAACGGCAGGGCGCGAAGACGCTGGGCCAGGGCATTGGCCGGCGGATCGCC
GGCGCGGCCGCCGGCGTGATGATCGAGACCGATATGGATCACACCGCCGAGGAAGGTGCCGGTGCACAGCACCACGGCGTCCGCCATGAAGCGGATGCCG
GTCTGGGTGACCACGCCGGTGACCCGATCGTTATCCACAACCAGATCGTCGACCCCCTGCTGGAACAGCGTCAGGTTGGGCTGGTTTTCCAGCAGCGAGC
GAATGGCAGCCTTGTAGCGCACACGGTCGGCCTGGGCACGAGTTGCCCGCACCGCCGGCCCCTTGCGTGAGTTGAGCACGCGAAACTGGATGCCGGCGAG
ATCGGTCGCGGTGGCCATGGCACCGCCGAGCGCATCGATCTCCTTGACCAGATGACTCTTGCCGATGCCACCGATGGCCGGGTTGCACGACATCTGCCCC
AGCGTCTCGATGTTGTGTGTCAGCAGCAGGGTGCGTGCACCGCTGCGGGCCGCTGCCAGCGCGGCTTCGGTGCCGGCGTGGCCGCCACCGATGACGATGA
CCTGAAATCGGTCGGGGTACTCCACAGTCCACCTCGTTGAGTGCTCGAACCGGCACCCGACAGTGAATGAATCTTGAAAACGGCTGCCCGGCGAGATTGT

TGACTCAAAGTCAAAGATCCCGCCGACAGCGGCGACCGGAGTCGCGAAAGCCGCCCAGTATAAACGTCCTGTGGGTAACCGTCAGGGGTTTTCCACACCT
CTCGACCTGCGCCAGCCAGGTCACAAGGTCATTACTAAAAAATAAAGAAATATAGAAAAGAAGTTCTGTAGTGATGGTTACTACTGGTGTGTACATATTT
**TGTGGAAAA**GGGACTTTTCCCTATAGAAAACATTCACTTGGATTGTTATTGGATGTATTGAAAAGGGCTTAGGAGGCTGCTGACAGACAGCGCTGAAGCT
GTGGACGAAAAGCGTCGTTATACACAGCGCCAGCTACCCAGCGCTTGTCCACCGCTGCATACGCCTGCTGTCACCGTGACGGTCCCCAACGCGGGCATCG
CGCTGGAGGACACGGCAGAGGCGGGCTGCGTCGAGATAGGCTGAAAAACTATCCACAGGCGAAAAAAACCGCTTGGGATAACCTTTCTGGCGCTATCGCG

GGCAGATCTAGAGACGATTCGAAGGTGATTCGGGGGTGATGCCGGGAGGAATCCGGTGCTCAGGGCGGGGAAAAGGGGGGCATGACGCGAAGACAGAGCG
GCGATAGGAACCGCAGCCAGCCAGGCGCGCGTGAGAGACGCTCGAAGCTAGCGCGAACGGGCCACCGTCGAGGTGGCCGCGTTTGCGCCGAGTGGGGGCG
GGCTCAGTGCTTGAGCACGCGGGAAAGGAAGGCGCGGGTACGCTCGTTTTGGGGATCATCGAAGACGGCCTCCGGCGTTCCCTGCTCGACGATCTCGCCG
CCATGGACGAATACCACACGGTCGGCGACCTCGCGTGCGAAGCCCATTTCGTGGGTCACGATCATCATGGTCATGCCGTCCTTGGCCAGCTCGCGCATGG
CATCCAGCACTTCGCCGATCATCTCCGGATCCAGCGCTGAGGTCGGCTCGTCGAACAGCATCAGCCGTGGTTCCATGGCCAGTGCCCGGGCCAGCGCCAC
GCGCTGCTGCTGACCGCCGGAGAGCTGGCTGGGGTACTTGTCAGCCTGGTCGGAGATTCCCACGCGCTCGAGCAGACGCTCGGCGGTGGTGTTGGCCTGC
CGGCGCGACCAGCCACGCACCTTGCGCGGGGCCAGGGTCACGTTGTCGCGAATCGACAGATGAGGGAACAGGTTGAACTGCTGGAACACCATGCCCACTT
CGGTGCGTACCGCCTGCAGCGACTTGCCGCTGGCGCCGTGGGGGGCCAGGGTCTTGCCATCGACCTCGATGCGTCCGGACTGGAACGGCTCCAGCCCGTT
GACGCAGCGAATCAGGGTCGACTTGCCCGAGCCGCTGGCACCGATGATCACCACCACTTCGCCCGGGGCCACGCTCAGGTCGATGTCGCGCATCACGTGC
AGCTCGCCGAAGTGCTTGTTGACCTTGTCGAGGCGCACGATCGGCGGCGCATCCCGCTGGATCTGCTTGTCGTCGTTCATCCCACCAGCCCCTTGCGTTC

>Salinicola endophyticus CPA92 [NZ_PZJO01000003.1:365462..365601], dnaA> ig dnaN>, 140 bp

CCCCATCGGCTCGGCAAGCGCCGCCAGGGGATTGCCCGTATCGCCCCGGGCAGCAAGAATGAACCCCAGCCGGCAGCGTCGCCGCGGCCGCTCGCGGCGC
CGACGTTTCTTTCAGGATCGAATTCAACCACGGAGCTCCC

WebSkew GCmin pos. 286665 (GCmin<>oriC1 38.9 kb; GCmin<>oriC2 13.7 kb; oriC2<>oriC1 25.2 kb)

>Salinicola halophilus CECT 5903 [NZ_PZJV01000006.1:c247540..248039], <rpmH oriC1 dnaA>, 684 bp

GCGCGGAGAGGGCGCCGTCGCGATCGCCATCGTTACCCGGGGCGGCGGTGGTGACTTCCGGCGCGGGGTTGTTGACGACGCTGTGACCGTAGTCCTGATT
CCACTGCATGATCAGCAGATAGGCGAGTAACGCCAGCGGAATCACGAATAGGAGTCGTTTGAAATCCATAGAAGGGTCTGCCCGGTAAAGGAATGGGTCG
ATGGGCGCCGGGCATCGGCGGTGGTCCGGTGGTGGACACCGAAACTCCCCGGCCGTGGGCGCGGGGATGTCCGATCAGGTCGAGAGCGACGAGCCGCGTC
TAGCGAGCGGACTTCTCCAGCCGCCGCCACATGCCATGGAGCTGGCGAGAGACGGTGGCGTTATCCAGCTCGTTCACTCCACGTCGCGCAAGCAAGACGA
TGTCCACGCCGGGCAGCTGATGCTGACGATGCCGGATGGATTCGCGGACGAGACGCTTGAGACGATTACGGTCGACGGCACGCCGCACGTTCTTCTTGCT
GAAGATCAACCCGATCCGCGGATGGCCCAGTGTGTTGGGACGAGCCAACGCCAGCATTCCTTTACCGTGGACCTTGAAGGGAGCGTGGTCGAAAACGTGC
TTGTAGTCCCCGGCATTCAGCAGTCGCAGCTGCCGGGGAAAACCTTGATCGAACACCCGAGGTCCGATCAGGCGCTGAGACGCTTGCGGCCTTTGGCACG
ACGACGGTTGAGGACCTGACGGCCGTTCTTGGTGGCCATGCGTGCGCGGAAACCGTGATTGCGCTTACGCTTCAGAACGCTCGGCTGAAAAGTACGTTTC
ATGGTGGCTGATTCCCAATTGGTTGCGTGTTAATGCCGCTGACTAGCCGTGCAAGTGACTGACTAGAAAGCAAAACGGGTTCGAAACGGCACGCCCGATG
GGTTCGCAGGGACCGGGCTCGCTCGGCTCAAGAGCGGGAATTCTAGAGAATCCTGGGAAGCGGTGCAATTTTTCTGTGGATTATCCACACCCCTTTTGAT

GTGAATTTCGCGCCAGCCTGTGGGCAAAATTGTGGGCAATATCGGATCGACGCAGTCGGCGCCATTCGGCTCTTGGCGCGGCTTTCGCGCCGCTGCGGCG
TCGCGTTTTCCACCCGGGAGTCGGTCACTGGCTCGGGTGGGGATGGGTACCAGAATAATTTTGTCAATCGCACCATGTGGTTAAAAGTGGGGACACAATT
**TGTGGTTAA**CTCCACTTAACCCAAAAAATTCAAAAACATACGGCTATTTATGATGTGTGCATAACCTGCTGGATAACGGTGAATCAAGCGTGGATGGCTG
TGTCCTGTGGGCGTTTCGATGATGAGCGGTGATCGACATTGAGAGAAGTTGTCCCCGCGGCTTCTACCGCTTGCCACTAGGTTATCCACAGGACGACTTC
TCGCTTACCTCGTGTGGATAACCTGAGCGCTGAGCGCTACAATGCCACCCCGATTTTCCCAACTCTTCAAATTCTCGTGAGGTCGCGTGTCGGTCGCTCT

CTGGCAACAATGTCTCAATGCGCTGCAGGATGAGTTGAGCTCGCAGCAGTTCAACACCTGGATTCGTCCGCTGCAGGCGGAGGACGATCCGTCGGACGAG
CTCTGCCTGCTGGCGCCCAATCGCTTCGTGCGCGACTGGGTCAGCGACAAGTACTTGAAGCGCATCAACGAACTGCTGCGGGAAGTTTCCCCTTCCAGCA
AGCCGCCGAAAGTGCGTCTGGTAGTGGGTAGTCGCGGCACGGGGGCGGTTTCGGCGCGCGGACTGGGGGCGCCGGTGTCGGCACCGACACCGAGTCAGCC
TGCCGCTCCCAGCGTGCAGACGACGTCCCGCTCTTCGAGCGCGAGTGACAATCGCGAAGAGGACCGTGCGCGCGAACAGGCGCCAGCGCGGGAGGAACCC
AGCCGGCGCGGTAACCGTGAGCGTCAGGTCCAGGTCGAAGGGAGTCTGAAGCACGGCAGCGGGCTCAACCCCAACTTTACCTTCGAGACCTTCGTCGAAG
GCAAGTCGAACCAGTTGGCCCGGGCGGCCTCGCGTCAAGTCTCCGAGAACCCCGGCGGCGCCTACAATCCGCTGTTCCTCTATGGCGGCGTGGGGCTGGG
TAAGACCCACCTGATGCATGCAGTGGGTAACCACCTCGCGGGGATGCGCGACAACGCCAAGGTGGTCTATCTGCACTCCGAGCGCTTCGTCGCCGACATG
GTCAAGGCACTGCAGCTCAATGCCATCAACGACTTCAAGCGCTTCTACCGCAGCGTCGATGCGCTGCTGATCGATGACATTCAGTTCTTCGCCGGCAAGG
AGCGCTCTCAGGAGGAGTTCTTTCACACTTTCAACGCCCTGCTCGAGGGCGGCCAGCAGATGATTCTGACCTCGGACCGCTATCCGAAGGAGATCAGCGG
CGTCGAGGAGCGCCTGAAGTCGCGCTTTGGCTGGGGTCTGACGGTGGCGATCGAGCCGCCGGAACTCGAGACGCGCGTGGCGATCCTGATGAAGAAAGCG

>Salinicola halophilus CECT 5903 [NZ_PZJV01000006.1:c272720..273219], <mnmG oriC2 <COG1126 (glnQ), 850 bp

GGTGGCTCGTCTTGTCGGCGAAGCGGTGAACCTTGTCCTCGATCGACGGGCAGTAGCGCGGACCGACCCCGCCAATAACGCCGGAGTACATCGGCGACCG
ATCGAGATTGGCGAGAATGATCTCATGGGTTCGGGCGTTGGTGTGGGCGATGAAGCAGTCCACCTGGCGCGGATGCTGTTCCCGTGAGCCGAGGTAGGAC
ATCACCGGCAGCGGCGTATCGCCCGGCTGACGTTCGAGCTGAGAGAAATCCACGCTCTTGGCATCGATTCGCGGCGGCGTTCCGGTCTTCAGGCGATCGA
CGCGAAACGGCAGCGCTCGCAGTCGTTCGGCAAGCGCGTTGGCGGGCGGGTCGCCGGCGCGGCCACCGGCGTAGTGATCCATGCCGATGTGGATAACCCC
GCCGAGGAAGGTACCGGTACACAGCACCACGGTATCGCCCATGAAACGGATACCGGTCTGGCTCAAAGCACCGATCACCGTATCGCCGTCGACGATCAAG
TCATCCACACCCTGTTGGAACAGGGTGAGATTCGTTTGGTTCTCCAGCAGATGCCGGATGGCTGCCTTGTAGCGAACGCGGTCCGCCTGGGCTCGAGTCG
CACGGACTGCCGGGCCCTTGCGTGAGTTGAGCACGCGAAACTGGATACCTGCAAGGTCGGTCGCCGTTGCCATGGCACCGCCGAGTGCATCGATCTCCTT
GACCAGATGACTCTTGCCAATGCCGCCGATGGCGGGATTGCAGGACATCTGACCCAGCGTCTCGATGTTGTGAGTCAACAGCAGTGTCTTGGCACCGCTA
CGAGCGGCTGCCAGAGCCGCTTCGGTGCCGGCATGGCCGCCACCGATGACGATGACCTCAAAGCGGTCGGGGTACTCCACGTCTCACCTCGTTGAACGCG
CATCGATGCGTTCGACAGCGAATGAATCTCGTTGATCGCTTCTGTGACGAGCCGGCGCGTTAGCGACGGCGGCGATCCGCGACCCGGTCTTCCTCGATTC

GACGTTGGGTGATGACACGCTCGTTGAATCAGGGCCGGTATCGCAAGGGCCGCACAGTATAAACGTCCTGTGGGTAACCGTCAGGGGTTTTCCACACCGT
CTCGCTATGCGACGGGTCAGTCATCAGTATCAATATTAAATAAAAGAAAGAATATAAAAGAACTTCTGTAATGGTAATGACTATTGGTGTGGATGATTTC
**TGTGGATAT**TCTAAAAAACTCTAATATTTCCAATAGCTTGTGCTGTTATTCAAAACCGTTGAAAGTTCTGACGAGGCTGGCGACAAGCGGTCCGGATCCT
GTGGATGAAAGCGTACATTGTCCACAGCCGTGATGGCGCACGGGTTATCCACCGATGCATACGCCCGCTGTCACCGTAGGCCTCAACAGATGATCGATCG
CGCTGTGACGACTGCCATGAGCGGGATGGAAGCGAATTCGTCGAAAAACTATCCACAGGCCCAAAAAGTCGATGTGAATACCGATTAGCAGGGTGCCATT

CTCACGATGAGCTCCTGATCGATGCCCGCAGGGCGGTCATGATCGAGCGCTGAGCGCGAATGCTGTGCCGTGCTCGCTGAAGGCGCGAGCGCGGAATCGG
CATCGAAAGAGCGAGCGAGAGTCGAGAAGAGAGAGCGTTGAGAAGAGAGGAAAAGAGGCTGTCGGTGCGAGGGAAGCGAGGAGACCAGAACAAGCGCGAG
CGAACTCCGAACGGGTCCGCTCGCGGGGGGCTAGTGCTTGAGCACGCGTGAGAGGAACGCCTGAGTCCGTTCATGACGTGGCTGGTCGAAAACGCGCTCT
GGCGGCCCTTGCTCGACGATCTCGCCGCCGTGGATGAAGACCACGCGGTCGGCGACTTCGCGGGCGAAGCCCATTTCGTGGGTGACGATCATCATCGTCA
TGCCGTCCTTGGCCAGCTCGCGCATGGCATCGAGCACCTCGCCGATCATCTCCGGGTCGAGGGCCGACGTGGGCTCATCGAACAGCATCAGTCGCGGCTC
CATGGCGAGCGCTCGCGCCAGCGCCACGCGCTGCTGCTGGCCGCCGGAGAGCTGTCCCGGATACTTGTCCGCCTGATCGGCAATGCCCACCCGCTGGAGC
AGCCGGTCCGCGGTCGCATCGGCCTGCTGACGCGTCATACCGCGGACCTTGCGCGGTGCCAGCGTGACGTTGTCGCGGATCGAAAGATGGGGAAACAGAT
TGAATTGCTGAAATACCATGCCGACCTCGGTGCGCACCGCCTGGAGCGATTTGCCACTCGCCCCGTTGGGCTCGAGGTATTTGCCATCGACTTCGATCCG
GCCACCCTGAAAGGGTTCGAGGCCGTTGACGCAGCGGATCAGCGTCGACTTGCCCGAGCCGCTGGCGCCGATGATCACTACCACTTCTCCCGGCGCCACA
TCGAGATCGATGTCGCGCATGACGTGCAGCTTGCCGAAGTGCTTGTTGACCTTCTGCAGGCGCACGATGGGCCGTGTGTCGGTCGCGCTCATCCCACCAC

>Salinicola halophilus CECT 5903 [NZ_PZJV01000006.1:c245924..246065], dnaA> ig dnaN>, 142 bp

GCGCACCATCGACTCGCCGCGCGGGGATTGCGGCCCGGGCACGGCGGGGCAAGAATGGTTCCAATCGCACCGACCAACAATCATCGAACGTCTCGGTATC
GAATCCACCTGACGCCGGATCGACATCTCTAGTGGAGCTCCA

WebSkew GCmin pos. 147312 (GCmin<>oriC1 31.1 kb; GCmin<>oriC2 6.1 kb; oriC2<>oriC1 25.0 kb)

>Salinicola aestuarinus CPA62 [NZ_PZJP01000004.1:c115964..116463], <rpmH oriC1 dnaA>, 683 bp

GCGCGGAGAGGGCGCCATCGCTTTCGCCGTCGCTCCCGGGAGCGGCCGTGGTGACTTCCGGCGCCGGGTTGTTGACGACGTTGTGACCGTAGTCCTGATT
CCACTGCATGATCAGCAGATAGGCGAGTACCGCCAGCGGAATCACGAATAAGAGTCGTTTGAAATCCATAGAAGGGTCTGCCCGGTAAAGGAATGGGTCG
ATAGGCGCCGGGCATCGGCGGTGGTCCGGTGGCGATCACCGGCACTCCCCGGCCGTGGGCGCGGGGATGTCCGATCAGGTCGAGGGCGACGAGCCGCGTC
TAGCGAGCGGACTTCTCAAGCCGCCGCCACATGCCATGTAACTGGCGTGAGACAGTAGCGTTGTCCAGCTCATTCACCCCGCGTCGCGCGAGCAAGACGA
TATCCACGCCCGGCAACTGATGCTGACGATGACGAATGGATTCGCGGACGAGGCGCTTGAGGCGATTACGGTCGACGGCACGACGCACGTTCTTCTTGCT
GAAGATCAACCCGATCCGCGGATGGCCCAGTGTGTTGGGACGAGCCAGCGCCAACATTCCTTTACCGTGGACCTTGAAGGGAGCGTGGTCGAAAACGTGC
TTGTAGTCCCCGGCATTCAGTAGTCGAAGCTGCCGGGGAAAACCTTGATCGAACACCCGAGGTCCGATCAGGCGCTGAGACGCTTGCGGCCCTTGGCACG
ACGGCGATTGAGGACCTGACGGCCGTTCTTGGTGGCCATGCGTGCGCGGAAACCGTGGTTGCGCTTGCGCTTGAGGACGCTCGGCTGAAAAGTACGTTTC
ATGGTGGCTGATTCCCAATTGGTTGCGTGTTTAAGCCGCTCACTAGCCGCCCAAGTGGTTGACTAGAAAGCAAAACGGGTTCGAAACGGCACGCCCGATG
GGTTCGCAGGGACCGGGCTCGCTCGGCTCAAGAGCGGGAATTCTAGAGAATCCTGGGAAGCGGTGCAATTTTTCTGTGAATTATCCACACCCTGATCGAC

AACCGTTTCGCCTCGCCCTGTGGGCAAAATTGTGGGCAACACGATGTCGTCCTAGCCAGCGCCATTCGGGTGTCAGCGCGACTTTGGCCATTAGCGGGG
CCGTGATCAAGACACGGCGAAAGGTGCACCGCTCGGCCCGCCACGGGGAGCATGCCGCGCTTGTCAACCGCATCATGTGGTTAAAAGTGGGGACATCATT
**TGTGATTAA**CTCCAATTAATACAATTAAATCATGTACCTATACGCAGTTTGTGACTGTGCATAACATGCTGAATAACGGTGGCCACAGGTGGATGACTT
CGCCTGACAGGGCGGGCTGTCCCGTGACCCGTTCCTCCCGATAGTGAGTTGTCCCGAGATCCTCTACCGCTTGTCACTGGGTTATCCACAAGGTGTCGTC
TTTCGTACCGCGTGTGGATAACCCGGACGCTGGACGCTACAATGCCCACCGCTTTTCACCACCCTTCAAAATTTCGTGAGGTCGCGTGTCGGTCGCTCTC

TGGCAACAATGTCTCAACGCGCTGCAGGATGAGTTGAGCTCGCAGCAGTTCAACACCTGGATTCGTCCGCTGCAGGCGGAGGATGATTCGACGGACGAAC
TCTGTCTGCTGGCACCCAACCGCTTCGTGCGTGACTGGGTCAGTGACAAGTACCTGAAGCGTATCAACGAGCTGCTGCGTGAGGTCGCCCCCCTCCAGCAA
GCCACCCAAGGTGCGCCTGGTGGTCGGTAGCCGGCGGACCGGCGCCGTGTCGACGCGCGGGCTGGGGGCACCGGTCTCGGCGCCGACGCCCAGTCAGCCG
GCACCGAGTCGGCCGGCGGCGCCCAGCGTGCAGACGACCCCCCGTTCTTCACACGCGAGTGACAACCGCGAAGAGGATCGAGCGCGCGAGCAGGCGCCGG
CGCGGGAGGAGTCCAACCGGCGTGGCAACCGTGAACGCCAGGTTCAGGTCGAGGGGAGCCTCAAGCACCAGAGCGGCCTGAATCCCAACTTCACCTTCGA
AACCTTTGTCGAGGGTAAGTCCAACCAGCTGGCCCGCGCGGCCTCGCGGCAGGTCTCCGAGAATCCCGGCGGTGCCTACAACCCGCTGTTTCTCTACGGC
GGCGTGGGCCTGGGTAAGACCCACCTGATGCACGCGGTCGGCAATCATCTCGCCGGGATGCGCGACAACGCCAAGGTGGTCTATCTGCACTCCGAGCGTT
TCGTCGCGGACATGGTCAAGGCGCTCCAGCTCAACGCCATCAACGACTTCAAGCGCTTCTATCGCAGTGTCGACGCCCTGCTGATCGATGACATCCAGTT
CTTCGCTGGCAAGGAGCGCTCGCAGGAGGAGTTCTTCCACACCTTCAACGCCCTGCTCGAAGGTGGCCAGCAGATGATCCTGACCTCGGACCGCTATCCG
AAGGAGATCGCGGGTGTCGAGGAGCGGCTGAAGTCGCGCTTCGGTTGGGGGCTGACGGTGGCGATCGAGCCGCCGGAGCTGGAAACCCGCGTGGCGATCC

>Salinicola aestuarinus CPA62 [NZ_PZJP01000004.1:c140951..141450], <mnmG oriC2 <COG1126 (glnQ), 800 bp

GTCGAGGCCCTCGGGCTCGATGAAGATCTGGTGGCTCGCCTTGTCCGCAAACCGGTGGACCTTGTCCTCGATCGACGGGCAGTAGCGCGGGCCGACACCC
CCGATCACACCGGAGTACATCGGTGAGCGATCGAGGTTGGCGAAGATGATGTCGTGCGTCCGGGCGTTGGTATGCGCGATGAAACAGTCGACCTGCCGCG
GATGCTGTTCGCGGGAACCGAGATAGGACATCACCGGCAGCGGCGTATCCCCAGGCTGACGTTCGAGCTGAGAGAAATCCACGCTCTTGGCATCGATCCG
AGGCGGCGTCCCGGTCTTGAGCCGATCGACCCGGAACGGCAGGGCACGAAGTCGCTGAGCGAGGGCATTGGCAGGCGGATCGCCGGCACGGCCACCGGCA
TGGTTGTCCAGCCCGATGTGGATAACCCCACCGAGGAAGGTACCTGTGCACAGCACCACGGTGTCCGCCATGAAGCGGATGCCGGTCTGGGTCAGCGCCC
CGGTGACGGTGTCCCCCTCGACGATCAAATCGTCGACGCCCTGCTGGAACAGCGTGAGATTGGGCTGATTTTCCAGCAGGTGACGGATGGCTGCCTTGTA
GCGAACGCGATCCGCCTGGGCCCGAGTGGCACGAACTGCCGGGCCCTTGCGGGAGTTGAGCACGCGGAACTGGATGCCGGCGAGATCCGTAGCCGTTGCC
ATGGCACCGCCGAGCGCATCGATCTCCTTGACCAGATGACTCTTGCCGATGCCGCCAATGGCCGGATTGCAGGACATCTGGCCCAGCGTCTCGATGTTGT
GAGTCAGCAGCAGGGTCTTGGCACCGCTGCGAGCGGCAGCCAGTGCGGCTTCGGTGCCGGCATGTCCGCCACCGATAACGATGACCTCAAAACGGTCAGG
GTACTCCACGTCTCACCTCGTTGAACGCGTTTCGAAGCGTTCGACAGCGAATGAATCTCGACGATCGTCGGGGCGCCAGGGTCTGCCCTTGGCCGGCGGC

GATCACTGCGACAGGGCGGGTCGACAACTCGATCCGATATCGCAAGGGCCGAACAGTATAAACGTCCTGTGGGTAACCGTCAGGGGTTTTCCACACTCGG
TCGCCTTGGGTCTCAGCCAGTCATCAGTACCAATATTAAATAAAAAGAAAAATATAAAAGAAGTTCTGTAATGGTAATGACTATTGGGGTGGATAGTTTC
**TGTGGATAT**GTCAAAAAACCGAATGAAATTCAGCTTTTTGGGCTGTTATTCGATGGAGTTGAAAGCTCTGGCGAGGCTGACGACAAGCGGTCTGGATCCT
GTGGATGAAAGCGCTTGTTGTCCACAGTCCTGATTCCGCACGGGTTATCCACCGATGCATACGGTCCTCGTCAGCGTAGTGGTCAACAGATGGGGAAATC
GGCTGCAGACACTGGCATGTGCGGGCGCGGACCGATTCCACCAAAAAAACTGTCCACAGGGCAAAAGGCGCGCGATGGATAACCTCAAAAGGGGCAAGCC

GACGTCGACAGGCGACAAAAAGCGGAGTGAGGCGTCGTCAGCGCAGTCATTATTGGCGCGGCTGTCGTCAGTCCGAATCGCCGGACGGAGAGAGAGGAAC
GTGAGAGGCGGGGACGAAAGGGACGGGGACGAGCGAATCCACAATGGATTCGCTCAAACGGCGACTTGAGACGTCGCCGGGATTCGGTACAGACCGGTGT
GGCGAGACGTCAGTGTTTCAGCACGCGGGAGAGGAAGGCCTGGGTGCGTTCGTGTTGGGGCTCATCGAAGACGCGCTCCGGGGGTCCTTCCTCGACGATC
TCGCCCCCATGAACGAAGACCACGCGGTCGGCGACCTCGCGGGCGAACCCCATCTCGTGGGTGACGATCATCATGGTCATGCCGTCCTTGGCCAGCTCGC
GCATGGCGTCGAGCACCTCACCGATCATTTCCGGGTCGAGCGCCGAGGTGGGCTCGTCGAAAAGCATCAGTCGAGGCTCCATGGCAAGTGCCCGGGCGAG
GGCCACGCGCTGCTGCTGACCGCCTGAGAGCTGCCCGGGGAACTTGTCCGCCTGGTCGGCGATGCCCACGCGATCGAGTAGCCGGTCGGCCGTCGCGTCC
GCCTGCTGTTTCGTCATGCCGCGAACCTTGCGCGGTGCGAGTGTGACGTTGTCGCGTATCGAGAGGTGGGGAAACAGGTTGAACTGCTGGAACACCATGC
CGACTTCGGTGCGAACCGCCTGCAGCGACTTGCCACTGGCGCCGTTGGGTTCCAGGGTCTTGCCATCCACTTCGATGCGCCCCCCCTGAAACGGCTCGAG
GCCATTGACGCAGCGGATCAGCGTCGATTTGCCGGAGCCGCTGGCGCCAATGATCACCACGACTTCGCCGGGGGCGACGTCGAGGTCGATGTTGCGCATG
ACGTGCAGATTGCCAAAGTGCTTGTTGACCTTCTGCAGCCTGACGATCGGCCATTTGTCGGTCGTGCTCATCCCACCACCCCCTTGCGTTCCAGCAGGTT

>Salinicola aestuarinus CPA62 [NZ_PZJP01000004.1:c114334..114475], dnaA> ig dnaN>, 142 bp

GCCTGCTCCCTGCTCGTCGCTCGGGGATTGCGGCCCGGGCGCGGTGGGGCAAGAATGGGTCCAATCGCACCGCCCAATAATCATCGAACGTCTCGGTATC
GAATCCACCTGACGCCGGATCGACATCTCTAGTGGAGCCTCA

WebSkew GCmin pos. 147519 (GCmin<>oriC1 25.4 kb; GCmin<>oriC2 1.7 kb; oriC2<>oriC1 23.6 kb)

>Chromohalobacter israelensis DSM 6768 [NZ_JQNW01000005.1:c121901..122400], <rpmH oriC1 dnaA>, 589 bp

TCGGTACGCACCTGGATGAGGCCGTTGGCGTCGGTCTGGGAGTTCTCGCCGGGCATGCCGGCACTGTCTGGCGATGCGTTATTGCCGGCGTTGGAAGCCG
AAGGCGCCGACAAGCCTTCACTGGCGTCATCGCTATCGGTGGTACCTTGCGCCACGCTGGTGGGAGCGCTGTGTTCGACATTGTTCTGCCCGTAATCAGC
GTTCCACTGAATGACCAGCAGATAGGCGAGCACCGCGAGCGGGATGACGAGTAAGAGTCGTTTTACGTCCATAAGGATTCTGCCCGGTGAAACATTCTGA
CGACACATCCCCACCGGCGGAGCGCACACGGCGGGCGTGGCGGGAAAGGCCGATCAAGGGGATGGGGTGGACGCGACCGACTGCTTGCGAACGTCTTTTT
CCAGGCGCCGCCACATGCCGTGCAGTTGGCGATGCAGGGTAGCATTGTCCAGTGCATCAACTCCGCGACGAGCGAGAACGATGATGTCCACGGCGGGGAG
ACGATGTGACTGGAGGCGAAAGGATTCACGGACGAGGCGTTTCAGACGGTTACGGTCGACGGCACGGCGCACGTTCTTCTTGCTGAATACGAATCCCACC
CGCGGGTAGCCGAGAGCGTTGGGGCGCGCAAGCACCAGAAGGCCCTTGCCGTGCACCTTGAAGGACGCATGAGCGAAGACGTGCTGATAGTCCCCGGCGG
TAAGCAGACGCTGTCGCCGGGGAAAGCCCTGATCGGACATTCGTTGCTCAATCAGGCGCTCAGACGCTTGCGGCCCTTGGCGCGACGGCGAGCCAGCACC
TGGCGGCCGTTCTTGGTGGCCATGCGAGCACGGAAACCGTGCACGCGCTTGCGCTTTAGAACGCTCGGCTGAAAAGTGCGTTTCATAGCTGCGAATTCTC
ACGTGGTTGTTTGGAAACGGTTGAAACTGAAGAGCCCGGCACGTTGGCACCAGGCGGTATTCGGCTCAAGACCGGCAATTCTAGTGAATCCCCAGGGCAG

GTGCAATTTTTCTGGCGCTTTTATCCACACCTGTTTTTGCGACGGCCCTCGCCGGGGGAGCGCGATCCGTCTCGAGACTCGAAGGTATGGGGACATCATC
ATGTGCTGAAGAGAAGGGGGCTTTCCACCGTCCAACAGCCACTTCCATTACTACAGGTTTGTTTTCTACTGATGTTGTTATTAATGGTGGGGACAAATTT
**TGTGGATAA**CCCAAAAAAAGATCAGGATGTCATGTAGTTGGCGTGTTCCTCTTTGTGTGGGGAAGTTGCGGGCGTCCCGTGGGAAAGCTGTCGATGCCGT
TGTGGGTAACACGTCGTCTGTCCACAGGCACGCCGGCGTCCCCGATGTATGGACAATCTGTCCCCGTGGTTGGCGGCCCGGTTTTCCACAGGAAAAGAGA
GCGGGGATCCATGTGGATAACCTCGGTCTCGAACGCTACAATACTCGGCCATTCATCAACGCTAGGTGAGGTCGTGTGTCGGTCGCTCTCTGGCAGCAAT

GCCTCAACTTTCTTCAGGACGAACTGAGTTCACAGCAGTTCAACACCTGGATTCGCCCACTGCAGGCGGAAGACGGTGACGCCAACGAGCTGTGTCTTCT
GGCGCCCAACCGCTTCGTGCGTGACTGGGTCAACGATAAATACCTGAAGCGCATCAACGAGCTGCTGCGGGACTTGGCATCCGGTAAGCCGCCCAAGGTG
CAGCTGACGGTGGGGAGTCGCCGCAACGTGGCGATGTCGTCGCCGCGTGACCTGGGCGCGCCGGTGTCGGCGACGACGATGAATGCCTCGCGCCCGACCG
AGGCGCCGGCGGTGCATGCCGCTCCGCGTGCCAAGGGGGACTACGCCGACGAGCAGGAAATCGATCGCCTTCGCGAACGCGAGGAAACTCCGCGGCGCGG
TGGTGGCGAGCGCCAGGTCCAGGTGGAGGGGAGTCTCAAGCACCAGAGCGGTCTCAACCCCAATTTCACCTTCGAGACATTCGTCGAGGGCAAGTCCAAC
CAGCTTGCGCGTGCCGCGTCACGGCAGGTCGCCGAGAACCCCGGTGGTGCCTACAATCCCTTGTTCCTGTACGGTGGGGTCGGTTTGGGGAAGACGCACC
TGATGCATGCCGTCGGCAACCATCTGGCGGGGCAGCGCGAGAACGCCAAGGTCGTCTACCTGCATTCGGAGCGTTTCGTCGCCGACATGGTCAAGGCGCT
ACAGCTCAACGCCATCAACGACTTCAAGCGTTTCTATCGCAGCGTGGATGCCCTGCTGATCGACGACATTCAGTTCTTCGCCGGCAAGGAGCGTTCGCAG
GAAGAATTCTTCCACACGTTCAATGCGCTGCTCGAGGGCGGCCAGCAGATGATCCTGACCTCGGACCGTTATCCCAAGGAAATCAGCGGGGTGGAGGAAC
GTCTCAAGTCGCGCTTTGGCTGGGGACTGACGGTCGCCATCGAGCCGCCGGAGCTGGAAACGCGCGTGGCGATCCTGATGAAGAAGGCCGACCAGGCCAA

>Chromohalobacter israelensis DSM 6768 [NZ_JQNW01000005.1:c145515..146014], <mnmG oriC2 <COG1126 (glnQ), 601 bp

CGAACTGCACGTCGAAGGGCAACGAGGTCGAGATCCCGTTGGGATAGAGCTCGTGGGTATCCAGCCCCTCCGGCTCGACGAAGATCTGATGGCTCGACTT
GTCGGCGAAGCGGTTGACCTTGTCCTCGATGGACGGGCAATAGCGCGGGCCCACACCGCCGATCACGCCGGAATACATCGGCGAGCGATCGAGGTTGGCA
TGGATGATCTCGTGCGTGCGCGCATTGGTGTGAGCGATGTGGCACGCGACCTGACGCGGATGCTGATCCCGACGGCCCAGATACGACATCACCGGGGTCG
GGTCGTCGCCGGGCTGGGTCTCGAGCACCGAGAAATCCACGCTCTTGGCGTCCAGACGCGGGGGAGTCCCGGTCTTGAGCCGGTCGACGTGAAACGGCAG
GGCACGCAGCCGCTGCGCAAGGGCATTGGAGGGCGGATCACCGGCACGCCCCCCGGCATGATGATCCAACCCGATGTGGATAACGCCACCCAGGAAGGTA
CCGGTGCACAGCACAACGCTCTCGCCGAACACGCGAATGCCGGTCTGCGTCACGACTCCGCGGACCGTGTCGTTTTCCACGATCAGGTCGTCCGCGGCCT
GCTGAAACAACGTCAGGTTGGCCTGGTTTTCCAGCATGCCGCGAATCGCTGCCTTGTAGCGCACCCGGTCGGCCTGGGCCCGCGTGGCGCGCACCGCCGG
TCCCTTGCGGGCGTTGAGCACCCGGAACTGGATGCCACCCAGATCGGTGGCATGCGCCATGGCGCCGCCCAGGGCATCGATTTCCTTGACCAGATGACTC
TTGCCGATCCCGCCAATGGCCGGATTGCAGGACATCTGCCCGAGCGTCTCGATGTTATGGGTCAGCAGTAACGTTTGACAGCCCATGCGAGCGGCAGCCA
ATGCGGCTTCTGTCCCCGCATGACCACCGCCGATGACGATCACATCAAAGCGGTCTGGGTAATCCACGTTGCACCTCGAATCGCAGCGTCTGGCTGCGGT

CGTACATGTCGACAGGCGTCATCGAGCGAGGACGGCGCCTGGGAAATAGCCGGACAGTATAAACCCCTTGTGAGTAACCGTCAGGGTTTTCCACACATGG
ATTGTCCGCCGGTGGTGTCTTCGTCCTTGGGTCTATAAATAATAATAAGTGTGTAGAAAGAAGTTCTGTTGCTGTTGTAACTACTGGTGTGGACCAAATC
**TGTGGATAA**GCGTTAAAAATCTTTCATTCACAATTGCTTGCGATGTTAGTCGAACGTTGTAGAAGCGGCAGACAGGCTGCGTGCAGGAAGGGGACGCCCT
GTGCATGAAAATGCCGGTTGTCCACACCGCGAAACGTGCACCACTTATCCACGGGTGTATACCTGATCGCCCCCCGTAGCCGTCAACAGTCAGGCACGCG
TCTCTGGAACCCCGTAGTGACGGGGGCTGGAGGCGATATTTCATGGGCAGGAAAAAAATATTGTCCACAGGCGTGAAATCGGCCTGTGAAGCGCCGTGAC

AACGCACGGCAACACCACGATCCGCGAGGGGATCGTGGCGTCAGTCGAGGGGGAGAGTCGGGAAAAGGTCAGTGCTTGAGCACGCGGGAGAGGAAGGCCT
GGGTGCGTTCGTGCTGGGGATGGTCGAAGACCTGGCTGGGCGAGCCCTGTTCGACGATGCGTCCGGCGTGGACGAAGATGACGCGATCGGCGACCTCGCG
GGCGAAGCCCATCTCGTGGGTGACGATCATCATGGTCATGCCTTCGCTGGCCAGTTCACGCATGGCATCGAGTACCTCGCCGATCATTTCCGGATCGAGT
GCCGAGGTGGGCTCGTCGAACAGCATCAGGCGCGGTTCCATCGCCAGGGCGCGGGCCAGGGCGACACGCTGTTGCTGGCCCCCTGAGAGCTGCCCGGGAT
ACTTGTCGGCCTGATCGCTGATGCCGACCCGTTCGAGAAGGCGATCTGCGATTTTGGATGCCTGGGCATCGTCCAGACCGCGGACCCGGGTGGGGGCCAG
CGTGACGTTGGCCCGTACCGACAGATGCGGAAACAGGTTGAACTGCTGGAAGACCATGCCGACCTCGGTCCGGATGGTCTGCAGCGATTTGCCCCCCTTG
CCGTACGGTGTGAGTTCGTTGCCATCGACGATGATGCGGCCGGACTGGAACTCTTCCAGGCCGTTGACGCAGCGGATGAACGTGGATTTCCCGGAGCCGC
TGGCGCCGATGATGACGACCACCTCGCCGGCGGTGATCTCGAGATCGATGTCCTGCATCACGTGCAGCTTGCCGAAGTATTTGTTGACCTTCTCCAACTG
CACGATCGGCGCGGCGGAAGCTTGTGCGTTATCACTCATGATGGGCGCCTCATTGTCCGACCAGACCCTTGCGTTCGAGCAGACGCAAGGCGATAGAGAG
ACTCAGGGTGATCACCAGATACAGCATGGCGACCATGAAATAGACTTCCAGGGCGGTGAACGTGGTGGCGATGTAGACCTGTCCCTGGCGTACCAGCTCA

>Chromohalobacter israelensis DSM 6768 [NZ_JQNW01000005.1:c120389..120437], dnaA> ig dnaN>, 49 bp

WebSkew GCmin pos. 3696649/0 (GCmin<>oriC1 0.1 kb; GCmin<>oriC2 23.8 kb; oriC2<>oriC1 23.6 kb)

>Chromohalobacter salexigens DSM 3043 [NC_007963.1:3696267..3696649], <rpmH oriC1 dnaA>, 589 bp

TCGGTACGCACCTGGATGAGGCCGTTGGCGTCGGTCTGGGACTTCTCGCCGGGCATGCCGGCACTGTCTGGCGATGCGTTATTGCCGGCGTTGGAAGCCG
AAGGCGCCGACAAGCCTTCACTGGCGTCATCGCTATCGGTGGTACCTTGCGCCACGCTGGTGGGAGCGCTGTGTTCGACATTGTTCTGCCCGTAATCAGC
GTTCCACTGAATGACCAGCAGATAGGCGAGCACCGCGAGCGGGATGACGAGTAAGAGTCGTTTTACGTCCATAAGGATTCTGCCCGGTGAAACATTCTGA
CGACACATCCCCACCGGCGGAGCGCACACGGCGGGCGTGGCGGGAAAGGCCGATCAAGGGGATGGGGTGGACGCGACCGACTGCTTGCGAACGTCTTTTT
CCAGGCGCCGCCACATGCCGTGCAGTTGGCGATGCAGGGTAGCATTGTCCAGTGCATCAACTCCGCGACGAGCGAGAACGATGATGTCCACGGCGGGGAG
ACGATGTGACTGGAGGCGAAAGGATTCACGGACGAGGCGTTTCAGACGGTTACGGTCGACGGCACGGCGCACGTTCTTCTTGCTGAATACGAATCCCACC
CGCGGGTAGCCGAGAGCGTTGGGGCGCGCAAGCACCAGAAGGCCCTTGCCGTGCACCTTGAAGGACGCATGAGCGAAGACGTGCTGATAGTCCCCGGCGG
TAAGCAGACGCTGTCGCCGGGGAAAGCCCTGATCGGACATTCGTTGCTCAATCAGGCGCTCAGACGCTTGCGGCCCTTGGCGCGACGGCGAGCCAGCACC
TGGCGGCCGTTCTTGGTGGCCATGCGAGCACGGAAACCGTGCACGCGCTTGCGCTTTAGAACGCTCGGCTGAAAAGTGCGTTTCATAGCTGCGAATTCTC
ACGTGGTTGTTTGGAAACGGTTGAAACTGAAGAGCCCGGCACGTTGGCACCAGGCGGTATTCGGCTCAAGACCGGCAATTCTAGTGAATCCCCAGGGCAG

GTGCAATTTTTCTGGCGCTTTTATCCACACCTCTTTTTGCGACGGCCCTCGCCGGGGGAGCGCGATCCGTCTCGAGACTCGAAGGTATGGGGACATCATC
ATGTGCTGAAGAGAAGGGGGCTTTCCACCGTCCAACAGCCACTTCCATTACTACAGGTTTGTTTTCTACTGATGTTGTTATTAATGGTGGGGACAAATTT
**TGTGGATAA**CCCAAAAAAAGATCAGAATGTCATGTAGTTGGCGTGTTCCTCTTTGTGTGGGGAAGTTGCGGGTGTCCCGTGGGAAAGCTGTCGATGTCGT
TGTGGGTAACACGTCATCTGTCCACAGGCATGCTGGCGTCCCCGATGTGTGGACAATCTGTCCCCGTGGTTGGCGGCCCGGTTTTCCACAGGAAAAGAGA
GCGGGGATCCATGTGGATAACCTCGGTCTCGAACGCTACAATACTCGGCCATTCATCAACGCTAGGTGAGGTCGTGTGTCGGTCGCTCTCTGGCAGCAAT

GCCTCAACTTTCTTCAGGACGAACTGAGTTCACAGCAGTTCAACACCTGGATTCGCCCACTGCAGGCGGAAGACGGTGACGCCAACGAGCTGTGTCTTCT
GGCGCCCAACCGCTTCGTGCGTGACTGGGTCAACGATAAATACCTGAAGCGCATCAACGAGCTGCTGCGGGACTTGGCATCCGGTAAGCCGCCCAAGGTG
CAGCTGACGGTGGGGAGTCGCCGCAACGTGGCGATGTCGTCGCCGCGTGACCTGGGCGCGCCGGTGTCGGCGACGACGATGAATGCCTCGCGCCCGACCG
AGGCGCCGGCGGTGCATGCCGCTCCGCGTGCCAAGGGGGACTACGCCGACGAGCAGGAAATCGATCGCCTTCGCGAACGCGAGGAAACGCCGCGGCGCGG
TGGTGGCGAGCGCCAGGTCCAGGTGGAGGGGAGTCTCAAGCACCAGAGCGGTCTCAACCCCAATTTCACCTTCGAGACATTCGTCGAGGGCAAGTCCAAC
CAGCTTGCGCGTGCCGCGTCGCGGCAGGTCGCCGAGAACCCCGGTGGTGCCTACAATCCCTTGTTCCTGTACGGTGGGGTCGGTTTGGGGAAGACGCACC
TGATGCATGCCGTCGGCAACCATCTGGCGGGGCAGCGCGAGAACGCCAAGGTCGTCTACCTGCATTCGGAGCGTTTCGTCGCCGACATGGTCAAGGCGCT
ACAGCTCAACGCCATCAACGACTTCAAGCGTTTCTATCGCAGCGTGGATGCCCTGCTGATCGATGACATTCAGTTCTTCGCCGGCAAGGAGCGTTCGCAG
GAAGAATTCTTCCACACGTTCAATGCGCTGCTCGAGGGCGGCCAGCAGATGATCCTGACCTCGGACCGTTATCCCAAGGAAATCAGCGGGGTGGAGGAAC
GTCTCAAGTCGCGCTTCGGCTGGGGACTGACGGTCGCCATCGAGCCGCCGGAGCTGGAAACGCGCGTGGCGATCCTGATGAAGAAGGCCGACCAGGCCAA

>Chromohalobacter salexigens DSM 3043 [NC_007963.1:3672645..3673144], <mnmG oriC2 <COG1126 (glnQ), 602 bp

GAACTGCACGTCGAAGGGCAACGAGGTCGAGATCCCGTTGGGATAGAGCTCGTGGGTATCCAGCCCCTCCGGCTCGACGAAGATCTGATGACTCGACTTG
TCGGCGAAGCGGTTGACCTTGTCCTCGATGGACGGGCAATAGCGCGGGCCCACACCGCCGATCACGCCGGAATACATCGGCGAGCGATCGAGGTTGGCAT
GGATGATCTCGTGTGTGCGCGCATTGGTGTGAGCGATGTGGCACGCGACCTGACGCGGATGCTGATCCCGACGGCCCAGATACGACATCACCGGGGTCGG
GTCGTCGCCGGGCTGGGTCTCGAGCACCGAGAAATCCACGCTCTTGGCGTCCAGACGCGGGGGAGTCCCGGTCTTGAGCCGGTCGACGTGAAACGGCAGG
GCACGCAGCCGCTGCGCAAGGGCATTGGAGGGCGGATCACCGGCACGCCCCCCGGCATGATGATCTAACCCGATGTGGATAACGCCACCCAGGAAGGTAC
CGGTGCACAGCACAACGCTCTCGCCGAACACGCGAATGCCGGTCTGCGTCACGACTCCGCGGACCGTGTCGTTTTCCACGATCAGGTCGTCCGCGGCCTG
CTGAAACAACGTCAGGTTGGGCTGGTTTTCCAGCATGCCGCGAATCGCTGCCTTGTAGCGCACCCGGTCGGCCTGGGCCCGCGTGGCGCGCACCGCCGGC
CCCTTGCGGGCGTTGAGCACCCGGAACTGGATGCCACCCAGATCGGTGGCATGCGCCATGGCGCCGCCCAGGGCATCGATTTCCTTGACCAGATGACTCT
TGCCGATCCCGCCAATGGCCGGATTGCAGGACATCTGCCCGAGCGTCTCGATGTTATGGGTCAGCAGTAACGTTTGACAGCCCATGCGAGCGGCAGCCAA
TGCGGCTTCTGTCCCCGCATGACCACCGCCGATGACGATCACATCAAAGCGGTCTGGGTAATCCACGTTGCACCTCGAATCGCAGCGTCTGGCTGCGGTC

GTACATGTCGACAGGCGCCATCGAGCGAGGACGGCGCCTGGGAAATAGCCGGACAGTATAAACCCCTTGTGAGTAACCGTCAGGGTTTTCCACACATGGA
TTGTCCGCCGGTCGTGTCTTCGTCCTTGGGTCTATAAATAATAATAAGTATGTAGAAAAGAAGTTCTGTTGCTGTTGTAACTACTGGTGTGGACCAAATC
**TGTGGATAA**GCGTTAAAAACCTTTCATTCACAATTGCTTGCGATGTTAGCCAAACGTTGTAGAAGCGGCAGACAGGCTGCGTGCAGGAGGGGGACGGCCT
GTGCATGAAAATGCCGGTTGTCCACACCGCGAAGCGTGCACCACTTATCCACGGGTGTATACCTGATCGCCCCCCGTAGCCGTCAACAGTCAGGCACGCG
TCTCTGGAACCCCGTAGTGACGGGGGCTGGAGGCGATATTTCATGGGCAGGAAAAAAATATTGTCCACAGGCGTGAAATCGGCCTGTGAAGCGCCGCGAC

AACGCACGGCAACACCACGATCCGCGGGGAGATCGTGGTGTCAGTCGAGGGGGAGAGTCAGGAAAAGGTCAGTGCTTGAGCACGCGGGAGAGGAAGGCCT
GGGTGCGCTCGTGCTGGGGATGGTCGAAGACCTGGCTGGGCGAGCCCTGTTCGACGATGCGTCCGGCGTGGACGAAGATGACGCGATCGGCGACCTCGCG
GGCGAAGCCCATCTCGTGGGTGACGATCATCATGGTCATGCCTTCGCTGGCCAGTTCGCGCATGGCATCGAGTACCTCGCCGATCATTTCTGGATCGAGT
GCCGAGGTGGGCTCATCGAACAGCATCAGGCGCGGTTCCATCGCCAGGGCGCGGGCCAGGGCGACACGCTGTTGTTGGCCCCCTGAGAGCTGCCCGGGGT
ACTTGTCGGCCTGATCGCTGATGCCGACCCGTTCGAGAAGGCGGTCCGCGATTTTGGATGCCTGGGCATCGTCCAGACCGCGGACCCGGGTGGGGGCCAG
CGTGACGTTGGCCCGTACCGACAGATGCGGAAACAGGTTGAACTGCTGGAAGACCATGCCGACCTCGGTCCGGATGGTCTGCAGCGATTTGCCCCCCTTG
CCGTACGGTGTGAGTTCGTTGCCATCGACGATGATACGGCCGGACTGGAACTCTTCCAGGCCGTTGACGCAGCGGATGAACGTGGATTTCCCGGAGCCGC
TGGCGCCGATGATGACGACCACCTCGCCGGGGGTGATCTCGAGATCGATGTCCTGCATCACGTGCAGCTTGCCGAAGTATTTGTTGACCTTCTCCAACTG
CACGATCGGCGCGGCGGAAGCTTGTGCGTTATCACTCATGATGGGCGCCTCATTGTCCGACCAGACCCTTGCGTTCGAGCAGACGCAAGGCGGTAGAGAG
ACTCAGGGTGATCACCAGATACAGCATGGCGACCATGAAATAGACTTCCAGGGCGGTGAACGTGGTGGCGATGTAGACCTGTCCCTGGCGCACCAGCTCA

>Chromohalobacter salexigens DSM 3043 [NC_007963.1:1581..1629], dnaA> ig dnaN>, 49 bp

WebSkew GCmin pos. 412111 (GCmin<>oriC1 25.3 kb; GCmin<>oriC2 1.8 kb; oriC2<>oriC1 23.4 kb)

>Chromohalobacter marismortui DSM 6770 [NZ_SOBR01000004.1:c386588..387087], <rpmH oriC1 dnaA>, 589 bp

CAAGAACATCGGTCCGCACCTGAACCATATGATCGGAACTGGTCTGTTCCTGCTTGCCCGGCATACTGGAAGCGCCTTGGGATGCGCCCCCGGCACTACT
GGGTGCCGAGAGTCCCTCTTCGGCGGCATCATTGCTGGCGCCGTTCTGGGAAACGCTGGCGGGGGGGCGTTGTTCGACGCCGTTTTGCCCATAATCCGCG
TTCCACTGAATGACCAGTAGATAGGCAAGTACCGCGAGCGGGATGACGAGTAGGAGTCGTTTTACGTCCATAAAGATTCTGCCCGGTGATACATTCTGAC
GTCACGTCCCGATCGGCATACGCCCACGGTGGAACGTGGCGGAGAAATGCCGATCAAGGGGATGGCGTGGACGGGACCGACTGCTTGCGTACGTCTTTTT
CCAAGCGTCGCCACATGCCATGCAGCTGGCGATGCAAGGTATCGTTATCCAGCTCATGAGCCCCGCGCCGAGCGAGAACGATGATATCCACGGCGGGGAG
ACGCTGCTGCTGAAGACGAAAGGATTCACGGACGAGGCGTTTCAGACGGTTACGGTCGACGGCACGACGCACGTTCTTCTTGCTGAAGACGAGTCCCACT
CGCGGCGAGCCGAGAGCATTGGGGCGCGCCAGCATCAAAAGGCCTTTGCCGTGCACCTTGAAGGACGCATGAGCGAAGACGTGCTGATAGTCCCCGGCGG
TCAGCAGTCGTAACCGCCGGGAAAAGCCCTGATCGGACATTCGTTGCCCAGTCAGGCGCTCAGACGCTTGCGACCTTTGGCACGACGACGCGCCAGCACC
TGGCGGCCGTTCTGGGTGGCCATGCGAGCACGGAATCCGTGCGCGCGTTTGCGCTTTAGGACGCTCGGTTGAAAAGTGCGCTTCATAGCTGCGAATTCTC
ACGTGGTTGTTTGGAAACGAGTTGAAACTGAAGAACCCGGTACGCTTGCACCAGGCGGTATTCGGCTCAAGACCGGCAATTCTAGTGAATCCGTGGGGCG

GGTGCAATTTTTCCGACGCTTTTATCCACACCTGTTTTGTTACGGCGGCGGGGACTTTGGCATGTACCGCGGCAGTCGACCGGCATGGATGACGTTTTGG
ATGCTTGTCGCTCGTCGTCGCCCTTGGGGAACGCGATACCGCTACTATCATTATTGAAATTATATTGATTGTTGTTGTTATTAATGGTGTGGACATTGTT
**TGTGGATAA**GTAGTTTTTGTTCTTTTTTTACAAACGCACAGCATGTTATTGAATATGTTGGCAACAGGTGAAGGAGCAAGGGGTAAGCTGTCGATAGTGG
TGTGGACAATGGGTGTTTTGTCCACAGCCGACAAGCCTTCCACGTTCCACGGACATGATGTCCGTTGAGTTATGGAGAGAGTTTTCCACAGGAAAAGAGA
CATGCCATCCATGTGGATAACCCGAGCCTCGAGCGCTACAATAGCCGGCCATTCATCAACGCTAGGTGAGGTCGTGTGTCGGTCGCTCTCTGGCAGCAAT

GCCTCAACTTTCTTCAGGATGAGCTGAGCTCGCAGCAGTTCAACACCTGGATTCGCCCCTTGCAGGCGGAAAACGGTGACGCCAACGAGTTGTGTCTGTT
GGCGCCCAATCGCTTCGTGCGTGACTGGGTCAACGATAAATACCTGAAGCGTATCAATGAATTGCTGAGGGAGCTTGCGTCCGGAAAGCCGCCCAAGGTG
CAGCTGACGGTGGGTAGTCGCCGCAACATGGCGGTTTCATCACCGCGGGATCTTGGTGCGCCGGTCTCGGCGACGGCGATGAATGCCTCGCGCCCAACCG
AGGCGCCGTCGGTACATGCCGCTCCGCGTGCCAAGGGAGACTATGCCGACGAGCAGGAAATCGACCGCCTTCGCGAGCGCGAGGAAACGCCGCGGCGTGG
TGGTGGCGAGCGCCAAGTCCAGGTCGAAGGCAGTCTCAAGCACCAGAGCGGGCTCAATCCCAACTTTACGTTTGAAACCTTCGTCGAGGGCAAGTCGAAC
CAGCTGGCGCGTGCCGCCTCGCGCCAGGTGGCCGAGAATCCCGGTGGTGCCTACAATCCTTTGTTCCTGTACGGTGGGGTCGGCTTGGGTAAGACGCACC
TCATGCATGCGGTGGGCAACCATCTGGCGGGTCAGCGGGAAAACGCCAAGGTGGTGTATCTGCACTCTGAGCGTTTCGTGGCCGATATGGTCAAGGCGCT
ACAGCTCAACGCCATCAATGACTTCAAGCGTTTCTACCGCAGCGTGGATGCCCTGCTGATCGACGATATCCAGTTCTTCGCCGGCAAGGAGCGTTCGCAG
GAAGAGTTCTTCCATACGTTCAATGCGTTGCTCGAGGGCGGTCAGCAGATGATCCTTACCTCGGATCGTTACCCCAAGGAAATCAGTGGGGTGGAGGAGC
GCCTCAAGTCGCGCTTCGGCTGGGGGCTGACGGTGGCCATCGAGCCGCCCGAGCTGGAAACCCGCGTGGCGATTCTGATGAAGAAGGCCGACCAGGCCAA

>Chromohalobacter marismortui DSM 6770 [NZ_SOBR01000004.1:c410041..410540], <mnmG oriC2 <COG1126 (glnQ), 614 bp

CTGGAACTGCACGTCGAAGGGCAGCGACGTCGAGATCCCATTGGGATAGAGCTCGTGGGTATCGAGGCCTTCGGGCTCGACGAAGATCTGGTGACTCGAT
TTGTCGGCGAAGCGATTCACCTTGTCTTCGATGGACGGGCAGTAGCGCGGTCCAACACCACCGATCACGCCGGAGTACATCGGCGAACGATCGAGGTTAG
CGTGGATGATTTCATGAGTACGCGCGTTGGTATGCGCAATATGGCAATTCATCTGACGGGGATGTTGCTCGCGATTGCCTAGATACGACATGACCGGTGT
CGGCGTGTCGCCAGGCTGCGTTTCGAGCACCGAGAAATCCACGCTTTTGGCATCCAGGCGCGGCGGTGTCCCGGTCTTGAGCCGATCGACGTTGAACGGC
AGGGCACGCAACCGCTGCGCCAGTGCATTGGAAGGCGGATCCCCGGCACGCCCGCCGGAGTGATTGTCCAACCCGATGTGGATAACGCCACCGAGGAAGG
TACCGGTACACAGCACCACGCTTTCGCCGTAGACGCGGATCCCGGTCTGCGTCACGACGCCTCGCACCGTGTCGCCATCGACGATCAGGTCGTCGGCGGC
TTGCTGGAAGAGCGTCAAATGGGACTGGTTTTCCAGCTGGCTGCGAATCGCGGCCTTGTAACGGACGCGGTCGGCTTGGGCCCGTGTGGCGCGTACGGCG
GGGCCCTTGCGGGCATTGAGCACACGGAACTGTATGCCACCGAGATCGGTGGCATGCGCCATGGCGCCGCCGAGTGCATCGATTTCCTTGACCAGATGAC
TCTTGCCGATCCCGCCGATGGCGGGATTGCAGGACATCTGCCCGAGCGTCTCGATGTTATGCGTCAGCAATAGCGTCTGACAGCCCATGCGGGCGGCAGC
CAATGCGGCTTCGGTTCCCGCATGGCCCCCGCCGATGACAATGACATCAAAGCGGTCTGGGTAATCCACGTTACACCTCGAATCGCAGCATCTCGCTGCG

GTCGTGCATGTCGGCGAGCTCAATGCATTGCAACGAGCTCGGCAAATAGCCGGACAGTATAAACCCCTTGTGAGTAACCGTCAGGGTTTTCCACACATGG
ATTGCCAGTGAGTGGTCCTTCCCCGGGGCTTTCTATTAATAACAATATAGAATTTAAAAGAAGTTCTGTTGTTGTTGTTATTACTGGTCTCGGCAAAATC
**TGGGGATAA**GTTAAAAATATACATTTTTTTCAGATGGTTATGGTGTTTGCCCTTGCTCGGAAAAAATGCCGACAGCCTGCGTGGGAACGGGGGACGTCCT
GTGGATGGAAGTGGTGATTTTCCACAGGGCGGATCGTGCACCACTTATCCCCAGCCTCATACCCGAGCCCCGCCCATAGTGATCAACATCCAGCAACATT
GCCTCTAAACCCGCGCATGACGGGGCTTGCGGCGAGAATAAGCGAGTCGTGGCATCCATGTTATACACAGGCAAAAAATGGCCCTGTGGAGACGATAGGG

GGGAGGCAGCGATTCGGAAATCGCTGCTTTATCACCAGGGGGAAGCGTTACTTGACGCTTGGCAATGCGCTTACTCGAGAACGTTAGTGCTTGAGTACGC
GGGAGAGGAAGGATTGGGTACGTTCGTGTTGAGGGTGGTCGAAGACGTCCTGTGGGGTTCCTTCTTCGACGATGCGACCGCCATGCACGAAGATGACGCG
ATCGGCGACCTCTCGAGCGAAGCCCATTTCGTGGGTCACGATCATCATCGTCATGCCCTCATTGGCCAGTTCGCGCATGGCGTCGAGCACTTCGCCGATC
ATTTCCGGATCGAGTGCCGAGGTGGGTTCGTCGAAGAGCATCAGGCGCGGTTCCATGGCCAATGCGCGTGCCAGGGCCACACGCTGCTGCTGCCCGCCGG
AGAGTTGGCCGGGGTACTTGTCGGCTTGATCGGTGATTCCCACACGCTCGAGCAGTCGATCGGCGATTTGGCGTGCCTTGCCGCTATCAAGCCCGCGTAC
CTTTTCAGGGGCCAGGGTCACGTTGGCATGCACCGACAAGTGAGGGAATAGATTGAATTGCTGGAAGACCATGCCGACTTCGGTGCGAATGGTCTGCAAC
GATTTCCCGCCCTTGCCGTGCGGTGTGAGTTCGTTGCCATCGACGACGATGCGTCCCGATTGAAACTCCTCGAGGCCGTTGACACAGCGAATGAGCGTCG
ATTTGCCGGAGCCGCTGGCGCCGACGATGACGACCACCTCGCCGGCGGTTATCTCAAGATCGATATCCTGCATGACATGCAGGCTGCCGAAATATTTGTT
GACCTTCTCCATATGGACGATGGACTGAGCCGTTGCCGGGGACGTTTCATGTTGGGGGTTGTCATTCATGGCATTGGCTCCTCATTGACCGACCAGACCA
CGGCGCTCGACCATGCGCAGGGCTAAAGAAAGACTTAAGGTGATTACCAGATAGAGCATGGCCACCATGAAATATACCTCGAGTGCGGTGAAGGTCGTGG

CGATATAGATTTGTCCTTGGCGAACCAATTCACCCACGCCGATCACCGAGAACAACGAGGTATCCTTGATACTGACGATACCTTGATTGCCAAGTGCAGG
AATCATCCGGCGCAAGGCTTGTGGCCAAATGATGTAGCGGAAGGTGTCGACTCTCGATAAGCCCAATGACAGGCCCGCTTCGCGTTGTCCTCGGTCGATG
GATTGCACCCCGCCACGCACGATTTCCGAGATATAAGCCCCCGCGTTCAAGGCGATGGCCGCGATACCTGCGGTGACGGCGTCGATGGGGCCGCCGAGGA
TTTGTGGCAAGCCGTAGAATATGAACAGCACCTGGACGAGGACCGGTGTACCACGGAATATTTCGATATAGGTGACGGCCGGCCACTTCAAAAGTCTTAA
CGGACTAATGCGCAACAGGCCAAAGAAAATACCGATTAAAAAGCCGATAGCAAGACCGCCGAACGAAATCAGTAGGGTCCAGGGGATACCTTCGAGCAAG
TAGGGTATGGAGGCGATAGCTGCCTGCCAATCGAACTGGAAAGTGACGTCCACGTCGGAATCTCCGGCGTGGCTCGCCAGCGTCCGGCGAGCCACGTGT

>Chromohalobacter marismortui DSM 6770 [NZ_SOBR01000004.1:c385076..385124], dnaA> ig dnaN>, 48 bp

WebSkew GCmin pos. 250500 (GCmin<>oriC1 26.3 kb; GCmin<>oriC2 2.7 kb; oriC2<>oriC1 23.6 kb)

>Chromohalobacter canadensis 85B [NZ_JAJQJH010000002.1:276560..277059], <rpmH oriC1 dnaA>, 590 bp

CGTACCTCGATCAAGCCGTCCGAGTCGGTTTGCCCGTCCGCACCAGGCATGTCGGCGGCCTCTTGCGATGAGGCACTGTTACCGCTGCTGTTGGACGTGG
GAACCGAGAGGCTGTCGTTGCCGGCGTTACTATCGGACGTATCCTGGGTAACGCTTTCAGGTGCGCGTTGTTCGACGCTGTTTTGCCCATAATCCGCGTT
CCACTGAATGACCAGTAGATAGGCGAGTACCGCGAGCGGGATGACAAGTAGGAGTCGTTTTACGTCCATAAGGATTCTGCCCGGTGATACATTCTGACGT
CACGTCCCGACCGGCACTGCGGCCACGGTGGAACGTGGCGAAGAAATGCCGATCAAGGAGATGGCGTGGACGAGACCGACTGCTTGCGTACGTCTTTTTC
CAGGCGTCGCCACATGCCGTGCAATTGGCGATGCAAGGTATCGTTATCCAGCTCATGAGCTCCACGCCGAGCGAGAAAGATGATATCCACGGCGGGTAGA
TGATGCTGCTGGAGACGAAAGGATTCACGGACGAGGCGTTTCAGACGGTTACGGTCGACAGCACGACGTACGTTCTTCTTGCTAAACACGAGCCCCACTC
GCGGTTCACCGAGAGTATTGGGACGCGCAAGCATCAAAAGGCCCTTGCCGTGCACCTTGAAGGCCGCGTGAGCGAAGACGTGCTGATAGTCCCCGGCAGT
TAGCAGTCGCCGCCGCCGGGGAAAGCCCTGATCGGACATTCGTTGTCCAGTCAGGCGCTCAGACGCTTGCGACCTTTGGCACGACGACGCGCCAGCACCT
GGCGGCCGTTCTGGGTGGCCATGCGAGCACGGAAGCCGTGCACGCGCTTGCGCTTTAGGACGCTCGGTTGAAAAGTGCGCTTCATAGCTGCGAATTCTCA
CGTGGTTGTTTGGAAACGATTTGAAACTGAAGAGCCTGGTACGCTCGCACCAGGCGGTATTCGGCTCAAGACCGGCAATTCTAGTGAATCCCTGGGGCAG

GTGCAATTTTTCCGGCGTTTTTATCCACACCTGTTTTGTTACGGCGCCGAGGGTTCGGCATGTGCCGAGGCGGTCCGCCCAAGCGAATACTCGTTTGGAC
GCTCACCGCTTGTCGTCCCGTCCCTGAGTGAAACGATAAAGCAATTATCATTATTGAAGTTATACTAACTGATGTTGTTATTAATAGTGGGGATATTGTT
**TGTGGATAA**CCAATTTTATACCTTTATTAACAAAGAGATGCCATGTTCTTGAATATGTGGAGAAAAGGCGTGAAAGCGGTGGGTAAACTGTCGGTAGCAT
TGTGGGTAAGTGGCGATTTCTCCACACCCTGAGACATCTCCACGTTGAGAGGACACGTTGTGCGCTGAGTTATGGATAGGGTTTTCCACAGGAAAAGAGA
CTTGGTGCCCATGTGGATAACCCGGGCGCCGAACGATACAATAGTCGGCCATTCATCAACGCTAGGTGAGGTCGTGTGTCGGTCGCTCTCTGGCAGCAAT

GCCTCAACTTTCTTCAGGATGAGCTGAGCTCTCAGCAGTTCAACACCTGGATTCGCCCCTTGCAGGCGGAGGACGGTGACGCCAACGAGCTGTGTCTGCT
GGCGCCCAATCGCTTCGTGCGTGACTGGGTCAATGATAAATACCTGAAGCGTATCAATGAGTTGCTGAGGGAGCTTGCATCCAGCAAGCCGCCCAAGGTG
CAGCTGACGGTTGGCAGCCGTCGCAACGTGGCCATGTCGTCGCCGCGTGATCTAGGGTCACCTGTATCGGCGGCGGCAATGAATACCTCGCGCCCCCCCG
AGACACCGCCCGCCGTACATGCACCGCGAGCTCGGGGCGATTATGCGGATGAGCAGGAAATCGATCGCATGCGTGAGCGTGAGGATTCCTCGCGCCGGAG
TAGTGGCGAGCGTCAGGTGCAGGTGGAAGGGAGTCTCAAGCACCAGAGTGGACTCAATCCCAATTTTACCTTCGAGACCTTCGTCGAGGGTAAATCGAAC
CAGCTCGCGCGTGCTGCTTCGCGCCAGGTTTCCGAAAACCCCGGTGGTGCCTATAACCCGCTGTTCCTGTACGGTGGTGTCGGCTTGGGTAAGACGCACC
TCATGCATGCAGTAGGCAATCATCTGGCTGGCCAGCGGGAAAACGCCAAGGTGGTTTACCTGCATTCAGAGCGTTTCGTGGCTGACATGGTCAAGGCACT
GCAGCTCAACGCCATCAACGATTTCAAGCGTTTCTACCGCAGCGTGGATGCACTGCTGATCGACGACATCCAGTTTTTCGCCGGCAAGGAGCGCTCGCAG
GAAGAATTTTTCCATACGTTCAATGCGTTGCTTGAGGGCGGCCAGCAGATGATCCTTACCTCGGATCGCTACCCCAAGGAAATCAGTGGGGTGGAAGAAC
GCCTCAAATCGCGCTTCGGCTGGGGGCTCACGGTGGCCATCGAGCCGCCCGAGCTGGAAACCCGCGTGGCGATCCTGATGAAAAAGGCTGACCAAGCCAA

>Chromohalobacter canadensis 85B [NZ_JAJQJH010000002.1:252995..253494], <mnmG oriC2 <COG1126 (glnQ), 605 bp

GGAACTGCACGTCGAAGGGCAGCGAGGTTGAAATCCCGTTGGGATAGAGCTCGTGAGTATCGAGGCCTTCGGGCTCAACGAAGATTTGGTGACTCGACTT
GTCCGCGAAGCGATTCACCTTGTCTTCGATGGACGGGCAGTAGCGCGGCCCGACACCCCCGATGACACCAGAATACATCGGAGAGCGTTCGAGATTGGCG
TGAATGATCTCGTGGGTACGGGTGTTGGTATGCGCAATGTGACAGTTCATCTGACGTGGATGCTGCTCGCGGTTGCCGAGATATGACATCACCGGGGTCG
GCGTATCGCCAGGTTGGGTCTCGAGTACCGAGAAATCCACGCTTTTGGCATCCAGTCGCGGCGGCGTACCGGTCTTGAGCCGATCGACGTTGAACGGCAG
GGCACGCAGGCGTTGCGCCAAGGCATTGGAGGACGGATCCCCGGCACGCCCGCCGGAGTGATTGTCCAACCCGATGTGGATAACGCCGCCGAGGAAGGTC
CCGGTGCACAGCACCACGCTCTCGCCATAGACCCGGATACCCGTCTGTGTGACGACGCCGCGCACCGTGTTGCCGTCGACGATCAGGTCGTCGGCAGCCT
GTTGGAAGAGCGTCAAATGAGACTGGTTTTCCAGTTGGCTGCGGATCGCGGCTTTGTAACGGACGCGATCGGCTTGGGCCCGAGTGGCACGCACGGCGGG
CCCCTTGCGGGCATTGAGTACACGGAACTGAATGCCGCCGAGATCGGTGGCATGCGCCATGGCGCCGCCTAAGGCATCGATTTCCTTGACCAGGTGGCTC
TTGCCGATCCCGCCGATGGCGGGATTGCAGGACATCTGGCCAAGAGTCTCGATGTTATGCGTCAGCAATAGCGTCTGACAGCCCATGCGAGCAGCAGCCA
GTGCGGCTTCGGTTCCCGCATGGCCCCCGCCGATGACGATGACATCAAAGCGGTCTGGATAATCCACGTTACACCTCGAATCGCAGCGTATGGCTGCGGT

CGTGCATGTCGGCAAGCCTCATCGTCAGGCGATGAAGCTTGGTAAATAGCCGGACAGTATAAACCCCTTGTGAGTAACCGTCAGGGTTTTCCACACATGG
ATTGCCAGTGGGTTGGCCCGTCATAGGGTCTCTTATTAATAACAATATAGAATTTAAAAGAAGTTCTGTTGTTGTTGTTATTACTGGTATCGACTAAATC
**TGGGGATAA**GTGTAAAAAAACAAGTTTCATCAGGGTTTTACACTGTTCATCGCTGATAGGAAAAAGCGCAAATAGCCTGCGCGGGAGCGGGGGATGGCCT
GTGGATGGAAGGGGGGCTTATCCACACGCCAGATCGTGCACCACTTGTCCCCAGGCGTATACCAGGGCCCCGCCCATAGTGGTTAACATTTGGCGGTTTT
GCCTCTTAACCCCCGCATGACGGGGCTGGTAGCGAGAATGAGCGGTTCGTGACATCAATGTTATGCACAGGCAAAAAATTGCCTTGTGAAGACAGAAATG

AAGCGCGGCGACCCTTGGAGGTCACCGCGTGATCGTGAGGTGGGAGCGCTACCTGGTGTTTGAGAACAAAAGCTAGTGCTTGAGAACGCGGGAGAGGAAA
GCTTGGGTGCGTTCATGTTGGGGACGATCGAAAACTTCTTGTGGCGTGCCTTGTTCGACGATGCGGCCACCGTGCACGAAGATGACGCGGTCGGCGACCT
CGCGTGCAAAACCCATCTCGTGGGTCACGATCATCATCGTCATGCCTTCATTGGCCAGTTCCCGCATGGCGTCGAGTACTTCGCCGATCATTTCCGGGTC
GAGAGCCGAGGTGGGCTCGTCGAACAGCATCAGGCGGGGTTCCATCGCCAATGCGCGGGCCAAGGCGACTCGCTGCTGCTGGCCCCCGGAGAGCTGGCTG
GGATACTTGTCGGCTTGATCGGTGATACCCACGCGTTCGAGCAGTCGATCGGCGATTTCCCGGGCTTTGGTGTTATTGAGCCCACGTACTTTCTCCGGGG
CCAGGGTCACGTTGGCATGCACCGACAAATGGGGAAACAGATTGAATTGTTGAAAAACCATGCCGACCTCGGTGCGGATGGTCTGCAACGATTTTCCGCC
CTTGCCGTAAGGTGTGAGCTCGTTGCCATCGACGACGATGCGTCCTGACTGAAACTCCTCGAGGCCGTTGACGCAGCGAATGAGCGTCGACTTGCCGGAA
CCGCTGGCACCGACGATGACGACCACCTCGCCAGCGGCGATCTCGAGATCGATATCCTGCATCACGTGCAGGCTACCGAAATACTTGTTGACCTTCTCCA
GCGTGACGATCGGAGTGGTGGAATCTTGCTGACTGTCATTCATGATAGGCGCCTCATTGTCCGACCAGGCCCTTGCGCTCGAGCAGGCGCAAGGCGAAAG
AGAGACTCAGGGTGATCGCCAGATAGAGCAAGGCGACCATCAAGTAGACCTCGAGCGCGGTGAAGGTGGTCGCGATATAGACCTGGCCCTGACGCACAAG

>Chromohalobacter canadensis 85B [NZ_JAJQJH010000002.1:278523..278571], dnaA> ig dnaN>, 49 bp

WebSkew GCmin pos. 1207808 (GCmin<>oriC1 36.2 kb; GCmin<>oriC2 2.5 kb; oriC2<>oriC1 38.7 kb)

>Kushneria konosiri X49 [NZ_CP021323.1:c1171334..1171833], <rpmH ig dnaA>, 332 bp

CGTTGACGGTTGCCCGCATGTCGACCTGAAGCGTCTTTTGATCATCGCTCAGCTGATAGCGGGTTTTTTCGGGGGTAAAGGTAATTCGCCCCTTGTGGCC
TTCCATCTGCAGCCCGGAGCGTGCGATGTAGCTGCGGCGCTGATCATCTGAAAGCAGAACGAAGGGTGCATCGCTATCCAGGCGCTGTAGATGCTTGGGC
AGGGCGGCATAGACGATATCGCCACCGCGCGGGTCGATGCGCAGGTCAAGGGCATCGGTGCGTACACTGATCAGGCTGGTGTTTCCGGTATCGGCGGCGC
TGTCGGTGCTCTCGGTAGTGTTGCCGGGCTGCGGGGTACTGGCGCGTTCACTGGCCGTGTCGGGGGCGTCGAGCGACTGCTCACCGGGTTCAACGTCACT
GCTTTGCTGGGACAGGGCGGGAGGCGCTTCATCCGGCAGGTCGTTTTGCCCGTGATCATTGCTCCATTGAATGATCATCAGATAGGCAAGCACGGCCAGC
GGGATGACAAAAAGAAGTCGTTTCAGATCCATTGGGATTGCCCGGTGGGAAAATGTCGACGCGGCGCAAGAAAACGACGGGTCGCACTCGACGGTTCGTC
GCGCCTCGTATTATTGCTGATGTTCCGGCTGGCGCTTGACGTCCTTTTCAAGACGTCGCCACATACCGTGAAGCTGTCTGGCCAGCGTTTCCGAGTCGAT
CTCGTGAACGCCCCTGCGTGACAGCACAACGATATCCACGGCTGGCAGCCACTGCTGGCGCAGTCGCACCGATTCTCGAGTCAAGCGCTTGAGACGGTTG
CGATCCACGGCTCGGCGTACGTTCTTCTTGCTGAATACCAGCCCCACCCGAACGTGACCGAGATCGTTCGGTGTGGCGAGTGCGAGAAGGCCCTTGCCGT
GAATTCTAAAACTGGCATGTTCGAAAACATGACGGTATTCCCCGGCGGTCAGAATTCTAAGACCGCGGGGAAATCCCTGATTGGTCATAAGCTGGACCGA

TCCTGTCGAGGGATCAGGCACTCAGGCGCTTGCGACCCTTGGCACGGCGACGCGAAATGACCTGACGGCCGTTCTTGGTGGCCATGCGAGCGCGAAAGCC
GTGGTTACGCTTGCGCTTGAGCACGCTGGGCTGGAAAGTACGTTTCATGGTTGCGATTCCCGAATTGAGACTTGAGTGATCGGTGACACCGATGCCCTCA
AAATCGAGGCGCACGGGGGTCAAGAGCCGGGATTCTAGAGAATTCAACGAAGACGTGCAATTTTATCCACGCAAGAAAACGATCATTTTGTGCATAACT
TTGTGCACAAGGCGATCGCATCTTTTTTTCAGGGCCCGCCGTGCGAAGCTGAGAGGCCAGAACAGCGAGCACAAGGTTACCCACAGGAGGTTTGGCGCGC
CATCACGGTGTGGATAACCTGGTGCACGGGCGTTACACTATCGCCCCATTTGCAAGACTCACCATCAATAGAGGTCGCGTGTGTCCATCGCTCTCTGGCA

GCAATGTCTTGACTATTTACAGGATGAATTGAGCTCTCAGCAGTTCAACACCTGGATCCGTCCCCTGCAGGCCGAAAGCGGTTCCAGTGACCACGAACTC
TCCCTGCTGGCCCCCAATCGCTTTGTGCGTGACTGGGTCAATGACAAGTATTTAAAGCGCATTAACGAACTGATGCGCGAACTGGCCCAGGGCCGTCCGC
CCAAGGTCTCGCTGGATATCGGTAGCCGCCGAACCGTGTCCAATCCTGCGCCGAAGCCTGCCGCCTCCGGCGAGTCAGGCAGCGCCGCGCCGCAGTCCGG
TGCCGGGCAGCCGGCACCGGAGAGCTGGTCTGCGGGCGTGACACGAACGTCCTCGGCGCCCATGCCGCCCTCGCCGCCGACTTCTCCGCGCGGCGACTAT
CAGGACGAGCGCGAAATTCCTGCGGCGCCGCCTTCGGGCAATACCACGGGCGGACGGCGCAGCGACAGCGGCGAACGCCAGGTGCAGGTCGAGGGCAGCA
TCCGCCATCAAAGCGGGCTCAACCCGGGCTTTACCTTCGAAACCTTTGTCGAAGGTAAATCCAACCAGCTGGCGCGGGCCGCGTCGCGCCAGGTGGCCGA
AAATCCCGGTGGTGCCTATAACCCGCTGTTTCTGTATGGCGGCGTGGGGCTGGGTAAAACGCATTTGATGCACGCAGTGGGCAATCACCTGGCCGGCCTT
CGTGATAACGCCCGGGTGGTCTATCTGCATTCCGAGCGCTTCGTGGCCGACATGGTCAAGGCGTTGCAGCTCAACGCCATCAACGATTTCAAGCGCTTTT
ATCGCAGCGTGGATGCGCTGTTGATCGATGACATCCAGTTCTTTGCCGGCAAGGAGCGCTCCCAGGAGGAGTTCTTCCATACCTTTAATGCCCTGCTGGA
AGGCGGTCAGCAGATGATTCTGACCTCGGATCGCTACCCCAAGGAGATCAGTGGGGTCGAGGAGCGTCTGAAATCGCGTTTTGGCTGGGGTCTGACGGTC

>Kushneria konosiri X49 [NZ_CP021323.1:c1210085..1210584], <mnmG oriC2 <PRK13623 (ErpA), 622 bp

AAGGGCAGCGACGTCGAAATGCCGTTGGGATAGAGCTCGTGAGTCTCGAGTCCTTCCGGTTCGATAAAGACCTGATGGCTCTGCTTGTCGGCAAAGCGAT
GAATCTTGTCCTCGATGGAGGGGCAGTAGCGCGGCCCGGTACTTTCGATGACGCCGGAATACATCGGCGAGCGGTCCAGATTGGCCCGAATGATGTCGTG
AGTGCGCTCATTGGTATGCGCGATGAAACAGCTCACCTGATCGGGGTGCATATCACGCCGCCCCATGTAGGACATTACCGGACGCGGCGTATCACCGGGC
TGCTCGGCCAGCTTCGAAAAATCGACCGTTCTGGCATCGAGTCGCGGTGGCGTGCCGGTCTTTAGCCGTGCCACCCGCAGGGGCTGCTCGCGAAGACGTG
CCGCCAGCGCATTGGAGGCCGGATCGCCGGATCGCCCGCCACTGTGATGTTCCATGCCGATATGGATGACGCCCCCCAGAAAGGTCCCGGTACACAGCAC
CACGGTCTCGGCCATGAAACGGATACCGCCCTGGGTCAGCACACCGCGCACGGTGTCACCTTCAACGATCAGATCGTCGGCGGCCTGAGCAAAGATCGAC
AGGTTGGGCTGGTTTTCGAGGATATGGCGAATGGCCGCCTTGTAGCGCACGCGATCAGCCTGAACGCGAGTGGCTCGAACAGCCGGTCCCTTGCGGGAGT
TGAGCGTTCGAAACTGAATGCCGCCCTGATCGGCGGCGAGCGCCATGGCACCACCAAGCGCATCGATTTCCTTGACCAGGTGGCTCTTGCCGATGCCACC
GATGGCCGGATTACAGGACATCTGTCCCAGCGTTTCGATGTTGTGACTGAGCAGCAGTGTCTGACAGCCCATGCGAGCGGCGGCCAGTGCGGCTTCGGTA
CCCGCATGACCACCACCGATGACAATAACGTCAAAGCGGTCCGGATAATCCACGTGAAAAACCTCGAGGGGTTGAAGATGAACGTGACACCCGACTGATC

AGCGCGTGTCGATACCCACATGCCCGTTCGGGGCATTGGGGATAAGCCGTATTGGATGGCGCGCATTATAACGTGGCTGTGAGTAAGCGTCAGGGTTTTC
CACACAGGGATTTTTCAAAAGGAGAAAGTCATTAATAAATAAAATAGTTTTTTAAATAAGAAGTTCTGTTGTGGTTGTGACTACTGGGGAAGCGGATTTC
**TGTGGAAAA**GCATCTTTTAGACTTTAAAAACAGAGGGCTGGGGTGTGACTGGAAACCCAAGCAAGCTGCGTGGTTCCCTGGGGCAGGTACTGTACGGGGT
GTGCATAAGTGTCCGGGTTACCCACAGCTCGATTTATCCCCTGTCCGTTAACCGGGCTCAATGAGGCTTCTCCCCATACTCATCAAGAATGACGATGTTA
TCGGGAAAAGTGATGAAAGGTACACGTGTGGCGGGCTGTGGAGCAGAAGGACACAAGAGACGTCCTGTTTTGTCCACAGGGCAGAAAGAGCAAAAAAGAT

CGGTGGAGGCGATGAGCGGACACAAAAAAGGCGTCTGTCAGAAGCTGACAGACGCCCTAGCATCCATGGGAAGCATCAGACCGTAAAGGATGCACCACAG
CCACAGGTTGTGGTGGCGTTGGGGTTTTTGATCAAAAAGCGGGCACCGGCCAGACCCGCCTCAAAGTCCACCGTGGAGCCCACCAGATACTGATAGCTCA
GCGGGTCGATCACCATGGCGACATCGCCATTTTCGATGACGGTGTCGTCTTCTCCAACGGTTTCTGCAAAATCGAAACCATACTGAAAGCCCGAGCAGCC
GCCTCCCGTTACGTAGACGCGCAGTTTAAGGGCCGGGTTGCGCTCTTCTTCGATCAGCGCCCTGACTCTGCCTGCCGCGGCATCGGTAAAGACAAGCGGT
GTCGGAACAAAGGATTCTGCTCCGCTCATCGCGGCCTCCTTTTGAGATGACACGTCCGGGACATGTCAGAAATGGATCAGCGCTAGGGCTGTCATGACAA
CCATTATCGACACCCCCTCTGTCGATGGTCAACCGCCGTGCGTCAGGGCCCGTCATAAGAGGCATGGAACATAAACAATGATAGGCTGAATAATGTGATC
CGCCATTGTGCTGCTGGGGAGGAAAAGCTTGCCGCGCCGTTTTCTGGATACGTCACGCCTGAGCCTTGAGAATTTTCGACGCGAGCTGGCAAGCGTTGAT
GCATTGCCGCAACTGTGTTTGCTGGCAGTGATCTCCGGGGCGTTGACCGGCATCATCATGATTGCCTTTCGAGAGCTGATCGATCTGGGCGCGTTGATAC
TGATGCCCAACGGTCATGGCGAGGCCTTCGAGAGCCTGCCCGTGGCGGTTCGAACAGGATTGCCGGTAGTGGCTGTTACCCTTATCGGGCTGGGGCTTGG
CTGGCAGAAAAAAAGTGCGCGGCGACTCGGCATCTCGCATGTCATTGAACGCTTCTCTTTCCATCAGGGCGTCATGGCACGTGACAACTGGCTCAATCAG

>Kushneria konosiri X49 [NZ_CP021323.1:c1169711..1169805], dnaA> ig dnaN>, 95 bp

< page 1 | #144 | Kushneria marisflavi SW32 = DSM 15357

WebSkew GCmin pos. 2079189 (GCmin<>oriC1 46.7 kb; GCmin<>oriC2 10.5 kb; oriC2<>oriC1 36.2 kb)

>Kushneria marisflavi SW32 = DSM 15357 [NZ_CP021358.1:2125617..2126116], <rpmH ig dnaA>, 332 bp

CGTTGACGGTTGCCCGCATGTCGACCTGAAGCGTCTTTTGATCATCGCCCAGCTGGTAGCGGGTTTTTTCGGGGGTGAAGGTAATCCGCCCCTTGTGGCC
TTCCATCTGCAGTCCGGAGCGCGCGATGTAGCTGCGGCGCTGGTCATCGGAAAGCAGAACGAAGGGCGCATCGCTATCAAGGCGCTGTAGATGCTTGGGC
AGGGCGGCATAGACAACATCGCCACCGCGCGGGTCGATGCGCAGGTCAAGGGCATCGGTGCGAACACTGATCAGACTGGTGTTCTGAGCGTCAGCAGCGC
TGTCGGGACTGTCAGTCGTGTTGCCGGGCTGCGGGGTGCTGGCACGCTCACTGGCCGTGTCGGGTGCATCAAGCGACTGCTCACCGGGTTCAACGTCACT
GCTTTGCTGGGACAGGGCGGGAGGCGCTTCATCCGGCAGGTCGTTTTGCCCGTGATCATTGCTCCATTGAATGATCATCAGGTAGGCAAGTACGGCCAGC
GGGATGACAAAAAGAAGTCGTTTCAGATCCATTGGGATTGCCCGGTGGGAAAATGTCGACGCGGCGCAAGAAAACGACGGGTCGCACTCGACGGTTCGTC
GCGCCTCGTATTATTGCTGATGCTCCGGCTGGCGTTTAACGTCCTTTTCAAGACGTCGCCACATGCCGTGAAGCTGTCTGGCCAGTGTTTCCGAGTCGAT
CTCGTGTACGCCCTTGCGTGACAGCACAACGATATCCACGCTCGGCAGCCACTGCTGGCGCAGTCGCACCGATTCTCGAGTCAAACGCTTGAGACGGTTG
CGATCCACGGCTCGGCGTACGTTCTTCTTGCTGAATACCAGCCCTACACGAACGTGACCTAGATCGTTCGGCGTGGCGAGTGCGAGAAGCCCCTTGCCGT
GAATTCTGAAACTGGCATGTTCGAAAACATGACGGTATTCCCCGGCGGTCAGGATTCTGAGACCGCGGGGAAATCCCTGATTGGTCATAAGCTGGACCGA

TCCTTTCGAGGGATCAGGCACTCAGGCGCTTGCGACCCTTGGCACGACGACGCGAAATGACCTGACGGCCGTTCTTGGTGGCCATACGAGCGCGAAAACC
GTGGTTACGCTTGCGCTTGAGCACGCTGGGCTGGAAAGTACGTTTCATGGTTGCGATTCCCGAATTGAGACTTGAGTGATCGGTGACACCGATGCCCTCA
AAATCGAGGCGCACGGGGGTCAAGAGCCGGGATTCTAGAGAATTCAACGAAGACGTGCAATTTTTATCCACTCATCGAAACGACCGTTCTGTGCATAACT
TTGTGCACAAGACGGTGCCATGATTTTTTCAGGGCCCGCCGTATGACCCGGAGAAGCCCGGACAGCGAGTATAAGGTTATCCACAGGGGGTTCCAGCTAC
TATCCCGGTGTGGATAACCTGATGCGCGGGCGTTACACTACTGCCCCATTTGCACGACTCACCATTAATAGAGGTCGCGTGTGTCCATCGCTCTCTGGCA

GCAATGTCTTGACTATTTACAGGATGAATTGAGCTCTCAGCAGTTCAACACCTGGATCCGTCCCCTGCAGGCCGAAAGCGGTTCCAGTGACCACGAACTC
TCCCTGCTGGCCCCCAATCGCTTTGTGCGTGACTGGGTCAATGACAAGTATTTAAAGCGTATCAACGAGCTGATGCGCGAGCTGGCTCAGGGTCGTCCGC
CGAAGGTGTCGCTGGACATCGGCAGTCGCCGCACGGTATCCAATCCCGTATCACGGTCGGCGGATGCGCCGCAGCAAAACGGTTCTGCACCACAGGGCAG
TACCGCTGCGCGCACTCAGGAGAGCTGGTCTACGGGCGTGACCCGTACGTCTTCGGCGCCCATGCCGCCTTCGCCGCCAACCTCGCCTCGTGGCGACTAT
CAGGACGAGCGTGAAATCCCCGCGGCGCCGCCTTCCGGCAACACCATGGGTGGACGACGCAGCGACAGTGGCGAGCGCCAGGTGCAGGTCGAAGGCAGCA
TCCGCCATCAAAGCGGTCTCAACCCGGGTTTTACCTTCGAAACCTTTGTCGAGGGCAAGTCCAACCAGCTGGCGCGGGCTGCATCGCGCCAGGTGGCCGA
GAATCCCGGTGGCGCCTACAACCCGCTGTTCCTTTATGGCGGTGTGGGTCTGGGTAAAACGCACCTGATGCACGCCGTTGGCAACCACCTGGCCGGTCTT
CGAGACAACGCCCGGGTGGTCTATCTGCATTCCGAGCGCTTCGTGGCCGACATGGTCAAGGCGCTGCAGCTCAACGCTATTAACGACTTCAAGCGCTTTT
ATCGCAGCGTCGATGCGCTGTTGATTGATGACATTCAGTTCTTTGCCGGCAAGGAGCGCTCTCAGGAGGAGTTCTTCCATACCTTCAACGCCCTGCTTGA
AGGCGGCCAGCAGATGATTCTGACCTCGGACCGCTATCCCAAGGAGATCAGCGGTGTCGAGGAGCGCCTGAAATCGCGCTTTGGCTGGGGCCTGACGGTN

>Kushneria marisflavi SW32 = DSM 15357 [NZ_CP021358.1:2089449..2089948], <mnmG oriC2 <PRK13623 (ErpA), 622 bp

AAGGGCAGCGACGTCGAAATGCCGTTGGGATAGAGCTCATGGGTGTCGAGCCCTTCCGGTTCGATAAAGACCTGATGGCTCTGCTTGTCGGCAAAGCGAT
GGATCTTGTCCTCGATGGAGGGGCAGTAGCGTGGCCCGGTGCTTTCGATGACACCGGAATACATCGGTGAGCGATCCAGATTGGCCCGAATGATGTCGTG
CGTGCGCTCGTTGGTGTGAGCAATGAAGCAGCTCACCTGATCGGGATGCATATCCCGCCGGCCCATGTAGGACATCACCGGACGCGGTGTGTCACCGGGC
TGTTCAGCCAGCTTCGAAAAATCGACCGTTCTGGCATCAAGCCTCGGCGGGGTACCGGTCTTGAGACGTGCTACCCGCAGGGGCTGCTCGCGAAGACGGG
CTGCCAGCGCATTGGACGCCGGATCGCCGGAGCGGCCACCGCTGTGATGCTCCATGCCGATATGGATGACACCACCAAGGAAAGTCCCGGTACACAGCAC
CACTGTCTCGGCCATGAAACGTACGCTGCCCTGGGTCAGGACGCCGCGCACGGTGTCTCCCTCAACGATCAGATCATCAGCGGCCTGAGCGAAGATCGAC
AGATTGGGCTGGTTTTCAAGGATATGGCGAATGGCGGCCTTGTAACGAACACGATCAGCCTGAACGCGAGTGGCCCGAACGGCAGGGCCCTTGCGGGAGT
TGAGTGTCCGAAACTGGATGCCACCCTGATCGGATGCCAGCGCCATGGCACCACCAAGGGCATCAATTTCCTTGACCAGATGACTCTTGCCGATGCCACC
AATGGCCGGATTACAGGACATCTGTCCCAGCGTTTCGATGTTGTGACTGAGCAGAAGGGTCTGACAGCCCATGCGAGCGGCGGCCAGTGCGGCTTCGGTA
CCCGCATGACCACCACCGATGACAATAACGTCAAAGCGGTCCGGATAATCCACGTGAAAAACCTCGAGGGGTTGAAGATGAACGTGACACCCGACTGATC

AGCGCGTGTCGATACCCACACACCCGTTCAGGGGGTTGGGGATAAGCCTTGTTGAATGGCGCGCATTATAACGTGGCTGTGAGTAAGCGTCAGGGTTTTC
CACACAGTGATTTTTCAAAAGGAGAAGTCATTAATTAATAAAAATAGTCTTTTAAATAAGAAGTTCTGTTGTGGTTGTGATTATTGGGGGCACCCATTTC
**TGTGGAAAA**GCCATTTTTAGACTTTAAAAACAGGAGGCTGGGATGTGACTGGAAACCCAGGCAAGCTGCGTGCTTCCCTGGGGCAGGTACTGTACGGGGT
GTGCATAAAAGCTCGGGTTACCCACAGCACGATTTATCCCCTGCCCGTTAACCGGGCTCAATGAGGCTTCTCCCCATACTCATCAAGAGTGACCATGCTA
CCTAAAAAAGTGACGAAGCGAGCTCATGTGGCGGGCTGCAGGGCAGATGGGCAAAAAGGGCGCTTTCTTTTATCCACAGGACAGTGGCATCAAAAAAGAT

CAGTGCAGACAATAAATGGGCATAAAAAAGGCGCCTGTCAGCTTTTGACAGACGCCCGAGTCCCGATAGGAAGTATCAGACCGTAAAGGATGCCCCGCAG
CCACAGGTCGTGGTGGCATTGGGGTTCTTGATCAAAAAGCGAGCGCCGGCCAGACCCGCTTCAAAGTCCACCGTGGAGCCCACCAGGTACTGATAGCTCA
GCGGGTCGATCACCATGGCGACATCGCCATTTTCGACGACGGTATCGTCTTCTCCAACGGTTTCTGCAAAGTCAAAGCCATACTGAAAGCCCGAGCAGCC
GCCTCCCGTTACGTAGACGCGCAGTTTCAGGGCCGGATTGCGCTCTTCCTCGATCAGCGCCCTGACTCTGCCTGCCGCGGCATCGGTAAAGAAAAGCGGT
GTCGGAACAAAGGATTCTGCTCCGCTCATCGCGGCCTCCTTATGAGATGACATGTCCGGGACATGTCAGAAATGGATCAGCACGCAGGCTGTCATGACAA
CCATTATCGACACCCCCTCTGTCTATGGTCAACCGCCACACGTCAGTGTCTGTCATGAGAGGCATTGAACATAAACAATGATAGGCTCATTGGTGGGGTG
TTCCATGATCCGCCATTGTGTCGCTGGGGAGGAGAAGCTTGCCGCGCCGTTTTCTGGATACGTCACGCCTGAACCTTGAGAACTTTCGACGCGAGCTGGC
AAGCGTCGATGCATTGCCACAGCTGTGTCTGCTGGCCGTCATTTCCGGCGCATTGACCGGTATCATCATGATTGCCTTTCGAGAACTGATCGATCTGGGC
GCGCTGAAGCTGATGCCCGACGGCCATGGTGAAGCCTTTGAAAGCCTGCCCGTGACGGTTCGCATGGGATTGCCGGTGGTGGCCGTCATTCTGATTGGCC
TGGGGCTTGGCTGGCAGAAGAAGAGTGCGCGTCGACTGGGGATCTCGCACGTCATCGAACGCTTCTCCTTTCATCAGGGGGTCATGGCACGTGACAACTG

>Kushneria marisflavi SW32 = DSM 15357 [NZ_CP021358.1:2127645..2127738], dnaA> ig dnaN>, 94 bp

WebSkew GCmin pos. 355636 (GCmin<>oriC1 47.9 kb; GCmin<>oriC2 4.9 kb; oriC2<>oriC1 43.0 kb)

>Kushneria phyllosphaerae EAod3 [NZ_ONZI01000001.1:c307454..307953], <rpmH ig dnaA>, 332 bp

CGCCGTTGACGGTCGCGGTCATGTCGACCTGAAGCGACTTCTGATCATCCCCCAGCTGGTAACGGGTCTGCTCGGGGCGGAAGGTGATCCGGCCCTTGTG
GCCTTCCATCTGCAGTCCGGAGCGCGCGATATAGCTGCGGCGCTGATCATCGGAGAGCAGAACGAAGGGCGCATCGCTATCGAGGCGCTGCAGATGCTTG
GGCAGGGCGGCATAGACGATATCGCCACCGCGCGGATCAATGCGAAGGTCAAGGGCATCGGTACGAACACTGATCAGGCTGGTGTTTTGAGCATCAGCGC
TATCGGGACTGTCAGTGGTGTTGCCGGGCTGTGGCGTGCTGGCACGGTCACTGCCGGTATCGGGTGCATCGAGCGACTGCTCACCGGGTTCAACGTCACT
GCTTTGCTGGGATAGGGCGGGAGGCGCTTCATCGGGCCGGTCGTTTTGCCCGTGATCATTGCTCCATTGAATGATCATCAGGTAGGCAAGTACGGCCAGC
GGGATGACAAAAAGAAGTCGTTTCAGATCCATTGGGATTGCCCGGTGGGAAAATGTCGACGCGGCGCAAGAAAACGACGGGTCGCACTCGGCGATTCGTC
GCGCCTCGTATTATTGCTGATGCTCCGGCTGGCGTTTGACGTCCTTTTCAAGGCGTCGCCACATGCCGTGAAGCTGTCTGGCCAGTGTTTCCGAGTCGAT
CTCGTGCACGCCCTTGCGTGACAGCACGACAATATCCACACCCGGCAACCACTGCTGGCGCAGTCGCACCGATTCTCGAGTCAAACGCTTGAGACGGTTG
CGATCCACGGCTCGGCGTACGTTCTTCTTGCTGAACACCAGCCCTACACGAACGTGACCGAGATCGTTCGGAGTCGCGAGTGCGAGAAGACCCTTGCCGT
GAATCCTGAAACTGGCATGTTCGAAAACATGACGGTATTCCCCGGCGGTCAGGATTCGAAGACCGCGGGGAAATCCGTGATTGGTCATGAGCTCGACTGA

TCCTTTAAAAGGATCAGGCGCTCAGGCGCTTGCGGCCCTTGGCGCGGCGACGCGAGATCACCTGACGGCCGTTTTTGGTGGCCATGCGAGCGCGAAAACC
GTGGTTACGCTTGCGCTTGAGGACGCTGGGCTGGAAAGTACGTTTCATGGTTGCGATTCCCGAATTGAGACTTGAGTGATCGGTGACACCGATGCCCTCA
AAATCGAGGCGCACGGGGGTCAAGAGCCGGGATTCTAGAGAATTCAACGAAGACGTGCAATTTTTATCCACGCACGAATATGACCGCTCTGTGTATAACT
TTGTGCACAAGCTGCGCACATCGTTTTTCCACACGCATCTGCACGCCTTGGAGAGGCCCTGGTAGCGGGGATAAGGTTATCCACAGCGGGTTTCGACCAC
TAACCCGGTGTGGATAACCCGTAGCACGAGCGTTACACTACCGCCCCATTTGCACGACTCACCATTGATAGAGGTCGCGTGTGTCCATCGCTCTCTGGCA

GCAATGTCTTGACTATTTACAGGATGAACTGAGTTCTCAGCAGTTCAACACCTGGATCCGTCCCCTGCAGGCCGAAAGCGGTTCCAGTGACCACGAACTC
TCCCTGCTGGCCCCCAATCGCTTTGTGCGTGACTGGGTCAACGACAAGTATCTAAAGCGCATCAACGAGCTGATGCGCGAGCTGGCTCAGGGCCGCCCGC
CCAAGGTGTCGCTGGACATTGGCAGTCGTCGCACGGTATCCAATCCCGTGTCACGTCCGGCCGCTCAAACAGAGCCCAGTGCAGCGCCTCGCCATGATGA
CACTGGCGATCATTCGCAGGCTGCCTGGTCTTCAAGCGTGACCCGGGCCTCGGCACGCGCGCCCATGCCGCCCTCGCCGCCGACATCGCCGCGGGGTAAC
TATCATGACGAGCGCGAAATTCCTTCCGCGCCGCCTTCCGGCAATACCACCGGCGGGCGCCGCAGTGACAACGGCGAGCGCCAGGTACAGGTCGAGGGCA
GCATCCGCCATCAAAGCGGGCTGAACCCGGGATTTACCTTTGAAACCTTTGTCGAGGGTAAATCCAACCAGCTGGCGCGTGCCGCGTCGCGACAGGTCGC
CGAAAACCCGGGTGGTGCCTACAACCCGCTGTTCCTTTATGGCGGTGTGGGTCTGGGTAAAACGCACCTGATGCACGCCGTCGGAAACCATCTGGCCGGT
CTTCGAGACAACGCCCGGGTGGTCTATCTGCACTCCGAGCGTTTCGTGGCCGACATGGTCAAGGCGCTGCAGCTCAACGCCATTAACGACTTCAAGCGCT
TTTATCGCAGCGTCGATGCGCTGTTGATCGATGATATCCAGTTCTTTGCCGGCAAGGAGCGTTCTCAGGAAGAGTTTTTCCATACCTTCAATGCGCTCCT
TGAAGGCGGTCAGCAGATGATTCTGACCTCTGACCGCTACCCCAAGGAGATCAGTGGGGTCGAGGAGCGTCTGAAATCACGCTTTGGCTGGGGGCTGACG

>Kushneria phyllosphaerae EAod3 [NZ_ONZI01000001.1:c350434..350933], <mnmG oriC2 <PRK13623 (ErpA), 620 bp

AAGGGCAGCGATGTTGAAATGCCGTTGGGGTAGAGCTCATGGGTGTCGAGCCCTTCCGGCTCGATAAAGATCTGGTGGCTCTGCTTGTCGGCAAAGCGAT
GAATCTTGTCCTCGATCGACGGGCAGTAGCGGGGACCGATGCCCTCGATGACGCCGGAGTACATCGGCGAGCGATGCAGATTGGCCCGAATGATGTCGTG
AGTGCGCTCGTTGGTGTGCGCAATGAAGCAGCTCACCTGGGCGGGGTGCATGTCACGCCGGCCCATGTAGGACATCACCGGACGCGGCGTATCGCCGGGC
TGTTCGGCCAGTTTCGAGAAATCGACCGTTTTGGCATCAAGTCGCGGTGGTGTGCCGGTCTTGAGACGTGCTACCCGCAGGGGCTGCTCGCGAAGACGTG
CCGCCAGCGCATTGGACGCCGGATCACCCGAACGGCCGCCGCTATGATGTTCCATGCCGATATGAATCACGCCGCCCAGAAAGGTCCCGGTGCACAGCAC
CACGGTTTCGGCCATGAAACGTACGCCGCCCTGACTCAGGACACCGCGCACTGTATCGCCTTCCACGATCAGATCATCAGCTGCCTGGGCAAAGATCGAC
AGATTGGGCTGGTTTTCAAGGATATGGCGAATGGCTGCCTTGTAGCGAACGCGGTCGGCCTGAACACGGGTGGCGCGTACGGCCGGTCCCTTGCGGGAGT
TCAGCGTTCGAAACTGGATGCCGCCCTGATCTGCAGCCAGTGCCATGGCACCGCCCAGCGCATCAATTTCCTTGACCAGATGGCTCTTGCCGATGCCACC
GATGGCCGGATTACAGGACATTTGTCCCAGCGTTTCGATGTTGTGACTGAGCAGCAGGGTCTGACAGCCCATGCGAGCGGCGGCCAGTGCGGCTTCGGTA
CCCGCATGACCACCACCGATGACAATGACGTCAAAGCGGTCCGGATAATCCACGTGAAAAACCTCGAGGGGTTGAAAATGAACGTGACACCCGACTGATC

AGCGCGTGTCGATACCCACATGCCCGTGCAGGGCGTTGGGGATAAGCCGTATTGAATGGCGCGCATTATAACGTGGCTGTGAGTAAGCGTCAGGGTTTTC
CACACATGGATTTTCCCGTACCCCCAAGCCTTTAATTAATTAAATAGTCTTTTAAATAAGAAGTTCTATTGTGGTTGTGATTACTGTGGGCGGCATTTTC
**TGTGGAAAA**GCGGCTTTTACACTTATAAAACAAAATGCTGCGGTGTGACTGGAAACCCAGGCAAGCTGCTTGCATCCCTGGGGTAGGCACTGTACGGGGT
GTGTATAAAACGGCAAGTTACCCACAGCTCGATTTATCCCCGTTTCGTTAACCGGGCTCAGTGGGGGTTCTCCCCATACTCGTCAAGAAAGTGAATGCGG
CAGGAAAATGAGAGGAGACTCGCGTGTGTGGCGGGCTGTAGAGGCGGTGGTAGAAAAGGGCGCTTTTTTTATCCACAGGGGGCCGGCGTCAAAAAAAGCC

GATTGTGGTGATAATGGACATAAAAAAGGCGCCTGTCAGAAGCTGACAGGCGCCTGAATATCGGCAGGCGACCTCAAACCGTAAAGGATGCACCGCAGCC
ACAGGTCGTGGTGGCGTTGGGATTTTTGATCAAAAACCGGGCGCCTGCCAAGCCGGCTTCGAAATCGACCGTGGAACCGACCAGATATTGATAGCTCAGC
GGGTCGATCACCATGGCCACATCGCCATTTTCGATAACGGTGTCGTCCTCACTGATGGTCTCTGCGACATCGAAACCGTATTGAAAGCCCGAACAACCGC
CTCCCGTTACGTAGACGCGCAGTTTCAGGGCCGGGTTACGCTCTTCTTCTATCAGCGCCCTGACTCTGCCTGCCGCGGCATCGGTAAAGACAAGCGGTGT
CGGAACAAAGGATTCTGCTCCGCTCATCGCGGCCTCCTTTTGAGATGACACGTCTGATGCGTGTCAGGAATGGATCAGCACGATGGCTGTCATGGCACCC
ATTATCGGCACCCCTTCAGCCGGTGGTCAACCGGCACCCGCCGGGCAGCAATCATGAAAGGCGATGACCCTAAACAATGATAGGCTGAAGGACGTGTGGT
CCCATGACCGCATTACCGTGCCGCTGGGGAGAAAAAGCTTGCCGCGCCGTCTGCTGGATATGTCACGCCTGAACCTTGAAAACTTTCGACGCGAGCTGGC
AAGTGTCGATGCCCTGCCGCAGCTGTGTTTGCTGGCGGTCATTTCGGGTGCCTTGACCGGCCTCATCATGATTGTCTTTCGAGAGCTGATCGGGCTCGGG
GCGCTCATGCTGATGCCAAACGGTCAGGCCGAGGCCTTTGAAAGCCTGCCGATGATGGTGCGCGTGGCCCTGCCGGTGATCGCGGTCATCCTGATCGGGC
TGGGACTTGGCTGGCAGAAAAAAAGCGCCCGGCGCCTGGGCATTACACATGTCATCGAGCGCTTCTCCTTTCATCAGGGCGTCATGGCGCGCGACAACTG

>Kushneria phyllosphaerae EAod3 [NZ_ONZI01000001.1:c305826..305922], dnaA> ig dnaN>, 97 bp

WebSkew GCmin pos. 627732 (GCmin<>oriC1 46.0 kb; GCmin<>oriC2 7.7 kb; oriC2<>oriC1 38.3 kb)

>Kushneria indalinina DSM 14324 [NZ_QRDJ01000007.1:673518..674017], <rpmH ig dnaA>, 330 bp

TCATGTCAACCTGCAGCGACTGCTGGTCATCTCCCAGCTGGTAGCTGGTCTGCTCGGGGCTGAAGGTGATTCGGCCCTGATGGCCTTCCATCTGTAGCCC
GGAGCGCGCGATATAGCTGCGGCGCTGATCGTCGGAGAGCAGAACAAAGGGGGCATCGGAGTCAAGACGCTGGAGATGCTTCGGCAGGGCGGCATAGACA
ATATCACCACCGCGCGGGTCAATACGCAGGTCAAGGGCGTCGGTGCGAACGCTGATCAGGCGGGTATTCTGAGCATTGGCATCGCTGGTATTGCTGGCAC
TGTCAGTGTCGCCGGGCATATCGCCGGAATCAGGCGCACTGGCGCGTTCACTGGCGGCCTCGGGGGCATCGAGCGACTGCTCACCGGGTTCAACGTCACT
GCTTTGCTGGGTAAGGGCAGGAGGCGCTTCATCCGGTCGGTCGTTTTGCCCGTGATCATTGCTCCATTGAATGATCATCAGATAGGCAAGTACGGCCAGC
GGGATGACAAAAAGAAGTCGTTTCAGATCCATTGGGATTGCCCGGTGGGAAAATGTCGACGCGGCGCAAGAAAACGACGGGTCGCACTCGACGGTTCGTC
GCGCCTCGTATTATTGCTGATGCTCCGACTGACGTTTGACGTCCTTTTCAAGGCGTCGCCACATGCCGTGAAGCTGTCTGGCCAGTGTCTGCGAGTCGAT
CTCGTGTACGCCCTTGCGTGACAGCACGACGATGTCCACACCCGGCAGCCACTGCTGGCGCAGTCGCACCGATTCTCGAGTCAAGCGCTTGAGACGGTTG
CGATCCACGGCTCGGCGCACGTTCTTCTTGCTGAACACCAGTCCCACACGAACGTGACCGAGATCGTTGGGGGTGGCGAGTGCCAGAAGACCCTTGCCGT
GAATCCTGAAACTGGCATGTTCGAAAACATGACGGTATTCCCCGGCGGTCAGGATTCGAAGACCGCGGGGAAATCCCTGAGTGGTCATGAGCTCCGCTGA

TCCCTTGAGAGGATCAGGCGCTCAGGCGCTTGCGGCCCTTGGCGCGGCGGCGCGAGATGACCTGGCGGCCGTTCTTGGTGGCCATGCGAGCGCGGAAGCC
GTGGTTACGCTTGCGCTTGAGCACGCTGGGCTGAAAAGTACGTTTCATGGTTGCGATTCCCGAATTGAGACTTGAGTGATCGGTGACACCGATGCCCTCA
AAATCGAGGCGCACGGGGGTCAAGAGCCGGGATTCTAGAGAATTCAACGAGGACGTGCAATTTTTATCCACGCACGAAAAGGACAGCGCTGTGCATAACT
TTGTGCACAAGCTGCGCACTTCATTTCCGGAAGGCCATCCATGCGACCCGGAGAGGCCCGGGTGACGGGGATAAGGTTATCCACAGGCCGTCTGAAGTCC
CAACCCGGTGTGGATAACCCGTTCCACGAGCGTTACACTACCGCCCCATTTGCACGACTCACCATCAAGAGGTCGCGTGTGTCCATCGCTCTCTGGCAGC

AATGTCTCGACTATTTACAGGATGAATTGAGCTCTCAGCAGTTCAATACCTGGATCCGTCCCCTGCAGGCCGAAAGCGGTTCCAGTGACCACGAACTCTC
CCTGCTGGCCCCCAATCGCTTCGTGCGTGACTGGGTCAATGACAAGTATTTAAAGCGCATCAACGAGCTGATGCGCGAGCTGGCCCAGGGCCGGCCGCCC
AAGGTGTCGCTGGATATCGGCAGCCGCCGTACGGTGTCCAACCCCGTATCGCGTCCGGCGACCGACAGCGAACCTGCGACCGGCCAGGCGCAAAACAATA
GCGGTGATCATGGCCAGGCGGCCTGGTCTTCAGGCGTGACACGTTCCTCGACGCGTGCGCCCATGCCGCCTTCGCCACCCACGTCGCCGCGCGGCGACTA
TCAGGACGAGCGCGAGATCCCTGCTGCGCCGCCCTCCGGCAATACCACCGGCGGGCGTCGCAGTGACAGCGGCGAACGTCAGGTGCAGGTCGAAGGCAGC
ATTCGCCACCAGAGCGGGCTCAACCCGGGATTTACCTTTGAAACCTTCGTCGAGGGTAAATCCAACCAGCTGGCGCGGGCCGCCTCGCGCCAGGTCGCCG
AGAACCCGGGCGGTGCCTATAACCCGCTGTTCCTTTATGGCGGTGTGGGTCTGGGTAAAACCCACCTGATGCACGCGGTCGGCAACCATCTGGCCGGGCT
GCGCGACAACGCTCGGGTGGTCTATCTGCACTCCGAGCGCTTCGTGGCCGACATGGTCAAGGCGCTGCAGCTCAACGCCATCAACGACTTCAAGCGCTTT
TATCGCAGCGTCGATGCGCTGCTGATCGATGACATCCAGTTCTTTGCCGGCAAGGAGCGCTCCCAGGAGGAGTTCTTCCACACCTTTAATGCGCTGCTGG
AAGGCGGTCAGCAGATGATTCTGACCTCGGACCGCTACCCCAAGGAGATCAGCGGCGTCGAAGAGCGTCTGAAATCGCGCTTTGGCTGGGGCCTGACGGT

>Kushneria indalinina DSM 14324 [NZ_QRDJ01000007.1:635195..635694], <mnmG oriC2 <PRK13623 (ErpA), 622 bp

AAGGGCAGTGATGTCGAAATGCCGTTGGGATAGAGCTCATGAGTATCGAGCCCCTCCGGTTCGATAAAGACCTGGTGGCTCTGCTTGTCTGCAAAGCGAT
GAATCTTGTCCTCGATCGACGGACAGTAGCGCGGCCCGGTGCTGTCGATGACACCAGAGTACATCGGTGAGCGATCCAGATTGGCGCGGATGATGTCGTG
AGTGCGTTCGTTGGTGTGTGCGATGAAGCAGCTCACCTGATCGGGATGCATGTCACGGCGCCCCATGTAGGACATCACCGGACGGGGGGTGTCACCGGGC
TGTTCGGCCAGTTTCGAGAAATCGACCGTTTTGGCATCCAGACGCGGCGGCGTGCCGGTTTTCAGCCGTGCTACTCGCAGGGGCTGCTCACGCAGGCGCG
CAGCCAGTGCATTGGAGGCAGGATCCCCTGAACGACCGCCGCTGTGATGTTCCATGCCGATGTGAATCACGCCGCCCAGGAAGGTACCGGTACACAGCAC
CACGGTCTCGGCCATGAAACGGATACCGCCCTGGGTCAGCACACCGCGCACCGTGTCGCCTTCCACGATCAGATCGTCAGCGGCCTGGGCAAAGATCGAC
AGGTTGGGCTGGTTTTCAAGGATATGGCGGATGGCTGCCTTGTAGCGCACGCGATCGGCCTGGGCACGGGTGGCACGCACGGCCGGTCCCTTGCGGGAAT
TGAGCGTTCGAAACTGGATGCCGCCCTGGTCGGTAGCCAGCGCCATGGCACCGCCAAGCGCATCAATTTCCTTGACCAGATGACTCTTGCCGATCCCGCC
GATGGCCGGATTACAGGACATCTGTCCCAGCGTTTCGATGTTGTGACTGAGCAGCAGGGTCTGGCAGCCCATGCGAGCGGCGGCCAGTGCGGCTTCGGTA
CCCGCATGACCACCGCCGATGACAATAACGTCAAAGCGGTCCGGATAATCCACGTGAAAAACCTCGAGGGGTTGAAGATGAACGTGACACCCGACTGATC

AGCGCGTGTCGATACCCACATGCCCGTGAAGGGGGCTGGGGATAAGCCGTATTGAATGGCGCGCATTATAACGTGGCTGTGAGTAAGCGTCAGGGTTTTC
CACACATGGATTTTCTCCGGAGAGAAAGTCATTCATTAATAAAATGGTCTTTTAAATAAGAAGTTCTGTTGTGGTTATGATTATTGGTGAAGCCACTTTC
**TGTGGAAAA**GCGATGTTTCCTCTTGGAAAACAGATTTCTGGGATGTGACTGGAAGCTCGGGAAAGCTGCGTGCTTCCCTGGGGCAGGCCCTGTACGAGGT
GTGCATAACCGGGTGACTTACCCACAGCTCGATTTATCCCCGTCTCGTCAACCGGGCTCAGGGCCAGTTCTCCCCGCAGTCATCAAGAAAGGCACCAGGG
CGTGGAAAAAGGGAAAGGCTCCCGCGTGTGGCGGGCTGTAGCGCAGAGGAAGCAAAAGGGCGCGGATTATTATCCACAGGGGGAGTGAGGCAAAAAAATT

GGCGAGAGTGCGCAAATGGGCATAAAAAAGGCGTCTTTCAGAAGCTGACAGACGCCCGAAGATCGCCAGGGGGCATCAGACTGTAAAGGATGCACCACAG
CCACAGGTCGTGGTGGCATTGGGGTTTTTGATCAAAAAGCGGGCCCCTGCCAGACCGGTTTCGAAATCGACCGTGGAACCAACCAGATATTGATAGCTCA
GCGGGTCGATCACCATGGCAACATCGCCATTTTCGACAACGGTGTCGTCTTCGCCGACGGTTTCTGCAAAATCGAAACCGTACTGAAAGCCCGAACAACC
GCCTCCCGTTACGTAGACGCGCAGTTTCAGGGCCGGGTTGCTTTCTTCTTCTATCAGCGCCCTGACTCTGCCTGCAGCGGCATCGGTAAAGACAAGCGGG
GTCGGAACAAAGGATTCTGCTCCGCTCATCGCGGCCTCCTTTAGAGATGACACGTATGATACGTGTCAGGAATGGATCAGCACGATGGCTGTCATGGATC
CCATTATCGACACCCCCCCAGCCGGCGGTCAACCGGCATGCGCTGGGCGGTGATCATCAAGAGTGACAGATAAAAACAGTGATAGGCTGATAGGCGTGGT
GTCCCATGACCCCGTTATTCCGCCGTTGGGGAGAAAAGCTTGCCGCGTCGTCTTCTGAATACGTCCCGCCTGAGTCTGGAAAACTTTCGACGCGAACTGG
CGAGCGTCGATGCACTGCCGCAGCTGTGTCTGCTGGCGGTCATTTCAGGTGCGCTGACAGGCGTCATCATGATTGTCTTTCGGGAGCTGATTGAACTGGG
AGCGCTGGGGCTGATGCCCGAAGGGCGTACCGAGGCCTTTGAGGATCTGCCCGTCATGGTTCGCATGGGCCTGCCGGTCATCGCCGTCGTGTTGATCGGG
CTGGGGCTCGGGTGGCAGAAGAAGAGCGCGCGTCGACTCGGCATTACCCATGTCATCGAGCGCTTCTCCTACCATCAGGGCGTCATGGCCCGTGACAACT

>Kushneria indalinina DSM 14324 [NZ_QRDJ01000007.1:675547..675643], dnaA> ig dnaN>, 97 bp

WebSkew GCmin pos. 249473 (GCmin<>oriC1 35.4 kb; GCmin<>oriC2 0.3 kb; oriC2<>oriC1 35.1 kb)

>Kushneria pakistanensis BKCTC 42082 [NZ_BMZM01000002.1:c213823..214322], <rpmH ig dnaA>, 330 bp

ACGCTCATGTCGACCTGCAGGGACTGCTGGTCATCACCCAGCTGGTAGCTGGTCTGCTCGGGGCTGAAGTTGATTCGGCCCTGATGACCTTCCATCTGCA
GTCCGGAACGCGCGATATAGCTGCGGCGCTGGTCATCGGAGAGCAGTACGAAGGGAGCATCGGACTCAAGACGCTGCAGATGTTTCGGCAGGGCGGCATA
GACGATATCGCCACCGCGCGGATCGATACGCAGATCAAGCGCATCGGTGCGAACGCTGATCAGGCGGGTATTCTGAGCATTGGCATTGTCGGCACTGTCA
GTATCGCCTGGCATGTCGCCGGACTGCGGCGCACTGGCGCGCTCACTGGCGGACTCGGGCGCATCAAGCGACTGTTCACCGGGTTCAACGTCACTGCTTT
GCTGGGACAGGGCGGGAGGCGCTTCATCGGGCCGGTCGTTGTGCCCGTGATCATTGCTCCATTGAATGATCATCAGATAGGCAAGTACGGCCAGCGGGAT
GACAAAAAGAAGTCGTTTCAGATCCATTGTGGGATTGCCCGGTGGGAAAATGTCGACGCGGCGCAAAAAAAACGACGGGTCGCATTCGACGGGTCGTCGC
GCCTCGTATTATTGCTGATGCTCCGGCTGACGTTTGACGTCCTTTTCAAGGCGTCGCCACATGCCGTGAAGCTGTCTGGCCAGTGTCTGGGCGTCAATCT
CATGCACGCCCTTGCGCGACAGCACCACGATATCCACGGCCGGCAGCCACTGCTGGCGCAGTCGCACCGATTCTCGCGTCAAACGCTTGAGACGGTTGCG
GTCCACGGCACGGCGCACGTTCTTTTTGCTGATGACCAGTCCGACCCGAACGTGACCGAGGTTATTGGGGGTGGCGAGTGCCAGAAGCCCCTTGCCGTGA
ATCCTGAAACTGGCATGTTCGAAAACATGACGGTATTCCCCGGCGGTCAGGATTCTGAGACCGCGGGGAAATCCCTGACTGGTCATGAGCGCAGCTGATC

CACTAATAAGGGATCAGGCGCTCAGGCGCTTGCGGCCCTTTGCACGACGACGTGCGATGACCTGACGGCCGTTCTTGGTGGCCATGCGGGCACGGAAGCC
GTGGACACGCTTGCGCTTGAGCACGCTGGGCTGGAAAGTACGTTTCATGGTTGCGATTCCCGAATTGAGACTTGAGTGATCGGCGACACCGATGCCCTCA
AAATCGAGGCGCACGGGGGTCAAGAGCCGGGATTCTAGAGAATTCAACGAGGACGTGCAATTTTTATCCACGCATCAATATCAGGCGGGTGTGCATAACT
TTGTGCACAAGCATTACACATCATTCCTGCAGGGGCAGCTGTACGGCCTGCAGGGGCCCTGACAGCGGGGACAAGGTTATCCACAACCGGTTTTTCACGC
CAACCCGGTGTGGATAACCCGCTGCGCGAGCGTTACACTATCGCCCCATTTGCACGACTCACCATCAAGAGGTCGCGTGTGTCCATCGCTCTCTGGCAGC

AATGTCTTGACTATTTACAGGACGAATTGAGCTCTCAGCAGTTCAATACCTGGATCCGTCCCCTGCAGGCCGAAAGCGGTTCCAGTGACCACGAGCTCTC
CCTGCTGGCCCCGAATCGATTTGTGCGTGACTGGGTCAACGACAAGTATTTAAAGCGTATCAACGAACTGATGCGCGAGCTGGCCAACGGCCGCCCGCCC
AAGGTGATGCTGGATATCGGCAGCCGCCGCACGGTATCCAATCCCGTGTCGCGCCCGGCTGGTAGCGGTGAGCACGATGCCACCGCGTCTCGGGATAGCG
CCAGCGAGCCCGCCGTGGAGGCCTGGTCATCGGGTGTGACGCGTGCGTCTGCGCGCCCGCCCATGCCGCCGTCCCCACCTGTTTCGTCGCGAGGCAGCTA
CCACGACGAGCGCGAGATTCCTGCGGCACCGCCTTCCGGCAACACGATGGGCGGTCGGCGCAACGACAACGGCGAACGTCAGGTCCAGGTCGAGGGCAGC
ATCCGCCATCAGAGCGGACTCAACCCGGGATTTACCTTTGAAACCTTTGTCGAGGGTAAATCCAACCAGCTGGCACGCGCCGCCTCGCGGCAGGTCGCCG
AAAATCCCGGCGGTGCCTACAACCCGCTGTTCCTTTATGGCGGCGTGGGCCTGGGTAAAACGCATTTGATGCACGCGGTGGGCAATCATCTGGCCGGACT
TCGTGACAATGCCCGGGTAGTCTATCTGCATTCAGAGCGTTTCGTGGCCGACATGGTCAAGGCGCTGCAGCTCAATGCCATCAATGACTTCAAGCGCTTT
TATCGCAGCGTCGATGCGCTGTTGATCGATGATATCCAGTTCTTTGCCGGCAAGGAGCGCTCGCAGGAGGAGTTTTTCCACACCTTCAATGCGCTGCTTG
AAGGCGGCCAGCAGATGATCCTGACCTCCGATCGTTACCCCAAGGAGATCAGCGGTGTCGAGGAGCGCCTCAAGTCCCGTTTTGGCTGGGGCCTGACGGT

>Kushneria pakistanensis BKCTC 42082 [NZ_BMZM01000002.1:c248931..249430], <mnmG oriC2 <PRK13623 (ErpA), 620 bp

AACGGCAGAGAGGTGGAGATACCGTTGGGATAGAGCTCATGCGTATCCAGCCCTTCCGGTTCGATGAAGACCTGGTGGCTCTGCTTGTCGGCAAAGCGAT
GGATCTTGTCTTCGATCGACGGACAGTAGCGCGGACCGATGCCTTCGATGACACCCGAATACATCGGTGAACGGTCCAGGTTGGCGCGAATGATGTCGTG
GGTACGCTCGTTGGTGTGCGCGATGAAGCAGCTCACCTGCTCGGGATGCATCCCGCGTCGGCCCATGTAGGACATCACCGGGCGCGGCGTGTCACCGGGC
TGCTCGGCCAGTTTCGAGAAGTCGATCGTTCTGGCATCCAGTCGCGGCGGGGTGCCGGTCTTGAGACGTGCCACACGCAGGGGCTGCTCCCGCAGTCGCG
CGGCCAGCGCATTGGAAGCCGGATCGCCCGAACGTCCGCCGCTGTGATGTTCCATGCCGATATGAATCACACCGCCCAGGAAGGTACCGGTACACAGCAC
CACCGTTTCGGCCATGAACCGCACGCCGCCCTGGGTCAGCACGCCGCGCACCGTGTCGCCTTCAACGATCAGATCGTCAGCAGCCTGAGCGAAGATGGAC
AGGTTGGGCTGGTTTTCAAGGATATGGCGAATGGCCGCCTTGTAGCGTACCCGATCGGCCTGGACGCGGGTGGCACGAACGGCCGGCCCCTTGCGGGAAT
TGAGCGTTCGAAACTGGATACCCCCCTGGTCGGTTGCCAGCGCCATGGCACCACCGAGTGCGTCAATTTCCTTGACCAGATGACTTTTGCCGATCCCGCC
AATGGCAGGATTACAGGACATCTGGCCCAGCGTTTCGATGTTGTGACTGAGCAGCAGGGTATTGCAGCCCATGCGGGCGGCAGCCAGTGCGGCTTCGGTA
CCCGCATGACCACCACCAATGACAATGACGTCAAAGCGGTCCGGATAATCCACGTGATGAACCTCGAAGGGTTTGAAGATGAACGTGACACCCGACTGAT

CAGCGCGTGTCGATACCCACATGCCCGGAAAGGGGCTGGGGATAAGCCGTATTGAATGGCGCGCATTATAACGCCCCTGTGAGTAAGCGTCAGGGTTTTC
CACACATGGATTTCCCGGGTCAAAAAAGTCTTTAATTAATAAAATATTCTTTTAAATAAGAAGTTCTGTTGTGGTTGTGATTAATGAGGGCGCCACTTTC
**TGTGGAAAA**GCGGTTTTTTGGTTTATAAAACAGCTTCCTGGGGTGTGATTGAAGACCCAGGCAAGCTGCGTGCTTCCCAGGGGCAGGCTCTGTACGGGAT
GTGCATAACCTGATGACTTACCCACAGATCGATTTATCCCCTTCTCGTCAACCGGGCTCAGGCGGGGTTCTCCCCATACTCATCAAGAAAAGCGTCAGCG
TGGAAAAAAGAAGCATGATGCCCGCATGTGGCGGGCTGCAGCGCAAGGCAGGGAGAGACGCGTAATTTTTTATCCACAGGGGATGGGCACAAAAAAAGCG

CCAGAGGATGCTTATAGGCATAAAAAAAGCGTCTGTCAGAGGCTGACAGACGCCGGAACATCACCAAGGGGCATCAGACCGTAAAGGATGCACCGCAGCC
ACAGGTCGTAGTGGCATTGGGGTTTTTGATCAAAAAGCGGGCACCTGCCAGACCTGCCTCAAAATCGACCGTGGAGCCGACCAGATATTGATAGCTCAGC
GGGTCGATCACCATGGCAACATCGCCGTTTTCAATCACGGTATCGTCTTCACTCACGGTTTCTGCAAAGTCGAAACCATATTGAAAGCCCGAACAACCGC
CTCCTGTTACGTAGACGCGCAGTTTCAAAGCCGGGTTGCTTTCTTCCTCTATCAGCGCCCTGACTCTGCCTGCAGCGGCATCGGTAAAGACAAGCGGTGT
CGGAACAAAGGATTCTGCTCCGCTCATCGCGGCCTCCTTTAGAGATGACACGTATGATACGCGCCAGGAATGGATCAGCACGGGGGCTGTCATGAACACC
ATTATCGACACCCCTCCAGCTGGCGGTCAACCGCCATGCAGGGGCTGCGATCATGACGGGCGCCATGTGTAAACAGTGATAGGCTGACCCCCTGTCGTGT
CATGAGCTTCGTTACCCTGTCGCTGGAGAGAAAAAGCTTGCTGCGTCGTCTTCTGGATCCATCCCGCCTGAGCCTTGAAAATTTCGACGCGAACTGGCA
AGCGTCGATGCCCTGCCGCAACTGTGTCTGCTGGCGGTCATCTCGGGGGCGTTGACCGGGCTTATCATGATTGTCTTTCGAGAGCTGATCGGGCTGGGTG
CACTGGTGCTGATGCCAAAAGGGCATGCCGAGGCGTTTGAAGCCCTGCCCATGGCGGTGCGCATGAGCATGCCGGTGATCGCTGTCGTGCTGATCGGGCT
GGGGCTTGGCTGGCAGAAAAAAAGCGCGCGCCGACTCGGCATTACCCATGTCATTGAGCGCTTTTCCTTTCATCAGGGGGTCATGGCGCGCGATAACTGG

>Kushneria pakistanensis BKCTC 42082 [NZ_BMZM01000002.1:c212197..212293], dnaA> ig dnaN>, 97 bp

WebSkew GCmin pos. 490590 (GCmin<>oriC1 50.4 kb; GCmin<>oriC2 7.2 kb; oriC2<>oriC1 43.2 kb)

>Kushneria avicenniae DSM 23439 [NZ_FOLY01000003.1:540766..541265], <rpmH ig dnaA>, 330 bp

GTCCACGCCATTGACGGTCGCGCTCATGTCGACCTGAAGCGACTGCTGATCATCGCCCAGCTGATAGCGGGTCTGCTCGGGCGTAAAGGTAATCCGGCCC
TGGTGGCCCTCCATCTGCAGTCCGGAACGGGCGATATAGCTGCGTCGCTGATCGTCGGAAAGCAGAATGAAGGGGGCATCGCTGCCAAGGCGCTGCAGAT
GTTTGGGCAGGGCGGCATAGACAACATCGCCACCGCGCGGGTCGATGCGAAGATCGAGTGCATCGGTGCGGACGCTGATCAGACGGGTGTTCTGGGCATT
GGCGTTATCGTCGCCGCCGGCAGCATTGTCCGACATGTCGACGGACGCTGGCGTGCTGGCGGTATCGGGGGCATCAAGTGACTGCTCACCGGGGGCAACG
TCACTGTTCTGCTGGGAGAGGGCAGGAGGCGCTTCATCCGGTCGGTCGTCGTGCCCGTAATCATTACTCCATTGAATGATCATCAGGTAGGCAAGTACGG
CCAGCGGGATGACAAAAAGAAGTCGTTTCAGATCCATTGGGATTGCCCGGTGGGAAAATGTCGACGCGGCGCAAGAAAACGACGGGTCGCACTCGACGGT
TCGTCGCGCCTCGTATTATTGCTGATGCTCCAGCTGACGTTTGACGTCCTTTTCAAGGCGTCGCCACATACCGTGAAGCTGTCTGGCCAGTGTTTCCGAG
TCGATCTCGTGGACGCCCTTGCGCGACAGCACAACGATATCCACCGCCGGCAACCACTGCTGGCGCAGTCGCACCGATTCTCGAGTCAAGCGTTTGAGAC
GGTTGCGATCCACGGCTCGGCGCACGTTCTTCTTGCTGAACACCAGTCCCACACGAACGTGACCGAGATCGTTGGGGGTGGCGAGTGCCAGAAGTCCCTT
GCCGTGAATCCTGAAACTGGCATGTTCAAAAACATGACGGTATTCCCCTGCGGTCAGGATTCTGAGACCGCGGGGAAATCCCTGATTGGTCATGAGCTGA

TCCTTGAAGGGGATCAGGCGCTCAGGCGCTTGCGGCCCTTGGCGCGACGACGCGAGATGACCTGACGGCCGTTTTTGGTGGCCATGCGGGCGCGGAAGCC
ATGGTTACGCTTGCGCTTGAGCACGCTGGGCTGAAAAGTACGTTTCATGATGGTGATTCCCGAATTGAGACTTGAGTGACCGGTGACACCGATGCCCTCA
AAATCGAGGCGCACGGGGGTCAAGAGCCGGGATTCTAGAGAATTCAACGATCACGTGCAATTTTTATCCACGCACAAAAATGATCCGGCTGTGTATAACT
TTGTGCACAAGCTCAGGGCTCTTCTTCCAGGAGGGCGGTAGCGGCATCAGGGAAGGCCCAAACAGCGAGGATAAGGTTATCCACAACCCTTTTCGCCACC
CAATCCGGTGTGGATAACCCCGGCGTCGAGCGTTACACTACCGCCCCATTTGCACGACTCACCATCAAGAGGTCGCGTGTGTCCATCGCTCTCTGGCAGC

AATGTCTTGACAATTTACAGGATGAGTTGAGCTCACAGCAGTTCAATACCTGGATCCGTCCCCTGCAGGCCGAAAGCGGTTCCAGTGACCACGAACTCTC
CCTGTTGGCCCCCAACCGCTTCGTGCGCGACTGGGTCAATGACAAGTATTTAAAACGTATCAACGAGCTGATGCGCGAGCTGGCGCAGGGTCGCCCGCCA
AAGGTGTCACTGGATATCGGCAGCCGCCGGACAGTGTCCAATCCCGCGCCGAAGCCTGCCGCCTCCGGTGAGCAGAGCAGCGCCGCGCCGCAGGGCAGCG
CCGGTCAGCCGGCACCGGAAACCTGGTCGGCAGGCGTGACCCGCACTGCTTCGGCCCCCATGCCGCCCTCGCCGCCGACCTCCCCGCGCGGTGACTATCA
GGATGAGCGTGAGATTCCGGCCGCGCCGCCCTCCGGCAACACGACCGGCGGGCGGCGCAGCGACAGCGGCGAGCGTCAGGTGCAGGTCGAGGGCAGCATT
CGCCACCAGAGCGGTCTCAACCCCGGCTTCACCTTTGAAACCTTTGTCGAGGGCAAGTCCAACCAGCTGGCGCGGGCCGCCTCGCGTCAGGTGGCCGAAA
ACCCCGGCGGTGCCTACAACCCGCTGTTTCTGTACGGTGGCGTGGGGCTGGGTAAAACGCACCTGATGCACGCGGTGGGCAACCATCTGGCCGGCCTGCG
CGACAACGCCCGGGTGGTCTATCTGCACTCCGAGCGTTTCGTGGCCGACATGGTCAAGGCGCTGCAGCTCAACGCCATCAACGACTTCAAGCGTTTTTAT
CGCAGTGTCGATGCGCTTCTTATCGATGATATCCAGTTCTTTGCCGGCAAGGAGCGCTCCCAGGAGGAGTTCTTCCACACCTTCAATGCGCTGCTGGAAG
GTGGTCAGCAGATGATTCTGACCTCGGACCGCTACCCCAAGGAGATCAGTGGCGTCGAGGAGCGCCTGAAATCGCGCTTTGGCTGGGGGCTGACGGTGGC

>Kushneria avicenniae DSM 23439 [NZ_FOLY01000003.1:497538..498037], <mnmG oriC2 <PRK13623 (ErpA), 623 bp

AAGGGCAGCGACGTCGAGATGCCGTTGGGATAGAGCTCGTGGGTATCAAGCCCTTCCGGTTCGATAAAGATCTGGTGGCTTTGCTTGTCGGCAAAGCGAT
GGATCTTGTCCTCGATGGAGGGGCAGTAGCGCGGGCCGGTGCTCTCGATGACGCCGGAGTACATCGGCGAGCGATCCAGATTGGCCCGGATGATGTCGTG
GGTACGCTCGTTGGTGTGCGCGATGAAGCAGCTCACCTGATCGGGATGCATCTCACGCCGCCCCATGTAGGACATCACCGGCCGCGGCGTATCGCCGGGT
TGTTCGGCCAGCTTCGAGAAATCGACCGTTCGGGCATCCAGCCGTGGCGGGGTGCCGGTCTTGAGACGTGCTACCCGCAGGGGTTGTTCGCGCAGGCGCG
CAGCCAGCGCGTTGGCGGCCGGATCGCCCGAGCGACCACCGCTGTGGTGCTGCATGCCGATATGAATCACACCGCCCAGGAAGGTGCCGGTACACAGCAC
CACGGTTTCCGCCATGAAGCGAATGCCGCCCTGGGTCAGCACGCCGCGAACCGTATCGCCCTCGACGATCAGATCGTCGGCGGACTGGGCAAAGATCGAC
AGGTTGGGCTGATTTTCGAGGATATAGCGAATGGCCGCCTTGTAGCGCACGCGATCGGCCTGTGCCCGGGTGGCACGTACCGCCGGACCCTTGCGGGAGT
TGAGGGTTCGAAACTGGATGCCCCCCTGATCGGTGGCCAGCGCCATGGCACCGCCGAGCGCATCGATTTCCTTGACCAGATGGCTCTTGCCGATGCCGCC
GATGGCGGGATTACAGGACATCTGTCCCAGCGTTTCGATGTTGTGACTGAGCAGCAGGGTCTGACAGCCCATGCGAGCGGCGGCCAGTGCGGCTTCGGTA
CCCGCATGACCACCACCGATGACAATAACGTCAAAGCGGTCCGGATAATCCACGTGAAAAACCTCGAGGGGTTGAAGATGAACGTGACACCCGACTGATC

AGCGCGTGTCGATACCCACATGCCCGAAGAGGGGCTGGGGATAAGGCGTATTGAATGGCGCGCATTATAACGTGGCTGTGAACAAGCGTCAGGGTTTTCC
ACGCACGGATTTTCACCCGACAGAAGGTCTTTAATTAATAAAACATCTTTTTTAAATAAGAAGTTCTGTTGTGGTTGTGATTACTGGTATTGAGCCTTTC
**TGTGGAAAA**GCATTTTTTCCGTAAATAAAACAGCTTTCTGGGACGTGAATGAAACGCAGGGCAAACTGCGGGTTTCCCTGGGGCAGGCCCTGTACGGGGT
GTGCATAACCCTGCGACTTGTTCACAGCTGAATTTATCCCCGCATGATCAACCGGGCTCAGGGAGGATTCTCCCCATACTCGTCAAGAAAGGCGTCATTG
TAGATTCGATCGTTGAAAAGCCCGTCACAGGCGGCTTTCAAAGCAGTGGGGCAGCCCGGGCGTCAAAAACTGTCCACAGGGGCAGGCGCGTTCCTGGTGC

AGGAGGAGGTAATCAATCGGGCATAAAAAAGGCGTCTGTCGAAAGCCGACAGACGCCTGGGCGTCATCGCGGGACGTCAGACTGTAAAGGATGCCCCACA
GCCACAGGTCGTGGTGGCGTTGGGATTTTTGATCAAAAAGCGGGCACCGGCCAGACCGGCTTCGAAGTCCACCGTGGAACCGACCAGATACTGATAGCTC
AGCGGGTCGATGACCATGGCGACATCGCCGTTTTCGATGATGGTATCGTCTTCGCCGACGGTTTCTGCAAAATCGAAACCATACTGAAAGCCCGAACAGC
CGCCTCCCGTTACGTAGACGCGCAGTTTCAGAGCCGGGTTGTGCTCTTCCTCGATCAGCGTCCTGACTCTGCCTGCAGCAGCATCGGTAAAGACAAGCGG
GGTCGGAACAAAGGATTCTGCTCCGCTCATCGCGGCCTCCTTCTGAGATGACACACATGAGATGTGTCAGGAATGGATCAGCCTTGTGACTGTCATGATC
ACCATTATCGGCACCCCCTCTGTCAATGGTCAACCGACAGGGCTCGGGAGATCATGGGGGCCAGCGGCATAAACAATGATAGGCTGATGCGCGTGGTGTT
TCATGACCTCGTCATTCTGCCGCTGGGGAGAGCAAGCTTGCCGCGCCGTTTTCTGGATACATCACGTCTGAATCTTGAAAATTTCCGACGCCAGCTGGCC
AGTGTCGACGCCCTGCCACAGCTGTGTCTGCTGGCGGTCATTTCCGGTGCGCTGACCGGACTGATCATGATCGCCTTTCGGGAGTTGATCGAGCTGGGGG
CACTGGCGCTGATGCCGGGCGGGGACGCCGAGGCCTTTGAAGCCCTGCCCGTGGCCGTTCGCACGGGACTGCCAATGGCGGCCGTGATCCTGATTGGCCT
GGGACTGGGCTGGCAGAAAAAAAGTGCGCGTCGCATCGGCATTACGCACGTGATCGAGCGCTTTTCCTACCATCAGGGCGTCATGGCTCGTGACAACTGG

>Kushneria avicenniae DSM 23439 [NZ_FOLY01000003.1:542792..542888], dnaA> ig dnaN>, 97 bp

WebSkew GCmin pos. 173236 (GCmin<>oriC1 66.9 kb; GCmin<>oriC2 6.4 kb; oriC2<>oriC1 60.5 kb)

>Kushneria aurantia DSM 21353 [NZ_KB907862.1:c106045..106544], <rpmH ig dnaA>, 396 bp

ATATCCACGCCGTTGACGGTCGCGCTCAGATCGACCTGTAGCTGGTTGCCGTCGCCGAGCTGATAGCTGTTCTGCGCCGCGCTGAAGTCGATGCGTCCCT
CATGCCCCTCGAGCTGCAGACCGGAGCGGGCGATGTAGCTGCGCTGCTGATTGTCGGAGAGCAGTACATAGGGGTCGCTGGCACCGAGCTGGCGCTGATA
TTTCGGCAGGGCGGCGTAGACGATATCGCCACCACGGGTGTCGATACGCAGATCAAGGGCATCGGTATCAACGCTGATCAGCCGCTCCGGCGCGCCTTCA
GCGGGTGCCTCGTCGCTGCCCGGCTGGCTGCCGCCCGGCTGTGGGGTGGTGCTGCCGCTGGCGGGGGCGTTGAGCCCGCCGCTGCCGCTGCTTTCGCCGC
TGCCTTCCTGCTGGGCCACGGTGGGGGGCGCATCGACGCCCGCCGGTTGCTGGCCATAATCGTTGCTCCACTGGATCACCAGCAGGTAGGCAAGCACGGC
CAGCGGAATGACGAAGAGAAGACGTTTCAAATCCATTGGGATTGCCCGGTGGGGAAAATGTCGACGCGGCGCAGACCACGGATGGCACGGGAAAAGAGAT
CGCGCCTCGTACGGCTATTGAGGTTTCGACTGACGCTGCACATCCTTTTCAAGGCGCCGCCACATGCCGTGAAGCTGCCGGGCCAGGGTTTGTGCTTCCA
GCTCGTGCACGCCGCGGCGAGAAAGCACCACAATATCCACCGCCGGCAGCCAGTGCTGGCGCAGCCGGATCGACTCGCGGGTCAGCCGTTTGAGGCGGTT
ACGATCCACGGCGCGCTTGAGATTCTTCTTGCTGAAAACCAGCCCCACGCGGACATGTCCCAGGTCGTTGGGCGTGGCGAGGGCCAATAACCCCTTGCCG
TGAACCTTGAAGCTGGCGTTGTCGAAAACGTGACGGTATTCCCCGCCGGTCAGAATGCGCAGACCGCGGGGAAATCCCTGTGAGGCCATGGGGCGCAGCT

TCCAGACGCCATTCAGGCGCTCAGACGCTTGCGGCCCTTGGCACGGCGACGTGCAAGAACCTTGCGGCCGTTTTGGGTCGCCATGCGGGCGCGAAAACCG
TGGACGCGCTTGCGCTTGAGCACGCTGGGCTGAAAGTACGTTTCATGATGGCACTTCCTGAATTGGCTACGAGAAAATTGATCGGTCGACCCGCGCCCT
GGCGTTAGGCGCACGGAGGTCAAGAGCCGGGATTCTAGAGAATTCAACCGGGACGTGCAATTTTTATCCCGACCGATCTGCAAGCCCCTGTGCATAAAAC
TGTGCACAGGCGGTCTATATCCTGCCGGCAGATAACGCAGGTCGCTGAAAGGCTTGTTTTATTGCTTCCGATCCGGTTGTCCACAGGCGTTGCGCGTCAG
GGAAAAGGTGTGGATAACCCCTGATGCGGACGTTACACTGAGCGGCCATTGGCAGCAGGAGGGCCCTGCCCGCCACGCTCACGATGCGCGGTGCGAAGGC

TCGGTGCACCCTCCTTCAAGACTCACCATCAACGAGGTCGCGTGTGTCCATCGCTCTTTGGCAGCAGTGTCTCGACTACCTGCAGGATGAGTTGAGTTCT
CAGCAGTTCAATACCTGGATTCGCCCCCTGCAGGCCGAGGGTGGCGACAGCGAGCACGAGCTTGCGCTGCTGGCGCCCAATCGCTTCGTGCGTGACTGGG
TCAACGACAAATACCTCAAACGGATCAACGAGCTGATGCGCGAGCTGGGCCAGGGGCGGGCCCCGCGCGTCAGCCTGGGGATCGGCAGCCGGCGCACGGT
ATCCAATCCGCGCGCTGCAAGCGCCGACGGGGCTGCTTCGTCCGGCGTCGAAAGCGCGCCGCCGGCGGCCGATATGGCGGCCGTTAAAGAGGCGCCCATG
CCGCCGCCGTCGCCCACCGCGCCGCGCGGTGAGTATCAGGACGAGCGCGAGATTCCTGCCGCTTCCGGCGGCAGCGCGACCGGGAGTTCGCGCCGTACCG
AGGGCGGCGAGCGCCAGGTGCAGGTCGAGGGCAGTATCCGCCACCAGAGCGGGCTCAACCCCGGCTTTACCTTCGAAACCTTCGTCGAGGGCAAGTCCAA
CCAGCTGGCGCGGGCCGCGTCGCGGCAGGTGGCGGAGAACCCCGGCGGCGCCTATAACCCGCTGTTTCTCTACGGCGGCGTGGGGTTGGGCAAGACCCAC
CTGATGCACGCGGTGGGCAACCACCTGGCGGGGCTGCGGGAGAACGCCCGGGTGGTCTATCTGCACTCCGAGCGCTTTGTCGCCGACATGGTCAAGGCGC
TGCAGCTCAACGCCATCAATGACTTCAAGCGTTTTTATCGCAGCGTCGATGCGTTGCTGATCGACGACATCCAGTTTTTCGCCGGCAAGGAGCGCTCCCA
GGAGGAGTTCTTCCACACCTTCAACGCCCTGCTGGAGGGCGGTCAGCAGATGATTCTGACCTCGGACCGCTATCCCAAGGAGATCAGCGGCGTCGAAGAG

>Kushneria aurantia DSM 21353 [NZ_KB907862.1:c166600..167099], <mnmG oriC2 <PRK13623 (ErpA), 964 bp

CGAGACCTTCCGGTTCGATGAAGACCTGGTGGCTCTGCTTGTCGGCGAAACGGTGGATCTTGTCCTCGATCGACGGGCAGTAGCGCGGCCCGGTGCTGTC
AATCACTCCGGAGTACATCGGCGAGCGGTCGAGGTTGGCGCGAATGATGTCGTGAGTGCGCTCGTTGGTGTGGGCAATGAAGCAGCTCACCTGCTCGGGG
TGCATATCACGGTGGCCCATCCACGACATGACCGGTCGCGGCGTATCGCCGGGCTGCTCGCCGAGTTTCGAGAAATCCACCGTTCTGGCGTCGAGCCGCG
GCGGTGTACCGGTTTTCAGCCGCGCGACACGCAACGGCTGTTCGCGCAGCCGTTCGGCCAGTGCGTTGGCGGCCGGGTCTCCGCAACGTCCGCCGCTGTG
ATGTTCCATACCAATGTGGATCACGCCACCGAGGAAAGTACCGGTACACAACACCACGGCGTCGGCCATGAAACGCACGCCACCACGGGTCAGCACGCCA
CGTGCGGTATCGCCTTCGACGATCAGGTCATCGGCGGCCTGGGAGAAGATCGACAGGTTGGGCTGATTCTCCAGCATGTGGCGAATCGACGCCTTGTAGC
GTACCCGATCGGCCTGAGCACGGGTGGCGCGCACCGCCGGCCCCTTGCGCGAATTGAGGGTACGAAACTGGATGCCGCCATGGTCCGTGGCCAGTGCCAT
GGCGCCGCCGAGAGCATCGATCTCCTTGACCAGATGGCTCTTGCCGATACCGCCGATGGCGGGATTACAGGACATCTGGCCGAGGGTTTCGATGTTGTGG
GTCAGCAGCAGGGTACGACACCCACTACGAGCGGCAGCCAGTGCGGCTTCGGTACCCGCATGACCACCGCCGATGACGATTACATCAAAGCGGTCGGGGT
AAATCACGTTGTCAACCTCGAAAAACGGGAAAGCAGCAGGCGATCAGCGGTAAGCGTCGGTCCTTTGGTCGGATCCGATTCCGTGAGCCCGCTGTTGGTA

GCGGTTATCGGTTGCCCACAGGGGTTTGAACCGGCTGTGGGCAACCTTCCTTGCGAATGGTGCGCAGTATAACGCGACTGTGAACAGGCGTCAGGGTTTT
CCACATTGGGCACCACACCTCATCGGGTTTCAATATTTAAAAACAGTCTTTTTAAATAAGAAGTTCTGTTGTGGTTGTGATTACTGGGGACGCTCGATTC
**TGTGGAAAA**GCCGATTTTTTGTAAAAAAATCAGTTGCTCACCATGTTGTCTCAACGGTTGTGAAACTGCGCGCGGTGCGGGCACAGTCTCTGCGAAGGCT
GTGCACAAGATCCCTTTTTACCCACAGGCGTTGCTGTCCCGGTTTTTTCAACCGGGCTGTACGGCAGTTGTGCCCGCAGGCGTTCAGTGCTGCCGGCAAG
TATCCGAAGCGTGCTGAAAGCCGCCGGTGACAGGAGTTGTAGGCAGAAAAGGAGAGGCCTGAAATTTTTTTTGTCCACAGGGGGAAAGCGGCATTCAGAC

CGGGAGCAGGAAGGCGAAAATAGCGTCAGGCGGACCTGGGGGGTGCCCGCCACGCAGTGGTGGCCCCAGGGGTGCCCGCCAGGGAGGGCAGGGCACAAAA
AATCGCCGGGAGCGATTTTTGACGCCGCAGGCGGCCCCAGGGGGGTGCCCGCCAGGGAGGGCAGGGCACAAAAAATCACCGGGAGTGATTTTTGACGCCA
CGCAGTGGCGGCCCCAGGGGTGCCTGCCAGGGATGGCAGGGCACAAAAAATCACCGGGAGTGATTTTTGACGCCACGCAGTGGCGGCCCCAGGGGTGCCC
GCCAGGGAGGGCAGGGCACAAAAACGCCTGCTCGAAGGCAGGCGTGTATTGAAGCGCTGACAGGCGGCGACTCAGACGGTAAAGGAGGCGCCGCAGCCGC
AGGTGGTAGTGGCATTGGGATTTTTGATCAGAAAACGCGCGCCGGCCAGGCCCTCCTCGAAGTCCACCGTGGAGCCGACCAGGTATTGATAGCTCAGCGG
GTCGATCACCATGGCGACATCGCCGTTTTCGATCACGGTATCATCGTCGTTGAGGTTGTCGGCAAAATCGAAGCCGTACTGAAAGCCCGAACAGCCGCCT
CCCGTCACGTAGACGCGCAGCTTGAGAGCGGCGTTGCCCTCCTCGATAACCAGCGCTCTGACTCGCGCTGCTGCGGCGTCGGTGAAGATGAGCGGTGTCG
GGACAAAGGATTGTGCTGTCCCGCTCATCGCGGCCTCCCTGTTGGATAGCGAACGGCACAGGGCCGAATGATGCCTCATTATCGTCAGCAAGGCGCCGAG
GGGTCAATCGCGGCAATTGATTCATGCGCGACCGACAATATCGGCGACGCCTGAAACCGCCAATAGTGATAGGCTTGTAAAGGCCATGGCCCGGACAGGG
CAGCGTCAGGGAGATCGGGTTTGCCACGCCTACCGCGTCGTATATCGCAGTGGAATCTGGAAAATTTTCGTCGCCAGCTGGCCACCGTCGATGCGCTGCC

>Kushneria aurantia DSM 21353 [NZ_KB907862.1:c104394..104480], dnaA> ig dnaN>, 87 bp

WebSkew GCmin pos. 67354 (GCmin<>oriC1 54.3 kb; GCmin<>oriC2 1.8 kb; oriC2<>oriC1 52.4 kb)

>Kushneria sinocarnis DSM 23229 [NZ_RBIN01000001.1:121382..121881], <rpmH ig dnaA>, 390 bp

GCAGCTGCTGCTTGTGCTGCGGCAATGCGGCATAGACGATGTCGCCGCCGCGCGGATCGATCCGCACCTGGAGCACATCGGTGGCGACACTGATCAGCGA
GCTGTTGTCCTGCTGACCGGCGCCGGAACCGCTGTCGGCCATGGCCTGGTCATCGGGCTGGCTGGCGCGCTCGGCGCTGCTGTCGGGGGCATCGAGGCCC
TGCTCGGAAGGCGCGGCATCGCTGCTCTGCTCGGAGAAGGCCGGCGTGTCAGTGGGCTGCTGCTGACCGTGGTAGTCCTGGTTCCACTGGATCACCATCA
GGTAGGCAAGCACAGCCAGCGGGATGATGAAGAAGAGTCGTTTCAGATCCATGAGGGATTGCCCGATGGGAAGATGTATCGGCTCGGTGACGGTGTCCGG
ACGGAACCGGAAGACGGCGCAACCGTGGCGTGATGCGGCCGGCAGGGGTACCCGGTAAAGGCGTGGGGCAGGGCGTCGGCCCGGCTGCCGAAGGGATGTG
ACGCTTGGATGGTAATGGCGTGATCTCCTCGAACCGGGCCCGGATTCACTCGGGTTTCACTCGGTCACGGCGATGCTGATCGCCGGGCGGCGACGAGTCG
GTCTGTTTGCCGACTTCCCTGCTCAGCCGCCGCCACATGCCGTGCAGCTGGCGCGTGATGGTATCGTTATCGAGCGATGAGGCATCGCGTCGCGCCAGCA
CGACGATATCCACCGCCGGCAGCCAGTGCTGACGATGGCGGATGGATTCGCGGATCAATCGCTTGAGTCGGTTGCGATCAACGGCACGACGTGCATTCTT
CCTGCTGATCACCAGGCCCACGCGTACATGGCCCTGCTGATTGGGGATGGCCAGTGCCAGAAACCCGCGTCCGTGAATCCGAAAGGCTGTCCGTTCGAAA
ACCTGCCGGTAATCCCCGCTGCTCAGCAACCGCATCCGGCGGGGAAACCCCTTGGGATTCATGGGGCTACCGCGATCAGGCGCTCAGGCGCTTGCGGCCC

TTGGCACGACGACGGGCCAGGATCTGACGACCGGCCTTGGTGGCCATGCGGGCACGGAAACCGTGATTGCGCTTGCGCTTGATAACGCTGGGCTGAAATG
TGCGTTTCATGGTGACATCTTCCGAATTGAAGCATGAGGAAATCGACTGGCCGCAGTGCTTGCCCTGACCATGGGCGCACGGGGGGCCGGGAGCGGCGAT
TCTAGAGAATTCGCGCCGGTGGTGCAATTTTTCTCTTGCCTGACCCCCGCTGACGACGCCATGGGGCAATCCATGTCGCGAAGTGCTGTGGATAAATCTC
TGGACAAGCTGCGGGCAACCCGGTGACAGATCTCCGAGAGCCCGCTTGCAGAGCCGCTGACAGGGGGCGCCCAGGGTTGTCCACAGCTCGTTCGGGGCTT
GGATCAGGTGTGGATAACCCGGCGGGGCAGCGCTACAATGCCTGCCCTCGCTCGCGTCGTTGTCGAGCGGCCATTTCCCGGATCACCACGGTGAGGCCAC

GTGTCGATCGCACTCTGGCAACAATGTCTCGAGTATCTGCAGGACGAACTCAGTTCGCAGCAGTTCAATACCTGGATTCGTCCGCTCCAGGCCGAGGACG
GTGAGGAGGCCAACGAGCTGTCGCTGCTGGCCCCCAACCGTTTCGTGCGCGACTGGGTCAGTGACAAGTATCTCAAGCGCATCAACGAGCTGATGCGCGA
ACTCTCCAGCGGCGGCAAGCCGCCCCGGGTATCACTGGGCATCGGCAGTCGCCGGACCGTCTCCAATCCGCGCATGGCGCCCGCCTCGAAGGAGCGCAGC
GCTCCGGCAGAGGCGCCGGCCCCGGCGCCGCCGGCTGCCCCGACCGCGCCGCGCGGCGACTACCGCGACGAGCGTGAGCTGGACTCCGCCGCCACGCCCC
GGCGCAGCAGCAGTGCCGGCAACAGCGGCAGCGGTGGCAGCGGCGAACGCCAGGTGCAGGTCGAGGGCAGCCTCAAGCACCAGAGCGGACTCAATCCCAA
CTTCACCTTCGATACCTTTGTCGAGGGCAAGTCCAACCAGCTGGCCCGGGCGGCTTCCCGCCAGGTGGCGGAGAACCCCGGGGGCGCCTACAACCCGCTG
TTTCTCTACGGCGGCGTGGGGCTGGGCAAGACCCACCTGATGCACGCCGTCGGCAATCAGCTGGCCGGCCTGCGCGACAACGCGCGGGTGGTCTATCTGC
ACTCCGAGCGCTTCGTCGCCGACATGGTCAAGGCGCTGCAGCTCAATGCCATCAACGATTTCAAGCGCTTCTATCGCAGTGTCGATGCGCTGCTGATCGA
TGACATCCAGTTCTTTGCCGGCAAGGAGCGCTCCCAGGAGGAGTTCTTTCATACCTTCAATGCGCTGCTGGAAGGCGGGCAGCAGATGATCCTGACCTCC
GATCGCTACCCCAAGGAGATCAGCGGCGTCGAGGAGCGCCTGAAGTCGCGCTTCGGCTGGGGGCTGACGGTCGCCATCGAACCGCCGGAGCTGGAGACGC

>Kushneria sinocarnis DSM 23229 [NZ_RBIN01000001.1:68928..69427], <mnmG oriC2 <PRK13623 (ErpA), 607 bp

TCCATCGAGCGTACCAGCTCGAGCTGGACATCGAACGGCAGCGAGGTCGAGATGCCGTTGGGGTAGAGCTCGGGGGTCTCCAGGCCTTCCGGTTCGATGA
AGACCTGATGGCTCTCCTTGTCGGCAAAGCGGTGAATCTTGTCCTCGATCGACGGGCAGTAGCGCGGCCCCACGCTGTCGATGGCGCCGGAAAACATCGG
CGAGCGCTCGATGTTGGCGCGGATGATGTCATGGGTACGCGTGTTGGTATGCGCGATGTAACAGCTGCGCTGGGACGGATGCATGTCGGGCGTTCCCATC
ATCGACATCACCGGACGCGGCTCGTCCCCCGGCTGGGCGGCCAGCTTCGAGAAATCGACGCTGCGGCCGTCCAGTCGCGGCGGGGTACCGGTCTTCAGCC
GATCCACCCGCAGCGGCTGTTCACGCAGGCGCCGCGCCAGCGCCGAGGCCGAGGGATCGCCGCTGCGGCCACCACCATGGTGCTGCTCACCGATGTGGAT
CACGCCCCCCAGGAAGGTACCGGTGCACAGCACCACGCTTTCGGCGAAAAAGCGGATGCCGCCCTGGGTGACCACGCCGCGGGCGGTATCGCCCTCGACA
ATCAGGTCATCCACCGACTGCTGAAACAGCGTGAGATTGGGCTGATGTTCGAGCGTGTGCCGGATGGCCGCCTTGTAACGGACCCGATCCGCCTGTGCCC
GGGTGGCGCGCACGGCCGGCCCCTTGCGGGAGTTCAGCACCCGGAACTGGATGCCCGCCCGATCGGTAGCCAGCGCCATGGCACCGCCGAGCGCGTCGAT
CTCGCGTACCAGATGGCTCTTGCCGATGCCGCCGATGGCGGGATTGCAGGACATCTGGCCCAGCGTCTCGATGTTGTGCGTCAGAAGCAGGGTCTGACAG
CCCGTTCGGGCGGCGGCGAGTGCGGCTTCGGTTCCGGCATGACCACCGCCGATGACGATGACGTCAAAGCGGTCCGGGTAATCCACGTATTGACCTCGAG

CACGTTGGCCACGGCCACGCCTGCCTGATCGGCTGCTGACCGTAACGGAAATGGGGAAATCAGGGTTCGCGATTATAACGCGCCTGTGGGTAACCGTCAG
GGGTTTTCCACACGAGCTCGCGTTTTCCTTCAATCAATCAATACAGGGTGTATAAAAAGGAAGTTCTGTTGTGGTTGTGATTATGGGGAGCCGCAATTTC
**TGTGGATAA**CCGATTCATTGCACCATGAATCAGTTATCTGGCGTGTTGATACGGTTGCGCACGAGTGGCGTATGAAGGGCGGATGACCGTTGCAGGAGCT
GTGGATGAAAGGCCTGGTTATGCACAGGGCAGCTTATCCCCCGAGTACTGCCCAGCCTGTCCCGATGGTTGTTACCGCACTCGTCCAGAGCAGGGCAGCA
GTCGGCCCCGGCAGCCACGAGCGCCGTGTGGCGTGGCTCGGAGCGACGCATGCAGGTGATGAGCGATGAATGTTATCCACAGCCATCGGAAGAGCGGATC

GGCGATCCGGATGGCGGATTCAGGGAGGGTGGACATGAAAAACCCCGGCACGCTGGCCGGGGTTCGAGGAGAAGTGATCAGGCTTTCGAAGGGTCAGACG
GTGAAGGATGCCCCGCAGCCACAGGTGGTAGTGGCGTTGGGGTTCTGGATCATGAAACGGGCACCCGCCAGCCCTTCCTCGAAGTCGATGGTCGAACCGA
CCAGGTACTGATAACTCAGCGGATCAACCACCATGGCCACCTCGCCGTTCTCGATCACGGTGTCTTCCTCGTCCACCGCTTCGGCAAAGTCGAAACCGTA
CTGGAAGCCCGAGCATCCGCCACCTGTCACGTAGACCCGCAGCTTGAGCGCCGGGTTGTGCTCCTCGGCGATCAGGGCCCGCAAGCGGCCTGCTGCAGCA
TCGGTGAAGATAAGCGGCGTCGGGACGAAGGATTCTGCCCCGCTCATCGCGGCCTCCCGAAGTATCTTGAATGAATGCCTGCATTATCGGACCCAGGCGC
AACCGGGGTCAACCTGGGTGGCCGCGGGAAAGGTCGTGCTCTTTGAATTGCACCGGGGGTTGTGGCAACAATGAAGGGTCGGCAGGGTGGCTGCCAGCGA
CAGCCGCCTCGTCAGCGTCAGGGATGTACGAGCAGCCGGCATGACATCCATTCCTGCGCCTGTCCGCTACCGGGGAGAACCCGACTTGCCGCGTTATCGT
CTCGGCATTACACGCTTCAGTCTGGATCATTTCCGCCGGCGTCTGGCGTCGGTGGATGCGCTGCCGCAGCTCTGTCTGCTTGCTGTGCTTTCCGGTGCCC
TGACCGGTCTGCTGATGGCAGCCTTTCGTGAACTGATCGATGTCATCACGCTGGGATTCATGCCCGGTGGTCGTTCGGACGATTTCGAGGCGCTGCCGCC
ACCGATCCGTGTGCTGCTGCCGCTGGCCGCGGTGGCCCTGATCGGGGTCGGCCTGTGGCGCCAGCCGGCCGCAGCGCGCAAGCTCGGCATCGGCCACGTC

>Kushneria sinocarnis DSM 23229 [NZ_RBIN01000001.1:123382..123464], dnaA> ig dnaN>, 83 bp

WebSkew GCmin pos. 3584136 (GCmin<>oriC1 40.2 kb; GCmin<>oriC2 4.9 kb; oriC2<>oriC1 35.3 kb)

>Kushneria phosphatilytica YCWA18 [NZ_CP043420.1:3624116..3624615], <rpmH ig dnaA>, 396 bp

AGCTGCTGCTTGTGCTTCGGCAGAGCGGCATACACGATATCGCCACCCCGCGGATCAATCCGTACCCGGAGCACATCGGTGGTGACACTGACCAGGGGGC
TATTGCCCTGCTGAGTGGCGCTGTCACTACTGCCCGCCATGGACTGCCCATCAGGCTGGGTGGCCCGATCGGCGCTGCTGTCAGGCGCATCAAGACTCTG
CTCGGAGGGTGAAGCGTCGCTACTCTGTTCGGAGAAGGCCGGCGTATCAGTGGGCTGCTGCTGACCGTGGTAGTCCTGATTCCACTGGATCACCATCAGA
TAGGCGAGCACAGCCAGCGGGATGATGAAGAAAAGTCGTTTCAGATCCATGAGGGATTGCCCGATGGGAAGATGTATCGGCTCGGTGACGGTGTCCGGAC
GGAGCCGGCAGACGGCGAAACCGCAGCGTGACGCAGTAGGCAGGGGTGACCGAAAACGGCATGAGACAGGGCGTCGGCCCGGCTGCCGAAGGGATATGAC
GCTTGAATGATAATGGCGTGATCTCCTCGAACCGGGCCCGGATTCACCTGGGCTTTAATCGGTCACGGTGATTCTGCTCATCCGGCGGCGCCGAATCGGT
CTGCTTGCCGACTTCCTTTGTCAGTCGCCGCCACATGCCGTGCAGCTGGCGCGTGATGGTGTCGTTGTCGAGCGATGAGGCATCGCGTCGCGCCAGTACG
ACAATATCCACCGCCGGCAACCAGTGCTGACGATGGCGCACCGATTCGCGAATCAATCGCTTGAGGCGATTGCGATCGACCGCGCGACGTGCATTTTTTC
TGCTGATCACCAGGCCCACTCGGACATGGCCCTGACGATTGGGTATGGCCAGTGCCAGAAACCCGCGTCCGTGAATCCGAAAGGCTGTCTGCTCGAAAAC
CTGCCGGTAATCCCCGCTGCTCAGCAACCGCATCCGGCGGGGAAACCCCTTTGGACTCATAGGGGTATTGCGATCAGGCACTCAGACGCTTGCGACCCTT

GGCACGACGACGCGCCAGAATCTGACGACCGGCTTTGGTGGCCATGCGGGCACGGAAACCGTGGTTGCGCTTGCGCTTGATCACGCTGGGCTGAAATGTG
CGTTTCATGGTGACGTCTTCCGAAATTGAAGCATGAGGAAATCGTTTGGCCGAAGTGCCTGCCCGAACAATGGGCGCACGGAGGCCCGGGAGCGGCGATT
CTAGAGAATTCGAGGCGTGGGTGCAATTTTTCTCTCCATGCCGTGTCCGATGCCGACGCAACAGGACAAGCCATGTCGTGAAAGGTTGTGGATAAGATTC
TGGATAAGCTGCGGGCAACCGGGTGACAGATACTCTGAAGCCCGCTTGCAGAGCCGCTAACAGGGCTTGTCAAGGGTTTTCCACAGCCAGGCGAAAGTGT
CACTCAGGTGTGGATAACCTTGCAGGGCAACGCTACAATGCCTGCCCTCGCTCGCGTTGTCATTTGTCGAGCGGCCATTTCCCGGATCATCACGGTGAGG

CCACGTGTCGATCGCACTCTGGCAACAATGTCTTGAGTATCTGCAGGACGAAATCAGTTCGCAGCAGTTCAATACCTGGATTCGTCCGCTCCAGGCCGAG
GATGGTGAAGACGCCAACGAGCTGTCGTTGCTGGCGCCCAACCGCTTCGTGCGCGACTGGGTCAGCGACAAGTATCTCAAGCGCATCAACGAGCTGATGC
GTGAGCTCTCCAGTGGTGGCAAGCCGCCTCGGGTCTCGCTGGGTATCGGCAGTCGGCGGACCGTCTCCAATCCGCGCCTGGCGCCCTCGGCGAAAGAGCG
CAGTGCGCCGGCGGAAAATCCGGCTCCCACACCACCCGCTGCGCCGACCGCCCCGCGGGGTGACTATCGCGATGAGCGGGAGATCGATTCCTCGGCCACC
CCGCGACGCAGCAGCACGAACACGGCCAGCGCCAACAGCAGCAACGGTAGCGGTGAGCGTCAGGTGCAGGTCGAGGGCAGCCTCAAGCACCAGAGCGGGC
TCAACCCCAACTTCACCTTTGATACCTTCGTTGAGGGCAAGTCCAACCAGCTGGCCCGTGCAGCTTCGCGCCAGGTGGCAGAGAACCCCGGTGGCGCCTA
TAACCCGCTTTTCCTCTATGGCGGGGTAGGTCTGGGCAAGACCCACCTGATGCATGCCGTAGGCAATCAGCTGGCCGGACTGCGCGAAAATGCCCGGGTG
GTCTATCTGCATTCCGAGCGCTTTGTCGCGGACATGGTCAAGGCGCTGCAGCTCAACGCCATCAATGACTTCAAGCGCTTTTATCGCAGTGTCGATGCGC
TGCTGATCGATGATATCCAGTTCTTTGCCGGCAAGGAGCGCTCCCAGGAGGAGTTCTTCCATACCTTCAATGCCCTGCTGGAAGGCGGTCAGCAGATGAT
TCTGACCTCCGATCGCTATCCCAAGGAGATCAGCGGTGTCGAGGAGCGGCTCAAGTCCCGCTTCGGCTGGGGGCTGACGGTCGCCATCGAGCCGCCGGAA

>Kushneria phosphatilytica YCWA18 [NZ_CP043420.1:3588794..3589293], <mnmG oriC2 <PRK13623 (ErpA), 601 bp

TTCCATCGAACGCACAAGCTCAAGCTGTACATCGAACGGCAGCGAGGTCGAAATGCCGTTGGGATAAAGCTCGGGAGTTTCCAGACCTTCCGGTTCAATG
AAGATCTGATGGCTGTCCTTGTCGGCAAACCGATGGATCTTGTCCTCGATCGATGGGCAGTAGCGGGGTCCGACACCGCCGATGGCACCGGAGAACATGG
GCGATCGATCGAGATTGGCCCGAATGATGTCATGGGTCCGGGCATTGGTATGCGCGATATGGCAGCTGCGCTGTGGCGGATGCATCTCGGAGGAGCCCAT
CATCGACATGGCCGGTCTTGGATCATCCCCCGGCTGCTCCACGAGTTTCGAGTAATCGATGCTGCGTCCGTCCAGCCGTGGCGGGGTACCGGTTTTTAAA
CGATCCACCCGCAGAGGCTGTTCTCGTAACCGTCGAGCCAGGGCCGAAGCCGAGGGATCACCGCTACGACCGCCACTGTGGTGTTCCAGTCCGATGTGAA
TTATTCCGCCGAGGAAGGTGCCGGTGCACAGCACCACGCTCTCGGCGAAAAAGCGAATTCCCCCCTGGGTGATCACACCGCGAGCGGTATCGCCTTCAAC
GATCAGATCATCCACCGATTGCTGGAACAGCGTGAGATTGGGCTGGTGCTCGAGCATATGCCGAATTGCCGCCTTGTAGCGAACACGATCGGCCTGTGCC
CGGGTAGCGCGCACGGCTGGCCCCTTGCGGGAATTGAGCACTCGGAACTGGATGCCTGCCTGATCGGTTGCCAGCGCCATGGCGCCGCCGAGAGCGTCGA
TCTCGCGCACCAGATGACTCTTGCCGATACCGCCGATGGCGGGATTGCAGGACATCTGGCCGAGCGTCTCGATGTTGTGTGTCAGAAGCAGGGTCTGACA
GCCCGTTCGGGCGGCGGCGAGTGCGGCTTCGGTTCCGGCATGACCACCGCCGATGACGATAACGTCAAAGCGGTCCGGGTAATCCACGTATTGACCTCGA

GCACGTTGGTGACAGCGCGTCAGTCTGAAGACAACGCACTGTAATGGAAATGGGGGAATCCGGGGACAGGATTATAACGTGCCTGTGGGTAACCGTCAGG
GGTTTTCCACACTTCAAGGCGTTTCCCTTCAATCAATCAATACAGAATGTATAAAAAAAGAAGTTCTGTTGTGGTTGTGATTATAGGGCAGCGCAATTTC
**TGTGGATAA**ACGCCTGGTTTCTCCACAATTCAACAACCTGACATGTTGATACGATTGCGCATGACAAGCGTATGACGTGCGAATGACCGTTGCAGGAGCT
GTGGATGAAAGGTGGGTTTATGCACAGGGCAACTTATCCCGTCACTGCTACCCAGCCTGTCCCGGTACTTGTTACCGCAGTCGTCCAGAACCTGGGTAGT
GAAGGTCACCATGGCGTAAACACCCGTATGCCATGGGTTTGACCCCGGCAGGCGCGTGCTTTTGGTGTCCGGTTGTCCACAGCCGGAAAAAGTGGAAAAA

ATCGTCTGGAAGGGTCAGCAGCGTGGAGAGCGCGCATGAAAAACCCCGGCACGAAGACCGGGGTTGAAGGCGGGAAGAGAGTTTGAAATCAGACCGTAAA
GGAAGCGCCACATCCACAGGTGGTTGTCGCGTTGGGGTTCTGGATCATGAAGCGTGCCCCCGCCAGACCCTCTTCAAAATCGACGGTCGAGCCAACCAGA
TACTGATAGCTGAGGGGATCGATCACCATGGCTACTTCACCGTTTTTGATGATGGTGTCTTCCTCATCCACCGTTTCAGCAAAGTCGAAACCATACTGGA
AGCCCGAGCAACCGCCACCTGTCACATAGACCCGCAGTTTGAGCGCCGGGTTGTTTTCCTCGGCGATCAGCGCCCGTAGGCGGGCAGCTGCGGAATCGGT
GAAGATCAGCGGTGTCGGGACGAAGGATTCTGCCCCGCTCATCGCGGCCTCCCGAAGCATTCAGAATGAATACTGATATTATCGGATGCGGTTGCAACCG
GGGTCAACCACGCTGGCCGAATGGGTCAGTAGTGCTCTTTGAATTGCAACAGGGGTTGTGACAACAATGTGGGGCCGTTGTGGAAGCCAGGTGAGCGACC
AGTGCATCGAGCATTCGGTCCGAGCTATTGCCGGTTTATCCGTAACGGCGCTCATCCGCCATTGGGGAGAGCCCGACTTGCCGCGTCATCGTCTCGGCAG
TGCCTTTTTCAACATGGATCAGTTCCGCCGGCGTCTGGCGTCGGTGGATGCTCTTCCGCAGCTCTGTCTACTCGCCGTTTTCTCTGGCGCCCTGACCGGG
CTGCTGATGACGGCTTTTCGTGAACTGATCGATGTCATCACTCTGGCATTCATGCCGGGGGGGCGTTCGGATGATTTCGAGTCCCTTGCGCCCGTCATGC
GTATGGCGCTGCCGCTGATCGCGGTCACGCTGATCGGGGTGGGGCTATGGCGCCAGCCGGCAGCCGCGCGTAAGCTGGGCATCGGGCATGTGATTGAACG

>Kushneria phosphatilytica YCWA18 [NZ_CP043420.1:1507..1589], dnaA> ig dnaN>, 83 bp

WebSkew GCmin pos. 646464 (GCmin<>oriC1 26.7 kb; GCmin<>oriC2 0.4 kb; oriC2<>oriC1 27.1 kb)

>Phytohalomonas tamaricis R4HLG17 [NZ_PVRV01000019.1:c619491..619990], <rpmH ig dnaA>, 458 bp

TGTGTTGGTGTCCGAAAGCAGTACAAAAGGACGATCCGATTCAAGGGTTTGCTTCTGCTGGGGCAGTGCGGCATAGACAATATCGCCACCGCGCGGATCG
ATACGCACATCAAGCACATCGGTATTGACCGCAATCAGCTGAGTATTTTGATTCGGAGTGTCTGGCGTGGACGGAGTAGAAGCACGCTCTTGTGAGGAAT
CAGGTGCTTGAAGCCCGGAGGGTTGTGGTGCGTCGCTACCCTGACTGGAAAAAGCCGGGGTGTCAGGCTCATTATGCGGCTGGTTATAGTCCTGGTTCCA
TTGGATCACCAGTAAATAAGCCAGGATCGCCAATGGAATCACTAAAAGGAGTCGTTTCAGATCCATGTTGTTGTGCCCGGTGGGGAGTGTGTCGGGTCGC
TTTAAATCCCGGCCGGGCGTCACCTGCAAGCAGCGTGCTCACCTATCGGCCGGTACAACACCAGACATGCGCCAAAGCCTTAAGCGATGGACTGCGTTGC
AGCCAAATGACGCTTCGCTTCCTGCTCCAGGCGCCGCCACATACCATGCAATTGACGTTGCACAGTATCATTATCCAGCTCTCCTGCTCCGCGTTTTGAC
AAGACTACAATGTCTACGGCAGGTAGGCGATGCTGACCGAGGCGGATGGATTCGCGTGTCAGGCGTTTTAGACGGTTACGGTCGACGGCGCGGCGTACGT
TCTTCTTGCTGAACACCAGTCCGACACGGGCATGGCCCAGCGTATTTGGACAGGCGAGCGCCAATAGGCCCTTGCCGTGGACCTTGAAACTTGCATTATC
GAAGACGTGCCGGTATTCCCCGGCGGTCAGGATTCTTAGACCGCGGGGAAATCCCTGATGGGCCATTGAGAGACCCGATCAAGCGCTCAGACGCTTACGG
CCTTTCGCACGGCGACGTGCCAGCACCTGACGGCCATTTTTAGTGGCCATTCGGGCACGAAAGCCGTGGACGCGCTTGCGCTTCAGAACGCTGGGTTGGA

ATGTGCGTTTCATAGCTGCTATTTCCCAGATGAAAAGATGATAAATCGGTCGCAGAGCTACCCTTTCAACGGGCTGGCTCTAGTCAAGAGCGGGAATTCT
AGAGAATTCGCCTGCACTGTGCAATTTTTCTTGACGTAATAAGCATTTTAAAATGCCTGGCTCGAGCAAGGCATCAGCCGTTACTTTAATCAATGGTGTG
CTAATGCAGTAGGTTCGCCCCATTATCGAGTGTTGATATAGGGCCTAAAGGCCGCTTTGACAGGACATGACGTTGTCGCGGCTACTGTGGATAACATTAT
GGATAAGCTGAGCGAAAAGCGGTGATAACCATAATGCCGCGGGTGAAATAATCCGGCTATTAGTAGGAAGTAAAGGTTTTCCACAAGGGGAAATAGAAGT
GCCTAATGTGTGGATAACCTTTGCCTTCATCGCTACAATAACGCCTTTGCTTATCTTGATGGCGAGGGCGTGTGTCGATCGCAATCTGGCAGCAATGCCT

TGAATATCTTCAGGACGAATTGAATTCGCAGCAGTTCAATACTTGGATCCGTCCTTTGCAAGCTGAAACGGTGGGTGACAGCAACGAGCTATCTTTGCTG
GCGCCTAATCGTTTTGTACGCGACTGGGTCAATGATAAGTATCTTAAGCGCATCAATGAACTGCTCAGAGAGCTTGCACCAGGCAAGCCGCCTAAAGTCA
GCTTGGGGATTGGTAGCCGGCGTACAGTGTCGAACCCGCGTGTGACCAATACGCCCAACACGTTACCCACCATGCCGCAGACGCCACCGCGTATGCCAAC
AGCCTCTGGGCATGGATCAATGCCACGTAACACGTCCTTCAATGGTAATGAGACATCCGATTCGGATGGCCAAGGCAGTGGAAATACGCCACGTCGTGGT
AATAGTAATGAACGCCAAGTACAAGTCGAAGGTAGCCTCAAGCATCAAAGTGGACTGAATCCCAACTTTACGTTTGAAACCTTCGTCGAAGGTAAGTCCA
ACCAGCTGGCACGCGCTGCATCTCGACAGGTAGCTGAAAATCCTGGCGGCGCCTACAACCCACTTTTTTTATATGGCGGAGTAGGGCTAGGCAAGACACA
CCTGATGCATGCGGTGGGCAATCATTTAGCTGGGCTGCGTGATAATGCACGGGTCGTCTATTTGCATTCCGAGCGTTTCGTCGCGGATATGGTCAAGGCG
TTGCAGCTCAATGCCATTAATGATTTCAAGCGCTTCTATAGAAGCGTCGATGCATTATTGATCGATGATATTCAGTTTTTTGCTGGCAAGGAGCGCTCCC
AGGAGGAGTTCTTTCATACCTTCAATGCCCTGCTTGAAGGCGGTCAGCAAATGATTCTGACTTCTGACCGCTATCCCAAGGAAATCAGCGGTGTCGAGGA
GCGTTTAAAATCGCGCTTCGGTTGGGGATTAACGGTGGCGATTGAGCCGCCCGAACTTGAGACCCGTGTCGCGATCCTGATGAAGAAGGCCGAACAGGCC

>Phytohalomonas tamaricis R4HLG17 [NZ_PVRV01000019.1:c646586..647085], <mnmG oriC2 <PRK13623 (ErpA), 623 bp

TGGCCGTGTGATATGAGCATTTTCAAGGCCACCGATCGAACGAACCACTTCAAGCTGCACATCGAACGGCAGCGATGTCGAAATACCGTTGGGATAGAGC
TCATGGGTATCAAGCCCTTCCGGTTCGATAAAGACCTGATGACTCTGCTTATCGGCAAAACGATGGATCTTGTCTTCAATCGAGGGGCAATAACGTGGAC
CGATACCCTCGATGACCCCGGTATACATCGGTGAACGATCAAGATTGGCACGAATAATATCGTGAGTTCGCTCATTGGTATGGGTAATGAAACAGCTTAC
CTGAGCGGGATGCATCGCTTGATTGCCCAGGTAAGACATCACTGGCTGCGGTGTATCACCAGCCTGTTCATCCATGACCGAGAAATCAACTGATTTGGCA
TCGATGCGAGGCGGTGTGCCTGTTTTCAACCTTGCCACGTTGAAGGGCAGTTCTCGCAATCGTGCTGCCAAGGCATTCGAAGGAGGATCGCCTGCACGCC
CACCAATGTGTTTCTCAAGGCCAATATGAATTACACCACCTAGAAAAGTGCCCGTGCATAGCACAACGGTTTCGCTTTTAAACCTTATCCCACCCTGGGT
AACAACGCCGCGTACGGTATCGCCTTCAACGATGAGATCATCTGCTGCCTGCTGGAAAAGCGTCAGATTAGGCTGGTTCTCCAGCATCCCACGTATTGCA
GCCTTGTAACGCACACGATCGGCTTGAGCACGAGTGGCGCGTACTGCTGGCCCCTTACGGGCATTCAGAACACGAAACTGAATGCCACCATGATCAGTTG
CCAGCGCCATTGCGCCACCAAGTGCATCGATTTCTTTAACCAGATGACTCTTGCCAATGCCGCCAATTGCGGGATTACACGACATTTGACCAAGTGTTTC
GATGTTATGTGTCAGAAGAAGGGTCTGACAACCCATGCGGGCGGCGGCAAGTGCGGCTTCGGTTCCTGCATGACCACCGCCGATGACGATAACGTCAAAG

CGGCTTGGATAATCCACGTTGTACCTCGGAAACACGACGGTGTTGGGGATAAAGAGGAATAGAAAGGCCGGCATTATAACGTGCCTGTGGGTAACCGTCA
GTGGTTTTCCACAACTCTTTTTTAACTACTAATAAATAAAGAAATAGTTTTATTTAAAAGAAGTTTTGTTGTTGTAGTGATTATTAGGGAGACTGAAAAC
**TGTGGAAAA**GAAAAATTAGTTGTTTATTAACATTATGTTGGGCTGTTCACAATTATGTATAAATAGCTTGCATAGCCTGCGTAGTAAGAGAGGACAGGCT
GTGGATGAAATGCGCATTTATCCACTAGCGTATTTTTCCATAGCTTGTTAAGAAATGCATACCTCAGTTGTTACCGTAGTCGTGCACGTTTGTAATAAAA
GAAGGCAGGCGGGAGATAGCGCGGGTTATGGCGTTAAAAAGAGGGGCAGCAGGAAAAATTTATCCACAGGCACAAAAAATATTTTTTATGCGCCTGAGAC

GGTTTTACACCCAATACGTCGGCTAATTGTCATCACCAACGGGTACAGATGCGCATGGAAGCATCAGGAAAGATAGGTATAGACATGAAAACGCCCCACA
AGAAGGCGGGGCGTAAAAGCGGTTATTCAGATTGAACTGATCAAACAGTGAACGACGCACCACAGCCGCAAGTAGTGGTGGCATTTGGATTGTGTATCAA
GAAACGTGCTCCCATCAGTCCTGTTTCGAAATCAACCGTTGAGCCGACAAGATATTGATAGCTCAATGGATCAACAACCAGAGACACATCGCCATTCTCA
ATAATGGTGTCATCGTCATTGATGGTTTCGGCAATGTCGAAACCATACTGGAAGCCCGAACATCCGCCACCGGTCACATAAACGCGCAGTTTAAGTGCTG
GGTTGTTCTCTTCAACGCTTAGCTGGCGAACGCGGTTCGCAGCAGCATCGGTGAAGATGAGCGGTGTCGGGACGAACGTTTCAGCCCCGCTCATCGCAGC
CTCCTATAGAAGATATTGTAAAGATGTTTGGTTGAAGTACCGACCATTATCGGCAAACCCCACTATTTCGGTCAAATAAAGCACTTTTTTCTGAATTGCA
GTAATCGTTACCGGCTATCAGTGGATTGAGCAATGCGCTTGATGCCGCATTAGCAAGGAAACTATTGCTGTAGGGTAGCAGGTTGTTGGACGTGATCAGT
GCTTACCCGCTTATACTATTATATTTAACGCTGTTGACCTTGTGCTCCAGGAGTCCGCCTTGCCGCGCCGCCGTCTGAGTGCTTCGCTTTTCAATCTCGA
TAACTTTCGCCGTCAGCTTGCTAGCGTCGATGCCTTGCCACAGCTTTGTCTTCTTGGTGTCCTTTCCGGGCTTCTGACCGGCATATTGATGGTGCTGTTT
CGACAATTGATTAGTTTTGGCGCATGGCTGTTCATGCCTGGCGGGTATTCTGAAGCATTTGAATCCATGCCCACGTATTGGCGTGCCACCATGCCGTTGA

>Phytohalomonas tamaricis R4HLG17 [NZ_PVRV01000019.1:c617945..618025], dnaA> ig dnaN>, 81 bp

WebSkew GCmin pos. 198755 (GCmin<>oriC1 46.7 kb; GCmin<>oriC2 1.2 kb; oriC2<>oriC1 45.4 kb)

>Carnimonas nigrificans ATCC BAA-78 [NZ_JAGO01000013.1:c151826..152325], <rpmH ig dnaA>, 494 bp

GAGTGTTGCTCGGCTTGCTGGCTGGTCTGCTGTTCCTGATCTGCCTGGTGGTAATCCTTGTTCCACTGGACCACCAGCAGGTAGGCGAGCACTGCCAGCG
GAATCAGGATGATGATGCGTTTGAAGTCCATGTAGGAATATGCCCGGTAAGAAGTGGCTTGGCGAATGAGCGCTTTGCCATGCTGAGGGAGAGCAGCCCT
GACTATATATGGGTGCGCTCGACAGCATAGCCGGCGATCAGCGTGCCATGTTATCGACCATCCCTTGGCGCCTCAAATGAAACTATTGGCGATAGGTACA
AAACACCTTTTACGTGTGTTCTGCAGGCAGAAGATGGCGGGAAGAAAGGCTGCTTTCGCCTACCTTGTTACTGCATTGCTGTCAGCGATTGTCGTATCTA
CCGCCATAGGGGGAATTACGGCAACTGACGTTATTCATCGAGCCAGCACGACCGAACCGCTGCATTCACACAGAATGAGCGACTTTCTCTGGTGAAGGCG
GGGCGGTCTGGCTGTGCAAAAGTGTGTTGGGCGAAAAAGCAAAAGCCGTCAGGCTGACGGCGCGGTAGCGAGAGTGCGCTTCTGCGCCTCTTTCTCTAAC
CGTCGCCACATGCCTGACAGCTGCTGCGCCAGTGTGGTTCTGTCCAGCTCGGTTACCCCGCGACGGGACAACACCACAATATCAACGGCCGGCAAGCGGT
GCTGGCGCAGGCGAATGGATTCCCGGGTCAGGCGCTTGAGCAGGTTGCGATCCACAGCGCGTTTCATGTTCTTACGACTGAACACCAGGCCCACTCGTAC
ATGTCCGCAGCTGTTGGGGCACGCCAGTGCCAGAAGCCCTTTACCGTGAATCTTGAAGCACGCTGCCTCAAAAACATGCCCATATTCCCGACTGGACAGT
ATTCTCATACTGCGCGGGAAGCGGCGATCTGCTGTTTTCTCTGCCATAGAGAGGCACATGCCCCTTCGCTTTACAACACTTAGGCGCTCAGACGTTTGCG

GCCTTTGGCACGACGACGAGCGATGACCTGACGGCCGCGTTTGGTCGCCATGCGAGCACGGAAACCGTGGTTGCGCTTACGTTTCAGTACGCTGGGCTGG
AATGTGCGTTTCATTGCGTCTTTCTCCTCAGCTAGGAAGGGAAGGATGGAAATATTCCGAGAGCAAAAGCGCCCTTCAGGAATGTCAAGAGCGGGGATTT
TAGAGAATCCAACGCAGCGACGCAATTTTTCTATGCATTATCGCGCGCATTTAGAGCAATTTTTTCATTCTTATTGTGCAAAATGCTGTGAGTAAGCTGG
TGGTAACCTATTGGCAAGCTGCTCATTTCATGTTCAAAAGTGGATAAATGCAGATTCTTCCACTTTTTTGCTTATGTTATCCACACCTGGATGATGTTGT
GCGGGTGTGGATAACTGGTGTCAGGGTCGATACACTAGCGGTTCCGCTCTGATTTTTACAGCGCAGGTGCGGCATTTATGCGAAACGTCGCCGATTCAAC

AAGGTATGCGCGGTGGGCGGTTCTCCTGAGCGCGACGGTATAACAGTGGGATATGTTAGCGTGGCGACAGGTACAGATCACTGGCAGTTATCAGGATGAA
GAATTAGCGTGTTGATTGCACTATGGCAGCAATGCCTTGAACGTCTGCAGGATGAGCTAAGCTCCCAGCAGTTCAATACATGGGTACGTCCGCTTCAGGC
TGAAGAGGGCGAAGCCGGCGATTTGGTGCTCTTGGCGCCCAACCGCTTCGTGCGCGACTGGGTAAACGATAAGTTTCTCAAGCGTATCAACGAATTGTTG
CGTGAACTCGCGCCCAGCAAGCCGCCCAAGGTAAAACTGGGCATCGGCAGCCGCCGCAGTGTTTCCAATCCTGCTGCTGTGCAGCGCCCTGCCCGACCAG
CGGCGCCTGAAGCGCCCGCCGTTATTCGCAGTCGCAGCCGTCCTTCTCGTGCTTCCTTCTCTGAAGATGTCGATGTCCCCGCCGCACCTCCCGAAGAGTC
GTCAGCGTCGCAGGACAAGGGACGAGATGTGCAGGTGGAGGGTAGCCTCAAGCACACCAGCGGTCTTAACCCCTCCTTTACCTTTGATACCTTCGTGGAA
GGTAAATCCAACCAGTTGGCCCGTGCCGCCTCGCGTCAGGTTGCCGAGAATGCTGGCGGTGCCTATAACCCGCTGTTTCTGTATGGCGGTGTGGGCCTGG
GTAAAACGCACCTGATGCACGCGGTGGGCAACCACATGGCCGGGCTGCGTGAAAATGCCAAGGTGGTCTATCTGCACTCCGAGCGTTTTGTGGCGGATAT
GGTCAAGGCGCTCCAGCTTAACGCCATCAACGATTTCAAGCGTTTTTACCGCAGCGTTGATGCGCTGCTGATTGATGATATTCAGTTTTTTGCTGGCAAG
GAGCGCTCTCAGGAGGAGTTCTTCCATACCTTCAACGCGCTGCTTGAAGGCGGCCAGCAGATGATTCTGACCTCGGACCGCTACCCGAAGGAGATTTCGG

>Carnimonas nigrificans ATCC BAA-78 [NZ_JAGO01000013.1:c197278..197777], <mnmG oriC2 <PRK13623 (ErpA), 855 bp

TGGGGTACAGCTCGTTGGTTGATAGTCCTTCCGGCTCGATAAACACCTGGTGGCTGTGCTTGTCGGCAAAGCGGTGAATCTTGTCCTCGATGGAGGGGCA
GTAACGCGGCCCAACGCCATCAATAACGCCTGAATACATGGGCGAGCGGTCAAGGTTGGCCCGGATAATGTCATGGGTGCGCTCATTGGTATGAGCGATG
AAGCAGTTGACCTGCTCAGGGTGCCATTCCTGCTGGCCCAGATAGGACATGATCGGTGTTGGCGTGTCGCCAGGCTGTTCCTGCAAACGCGAGAAATCGA
CACTTTTGGCGTCGATGCGAGGAGGCGTGCCGGTCTTGAGCCGCGCCACATTAAACGGCAGTTCACGTAAGCGTTGGGCAAGCGCAGTGGCACTCGGGTC
CCCTGCACGACCGCCGCTGTGGTGCTCAAGGCCGATATGGATGACCCCTCCGAGGAAGGTACCGGTACACAGCACCACGGTGTCAGCATGAAAACGGATG
CCGGCCTGTGTTTCGACACCGCATACCCGATCACCCGCCACGATCAGATCACTGGCCGCCTGCTGAAAGATTGCCAGGTTCGGCTGGTTTTCCAGCATTT
CACGAATGGCCGCCTTGTATAGCGCACGATCAGCCTGGGCACGGGTAGCGCGCACCGCTGGCCCTTTACGTGAATTGAGTACACGAAACTGGATGCCAGC
GCGATCGGTAGCCAGTGCCATGGCGCCGCCCAGTGCATCCACTTCCTTGACCAAATGACTTTTACCAATCCCACCGATGGCGGGATTGCAGGACATCTGG
CCCAACGTTTCCACGTTATGGGTCAGCAACAGGGTATTAACGCCCATGCGAGCGGCGGCGAGAGCGGCTTCAGTGCCGGCGTGTCCACCGCCGATGACGA
TTACGCCAAAGCGGTCAGGGTAATCCACGGTACAACCTCGGATAGCAGAACGCCCGACACGCTTTCAGGTGGACGCTTCCGTTAGCGTCAATGTGCCAGG

CGAAAAATAGTGTGAAGGTGCTGAAGCGTCAGCACGGTGTTGATCAGGGCGAGCAGTATAACGTGCCTGTGGGTAACCGTCAGTGGTTTTCCACACTGAT
CTCTGAAGCGCGCATTCAAGAGCGTTTATCTAAATAAATAAAAAAGAATGTTTTTAAATGAAGTTCTGTTGTGGTGATGATTATTGTTGCGGCGATTTTT
**CGTGGAAAA**CTCAACTTAAACGATAAAAATCAGGCAGTTACGCTGTTATCTTGTGGAGGGATAAGAGGCGTATGAGCTGCGCAGTTCTGGTGGGTAGCCT
GTGGAAGAAAGTACCGGATATCCACAGCCAGAGTTGTCCCCTGTTTATCCACCCGTGGCTGACGTTGTTGTCTGTGTAGTTGTTAACGTTTACGTCAAGG
TCGCAGGGCGCCAGAAATGGCGGACTCCCTAGCCCGATTTGCAGAGGATAACCGTAAGAGTTTTCCACAGCCATAATGGGCAGTAAAGCGATGTCTCTAT

CAAGGCGGGAAAAGACTCTGCAACCCGCATGGTTGTCAGCTTTCAGGCGACTATGCGGTCTCTTCTGAATGAAATGTGAAAATGCAGGTTTCGTCACTCA
AACTGGGCATTGATTGGCTGTTCAAGAGAGTAATGCGGCGGTGTTGACGATCGTTATGGCGATGCAAGAGAAAGGAAGCAGACTACTTTACAGGGGATTT
TTAGCAGTTGGAGGGAGTACAAACGACAATGCCGCCGCTGGAGGAAAACCCCGCGACGGCATTGGCAAGGAGGGGGGCAAATATCAGACGCTGAAGGATG
CGCCACAGCCACAGGTAGTGGTGGCATTGGGATTTTTGATCAAAAAGCGTGCACCTGCCAGACCTTCTTCGAAATCCACGGTAGAGCCGACCAGATATTG
ATAGCTAAGCGGGTCGATCAGCATAGTGACGCCATCTTTTTCTACGATGGTGTCCTCTTCATCGACGGCTTCGCTGAAATCGAAGCCATATTGAAAGCCA
GAGCAACCGCCACCGGTGACATAAACACGCAGCTTCAGGTTCGGGTTATTCTCGGCTTCTGCGAGGGTTCGCACGCGTGCGGCCGCACTCTCGGTCAGGT
AAAGCGGAGACGGCACGAAGGATTGGGCGCCACTCATGGGTAAGAGCCTCCTGGTATCAACCTGCTACTGCTGTTTGCCTGGTTATGTGAACGCTTCTCA
CTGAGTGAGTCGCGCAAGCCTGGCAGTGGCAGGTTCTATTGGGTCATTTCACGGCCTGTTGGCAACGCCGTTGGTACGGTTGATTGCGTGAAGCCTGGAG
AGCGATGTGTGACTGCGCCCCTCTGCGTTACGGTAAAAACGTCTACGGCAAAAGGATCAGCGGCACCGCATGGATGAGTGCTAGTGTACCATTTCGATCA
AGAGGGCGGCGATGTCGGGAAAAGAAGGCATACTCGCGGCCTGACTCGTGGTGTTGTGTCGGGTGTAATGATGTAGCGGTGGGGCATGTTGAGGAAATCC

>Carnimonas nigrificans ATCC BAA-78 [NZ_JAGO01000013.1:c150138..150250], dnaA> ig dnaN>, 113 bp

WebSkew GCmin pos. nd (GCmin<>oriC1 nd; GCmin<>oriC2 nd; oriC2<>oriC1 nd)

>Halotalea alkalilenta DSM 17697 [NZ_KK211226.1:c305671..306170], <rpmH ig dnaA>, 428 bp

ATGTAGCTGCGCGAGCTGCTGTCGGAGAGCAATACGAACGGCGTCTCGGAGTTGAGCGCCTGCTGCTGTTCCGGCAGTGCGGCATAGATGATGTCGCCGC
CGCGCGGGTCGATGCGCAGGTCGAGCACGTCGGTGTCGACGCTGATCAGCGAGCTGGAGGCGTTCTCCTGCGCTGGGTTGGCGCTTTGCCCGTTCTCCAG
CGGAGAGGTCGCGCGCTCGCTGGTCGATTCGGGCGCGTCGAGCACATCGCTGGGCTGCGTCGCGTCGCTCGAGTTGGCGGCGAGCGTCGGTGCCTGCGGC
TCAGGGGTCGGCTGGTGATAGTCCTGATTCCACTGTACGACCAGCAGATAGGCGAGGATCGCCAGCGGAATCAGGATGATGATGCGTTTCAGATCCATGA
GGGATGAGCCCGGTGAAAGGGTGTCGTGGCGTATCTTTGATGAATGCTGAAGACGGCTCGGCGCGTCGTCTCGAGCGGGAGGCCTTTCACCGGCGGCGTG
CGCGGGGAGATGTCTCCCTTTGCACTCCATCCCTCGCGCCGGGAACGGGCAGGGACGGGATCAGGCGGGTTCGCGATCAGGCGCGGGGGCTCGTCGCTTC
ACGTCCTGCTCCAGGCGGCGCCACATGCCGGCAAGCTGCCGACTGAGGGTCTCCCGGTCGAGTTCGAGAACTCCTCGGCGCGACAGTACCACGATGTCGA
CGGCTGGCAGCCGGTGCTGGCGCAGGCGTATCGACTCGCGGGTCAGGCGCTTGAGCAGGTTGCGATCGACGGCGCGGCGCACGTTCTTGCGGCTGAACAC
CAGGCCCACTCGAACGTGACCATGCTGGTTCGGGCAAGCGAGCGCCAATAGCCCCTTGCCGTGAACCTTGAAGCACGCATGCTCGAAGACGTGGCCGTAC
TCCCCGCTGGTCAGGATTCGCAGACTGCGGGGAAAGCCGTGGATCGCGTCCTGACCGGCCATGGTGGCCTCTCTTGAAATCAGGCGCTCAGGCGTTTACG

GCCTTTGGCGCGACGACGAGCCAGGACCTGACGGCCGCGTTTGGTGGCCATGCGAGCACGGAAACCGTGGTTGCGCTTGCGCTTCAGGACGCTGGGCTGA
AAAGTGCGTTTCATGACTGCTTTCCTCGAGACGGAGGGAGTGGTGGAGAGATGGAAAAACGCCGGTGCAAACACCTGCGGTCCCGTCAAGAGCGGGAATT
CTAGTGAATACGCGCAGGCGGCGCAATTTTTCTCTTACCAAGTAGGATTTTTTTTGCTAAGGGAGCCTTCGGCTGTGTGCAAAAAGCTGTGGATAGTCGG
GGGGAAAGCTATGGGCAAGCTGGGGAGTGGCTGTGTCATAGGGCTTGGAAGCAGTTTCAGGGCGCTTTTTACCTGAGTTATGCACAACCTCTTCAATCGC
AGAGATGGTGTGGATAACTCCTGCGGCCCCCGATACACTAGCGGGTCCGTAATATAGCCATTGAGTCCGAATCGCCTTCCGCCCTTCCGGCGCCGGAAGT

CCGCGTGCATGGCCCTCCACTTCGCAAGGATGACGAGTTCGCGTGTCGATAGCACTCTGGCAGAAATGTCTCGAATACCTCCAGGACGAGCTGAACTCCC
AGCAGTTCAACACCTGGATACGTCCGCTGCAGGCTGAGGAGGGCGAGGCCAATGGCCTCGCGCTGCTCGCGCCGAACCGTTTCGTGCGCGACTGGGTCAG
CGACAAGTACCTGAAACGGATCAACGAGCTGCTGCGCGAACTCGCGCCCGGCAAGCCGCCCAAGGTGTCGCTCGGGATCGGCAGTCGGCGCAGCGTCTCC
AACCCCCGGCCGACCGAGCCGGTGCGCCCGGCCGCGCCGATGATGCCGCGCAGCGCGCCGTCGCGGGCGTCGTTCTCCGGCGAGGCGGAGGATGCGGTCG
AAGCGGCGGGCGAGACGGCGGTAGAAGAAGAAGGCAGTGCGCGGCGCCCCAACCGGCAGGTGCAGGTCGAGGGCAGTCTCAAGCACCAAAGCGGGCTCAA
TCCGAACTTCACCTTCGAGACCTTCGTCGAGGGCAAGTCGAACCAGCTCGCTCGCGCCGCCTCCCGCCAGGTCGCCGAGAATCCGGGCGGAGCCTACAAT
CCGCTGTTCCTCTACGGCGGTGTCGGCCTCGGCAAGACCCACTTGATGCATGCCGTGGGCAATCATCTCGCCGGGCTGCGCGAGAATGCGAAGGTGGTCT
ATCTGCATTCCGAACGCTTCGTCGCCGACATGGTCAAGGCGCTGCAGCTCAACGCGATCAACGATTTCAAGCGCTTCTATCGCAGCGTCGATGCGCTGCT
GATCGACGACATCCAGTTCTTCGCCGGCAAGGAGCGCTCCCAGGAGGAGTTCTTCCACACCTTCAATGCGCTGCTCGAAGGTGGCCAGCAGATGATCCTC
ACCTCGGATCGCTATCCCAAGGAGATCTCCGGCGTCGAGGAGCGTCTCAAGTCCCGCTTCGGCTGGGGGCTGACCGTGGCGATCGAGCCGCCGGAGCTGG

>Halotalea alkalilenta DSM 17697 [NZ_KK211229.1:c47040..47539], <mnmG oriC2 <PRK13623 (ErpA), 661 bp

ACGGCAGCGAGGTCGATATCCCATTGGGATAGAGCTCGTGGGTGGTGAGGCCTTCCGGCTCGATGAATACCTGGTGGCTCTGCTTGTCGGCGAAGCGATG
GATCTTGTCCTCGATCGACGGGCAGTAGCGCGGTCCGATGCCCTCGATCACTCCCGAATACATCGGCGAGCGGTCGAGGTTGGCGCGGATGATCTCGTGG
GTACGCTCGTTGGTATGGGCGATGAAGCAATCGACCTGGCGCGGCTGCATCGCGCTGGTGCCGATGTAGGACATCACGGGGGTCGGCGTGTCGCCCGGCT
GCACCTGCATGCGCGAGAAATCGACGCTCCTGGCGTCGATTCGCGGCGGTGTACCGGTCTTCAACCGCGCCACGTTGAACGGCAGCTCACGCAGCCGCGC
CGCCAGCGCGGAGGCGCTGGGGTCGCCTGCGCGGCCACCGCTGTGGTTTTCGAGCCCGATGTGGATCACCCCACCGAGGAAGGTGCCGGTGCACAGCACC
ACCGTCTCGCCATGAAAGCGAATCCCGGCCTGGGTCACCGCACCGCGTACCTTGCCGTCCTCGACGATCAGATCGCTGACCGCCTGCTGGAAGATCGAAA
GGTTCGGCTGGTTCTCCAGCATCTCCCGGATCGCTGCCCTGTACAGCGCTCGGTCTGCCTGGGCGCGCGTCGCACGCACCGCTGGCCCCTTGCGTGAGTT
GAGCACACGGAACTGGATCCCGGCGCGATCGGTGGCCAGCGCCATGGCGCCACCCAGCGCATCGATCTCCTTGACCAAGTGGCTCTTGCCGATCCCGCCA
ATGGCGGGATTGCACGACATCTGGCCAAGGGTCTCGACATTGTGGGTCAGCAGCAGGGTGTCGACGCCCATGCGAGCGGCAGCGAGCGCGGCTTCGGTTC
CGGCATGGCCACCGCCGATGACGATGACGTCAAAGCGGTTGGGGTAATCCACGGTAGTACCTCGAAAGACTGTCGGGGCGCTGGAGCATGCCGGAGAGGG

TCGATCCTGCCGTTCGCGTAAGGCGGAAGGCATGCGGTGCGCTGGGGATTAGGCCGCGCATTATAGCGCGCCTGTGGGTAACCGTCAGGGGTTTTCCACA
CCTCGTCTCGAGCTTGGATTCATTTACTAAACAAAAATAAGAAAAAGGATTTATTTAAAGAAGTTTTGTTGTAGTAGTGACTAACTGAGCAGCGGTTTTTC
**TGTGGAAAA**CCGACTTTTATCCTTTCTCTACATAAACTTAGCCTGTTCGCTACAGTGATGGAAAGTGGCGCACCGGCTGGGTGCGGCCGGTGTGCGGCCT
GTGGACAAGATGCATGGTTGTCCACACCCCGAGTTGTGCCCCTGTTTTCAACCGGCGCATACCCTGCTTGTCATCGTTTTCGTTCACGTCCACGTCAAGC
TGGTACGCGGCCAGCGCTGCCGTGGGCTGTAGGCCGATATCGGGGGTCGGGGAAAAAAACTTGTCCACAGCCGGATATCCGCGGATTCCGGGCTGCGCTG

GATCGAGCCGACGTAAGCCGGGCGTCTTGAGGCGCTGAGTCTTGAGAAGTTAAGAGAGGAAAGACGAGGGTACTCGACGCGGGCATTGCGCCCGCGCGAT
GGGTAAGAGGGGATCAGACGGTGAAGGAGATCCCGCAGCCGCAGGTGGTGGTGGCGTTGGGGTTCTGGATCAGGAAGCGGGCACCGGCGAGGCCCTCCTC
GAAGTCCACCGTGGAGCCGACCAGGTACTGGTAGCTCAACGGATCGACCAGCAGCGAGACGCCGCCGTTTTCGATCACGGTGTCGTCTTCCTGGACCTCG
TCGCTGAAGTCGAAGCCGTACTGGAAACCGGAGCAGCCGCCTCCGGTGACGTATACGCGCAGCTTGAGCGCGGGGTTGTGCTCTGCTTGCGCCAGCTGGC
GCACCCGCGCCGCGGCGCGATCGGTGAGGATCAGCGGAGTCGGGATGAAGGTTTCGGCGCCACTCATGCGTAGCCTCCTGGACAGGTGGTGCGATGGCGC
TGGGCGGACGTCGCTGCGGCCTCATCCATCGCAGAATGGGAGATAGTGTAGCATTTCGCTGCGCCGGCTGGCGTCATCGTGGCCGCCGGGCAGGTCGCTT
TCCGTTATGATGACGAGGCGCGCAGCCCGTTCATCGTCGGCCGTGTTCCCAGGTCGAGCGCGGGCGCGGCAAGCGCGTTGGATTTCGATAAGACCGGCGG
TACGCCGGCGTCACCGACCGTTGGTGCTGGACCGATACTGGATTGGGAGTCCGAATGACCGCTTGGCTTTTGGCATTGCACATCCTGGCGCTGATTCTCT
GGTTCGCGGCATTGTTCTACCTGCCGCGTCTTTTCATCTACCACGCCTACGCGCTGCGCCAGCAAGACGAGCAGGGCGCCCATCGCTTCGAGGTGATGGA
GCGCAAGCTCTACCGGCTGGCGATGAACCCGGCGATGATCGTCACCCTGTTGCTCGGGATTTCGCTTGCGTTGACCAATCCCGGCTACTTCCTCGGCTCG

>Halotalea alkalilenta DSM 17697 [NZ_KK211226.1:c304092..304164], dnaA> ig dnaN>, 73 bp

WebSkew GCmin pos. 469409 (GCmin<>oriC1 116.1 kb; GCmin<>oriC2 13.8 kb; oriC2<>oriC1 102.3 kb)

>Halotalea alkalilenta IHB B 13600 [NZ_CP015243.1:585236..585735], <rpmH ig dnaA>, 428 bp

ATGTAGCTGCGCGAGGTGCTGTCGGAGAGCAGTACGAACGGCGTCTCGGAGTTGAGCGCCTGCTGCTGTTCCGGAAGAGCGGCATAGATGATGTCGCCAC
CGCGCGGATCGATGCGCAGGTCGAGCACGTCGGTGTCGACGCTGACCAGCGAGTTGGAGGCGTTCTCCTGCGCTGGATTGGCGCTTTGCCCGTTCTCCAA
TGGAGAGGTCGCGCGCTCGCTGGTCGATTCGGGCGCGTCGAGCACATCGCTGGGCTGCGTCGCGTCGCTCGAGTTGGCGGCGAGCGTCGGTGCCTGCGGC
TCCGGGGTCGGCTGGTGATAGTCCTGATTCCACTGTACGACCAGCAGATAGGCGAGGATCGCCAGCGGAATCAGGATGATGATGCGTTTCAGATCCATGA
GGGATGAGCCCGGTGAAAGGGTGTCGTGGCGTATCTTTGATGAATGCTGAAGACGGCTCGGCGCGTCGTCTCGAGCGGGAGGCCTTTCACCGGCGGCGTG
CGCGGGGAGATATCTCCCTTGGCACTCCATCCCTCGCGCCGGGAACGGGCAGGGACGGGATCAGGCGGGTTCGCGATCAGGCGCGGGGGCTCGTCGCTTC
ACGTCCTGCTCCAGGCGGCGCCACATGCCGGCAAGCTGCCGGCGGAGGGTCTCCCGGTCGAGTTCGAGAACTCCTCGGCGCGACAGTACCACGATGTCGA
CGGCTGGCAGCCGGTGCTGGCGCAGGCGTATCGACTCGCGGGTCAGGCGCTTGAGCAGGTTGCGATCGACGGCGCGGCGCACGTTCTTACGGCTGAACAC
CAGGCCCACTCGAACGTGCCCATGCTGGTTCGGGCAGGCGAGCGCCAATAGCCCCTTGCCGTGAACCTTGAAGCACGCATGCTCGAAGACGTGGCCGTAC
TCCCCGCTGGTCAGGATTCGCAGACTGCGGGGAAAGCCGTGGATCGCGTCCTGACCGGCCATGGTGGCCTCTCTTGAAATCAGGCGCTCAGGCGTTTACG

GCCTTTGGCGCGACGACGAGCCAGGACCTGACGGCCGCGTTTGGTGGCCATGCGAGCACGGAAACCATGGTTGCGCTTGCGCTTCAGGACGCTGGGCTGA
AAAGTGCGTTTCATGACTGCTTTCCTCGAGACGGAGGGAGTGGTGGAGAGATGGAAAAACGCCGGTGCAAACACCTGCGGTCCCGTCAAGAGCGGGAATT
CTAGTGAATACGCGCAGGCGGCGCAATTTTTCTCTTACCAAGTATGATTTTTTTTGCTAAGGGAGGGTTTGGCTGTGTGTAAAAAGCTGTGGATAGTCAG
GGGGAAAGCTATGGGCAAGCTGGGGAGTGGCTGTGTCATAGGCCTTGGAAGCAGTTTCAGAGCGCTTTTTTCCTGAGTTATGCACAACCTCTTCAATTGC
ATAGATGGTGTGGATAACTCCTGCGCCCCCCGATACACTAGCGGGTCCGTAATATAGCCATTGAGTCCGAATCGCCTTCCGCCCTTCCGGTGCCGGAAGT

CCGCGTGCATGGCCCTCCACTTCGCAAGGATGACGAGTTCGCGTGTCGATAGCACTCTGGCAGAAATGTCTCGAATACCTTCAGGACGAACTGAACTCCC
AGCAGTTCAATACTTGGATACGTCCGCTGCAGGCGGAGGAGGGCGAGGCCAATGGCCTTGCACTGCTCGCGCCGAACCGTTTCGTGCGCGACTGGGTCAG
CGACAAGTACCTGAAACGGATCAACGAGCTGCTTCGCGAACTCGCGCCCGGCAAGCCGCCCAAGGTGTCGCTCGGGATCGGCAGCCGGCGCAGCGTCTCC
AATCCCCGGCCGACCGAGCCGGTGCGCCCGGCCGCGCCGATGATGCCTCGCAGCGCACCGTCGCGGGCGTCGTTCTCCGGCGAGGCGGAGAACGCTGTCG
AAGCGGTGGGCGAGACGGCGGTCGAAGAAGAGGCCAGCGCGCGGCGCCCCAATCGGCAGGTGCAGGTCGAGGGCAGTCTCAAGCACCAAAGCGGGCTCAA
TCCGAATTTCACCTTCGAGACCTTCGTCGAGGGCAAGTCGAACCAGCTCGCTCGCGCCGCCTCTCGCCAGGTCGCCGAGAATCCGGGCGGAGCCTACAAT
CCGCTGTTTCTCTATGGCGGTGTCGGCCTCGGCAAGACCCACTTGATGCATGCGGTGGGCAATCATCTGGCCGGGCTGCGCGAGAATGCGAAGGTGGTCT
ATCTGCATTCCGAGCGTTTCGTCGCCGACATGGTCAAGGCGTTGCAGCTCAACGCGATCAACGATTTCAAGCGCTTCTATCGCAGCGTCGATGCGCTGCT
GATCGATGACATCCAGTTCTTCGCCGGCAAGGAGCGCTCCCAGGAGGAGTTCTTCCACACCTTCAATGCGCTGCTCGAAGGTGGCCAGCAGATGATCCTC
ACCTCGGATCGCTATCCCAAGGAGATCTCCGGCGTCGAGGAGCGTCTCAAGTCCCGCTTCGGCTGGGGGCTGACCGTGGCGATCGAGCCGCCGGAGCTTG

>Halotalea alkalilenta IHB B 13600 [NZ_CP015243.1:482976..483475], <mnmG oriC2 (locus_tag="A5892_RS02175" ignored) <PRK13623 (ErpA), 663 bp

CGGCAGCGAGGTAGATATCCCGTTGGGATAGAGCTCGTGGGTGGTGAGGCCTTCCGGTTCGATGAACACCTGGTGGCTCTGCTTGTCGGCGAAGCGATGG
ATCTTATCCTCGATCGACGGGCAGTAGCGCGGTCCGATTCCCTCGATCACCCCCGAATACATCGGCGAGCGATCGAGGTTGGCACGAATGATCTCGTGGG
TGCGCTCGTTGGTGTGGGCGATGAAGCAATCGACCTGCCGCGGCTGCATCGCGCTGGTGCCGATGTAGGACATCACTGGGGTCGGCGTGTCGCCCGGCTG
CACCTGCATGCGCGAGAAATCGACGCTCCTGGCGTCGATTCGCGGCGGTGTACCGGTCTTCAACCGCGCCACGTTGAACGGCAGCTCACGCAGCCGTGCC
GCCAGCGCGGAAGCGCTGGGGTCGCCTGCGCGGCCACCGCTGTGGTTTTCGAGCCCGATGTGGATCACCCCACCGAGGAAGGTGCCAGTGCACAGCACCA
CCGTCTCGCCATGAAAGCGAATCCCTGCCTGGGTCACCGCGCCGCGCACCTTGCCATCCTCGACGATCAGATCGCTGACCGCCTGCTGGAAGATCGAAAG
GTTCGGCTGGTTCTCCAGCATTTCCCGGATCGCCGCCTTGTACAGCACTCGGTCGGCCTGAGCGCGCGTAGCACGCACCGCCGGCCCCTTGCGTGAGTTG
AGCACGCGGAACTGGATCCCGGCGCGATCGGTGGCCAGCGCCATGGCGCCACCCAGGGCATCGATCTCCTTGACCAGGTGGCTCTTGCCGATCCCACCGA
TGGCGGGATTGCACGACATCTGGCCGAGGGTCTCGACATTGTGGGTCAGCAGCAAGGTGTCGACGCCCATGCGAGCGGCGGCGAGCGCGGCTTCGGTTCC
GGCATGGCCACCGCCGATGACGATGACGTCAAAGCGGTTGGGGTAATCCACGGTAGTACCTCGAAAGACTGTCGGGGCGCTGGAGCATGCCGGAGAGGGT

CGATCCTGCCGTTTGCGTAAGGCGCAAGGCATGCGGTGCGCTGGGGATTAGGCCGCGCATTATAGCGCGCCTGTGGGTAACCGTCAGGGGTTTTCCACAC
CTCGCTTTTGACCTTGGATTCATTTACTAAACAAAAATAAGAAAAAGGATTTATTTAAAGAAGTTTTGTTGTAGTAGTGACTATTGAGTAGCGATTTTTC
**TGTGGAAAA**CCAACTTTTATCCTTTTCATACAGAAAGTTAGGCTGTTCGCTACTGCGATGGAAAGTGGCGGACCGGCTGGGTGCGACCGGTGTGCGGCCT
GTGGACAAAATGCATGGTTGTCCACACCCCGAGTTGTGCCCCTGTTTTCAACCGGCGCATACCCTGCTTGTCATCGTTTTCGTTCACGTCCACGTCAAGC
TGGTGCGCGGCCAGCGCTGCCGTGGGCTGCAGGCCGATATCGGGGGTCAGGAAAAAAAACTTGTCCACAGCCGGATATCCGCGGATTCCGGGCTGCGCTG

GATCGAGCCGACGAAAGTAGGGCGTCTTGAGGCGCTGAGTCTTGAGAAGCTAAGAGAGAGAAAGGCGAGGAGACACGACGCGGGCATTGCGCCCGCGTGA
TGAACAAGCGGGGATCAGACGGTGAAGGAGATCCCGCAGCCGCAGGTGGTGGTGGCGTTGGGGTTCTGGATCAGGAAGCGGGCACCGGCGAGGCCTTCCT
CGAAGTCCACCGTTGAACCGACCAGGTACTGGTAGCTCAACGGATCGACCAGCAGCGAGACGCCGCCGTTTTCGATCACGGTATCGTCTTCCTGAACCTC
GTCGCTGAAGTCGAAGCCGTACTGGAAACCGGAGCAGCCGCCTCCGGTGACGTACACGCGCAGCTTGAGCGCGGGGTTGTGCTCTGCTTGCGCCAGTTGG
CGCACCCGCGCCGCGGCGCGATCGGTGAGGATCAGCGGAGTCGGGATGAAGGTTTCGGCGCCACTCATGCGTAGCCTCCTGGACAGGTGGTGCGATGGCG
GTGGGCGGACGTCGCTGCGGCCTCATCCATCGCAGAATGGGAGATAGTGTAGCATTTCGCCGCGCCGGCTGGCGTCATCGTGGCCGCTGGGCAGGGCGCT
TTCCGTTATGATGACGAAGCGTGCAGCCCATTCGTCGTCGGCCGTGTTCGCGGGTCGAGCGAGCGGGCGCGGCAAGTGCGTTGAATTTCAATAAGACCGG
CGGTATGCCGGCGTCACTGACCGTTGGTGCTGGACCGATACTGGATTGGGAGTCCGAATGACCGCTTGGCTTTTGGCATTGCACATCCTGGCGCTGATTC
TCTGGTTCGCAGCGTTGTTCTACCTGCCGCGTCTTTTCATCTATCACGCCTACGCGCTGCGCCAGCAAGACGAGCAGGGCGCCCATCGCTTCGAGGTGAT
GGAGCGCAAGCTCTACCGGCTGGCGATGAACCCGGCGATGATCGTCACCCTCTTGCTCGGGATTTCGCTTGCGTTGACCAATCCCGGCTACTTCCTCGGC

>Halotalea alkalilenta IHB B 13600 [NZ_CP015243.1:587242..587314], dnaA> ig dnaN>, 73 bp

WebSkew GCmin pos. 3019977 (genome 3023483; GCmin<>oriC1 3.2 kb; GCmin<>oriC2 25.7 kb; oriC2<>oriC1 22.5 kb)

>Zymobacter palmae IAM 14233 = DSM 10491 [NZ_AP018933.1:c2942248..2942747], <rpmH ig <PRK00073, 234

CCGCGAGGGTCAGACGATACTGGCCATCAACGAAGACGTTCGACTGAGCCACGTAGTACATAGAGCTGCTTGTCGACATCAGCACGAACGGTTCTTCGCC
ATGCAGCTGGGCTTTCTGCTGCGGCAGAGCGGCGTACACGATAGCGCCGTCGTTCGGATTGATCCGAACGTTGAGAACGTCGGTGTTCACGCTGACGAGT
GCTGACGAATTGGCCGCGCGGGCGTTATCGGCAGCAGGAAGTTCGGAGGCGGTCGCGGGGGCCTGTGTCACGTCGTCAGGCTGCTGGGCATCGCTCTGGG
TGGCCGTCAGGACACTTTGAGGCGCCTGCGACTGGGGAGTGTCCTCGTTCCACTTGATGACGAGCAGGTACGCTACGATCGCCAGTGGAATGAGCAGGAT
GATGCGTTTCACGTCCATGAGGGAAAACTCGGTGGAATAGAGTGTCGTATCAACGTGATGCCTGACGTACAGGCACGAACGCCATGAGGGTTATCAAGAC
AGTCGCTGTTCCGGCGGCTTCCGCTGGGCACAACGCAAGAAAGCCCGTCGGCACCGGGCCGTCAGGCAGAAGTCTTGGCGTCGGCTGCGCGCTTCACGTC
CTTTTCAAGACGTCGCCACATACCATGCAGTTGACGTGCGAGGGTCTGGCGATCCAGCTCGAGAACACCACGTCTAGACAGAACGACTATGTCGACCGCG
GGCAGCCGATGCTGCTGCAGTCGAATTGACTCGCGCGTCAATCGCTTCAGCAGATTACGATCCACTGCGCGACGTGCATTCTTCTTGCTGAATACCAGAC
CGACGCGTACGTGCCCCACATCATTGGGGCAGGCCAGGGCCAGAAGACCCTTGCCGTGCACCTTGAAGCACGCACTGTCAAAGACATGGCGGTATTCGCG
GCTGCTGAGAATACGCAGTTCACGCGGAAAACCGAAGGTCGGTGTCGAACCGCCCATGCCGATAATGATCAGGCGCTCAGGCGTTTGCGGCCTTTGGCAC

GACGACGTGCCAGAACCTGACGGCCACGCATGGTAGCCATGCGAGCACGGAAACCGTGGTTACGTTTGCGTTTCAGAACGCTGGGCTGGAAAGTACGTTT
CATTGGATCGAATTCCTGTTGTAGAAGGATGTCGAGTCAGCTCGGCTTCCCTCAAACAGGGAATGACACGGGCTGTCAAGAGCGGCGATTTTAGAGAATC
CCCGCCACAGGTGCAATTCCTGTTGTGTCATGGACACATCAAAATTATCACCTGCCGATGCAAGCTGCCCATCGGTAGAGCCGATGGGCAGCAGGGCAAG
GGACCCCAAGCCAAGCTTGGAATCCCTTAGGCAGCAATTACTTGGCTGCGTCTTCCAGTGCTTTTACAGCTGGAAGAACTTTGCCTTCGACGTATTCAAG
GAATGCGCCGCCGCCAGTAGAAATGTAAGACACTTGGTCTGCAATGCCGTATTTGTCGATAGCGGCCAGCGTGTCACCACCACCTGCGATGGAGAACGCT

TTGCTTTCTGCAATCGCTTTGGACAGTGCTTTGGTACCTTCTGCGAACTGGTCGATTTCGAATACGCCGACCGGACCGTTCCACAGGATGGTGCCAGCGC
TTTTCAGAGCGTCAGCGAAAGTAGCGGCCGTTTTCGGACCTACGTCCAGAATCAGGTCGTCATCAGCAACGTCGGCAACGTCTTTGACAGTTGCTTCGGC
TTTGTCGGAGAACTCTTTGGCTACGACAACATCGACCGGTACCGGGATGTTGACTTTGCCCATCAGGTTCTTAGCAGCATCGACCAGATCAGCTTCGTAC
AGGGATTTGCCCACGTTGTGGCCAGCCGCTGCGATGAAGGTGTTGGCAATACCACCGCCGACGATGATGGCGTCGCATTTTTCGGACAGCGCGTTCAGCA
CTTCCAGTTTGGTGGATACTTTGGAACCACCAACGATAGCCACCATCGGGCGTTTCGGTTCAGCCAGCGCTTTGTGCAGAGCGTCCAGCTCGGATTCGAG
CAGCGGACCTGCAACGGCTTCAGGCGCGAAACGAGCTACGCCATGAGTAGAAGCCTGGGCGCGGTGAGCGGTACCGAAAGCGTCCATGACGAAGATGTCA
CACTGGCTAGCGTATTTCTTGGCCAGCGCTTCATCGTCTTTTTTCTCGCCTTTGTTGAAGCGAACGTTTTCGAACAGGACGATGTCGCCATCGTTCAGTT
CGACTTTCTTGTCGAGGTAGTCTTTTTCCAGACGAACCTGAACGCCGAGCAGTTCGCTCAGGCGGTTAGCAACCGGTGCCATGGAAGAAGCTTCATCCGG
TTCGCCTTCAGTCGGGCGGCCCAGGTGGCTCATCAGCATCAGTTTGGCGCCAGCGTCGTGGGCCATTTTGATGGTTTTGAGGCTTGCGCGAATACGCGCG
TCAGAAGTGACACGACCGTCTTTGACCGGAACGTTCAGGTCTTCACGGATCAGGACGCGTTTTCCTTTGAGGTCCAGCTCGGACATCTTGCGCACTGTCA

>Zymobacter palmae IAM 14233 = DSM 10491 [NZ_AP018933.1:c3022921..3023420], (SdiA) PRK10188> ig dnaA>, 345 bp

AGAGATTTTGGCTTGTCTTAAAGTACAGTATTACGTGGCTAATTCTGATGATACCTGATTATAACTCACCTGACCTATCGTGTAGTAACAGCACGAAATA
CAATGAGATAAGGCATAATATGCAAAAAACGGTCAAAAAAACGGGTTTATTCCAATAAATACACCCGAGCGCTCAGATAAGCATTTTCCTTTCAGTGCGC
TTACCACCCTAATTTATTTTTCCTAATCAAAAAGATAGATAAGGGAAATTCTTATCTTGATAATTGTGGTCACGTGTTATTATATGGTAATACTGCACAT
GTTATTATGTGCATAACTCTACCAAAGGTGAACGCCGTCATGTCCGACTTGCGACGGGTGTGCTTTACATGGCACTCTTTCAACTCGAAATATCTATAAA
AAACACCAGTTCTTCTCTCAAACTGACCTTTCAGTCAGTGTTTCTTACTGAATCCTTTTTTTACCGTGAGGAGAACTTTTCAGTCATCGCGATACAAATG
AGCATCGTGGGATATGAATAATTGGCAAATCAACTATATGACCGAGCTGAAGAAGGCATCGCTTCTTCAGGACGTGCTCGACATCGCGCAGCACGCCATC
CGCTCGCTCGGGCTCGAATATAGCAGCTGGATTGCCTACAACCCTTTCCCACTATCACGCAGGGACATGTTCGCGCTAACGACGACGGACGACGAATGCC
ATCGCCGCCTGAAAGCCGGCGAGTTCAACGACAGTCCGATCTTTAGGGCGTTGGAAGAAACACAGCAACCCATCTGCTGGCAGGGCGATGAAAACGACCA
GCTACTGGGACAGCAGCCGGAATTCCTCTCCGATTACCGCAGCCTCGGTCGTCGCAGTGGCTGGGCCCAGACCACGATCGACAATACCGGTACGCGCAGT
TTTTTCCTCAACTGCTGCACGCGCATTCTGTCGCTGACCGACCTCACCAATATCAGCATCAATATGCAGTGGCTGGCCGCGTCGGTTCACTACAATATGC

TGCGTGTCAAAGATCGCGCTGGCGTAAACCTGTCGCTGCGCGAGCGAGAGATCTTATGCTGGACCGGCGATGGCAAGACCGTCAGCGATATCGCGGAAAT
CTTGGGGCTTAGCGAGAGCACAGTAAATTTTCATTTCAGAAACATCATGAACAAGCTGGGTACCCCCAATAAGACCTCCGCCGTAGTTAAAGCGATCTTT
CTTGACCTGCTCTTCTGACGTCCTTCTGCTCGATCCATCCTGTCTTTATCCTCTGTTCCCCCGCGCACCTTACCCACATCCCTGTTATGATGCCCCCTCG
TTTCCGTAAGCGTCAGCCATGTTCATGACACGCTTCTGTTGCCGTTACAGGCAAGGCGAAACGACACGCCCCGCTCAAGTTATCCACACCCCTCTGTCAC
CAGTCGGGTGTGGATAACTTGTTTCGGGGCCGCTAGACTAATGAATCCCGCGCGCAGCGTCTTCGCTGCCACGGTCAGCATTCCAGGCGCTGCGCCCAAC

GCGCTGTGTCTCTGTTTTCGATCACTCCTGATGACCTTCGTCTACGAAGGATGGTGAAGTCGCGTGTCGACAGGTCTTTGGCAAAAATGTCTCGCATACC
TTCAGGATGAATTGAGCTCTCAGCAGTTCAACACCTGGATCCGCCCGTTGCAGGTGGAAAGCGATATGGACAGCAATACGCTTGTCCTCTTGGCTCCCAA
CCGTTTCGTACGGGATTGGGTCAGTGACAAGTACCTCAAGCGCATCAATGAACTGCTGCGCGAACAGACCTCTGGCAAGCCTCCAAAAGTCTCGCTGGGT
ATCGGTAGCCGCCGAGCGGCCGCCGCACCGGCAGCGGCTCCCAAGGAGCGCCCGCGTGGGGTAGCCACGCCGAGCGCGCGCCCTGCTGCGCCGCGGGTTG
AGTTCGGTCAGAACAGCGAAGACAACGCTGCTGCGGACATCGACGAAGCGCCGGCTGTCGTGCGTGGTAACGCTGCGCGCCACGTGCAGGTCGAGGGGGG
GCTCAAGCACCAGAGCGGCCTGAATCCCAACTTCACCTTCGACACCTTCGTCGAAGGTAAGTCGAACCAGCTGGCCCGCGCCGCTTCCCGCCAGGTCGCC
GAGAACCCGGGCGGTGCCTACAACCCACTGTTCCTGTACGGCGGTGTCGGTCTAGGTAAGACCCATCTGATGCACGCTGTCGGCAATCAGTTGGCGGGGC
TGCGTGAGAATGCACGCGTGGTCTATCTGCATTCCGAGCGTTTCGTGGCGGATATGGTGAAGGCACTGCAGCTCAACGCCATCAACGATTTCAAGCGTTT
CTACCGCAGCGTCGATGCTCTGCTGATTGACGATATCCAGTTCTTCGCTGGCAAGGAACGTTCGCAGGAAGAGTTCTTCCACACTTTCAATGCGTTGCTT
GAAGGCGGTCAGCAGATGATTCTGACCTCGGACCGCTATCCCAAAGAAATCGCGGGCGTCGAAGAGCGTCTGAAGTCGCGCTTTGGTTGGGGGCTGACCG

>Zymobacter palmae IAM 14233 = DSM 10491 [NZ_AP018933.1:c21994..22493], <mnmG oriC2 <COG2859, 678 bp

AACGGCAGCGACGTTGAAATGCCGTTGGGATACAGCTCGGGCGTATCCAATCCTTCGGGTTCGATGAAGACCTGATGGCTGTTCTTGTCGGCAAAGCGGT
GGATTTTGTCTTCGATCGACGGGCAGTAACGCGGACCAACACCTTCGATGACACCGGAGTACATCGGCGAACGGTCGAGGTTATTGCGGATGATGTCGTG
GGTGCGTTCGTTGGTATGCGCGATATAGCAGTTGACCTGCCGCGGGTGCATTTCACGATTGCCCAAGAAGGACATCACCGGCACCGGCGTGTCACCTGGC
TGAACCGCCAACTTGGAGAAGTCAATCGAACGTGCGTCCAGACGCGGCGGCGTTCCGGTTTTCAGGCGCGCCACGTTGAACGGCAGTTCACGCAGACGTT
CCGCTAGAGCAATTGCACTAGGATCGCCAGCACGTCCCCCTCGATAGTGTTCCAGACCAATGTGAATCATCCCGCCCAGAAACGTTCCCGTACACAGCAC
TACTGAGGTGCTGTGAAACCGAATGCCTGCTTGCGTCACCACACCTTTTACGTCATCGCCTTCAACGATCAGATCGCTGGCTGCCTGCTGAAAAAGCGAG
AGGTTGGGTTGATGCTCGAGGGTATGGCGGATGGCGGCCTTGTAAAGGACGCGGTCCGCCTGTGCGCGCGTAGCGCGCACCGCCGGCCCTTTTCGGGCAT
TCAGGGTACGGAACTGAATACCGCCTTTATCGGTGGCCAGCGCCATGGCTCCCCCGAGGGCGTCGATTTCCTTGACCAGATGGCTCTTGCCGATGCCACC
GATGGCCGGGTTACAGGACATCTGTCCCAGCGTTTCGATGTTGTGAGTCAGCAGAAGCGTTTTCGCACCAGTGCGCGCAGCGGCCAATGCGGCTTCTGTT
CCGGCATGACCACCGCCGATGACGATGACGTCGAAGCGGTCTGGATAGTCCACGTGTGTAACCTCGATGAATGGAGCTTTTGGGGTAACGGCAACGTCAG

GAAAAAATCGTTTCCATGCGAATGCGATGGGGGCAATCCGTCGGGGAGGCTGCGTTGCGGGCCGCATTATAACGTGGCTGTGGGTAACCGTCAGGGTTTT
CCACATGGGCCAAGTCATAACAGTGATTCAGTACATATCTAAAGAAAAGTATTTAAAGAGAAGTTCTATTGAAGTAATGATGATTAGTCAGACGTCTTTT

**TGTGGATAA**CGCACTTTTGTCCATTGAATACATGAATTTATAGCCTTCATATCCTTCTGGGGCGATGTTGTATGGACTGGGGAGATGATCTTCATGACCT
GTGAACAACTTTCGTGATTATCCACATGTCCATCCGTCCACGGTTGCTAACGGATCTGTACGTAAGTTGTGCCCGAGGTTATTAACGTATACGTCAAGG
TAGTTCGAGTTGGCATGCCACTAGGGATGGCAAGGAGATAGGAGTGGATAACAGTGATTTTTATCCACAGCCCAATTTCCTTATTTCCAATGCTGTTTTG

AGGGGCTTTACATGAAGGCGGAAGGTACTGCATACGATCGAGGGGCCATTGCAAGCGTCATTTGCTTGACGTTAAAAAGCCTGCCGTGTGGCAGGCTTTT
CTGCATGATGCTTGAGCAGCGCCGAGGACGTTTAGTCCTTGAGGGTGTAATCGATGTTGGTGACGACGCGGACTTTCTTGATGTCAGGCGTGTAATTGTC
CAGATCTTCTACCGTGAAGTACCCCTGTGAAGCGTTCTTGATTGCGCCAAGGTGGCTTTCCGAATCCGACGCAAACTGCTCTGCCGCTTTGCGTGCGTTG
GCCGTCGCTTCGCGGATCATGTCGGGCTTGATGTCATTAAGCTTGGTGAACTGGTACTGCGGGGCGTAATCGTTGCCATCGTACAGCCCGGATGACAGCA
GCACTCCTTGGCTGACCAGATCACCTGTATGGCGCGAAGCCGTGCGAACGGTTTCTACCTTGGATGTGCGCACGACGACAGCAGCAGAGCCGGTGTAGCG
GTTGATAATCGGTTTGTCTGAAGAACCGTATTCGTTGCGATCGCTGACCTTCATCGGTGAGAGGCTAAGCTCGCTGTCTTCGAACCCTTGCTGGCGGAGG
AAGGCTCGGGTGAGGTTTTCACTTTTCTGCAGCTCGATCTGCAGCGAGGCCAGATCCTTGGCTTCTACACGGAAATAGACGGGCCAGAGGGCCATGTCAG
CTTTCAGTTCGCGTTCGGCGACCCCTTTGACGGAGGCGATGCGGTTAGATTCCTGCCAGATCTTGCCGACCGTGCGTAGGTAGCTGCCGCTCCAGACCAA
GCCTACGGCGATTAGGCCGCCCAGCACGGCACCTGTCAGTAGGTTGCTTCGTGAATGACTCATTGAAATATCCCTGTGAGATAGGAGGGGTGCGAGGAGG
GTTCCTCTCTTCTTTTGTAGCGCAGAAGAAGTGTATCGCAACGTGTGTTCATGAAAGGTGAAGGCAGTGATGAAAATGCCCTTGCTTCCAATAAGGCAAG

>Zymobacter palmae IAM 14233 = DSM 10491 [NZ_AP018933.1:c1468..1655], dnaA> ig dnaN>, 188 bp

GGCACACGGGGTAGACCGTCTGCCGCGGGGAACATGACCGTTTCCGCAAGGCCGATGGCATAGGTCACAATGGCAATAGGCGCAGGTCGTCAAACGCGCT
ACAGTCATGCCATGATATCGTTATGGAATCACACCGCAGGTGTGTCTTCCGACAGAAGTTTCAGCCCAGATACGAAGAGCGGAATGCC

WebSkew GCmin pos. 364812 (GCmin<>oriC1 34.6 kb; GCmin<>oriC2 1.4 kb; oriC2<>oriC1 33.2 kb)

>Larsenimonas salina CCM 8464 [NZ_JAMLJJ010000003.1:329968..330467], <rpmH ig dnaA>, 405 bp

AAGGTGATCCGGCCCTTGTGATTGCTGAGCTCGAGGCCGGACTTGGCCACATAGTTCAAGGCCTGTTCGTTGGCGAGCAGCACAAAGGGCTGATTGCCGC
CGAGTTGCTGGGTGTGCTTGGGCAGTGAGGCGTAGACGATGTCGCCGCCACGAGAATCGATGCGCAGATTCAAAACGTCAGTGTCGACGGAAATCAGCGT
GCTGCTGCCGGAGGAGGCAGGTTCTTTGGTCGCATCTACGCTTAGTTCACTGGTCGTTGACGGTGCATCGAGGCTGTTGGATTGCTTGGCGTCGCTGCCC
TGGCTTGCAAGCGCCGGCGTTTCGGGCTGCGGTGTCTTGGGCTGGTGATAGTCCTGATTCCACTGGATGACCATGAGATAGGTCACCACGGCGAGAGGTA
TCACGAAAAGGAGTCGTTTCAGATCCATGCGTGTTTTGCCCGGTAAGGGTGTGTATGTCGGGTCCAGACGAAGAACCGCCCGAGCCAGTCATTGCAACAT
GCTGGTCGGGCGGAGTCTCTGACGTCTTGTGGCCATGACCTTCGTACTCGCATAATCAAGCGGACTTTTTCGAACGGTTCCGTTTGACGTCCTGTTCGAG
TCGCCTCCACATGCCGTGAAACTGGCGCTTGAGGGTCTCGTTGTCCAGGTCTGCAACACCACGCTTGGAAAGAACTACGATATCCACAGCGGGGAGCCAG
TGCTGACGAAGGCGAATGGATTCGCGTGTCAGACGTTTCAATCGATTTCGGTCAACGGCGCGACGCATGGTTTTCTTGCTAAAAACCATTCCCACACGGG
CATGGTTCAAATCATTGGGGAAGGCGAGGGCGAGGACGCCTTTGCCATGCACCTTGTAACACGCATTATCGAATACGTGCTGATACTCTCGGCTACTGAG
TAGCCTTAGAGCGCGTGCAAACCCATGCGACTCCATCCAAGGACCGATTAAGCGCTGAGGCGCTTACGACCTTTCGCGCGACGACGAGCGATGACCTGAC

GGCCGTTTTTGGTTGCCATGCGGGCACGGAAACCGTGGTTGCGCTTGCGTTTGAGCACGCTGGGCTGAAATGTACGTTTCATAGCTGATTCCCAGGAAGT
GGTTGGTATGAAAACAGGTGATTAGCGGCACTTGCCTTTTTGCAAATTCCGATCAAGGGCGAGGATTCTAGTGAATTAGACGCAGCCGTGCAATTTTAAT
TCGCTGTTAATTCACTGATCGATACGTGATTTTTTCCGCTGGGAGGCGCGTGTTCAGCACACCCTGGTGCTTAGACCCTGTGGACAGTTTTGTGTACTAA
AAAAGGAAAACTCTGGAAGGGTGTGCGTGTAAGCTCAATGATTGTTAGCAGAATCATAAGGTTGTTCCCAAATTAATTATCCACACCCCGAGCCGGTGCG
CGACGGGATGTGGATAACCTGAGTGTGAATCGCTACAATGCCGGCTTTGCATCGGTCAAACCCTGGTGAGGTCGTGTGTCTATCGCTTTGTGGCAAAAAT

GCCTTGATTATCTGCAGGACGAGTTGAGCTCACAGCAGTTCAATACCTGGATTCGTCCATTGCAGGCCGAGGCCGTCGAAGACGCGAACGAGGTTGAGCT
GCTTGCGCCCAACCGGTTCGTGCGCGATTGGGTCAACGACAAGTATCTAAAGCGCATCAACGAAATGATGCGCGAACTGTCGCCGGGCAAGCCGCCCAAG
GTCAGCCTTGGTATCGGCAGCCGGCGCAGTGTATCAAATCCTGCTGTGACATCTGCAGCGGCGGGTTCGACGCGAACGTCAAACGATGAGGCACCCGCCG
CGCCGGTTCCCCCCAAGATGCCATCCGCCGGGCGCATGAACATGTCAAATCGGCAGGACGCCCCACGGCCGGCCAGCGAACCGGCAATGCCGTCGGGCTC
GGGCACGAGTGGCCAGAATCAGCGCTCCTCAAGCGGCGAGCGCCATGTTCAGGTTGAGGGCAGCCTCAAGCACCAGAGCGGCTTGAACCCCAATTTCACC
TTTGAAACCTTTGTCGAGGGCAAGTCAAACCAGCTTGCGCGCGCGGCGTCACGCCAGGTCGCCGAGAATCCGGGGGGCGCTTATAACCCCTTGTTTCTTT
ATGGCGGTGTTGGCCTGGGTAAAACGCACTTGATGCATGCGGTCGGTAACCACCTGGCTGGACTTCGTGATAACGCTCGCGTGGTGTATTTGCACTCCGA
GCGCTTCGTGGCGGATATGGTCAAAGCGCTTCAGCTCAATGCCATCAACGACTTCAAGCGCTTCTATCGAAGCGTCGATGCGCTTCTGATCGATGACATC
CAGTTCTTTGCCGGCAAGGAACGCTCGCAGGAAGAGTTCTTTTATACCTTCAATGCCCTGCTTGAAGGCGGACAGCAGATGATTTTGACCTCCGACCGCT
ATCCAAAAGAGATCAGCGGCGTGGAGGAACGATTGAAGTCTCGTTTCGGGTGGGGGCTAACGGTCGCGATCGAGCCGCCGGAGCTCGAGACTCGCGTGGC

>Larsenimonas salina CCM 8464 [NZ_JAMLJJ010000003.1:363150..362651], <mnmG oriC2 <PRK13623 (ErpA), 562 bp

GCGCGTTTTCCAATCCCTCGATGGAGCGCACCAGTGCCAGCTGCACATCGAAGGGTAACGACGTCGAAATACCGTTCGGATAGAGCTCATGGGTATCGAG
CCCTTCGGGTTCCAGAAAGATCTGGTGGCTCGTCTTGTCAGCAAATCGATGGATCTTGTCTTCGATCGACGGACAGTACCTCGGCCCTACGCTTTCGATG
CTGCCCGAGTACATCGGAGAGCGGTCCAGATTCTTACGAATGATGTCGTGAGTGCGCTCATTGGTGTAGGCAATGTGGCAGCTGACCTGAGCCGGATGTT
GATTCCGCGAGGCCATGTAGGACATGACTGGGACGGGTGTATCGCCCGGTTGCTCGGCAAGCCTTGAGAAATCCACGGTACGGGCGTCGATTCTGGGCGG
CGTGCCTGTCTTGAGACGACCGACGTTGAACGGTAATGCTCTGAGTCGTTGTGCCAGTGCATTGGCTGGAGGGTCGCCCGCGCGCCCCCCGGTTTGACGC
TCCATACCGATATGGATAATGCCGCCTAGGAAGGTCCCGGTGCAGAGCACAACGGTTTGACTTGTAAACCGAATGCCGCCGCTGGTCACAACGCCTTTGA
CACTTCCGCTTTCAACGATCAGGTCATCGGCAGCTTGTTGAAATAGCGTCAGATTCTCTTGGTTTTCCAGCATATGCCGGATTGCTGCCTTGTAACGCAC
TCGATCGGCCTGAACGCGCGTTGCACGAACAGCAGGGCCCTTGCGTGCGTTCAGTACACGAAACTGGATACCCCCTTGATCCGTGGCAAGTCCCATGGCG
CCACCGAGCGCATCGATTTCCTTGACCAGGTGGCTTTTTCCGATGCCCCCGATAGCAGGATTGCAGGACATCTGTCCGAGCGTCTCGATGTTGTGTGTCA
GAAGTAGGGTTTTGGATCCTGTACGAGCGGCGGCAAGTGCGGCTTCGGTTCCTGCATGGCCACCGCCGATGATGATGACATCAAAGCGGTCCGGATAATC

CACGGCGCGACCTCGTCAATACGAATGGGAGTGGGGATAACTGTGTACCTGAATAACGCGCATTATAACGTCCCTGTGTGTAACCGTCAGTCTTTTCCAC
ACCTTCCTGGCTCTACGTCTGAACGACCTTTCTATTAAATATAAAGAGAATTATATAAAGAAGTTTTGTAGTGGTTGTGATTGTAGGGGACGCTATTTCT
**TGTGGTTAA**CTCCCCTTTTTACCCACATAACAATGGATTAGACTGTTCAAAATGAGGCAAAAAACTGCGTTAAGCCCTGGTGAAGACGTTCTAATAAGTT
GTGTATGAGTTGCCGTTTCATCCACAGGAAAATTGTCCACGTGTTTATCAACGTGCACATACCATGTCTGTTACCAGGGTTTTCAAGAATTACGAGTATC
CTGAAATCGGGGCTGCATCGCCTGTATGGCGCGGGAAATACGAAACACGCCTGAAAAGTTATCCACAGGGGGTTTGCGAAATAATTTGCGAAAAATGAGG

AAAAAATAAAAAAAGCGGGCCATAAGGCCCGCTTTTATATGCGTACTTGAGTCTACCGATGAGGGTTAGACCATGAATGAAGCTCCACAACCACAGGTCG
TGGTGGCATTGGGGTTCTGGATCAGAAACCGCGCACCGGCGAGCCCTTCTTCGTAATCGACGGTAGAGCCAACTAGATATTGATAACTCAGTGGATCAAC
TGCGAGTGTTACGTTCTCGCGTGTAATTAGCGTATCGTCTTCGGCGGTCGCCTCGGTTAGGTCGAAACCGTATTGGAATCCAGAGCAACCCCCACCAGTT
ACGTACACTCTAAGCTTGAGAGCGACATTTTGCTGTTCTTGCATCAGCAATTGGACGCGGGCAATGGCCCGATCCGTTAATATCAAGGGCGTAGGAACAA
ATGAACTTACCCCGCTCATTGCAGACTCCTGCGTTCGCGTGGAAATGATGAACATTATCGTGGGCTTTGTCGCAGTGGGTCAAACTGGATGCCAGATAAT
TTAGTACGTCGGTCACGTGTCCGGCGTCAAAGGAGAACAGTGTGCGTCGCCGCCTTAATAGGTCCTATTCCTTCACGCTGGATACGTTCCGGCGACAATT
AGCCAGTATCGACGCGCTTCCCCAATTATGCATTCTCGGCGTGCTTTCTGGTGCATTGACAGGCGCCATCATGGTCGGTTTTCGAAAGGTCATTTCGTTC
GGGGCCGCATTGTTTATGCCCAATGGCAGCTCAGAAGCATTTGAGCTTTTGAATCCATATATCCGCATCGGACTTCCGATTCTTGCTGCACTACTCATCA
TGACCGGTCTTGCCTGGCAGCCTATGAAGCACCGAAAGCTCGGCATTGGTCACGTTATCGAACGCCTTACTTATAACCAAGGCCAAATGCCGAGCACTAA
TTGGCTCAACCAGTGGTGGGTGGGCGTACTGTCTGTACTAGGTGGTCTCTCTGCGGGCCGCGAAGGGCCCGCCATACATTTAGGCGCGGGTGCGTCGAGT

>Larsenimonas salina CCM 8464 [NZ_JAMLJJ010000003.1:328416..328474], dnaA> ig dnaN>, 59 bp

WebSkew GCmin pos. 39200 (GCmin<>oriC1 30.0 kb; GCmin<>oriC2 1.7 kb; oriC2<>oriC1 28.2 kb)

>Larsenimonas suaedae DSM 22428 [NZ_JARWAO010000001.1:68948..69447], <rpmH ig dnaA>, 392 bp

GAAGGTGATCCGGCCCTTGTGATTGCTGAGCTCGAGCCCGGACTTGGCCACATAGTTCATGGCCTGTTCGTTGGCGAGCAGCACAAAGGGCTGATTGCCG
CCGAGTTGCTGGGTGTGCCTGGGCAGTGAGGCGTAGACGATGTCGCCGCCACGAGGATCGATGCGCAGATTCAAAACGTCGGTGTCGACGGAAATCAGCT
TGCTGCTGCCGGAGGAGGCAGGCTCCTTGGTCGCGTCTACGCTCAGTTCATCGGTCGTTGACGGTGCATCGAGGCTATTGGACTGCTTGGCGTCGCTGCC
CTGGCTTGCAAGCGCCGGCGTTTCGGGCTGCGGTGTCTTGGGCTGGTGATAGTCCTGATTCCACTGGATGACCATGAGGTAGGTCACCACGGCGAGAGGT
ATCACGAAAAGGAGTCGTTTCAGATCCATGCGTGTTTTGCCCGGTAAGGGTGTGTATGTCGGGTCCAGACGAAGAACCGCCCGAGCCAGTCATTGCAACA
TGCTGGTCGGGCGGAGTCTCTGACGTCTTGTGGCCATGACCTTCGTACTCGCATAATCAAGCGGACTTTTTCGAACGGTTCCGTTTGACGTCCTGTTCGA
GTCGCCTCCACATGCCGTGAAACTGGCGCTTGAGGGTCTCGTTGTCCAGGTCTGCAACACCACGCTTGGAAAGAACTACGATATCCACAGCGGGGAGCCA
ATGCTGACGAAGGCGAATGGATTCGCGTGTCAGACGTTTCAATCGGTTTCGGTCAACGGCGCGACGCATGGTTTTCTTGCTAAAAACCATTCCCACACGG
GCATGGTTCAAATCATTGGGGAAGGCGAGGGCGAGAACGCCTTTGCCATGCACCTTGTAACACGCATTATCGAATACGTGCTGATACTCTCGGCTGCTGA
GCAGCCTTAGAGCGCGTGCAAAACCATGCGACTCCATCCAGGGACCGATTAAGCGCTGAGGCGCTTGCGACCTTTCGCACGACGACGAGCGATGACCTGA

CGGCCGTTTTTGGTTGCCATGCGGGCACGGAAACCGTGGTTGCGCTTGCGTTTGAGGACGCTGGGCTGAAATGTACGTTTCATAGCTGATTCCCAGGAAG
TGGTTGGTATGAAAACAGGTGATTAGCGGCACTTGCTTTTTGCAAATTCCGATCAAGGGCGAGGATTCTAGTGAATTAGGCGCAGCCGTGCAATTTTAAT
TCACCGTTAATTCACTGATCGATACGTGATTTTTTCCGCTGAAAGGCGCGTATTCAGTACGCTCTGGCGCTCAGCCCCTGTGGACAGTTTTGTGCACTAA
AAAAGGAAAACTCTAAAAGGGTGCGTTTGTAAGTCCAAGGATTGTTAGCAGAATCATAAGGTTGTTTCCAAATTAATTATCCACACCCCGAGTCGGCACA
TGACGGGATGTGGATAACCTGGGTGTGAATCGCTACAATGCCGGCTTTGCATCGGTCAAACCCTGGTGAGGTCGTGTGTCTATCGCTTTGTGGCAAAAAT

GCCTTGATTATCTGCAGGACGAGTTGAGCTCACAGCAGTTCAATACCTGGATTCGTCCCTTGCAGGCCGAGGCCGTCGAAGATGCGAACGAGGTTGAGCT
GCTTGCGCCCAACCGGTTCGTGCGCGATTGGGTCAACGACAAGTATCTAAAGCGCATCAATGAAATGATGCGCGAACTGTCGCCGGGCAAGCCGCCCAAG
GTAAGCCTTGGCATCGGCAGCCGGCGCAGTGTATCAAATCCTGCGGCGGCATCTGCAGCGACAGGTTCGACGCGAACGTCAAACGATGACGCACCGGCCG
CGCCGGTTCCGCCCAAGATGCCGTCTGCCGGGCGCATGAACATGTCAAACCGGCAGGACGCTCCGCGGCCGGCCAGCGAGCCGGCAATGCCATCAGGCTC
GAGTGTGAGCGGTCAGAACCAGCGCTCCTCGAGCGGTGAGCGCCATGTTCAGGTCGAGGGCAGCCTCAAGCACCAAAGTGGCTTGAACCCCAATTTCACC
TTTGAAACCTTTGTCGAGGGCAAGTCGAACCAGCTTGCGCGCGCGGCGTCACGCCAGGTTGCCGAGAACCCGGGGGGCGCTTATAACCCCTTGTTTCTGT
ATGGCGGAGTTGGCCTGGGTAAGACGCACTTGATGCATGCGGTCGGTAACCACCTGGCGGGGCTCCGTGATAACGCTCGCGTGGTGTATCTGCACTCCGA
GCGCTTCGTGGCGGATATGGTCAAGGCGCTTCAGCTCAATGCCATCAACGACTTCAAACGCTTCTATCGAAGCGTCGATGCGCTTCTGATCGATGACATT
CAGTTCTTTGCCGGCAAGGAACGCTCGCAGGAAGAGTTCTTTTATACCTTCAATGCGCTGCTTGAAGGCGGACAGCAGATGATCTTGACCTCCGACCGCT
ATCCAAAGGAGATCAGCGGTGTGGAGGAGCGATTGAAGTCTCGTTTCGGGTGGGGGCTAACGGTCGCGATCGAGCCGCCGGAGCTCGAGACTCGGGTGGC

>Larsenimonas suaedae DSM 22428 [NZ_JARWAO010000001.1:40697..41196], <mnmG oriC2 <PRK13623 (ErpA), 568 bp

GTGCGTTTTCCAATCCTTCGATGGAGCGCACCAGGGCCAGCTGGACATCGAAGGGTAACGACGTCGAAATGCCGTTTGGATAGAGCTCATGGGTGTCGAG
CCCTTCAGGTTCCAGAAAGATCTGATGGCTCGTCTTGTCGGCAAATCGATGGATCTTGTCCTCGATCGACGGACAGTACCTCGGCCCTACGCTTTCGATG
CTGCCCGAGTACATCGGAGAGCGGTCCAGATTCTTGCGAATGATGTCGTGAGTGCGCTCATTGGTGTAGGCAATATGGCAACTGACCTGAGCCGGATGTT
GATCCCGTGAGCCCATGTAGGACATGACCGGAACGGGTGTATCGCCCGGTTGCTCGGCAAGTCTTGAGAAATCCACGGTGCGGGCGTCGATTCTGGGCGG
CGTGCCTGTCTTGAGACGACCGACGTTGAACGGCAGTGCTCTGAGTCGCTGCGCCAGTGCATTGGCCGGGGGATCGCCTGCGCGTCCCCCGGTCTGACGC
TCCATACCGATATGGATAATGCCGCCCAGGAAGGTCCCTGTGCAGAGCACCACGGTTTGACTTGTAAACCGAATTCCGCCGCTGGTCACGACTCCTTTGA
CACTTCCGTCTTCCACGATCAGGTCATCAGCAGCCTGTTGAAATAACGTCAGATTTTCTTGGTTTTCCAGCATGTGCCGGATCGCGGCCTTGTAACGCAC
TCGATCGGCTTGAACGCGTGTTGCACGAACAGCGGGCCCCTTGCGTGCGTTCAGTACGCGAAACTGGATACCCCCTTGATCCGTGGCAAGTCCCATGGCG
CCACCGAGCGCATCGATTTCCTTGACCAGATGACTCTTGCCGATGCCCCCAATGGCGGGATTGCAGGACATCTGCCCGAGCGTCTCGATGTTGTGTGTCA
GAAGTAGGGTTTTGGATCCTGTACGAGCGGCGGCAAGTGCGGCTTCGGTTCCTGCATGGCCACCGCCGATGATGATGACATCAAAGCGGTCCGGATAATC

CACGGCGCGACCTCGTCAATACGAATGGGAATGGGGATAACTGTGTTCCTGAATGGCGCGCATTATAACGCCCCTGTGTGTAACCGTCAGTCTTTTCCAC
ACCTCTCGGGCTTCACGCTCAAACGACCTTTCTATTAAATATAAAGAGAATTATATAAAGAAGTTTTGTAGTGGTTGTGATTGTAGAGTGCGCCTTTTTC
**TGTGGTTAA**CTCACATTCTTACCCACAGAACAATGGGTTGAGCTGTTAAAAATAACAGTGAAAACTGCTTTAAGCCCTGGTGAGGACGTTCTAACAAGCT
GTGCATGAGTTGCCGTTTCATCCACAGGCGCATTGTCCACGCGTTTATCAACGTGCGTATACCGGCTCTGTTACCAGGGTTGTCAAGAAATACGAATATC
CAGACATCAAGGCGTACCGCCCGTATGACGGGGGTAATATGAAAATCTTCTCAAAAGTTATCCACACCCTTTTTGTGAAAAAAAACCGAGAAAAAAAGGA

AAATAAAAAAAGCGGGCCGTAAGGCCCGCTTTTTTTATGCGCATTTGGTTCTGGCGATGTATCGGTGAGGGTTAGACCATGAAGGAGGCTCCACAACCGC
AGGTCGTAGTGGCATTGGGGTTCTGGATCAGAAAGCGAGCGCCTGCTAATCCTTCTTCATAGTCAACGGTCGATCCGACCAGATACTGATAGCTCAACGG
ATCGACGGCGAGAGTCACATTGTCACTGGTAATCAAAGTGTCGTCTTCTGCGGTGGTGTCGGTTAGATCGAAGCCATACTGAAAACCTGAGCAACCACCG
CCGGTAACGAATACCCTGAGCTTCAATGCTGTATTTTGCTGCTCGTGCATCAGCGATTGGACGCGAGCAACGGCACGATCCGTTAGTATCAAGGGAGTGG
GAACAAAAGAACTTACCCCGCTCATTGCAGACTCCTGCGTTCGCGTGGAAATAAAGACCATTATCGTGGGCTTTGTTGCAATGGGTCAAACGTGGATGCC
CAATAATCCACCATGGTTGATGTGTTGATCGAACGCCAAAGGAGAACGACGTGCGTCGTCTGAGCAAGCCTCATTTCACGCTGGATACCTTTCGTCGCCA
GCTGGCCAGTATCGACGCACTGCCCCAGCTTTGTATTCTCGGAGTGCTGTCTGGAATCCTGACCGGTGCCATCATGGTCGGGTTTCGTCAAGTTATCTCC
ATAGGCGCGGCATTGTTCATGCCAAATGGCAATTCTGAAGCCTTCGAGCAGTTGACACCCTATATCCGTACAGGGCTTCCGATCGTGGTTGCCTTGGTTA
TCATGGGAGGGCTCGCCTGGCAGCCTCTTCGCCGCAGAAACCTCGGTATCGGTCATGTCATCGAACGCCTGACGTACCATCAGGGGCAGATGCCCAGCGC
CAACTGGCTCAACCAATGGTGGGTTGGCGTCCTGTCGGTGCTTGGTGGACTTTCCGCTGGCCGTGAAGGCCCGGCCATCCACCTTGGCGCCGGCGCATCA

>Larsenimonas suaedae DSM 22428 [NZ_JARWAO010000001.1:70941..70999], dnaA> ig dnaN>, 59 bp

WebSkew GCmin pos. 4176400/0 (GCmin<>oriC1 37.8 kb; GCmin<>oriC2 5.9 kb; oriC2<>oriC1 31.9 kb)

>Cobetia marina JCM 21022 [NZ_CP017114.1:c4138362..4138861], <rpmH oriC1 dnaA>, 841 bp

CCGGTGGAGGGTGAATCAATCTGGCGGGAGTCAGATGAGTCCGGGTACGAACGGTGGGGCGCATAACGAAAAAAACGCCACATGCCGGACGCACCGGAGA
GGTGTCGCGTCAGGCAGGTGGCGTTGACTTGGCCGCTTTCAGACGTCTGGCATCCTTCTCCAGACGAGCCCACATGCCGTCAAGCTGGCGATGCAGTGTC
TCGTTGTCCAGCTCGTTCACACCTCGCTTGGCCAATACGACGATGTCGACGGCCGGAAGGCGATGCTGGCGGAGGCGAATGGATTCGCGTGTCAGACGTT
TGAGGCGGTTGCGATCTACGGCACGGCGCACGTTCTTCTTGCTGAACACCAACCCCAGGCGGGCATGTCCCAGGTCGTTCGGACACGCCAGAGCCAGAAG
CCCCTTGCCGTGTACCTTGAGGGTGGTGTGATCGAAGACGTGCCGGTAATCCCCGGCAGTCAGCAAGCGAAGCTGCCGGGGAAATCCCTGACGGGACATC
CGGGCCTCGATCAGGCGCTCAGACGCTTACGGCCTTTCGCACGACGACGCTGGATGACCTCACGGCCACCTTTGGTCGCCATACGAGCACGGAAACCGTG
TGTGCGCTTGCGCTTCAGGACGCTCGGCTGAAATGTGCGTTTCATAGCTGCTTCTTCCCACGTGGTTGTTTGGAAAAGGTCAACGAATGTTGCCCTGTAT
CCACATGAGGACAGGGCAGATTGGTCAGGGGCAGGGATTCTAGTGAATCTGCAAGTCGCTTGCAATGTCGCGTGAGAAGTTGTCCACAGCCGTTTCCCGA
CATTTCACTGCTGATTCGTGCCACGCCGCCCGTCAGCTGGCGTTTCTCACGGCTATCGAGCCAGCCACGGAGCACGTGTGCAGGGCCAAAAAAATCCTTG
TCGAGAAAGGAGAGGAGAAACTGCGCCTCATGCTGACCGAGGGGAAGACTTGCGAAAGGGGGGGGCGGCGACTCGTGCATCGGGTCCCGCTCTTTCGGGG

GATAGCCCCTGCCTCCCGAGCTGTCTTGCCCGACCAGCGAATGACAGATTCGCGGCAGGAAAGCACCGGTGGATAACCTTGCCTGTGCGTGCGTCGCCAC
CTCCTCGAGAGGGAAAGCGCATTCCCGAGCAAGGCCCTCGCTATCATTGCTACAGAAAGGTAGTTCTAAATATACAGTAACTATAAAGGAGGGAAGATTC
**TGTGGATAA**CCTGTTTTTGTCGAAGAAGTTCAATCTACTGCCCTCAGGATATCCCTGATGATAAAGGGTGAAGAACCTGAGTTCAGCCTGTGCATGACAC
TGTGCATAAGTATCGCCTTATCCCCAGCGGTCCAGATGGCCTCCATCACTCCCCAGAATGTCCAGATGGCAGAAGAATCTTGATACCCAGGTTGTCCACA
GTGGATTTCCCTGCCGATGTCGTGTGGATAACTCGTGGCTTGGCCTATACAATGGCCGCCCCTTTGATCACTGGATGTGAGGTCGCGTGTCGCTCGCACT

TTGGCAACAATGTCTTGACGCCCTGCAAGACGAGCTGAATTCACAGCAGTTCAACACCTGGGTTCGTCCCTTGCAGGCTGAGGAAGGTGACGGCGGAGAG
CTGTGCCTGCTCGCCCCCAATCGCTTCGTACGCGACTGGGTCAATGACAAGTACCTCAAGCGGATTCATGAGCTTTTGCGTGAGCTTGCTCCGGGCAAAC
CGCCCAAGGTCGAGCTCAGCGTGGGAAGCCGCCGCACGGTCTCCAATCCGCGGCCAGAGGGGGCATCGACTTCATCGGTGTCCGCTCAGGCCCCTGCGCC
CGGCCAGATGCCGAGCCCGCGTGCGCCGTCAGTGCATACGCCCCCGCGTGGCAATTACGCAGAAGAGCGTGAGATGGCGGGTGACGGTGCGCAGGAGGCG
GCACCGCGTCGCAATGCGCGGCAGGTGCAGGTAGAAGGCAGCCTCAAGCATTCCAGTGGCTTGAATCCGAATTTCACCTTCGAAACCTTCGTCGAAGGCA
AGTCCAACCAGCTGGCACGTGCCGCCTCCCGTCAGGTCTCTGAGAACCCGGGTGGCGCCTACAACCCGCTGTTCCTCTATGGTGGCGTCGGTCTGGGCAA
GACCCACCTGATGCATGCGGTGGGCAACCATCTGGCCAAGCAGGGTGGCGAGAATGCCAAGGTGGTCTATCTGCACTCCGAGCGCTTCGTGGCTGACATG
GTCAAGGCGCTGCAGCTCAATGCCATCAATGACTTCAAGCGTTTCTACCGCTCGGTGGATGCGCTGCTGATCGATGACATCCAGTTCTTTGCCGGCAAGG
AGCGTTCTCAGGAAGAGTTCTTCCATACCTTCAATGCCTTGCTGGAAGGGGGACAGCAGATGATCCTGACCTCGGATCGCTATCCGAAGGAGATCAATGG
TGTCGAAGAGCGCCTGAAGTCCCGCTTCGGTTGGGGGCTGACAGTGGCCATCGAGCCGCCGGAGCTTGAGACTCGTGTCGCGATCCTGATGAAGAAGGCG

>Cobetia marina JCM 21022 [NZ_CP017114.1:c4170256..4170755], <mnmG oriC2 <COG1126 (glnQ), 1388 bp

CGTCACCCGGCTGAGCTTCCATCTTGCTGAAATCAACGCTCTTGGCATCGATGCGCGGCGGGGTACCGGTCTTGAGCCGATCAACCCGGAATGGCAGCTC
GCGCAGACGTGACGCCAGGCCATTGGCGGGCGGATCCCCGGCACGTCCACCGGCATGGTGGTCCAAACCGATATGGATCACGCCACCGAGGAAGGTCCCG
GTCGCCAGCACCACCGTGGTGGCGTGGAAACGAATGCCGGTCTGAGTGACGACACCCCGTACGGTCTCGCCTTCGACGATCAGGTCTTCCGCACCCTGCT
GGAACAGCGTCAGGTTCGGCTGGTTCTCCAGCACGTTGCGAATCGCCGCCTTGTACAGCGCCCGGTCGGCCTGTGCTCGAGTGGCACGTACCGCCGGGCC
CTTGCGGGCATTCAGCGTGCGGAACTGGATACCCGCCATGTCGGTTGCCAGGCCCATGGCACCGCCCAACGCATCGATCTCCTTGACCAGATGGCTCTTG
CCGATCCCACCGATGGCGGGATTGCAGGACATCTGCCCCAGGGTCTCGATGTTATGGGTCAGCAGCAGGGTCTGACGGCCCATGCGGGCAGCCGCCAGTG
CGGCTTCCGTACCGGCATGACCGCCGCCGATGACGATCACGTCAAAGCGGTTCGGGTAATCCACGTTGGCACCTCGGGGTATCACGAAACTTGGATGAAG
CGCTGTTCGCATGGTGGCGAATCAGGCAAATCGGCCAGACACGACGATGACTCGATGTCTGGATGAATCCGGGACTGACGAACGTCATGGCGAAGACAAG
GCGCCGCAGACGGCATGAGCTCGACCCGATGGGGAGCCTGCAAGCCAATCGGATGACGTTCGGGCCGCACAGTATAAACCGCCTGTGGGTAAGCGTCAGG
AGTTTTCCACAGTGCCATGGCAGGACGTGCCATGCAGTGGACTCGCGGATGTTTCTCTCAGGGCATCAATCACGACAGTGAGTCGAGCCCGTGTCGAGTG

TGTGTGGCGTAGGGGATTTTCCAGAGGACTGGGTCGGATCGCCCTGCCGTTCGTTATCCCGAACACCTCGGGAAGTGAGCTTCCTCGAACCCGGTCTTTC
GGGGTGTCCGAACGTCCTTCGGGTTTCCGGATAACAAGATTATATAAAGAAAAGAAAAAGGGGTTCTGTTGATGTTGTAACTACTGTTGTGGACAAAATC
**AGTGGGAAA**TCGACTTTATCTCTTAAAAATCAGTATCTTACGATGTTGATATAGTTGGGGAGAAAATGCGTGCTACCTGAGGATGCCACCGGTGGCTTCC
TGTGGATGAAACACCGCCTTGTCCACAAGCCCAGTTATCCTCGTTTTCATACCGTGCTGGTACCGGACTTTTCTGCCCTGCTGTGCACAAGTGGCCAGTT
TACCCACAACCCCTAGTGCTGCAAGGGCTAACGTACAGAAGGCCGCGGATGATGCCGCAAAATGTCCACAGAGGCAAATCGTCTTCACCTCGACACGCCA

ACTCTGTAGATAGTACGGGGATAAGCCGTGAGTGGATGGCTACGCTGCCCAGCAGTGGTGCCCTGAGAGATACGATGATACTGCTGTGGATAGTTGTTGG
CGTGATTGAGGCTTGCTTACCGGTGCCTGGCCGGCCAGACGACTCATGGGCTGACATCAGGCGTGTGTTATCACCAGCCACGGCCCCTTGCAGGCGATGT
TTCGAGCGCCTTTCGCGTCAGGACATGCGGTGTGAAAGACGGGTCCTTCGGCGACTGATGAAGTGAATCGAGAGCCACAGCGAGGCTGGGGGCGTGACGT
GAGAGGCAGCCAAGGCGGGGCGAATCATCTGATTTCATTGCATGAAAAAAGCGCGAAATCGTCAGTCGAGATCGAAAAACTTCATTTCGAATGCCAGAAC
TCCCGTCATTCGGGCCACAACAGACACGTCTATCGCCAGGTGATGCTCTGGATTGGCCAGGAATTGCCTCTTCACGGCGTGGCATTCGGGCAAAAAAAG
CCCCGACCTGACGGTCGGGGCATGGCCTCTCTCAAGAGATGGCAGCGGGGGTTCAGTGCTTGAGAACGCGCGCCAGGAAGCTCTGGGTGCGCGGATGACG
AGGGGTGTCGAAGACCTCGCTCGGAGTGCCTTCCTCGACGATGCTGCCCTGGTGGACGAAGACGACGCGATCCGCGACTTCACGCGCGAAGCTCATCTCA
TGCGTGACGATCATCATCGTCATGCCGTCGCGTGCCAGCTCACGCATCACGTCCAGCACCTCCCCGATCATCTCGGGGTCCAGTGCGGAGGTGGGCTCAT
CGAACAGCATCAGGCGGGGTTCCATCGCCAGTGAGCGGGCGATGGCGACGCGTTGCTGCTGGCCGCCTGACAGCTGGCCGGGGTACTTGTCGGCCTGCTC
GAGAATATTGACCCGCTCGAGCAGCGTGTCGGCCTGTCGCTCGGCGGTCGCCCGGTCACTGCCGCGCACCTTGATGGGGGCCAGGCTGACGTTGTCACGT

>Cobetia marina JCM 21022 [NZ_CP017114.1:c4136740..4136887], dnaA> ig dnaN>, 148 bp

TTCAGTTGCCACGCAGCGCCGAGAGGGTGAGGTCAACCTGTCCGGCGCTTAGTGCACCGGGCCCTGTCGTGCGACGGCCCGGAATATTTGATGAGTGTGT
GCTGGTCACCGGCAAGCACTTCTCCCGTTTCCAGAGAGTGGAGTCTCC

WebSkew GCmin pos. nd (GCmin<>oriC1 nd; GCmin<>oriC2 nd; oriC2<>oriC1 nd)

>Cobetia crustatorum JO1 [NZ_JEMF01000015.1:31525..32024], <rpmH oriC1 dnaA>, 943 bp

GGCAGGTGGCGTTGACTTGGCCGCTTTCAGACGTCTGGCATCCTTCTCCAGACGAGACCACATGCCGTCAAGCTGGCGATGAAGGGTCTCGTTATCCAGC
TCATTTACGCCTCGTTTGGCCAGAACGACAATGTCGATGGCCGGAAGGCGTTGCTGGCGGAGGCGAATGGATTCTCGCGTCAGACGTTTGAGGCGGTTGC
GATCTACGGCACGGCGCACGTTCTTCTTGCTGAACACCAACCCCAGGCGGGCATGCCCTAGGTCATTCGGACAGGCCAGGGCCAAAAGCCCCTTGCCGTG
TACCTTGAGGATGGTGTGATCGAAGACGTGCCGGTAATCCCCGGCAGTCAGCAAGCGAAGCTGCCGGGGAAATCCCTGACGGGACATCCGGGCCTCGATC
AGGCGCTCAGACGCTTACGGCCTTTCGCACGACGGCGCTGGATGACCGCACGGCCACCTTTGGTCGCCATACGAGCACGGAAACCGTGTGTGCGCTTGCG
CTTAAGGACGCTCGGCTGAAATGTGCGTTTCATAGCTGCTTCTTCCCACGTGGTTGTTTGGAAAAGGTCAACGAATGTTGCCCTGTATCCACATGAGGAC
AGGGCAGATTGGTCAGGGGCAGGGATTCTAGTGAATCTGCAAGTGGCTTGCAATGTCGAGTCGAAAAGTTGTCCACAGCTGGTTCGTGGCATTTCAGGGT
GACGATGCCTCGCGCCGCCCGTCAGATGGCGTTTCCTGTCTACCGGTGACGGGCGGAATTTGGACGTTGGCAGGCAAAAAAAATCAGTTTTTCTGACAG
GCACTGCCTGGGGAGCTGAGGTGTGGCCAGAGAGACTTGCATTCCAAGCGAGGAATACACGAAACCCAAAGTGGCGAGACTTGAAAGACAAGACCGAAAA
ACTGAAGTGGTTTCTCATGCTAAACCAGCAGATATTTTTGCTGCGCTGCGGTTTCGTAAGGTGCTGACTTCGCCCATCCGGCATGAGGCAGAGATGCTCG

TACGACCCCACGATTTGTGTAGCAGGCCAGTCAGCGATGTATGCATCATCGAGCATCAAAGGCCCAGTGGATAACCTTGAGTCGACCTGATCAGGAGTGC
AGCGCTGAGCTTGATCCGAGACGCATTTTTAGCATTGCCCCTATCATTACTATAAAGAGGTAATTCTGAGTAACTAATAACTATAGAGGGAGGGAGATTC
**TGTGGATAA**CCTATTTTTCTAGAGTTAGCTCAATCTTCTGTCATCAGGATATCTCTGACGATAAGCAGTGGAAAACCTGATATTACCCTGTGCATGGCTG
TGTGCATAAGTGTTGACTTGTCCCCAGCGTCTGAGTCGTCTCATATTGCTCCCCAACCTTTCCCAAAGTAAGAGGGTCCTCAATACCCATGTTGTCCACA
ACAGCTCATGACGCAGATGTCGTGTGGATAACTCAGGGCTTGAGCTATACAATGGCCGCCCCTTCGACAACTGGATGTGAGGTCGCGTGTCGCTCGCACT

TTGGCAACAATGTCTTGACGCACTGCAAGACGAGCTGAATTCACAGCAGTTCAACACTTGGGTTCGTCCTCTGCAGGCTGAAGAAGGTGACAGCGGAGAG
CTGTGCCTACTCGCTCCCAACCGCTTCGTGCGTGATTGGGTCAATGACAAGTACCTCAAACGTATCCACGAACTACTGCGTGAGTTGGCACCGGGCAAGC
CGCCCAAGGTCGAACTCAGTGTGGGTAGTCGACGTACGGTGTCCAATCCGCGACCGGAAGGAAGCCCCGCGTCTTCTTCATCCACTTCTTCATCTGCTCA
GGCACAAGCGCCTGGCCAGATGCCGAGCCCGCGTTCGCCACATGTGCATACGCCACCGCGCGGTAATCTCGGCGAAGAGCGCGAGATGGCTGGTGATGGC
TTGCGCGATAGAGCCCCGCGTCGTAATGCACGCCAAGTGCAGGTCGAAGGCAGTCTCAAGCACTCCAATGGTTTGAACCCGAACTTCACCTTCGAAACCT
TTGTCGAAGGCAAGTCGAACCAGCTGGCGCGTGCAGCATCACGCCAGGTATCCGAAAATCCTGGTGGCGCTTACAACCCGCTATTCCTTTATGGCGGCGT
GGGCTTGGGTAAAACCCACTTGATGCACGCAGTGGGCAATCATCTCGCCAAGCAAGGAGGCGAGAATGCCAAGGTGGTCTACCTGCATTCAGAGCGCTTC
GTGGCCGACATGGTCAAGGCGTTGCAGCTCAATGCCATCAATGATTTCAAGCGCTTCTATCGTTCAGTCGATGCGCTGCTGATTGATGACATTCAGTTCT
TTGCGGGTAAGGAGCGCTCTCAGGAGGAGTTCTTCCACACCTTCAATGCGTTGCTTGAAGGCGGCCAGCAGATGATTTTGACATCGGATCGCTATCCGAA
AGAGATCAATGGCGTCGAAGAGCGACTCAAGTCCCGCTTTGGCTGGGGGCTGACTGTCGCCATCGAGCCACCTGAGCTAGAGACGCGCGTCGCGATCCTG

**oriC2** not in genome sequence, only flanking regions

>Cobetia crustatorum JO1 <mnmg ig | [NZ_JEMF01000037.1]

```
     gene            complement(19749..21695)
                     /gene="mnmG"
```

.............................................................ATTCAGCACGAGACGTGAACATGCGATACGGCTCGCGTG
TTCCGAGCGTAATCAGGTCATCGACCATCACGCCGGTATACGCCTCATTACGTAGCGGATGCCAAGCTTCGTCGTCTGTGGCACGCAGCGCCGCATTTAC
ACCGGCCAGCAGACCTTGTGCGCCAGCTTCTTCGTAACCAGTGGTGCCGTTGATCTGCCCTGCAAAGAACAGATTCTTGACAAAGCGCGTCTCAAGAGAG
TGCAGCAGATCCTGGGGGTTGAAAAAATCATATTCAATGGCGTAACCAGGACGCGTGATGTGGGCGTTTTCCATGCCACGAATCGAACGCACTACCTCGA
GCTGCGTATCAAAGGGCAGCGAAGTAGAGATGCCATTGGGATACAGCTCATGAGTGTTGAGACCTTCAGGCTCAATGAATACCTGATGGCTGTTCTTGTC
TGCGAAACGGTGCACCTTGTCCTCGATGGACGGGCAATAACGCGGGCCGACACCACCGATCACACCTGAATACATCGGCGAACGATCAAGATTGGCGAAG
ATAATCTCGTGAGTCCGCTCATTGGTGTGAGTGATGTAGCAGTTGACCTGCTCAGGATGCAGCTCACGACTGCCCATGTAAGACATCACCGGACGTGGCG
TATCGCCGGGCTGCGCTTCCATCATGCTGAAATCGACACTCTTGGCATCGATACGCGGCGGGGTTCCTGTCTTGAGACGATCGACCCGGAACGGTAGCTC
ACGAAGTCGAGCGGCCAGACCGTTTGCCGGCGGATCCCCTGCTCGTCCTCCGGCATGATTATCCAGACCGATGTGAATCACACCGCCAAGGAAAGTACCT
GTTGCCAGAACCACCGTGGTGGCATGGAAGCGAATACCGGTCTGAGTGACCACGCCCCGCACGGTATCACCCTCGACGATCAGATCTTCAGCGCCTTGTT
GGAACAGCGTCAGATTCGGCTGATTTTCCAGCTGACTGCGAATGGCGGCTTTGTACAGCGCCCGGTCTGCTTGAGCACGAGTGGCTCGAACCGCAGGCCC
CTTGCGGGCATTCAACGTACGGAATTGGATACCCGCCATGTCGGTGGCCAGACCCATGGCACCGCCCAGCGCATCAATTTCCTTGACCAGATGGCTCTTG
CCAATCCCGCCAATGGCCGGATTGCAGGACATCTGGCCCAGCGTCTCGATGTTATGAGTCAGCAGCAGGGTCTGGCGACCCATGCGGGCAGCAGCCAGTG
CGGCTTCCGTACCGGCATGACCGCCGCCGATGACGATCACGTCAAAGCGGTTCGGGTAATCCACGTTGGCACCTCGGGGTATCACAAAACTTTGGGATGA
AGCGCTGTCTGCACAGTGCAGATCAGGCAAAACGGCCAAATCCAATGTTGGATCGGCGTCTGGCAGACGGCTGTGACGGATGAATGATCGTCACCGATAG
GACCTGCTCGACCCTGGTTGGGGAGCTGATGTTCAGAATCGAGATGACGTCTGGGCCGCACAGTATAAACCGCCTGTGGGTAAGCGTCAGGAGTTTTCCA
CAACCGCCGGGCTAGACCGATGTGATGTGGACAGCCTCCACGGTGCCGATACCGCACCAATCGGTCAATCATCGGCTATCGCTTGAGGCTTCGATTTTCG
AAGTGAATCTATCTGCGTATCGAATGAATCATCCGGGTGTTCGTTATCCCGAACAGATGCATGGCGAAGAGATCATATTGATCCCGATAGG

>Cobetia crustatorum JO1 | ig <COG1126 (glnQ) [NZ_JEMF01000015.1]

```
     gene            complement(573..1319)
                     /locus_tag="BF12_RS0103205"
```

CCCTAGTGCTGCATGGCCTAACGCCAAGATGGTCGCGGATAATGCCGCAAAATGTCCACAGAGGCAAATGACATTCTTTCTTCATGGCAAAGTCTGTAGA
CAGTTCTGGGATAACCGGTGAGTGGAGTGCTGTAGACCGCTTGAATGGTGCCGATTGAGCGACGACAAGTATTTTGTGGATAGTTTTGGGATGTGATTCG
GCAAAGCACTATCGTGGATGCCGTGATTCCAGGATGGGAGCGTAGGAAGGGCGTGTGCTTTACCGTCCTCACACGCTGAATGGCGATCAATGTGGCTCTG
CTGCGAAGTGATAAAGCGAGCGTGGAGGCGAGTAAACAGGCGCCCGACATGCGAAGAGCGCCATCTGACACGTGACGATAGACGAGTAAGGCAACGTTTG
TTCGTTCGTTTTTATTGGCTGAAAATAAAAAATATATTTAATAGAAGCGAAAATATTCATGCGAATGACGAGGTTTACGCCAAGGGGGAGCCCCGCTAGC
GCGCAAAAAAATGCCCCGATCAGGAATCGGGGCATCAGATGGTCCTCATCTGGGATGTTTTCTCGCGGCGACCTAGTGTTTGAGCACTCGCGCAAGGAAG
CTCTGAGTTCGAGGATGTTGCGGATGATCGAAGATTTCTTCCGGCCTCCCCTGCTCCACGATAGCGCCCTGGTGGATGAATATGACTCGGTCAGCGACTT
CGCGGGCAAAGCTCATCTCGTGCGTCACGATCATCATCGTCATGCCTTCACGGGCCAGCTCACGCATTACGTCGAGGACCTCGCCGATCATCTCGGGGTC
GAGTGCTGAGGTGGGCTCATCGAACAGCATCAAGCGGGGTTCCATCGCCAGTGAGCGTGCAATCGCCACCCGCTGCTGCTGTCCACCGGACAGCTGTCCG
GGGTACTTGTCAGCCTGCTCAAGGATATTGACCCGCTCCAGGAGTGCATTGGCCTGTTTTTCGGCTGCGGGGCGATCAGTGCCACGTACCTTGATGGGTG
CGAGGCTGACGTTGTCGCGTACGTTCATGTGAGGGAACAGATTGAACTGCTGAAAGACCATGCCAACTTCGGTTCGTACTGCCTTGAGGGCATTGTGACT

>Cobetia crustatorum JO1 [NZ_JEMF01000015.1:33511..33661], dnaA> ig dnaN>, 151 bp

TCGCATTGCAGCGGCGGGTAGGGTTGGCGACAATTGTCGATTTGACTCGCTGTGCGCATTGGGCCTTGTATCCCGTTGGCCCTGAATCTTTTTTGAGTGC
GCGCTGCTGTGGCAGCGAGCACTTGCCCCGTTTCCAGAGAGTGGAGTCTCC

WebSkew GCmin pos. nd (GCmin<>oriC1 nd; GCmin<>oriC2 nd; oriC2<>oriC1 12.9 kb)

>Terasakiispira papahanaumokuakeensis PH27A [NZ_MDTQ01000001.1:c3609151..3609650], <rpmH oriC1 dnaA>, 568 bp

TTCACCCGTAGCTTATGATCAAGCAGCGCGGCATAAACAATATCTCCGCCTTTGGGTGAAATTTTCAGATCTAACAAGTCAGTCCGCACATTAATCGTGG
ACTGATCAGTCCCCGCTGCGCTGCTGTCTTGAAGCTCTGTTGTCGATTGAGTCTGACCAAACTGGAGCGTCTCATCATCACTGCCCGAAGGCTGTGCCGC
GCTGCCATTGTCGGTCTGTACTTCAGCACTCGTAGGCGCCTGGGGCTCGTGATAATCCTGATTCCATTGGATTATCATCAAATAGCCCACTACGGCCAGT
GCCAGATAGAGAACGATGCGTTTCACATCCATGGGCTGACTGCCACTCTTTTGCTGGGTCTAGTAATACACGAAAAGGCCGAACATCGGGGGATGTTCAG
CCTCTTCAAGCCTGATGCTTGGCTGCTCGATTCCTTGTCTTGGGCGCTTGACCATCCTGGTGAGACTTTGTCTGCGCTTTTCGACAGAGCTGTCGCCATG
CCTTATCGAGTTGACCATGAAGTTGCTGCGTATCGATGTTCGCAGCCCCTTTCACAGCTAACACAATAATATCCAAAGGAGGTAAATCATGTTGATGGTG
ACGGAAAGAATCCCGAACCACTCGTTTAATACGATTACGATTCACCGCCAGGCGGACATGCTTTTTACCTATGACAAACCCTAAACGGGTTTCTTCCGTC
TGAGACGGAGCGGCTAACAGCAACAAGCCAGGACAATAGGCTTTATAGCAGGCCTGATCAAAGACATGACGAAAATCCCCGGCACTGAGTAAACGTCGTT
GCCGGGGAAAGCTGAATGAGACCACCCGAAACTGGGGATCAAGCACTCAGACGATGACGGCCTTTGGCACGACGACGAGCCAGGACTTTACGGCCATTTT
TGGTGGCCATACGAGCGCGAAAGCCGTGAGTACGCTTACGCTTAAGAACACTAGGTTGAAAAGTGCGTTTCATGACGTCTAATCCCACTTTGAGGTTGAT

GTAGCCACGCCCAAAATGGGCGTAATTAGCTCATGCAAGGGCGAGGATTCTAGAGAATAACTGGACTGAACGCAATTTGAACGTCACTTCAATTGCAATT
TTCTGTATTTCAGACCCCTTGTGCACAACCTACTATAAACATAGATAAGAAAAGGTTTAAAAGCTTATTAGTTATGACTATTACTAGTGCTTTGCATTGG
**TGTGGATAA**GCATGAAAAAACCTTTGAATTCTTCATGTTATAGAGTTATTAACACTCTTGATGACAACCGTATAGGCTGTGCTGCAACTGTCGGAGTGCT
GTGGATAAGCAAGCTACTTATGCACACCTCAAGGTATCCCTCTATCTATACACAGTGCTAAAGGCGGTTTTCAAATAGGGTTGTCCACACCCTAAAAAAA
TTTTAGAAAACGTCGTTTTAAGGCCTGATGTGATGCCCAATTGATTTTTTAAGAATAAAAATGACCTGGAAGTGATGGTGCGCTGTGGATAAGCCGGGGT

GGGTCACGTAGAATGAGCCGGTTAAATTCAGGAGCCTTGATGTGACGTCAGTCGATCTATGGAACCGCTGTACCGCCTACTTGCAGGATGAACTGCCCGA
ACAGCAGTTCAATACTTGGGTCAGGCCTTTGCAGGCAGAATTGGAAGGTCAACAACTTGTATTGTTGGCACCGAACCGTTTTGTGATGCAGTGGGTTGCC
GATAAATACGCACCGCGTATTCGTGAGTTACTTAAAGAGCTGGGCGGGGATCAACCCCTTCAGCTTAAGCTCGGTGTGGGTAACCGACAGCCTCGGGCAC
CGGTTACGCCACCGGCACCTCCGGCTGAACCAAGTTCTCCGACACCTTCTGCCAGTGCGCCAGAAGATTATCGCGATGTCATGCATAAAATGGCTGGTGA
AATTGGACAACCTGCGCCTTCTTTGGCCGCACAGCCCGTGGCCTCTGATATACCTACTGAGCCTCGAGGACGAGATAGTCGAGAGCCGTCAACCAGTCGT
GAAGAGCGTCCGGTTCAGGTTGAAGGGGCGATTAAACATAATTCGACTTTAAACGGCCAACATACTTTCGATACCTTTGTTGAAGGTAAATCAAACCAGT
TGGCTCGAGCGGCTTCGCGACAGGTCGCAGAGAATCCGGGAGGGGCTTACAACCCACTCTTTATCTATGGCGGTGTTGGGTTGGGTAAAACTCACTTGAT
GCATGCTGTTGGTAATCAGATCCTGAAGCAGAATCCTAATGCCAAGGTGGTCTACCTGAACTCTGAACGCTTTGTGCAAGATATGGTGAAAGCTTTGCAG
TTGAATGCCATTAATGAGTTCAAACGCTATTACCGAGGCTTAGATGCTTTATTAATTGATGACATCCAGTTCTTTGTCGGCAAAGATCGCTCTCAGGAAG
AATTTTTCCACACCTTTAATGCGCTATTGGAAGGCGGCCAACAAATTATTCTGACTTCGGATCGCTTCCCCAAAGAGATTGGGGTAGAAGAGCGACTTAA

>Terasakiispira papahanaumokuakeensis PH27A [NZ_MDTQ01000001.1:c3622074..3622573], <mnmG oriC2 COG3713>, 541 bp

ATGGCATATCCCGGACGCGTGATATGAGCGTTTTCCATCCCTCGAATCGAGTGCACAACCTCAAGTTGCACATCAAACGGCAATGAAGTTGAAATCCCAT
TGGGATAAAGCTCTTGGGTATTTAAACCTTCTGGCTCAATAAAAACCTGATGGCTGTCTTTATCGGCAAACCGATTCACTTTGTCCTCAATCGAGGGACA
ATAACGCGGCCCAACACCTTCAATGACCCCGGCATACATGGGGGATCGATCCAGGTTAGCGCGGATGAGCTCATGGGTTTGAGCATTGGTATGGGTAATC
CAGCAATTAACCTGCTCAGGATGTTCTTCAGGACGTCCCATAAACGACATGACCGGCATCGGATCATCGCCAGGTTGGGCATCCATCACAGAGAAATCGA
CCGTTTTCGCATCAATTCGCGGAGGCGTACCTGTCTTCAAACGCCCAACGGGTAAATCAAGTTCACGTAAACGATCCGCTAATGCGATAGACGGCGGGTC
ACCGGCACGACCACCGCTATGGTGCTGCATACCGATATGAATCACGCCGCCAAGAAAAGTCCCCGTGGTTAAGACCACGCTCTTAGCGCTGAACCGAATG
CCGCCTTGTGTCACCACACCGCGAACTTCATCACCTTCAACGATGAGATCATCCGCAGCCTGCTGAAACAGCGTTAAATTCGGCTGATTTTCTAGAATAT
TTCGGATCGCTTGGCGATACAGGGCGCGGTCTGCTTGGGCTCGGGTCGCTCTGACCGCAGGTCCTTTGCGCGCATTTAACACACGAAACTGAATACCGGC
TAAGTCAGTCGCATGCGCCATAGCACCACCTAGGGCATCTATTTCGCGCACCAAATGACTTTTACCAATCCCGCCAATAGCCGGGTTACAGGACATTTGT
CCAAGGGTTTCGATGTTGTGGGTTAACAGGAGGGTATCAGCCCCCATACGTGCTGCGGCCAGCGCAGCTTCCGTGCCTGCATGGCCGCCGCCGATCACAA

TGACATCAAAGCGGTTGGGATAATCCACAGCCACACCTCAAGAATTCGAGCAATTGGGAAAAGGGACAACAGTATAAAGATTCCTGTGAAATAAAAAAGT
ACCCAAGTCGGTGTGGATAGCTTTATCCGATATCCACAATAACAATCAATAGCTGTTTTAAAAGATTACTACTGTTATTATTAATAGTGTGGATACTTTC
**TGTGGATAC**TTTGCTTTATCTCCCTATA**C**AGGAGGCTTCGTGGCTTGGGATATGTTTGTTGTTAACTGCCTTATGAAGCGTGCGTAACTCGGTATAGACC
TGTGTATAACACCTTGTTCCATCCACAGCTCAAACTAGACCCAACTTATCTTCAGGGCTTAGGCCTATTTCTACCTCCAGATATCCACACCCATTTCGAC
ACACAACGACAGTATTGTTGTCACCCTATAAGCAGTATCTTGTCTCTTGAATGAGCTTGTATAAGCAGCTCATCAATTCTTCTGTTCGATCACATTGATG

ATGTATTAAAACAATCACCCATGTGCTCAATGCGACATGTTCTTAATTCAACATAAGGAGTACACCCGTATGACGACAGCCTATTGCCCTCGACGTCTGA
TGTTTTTAGCGGGTCTTGGTATGACAGCCTGGGGGTATGCCACAGGGGCCTATGCCGATACAGCCTGGACAGGATCAATCGGTGCGGGAGGCGTTTGGAA
ACCCGATTATCTCGGCAGTGATGATTATGAATTTCGAGGCTTTCCTAACTTAACGCTACAGGCCAAAGGCGTGGGTTACTTAAGTCTCAAAGACGGTATT
GGCACTTATTTGACAGAAAGTGACCATTGGCAAACATCGATATTCCTGACTTACTTATTTGGCCGTGACAATACAGGCGATCTATCGGATTTTGAGGAGG
TGGATGGCGGTATTGCTGGAGGGCTTCGACTGGGTTACAACCAAGGCGCATGGCAGCATAACTTATCCATTAAAGCCCCCATTGGTGGGGATTTAGAAGG
TTATCGAGTCGATGTATCGACTCGATGGAAAACAGCATTGTCTCAGCGCTGGTCCTTCTTTATTCAGCCGGGTATTGGCTATGCCGATAAACAATGGAAT
CAAGCCTATTTTGATGTATCCGCAGAGGATAGCGCTGCCAGCGGTGTTCGCGCTTATCGTGTTAGCGATGGCAGCTGGGATGCTCATTTCACGACCGGCG
TTAATTTCCAAATAACAAAAGAATGGCGTTTATCAATGGCAGGTCGAGTGAGCCGGTTGACCGGTGATGCCGAGGATAGCCCCATTGTGGATGAATTAGG
TGATGCGACTCAAGGCAGCTTGATGATTGGTACAGCCTATTCGTTTTAAAGCGATACGTTTTAAACTGACAGATTTTAAATCTATGGGTTTTTATCTAAG
ACCCCATTAAACGACTCCCATACAGAGTCTCTCAGTGTATTATCCGGTGTGTCACGTCACACCGGATAATGCAACATACAGTGGAAGCGCACAGGGACAT

>Terasakiispira papahanaumokuakeensis PH27A [NZ_MDTQ01000001.1:c3607560..3607594], dnaA> ig dnaN>, 35 bp

WebSkew GCmin pos. 64860 (GCmin<>oriC1 18.6 kb; GCmin<>oriC2 1.4 kb; oriC2<>oriC1 17.2 kb)

>Halovibrio variabilis DSM 3050 [NZ_BJXV01000011.1:c46049..45550], <rpmH oriC1 dnaA>, 534 bp

GAGGACATCGGTAGTCACCGCAATGAAATCGCGGCTAGACGTCTCGCTTTCGGTATCGCCGGGGATACCTTCACCCACGTTGCCTTGTGGCGCCGAGGTG
CTTGGCACCGCTAAGCCCTCGCTATCCTCAACGTTGCTGGGAGGCGCCGAGCTGCTGCGTTGCGTCGTTTCAGGTACCGAATCGTAATTCGGCTGCCCGT
AATCCTGATTCCACTCGACGACTAGCAGGTACGCCAGTACGGCTAGTGGAATCAGTAATAAAAGTCGTTTAACGTCCATGAGGGTTCTGCCCGCTGGGTG
GTGAACATGCTAATGGCGCAGTGTCATTTACTTGACTCTTGACACACGAAAGCCTGCCAGACGGCAGGCCGAGGTCGAGCGGCAAGGGGTGCACCTAATG
TCACGTGTTACGCTCCGGCGTTGGTGCTGCGATCATCGCCTGTGCGTCACGCTGAAGCCGCTTCCACATACCATGTAACTGGCGATGCAAGGTGTCGTTG
TCTATATCCTGAACGCCGCGCCGGGCAAGCACCACAATATCCACAGAGGGTAAGTGATCTTGACGAAGACGTAAGGATTCTCGAACGAGACGCTTAAAAC
GGTTGCGGTCGACGGCGCGCTTAACATTTTTTTTGCTGACCACCAGTCCAAGGCGAGGGTGTCCCCGGGTACTTAGGCGAACTAATGCCATCATACCTTT
GCCATGTACTTTTAGATCGGCTTGCTCAAACACGTGACTGAAGTCCCCAGCATTTAGTAACCGTAATTGCCGGGGAAAGCCTTGCTGCGGCACACGCAAT
CCTAGATCAAGCGCTCAGACGCTTGCGGCCTTTGGCACGACGACGTGCCAGGATGGCGCGACCATTTTTGGTAGCCATACGAGCACGGAAGCCATGCGCG
CGCTTGCGCTTAAGAACGCTGGGTTGAAAAGTGCGTTTCATAACTCGTTATTCCCACGTGGTTTAGGACGGAAAGTGAATTCCAAAAAGCCCGGTCCAGT

GAGGAACTAGCCGGGGAGATTCAATTCAAGACCGGAAATTGTAGAGACTTCCTCCGCTGCGTGCAATTTTTCCTTGCACAAAAAGCGTTGTCACCTCCCA
GCCAGCTTGCGCGGCCGGTCTAGAGTAGTCGCCCATGTTGTCACTATTATTACTAATAGGTAGTTCTAAACCTGTTGTTATTAATAGTGGGGATACTATT
**TGTGGATAA**GTGGTTTTAGTGTCTTTATTTCAACAACCTATAGGTATAAAAAATCTGTGTGTAAGTCGCGTATAAGCAGGAGATAAGCTGTTATTTTGAT
GTGATTAACCGCTGTGTAAGCAGTGATATTGAGGTTTATCCCTATCTATTCACAACGTTTTCCCCGTGGCTTATGACAAGTTGTCCACAGGGCGATATTG
ATGTCTAAACCTGTGTGGATAACCGCGGCTTTGGGCGCTACAATGGTCGGCCGTTTTCAACCGATGGTGAGATGAGTGTCACTGACGCTTTGGCAACAGT

GCCTGGACAACCTGCAGGATGAACTGAATTCACAGCAGTTCAACACCTGGATACGCCCACTGCAGGCAGAAGAAGGAGAGTCCAACGAGCTGCGCTTATT
AGCGCCGAATCGCTTTGTGCGCGATTGGGTGAGCGATAAGTATGCCAAGCGGATTAGTGAGCTGATGCGTGAGCTGGCACCGGCCAAACCACCTAAAGTC
AGTTTGACGGTAGGTAGTCGTCGCCCAGCTAACCACGCACCTCAGCATCGTGATTTGGGCAACCCCGTATCGGCGTCACCATCGCGGCCTTCAGCGCCGG
CGGTGCATACGCCGCCGCGTGGCGATATACCCGATGAGCGCGAATTAGATAAGCAACGCGAAGCGGCTTCGGGTCGGCGTACCAATGAGCGTCAGGTACA
AGTCGAAGGCAGCTTGAAACACGGCAGCGGTCTAAATCCAAACTTTACCTTTGACACCTTTGTGGAAGGTAAGTCGAATCAGCTCGCCCGCGCGGCGTCG
CGCCAAGTGTCAGAAAACCCCGGTGGTGCTTACAACCCCTTGTTTTTATACGGGGGAGTGGGCCTGGGTAAAACCCACCTAATGCATGCGGTGGGTAATG
CGTTAGCCACCCGGCGCGAAAATGCGCGGGTTGTCTATCTGCATTCAGAGCGTTTTGTGGCGGATATGGTCAAAGCACTACAGCTAAATGCGATTAATGA
TTTTAAGCGCTTTTACCGTAGCGTCGACGCCTTATTAATCGATGATATTCAGTTTTTCGCTGGCAAAGAGCGGTCGCAGGAGGAGTTCTTTCACACCTTC
AATGCGCTTTTGGAAGGCGGCCAACAGATGATTTTGACCTCTGATCGCTATCCGAAAGAGATCAGTGGTGTTGAAGAGCGGCTTAAATCACGTTTTGGTT
GGGGCCTGACGGTAGCGATTGAGCCGCCGGAGCTTGAGACCCGCGTGGCTATCCTGATGAAAAAGGCTGACCAGGCGAAGGTCGATTTGCCCCACGATGC

>Halovibrio variabilis DSM 3050 [NZ_BJXV01000011.1:c63206..63705], <mnmG oriC2 <COG1126 (glnQ), 617 bp

CGCACCAGCTGAATTTGCACGTCAAACGGCAGCGAGGTCGAGATCCCGTTGGGATACAGCTCGTGCGTATTTAAGCCTTCGGGTTCAATAAACACCTGAT
GGCTGGATTTATCGGCAAAGCGATGAACTTTGTCTTCGATTGACGGGCAATAGCGCGGCCCAACCCCTTCGATCACGCCCGAATACATCGGTGAGCGATC
AAGGTTTGCCAGAATCAGATCGTGGGTCTGCTGATTGGTATGCGCTATGTGGCAGCTGACCTGCTGGGGGTGCCGATCTCGCGAGCCCAGGTAGGACATC
ACCGGCGTTGGCGTATCGCCTGGCTGCTCTTCCATCTGGGAAAAGTCAACGCTTTTGGCATCAATGCGCGGCGGCGTGCCGGTTTTTAACCGGTCAACGC
GAAACGGTAATGCCCGCAGCCGTTCGGCCAGCGCATTGGAAGGCGGGTCGCCCGCTCGGCCGCCTTTGCTTTGGTCCATGCCAATGTGGATCACGCCACC
TAAAAACGTCCCGGTACTCAACACAACGCTGTCGGCGTGAAAGCGAATGCCGGTTTCCGTAACAACGCCACGCACGGTGTCGTTATCCACAATGAGATCG
CCAGCGGCCTGTTGGAAAAGCGTTAAATTTGGCTGGTTTTCCAACATGCCACGGATAGCGGCTTTGTAAAGAATACGGTCAGCCTGAGCACGGGTTGCCC
GAACGGCTGGTCCCTTGCGGGCATTGAGCACGCGAAATTGGATACCGGCTAAATCCGTGGCAAGCCCCATGGCGCCGCCCAACGCATCAATTTCCTTGAC
CAAATGGCTTTTGCCGATGCCGCCAATCGCGGGGTTACACGACATTTGACCCAGGGTTTCGATGTTGTGGGTGAGCAGCAGGGTCTGACAGCCCATGCGA
GCAGAGGCCAGTGCGGCTTCAGTTCCCGCATGGCCACCGCCGATGACAATCACGTCAAAGCGGTCGGGGTAATTCAAGAGACACCTCGTCTTGTGCCTAT

CGGCACGGATGATGTGAGCCGGTGCAGTCAGCTCCAGACATGAGCACCGTAGATCAATGAGCGGGGCAGTATAAACCCCCTGTGGGTAACCGTCAGGTTT
TTCCACACCCGGATTCACCAGAGAGGCTCTATTAATAACAACATAAATATAAATTAAAAGAAGTTCTGTAAATGTTGTAATTACTAGGGTGGACAATTTC
**TGTGTATAA**GCGATTTATTGCTATTTTTTTCAACAGTTTACTTTGTTGACTTCTACATGTTTGAAAGCGGGACTGCCTGGGTAGAAGGGGGGTTAAGCCT
GTGGATGAAATAGCGCTTTATCCACAGAGATAGTTAAGCACCTTTTATCCACCGGCCAGTACCTGGGTTGTTACCGCACCTGTGAACAACCGTGGCGGGA
TGTGGAGCTAACGCTAAGAGACTCGCAGGATAGGCCTTACAGAGGTATAGAGGAGCAAAGTAAAAAAAGTTTATCCACAGGCAAAAAAATGGCCTGTCAC

GTGGACAGGCCGGTAAAAAACGCCATTTAGCAGGGTCAAACATATTATTTTGGTGCCGAGTGCAGGTCTACGCGAGGCGATTAAGTGATGACATTTAGTG
TTTAAGTACCCGTGCAAGGAAGGATTGGGTGCGCTCATGTTGCGGCGTATCGAACAGTTTTTCAGGGGGGCCCTGTTCAGCAATTTCGCCTTTATGAATA
AAAATAACCCGGTCGGCGACTTCGCGGGCAAAGCCCATTTCATGGGTAACAATCACCATGGTCATGCCCTCTTTGGCCAGCTCGCGCATGGCATCGAGCA
CTTCACCGATCATTTCCGGGTCAAGCGCCGATGTGGGCTCGTCAAACAGCATAAGGCGCGGTTCCATGGCCAATGCGCGGGCTAGCGCCACACGCTGCTG
CTGGCCGCCTGAGAGATGGCTGGGGTACTTGTCGGCTTGGTCGGCGATACCCACACGGTCAAGCAAGCGTTTGCCGCTACTGATGGCGTCATCACGACTC
AAACCGCGTACTTTCATGGGGGCTAGGGTAACGTTGTCGAGCACACTGAGATGGGGGAAAAGATTGAACTGCTGGAACACCATACCCACTTCGGTGCGAA
TCTTTTGCAGCGCTTGGGTGCTCTTTCCATAGGGCAGCAGTTCGCTGCTATCGACGACCAGATTGCCCGACTGAAACTCTTCCAAGCCGTTTATACAGCG
GATCAATGTCGATTTACCCGAGCCACTGGCACCGATAATTACGACCACTTCGCCAGGCACAATGTCGAGGTCGATGTTGTTGAGTACGTGCAGGCTGCCA
AAATGCTTATTGAGCTGCTGCATACGCACAATTGGCTGCGTTGTATCGTTCATTACGCGTGTCCTTATTGTCCAGCCAGGCCTCTGCGCTCTAGCTGGCG
CAGCAGGATTGAGAGCGTCCAGGTGATCGCTAGATACATCAGCGCGACCATGAAATAGACCTCAAGGGCGGTGAAGGTAGTGGCGATATAAATTTGTCCC

>Halovibrio variabilis DSM 3050 [NZ_BJXV01000011.1:c44048..44107], dnaA> ig dnaN>, 60 bp

WebSkew GCmin pos. 505946 (GCmin<>oriC1 5.4 kb; GCmin<>oriC2 3.9 kb; oriC2<>oriC1 9.3 kb)

>Halovibrio salipaludis YL5-2 [NZ_NSKD01000001.1:c500303..500802], <rpmH oriC1 dnaA>, 626 bp

ATCGCTGCTGTCGCTACTGTCCGGAGGCTGTTGGCTCTGGGTTCCACTGTCAGCGGGAGCGGACTCCTCGGCTGCCACGGGTTCAGTGGAGACGGATTGC
TGGTGATAATCCTCGTTCCAGGCCATGACGAGGAGGTAGCTGACTATGGCCAGACCAAGCCATATGGCGACGCGGGGGATATCCATGGATTGGGTCATCC
TTTGCGTGAACAGGATTGATGGGTCGGGGGAACCCCTGGAACCGGATCATAACCGGATGTGCCCCAGGGATGGCATCGGAAGAGACGGTGCAAGGCGTAG
TAACCTCCCCGGAGCATGCCGTGCTGTTGGATCGCGCTCGCGGCGTATTCGGAACAGCTGGGCGTGTGCCGGCAGTGAGGAGGCATCAGCGGACTGATCC
CGTACTGGTAGATGCGAATCAGCAGAAGAAGCAGCCAGCGCATTGGGTTATCCCGTTTTTACCGGAGACCGGGAGGTCGCCTGTTGGCCCAGCTTCCGCC
ACGCCCGGTCGAGTTCCCGGAACAGTTCGGCATTGTCCATGGTATTGCCGCCAGGCCGGGCCAGCACGATGATGTCCAGAGCTGGCAGATCATGCTGTCG
GTGCCTGAAGGATTCCCGTACGGCACGCTTGAACCGGTTGCGCGCCGTAGCCTTGGGCAGATGCCGACGGGCCACGATGATGCCCAGACGGGGCAATGCC
CGGGATTCGTCGGAACTCCGGGCAAGCATCAGGAATCCGCGGCTGCCGAACCTGAACTCGACACTGTTGAATACCCGCGTGTAATCACGACTGTTGAGCA
GTCTTGCCTGGCGGGGAAATCGGAGGTCAGGCGGCAACTGGACGGGTTCAGACGCTGAGGCGCTTACGACCCTTGGCACGACGGCGCGCAATCACTTTGC
GGCCATTCTGGGTTGCCATGCGGGCACGGAAACCATGGGTCCGCTTGCGCTTGAGGTTACTCGGCTGAAAGGTTCGTTTCATGGTGATGGTCCTGTTCAG

GGCTGCGAATGTCGGAACAATGGTTTCGGCCGATGGTCCGAACACGGAGCTGCCACCGGGATGCGGGATTCTAGTCAATTTGTGGCGTTCAGACAATGGG
CAGCGGTGCCGGAAAGGCCTGTCAGTTACAGTAGAAACAAAGTTTTTATCTTTAAGGAATCAATAAGACTGTTGTTACTGTTATTAGGGAGGCGGTTTTC
**TGTGGATAA**CCGGTTTCGGGCGGCTGGTGGCGCCAATCCAGAGTGGGGACGCTTCTGTTCAGTTGCGCTTGGTAACCTGGGACTAACAGGGGGTATGGAA
TGGGTATAACAGGGGGTGGCTGACCGGGGGGCGGGTTATCAACGGGTTCGTCCGGGGGTTATACGGTCGTGAACACACGGTGTTTGCACAGGAGGTTGTG
GATAAGCTAATTCAAGGGGAATTGATCTGTAATTTTGTGCACGACTCTGTTTGCAAGTTGGTAGTTTTCTGAATGAAAACGAAAAAGTTATCAACAGGGT

GTTTATATCGTTGCTGTGGGGGCTTTTACGGGCGGTGAGCGGCGCGTAGAATACCGCTCTTGCCGGGGAGAGACCCGTCCGAAGAAGAACAATGCGAGGT
CAGCAGTCGTGGCACAGGAAATCTGGCAGCAGTGTCTGGAACTGTTACAGGATGAGTTCCCCGCCCAGCAGTTCAATACCTGGTTGCGTCCGTTGCAGTC
GCGCAACGATGGGGGGGCTCTGGTACTGCTTGCACCCAATCGGTTTGTGCGCGACTGGGTGAGCCAGAAGTATCTGGATCGTATTGAAGAGGTCATCCGG
GACCTGAATGGCGCCAAGGGGCTGCAGGTCGAGGTCAAGGTGGGATCCGCGGAGGACAGACAGCCCCGGTCGAGTCAGCCCCAGGCCGATCGTGGTGACA
GGGCGGTGAAACGCCAGTCAGCTGACAGCGCGGGCTCCGAAGTGGGATCGGCCAGCGCGGAAGTGGCACCGCCGGTTAATGATGCGGAACTGAATACAGC
CGATGAGCGCCCGGTTGAGGATTCACCCGTGGCCGCGCGTATTGCCGGGCGGGCCAAGGGTGGAGGCGATGATCGCCGGGCCAATGTCGAGGGTGGCGTC
CAGCACAGCGAGACCTTCCTGAACCAGCAGTTCACGTTTGGCACCTTCGTTGAGGGTAAATCCAACCAGCTGGCGCGGGCTGCCTCGAGCCAGGTGGCGG
AAAACCCGGGCGGTTCCTACAACCCGTTGTTCCTCTATGGCGGTGTGGGTCTGGGTAAGACCCACCTGATGCACGCCGTGGGCAACGAGATCGTGCGCCG
CAATCCCGGGGCACGCGTGGCCTACCTGCGCTCCGAGCGCTTTGTGGCGGACATGGTCAAGGCGCTGCAGCTCAACGCCATCAATGATTTCAAGCGCTTC
TACCGGTCCCTGGATGCCCTTCTGATCGACGATATCCAGTTCTTCGCGGGCAAGGAGCGCTCCCAGGAGGAGTTCTTCCACACCTTCAATGCGCTGCTTG

>Halovibrio salipaludis YL5-2 [NZ_NSKD01000001.1:c509647..510146], <mnmG ig <DUF3833, 286 bp

CGCGCGGCCAGGGCCGCGTTGAGCCCGGCCAGCAGGCCCTGGGCGCCGGCTTCCTCGTAACCGGTGGTGCCGTTGATCTGGCCCGCCAGATAGAGCCCGC
CGATAAAGCGGGTCTCAAGCGTGTGATGGAGGTCGCGCGGGTCCAGGTAGTCGTATTCGATGGCGTAACCCGGCCGGGTGACGTGGGCGTTCTCGAAGCC
CGGGATGGTCCGGATGAAGGCCATCTGCACGTCGAACGGCAGGCTTGTGGAAATACCGTTGGGATAGAGTTCCTGGCTGTTCAGGCCTTCCGGCTCCACG
AAGATCTGGTGGGAATCCTTGTCCGCGAAGCGGTGGACCTTGTCCTCGATGGAGGGGCAGTAGCGCGGCCCAACGCCTTCAATGTCGCCAGAGAACATGG
GCGACTGATCCTGGGCCCCCCGGATGATGTCGTGGGTCTCCGGCGTGGTGCGGGTGATGTAACAGCACATCTGCTCTGGATGCGAATCCGCACTGCCGAT
GTAGGACATCACCGGTTCGCGTTCGTCGCCGGGCTGAACGTCCATCTTCGAGAAGTCCACGGTCTTGGCGTCGATCCGCGCGGGTGTGCCGGTCTTGAGC
CGGCCCACGGTGAACGGGAGCTCCCGCAGCCGTTTGGCCAGGCCGATGGTCGGCGGGTCGCCGGCGCGGCCGCCCTGCTGATTCTGCATGCCAATGTGGA
TGACCCCCCCGAGGAAGGTACCGGTGGTCAGCACCACCGTGGGTGCCCGGAAGGCGACACCGGTCTGGGAGACCACGCCCTGCACCTGGTCATTCTCGAC
GATCAGATCCTCCGCCGCCTGCTGGAAGATATGCAGGTTGGGCTGGTTTTCCAGCATGTTGCGGATGGCGGCCTTGTACAGGGCACGGTCCGCCTGGGCA
CGGGTAGCGCGTACGGCCGGGCCCTTGCTGGAATTGAGAACCCGGAACTGGATGCCCGCCTGGTCCGTGGCCAGGCCCATGGCCCCGCCCAGCGCATCGA

TCTCCTTCACCAGGTGACTCTTGCCGATACCGCCGATGGCGGGATTACAGGACATCTGGCCAAGGGTCTCGATGTTGTGCGTCAGCAGGAGGGTATCGCG
CCCCATGCGGGCCGCGGCGAGGGCCGCTTCGGTGCCCGCATGGCCACCGCCGATGACGATGACGTCGAAGTGCTTGGGGTATTCCACGCTGGACCTCGGG
GTTTGCCGGTATCTCTGAATGAACCGCGTAGTATAACCCGAAGCGTCAAGCCTGTGCAGGCGATCCGGACAATTTCTGAACAGATACCATCTATCCCGGA
ATAACAGACACCTGCAGTGCTTTCGTCACAAAGCACTCAAAAAACTGTCCACAGATTGGAGGCGCCTTAAAAAGCCTTATTTATTACCTAGAATGCTGAC
GTAAAAGTATTTTTTAAATTTTCTACAGATCTCTCCCTATTCTGTAAACAGAGTTTTCCACAAGGAAGGCGACCTACTGTCTCTGGATCACCAGGACCAC

CTCGCCCACACGGAATCCCCACCAGGTCATTTCCGTTTCATTGATGACGGTGTCCTCGGTGACACGGTACATCCAGTCATCCATTCGGACATTGTAGGTG
CTGCCTCCGAAGGACACTTCCAGCACATAGTTCATGTGGATGGCATTGCCCTCGAACTCCATGAACCCCGGCTCTGTCACATCCCCTGCCGTGGCGTGAT
AGCCGCCGTCACGGGGTTCCAGGGTCCAGACGCGCGTGTCGGTCTCACCATCGCTGTACGTGAACACCTCGTCCAGGGTACCCACGCCATCCTCATCCCA
TGACGCATCAATGGTGGCATCGAAATGGCGGATCACCTCGCCGGAACGGTTCTTCACCACACCACGGGCGGTCAGCTCACCCTGGAAGAACTCACTGGGC
ACCAGCTCCGGCTCCTGGCCCGCATAGTCGTTGATTGCGGGGCCGGAGCAGGCCGCCAGAAAGGGCAGGACCAGAAAAAGAAGCCCGTAGACGCGGCGGG
CTGGATGTATGACTGCCATGGGCTGAACCTCCATCGGAATGAGTGACTGTTACTGGATACGTGGGTTTTCCCGGGAGGGTTCATTGGGTGGCCGCAGTGA
CGTACACAGATGTACACGTCGCCATTGGCTCGAATGACGGAGCTGTCAGAAACTGTCTTCACAGTAAACAGATGGGAGGGTCCGAGATGGCAAGCTACAA
GGCGCCTCTGACCGACGTGAATTTCCTGCTCAACGAAGTGTTCGACTACCCCAGTCACTACCAGAGCCTGAGCAACGGGGCCGAGGCGACACCGGACATG
GTGGACGCCATCCTCGGTGAATGTGCAAAGTTCTGTGAGAACACTCTGTTTCCGCTCTACCAGAGCGGCGACCAGGAAGGCTGCAAACTGGTGGATGGCG
AGGTTAAGACCCCGGCCGGCTACAAGGAAGCCTACGACGAATACGTGGCCGGTGGCTGGCAGGGCCTCTCCAGCCTGGAGGAATACGGCGGCCAGAACCT

>Halovibrio salipaludis YL5-2 [NZ_NSKD01000001.1:c498630..498673], dnaA> ig dnaN>, 44 bp
