## Supplementary material for "Dual chromosomal origins of replication (*oriC*) in the genomes of the Halomonadaceae – a prediction study": Suppl. Table 3_v02.zip: Suppl. Table 3_v02.pdf



| # | Species / Strain | RefSeq / Nucleotide | 5' gene | ig | oriC2 | 3' gene | oriC2 Δ oriC1 [kb] | 5' gene | ig | oriC1 | 3' gene | DoriC 12 | RefSeq / Nucleotide | oriC2 | oriC1 |
|---|---|---|---|---|---|---|---|---|---|---|---|---|---|---|---|
| | Halomonas (sensu stricto) | | | | | | | | | | | | | | |
| 001 | Halomonas elongata DSM 2581 | NC_014532.2 | < mnmG | 535 | | < glnQ | 18.2 | < rpmH | 504 | | dnaA > | ORI97226808 | NC_014532.1 | | |
| 002 | Halomonas eurihalina DSM 5720 | NZ_JARWAH010000005.1 | < mnmG | 536 | | < glnQ | 18.1 | < rpmH | 659 | | dnaA > | | | | |
| 003 | Halomonas caseinilytica JCM 14802 | NZ_BDEP01000014.1 | < mnmG | 536 | | < glnQ | 19.2 | < rpmH | 514 | | dnaA > | ORI97099728 | NZ_BDEP01000014.1 | | |
| 004 | Halomonas halmophila NBRC 15537 | NZ_BJOC01000021.1 | < mnmG | 582 | | < glnQ | 19.2 | < rpmH | 515 | | dnaA > | ORI97035787 | NZ_BJOC01000021.1 | | |
| 005 | Halomonas sabkhae CECT 7246 | NZ_JAUFPM010000008.1 | < mnmG | 583 | | < glnQ | 19.4 | < rpmH | 505 | | dnaA > | | | | |
| 006 | Halomonas almeriensis CECT 7050 | NZ_JAUFPO010000005.1 | < mnmG | 537 | | < glnQ | 19.3 | < rpmH | 597 | | dnaA > | | | | |
| 007 | Halomonas halophila NBRC 102604 | NZ_BJUS01000022.1 | < mnmG | 537 | | < glnQ | 25.7 | < rpmH | 596 | | dnaA > | ORI97034337 | NZ_BJUS01000022.1 | | |
| 008 | Halomonas salina DSM 5928 | NZ_JASFAS010000003.1 | < mnmG | 537 | | < glnQ | 25.7 | < rpmH | 598 | | dnaA > | | | | |
| 009 | Halomonas smyrnensis AAD6 | NZ_AJKS02000002.1 | < mnmG | 537 | | < glnQ | 25.7 | < rpmH | 515 | | dnaA > | ORI97106243 | NZ_AJKS02000002.1 | | |
| 010 | Halomonas borealis ATF 5.2 | NZ_SDMO01000002.1 | < mnmG | 525 | | < glnQ | 25.6 | < rpmH | 515 | | dnaA > | ORI97054620 | NZ_SDMO01000002.1 | | |
| 011 | Halomonas organivorans CECT 5995 | NZ_JACHXM010000012.1 | < mnmG | 529 | | < glnQ | 25.9 | < rpmH | 516 | | dnaA > | | | | |
| 012 | Halomonas koreensis Lim DSM 23530 | NZ_JARWAK010000004.1 | < mnmG | 527 | | < glnQ | 25.0 | < rpmH | 557 | | dnaA > | | | | |
| 013 | Halomonas stenophila CECT 7744 | NZ_JACHXR010000003.1 | < mnmG | 537 | | < glnQ | 25.9 | < rpmH | 515 | | dnaA > | ORI97171330 | NZ_JACHXR010000003.1 | | |
| 014 | Halomonas maura CECT 5298 | NZ_JAUFPQ010000013.1 | < mnmG | 536 | | < glnQ | 20.4 | < rpmH | 514 | | dnaA > | | | | |
| 015 | Halomonas nitroreducens 11S | NZ_RXNS01000004.1 | < mnmG | 536 | | < glnQ | 25.7 | < rpmH | 515 | | dnaA > | ORI97096104 | NZ_RXNS01000004.1 | | |
| 016 | Halomonas halocynthiae DSM 14573 | NZ_AUDZ01000013.1 | < mnmG | 542 | | < glnQ | 17.3 | < rpmH | 521 | | dnaA > | ORI97203728 | NZ_AUDZ01000013.1 | | |
| 017 | Halomonas cerina CECT 7282 | NZ_JACHXP010000005.1 | < mnmG | 531 | | < glnQ | 20.4 | < rpmH | 505 | | dnaA > | ORI97151594 | NZ_JACHXP010000005.1 | | |
| 018 | Halomonas huangheensis BJGMM-B45 | NZ_CP013106.1 | < mnmG | 643 | | < glnQ | 20.2 | < rpmH | 616 | | dnaA > | ORI97124172 | NZ_AVBC01000035.1 | | |
| 019 | Halomonas cupida DSM 4740 | NZ_FRCA01000005.1 | < mnmG | 546 | | < glnQ | 20.8 | < rpmH | 632 | | dnaA > | ORI97205276 | NZ_FRCA01000005.1 | | |
| 020 | Halomonas litopenaei SYSU ZJ2214 | NZ_PXNS01000004.1 | < mnmG | 540 | | < glnQ | 20.9 | < rpmH | 601 | | dnaA > | ORI97068460 | NZ_PXNS01000004.1 | | |
| 021 | Halomonas urmiana TBZ3 | NZ_VBUI01000001.1 | < mnmG | 565 | | < glnQ | 19.4 | < rpmH | 563 | | dnaA > | ORI97067155 | NZ_VBUI01000001.1 | | |
| 022 | Halomonas saccharevitans CGMCC 1.6493 | NZ_FPAQ01000016.1 | < mnmG | 576 | | < glnQ | 19.4 | < rpmH | 573 | | dnaA > | ORI97155334 | NZ_FPAQ01000016.1 | | |
| 023 | Halomonas denitrificans DSM 18045 | NZ_PYVX01000003.1 | < mnmG | 553 | | < glnQ | 19.6 | < rpmH | 553 | | dnaA > | ORI97105956 | NZ_PYVX01000003.1 | | |
| 024 | Halomonas aestuarii Hb3 | NZ_CP018139.1 | < mnmG | 532 | | < glnQ | 19.4 | < rpmH | 530 | | dnaA > | ORI97227175 | NZ_CP018139.1 | | |
| 025 | Halomonas ventosae CECT 5797 | NZ_SNZJ01000018.1 | < mnmG | 531 | | < glnQ | 19.1 | < rpmH | 510 | | dnaA > | ORI97169281 | NZ_SNZJ01000018.1 | | |
| 026 | Halomonas fontilapidosi CECT 7341 | NZ_JACHXQ010000006.1 | < mnmG | 535 | | < glnQ | 18.7 | < rpmH | 508 | | dnaA > | ORI97149646 | NZ_JACHXQ010000006.1 | | |
| 027 | Halomonas daqiaonensis CGMCC 1.9150 | NZ_FOBC01000008.1 | < mnmG | 533 | | < glnQ | 19.0 | < rpmH | 507 | | dnaA > | ORI97174626 | NZ_FOBC01000008.1 | | |
| 028 | Halomonas lysinitropha 3(2) | NZ_CABVOU010000015.1 | < mnmG | 530 | | < glnQ | 18.0 | < rpmH | 508 | | dnaA > | ORI97164369 | NZ_CABVOU010000015.1 | | |
| | Halomonas_G | | | | | | | | | | | | | | |
| 029 | Halomonas marinisediminis 204 | NZ_SLTR01000012.1 | < mnmG | 569 | | < glnQ | 27.3 | < rpmH | 578 | | dnaA > | ORI97044454 | NZ_SLTR01000012.1 | | |
| 030 | Halomonas icarae D1-1 | NZ_WUTS01000001.1 | < mnmG | 569 | | < glnQ | 27.3 | < rpmH | 578 | | dnaA > | ORI97195746 | NZ_WUTS01000001.1 | | |
| 031 | Halomonas halodenitrificans DSM 735 | NZ_JHVH01000002.1 | < mnmG | 554 | | < glnQ | 20.0 | < rpmH | 568 | | dnaA > | ORI97150221 | NZ_JHVH01000002.1 | | |
| 032 | Halomonas alimentaria DSM 15356 | NZ_WUTT01000001.1 | < mnmG | 543 | | < glnQ | 18.1 | < rpmH | 569 | | dnaA > | ORI97151551 | NZ_WUTT01000001.1 | | |
| 033 | Halomonas shengliensis CGMCC 1.6444 | NZ_FNIV01000006.1 | < mnmG | 532 | | < glnQ | 18.2 | < rpmH | 580 | | dnaA > | ORI97154447 | NZ_FNIV01000006.1 | | |
| 034 | Halomonas ramblicola CECT 7896 | NZ_JAUFPL010000032.1 | < mnmG | 535 | | < glnQ | 19.0 | < rpmH | 521 | | dnaA > | | | | |
| 035 | Halomonas alkalicola M2 | NZ_CP130143.1 | < mnmG | 530 | | < glnQ | 19.1 | < rpmH | 545 | | dnaA > | | | | |
| 036 | Halomonas mongoliensis DSM 17332 | NZ_JARWAL010000011.1 | < mnmG | 532 | | < glnQ | 19.0 | < rpmH | 556 | | dnaA > | | | | |
| 037.1 | Halomonas campaniensis 5AG Ga0075189_04 | NZ_JACHZF010000004.1 | < mnmG | 528 | | < glnQ | 19.2 | < rpmH | 520 | | dnaA > | ORI97196029 | NZ_JACHZF010000004.1 | | |
| 037.2 | Halomonas campaniensis LS21 | NZ_CP007757.1 | < mnmG | 565 | | < glnQ | 21.5 | < rpmH | 534 | | dnaA > | ORI97227523 | NZ_CP007757.1 | | |
| 038 | Halomonas urumqiensis BZ-SZ-XJ27 | NZ_PYUD01000002.1 | < mnmG | 574 | | < glnQ | 24.9 | < rpmH | 516 | | dnaA > | ORI97154686 | NZ_PYUD01000002.1 | | |
| 039 | Halomonas heilongjiangensis DSM 26881 | NZ_PNRE01000019.1 | < mnmG | 563 | | < glnQ | 19.0 | < rpmH | 519 | | dnaA > | ORI97154067 | NZ_PNRE01000019.1 | | |
| 040 | Halomonas korlensis CGMCC 1.6981 | NZ_FPBP01000003.1 | < mnmG | 546 | | < glnQ | 23.4 | < rpmH | 565 | | dnaA > | ORI97164687 | NZ_FPBP01000003.1 | | |
| | Halomonas_H | | | | | | | | | | | | | | |
| 041 | Halomonas sulfidivorans MCCC 1A13718 | NZ_CP053383.1 | < mnmG | 536 | | < glnQ | 26.7 | < rpmH | 472 | | dnaA > | ORI97022290 | NZ_CP053383.1 | | |
| 042 | Halomonas antri Y3S6 | NZ_JAHYCA010000007.1 | < mnmG | 534 | | < glnQ | 26.7 | < rpmH | 473 | | dnaA > | ORI97152511 | NZ_JAHYCA010000007.1 | | |
| 043 | Halomonas sulfidoxydans MCCC 1A11059 | NZ_CP053381.1 | < mnmG | 538 | | < glnQ | 27.1 | < rpmH | 462 | | dnaA > | ORI97021701 | NZ_CP053381.1 | | |
| 044 | Halomonas lactosivorans KCTC 52281 | NZ_QHGY01000015.1 rpmH<br>NZ_QHGY01000013.1 mnmG | < mnmG | 529 | | < glnQ | | < rpmH | 473 | | dnaA > | ORI97196047 | NZ_QHGY01000013.1 | | |
| 045 | Halomonas bachuensis DX6 | NZ_JAAQTO010000018.1 | < mnmG | 530 | | < glnQ | 22.1 | < rpmH | 485 | | dnaA > | ORI97056881 | NZ_JAAQTO010000018.1 | | |
| 046 | Halomonas diversa MCCC 1A13316 | NZ_CP053382.1 | < mnmG | 529 | | < glnQ | 25.2 | < rpmH | 486 | | dnaA > | ORI97024597 | NZ_CP053382.1 | | |
| 047 | Halomonas tianxiuensis BC-M4-5 | NZ_CP035042.1 | < mnmG | 539 | | < glnQ | 38.4 | < rpmH | 499 | | dnaA > | ORI97031521 | NZ_CP035042.1 | | |
| 048 | Halomonas zhangzhouensis MCCC 1A11036 | NZ_JABFTT010000001.1 | < mnmG | 537 | | < glnQ | 32.2 | < rpmH | 499 | | dnaA > | GCF_021404465. | NZ_JABFTT010000001.1 | ? | ? |
| 049 | Halomonas desiderata FB2 | NZ_JAAQTN010000031.1 rpmH<br>NZ_JAAQTN010000055.1 mnmG | < mnmG | 536 | | < erpA | | < rpmH | 513 | | dnaA > | | | | |
| 050 | Halomonas daqingensis CGMCC 1.6443 | NZ_FNVC01000006.1 | < mnmG | 536 | | < erpA | 40.7 | < rpmH | 513 | | dnaA > | ORI97201095 | NZ_FNVC01000006.1 | | |
| 051 | Halomonas ethanolica MCCC 1A11081 | NZ_JABFTX010000004.1 | < mnmG | 534 | | < erpA | 35.9 | < rpmH | 513 | | dnaA > | ORI97070472 | NZ_JABFTX010000004.1 | | |
| 052 | Halomonas aerodenitrificans MCCC 1A11058 | NZ_JABFTV010000003.1 | < mnmG | 533 | | < erpA | 40.7 | < rpmH | 513 | | dnaA > | GCF_021404405. | NZ_JABFTV010000003.1 | ? | ? |
| 053 | Halomonas kenyensis DSM 17331 | NZ_JACEFT010000002.1 | < mnmG | 541 | | < erpA | 29.3 | < rpmH | 461 | | dnaA > | ORI97066006 | NZ_JABFUB010000004.1 | | |
| 054 | Halomonas saliphila LCB169 | NZ_PJRN01000005.1 | < mnmG | 555 | | < glnQ | 26.0 | < rpmH | 509 | | dnaA > | ORI97072400 | NZ_PJRN01000005.1 | | |
| 055 | Halomonas pellis L5 | NZ_VTPY01000006.1 | < mnmG | 521 | | < glnQ | 25.0 | < rpmH | 508 | | dnaA > | ORI97135683 | NZ_VTPY01000006.1 | | |
| 056 | Halomonas chromatireducens AGD 8-3 | NZ_CP014226.1 | < mnmG | 542 | | < glnQ | >1.2 Mb | < rpmH | 501 | | dnaA > | ORI97227308 | NZ_CP014226.1 | | |
| 057 | Halomonas montanilacus PYC7W | NZ_QPII01000003.1 | < mnmG | 542 | | < glnQ | 29.5 | < rpmH | 502 | | dnaA > | ORI97200287 | NZ_QPII01000003.1 | | |
| 058 | Halomonas endophytica MC28 | NZ_PNRF01000027.1 | < mnmG | 528 | | < glnQ | 21.8 | < rpmH | 500 | | dnaA > | ORI97203917 | NZ_PNRF01000027.1 | | |
| 059 | Halomonas campisalis A4 | NZ_JABFUC010000002.1 | < mnmG | 733 | | < erpA | 17.0 | < rpmH | 506 | | dnaA > | ORI97075301 | NZ_JABFUC010000002.1 | | |
| 060 | Halomonas azerbaijanica TBZ202 | SBLD01000003.1 | < mnmG | 525 | | < glnQ | 23.7 | < rpmH | 388 | | dnaA > | ORI97155081 | NZ_SBLD01000003.1 | | |
| 061 | Halomonas gudaonensis CGMCC 1.6133 | NZ_FNES01000002.1 | < mnmG | 523 | | < glnQ | 40.0 | < rpmH | 377 | | dnaA > | ORI97197340 | NZ_FNES01000002.1 | | |
| | Halomonas_I | | | | | | | | | | | | | | |
| 062 | Halomonas aquamarina 558 | NZ_FODB01000017.1 rpmH<br>NZ_FODB01000057.1 mnmG | < mnmG | 522 | | contig end | | < rpmH | 534 | | dnaA > | ORI97156283 | NZ_FODB01000057.1 | | |
| 063 | Halomonas meridiana ACAM 246 | NZ_FSQY01000001.1 | < mnmG | 579 | | < glnQ | 49.9 | < rpmH | 534 | | dnaA > | ORI97051522 | NZ_FSQY01000001.1 | | |
| 064 | Halomonas axialensis FXH-124 | JAJGND010000025.1 rpmH<br>JAJGND010000012.1 mnmG | < mnmG | 579 | | < glnQ | | < rpmH | 534 | | dnaA > | ORI97061047 | NZ_JAJGND010000012.1 | | |
| 065 | Halomonas lionensis RHS90 | NZ_MWVI01000087.1 rpmH<br>NZ_MWVI01000003.1 mnmG | < mnmG | 579 | | contig end | | < rpmH | 534 | | dnaA > | | | | |
| 066 | Halomonas piezotolerans NBT06E8 | NZ_CP048602.1 | < mnmG | 578 | | < glnQ | 21.1 | < rpmH | 534 | | dnaA > | ORI97031968 | NZ_CP048602.1 | | |
| 067 | Halomonas lutescens CGMCC 1.15122 | NZ_BMHM01000004.1 | < mnmG | 576 | | < glnQ | 21.9 | < rpmH | 534 | | dnaA > | ORI97063569 | NZ_BMHM01000004.1 | | |
| 068 | Halomonas stevensii S18214 | NZ_AJTS01000041.1 | < mnmG | 542 | | < glnQ | 19.1 | < rpmH | 533 | | dnaA > | ORI97062038 | NZ_AJTS01000041.1 | | |
| 069 | Halomonas johnsoniae KCTC 22157 | NZ_BMXO01000019.1 | < mnmG | 542 | | < glnQ | 21.3 | < rpmH | 533 | | dnaA > | ORI97105934 | NZ_BMXO01000019.1 | | |
| 070 | Halomonas hamiltonii KCTC 22154 | NZ_BMXN01000011.1 | < mnmG | 542 | | < glnQ | 21.8 | < rpmH | 533 | | dnaA > | ORI97159582 | NZ_BMXN01000011.1 | | |
| 071 | Halomonas humidisoli WN018 | NZ_NSKA01000003.1 | < mnmG | 565 | | < glnQ | 21.4 | < rpmH | 534 | | dnaA > | ORI97100398 | NZ_NSKA01000003.1 | | |
| 072 | Halomonas alkaliphila DSM 16354 | NZ_JABASV010000002.1 | < mnmG | 565 | | < glnQ | 21.6 | < rpmH | 534 | | dnaA > | ORI97029697 | NZ_CP024811.1 | | |
| 073 | Halomonas venusta NBRC 102221 | NZ_BJUL01000002.1 | < mnmG | 566 | | < glnQ | 21.5 | < rpmH | 535 | | dnaA > | ORI97082281 | NZ_BJUL01000002.1 | | |
| 074 | Halomonas hydrothermalis Slthf2 | NZ_AP022843.1 | < mnmG | 565 | | < glnQ | 21.8 | < rpmH | 535 | | dnaA > | ORI97024030 | NZ_AP022843.1 | | |
| 075 | Halomonas andesensis DSM 19434 | NZ_RZHG01000019.1 | < mnmG | 578 | | < glnQ | 21.5 | < rpmH | 533 | | dnaA > | ORI97206017 | NZ_RZHG01000019.1 | | |

Suppl. Table 3



| # | Species / Strain | RefSeq / Nucleotide | 5' gene | ig | 3' gene | oriC2 Δ oriC1 [kb] | 5' gene | ig | 3' gene | DoriC 12 | RefSeq / Nucleotide | oriC2 | oriC1 |
|---|---|---|---|---|---|---|---|---|---|---|---|---|---|
| | Halomonas_I (cont.) | | | | | | | | | | | | |
| 076 | Halomonas zhaodongensis NEAU-ST10-25 | NZ_JACCDD010000002.1 | < mnmG | 564 | < glnQ | 21.5 | < rpmH | 534 | dnaA > | | | | |
| 077 | Halomonas nigrificans MBT G8648 | NZ_NWUX01000004.1 | < mnmG | 554 | < glnQ | 22.1 | < rpmH | 534 | dnaA > | ORI97170423 | NZ_NWUX01000004.1 | | |
| 078 | Halomonas malpeensis YU-PRIM-29 | NZ_WHVL01000008.1 | < mnmG | 565 | < glnQ | 21.6 | < rpmH | 532 | dnaA > | ORI97134804 | NZ_WHVL01000008.1 | | |
| 079 | Halomonas populi MC | NZ_RZHE01000015.1 | < mnmG | 555 | < glnQ | 19.7 | < rpmH | 534 | dnaA > | ORI97066468 | NZ_RZHE01000015.1 | | |
| 080 | Halomonas nanhaiensis JCM 18142 | NZ_RZHF01000015.1 | < mnmG | 554 | < glnQ | 19.6 | < rpmH | 534 | dnaA > | ORI97175004 | NZ_RZHF01000015.1 | | |
| 081 | Halomonas songnenensis CGMCC 1.12152 | NZ_PVTK01000004.1 | < mnmG | 556 | < glnQ | 23.0 | < rpmH | 534 | dnaA > | ORI97131894 | NZ_PVTK01000004.1 | | |
| 082 | Halomonas zhanjiangensis DSM 21076 | NZ_KB898653.1 | < mnmG | 574 | < glnQ | 26.4 | < rpmH | 534 | dnaA > | ORI97178015 | NZ_KB898653.1 | | |
| 083 | Halomonas titanicae BH1 | NZ_AOPO01000001.1 | < mnmG | 617 | < glnQ | 21.7 | < rpmH | 533 | dnaA > | ORI97078860 | NZ_AOPO01000001.1 | | |
| 084 | Halomonas sedimenti QX-2 | NZ_JACCGK010000004.1 | < mnmG | 617 | < glnQ | 21.7 | < rpmH | 533 | dnaA > | ORI97036809 | NZ_JACCGK010000004.1 | | |
| 085 | Halomonas alkaliantarctica CRSS | NZ_SOCZ01000018.1 | < mnmG | 616 | < glnQ | 23.5 | < rpmH | 533 | dnaA > | ORI97168398 | NZ_SOCZ01000018.1 | | |
| 086 | Halomonas neptunia CECT 5815 | NZ_JAUFPP010000021.1 rpmH<br>NZ_JAUFPP010000005.1 mnmG | < mnmG | 616 | < glnQ | | < rpmH | 533 | dnaA > | | | | |
| 087 | Halomonas olivaria TYRC17 | NZ_JALPZO010000066.1 | < mnmG | 552 | < glnQ | 17.0 | < rpmH | 534 | dnaA > | | | | |
| 088 | Halomonas boliviensis LC1 | NZ_JH393257.1 | < mnmG | 595 | < glnQ | 21.8 | < rpmH | 534 | dnaA > | ORI97071426 | NZ_NPEY01000006.1 | | |
| 089 | Halomonas maris QX-1 | NZ_JABWCV010000006.1 | < mnmG | 591 | < glnQ | 23.5 | < rpmH | 535 | dnaA > | ORI97089065 | NZ_JABWCV010000006.1 | | |
| 090 | Halomonas glaciei DD 39 | NZ_JACCDE010000017.1 | < mnmG | 594 | < glnQ | 17.4 | < rpmH | 534 | dnaA > | ORI97076833 | NZ_JACCDE010000017.1 | | |
| 091 | Halomonas sulfidaeris SST4 | NZ_QNTU01000008.1 | < mnmG | 593 | < glnQ | 20.9 | < rpmH | 533 | dnaA > | ORI97108502 | NZ_QNTU01000008.1 | | |
| 092 | Halomonas utahensis NBRC 102410 | NZ_BJXV01000011.1* | < mnmG | 617 | < glnQ | 17.7 | < rpmH | 534 | dnaA > | ORI97064817 | NZ_BJXV01000011.1 | | |
| 093 | Halomonas profundi MT13 | NZ_JAHRFY010000004.1 | < mnmG | 592 | < glnQ | 21.6 | < rpmH | 534 | dnaA > | ORI97142013 | NZ_JAHRFY010000004.1 | | |
| 094 | Halomonas subterranea CGMCC 1.6495 | NZ_FOGS01000001.1 | < mnmG | 557 | < glnQ | 22.8 | < rpmH | 532 | dnaA > | ORI97155462 | NZ_FOGS01000001.1 | | |
| 095 | Halomonas janggokensis DSM 18043 | NZ_JARWAJ010000002.1 | < mnmG | 557 | < glnQ | 22.7 | < rpmH | 532 | dnaA > | | | | |
| 096 | Halomonas gomseomensis DSM 18042 | NZ_JARWAI010000003.1 | < mnmG | 558 | < glnQ | 21.3 | < rpmH | 532 | dnaA > | | | | |
| 097 | Halomonas arcis CGMCC 1.6494 | NZ_FNII01000021.1 rpmH<br>NZ_FNII01000001.1 mnmG | < mnmG | 557 | < glnQ | | < rpmH | 532 | dnaA > | ORI97160114 | NZ_FNII01000001.1 | | |
| 098 | Halomonas salicampi BH103 | NZ_JACCDF010000018.1 rpmH<br>NZ_JACCDF010000008.1 mnmG | < mnmG | 551 | < glnQ | | < rpmH | 524 | dnaA > | ORI97132472 | NZ_JACCDF010000008.1 | | |
| 099 | Halomonas vilamensis DSM 21020 | NZ_JARWAN010000010.1 | < mnmG | 550 | < glnQ | 23.6 | < rpmH | 534 | dnaA > | | | | |
| 100 | Halomonas massiliensis Marseille-P2426 | NZ_LT699745.1 | < mnmG | 551 | < glnQ | 26.7 | < rpmH | 522 | dnaA > | ORI97198024 | NZ_LT699745.1 | | |
| 101 | Halomonas azerica TBZ9 | NZ_JABFHI010000006.1 | < mnmG | 584 | < glnQ | 16,9 | < rpmH | 577 | dnaA > | | | | |
| 102 | Halomonas jeotgali Hwa | NZ_AMQY01000003.1 | < mnmG | 707 | ydfG > | × 33.7 | < rpmH | 728 | dnaA > | ORI97118047 | NZ_AMQY01000003.1 | | |
| 103 | Halomonas subglaciescola ACAM 12 | NZ_LT670847.1 | < mnmG | 652 | ydfG > | 38.8 | < rpmH | 544 | dnaA > | ORI97019619 | NZ_LT670847.1 | | |
| 104 | Halomonas rituensis TQ8S | NZ_QPIJ01000013.1 | < mnmG | 566 | < glnQ | 21.9 | < rpmH | 543 | dnaA > | ORI97199216 | NZ_QPIJ01000013.1 | | |
| 105 | Halomonas zhuhanensis ZH2S | NZ_WTKP01000010.1 | < mnmG | 565 | < glnQ | 23.7 | < rpmH | 544 | dnaA > | ORI97200149 | NZ_WTKP01000010.1 | | |
| | Halomonas_F | | | | | | | | | | | | |
| 106 | Halomonas pacifica NBRC 102220 | NZ_BJUK01000023.1 rpmH<br>NZ_BJUK01000013.1 mnmG | < mnmG | 544 | < glnQ | | < rpmH | 556 | dnaA > | ORI97103725 | NZ_BJUK01000013.1 | | |
| | Halomonas_B | | | | | | | | | | | | |
| 107 | Halomonas salipaludis WRN001 | NZ_NSKB01000003.1 | < mnmG | 544 | < glnQ | 26.3 | < rpmH | 501 | dnaA > | ORI97058627 | NZ_NSKB01000003.1 | | |
| 108 | Halomonas qiaohouensis DSM 26770 | NZ_JARWAM010000001.1 | < mnmG | 544 | < glnQ | 27.7 | < rpmH | 502 | dnaA > | | | | |
| 109 | Halomonas pantelleriensis AAP | NZ_FNGH01000005.1 | < mnmG | 547 | < glnQ | 26.3 | < rpmH | 501 | dnaA > | ORI97152368 | NZ_FNGH01000005.1 | | |
| | Halomonas_E | | | | | | | | | | | | |
| 110 | Halomonas taeanensis BH539 | NZ_FNCI01000005.1 | < mnmG | 615 | < glnQ | 26.6 | < rpmH | 568 | dnaA > | ORI97158915 | NZ_FNCI01000005.1 | | |
| 111 | Halomonas niordiana ATF 5.4 | NZ_SDSD01000004.1 | < mnmG | 617 | < glnQ | 26.6 | < rpmH | 557 | dnaA > | GCF_004798965. | NZ_SDSD01000004.1 | ? | ? |
| | Halomonas_A | | | | | | | | | | | | |
| 112 | Halomonas anticariensis FP35 = DSM 16096 | NZ_KE332393.1 | < mnmG | 549 | < glnQ | 27.8 | < rpmH | 547 | dnaA > | ORI97158007 | NZ_KE332393.1 | | |
| 113 | Halomonas qijiaojingensis KCTC 22228 | NZ_BMXS01000003.1 | < mnmG | 549 | < glnQ | 28.3 | < rpmH | 565 | dnaA > | ORI97130853 | NZ_BMXS01000003.1 | | |
| 114 | Halomonas rifensis CECT 7698 | not yet in RefSeq | | | | | | | | | | | |
| 115 | Halomonas xinjiangensis TRM 0175 | NZ_JPZL01000003.1 | < mnmG | 537 | < glnQ | 25.5 | < rpmH | 531 | dnaA > | ORI97053782 | NZ_JPZL01000003.1 | | |
| | Aidingimonas | | | | | | | | | | | | |
| 116 | Aidingimonas halophila DSM 19219 | NZ_FNNI01000003.1 | < mnmG | 550 | < glnQ | 33.1 | < rpmH | 447 | dnaA > | ORI97159950 | NZ_FNNI01000003.1 | | |
| 117 | Aidingimonas lacisalsi XHU 5135 | NZ_VRYH01000001.1 | < mnmG | 559 | < glnQ | 33.5 | < rpmH | 459 | dnaA > | ORI97203524 | NZ_VRYH01000001.1 | | |
| | Modicisalibacter | | | | | | | | | | | | |
| 118 | Modicisalibacter tunisiensis LIT2 | NZ_JAGXFD010000001.1 | < mnmG | 554 | < glnQ | 33.6 | < rpmH | 607 | dnaA > | | | | |
| 119 | Halomonas coralii 362.1 | NZ_QWBW01000012.1 | < mnmG | 564 | < glnQ | 22.4 | < rpmH | 567 | dnaA > | ORI97036383 | NZ_QWBW01000012.1 | | |
| 120 | Halomonas zincidurans B6 | NZ_JNCK01000001.1 | < mnmG | 587 | < glnQ | 88.9 | < rpmH | 569 | dnaA > | ORI97111763 | NZ_JNCK01000001.1 | | |
| 121 | Halomonas radicis EAR18 | NZ_CAAHFN010000048.1 | < mnmG | 638 | < glnQ | 22.4 | < rpmH | 572 | dnaA > | ORI97084766 | NZ_CAAHFN010000048.1 | | |
| 122 | Halomonas lutea DSM 23508 | NZ_KB899996.1 | < mnmG | 573 | < glnQ | 18.8 | < rpmH | 570 | dnaA > | ORI97195236 | NZ_KB899996.1 | | |
| 123 | Halomonas xianhensis CGMCC 1.6848 | NZ_FOPY01000006.1 | < mnmG | 586 | < glnQ | 41.9 | < rpmH | 570 | dnaA > | ORI97157485 | NZ_FOPY01000006.1 | | |
| 124 | Halomonas ilicicola DSM 19980 | NZ_FQUJ01000004.1 | < mnmG | 591 | < glnQ | 31.0 | < rpmH | 571 | dnaA > | ORI97182163 | NZ_FQUJ01000004.1 | | |
| 125 | Halomonas muralis DSM 14789 | NZ_FNGI01000004.1 | < mnmG | 611 | < glnQ | 19.9 | < rpmH | 564 | dnaA > | ORI97185044 | NZ_FNGI01000004.1 | | |
| | Salinicola | | | | | | | | | | | | |
| 126 | Salinicola halimionae CPA60 | NZ_PZJQ01000002.1 | < mnmG | 1026 | < glnQ | 28.9 | < rpmH | 594 | dnaA > | ORI97157562 | NZ_PZJQ01000002.1 | | |
| 127 | Salinicola halophyticus CR45 | NZ_PZJN01000009.1 | < mnmG | 975 | < glnQ | 30.3 | < rpmH | 617 | dnaA > | ORI97196575 | NZ_PZJN01000009.1 | | |
| 128 | Salinicola socius SMB35 = DSM 19940 | NZ_MSDO01000012.1 | < mnmG | 911 | < glnQ | 28.6 | < rpmH | 604 | dnaA > | ORI97075148 | NZ_MSDO01000012.1 | | |
| 129 | Salinicola salarius DSM 18044 | NZ_PZJT01000012.1 rpmH<br>NZ_PZJT01000021.1 mnmG | < mnmG | 761 | < glnQ | | < rpmH | 606 | dnaA > | | | | |
| 130 | Salinicola lusitanus CR50 | NZ_PZJM01000009.1 | < mnmG | 749 | < glnQ | 26.8 | < rpmH | 605 | dnaA > | ORI97176203 | NZ_PZJM01000009.1 | | |
| 131 | Salinicola acroporae LMG 28587 | NZ_PZJW01000003.1 | < mnmG | 748 | < glnQ | 26.8 | < rpmH | 604 | dnaA > | ORI97203256 | NZ_PZJW01000003.1 | | |
| 132 | Salinicola corii L3 | NZ_VTPX01000004.1 | < mnmG | 761 | < glnQ | 26.9 | < rpmH | 610 | dnaA > | ORI97101956 | NZ_VTPX01000004.1 | | |
| 133 | Salinicola peritrichatus JCM 18795 | NZ_PZJU01000083.1 rpmH<br>NZ_PZJU01000033.1 mnmG | < mnmG | 588 | < glnQ | | < rpmH | 618 | dnaA > | | | | |
| 134 | Salinicola rhizosphaerae KCTC 32998 | NZ_BMZI01000002.1 | < mnmG | 734 | < glnQ | 27.1 | < rpmH | 621 | dnaA > | ORI97174408 | NZ_BMZI01000002.1 | | |
| 135 | Salinicola tamaricis F01 | NZ_CP023559.1 | < mnmG | 795 | < glnQ | 29.2 | < rpmH | 622 | dnaA > | ORI97033057 | NZ_CP023559.1 | | |
| 136 | Salinicola endophyticus CPA92 | NZ_PZJO01000003.1 | < mnmG | 778 | < glnQ | 29.2 | < rpmH | 624 | dnaA > | ORI97178573 | NZ_PZJO01000003.1 | | |
| 137 | Salinicola halophilus CECT 5903 | NZ_PZJV01000006.1 | < mnmG | 850 | < glnQ | 25.2 | < rpmH | 684 | dnaA > | ORI97178279 | NZ_PZJV01000006.1 | | |
| 138 | Salinicola aestuarinus CPA62 | NZ_PZJP01000004.1 | < mnmG | 800 | < glnQ | 25.0 | < rpmH | 683 | dnaA > | ORI97168234 | NZ_PZJP01000004.1 | | |
| | Chromohalobacter | | | | | | | | | | | | |
| 139 | Chromohalobacter israelensis DSM 6768 | NZ_JQNW01000005.1 | < mnmG | 601 | < glnQ | 23.6 | < rpmH | 589 | dnaA > | | | | |
| 140 | Chromohalobacter salexigens DSM 3043 | NC_007963.1 | < mnmG | 602 | < glnQ | 23.6 | < rpmH | 589 | dnaA > | ORI97226780 | NC_007963.1 | | |
| 141 | Chromohalobacter marismortui DSM 6770 | NZ_SOBR01000004.1 | < mnmG | 614 | < glnQ | 23.4 | < rpmH | 589 | dnaA > | ORI97187899 | NZ_SOBR01000004.1 | | |
| 142 | Chromohalobacter canadensis 85B | NZ_JAJQJH010000002.1 | < mnmG | 605 | < glnQ | 23.6 | < rpmH | 590 | dnaA > | | | | |
| | Kushneria | | | | | | | | | | | | |
| 143 | Kushneria konosiri X49 | NZ_CP021323.1 | < mnmG | 622 | < erpA | 38.7 | < rpmH | 332 | dnaA > | ORI97227543 | NZ_CP021323.1 | | |
| 144 | Kushneria marisflavi SW32 = DSM 15357 | NZ_CP021358.1 | < mnmG | 622 | < erpA | 36.2 | < rpmH | 332 | dnaA > | ORI97227296 | NZ_CP021358.1 | | |
| 145 | Kushneria phyllosphaerae EAod3 | NZ_ONZI01000001.1 | < mnmG | 620 | < erpA | 43.0 | < rpmH | 332 | dnaA > | ORI97048201 | NZ_ONZI01000001.1 | | |

Suppl. Table 3



| # | Species / Strain | RefSeq / Nucleotide | 5' gene | ig | | 3' gene | oriC2 Δ oriC1 [kb] | 5' gene | ig | | 3' gene | DoriC 12 | RefSeq / Nucleotide | oriC2 | oriC1 |
|---|---|---|---|---|---|---|---|---|---|---|---|---|---|---|---|
| | Kushneria (cont.)) | | | | | | | | | | | | | | |
| 146 | Kushneria indalinina DSM 14324 | NZ_QRDJ01000007.1 | < mnmG | 622 | | < erpA | 38.3 | < rpmH | 330 | | dnaA > | ORI97151554 | NZ_QRDJ01000007.1 | | |
| 147 | Kushneria pakistanensis BKCTC 42082 | NZ_BMZM01000002.1 | < mnmG | 620 | | < erpA | 35.1 | < rpmH | 330 | | dnaA > | ORI97128315 | NZ_BMZM01000002.1 | | |
| 148 | Kushneria avicenniae DSM 23439 | NZ_FOLY01000003.1 | < mnmG | 623 | | < erpA | 43.2 | < rpmH | 330 | | dnaA > | ORI97110568 | NZ_FOLY01000003.1 | | |
| 149 | Kushneria aurantia DSM 21353 | NZ_KB907862.1 | < mnmG | 964 | | < erpA | 60.5 | < rpmH | 396 | | dnaA > | ORI97163849 | NZ_KB907862.1 | | |
| 150 | Kushneria sinocarnis DSM 23229 | NZ_RBIN01000001.1 | < mnmG | 607 | | < erpA | 52.4 | < rpmH | 390 | | dnaA > | ORI97128671 | NZ_RBIN01000001.1 | | |
| 151 | Kushneria phosphatilytica YCWA18 | NZ_CP043420.1 | < mnmG | 601 | | < erpA | 35.3 | < rpmH | 396 | | dnaA > | ORI97037491 | NZ_CP043420.1 | | |
| | Phytohalomonas | | | | | | | | | | | | | | |
| 152 | Phytohalomonas tamaricis R4HLG17 | NZ_PVRV01000019.1 | < mnmG | 623 | | < erpA | 27.1 | < rpmH | 458 | | dnaA > | ORI97076951 | NZ_PVRV01000019.1 | | |
| | Carnimonas | | | | | | | | | | | | | | |
| 153 | Carnimonas nigrificans ATCC BAA-78 | NZ_JAGO01000013.1 | < mnmG | 855 | | < erpA | 45.4 | < rpmH | 494 | | dnaA > | ORI97048426 | NZ_JAGO01000013.1 | | |
| | Halotalea | | | | | | | | | | | | | | |
| 154.1 | Halotalea alkalilenta DSM 17697 | NZ_KK211226.1 rpmH<br>NZ_KK211229.1 mnmG | < mnmG | 661 | | < erpA | | < rpmH | 428 | | dnaA > | ORI97154603 | NZ_KK211229.1 | | |
| 154.2 | Halotalea alkalilenta IHB B 13600 | NZ_CP015243.1 | < mnmG | 663 | | < erpA | 102.3 | < rpmH | 428 | | dnaA > | ORI97227272 | NZ_CP015243.1 | | |
| | Zymobacter | | | | | | | | | | | | | | |
| 155 | Zymobacter palmae IAM 14233 = DSM 10491 | NZ_AP018933.1 | < mnmG | 678 | | < COG2859 | 22.5 | < sdiA | 345 | | dnaA > | ORI97022155 | NZ_AP018933.1 | | |
| | Larsenimonas | | | | | | | | | | | | | | |
| 156 | Larsenimonas salina CCM 8464 | NZ_JAMLJJ010000003.1 | < mnmG | 562 | | < erpA | 33.2 | < rpmH | 405 | | dnaA > | | | | |
| 157 | Larsenimonas suaedae DSM 22428 | NZ_JARWAO010000001.1 | < mnmG | 568 | | < erpA | 28.2 | < rpmH | 392 | | dnaA > | | | | |
| | Cobetia | | | | | | | | | | | | | | |
| 158 | Cobetia marina JCM 21022 | NZ_CP017114.1 | < mnmG | 1388 | | < glnQ | 31.9 | < rpmH | 841 | | dnaA > | ORI97227378 | NZ_CP017114.1 | | |
| 159 | Cobetia crustatorum JO1 | NZ_JEMF01000015.1 rpmH<br>NZ_JEMF01000037.1 mnmG | < mnmG | seq gap | | < glnQ | | < rpmH | 943 | | dnaA > | ORI97078242 | NZ_JEMF01000037.1 | | |
| | uncertain | | | | | | | | | | | | | | |
| 160 | Terasakiispira papahanaumokuakeensis PH27A | NZ_MDTQ01000001.1 | < mnmG | 541 | | ompV > | 12.9 | < rpmH | 568 | | dnaA > | | | | |
| | not Halomonadaceae | | | | | | | | | | | | | | |
| 161 | Halovibrio variabilis DSM 3050 | NZ_BJXV01000011.1 | < mnmG | 617 | | < glnQ | 17.2 | < rpmH | 534 | | dnaA > | ORI97064817 | NZ_BJXV01000011.1 | | |
| 162 | Halovibrio salipaludis YL5-2 | NZ_NSKD01000001.1 | < mnmG | 286 | | < DUF3833 | 9.3 | < rpmH | 626 | | dnaA > | | | | |

* see de la Haba *et al.* (2023) Supplementary Fig 1

× inverted gene order

strong DUE

weak DUE

no DUE

no oriC

Suppl. Table 3



| Species / Strain | RefSeq / Nucleotide | DoriC 12 | RefSeq / Nucleotide | oriC2 | oriC1 |
|---|---|---|---|---|---|
| Species / Strain only in DoriC 12 | | | | | |
| Halomonas sp. KO116 | GCF_000734975.2 | ORI97227583 | NZ_CP011052.1 | ■ | |
| Halomonas sp. 'Soap Lake #7' | GCF_002009255.1 | ORI97227511 | NZ_CP019915.1 | ■ | |
| Halomonas sp. GT | GCF_002082565.1 | ORI97227437 | NZ_CP020562.1 | ■ | |
| Halomonas sp. SF2003 | GCF_003032495.1 | ORI97025281 | NZ_CP028367.1 | | ■ |
| Halomonas sp. 'Soap Lake #6' | GCF_003031405.1 | ORI97026512 | NZ_CP028367.1 | ■ | |
| Halomonas sp. 'Soap Lake #6' | GCF_003031405.1 | ORI97026512 | NZ_CP020469.1 | ■ | |
| Halomonas socia | GCF_010977575.1 | ORI97029328 | NZ_CP048812.1 | ■ | |
| Halomonas sp. PGE1 | GCF_013000985.1 | ORI97027731 | NZ_CP053032.1 | | ■ |
| Pistricoccus aurantiacus | GCF_007954585.1 | ORI97028094 | NZ_CP042382.1 | ■ | |
| Marinobacterium sediminicola | GCF_022343685.1 | ORI97028124 | NZ_CP092123.1 | ■ | |
| Bacterioplanes sanyensis NV9 | GCF_002237535.1 | ORI97028163 | NZ_CP022530.1 | ■ | |
| Vreelandella aquamarina SCSIO 43005 | GCF_009846525.1 | ORI97029489 | NZ_CP024621.1 | ■ | |
| Vreelandella titanicae S6 | GCF_014697095.1 | ORI97029687 | NZ_CP059082.1 | ■ | |
| Halomonas sp. HG01 | GCF_000968715.2 | ORI97029747 | NZ_CP022134.1 | ■ | |
| Halomonas sp. THAF5a | GCF_006542315.2 | ORI97030651 | NZ_CP039374.2 | ■ | |
| Vreelandella titanicae ANRCS81 | GCF_006542315.2 | ORI97030651 | NZ_CP039374.2 | ■ | |
| Halomonas sp. SS10-MC5 | GCF_015992245.1 | ORI97030758 | NZ_CP065435.1 | ■ | |
| Halomonas sp. FeN2 | GCF_020595105.1 | ORI97030989 | NZ_CP074200.1 | ■ | |
| Billgrantia tianxiuensis BC-M4-5 | GCF_009834345.1 | ORI97031521 | NZ_CP035042.1 | ■ | |
| Vreelandella piezotolerans HN2 | GCF_019443265.1 | ORI97032485 | NZ_CP080328.1 | ■ | |
| Vreelandella aquamarina Eplume2 | GCF_011764625.1 | ORI97032574 | NZ_AP022869.1 | ■ | |
| Halomonas sp. THAF12 | GCF_009363675.1 | ORI97033296 | NZ_CP045399.1 | ■ | |
| Cobetia sp. 4B | GCF_018831605.1 | ORI97033459 | NZ_JAIQZV010000004.1 | | ■ |
| Cobetia sp. 4B | GCF_018831605.1 | ORI97033459 | NZ_CP059843.1 | | ■ |
| Vreelandella populi PC 12 | GCF_003989805.1 | ORI97034820 | NZ_RZHC01000012.1 | ■ | |
| Halomonas citrativorans JB380 | GCF_900163645.1 | ORI97037180 | NZ_FUKM01000020.1 | ■ | |
| Kushneria phosphatilytica YCWA18 | GCF_001854625.1 | ORI97037491 | NZ_MKZT01000019.1 | ■ | |
| Halomonas sp. KM-1 | GCF_000246875.1 | ORI97038789 | NZ_BAEU01000018.1 | ■ | |
| Halomonas citrativorans FME16 | GCF_014861475.1 | ORI97039570 | NZ_RRZC01000003.1 | ■ | |
| Halomonas caseinilytica DSM 18067 | GCF_001662285.1 | ORI97042319 | NZ_BDEO01000008.1 | ■ | |
| Vreelandella aquamarina IOP_44 | GCF_021025935.1 | ORI97042647 | NZ_JAIQZV010000004.1 | ■ | |
| Halomonas sp. McD50-5 | GCF_022348125.1 | ORI97043926 | NZ_JAKRFH010000013.1 | ■ | |
| Vreelandella alkaliphila LS44 | GCF_010179695.1 | ORI97045443 | NZ_JAAEHK010000006.1 | ■ | |
| Halomonas sp. G15 | GCF_021282295.1 | ORI97045681 | NZ_JAJSPA010000026.1 | ■ | |
| Chromohalobacter israelensis MP25462 | GCF_013395975.1 | ORI97045860 | NZ_QEQS01000002.1 | ■ | |
| Vreelandella aquamarina NBRC 15608 | GCF_006540125.1 | ORI97046550 | NZ_BJOI01000029.1 | ■ | |
| Halomonas sp. DP5N14-9 | GCF_019802175.1 | ORI97047500 | NZ_JAHVHR010000001.1 | ■ | |
| Halomonas sp. ALS9 | GCF_001651035.1 | ORI97047817 | NZ_LXRG01000011.1 | ■ | |
| Halomonas sp. HAL1 | GCF_000235725.1 | ORI97048697 | NZ_AGIB01000080.1 | ■ | |
| Halomonas sp. ATBC28 | GCF_005937175.1 | ORI97048836 | NZ_VCQL01000014.1 | ■ | |
| Halomonas elongata MH25661 | GCF_003287205.1 | ORI97050223 | NZ_QJUB01000004.1 | ■ | |
| Halomonas sp. YLB-10 | GCF_003856855.1 | ORI97050539 | NZ_RRCU01000005.1 | ■ | |
| Salinicola sp. MIT1003 | GCF_001858555.1 | ORI97050884 | NZ_MCAP01000044.1 | ■ | |
| Cobetia sp. QF-1 | GCF_002213105.1 | ORI97053081 | NZ_NBVS01000005.1 | ■ | |
| Halomonas sp. CUBES01 | GCF_020991005.1 | ORI97053544 | NZ_JAJLLW010000001.1 | ■ | |
| Halomonas alkalisoli M5N1S15 | GCF_021412645.1 | ORI97053546 | NZ_JAKCMJ010000028.1 | ■ | |
| Halomonas sp. Choline-3u-9 | GCF_002836495.1 | ORI97055119 | NZ_PJBI01000057.1 | ■ | |
| Halomonas sp. S2151 | GCF_000967645.1 | ORI97055273 | NZ_JXYB01000041.1 | ■ | |
| Halomonas sp. MMH1-48 | GCF_022348105.1 | ORI97057459 | NZ_JAKRFL010000010.1 | ■ | |
| Vreelandella aquamarina ACAM 239 | GCF_900128865.1 | ORI97057563 | NZ_FSQX01000001.1 | ■ | |
| Halomonas sp. IOP_6 | GCF_021151565.1 | ORI97058115 | NZ_JAIRBO010000004.1 | ■ | |
| Halomonas casei FME64 | GCF_014897975.1 | ORI97058682 | NZ_JABUYY010000004.1 | ■ | |
| Billgrantia desiderata MCCC 1A05775 | GCF_021404445.1 | ORI97058704 | NZ_JABFTR010000018.1 | ■ | |
| Billgrantia desiderata MCCC 1A17486 | GCF_021404365.1 | ORI97059874 | NZ_JABFTY010000005.1 | ■ | |
| Halomonas elongata HEK1 | GCF_001678785.1 | ORI97062006 | NZ_MAJD01000002.1 | | ■ |
| Halomonas eurihalina MS1 | GCF_008274785.1 | ORI97062447 | NZ_VTPU01000015.1 | ■ | |
| Halomonas salina CIFRI 1 | GCF_000758175.1 | ORI97064600 | NZ_JOKD01000014.1 | ■ | |
| Vreelandella neptunia HM116 4 | GCF_022489065.1 | ORI97065128 | NZ_JAKVTW010000004.1 | ■ | |
| Halomonas colorata FME66 | GCF_014897985.1 | ORI97065348 | NZ_JABUZA010000006.1 | ■ | |
| Vreelandella venusta DSM 4743 | GCF_016107635.1 | ORI97068595 | NZ_JABASW010000003.1 | ■ | |
| Chromohalobacter israelensis voya40th042 | GCF_020076335.1 | ORI97068599 | NZ_JAIQWZ010000002.1 | ■ | |
| Halomonas hibernica B1.N12 | GCF_013183775.1 | ORI97068873 | NZ_VIFN01000013.1 | ■ | |
| Cobetia marina NBRC 15607 | GCF_006540105.1 | ORI97069072 | NZ_BJOH01000004.1 | ■ | |
| Halomonas sp. IOP_14 | GCF_021021315.1 | ORI97071669 | NZ_JAINWD010000007.1 | ■ | |
| Halomonas sp. PBN3 | GCF_000475415.1 | ORI97072454 | NZ_AXCA01000166.1 | ■ | |
| Halomonas elongata BLLS135 | GCF_014926355.1 | ORI97072854 | NZ_JADDHN010000004.1 | ■ | |
| MAG: Halomonas sp. isolate coassembly_bin.43 | GCF_020832125.1 | ORI97072864 | NZ_JAFIFI010000014.1 | ■ | |
| Halomonas litopenaei LZD012 | GCF_018424645.1 | ORI97074117 | NZ_QTKW01000013.1 | ■ | |
| Halomonas sp. DP8Y7-3 | GCF_019801675.1 | ORI97074723 | NZ_JAHVIO010000004.1 | | ■ |
| Halomonas denitrificans AmN35-1 | GCF_020171695.1 | ORI97074968 | NZ_JAIVAB010000003.1 | ■ | |
| Halomonas sp. 1513 | GCF_001971685.1 | ORI97227586 | NZ_CP019326.1 | ■ | |
| Halomonas sp. R57-5 | GCF_900070345.1 | ORI97227463 | NZ_LN813019.1 | ■ | |
| Halomonas beimenensis NTU-111 | GCF_002549795.1 | ORI97227386 | NZ_CP021435.1 | ■ | |
| Halomonas hydrothermalis Y2 | GCF_002442575.1 | ORI97227085 | NZ_CP023656.1 | ■ | |
| Modicisalibacter luteus KCTC 12847 | GCF_003989795.1 | ORI97206017 | NZ_BMXD01000006.1 | ■ | |
| Vreelandella piezotolerans NBT06E8 | GCF_009660035.1 | ORI97205467 | NZ_ML762934.1 | ■ | |
| Halomonas sp. MCCC 1A17488 | GCF_022097085.1 | ORI97205463 | NZ_JABEBV010000003.1 | ■ | |
| Modicisalibacter xianhensis 6_TX | GCF_004366455.1 | ORI97203341 | NZ_SOEC01000009.1 | ■ | |
| Salinicola sp. DM10 | GCF_017306975.2 | ORI97200548 | NZ_JAFLEN020000003.1 | ■ | |
| Halomonas sp. 1M45 | GCF_014897945.1 | ORI97195523 | NZ_JABUZC010000013.1 | ■ | |
| Halomonas sp. ISL-56 | GCF_018612805.1 | ORI97200069 | NZ_JAGGRO010000028.1 | ■ | |
| Halomonas sp. ISL-104 | GCF_018612735.1 | ORI97196496 | NZ_JAGGRM010000009.1 | ■ | |

Suppl. Table 3



| Species / Strain | RefSeq / Nucleotide | DoriC 12 | RefSeq / Nucleotide | oriC2 | oriC1 |
|---|---|---|---|---|---|
| Species / Strain only in DoriC 12 | | | | | |
| Salinicola sp. CPA57 | GCF_003206035.1 | ORI97199546 | NZ_PZJR01000010.1 | ■ | |
| Halomonas sp. JSM 104105 | GCF_010993975.1 | ORI97195403 | NZ_JAACZO010000058.1 | ■ | |
| Halomonas salina B6 | GCF_000821105.2 | ORI97202382 | NZ_LN651347.1 | ■ | |
| Halomonas caseinilytica CGMCC 1.6773 | GCF_900109905.1 | ORI97199611 | NZ_FOBT01000007.1 | ■ | |
| Vreelandella halophila P-halo | GCF_900162675.1 | ORI97197809 | NZ_LT727794.1 | | ■ |
| Vreelandella titanicae DSM 26667 | GCF_900099955.1 | ORI97195179 | NZ_FNEF01000009.1 | ■ | |
| Modicisalibacter sp. 'Wilcox' | GCF_010993675.1 | ORI97194807 | NZ_WNXF01000015.1 | | ■ |
| Halomonas sp. QHL1 | GCF_001882345.1 | ORI97194451 | NZ_MINL01000001.1 | ■ | |
| Halomonas sp. ISL-60 | GCF_018612795.1 | ORI97194372 | NZ_JAGGRP010000047.1 | ■ | |
| Salinicola sp. RZ23 | GCF_003206675.1 | ORI97194022 | NZ_PZJK01000001.1 | ■ | |
| Halomonas sp. KRD171 | GCF_017168345.1 | ORI97193038 | NZ_JABFAT010000004.1 | ■ | |
| Salinicola sp. CR57 | GCF_003206585.1 | ORI97192861 | NZ_PZJL01000001.1 | ■ | |
| Halomonas alkaliantarctica FS-N4 | GCF_000712975.1 | ORI97192038 | NZ_JHQL01000002.1 | ■ | |
| Cobetia crustatorum SM1923 | GCF_007786215.1 | ORI97191144 | NZ_VNFH01000003.1 | ■ | |
| Halomonas sp. KAO | GCF_015595025.1 | ORI97190440 | NZ_JADNRL010000019.1 | ■ | |
| Halomonas sp. 113 | GCF_904067645.1 | ORI97190362 | NZ_LR881453.1 | ■ | |
| Halomonas sp. NO4 | GCF_010994375.1 | not accessible | NZ_WHMA01000003.1 | | |
| Halomonas sp. BL6 | GCF_006228115.1 | ORI97187360 | NZ_VDGL01000002.1 | ■ | |
| Halomonas sp. ISL-106 | GCF_018612775.1 | ORI97186890 | NZ_JAGGRN010000010.1 | ■ | |
| Vreelandella profundi MT32 | GCF_019504665.1 | ORI97185492 | NZ_JAHRFZ010000004.1 | ■ | |
| Chromohalobacter canadensis USBA 855 | GCF_900221025.1 | ORI97185078 | NZ_OBQJ01000001.1 | ■ | |
| Halomonas sp. 15WGF | GCF_005218165.1 | not accessible | NZ_SSXT01000033.1 | | |
| Chromohalobacter japonicus CJ | GCF_000821045.2 | ORI97179604 | NZ_LN651366.1 | ■ | |
| Salinicola socius DSM 19940 | GCF_003206115.1 | ORI97179348 | NZ_PZJS01000015.1 | ■ | |
| Vreelandella populi RC 4 | GCF_003990195.1 | not accessible | NZ_RZHD01000004.1 | | |
| Onishia taeanensis As1101 | GCF_013349955.1 | ORI97176489 | NZ_JABVXC010000013.1 | ■ | |
| Halomonas sp. MCCC 1A17488 | GCF_021404315.1 | ORI97175080 | NZ_JABFTZ010000003.1 | ■ | |
| Halomonas sp. ANAO-440 | GCF_019915905.1 | ORI97172955 | NZ_JAHXBV010000020.1 | ■ | |
| Halomonas sp. 59 | GCF_904067605.1 | ORI97172808 | NZ_LR881455.1 | ■ | |
| Onishia taeanensis USBA-857 | GCF_003268885.1 | ORI97172098 | NZ_QLSX01000009.1 | ■ | |
| Halomonas sp. G11 | GCF_001663515.1 | ORI97171424 | NZ_LYXG01000003.1 | ■ | |
| Aidingimonas halophila KCTC 12885 | GCF_014651655.1 | ORI97169673 | NZ_BMXH01000004.1 | ■ | |
| Halomonas sp. LBP4 | GCF_003202205.1 | ORI97169327 | NZ_PDOG01000004.1 | ■ | |
| MAG: Halomonas meridiana M30B31 | GCF_018402055.1 | ORI97166738 | NZ_JAGYJC010000010.1 | ■ | |
| Vreelandella aquamarina IOP_19 | GCF_021021635.1 | ORI97165635 | NZ_JAINWP010000004.1 | ■ | |
| Halomonas sp. 19A_GOM-1509m | GCF_000620585.1 | ORI97165439 | NZ_JIAY01000018.1 | ■ | |
| Zymobacter palmae DSM 10491 | GCF_000620025.1 | ORI97165347 | NZ_JHVG01000001.1 | ■ | |
| Halomonas urumqiensis CGMCC 1.12917 | GCF_014653135.1 | ORI97164549 | NZ_BNAE01000002.1 | ■ | |
| Halomonas sp. SL1 | GCF_003030865.2 | ORI97161638 | NZ_PYUQ02000018.1 | ■ | |
| Halomonas sp. ML-15 | GCF_014779595.1 | ORI97161495 | NZ_JACXZT010000019.1 | ■ | |
| Halomonas sp. TD01 | GCF_000219565.1 | ORI97159101 | NZ_GL949756.1 | ■ | |
| Cobetia marina 402 | GCF_013350055.1 | ORI97158401 | NZ_JABVXD010000004.1 | ■ | |
| Halomonas sp. 23_GOM-1509m | GCF_000518325.1 | ORI97158249 | NZ_JADJ01000020.1 | ■ | |
| Halomonas sp. 3A7M | GCF_014897875.1 | ORI97157935 | NZ_JABUZD010000110.1 | ■ | |
| Halomonas sp. BC2 | GCF_002078115.1 | ORI97157436 | NZ_LFGT01000021.1 | ■ | |
| Halomonas flagellata EGI 63088 | GCF_022487945.1 | ORI97155980 | NZ_JAKVPY010000008.1 | ■ | |
| Halomonas heilongjiangensis 9-2 1262 | GCF_003202165.1 | ORI97155648 | NZ_PDOH01000014.1 | ■ | |
| Halomonas ventosae USBA 854 | GCF_003002995.1 | ORI97155527 | NZ_PVTM01000009.1 | ■ | |
| Halomonas sp. C22 | GCF_007655045.1 | ORI97155168 | NZ_VBVC01000008.1 | ■ | |
| Chromohalobacter israelensis 40a_TX | GCF_003182525.1 | ORI97155078 | NZ_QGTY01000011.1 | ■ | |
| Halomonas sp. 3D7M | GCF_014897885.1 | ORI97154388 | NZ_JABUZF010000091.1 | ■ | |
| Vreelandella titanicae isolate Halomonas sp. 153 | GCF_902506245.1 | ORI97153783 | NZ_LR733721.1 | ■ | |
| Halomonas campaniensis SK03 | GCF_002211105.1 | ORI97153329 | NZ_JPUA01000001.1 | ■ | |
| Halomonas sp. A11 | GCF_003182195.1 | ORI97153068 | NZ_QGTM01000005.1 | ■ | |
| Halomonas sp. BN3-1 | GCF_003056325.1 | ORI97152912 | NZ_PYVV01000012.1 | ■ | |
| Vreelandella titanicae SM1922 | GCF_007786325.1 | ORI97152539 | NZ_VNFE01000001.1 | ■ | |
| Halomonas sp. KHS3 | GCF_000830495.1 | ORI97151973 | NZ_JWHY01000001.1 | ■ | |
| Vreelandella titanicae TAT1 | GCF_013371485.1 | ORI97151612 | NZ_JABWTB010000001.1 | ■ | |
| Halomonas sp. DQ26W | GCF_003339685.1 | ORI97150949 | NZ_QPKI01000002.1 | ■ | |
| Halomonas urumqiensis BZ-SZ-XJ27 | GCF_002879635.1 | ORI97149963 | NZ_PNRG01000013.1 | ■ | |
| Halomonas sp. 156 | GCF_904067615.1 | ORI97149868 | NZ_LR881458.1 | ■ | |
| Halomonas ventosae 1_TX | GCF_004361885.1 | ORI97149522 | NZ_SNWH01000013.1 | ■ | |
| Halomonas sp. BC1 | GCF_002078135.1 | ORI97149070 | NZ_LFGS01000017.1 | ■ | |
| Halomonas sp. N3-2A | GCF_002216165.1 | ORI97017824 | NZ_CP022286.1 | ■ | |
| Cobetia sp. ICG0124 | GCF_004006355.1 | ORI97017898 | NZ_CP024893.1 | ■ | |
| Halomonas sp. JS92-SW72 | GCF_003547075.1 | ORI97018128 | NZ_CP032147.1 | ■ | |
| Halomonas sp. PA5 | GCF_013000965.1 | ORI97018643 | NZ_CP053025.1 | ■ | |
| Halomonas sp. TA6 | GCF_013004025.1 | ORI97018645 | NZ_CP053087.1 | ■ | |
| Halomonas sp. TA22 | GCF_013009075.1 | ORI97018646 | NZ_CP053108.1 | ■ | |
| Vreelandella venusta 4743 | GCF_016859375.1 | ORI97018926 | NZ_CP066539.1 | ■ | |
| Halomonas sp. hl-4 | GCF_900215475.1 | ORI97019632 | NZ_LT907845.1 | ■ | |
| Halomonas sp. A3H3 | GCF_000982975.1 | ORI97019738 | NZ_HG423343.1 | ■ | |
| Vreelandella titanicae GPM3 | GCF_013347205.1 | ORI97020859 | NZ_CP054580.1 | ■ | |
| Halomonas sp. NyZ770 | GCF_020616575.1 | ORI97021057 | NZ_CP085143.1 | ■ | |
| Vreelandella venusta PBH | GCF_015689495.1 | ORI97021180 | NZ_CP065135.1 | ■ | |
| Halomonas sp. SH5A2 | GCF_014263395.1 | ORI97022234 | NZ_CP058321.1 | ■ | |
| Cobetia sp. L2A1 | GCF_009796845.1 | ORI97024012 | NZ_CP047025.1 | ■ | |
| Cobetia sp. AM6 | GCF_009617955.1 | ORI97024527 | NZ_AP021868.1 | | ■ |
| Vreelandella venusta MA-ZP17-13 | GCF_003950215.1 | ORI97024745 | NZ_CP034367.1 | ■ | |
| Halomonas sp. GFAJ-1 ("the arsenic bacterium") | GCF_002966495.1 | ORI97024966 | NZ_CP016490.1 | | ■ |
| Halomonas sp. TD01 | GCF_923868895.1 | ORI97025071 | NZ_OV350343.1 | ■ | |
| Cobetia sp. cqz5-12 | GCF_016495405.1 | ORI97025675 | NZ_CP044522.1 | | ■ |

Suppl. Table 3



| Species / Strain | RefSeq / Nucleotide | DoriC 12 | RefSeq / Nucleotide | oriC2 | oriC1 |
|---|---|---|---|---|---|
| Species / Strain only in DoriC 12 | | | | | |
| Halomonas sp. 18071143 | GCF_019211725.1 | ORI97025982 | NZ_CP078120.1 | ■ | |
| Vreelandella aquamarina Slthf1 | GCF_011398715.1 | ORI97026070 | NZ_AP022821.1 | ■ | |
| Vreelandella profundi MT13 | GCF_019722725.1 | ORI97026104 | NZ_CP077941.1 | ■ | |
| Halomonas sp. A40-4 | GCF_015767635.1 | ORI97026145 | NZ_CP065230.1 | ■ | |
| Cobetia marina GPM2 | GCF_009931455.1 | ORI97026217 | NZ_CP047970.1 | ■ | |
| Halomonas sp. YLGW01 | GCF_014840935.1 | ORI97026510 | NZ_CP062005.1 | ■ | |
| Cobetia amphilecti N-80 | GCF_020217465.1 | ORI97027116 | NZ_CP084115.1 | | ■ |
| Billgrantia desiderata MCCC 1A05748 | GCF_021404505.1 | ORI97148525 | NZ_JABFTQ010000018.1 | ■ | |
| Halomonas sp. 3F2F | GCF_014897805.1 | ORI97148248 | NZ_JABUZE010000079.1 | ■ | |
| Litchfieldella anticariensis FP35 = DSM 16096 | GCF_000428505.1 | ORI97148114 | NZ_AUAB01000005.1 | ■ | |
| Vreelandella halophila DSM 3051 | GCF_019903425.1 | ORI97144297 | NZ_JACHUZ010000013.1 | | ■ |
| Chromohalobacter salexigens KG13 | GCF_013393405.1 | ORI97144202 | NZ_JABEVP010000007.1 | ■ | |
| Halomonas sp. I3 | GCF_904067555.1 | ORI97141763 | NZ_LR881461.1 | ■ | |
| Kushneria marisflavi DSM 15357 | GCF_003610515.1 | ORI97140413 | NZ_RAPM01000004.1 | ■ | |
| Halomonas caseinilytica ALO Sharm | GCF_900142315.1 | ORI97139928 | NZ_FRAL01000006.1 | ■ | |
| Vreelandella titanicae isolate Halomonas sp. 98 | GCF_902506335.1 | ORI97139041 | NZ_LR733707.1 | ■ | |
| Billgrantia desiderata MCCC 1A05776 | GCF_021404475.1 | ORI97138854 | NZ_JABFTS010000006.1 | ■ | |
| Cobetia marina MM1IDA2H-1AD | GCF_900119965.1 | ORI97137412 | NZ_FPLA01000102.1 | ■ | |
| Halomonas sp. DP8Y7-1 | GCF_019801685.1 | ORI97135248 | NZ_JAHVIN010000014.1 | ■ | |
| Halomonas sp. MES3-P3E | GCF_002836035.1 | ORI97133776 | NZ_PJBT01000002.1 | ■ | |
| Halomonas sp. FME65 | GCF_014897995.1 | ORI97130322 | NZ_JABUYZ010000042.1 | ■ | |
| Vreelandella aquamarina ACAM 255 | GCF_900142865.1 | ORI97129063 | NZ_FSRR01000001.1 | ■ | |
| Billgrantia kenyensis DSM 17331 | GCF_013697085.1 | ORI97128851 | NZ_JACEFT010000002.1 | ■ | |
| Vreelandella azerica TBZ9 8 | GCF_013112225.1 | ORI97128326 | NZ_JABFHI010000008.1 | | ■ |
| Halomonas alkalisoli M5N1S17 338 | GCF_021412585.1 | ORI97126831 | NZ_JAKCMI010000009.1 | ■ | |
| Halomonas sp. McD50-4 | GCF_022348135.1 | ORI97125947 | NZ_JAKRFG010000013.1 | ■ | |
| Halomonas sp. JB37 | GCF_002332255.1 | ORI97124659 | NZ_NRGW01000001.1 | ■ | |
| Cobetia marina T1 | GCF_005144735.1 | ORI97122741 | NZ_SWFN01000005.1 | ■ | |
| Halomonas sp. DSH1-27 | GCF_022348215.1 | ORI97122340 | NZ_JAKRFF010000010.1 | ■ | |
| Vreelandella titanicae AGSA3-1 03 | GCF_021395175.1 | ORI97120013 | NZ_JAJVKU010000003.1 | ■ | |
| Halomonas sp. MG34 | GCF_009898855.1 | ORI97119728 | NZ_WJPH01000008.1 | ■ | |
| Halomonas jincaotanensis TRM 85114 | GCF_018448785.1 | ORI97118230 | NZ_JAHCLU010000006.1 | ■ | |
| Cobetia sp. 5-25-4-2 | GCF_013374075.1 | ORI97118140 | NZ_BLWK01000004.1 | ■ | |
| Billgrantia desiderata SP1 | GCF_002151265.1 | ORI97117110 | NZ_MUMZ01000080.1 | ■ | |
| Halomonas sp. 141 | GCF_002810305.1 | ORI97116497 | NZ_PINZ01000007.1 | ■ | |
| MAG: Halomonas sp. isolate M30B70 | GCF_018401555.1 | ORI97116265 | NZ_JAGYKP010000081.1 | ■ | |
| Halomonas sp. TG39a | GCF_000744395.1 | ORI97115713 | NZ_JQLV01000001.1 | ■ | |
| Bisbaumannia pacifica CARE-V15 | GCF_016056335.1 | ORI97115197 | NZ_JAEDAF010000010.1 | ■ | |
| Vreelandella boliviensis kknpp38 | GCF_018310235.1 | ORI97114000 | NZ_JAGWDL010000005.1 | ■ | |
| Vreelandella venusta YYYZ-3 | GCF_013177515.1 | ORI97113911 | NZ_QDKN01000010.1 | ■ | |
| Halomonas sp. MCCC 1A11062 | GCF_021404355.1 | ORI97113122 | NZ_JABFTW010000001.1 | ■ | |
| Halomonas sp. DP4Y7-2 | GCF_019802305.1 | ORI97111453 | NZ_JAHVHM010000003.1 | ■ | |
| Halomonas sp. DP3Y7-1 | GCF_019800785.1 | ORI97110302 | NZ_JAHVKI010000003.1 | ■ | |
| Vreelandella neptunia DSM 15720 | GCF_019903445.1 | ORI97108761 | NZ_JACIVL010000005.1 | ■ | |
| Halomonas caseinilytica K4 | GCF_001722495.1 | ORI97108265 | NZ_LWGO01000009.1 | ■ | |
| Halomonas sp. CSM-2 | GCF_002119345.1 | ORI97106997 | NZ_NBYR01000012.1 | ■ | |
| Billgrantia desiderata MCCC 1A17499 | GCF_021404345.1 | ORI97103381 | NZ_JABFUA010000002.1 | ■ | |
| Halomonas citrativorans FME63 | GCF_014898005.1 | ORI97103170 | NZ_JABUYX010000009.1 | ■ | |
| Cobetia sp. 5-11-6-3 | GCF_013374055.1 | ORI97102600 | NZ_BLWJ01000005.1 | ■ | |
| Halomonas sp. MM17-29 | GCF_022348065.1 | ORI97101101 | NZ_JAKRFJ010000010.1 | ■ | |
| Halomonas sp. 328 | GCF_015645095.1 | ORI97099947 | NZ_JADOTW010000024.1 | ■ | |
| Halomonas sp. SYSU XM8 | GCF_003045765.1 | ORI97097831 | NZ_PXNT01000008.1 | ■ | |
| Halomonas sp. M20 | GCF_020622265.1 | ORI97096645 | NZ_JACNMQ010000003.1 | ■ | |
| Halomonas sp. SBBP1 | GCF_020428305.1 | ORI97086863 | NZ_VOGA01000004.1 | ■ | |
| Halomonas sp. DP5Y7-2 | GCF_019802025.1 | ORI97085615 | NZ_JAHVHY010000003.1 | ■ | |
| Halomonas colorata FME20 | GCF_014861465.1 | ORI97084334 | NZ_RRZB01000003.1 | ■ | |
| Halomonas denitrificans DP5Y7-1 | GCF_019802095.1 | ORI97084242 | NZ_JAHVHX010000002.1 | ■ | |
| Halomonas hydrothermalis MTCC 5445 | GCF_000800185.1 | ORI97084071 | NZ_JTDR01000006.1 | ■ | |
| Halomonas sp. KX33721 | GCF_001596805.1 | ORI97082208 | NZ_LVIA01000035.1 | ■ | |
| Halomonas sp. DP1Y21-3 | GCF_019801245.1 | ORI97081595 | NZ_JAHVJM010000003.1 | ■ | |
| Halomonas sp. DP4Y7-1 | GCF_019800705.1 | ORI97080740 | NZ_JAHVKS010000003.1 | ■ | |
| Chromohalobacter japonicus SMB17 | GCF_001937175.1 | ORI97079746 | NZ_MSDQ01000007.1 | ■ | |
| Cobetia sp. MM1IDA2H-1 | GCF_002916775.1 | ORI97078832 | NZ_MUEJ01000085.1 | ■ | |
| Halomonas sp. Cn5-12 | GCF_021532545.1 | ORI97077413 | NZ_JAKGAY010000014.1 | ■ | |
| Halomonas denitrificans DP3Y115-1 | GCF_020171385.1 | ORI97076644 | NZ_JAIUZN010000003.1 | ■ | |
| Halomonas sp. DP3Y7-2 | GCF_019800795.1 | ORI97076435 | NZ_JAHVKJ010000001.1 | ■ | |
| Halomonas sp. MCCC 1A11057 | GCF_021404425.1 | ORI97076374 | NZ_JABFTU010000007.1 | ■ | |
| Halomonas cupida NBRC 102219 | GCF_007991155.1 | ORI97076262 | NZ_BJXU01000060.1 | ■ | |
| Halomonas sp. PR-M31 | GCF_001039575.1 | ORI97075700 | NZ_JIBT01000076.1 | ■ | |
| Halomonas sp. JCM 19031 | GCF_001310335.1 | ORI97088422 | NZ_BAWX01000001.1 | ■ | |
| ? = not accessible: 2025-07-28 – 2025-07-30 | | | | | |
| Halomonas casei FME1 | GCF_014861495.1 | ? | NZ_RRZD01000007.1 | ? | ? |
| Halomonas salinarum G5-11 | GCF_012062845.1 | ? | NZ_WWNB01000012.1 | ? | ? |
| Halomonas alkaliantarctica 22509_21_Filter | GCF_009856325.1 | ? | NZ_WMEV01000004.1 | ? | ? |
| Halomonas sp. McH1-25 | GCF_022348165.1 | ? | NZ_JAKRFI010000002.1 | ? | ? |
| MAG: Halomonas sp. coassembly_bin.77 k141_1516 | GCF_020832045.1 | ? | NZ_JAFIFK010000088.1 | ? | ? |
| Halomonas sp. ND22Bw | GCF_003011945.1 | ? | NZ_PXWD01000003.1 | ? | ? |
| Salinicola sp. MH3R3-1 | GCF_001937155.1 | ? | NZ_MSDP01000007.1 | ? | ? |
| Vreelandella aquamarina ACH-L-8 | GCF_001576535.1 | ? | NZ_JYFV01000019.1 | ? | ? |
| Vreelandella aquamarina R1t3 | GCF_000943375.1 | ? | NZ_JZEM01000006.1 | ? | ? |
